# Extreme GC3 Codon Bias in a Novel Brown Seaweed Virus Results in Pseudoambigrammatic Characteristics

**DOI:** 10.1101/2025.03.14.643323

**Authors:** Rob J. Dekker, Wim C. de Leeuw, Han Rauwerda, Marina van Olst, Wim A. Ensink, Selina van Leeuwen, Job Cohen, Klaas R. Timmermans, Timo M. Breit, Martijs J. Jonker

## Abstract

Although viruses are often studied in relation to disease, they can also offer valuable insights into RNA functionality. Due to their vast range of hosts and high mutation rates, viruses can quickly adapt to changing environments. It is therefore expected that viruses exploit all possible RNA functionalities to develop strategies for host infection and propagation. Our aim is to discover RNA viruses with survival strategies that reveal unknown RNA characteristics and functionalities. We have chosen seaweed as host organism because relatively little is known about RNA viruses in seaweed. Additionally, seaweeds are evolutionary distant from terrestrial plants and inhabit environments with entirely different conditions, increasing the likelihood of discovering new RNA viruses with unknown survival strategies. Here we report a new, unclassified negative-stranded single-strand RNA virus from seaweed *Saccharina latissima* that showed a unique genome characteristic. The novel virus, named SalaUV-NL1, contains a large segment of 7.065 nucleotides coding for a 2.296 amino acids protein and a small segment of 2.120 nucleotides coding for a 328 plus a potential 315+ amino acids protein. The large segment also appeared to have an additional large open reading frame (ORF) on the negative strand, suggesting it could be an ambigrammatic virus. However, the virus displayed an extreme GC3 codon bias (98%) in the (+)ORF, which differs from the typical avoidance of just the reverse-complement stop codons seen in ambigrammatic viruses. Consequently, the (−)ORF appears to occur accidental and is unlikely to encode a functional protein. Therefore, we have termed our virus to be *pseudoambigrammatic*. Several other viruses with the same pseudogrammatic high GC3 codon bias were identified in Genbank, most without a (−)ORF. The SalaUV-NL1 virus codon bias surpasses the high GC3 codon bias of the *S. latissima* host genes, however other species that host pseudoambigrammatic viruses do not display a high GC3 codon bias. Comparison of several variants of SalaUV-NL1 showed that the relative nucleotide change rate, particularly for silent changes from A/T to G/C, was extremely high (up to 81%). This novel virus demonstrates a new survival strategy of RNA viruses that is not yet understood.

## Introduction

RNA viruses are frequently studied in the context of disease, but they also offer valuable insights into RNA functionality (Scholthof and Scholthof 2023). RNA is a versatile molecule that performs an exceptional set of cellular functions (Cech and Steitz 2014, Caprara 2000, Nilsen 2000, Haseltine and Patarca 2024, Cao *et al*. 2024). Despite the many RNA functions and characteristics discovered over the past decades, new findings continue to emerge. To contribute to the understanding of RNA, we chose to study RNA viruses through observational research, as they are among the smallest operational units in nature in which RNA plays a dominant role. Viruses infect all sorts of life and display a remarkable array of strategies to enter their hosts and to propagate (International Committee on Taxonomy of Viruses, ICTV 2023). This is possible because RNA viruses have a high mutation rate which, combined with natural selection, allows for quick adaptation to changes in their host environment (Wolf *et al*. 2018). These adaptations utilize the full capacity of known and unknown RNA functionalities. There is a saying: “Anything you can think of happening to RNA, there is a (plant0 virus somewhere doing exactly that” (Scholthof and Scholthof 2023). However, we believe that it should include, “and many things you would never think of.” Of course, the challenge remains to find RNA viruses with new characteristics and strategies that will elucidate or hypothesize yet unknown RNA functions. To optimize our chances for such discoveries, we turned to a group of host species where the knowledge about associated viruses is still limited, which is evolutionary distant from mammals and plants, that lives in an entirely different environment than terrestrial species, and is relatively easy to obtain: seaweed or macro-algae (Lachnit *et al*. 2016, Waldron *et al*. 2018 and 2020, Schroeder and Mckeown 2020, Chiba 2020, van der Loos *et al*. 2023, Dekker *et al*. 2024, Zhao *et al*. 2024).

A contemporary approach to discovering (parts of) new non-pathogenic viruses involves large-scale metagenomics experiments that often analyze pooled samples from various sources (Waldron *et al*. 2018 and 2020, Batson *et al*. 2021, van der Loos *et al*. 2023). In our studies, rather than relying on these large-scale metagenomics experiments to discover and broadly describe as many variant or novel viruses as possible, typically based on RNA dependent RNA polymerase (RdRp) protein homology (Yin and Fischer 2008, Edgar *et al*. 2022), we employ a more reductionistic approach (Dekker *et al*. 2024). Our goal is to discover the complete genome of new RNA viruses, including hard-to-recognize segments, using creative bioinformatics analyses to facilitate their discovery and delineate their genomic characteristics.

By studying RNA viruses in seaweed, we aim to uncover novel RNA functionalities that could contribute to a better understanding of RNA biology. In this study, we analyzed several species of green and brown seaweed, both with and without possible disease phenotypes. We discovered a previously unknown RNA virus with a bipartite genome. This virus exhibited an extreme codon bias in their genomic sequences as well as largely overlapping ORFs on both RNA strands, suggesting an ambigrammatic nature (Cook *et al*. 2013, deRisi *et al*. 2019). However, further analysis revealed an extremely high GC3 codon bias in this virus genome, leading to the discovery of a yet unknown pseudoambigrammatic characteristic. This unique feature was subsequently identified in viruses from other species as well.

## Material and methods

### Samples

The 24 S1 seaweed samples are essentially the same as in the smallRNA-seq experiment described in Deller *et al*. 2024: 8 *Ulva lactuca* (sea lettuce, Chlorophyta, green algae) samples (S1-1 to S1-S8), 12 *Saccharina latissima* (sugar kelp, Phaeophyceae, winged kelp, Phaeophyceae, brown algae) samples (S1-9 to S1-16, and S1-26 to S1-28) and 5 *Alaria esculenta* (brown algae) samples (samples S1-21 to S1-25) supplemented with two *S. latissima* samples S2-1 and S2-2. The samples originate from the Royal Netherlands Institute for Sea Research (NIOZ, location Yerseke, in the south of the Netherlands) and from a commercial development facility in the north of the Netherlands (Table 1).

**Table 1.**
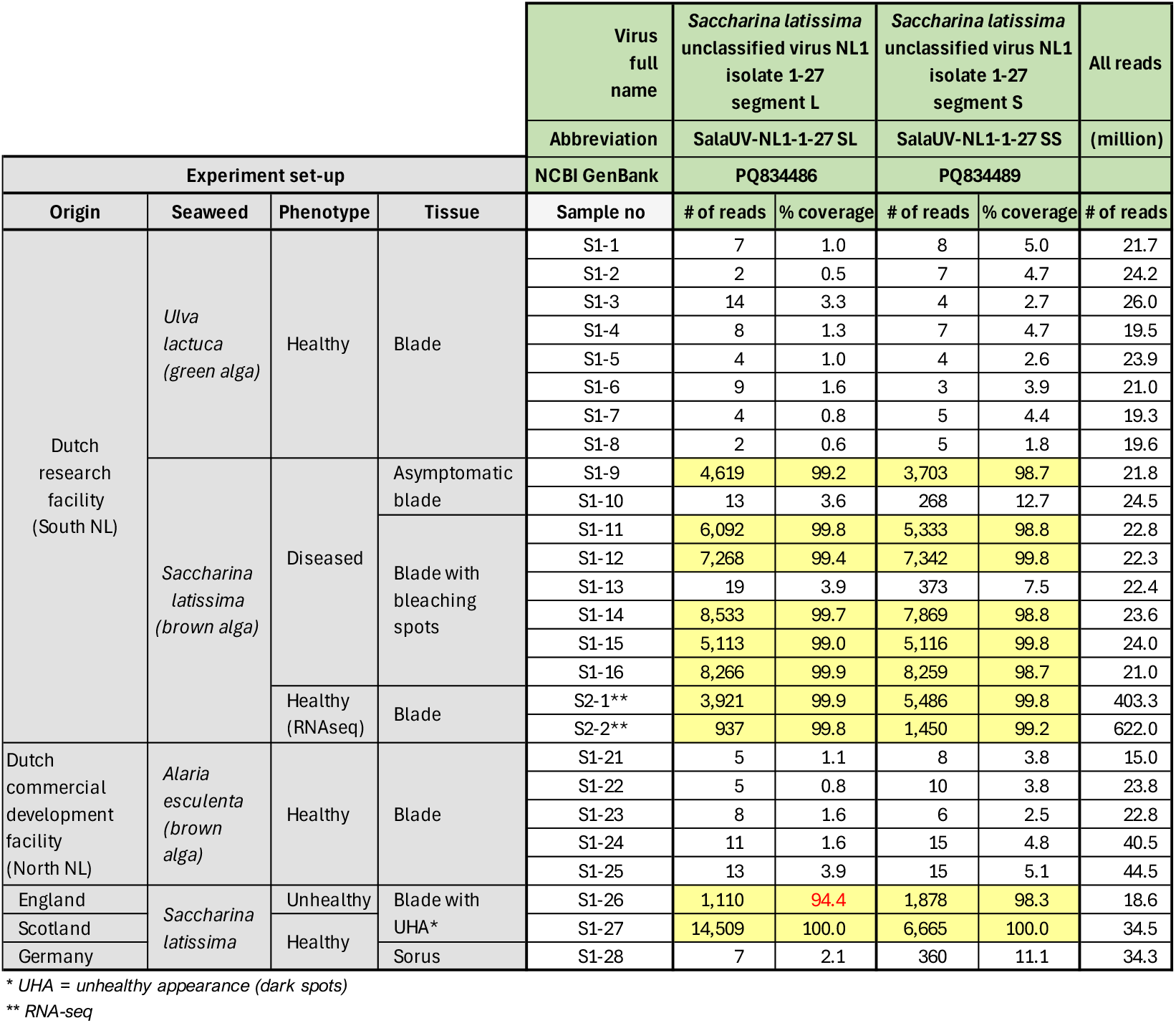
SalaUV-NL1 virus-derived siRNA presence in seaweed samples

### RNA isolation

RNA was isolated using a CTAB extraction buffer as described previously (Sim et al., 2013) with a minor adaptation of method 5, using a final concentration of 2.5 M LiCl. The concentration of the RNA was determined using a NanoDrop ND-2000 (Thermo Fisher Scientific) and the RNA quality was assessed using the 2200 TapeStation System with Agilent RNA ScreenTapes (Agilent Technologies).

### RNA-sequencing

Barcoded smallRNA-seq and RNA-seq libraries were generated using a Small RNA-seq Library Prep Kit (Lexogen) and a TruSeq Stranded Total RNA with Ribo-Zero Plant kit (Illumina), respectively. The size distribution of the libraries with indexed adapters was assessed using a 2200 TapeStation System with Agilent D1000 ScreenTapes (Agilent Technologies). The smallRNA-seq libraries were clustered and sequenced at 1×75 bp on a NextSeq 550 System using a NextSeq 500/550 High Output Kit v2.5 (75 cycles; Illumina). The two RNA-seq libraries were clustered and sequenced at 2×150 bp on a Novaseq X Plus System (Illumina).

### 5’ & 3’ RACE and Amplicon-seq

To generate amplicons, reverse transcription was performed using SuperScript IV Reverse Transcriptase (Thermo Fisher Scientific) with the Rv primer, following the manufacturer’s instructions. The cDNA was amplified using Q5 High-Fidelity 2X Master Mix (New England Biolabs). For the 5′ and 3′ RACE, 300 ng of total RNA was first polyadenylated using E. coli Poly(A) Polymerase (New England Biolabs). Subsequently, cDNA synthesis was performed using SuperScript IV Reverse Transcriptase (Thermo Fisher Scientific) and the oligo(dT)-adapter primer following the manufacturer’s instructions. The synthesized cDNA was used to generate amplicons with Q5 High-Fidelity 2X Master Mix (New England Biolabs), according to the provided protocol. Amplicons were purified using CleanNGS beads (CleanNA) at a 1.8x volume ratio. Size validation of the PCR products was performed using the 2200 TapeStation System with Agilent D1000 ScreenTapes.

The used primers are: F1_Fw-TCGACGCTGTAGTCCTCC; F2_Rv-AGTTGAGGTTGCACGTGAAG; F3_Fw-GCATCTCCTTCTCGTGCAAG; F4_Rv-GCACCCGTTCCTGATGAAG; BaNe3_Fw-TTCTGCGCGTAGAACTCCTC; BaNe5_Rv-GAGTTCGAGGCGGAGGTG; RACE-B_Rv-GCGAGCACAGAATTAATACGACTC; dT-B-GCGAGCACAGAATTAATACGACTCACTATAGG(T)_30_VN.

The purified PCR products were prepared for Nanopore sequencing using the Native Barcoding Kit 24 V14 (SQK-NBD114.24) and sequenced on a Flongle flow cell (Oxford Nanopore Technologies).

### Bioinformatics: mapping and assembly

Sequencing reads were trimmed using trimmomatic v0.39 (Bolger *et al*. 2014) (parameters: LEADING:3; TRAILING:3; SLIDINGWINDOW:4:15; MINLEN:19). Mapping of the trimmed reads to the NCBI virus database was performed using Bowtie2 v2.4.1 (Langmead *et al*. 2012). Contigs were assembled from smallRNA-seq data using all trimmed reads as input for SPAdes v3.15.5 De Novo Assembler (Prjibelski *et al*. 2020) with parameter settings: only-assembler mode, coverage depth cutoff 10, and kmer length 17, 18 and 21. Assembly of contigs from RNA-seq data was performed with default settings. Assembly of contigs from RNA-seq data was performed with default settings. Scanning of contig sequences for potential RdRp-like proteins was performed using PalmScan (Hou *et al*. 2024) and LucaProt (Babaian and Edgar 2022).

### Bioinformatics: determining variant virus sequences

Variants of the SalaUV-NL1 sequence in other samples were identified by mapping smallRNA-seq reads of each sample to the SalaUV-NL1-1-27 reference genome using Bowtie2 (v2.4.1), followed by variant detection with FreeBayes (v1.3.2) (Garrison and Marth 2012). Detected variants were incorporated into the reference to generate an updated sequence. This process of mapping, variant detection, and reference updating was repeated iteratively until no further changes were observed.

### Bioinformatics: determining phylogenetic trees

The phylogenetic trees of SalaUV-NL1 variant RNA and protein sequences were generated using CLC Genomics Workbench 24.0.2 (QIAGEN Aarhus A/S) using the ClustalO alignment option.

### Bioinformatics: co-presence analysis

The trimmed read reads of all samples where mapped using bowtie2 v.2.4.1 to all contigs (>200 nt) of each sample assembled by SPAdes by previously described method of sample S1-27. The read count for each contig in each sample was calculated using samtools idxstat v. 1.9 (Genome Research Ltd). This read count was normalized for each sample by dividing it by the total number of reads in the sample. Co-presence was estimated by calculating the spearman correlation between the normalized presence of a contig over the sample and that of the sample S1-27 contigs of interest (NODE_1 and NODE_6). The 20 contigs with the highest correlation with the contig of interest were selected.

### Bioinformatics: finding (pseudo)ambigrammatic viruses in GenBank

On 18 Oct 2024 a selection of virus sequences, with a non-human host, were downloaded from NCBI Virus (https://www.ncbi.nlm.nih.gov/labs/virus), which have a length ranging between 2,500 nt and 29.600 nt. To ensure the exclusion of artificial or modified sequences, any entries containing the terms “Modified Microbial Nucleic Acid”,” vaccine”, “Patent”, “recombinant”, or “Expressions System” were removed. Subsequently, for each non-phage remaining sequence, the longest non-stop-ORF in an arbitrary frame without a stop codon was identified in both forward and reverse frames using R (v4.3.2). The overlap between the longest forward and reverse ORFs was also calculated. Finally, for these ORFs, the GC content at the third codon position (GC3) and the overall GC content were determined.

Potential ambigrammatic virus sequences were identified based on two criteria: an overlap greater than 75% between the (+)ORF and (–)ORF, and a C/G content below 70%.

Potential pseudoambigrammatic virus sequences were identified based on two criteria: a (+)ORF GC3 percentage over 90%, and a C/G content below 65%.

### Bioinformatics: calculating the CG3 of host organisms

Protein-coding genes of the target organism were retrieved from NCBI using e-utils (v15.6) with the query “txid<tax_id>(Organism:exp) AND biomol_mrna(PROP)”, where <tax_id> was replaced with the organism’s NCBI taxonomy ID. The longest open reading frame (ORF) for each sequence was identified using the ORFik R package (v1.22.2) (Tjeldnes *et al*. 2021) in R (v4.3.2). Sequences were then split into codons, and the last nucleotide of each codon was tabulated using standard R functions.

### Bioinformatics: Saccharina latissima transcriptome assembly and mapping

Paired-end RNA sequencing data was downloaded from the SRA (run ERR4387699). Quality control was performed with FastQC (Andrews 2010). Reads were trimmed with Trimmomatic (Bolger *et al*. 2014). High quality reads were assembled with rnaSPAdes (Bushmanova *et al*. 2019). Using the ORFik R package (Tjeldnes *et al*. 2021) the longest ORF was determined in each contig. The resulting *S. latissima* transcriptome contigs can be found in Supplemental Document SD2.

Expression levels were calculated by mapping and counting the reads of sample S2-1 to the ORFs. The CG3 percentages and codon usages were calculated using standard R functions.

## Results and discussion

### A new unclassified virus species in Saccharina latissima (SalaUV-NL1)

To investigate the presence of unknown viruses with potential novel RNA-based survival strategies in edible brown and green algae, we analyzed the virus-derived siRNA presence in eight *U. lactuca*, 11 *S. latissima*, and five *A. esculenta* samples, with and without phenotypic abnormalities, from an academic and a commercial research facility in the South and the North of the Netherlands, respectively (Table 1). Previously, we reported on the discovery of four new viruses in these *S. latissima* and *A. esculenta samples* (Dekker *et al*. 2024). The sRNA-seq yielded roughly 641 M reads, resulting in an average of approximately 49 M reads per sample. Additionally, we analyzed two *S. latissima* samples with RNA-seq which yielded on average 513 M reads per sample (Table 1).

After assembly (Supplementary Document SD1) plus additional amplicon PCR and 5’/3’ RACE analyses, followed by sequencing, we identified a virus sequence of 7.065 nucleotides (nt) (Figure 1A). This sequence contains large ORFs in both orientations: one ORF on the positive strand (+)ORF), coding for a 2.296 amino acid (aa) polyprotein), and one ORF on the negative strand (−)ORF), putatively coding for a 2.317 aa polyprotein (Figure 1A). This is characteristic of ambigrammatic viruses (Cook *et al*. 2013, deRisi *et al*. 2019). RdRp analysis (https://serratus.io) revealed an RdRp domain in the protein from the (+)ORF, indicated by the presence of a RdRp conserved RdRp palmprint (palmID, Figure 1D). This analysis also identified the virus as possibly belonging to the phylum *Negarnaviricota* (realm: Riboviria, kingdom: Orthornavirae, Kuhn JH, *et al*. 2023), which comprises negative-stranded viruses with a single-stranded RNA genome. Supporting evidenced arose from a noticeable feature of the new genomic virus sequence i.e. the reverse complement similarity between the first 54 nt, which can form a panhandle RNA structure with a characteristic protruding cytosine at position 11 in the 5’ end sequence (Figure 1C) (Ren *et al*. 2020, Malet *et al*., 2023). Such panhandle structures are often present in *Negarnaviricota* viruses, for instance in Bunyaviruses. Furthermore, the size distribution of the virus-associated sequencing reads shows a peak at 21 nt, indicating the presence of siRNAs (Figure 1H). These 21-mer reads are distributed across the entire virus sequence (Figure 1E). The presumable siRNAs also shown a bias for their 5’ first nucleotide, being uracil (84%) and 3’ final nucleotide being cytosine (42%) rather than adenine (8%) (Figure 1G), which is a hallmark of siRNAs in animal, plant and fungal species (Kim 2008, Waldron *et al*. 2018).

**Figure 1.**
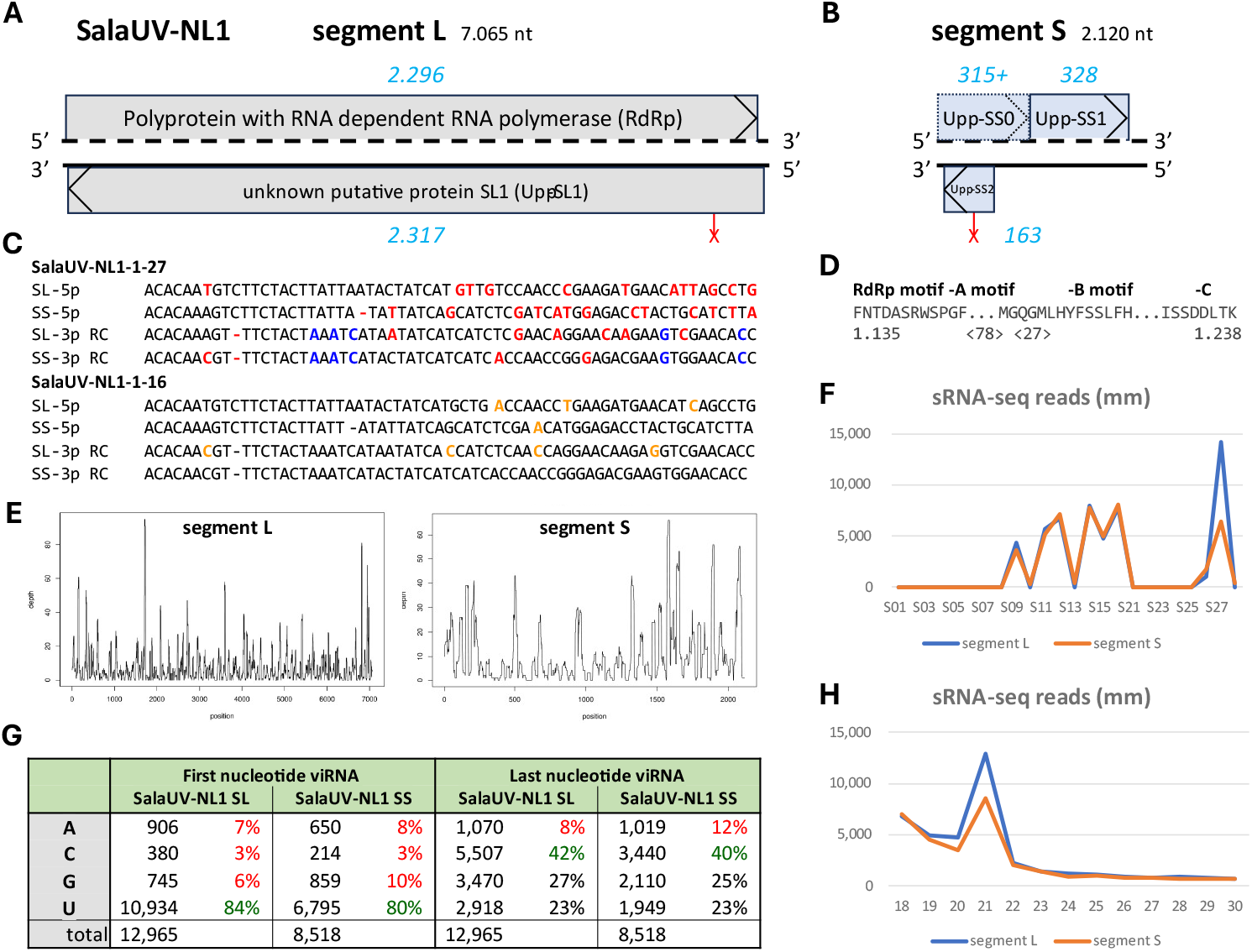
Schematic representation of Saccharina latissima Unclassified Virus NL1 and characteristic of related siRNAs. A) Schematic representation of the genome, the polyprotein, and the mature proteins of the Saccharina latissima Unclassified Virus NL1 isolate 1-27 large segment (SalaUV-NL1-1-27 SL). Indicated are the proposed polyproteins (grey), protein sizes (blue) and variant stop codon (red X). (RdRp). B) Also, for SalaUV-NL1-1-27 small segment (SS). C) Similarities between the 63 nt 5’ and 3’ terminus (panhandle) sequence motifs of the two RNA segments of the SalaUV-1 virus from two samples. To facilitate interpretation of the sequence complementarity, the reverse complement (RC) of the 3’ terminal sequences are presented. Nucleotides that differ from the preferred nucleotide are highlighted in red or blue if they are similar in the 3p sequences. Nucleotides that differ between variants 1-27 and 1-16 are highlighted in orange. D) The motifs of the RdRp Palmprint. E) SalaUV-1 related sRNA-seq read counts per virus segment position. F) Sample distribution plot for SalaUV-1 related sRNA-seq read counts. G) Summary of nucleotide occurrence at the ends of SalaUV-1 related sRNA-seq reads (viRNA) from all samples. H) Size distribution plot for SalaUV-1 related sRNA-seq reads.

A GenBank BLAST search of the RNA and both polyprotein sequences revealed that the most similar known virus sequence is “Barns Ness serrated wrack bunya/phlebo-like virus 1 (BNswBPV1) isolate C7, segment L” (MF190051.1) (Waldron *et al*. 2018). The RNA sequence identity is 68.2% (at 33% coverage) and the protein sequence identity (RdRp ORF) is 34.5% (at 94% coverage, Supplemental Table ST1). The (−)ORF protein did not show any protein hits in GenBank, besides a wrongly annotated ambigrammatic virus RdRp sequence (QJW70364.1). Further analysis into the aa composition revealed that the (−)ORF putative protein virtually lacks (<1%) ten of the 20 possible aa types, rendering a functional protein less likely (Supplemental Table ST2). Mapping the sRNA-seq reads and RNA-seq reads of all samples to this new virus sequence, revealed presence of this new virus in ten samples originating from both facilities (Table 1 and Figure 1F). Given that the relatively dissimilar virus BNswBPV1 is already classified as just bunya/phlebo-*like* and the new virus was found exclusively in *S. latissima* samples (Table 1), we named the new virus *S. latissima* unclassified virus number 1, from the Netherlands (SalaUV-NL1, King *et al*., 2012).

### Segment S of *Saccharina latissima* unclassified virus 1 (SalaUV-NL1 SS)

Another characteristic of *Negarnaviricota* viruses is their multisegmented RNA genome (Kuhn JH, *et al*. 2023). Given the considerable size of the new virus sequence, as well as the presence of a RdRp domain, we assumed it to be a large segment (SalaUV-NL1 SL). Next, we set out to find possible other, smaller genomic RNA segments of the new SalaUV-NL1 virus. As we did not have any virus sequence to identify such an RNA segment, we first used the 5’ and 3’ panhandle sequences to search for similar sequences at the ends of contigs from the sample with the most reads present for the new virus sequence (S27). This yielded an RNA contig of 2.120 nt with almost identical panhandle sequences for the first 15 nt at both ends (Figure 1C). We also applied an alternative approach by mapping all 21-mer reads of all sRNA-seq samples to all assembly contigs of sample S27 (Supplemental Document SD2) and identifying the contigs that showed a presence distribution over the samples which was almost identical to SalaUV-NL1 SL (Table 1, Figure 1F, and Supplemental Table ST3). Interestingly, the contig with the best read-distribution correlation, was the same one identified using the panhandle-sequence approach. Furthermore, sRNA-seq reads associated with this contig show the same 21-mer peak in their size distribution (Figure 1H) and the same prevalence for the first and last nucleotide as the SalaUV-NL1 L segment (Figure 1G). Hence, we concluded that, considering its size of 2.120 nt (Figure 1B), this contig presumably represented the small segment of the new virus (SalaUV-NL1 SS). As we were unable to find any other contigs or sRNA-seq reads that showed obvious similarity to the observed four panhandle sequences (Figure 1C) of the new virus, and we did also not find any contigs with conclusive co-presence over these samples, we assumed that there is no Medium (M) segment, although there have been viruses reported with panhandle sequences for the L and M segments, but not the S segment (Ren *et al*. 2020, Malet *et al*., 2023). Also, we were unable to identify a coat protein by similarity searches. The small RNA segment SalaUV-NL1 SS contained an (+)ORF for a putative protein of 328 aa (Upp-SS1) and an (−)ORF of 163 aa (Upp-SS2, Figure 1B), probably again not coding for a putative codon as it virtually lacks six aa types (Supplemental Table ST2). Both proteins showed no convincing similarity to any GenBank virus protein sequence.

### SalaUV-NL1 variants

siRNAs related to the newly identified virus, SalaUV-NL1 were detected in eight samples: six from the NIOZ seaweed cultivation facility and two from the NL-North facility involved in this study (Table 1). The siRNA read coverage for both SalaUV-NL1 segments from the six samples of the NIOZ facility averaged 99.6% (Supplemental Table ST4). To identify substantial virus sequences differences between the infected seaweed samples, we adapted the original SalaUV-NL1-1-27 virus sequence, based on siRNAs present in each sample (Supplemental Document SD2). The RNA and protein sequence similarity between the NIOZ facility sample variants was high: 99,8% and 99.9%. In contrast there were quite some differences between NIOZ facility virus variants as compared to the North-facility/Scotland-originating SalaUV-NL1-1-27: 97.2% and 96.0%, respectively (Supplemental Table ST4). These differences are clearly illustrated by the phylogenetic trees of all SalaUV-NL1 variant sequences (Supplemental Figure SF1). Furthermore, these RNA sequence differences led to an early stop codon at position 526 on the negative strand of the large segment in the SalaUV-NL1 variants from the NIOZ facility, which effectively truncates the putative protein encoded by the (−)ORF from 2,317 aa to just 165 aa (Figure 1A). This suggests again that the putative protein from the (−)ORF is non-essential for this virus (deRisi *et al*. 2019). As a surprising coincidence, we also discovered a truncating stop codon in the small segment-encoded short (−)ORF, Upp-SS2 (Figure 1B). This small putative protein, originally 163 aa long, is truncated to 64 aa, due to this stop codon at position 1,760 of the S segment in the SalaUV-NL1 variants from the NIOZ facility.

### The SalaUV-NL1 host

The genomic sequence of the virus SalaUV-NL1 was determined through metagenomics, making it challenging to identify its actual host. While S. latissima is the likely host, it is also possible that the host could be another organism, such as a fungus, infecting the seaweed. To address this uncertainty, scientists often examine their metagenomics data for genomic traces of other organisms. However, while some scientists, if they do find another pathogenic organism, conclude that the virus originates from that organism (Waldron *et al*. 2018, Batson *et al*. 2021), we argue that this approach is less effective in non-pooled samples. Instead, we applied a co-presence analysis across our experimental sRNA-seq samples. SalaUV-NL1 was present only in eight out of these 24 samples, so we determined contigs with similar presence patters. As expected, the SalaUV-NL1 contigs were the highest co-presence contigs. Of the remaining top 10 co-presence contigs per SalaUV-NL1 segment, 16 (80%) showed similarity to seaweed sequences in GenBank (Supplemental Table ST3). This outcome strongly suggests that *S. latissima* is the host of the SalaUV-NL1 virus. Remarkably, the regions that were consistently found, were two adjacent S. japonica U-linked SDR genomic sequences (Luthringer *et al*., 2015, Du *et al*. 2022), suggesting that this region may be involved in seaweed virus infection.

### Ambigrammatic viruses

For known ambigrammatic viruses, it is understood that the reverse-complement codons (UUA, CUA, and UCA) of the stop codons (UAA, UAG, and UGA) are absent from the RdRp (+)ORF (Cook *et al*. 2013, deRisi *et al*. 2019). This absence ensures that there are no stop codons in the (−)ORF, resulting in a non-stop (−)ORF of at least similar size of that of the (+)ORF. The presence of an AUG codon (M) early in the sequence will render it a regular ORF, and thus ambigrammatic virus. There have been several reports of ambigrammatic viruses (Cook *et al*. 2013, deRisi *et al*. 2019, Dinan *et al*. 2020. Batson *et al*. 2021, Zhang *et al*. 2023, Abbo *et al*. 2023, Cheng *et al*. 2024). To get an overview of known ambigrammatic viruses, we analyzed all available GenBank virus sequences, excluding those annotated “human”, for largely overlapping ORFs on both strands. About 114 out of 221K virus sequences showed an overlap between (+)ORF and (−)ORF higher than 75%, while having a C/G content <70% (Supplemental Table ST5), many (40) being Culex narnavirus (Goertz *et al*. 2019). For these ambigrammatic viruses, keeping the non-RdRp ORF intact appears to be important (Wilkinson *et al*. 2021, Retallack *et al*. 2021, Dudas *et al*. 2021). This feature makes the ambigrammatic nature of SalaUV-NL1 somewhat puzzling, as the virus does not seem to require the putative protein coded by the (−)ORF, demonstrated by the virtually lack of about half of the aa types (Supplemental Table ST2), as well as a variant from the NIOZ facility containing a dramatically truncating stop codon (Figure 1A). The most similar virus sequence (BNswBPV) also has no protein coding (−)ORF, although this is caused by the absence of a start codon for the non-stop ORF. We therefore investigated the origin of the ostensible ambigrammatic nature of SalaUV-NL1.

### Pseudoambigrammatic characteristics of SalaUV-NL1

A quick analysis of the codon usage in the SalaUV-NL1 LS RdRp ORF revealed not only the absence of the reverse-complement stop codons, but also that almost all codons have a G or a C at the 3^rd^ position of the codon (A, 1.0%; U, 0.9%; C, 47.0%; G, 51.0%, (Table 2 and Supplemental Table ST6). Hence, the (+)ORF of this virus has a third-base G+C percentage (GC3) of over 98%, which is also true for the (+)ORF of BNswBPV1 (Table 2). The most extreme GenBank example of a virus sequence with such a high GC3 is “Trichoderma gamsii negative-stranded virus 1” (Table 2), with only one codon ending with a U and two with an A, for an (+)ORF of 2,268 codons and a non-stop (−)ORF of 2,300 codons with no start codon at all. Thus, it appears that the absence of reverse-complement stop codons, all having an A as third base, is merely a “byproduct” of the fact that virtually all third-base A and U are avoided (Supplemental Table ST6) and is not intended to produce a (−)ORF functional protein. We therefore propose to characterize our SalaUV-NL1 virus as *pseudoambigrammatic*.

**Table 2.**
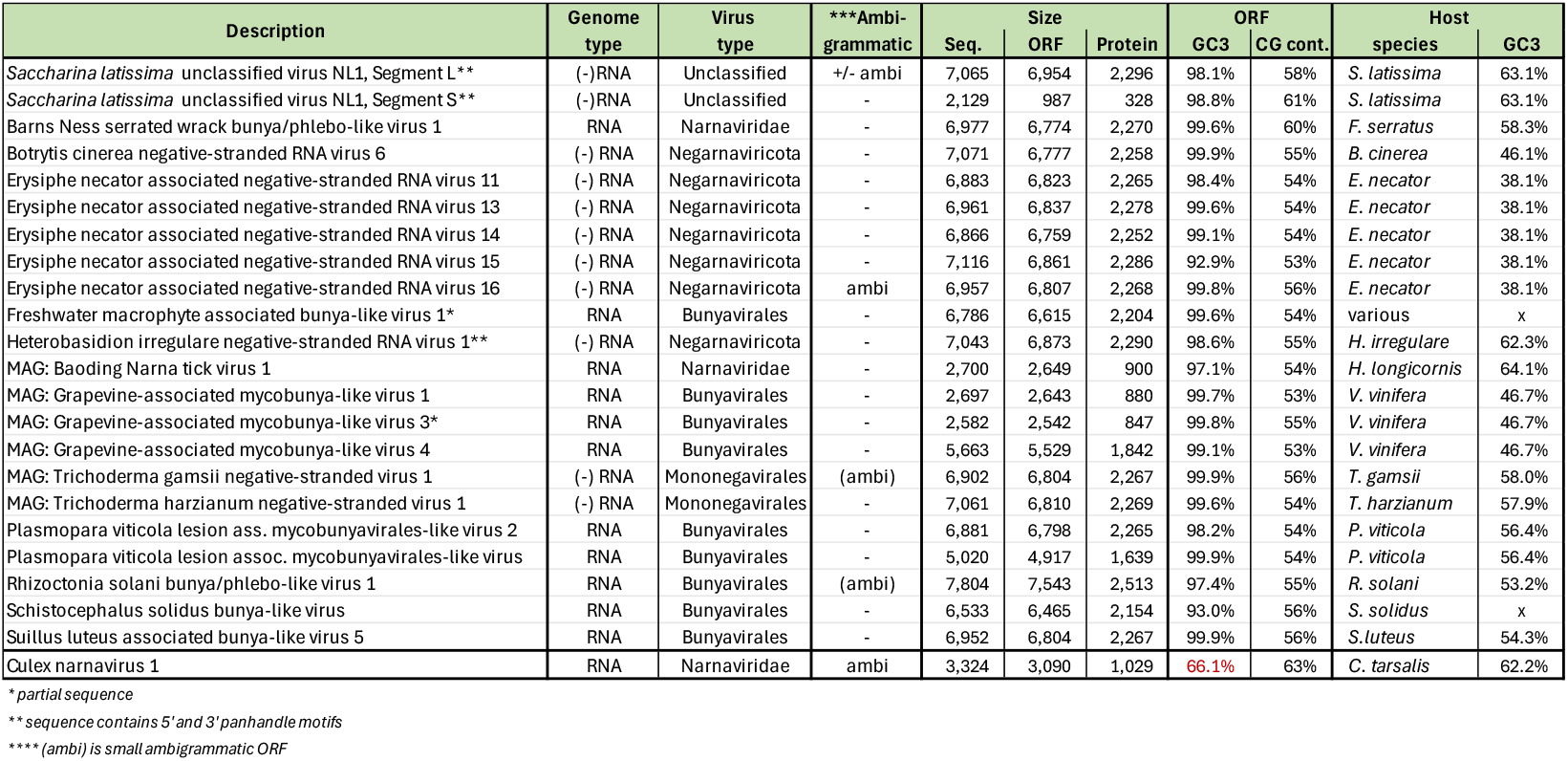
Summary of GenBank pseudoambigrammatic virus sequences with extreme GC3 codon bias

To emphasize the biological relevance of the high GC3 percentage, the utmost 5’ and 3’ ends of the SalaUV-NL1 large segment displayed a noticeable low GC3 percentage (21% and 25%, respectively) (Supplemental Figure SF2). Moreover, the small segment of SalaUV-NL1 also exhibit a high GC3 percentage (98.8%) (Table 2), as well as the low GC3 percentages (13% and 21%) at its 5’ and 3’ ends, respectively. This is remarkable since there appears to be only a relatively small (+)ORF present in this small virus segment (Figure 1B). There is however, a non-stop (+)ORF (315 codons) present from the start of the small segment (+) strand until the stop codon right before the ATG of (+)ORF Upp-SS1. Given that SalaUV-NL1 is a negative-strand virus, it is possible that this segment uses cap-snatching including a start codon to achieve another protein coding (+)ORF (Upp-SS0, Figure 1B, Ho *et al*. 2020), which would explain the presence of the high GC3 percentage in this virus segment. Despite this, we noticed that the GC3 high region in both virus segments extended beyond of the proposed (+)ORFs. This is intriguing, as it raises questions about whether the GC3 pattern is truly a result of just translation processes.

### Other pseudoambigrammatic viruses

Pseudoambigrammatic viruses are relatively easy to discover in databases, because they display a signature high GC3 percentage, while having an average C/G content. It is important to note that an overall high C/G content also can result in the absence of reverse-complement stop codons, as all three have at least one A and one U. Hence, we scanned the GenBank database for virus sequences that displayed a high GC3 percentage (>90%), but an around average overall C/G content percentage (<65%). The database search yielded 20 such virus sequences, which are, except for two, categorized as Negarnaviricota, a phylum of negative single-stranded viruses that commonly have a segmented genome (Table 2 and Supplemental Table ST7). The high GC3 percentage likely results in a codon bias, which can be defined by the “effective number of codons (ENC)” (Wright 1990). The pseudoambigrammatic virus ORFs showed an average ENC of about 27 (Supplemental Table ST7), whereas the true ambigrammatic sequences have an average ENC of 49 (Supplemental Table ST5). Comparing the codon usage of these pseudoambigrammatic viruses revealed substantial correlation between the SalaUV-NL1 SL (+)RdRp ORF and that of the other pseudoambigrammatic viruses, which is unsurprising given the extremely high GC3 bias (Supplementary Table ST8). Only one A-ending codon (ATA) was substantially present (>10% codon frequency) in 10 out of 21 (48%) virus sequences. Surprisingly, the GC3 bias also applied to the stop codons, where 16 out of 18 (89%) were TAG. Furthermore, a protein sequence comparison revealed that, with some exceptions, there is little identity between these pseudoambigrammatic virus protein sequences (Supplemental Table ST1). On the other hand, there is a substantial correlation between the SalaUV-NL1 SL RdRp protein aa distribution and that of the RdRp polyproteins of the pseudoambigrammatic viruses (Supplemental Table ST9). The lowest correlating virus protein (QXN75452.1, 79%) aa distribution is correlating much stronger than the RdRp protein aa distribution of a true ambigrammatic virus (QBR53296.1, 43%).

### Origin of codon usage bias of SalaUV-NL1

High GC3 codon usage bias is known to occur in highly expressed genes (Quax *et al*. 2015, Wint *et el*. 2022, Parvathy *et al*. 2022). However, most of the *S. latissima* transcriptome shows a relatively high GC3 codon bias (average 63.1%, Figure 2A and 2B, Table 2) (Luthringer *et al*. 2015) that does not seem to correlate with gene expression (Figure 2C). Even so, the GC3 percentage in SalaUV-NL1 (98%) is higher than that of almost all *S. Latissima* transcripts (Figure 2A). Additionally, there is an obvious correlation between the GC3 percentage and the GC3 codon bias of SalaUV-NL1 (Figure 2D). Together, these findings suggest that the GC3 codon bias observed in SalaUV-NL1 may originate from the virus mimicking the codon usage of the majority of genes of its host organism. To investigate whether this holds true for other pseudoambigrammatic viruses, we examined the GC3 codon bias in the host organisms from other pseudoambigrammatic viruses (Supplemental Table ST7). The GC3 codon percentage in these organisms ranged from 38% to 64% (Table 2), indicating no relevant relationship between the GC3 codon bias of pseudoambigrammatic viruses and the GC3 codon usage of their host organisms. Yet because the SalaUV-NL1 values in a GC3 percentage and expected number of codons plot are well below the neutrality line (Figure 2B), the GC3 codon usage is non-random and caused by selection pressure (Zhou *et al*. 2015).

**Figure 2.**
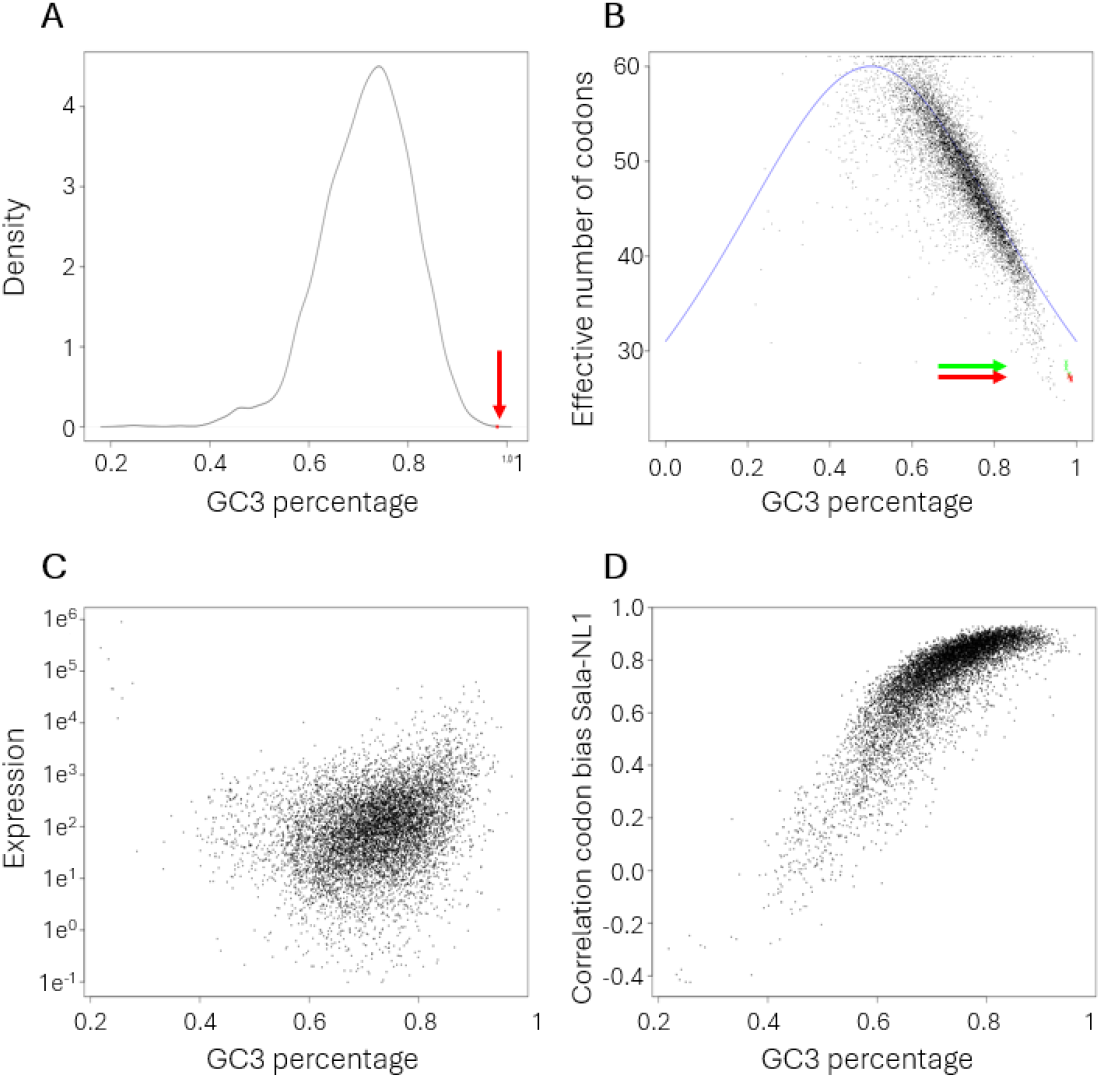
Feature related to GC3 percentages of S. latissima transcriptome ORFs. A)Density plot of GC3 percentages of S. latissima transcriptome ORFs, the red arrow and dot indicate the GC3 percentage of SalaUV-NL1; B)GC3-ENC plot of s. latissima transcriptome ORFs (black dots) with the neutrality line (blue), plus the SalaUV-NL1 SL (red arrow and dots) and Sala-NL1 SS (green arrow and dots); C) Scatterplot of the GC3 percentages and expression levels S. latissima transcriptome ORFs in sample S2-1; D) Scatterplot of the GC3 percentages and the correlation with the codon bias of SalaUV-NL1 LS of S. latissima transcriptome ORFs.

### Characterizing the sequence differences between SalaUV-NL1 variants

In this study, three distinct SalaUV-NL1 variants: 1-16, 1-27, and 2-1 were identified. Given the extreme GC3 bias, we investigated whether the RNA sequence differences between these virus variants were related to their base position in the ORF codons. In total, there were 164, 93, and 39 differences between the virus variants for segments L ORF RdRp, segment S complete, and segment S ORF Upp-SS1, respectively (Supplemental Table ST10). RNA sequence differences can have various effects, such as resulting in a protein sequence difference (missense mutation), or no protein sequence difference (silent mutation). This distinction revealed a noticeable difference between the percentages of A or T to G or C (A/T> G/C) nucleotides at the third base position that leads to a silent mutation between the SalaUV-NL1 variants (Table 3). While most other differences ranged between 0.04% and 3.5% of the involved nucleotides, the 3^rd^ base A/T>G/C change between variant 1-27 and 1-16 ranged from 41% to 80%. This was echoed by these differences between variant 2-1 and 1-16 with a range from 38 to 50% (Table 3). Hence, there seems to be a selection pressure to change the final 3^rd^ base A/T nucleotides to G/C. This is reflected by the higher number codons that are totally in segment L of variant 1-16 (22) as compared to that of variant 1-27 (14) (Supplementary Table ST6). There appeared to be one exception: of the 24 codons in the segment L of variant 1-27 that end with an A (16 are ATA), all 8 non-ATA codons changed into G/C ending codons and only two ATA codons changed to ATC codons as compared to the large segment of variant 1-16. ATA is a distinct codon in that it is the part of the only aa that is coded for by three codons. These observations shows that the codon selection pressure is likely at the tRNA level and not the protein level.

**Table 3.**
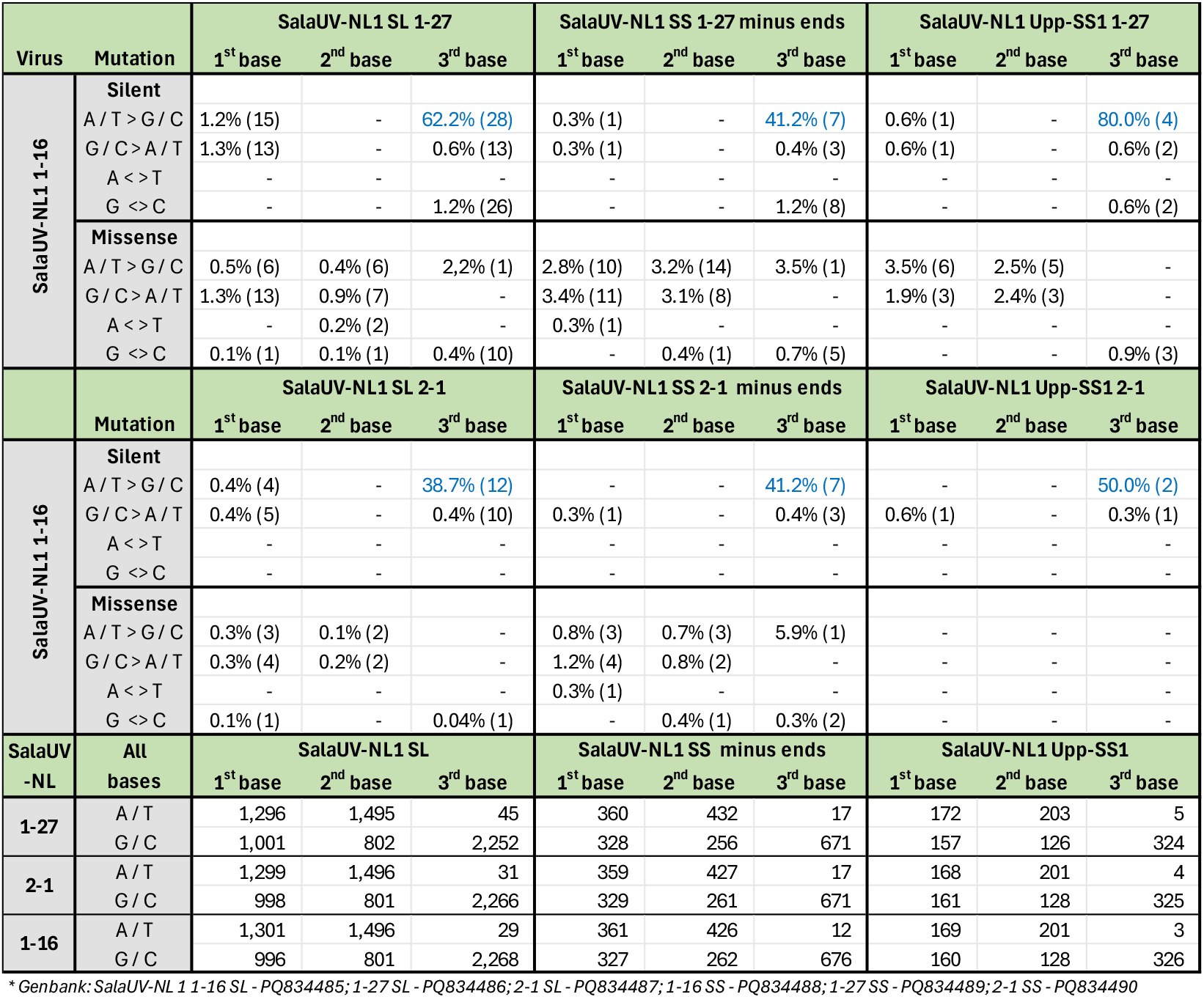
Sequence changes between SalaUV-NL1 variants related to codon position*

## Concluding remarks

In our quest to discover new RNA viruses with yet unknown RNA functionalities in seaweed, we encountered a novel multisegmented RNA virus in *S. latissima* samples. The virus, named SalaUV-NL1, consists of a large segment and a small segment and shows no substantial similarity to any known virus sequence, yet possesses remarkable genome characteristics. Initially, it appeared to be an ambigrammatic virus. These viruses avoid, next to the stop codons, also the reverse-complement codons of the stop codons in their (+)ORF, resulting in a non-stop (−)ORF that, with an early start codon, can code for a protein similar in size to that encoded by the (+)ORF (Cook *et* al. 2013, deRisi *et al*. 2019). However, this was not the case with our new virus. We observed an extreme codon bias in the (+)ORF of segment L, which codes for a large polyprotein encompassing a RdRp domain. This codon bias avoids most codons ending with an A or T base, indicating a high GC3 bias. Since all reverse complement stop codons end with an A, this means that there were no stop codons in the corresponding (−) strand, resulting in a putative (−)ORF somewhat larger than the (+)RdRp ORF. However, unlike ambigrammatic viruses, we do not believe that this (−)ORF is translated into a functional protein (Wilkinson *et al*. 2021, Retallack *et al*. 2021, Dudas *et al*. 2021), as it virtually lacks half of the possible amino acid types. Additionally, the variants of this virus we discovered in the NIOZ facility carried a stop codon drastically truncating the (−)ORF. We are convinced that the (−)ORF is a byproduct of the extreme GC3 bias in the (+)ORF and we propose to call this characteristic ‘pseudoambigrammatic’. A Genbank search revealed the existence of several other pseudoambigrammatic viruses in different host species (Table 2). Although pseudoambigrammatic viruses probably do not code for a functional (−)ORF protein, this does not exclude the possibility that the ribosome protection theory applies to them (deRisi *et al*. 2019, Wilkinson *et al*. 2021, Retallack *et al*. 2021, Dudas *et al*. 2021) or other factors (Plotkin and Kudla 2011, Cardinale *et al*. 2013).

If the observed CD3 codon bias is not likely caused by the produced (−)ORF proteins, it may be related to the availability of tRNAs in the host. This is supported by the observations that (+)ORF of almost all pseudoambigrammatic viruses use the only G ending stop codon (TAG) in their (+)ORF, and that biased GC3 codon usage often occurs in highly expressed genes. Another argument to consider the (−)ORF irrelevant is that the virus seems to mimic the codon bias of highly expressed genes. Since these genes only produce a sense mRNA strand, the antisense strand has no influence on the creation of the GC3 high codon bias. However, complicating the explanation of the pseudoambigrammatic characteristic of the new virus, the underlying high GC3 bias was observed not only in the relatively small (+)ORF at segment S but also in the segment S complete rest sequence, except for the utmost 5’ and 3’ ends which showed a conversely high AT3 bias, similar to the segment L ends. It might be that there is an unrecognized ORF in this RNA sequence, possible be created by so-called cap/ATG snatching (Ho *et al*. 2020), as the glutamic acid-rich region in the GC3 rich frame bears some resemblance that of the Latency-associated nuclear antigen (LANA) in herpesvirus (Verma and Robertson 2003). However, the high GC3 presence in both SalaUV-NL1virus segments extended well beyond the identified (+)ORFs. Thus, if no translation is exclusively the cause of the almost maximal GC3 percentage in this virus, the GC3 codon bias might occur due to selective forces that stabilize virus RNA, much like in human mRNA (Hia *et al*. 2019).

Another puzzling observation was made by comparing the variants found in the samples obtained from the two facilities. The comparison revealed a high relative percentage of A/T to G/C differences between nucleotides at the third base position, leading to silent differences between the SalaUV-NL1 SL variants. This increased the GC3 percentage from 98.0% in variant 1-27 to 98.7% in variant 1.16. In total, there were 45 A/T-ending codons in variant 1-27, compared to only 29 in variant 1-16. Although, we do not know the relationship between the samples, which originated from Scotland and the Netherlands respectively, it is remarkable that such a high percentage (62%, 28 out of 45) of 3^rd^ base A or T bases in variant 1-27 were silent C or G bases in variant 1-16. In contrast, the reciprocal difference was only 0.6% and the missense C or G bases were present only in 0.2%. This skewed rate in genome differences was highlighted by the facts that 67% (16 out of 24) of variant 1-27 codons with a 3^rd^ base A are codon ATA and all remaining A-ending codons (8 out of 8) were C or G ending in variant 1-16. ATA is a distinct codon as it shares a first and second letter codon position with the start codon ATG. It is unclear what is driving these specific genomic changes. To unravel this pseudogrammatic phenomenon, we will need to collect more variants of the SalaUV-NL1 virus and determine and compare their different genome sequences.

## Data availability

The raw sequence reads have been deposited in the NCBI Sequence Read Archive under BioProject accession numbers PRJNA1089059 and PRJNA1195823. The following virus genome sequences have been deposited in NCBI GenBank: SalaUV-NL1 1-16 SL (PQ834485); SalaUV-NL1 1-27 SL (PQ834486); SalaUV-NL1 2-1 SL (PQ834487); SalaUV-NL1 1-16 SS (PQ834488); SalaUV-NL1 1-27 SS (PQ834489); SalaUV-NL1 2-1 SS (PQ834490).

## Supplemental information

Supplemental Document SD1: sRNA contigs from sample S27

Supplemental Document SD2: NODE_1_length_15848_cov_37.681204_g0_i0

Supplemental Document SD3: SalaUV-NL1 variant sequences

Supplemental Figure SF1: Comparison of Saccharina latissima unclassified virus 1 (SalaUV-NL1) variant sequences

Supplemental Figure SF2. Codon comparison of the 5’ and 3’ ends of SalaUV-NL1

Supplemental Table ST1. Protein comparison of all pseudoambigrammatic viruses

Supplemental Table ST2. Amino acid composition of putative SalaUV-NL1 proteins

Supplemental Table ST3. Annotation of the sRNA-seq contigs with the highest co-presence compared to the SalaUV-NL1 contigs

Supplemental Table ST4. Comparison of sRNAs coverage and RNA plus protein sequence similarities of SalaUV-NL1 variants

Supplemental Table ST5. Examples of true ambigrammatic viruses

Supplemental Table ST6. Codon usage for SalaUV-NL1 segments and variants

Supplemental Table ST7. Extended summary of GenBank virus sequences with extreme CG3 codon bias

Supplemental Table ST8. Codon usage bias (CUB) for CG3 rich virus genomes that display pseudoambigramatic characteristics

Supplemental Table ST9. Amino acid distribution for RdRp proteins of CG3 rich viruses that display pseudoambigramatic characteristics

Supplemental Table ST10. Sequence changes between SalaUV-NL1 variants related to the codon position and the effect on aa coding

## Acknowledgments

This research was directly and indirectly funded by the Swammerdam Institute for Life Sciences of the University of Amsterdam.

