## Supplemental Document SD1 for "Extreme GC3 Codon Bias in a Novel Brown Seaweed Virus Results in Pseudoambigrammatic Characteristics"

### Supplemental Document SD1: sRNA contigs from sample S27

>NODE_1_length_6426_cov_34.462295

AAGTCGATGGCCATCCCGCCCATCTCCTCCTTCCCGATGGAGGAGAACATCTCAGCGTTCAGCTTGGTGG

CCATGGAGGACGAGGATATCCCGAAGTCCGTGGCGCTGATGGCGCCCATGACGTTCTCGACGCTGTAGTC

CTCCACGGCCTGGGACACGTCCATCATCTCGTATATCGTGGAGTCCACCTCCGCCTCGAACTCGGCCTCC

TTGACCCTGTTGTCCGGGGCGAACATGGAGTTGTCCGCTATCCCCAAGTCGTTCAGGATCTCCTCCACGT

CCGACAAGTCCAACTTGCTCGAGACGACGAGCTCCGAGAACTTCCTGATGTTGGGGTTCTCGGAGTCCTC

GTTGAACGGCATCTTGTAGCTGTTGGTGCCGATGTGGAAGAACACCTTCCCCCCCAACTCCCTCTCGTTG

AACATGGCCAACACGGTCCACTTCGTCATCCTCCTGGTCAGGATGAGCTCCGTCTTGGGCCAGATCCTGT

AGACCTTGATCCACAGGTTCCCGATCTTGGCATTCTTGTCCACCCCCAAGTTGTCCGTCCTGGGGGCCCA

GGGCCTGATCTGGATGGTGGAGAGGTCCTTGTGCATCAGCCTTGTGATGTTCCCCAACTCCACGTCGGAG

CTGCTGACCTTGTACATCCTCACCGAGACGGTGTGCGGGGACGCCCCCCTGTCCTCGTGGATGATCCCGA

GGTGCACCAAGTCCGTGTAGTACCTGGTCTTCATCTTCCCGTTCACGTCGACCTTGTAGAACACCTTGAG

GGACTTGGCCCCCTGGAAGGAGTACTTGTCCTCGTCGGTGGCCGAGACGGCCATCTTCGCGAACCTCACC

TCCAAGGAGTCCGACGGGATCACGGTCAGCAGGGGGTTGTCGGTGCCCCAGGGGTCCCTCTCCAAGTTCT

CCATCGGGCCCTCCGTGAGGTCCCCCCTGAACTCCCCCACCTCGAAGCGGTCCACCCTCCCCCTCATCGA

CAAGGCGCTGTTGAACCTCTCCTCCGACACCACCCTGAGGGACTTCTTCGGGACGAACATCTGCAAGATC

CCGCAGTTCGAGTACTTCTCCAAGTAGAGCATCCTCATGTTGGTGAGCGCCTCGGAGTTGTCCGGGAAGG

AGACCATCATCTCCATGTTGAGGTCAGAGGTCATCTTCAGCCAGTACATGATGTAGCTCTTCAGCATCGT

CAAGGGTCTCCTCACCCCCGGGAAGGCCTTCTTGATCTGGTATATGGGGTACTCGATGAACTTCTCCAAC

TTGACCCCCAACGACTCGGAGATCTTGTTGGCCTCCGTCATGGCCGAGTTGCTGGCGTTGGAGGTGGCCC

TCATCATGTACCTGAACACCTCCATGTCGGACACCTTCACCCTGTCCTTCATCGGGGTGAAGTTGATCCT

CCTGGACTTGTTGTGGCTGTAGGAGTGGGAGGGGATGGCGATCTCCGCCTCCTTCTCCGAGATCTCCCCC

ATCTCGCAGATCTCCTTGAAGGGGGCGGTGAGGTGGAACTTGGACTTGGTGTCGTCGTTCTTCAGGATCT

TGTCCACCACTCCCACGAAGTCCGTGGCCACCACCCTGGAGGAGGAGATCAACGCGATGTCCTCCCCCGC

CATGGCCAAGGCCCTGATCAAGATGTGGACCCTGATGGCGTCGTTGAACCCGAACTTCCTGTCCATCTGG

AACATGTAGGTGTCCATGTTGGAGAAGAAGCTGGCCGAGTCGCTGCGGTCGGAGAAGGACAAGGCGGTGT

TCCTCTCGGCCAGCTTGTCGATGTCCTCCCTCGTGATGGACCTCTTGTTCATCAGCCTCTTCTTCATCTC

CTTGATCTTCGTGTCGAACCTGAAGGGAAGGGTCATCTTGGTCTTGCCGTTCATGGAGGCCATGTAGTCC

CTGTCCTCCTCCATCTCCTCCTCGGTGTAGTCCAGGGAGGCGGCCATCTCGACGTCGGTCATGTACGCGG

CGTAGTACTTGACGTAGAACTCCATCACCTTCTCGGAGTTGCCCTCCTTGTACATCAGCACCTCCGGGCC

GGCGATCATCTGCTCCACCACCCTGTCGGTCGACATGTACCCGAGTTGGAAGGGCAGGGAGTTCCTGTCG

CAGTCCAGCTTCCTCATGATCTCCTCCTTCCTCTCCCCGTCGATGGCGTAGATGGACGAGAGCGCCACGG

ACTGCAGCTCGAACATCACCTTGATCGTGGGGATCTTGACCCCGTTCTCGAACGCGTTCCTCATCTGGGA

TATGGAGGACTTCACCGCCTTCTCCGGCTCGGTCATGTCCACGATGTTCTGGGAGTTGTAGACGGCCTTC

AGCACGCTGTCCGTGGCCCTCTTCCCGGTGGAGAAGTAGGAGTTGAACTCGGTCAAGATGGACAAGGCGG

ACTTCTTCCAGTTGGTCGTGATGTTGGCCATCCTGTTGGTCATGTCGTGCACCTGGAGCATGGACACCAT

GAAGGCCGTCAAGTCGGACATCCTCGTGTACCTGACCAGGAAGATCTTCGTGAGGTCGTCGGAGGAGATG

AGCGTGGTGCACGCGGCGTCCCCGTACGACTTGGAGAACAGGAAGCACATCACCGAGTCCATGACGTCGT

CCTTCGCGCAGTGGAACAGGGAGGAGAAGTAGTGGAGCATGCCCTGCCCCATCCCGGACTTGAATATGGC

GGTCCCCCTCGAGGAGACGAAGTTCTCCTTGAGCCAGTTTATGTTCTTGTCGTCGGTCTTGTCGTTCCAC

AGCTTGTCCGTCTGCCACCTCCTGGTCAGGGTGGGCGGGAGGAACATGGTCTTCGTGGCGAAGGCCCTGA

TCACGGTGACGCAGAACTCCTTCAACTTCACCGGCAAGGGCATCATCAAGCAGAAGTAGATGAAGCTATC

CATCACGAACCCGGGGGACCACCTGGAGGCGTCCGTGTTGAAGGACAGGATGATGCCCGAGGGGGAGGTC

TCCGACTCGTCCTTGTGCTTGTTCATCTCCAGGATCATCTTCTTGTACACGCTGGTGGTGTTGGACTGGA

GGTCCTTCTTCACGTCCGCGTGCGTAAGCATCTCCTTCTCGTGCAAGGAGCAGAGCTCGAAGAACATCTT

CTCCAAGAACTTCACGTGCAACCTCAACTTGGCCGCCTGTATCAGGATCTCCCTCGGCCCCCCAATCTGG

GACTTCGGGAAGAGCTTGAAGACGGCGTTCACCACGTCCGTGCAGGTGACCATGTCCATCAGGAAGATGG

ACTCCAGCTCCTCCACCTGCTCGCACATCGTCTCGAAGGCGGTCGCCGTGGTCAGCTTGTCCGAGTAGTC

CACCGACTCCTCGTACAGGGGGCACGACTTCAAGGACGACCTCATGGACATCGTCTCCCTTATGGTGGTG

GTCATCGCCCTCATCAGGCCCTTGTTCAAGTCGGGGGAGTGGATTATCTTCATCCTGAGCTGGTAGCCCA

TGTACATGACGAACTTCTTGTTGAAGGTGTGGGTCTGCATCTTGTTCTTCGCGAGGGCGTCGAAGTCCGC

GTCGCCCGTGGAGGCCCCGGGGAACAGCTCCGACATGGACAGGTAGTGCCTCTCCTGCTCGGACATCTTG

GTCATGATGGGGACCAACCTGTGGCCCGCCGTCCCCTTCTCCTTGTTGAAGAGGTTGCTCATGTAGATGT

CGTTCATCACCTCCGCGAACTCGACCAACATCCCGGCCACCTTGAACGAGGGCATGTATATCCTGTCGTA

GTCCGCGAACTCCGAGGCCATCTTCTCCGCGTACTGCTTTATGATATCCGGGGCGGCCATGGTCAGCCTC

TCGGCGTAGTCCATCTGCATCGCCCTGATGGAGGCCTCGAACATCGTCCTGACGGGGTCCTTCATGATGT

TCTTCACCATGCTCACCTTGTCCGAGGCGAACGCCGTGAGGCCGTGCATGATGTACCTGTTGTACTGGAG

CGAGGTGGACACCCCCCTCTTGTGGCACATCATGACCAGGATGGGGACCATCACCATGGCGGACCTCCTG

TACTCCTCCCTGGTCATCTTCTGGGCCCTGGTGCTCTCGAACTCCTCCCTGTCCGCCATCACCTGCGAGT

GGATGGCCGCGCTCCTCTCGTAGAGCATGGAGTAGTGCTCCACGTCCGCGACCGAGACGGTGAGCCACTC

CGTCTGGTACACCTCGGAGTACTCCTTCGTGTACTGGTTCCTGGCGATGGTCACGTCCATGTAGGACTTG

CAGTCCTCCTTGCTCTTCAGCTTCTGGCACTCCATCCCCTCCATGTGCTTGTAGGGGGAGAATATCTTGA

CCCTGATCTGCTTGTCCTGGGTCAGGGCGGACCCGGCCTTCACGGCCAAGCAGTAGTCCCCGAACTCCTT

GATGACCGTAAAGCGGGACACCTTCTTGCCCGAGTCCCCGTTCCTCTTGGCCCTCCTCCTCCCCTCCAGC

CTGGCGATGTTCCTGTAGAGGTCCGTCCAGAAGTGGGTCACCTCCGCCACGGCCAGCTCGTTGAACATCT

TCATCTGGGGGTCCTTGTACACCTTGGCCATGTTGTTGGTGAACTCGAACGTGGCGTCCCTGGACATCTT

CTCCACCATGTTCTCCATGAACTTCCTGTCCAAGAGGTTGGCCCCGATGCCCGCCATGTTCTCCTTCTTC

TTGGTGTTCCCCGCCTTGGTGCAGTCATCCAAGAAGATGGTGCCGTCGAAGAGGGAGCCCCCCTTCGAGT

CGGTCACGTGCATCTTGACGTCCTTCATGTCGTTCAGCTTCTGGGGGATCGGCATCGGGAAGGATTTCCC

GTAGTCTTTCTTGGCCCTGTTCATCTCCTCCGTGGGGGACAGCTCGTCCATCTTGTTCAACTCGGAGACG

ATCTCCTCCTGCGAGGTGCAGACCGCCTTCTTGATCTCCTCCGTGTACTCCGCGTTGGTGAACATGTCCT

TGAAGTCCGAGACCAGCTCCTCCCTGAAGGCCTCCCCGTCGAAGTTCTTGCAGTCCTCCTTCGCGTCGGC

CACCATCTTGTCCTCCTTGTTCTTGACGAACTCCTCGCTCCTGTAGAAGGTCTTGGCCTCCGCCCTCATC

TTCTTGGCCCTCATCTCGTCCAGGGACTCCACCTCCGAGAAGATGTCGGTGCACACCTCGATCATCCTCT

GCACCTTCTCGTCGGCCTTCTCGTCCACCTCCTCCCCCTCCCACTGGGTGTGCAAGGCCCTGTACCCCTT

GTAGGAGTTGAAGCCCCTCATCATGATGTCCGCCACCTGGATCTTGCTGGCGGCGGACTCCAGGGAGGCC

ATCACGTGGGCCGTCTTGTACTTCCCGGGGCACATGATCTCGTTGGTCTTGGTGGAGTACACCAGCACGT

CCACGGTCCCCCTGGTGCAGATGTTCTGGGCCACCATATCCTTCATCACGGGGTCGTACTTCTCCCTCTT

GGACAGGGCCCTGTCAGAGCAGATGCCGTCCGTGACGGCCACGTCGATCATGTCCAGGCAGTTCCCGGGG

GCCGAGAAGAGGATGTCGGGGGTCTTGTTCGCGAACTTGCCGTGGTAGTTGGCCTTCTTCAAGTCAACGT

CCATCTCCCCCCTCTCGCACCCCACCGAGGTCATCCCCAGGGAGTCCAGGAAGGCCTTCCCCAACAGGTC

GTGCCTGGACCTGAGGTAGACGTCCACCACCTCCACGGACATCTCCGGGCCGTCCTTCTTGTTGATGGAG

TCCACGTACTCCACCAGCTCGTCCAGGCTCTCCATGGGTTTCACCTCCTTCCTGACTTCCTTCACCTTGT

CCCTGGACATGTGCTCCTCCACCTCCTCCAGCAGCTCGCACACCTTCTCCACCTTCAACATCTTCTGCAT

CGCCCTGGCCCTCTCCTCCGTGTTCGTACCCACCCTGCCGGAGAAGGGGTACAGGATCTTGTCCCCCTTC

ATGCCAGACATCTTCTGCGCGTAGAACTCCTCCCCCGTTATCACGAACCTGGCCATGTAGGTCCCCCTGA

TAGTCGTCGGGGTGGAGAACAGCTCCAAGTACTCCTTGTAGGAGATCTCCTCGAACTCGTCCGTCTCCAT

CTCGGTCAACTCGAACGCGTTGGGCGAGCGGCTGAAGGACTGCTCGGAGTTGGGCATCTCTATAGCGGTC

TTCATGGACTCCGTCAACATCCTGACCACCTTCTCCTTCGAGTGGTTCTCCAGCATGTTGGTGTAGATGC

GGGAGGGGGTCACCTGTATGTCCCCGCCCATGGAGTTGATCACGCCCGAGAGGCCCCTGTGGAAGGTGAC

CCTGTAGGGGACCGACTCCATGACGACAGAGTAGGAGACCTTCATCAGGAACGGGTGCATCGGCCCCTGA

AGGAACTCCTTCATGTTGATGCACGGTGATCTCCACGATCGCGAAGCTGGTGTTCGACTTCTTGTTCCTG

TTCGAGATGATGATATTATGATTTAGTAGAAACTTTGTGTTGGAATTCTCGGTGCC

>NODE_2_length_2998_cov_16.467585

TGAACCTTGCACGGTAAAAATGACCTCCCTGGAGGAAAGTAACCAGTACAGCAGCGGGGCAAGAAGATAG

AAGATACGGTCGTGGGACGCCAGGATGGTTACCATGCAGATCGCCTGACACAAACGTGGGGTACTTGTGT

TAAGGGTGTTGAACTTGGCTCCAGGCTACGAGCCCGTGTTGGAGACACTATTTAGTGTGGTGCGCCGGCA

AAAGAACTGAAAGACACCAGACCCTCAACGAGTACTAGGATTGGGAAGTCCCAGTAACTTGAACATTTCC

CACTACAAATATGAACAAAAACAAGCTGGAGATAGTCTCGGAAGAGTTTGGGAGAAAGACGGTGAGGCAG

GGTGGGTGGAGTAAGAAAGATTGGAAGGGTAAAGTACTAGTATTGCTGGTATTCGGAGTTCCTAGCGACA

TGACAAGAATAGAGTTCAAGAGGAAGTGCGACGAAGTAGGACTATACTGCTTTGCATTAGGACAAGTAGA

AAGGGAGGGAGAGCATTTTCGAATCACGCTTAAACCGAAAGCGAGCATGGAGTGGCAGAGGTTAGAGACA

GAAAAAGTGTCGGCATGGGTTAGAAGACTAGGATGGAGGGTATGTACAGCTAATGGGATAGAAAATCTAA

GACGTAGTCCTAGGGTGAACGGAGGTAAGGGGAGGCAGGAAACAGTAAGTGGCCCAAAGGGAACGGGAGA

GAAACACAGCGGCGTAGAGCAAGAAGACAAAACCGGAAACGGAAACAGTAAGAAAGCTCAAAGAAGAGAA

CACAGAAGTAGCAACGAAGGAGAAAAAACGAAAGACAAGAGTCGAGCGAACAGTGAAACGAAGAAGAAGC

TGAGAATAGGATCATGGAACGTATGTGGATTCGCCACGGAAGAGCGCAAGAGGATAGAAATAGCCGAACA

GGTAAGTGAGCGCGACCTAGACATAGTGGGAATTCAAGAGTCGTGGGAAAAAGAGGGGGGGGAAATAGGA

TGCACAATTGGAGAGTACGCATGGATAGGGAAAAAGAGAAAGGGACAGAACAAGAAGAATAGGGGGGCGG

GGGGAGTAGGGTTCCTAGTCAAAGAATACCTGTGTGACATAATAGAAGTGATCAAAGACACGAAATTCGA

CGAAAGCATCTGGATAAAGATACCAGGGGAGAGGGGAGCGAAAGATTATTTCTTAGGAAACATCTATATG

CCCCCAGAGTCGAAGAGTACGGTAAAAGATATACAGAAGAAGTTCGGAGAAATAGCAGTAGATGTACAAA

AATATAAAAGACAAGGAGAAGTAGTACTAGTCGGAGATTTTAACTCAAGAATTGGGAAGGCAAGCAACCC

GAACGAGAACATTGGACAGTATGGGGAAGTAACAAAGAATAAGAACGGGGAAGAGATGCTGAAGTTTCTT

AAAAACAATGAGATGAAGACTCTGAACGGTAGAGTAAATAAGCCAGAGCCAGAATGGACTAGACAGTGCA

TACAAAAGGGAGAAAGCTCCGTTCTTGACTTTATAGTGGTAGAAAATGGAAGTAGTAAGGAAACGGAAGT

ACATGTATGCGCGGCGGACGTAGGAACTACAGATCACTGCCTAATATGGACAGACAGCCAGCAGACGAGA

GTTATCAAAAATAGGCGCGGTAGGAAATTGTACAGATGGCGAATAGATAAACTAGAAGTAAAGGAAAAAC

GGGAAGAATTCCAAGAAGAGATGACTAAAAATGCAGAGCATTTTTCCGATCTGCTGGAAAGCATAGGTAC

GACAAAGAACAATATAGAGAGAGATAGTGCAGGAGCGAGGATAATAGAAGGATGGGAGAAATTGGTCAAG

ACCACAGCAAGTAAAGTAATAGGGAAGAAGTTGATAATATGCAATAGAGCAGTAAAATGGTGGGATGAAG

AAGTAAAAGAGGCTATAAGAGTAAGGAGAAAGGCACACGCAAGATACACATCGAGCAAAACTACGTCAGG

ATGGGAGGAGTATGGTATTGCTAGAAAGAAAGTAAAACAGGTGGTTGAGAAGAAGAAAAAAGGGGTATGG

AAAGACGTAGTTAATAAAACGAATGAAGACTTTGATGGCGGGATGAAGCAGATGTGGGTAGGGATAAAAG

GGATACTAGGTAAACAAGCAGGAGAGGCAGATACGGGGATAGCTACATTAAGAGCACAGAACGGTAAAAT

GGTTAGTAGTTCGAAGGGGAAAAGGGAAGTACTTGTAGAGCATTACCGCAAGCTAGGGACGCCCACAACC

AACGAAACGTTCGACGAAGAATTCGAGAAGGATATCAACGCGTGGGCAGAAGCGAACGTAGATGCATCAG

AAAGGGAAGACAGTGGTTCAGAAGGGTTACAGGGAGAGTTCACAAGAGAAGAAGTAAAGAAGTGTGTAGC

TAAACTTAAGAATAGAAAGGCAGCAGGGGCAGACCAAATAGTTAATGAGTTTATGAAGTACGGGGGAGAA

GGAATGCTTACCATGATGGTTAAGTTGTATAACTGGATATGGAAAAACGAGTACGCACCTAAGAGATGGA

GAGAAGGAGTAGTAGTAAACTTGTTTAAGAAAGGAGACAAGGCTGACCCGGGTAACTATAGGGGGATAAC

GCTACTAAGCACTGTAGGTAAAACGTTCTGTAAGATTTTGAACGATAGAATGGGAACAATGATGGAGAAG

GAAGAAAAAATTAGCGAAGGGCAAGCAGGATTTAGGCCAAACCGTAGCTGCGTCGACCATGTGTACACAT

TAGGTAAGATAATCCAAGGCAGGAAAGACGCGGGGCTAACAACGTACTGTTTCTTTTTAGATGTACAGAA

GGCCTACGATACAGTATGGAGAAATGGGCTGTGGAAAAAGTTGTGGGAAATAGGGATAAGAGGAAAGATG

TGGAGAATGATGAAGAAGATGACGGAATGTGCGAGAAGTGCTGTGATGTTGGACGGGGAAATATCGAAGT

ACGTTGATATCCTACAAGGAGTTGCACAGGGATGTACGCTATCACCTAACGTATTCAA

>NODE_3_length_2662_cov_40.500568

ATTAGCTGATTAGGTAAGTCTGATATTAATCATTTTATTTAAAATTTTTCTATTATCTACTTTATTATTC

CTTGCAGTTAATACTTAGTAATTAACGTATGACTATTGCAATTGGACAAACAGAGCGTAGTGGGTTATTT

GACCTTGTAGATGACTGGTTAAAAAGAGATCGTTTTGTTTTTGTAGGCTGGTCAGGTTTACTTCTGTTTC

CTTGTGCTTATTTATCACTTGGGGGCTGGTTTACTGGAACAACTTTTGTTACTTCATGGTACACACATGG

GTTAGCAACTTCGTATTTAGAAGGCTGTAATTTCTTAACTGCGGCAGTTTCAACACCATCTAATAGTATG

GGACATTCCCTTTTACTTTTATGGGGACCAGAAGCTCAGGGTAATTTCACACGTTGGTTCCAAATTGGTG

GTCTGTGGGCTTTTATAGCACTACATGGATCATTTGCACTAATTGGGTTTTGCTTACGTCAGTTTGAAAT

TGCTCGTTTAGTTGGGATTCGTCCTTATAATGCGATAGCTTTTTCTGGACCGATTTCGGTTTTTGTTTCT

GTTTTCTTATTATACCCATTAGGACAAGCGAGTTGGTTTTTTGCTCCAAGTTTTGGGGTAGCGGCGATAT

TCCGTTTTTTATTATTTTTACAAGGGTTCCATAACTGGACATTAAACCCATTCCATATGATGGGTGTAGC

TGGTATCTTAGGTGGTGCATTATTATGTGCTATCCATGGTGCTACTGTAGAGAATACTTTATTTGAAGAT

GGAGATGCTGCGAATACTTTCCGTGCATTTACACCAACACAATCAGAAGAAACTTATTCTATGGTAACAG

CAAACCGTTTTTGGTCACAAATTTTTGGAGTAGCTTTCTCAAACAAGCGTTGGTTACATTTCTTTATGCT

TTTTGTACCTGTAACAGGTTTATGGACAAGCGCTATCGGGATTGTAGGACTTGCTCTTAATTTAAGAGCT

TATGATTTTGTATCACAAGAGCTTCGTGCTGCTGAAGATCCAGAATTTGAAACATTCTATACTAAAAATA

TTTTATTAAACGAAGGTATCCGTGCTTGGATGGCTGCACAAGATCAGCCACATGAAAACTTTGTATTCCC

TGAGGAGGTTCTACCACGTGGAAACGCCCTTTAATATTGTAGCAGGTAGAGACATTGAATCTACAGGTTT

TGCTTGGTGGTCTGGTAATGCTCGTCTTATTAACGTATCTGGTAAATTATTAGGTGCACATGTAGCCCAT

GCAGGTATCATGGTTTTTTGGACAGGTGCGATGACGTTATTTGAAGTTTCTCATTTCATCCCTGAAAAAC

CTTTATACGAACAAGGTTTTATTTTAATCCCTCATTTAGCAACTTTAGGTTGGGGTGTAGGCCCTGGTGG

AGAAATTGTAAGCACTTACCCATACTTTGTGGTTGGAGTTGTACATTTAGTTTCTTCAGCTGTACTAGGT

TTTGGTGGTATTTACCATTCATTAATTGGTCCAGACACACTAGAAGAATCATTCCCGTTTTTTGGTTACG

ATTGGCGTGATAAAAATAAAATGACTTCTATCTTAGGAATCCATTTAATCTTTCTTGGTTTAGGTGCTTT

ACTATTTGCTTTCCGAGCTATGCCAGGTAACCTATTTAGCTATGGGTTATATGATACTTGGGCACCAGGT

GGTGGAGATGTTAGGTTTATTGATAACCCAACTATAAACCCTTTTATTATTTTTGGATATGTATTCAAAT

CACCATTTGGTGGTGATGGTTGGATTGCTTCAATTGATAATATGGAAGATCTTGTAGGTGGCCATATATG

GGTTGGTGCGCTTTGTGTTATTGGTGGTGTATTCCATATCGTAACTAAGCCATTTGCCTGGGCACGTAGA

GCATTTGTTTGGTCTGGGGAAGCTTATTTATCTTATAGTTTAGCTGCTCTATCTATTATGGGTATAACTG

CTTCTATTTTTGTATGGTATAACAATACTGCTTACCCAAGTGAATTTTTTGGTCCTACTGGTCCAGAAGC

TTCTCAAGCCCAAGCATTTACTTTCTTAGTAAGAGATCAACGATTAGGTGCAAATATTGCTTCGGCACAA

GGACCTACTGGTTTAGGAAAATATTTAATGAGATCACCTAGTGGTGAGATCATATTTGGTGGTGAAACAA

TGCGTTTTTGGGATTTACGTGCTCCATGGGTAGAACCTTTACGTGGTCCTAATGGTTTAGATATTAACAA

AATCAGAAATGATATCCAACCTTGGCAAGAACGTCGAGCAGCGGAATATATGACTCATGCTCCATTAGGG

TCTTTAAATTCTGTTGGTGGTGTAGCTACTGAAATAAATTCTGTAAACTATGTATCACCCCGTTCTTGGT

TAACTACGTCTCACTTTTTCTTAGGTTTCTTCTTATTAGTTGGTCATTGGTGGCATTCTGGTCGTTCAAG

AGCTGCTGCTGCAGGTTTTGAAAAAGGTATAAACCGACAAACAGAGGCTGTACTTTCAATGCGTCCAATT

GATTAATCTAATTTTTATTTCTTATAATAATAATTAAAGCTAAGTATAAGGAAATTTATTAAGTTTTCTA

TACCCAACTTTAATTATTGTTCAACATTATAATTATAAATGTTAAAAAACTTTACCCATTCCTATTAATA

AA

>NODE_4_length_2463_cov_82.890254

TCCAGGATGTGTTGAGCTTATAGTCAAGTATATGTTACCATATATCCTGCTACTCCAAAACCGTACGTGA

ATGTCTCCATTCATACGGCTCCTACCGAGCAGAGATTCACATCTGTAATCTAAACAGAATGAACCTGGTC

ATGGCAGTGCCCGTGAACCCATTGCAATTGTTGCTTTGTTTCTCCCCCTTGGCATAGACGTTTTGGGACA

ACATGGTGAAGTTCCAATACATCCGTTGGGAGGAAGACTTGGCCACAAATTTTGCATCTGTGATCTTGGG

ACTTTAATAACCTATTTAATCTTTTAATTCTCGAATTACCGAGTGATAGTCTTTTGGCAAAGTATATTAA

GTGTCCGTCATATATCGAAGCGCCATGCCGGATCTTTATATGATTGCGTGTCTTAGTCTGATCATGTCTT

TTCAGTAGGATATTTAAACCGTCAGTTTTGCTAACGTATCCCAGACTCCACCGATTCGCTGTTGACCAGC

TCTTAGTCACTGCGGCCTTCCTGCTTCCTTGATGTCTCTTTACCGACCACTTCCATAGAATCTTCCACAG

GCGAAAGTCTATATAGGAGAAGGTTTTTAAACAAGTGGATATACAGAAATAGTTTGTCCAGCCTGTGATT

TTGGCACTAAGTACCGAAACCATCGACTCAAGGGGAGACACCTTATACTTCCTTACAGAAGCCTTCAGAG

CCTCTATATGGCTTTTCACAGCCTTTCGGCTTGGTGTGGTCCGCTGTATAAAGGTTTGCTTTTTCCCTTG

CGTACTCTTTACCCCTCGATGTACAGATGTTTTGTAGTTTCTAAAGTGAAAACCTAGAAAATCGAATCCT

GCTTCTGAGCCTTCATAGACAAACATAGAGTGACCTATGCGGGTCTTTTCCAAATTGAGGTTTAGTCCAA

TTGGTTTTAAGAAAGCTGAGGCGTGGTCTTTCGATTCCTTAATAGCCTCTAAAGAACTACTAAGCACCAC

AAAATCATCTGCATACCGAATCACCTTAATCTTATTCTGTGCCTTTTCCATGAGGTAGGTTTCCATACCA

TGGAGGGCTACGTTCGCAAGAAGAGGAGAAAGAATACTGCCTTGTGGCGTACCCTTAAGGTTCTCGAATT

CTTCGTGTTCTTTCAGTGGCTCTTCCAGTATTTCGGCCTTAAGCCATGCCTGGATTTGACTCGCGATCAT

GGGTGACGTTTTTAGTTTATCCAAGAGGTATACATGTGAAATATTATCAAAGCAACCCTGGATATCCGCA

TCTAGGAAATATTTGGGAGAACCCTGAATCTGACGCGTCACTGCCCATTTGGCGTCGGCGCAGCTATACC

CGGGTCGAAACCCATAGCTGTTAGGTTCAAATCTAGCTTCCCATTCGGGCTCAAGCGCTATCTTTATCAA

GTTTTGCTTGGCCCTATCCTTAATAGTAGGTACTCCTAAGGGTCTAAGGTTGCCGTTAGCTTTGGAGATC

CACACCCTTTTTACAGGGGATGCACTTCCATCGATGATCAGTTCCTTTGCTAGTCGTAGGCGTTCTGCCC

CGGTTTTAACCGTTGCTCCGTCTACTCCTGCCGTTGCCTTCCCACGATTATCTTGAGTGACTCTTCTTAC

AGAAAGAAGTCGCGCTTCCATGCACTTAATTAGAGATTTTTGCAGATTAAATATTTTAGATTCATTACCA

CTTATTTTACTTAGTTTAAAGATTTTAGACTGTAGGTGATATACTTTTTGTTCAACCTCCGACCACTGAA

TTTTACTCCAATCCTTTGGCATGTTACTTTTTAGCATACAACAATACAAGCAAAGCAGATCATTAACCCG

TACAGTTCTTTAAGCTATCTACTTCGGCTCCACACGTGGGCATATGATACACATTACATGTACCCATTCG

CTTCTTGTGGAATCGTTATGAGGGAGGGCTTGTAGCGGACCCCACCAATAAATTGGAGCTCCTCCCTCAC

TTATTCTGTTCCAATAATGTATAAAACTGAACCTGACCCTGACTCTACGCCTCTTCTCTAGAGATTCTCT

AAGGACTCCTGTCGTACGGAGACCCTCCTCTGTGTGAGAAGATTCCACCTATTCGTGGGCCTGGGTTGGC

TTACGCCTAACGTATCATTATACCCAGACTTCCTCGTTGAAGACGTTTTATTACATCAGTTCCCTTTCCG

TAGTCTTGTTCAGTACATTCTCATAAATATCTCTTTAATACGTAAAAAGATATTTACTTTCACCCTCGCT

TCCGAAATCTCTGTGATCGCGCATCAGAAATTCCGTGAGGACGCTCCCCTGATAAATGGGGATGTGCGAT

CAATATCCTTTGAGAATTGGATATGCGGGAGTGGGTACCCGCCTACATTATTAGTTAAATGGTTCTTACC

TAGAGAACACATTCGGTTTTACATAAGATATCAATTTCTTACTTTACACCAACAGCTCGCACTCGACATC

GAGGTGCCAAACG

>NODE_5_length_2388_cov_46.078158

CCCTGCGGTTTCTAGAAATGGATTCAGTAGAAGCGGGGTAAAAAGCCAACTGAGGAGACCTGTTAACTCC

GACAAGGTTTGGCAAGCTACAATTACTATGATTAGGTTTACAAAGTTGATACGAGTACTTGGCCTCACAC

TGGGCGGTGGACGTAACATATCGGGTAACGGTCCATCTTTAATGTGGGTTCCTTCCAGAATGTATGGGAG

CCCTAGGATAAACATTGAACGATTCTGCAGTATGGCAGCAAAAATAATCTCGCAACGATCTAAGGTAATT

CATGACGTGTTATTGGGAACTATGGGAAACCCTAAAAGACTAAAGTCTCATGGGTTCGGAGGTCTCGTAT

TAGGGAGCTTAGTTTATCGCTCGCGAAGGGGACCAGAAATTCGACAACTCCATTCGGAGAGGGGCATTAA

CCTTAACGGGGGCGCCCCCCTAAAGTCTGGGGCAGATATTGCAACTGGTAGGATCACCAAGGTGATCCAC

GATATCGCATCTTTAAAAAATATTACTAGAGCTTACGAGTCTATTAAAAGTAAGCCTGGGAACATGACAC

CTGGAGCTAGCCTCGAGACCTTGGATGGTATCGGACTGGCGTGGGTGGTCAAAGCTTCGAACGACTTGAA

AGCGGGAAAGTTTTCCTTCAGCAATGCTAGAAGGGTTAGGATCCCAAAACCAGGAAAGTCGGAGTCTAGA

CCACTGGGTGTCGTTAATCCCAGAGACAAAGTTATCCTGACCGCAGCGCTACAGGTCTTGGAGCCATTCT

ATGAAAATAAGTTCCTTGATTGCTCGCATGGTTTCCGGCCATCCAAGGGTTGCCATACTGCGTTAAATTT

TATACAATTTCGCTTTGGACACTCTAGTTGGGTTATTGAGGGGGATATAGCACAATGTTTTGACGATATT

GATCATAATATTTTGCTGGGAATCCTCAGACGGGACATTGATTGTGAAAAAACTATAGCACTTATAAACA

GGTCCTTAAAAAATCCGTTTGAGGATCAAGGGATAATAGTTAAACCTAAGAAAGGGACTTTCCAAGGGAG

CCCTCTCAGCCCTTTGCTTTGTAACATCTATCTTCATGAAATGGATTTATTCATTGAAGGTCTTAGCGAA

GTTTATAACTCGGGCGTAAGAAGACGGAAGTTTCCGAAATATAGGACTATACAATACGAGCTTACAAAAT

CTTTAGGTGTATCCGAAAGGAAACGCCTAAACGTTCGTCTTCGAAGCACCCCAAGTAAGGACCCGTTAGA

CCCAGATTTTCGGAGATTTAATTATGCCCGGTACGCTGATGACTTCGTTATCGGTATCTCCGGGCCTAAG

GAAGACTGTGAAGTAGTTCGAGACAAATTAAAGGCATTTTTAACGAAAACCTTAGCCCTGGAGCTAAGTA

TGGAAAAAACAATAATTTCGCACTTTAACCAAGATGGTATGAACTTTCTAGGGACCCATATTAGTGGTAA

CAAAGAGAGGGAAAAAGTTATCCGAAAAATCTCAAGAGGGGGTAAGGTCTTAAAAGTCCGTACAACCTCT

AGGGCCAGATTGGAAGCTCCTATCGAGGTTCTACTCGAAAGAAGTAGGGCAAATGGCCTCTTTAAAAGAC

TCCAGAATGGGAAGGTTGTGCCAACAGGCTTAGGAAGAGTAGTTAATCTAGATCACTCGGACATCCTGAG

GTTCTTCAATCATAAAATTCGGGGTATTTTAAACTACTATTCCTTCGCAGATAATGCGAAAAGCCTAGGA

ATTATTGTTCATGGAATGAAGCACTCTTGTGCGTTGACCCTAGCCCTTAAATTTAAACTAAGGCATAGGG

CCAAAGTATTCAAAAAGTTCGGCAAGTTCCTAGAGTGTAAAAAGTCTGGTTCTAAACTCTTCATTCCAAG

TTCATTTTCACCGACGAAGAAGTTTCACATTAACCCACCAACTTCAGAAGCAGTCTTGGACGCTAAGTGG

AATAATAAATTGACACGCAGTAACCTGAACAAGGGGTGTCTCGTTTGTGGGGAAACCCCTAGCGAAATGC

ACCATGTGAGGAAAATTAAAGACCTTAAATCGCGCTACGCACGACAAGATATCGATTTTTGGACTCTTCA

GATGGCTGCTGTAAATCGGAAGCAAATACCACTATGCAAGGTGCACCATTTGGCAGTGCATAGAGGTACC

CTCACCCAATTGGAAAAGGAATTGCTTAAGGAGTCAATTGAATCTTTGAAGTAGTCTTGTATATTTATAC

AAAATAAAGATATAAAAAGTATAAATAGTTATTTTTTAAAGGAATTTTGGAAAGCCGTATGAGGGGAAAC

TTTCACGTACGGTTTGGAGGAAAGTCAGGATATCCTAAGGCGAGAGCTACGGATACCGGGCTCATCCTAC

TAATTTTA

>NODE_6_length_2136_cov_44.837825

GGCCCCGAGAATTCCAACACAACGTTTCTACTAAATCATACTATCATCATCACCAACCGGGAGACGAAGT

GGAACACCTACATCAGCTTGGCCATCACCGTGGCCATCTTGTTGAGCTCGAAGTCGAGGAAGAAGGTGGT

GAAGGAGACGGAGGAGAGAGAGGAGGTGTCGTCGGAGGGGGTCAAGGTGGCGGGCAAGACGGGCTTGAGC

GACAAGTCGGATACCGCGAGCAAGACGTCCTCCACCACCCTCAGCAAGAGCGCGAAGAAGAGGAAGCAGA

AGGAGAAGAAGGAGGAGGAGGAGAAGGCGTTGGCCAAGGCGTTGGAGGCGACCAGGCTCAAGGAGGTGGA

GGCCGCCAACAAGGAGGAGGAGACCCACGACAGGGAGGAGGAGGAGGGCCAGAGGACCACGGCCAGGTCG

TTGAGGTCCCAGACGGTGGAGGAGGGGGTGGACCTGGGGGAGAGCTTCAGCGGGAGGACCGCGTCCGTCG

TGTGCAGCGAGGTGTTGGCGAGCAGGGTCGAGGACAACGCAAGGAAGCTGGAAGAGGTCTCCGAGAAGTT

GGACAACATCTTGTCCAAGCTTTCCTTGCTGAGCCAGGCGCAGGGGGCCTCGTTCGCCAAGGAGAACACC

TCCCAGTTGCAGGAGATCTTCGCGGCCGAGCAGGAGAAGCACAGGAAGGCGGCGCAGGAGCTGGTCAACA

GGAAGAGGGAGGTGGAGAAGAAGAGGGAGGAGGAGGCGAAGAGGGAGAGGAAAGCGGAGGAGGAGAAGAA

GAAGGAGGAGGAGAAGAGGAGGGAGAAGGAGAAGAGGGAGGAGGAGAAGAGGGAGGCGGAGAAGAGGGAG

GAGGAGGCCAGGGACGAGGAGGAGGGCTTGAAGGAGGAGAGGGGCGTGCAGGAGAGCAAGGAGGAGAAGG

AGACGGAGGCGGAGAAGAGGGCCAGGCTCCTGAAGAAGTTCAGGTCGCTGTGCTAGATGGCGACGGAGAT

GGACAAGAGGTTGTCCGACTTCCAGTTCCAGGTCGACTCCTCGGGGAGGCTGTCGGCCCCCGTCATAGCG

TCCCCGCAGACGGTGCTCCCCTCCTACAAGCCGAAGAAGGGGGACATGGACATCATCTCGGACGCGTCGG

AGTTGGGCACCCAGGCCGCCTTCTTCTGCAAGGAGATGGTCTCCAGCTTGAAGAACGCCATGGCGAACCT

CACCAGGCTGAAGCCCGAGTACTTCCCGAGCGACATGGGGATCTACACCAGCCACACCCACAGGGCGAAG

CTCTTCAACCTGATCAAGGCCGAGGGCATGGACCCCTCCACCCAGCTGGCCCTGTTCGCCATGGCCGTGA

GGATCGAGGACGTGGACAGGATGAAGATCTGGCTGAACGGGGCCGAGTGCTCCTTCGACGTGGGGATGAA

GAAGAGCCTCCAGGACTTCGTCAACATGCGCTTGGTGCAGTGGAAGAAGGAGGCCAACAACTCCCCCACC

AAGTTCTGCGTGAAGAATATCTGCTCCTCCTCCCCCCCCATGGCAGCTTTGGTGTGGAAGATGATCACGC

CGTACGAGGAGAGGACGGTGGGGAACTTCCTGGCCAACAAGTGGGCGGCGCAGTTTAGGTTCGACGAGGA

GATCGAGACCGCGCACCTCTGGTGGGAGCAGTCCTACTGGAACGTCACCATCGCCACGTCGAACAACCCG

GACCCCGACGTCTTCGAGAGGGGCTTCCAGGCCAAGTACTACAAGACCAAGCAGGGGGACAGGTACCTCT

TGCCCGTCGTCAAGGACACGGAGATCGTCTACATGAAGCCCGACGACATGGTCAGGACCAGCTACACGGG

CTCCAGCCTGGTCAAGTGGCTGAGGTCCTCCGACGTGGAGGTGGCGCTCTGCCCCACCGACGACCTCGAC

GCCCTCTTCGAGATGGTCAGCGACAGGATGGCCAAGCAGGAGGAGGAGGACCTCGGGAAGTAGGTCGAGT

TGGAATGGGTCTTGTTCACCGTAATCTACACGGACTGGAGGTGGAACATGTAATTTAGGTCCCACTAGTT

TAAGCCCAGGTAGTCCGCCCTGGCGCACAAGGTCTTCTACGGTAAGATGCAGTAGGTCTCCATGATCGAG

ATGCTGATAATATAATAAGTAGAAGACTTTGTGTAT

>NODE_7_length_2105_cov_8.790307

CGATGGAAGAAAACAAGCAGCAAGTGCAAAAGAACAAACGCAGCTCCAATACACAAGGGAACGGCAGTAA

CGAGCTGTGGTACACCATCCCGTCTCACTTGCAACCAAAGAATGCCAGCAGCTATACGGGTATGTACGAA

GCAGGGTAAACAGCAAAAGAACACACACAGACCAAGAAAGAAAATGATGAAACACGCAACAAAATACACA

TGCGCTCAAGGCCTCAGGCCAAGTCCCGATGGGAAAAAGTTGTCAACTGCTGGTACTATCAGTCTTATGA

TTCGTCCAGATCATGGACCACCTATAGGACGACAAAAAACGACCAGACTGACGACAAGACAAGACCAGGT

CAGAGACGAAGAGCTAATACCCAAAAGCCGAGCAGTCCTTTTATGTCGAGCTGGTGGGCCAGGCAGACAG

CAAGATGCCATGTACAAAGGGGACTAAAACACAAAAGAGCAAAGACAAGAAGGTCACAAAAGCAAAAGCA

AAATTAACAATAGGGTGACTGACGCAGAAGGTCCCAAAAGAAAATTCAAAATAAACAATACAACGCAGAG

TACGGGGCGGCGGCTACCATTGACTTGGCAGGGAAGAGGGCAACGCAACGAGGAGCAGGGGTTGTAAAAC

AGAAAGCAAAGGTCGCTGGGCGTCTCAAGTAGAGAGACGGCCAGTGCCACGTTTCAGCGGTTGCCTTCAA

TTTCGTAAGCGGCCAAGTTTCTGTTGCTACGCTTCCTTCAACGCGGGTCCTCTTCTTCCCTCCCCCGGGC

GTTGGGCGCAATCGCGTTTTGGGTTTCTGCGTAGCTCACACCCCAAGCGATCGCCGTTTTGCTTCATTCC

CCCTTCTCCGTAACGAACTACAGCAGCGCTTCATGTACAGCTCTGCCGCTTTACTCTCAACACGTCTCAA

CCACTTGCCCGATTTCTTCGCTGCCAGCGCCCATTGTTTCACTTCGGTGGGCAAGCTGAACAACGATAGA

GAACGTTCGAAACAACTCGTCCAGTCCTGATCTTTTGCCCACCTAAGCAACCCTTTCCTCCCTCCACTCC

CCAAACATAGCTCGTTTTGGTAGCCTCCCATTACCCACACGAGCCAAGAACCCCACGACAAGTATCCTTC

ATGTTTGCACCGTTGTCTCGACGTTCTGACAGACGGTCTTGGCGCACGCGCCTGAGTAGGATATATTATC

ATGTGCGGAGCGACGTTTCCTCTTCCATCCAACGCCGCGGAGGAGCAATCTTAACCGGTGTTGACTGGTT

TCCCGTGGTCCATAAACCTCTTGTTGTCGTAAGGCGTGCCGACCATAGATAGCCATTACCCGCCGATCTC

CTCCAATCAGCACGTCATCATGTTTATGGAAACCCCGCTGCTCCTCAAAAGTGCTACAACTAGCTGTTTC

AAGCTTTAGACGTAGCAGCTACAGGCAGTATACGTGTTAGAAGTAGCGGCGGTGTCCAGGGAGACCACGC

GGTATCATCTCCGGAATATCGCGAGCCACTTGCACCCACGTAACGTCGTCTACCACCCGATACACCATCA

GTCCAACGTGTTGGGCTCGCTCAGGCATCGCCCAAGACCACGTCTTCAAGAATCTCGTCTTCTGGTCGCG

GGCGGGCAGGTTAGCCATGACCCAGCGCATGTAATCAGTCATGTCCGAGCTCGGTGTGGCGTTCCAGCTG

TCCCGAACCAGATCCTTAGTCGTCTTCATTGGTAGATACTTGCGGGCCACAGGGGCGGAGAAGAACAGCT

CCTTGTGGTCCAAGTGGTCCATCAGCTGTTTTTGGAACTCGCCCAGGTTATATTTCAGCGCGGCCAACAG

CGTAGCGCGGCAGTCTTTGCCATCTACGCCGCCAGAACAGCTGAGGAGGGTGTCGAGCGCATTCTGAATC

TTCTGCATGAGCTGCTTGTGCTCGTCATGGTTTACGGACGCGATAATAGTCTGTCCAGGGAAGAACGCCT

CCAACGCCGGGAAGATGACCTTGTCTTGGTGCATGCAGTGGATGCGGTGCGCATGAGAGATGGCCGACCA

GGACGAAGCGAGTTGACGGAGGGCCGAAGCGTCGTCCCCCACGGGCTTGACCGCCCGTTGTGGGAGCCAA

GACAT

>NODE_8_length_2045_cov_19.686759

GGTTTAATGTGAAGAATACACGCACACACACACCATGCCAGTGACAGTAGCCGGGCGTAGGCTGTACTTT

CTGGAAGCGTCCTCATTCCGCCAAACCCTGGTATACTTGACAGGTAGTAAATGTCCGCGTCGTACCAACC

AGTCGACCCATCAGGCCTCGGCACGAAGCGCAACGTGCAGTTATCTAACATGTAAACATGCAACCATTAT

CTTACCACCACGCTAAACCCAGTCGGAGACACAACGAAGCACAAATGCAGATATATGCTGAAATGTTGGC

GCGCCCACACGCAACCTCGATACCCACGGCCTCTTGTACCAGAGTTCAAAATGATACACATTTTCGCGAC

CATATCGCCGTTGGTTGTGCGCTGCGAAGAAGCCGCGTCCCAACTTGATTGGCCACATCCGACGGATCCT

CATCGCCACAAATTCCATGTAATTTTCCAGAAGCAGATAGATCCGACACACACAGCGGAACACGCTACTC

GCAGCGAGTCGTAGATCTACGTATCATGCAATTCCTCTCTCAGGCATGGTCGAATCGCTAGAACGAGGTA

CAACGCGCATGGCACACACGCCCCCCCACTCCCCCCCCCGCGACCAAAACAACCACCACCACCGGCAAAT

GTGACTAAAAACGCGCAAAAATACAAACACACATATTTGTTTTCTTCGCCCTCGGCTTCTCCAACCCTTT

GTCGTCTTCGTTTAGAACCGAGACGTCTGTTTGATTCACTGCTTAATCGCCGTCAGCGGCAAGACCCTCC

TCGATTGGATTCCATCCTCAACTATAGCATGTTGTTTATCGCAAGAACTTGAGCGGTTTGGGACGCTTCA

TCACCGATGTGACAAAGAAATAACGTGCGAAACAGAAACACAAAATAAGTTTAGTTTGTGTTCCCTGGGC

TTCTCCATACCCGTGACATCTTGGTTCAGAACCGAGACGACTGTTCTATTTAGTACTTGGTTGCCGTCAG

CGGGAAGACCCCTTATGCCTGCTCGATCGCATTCCACCCTCAACTCTAGGATGCTGTTTATCGCGAGAGT

TTGAGCAGTTAAGGATGTGGATCGGGGCTGCCAACAGTGAATCGCCGACAGCGGAAACCCGAGCAGAGAC

AACGCGGCGAGCGACGTTCCTTCGTTTTCGTACGGTGACCAGTCAGGAAGAAATATACCGATATGGACTA

ATAAGACAGGAACACACATCAGCACCAAAACGCCTCCTCCCCGCGTCTACAAATCATTGCATACATATAT

GCATATGTATGTATTGGCGTGTTATTGTATGCACACAGCGGTGGCCTGGGACGACGACAAAGCAGGTACA

CTATCTCTAGTCGCATAGGGATGGAGAGAGCCCAGCGAATATAACAACAACAACAAAACAAATCCCCAGA

AAATCGCGGGCGCTCTCGTCTGCATTCACTACGTACATACCACGCAAAACGTAAAGGCACGTTCTGTGCA

GTAGTGCGGTGCGAATCTGGTAGCAACAAAACGAACTTCGGTCCCTAGGGCGCTCTCGTCTATCGACTAC

GGTACACAAATAGTAGCAGAACCGTGCTGCAGCAATAGGGCCGGAACCTATCAGCAACGAAAATGAACAT

AACCGACCTCATGTGCAATCTTGTCTATCCACTACGATGTCCACAGTAGGTGTACGACGATAGGAGTTAC

GTGTATACTGCCATAGCAGTCCTGTAGGTTGGGTGAGTAAAACCCCGTTGGAAACCTCCGGATTCGCACC

GAAAACGGACGATGACATGCGGAGGCTCTCAACACCTTTTCATCATCTGCCGTCGGATGCAACATGTAGG

GGGTGCCTCTGCGCACCTCCCAATATAACCTCTCGATAGAATCGCGGCAAGCCGCAAAGGAGGAGCACAG

CAAGCAAGTGTGATCGAGCTGACACTTCGACGTCTGACGTTTTGTTTTGTCAAATCAATACACGCTTGTG

ATGATGGATCCGGTTGGAACTTGGCGACGTCCTGTGGCCTCGCGGTGCAGTCCACGGTTGTAGACCTCTA

CCCGAGAGGTGGAAG

>NODE_9_length_2021_cov_16.744500

AAGGCTAGTTCGTATGAACAAACAAAATCTAGTAAATTTTTAGGCATTAAAAGTGACCCGTTAGTAAAAA

AAATGATTAATCTTTTAATGCGTGATGGTAAGCGTTCAAAAGCGGAAAATGTTTTAAATAAAGCTCTTCA

ATCTCTTGATCGTGATTATCCTGGACGGGCTATACATATTTTTTATTTAGGTATTCTTGCGGTTAGGCAA

GATATAGGTGTTCGTCTTAAACCAAAACCGACGACTCGTCGTAACAAGCAGTCTTTTAATGTATATTTAC

CACGAGTAATTTCCCCTATTTGTGGTCTTAATCTTGGCATGAGGTCATTGTTAAAAGCAAGTAGAGACCA

ATCTATAACCTCCCCTTTGTGGGAAAATTTAAGTAAAGAATTGCTTAAAGCCTCGCAGAATAAAGGAGAG

GTTGTTAATAAAAGGTATAGTTTAAATAAGTTGGCTGTGTCTAATAAACGTAGAATTCATTTTGGTGTTC

GTTATTGATATTTTAATTTTCAAAAATTCATTATGGATTTTATTCATTTTTTCTTCGTCGCTTCTTTATT

GTTTGTTATAGGTGTTGGGGGTGTTATTTTAAACAGAACAAATATTGTTGTTGTTCTTATGTCTTTAGAG

CTTGCTCTTCTTTCAGTAAGCTTAAATTTTATTGTTTTTTCTGTATGTTTAGGGGATTTAATGGGGCAAA

TCTTTGCTATTTTTATTCTTATAATTGCGGCTTGTGAATCATCCATAGGTTTAGCTATTATTTTAGTTTA

TTTTCGGGTTAGAGGTAGTATACGTATTGATCAAGCTTCATTATTGAAGTCTTAATGGGTAGGCTTCCAT

GGTGAAATTGGTAGACACGGCAGATTCAAAATCTTTTGTCTTTTGACGTGCTGGTTCGAATCCGGCTGGA

AGTAGTAAATATGATTAGAACCCAAAGTCTTCTTAAAGTTGTAGATAATTCAGGAGCCAAACTGGTTAAA

TGCATAAAAGTTAAAAAAAAAAGTGGTAAGGCACACGCCAATGTGGGGGATGTTGTCCTTGTAGCAGTTC

AGAGTCTTCGACCCCGTTATGGTAGAAGGGTTAGATTAAAAATGAATAAATCGGAAGTCCATCATGGGGT

TATTGTACAAACTCGTAGACTTTTTCGCAATAAAGCTGGCGTAGGTGTTTCGTTCGCATCCAATTCAGTA

GCTCTTATAACACCACAACATAAACCTCTTGGAACTCGAATATCAGCGATATTGCCGAGTAGATTAAGGG

GGTTGAAATGGGCAAAACTAGCTGCGATGGCTAAAAAAATTATATAAGTGTCTCTATGCATCCTTTTGAA

AAACATTATTACGAGATTGTAAAAAAAGATTTAATTCTGAGTGAAAACATCTCAACGGTTTCGTTGTTAT

CTGGTCCAAAAAAAATATCTATTTGTTTAGGTGGGGAAAGTTCCGACGAAAATTATGTTGTTGCGTGTCT

TGGGGCTTTGAATATTGTGGGAGGTCAAAAGCCTTATTTTATACAGCAAAATCCTTCTCAACAAAGAAAC

AGTGGCCCAAGAGAAGGGGTCGGGGGTAAAGTTACTTTAAGGAAAGGGGCTATGTACCTTTTTTATTATA

AATTATTATTTCATGTGTTGCCACGTGTACGCCAATTTGAAGGTCTACGAGCACCAGCTCATGATAATAT

ATATTGTTTTGTTTTAAAAGATATTTTTTCTTTTGAGGAATTGGTACCATTGTTTCCTTATTTTGAAGAG

GTTGGTTCTCTTCAATGTCAATTTCATTTTACGACAAAAGATAAAGCTGAGACTAATGTTTTAGGGCATG

GTTTACAGGTTTGCTTTTTTTCTAGTGGAAAATAGGAAAAGCGGAAGTAACTCAATTGGTAGAGTAAGCT

TGCCAAGCTTAAAGCTGAGGGTTCAAATCCCTTTTTTCGCTTTAGTTTTTAATTATACAGTTGGAATAGT

GTCGAGTTTTGTTATATTGCGGGTAATTAAAGGAGGAGTTTGAACTTCATTAATCTTTATT

>NODE_10_length_2010_cov_21.559075

ATTCCGGGCGGTTTTCACTTGGCTCCCAGTTGACCCATGCTTCTCTCTTTACACTCGCCCATGCGGCGCG

CTATTGGGGCTTTCCTGTATGCGCAGTTGGTGCTGACGCTCACATCGACGTCGACAAGCGCCCAAGACGA

TACCACCTTCTCGTGCGATGGCCTCGTAGTGGCTGGCGCGGGACCGGGAGGGTTGTACTCTGCCTGGCGC

CTCATTGAAGCTGGTTTGGTCGACCCGTCCAAGACCTGCATTTTCGAGCAGACCCAACGTATTGGTGGCA

GGATGCGGTCGGTACGGGGTGACGAGATGCCCGAAGAACTAACCGCTATTGGCGCCAAGAGCCTCGATGT

TTCTGCTTACCGCGCGTGGCCTGGCTTGCATCCGTTTGTCATCGCGGCCTGGGCGGAGCTCGGGGTGGAG

ACCGGCTGCTACCATGACATCGCTTTCCGCACGGGGAACTGTGTTGACGAGTGTGATCCTTCCCATAGAA

TCCACAAGTACCTCCGAGACAAGTACATCCCCTTCGAGGACACACTGGACTCCACCAACGACTTGCCCTA

CAACTTTGGGCCTAGCAACTCGTGGAACGTCAACTCCACCTATCTGAGTCCCTACGACTGGCTGTTTGGG

GCGTACGCCGCTCCTTTCTGGGCGGACAACTTCGAAGCCCTGGCCATGAGCGAGGTCTCGAAAGTCAGGT

GGGCCGCACGCGACGCAATCGTGGCAGCGTCCCGGGAGTATGAGATCGACGGGACCCCATCCACCATGCT

CACGCTCGAGATCCTTGCAATGAATGGCCACAACAGCGACGGCGTATACATGACCAAGGAAGACTACGAC

TTTTTCCTGGACGGTGACTGCAGCGCGGGGGAGCACGTGACGACGCGCATCAGCTTCCACGACATGGTCG

TGGAGTTCGCGAACAACTTTGCTAACGGCCTTGGCCAGGGTCTCGGAGGGTATTTCTTGGTTCCTGTGGA

TTCAGACGGGAAACAAATAGGCTTCGACATCCTCGCCATTCTCATCCGGGAAAGGCTTGAGGCCAAGGGG

GTAAAAATTTACTTCAAGCACCAGGTGTACGGCACGCGCCCGAACCCGAAGAGGGGCAAGAACGCCGTCG

TCAGGGTCCAGAATTTGGAAAGCGGCGAGAAGTTCAAGATCAACACCGAGCACGCCGTGCTAAACGTCCC

TCATGAGTTCATCCTGGAAGTGCTCGGCGACGGTGCCATCGAGAAGTACGGCCGAGATGACCTCTACGAC

GTGTTTCTGGCTTCTGAGGAGTGGAACTTCGTAAAGGCTTATCCGTACTACGAAGATGCCTGGTGGGCCA

AGCTGGGCTATGTCAGCGGGCGAATGAAGACAACACAGCACATCTATCACACCAGGTACCACGACGGGTA

CTCCAACTGCACCGACGACACGTTCACCGACTGCAGCGGTGTTCTCCTCGCTACCTACACTCAGGGAGAC

AGGGGAGAGCAGTGGATGCAGCAGTTCTTCCCGAAGAAGCCTTACGCCGAAAGTGCCGACGGTCTCACCA

CCATCCGACGTGGCCAGGGCCCCATCGAACAGGACAACTTCCTGGACGCTCTTCACGCCCAGAACATGGA

ATCACACAAGGACGCACTGGCCGCAGTCGGGATCGACACGGCGGACATTCAAGTCCCCGATATGTGCCTT

GTGGGCGTCTATATCGGCTCCGTCGCTGGAGCGCACGTTCGCCCCACCTACACCCTCGCTCCTGGAACTA

TCGACGAGGCTGTTAACGAGCCTTTCCCTGGCTACAACATCCATCTAGTCAATGAAGCTTACGGTATCCC

TGGCGGCTGGCTCCAGGCATCGCTCTACAGCTCGGAGCGCATGCTTCACCACTACTACGGCCTCGGCAAG

CCCTCTTGGTTCACTGACGACATCTACGAGGGCTTCGACTCCGACAGTTTCTACAACTCGGCCATCGGCA

ACTCCAACGGAAACGGCAAGTAAAACCGTAAGTGGAATTCCGGGTGCCAA

>NODE_11_length_1967_cov_22.061151

CACCCGGCATACCGCCAGGCATGCCGCCCCCAGCAGCCGCCATCTTTTGGATGATGGGGTTGCAGACGGC

CTCGAGCTCCTTCTGCTTCTCCTCGAACTCCTCCTTCTCGGCGGTCTGGTTGGCGTCGAGCCAGGCGATC

GTCTCGGAGATTTTGGTCTCGACAGTGGTCTTGTCGGCCGCATCCAACTGGGCGGAAACCTTCTCCTCGT

TGATGGTGTTCCGCATGGAGTAGGCGTAGTTCTCCAGGGCGTTCTTCCCCTCCACGCGCTGCTTTTGCAC

GTCGTCCTCGGCCTTGTAGCGCTCGGCCTCCTCCACCATGCGCTCGATGTCGTCGGCGCTCAGGCGTCCC

TTGTCGTTGGTGATGGTGATCTTGTTCTCCTTGCCCGTCGACTTCTCCACCGCGTGCACGTTGAGGATAC

CGTTCGCGTCGATGTCGAACGTCACCTCGATCTGCGGCTGTCCCCGGGGCATGGGCGGGATACCGTCCAG

GTTGAACTTGCCCAGGAGGTTGCAGTCGCGGGTCATCGCGCGCTCACCCTCAAATACCTGGATGAGCACG

CCGGGCTGGTTGTCGGCGTAGGTAGAGAAAGTCTGGGACTTCTTGGCGGGCACGGTGGTGTTGCGCTTGA

TGAGGGTGGTCATCACGCCTCCAGCCGTCTCGAGACCCTGACCGAGAGGGGTCACGTCGAGCAGAAGCAG

CGAGTTGAGCTTCTCCGACTTGTCCTTTCCGGACAGGATAGCCGCCTGGACGGTGGCGCCGTAAGCCACT

GCCTCGTCGGGGTTGATCGACTTGCAGGGCTCCTTGCCGTTGAAGAACTCGGACAGCATCTGCTGGACCT

TGGGGATACGGGTGGATCCGCCGACAAGAACCACCTCGTGGATTTGCCCCTTGGACATCTTGGCGTCCTT

CACCACCTTCTCCACGGGGGTGAGGCACTTCTGGAAGTAGTCCTGGTTCATGTCCTCGAACCTGGCGCGG

GTGATGGTGGAGTTGAAGTCGATGCCCTCGAAGAGCGAGTCGATCTCGATGTGAGCCTGGGTGGACGACG

AGAGGGTACGCTTCGCGCGCTCGCACGCGGTGCGCAGGCGTCGGAGCGCGCGCTGGTTCGACTTCATGTC

CTTTCGGAACTTGCGCTTGAACTCCTGCAGGAAGTAGTCGACCATCCGGTTGTCGAAGTCCTCGCCGCCC

AAGTGGGTGTCACCCGCGGTGGCCTTCACCTCGAAGATGCCCTCCTCGATGGTCAGCACCGACACATCGA

AAGTCCCGCCCCCGAGGTCGAAGATGAGCACGTTGCGCTCCTCACCCTTCTTGTCCAGGCCGTACGCGAT

AGCGGCGGCGGTGGGCTCGTTGATGATGCGCTTCACCGACATTCCCGAGATCGTTCCGGCGTCCTTGGTA

GCCTGGCGCTGCGCGTCGTTGAAGTAAGCGGGCACGGTAACGACGGCGTTGCCGATCTCCTTGCCCAAGA

AAGCCTCCGCGATCTCCTTCATCTTCACCAAGACCATGGACGAGATCTCCTCGGGCGCGAACTGCTTCTT

CTCCCCCTTGAACATCACCTCGATGGTGGGCTTGCCCCCGTCCCCCTCGACAACCTCAAAGGGCCAGTGC

TTCCTGTCGGAGGCCACGGCCGGGTCGCTGTAGCGGCGACCGATCAGACGCTTGGCGTCGAAGACGGTGT

TGGTGGCGTTCATCGCCACCTGGTTCTTGGCAGCGTCACCGATAAGACGCTCGGTCTCGGTGAAAGCGAC

GTACGAGGGCGTGGTACGGTTGCCCTGGTCGTTCGCGATGATCTCCACGCGGTCATTCTGCCACACCCCC

ACGCACGAGTACGTCGTGCCCAGGTCGATCCCCACGCTCTCTCCCTCGACGGCTGCCATGTTATCTTTCG

TGTGGTGTTTTGCGCTCTGCTTGGTTTATTCTCGTGTCGTCGCAGTTGTGTATCGCTGTGTGATGCTGTG

CTCTTGC

>NODE_12_length_1776_cov_14.884900

AAGTACCGCACCCTCAGAGAACGCGATACGATAGGTCCCGAATAACTTATCACCAATTGCCAGGTAAAAC

GGCAAGTCCCAAATTCCATAATAAATCATCGCCAAGGTAAAACGGCAACCCCAAATTCCATCCTTCATCA

TTGCCAAGGTAAAACGGCAATCCCAAATTCCATCCTCAAGAATTGCCAACGTAAAACGGCAATCCTGATA

CCAAATCTCTCAATCTCGATTCCAGTATATATCTCCTAAACTATAGCGAGACAGTAACTACGGGAAGGGG

AAAATCCAACAAACGTGAGTATGCTGGTAAAACCTTCAAACAAAATCAATAACACCGTCTTCACATAAAT

ACCGTACCGCCCGATAACATCGGCCCCCCTGTTGTATACTCTGTGAGAGCACAACAACAGAACATCTCTG

ATATAGAACACCTCAAATTAAATCATAGCAGCCCCCCTGCTTGACACTCCCCTCGGAAGTATACAAGAAG

ACAACAATCCGCATCTGAATCCAAAAGTACTATAGGTATCAGTAATTGGTTCCCCTGTACGTTGTGCGCA

GTTAACCAATAGCCAAAGCGTACGGCACGATTCATCAAGACCATCGGCATACTCTATTGGCGTATTAACC

TTATTCAACGTATGGGTACCTCCCAACACTTTCTCCATTTCCTTAAGAACTCCTGAAGTTGCACGTTAAT

TCTTCAAGTACTTTCCAAAAACATCCCATGCACATCGAAAACTCCTACTATTATCCCCCAAAACCAAATG

CCCCAAAACCCCCGGAAAAACAAACAAACGAACCGGTGGATGAAATCACAACAAGTTCATCACAGAGTCA

CGGTCAAAGCGAAACGTCTCAGCGTGGAACGGTTTGATGAGAAAGTCGGCATGCTGCTCTGCCGACCGTA

CGTGTTAATCTCAAACTACCCGTTAGACACATGCTCCAGACATCTACGTCCTATAATCAAACCATACACA

GACACCAGCGCGCATAACTGCTCCACCAGAGACAGACCTCCGGTCAGTAGTCTTGCGTGCATAGTCCGCA

TGCACGTATGATATAACCCGACTCCACTATCTGCGATGGAAAGTCTTCGTCGTGTTACCGAGTACAGCTA

CTATCTCCCCCCTGCTCGGTAGACGACAAAACATTCATCGCATGATTGATAGTAAAATGCTGCGATATCA

TTGTGACTCTAAAATCCCTCAACCATAAAATCAAAAACCAAATTCAAATCCAAACCCACACACCAGAAAC

ATCAGCGGTCAATGATAAATAAGCACCGTAGACAACGAAACAAACAGAACGTCGTTGTCGTTGTATGCGT

GCACTCACGTACAAGACTGCTCGATCCAGCGTTCAAATTCAACTATCCTCTTACCAAAACAAGCATTTAG

GCCCTCGATATACCAGGTAAAGCATCGTCTCTCAATCATACCCCAGGTCATGATAAATGGACTGCCATAA

CCGAGATACGTGACAATATACAGCTGTTGATGCTGTTATTTTATAGTATTCTGACATCGTACTTCAGAGG

TCGTAGTACGCAATTCAGAATGCCAAATATCAGATACCAGTATACCAGGTGCATATAACTCTACTCGTAT

ATCCAATGCAAACATATTACAATCCAAGCGTCATAAATATTTCGTCTCGTCAATTCCTACTGTAACAAAC

GATCCAAAAACATTTAATCTACTGTAGGCCATTGCTACTAAACTACCCCAACCCCACGCGCTCAACCCCC

GTCTCCTGCTCCCTACCCCACGGACT

>NODE_13_length_1749_cov_413.247106

TAGTTGACGGAGCTGCTTCTCTTTAGCCTATTCTCTTAGTTTAGAAAACTTGTTGTTGTTATATTCAGTA

TTGAAATACCCGTATAATGGTTTATCTTTAATATACTAGTTTAGTGTGGAGTGAATGCCTTGGAAAAAAA

TATAGGCGCGGACAATCGCGAAAGTGTTAACACGAGGAAAGCCTGTGACGTTAACCAACCTAAAATGGGA

AACCGGGGCTAAAAAGCCCGCTGTTTAGGCAGTATCAAGGGGAAGTGAAACATCTCAGTACCCTGTAGAC

CAAATCAACCGAGATATCGTTAGTAGTGGTGAGCGAAAGCGATAAAAAGGCAAAACGATATTATTATCAA

GAATGAAAAGAAATTCTATTTTTAAAGAGGATTTCTACCTTAGAAGGTGTTAGTCCTGTAATTCTTTTTT

TATTCGTGAAAGTAAGACGAGGCAAGTTTACCTTGTCTGAGTATTGGGGGACCACCCTCAAAGCCTATAA

TTATTTTTTTTACCGATAGTGAACGAGTACCGTGAGGGAAAGGTGAAAAAGAAGTTGCGACAGTGCAAAG

ATCCTGAAACTCCATATTTGAGTCGAAGCTGTAAAAAGCAACGGCATACCTTTTGTATAATGGGCAAGTG

AATCTTAATAAGGGGCAAGCTTAAGCTAATAAAAAGTGTAGGCGAAGAGAAATCGAGTCCTATGTGGCGC

TTTAAAGTCCTTTGTTAAGTACCCGAAACCGGCTGATCTAAACTTGAGCAGGCTGATATTGTAGTGGCAA

CATTCTTTGGAGGGCCGAACCCGTGTATGTGGCAAAATACTGGGATGACTTGAGTTTAGGGGTGAAAGGC

CAATCAAAGTCGGTGATAGCTGGATTTCTGCGAAATCTATTGAAGTAGAGCGTCCTAAACAGTGAATTTG

GGGTAGAGCACTGTGTAAGCAAGGGGGAGATCTTCTCTTACTAAACTTTAGCAAACTCCGAATACAAATT

TTATATTTAGGGCAGACAGAGTATGTATGCTAAGATTCATACTCAAGAGGGAAACAGCCCAGACTGTAGG

CTAAGGTCCATAAAATAGTTTCAGTGGTAAAGGTGATTTCTGCTCTGAGACCGTCAGGAGGTAGGCTTGG

AAGCAGCCATCCTTTTAAGACAGCGTAACAGCTCACTGACCTAGAGTAGAAGTCCCGCCAATTTTGAGGG

CTTAAAACTATTACCGAAGCTTCAGACAAAGATGTAAAAATTATAATTTTTTACATCTTTGTGGTAGCAG

AACATTCCGTAGGTCGTAGAAGTTTTGATGTAAGTTAAGATTAAGATATCGGAAGTGAGAATACTTGCAT

GAGTAGCATAAAACAATGAACTTAAAGCATTGTGGCCGTAAGTCCAAGGTTTACGATCTTAAGTAAAACT

GGGTCGTGAGAGGGTAATACCTCGATTTAAAACGGTCCCTAAGCTGGTAAGCTAAGGCTTACAGTAATGG

GTAAGATTAGACTGTTAAAAGTTGCTGACGAAAACTTCACCTTATTCTTAAATTTTGTTAAGGGTGAGTT

ATTTTTTCCAAGAAAAAGGCACTTGATTTATACCGTACCTTAAACCGCCTCAGGTGGACGGGTGGAGAAC

ACTAAGGCCTTGAGATAACCCTGTTGAAGGAACTCGGCAAAATGACTCTGTAACTTCGGAATAAGGAGTA

AAGACATATAGCGCAAGCTTTTGTCTTTGTCACAAAATCGGGGGTGGCGACTGTTTATTAAAAACACAG

>NODE_14_length_1719_cov_19.644876

TTGAGTATTAATAGGGTTTTTTTTATTTTGTTTTTTTTTTGTTATTTTACCAGGTGTACACTGGAATGAT

ATGCCAATATTATGACAGTACTCGTACGTACTATATCGTCCCGTCAATGTCGTACCCTACTGAGCACATT

TATCTTATATGATAGAATTTACGACACCTCCGCAGTAGGGGGTTGGCGCTCCTACTGAGTATGCCGATGT

GATCTTGGACAGTCGACCAAGGGCATGCTTCGACCTATCCACTTGAAGGAGGCCCCAATGCAACCATTGC

GCCTTGACCGACCAATGCGCAATCCGATGAACAGCGTGGAAACAATAACCCCCCACCCAAGCTGTTTGGC

TCGCCCCTATCCATCGTACACCTATTATCTCAAAAACGCTCGAGTTGTCACGTCTGGTTACACCATTGCA

ATCCTCGTGTAGACAGTCCTTCCTGGACACAGGGTAACTCACCCTTCCTTCAACATTTGATGTCGTGGAG

CGTGTACTGGACATGAGAGTGCGAAGCGCAACCCTATGCTTGTCTAGACAACGGGAAACGAGCAGTCTTT

TCCGGAAACCGTCAACAGTCAGGAAGAACATCGTTTACCGAACTGCATATTTGTACGTGGGCTTTCGGAA

TTTACACACGTGCGGCAATGAATTCACTGATCGTGTGTTTAGCTGAGCCTGACGTCGTCAGTACGACATA

GGTATCACCCTTCACAAATAATTGCAGTATGTTTTCGCGTGCCCCTTTCTGATTTTACTGAGCTCGTCAT

CATGCGCCTCTCGTCCTCTCGATAAGAGAAAATCGCTGGGGAACATATATAACTCAGCCACCCGTTAGTG

TTTCTACCGGGTCCCCCAAATCTCCCGTACGCAATGAAAACAAAAATAACACATATTTGGGACCTTCGAA

CAGGCCACGCGACACCAATCGAAACACGATTCCCAACCATTTTACGTATCCCATATCAATACGCTGCGGC

AGAACCTCAACACTGGTTCACGCATCTCGAACGGTTACCTACTAAATACACACATATGATACCCCGGACG

CCCGCCACAAAATATACACATATGGTGCACCAGACGCCCGCCACGGCAGGAGGCGAACCCTTGGAAGGTA

TGAAGTTCTGCATCTGTGGAGCTTCATCGCTGAACTCAAACCACAGGTTTGACAACCATTGTACGCTGGA

CTCAATGAACAGTCCTCGTCGTATGCGTAGGAGCGTCGCGACCAACATCACAGACGACAAAGGCGATACT

AAAAGCATGACTGTGATGATAAATCCCGTCGGTGGGATGTTCATGGAGCCAAACGTACCCCGACTATCGA

CGCAGGGCTCGTCTCATCGTGTAGCGGATATCACGTGGAACAAACAGTTCACGAGTCGAGCACGATTTGA

GAAGGGGGCAGGGGACTCTGCGCGGGTGTCTTGGGCAGTTCAAGTCCTTCAAAGCTCGGTGTGGATCGAG

AAGGTGGCGTGTTTCTGGACAATGTCTTCATCGGAGTCCGACGCGCTCTCCGGTACCTCGTTGCAAAAGT

CGTCGCCGGTGTAGGCCCCCGTGCAGAGCTTGCCATCCAACTTGTACAGGTTCCTGCCGTACAGCTCGAG

CGTCTTGCCGCCGTAAAGGCGGAAGCTGTACGTGGCCGGGGGGGTGGCCTTCTTCCCTGCGTCCTCCACA

ACGCGGGCCTCCGAGAGTCCGGGCAGTTCCTGCTGCAAG

>NODE_15_length_1620_cov_11.107567

CACCACACCGCAAGCCCCACCGGACTGGCTGAGCTTCTCACTGTGAGCTCTCTCCTCCCACAACAGTACC

TCGTTTATCGTCCGACGCATCTATCTGCTACTGCCGGGTCGGGCTTACCGCGGTTCTGGCAATTGCTGAC

TAGCTATCGACCTCCTGAGCGAGACCATCATAATAGCCGTCATTGCCCGACCGATAGCGCTTCGACTATG

GAACGCCGCGGATCCTTCGCATCTAGCGCTACGCTCCTCTTGCTGTTCTCAAGCGTCTGCCACGGACTGG

TCGAAACGTGCGAGCAGCTGCAGGCAGCATTCGACTTGACCAAGACCCAAGATGTCGTCGTCGAAATGCA

CCCCAGCCAACGTATCAACTGCGTAACTGCCACCACGATGAGCATGGACTCCAACACGCTCACCGTGGAG

TCTAGTGAGGACCTCGAAAGATTCGGCGGAAACTCAGATCTGTTCCAGGTTCGGTTTGAGGTGACGAACG

GCGCACAGTTGATCTGGGAAACGAATGTAGAGTTCCACGGGCCGACGGGGCAGGACGTTAACGGAGGAGC

CATGTTCGTAGGAGAGGGCTCGAGCGTCCGCTTCTTGAACGACCTATTGATGGACGAGGTCGGCATCTCT

AGCGTGCCCGAGGAGAACTCCGATTTTGCCCAATACGTTCTAAACGGAGGTTGTGTCTACAACGACGGCA

ACTTCAGGGTGGATGGCGACGCAACGTTCACCAACTGCGATAATTCGGGTGGCGGGGAGAGCCCACCGGG

CGATGGCGGTGTGCTGTACGTCGGCGAACAGGGTTCCGCGCAGTTTAACGGGGCGTTGCAGATGAAGTAC

ACTTCCATTACCGACGACGGCGGGGGCAACGGAGGAGGCATCTACAACGCGGGCAAGGTCGACATCAAAG

GAAACGCCAAGTTCGAATCTCTCCGGGCTGGGGACGGTGGAGCCCTCTACAACGCGGGCGGTGCCCAGTT

CAGGTTCACGGATCAGGCGACCGCCCTGTTCATAGACTGCACGTTGCGCGACGGCATCGGTAGCGCATTG

TACAACGCAGGGAGCTTCAAGTTTTCTGGTCCCGCTCTTTTCGTCGACCAGGAAACGCCAACCATCTACG

TTTCTTCTACCGGCGAGACTGCGCTTTCTGAGAACTCGGTGTTCTGGGGTGACGGACCCCGCGACGAGGT

CATTGGTGTTGCCTCAGGGGGTGTGATCGATATCCCTAGCTCTGTTTTGTTCGTCTAGACGACGCTGAAT

CTGACTGCAGCACGGTATACTATGTGGAAGCCGAAACGTGCCTAGAATGAGGTGTAGCGTCGTGAAATTG

CAGATTGAGGTCCTGCTGCCCGTGTGTTACATGCTTGGCAGGTTTCTTTCGTCGATAAACCACAAAGTCG

ACTCTGCGCGGTGGGCCTGGTTTGCCCGTATTCCTGTGTGTGGGGTCGGCATATGTTTGTTTCGTTCTGC

CCATGGTGCGATGGATTTCGTTGTGAGGGGGGGCTCGACTCGAGGGTGCTGCAAGGAGTGCTCTGAAATC

GATATGGCGGTGGTAGTGCTACCGGGCGTGGGTAGTGACGTTCTCGAGAATCATTGCACACGGGGGTTCA

GTTGACCGTG

>NODE_16_length_1531_cov_10.172848

ACCTAGCGCTACTTGGCCTACTTTTTCATTGGCTAACCCTTGGGCACGACAGCCGCATGGGGCAGCACAT

TGGCAGAAGCGAATCCTAACACTATAATATAATACCAGGTACGGTGGTTGTTATACTTACTGTTGTTGTA

GTATTGATAGTAGTCCTCCCTACAATTCTACCAATGTAAATGTGTGGTGTAGTTTGTACCCAGATATCTG

TGAATGCCAAGCACGAAACATATATGCCTACGAACCGCCCCCCCTGTATGTTTGGCCTTGTTGGAGAAAA

AAAAACAATGGGATGATACTAAAAACTCTGGAGTCAACGCTACGATTCATATGAATAACAGCAGTCAGGG

TAACCACAGCTGTTATATTAATGGACGCAATCGCATCTAATACATATTAATAGACACCAATACCAACCAT

GATCTAAACCAACGAACTATTTCAGAACACGACCAAAATCGGAAACAAATATAAAATCCCGAAACGCAAT

AATGTCATGTCACGCGCACAGATTAGTTGACCCGTCGTACAAAAACAAAGGCTGTTTCTTATATATATAT

GTGTTCCTTATCTTGTTGTATTCCACTGCAGTTGTTGTACTTTTTAGCTTTCATAGCGTATGCCTCGTCC

TAACAATGGCCTGCGCGTGTCCCAACAAAAGGGTTTGCAGTACTAAAGGGGTGATAAGCCAAAAATAAAC

AAACAATCTCACCACGGGTGGGACCGACCCGTAGGATCAACTCTATTGATAGTCCAATACTCGAGGGTAG

CTACAATTGACCACCGGCCCATTTGGATCAAAAGAACTTATCATACTGATAGGAGTCGTAGTGTGATATC

ATGAACCGTCTATACTTTATTATTGATAGTGGTGGGTCCAGTCTATTATATAATTCATGTCGTGCGCGCC

GTACGATGGATATACATATAAATAGACCGCTGTGTTCTGCTTTGTTGTAGTTGTAGTAGTATTATGCGAT

ATCTCATTCCTACTATGATGACTCTCCCCCCCCCAAAAAAACAGATGCTCTCTGCCAACAAAAACAATAC

TGGTGATAATACAGGTGCCTTCACTTAAATTGTAGTAGACTATAAGCATTCATCTTGTATTCATCCTTTC

TCGCACAGTCTTGGTTCCGTTATATGGCTACCTATAATACGGTAATAATCGTGACAAGCGGCTTGGGCCT

ACTCCTGTTGTAGTTCTCGTAAAATCCACCTATTCATAGCTCTCATAGTTAGCTCTCGGCATCCATGATA

AGACACAAGCTCTGGGGCCATAAAACTACAACATAGAGACAATACCTTGTAATGGAGAGTGCGGGCGTGT

AGTTCAGTTTTGGACACGCATAACATGCTGTAGGGATTTTAGGTGTACCTCTTCGGACTTGTGTATCCTT

TCCCTGCTAAGTGTGCGTGTTCTTTAGGATTCTCTGATGGCTCTAGCACTTTTGTGACCACGGGCTGTGA

TCTTGCTAGATCAGACTATAGGGATTAGCAACTGAAACGAGTTTAATGTGAAGACAACAAC

>NODE_17_length_1511_cov_18.026174

TTCAAGGTATATATTAATGACATGATAGTAGCAGTAGAAGCAGCATCGCAGGGGGTCACGGTGGGGGAAG

ATACGGTGTCGGGATTGATGTTTGCGGATGATTTCGTCTGCATATCAGAAACACCCGAAGGACTGCAGAC

ACAAATAGAGAAGGCACTAGAGTACACCAGGAAGTGGAGAGTGACGGCGAACGTAAAAAAGTGCGCAGTA

GTTGTATGTAACGAAGATAAGGAGAGCCCGGTAACTTGTAAGTGGAAGTGGGGAGAAGATGAATTACCGA

TCGTAGACCAGTATACGTACCTTGGCGTAGAGATCTCAAAAGATTGCTCTTGGGATGCACACATAAATAA

AGTAATAGGCAAGGGTAAGGCACACGTAGGCAAGATGGAAGCAGTCCTGACAGACTCGCACCTTGACACT

AGAATTAAGAGATGTATACTGATGAATGTGATTGTACCAAAGCTAGAATATGCAGGAGAAGTATGGGAAG

GGAATGCGAAGCTCGTAAAACAGCTGGAAACAGTACAGATGACAGGAGCTAAAAAAGTACTAGGATGCTC

AAGTACGACGAGTAATACAGTTTTGAGAGCAGAACTGGGAATGTACCCACTTAAGACAAATAGAGACGTG

CGTAAGTTGAAATGGCAATATAAAGTAAAGAACATGCCAAATAAGAGGTTGCCAGCCATAGCTGATAAAG

CTGTATGGGAGAAAGTAACCAAAGGGCGAGCTGGAATAAGGTGGGATAGCGTAGTCGAGAAAGTATGGAA

GGATATAGGAAACCAAAAAGAGATGCTAGCCATAGAGAAATTTAGCGGATACAAGACAGAAGTAAAAGAA

AGGCTAGAAAGAAGGGAAAGGCTAGCGCTAAGAAAGAAGGTGGAAGGGGAAGAACACTTAGAGATATACG

GGGGGTTGAAAGAAGGAATAGGAATGAAAAGGTATCTGCACGGCCCAATGGACTTCGCGAAAAAGCTGAA

ACTGCGATTTCGTGTAGGGGACCTGGACCTGCCAGAAAGAAGAAGGAGGTATACCAGTAGCCGGGAGGAG

GAGGAAGTAGATGCACAGATGTGCCCATGTGGCAAAGCCATAGAGAGTAGAATTCACATAGTGGGAGAAT

GTGAAAGGTACAAGGAGGAAAGGGATGTGTTAGAGGAGGAGATGAGAAAAATAGACGAGTGTGACATGGA

GCAGTTTGGTACACTAAATAGTAGCGAGAAAACGATCGCTATCTTAGGAGATAGATGGTGGCCACAGGCG

GCGAAACAGGAAGGGGATAAGATTAGCAAAAAGTTTCTTTGTAATATATGGAAACAACGTAATGAGCGCC

CAAGTGTTGGAGGTGTCTCTATTAGGAGTAGGAACGGTGCTCCGTCTCGAAAGGGATGCGTGGTCAATGG

TCAAATGACTAAGGCAAGCAACGAATGAGCACTTCCCCCGTCCCTGGAAGTGGCCACATCGGAAAAAAAA

GTCAATGAATTCGACGCTGTCATGGAATTCTCGGGTTGCCA

>NODE_18_length_1510_cov_17.672935

GGCGCGTCAATTATGCCTACTTTGGGATACCTTAACGGAATGATTTATTCAATGAGACAAGCTACAATAT

ACACGTATGACGACTTAATTTACTCCAACATCAAGGCGGCAGGGCGAATGCTCTGCTTGCCAAGTCGCTT

TCCCGGCTACAGTGAGTGGACGTACACATAGAGTCCCCCCTTACCCACAAAAAAACAAAAACTCTCCCCT

AACGGGAAGGACAAACAGTGTACTGAGACCTTGGAAAGAGGATTTTCCATCCAAGGCGTGTCCTTCCACC

TACGACGATACGTCAAGCACTCACGTCAACAGTCTATCTTGGATTTCTCGGGTGTCAGCACCGTGGACGA

CGATTTTGGCGGCATTTCTAGTAACGTACGTACCGTGTAGGCGTTATAAACCTCAACGAAACCAACGCGT

GTCCACACGCACTCAACTCCAACCGAAATACGAAATGAGGTCCAATCGGTTCGCACACGTTTCCTAGGAA

GAAAGCGATCAAGATCGCTAGCTGACGAAGTCAATAGGATAAAGCCCGCTACCGCTCCACGGTGTGTAAC

TCCTTGTTGATGATATCACACCAATTCACCCGTACAGGCGTGTCTGAGTCATGAGCGACAACTGCAAACA

CTACGACCCGGTCCAATTCCAACGTTCCACCAAAGCCCTTTCCGTCGTTGCAAGAGGAGTATACAGAGCC

TTAGCCTGGCTTCTTAATCACTCTGGATACTGTCGGCCCTACGAGTGGACCCTCGAAGGCTTCGAGATGA

ACACCTCGGCGACAGGGTTCCTCCAGTCGTAGGCATCAGCCGCAAGTTCTCCGCCGATTCCGTGGAAGCC

ACGGTGCACCTGGACGAACCCCTCCCCCGATTCAGTCGGAAGGTTGCCGGATGTGGCGTCGCACGCTGGG

CCCGGGATGTGAGAGCAGAGCTCGTTGTTCTCCTCGGTGCCGGCGTCGTAGGCAGGGGGGTAGACCGTTG

TCGATCGCTTCTTGTCCAAAGGCATAGTGTCTATGCCCACAAAAGCATCATTGGTGTTGACGAGCATACT

GGCCATGGAGAAATAAATGTCGCCTTTGCAGTGTTTCTCGTCGACGAGCAAGGTGAACCACTCCTCTCCC

GCGGGGCCCAAGGCGTCTTCGCCCAGCGTGACGCCGCACACGAAGGGTGCCACGTCTTCGTCGTCTGCTC

GTGCCAGGAGTTCAGAGGCGTCACCGTCTTCAGCTAGTGGCCTGAACTCTGGGACAGCCTCTCGGCCCAC

TTGGAACGCTTGTATGTCCTTGGTGTGCGTGAACACCATCATGGGAGAGAAGGGCTGGAGGTAGGTCAAG

TTCCTCACGCGGATCTTGTACTCCAAGGTTGCCTCGGAGAAGATGGGAGTAGAGGCAGTCAGCAGCAGCG

CGGTCGAAGCGGAGACGATGCGAACCATGATGATGATGCTGGTGATGAGGAATAGGTACTGGACGCTGCG

AAACTTATTCAACAGAAGAAGTATTGACAGGTTGACGGAA

>NODE_19_length_1456_cov_21.039721

GGCTTGCGAGATGTTCATCCTGGAGCTCACTCTCAGGTCTTGGTGCTACTCGGAGAAGAACAAGCGGCGA

ACGCTTCAAAAGGAGGACATTCAGGCGGCCATCCGCAAGACGGAGATCTTCGACTTTCTCGTCGACGTTA

TCCAATAAATAGAACAAGAAGCAACAAACAGAAGCTGCAGTTTTTTTTGTTTTGTTTTCACTTTTATTTC

TCGAAGTGTTGACTCTCGGCTCCGACTCTCGGAGGCCGTCCGACTCTCGGAGACCGTCCGTTCGCTCGTT

AACCGATGAGTTTTTTGTTTTGAATTTTGTTTTGAATTTTGTTTTTGAGAGTACACATGGTTTTTCGCGA

TTGAAACACTTTCTCGCACGCATCTTGCGTGTTCGAGTTGTCGCTTGGCGCGATTATTGTCCTCCCGTTT

TGGTGGGTGTGCGCGAGCGGCGCGCGCAAAGCGCCTTTTTATCACCGCCCGCCATAAGGTTCAATTTCTA

CGTGAAATAGAAACATGTTCAGGTATTGCCAAAGTGGTGGGATGTACAACAAGGGTATTACTAGCATGTG

CCCCCGTCCACTTCTCGCCTGGTATCATAGGTATGACTCTTTGATTCTGCTACGATTGACTCGCTCTATT

GTGGCGGCGGCAACGCTCCACGATGGTGGAATTGGCTCGGGTGGCGGGGTCGTATTGCTTTATTGATTGG

ACGCTGCGCGAGGGTGTGGATTGGGTTGGGTGGCAGTGGCATATGTGATTATTGATTGGTTGCTGCGCGC

GGGAGGACTTTGATTGAGTGGCAGTGTCACGTTTTATTATTGATTGGTTGCCTGACGGGACGGAGTGACA

GCCGGAAAATAAAAGCCCTGTAGGTTGTGTTTTATATACCGGGTACCCACCTGGTGAGGGCCTTGTTTGT

CTGATCTCTTTTCTGTATTTTGGTGTTGGAGTCTTTCTCGCTGAGCTGCTCTCTGTTATGTTATACCTGG

TACCAGGTATATACGAGCAGGTCTTGCGACTGCTCTCGGGTGTGTTGCGCCTTTGGGTTGGAATTTTGGC

TTGATTGCACTCTCTTGCGTTGGGCCTTGCCTGACTGCTCTCTCTCTCTCGTTTGAGGCAATCGGGATAG

ATGAGGACCAAACGTGAACACCATGCTTGCCTGCCAAGAATGGGCGACCCAAAAACGCGCTCAGCAATTG

AATCGCTTGCATCCGGGGCAAGGCTGTGTAGAAGGAAGGTGTGTTTCGACTCGGCCGGCCTCGGAGGCCG

CAAGATTTCGATAAAACAAACCGTGTAGACTTGAATGATGATGCAGTATGCGGGGTACCGCTAATAATGC

GTTCTGCGTTCTGCAATATAAGCAATAACCGGTAAGACAAGAGTAAAACACAACGGCAACACTGCCCAGT

ACACGACGGACTCCAGCCGGTAACCACACACCACGCCGACACGCCTAATTTCACCC

>NODE_20_length_1446_cov_35.134737

CACCCACCCACCCAGCCAAGCTAAGCCTTAGGCTAACCTAACGTACCCCGCGACTGCGAACCGAACCTTT

TTTCCGTCATGACAAGCTATGGAGAAGCGCGCCTGATGGCCGCGCCTCGCGCAAACCTCGTGGATGACCT

CACCACGACGGCCGGCCTGATCCTGTGCTTCGGCCTTCTCGCGTTCATTCCCGGAAACGGCGGCCTGATG

AATTCCATTATGGACGGCGACGTCGACTGGAACGGCTCCGACACCGTCTTCCCCCCCATTGTAAGCCTGC

TGGGGGATATCACGGTCACGCTGTTCGGCATGTCCTCGGTGTTTATAGGCTTCCAGTATCTGGCATGCAG

ATGGGGTTCAAAGACGTCCTCGCTATTTGGCCTCGCGATCACCCTCGCGGCGTGGTTCCCCTTCTTGGTC

AACATCGCCAGGATCTCTTTCCTGGCACACAATAAGATCGTCAGCGGGCCTCCCCTCGTCGTCGTGTTCC

CTGCTTCGAAGAACGATATTGTGGCGATTGCTATCATGGGCATCATCGGCTTCTACGGCTACGCAGGCAC

TGTTATCGGCGCCCTGACGTTCCTTCACTGGAAGATGTACCGCTTCTTCGCTGGCTACGCCCACACCTAC

ACCGCCGCCTACTACCGCAGCCGCCAGAACTACTACGGAGGCCTCACCGTCATTGTTGGCGTCGTACAAC

TTGCTCTCGGCTTGTTTGTGCAGGACAAGTTCGGAGACGGCCAGCTGATGGGACCCGTAGCCAGTGGCCC

CTTCTTTATCCTCTGGCCCGAGCTTAGCATCACCGTGGGTGTCTGCCAGATCCTCTTCGGTGCCCTCGTC

ATCTACCGCTCGCACACCGCCGGAGTCGGACCTTCCACCGGCAACAAGGACCATGGGTTGTTCATGTTCC

AGTGCGTAGCTTGGCTTGTGTTCCTGATCAGCTACGGGCTCCAAATGATCGGCCAGGTCTCGCTCCACTC

CGACAACATGCCGTCGGCGTTCCTCGTAAATATCATCGGGTTCGCCATCTTCCCTATCTACCTTGACGTG

ATGGCGCACACCACGCCAGAGAACGTAACCAGCGACATGTTCAGCATGACCAAGACTCTCTCATCCGGCG

TCTAATTGTAATACGCCCTGCGAAAAAACTTTTTTTTTCGAGCTGGAGACAAAAGGCGAGGGGAACGGGG

GGGGCAAGGAAGCAAGAAGGCGGACTGAATGACGTGAAGTATTTTTTTTAACGAATATAGTAACTCTTCT

ATCTGCGACGTTTTGCGTGCGCCGCGCCGCCTCGCCTCGCTCCTTGTGAAAAAAAAAAAAACGTTTTTTT

CGGTGACTGATTTATATGAAATATAAGCAGTGGGCATCGATGAAGCGCGCGGGATAACTCATCTGTGTAC

CTACTACAGGGCATCTGTGTTTTCTAGTACACGCAGGCGGTGGGCG

>NODE_21_length_1445_cov_18.724719

CCCGTTCTTAAAAAAGATGATTTAAGTCAAGTTGAAATAAGCTATAAGGCTTTAAAAGATATGGCTGATT

TGATAGAGTACAGCCGTGTAGAGGTAGCTAAAGGTGAGAAACAAACTCAAATCGCTATAATTAAAGAGTT

TTTGGGATCATTTGAAAAAGATATGGAAGATGTTCCACAAGGCGCTGCTGATATGTTATCTCGGCGCTTA

GAATGGCTGGTTATAAATATTTTAGGGAACGAGGGTCGCAAAGAATTTTACGGTCCGTTAGAATCACATT

TAACAGGGGAAATGGTATCTATTTTATCTGGAGTTAAAGAACAGTTATCAAAGGGTTCACTAATTAAAGA

ACAGCAAGATTTAGTTATTAAAGCTTTAAAAATTTTTAAACAATATATAAGGATTGTTAATGAAACTGAG

CTTACGGCAAAAACAATGGATTTATTCAAGTCTTTAGATGAAATTAAACCAGGTAAAGGTAATACACCTT

CAAGTGATGATAGTTCTGGAGGTTTAGATTCTAATATAGGGTCTATTGTTGAGGAATTAAATAAAGTGGA

TGAGGCTAGTTATAAATGGTTAGAACTTAGGTCAGCTGAAAATTTTTATGAAGGGGCTGAGACGGAATTA

AGTAGAGCTTTAACACAAGTTTTTGAGGCAGAGTGTGTGTGTCTTAGAGCCCTTGCCCGTGGTGAGATAA

AAATACCTATAGAGGAAATGATGGCTAATTTAGACGAAGGCAGAATAAGACTTGACAAAGCCTTGGTAGA

AGCAGAAATTGCGGAAAATAATTTAATCGATACACAGTTAATTGCCCATCAATTAAGATTAACTAGACCG

GAAAGAGAGGGTTATAGTAGTGAGTTTGAGTGGGATACTGCTAATGTTGAGTTTAAGTTTTCTGAGGCGG

AATTAGAAAATGCGAAACTGGAGCTAAATAATGCTAAAGCTAGTTCCAAATGGGCGAAAATTGACTTGCT

TGCCTCACAGGTTAACGCGTTAACTGCAGAGTATTTTCAAGCAGATGCTGAATGGGCTTCAAAAGATCCA

TCTATAATTCGGAGTAAAGTATTACTTAAAGATGGTTATGATGGTTGGAATGATAAATTAAAATCAGATT

CAAAGAAAATTTTGGATAGTCACGAAATTAAGTTTATGCTTGCTCACGAAGAGTTGAAAAGTGCTAATAC

AATTTTAGATAAATTTCAATCTGGATTGACTGCAGAAGAATTGAGTAAGTGTCACTCCATTGCGGATAGT

GCAGAGGTTGAATTTATGTCTCTTGAAAAACAAGAAATATCAGTTGCAGAGGAATTTGAAAGAGGTAAAG

ATATTTTAGCAAATGCCAAAGAGGAACTAAGCAATTATAAAGCAGTGTCAAATAGAGCTGATCTTCGGTT

AGAACGGTCTCAAATTGCGTTTGACAATTATAAAGCAGTGTCAAA

>NODE_22_length_1443_cov_10.796062

AAGCTTGAATTAAAGCCACTGCAAGCTCAAGCCCAAAAAGAATTACTAAGACTGCTAATGGTACGATATG

GGCAACTAATAACAAACCACCACTACCCATCATAGCCCATGCAAAGCCGGCAATTACTTTTAGTAATGTG

TGACCTGCCATCATATTGGCAAAAAGACGAACGGCCAAACTAAGAGGTTTAAATACGTACGAAACTATTT

CTATAGGGACTAATAAAAAAGCTAAAACCATAGAAGTTCCACCAGGTAAAAATAAACTTAACATGTGAAA

ACCATGTTTAGAAGCACAAATAATCATCACACCAATAAATACTGTCAAAGCCAAAACCATTGTCTGGATA

AGATGACTTGTTATAGTAAAAGAATATGGTACAAGTCCTGAAACATTCGATATCAAAATAAAACTAAAAA

TGACAAATAAAAAAGGAAAATATTTTTGACCTTGTTGACCAACACTATCCCACACAAGACCTGCAGTACT

TAAATAAAGACTTTCTAAAACTAATTGCCATCTAGTCGGGAAACAACTTAATCCATAAACTGAAAGACAA

TAATAAATTAAAACAAAAAAGGCTAAACCGACGCATGTTGTTACCATTGCGTTAGTTATTGAAAAATCAA

ATAAACCAACACCGAGACTTAAAAGAGGAAGTATTTGGAATTGTTCTAGTGGGCTGTAAAACATGTATAA

ATTAAAAATATTATAAGCTAAATAGTTAGATATCAACCTGTTAAAATATGTAAAAAAGTTTATTTTTCAG

ACTTTTTGTTAATCCATGGGGAATATACCCGAAAGGGGATCCAATTATTCCTACGGTGCTTGAAAAAATT

TCTTTATAAAGTACCATAAATGTTTTTCTCGTATTCAAAAAACAAAACTATCAAAATATACACTTTTTTA

GAATCCAAAACAGTAAAAGTATCCTTATAGAAAAATTCTACTAGAAAATTTTATTGTCAGTTATAACCAT

ACCATTTTTGGAGACCCATACCAGTAACGTCAAATAATTTTATATAAAGTAACTTTATACTCTAAAAAAA

AATATTAGCACTTTCACGGTAAATACCTATCATTTTTTTTTTTACAATAACGAACTAAAACCTTAGACGA

CCACAATAAAACCTGGAAGTCATAATTCCAATTATCACTATCCTTTATTGTAGCATGAAAAGATAATTTG

TTAGGTAATTTATAAATACATTTCGTAAAAGTATAAAAACACCAACTATATCTAAGCAAGAACGAAGAAC

ATACTTTTAAATAATATTCTACCCCTTTATAAAATTGTAGATATTTAAACACTTCAGAAATTTTAGAGAC

CTCAATTACAAGATAATTAAAAGTTTGTGGGGGTGTCTTTTTATGGCTATTAATTATTAAATGACCCCAA

TTATACATCGTATCCTCTAAAAAAAAACTTAAGATATTTGGTT

>NODE_23_length_1441_cov_14.319014

AACCTCCTCTGACACCACTCCTCCTACCTACCGCTTTCTTCCAGAGTCCTTGTTGAATATGACGAAGTCT

TGGACTATGCTCGCACTGGCGACGTCGGCGCTCATCTCCACGGCCAGAGCTGCATGCGTGGACATCAACC

AGTACCTCGACATACCCTCGCCAATGACGGGCGACATGGAGATCAACCTCGACGCGGTGGACGCGATTGA

ATGTCAAGAGATCGAGGCATTCGACGAACCAATAGAATTGGAGATTGTCACTGTCGACGGTGGAACCCTC

ACGCTCAAGAGCTCCAACAGTGTCCGGTTTGTAAACATGGGCTTCACGGTGAAAAATGGAGCCAGCATCG

TGTTTGACATGCCCAAGACGAAGTTCGGGCCCAACACCTTCGGGAGTCATCGCGGACGACACTACAGCGT

TACTGTGGCAGTTGAGGAGGGCGCGAGCGCTACTTTCGTCGGGCAAGTTTTGACGGAGGAGGTGACCAAC

GTCTTCTCCCTTTTCACGAACGCCGGCAACATTGAGTTTAAGGACAGTTTTGTGGTAAAGGACAGCGGGA

ACGTGTTCAGGCTCAACACTGGGGTCACGAAGTTCGGCGGTGATGCCACCTTCAAGGACAACAGGTACCT

CGCCCTTCAAACCCTGGACTCCGCATACGTCACGTTTGAAAAAAAAGCAAGCTTCGACAATAATGCTCAC

AACTTCGACGGAGCAGGTGGCTGCAGTGTGACGAACGAAGATACTTCGACAGTCATCTTCCGAGGAGACG

CAACGTTCCTGAACAACGCGTGCAAGGAGAGCGCAGCCGTTTCGAACGAGGGAACTATGACGTTTGAAAA

ACAGGCGTACTTCAACGATAACTGGAACAACTTCGACGGTGGCGGTGTAACAAACTGGAGCGGGACGATA

TTGTTCAAAGAGAATGCACAGTTCAACGACAACACTGCGAGCGAAGGGGCGGGCATTTATGTCAGAGATG

GCAAAGTAACCTTCAACAAGGCGGTTCTTTTTGACGGCAACACAGCTAAAACGGGCGCCGCCTTTGCCAT

CGGAGACGGTGGTTCGCTCACGTTCAAAAAGCCGAGCGACGTCATCTCCAAGAACGCCATGCTGAGCACG

TATTCGAACGAAAAAAGCTGCACCTTTGGAGAGGTCTCCGACGGAGCGACTCTGATCGGCTTCACCGGCG

ATGACGTCTGCATCCCTGCTGGCGATGCGGCTTAGACCCCGCTCACGACTCCCACAGTAGTGAAGTTGTG

GTACTGGTATTCCTCGACTGTTGCGGCTGCTCACTAACAACGCAGAGCCTTCAGATGTCTATTTGGATTA

GCGGTCAACTTGGACCGGGTGGTCAATTTGTTTTAGTCTGCACAGCCGATTATGTTTGCTCGGACTCTTG

AATATCATAACCGAAAGAGCTTCCTACGACGTTTGCAAACG

>NODE_24_length_1357_cov_18.514970

ACCGGTCAGTTTGTTGTCTGGCGTGCTCAACAATCTTGCACGCATGAAATTCGCTTTAATATTCCAAAAT

GACACTGCGGCTTTTTTGCCTGAGTTGTTTCTTGCCATCTCTATATTAGTTATTTTATTACATGGTAGTT

TTTTAGGGGTAGCCGCGGCTTCTCTTTATACTTATTTAACTCCAAGTATGGTTAGATTAACGTCAACTAT

TTTGTTTGTAAGTTTGCTTTTAGTGTTGAATAATCCTGTTGAATCTCAAACTTTATGGAATGCTATTTTT

GTTACTGATAATATAGGTACTTGGTCGAAAGCAATTGTTTTTTTAGGTCTTATTTTTTGTGTGGCAATAA

GTGAATCTTATATGTTTTCAGCTCGTTTTAGAGCTTATGAGTTTTTTGTTTTTGTTATTGGGATTGGTTT

GAGTTTGTGCTTATTGATTTCTTCATATGATCTTCTTTCTATTTATTTAAGTATGGAATTTTTGTCGCTT

ATTTTTTATGTGTTAGCAGCATGGAAAAAAAATTCGTATTTTTCTGCTGAGGCTGGGTTAAAATATTTTA

TTTTAGGGTCTGTTGCCTCTATTTTTTTTTTGTTTGGTGCGTCTTTAATTTATTTCTCTATGGGTACAAC

TAATTTAGGTTCTTTAACCCTTTTAACAGAGAATTTAATAACATCATCTCCTTTTATATATTTGGGACTT

ATATGTGTGGTTTCTGCTTTATTGTTTAAGATAGGGGCAGCTCCTTACCATATGTGGATAGCGGATGTAT

ACGAGGGGGCCCCTACTTTAGTAGGTTTTATTTTTGCTGTTATACCTAAATTCGCTTTTTTTGTTGTAAT

ATTGCGTTTATCATTTACAAGTTTTTGGTCGTTTTTCCCGAGTCTATGGGAAAACTTTTTTTGTATCTGT

GGGCTTCTTAGTCTTTTTATTGGTTGTGTTTGTGGTTTAGGGGAAACAAAAATAAAGCGCCTTCTTGCCT

TTAGTAGTGTCGGTCATGTAGGTTTTTTATGCCTTGGGTTAGCTAGTGGTAATATAGAGGGAATACAAGC

AGTATTATTTTATCTTTCGATTTATATGTTTACAGCAGCATTTTTATGGGTTTATATAGTGCATCTTGAT

TTATCCTCTACCTCTGGAAGTACTCTTTTAACGTTTTCCGATTCTATAGGCTTGGTTCGATCGAATCCAC

TTATGGGGTTCGGGGTTGCATTAATGATTTTTTCCTTAGCCGGTATACCCCCTTTTGGTGGTTTTTTTGC

TAAATTAAATATTTTTGTGAGTTTAGTGGATTCTTCTTTTTATATTATAGCGATTTTTGTTGTTATTACT

AGTGTTATTTCAGCTTTTTATTTGGAA

>NODE_25_length_1336_cov_602.565779

AACAGGTTATTATTAACTTGTATCTTGGTAAAAATTATAAATATTTTTGAAAGATATAAGTTACCTGGGA

AGAATTAAAACTTATTTTAAAAAAACTATTATGGTTGCAACTTTAGAAAGACGCGAAAGTGAAAAAAGAG

ATTGGGGAACATTTGCTACATGGATTACTAGTACTGAAAACCGTTTATACATTGGTTGGTTTGGTTGTTT

AATGATTCCTACATTATTAACAGCAGCTTCTTGTTACATCATCGCTTTTATCGCAGCACCTCCTGTAGAT

ATTGATGGTATTCGTGAGCCAGTAGCCGGATCTTTATTATATGGTAACAACATCATCAGTGGTGCTGTTA

TACCTAGTTCAAACGCAATTGGTATTCATTTCTACCCAATTTGGGAAGCTGCTTCAGTTGAAGAATGGTT

ATACAATGGTGGTCCTTACCAATTAATCGTATTCCATTTCTTAATTGGTGTTGCTTGTTGGATGGGTCGT

GAATGGGAACTTAGCTACCGTTTAGGTATGCGTCCTTGGATTTTCGTAGCATTCTCTGCTCCAGTAGCAG

CAGCTTCTGCGGTATTCCTAATTTACCCTATCGGTCAAGGTAGTTTCTCTGACGGTATGCCTTTAGGTAT

TTCTGGTACATTTAACTTCATGATCGTTTTCCAAGCTGAACATAACATTTTAATGCACCCATTCCATATG

GCTGGTGTTGCTGGTGTTTTCGGTGGTTCATTATTCAGTGCTATGCATGGTTCTTTAGTAACTTCAAGTT

TAATACGTGAAACAAGTGAAGTTGAATCTGTTAACTACGGTTACAAATTCGGACAAGAAGAAGAAACTTA

CAACATTGTAGCTGCACATGGTTACTTTGGTCGTTTAATCTTCCAGTACGCTAGTTTCAACAACTCTAGA

GCTTTACATTTCTTCCTTGCAGCATGGCCTGTAGTTGGGATTTGGTTAACAGCTTTAGGTGTTAGTACTA

TGGCATTTAACTTAAATGGTTTTAACTTTAACCAGTCTGTTGTAGATAGTGAAGGTCGTGTTATTAACAC

ATGGGCTGACATCATCAACCGTGCAGATTTAGGTATGGAAGTAATGCATGAACGTAACGCTCATAATTTC

CCATTAGATTTAGCATCTAACGAAATTCTTCCTGTTGCGATTTCTGCTCCTTCTGTTGTTGGTTAATTCA

ATTAAAATAATCAGATTTATTTCTGTATTATTATTTTATCTCTAACTACCTTTTTTGGTAGATAAGAGAG

ATAGGTCGAAAGATCGCTAGAGTAATAAGAAGTTCTATACTTCTTATTACTCTATTTGGAATTCTCGGGG

TGCCAA

>NODE_26_length_1326_cov_15.268966

CTTGCAACGCAAAGGGTCACCACACGCCTCCAGGCCGCCTGGCGCTCCATGAGCTGAGCGGTGACGAAGG

GGTCGACAAGAAGGTTGTCGAGGTTGGCGTTCTTGATGTATGCGTTCTTGATGTCATCAAGGAACGCAGC

CCGGATGATGCAGCCACCCTTCCAGATACGGGCACACTCTCCGAGGTCCACGCCCCACTCGAAGTGCTCG

GATGCGGCTTTGATGAGGTTCATGCCCTGGGCGTACGAGCAGATCTTGGAGCAGTACAGGGCGGCCTTCA

TGTCTTCAATCAGCTGCGCCTTGGGCACCTTCGGGATCGTGCTGGGTCCCTTAAGAGTCTTGGCAGCCTC

TACGCGCTCGGCCTTACGTCCGCTCATGTAACGGGCATCAAGAGCCGCCGCCATGGTAGGCGCGGCTGTG

CTGCGTTCGGCGGCTTCCTGAACCGTCCATCGCCCGGTGCCCTTCATGCCTGTCTTGTCCAGAATCTTGT

CCAAGACGTGGCCGCTGTCCGTGATGTCGTCTTTCCTCGCGAAAATGACGGCCGTGATCTCGATCAAGAA

GCTCTGGAGCTCTCCGGAGTTCCAGTCCGTGAAAGTGGCGGCAAGCTCGTCGTTGCTCATGCCAGCCACG

TGCTTCAGCACGTCGTACACCTCGGCAATGAGCTGCATGTCTCCGTATTCGATGCCGTTGTGAACCATCT

TGACGTAGTTTCCCGAGCCGATGCTGCCGAGGTAGGTGGTGCACGGTCCGTCGACCTGAGCAGCACACTT

GGAGATGATGGGGGAAAGCGCGTCGTACGCCTCCCTAGGCCCACCAGGCATCAGGGAGGGGCCGTTGCGC

GCGCCCTCTTCTCCACCAGAGACACCCATGCCTACAAACATAATCTTCTTCGGCTCCAGCTCCTTTGCAC

GGCGCTGGGAGTTGGGGAACCACTCGTTGCCACCGTCAACGAGGATGTCTCCTTCCTCCAAGAGCTCGGA

AAGCGTAGCGATCGTGGCGTCTACGGCTTTGCCGGCCTGCACAAGAAGGACGATCTTGCGAGGCTTCGAC

AGCGATTCGACGAAGCTCTTGGGGTCCTTGTAGCCCTTCAACGGGAGGTCGCCCTCCTCCTTGGCACGCG

CGACAGTTGCGTCAACCTTCTCGGGCGACCGGTTGCTGACAGACACGGAGAAACCGTGGCTCGCCATGTT

GAGGGCGAAGTTCTGGCCCATTACCGCCAAGCCATAGAGACCAACGTCGGCCAACGGAGAAGCGGACATC

TTGGATGGGTTGCGTAGAACTTCTCTCGATGCTTAGCGCTTGTAGTGGAGATAAATTGAGGCGCTG

>NODE_27_length_1309_cov_10.083075

GGAAGCCCTTGATGCCCATCTTGTCCTTGAAGGCCTTGTTGTCCTCCTCCGTGCAGTCCATCTTGGCGAA

GACAAGCTTGTCGTCCACATCTTTGATCTTCGCGGCCGCCTCCGACAGCGCCGGTGCCAGCTTCTTGCAG

TGACCGCACCACGGCGCGTAGAATTCTACGAGCACGTCCTTGTCGTTCTCAAGCACCAGCTTGGAGAAGC

TCTTCCCCTTGAGCACCTTCACGGGCTGGGCGGTGTCCTCGTCCGAGGGCTCCTCGGACTTAAGCGAGGG

CGACAGGTCACCCTTGAAGAACTTCTCCTCGAACGCCACGAGGCCGGCCTCGTTGATCTCACCGTCGTAG

ATGTACTTCTTGATGGCGCTGTTGGAGGTCATGTCGGCGATGACCACCATGGGCAGGTTCTCTTCCTTGG

CGCCGAAGTACTGGAGCACGCGGTCCTCGGTGTGGGGCACGTGCACGTGCAGGAGCTTTCCGCGGTTCTT

CTCGGCCACCTTGGTGAGGGATGCGATGTGTCCCTCGAAGGGGGCGGAGGTGGCGTCGGACATGTAGAGC

ATGTGCACCTTGATGTTACCGCCGAAGATAGCAGCGGACGACTCGGCGGAGAAGGTGGTGACCAGGCGCA

TGGAAGAAGCCTCCACGAATTCCTCGATCTCCGTCGCGGTGGTGGCCTTCTTGGCGTCCATGGTGGCCTT

CTTCTCGTCGAACTCCTTCATCACCACAACCTTGTCCACCTTAGACGCCTCGTACGCCTTGAGCACCTCC

TTGGTGGAGGAGATGGCGAAGGGAAGGCGGTCCACGCCGGAGGCGGCGGACATGAAGGCCTTGGCCTGCT

CGGAGTCGATCTTGGAGAAGAGCCCGATGACGACAGCCTCGCCCTCCGCCTCGAAGGCCTTAGCCTCCTC

CACGGTGGTGATAGTCTTGGCAGCGGGTCCGGACTTCTTGAGGACGAACTTGACGATGTCCTCGGTCGTT

CGGCCGCCGTTGTAGTCCTGAGGCTTGCCGTCCTTCAGGAGCTTGAGGGTGGGGAACCCGCTCACGCCGT

ACTCCTTTGCGATGTCCTTGTGCTCGTCGCAGTCCACCTTCGCGATCTTGAGGTCGGACTTAGCCAGAGC

CTCCGCCGCCTTCACGTACTCCGGGGCGAGCTTCTTGCAGTGCCCGCACCACGGAGCGTAGAACTCCACC

AGGAGGGTGGGGTTCTGTGCGACGGCGTCCGCGAAGTTGGACGAGTCGAGCACGAGGACACCATCCTCCT

CGGGGATGTCTGCCGCCAAAGCACTGGAGGCAACCACCAGCAATGCCAC

>NODE_28_length_1303_cov_35.807332

GGTGTGACTGGTGTTGCTTGGCTCGCGAGTTTTCCATTTCCGCCACATCCCAGCGATCAACGCGCTACTG

GCGTTGCGATGCGAAGCGGCCGCCACGCACGACTCTACACGCACTTGAACACGGAAGCATTGCTGTCCGG

TTTGCCGTACGACGCCGAACGTGAACAAATCACACTATTAATAGTATTTTCGCGGCCGAAACCAAAATAC

CGCCGCTCGGGGGCGATTGAAGAACGTCAGCCACTTTACCAAACCACAAGAAATGTCCCTACTGATATGT

CCTGAAGTCTGCCTATAACCCACCAGTAGTGTCCCGTTCTCGAGGCTTTACGAACAACTGTCGTAGCTTG

ATGAGATCCGTCCGCAAACCACTCCCGGTGTCGAGATACGCTTCGCTGTTGGGTACACGAAGAGCTAGCG

AGCAAGCACGCCTCGCGGCAAAGTCTTTCCCATGGCAAAGGCCCCTTCTATACTATTGATCGACACTGCG

GCGTCGTTTTTTCCCCGTCCTCGCGATAAAAGTGACCATCTATATAGTTTGGGGGTCATTCCAATAACAC

CGGGGTCATCCAAACGTTAACCCTAACCCCCCCCCCGTTCCGCCTCCTTCTATTGTTTTGGACAGGCACC

GTTGCCTCGTCCCCCCTGCCGTGTGCCTCTTACTCGTCGGCGGTGTAGTACGCCTTCACCTTGGGGTGTT

CCATTACGGCGTCGTCGACGGCCTTAAGGGAGGGGCAGATGTCCTCCACCATGGTCGTGGGGATGCCGGT

CATGAAGCCAGCCTGGACGGTGGCGAACACGGCGTGGATCGTGATGTCGGCGATGGTGAGGGAGTCTCCG

ACGCAGAAGGCCGAGCACTTGTTGGCTTCCACCTTCCTGGCGATGTCCTCCAGCAGCTCCTTCACCTTGC

CCGCCATCAAAGCCTGGCACTTCTCCAACTTCTTGGCCTCGTCGCTCTCTTTCATGGTGGAGAAGAGGTT

GATGAACACGTCCTCCGCGATCATCACGATCTCGTCCACCTTGAGGGCCTTGAGAGCGCATTCCGGGTAC

GTGCCGCCGAGCTTTCCGGCGTAGCGGAGCAGAGCCGTCGACTGGGTGTAGTCGGTGCCGTCCATCTGAT

AGACGGGCAGGGTCTTGAAGCCGCAGTCACCGAACTCCGCGAACGAGATGTGCTTGTCCTCGAAGTCAAC

GCCTGCCACGGCCAGAGCGACGCGGGTGGCCTCGCCACGGGCGGGGATGCCGAAGTAGTTGAATACGGGA

GCCATGGTGATAGTGAGTGGTGGTGGTAGTAGTGGTAGTAGTG

>NODE_29_length_1260_cov_24.264730

CCCCGAGAATTCCAGGTTACGCTCTTCTAGAGCCACGAGAAACGCCAACAGAACAAAGCACCTATGGCAG

ACGATCATATGGCAACAAAGCCGCACCGCTCAACAAAGAGACTCCACCCGGACAACTTTTACGTTACACT

GTCCGGTGCGCCTGCCACAAAACGAGGAGCCTTCAACCGTTCACAAGACGTTTTGCACGACGGAGAGAGA

AATCCCCGTCGCTCGGGAACGATTCAAAAACGCCACTACTTATCCAGGGCTGCGCGACAAAGGCACGCCT

TGGCTGGGCTCACACTGGCTACCACTCTGTCTAATATATAAGGACCGTTTGCATACATTATTAATATACA

AACGATACTTATATTTTCGATACGCCACGGCTAGTGCTGTGCCAGATGGTACCTCGTAGTCCCTCGGAGA

AAACACACTAACTGGCTTGGTTTGTTATCACTGAAGAACCACAACTTCAACCCAACAAAAAATCAGCAGT

CAGCGCGATACAAACGAGAGTGTGAATGAACTCTCCATACTTACTCCACCCAACTAACGTGTCACCTTAT

CCAACTATTAACTAGTTGGCGACGTGGAAGCCCCGGTCAACCCATACCCTTAACACTAGAGCCTGCCACA

ATGCTACAGTAGTCCCAGTTTCAAGCGGATTTTGTCGCTAAATATACCATATGTTTGGAGCCAGTTCTCA

AAGGCGATTCAACGGCCTAGTCGAGGTCGACCAGAGTCCGAGCAAAACTTCGCCCGAGAAGAACCTCAAA

CACCTGCACGAATCATCTTACTCCTTACTCCAATTCCAAAGTGGCTTTGCCGCCAAACCATGGAGATACA

AGAAGTTCTGAACGCGTTTGTTTCCAACGATCAAGTCGATCGAGCAGCCCGAGCTACTCGCCCGAGGGAA

CACACGATTCCACACTTGATTATAATGAACCTATGTTAATTCCAAGTGGTTTATCCTCGAAAGCGATGGT

ATCAGTTCTCTATATGTACGAGGCGTTACGAACCGTCGGAACAACAACTCGCTTACGTACGCCTGGGATC

AACTCCCCGCATCGAGGTACGCCTCCCTGGTGGACACACCAAGAGGTAGCGGGACAAGTGCACCACGCTG

CGTTGCCTACTCCGTGCCTCTACCCTGCCTGTGGTCGCTCGAACAAAATTGTGGGTCACCGTTCCTTCCC

CCCCCGTCTTCAGTTCGGTTTCGAGCGAAGAAAATCGTTTTTCTATGTGTGTTGGTCGTCCAATCATGAC

>NODE_30_length_1253_cov_337.237825

AAATAAGTATTGTATTAAAGATAATTACAATAAAACCCCACTATAACGTAATAAAACGTAACTGTGGTAT

TAAAAAAAATGAGTTTGATCCTGGCTCAGAGTGAAGGCTATAAGCCTGCTTGAATACATGCGAGTTGAAT

GGTTAAAACCATAGCGTACGGGTGAGTAATAGGCTAATATCTGGCTTCAACTTCGGAATAATCCTATCCC

CCAGGGGAGAAGGCAACAGGAGAATACCGGATAAATATATAAAGGTCCGCCGGTTGAAGATGATTAGGTT

CAGTATTAGGTAGTTGGTGAGGTAAAGCTTACCAAGCCAATGATGCTTAGTTGGTCTGATAGGACGATCA

ATCACACTGGGACTGAGACAAGGCCCAGGTTTCATTTGAAGGCAGCAGTAAGGAATATTGGACAATGAGC

GAAAGCTTGATCCAGCAAGGCTTGGTGAGGGATTGAAGGCTTAGGTTGTAAACCTCTTTTTTAATTGAGG

ATAATGACAGTAAATTAAGAATAAGTCCCGACTAACTTCGTGCCAGCAGTCGCGGTAATACGGAGGGGGC

AAGCCTTATTCGTCATAACTGGGCGTAAAGGGCCCGTAGGTGGTTTTCTGATATGTTATTTGTCAAAAAC

TGTAGCTTAACTATAGATCGGCATTTAAAACAGGAAAACTTGAGTCTGACAGAGGAATATGGAATTTTTC

AATTAGGGTTAGAATCCAGATAAGTGGAAGAGAACATCATGTGCGAAAGCGATTTTCTTGTTCAGTACTG

ACGCTGAGGGGCGAAGGCGTAGGTAGCGAACAGGATTTGAGACCCTGGTAGTCTACGCTGTCAACTATGA

GTGCTAGCTTTTAGAAATCTAGAAGACATAGCTAAGGCATTAAGCACTCCGCCTGAGGAGTACGAGCGCA

AGCTTAAAAATCAACGGAATTGGCGGGGGTTCAAACAAGTGGTGGAGCATGTGGTTTAATGCGATAATAC

GCGCGTAACCTTACCAGTTCTAGTAAGTCTGTCGTTCTTCCGGAGACGTTAAGAATTTTGGGACTTTCAG

GTGTTGCATGGCTGTCGTCAGCTCGTGTCGTGAGATGTACGGTTAATTCCTGTAACTGAGCGCAACCCTT

ATCTCTTTTATGTTTTGAAATGGTCTTTACCAATAAATAAATTGAATAGACACTGCCAGCGCTAAGCGGG

AGGAAGGTAGGGATGAGGTCAAGTCTTCATGGCCCTTATGAACTGGGCTACACACGTGCTACA

>NODE_31_length_1240_cov_12.382281

CCAAATAAGAAGCTCATTAATTGATATATTAAGCATAATTAGGGTAAAACTTCTAAATCGCACGGATTGT

TAGGTTTAAAAAATGGTTTTTCTTTTATAGAAAAAACAGCATTTTTAGATTCACGACCTAACTGCCGGTA

TAAAGATTCAGGACAATTTTTTTGTAATGTTAAACTAATAGCTCCTACCATAGCTATTAAAAGAATGATT

CCAGCTATTATTAGTAATATTGCATGGGTTGAATATAAAAGATAACTAGGATCATTTAATGCGCAAGAAG

AGTCAAATTCTATAAACCATATTGTATTTAACCCATAAGACCCTTCTACACTTTCTTTATGAAACAATAA

TCGAAGAAACTCTGCTAACCCTTCTGAATCACATAAATGCTGCCACATTTCAATTCTCTCCTCCATAAAA

TCTTTAAAAGATTTTTTAGCTTCTTTAACCGACGGCTCAGGTTCACCTTTTCTTTTTCTTTTACTTCGCT

CCTGGAATCTTTTTAAGTTGTCATTAACTGTTTTGTTAATAATTACCGGGTGAAATAAATTTTTGATCTC

TTCTTCGTAAGATACTAAACTAAAAGAAACCAATGAACTCATATAAACTGTTGACACCATATTCATTAAA

TTTTGCAAATCTTTACTATATATTGCAGCACTCAAAAAAAATGCAAAAATGAAAGCTCCCAAGTACAACA

AACGTTTTTTTTTTGCAGACCATGGGCTTTCTATAGCTTTTACATCTAACATCATAACAACGAACAACAC

CAATACTGCGATAGCACCGGCGTAAACCAATAAAAAAAGTAAAGCCATAAATTCTAAACCCAATAAAAAT

GAAATAGCGGAACCCAAAAAGAAAAGTAATACGAGATAAAGACCACTATATACTGGATTTTGTGTACTAG

TAACTTTAACCCCTAGGGTACCACCAAGAGTTGCAATAATAATAAAAATTATAGGATCGACCATAATACA

ATAAAGGTTAGCAGCGCTAAAACACGTGAAAATTGAAAACCAAACACGAAATATATTCCAAATAAAAAGT

TACTCCAACCTAAAATCACTAAATAGTGATATAAATAACCTGAATGCCATGCACGAGATTTATTTACTTG

AGCAATAAGTACGTTGGATATCCCAAAAGGACCTAAACTTTCAATAAGACCACGATCAATGGATTTATAA

GTAAAATGATAACCAAAGTCTAGTAAATTTTGAGTTATAACTTCATTATA

>NODE_32_length_1230_cov_8.396195

GTGTCGAAGATGGCGGAAACCCCGGAGGAGGCCGAGAAGGCGTGCGCGGACATCATGGGCGGCGACGTGA

TGAAGGACCTCGGTGTCAAGTTCGTCCGCCCCGCCATACCTTCCAAGATCGGCTTCGAGGCAGCGTCATA

CTGCGCCACACAGAAGGACCTGGACGAACCGTGCGATTGCTGAAAAAGCTTAGGATGGTCGCCGGTGTTC

CGCAGGCCGTGTTTGGGTCGGCAGCTCTTCTTCTCGCAACGACCACTACTACGCTCGTGCAATCAGCTGG

CGCAAGCTCGTCCTCATCGTCGACCACGTTTTTCGATCCGTCCACCGTTACCTTATATTCTTCCACGTTT

TCTGAAGCGTGTGCAGACGAACAATTTTCCTGTATTATTAGTGACACCTGCTACAGTTGCTCTTACTTGG

ATGAGGTGGATACCAGGCAGCTCGCGGCATGCGTCGACGATATATCTGTAGACGACCCTGACAGCTCCAC

GGAAAACTGTTCCGATATCGTGGCCATCATTTGCTGCTTCGACAAACTTTCGGATAACGATTGCGTAGAT

AACGATCAATTTGTTGACCTGCAGCTGTGCAACTTGGCAAGCCAAAGCTGTTCCTTCGACGATATCACGT

GTGACTACAACGGAGGCGCGGCGGGTGCAAGAAGCAACGCGGCGGTCTCCGTCGGCTCTGCTTCGTTTGC

CGCATTTGCCTGTGCGTTGTTGGTACATGTCCTACCTCTTCTCTAAGGGTTTGAATACGGACGTGTGAAA

GACGTGATTCACTCCAATGCCCTGTTTGTGATAGGTGTTTGAACGTTGTCTATGGGTAGATGGCCTTGCG

TTTCGGAATAGTGTCCACGTGAATGGTAAGGAAATCACACATACGCGGGGTAGGTACCGGTCGAGCCACA

TGATAAGTTTTGCGTTGTTCAGGGATGGTGTCTGTGCTGAACCAATTCGGTAAAAAGGGGATTAGGCGGC

CACCTCCGTTCGTGGAGAGCGTAATAGGTGGACTTCGGAGGTGGAAAGACTCGGCGAACATTGGCTGCCT

TAGTTACACGGGGGACTCTAAATACGGAGCACAGTTTTCGAGACAAAGGCAGTCATGGCTTATGTCAGTA

TCATCTTCCGTCGCTGACGCACCGTTTGGGGTGCAAGGGTTGTCCACTGCTTATACGGGTTGTTTGGCGA

CATACTCATGAACCCTCAAGGTACATGTTCAAGGTGGAAT

>NODE_33_length_1215_cov_40.130653

GGACGGCCAGCTCTGCAAGGGGGCATACACCGGGGACGACTTCTGCGACGTGGTGGACGAGGACAAGTTC

GAGACTTTTGAAGACATCGTCAAGCACAGTGCCACTTTCTCCTTGCACACCGAGCTGTGAAAAACGTCAC

GGACCTAAGGGCTCCCCAGTTGCCTTCTATAGTGCTTTCAATTGGCCCCGTGGCCCGTTGGACCTTGCGA

GATGCTGGATCAGATAAAACACTCCGGCGCACCGAACGTACCAAATTGTTTCCACTGGTCTGGGGGCGAG

TTGGGCATTATAGAGGCTGTAAGAAAGTTTGCCGATATGACCTGTATATTTATGTGACGGAAGAGTTTCC

GTTCGTGACATGTCCATCATTTTCTACAACAGGGTTCGATGAGTTTGTGCTTGTATTTGAATGTGCCTGC

ACGTTGTGGTGCTCTACTGCCTTTGCCTCGTTGTAGCACGTCACTGGAGGCTGACGAGAGAGGGAGAAAT

GCACATGTGTTCATTAGCGCACATTACTGTGCAGCTATATACGTCGGGTATGCCAAATAGCCCAGGCCCA

GCAGAAGAAGTGGGCCAAGTACTTCGCACGTACGAGAAGGCGGGTTTGTTTGCAACGGGTTCGTTACATT

GCAGATAGCGTGCACAATAAAACACTGCAGCAGTGGCATCGATTCGTAGCGTTAACAGGGAAGCCCTTCT

ACTTTGTTTGTCTTGTCAAAAGTTCACGTAACTCCATGAAATGGGAAACAGTTAGAGCCAACCCGGGTTT

CGTACTGTCGTTGGTTCTGAATTCCGCGATATACACCGTCCGAGGTGGAGTCCTGCTGCTGCATAGCCAT

CGGGGGTTTGCTGTCCTTGACATTCTTGGGTATGGCCGGGTTTCTGTCCGAGTTAATATCGCTAGCGTCG

GGTTTGGCGGCGAGGTTGGAGATTTGTGCTGCCCGAGCTCTGTATTTTGTCTGTTTGAGTGAGCGTTGTC

TATGCGAGTGAGAGTTGGCAAGCAAGGGTCAGTTTTGTGTGTGTGCTCCTTGCTGCCACTCACGTGTCGG

TGTTGAATGTAACGCCTCGCTTCGGTTGTCTGCGAACGCGCGGTGTTGGAGCGTGTGAGAAGAGTACTGC

CCGTAGCGCTTTTCCTGAACGTGTTGCGGAGTTATAACGAATGTCAAGGACAGGCAACCTACTGTCTGTG

AGGGAAAAAAAAAAATTTTTGGAAT

>NODE_34_length_1203_cov_54.741963

TTAGTAGGATGCAGCTCACCCCAGACACCACTCGCTCGAACACCTCAATGATTCTTCGCGCTGGGTGCGG

GCAATGTCTACGTGGTATGTGAAAACGCAACGTGCTCCGTCCCGGCGTTGTAGGGGTGCGCCGGTCTATG

TGGTCATGCGACGACTACGGTCATGCATGACATAAGTACATCGCGCCCCTTCCGCCAGGCGCCAAATGTG

TCCGAATACCGATCCTGGCTTTCCTTAAGCCTACGTACAGAATACAACCAAACGAAAAAGCCAAAACGGG

TTTGTACGACCAAAACCGGCATTTTAGGTAGTTCGATGCGGTTATTTCTCCTTCTTCACCCAACTGCAGG

TCTCGACTTCAAGATTCGGGCGCATCTTTAAGCTCTCCGTACTTCTCTTGGACGTGCTTGTTCAGGTCAG

TGATGGTCGTAATGAGGGTTTTGAGGTCATCGATCGATCCATTGTTTTCTGCCAGCTTGCTGGTCAGGGT

CTTAAGAATCTTGAGGTTGATGGCCAGCTCCTGCACGATGGCGAAGTCGTCCACATCTGCAGCGAGGGCC

TCCGAGAGCAGAAAGTCGATCTCGTCCGGCTCCAGCTGAGCCATGGGGATGACCGCCATGGCGCACAAGT

ACCCGATCACCGTCTCGCTCGCCTCGGCCCCTTGCGCTTTGGCTGTGAGCGACCTCTTCCAGAGATGGGT

GTCGACGCGCACGAAGGCGTTGAACAGGAAGATGACGAAGAACTCGTGGGCCATGCTCTCTCCGGCTTGC

GTTGCTCCCAGGAACGAGTTCAAGGCGCCGATGACCATTTGCTTCGCGCCCTGGATGGCAGACTGCTTAG

CACCTTGATTGTTTCCAGCGGCACCCTCAGAGGCAGACGTGAAAAAGTCACCGACGGCTGCTTCTATGTC

CTTGTCTAAGCCAGCACTGGTCGTGACCTTCACGAACTCCTTGAATCGTAGCGTGCGGCCGCCCGTGACC

TCGGTCTTGCTGAGGGCTGCCTCGTTTCCGCGGGTCTCCTTTACCTCCGCCCTGAAGATATCTATCTGCT

TCATCGCCAGGTTTGACAGATTGTCGAGTTCTTCGGAAGCGTCTGCCAGTCCCTCCTTCGCTTTCGACGT

TGACTGGCCCATTGTCTTTTTGGTTTGAAAGGATCAGGGGAAAAGTCTTGATCGGCTGAAAGCTCTAGAT

GGGCACACGAGAA

>NODE_35_length_1203_cov_9.816413

CTACAGGTGGAGCGAGTCAAGCGCGTTCTCTCCACCCAGCACCAGGCCCGTGTGGAGATCGAATCTCTCA

TCGACGGCGAGGACTTCTCCGAGACGCTCACCCGCGCTCGTTTCGAGGAGCTGAACATCGACCTGTTCAA

GAAGACGTTGGGCCCCGTCCAGAAGGTGATGAAGGACGCCGACCTCAAGCTGTCTGAGGTGGACGAGATC

GTGCTCGTCGGCGGCTCCACCCGTATCCCCAAGGTCCAGGAGCTCCTCAAGAACTTCTACGAGGGCAAGG

AGCCGTCTCGCGGCATCAACCCCGACGAGGCTGTGGCTTACGGCGCGGCTGTGCAGGGCGGTATCCTCTC

GGGCGAGGCTTCCGACGCGACTAAGGACCTCCTCCTCCTCGACGTGACCCCCCTCAGCCAGGGTATCGAG

ACTGTCGGCGGCGTGATGACGAAGATCATCAGCCGTAACACCGTTATCCCGACCAAGAAGGCGCAGACGT

TCTCCACCTACCAGGACAACCAGCCCGCGGTGCTCATCCAGGTGTTCGAGGGCGAGCGTTCGATGACCAA

GGACAACCACGTATTGGGTAAGTTCGAGCTGACCGGCCTCCCGCCGGCGCCGCGAGGAGTACCCCAGATC

GAGGTGACCTTCGAGATCGACGCCAACGGCATCCTTCAGGTGTCGGCGCAGGACAAGGGAACCGGCAAAG

CCGAGAAGATCACCATCACCGCAGACAAGGGGCGCCTGTCGGAGGAGGAGATCGAGCGAATGGTGAAGGA

GGCCGAGGAGTACGCGGAGGAGGACAAGGCGGTGAAGGACCGCATCGACGCCAGGAACTCGCTCGAGTCG

TACTTGTACAACATGAAGAACTTGTTGGACGACGAGGAGAAGGGGGTGGCTAGCAAGATCAGCGAGGCGG

ACAAGGAGGAGCTCGAGTCTACCATCAACGAGGCGTTGGATTGGCTCGACGAGACCCCCGACGCGGAGCT

CGAGGACTTGGAGGACAAGAAGAAAGAGGTCGAGCAGATCGCGAACCCGATCATGAAGGACCTGTACCAG

CAGGCACCTCAGGGCGGAGGGGAGGAGGACTTCGACTTCGGAGACGACGAATTGTAAAATCTGAGTTGCG

TTAATATGGTTTAGGTCTCTTTTTTGTCTATTTTCGTCGGGGAGAAAAGAGACGGCGGCGGCGGTGGTGG

TGGTGGTGGTGGT

>NODE_36_length_1196_cov_12.812766

CCAGCTGTAGGGGCTTCTACATGGGCTTCAGGAAGCCAAATATGTACGGGTAACATAGGAACTTTAGCTG

CAAATGATGCAAAGAAAGCAAGCCATAGCCACTTTTGATCATAAGGTGAAAAATTTATAACTGACAATAT

TTCGTAGTTTGTACTCCCGGCGAATAAATAAATATAAACTATACCTATTAACATGAATAAAGAGCCTAAA

AGGGTATACAAAAAAAACATATAAGCCGCCCGTATTTTTCTTTCACGTGATCCGTATATACCAATAACTA

AAAACATCGGGATTAATACAGCCTCAAAAAAAATATAAAAAAACAACAAATCTAAATTAGTAAAGACTAA

AAGCAAAACAAGTTCCATACCCAAAAAGCTTATTAAATAAGTTTTAACTTCCCCTTTATCAAAAGACCAC

GAGGCTAATAAACAAAGAGGGAAAATTAATGTTGTCAAAATAACAAAAAAAAGAGAAATCCCATCAATAC

CTAAAATTAAATTAATATTAGCTATAGGCAGCCACAAACTATCAACAACAAATTGAAATTTAGGTGTTGA

GTGATCAAAACCTAACCATAAAAATAATGAGGAGACAAAAACCGCCGACGTAATACCGAGAGCAAATTGC

CGCATAATTAACTTTTGACTAGAAGGAATAAAAATTAAACCAACAATTCCTAAAAGAAGAAGAAAGAATA

AAAAAGTTAAGAGCATAATCTTTTACCAACCCAAGTTAAATACTCGTCTTGGTTAACCGCTTGAACTACA

ATAGGCATAAACCCATGGTTAACACCACAAAGTTCACTACATTGACCATAAAAAACCCCTTCTCTTTTAA

TAAAAAGGAAAACTTGGTTCAATCTACCCGGACACGCGTCTACTTTAACCCCTAAACTTGGTACTGCCCA

AGAATGAAGAACGTCAGCAGAGGTTACAAGAACACGGATATGAGTGTTGGTTGGGACAACTAGACGATTA

TCAACCTCAAGAAGTCGGAAAACTTTACCAGCTTTTCCGGTACTTGATACTTCAGGAATAACCAAATCAT

CATCCGCAATAAGATATGAATCAAAATTTATACGTTGTCTTTCTGTTATACCTTCTTCAACATCTTTGTC

TTTATGATGCTCAATTAAATCATGAATCATGGTAATTTGAGTTTTTAAAGATTTGTTTTTTTCACCTTCC

TCATTA

>NODE_37_length_1162_cov_15.287467

CGTTAATTCCACTTTTAAAATTTTTTTTTTTTGTCAAAACTTTCGTTTTTTTAGTCCTCGTCATGGGAGG

GAGCGGCCCTGGCAGACCCGCCCCTCCCGGCGTTTGATTGTTCATCATGTCGTGCACAAAGGCAAAAGAC

ACTCTGCAAGCTGTTTAGCTCCCAAACCCTCCCCACGCTCGCGCACCCGGTTTTACCTACTCGCTAAAGA

TGCGGCGCAATCGCTTTTCCGGTGAAAAAAGACGCAACACGCAGCCTTAACAAGCAGCGGGCACGAAACG

TGGGTCGTGAATGGTAAGAGTAAACAATTATCGGCTGTAGGCCCTTCCGAAAAGCGCCAGTTGAGAGCAC

CACCGGAACCACCCCGAAGGAGGGGGGCGGACCAGGATCAGGAGAGACAGCATCGTCATCTTCAACAAAG

AAGCTTCTAGGAAACTTCTTAGAAGAAGATCTGGTGAACGGGCCCCATGGTGCTCTTGACGTACATGACC

TTGATGTTCTGCCAGTTCTTCTTGAGCAGAGAGGCCAAGAAGTTGGCGGCGAGCTGCACGTTGAGCTCGA

TCTCGTTCTTGGGCTGCTCCACGTTGGCCACGGCGACCGAAAGGCACATAACCTTCTTCATCTGGAACTT

GATCGTAGCGCGCACGGTGTCCACCTTGCTCTGTAGGTTGTCGCCGGCGTTCACGAGGGTGGGGAACTTA

CCCGCCTTGGTCAGCCCGGGGCCGAGCAGACGAGGGATCTGCTTGATCAGGGAGTCCGAGGCGAGGAAAG

CGTCGTACTTCTTTGCGAACTTCTTGATCAAGTTCTTGTTCTTGTTGAACTTCTTCAGGTCCTCCACCGA

CATGCGGTCAAGCCCGATCTGCTCGGCCTGTTCGCAGTGCACGGCGTTTCCGAGCATGCAGAGCTTCATC

CTCGGGCGGGGGATGGTGGGCAGCTTGAAGGTACCGGAGAAACGCTTGTCGCGCTGGGGGTCGTAGTTCT

TCAGGGTCACCTGGAGTTCGATGGATTCCGTGAACTTGCGCTTCTTGCCCACGACCTCGGCATCGCCCTT

CTTGATGGTCTCACCAGCCGAGTAGGCTAGGATGTCTTCCACAGCCTTGGTTACGAGTTCGCCATTCAGC

TTGCTAGTCATCTTTGCCAGACGTTGCTCGTCAAGAGCGAGG

>NODE_38_length_1152_cov_20.771883

ACTGTAGTTCCTTGGCAACCGTGATTTCCAGGGGTAATTTTAGAATCATCAACTTTAACACTTCCTTCAG

GCATTTTATTCAAACACTGATATATTATACTAAGGGATTGTCTCATTTCCTCCACCCGTACAAGATAACG

GTCATAGCAATCCCCTGTTTTACCAACAACCCCCTGGAAAGACAACTCTGAGTAGGCATCATACGGCTCA

TTTTTACGTAAATCCCAACGAATGCCAGACCCACGTAACATAACACCAGAAAACCCCCAATCAATAGACT

CCTGCGCATTTACCACCCCGATATCAACTAATCTTTCTTGCCATATACGATTATCAGTTAACATTTCTTC

AATTTCATCTAAACGTTGCCCAAATTGAGCAGCAAAAGAAAAAATATCATCTATTAAACCAATAGTGAGA

TCTTGACTAACACCTCCAGGTCTAAAATAACTAGCATGCATACGTGCACCACTAACCCTTTCATAAAATT

CCATTAACTTTTCTCTCTCCTCGAATGCCCACAAAAAAGGTGTCATAGCCCCTACGTCCATTGCATGACA

ACCCACAGCTAAAAGATGATTTAAAATGCGAGTTATCTCCGCAAAAAGAACTCTTATATATTGAGCCCGC

TTAGGTACTGAAATCTGTAATAAATTTTCAACAGCTAAAGCATAAGTATGTTCTTGGCACATCATTGAAA

CGTAATCCAAACGATCAAAATAAGGCAAAGATTGAAAATAAGTTTTAGCCTCTATTAACTTTTCAGTACC

TCTGTGCAGTAACCCTATATGGGGATCAGCTCTTTGAACAACCTCCCCATTAAGCTCAAGAATAAGACGT

AAAACCCCATGAGCCGCCGGATGTTGTGGACCAAAATTTATTGTAAAATGCTTAAGTTTTTTGTTAAATT

CTTTTTTTTTTAATGACATAATAAAGTTTGTAATGGGTTTTTGTGTATTACCAAACTACACGAACAAAAG

AATCAAAATATAAATTATCCCCGATACTAAAAAGTCTTTAAGATTAGCGCCAGAAGGATTTGAACCTACG

ACCTTCAGGGCATGAGCCTGATGAGCTAACCTTACTGCTCTATAGCGCAACCCTGTTAAAATTAAAAATT

ATTGCTTTTATGAGCCATGGTTCCCACAAAAG

>NODE_39_length_1135_cov_10.922801

TGGTGGCGTTTGCCACGCAAGGATATACGAGCCAACAACAACTTCAACGGCTAACCCAATGAACACATAC

AAACTGAATACAACCAAACACGCGGAGGGAGCCAAGGGTTCCATAGTTCGAATTGGAGAAACTTGAAATA

CTTCGTTGGTCATGCGCATTGTGTGCAGAGAGCCGGAAGCTATGCCGCCTGCGATGCGGTTCCCCCGCAA

AACGTGAATACACCCACCGTCCCACCCTAACGTATCGCATATGCCAACCTGTCCACACAAATTATAACTA

CCCTGCGCGCCTGCTCTATGCAGATCACATCAAAAAGCACCTGGCGTGCCAGGCAATGAAATGTGTGCAC

AGTCGCTCCGTTGCCAAATGACGTCAACAAGTAGGCTCTCCGATACGTCCGCTTGTGATGTCACCCACGC

TGTTGGGGAAAATGAAGGCTTGCCTACAGCTCCACCACCCCCCTCCGTACCGTCGACCACATCCACTATA

ATATACGATAGGGACCAATTCCACATATGATACACTACCAATGTCACGAGTTAGCGAGGGCATCCTGATC

AAGCCGTCATCGGGATGCGTCGATTCATACACCTTACACTGCACCGTACGGACCGAACCCCGTAGCTAGC

ACAAGCCCCACGAATCCACTGAGTACACGCAAGGTGCGACCAACTGCTAGGCCCACACCCCCCCCCGTCA

AGGAGGATCCCCTGCCGTCGTCAGAATCGCTGTCCCCGCTGTCGCTGTCCGAACTTTCACTGTCACTGTC

GCTGTCGCTGTCGCTGTCACTGTCGATACCGGGTACTCCGTTGTCGTCCTCAGATGGAGAGGTGTCAGAA

CTACCCGAGGTGACCCCTCGAGGATTGGCGGACGAGGACGACGCGGCCCCACTGCAGGTCGTGGCGGCGT

TTCCGCAGTTTACCTCGAATAACCCAGGAGTGTCGACCGTTTCAGACGGAAAGTAGCTGATAAACGTGTC

CCCGTCTACGTCACATGACCACGACTCGTCGCTCGGGTGCTTCAAGTTGGCAGAAGGGGTAGATGCGCAC

GGTCCGAGACTGGTTGAGCCCACCGCCAGGCCCAGGTTCCAGACGAGAGACTCGGTCCCGGTGGTTTCGA

GGAAACGGTTTGAAA

>NODE_40_length_1117_cov_4.483577

AGACCCTAGCGTGTGTATGCGCGAATCGACGCGGCTGTACTAGTGCACGAGATAAGAAATGTCTGAGCCC

TTTTAGCGGAGACAGAAGGACGACGGTGAAGTACATGATGGGACGGGAAAAACAATAAAATGACCAGGGG

TAGTAGTGTAGTCAACTTCAAAGGCTACTAAAGTCGGGAGGTCGGGAGGTCGAGCCCCCCCCGAGGAGCG

TTTTAATTGGGCAGCAAATCGGCCCTGCTGAAAGCGCCTGACACCTGCTACGATAGGATTGTGCACCCTG

AATTGCCGGAGCTCCGCACTTGAGGAGCTCCATAAACCACCAACAAACACAATTCGGAGGAAAATAAAAG

TTCTACCCGATGAACAACCTCATTACCTAATTACAGTATACCCTGGAAAATATTGAAACACAAGACTGAA

CGTCGGACGGTTTTGGGTTAAGGAGCCCCGTCATTCAGAAAACGTTTCAGCCAAACAGTGCTACCTCCAT

CGTAGCACGGGCCGTCGATCGCAGGTTTAGCAGTGCATGGTGTAGTGGCGGATACAATACCGATCGCTAA

GGGTGTCCACACCCTTAAAAATTAGACTCAACTATAGTCTATTGATTCAACGGTGGAAACCCTATGGCGT

GTTTGTCTACAGCACAAGAGCTCTTGGTGCGGTTTGCATTTTGCACAACGACGACTAAGTTCCACAAACG

CACTGCTTGTAGGAACCAGAAACGTCGAGTGTTGCCTGTTGTCTGGATAGGGGAGCAAATCAAGCTCGTG

GCGCAAGAACGTCCTACAATGAATGAATGACTTCGATTCTCCCGGCTAACGTCGAAACGTTGGGTTTCTT

TGGAGGAGCAAGCTTAGTTAGGGTTTTGCAGGAGTTGTTCCGTGTCGAGCGTTGTCGTTGGGGGGGACTT

GATATAGGTATCGATCGCGACGAACCTCGCTTCTCGTTGAGGCGCCAGAACACGGAAGGCATCGCTTGTG

CGACCGAGGGAGGAGCAACGCACGCGCGCTAAGATTAGAGGTATCACTGTTTGGCTATTTTTGTTTCTGA

ATGGCGGGGCTTCTTAACCCAAAATCGTCCGAATTCATTCTTGTGATGCAATCTGTTTCAGTTTGTA

>NODE_41_length_1079_cov_14.616257

TTCATTTAACTTTACGACCTCCTCAACGTTTATACCGTGCATTACGAATGGGTAATATCGAAACAGTATT

ATCTAGTAGTATTGCAGCTGTCTTCTTTGCAGCTTTTATTACTTCAGGAACTATGTGGTATGGTTCTGCA

ACAACTCCAATTGAATTATTTGGACCAACAAGATACCAATGGGATAGTGGGTATTTCCAACAAGAAATTG

AACGTCGTGTTGAAGTTTCATTAAATGAAGGTGTATCTGAAAGTGAAGCTTGGTCAAGTCTTCCTGATAA

ATTAGCTTTCTATGACTATATTGGTAATAATCCAGCTAAAGGTGGTTTATTTAGATCTGGTCCTATGAAC

AAAGGTGACGGAATCGCAGAAGCTTGGTTAGGACATCCAATTTTTAGAGATCGTGAAGGTCGAGAATTAG

CTGTACGTCGTATGCCAGCTTTCTTTGAAACTTTCCCTGTAATTCTAATTGATAAAGATGGAATTATTCG

TGCAGATATACCATTCAGACGTGCTGAATCTAAATATAGCATAGAACAAGTTGGTGTTAACGTTAGTTTC

TATGGTGGTAAATTAAATGGGCAATCATTTAAAGATGCACCAACAGTGAAAAAGTTTGCTCGTAAAGCTC

AACTTGGTGAAGTTTTTGAGTTTGACAGAACAAGTTTAGAATCTGATGGAGTATTCAGAAGTAGTCCACG

TGGTTGGTATACTTTTGGTCATTTAAATTTAGCATTATTATTCTTCTTAGGTCATTTATGGCATGGTTCA

CGTACTATATTCCGTGATGTGTTTACTGGTATTGGTTCAGAAGTTACTGAACAAGTAGAATTTGGTGCAT

TCCAAAAATTAGGTGATGAAACAACTCGTAAACAAGGTGTAGTATAATTTATTTTTTTAATTAATTAAAA

GGAAAAAATATGGGCATAGACTTTTCACAACTTTCACAACCAGCATTAGTGTATGCAACTTTATTGATAG

GAACACTAATGGTTCTCTTTTTTGCTGTTTTCTTCCGTGATCCACCTAAAATACTTAAAGAATAATTATT

TATAGTTTATCTTTTCTATCTAGATGAAA

>NODE_42_length_1046_cov_14.912195

AACAACTTGGGGACCATCGCCAAGTCCGGAACTAAGGCTTTCATGGAGGCCCTTAGCGCCGGCGCTGACA

TCAGCATGATCGGTCAGTTCGGTGTGGGTTTCTACTCTGCCTACTTGGTGGCGGACAAGGTGACCGTAAC

ATCAAAGCACAACGACGACGAGCAGCACACCTGGGAGTCTTCGGCCGGCGGATCCTTCACCGTAACACAG

GATGCCCCGGACGCCAAGCCGCTCGGACGCGGTACCCGCATCGAGCTTGTCCTTAAGGAGGACATGGCGG

AGTTCCTCGAGGAGAGAAAGATCAAGGACCTGGTGAAGAAGCACTCCGAGTTCATCGGCTTCCCGATCAA

GCTGTACACGGAGAAGACGACCGAGAAGGAGGTCACTGATGACGAGGACGAGGAGGCGGAGGATGAGGGG

GACGACGACAAGGAGCCCAAGGTAGAGGACGTTGACGAGGCGGAGAGCAAGGAGAAGAAGACCAAGAAGA

TCAAGGAGGTCTCGCACGAGTGGGAGCACTTGAACCAGCAGAAGCCAATCTGGATGAGGAAGTCGGAGGA

GGTAACGCACGACGACTACGCCGCGTTCTACAAGTCCCTCAGCAACGACTGGGAGGACCACGCCGCCGTG

AAGCACTTCTCGGTGGAGGGCCAGCTGGAGTTCCGGTCTGTGCTGTACGTGCCCAAGCGCGCTCCGTTCG

ACATGTTCGAGGGAGGAACGAAGAAGAAGCACAACAACATCAAGCTTTACGTGCGGCGAGTGTTCATCAT

GGACAACTGCGAGGACTTGATGCCCGAGTACTTGCAGTTCGTGAAGGGTGTGGTTGACTCGGAGGACTTG

CCTCTCAACATCTCTCGCGAGAGCTTGCAGCAGAACAAGATCCTCCGCGTCATCCGGAAGAACCTTGTCA

AGAAGTCGGTGGAGCTGTTCAACGAGCTCGCCGAGGACGCGGAGAAGTACAAGAAGTTCTACGAGGCGTT

CGCGAAGAACCTCAAGCTGGGCATCCACGAGGACTCGGCCAACCGCGCCAAGCTGGCCAAGCTCCT

>NODE_43_length_1025_cov_8.599602

CTTTGTCATGCTAAGAACTTAAACAAGAAAACAAAACTGGTTCAACAGTAGTCCATGTTACTTAATTATC

ATCCTGCCTGAACACACATAGACGGACATAGACAAAAAGACATCGGGACGGGGGGATAATTGGTTCTGGT

TTTCAATTATGTACAGAGAATGTAAGTCTTTCTATAGTTGCTCTTTGTCATGCTAAGAACTTACCGGAAT

TGGAAACGTGGTTTAGAGATATTATCATAGGGAGTCGGGAACGGGGGGAACTCGATAACAGCACGTTGGA

ATCTTGAGGATTTTAGAGAATACATCAATATTAATCAATGGAAACAAAATAATATTAAAAATAATATTTC

AGATGTCAGCGGAACTATACATGTCATGGAAATTACTACATACAGCAGAGCTACGCATACGACTAATCTA

CATACATCTTGGAAACTACTATACAAACACTTTCAAGACTGCAAAGGAAATACTACGAATGGTTACCTCG

TATGTATGAAAATAAGTAGCCCTGAAACCTTTATCGAAGTGTTACTCTATGAATGGATGTTGATATCTCA

ATAGATGTTTACAAAATGATAGATTCTGCTGTTCCAGATTCATTGAGATAATGTGTAAAGGGTGTACAAC

AATGACGGATATAGCTAATGCCTACAATTAGATTGATGTGTCTTAGATACCTAGGGGATTCCATACGTTA

CTCTGTTAGTGAAGACTCGAGCTTTCGAAGGGTTGCATATATATATATTATGATAATTGAAAAACTTTCA

ATAAACCAGGAAGGTAGATATACATGTACTCGAAACCTTTTCCCACCCACTTTGTCCCGAGCCACCGTTT

ATGCCCGTTTGATGGAGACAACCCGGTGTTACTAGCCTCCATCGATCCCCTGGACTCCACTCGTCTCCCT

GGACTACATCGTTGGACACTATCTTCGCCTTGTACGACACAGCGTCGTCGAGTCCGTCGTGTACTGCGCA

GCTTCGTCGAGTACATCGGGTACAACACAGCGTCGTCGAGTCCGT

>NODE_44_length_1022_cov_9.076923

TTTGATTTATTGTATTGCACATGTGCACACGTACTGTAGTGCAAGAATGTCACCGGTATGTCAACTGTAT

CCTTGCAGCAGGCCCCCCGCCCTTGACATTCGTTATTACACCTCGACATGTTTAGGGTAGCACTACAGTA

TATTTCGTACATGCTGCAACGCCACGCCGCTTTCGCCGACAACCGAAGCGAGGCACTACATCCAACTCAA

ACACGTGAGTGGCAGCAAAGAGCACAAAGAAAAGCTGATCATTGCTTGCCAACTCTCACTCGCATAGACA

ACTCTCATTCAAGCCGACACAATACATAGCTTGGAGAGCACAAATATCCCCTTCGCTGCCAAACTCGAAG

CTAGCAATATGAACTTGGCCATGCACAAGAATGTTAAGGACAGCGAACCCCCCATGGCTACCCGTATGCG

TCAGCGGGGTCTTACCTCCGACGACATATATCGTGGATTTCGGCATCAACGACAGTACGATCCCGATCGG

GGCTTGCTGTAGCCCATTTTACCGAGTACATGTATTTGTGACAAGACAAACAACATAGAAGGACACCCCT

GTTAACGCTACGGATCGACACTTCCTCAAGGTTTCGTTGTACACGCTATTTACAGTGTAACGAACCTATC

GAAAAATGATGCGTCTTCTACGTACGAAGTACTTGGCGCACTTCTTCTGCTAGATCTTAGCATTTTGGCA

AACCCTTGCTTCACAGCAGTGTGCGCCAATGAACATAAGTTTCACTCTCCCTCTCCTCAGCCTCCAGTGT

CTTGCTACAACTAGATGGAAGGGAAACACCACAACCACGTACACGCATATATAATCGTCAATCACAGACA

AACCTATCGAACCCTGCTGGAGACAAGGATTTTACATCATACAAAAGGAAACTTCCCGTCACTGAGGTAT

ACAGGTCATATCGATGCACATTCTTACAGCGTGCCCAACGCAGTGGACATTTCCTGCTACGCTAGGTGCG

CCCGATGGTGATCTCCTATCCACCACCATGCAAAGCGCAACG

>NODE_45_length_1012_cov_20.510595

CAGCACACAAGCCTTCCTCCCTCCTCCTCTTCTTCTTTTTCAAAAACCTCAATCATGGCAGGCAGGCCGG

CACCCGAGCAACCCCCTTTCCCGTGGAGGAGCGACATGCTCGCGGGAAAGAAGGCCATCGTTACTGGCGG

TTCGGCAGGCATCGGCGCTGCTATCACCGAAGCTCTGCTTAAGGCGGGATGCGCCGTGGTGGTCCTGGCG

AGGAGCCGCGCGAAGTTCGACTTGCTCGTGCCCATCCTCCAGGCGAAGGAGTTGTCCACGAAGGACCTGG

ACTTCATCTCCATCGACCTGTCGAGCACCGACGACATCAGGAGGGCCGCGGCTGAGGCGAACGAGTGGGC

GGGTGGTTGCGCGGACATCCTCGTCAACAACGCTGGGGTGGCCACCATCGCCCCTCTCGTGGACGCGACC

GTAGAGGACTGGGACTGGACCATGAACGTGAACGTGCGGGCACCGTTCATCTTGGCGCAGGAGTGCGCCA

GGCGGATGATCAAGCGCGGCACCGGGGGCAAGATCGTGAGCACGTCGAGCACGGCCTCCCTCTTCGCCCT

CCACGACCACGCGGCTTACTGCACGAGCAAGGCCGCCATCAACGGCCTCACACAGGTGATGAGCGCGGAA

TGGTCCAAGTACGACATCAACTGCAACACCGTCGGCCCTACGATCGTATTGACCGACATGGGCGCCAAGG

CCTGGGGTGAGCCCTCCAAGGGTGACCCCATGCTCGACCGCACCCCCCTCGGTCGCTTCGCCGAGCCCTG

GGAGATGGCGCACGGCGTGGTTTTCCTGGTTAGCCCCTCGTCGAGCCAGATGTGCGGGCAGATGCTTCTC

CTCGACGGCGGTATCTCCACCACCGGCACCCCTTGCGGAGGAATCTAAGTATACATCTCGCCGTGAAGGC

GTTTTGTCTCGGGGTTCTGTTTGCGGTAGATTTTGCCACGCTACGACGAGTGTGGTGGGTAGGCGCTTAG

CTTAGCCACGCCTGGTGGTGGGTGACGGGTTT

>NODE_46_length_1012_cov_11.432896

TAAACTGGCCAGGGGAAAACACTCCAAACGAACGAGTCAAGAAGCAAACTCACCCTTCAACGCCAACACC

CGCCATGAGGAGTGCTTTCGGATTCTTCGGGTCGGTATGCATGGCCATCTCTGCGTCCGCAAGTACCACT

ACGATCAGGACTCGCCCATCGTTTGGGGAAGAGAGCGCACGCATGGCATCGGCCGCGTTCGCCACGAGCC

GAAAAAGCAACTATACCGGCGGCCTCCGTCAACTGCAGACCACGTGCTCGAGCGATGCAGTCACCATTTC

AGTGCTGGAAAACCCCAGTTTTGACTCTTTGGCTGGATGCTACCAAGATACCGGGCTCCTTTTTTCTGGG

CTCCCGTACTACACTCCGTCTGGTCTGGACCTGCCGAACACGTATATCATAGGCCTGGAATTCGGCACAA

CGGGTTCCGACTCCTCTTCGACAATCGTACGGTGGACTATTGAGGACTACGAAAAGCCTGAGATACTGTG

CGCCTCCAATGAGCCCGCCGGGAGCATCCACCCAGCGGAAGCCACCTGGGGTTGCCTCAGCGATGACAGT

GTAGTCGAATTGGTATCTTCCGAGTGCGGCGGTGATTGTGAGGACAGCGACCCTGGCGAGACAACATGTT

CGAGCGAGACGGTTTTCGTAACGTCGTCGGCGTTCCCGGAGCTTGAGGGTTGCTACGAGCCAATTGGACA

AGAATCCGAAGGCCAACCCGTGTACTCCATGACAGGAGAGGTGGTCGAGGGAGAGCCACTGATGTTTGTT

GAAAACCACATGAGCTTCGACGGGACGGAGCATCCGGTCTGGAATCTGGGCCTGGCGGGGACCCCAACGG

AAGACGTCCCCTGCGCGTCTACCCTTTCTGGCGACATCGGGCACCCGAGCGACGAGTTGTGGTCATGTGA

CGTAAACGGGGACACGTACATCGACCAACTTCCGTCTGAAACGGCGGGCACACCTGGGCTATTCGAAGTG

ACCTGCGGAAACGCCGCCACCACCTGCAGTGG

>NODE_47_length_986_cov_10.649741

TGTATCGGCACGAACATGTTGTTATCGCAGGTTACTCGCAGCTGGTGTCGGTAACGAAACGTTTGTCATT

AGACTACGTGCTAGCTACGGCATGGAGCCAGAACGCACAGTTGGACGAAGCCTAGCACGATACGAAAAAA

TCAATTGAAGACGTGGACACAAATTATACAACCTTAACCTGGCGCTCGCCAACTCTCACCACGCTACGCC

ACCTTCACGGACAACACATATAAATAACGCCTCCCAACAAGAGCTTCTATTGTACAGGCCGGGATAAGTG

CCACCTTATGGAAGACTACCCCCCCCATCTCGTACATTATGACCTTGCCAAACAGTCTCCTCGACAAAGC

ATACAACGTAGACGGCCATTCGTAGTTGGATTTTATGCACCATAATTTGTCAGCGTCCACAGAACAAACG

GGAAATCTTACTCCACTGAAAACGCGTGGATGGGTAACGCCTTTGGCACTATATGTAACGCGCGCCCCCC

AGTCACTGTGATCACGTGGTGGCGTCACCTCTCTGCAGCCTCGAAACCCTGTGTTATCTGTCCACCCTCT

CGCCCGGCTCAACCGTTTTGGGGGCACTTTCCAAAATAAATGTTGCTCAAGATACCAAGAGGCACTATCC

AGCTTGTGTGGTGTTCGCATAGGGACAGCCGATATCTCGTCTGCAATGCGAACACTAACAACACACCTAT

CTATAGAACTGCCGAGGCATATACCGCACTACCCAAAGGTACCTAGTTTCTACTACACCGATGTGTAACG

CAGGAACCAGGACTCCACGACACAGTGCAGCTCAGCCTATTCCGTGACCGTAGTTCCAACACCTAGACCA

ACCTTGGGCTCCGGTGGGGACCTATCCCCATCTCCATTACCAAGATATGGCCACTCGGGCGAATGCCTAT

GTACCTTACTTTCATATATAACTTTCATCGATTATGTGGGCGCAGCATTACAAGTAATATCATGGCATTG

CACATC

>NODE_48_length_986_cov_6.895337

TTCCGCGTTGGGTCCCAGATCGGTACTGCTGGACTTGCTAGAATTTCCCATCAAGCGCACCAGCTGAAAG

CCTCTTATCCAACTACTCCCCGGCTTTTCCCGTTAACACGGGTAGGCCGAGGTCTTGATTCTGCTCGCCA

GTCGATCTTCGGCTCGACCTTCGGCTAACGTCCTCTCCCGTTCTCCATCGAACTCCACTTCACTTCCTCG

CGTTTACAGGGAACCGGATTACGAATTTCCAGCTCTCCTAGCTTTCAGGATCTCACGTTCGTTGAGTTCC

TCCCAATCTCCGTCTGAGTACCGAACCCGCCAGTACGGTGACTTGTAGTCGAAGATCTTCGAAAGGTGAC

GCTTCTTATTTCCTTCAGCGTCCGTGAATTCCCTCCAGATCACGGTCCCCACGGGGTGTTGTTTCAACGC

ATGTTCACGGTCGATTCCATCTCTTTCCTTCCGTGTTAGCATCGAAGTCGGCCTTGCCCGACACTCCGCT

CCGTGATATGTTTCCCACAGTGCATGGAAAACGTCCAGCTGTAAGGGGGTGAAACTGTCTCTCGCCTCCT

CTTCGGACAGCCACTGCGACTCCGTGCCATCCACGAATCGTCCCCTGTATCGCCACTTCCAAGTTTTGCC

TTGACCCATCACAGCCTCTCTGTCTATCAGGGTATACATAGGCGACGCTACAATCGACGGCTCCGCAAGT

CCGAAGTCAACTCCCCATGCCAGGTGAGCAAACTCGTTCTCGAAGTCGTGACGGAGGTCGTTCGGACGTA

AGTGGAACTTCTTGATGTTCGCTGCTGACGCCCGCCGTCGGCGAATCCCTCGCCCGTTCATCGTGACGAT

GTAGCTGAGTCCTGGTAAGACAACTTCAGTCACTTCCCAAGGTCCGGTCCAGTGCTCGTGGGCAAGTTTA

TGATGTATTCCCTCCCTTGCCATCACAGACTCCGCCTCCTTCACCAGCACCTTGTCTCCGACAACCACAC

GGGTTC

>NODE_49_length_982_cov_13.209157

AATTAGTATAAGAGTTATTTTTATTGATTTTAAATAAAATGTTCAGTTCTTTGATTGTATATGCTACAAG

TCTTACGAGGCTTGTACCTGTTCAAACTTTTCAGGTGTTTGGGCATGAGATTGTTATTAGCGTATCCAAA

AAATATTTGTTGGCTACATTAACTTTTTTACACCATCACAGTAATGGTAATTTTCAAATGCTTACTTCGA

TTTCCGGTGTAGATTATCCGGAGAGGGAAGAACGATTTGAAGTTGTCTATGATCTTTTAAATGTAAGGTC

TAATTGTAGACTTAGAATTAAAACACATACAGATGAAATAAATCCCCTACCTTCTGTGGTAAGTCTTTTC

CCATCAGCAAATTGGTGGGAACGTGAAATTTGGGATTTATTTGGAGTTTTTTTCTCAAATCACCCAGATT

TACGTCGTATTTTAACAGACTATGGTTTTGAGGGTCATCCGTTACGAAAAGATTTTCCTTTAAGTGGGTA

TTTTGAATTGCGTTATGATGAAGATCAGCGAGTTGTTGTTTGTGAAGCTATTGAGTTAAGTCAGGAGTTT

CGTTCATTTAATTTTCATGTCCCTTGGTCACAAAATAATTTACTAAGTTGATGTCGAGGTTTAATGTAAA

TGCTATACTTAGTTGGGTAGATTCTCATATTATTGATTACCCAACGGCTATGAATTTAAATTATAATTGG

AGTTTCGGTTCAACCGCGGGGTTTTGTTTGGTTATTCAAATCCTTAGTGGAGCTTTTTTAGCAATGCATT

ATGCTGGGGAAACTCATTACGCTTTTTCAAGTGTTGAGCATATTATGCGTGATGTCAAAAGTGGTTGGTT

AATTCGTTACATGCATGCGAATGGAGCTTCTTTTTTTTTTATTGTTGTCGTTGTTCTCACATTTAATAAC

TCATCGCCGTAAATCTACTCCGTCTGGCCACGAGGAGCGATGGCTTCCAACCATTTTGGAAGGAAAAAAG

AA

>NODE_50_length_963_cov_4.884289

TGAGCCAAAATCAGGTATATACCACCGCGCGCCACGAGCCGTGTTTTTCAGGACTTTTTCGCTGAGATGT

AGGGCAACTGCCAAAATCAGGTATACCTTTTGAGCCAAAATCAGGTATATAATAAAACCAAGTAAAAAAC

GCACGCCACGAACGGTGTTTTTCAGGACTTTTTGGCTGTGAGAAATAATCAAACATACTTTTGAATTTGC

GCCTGAACAGCAGAATGATAGCGGATGTGTGTGTCCTCCAAACTCTTGATAAATGTTGTATGTTATTTTC

AGTACGGGGATTAATATCTGTCGGATTTCGTGATCACACTCACTCCTACGACGTGTATTTCCGTATTCTT

GTACCAGGTATGCGTATGTATATACTCGTAGTCGTAGCCACATTGATAGCATACGAACAAAACCTCGATT

TAATTTGAGACTGAACTGCTCGCTTTCCGTACCGTGTCAGCAAGCAGCAGTAGTACGTAATACCTGGTAT

ATTACAACAGGTCCATTGTGCTGTATGCGCGCGTATGATACGTTCTGTAGTCGTTGTGGCGGGTCGTATG

TACGTAGTGTGGTGATGTCTTCGTGTCGGGCGTATAGGAGACAGTGTACGTATGGCCTCTGATTGGCTCA

AAATTCGCCAAAATCAGTTGAAAGCTAGACTAAGTAGACAAATCCAGGTGTACCTGACCATTTTCATGCC

AAAATCAGGTACGATGTGAGGCTGGCCAATGAGAGGTGTCCTAGCGTGTGATTTGTACTTGGTTTTAGTG

TAGTACCTGTTTTTGGCACCAAAGTGACCAGGTGTACCTGATTTTGGCTAACGCCCTTAATAAAAAATTT

AGCACGTCGCGCTCGCTCCCCATCCAACTAACATGAGGGCAGTTTTTTTGTAGCAAAAACTTGATACCTA

AAGAAGAAGAGAGGGTTGCAATACCACTACTACCACTGCCGTGCCGTGGTCCG

>NODE_51_length_953_cov_13.815451

CTGCGCCGCATGCTCGAGGAAAACCAGGACGCCATATGCACCGCCCTGGCCGAAGACCTCCGCAAGCCCT

TCGCCGAGGCGGCGCTCGGGGAGGTGTACGACGTGTTGGGGCAGACGTCATACATGGACAAGAACCTGGA

GAATTTCACCAAGGTCGAGAAGATCCCGACGCCGCTGCACCAGAAGCCGCTGTCTTTCGAGGCGCGGAGA

GAGCCGTTCGGTGTGGTGACTGTCATTGCCCCGTGGAACGTTCCGGTTGCCCTGCTGCTGAGCCCCCTGG

CCTCGATGCTGGCTGCCGGGAACGTGTGCGTCTTGAAACCGTCGGAGCTCAGCGCCGCCTCCAGCCGCCT

CTTGGAGGAAATGATCCCTAAATACTTCGACCCTGAGGAGGTGACGGTCATCACTGGAGGCGTTCCCGAG

ACCGCCGCTTTGCTGGCGCAGAAGGTCGACATCATCCTGTACACGGGGGGAGGGGAGGTCGCCAAGATCG

TCATGGCGGCCGCGGCCAAGAACCTTACGCCGGTCGTTCTCGAACTGGGCGGCAAGTCTCCCGCCTACGT

GGACGAGAGTGCCAACATCCAGGTCGCGGCCAGGCGAATCGTGTTCGCGAAAAACATGAACGCGGGCCAG

TTCTGCGTCGCCCCAGATCACATCCTCGTCCACGAAAAAGTCCGCGACTCCTTCCTCGCCCACCTCAAGA

CAGAGCACGAGAGGCTCTACCCGGGCAACGAACAGGACAATGCGGATAAGGGAAGGATTGTGAACCAGCG

CCACTTCGAGAGGCTGCAAGGCTTGATGAAGAACCACGGAGGACAGGTGGTGCAGGGGGGGAAGGTGGTG

GACGCGGAGGAGCGTTACATGGAATTCACCGTCATCACGAACCCTGACCCGGGGAGCATGCTCATGAAGG

AGGAGATATTTGGGCCCATCCTTCCGGTGATCACGGTCAAATC

>NODE_52_length_941_cov_20.528261

AATTTCTAACCAATATAAACCAAATAAGATTTTAATAGGAATGAGAAATAACTTTGTTATACTTATAAAC

AAGAAAGGTATAATATATTTTTATAAAATGGAGAAAGTTTTAAAATGTCTGGTGGTTCAACAGGGGAACG

TCCTTTTTCTGATATCATAACAAGTGTACGCTATTGGATTATTCATAGTATTACAATTCCTTCTTTATTT

ATTTCTGGTTGGTTATTTATAAGTACTGGTTTAGCCTATGATGTTTTTGGAACTCCTAGACCTAATGAAT

ACTTTACACAGGACAGACAACAAGTTCCGTTAGTAAATGATAGATTTAGTGCTAAGCAAGAATTAGAAGA

CTTAACAAGAGGATTATAAAAAAATATATAAAATTTAGGAGACATACGTGACTAGAAATACAAATCAACC

TGTAGCTTACCCTATTTTTACTTTTCGTTGGCTTGCTATACATGGTTTAGCTGTTCCAACGATATTCTTT

ATAGGTGCAATTACATCAATGCAATTTATCCAACGATAGGAGTTTATTCAATGACAGGTCCAAATCCTAA

TAAGCAAGAAGTTGAAATAAGTCGTACATCTTTATACTGGGGTTTATTATTATTTTTCATATTAGCAGTA

CTTTTCTCTAGTTATTTCTTTAACTAAAGAATTTTGTAATTACTAGGTAGAATTTTTATAAGGTTAAGAA

AAGATATAGGAGCAAGAGAATATTATGGCAGGAACAGGAAGAATACCACTTTGGTTAGTCGCATTAGTTG

GTGGCCTAGGAGTTATAGTCGTTGTCGGTACTTTTATTTTTGGTTCTTATTCTGGGTTAGGGTCTTCACT

TTAACAATCAATTACTATGGGCCTATTATTTATCTACATTGAAAAAGACACTATTGAATAACTTTCGAAA

TTTCCAAGGTTCTCTAGAGAACCTTATGGAA

>NODE_53_length_940_cov_8.952122

CGCACGAGGCGAGTATGATTTTGTTGCTTGCTGCTGATGATTTACGTTCTTGGATTAATTTTATGTTGAT

TTAAGAATGATGAATACCCCCACATTTGAGATTTGAGAATAGAGCATGTTTTGGAGATTTTGAGGGTGGA

ACGGAGGTATGACACACAGGCGTATTTGTATAGCTTCTCTCACTGAAGTTTTCCTGTAGAGACTGCTGTA

GCGTACAAGTACTACTGCAGGACAATGGCTGTCTGTTTTGTGGTTGCCTGATCGGACACTCTGGCTGGGT

ACCAATTCACGTACGAGTAAGACGTAGATTTTGAAAGTGCAATATGAGGAAATTATGAAAGAATGAGTTG

AACTTTCAGCGCACACACGTACATCATCGGTACTGCTGCTGTATCGTATCAACACAAACCCCGAAATTGA

ATACGAGTACGAAAGTAGTAGTAGTAAAGTACTGCGAGTACAAGGGTATACGCAGAGTTGAGTACGATAC

TACAGTACATCGTATATATCTGGACTGCAGAGTAGCACAACCCCCTTTGCGATATGTCCACCCTGCGTTG

TCGTGCCTTGTCATGCACAGAGCAGGGGGGAGGGAATTTGTATGGTCGTATGTATTGAGCAGGGGGGATG

TTGACTGTGCTAAGAGATACGACGGAGATGATCCATTTGGTAGTAGGGAGTTCCATGTATTGATGAGGCA

TTATGTGTGAGGGGAACATCTACATGGAATCAGCGGGTATATTGTCCGTTACCCTGGTATGTACTGCTGT

TGATGGTAGAGTTGTGAGTATGAGCTGAGGAAATACTTTGCACGGTACTGCTGTACATGTTCACCAAGTC

GGAGGATTGAGCGAGATACTACGCATACGTACAGCAGTATATAGTGATCGGTAATTCATGGATGCGACAT

GGTATAGGATATCATGTGGTATCATCGGTA

>NODE_54_length_918_cov_15.914158

GTACAGTCCGCCGAGCTTGCCTATAGTAAAAAGAAAAAGAAAACCAATTGAAAATAACATAGGGGTTTTT

AAACGAATAGAACCCCCCCATAAAGTAGCAATCCAACTAAAAATTTTAATACCTGTTGGCACCGCAATAA

TCATGGTAGCCGCTGTAAAATAAGCTCTTGTGTCGATGTCCAAACCTACTGTAAACATGTGATGAGCCCA

AACAATAAAACCAAGTATACCAATTGAAAGCATAGCATAAACCATACCTAAATACCCAAATACAGGTTTT

CTAGCAAAAGTAGATAGTATATGACTAACAATACCAAATCCAGGTAAAATTAAAATATAAACTTCAGGGT

GACCAAAAAACCAAAATAAGTGCTGATACAAAACAGGATCACCACCACCAGCCGGATCAAAAAAAGTAGT

ATTAAAATTCCTATCGGTTAATAACATTGTAATACCACCTGCTAAAACAGGAAGTGATAACAGTAATAAG

AACGCTGTGATTAAGACAGACCATACAAAAAGAGGTAACCTATCCATCGTCATACCAGGTGCTCTCATAT

TAAAGATTGTTGTAATAAAATTTATAGCTCCCAAAATAGAAGCAGCACCTGAAAGATGGAGACTAAATAT

AGCTAAGTCAACAGAAGGTCCTGAGTGTGCCTGAATACCACTAAGAGGTGGGTACACCGTCCAACCTGTC

CCAGCCCCAGATTCTACCAATGAGGACGCTAGAAGCAAGATTAAAGAGGGGGGTAATAACCAAAAACTAA

TGTTATTCATACGGGGGAAAGCCATATCAGGAGCACCGATCATTAAAGGTACAAACCAATTACCAAACCC

ACCAATAAGAATTGGCATAACCATAAAAAAAATCATTAAAAAAAATCCGCTTACAATGAACACAAGTCAC

ACCAACAT

>NODE_55_length_904_cov_6.908267

TTGACAACCCCAGCGAGTACGGACCCTTTTGCGAGCTCCCATGGTAGTCATCGTCGTGGAAAATCCGTCA

CGGTTGCGCAGTCGTCGGACCATCCTGTGACCACGTATATATTAATGTGTAGAGGGCTTGGAACCTGCGG

GATCAGGCGCGGGGCCGGCGGAGGGCACCGTATGTAGAAGTGCTTGTGCGCAGGAACAAGGGTGTTGGAA

AGTTTTTCGTCGTAGCGCTGGCTCAAGAGAGGAAGAGGGGTACTAGCAGCCAAACCGCGTCTGCTGTGGG

ATCGTCCCACCGAAATTGGACCTTGGCGCGTTACCACATCCGCACCAGTTGTTGGGTTGGACGCGATATG

TTTGCTGCTTACTGCGACTGCTGCGGTGGAAACGCAGCTACGAAAGGCGACGGCAGTAGTATTTGTTGTG

TACATGTGCGATATGACGAATCTCGTGTTTTTTTGTTGGTGAGAGGGGCCGTCATTGTCAACGGCGGGCT

TTGTGTTGGACGGCGCGGACTGGTGGTATGTGACTTTTAAGTGTTGTGCTTCGTATTGGATTATAGATTA

TAGATACGAGGTACTTGATATTGGTCAGCAGTCAGCAGTAAGCGGGCCCTGTTTAAGGCAGTTCTGGATT

GTACAGTACCAGGGAGGGACTCGCAAGCGTGTGCATCCTTCATACACTGCTGTAACCGCCTGCTTTCCTG

TTCGCACGGCTCGTAAGCGATAGGCGACCATTACGCGGAGCGGTTGTACTATCATTGAAGAAGGATATCG

GTATTAACTTGGGCCCCAAACGCCAGGCAGAGATGGAGATATCGCTGTACGTAGTCAGGTGGAGCAGCAG

TGCACTCACGAAAGGAAGGAGATGTTAGTGCTCGAGTCGGGGTGAGCCCTCCCCAAACAACTCG

>NODE_56_length_894_cov_35.605956

TTGGCTCCCACAACGATGAGGGGTAACGCAGATTGAGGAACGAGCACAAGACCCCACATTTGAAGAGTGT

CGTCGTTTCTCTTATGTATGTTCTCCCTGGTCTGGTTTGGACTTCGTTCCCCATCTCAACAACTGGGATG

TACGAGAGAGGTTGTTGTTACTGTTGTCATGGAAACTGGTTGTTGTTACCTTGTTTGTTCTTTCGAGTCC

AGCGCTTCTCCGTGTGGACTGTCGCGCATTTTCCGTCGGCTACAACAACAGCGGCGAACCACAAACCGAG

AGCGTTTGGAGGGAGAGAGAGGGCGGGAATAGAAAGAACGGGCTCGTCTCGTGTAAAATGCAAAGAGCAC

GAGAATAGACTCAATGATGATGTGTGTGGGCGGTACAGTATAAAGAGTGGACTCATTTGACTTTTAGGTC

TTTCATTTTCTTTTTTGTTTGAAATTTGGAATTTTCTCATCATGTGGTGTGCACCCCGCCCGCAAAGAGG

AGATAACTAGACTGCGCTCAAGAACGGAGGCGACCGATCGTTCTTGCATAGCGCGTGTTCTTTATTTTTG

TGTTTTTAGGGCAGAGCTATTAAACCAGCTACCAGCTTCGCAACTTATCGTACTTATGTTGCGGCGGCTG

TTCGTTTGTCTATACATCAAAATATGCGCACACCTGGGTATAGATATATGTGTGTACGTATAGCATACGC

GGGCCTCCTCTTGGTCAGTTGGTGCTGGTTTGTAGAAGAAGAGCGGATTGCATGTTTTGGCCTCACTAAC

TTATATAACTTAATAGTTTTTTTTTGTTATCCTTGTCGTGCCACTCATCATGCAGTCAGTAGCAAGGAAG

TAGCAATCAGCAAACAAACAAGAACGTAAACTTAAAAAAAAAAAAAAGTGGAAT

>NODE_57_length_878_cov_63.110852

AGTCACGTGACTGCGTGGTTCAGTGGGTAGTGCTATATAATTCCCTAGGGAGGGTCGCGGGTTCGATTCC

CGCCGATTTCATAATTTGTTCTTTTCGAGCGTGCGCGCTGGAGACATTGTGAATGGTTATGATAGTGTGC

TCCTGGACACTAGTCCGTGGCGAGGGGAGTGCCCCGGAGATCAATGCAGCATGGTGAGAATCAACGAACC

CGAAAGTATATATATATATAAATATTTAAAGGTCACCTAAAATAATTAAAAAAACTACCCGTTCCAATAA

TTTATTATTTAAAAATAAAGCTATTCCTCTTATTATTAAAGGTATATCTATTCCAGAACCTAAACTTTAC

GTACACTTTACCCAGAAATAATACTTTTATTTACACAGGTGTTGAAAATATTACCAACATGCTTTACGTA

AACCTGAAAAAGAACCTTTTGATGCTAAACGTCTAAATTGAAGACGGGATAACTTGTAAAAACTTATAAC

CGACCTTGATCTATTAGTATCAATGCAATGGTTGTGAACTCTACTTAAGCTACTTTTTCTAGGTAATTTA

GATTTTTCTAATTGAGCCCACCAACGTAAAGAAATCTCTAAATTTTGATTTGTTGCTACGGCTTGCAATG

CTTTACGACGAGACTCGTATAAAGCAAAAGATTTACGACGTTTGTAATCCATACCTCTCATTCCCAATCT

AGACTGCCAATTTGCCAAGCATATACAAACCCAATAACTAACTCAAAAAGAAAATCCATCATGGACCAAT

ATCCGATTGAAGGCAACTCACTCAAGGAAACTGCCCACGGAAAAAGAAAGACGGTTTCAATATCAAATAA

AAGAAACAAAATAGCAAGAATAGAAAGAATAGGTGGAA

>NODE_58_length_872_cov_9.937720

CACTTTCCGAAGCAAGGGACAATAATCGTGGGGTCGAGAATCTTCAAACCTCCAACTCTACAAGTCCAAG

ACGTCTCAGCTACAGGTAGTGCAGTAGTCTGCGCCAGATGCAGGATATCGACGAATAGTGTCATGATTCG

AAACTCCAACTCTACAACTCCAAGACGTCTCAGCTACACGTAGTGCTGCAGTCTGTGCCAGATGCAGGTG

TATATCAAATAATAGTTTAATGAATTACCGGAATGCATTGTCCGAAGGTATCCGACCTCATATCAACCCC

CTGTACCGAAATGGCACCCTCTGCGACATGCCCTACGGTGGTGGCAGTAGTAGTAGGGTGCACCTAAGCC

CTTAACCCCTCTGATGCTCCTCACTTTGTTATAGTGTAGAAGTTCAATGCCAGGCCCAAAAATGGACTTT

GAAAAAAAATGTACGCATAACCCGACGAAACATCCATCGCACTGCCCAAGCATCACCCATTGTAGGTCAA

CACGCTATCTACGGCTTGCTTTACACGCGCAGATTAAACCACCCCACCCTCCCGACTCACTCGGATTACC

AACATTGTTTCCAGCATACACGACGATAACAACGAGAACACGCGCAGTCGCAGTTATGCCCTACTCAATG

GGAACATCCGTGGCAGGCGCTGGCTACGTTTTTTACAGGCACGACGAGTTAGGTGAGTGCCAACATCAAC

ATCCGAACTATGCTGGAGTATGTACGAACTACAATACAGTCGTACATAGAATATACACACACGACGAGGA

GCATATATCCTTCGATCGATTCAATACCCTGAAGATTCCTGATCACTTCATGGCATTTACGCGTTCGTCG

ACTCTATATGATACACAGCCTTCAGACTGCCG

>NODE_59_length_859_cov_12.847255

GGCGAAGGTCACCACCATGCCGGGCTTGAGGGTGCCGGTCTCGACACGGCCCACCGGCACAGTGCCGATG

CCGCCGATCTTGTACACGTCCTGGAGGGGCAGGCGGAGGGGCTTGTCCGTGGGGCGCGTGGGCTCCTTCA

TCGTGTCCAGCGCCTCCAGCAGGTAGGGGCCCTTGTACCACGGCATGTTGGTCGACTTGTCCACCATGTT

GTCTCCGGCCCAGCCCGAGATGGGCACGAACGGGATCTTCACGGGTTTGTAGCCCACCTTCTTCAGGTAG

ATCGCCACCTCTTCCTTGATCTCCGTGTACCGAGACTCGCCGTACATCACCGACGAGTCGTCCATCTTGT

TAATGCACACGATCATCTGCTTCACGCCCAGGGTGAAGGCAAGCAGGGCGTGCTCGCGGGTCTGCCCGTT

CTTGGAGATACCGGCCTCGAACTCTCCCACTCCGGAAGCCACCACGAGCACAGCCACGTCGGCCTGCGAG

GTTCCCGTGATCATGTTCTTGATGAAGTCGCGGTGTCCGGGGGCGTCGATCACCGTGAAGTTGTACTTAG

GGGACTCGAACTTCCACAGGGCGATGTCGATCGTGATACCGCGCTCGCGCTCGGCCTTAAGGTTGTCCAG

CACCCATGCGTACTTGAAGGACGACTTTCCGAGCTCGGCGGCCTCCTTCTCGAACTTCTCGATGGTACGC

TTGTCGATACCTCCGCACTTGTAGATCAGGTGACCGGTCGTCGTCGACTTACCGGCATCTACGTGCCCGA

TAACCACCAGGTTGATGTGCACCTTCTCCTTGCCCATTTTTGCTGGTGGATTTCAAGGCGAAGGGTCCTA

GGAGGATGGGATGGGTTGA

>NODE_60_length_848_cov_21.735187

GGCCTACACCGGCGACGACTTCTGCAACGAGGTAACGGAGGACGGGTCCAACTCCTTTGAAGACCTTGTC

CAAGAACACGCTTCGTTCTCGTTCCGCGCCGAGCTATGAAGGACTTGACCGAGTCACAGGAGCCGCGTAG

AGTCCGCTCTGTGTTATCAAACCATGCTCACCTGGTGCGCTGCTTGTTCCGCGTGGTACTCAATGTGTCA

TGCGACGAGACAAGTGCTGACAGGAGGGGGACGTCTGACTCCAAGCATCTTCCCCTGCCGGGAGTTCTCC

ACTAAGGCACTACCCAAAACTTTTGGAATCCCCCTTTTGTATAAGCAATGTTTGTCGTTAAACCCCTGTA

TGTGAAGCCGTGGGGCTGTTCATTGAGTCGAGCGGTACAAAGATTCTCAAACCGTGATTTGGGCGGACCA

TATATATCATTTGATGCGAACCTCCAGCGCAGTGCAAGGTCCACCCGCTGCCCAGCTGTGTCTAGTTGTA

TGTGTGTGTGGTGAAGGTCGGTGTTCAAACCGCGTTAGCCAGTGTAGAAGCTGTAATGCTGTGTATTGAT

GTGGGATACATATAAGGTTGGTAATCGCGTTCCAATTGTTATCATGCGGCCTGTCGCGGGGTCCAATAGA

TCGTTGTTCGGTTTTCATTGCGTACGAGAGCTCTGCTGCCCGGTGAAGGTGCTAACGGTTAACTGAGTTT

TCTCACCGACTATATTGAGCCTGGCAAGAGGGAGAGAGGAACACAGTACGTAGTGAGTCCGTTAAAAGAT

TAAAGGCGACGCGCCAGCGTACAGAGATTGTTTCAGAAGGGCGACACACATGGCGTGCCTGCGACAGTGT

CCAGCTGT

>NODE_61_length_845_cov_27.264563

GGTAAAACAAGGGCCGTCGCAGATGAGACTCAAGTGCTCGAAAGCAAGATTGTCGGCGAACTTATCGTCA

TTCGGCGCTACCGATTTGGCTCGAAAACCAACGACCAAGCGGAACTGCGACAGTACCTCGGCGACATCAA

CGTCAGCACTACCATTCAAACGTGGAAGGCTACGGACATCGTTAGGCGTGAAGGTCACAGCCTTTGGTGC

GGACAGCACTGCTGCATACGAAGGGACAGATTGGACAGGGGGGACAGGGTTTGAGTTCGACTCGGACAAG

GTGACAGACGCTGCGGGACTACTGGTAACGCTTAGACTTGCCAGTTGTTGAAGCATCGAATTGTGCTGCT

CCGACAGCACTTCAATCTCGCCTCGCTCCATCTTCTCCAGCTCGCGGTGGACGATGCGGGCCAGATGGTC

ACGGATGGAGGCGATCGGAGCTCCAGGGTTGAAGGGTGCAGGAGGATCCGTGCTCCCGTTCTTGGCTTCT

TGAGCTCCGCCGGATGCTGTTGGTGTCTTCTCTGCTGCAGAGGTGTCGACGATCTCAGATACTCCTGACA

TGATTGCTCCTGACAACTTTAGAATCGACACTGCTGTTGATGATCAACTGCCGTCCGACGCCAATAAGAT

TCTGTAGTCGTCACCGCCGGTGGGTCGTGATCCCACTCCTCTCTGCGTGAGCAGAGTGTTCTAACTGGTA

AACTACGACGGCTGGAGTGTGTGACAACTCGGAGTGAGACAGCTATCCGATGACAAGGGAGAGAGGAGAA

GTCGAAGACAAGTAGACAAGTGAGAGCGAGGGGCGAGCAGTTGTTGTTGCTGCCGCTGCTGCTGCTGTTG

CTGCA

>NODE_62_length_844_cov_27.179830

GGGAGGACAAGAGGGTTTCCTTCCAGGAGTTTCCCGCTCTCAAGCCGGGTCTCCCGAACGGACAGGTGCC

TGTCCTGGAGGTTAACGGCAAACTGCTCCCCCAAAGCGGTGCTCAGCTTCGCTACGTCGGAAAGCTTGCC

GGACTGTACCCTGAGGACCCTCTTGAGGCAGCGTTCGCCGACGCGGCTGTGGATGCTGTCGGTGACTGCC

ACATGAAGCTGGCCCCCACCGTGCAGGAGAAGGACGAGGACAAAAAGATTGAAATGCGGCAGAAATTGGT

CAGCGACTTCCTTCCCACTTGGCTGGGAAACCTCGAGAAGGCCCTGGAGGCGGCTGGGGGCAGCTACTTT

GCCGGCGGAAAGCTCTCGATCGGCGACATCGCGGTAGTGGCCAGGCTGGTTTGGCTCAACGAGGGCAGTC

TCGACGGCATCCCCAGCACCATCGTGGACGAGTACCCTCTCCTCTCCGCTTTGATTGAGCGCGTCAACGC

CGAGCCAAAGATCGCGGCGTACCACGAGAAACGATCGGAGAGCAGCACCAAGTAACATCACCAGCATCGT

GGCTGTTACAGTAGTTATGGCATCGAGATAGCGCGGCGATGTAATCAAGGAGGGAGCTAGGTATTGAAAC

CTCTTCTGCGTATATAGTGTTGCGTGTCGTTAGTTAGCGTGCCCGCGAGATCAAATAGGAACATGGCGAA

GGAAACGGATGTATACTCGGGAAGGTGCCCGAGCACCCCACAATATACGCCATACGTTAGTAGGGAATGC

CCTCCGGGGTTACGGGAAATTAGGAACCGTACCCCAAACCGACACAGGTGGTTAGGTAGAGAATACTAAG

GCGC

>NODE_63_length_844_cov_6.859052

ACCACCACCACCACCACCACCCTGAGGAAGGCCTCTACCTGGGGCTCACAGCCATCGATCAGTACCGTCA

GGGATTTCTACATAACCACCGAACTTGTTGAGTGGGACTACGCACCGACGGGCGATGATCTGATCGATGG

CGTCCCTCTAGAGGACAACGAGAATGCCGTCATCTTCATCGAGGACACACCGTCGCTCATTGGACAAAAA

TACGTAAAGTCCCTGTTCAAAGAGTACACGGATGGCACATACACGACGGAGAAGAAAGCGCCGGGGTGGG

CGGGGAACCTCGGGCCCATCATAAGGGCCGAGGTCGGGGACACGATCCGGGTCCACTTCAAGAACATGCT

CCCTGACCTCCCTGCTTCCATGCACCCCCATGGAGTGCTCTACACGCCCGAGAACGAAGGGGCCTGGTGG

AGGCTCAACAACTTTCCGGGGTCTTCTGTCGCCCCCGGAGATAGCTTCGTTTACGAGTGGACCGCGCGCG

AGGACTCCGGCCCAACCGGGGACGACAAGAGCAACCTCTGGTTCTACCACGGCCATGTGGAACCAGTCAT

GGACATGTACTCCGGGCAAATCGGGCCGCTGATCATCTACGAGCAGGGGCTCCTGAATGGGCACGGACTG

CCCACGGACGTCGACGAGGAATTTGTGATCCTGTTCATGGTGTCTGACGAGAACCAGTCCCACTACTTGC

AGACCAATATCGACATGTTTGGCGGAGACCCCGACGACGCGGACTTCGAAGAGTCTAACTTGATGCACTC

GATGAACGGGTTCCTGTACGGTAACCTGCCCGGCTTGGAGATGACGGAAGGAGACACCGTGCGCTGGCAC

GTCG

>NODE_64_length_842_cov_42.300853

AATTTTAGCGGGTATTTTACTTAAATTAGGATCTTATGGTTTTTTGCGCTTTTCTTTAGGTCTTTTTCCT

TTAGCCTCTGTTTATTTCACCCCCTTTATTTTTAGTCTTAGTTTATTGGGTGTTGTGTATACTTCATTGA

CTGCTATTCGCCAGACAGATTTAAAACGTTTGATTGCTTATACTTCGGTAGCTCATATGAATTTAGTAAT

GATAGGCTTGTTTACAGGGACAGTAATTGGGGTGGAGGCAGCTATTTTACAAAGTTTAAGTCATGGTTTT

GTGTCTTCTGCACTTTTCCTTATCATTGGGGTACTGTATGATCGTTGGCACAGTCGAGTAGTTAAATATT

ATGGAGGCCTTACGCATACTATGCCAATTTTAATAATTATTTTTCTTTTTTTTACTATGGCTAATTTAGG

TCTTCCCGGCACATCAAGTTTTATAGGGGAATTTCTTTTATTAGTTTCGGCTTTTGAGGCAAATAGTACT

GCTTGCTTTTTTGCAGCTACTTCAATGATTTTAGGTGGTGGATATTCTCTTTGGTTGTTTAATCGTATAG

CTTATGGTAATATTAAAATGGTCGGTCTTGATTTAAGCTTCCGAGAATTTATGATTTTTTTACCTCTTGT

TTTAGGTACTCTTTTTATGGGTATTTACCCTAATTTATTCTTTTCTGCAATGCATGCAACAGTTTCAAAT

TTGGTATGGGTTGATTTGTGGTTGTAGATTGTAGTAGTGAAAAAGTTCTTTTAGTCTAATGTGTAGTTTA

ATATCTAGAAGGATAGTGCCATTATTTATGGTAAAGGTACGGGTTCAAGTCCCGTAAAGGACTCCATGGA

AT

>NODE_65_length_840_cov_6.433455

CTGGCTGAGCATCACCGAGGTCCAGTTCTGGGTGGAGTAAAACCTCCAAAGATTCCTGCTTCAGGGCCTG

GACAACGTCAACAAGGGAAAACACCATGGTTCATCTGAGGTTACTCTGTATGTCGTCGGCTTGTGTCCGG

CACATAGTTTGTTGTCATTGTGAAAGGAAGATAGAGCATAAAGTTGGATGTGACGTGCGAATCTGGCTGC

CGATGGGTTTTGTTCGAAGGAATGCACACAACAGCCCCACATATAGTTGTGTGTTGGCGTCTGTATCTGA

TTTTAGTGTCCCATCAGTGGCTGGAGGTGAAAACTACGCCTAGAGCGGTGTCGCTGTCTTTGTAGCTGGG

GTTGTCACAGAACGAGTACCGTCAATTCAACATATATCCGGGGAGAACTTTAGTGCTATCATGGCGGAGT

CGAGACAACCGCCCTAAAGGTCGCGTGTTTGAGATGATAATGTCCGGAGGATGGGTGACCCGTTCCAGGG

GATACCTAACGAACATGTCCCTGTGGCATATCCAGCGAACAGTTGGCTATCGTAAAGGATGTAAAAATGG

CGTGTGTGTAACATGTTGAGGGAGGAAGTCACCGCAGGTAGTGCGCGTGTGTGGTGTTCAGTACTATTGT

CGCAGTCAAGGGGAGGAGTGCTTGGTGCGTAGTCGGATGAAGTATAGACTGTATCGGGGTTTCGATGTTC

ATATGTGGGGCTCAAACTGAGAACATAAAATAAAAAAAAATAACAACTAAAAAACTAATAACAACATCAT

ATCTAAAACTAGCGAACCAACTAAAACAAATAACAAATAAAAAAAGTATGAATTCTACTGATACAAAAAA

>NODE_66_length_813_cov_15.582071

ACTCTACTCTACTCTACTCTACTCTAGGCGTAGGCTGGGGAGACGTGATTGAAATTGGATCTCAGTGCGC

AAGAAGGGTCGTCCTCTCCTCCTCCGCTTCACTCCCCTGATGGCGATGTCCCAAGTGGCACCATGAAGGC

CGATGACGTGTAGAAATCCCGACCTAGAGGTGGTTTGAGCCTCAACTTGAACTTCCAAGTTGACCTCAAC

TTGGAACATCAAGCCTAAGAGCCGGTCGAGCCTCAACTTGAACAGTTCCAAGTTGGTCTCAAGCCAACTT

GAACTTCCCACAATCGGTCGTACTTTATAAAGGCAGTAGCATATATATACGCATTTTTGTTTTGTTTTCT

GATTTTGTATTTTTTTCCCCTGTTTTTTGCTCATTGTTTTTCAGAGAAAAGAGAGTACACAGAGTAAGGT

AAGGTAGACTGCAGTTATTTGCCGCAGCTGTGGCTCTCGTCCCGTCAATCCGGGCCGGCCACCACGAGAC

TGTCACGAGACTCTCACGGGAATGGCAGTAGGGCCACCGCTCTCGTCTCACCGGGCGTAGCCGTCGCTCG

GGCGAACGCGCTTTCAGGTACATATCCCTTTAGGCCTTGAAAGGCTGCACTCACTCCCGCTGTGGCGACA

AAAAACTTGGAATTATAGTCAGATTCGGGTTTAAAAAACAGCAGTGCACTCTTAGCAAAAAAAGGGTTTC

CATAGTTGACGGAGGCGGGCGATGGGTTTCTACGAGGGGAGAATTCATTTTTTATTTATTTTTGTTTTTG

CGATGAAACGGAGTTGGAATTCTCTCGTGCCAAGGAACTCCAG

>NODE_67_length_813_cov_7.138889

AAGCGGATAAAGCAATTTACGCCAAACAGTCTGATTGCGTCGACTTCCTTGACCCATCATGTCGAAGCTA

ACAACCATTTAACTGACAAGTGTACAACGTACCCCACTGTATGCCCAGCTACCAAGTTCTTCAGCCGTAA

GACCAGCGGAACCAGCAGAACAATCTGCATGGCGGTCACTCTCTACTACATCCTGGATGTCGTCTTAAGG

AACAACTGACGAAACTGAATTGATAATCAGACGCACACATAGTCGATGTGGAGCAAATATCAAAATACCA

ACAAGAACGGTAGCTCAAATCCAGCAAACATCTCAGCGCGTACCATACATATGAGAAGAACCAGGTATCT

AGCGACCCCGTCTGTGCAAACCGACAACTGATATATAGTACGGAGATAATATCTGAATTCGTACCAACAG

GGGGCGGACACGTGTATCTCCGCGCCACATATGCAAATATGTATCGATACAGAGACAACGTTGTTGGAGA

GGTCAACGCCAGTCAGATGACCCAAGTCAAAGGCGCTCGAACGATAGCTATAATACCATGAATTCCTGCA

CACTTCTTCAAGGAAGCTGTGTGTACGATCACAGAAGGTTCACGTGTGTGAGTTCGATCAAAGCTTATCC

AAGTGAGCTCTCATAACCCCATTAAGTGTGCGTGGTGGACGACTTTGCACACTGGGGGTGATGTCGTACT

CCGCACCAAACCCGCACTTGAACGCCTTATGTCTTCAGAGCAATTGTTGAGGTCGTTCCATTCTTCAAAT

CTTATCGCCCGCAATTAGCCGTATCAGTGATAGTTGCAGCACT

>NODE_68_length_807_cov_4.769720

GTGGTGTGCCCATCGTCGGGTCACAAGCTGCACCTGGTGCGCAAGGGAATGTGCTCACGGAAGATTGCGC

GTGAACGGCAGCTGTAAATCGTGGCTTTATCTGTGGTCCCATTTCCGTCGCATTTTCGGTGTAAGTCGTG

GCTTTTATCCGTGGTTCCGTCGCGCATTTTAGGCGGTTTGTTTGTTTTCGTTACGAGTTGAGTCAACAGG

AGGTTTGTTCCTGGTGACATGAACACCGTGCGGTTTAGCCGATACGCTCGAGGTTTAGCGGTCGACTCAC

GTGCTGGCTGACAGTTTCAAGTCTTGGGGCTTTATGCAGGCAGTTTTATGTAGGGCGTGGGGGGGAGGCG

AGAAAGGTACGCAGTAGCGCCCGATCTTTCTGTACCTGTTATTGGTTGGCGCGCCAGTGCAACCATCAGG

TGTGCCAGAATGGCCCCGGATGATCGTTGTGGAGAAAGGCTCGAGGATCCGTTGCGTGCAAAGGTCTTGC

CTGTTGCCGGTATATGCTACGGGAGGTTTGCAGGAGGTCTTCGCCGGAGCCATGCCGGTGTGTACGATAG

TGAACGCTGATTGGTGGGCATGACAAGGTTTTCGTTTTTGGGGGGTGTGGTGACCGCCACGCTGTCACAA

CGGTCTCTTACCTGGTGCTATCGTAGGGGACGTTTGGAGGACCTCCAAGTTCCACGCTGGGGTCATGCTG

AATGGTGGGCATGGTACCTAATTTTGTTTGCGTGGATGGTGTGGTTGCCGCCACACTGCTGCCACGGTGA

AGGATTAAGGCAGAAAGCTTGGAATTCTGGGTGCCAA

>NODE_69_length_800_cov_5.890886

CGGCCAAGCAAGGATCAAGAAAATCGCAGCGCTGCAGACAGCAGTACGAGGTACAGCCACTTCAACCAGT

TGCATGGAACACGAACGCAGGCACACGCACCCACGCACGCACGCACGTATACAATAACCACTTACTCAGC

GGAGAAAAAAAAGAAACAGTAAAAAAACGCGGCCCCCATTTTTTCCGACGCGCACCATCCTACATACAAA

GAAAACCCCGAGAGATTTTGTTTTTTGTTGATAAATTCGCCACGCAAAAAGTGCAAATAAGCAAAACAAA

AGTCCATTAATCCCAACACACCAGCAGTATATAATAATAAAAAATAGCACAGCTGGGCAAGTTCAGAACA

GAGACTTTCCCTCCTCCCTCCTTTTTGTTGTCATCATATTGATTTGCCCCGTTCGACTTGTTTCTTATCC

CCCAACCCCTCCACTCCGAAGACGCGGGTTTTCCAATATGTTCCACGACACGCACCCAGGAGCGCAGAAG

TCCTTTACAAGACCCCCCCCCCGTCTCCATTCCCCTGAGAAATCAAGCCCTCCGGCACAAGCATTTTGTT

CCGAGAACAAAAACGAAGAAAAACAAAACGAACCAAAACGAAACGAAACGAAAAAAAAAACATGAATCAA

ATTCAATTCCCTATAACCTCAATCCTCCCCTGCCAAGTCAACGCCGCACGTCTTTACTTCTGCCTGTCTC

TCTCTCTCTCTCCCTCTCCGTCTGTCTCGTCCCCCCCCTATAAATATATGATATATTTATAGGTTGGGTT

GGCGTCGGTTGGCAACGACGCTGTAAAACA

>NODE_70_length_799_cov_6.993573

TTGGTTGGCTAGATTTAATTGCAAAATTTTTTTTTTTTTGATAAAAAGTGTATTTTATGGTTCACAGCGA

TCCCTGAACGACGCCAAGCCGCTAGCCAGAGGGACCCTCCTAGCAGACAACCATTTTCTCTCCAAACCGA

CGTAGATACTTCAAAGCCGCAAGTGTTCCTCGCGCCGTTGACAAAATAACAAAACGGCACATCAATGCAA

CTCAATCCATCCAATGCAATCCATCCAATGCATGCCTTCCAAGAAAAATCAAGGAAAGGCACACAGTTCA

AGGCAAGCAATACAAGGGACACTCAATCGGCCTTGCAGTCTAAGCCCAGCCCAGCCAAAACATCTTTTCA

ACTTTTGAGCACCATCAACTGCCGCCGTCGCTACCGGTACGTCCGGGAAAAACAAACTTACTACTATATC

ATGCGGGAAGATACAAACAATTAGGTACATTCATCCACCGCCCCCCCGGCGTACGAGGCCACCGACTACT

ACGCCCAGAACACTTGTCACAGCAGAAATGCTGCCAGCACCTGTCGCAATATAAATACTGCCAACAACAG

AGCGTAGTACCGATCTCCCCCGCTCCATTATATTGCCAGCCAGCTGCTCTATTCCATGTGTTCTAGATTA

GATCATTCGACACCAAATATTGTAGCACTGTGGCAGTAGTGACAAGATCTACAGAGAAGACAAGACACCC

AAAACAATACACGACAAAACAAAACAAAAACAGAGGGAAACACATGCAGCCAGACTTGCTATTCGGGGCA

ACTCGATTGGCAGGCCCGCATCAGTTGGA

>NODE_71_length_791_cov_242.764935

AAGCAGCCACTAAAGGCCAGGGACTAGGGTCAAATTGGGATAGGTAGGTCTTCACAGACTTTCCCCCCCT

CACAACCGTGCGTGCGACTTTCACCGCACACGGCTGAAGCATATTATTCTTAATATTTGAAGCAGAAAAA

TACGCTTCTACTATTATGTTCAGTAAAGAAACCGCCACCAATTATCCTGCAAGTATATATAGTAATAAAT

ATGTGACTAAATCTCATGATTGATTACTATGTTACAGGTTTTGTTGGCTAATACACCTAGATAAGTTAGC

AGACTTTCGTCTTTAAGGCACAGAGCCTCTATTTTGGATGTTAATTCCAAACATTGGCTTTTTATCTTCT

CTTCTACCCTGCACCCGATAATTAGAATTACTTCTAATCCGCTCTATTTGAGAAGATACAGGGCTTACCA

CGTTCCCTATCATCCACACTTAATTTATCCTTAGTCTCCAGCTTTACTCCGGTGGTCATACAATGTTACA

AACTGATGAATTTAGAGCATCAGTTTTAACCACTCTTCTAGAATATTAAGACGGTTTTCTAGATCCCATT

TTACGGAGCTTCAAACACTGATTCACATTACATTAGACATAGATAATCATTCGAGCACCTTATTACCAAA

CCGATTTGGCTTTTAGGTTACATTTGTCTCTCGAGCTTCAGACCCCCAAATTACTTCGGACGCATGTCAA

GATAGAATACTGACGTAACACAGTCAGTTCCGGATCTCACGGAAGTGTATGACACAGCCTCGTGTCGCAC

CTAAATCTCATGATTGATTAC

>NODE_72_length_787_cov_6.668407

GTTGGTGCATTTGACCACAATGGATTCCACCAGAAGCTACTGGAACACATTTTCTTAAAGATGCCCAATC

CATGTCGAAGAATAAACCTTCAGGTAAATTGATTTTAAGTTGAGTTAATAATAAAGTGTTGTAGAATCCT

TTAACCATTAAAGGATCTCCTTCTAATTTACCAACAACTGTACCAGCATGGATATGGTCTACACCACACA

TACGCATCCATTTACAGATAACTCGGAAGTTAATACCATGGTTTTTTTGACGAGCATAAGTAGAATTACC

TGCACGATGTAAATGTAAAATCATTTCAGCTTTTCGTGCCCAGATAGCCATAGTTTGAATAGCAGTATAA

CCGATAACTAAATCGATCATTACAATAACAGTACCAATTGAATGAGCGTACTCTGCACGTTCATACATTT

GTTCTATTGTTGCAGCAGTAATGTTAAGGTAAGAACCTTTAACTTCACCTGTTGCAGCAGCGGCACGGTT

AACACCTTCCATACAGTATAAGAAACGTTCTTTCCAACGCATGAATGGTTGTGAGTTAATGTTCTCATCA

TCCTTAAGGAAATCAAGACCACCAGCTAAACCTTCATAAACTACACGTCCGTAGTTTTTACCTGAAAGAC

CTAATTTAGGTTTTACAGTAGCACCTAAGAAAGGACGACCAAATTTGTCTAATCTTTCTCTTTCCACAAC

GACACCAGTAGCTGGACCTTGGAAAGTTTTTAAGTAAGCAAAAGGAATTCTCATGTCTTCTAGACGTAAA

GCTTTAACGGCTTTAAA

>NODE_73_length_784_cov_12.855832

TAAAGCAGTGTCAAATAGAGCAGAAGCGGTTTCAAAGGGTGCTGAGTCCTTTTTTAATACTTCTAAAGCA

AAATTGACTGGTCTTGAGCTAGGGTCTAGTTCTATTACCCCTGATATGACGTTGGACAGTATTGGTGAAA

AATATGACAGTGTAAAATCTGAATTTGATAAAGCATCTTCTTTAGTCAACACAGCTAAATTAGATTTAGA

TAATGCGCAAGATAGGTTAGAAGTTGTTAAGGGGTATGTCGTCAACACAGAGAAAAGACTTGATGACACC

CCAGTTATTTATGAGTTACCTAAAGTAACAGACAAACCCAAAGAATATTTTGATAATTTTTTATGGGAAA

TACATAATGAAGATCTTTTAACAGTAAAGGCCCAGTTAAAAGGTTCCGAAGCGGTGTTAGCAGAGTCGAA

AATAGCGTTGTTTAATGCGGAACAAGCCTTGACTGAATCTTTTATTAAATTAATTGAAATAGAGTTGAAA

AAGGGAAAAGCGTTGATAAATTTATTAAATTTTGATGTAGATTCTACTTTGGACCTCACAAGAAAAGAAA

AATTAATAGCTGTTGAGACATATATTGGAGACCTTCAACTAAATTTAGGTCATGCTCTTCGGGAAAAAGT

AATCTCTACGTTGAATGGTGTAAATTCTGATTGTCTTAAAGTTATACTTACTATGATAGATAATGATCTC

TTTAAAGCATCTTCTATTTCAGGATCGGTTAAACCAGCGTTTATTAATGCAACATCTGTTTGTGATATTC

CAGGTTTAATTGAA

>NODE_74_length_784_cov_11.652687

CGATACACGGGAGGTGTATATTTTACCGACCTCGCCTATTGGTTATTTGCCTGCGAGTTTGCGTATGGGG

CTCAGGCGTTTTGTAGGATTGCGACGGTTGTCGTTTAAAAGGGATTCCCGATTTGTTTTTGCCCTGGATC

TGTGGTAGAACCGTCACCTCCCAGTAGCTCGACGACCCTTGCGGGGTACAGTATATGCATATTATATTGA

TGTGTCTGTGGAATAGGCCTTGCGCCCTAGGGAGAGTCGTGAGCGCAACATACACGGGCTGATGGGGGAA

CCATTCGTGTGGAGCGGGAGAGATAATTGTATCACCACTCCAGCGGGGAAAAATATTCCTTGGTAGGACT

TAGGTGACCATGTCAAGAGGGGATTATTGTTCGCGGTCAACGCGGTCTTGAATGTTCGCTATCGCTGCTT

CGTATCGACGAACCCTGTAAGCTGGCGGGGGGACACCGGCGCTCTTCTCTGTGTCAGGTCGGGCCAGTGG

AGTCACTTACTTCTGGATAATAATTATGTGTCGCGTTCAACCAATTGGAGAACGCCTTTCGTTTTCTGTG

ATCGAAGGCCACGGCTACAGAGTGGTGTGGCTTGTCCAGCGGACGGTAGATGATTGAGGTTGGGGTGCTC

ATCGGAAGGGTTGGGTTTTCCAACTGGAATCGTCGCCGTTTCTCGTATAAAGTTTTGGTGGTTGGCGGGA

AGGAGTTGAAGGATGCATCAGATTATGCCATTACATGTTGTAAAAACGTTGATTGGAGTTGCTGCCGCTC

ACGGCGGGTTGAAG

>NODE_75_length_781_cov_8.963158

GGAAGCTGATTTCGGAGGTGGGGTCGTTAAGGACGTCGCATCCTTTAAAGCCTTCCAATTGGCTTCCGTT

GTCGTCGAAAGTCACGGTCTTCGAGAACCTCACGAATCCACCACTGGAGTTATCGATACCGAGACCATCA

TTTCCTCTGAACGTAGCATCGCCGCGGAACTTTATGGTACCAGAGTTTTCCGGGATCGCATTCTCGCTAT

TCTCGATAAGCATCGTCTTCTTGAACTCTAGGTCCCCAGAGTTGTCAATGACGATGTTTACGTCGCTGAC

CTCCTGAATTGTGAGGTCCCCGTTGAACGTGAGGCTGGTTCCCTCGCTCACCTCGAGCAGATTTCCCCGA

GGAAAGTCAAACGACGATCTATCGTTAGGTCCGAACAGAGTTTCGGGCATGTCGAACACGAGGGCGGCCC

CAGGCATAACGAGGAAGGCCACGTTGATAAACTGCACGGCGTTGTCGCTTTTGATGGTGAGAGTGCCGCC

TTCGACGATGATCTGCTCCAATTCCAACTGCACTGCACTCGTGTCCCTATCCACTTCTGGGCAGTCGATG

ACTCCCACGCTCAGGGTGATAGTGGTGTCTCCGGTCAAGGGCGAAGGAACGTCCAGGTAGTTGTCCAGGG

CGTCGCAGGCCGCCCTAGCAGTGGACAAGGCGCCCAACGCCAGCACAGTAGATCCCATCATAGCCCAAGA

TTTGAACATGGTTATGTTTGTATTGGGCAGCAATTTGGTGACGACGGTGGTCAGCGGCGGCAGGTTGGCT

GTCGTGTTGCT

>NODE_76_length_764_cov_10.779273

CGGCAGCTTCGGTGCCTGACGTTGTTCGAGCCCCTTGCGCGCAGGATTCCAATAATATTTCGCCCTTTCG

ATGTTTCATTGTGAGGCTTCGTAGACTCGCAATGCAGGGGTGTGCTCGTGGGTTTCGAGGAGCTACGGAA

CCAACGCCCCAACGGGTAGAGTGGGTCGTAAACTGGCCCGAGTCCAGCGGGTAGGCAGGCAGCATAGCCT

ACTACTGTATGTTTTTGTTAGTGGGCGAGCTCGCGCAAGTTCCAGCAGCCAAAGTTCACTTCGTGTTTTA

GAGATAGACCCTGAAATTTGGCACACATATTATTTGAATCCCTGCAGACCCGTTACTGGTTGAGGTTGTT

CTGGGGGGGGGGTCCGCCAGATAGTTTGGTTTTGATTTCGTTTGGGAGATTTGGGTCTTAGGGATCTCGA

ATTTTCAGCAACGAAAAAAAACAAACAACGTTCGTCAATGGTTCGGCAGGGGCACAGAACACGTGTGCAA

AAGTTCAGGGTTCAAATAATGAAAACGGCCTGAAAAATTGGACCTTTGTGAGGTAAAACGTGCAAATCCC

GGCATCGCATCGTAATCGAACTCGGTTTCAAGACATAATCGATTTTTGGCGTTAAAAATGACTTAATATT

AATCAAGCGCAGTCAGATCTTCGAATATTTGCGTGAAACTTTGTACAAACATGCCTTGGACCACCTGGAA

GCGACTCGTTCAGAAAAAAAAAATGGAATTCAAAGGAGCCAAAGAACTCCATTCACACCTACAT

>NODE_77_length_753_cov_9.957650

TTCGTCGAAACTCATCTTTGCTAGCTGAGTGTAATACTTCGATCGCGCGTACTTACGGGTTTTATATGTC

CTCATGTAGGATGTACATCCCGTATGAGGGGGACACAATCTGAGGCTGTAGAATTCAGGAAGTGTGCGTC

TATTTGTGGGGGGGGGAGAAATAATTAAAATAAATTAAAATTATAATTAAACAGTCTGAAATAAACATAC

CAGGTATACTATAGTGTTCCGACGAGTTTACGTGATGGAGTGGGTCTACTATTAGTAGCGCAAGCCACCT

GTCGTCGTTATATATCTGTATGGGGTATTTTTATCGTGCGTACTCTGGTTTTTATGTGTATTACTTATTT

GTTTTTGATACAGAATACTATATATCTATGTTGTTGCACATAGAACCAACCGTGGTAAAGTCATGTGCAT

ATCTTCGTACTGTGCTCTTTATCATAGCTTTGTTGATAATTTTATAATATATATTTAAAAAATTAAAAAA

ACAACAGCTGTGAGATCGAGGATTTCGTCGAAACTCATCTTTGCAAGCTGAGTGGGATACTTCGATCGCG

CGAACTACGGGTTTTGAGGGGAACAAAATCTGACAGCTGTGGAACTCGAGGGAGGTCACATATATACGAC

AGAAATCGTATATACCTGGTAAATTAATCTCGGTTGGCGGTCGTAGTAGAACCGGCTTCAGGCAGCTCGA

AAGGGGTACAGTACCTTTTTTTTGTCAGCTTCTACATGTGCGTTGGTTGATTT

>NODE_78_length_747_cov_20.337466

AAATTTTTTTTTTTTTTCAAGTTTTCGCCGTACATTATGTCGCTTTTTCTTGGCATGTCAATTGTTTCCC

TTACGGTAGGTCGCCCGTCCTTGGCATTCGTTGTTCCCTCGCAACACGGTTAGGGTAGGCGCAACTATCA

CGTACTTGCTGCAACACAATGCCATACTGAAGCGAGGCACTATTGAACGCAAATTACAACGACGGCAGCA

AGGAGCACGAAGACAAACTGACCCTTGCTTGCCAACTCTCACTCGCTCACTGTAGACAACTCTCAGTCAA

ATAGACAAAATACAGAACTTGGACAGTAGGAACACCCCCCCTGGCTGTCAAAGTCAACGCTAGCATTACG

GATTAAGACGCAAACCCAACTGTTCCCGTCTGGCCATATCATATATAATGTTAATGACAGCAAGACCCCG

ATGGCTATGCAGCAGCGGGATCCTACCCCCAACGGAATCTATCGTGAACTTCATAACCATCTACAGTACG

ACCCCGAGCGGCTCGTACTGTGGCCCATTATACAAAATACCTAATACGTATATTTTTGACAAGACAAACA

AAGTAGACGGACGTCCCTGTTAACCCGACGGATCGATACTACTGCTGCAGTGTTTCGCTGTACACGCTAC

CTACACGGTAACGAACCCGTCGTAAACATGTGCGCCCTCTACACCCGAAGTACTTGGCGTACGTCTTCCG

CTAGATCTTAGCATCTTGGCGCACCTTTGCTTCACAGCAGTGTGCGC

>NODE_79_length_747_cov_3.126722

CGAGAGGATTGGACACCGAAATACTGCGAGTACACAAAGTACCTCAAGTATTTTTTCTCTAAAATGCTTC

AATTATACTTCCAGGTAATGGGAGCATCTGTGAATTTTGAAATGCTATAAACGCCAGTCACATGCTCCCA

GAACCTGGGAGGAATGACGGCCGAGTCTTTAAAATGATGCAAAGATTCTCCTAACATACTTCACACTCCG

AGTATACTCCGAGTATCGAGTACATTTCGTACTCCGAGTTTGGAGTACTAGCGGCCTGAATACTCGGAGT

ACTTGGAGTATGTGAGCAGTACTGAAGCCCCAAATACTCGGAGTACAAGGAGTACCCGCAGTATCGGACC

CCGAAATACTCCGAGTACAAGAAGTATCGAGATCGTCGAGAGTGTTCATACCCGAAATACTCCGAGTACG

AGCAGTATCCCGGAGTATCGAACCCCGAAATACTTCGAGTACACGAAGTACCTGAAGTATTTTATCTCCA

ATACTCCTTACTTTACTCCAAGGTACTCGGAGCATCTGTGCAACATTGAAGGCAACGAGGAACTACCCCT

GAGCCCGCGGGAAAAACGCCGGACAAAGACTTGCATTTTTTTCAGGACTGGAGACTAGCAACTAGCAACA

CGAATGATACTCATTCGATTGCGATACTAACATACTGACACCTGGACAAAACTTGCTCAATAATAATGGC

CCTTTCTCAACACCTTATTTTTTAGTTAGACCGGAGACTAGCAACGC

>NODE_80_length_746_cov_6.386207

CCTCCTTCTTGGAGCCAGACTCCTTGACGGCCGCGAAGTGCTCCGTCACCTCCGCCGGGACCGTGAAGTG

CTCTTCCGGGTTGAAACCGAACGTCTTCTTCACGTGGGCGACGTCCGCGACTCCGAGCGGCGCTCCGTGG

GTCGTCTCCAGGCCCTGCTTCTGGCTGCCGTATCCGATGATGGTCTTGGTCTTGATGATGGAGGGTTTTC

CGGTCTCTGCCTTGGCGTTTTCTACCGCCGCCATGATCGCCGCGGTGTCGGTGCCGTCAGCGACCACCTG

AGTGTGCCACCCGTACGCTTCGTACCTCTTGATAACGTCCTCAGTGAAAGAGATGGCCGTGTCGCCGTCG

ATGGTGATCTCGTTGTCATCGTACAGGACGATAAGCCTGCCGAGCCCCAGGTGCCCACCGAGAGAACACG

CCTCAGAGGAAACCCCCTCCTGCAGGCACCCGTCGCCGCAGATCACGTACGTGAAGTTGTCGATAACCTC

AAAGTCGGGCTTGTTGAAAGTCGCGGCCAGGTGCCTCTCGGCCATAGCCATGCCGACGGCGTTAGAGATG

CCCTGTCCCAGAGGTCCGGTCGACACCTCGATACCGGCGGTGACGAAGTTCTCGGGGTGGCCAGGGGTCA

CCGAGTCGAGTTGCCGGAAGTCCTTCAGGTCGTCCATGGTGACACCGAACCCGCAGAGGTGCAGCATGAT

GTACTGCAGGGCGCAAGCGTGTCCGTTGGAGAGCACGAAGCGGTCG

>NODE_81_length_742_cov_10.285714

ATCCAGTAGTACGAACTTCAGATTCCAAAATCTAAACCATACTGGCTCAAACTTCAACAATCTTCCAAGA

CACTCAGCTGCCGAAACTAGGGCATAAGGCCTCGATAATCAGACGGCCATCAGCTCGGTGTTAATAGGTC

GATACAAGTTTATCCAGTAGTACGAACTTCAGCATCACTGCACGTTGCAAGAATCTCAAAACCTGAAAAA

TCCTCAATACAAATCTCAATACCGTACAATGTATTCTCGGAACTACCAGACCAGCCGTTATCATCATTGA

ATATCTCCCTGACAATCAACAATACAGCTCCCGATCACGGTTCGACCAAAAACAGTATATCCGCACAGCA

GTATCCACTATTTTAATCAGTAAACAGGCAGAATTTTCAATGCCCGAAAACCATCAATCAAAAACCACAA

CCCAACCTTAACCCCACCAGACCAATCAGAAACTCAATCCAAATCCTCCATTCTCGAATCCTCGTTCAAT

CAGTCAACCAGTCATCAATGTATTTTCCGAATTACCAGAAAAGCCGCCCAGGCCAGACAGCCAGAGACGG

TATCACCATTGAGATTGACTCTAAAACTAAACTAATAAATCTCAAATCCTCATTCCATACGCAAACGTAA

TTTATCCGATCGCGTGTTATGGTTTCAAGCTTACAGCAGTAGTTATTATGCCAACTTATCATCGGGGAAA

CGCGATTTCCCCTTGCCACGAGCTGCTGGTATAATCAAGAAC

>NODE_82_length_739_cov_8.121170

CTGCTGCTGCTCGCTCTGGTTCCCTCCTCCTGTGTCCAGTGTCCGGCACACCGTGGTCTTCTACGTGGCG

AGCGAGCGCTCTCTGCCTTCTGATTGATACCCTGAAACACCGAAACAGCCCACTATCAAGATGTCGAAGG

CCCTCGGGAAAATTGCCCTCATGAGCATGCTCGCTCTCAAGCTAGCCGCCGCGGCAGACACCGAGTCCCC

GAACGACCCCATCCCCAAGCCGGAGTGCGACTCGAGCCAAGCGGTAACGATCCGATACTCCTCCACGTCC

GCGCGGCTGTACCTCGAGTCCGGCGACGGCAGCACCCGCGGCGGCTGTGTGACACTGCAGCAGATCTGGG

AAAACCGTGGGGGTAAGGAGCCCTTGTACGCCGTCGACCCAGACAGCGGTGACATCAGTGACACCGCGAC

GGGGACTTGGTTGCTCACCGAGGAGCTGTTCGTGGAAGACGGCATCACCCTCCAGGTGCACGGAAAGTCT

GCCGGGGGTAAAGCCGACGAACTACGCCTGCTTAGCACGAGCAAAGACTTCATCAACTTGCGTGCCCACG

GGGGGAGCCTCGATTTCGTCAGCACCAAGGTGTTCGCATGGGACACATCCAAAAACAAGGTACGCACAGA

CGATAGCAAGGGCCGGTCGTACATCAGCGCCGTGTCGGAGGTCATCACCGACAGCAAGGAGAAGTGCAAC

GGCAAGGCGAAGAAGAACATGGGAGAGGCGCGGATGGAC

>NODE_83_length_737_cov_4.995810

TCCGGTTCAACTATGCGGGCAACTCGCACCTCATATACGAGACGGACACCGGTCGCTCCGGCCTGGTGGG

GTGCGTTCTGCTGCTTCCCATCGTGCTGCCCATGAAGTATGAGTGGATCAAGAGGAGGATGTCCTTCCAG

GCCAGGAAATACCTGCACATGACGTTCCTGCCGCTGATGATCGCGCTCTGCTTCCACAGCATCGTATTCC

GCTACGTGGGAGCCATTCTCATCGTGTGGTATGTCCTCGACCGGCTGTACTTCACCACCAAACAGTCGTT

CCTGATCACCAACCCTGTTTTCGAAGCGGTCGGGCGCGGAACGATGGTGACTGTGGACCTGCCCTTGGGT

TACACGTTCAAGGCTGGGTCGTACATCTACGTCAACTCACCCACGATCAGCCGCAGTGAATGGCACCCGT

TCTCGATCATCCAGGTGCCCGGCAAGGTGCCCAGGGCTGCTTTCTACGTCGAGGCGGTGGGAGACTGGAC

GCAGGACCTGTTCAGCCTCGGCCTGGAGCAGCCTCGCCTCCCGCTGTGGATCACCGCCGCGCAACCCAGC

ATCATGGAGAAGTCCATCTACTTCGACAACATCGTGCTGGTATGCACTGGGGCTGGAATTACACCCGCTG

TATCTATCATCGCGAGGTTTGCCAAGAGGAAGAACATCCACCTGTTGTGGATGTCGAGGGACGGTGGCCT

GGTAGCCCTCTTCGAAAAACAGCTCCGCCAGGTGAAC

>NODE_84_length_736_cov_12.534266

CTCGACGCTGCAGCCAGCGCCGCAACGACCTTCGGGTCCATGTCCTTGCTTCCCCCCTCTCCCAGGTAGC

GCTCCAACCCCTTGGTAATGTCCACTCCGGTCAGAGCCTCGACGGTTTCCGGTAGCTGGGACAAAATGGT

ACTGACGTCTTGCGTCAAACGACTGCCGGCGCCTCCACCCTCTCCGGAAATGAACACCATCTTTTCGGTC

TTCGCGAGCGGGGCGGCGATTTCTCTGGCGATGTCGGGCAGCCTGTCCACGATGCTCTGCACCACGGCCG

CGTTGCCGAAGTGCTGGAAGGAGTCACCGCGCATAGCGAGAACCTTAGCCTCCGCCTCTCCCTTCGCCAG

GACGGCCTCCGCTTCCGCGATGAGAACCTTCGCTCCAGCCTGTCCCTTTGCGAGGATGACATCGGCCTCC

GCCGTTCGCACCTTCGCCTCGGCCTGACCCCTCGCCACGATGGCAGCGGCATCCGCGATACCGTACTGGC

TGATCTCGTGCGCCTTGGCCTCGGCGTTGACACGGACGGCCTCGGCCTTGTTCTTCTCCTCGATGAGCAT

GGCGAGAGAGCCACCCTCTTTCTGCTTCTGCTGTCTCAACGCTTCCGTGCCCTGCACTTCCAGCAGGATG

ATCGCCTCCTCGGCCTTCTGTTTCGTCCTCTCCCTCACCACCGTCTGCCCCTGTCTGGCCTTCTCGATCT

CGAAAGCGACCTCTGCGGTAGCTTCCGCGTGGTTGA

>NODE_85_length_733_cov_10.672753

AATTTTTCAAGTTTGACATCGATCGACAAACCCAACGACTGCGTCATGAACTCTGAACGCATCGCGCTGG

TGAGGACCTCCGATCCACCCTAGTGTCACAGTTCTCACGCCTCGGTCGATGATATGCGGAGGGCGCCGAG

CAGGCGTCTGATGAAAACACAGTTGGCAGCTGATGAGTTCAAAACAGAACACGAAAACTCTCTCCGATCG

CAAGAACGAGTGGTTTTATGGTTTTTCGACGAAGACCGTCGGACGCCGCAAGCAACCGCTCGCCCCCTAG

ACTACGAATCGTAATGATACCTTCGAGGCCGCTATGTACACTTGCTGGCCTTGCCCGGTACGAAGTTCGT

TAGTCGTGGACTGAGTCTATGTCGAGATCGAAGGTCAATCGTCGCATGCAGAATATATGTTCTATTTGTG

GTTGATTTGAGAATTGGGCATCAAAGCGAACTGCTGTAGTTTTTTTTTTTTTTCGGCACCCATTACCAGC

TTATCCATTGTCTTCTGCCTTCAATTGCCTACAGTACTGCCTAGCGTCTATTCCCCCAAAAAACTCCGAT

TCTCCAAAGACGACAAACATGATGATCAACTCCTTGTTATATACTGTAGTCCCATCTCTTCTAGGGAATG

AGGACACACACTGGGCGGTTATATGGCAGTCACTGCTGTGCGGACCGAGAAATTGCACTGACTGACGGAA

GCACCACAAGCTTGTACGCGTACGGTGGGAACA

>NODE_86_length_725_cov_12.500000

GTTATATGTAATATGCGATGAAGGCCGTTGTTTTTTTTTCTCCTCCCCGTCGCGTCTGGCTTCACTTGAG

CTCTTTTTCCCTCAAGGCTCGGGGTGGGTGGTACCACTGCATGTGGTGTTGGCGTGTTGCGTAAGGGGCA

TGCGCGTCATTTGGTGCTGTTGTATTTTTGCTATTCTTTTTTTGTAGGTACATTAATGACATGACGTGCA

CGCTTTTGATTCAACTCTGCCTCCTCTCCGTGAGTCCGCGTCTATTTTTTCTTGTTTTTTTTTCCCCCTT

GTTAGCTCGTGTCGTGCACAACCCCTGTTGTCAAAAAAGGAGGGTGGCTTAGCCGCGAGTTTTGTGGAGG

CGTATTGATTAGTCTTTGTGTTGGGGTTTGCGGATGCTGTACAGGGATCGGCAAGACGAGGCTTTGTGCG

TTCGTTTGTGTCTGCATCCCTCCCGTCTTGGAATGCTTTTGGCGATCCAACCTCTTTTTGCCTCCGGGTT

CTTAAGTCTGGGGCGTGGGGAGCGGCGTCTGTGTTGTGATTAGCGATAAGACGGTCGGTAAACGTTGTGG

CGTGCTGTGGGCGTGCGGTTTTGAGGCTCTGCGTTCGTTTTGGCCACAACCTCGTACGTATAGCGACAGC

AGCAGTGTTGTGAGTGGTGGACGCGAGGTTTAAGGCTTAGCGTTGACGGCAAACTGCTCTATCAAATAGT

CTCGTGCTCTCGTAGTGCCCCCCCC

>NODE_87_length_724_cov_7.183499

CTCTGGAGTCAAGATGGCGGAGGAGACTTCAGACATCATGGCTGCGCGGGAGCGCATGATCGCGAAGCGA

TTCGGAGGGGCGAACGCGACTCGGACAGGAGGCGTACGGAGGAAAAAGAAGAACGTCCACAAGACTGCCA

CGTCCGACGACAAGAAGCTGACGACGACTCTGAAGCGGATTGGCGTGACGAACATCCCGGCGATCGAGGA

GGTTAACATGTTCAAGAACGACGGCGAAGTGATCCACTTTGTTGGACCCAAAGTCCAGGCATCTGTGGGC

GCCAACACGTACGTCGTCAGCGGACAATCCGAGACCAAGAAGTTGCAGGAGCTGCTTCCCGGCATCATCA

ACGAGCTGGGCTCAGAGAACCTTGACAACCTCAAAGAAATCGCGGGCAACTTCGGTGGGGCAGAATCGGC

GGCGGCCGATGACGACGATGACGTGCCGGACCTCGTCGAAAACTTCGAGGAGGCTTCCAAGTAGTTCAAA

GCAAACCTTGGACAATGTTGGCAGGCGGTAGTAGCTAAAGTTTCGGTCGTTGTTGCTTCATTGCCATGTA

GGTCGGTCGTCGTTGCTTCATGGGAGTTAGGTCTGTCTCCCTTGCTTTTGCGAGGGTGTCGGGGAGGCTG

TTGCTGTTGCTGCTGCTGCTGTTGTCCATACGACCGCTGCTGCTGCTGCTGTGGCCCGACGCCTCCATAG

TGCCGCTGCTGCTGCTGCTGCTGC

>NODE_88_length_723_cov_19.341880

CGAGGTGTCCCAGTGCTGCGAAGCGTTCTGCTCGTACCACGCATGCCCCAACAACTTCGTTCCGGTGGAC

GGCGCTGCGCACATCCAGTGCCCTACCTCCGGGTGCACCGACTACCTGTGCTGCGAAGATGGGCGCCACC

ACTAACTGCCCTCCGGAGACGGCAAGGACTGAAAAATGACATATCAAGATATACAGCGGCACCCCGAGCG

TGATGTTGCCCAACTGTACGACACGAGCTGTGGATACGGTTGTTAAATGAAGTATGCCTAGGAGGAGCGT

AAGTGCTTGAAGAAAGCTCTTCAAGCCATTAGTTAGAGGCACCACCTCTCAATAGCAATGTGTGGCTCTT

GGAGCTCATGGTATAGCTTCAAGTTCAACCGTGCCCGGATAGTGTTGTGGAACAAGTATATCGTATTGCT

AGATGTCTACCGCAGCTTTGTCGAGATTGCTGGCGGTGATTCCGCTTGTTGTCTGTCGCCGGCTACTTGC

CTGGCTTCGTTGGACGGCAGCAGGTTATGTTGGTGTTGGTGACTTCACGTGATGGTAGCATTGCTACTAT

CAGCTTGGTGACTTAGTTGAATGGATTATGTCGTCCTGTGGGAGACTCATCCAGTGTTTTGCTGGTGGGT

TGGAATATCAGTTGGGCTTTGGCCATGATTAGGGGACGGACTGGTTCGACTTCAAAACACATTAAACATT

TCAAAAAAAAAAAAAATTTTTTG

>NODE_89_length_720_cov_4.643777

AGGGGTCAGTCACACGGAAGTTGATTCCAAGTTTGACCGCACGAACGATGAATCAGTGATAAGAATAGGC

GCATAGTATCATGTACCACTCTGTACAGAAATAAAAGCACCGTTAATATTGAATATTTATGTACAAACGG

TGCTTATATTTATGGACAGAGTGGTAGGGGATTAACCAGTGTGGCAGTGTGCGGAGTAGTGCCGGTTGCC

GCTATAGTGCCGGTATTGTTGGCTGGCGCATACCGGTACTATAGAAGGGCCATTACCGTATAGTGAAATT

AATACCCGGGCGGCCCTTAATTTTTCGGACAGAACGGTATATATATATATACTACGGTAATACGTACATG

GAATACGTTTTTCTGCGTTCTCCTGTTGATCTTGCTATAAGATTAACTCCTGGGGATTGGCTTGCTGTGT

TCGTCGTAGTCGGTACGATCTGGGTTGAATTTTGTTTTGCTGGCGAGCGGGCGGGCTGCGGGAAATCCTT

CAGTTTTTGTCTCTGACTTTTGACGTCGGCGCCTTATGTTATGCATCTTTGGTAGTAGAGTGCCTCTCTG

AACGGAATTGTAATGACCGCTTCGGGTGTTAATATTGACACCAAAAACGGGTCCTTCCATTTCCGGACAG

CGTGTAGTAGCACCAGGAGGGAAAATAACGAAAAAAAAAAAGTCATAAATTAACCTCAAACAAATAAATA

AATACACACCAAAATAACTA

>NODE_90_length_719_cov_7.590258

GCGTGCCCTCGCCAAGCGCGAGGACATCAAGGAGGTCTACTCCAAGATGCGCGCTCGCCAGGCAATGACC

TAAACCGGAACGCTGCTGATCTAGTGTGGAGTTCTGGCAGCTCAGAGCAGAGGCGCATGCATGCATGGAC

GCGCGGTGCAGACATCCCGATTCGTCGCGGCATGTCTCTCCCTGTTGTCCGTGGAATCACCGACGTGGTG

GAATATTTTGCGTAGAAATGGGTTGGTGCTTGCGTCTGTCACTCTGATAGATAACCGCCTATTTTTTTCG

TTTTTGTTGCACTAACGTGTGGTGGGATATGCATTCGTGGGAAGGGTTCGCGTGCGTTTTGTCGTGTGGA

TGGGATATGCTTGCTATATACCGCCCCTTTTTTTTCGTTTTTTCGCAACTGATTGGTGTGGGAGAAGCGT

TGGAAGGGGGGTACAAGTGGGCTCCTCTATGGTTATGATGGATAGTGCAAGAGCCTTTTTTCCCCTTCGT

CTCTTAGTGTCTGTGTGCTTACGCACGTCTCTCTTCTCGTCGAATTGAGCGTGTATTGGCATCGCAGGCG

TTTCTGCTGCGTCGGGGGGCGGGCTGTGGGATGGGGTGCGGGGTGTGTACCGCCTTGTTTCTGTTGGAAC

TCCTTCGGGGGCTAGAAAAGCGATCAAGAGAAAAGGAGCAAAACTACTGCTGTGTGGCGCGGCGGCGTCG

CCAAAAACAGAAAACAAAA

>NODE_91_length_711_cov_6.779710

AGGGGGGCCCAACCCGTATAGCGCTGAAGGGCCGAAACTGAGGTACAACCGGTTCTCTTGGATGTCGCAT

TGTGCTTGTACTCCTCGTGTGAGGGCTGTCGTTGCATGCTTCCGCGTAGTTCCGAGTCTCGTTCAGCTGG

AATCCGTCGCAGGTTCAGTAGCAGTGAGCCTCGGGCGGGCATCGGCCACGCTCGCACTGTCCGAGGAACA

CCAACGTGCAGGGAAGTCCATCAACACACGGGCTCGGGTCGCACACATCGGAGCAGTAGTACGGATCCGC

GAGATTGTCGTCGTATTTCAGGCACTCCTGAAATTCGCCACAAAGTCCATAGCACGGGTCTTCATCCTCG

CACCCAGTCTCGAGGGAGCGGATGTCCTCATCGGTGACGAACTCCCAACCCCTCGATAGGCCGCACGTGT

TAGCCGAGAAAGAGCCCTCGCCGTTATGGTGGAGACCGACGAGGTGCTCGCCGACCAGATTCAGGCTGAT

CCCGCACGAGGCGCCGCTGGCGTTGGTGTCGAAGGTTATTTCCTGGCCTGGTTTGACTAGCCCTACCCCC

TTGTAGATGGTTTCGGGGTCCAAAATGTACGTTGTCCTGCGGTCCACGGGCCCCGTCCTTGACAGCGCGG

TTGCACGCACCACGACGTCTGCTTCCTCGAATTTCTGGCACAGGCTCCTAGTCACGCATCTGCAAGCGTC

AGCCTGTACGG

>NODE_92_length_710_cov_7.875181

GTGGTGACCTTCGCCCCCGTCATGCTCGAGACCGAAGTCAAGTCCGTGGAGATGCACCACGAGGCGCTGA

CGGAGGCCGTGCCCGGTGACAACGTTGGCTTCAACGTCAAGAACGTCTCCGTCAAGGACATCCGCCGCGG

ATACGTCGCCGGTAACTCCAAGAGGGACCCCCCCAGGGGCGCTGCCATGTTCAACGCCCAGGTTATCGTG

ATGAACCACCCCGGGCAGATCTCGAACGGTTACGCCCCGGTGCTCGACTGCCACACCGCCCACGTGGCGT

GCAAGTTCAAGGAGATCACCCAGAAGATGGACAGGCGTTCGGGTAAGGTGATGGAGGAGAACCCCAAGTT

CGTCAAGACCGGCGACGCGTGCATGGTGAACATGGAGCCCTCCAAGCCCATGTGCGTGGAGTCCTTCCAG

GAGTACCCGCCCCTCGGACGTTTCGCCGTGCGCGACATGCGTCAGACGGTGGCCGTGGGTGTCATCAAGT

CCGTCGAGAAGAAGGAGCTCACCAAAAAGGGCAAGAAGTAGCTATCTTACCAACCTTACTTGGATGTTAA

CACTTTGCTTGGGGAGCTGACTGGCGGCCGGAGACGGCCGCTTTCCGCTGGCTCGACCGATGCTTTATTG

GCTTTGCCATCTCTCGTTACATCCCCTCTTTGTTATCAGTCTATCCCGGCTTTTGAAGGGGAGGGGGCGG

CAGCGATTTC

>NODE_93_length_705_cov_5.634503

CATGCTGTGAGACAAACCGTATATCCCACTCTAATTCCAAGTAGATAGTCCCAAACCCCCCAAAAAGACG

CGTGCAGACATCAACGGGTCAAAAACGTCGTGCAAAACAAGCCGCCAGTTTCTTCCCCGCACGCGTCCTC

TATCATCCCCGCACGGGTCCACGCCTCCTTATCTCCCACCGTGACCGTACAGACGGAATTTTCTTACGTT

ATTTAATCCCGCCGCGCTCCGGCCTACGATTCCGCAAGGAAACCTTACTAGTGTGACATGACACAACCGA

GCGACCCGGACTAGAACCCCCGAAAAACGAAAACACATATATCAGCACGCCACGACTCGTATATATATAT

ACGAGAAAAGAAAAGGGTTTGGACTCCCCCACACACGATCAACAAACACGATGTTTTTTGTTGTTGTTGC

AGATTCATTCGATGTTGACACGCTTGGGCTCAGCGGCCTTCCCGACCGTGATCTCTAGAGCCGAAACGGC

TCCCACAGCGACAGAATTCTCCCCGCTCTCGCCGGCCCTCTCCTCCTGCTCGGCCTCGAACTTTGCCAAG

ATCTGCGTCAGGTCTGCGTCCTTCGGCAGGTTTACGTCCTTGAAGAGCGACCGGCGGCCTTCCGAGGCGT

AGGACACGCGGAGGGTGTTCTTCGGGGTGACGACGAGGTTGATGCCGTCGCGGGTGGCGTCCTCGGGGAG

GGAGA

>NODE_94_length_704_cov_6.434846

TTTGGCGACGTTCAGCAATACCAACTGCAGGAGCACCGAAGGAACGTCGCTATACAACGCCGGAACCATG

ACATTCTACGGAGAGGCGCGCTTCAATGATATCATTGCCCGTAACAACGGCGGCGCTGTTCTGAACGAAC

TGGGATCAATGACCTTCCGAAGCGACGTGGAGTTTAACAACAACGATGCCAGCAACTTCGGTGGTGGTAT

TGCTGTTGAGGGTGGAGACGTAACATTCCGGAAGGCGGTGAAGTTCATCGGGAACACTGCCGACTCTGGT

GGCGCTTTCGCTGTCATGTTCAAGGAGATGTTCATCCGTGGCGTGTTAATTAACTCTGGCTTTGGCTCGC

TTACCTTCGAGACCCCGTCTGCGGTAGTCTTCAAGGACAACACCGCTGATACCCGCTACTTCGACGACAA

CTGCACCGTGGGCGACGTTGAAGAGGGAACTACGCTGATAGGCTTTCCTGGTGACGACGTGTGTGTCGCA

GCTGTGTGATTCGTCGTCAGCGACATGTTGGGTGAAATAGTTTCCGGAAGGACCTTGTTTGTCTCGGAGC

TTCGTTCCGGTATCGTATGGCCCATTTTTGGTCGAAGCACAGGATGACAAGTAGTTCAACCCGTCAAGCT

GTCGGATATGCTAGCCCGGAGGTCCGTGATTGGTGCTCGGAGTGTTTATCGCCCAAATAAGTACTTGATA

TACC

>NODE_95_length_703_cov_5.875367

TGGTATCATCGGTATGTTGTCGTTGTCAAGTAACCATTATGCAGTGGCATTTGAGATAATTAAATTGCGG

TAAAAAGAGAGATTTGCAAGCTGATACGTATGTATGAATGCATGAAAAGAGCAAAATACCCGTGACCACC

CTGATTTCTGGTCCGCACGATTTGTCCCCTGTTGTTTGTTTCGAGTCTATGCTGTCGAGGTGACTGCTGA

TTTTACTCAGGGGGATGAGCTGGGTGACCCGAGCTGCGGCGACGCTGTGAGGCCGGAAAGAGAGGCAGCC

GCAGCGGAGGCAGCACCAGCAGTAGGAATGTTTGGGGCATTATTGGGACTGTGGGTAGGTTTTAAGTTTT

TGAATGAAACGTGATTGCCTTATTCTTACGAGCGATTTCGTTGTTTGAGTTTTTGCTGACCGATTTGCTT

GATGTGTATTGCGTTGAGAGTTTGACTGTTGAAGATGAAATAATCTCAGTCTACGACAAAATGACCACGA

CTGAAGTGACTTTGAACTGCACCAGAGGGCGCGTTCGGTCCAGATTAAATGTTCTTGGTTTTGAGCTATA

AGTCGTCTTTCTTTGCGTTTGTTATGCGTTGGATCGAGCGATATTGGGTCTGCATACTTGAAGTAATGTA

GAATACACGTGGATGGAGTTTCTGCGGTTACGTTCGTATAGTACGAATGGAATTTCATTTTTCCTGGTTC

AGC

>NODE_96_length_696_cov_8.583704

CACGCGCTGAGAAACGTTCCATTATTGCACCCGGTTGTAACGTAGAGATGAAAAGACATTTACGACATAC

TCGCGTGCGAACCGCAGCTCATTCATCAGCGCAAGAAGGTAAGAAAGCAGAAAAGGTAGCAAAAAGCTCG

CCAACCTAGCGAACCCCCCGCGCTCAATCCTACACCCCGACCCTTCTCATCCTGCACCGCCCCTTTTTTT

TTTTCACCCCATGCGTCCATTTTCCCTTCAGCACCCCCACCCCGAAGAGATGCGATGCGTCAGACTGGGA

AAAGAAGAAACGCGACTTAAGCCTAAGAAAAGTTGAATTGAAGAGAGAACGTGCTACGTTTCACAGCAGC

ACCATTCCTCGTCAGCTGCAACTCTACTCTTCGACGTTAGTCGCGCGCGTGTAGATGACCTGCGCGTGGT

GCGCGTCGACCTTCCGGATGAGCCCCTTGCCCTCGAGCTCGCGGATCGCGGCCCGTGCCAACGATCCGTT

GACCTTCAGGCGCTCTACCAATCCGGATGCCGTGATGAGCTTCATCTTGGGCACCTCCGCCAGGCAACGG

ACGTACGTGTCCTCGTTGAAAAGGACCTGGTTGGCCACTTTCTCCCTGACCTTTCCCTTGGACCACTTCT

TTTTGCGGCCCTTGCCACCCGCCATGGCGGCGGCCATCTTCTGCTCCTTCGACTTTTGCTGCGTCT

>NODE_97_length_694_cov_8.019316

AGAGTACGGTTGGTCCGCCAACATGGAGCGGATCATGAAGGCGCAGGCCCTCCGCGACTCGTCAACGTCT

TCCTACATGACGTCCAAAAAGACCATGGAGGTCAACCCCACCAACTCCATCGTTGTCGCGCTTCGGGAAA

AGGCGTCCGCGGACCAGAGCGACAAGACAGTGAAGGACCTCATCTGGCTCCTTTACGACACGTCGCTGCT

CACCTCGGGGTTCAGCCTCGACGAGCCGACCACCTTCGCTGGTCGCATCCACAGGCTGATCAAGTTGGGC

CTTTCGATCGACGAGGACGACGCGGCAGCAGACGACGGCGAGGACGACATCCCCGACCTCGACGACGCGG

ACGGGGACGAGGAGTCAACCATGGAGCAGGTCGACTAACCGATGTCACTGTTAGCAAATGGTTGAGAGGT

GGTGGTCGTTAGGGACTGGTCGTTAGCGGGGAAAAAAGCAAGGTGTAACCATCGCAAATGGTTGTCAGAG

GGCGGTCGTCGTTGGGCGCGAAAAGAGAAGCAGGCGGGTGGCTGTTGGCGAGAGGAAGACTCAAAGGCTC

TTTTAGTCATTACGTAGTGCACGTTGGTAACGAATTGACGTCGTCCAGAGCGGTAGTGTGAGGGGGCAGC

TTTTTTCTGGTTCTGTCGTTGTTTGGTGGTTTGGTTGTGAGAGGGGGGTCTGTCTGGCGGAGGG

>NODE_98_length_692_cov_30.845007

GGATTGCCATCGACTTCACCTCCCTCGCCGGGCTGGCCGGGGCCTTCTCGGGGTTGAAGGAGATCGACAC

CACCGCCTACATGGAGGAGGAGCCCCTGGAGGAGGGGATGTCCTTGAGGGAGTACATGGGCACGGTCTAC

ACCTCGGTGAAGAGGGTCTTCGAGAAGTTCTACTCCTTCGACCAGAGGCTGGTGATGAACACGGTCAACG

ACATCCAGAGGAACTTCATCGGGGACGAGAAGAAGAGGAAGCTGGCGGTGGTGTGCAACATGGTGTACAC

GGCCGTCTACATGAAGATGACCGGGACGGTCAACCCCACCACGGGCATGTCCGGCCTGGCCCAGATGGAC

GCCTTGAGGGACTTCAAGGAGAACAAGGGGTACAACTCCGCCTCGGACACCCTCGTCGTCATGGTGCAGA

TGGTCACGACGGGCTTGATCACGGAGATCTTCTCGTCCATCGGGGAGAAGGCGCCCACTCTCAACTCCCT

CATCAAGCTGGACATCGCCTTCAACACGAAGAGGGCGGTGGCCTTGGACAGGATGGTGACCAAGGTTCAG

GACAAGGGGAGGAGCTTGCTGTCCATCTTGTAGTTCCTGTACAGGGAGCAGGCTAATGTTCATCTTCGGG

TTGGACAACATGATAGTATTAATAAGTAGAAGACATTGTGTTGGAATTCTAGGGTGACAAGG

>NODE_99_length_692_cov_12.229508

TGTATTAATATGAGTTTTAGGTGTGTATTGAGTTGTTTTTATCATGTTTTTTATTCCAATGTTTCAACGG

AGACCCTCCATTCCTGTAAGGCGTTCCTCAGTCGTCGTCTGCTACAGTCCTGTGAAGATGTCTACGTACT

GTAGAATCAACTAGACTTATCCAATCACGATGCCCTCGAACTTCGCCTGGAACCACTTGACGGCGTCAGT

CTTGGTGACCTTGTGCTGGTAACCGATCTTGGTGCACTTCGCCTTGCGCTTCGCCACGCGGTATCCTGGG

CGGTTAAGTACGACGTAAAAGTCCATTCCGTAGATACCGGTAGATGGGTCGTACTTAAGGCCGAGATCGA

TGTGCTCCTGGATTCCGAAGCCAAAGCTGCCCGTGTTGGAGAAGTTCTTGCGCTTAAGCTCGTACTCCTT

CACCTTAAGCCCTCGCTCAAGGATCTCCAACGCCTTGTCTCCGCGGACCGTGACGTGGCACGCGATCTTC

TCGTTACGGCGGATGCCGAACGAGCGAACCGTGTAGCGTGCCTTGGAGTAGACAGGAGACTGGCCCGTAA

GTTGCTCAAGTACCTTCGCCGCGCGCGTGAGCCTGTCACCCGACTCTCCGACGGAGCAGTTGAGCACCAG

CTTGTCGACCTTGAGGTCCCTCATCGGGTTGGTCTTAGTCTCCGCCATCACGCCTGAGTATG

>NODE_100_length_689_cov_10.476048

GCAAATATAAACTTTTATCCGCACATTCATGTAAAAGATTTAGTAGCTTTTTTTGTTTTAATGTCCATGT

TATCTATTGTTATATTTTATTTTCCTAATAGTTTAGGAGACCCTGATAATTATTTAGAAGCTGACCCTTA

TGGGACTCCAGCGCATATTGTTCCTGAATGGTATTTTTTACCTTTTTATGCTATTTTGCGGGTAATTCCA

AATAAAGTTGGAGGTATTCTTACTATGGGTGGAGCGATCGCTATAATTTTTATATACCCCCTTTTAAATA

CATCGATACTGCGAAGTGGTAATTTCCGGCCAATTTATGCTGTAGTTTTTTGGCTTTTTGTTGCTGATTT

TTGCTTATTGGCTTGGTCAGGAACTACCCCAGTAGATGATTATTTTAAAATAGTTGGGGCTGTTGTTTCT

ATAGGGTATTTTGCTTTTTTTGTTTTCGGTGGGTTATTTGTAGGTTGGTTGGAAAAAATGCTTGCTGTTC

GGGTATTAAATATGCGTTCCACTAAAGGGAAGTAACAATAAAAAGGTATTAGGATAAAATGTTGGTTTTT

GTTTAATTGTGCTCAGATCATGAAATTTTCACATAAAAATATCATTAGAATAACTTCAATTGTTTTTTTT

TCGGGGTAGGCGGGAATGGTTTAGGCCCTGTATAGCCACGGGGTGCCGCGCGTTGACCC

>NODE_101_length_688_cov_15.203898

GCAGCAGCAGCAGCAGCAGCGAAGCATGGGAAACTGCATCGTCACCCCGCCGTCCCAAGTCGCGGTAATC

TCCGGACCCGGGAAGTCGCGCATGGTCATCGGAGAGTGTTCTTTCAAGAAATGGGCTATTGAAAGCGTGG

AATACCTGAGCCTTGAGCTGATAACCTTGTTAGTAAGGAGCGTGGAGGCCGAGACCGTGCGAGGCGTTCG

TGTAACGGTCGGCGGTACGTGCCAGGTCAAGGTGGACGCCTTCACGCACGGCGACCTGTCCCAAAACCTG

CCGCAGATAACCCTCGCTTGCCAGCACTTCCTTGGGAAGAGCTCTGAGGAGGTGCACCAGGCCTTGCTAA

AGACTCTTGAGGGTCACCAGCGCCAGATTCTCGGCACCCTCACAGTGGAGGAGTTGTATAAGGACCGCGC

TGCGTTTTCTCAGCGCGTTCGGGAACACATCAAGGAGGACCTGAACAACATGGGTTTTGCTCTCGTGTCT

TACACGGTCAACCAGGTCACCGACTCGCAAGGATACATGGAAGCGCTCGGTGCTACGCAGACCGCTCTTG

TCAAGCGTGAGGCTGCGGAGGGTGAGAGCAAGAACGTCTCCGAGGCGAAGAAGCGCGTGGCGGAAAACGA

GAGCTCCGCGAACATGGCCGAGGCTAGCTACCGCGCATGGAATTCTCGGCAGCCAAGG

>NODE_102_length_685_cov_2263.171687

CCAGTGAGCTATTACGCACTCTTTAAAGGATTGCTGCTTCTAAGCAAACCTCCTGGTTGTTTAAGTCTTC

CCACCTCCTTTACCACTTAGTAATTATTTAGGGGCCTTAGCTGGTGATCTGGGCTGTTTCCCTCTTGACA

ATGGAGCTTATCCCCCATAGTCTTACTGGTTAGCTATACACTTAGTATTCTTAGTTTGCGTCGATTTGGT

ACCGAATTACCAGCCCGCACCGAAACAGTGCTTTACCCCTAAGCTATAACGCTAACCGCTGCGCCTAAAC

ACATTTCGGGGAGAACCAGCTAGCTCCGAATTCGATTGGCATTTCACCCCTAACCACAGCTCATCTGCTG

ATTTTTCAACATCAGTCAGTTCGGACCTCCACTTAGTATTACCGAAGCTTCATCCTGGCCATGGTTAGAT

CATCCGGGTTCGGGTCCGTAATTGGTGACATAAGCGCCCTATTAAGACTCGTTTTCACTATGGCTCCAGT

ATTTCTTACCTTAACCGTGCCACCAACTACAAGTCGCCGGCTCATTCTTCAACAGGCACGCAGGCAGACG

TTAAAAGTCGTCCTTCTGCTGCTTGTAAGCTTACGGTTTCATGTTCTTTTCACTCCCCTCCCGGGGTTCT

TTTCACCTTTCCCTCACGGTACTATTACACTATCGGTCATTCAGTAGTATTTAGC

>NODE_103_length_678_cov_18.847793

TGGGGGGGACAAAAAGTGTTGATACGTGGCTCTTAAAACACGCGATTAGATGTAGGGGCTTAGGGGGGGG

GGCGGAAGCAAAACAGACTGGCACATCCAGTAGGTGAATGATAGTTCCAGCCCTTGTGAGTCGGATGAGT

TCGGCAGTCTTGATAAGTTCGGCAGCATAGAGCAAGAAGATCCATCTCAAAAACCATTTCGGCCTCTTGT

GGTCACGGGAAACTTTTGCTGCGGAAAACGTGGCGAGGGGGAAGAGGGACCCGCGGATTCTCGGTTGGCT

GGTGTGGCACACATGATGAATTGGATGCTGCCCATATTCAGTAATAATGTAGTGTAAGAACACTGGTATA

TACGTATATACATAATAATAACAAACCTGTTTTTTTTTCTCCCTACGTTCTAGTTTTTGTGTTGTCTCTT

AAATACGTACATCTTCTCGGAAGTGGAAGCGCATTCTCTGAGTATTTTCTGCCGATCGATACTACTCTCG

ACACGAAAGTTTTATGTTAGTATGCTGGCGTTCCACCCAACGCGCGAATAGGGTGCGTTCATTTTATGAA

ATATACCTGGCATTATTTACTGTAAAAAAAAAAAATAAGAAATGGAATTCTAGTTTTCAAATTAAATAAA

GTAAAACATACATCTCTTATTACTTCTTATTCTTGAAAAAAAAAATTG

>NODE_104_length_668_cov_8.013910

CATAGCTATGTCGCTCGTTATCTCGCCTGAGTTCCAGCACATCCTGCGTATTCTCAACACCAACATCGAT

GGCCGACGCAAGGTCCAGTACTCCCTGACCTCCATCAGGGGTGTGGGACGCCGTTTCTCCAACCTCATCT

GCAAGAAGGCTGAGGTTGACCCCAACAAGAGGGCCGGAGAGATGACCGCTGACGAGATCGAGAAGCTCGT

GGCGATCATCCAGAACCCCCGCCAGTTCAAGATCCCCGACTGGTTCCTTAACAGGCAGAAGGACTGGAAG

ACGGGCAAGACGACCCAGATCTCTTCCCAGACGATCGACCAGAAGCTCCGAGAGGACCTCGAGCGGCTGA

AGAAGATCCGCTCCCACCGTGGTATCCGCCACTACTGGGGTGTCCGCGTGCGCGGGCAGCACACCAAGAC

CACCGGACGCCGCGGCAAGATCGTGGCGGCGGGTGGCCGGAAGTAGAGAGGGGGCAGAGTTTTCTCATGG

CCGTCTTTTTGACTTTTTGCTGTCACGGCATGGGCTACTATTGCTTGCCGTCGTCGCCATGCGCGTTGTC

GTGTGGTAGCGGGGGGTAGTAGAAGCGGATGCGCAGGCAGGGAAGGTTTACACCGGAGTGAACGCAAGGG

AGATTAAAAAAATCCGACACAAAAAAAAAAAAATTTTT

>NODE_105_length_664_cov_5.122862

AGCGCGGAACCTCCCTCCTCCGCGACGATCTTCTTGAAGCAGTCGGCGGTGCCCTTGTACATCCACTCGG

CGGGGGGCTTCTCGGACTGCATCTGGAGGCGACGGCGGATCGTGTCGAAGGGGTAGGAAGCGTACCCCGA

AAGGATCGCAACGGACTGCGCGATCAAGAACTTAGATCCCAGGCCCTTCACGCCCTTGTCGTCCTTGTAG

GGGTTCTTCTCACGGAAGGTGTCGTACAGGCCGAAGTACACACCGCGGTAGGGGATGATACCCATCACGG

AAACGCCGAAACCGTTGTAGAGACCCATCACGCCCTTGGGGCCCTTTGCCGTCCGGCTAAGGCAGTCCCA

CAAGCCGTTGAAGTCGCGCTTTCCGGTGCCTACGTCGGACGCGAGACGGGTACGCGCGTAGTCGAGGGGG

TACACGAAGCACAGCGAACCGGCGCCGGCAAGACCGCCGGAGGCCATGTTGATGGCGAAGAACTTGCCGA

ATTCCGTCTTGGGGTTGGCCTTAGGGAACAATCCCTTGATGGTGTCCTTGAAAGCGAAGTTAAACGCCTG

GGTGGGGAAGTACCGGATGATGTTGGTCATGTTCCCGCGCCAGAAGGCCTTGAGCCCCTGCTCGCTGTAC

ACGCGCGTGAAGCAGTTCCCGATGCCGGTGTAAC

>NODE_106_length_656_cov_191.557480

ACTACTACTACTACTACTACTGTATTTTTATTATCTTGTCTTGGTTCCATGTATTATTCATCTTTTTTTT

TCTTTGATCCAAATCCACCACCAACGTGCGTGCTCCACGGCTCAGGCGTGCGGGAAAAACACCCGCAGAC

TGAATTGATAAGAGCACAACTCGAACCAACCCAACCCTACTCGCTCCCCAAATGCGCTGCGTACGCTCGA

ATCCCCAGCCAGTTCCCACCCACGTCCCCTTCCCACTCGTGGAGAAAAACGGCTTTCGCAAGAGGAAACC

CCAACATGTCACTTAACTACAACCAACCGCAGGCCAAAGACAAGCGGTTGATGCAGGGAGTCAATTTTGA

GGCTCAATTTTCGGAGCGGAGCTAAGACACGATATTAGTAGAAGCCGATTCCACCCCAGCCTGAAGACGA

TTCGTCGAACCCTCAAGAGAAGACAGGAGCGGCAAACGCTTTCAATCCCCGTAGGGGGGGGGGGCTCCAC

CACATACAGCGCTGCGGACACAAAGCGACCCCGAAAACTAACTACGGAGACGGCTGGCGAATAGCTGGAA

CGGTTCTCGGCGGACCGTCCATCCCTTAGATCTGGGACGGGTAGAGCATCACTGTTGTCCCCCACCTCAA

GGCATGTTCCCAAGCATTACTCACCC

>NODE_107_length_655_cov_9.686120

CTGCGCGTTGAAACATGCGCGTAGCAGTTATAAACCAACCATATTTATGGACAGTCGGTTTTTTGTAGAT

GCTTGTTCGGGAGTACTTGGGAAGGAGGACTGGGCCCTTGCGGTTGGAGGTGGCAGGGTCACGGGTACGG

GAAGTAGTGGCAGTGGTAGTATGTGGGGGGAAGGGTTGGAAGAGGTAGAAGAGCCACAGAGAGAGCGAGA

CGGCAGGGACTGTCTCCTAGGTACTACCGATTACAGTGTAGACCAGCAACACCGCACGACCTTCTGCGCG

TTGAAACATGCGCGCAGCAGTTATAAACCAACCATTTTTATGGACAGCGGGTTTTTGGTAGATGCTTCCT

CGGGAGTCAACAGGGAGGAGGTTGAAGGTGTAGGGCTAATTGAACACGGAGGTGGGCACCTGGTTTTCGT

AGTATTCGGCGGTTGCAAATCACCGTTTTAGGTGTGGACTTATTGGGGCCTGGGTGAATCATGTTGGCAC

ACAGCAGCAGCAGTAATTTTCCAGGGTTCTTTACGGTTAACCGTCCGTGCACGGTGCGTTTTTTTTTTTC

GAAAATCCTACGGTCCGGTTCGGTGCGGTTTTTAGATTTAGTAAATCCTACGGTGCGGTTCGGTTTTTTC

ATGCATCCTACGGTGCGGTTCGGTG

>NODE_108_length_653_cov_13.409810

GGGGAGGGTAGTCCGGTAGGAGATCGACATCCACTAGTTGGAAACCCCGTCGGCACCGTGTGATTCGTTT

TATTTATAGACAGGTAAAATATAATAACGCAGAGTGGACGGGGGGGGGTAGCAGCAGGCTACCAACCCCA

TGGTATGAGGCTAGAAATCGCTTCAATTTCCTTGGTCGGGTTTGTACGTTGTGAAAATTGATCAGGCGGG

TTTCGTATCGCTTCCCTAGTAGCTATAGCCTCGGGGGACGGGAGGGGGAAGGGGGCACATTGCTTGAAAT

TGACGAGCATATGGTTGTATCATAAGAGAGAATTCTCTTCCTTGAGGCGGAACTACTCAAGTGCATTTTC

TGAAAATGACGAGCGTTGGTAAGATTTTTCCTGGAAATTTCATGGGATTTTTCCTGGAAAATTCCTCGTC

TTTGAGGCGGTATGACGCGGGTTGGAAGGAAATTTCATGTATGGAGTTTTCCCTTTTTTCAGGCGGTGTT

ATGTTCGTCGGTACTTGCGTGTAATGCGTTGCTGTAGTGGGGACACAGGGGTGAGGCGGGGGGTGGCTTG

ACGGCGAAAGAGCCGTTGGAAAAGCTGTTGAGGATTCTGCCCGCGCGTGTATTAAGCAAAAAGCCCAGTC

CGACTGAAAAAAAAAAAAAATTT

>NODE_109_length_653_cov_8.691456

AATGGGATTATCACTCTGAGTTTACGCACAAACGTGCGGTTCCATCTGGTTCGGAATGTGTGCGAAACAT

GATGGCGAATTACGCGCCGGGTTTGCTGTAAGGGGTCTGACTCTAAGTATGTGTAACGAAGGATTCGAGC

TATTTTAGGCGTTCGCACCCTAGTCTGCAATTCCTTGTCCCCAGGCTCTCACCGCTTCGTAGTTCGTTTC

CTCCGTGGGTATGCTTGCTGAGCGATTCCTCCGAGTTCTTGTCACAGATGATAGTTTCCTGCGCAGACCA

GTCTTTATATCAGCTGCCTGCAAATTCCAGTCTCGCCTCAAGCCGTCCTCTTCTGCTTCGCAGGCAGATC

TCTGTATCAAACCTGCAACAGTGCCTTGCTTTAACTATCGCGGTACCCTGTCTGGTTTCTCTGTCACTCA

AGTTCAATCAAGTTCACGTTTTTCCCTCGGGATTAGTTTTGTGTGTTTTTGGTTTTTGGTCTGACCTAGG

CGGTGCTTCCTTTGGAATGGCATGGAGTTGCACATATCCTAGGAAGCTTCCGCTACGTACGTATTTTACG

TTACGTATGCCAATTCCGCCGCAACACCCCCCCAGCACGGTTTCCTGAGAATACCGTGCTAACCGGGCTG

CGGACACACTGGAGTAAGAGTGA

>NODE_110_length_647_cov_9.124601

CAGAAAGCGCAATAAATAGAGCTTTCTCCAGATATGCGAGATACCCGGAAGGGGATTCTTCAATAGAGTC

TAAGCGTACACGATACACCTTTTTGTGAAATGCTACCGTGTGTGGCTACGCATGTCGGCCCAAACAATCA

GCGCCCACCGTTCACAGTGCACAGGCCTCGAAATGTTTTCCAAACGTATTTACAATTGTTTCGAAGTCTG

ACTGTTTGGGTATCCCCGCTCCGGCCTAACTTACAGGTGCCGATCCACCACGTTCCCTCTAGCTCACAAT

ATCCCAATGGCGATGCAGCTGCGCTCCAGGAAGGCATGACATCAAAGGTAAGCAGGTAACGCCACAAATT

TCTACCAGCGCATATGTGCGAACATGATGCATTTCGTATTTAAATCGTGTCCATCTCCGTACAGACCCCC

CCACACACACCCCCACACACCCACTTTCCATGGGTGCGGGATGTTGTCTATGCGCCGCAGAACAGTTGTA

CACTGGTTCTTCCTAACAAGAGAAATCACATAGTATGACCAGTCTCTTTGCACAAACCGGACCCCTACAT

CAAAGCGCTCCGCTGCTACATGTAACCAACACGTGCGTATTGCTCTCTAAATGTATCTAGTTATTTAATC

CAAGTGAAATCGACAAC

>NODE_111_length_646_cov_24.292800

GCACCCCGAGAATTCCAAGAATACCAAAGTCGAATTATTATTTGACTACGAAAACCGTAGCTGAACCCCC

TCCTTCAACACGAAGTACGCGTTGCCCTTCCCCCTTGACCCCTCCGGGATCGCCACTACGTGTATCTCCC

CGTTGCCTCAGCGTGAACGTGCTGGGGGAAATTCCGAGCCGGCCGACTTAACTTACAAAGCAAAACATAC

CCCGCTATCTTTCAAGAACAACCAAAACACATCCTTCGTACCCCGTAGCGGTCACGGAGGAGTGTGTACG

AACCAAGTTCCCATCAACTACGTACTACATGAGATCTCGACGTACTGTAGCATTGCCTCTACGGCACCAG

CGGCCAAGGCACCGCGTCATTGACCGAGGCAACTGTGCGCTCAATGGGTGAGATGGCTATTCCAGTCTTC

GGTCGGTCTCCTCGTGGATGTCGTGAGTCCGTCACGGTGCTTAGAGCTCAGTATGGAGCGAGAACTTGGC

AAGGTCCTCGACGATGTCGTCGAACGATTCGTAGTCGTCCTCCTGCACCTCGTCACAGAAGTCGTCGCCT

GTAAAGGCCCCCTCGCAGAGCTTGCCGTCCAGCTTGAACAAGTTCGCGCCGTACAGCTCGAGCATCTTGC

CGCTGTAAAGGCGGAA

>NODE_112_length_641_cov_8.219355

CCTTGGCACCCGTTAATTCCACTTTTTTTTTTTTTGAAGGTTCCCTCTCTTTTTTTATTTTTTGTTCCCG

CGGCCGGGGAAGGACCACATCTTCTGGGGGGGGAGCAAGGACCAACCAAAACGAAGACAGAGCACACAGA

CGAAAGCCGAGCCAGACAAAAAAAAAAACAGGCTGACAAAAAGAGTTGCCTCGCAGCGACAGCATCCTTT

CTTATGCCGACTCAAACCAAAAGAAAACAAAACGTGTGTGCCCGAAAACAACACGTCTTGCGTACCGACG

ACAACCGACCGGGCAGACAAAAACTGATCTAACCGGCCCACGCCTAATCGACAGAGGCGGCAACGCCTCT

ACCTCATCGAAAACCACCCATACCATTGCCACCGATCGAACAGGCGACGCAAGGAGAAAAAGACAAGAAA

AACCCAGCTGCTTTGAGTCAACAAAAACCGGCGGATGGATCCCCCTGCCCCCTTCTTTTTTCATGGACAG

TTCGCTGTCTTCGTGTCGAACGCATTTTTCATGTGCTTGCGCTTTTCAGTCCACGCTCGCCATCATTCCT

CTCCCTCTGCTTTGGTGGCGAGCGCGCGCTCCGCCGTGTTTTCTGCTTCGTTTGTTTTCGAGACTAAAAC

GGTTTGCGTTT

>NODE_113_length_641_cov_7.922581

CTTAGACGTCAACAGGTGCCCGCACCGACGCTGTCGCGCGCGGGTACACCTTATTCTCGGTCTCTTCTTA

GCCGTCAACAGATATATGCCCGCACCGACTCTGTCGTTCGCGGGTTGTCCGGTTCCTTCCTAGTCACAGA

TGACCGCATCGACTCTTGGGTACATATTTTTGGTTTCTTCTTCTCTTAGACGTCAATAGATGCCCGCACC

ATCTCTGTGGGGTACGGGTCCATGTTTTGTCGGGACTCTTGTCGGACGTCAACAGATGCCCGCATCAACT

CTGTGGCAGGCGGGGTACATCGTGTTTTTCGGTTTCTTCCTAGACGTCAACATATGTTCGCGCGGTGTGT

TCCGTCGCCTTGACTAATGGCTGGCGAGCTCCACGGTCTTGCCGCTAGCCGAAGACCACTTGAGCAAGGC

ATCCAGCATGGCCTGTTTTGTCGTGGCCCCGTCGCTCGTGACCACCACCTTCTTCGCCTCCACGTCAGCC

TCCACCGCGGACACGCCTTCCATTTTGCCGAGGATCCGCTTGGTCGCGTTCGCGCACCCTTCGCACGTCA

TCCCCACGTTGAAGACAGTCGTGTCGGCCATGGTGGTTCGATGTGGTGAATCTGGCAGAGCTTGTGTTCG

TGCAGGTCAAG

>NODE_114_length_638_cov_4.675851

TGATTTTGATGTTCTTCGTCCAGTCGTGACCAGCCCTTTCGCGGTAAAGGCGGATTTGCACCGCCTTCGC

GTTTTCGAGGCTTTTTTTCACATATATACCAAGCTAAGAACAAGCTCGTACGTCGTATTAGGGCAAATAC

AATCTCGGGCGATTTTGCTTTTCCTCTTGGAAGCCGTTTTCGTGATAACAGGGCTAATTTTTTTGCACAG

CCCCTAACGAAATACATGATAGGGTGCTTGCTCGAAGCTTGTGCATACATACGTACGTACGGAGCTTAGA

TCTGTCTATCAATAGGCGGAGAGCCTCTGCTTCAAGCAGCGCGGCTGAATTGGAGTGTCACGGATGTTGA

TGTTGAAAAGAGATATTTTTTTTCCCGACTGGACCGAGAGACGGGTTAGGGTGGTCGCTCTTGCTCGTGG

AGTGGATGAAAACCTCACGCGCGGGCTGTCGTGGAGGGTGTGTGTTTTCTGTTTTCTGTTTTGTTTTTTT

GTTTTTGTCTTCCATCGTTCAAGGCCTTCGTGATAAGAAGGGCTACTTTTGTACAGCCATCATAACGTTT

TCGAGGCTTCTCTACCTCTACGAAGCTTTTGAATCGTCTGCAGTAGCCGAGGACAATGTAGTGTCGATCT

TCATCGTT

>NODE_115_length_636_cov_3.505691

GTAGTGTCGCTTGCAGAATATTATGCGCTGTTTGTGGGAGCAATAAGGTGCATCCCAGAAACGTTGGTCA

TTGAACCAACACAATCTACACCAATTTCCCACCACACTATCCCCATAGGACACGAACAGCACGGCCTGCC

GTATGTACTAGGACGAAATAACACCCATATTTGGGTACACAGACGATTATGAAGCACACTCCATGCGTTA

TGGGTGCGTACACGGTTCGTGCTGCTTAGCCCTGCTGCGCCGGTATGTTGCCCCCTATTTCCGAGCCGGT

TCACAACGCCAAACGCATCATGCCGCTTCAAACTAAACCCGTCTCAGCCGGTTGCACTTTGCAGTTTTCT

TGCTCGTGGGCAAGACAGGCAAAGTCATAATGCCGTAGAAGGAGTACCACCCCATTGGGTAGCGTCTCGG

TTGACTAGCCAGTCAGACAATGTTGGGGCCCAAAATAGCCCCCGCCCTACCGAATAACGCCTTCATACGG

TATGCGCATACAGCCACGAATATACCACTCGCCGTCTACCAGAGGTCTCAACCTTGCGAGGCTCATACAA

CTCTCGCGCAATTACACCGTACTTCACCGGTCGGGACTACACCCAAGAGACTTGGCTCATTGTCCAGGTT

ACACGG

>NODE_116_length_632_cov_8.471358

CGTCCGGGTTGCCAATCTGCCCCTGCAGGTAGCCCTCCTTGTCCTGGTAAAAGTTCTTGATGCCCTCCCA

GTACGTCCACTGTGTGTCAGTGTCGACGTTCATCTTGACCACGCCAGCCTTCACGGCAATTTGGATGTCC

TCCTCGGACGATCCCGACCCCCCGTGGAACACGAAAATGCACGGCCGCTCGTCCGCGCCCTCTGCGACGA

GGGCGGCGTGCACCTAATCATGGTGGACACCCGCCTGGAACTTGGACAGTGGGCTGGCCTCTGCAAGATC

GGCCCTGAGGGAGAGGCTAACAAGATCGTCAAGTGCTCTTGCGCTGTGGTGACCGACTACGGAGAGCAGA

CCCAGGCCCTCAACGTCCTCCTTGAGCACCTCAAGAAGGGCGGCGACTAAAAAAAGCCTAGGCGTGCTTT

TGCTGCTGTTGATAGCAGTGTAGTAATCGTATCGGGTTTTTTTTTCCCGGGGGTTTGCTCTACGGCTGTT

TTGGTAGCTGGTACGCTGTAGTAGCTGGTCGTGGTTTCTGCAAGCAGGATGGCTGCTCTTGAGGAGTGGA

CATCGGCTGTTACAGGCTTGTTGCTTGCAGGAGGAGTCGGTGTTGTCGAGTGATCTCACGGAGCTAATCG

GT

>NODE_117_length_631_cov_5.063934

GATGGGAGGGGACTGCCTCAACGTCGGCTGCGTACCGTCTAAGGCGCTCATTGCGTGCGCGACCAGGCTG

CACGAGGTGAAGACCTCTGCGGAATTCGGAGTGGAGATTCCGGGGGAGGTAACCGTAGACTTCGGCAAGG

TGATGAGGAGGATGCGCCGGATACGCTCGGACATATCGGAGCACGACTCCGTCGCCCGGTACTCGCGAGA

CTTCGTCAAGCACATCTTCCTGGGACGAGGCGTGTTTACATCGGCTGACACAGTAGAGGTTGCCGGGAAG

ACGCTTAAATTCGACAAGGCGATGGTGGCGACGGGGGCTTCGGCATCTATCCCCCCTGTGACCGGCCTGA

AGGACATGCCTCACATTACCAACTCCAACTTCTGGAACCTGGAGGAGCTGCCGCCTCGCATGGGCGTGAT

CGGCGCCGGGCCCATCGGCTTGGAAATGTCTCAGGCTATGCAGCGGTTCGGGTGCAAGGTAACTTGCTTC

GAGTACGCCGGCCAGCTCCTGCCTAGGGAGGACCCGGACGCCGCCAAGTGCCTTGAAGGGGCTCTGCTCG

AGGACGGGATAGACATCAGACTAGGTGCCCGCATCAAGCGTGTCGAGTGCGACGAGGGAGGGGGGCAGTT

G

>NODE_118_length_630_cov_6.110016

CCCCAGGCCATCTACCCCGACCCGTTCCGCGGCGGGGACAACATCATGGTGATGTGCGACTGCTACAAGC

CTGATGGCACCCCCATCGACAACAACACGAGGAAAGCGTGCGCCGAGATCATGGCTAAGACCGAGGCCCT

TGTGCCGTGGTTCGGTATCGAGCAGGAGTACACTCTCTTCGAGGCGGACGGCGTGACCCCCTTCGGATGG

CCCAAGAGCGGCTACCCCGGACCCCAGGGCCCGTACTACTGCTCCATCGGGTACGAGAACGCCTTTGGAC

GCCACATCATGGAGGCTCACTACAAGGCGTGCCTCTTCGCAGGCGTAACCATCTCGGGGACGAACGGGGA

AGTCATGCCGGGACAGTGGGAGTACCAGGTGGGCCCCTGCGTGGGCATCTCGTCCGGAGACGAGCTGTGG

ATCTCACGATACCTACTCCAGCGCGTGTGCGAGCAGTTCGGTGTCATCGTTTCGTTGGACCCGAAGCCCA

TCCCCGGAGACTGGAACGGGGCTGGATGCCACACGAACTACTCCACGAAGGAGATGCGCGAGGAGGGTGG

CTACAAGGCGATCATTGACGCCATCGAGAAGTTGGGCAAGAAGCACGACGTACACACCGCCCGTCGCACC

>NODE_119_length_629_cov_22.223684

TGTGTGTGTGTGTGTGTGTGTATTCTTCACATTAAACCTTCCTTTCTACCCGTCGACATCCAGCACACCC

AGTGTCCGTGCCGATTCTTCATCCTAAAAAGGTACCAGTTCTGTGGTCACAAAACTACGACACAGAGATA

CTACGACGCTCAGTGCAGTAAGCAGAAGAGACAATGCAATGCCATATACGACAGGCAATACCTACTCTCC

TAGCAATCCAGGTGCATAAGAGAGAAAAGGATAAAAACTGAAAAGAAAAAAGAAAAAAGAAAAAGAGAAT

AAAAGTAAACAAAATGAAGCAAATCAACAATCAAACAACCAAACGAAAGCATGACAAAATAAAAGAAAGA

AAGTGTCGTGTTTCTCCCAGAGGAGGTGCGACAGCCAATAAAATCAAATTAAATGAAACGAAATGAAATG

GACTGAAACACAGCGTACACAGGGCAAGACTAATAACGACAGGAAAGGACAGGCAGCCGGGGAACACCAA

AAGCACCTTCCAGGCAGCCGGGGCAGGTTTGTGCCCGTAGTACAGACAGGTTCTACCAGCTATTAATAGT

CGGTCTGGTACCTACTCTGTCTGCCGTCTCTTTCCTGCTTTCATCCGCCGTGGTTTCCCTACGAACCCC

>NODE_120_length_629_cov_7.075658

CCTTGTCACCCTAGAATTCCATAACACAAAGTCTTACTCACACATCGGGATGCCCCGTACGTCGCTTCCT

CCATCGTATGTACGTTTCACCATGGGTAGCTACCCCACTAATGCTCCCGAGTAGACCTTCGCTGCCGAAA

GCTTGGCCAACGATCCCCCTACACTTTGTACCGGCTTCTATCCTTCAAGATATCAAGATATCTACGTACC

GCTTGGTGCTCCGTCTCTTTGGTCGTGTGGTCGGTCAGTGGCAAGGCGTTTGTTTCATGAGTTCAATTGG

CCTGACTTAATGTCATGCATCATCCACCCCTCCCCTTCATTTGGCGCATGGTATGTGCACCAAGTTTCAG

GGGTTTAGGACGATGTCGGACCAGACTTCCACCGCCCAATGGAAATCGGTAAATTTTTCCGGGTATCTCA

CCACGAGACCCCAACCTTCATGTGAATTGAACATGTCGGGGTGACCTTGCTGTATCTATGTGTGTGCATG

TAAGGTGGTCCTCCTTCATGCTAGGTTTTTTGTCTATGTTTCAACGTTTTGCTGTGTATTACCTGGTACG

ATATGGGAGAAGATATACACTAAGTGTGTGGTTTGGGTTTTGATATGTGGAAAAAAAACTGGTTAGTTT

>NODE_121_length_624_cov_14.245439

AGATAAATAATAGGCCCATAGCAACGGCAGGAAAAACTAGCCCAACTAAAGGAACTAAAATTGAAGGTAA

ATAATTAATTAACATAATATATTTTATTAATAAATTTTGTAGTAAACGATACTAACACATAGATTTAAGA

TTAAATTTAAACTATACAAATGCATTCAACAAAGAGTGAAAATTTTTATAATCGTCTAAAAGAAATGCAT

AACTTTGCAGAAATCTACGCTAAACAAACTAATACCTTTTTTTGCTTAGACCCTTCAATAACTTCTGTAA

TTATTACAGGGTTAGCTACTTATAAAGATAATTATGGAGTTCCGTTATGTCCTTGTAGAAATTTTCGTAG

TGAAAAAGCTGAAATTGAACTAAATTATTGGGTTTGCCCTTGTGTTTCAATGCGTGAACGGAAAGAATGT

CATTGCAAATTATTTGTACAACCTAATGACTCAAGCGCCTCAGAAACTCAAAAAATTAGCCTAGATACTA

TTTATAAAAATTTATAAATCAGCTTTTTTTGCTATAAAAATAATAAGTGGGCCTGATACAATAATTAGTG

TTGCAGTAATTACTTGACCAATAACTTGCCAATTCATATGATTGTTCTCCTGAATAATTATCTA

>NODE_122_length_623_cov_13.408638

TGAACCTACATTGGTATCTTCATGAGAACTATGTTTTGCCAATTAAACTACAAACCGCCGTACTACAAAG

CACTTTTAACCACATCTTATAAAAAAGTGAACATATTTACTTAGCACTTTTTTTATAAGCTTTGATTTTT

TGTTTAACCCCCTTAATAAGTTGTGGTAAGTCAGCAAAAAAAAGAAAACCAAGAAAAAGTAAGAATAAAA

ATTGCAGAAAACTGATTCTCATATTATTTAGGACTTAAGTAGATAAACAGAAAGAGAGGGACTTGAACCC

CCAACCCTTGGCTTTGGAGACCAAAGTTCTACCAATTGAACTATCTTTCCTTGACTCTTCTTTTTTTTAT

GAACAGACTTCAAACCTCAATATATTATTTACAAGAACTTAAATACCGATTAGCTTACGCTATTTTCGGA

ACAATTTTTCTTTTCAGTACGACTTATACCTATAAACAAGGATTGATTTTTATATTTTTACCTAAAGGAT

TATCACACTTTGTTAGTACAGGATTAACTGAAATTTTTTTTACATACCTACAGATTTGTACTATTTTTAG

CTTTAGTTTTGGTCTCGGTATAACACTAACACAATTATACCTTTTTTTACGTCCTGGATTGTA

>NODE_123_length_623_cov_6.327243

TATAACCAATATAACCTGGGGGTGACCCAATTAATTTAGCTACAGTGTGACGCTCCATAAATTCTGACAT

ATCTAAACGAACCATAGAATCTTCTGCTCCAAAAAAGAATGACGCAAGAGTCTTTGTTAATTCAGTTTTT

CCTACACCAGTAGGACCTGAAAAGAAGAAACTTGCAATTGGTCGTTTCATATTTCGCATCCCAACCCTAG

CTCTTCGTACAGCTCGAGCTACAGCTGATACAGCTTCTTTTTGGCCAATTACTCTTTGGTGAAGAACACT

TTCTAATTCTAGTAATCTAGTATTTTCATCCTTGGTAATTTTTGTCACTGGAACACCAGTCCATAACGCT

ACAATTGAAGCAATATCTTGGGCTCCAACTTTTAATCTCTCTAATACTCCTTTTGGTACAATTCTTTTTT

GTGCTTTTATAATAGCACCCATTTGGGCACGCAAATTTAATTCATCATCTCTAATTTCAGTCGCTGTTTC

AAATCTTTGTTCTCGAACAGCTAAATCTTTATTATCTAAAATAGTTCGCAATTCTTTATCTAAAATATGT

ACAGCTTCAGGAAGACGATAATTTACTAAACGGACTCTTGCACTACCTTCATCAATTAAATCA

>NODE_124_length_619_cov_12.316054

ATTCCAGGGAGGTTGGACACGCGCGACGCCGTGCACGAGGTGTTGACGACGATAGAGCCCTTGTCGCCGC

TCTTGGCCATAGCAGGGATCTGATACTTGAAGCAGTAGACGAGGGACTTGAAGTTGACGTCCAGCATCTT

GTTCATCGACTCCTCCGTGATGTCGGCGAACGTGCCTACCGCGAAGGTTCCCGCGTTGTTGAAGGCCACG

TGCAGCCCTCCGAACTTTTCCACCGCGGCATCCACCATTCGTTTGCAGTCGGCCTCCTTGGAAACATCGC

CCGCCACAACGATGGCTTCGCCCCCCGCAGACGTGATCTCATCGGCAACGGCCTGCAGCTTCTCCTCTGT

CCTGGCTGCCAATACAACCTTGGCTCCCTCCTTGGCGAAGGCGATCGCCGAAGCCCTCCCGATGCCTGTC

GACGCTCCAGTCACGAGCGCCACCTTCCCTTGAAGCTTTCCAGCAGACATCCTGATGATTTTGTGGATGG

GACGTGTAGGGCGACTATATGTTCGATGAAGGTACTGCTGCTGTGGTACTGGAGAAAGGCTCTGATCTGG

TCGATGGAACAATGTAGGCAAGGGAAGTCGGCAAAATGGATCCGTAACCTCGGGAAAAG

>NODE_125_length_614_cov_423.448567

GCATTTCGCACAGGCAGGCGTCGTGGTGGCTCAAGACAGCAGTAACACAGAAACTCTCGTATGCATTATT

CCCATTGCTGTCGAGCTGGCTCGGCCCAAACCTCGAGCTGACTCGGCCCACGCATCGAGTGACATCGAGC

GGCTCATAAAACATCCCGATGTCCCATATCCCACTACAATAATTCCTATCAATCATGTGGAACTCCTACT

ATACGATGGATCTTATGCGTTGTGATACTGAGTACAATCAACATCCCCCCTGCTCATTATACACAACCAT

AATAATTCCCCCCTGCTCGGTGCATGACAGGGCACGACATTGGAGGTTCCATACATCGCAAAATCATAAT

GAACTCATACCAAACTCGGAAACTCAAACTCAAATCTGCAACTCCTCAAATCAAGACTCATACGATCTCA

AGTCCAAAACACATCAAATTAAGAGGAAACTCATCCCGACGACAAGTGAAAACCTCTCCCTCAACCCCAG

ACAAAAACAATGCAACTATGGAAAATGTGACAGTACGTAGTTACACCAAACAAATAAAAACTAAGAAAAA

GATAAAATGCCAGATGAAAATCAAAATCAAGTTAGCTTTTACCCTTTTGTTCTA

>NODE_126_length_613_cov_9.150338

TTCCAGACAATACACATCAGTAAATCCCGCTCAAAATAATTTTGAATATATCTTTGAAAATATTCTTCCA

CAAAAATTTCTATTTCTTCATGAGGTTCCTCAAAACAAAAAAAACAGTTGTTTTGGAACAACAATAAAAA

ATGATAGTTAATGTACTTGAAAAAACCTGATTTTATAAACTTCAAAAATTTTGGACTGTTTTGGAGAGTT

AATAAAAAAGACTTACTAAAATATTCTAAAATATATTTTTGAGTGCATCCAAAAATATATTGAGAAATAT

AAACCTCGCAATATTTAAGATTATAGTACTCACTAAAATTCTCTGACCAGAGGTGAAGCTGCGAATATTT

CTTAAACACAGGACTTACCGAATAAAAGTCGGCTAAATCAGCTTCACTGTATCTTTTAGCCCTCACCATA

AAACAAAACTAGATGATTACATTAAACCCCCACCAATAAATGGTAAGAAAAAGGAATAACCACACAACAT

CAACAAAATGCCAATACCACGCAGCAGCCTCGAAACCAAAATGGTGTTGACGGCTAAAGTGATCCCAATA

CAGCCGTAGAGTACAAATGGCCAAAAATAAAGTTCCAATTAACTCCAGACAAA

>NODE_127_length_610_cov_9.290323

GCCACCAAAACTCGCTTGCGCGCGAATGCCGCGTGCCCTTACACCCGCCTCCTCTTCTTCAGATGCATTG

ATTGCATTGGCTGCATCAAGGAATGCATGAATGTGCATGTTCGTTGTACCACAATACTGCGTTGTATGGG

CCCTCATCGATTCTTAGTGAGCTGCTTTAGACCTTCTGGACTTGTGACTTGATGTACTCAAGCTCGAGCT

CTCCCTGGTCCGTCGCCTGAAGGTGGATGCCGATGCCCATGAGGCCGGACTTGATCGCCTCCTTCGACGC

AGCGTACACCATCTTGTTCTTGATCTTGGCCGTGTCCGGGGACCACGAGATGAAAACCAGCTTGTTTCCC

TGGCGCCCATCGAGCGTCGTGAAGTCCAGGTCGAACACGCCGTAGCGGCACTCGTTCGCGGGCAGGCACG

CCGTGAAGTCGTCGTAGGTCTTGCTCTTGTCGCCGAAGGTGTCCACGACGATCTCCGCGTCGTTCTCTAT

CTTGTAGACGAAGTATCGGTTGTCGTGCGGCGCGCGGTGGAGCTTGAAGGCGTTGAACATCTCCACCGCC

TCATCGTTTACAGTGACACCGGTCGTCATTTTACTGTCCGTGCGTTGAAC

>NODE_128_length_608_cov_18.076661

CGACTACCTGTGCTGCGAAGACGGACGCCACCACTAGTTGGCCTCTGAAGTCGGCATGGACTGCACGATC

ACCTCTCAAGATAGAAAGAGGCCTCCCGAGTACGACACGGGTTGTGAAGACAGCTGCAAAATCACGTCTA

CGTAGTGGAGCGTAGCGCTTGAAGACAGTTCTGCGAGGCATTGGTTGCAGGGACCGCTTCTAGAGAGCAT

TGGGCGGCTCTGAAGACTACTGGTATATTCAGGCCCTACCGTGTCCGGCTAGTGGTGTCGAAAACGTATA

TCGTATTAGTACACGTCTACCGCAGACTCAGTCGACATTTTCTGCATATTATTCGGCTTGTTGTCTGTCT

GTCGCCGGCTACTTGCCGGGTGTTTCTTTGGACGGCAGCTGCTTATGTTTGTATTGGTGCCTTCGTATGA

TGGCTGTATTGCTACGGTAGTTTGGTGGCTGTGTTGAATTGGTTTGGCGTTCGGTGGGAGACGCGTCCGG

TGTTTGGCTGGAGGATTGGGTGGTCGTTTGGGCTTTGGCCATCATCATCATGGGTCTGGCCGGCTAAAAT

TCAATACAACATCAAACATTTCAACTCCAAAAAAAAAAAAATTTTTTT

>NODE_129_length_607_cov_12.107509

AAGCAGGCATGTTATATATTCCAACTTTCTAGATGGTGGATTCAGTGGATTGGATTTTCTTGGTCTGTTG

TTTTGTTTTTGTGCTTCTCTGTGTCTTGGCCGGCATATTTGAGCAATGACTGCTGTAGTTGTGGTGAGCT

TGTACCCTGGTATGATGGATCGCAGCTATGTGTGTGGTTGGATGGGAACTGGGTAGGAGAGCGGACGTTC

CCGTTTGACCCAGGCAAGTCAACAATTGGCTGGTATGTGTCGGGCGAAGACTGCTGTAGTTGTGTGATTG

TACCCTGGTATGATGGTGCACAACTATATTGTGTGTCTTGCTGTTGGAATCACGGGGGGGGGGTTGTGAT

GCCATGATTCTACAGCCAAGTGCACTGCTGTATGGAACCAGTTTGTGATGGACTGATTATTCAGCAAGCA

GCAGTGTATGATAATTTCTGACTGTTTGATTCTACAGCAGCAGTGCATGATTTGTTAGACTGCTGCTGTG

TACTGGTATACCGTATTTTATGTATACGTGGATGCATGACACCCGTACTGCTGTGTGTTGCTTGAAGCAT

AACAGGGGGTGGGTTGCCAGCTTGGTTCTCAAGTACATGATATTTAA

>NODE_130_length_606_cov_63.811966

GAAAGGCCAATCAAACTCGGAAATAGCTCGTACTCCCCGAAATGCATTTAGGTGCAGCGTTGATTTATAG

TTTTATAGAGGTAGAGCTACTGATTGGATGCGGGGGCTTCACCGCCTACCAATTCCTGACAAACTCCGAA

TGCTATAAAATGTTACTCAGCAGTGAGGGCATGGGTGCTAAGGTCCATGTCCGAGAGGGAAAGAACCCAG

ACCATCAGCTAAGGTCCCCAAATGTATGTTAAGTTGAAAAAACGCGGTTGGATTGCTTTGACAGCTAGGA

TGTTGGCTTGGAAGCAGCCATTCATTTAAAGAGTGCGTAACAGCTCACTAGTCGAGCGGACGAGCATGGA

TAATAATCGGGCATAAACATATTACCGAAGCTATGGATTTAGACGCAAGTCTAAGTGGTAGGGGAGCATT

CTAACAGGGTTGAAGGTGTGTCGTAAGGCATGCTGGACTGGTTAGAAAAGAAAATGTAGGCATAAGTAAC

GATAATGCGGGCGAGAAACCCGCACACCGAAAGACTAAGGTTTCCACAGCTATGCTAATCAGCTGAGGGT

TAGTCGGGACCTAACGCGAACCCGAAAGGGGTAGTGGATGGACAAC

>NODE_131_length_606_cov_7.094017

TCCAAAAATTTTTTTTTTTGTGCGGAAAAAACGAGAAGAAAGTGATATTATTATGCCGCTTGCCGTCCTT

TTGTCTCGGCATCCATCCCCATCCCTACCTGCCTGCCACCCAAGATTAAAAAAACGAAAAAGTGAGTCCA

AAACAACCGAGCGTTGTTACGATCGACCTTCCCCTCCTTAGCTAATTACAGACAGCTATAAAGTCTGTCT

TTTCTGCCTGTTGTCCGGTCATCTGTTCGTCTAATACAAGTACAAAAACCCCCGACATCAAGCAAATGTG

TTTTTAAGACTGAGCCGCGAGTGGTCCCGACTGACCGCCCCCTACGCTAAAACTAGCATATTAAGGCCGT

CAAAAGAGGCTAGGAGACCGCGGTGTGACTTTCTTGACTGACGGGTCCTACACTACCTAGCATAATAAGG

CCTCGGCGCGTCCAATTTGATCGACCGACAGCCCTTACACTATAACTAGCATAATAAGGCCTTGACGCGT

CTAAAACGACTATGACCCCCCGTGACAGCCTAGTTGCCCGTCAACCCACCGCCTCATGCAGTAGTTCGTC

GTGTACAGGTGACAACGCAGACCATACACCGAGCCCGTTTTCTCGA

>NODE_132_length_605_cov_4.890411

CTAAGGTGCTCCGAAGATGCTTGAATGCCGAGTACCACAAATCGCGAAATTTTGCAGAGTATCCATGTTG

TTGATTACACCAACGGGGAGCGCCATAGCTTCACACCGTTGGTAACTGGAGTCCAGACAGGCAGAGGCCG

TGCCGCGTTCATTCTGTGAACGAAGCAATAGTTCTACGGGTTCTAATGAAAACTCTGTGTGAACATGCAA

ACCGCTATCAATTACTCTCGGAGTACAATCGCGGTGTGAAAAATCTACCGGTGGAGCCAAATTTCCTTTT

ACCGTACATGCAACAGTACCGTCATAATACACCTACTCCGACCCAAAAGCCTGGCAAGCTGAAAACAAAA

CCTCATACACCCACCAGCTTACTCAATCACCCTGACCGAAATGGGTTCAAGTCGACCCAGACCTAGTCAC

AGCAGACAGCGTGTGTCCGTCTGGTCTAACCGAGTTGCCATCAGATAAAAACAGCGGTTGCCCTGCGAGG

GGCAGGGTCAGGCCCGATTTCTCACTCTTGCCCCACCGGGGCACAACCACGCCCCGTGAGATGAGACCGG

CAGGCCTAAGATTTTCGACTCTTCCCTGCAGTATGAAATACACCA

>NODE_133_length_604_cov_172.473413

TCATAGTGATCCGGTGGTTCTTTGTGGAAAGGCCATCGCGCAATGGATAAAAGGTACGCCGGGGATAACA

GGCTAATGATCCTCTAGAGCCCCTATCGACGGGTTCGTTTGGCACCTCGATGTCGACTCATCACATCCTG

GAGCTGGAAAAGGTTCCAAGGGTTCGGCTGTTCGCCGACGAAAGTGGTACGTGAGTTGGGTTTAGAACGT

CGTGAGACAGTTTGGTCCCTATCTTCTGTGGGTGTTTGAAAATTCCGAGGACTAGACCTTAGTACGAGAG

GACCGGGAAAACTCGATCCCTTGTGTTCCGGTTGTTCCCCAAGGGGCAGCGCCGGGTAGCTACATCGAGA

AGATATAATCGACCATTAGGCGAGAAACTCGCCTCTAGTAAAGTTTTCGCAAGGGGGTGGAAGATTACCA

CTTAGATAGGCGGGAGGTGTAAGAGCTGTGAGGTTTTCAGCTGACCCGTACTAATCCCTAAATAAAATTT

TTATTTACAGACTAAAGTTTAACAACTTTTCGGGCGTATAGCTCAGTTGGTTAGAGTGGTGGACTTTTAA

TCCGAGGGTCTTGGGTTCAACTCCCAATACGCCCACCATGGAAT

>NODE_134_length_600_cov_12.977547

TGGAGTTCCTTGGGAGCCGTGTATTCCAGGGGGGGGAGATACTACTGTAGAATAATGAGAAGAAAAAAAC

ATCTCTATTTTGAGAGATTACAGCAGCTTCATCGCCATAACAAGGGCGAATTTCGCATCGTAAAACTCTC

GGGACGACGGCAGTGATAACGCGACTAAACAAGACCAGGCCCGGCACCGACTGACAGTGCTTGCTGCGGC

TTTGCTTGAAACGTTTCTATTTGCTTTTGACGCCGAAAAAGCTGCACACGCGGGTACGTACAATCTATAA

AACTCCCCGCGCGCTGAAACCCCGTCGACGCCGATGCCCAAAAACTCCCAAGACTGCTACATACGATGTT

AGGGTGGAGGTACGCGAGATACATACAGCGAGTGCTCACACACCCGTTTCAGCGTCCGCTTAACCTGTAC

ACGCACTATAGCCGCGCCAAGAGAGGCGCTATCGAGTGTTGCCTATCACACCAGCGTGGAGGTTCGAGAA

GAAGGCCGAGCCTTCTTATGCCGCGCTGCCACCAACAGCAAATATATATACCCTAGCGCTCCCACACGCC

CTCCTCCCCCCTCCCAAACCCAGGAAAACGTGAAAACCCG

>NODE_135_length_600_cov_7.153713

CAAGACACCGCCTCCACCGGAGTACACTCGCATGTGCCATCCCTGATGTCCTCAGAGTCGAGCAAGACGT

TAGTAGCAGTGCTGGAAGGGAGGCACTGGAGGTCTTCATTGTCGAAGAGCACAAGTCGCTCGAGGTTGAC

CAATGGGGAGAAAACGCCAGCAGGCAAGGTGGTCAGGCTGCTGCCTTCGAGATTCAGGTTGTCCAGGTTC

ACGAGCCCTGTGAAGGAGTCTTCGGTCAGCAGAAGGTCGTTGAAACTCACCAGCAGCGAGCGAAGCCCAG

TGATTCCCGACAGCGAATCTCCAAAGCCCGTCAGGTTGTTGCCGGTCAGGTCCAGGGTACGCAGTGCCGT

AAGACCATCAAACACTCCGTCGGGCAAGTTGTTGATGCCGTTGAGGAAAAAGCTCAGGTCCTCCAGCTTG

GTGAGTCCTTTGAACAGTCCTTCGGGCAAAGTCACTATCCCATTTTCTCCGAGACCCAAGTCGACGAGCT

CCGTTAGGCCGTCAAACAACCCCACCAGCTCGGTCAAGTCCCGCAGATTGTTGCCGTCAAGGAACAGTTC

TCGAAGTTGAGGAGTCCCAGAGAAGAGACCAAGGGGCAGG

>NODE_136_length_596_cov_8.478261

ATGACGCAGCGCGTTTTGCGCTGATGAGCTGGTTTTTTGAGGAAAACCTCACTGATTTCACTCCCTACCA

ATCAAGATGCCCCGCCACGAGGACGGAGAGGAACACAAGGGCGGAGGGGCCGGCAGACACGGCAAGGGCC

ACGATGACGAAGGCCACGGCAAGCCCGGACACAAGCACGAGGAGGGAGAAGGTAGGCACGGCAAGCACGG

CCACGGCGGCGATGACGACGACGACTGCGACGACGACGACGACGACGAGGAAGACGCGAACGAGGAGGCC

AACGCGGAGGAGGCCAAGCACCCGGGCTGGAAGAGGGCCGGCAGGATCGCCTCCAGGAAGTGGAAGCGCC

AAAACGGCGTCGACCCCAAGACGCACATGAAGGACCTCGAACCCGAACAGAGGAAAGAGATCAGGGTTGC

CAAGAGGGTCGCCAGGAGGCACTTCGTTGCTGCCCAGGAAAAGGCCTAAATGCTTTCCTTCTCGGACACG

CAACATAAGTCTAAAACTTCGGCGATGAAACTGATCGGCGGCCAATCGAGTCTGCCCAACGAGCAAGCAG

TAGTAGTAGTGGTAGCAGTAGTAGTAGTAGTAGTAG

>NODE_137_length_596_cov_3.728696

GGGCAGGCGGGGAAAATCCCCATAAAATGGAACAACGCGGCTTCCCGTTTCAGGAAAGCCGCGCTGCGCA

ATTAGCCCCATTGAAAACATCAACTCCGGGGAGAGACGAGACCTTTCGCATCGGCAGATGGTGCCACAGG

GGCACGACTACGCGCCATGAAACCGGTCTGACTTAAGCCATTCCAGGCTTTCCTACAGTATGAAATACAC

CACCGGTGTTTACCTGCCACCGGCACACAGCTAATGACGTGCCCGACGGTGCGGTTGTGAAACTACCAAA

TAGTGCACGGGAAATAGCAGAAGGGCCTTCCGGCGGCAATGTACAAGATTAACAAGGAATATTTAGTGAA

CGCACTTCCAAGACTTTAGAAAACAAAGGGTCTATAGCTCCACGCCAGCAAGCAGTCCGTCCGTCACAGG

GCATAACAAACACAGCAGTGGCTTCGAAAAAAATGGGTTGGGTCGGTGAATGGTCACTCACGCGGCAGCA

GCAGCTGCTGTTTGCTGATAAATGCTGCACGTGGTGGATTTGATTTTCTTTCTTTTTTTTGTTCGTCTTT

TTCTTGTTTTCTTCCGTGTTTAGTTGGAATGTAGTT

>NODE_138_length_594_cov_23.785340

AGCGCGGAGGCGCGCCAAGGTAATGCCTCTAGCGAAGGGCAGGACCAGTGCTACTGTCGTTGTTATCGTG

GTTACTGTTGCTGCTGCTGTGATCACTACTGCCTTTGTTGTTGTCATTGTTGTTGTGGTGTAGGGTGGTC

CCTAAGACGGGCCCAGAGCTACCCCTGTTGTGATGTGCGTAGGGATATCCTTAGTTATCCTTAGATGGTG

AAGCCACCCTCGGGCTCCCAGACCACTCCAAGCTCAGAGAGCTTCTTGTGTGCCTTCTTGTAGTCCGCGA

AGAACGCGTCCTGGCTGTCCTTGTACTTTCGGGCAAAGGGCTTGAAGCCGTCGTCTGTGAACAGGCACTT

GTCCGTCGCCAGCTTGAGGATCTCGGGGTCGGAGCCCTCGTCCGGCATTTCCGAGAAGTAGCTGTTGTCA

AACTTGAGCCAGTTCTTGGTCCAGGCACTGCCTCCCGGGGTTCCTACGCCAGAAACCCCCGCACCCCTCG

GGCACATGCCTTCCGAGGTGATCCTGGTGCACTCCGCCCCCTCACCGCTGCGGTCCTTTTTGGCCCGTCC

GATTGTGTGGGCGGAGCACCTCAGATTTCATCGA

>NODE_139_length_594_cov_8.155323

CTTGGCCCCGAGAATTCCACATCAACTACTTCATCGTGCCGCTCTCCATACTGGGTTTGGGCCTCCCCTG

GGCTATTCTCGCCAAAATCATTGGAATGGGATGGATGGGGACCGTGGTGTGGCAGACGTTGGCATGGATA

GGAGTCAGCGTGGACGTCATGCTCGGGGCTCTGTACATCGCCAAGTGCTTCCTCTACCCCTCGAAGGTGA

AGAAGGAGTGGTTCCACCCCATACACCACAACTCCTTCGCGATTCCGTTCATCACCCTCATCTTCATGGC

GTGGATGTCGACAATGTGGTCGGGAGACCCCACTGGAAAGATCGCCATGGTTTTGTTCTGGCTGGGGGCG

ATCCCTCTGTCTTCGATCACCATTTACACCATCGCCAGCTGGATTGCTTTCCCTGTGGACGAAGAGCACA

TTCACCCCAGCTGGATGGTCATCCCGGTCGGAAACCTTGCAGGGGCCGTGACAGCAATGCTCGTTGACCG

CGACTACATCGATTGGGGCTGGTTCCAGTTCAGCCTCGGCACCCTGCTTTGGCTCGCGCTCTGGCCGACC

ACCTTCCGCAAGGCTGTCTTGGGCCACAACAGCA

>NODE_140_length_593_cov_4.924825

GCACCCAGTTCATCGACAACAAGTTCGAGGATGTCACGACGATCCGCTTCGATGACGCTACGGGGATCGT

CATGTCGGGAAACACCGGCCTCGACGACGTCGAGCTGAAGGTCGCCAACGGCGCCTGTTTCGACGAAGCT

TCCGACTCCGAATTCACCCCCACTTGCTAATTCGTGTTTAGCAAGCTGCCGGTATAAATAGTTGGTCTCG

GCACAATGTGCCACTCTCCGGCGAACAGCATCCGCGTTTCGCATTTTTTTGGTTCAGTGGTAAATTTTCC

CCGTTGCCTGTTGTTGTGTCGAGTTGTGTTTATGGGAACGCATTTTTTCAGTTTTCGGTTCGTTGGTTTC

CCCCGTCACCCCTTTTGTTCGTGTGTTGAGTTGTGCTACTGAATGTGTTTTGTGTGTGTGCTCGTTATGT

TGTTCATTGATTGGCATCCCCGGCATATATATATATACACTGCTGCTGTAGTAGTCGTGCTGCTGTCAGC

AAAAAGTGAAAAGCTAAAAAGCTATTTCGAAGAGCTGTCTATAGAAAGTTGGGCCGTTCGTCACGAAGAA

GAAAGAGAGGTGTATTACGGTCCACTCATAACT

>NODE_141_length_592_cov_326.914186

TTCTATGTCCGACGGTCGCGAGCGCCCCAAGTGATCCTGCGTATTAGTGCGGAAAATCGCGAAGAGGGAG

CAAAAAAGGATTTGAAAGGCTTGTTCCTTTCGTATCCGCTGGTATTTCGGTGAGGTTTGCGTAGGTTGTC

GGCTTCCGACATCGCTAGTAAAACGCGCAAACCACCTGGTCCGTACTACGTTCGCTCGCCGTCCCGACGG

AAGGCGAACGCAGAGGTCCAACCGTGGGCCCTTTTTGGCACGTAAGATTTGGAATTGCACGCCGAGAATC

TGATTAGTCGGACGCGTACGCCTTGTGAATCTATCTGAACGTGTTACTGAGGGCTCGTGGGTAAGCAAAG

GGAATTCCATGGCTGCGTGCGAGTTCGACGGCTGCTAAAGCTGTCTGGGTTCGTTCCGCAGTCTTCCTTT

TCGGTGTCATGTCTCGGCATGCATCGATTGGTTTGAGTCGAGAAATATAAAACATGACTCCTCGGTGCGC

GGTGCTTCGGCGCCGCGTCCCCTGCGTCCCCGTGGCTACTGATGAATTTCATGTGCTGCCAGGGCGAAAT

CAAGTTGGCAATCTGGTTGATTCTGCCAGTAG

>NODE_142_length_590_cov_223.630931

GTTTCAGTTCCCCTCGTTTGCCTCTACAACCTATGTATTCAGTTGCAGATAATGCATAAATGCACTGGGT

TCCCCCATTCGGACATCTGTGGATCAAAGCATTTTGTCGGCTCCCCACAACTTTTCGCAGACTTACACGT

CCTTCATCGCCTCTAACTGCCTAGGCATCCACCGTATACGCTTAGTCACTTAACCATACAACCCTAAGTA

GTCTTTTCTAATAAAAGAAGCGAAAACAAAGTACACCGTGGAGACTAATCCACAGTAATTGTAAAGTCTG

ACATTTTCACGCACACATTGTGTGTCTTGAATAAGAGTTGATATATTCTTACGAATATATCGGGTCGATT

ACATTTCCGAGGAGGAAACATAATCACCATTATTACTTCCCGATAAAAGAAGTAACAACAGCTTGGTATT

TATAACGTACAAACAACAGCGCGACACCGTGTTTGTCATATAAATACCGATATTTATATCAGCTTTCCAG

ATTGTTAAAGAACTAATTTTTAACTCACATTCGGTTAAAAACTTGATTACCTCTCCTAATGGAGAGTAAA

AACCAAATTTAAAAATCTCGTGACAAAAAA

>NODE_143_length_586_cov_3.734513

TAGGACGGGGTGATGACCTGCTTCAGGGAGTACGTCTCCCCCGGAGCAATGCTGTGGAACTTGCTGACGA

TTCCGGGCTCGATGCAGACCATCTTGTGGTACTCCTCGTCGCCGAAGTCTCCCAGGCTCTTGGCTTTAGC

GATCCACGGGTTCCACAGCACCACGTCCACCGGCACACACACCTCGGAGCCCTCCGACGCTTTCTTGACG

CAGGAGTTGTTGACCTTCGTGCCGCCCGCCACGCTGACGTCACCGATCTCTCCCGTGTAGATTCTGTCCA

CTTCGCGCGCGAAGACGGCCTGGGCTCGCTCGTCCGTGCTCACAGCGTTGCCCTCCAGCTGGTCGATGTA

CTGAAGGTTCTGGAAGCCTTCGATCTTAACATCCTCAATCTCTGGCACCGCGATGTACGTGTGGAGCAAG

CTCTGGAAGTCGAATGCCTCCTCTCCGGCGTTGGTGATGTGCAGCGTCACTCCCAGGACATCCTCGGCAA

GGTCCACCTCGTAGCGCAGGCTGAATTTGTGTGGCCATTTGGCGAGCGTCTCGTCCGAGTGCCCCAGGAC

AAAATTGGCCTGAAAAGACAACCGAC

>NODE_144_length_583_cov_9.478648

TTTTAACTTCACCAGCTAGTACTCCACCTTGCACAGCTGCACCAATTGCAACAACTTCATCTGGATTTAC

AGTCTGGTTTGCCTCTTTTTCTACTATCTTATAAACCAAATTTTGAACAGCTGGTATACGTGTTGAACCA

CCTACTAGTACTAACTCATTAATGGTATTTCGATCAACACGAGCATCTTTAATTGCTGCTTCTACAGGTA

ATCGACAGCGGTTTATTAAATCTTCACAAAGCTCTTCAAATTTACCACGTGTTAATATTTCCTCTATATG

CTTAGGCCCATCTTGATTTGCTGTAATAAAAGGAAGATTTATAGTTGTTTCAGATAGATTGGATAATTCA

ATTTTAGCTTTTTCAGCTGCCTCAGTTAATCGCTGTAAAGCTTGAGGGTCAGAAGCTAAGTTAATACCTT

CTTCCTTTTGGAATTTTGCAAGAATAAAATCAACAATTTTTCTATCAAAGTCATCTCCACCAAGTTTAGT

ATCACCGGATGTTGATAAAACTTCGAAAACACCATCTCCAACTTCTAATAAAGAAACATCAAATGTACCA

CCACCTAAGTCAAAAATAATAAC

>NODE_145_length_581_cov_5.808929

ATATAAGGATACTTCTCCGAATTCAAAATGGATAAGAATATAAAGCGTTTTAGTTGAGCATATTTTGATA

ATTCGATTACAATTAATTTACCATACCAGTCTACATGTATTAATTCAGTATTATCTTCAGGTTTTGTAGT

CGATGCATCAATTATAGCTGTCACACCTTGAAAGGAAAGTGGTAAAGTTTCTGGTATTGTTAAATCACCA

TAAACTAACTCAGCACCCCATTCTTTTAAAAAATTGGCCGCTCTTTTGTTACGAACAATACAACGAACTT

GAAAACCATTTTCTAAAGCTTTCCTAACAATTTGTCGGCCTAAGGTTCCGGTTCCTCCAAGAATAAGTAG

TGTCATAAATAGGTTATTTTTTTGTAAATTTTTAAGACTTGTGTCAGATATAAGATAGAGTTATAGTATA

ATACTATATTACTCTCTATATAACAAAATATAAAAGTCTATACTTTTTATACTTTACTATATAACCATTT

GTAGAATTCAATATAAATATCGATTTTTATTTTATATATTATAAACTATATGCTATAGGAAAACAAATTA

GTTCTAAATACGATTATATTG

>NODE_146_length_579_cov_4.616487

GGATGACGGTACCTATTTATATTGGCAAGAGGTCGATGACGACGCTTTTGGCTACCCAATACTCCTATAA

TACTATCCCACTTAGGACGAAGAAACGTTTTCTCTTTAGATAACAAATCGCCCCAAATATCAGTTCTAGA

CCATAAACAAGATTTATATCTATGTTTAAATCGCATTATTTATTATTACTTTCCCTTTTAAATTTAAGGG

AATTTTTAGGTTCTTTTTTTACTTCTTTTAACTGAGCCTTACCCAACAGACTAGAAAACACATGTCTTTT

TGTTGTTGATTTCATACGGTTACCTTTAACACAAGTCATGTAAATTAAACTAAAAAACCAAGAAGATAAC

AAAGGGGATTCAATATTTAAATTAAAAGGATAAGCTACTTTACTCATAGTAGGGCTCCCAACCCCAACCA

ACAACAAACCTTTTCTATGGGCCTCCACTAAATTTTCTTTTTCTATATCACTTAAAAACACCAAATCAAC

ATCTTTTGATTTAGCTAAATAACTACGTTTCCATTTTATATAACTTATATTAGGTTGATGAGAAAGACAA

ATTTTTAAAAGAGGAGAAT

>NODE_147_length_577_cov_23.908273

AGTGACGAAGCTGGCTAGGTGGAGTACTCCCTGCGCTATAGGATACCACCACAGGCATTCGAAGACACAT

CAAACGACGTCGCGTCCCTTGGTATGCATTGCCGCCAAAACGAGAGCCCTCTCTAGGCGAACCGAAGCAA

AGTGTTGCGATCCGGAAGCCGACCGTTCGATTGCGCTTTGTCTCTTAAGGACTCGTTGCATATCACGGCC

GAACCTCCCTGCACCGACTGCTCAGCCTTCGATTGCCCCGCCCACGAACATAAACGCCTCCGCGCAAACA

CTCAACACCGCCCGCGGCGGACACCGTACGTCCACGCGTTTGTGGTTGCTTTGGCCCCGATCCTTCACGA

CGCGGGTCTGCGCGGTCCGCATGTGTATCGTTCCAGCATCAAGCCTGCTCAGCCTTTCCATCACCTATAT

TTCCTGAGGCAGTTGGGCCGGTCTTTCTGAGTCTTCTTGGTGTTCTCGTTGTACCAGACTTTCCCTTGCT

TGGAGTCGTCGATCTTGAACCACTGGTCCTGTTCCCTCTTCTTGGTTTTTCTCTCCGGCGTTTTTGGCGG

CGGGGGCTGCGTTTTTG

>NODE_148_length_576_cov_4.508108

CTTCTTCTTCCGGGGTGAAGTCGTTCACGATGTTGAAGGTTTTGCGAATCTCTTCCGCTGTCTTGCCCTT

GATCATAGAGGCAACGGTCGCGCACGTAAGGTCAAGGAGCGGCTTGATGTCCATGTAATTCGCCGCCAAG

ATCAGTTCGAACAGGGTCTCCTGCTGCACCTGCACAAACTTCGCGTACCACTCCTGCACGACCTCGTGCA

TGTTTGCGCTCTTCAACGGCTTCTCGATCTCGTTCATCGGGTCCTCTTTGTAGTGAGCGCAAAACTCGAT

CACCTTTGACAGCACCGACGCCTTTACGTTAGGCAAGGGGATCTCCTGCGCGTCCCCGTCCTCCTCAGCG

TCCCCGTCGATCATGGTCTCGACGAGGTTGCTCATCTTGGCCACGCTAACCTTGATGTCGAAGCTCTCTC

CCTCCTGCGACACGAGATGTACCGAGCGGTTGTCCTCTGACTCCTCGTTGGCCGGCGCGTCGGCCGCCTG

AGACTTCTCCTCCAGTGCCGACATTGTGAGTGAGGTTTTTTTTCTATCGGTTTTCACTGAGGTGGAGGGG

AAGGCGACGACGGGTT

>NODE_149_length_574_cov_4.862568

GGTTTCGGAAAAGAAAACGGAAACGATGATCCTACGAACACGAGACCAGACACCCCCTCCTCCGCCGCTC

GTTATCGAAGCAGCTGGTCAGAGGTATAGACAGACGACCCAGTTCTTGTACCTCGGCGGCATTATCCACG

AAGATGCTGACCTCTCACTTGAAATCGAACGACGGATCCGTCTCATGTGGGCGTGCTTCAAACGGTTCGG

ACCAGAGTTGTACGACATGAAGACGGCTCGACTTAGCTTGAAAGTCCGCATGTTGAAGGCCGAGGTGATA

GAGACCCTGCTGTACGGATGTATGACGTGGACCCTCAGTGCACAACACTTCGCCAGGCTCCGATCGGTAC

ACCACCAAATCCTCCTGCGTGTCATTGGCTTCCAGCGTCGGCAACGTGGCGACTATAGGACCCTCTCGTA

CGCAAAGGCTCTCAAGCAGACACGATGCGAGAGCATCGAAACGACGATCCGTAAACGATCGTCGTAGATT

GTCCGTTCGTCATGGCTGCTCTGCTCTGCTCTTGCTTGTTGTTTGTCTCTGATATGCTGCTGCTGTTGTG

CTGTCCTCGTTGTG

>NODE_150_length_574_cov_4.582278

TAGTGCGACCATGAAAGAGTGTTGTGCTGTGGTTTGGGCTGTTACCCATTGGCGCCCATACTTATGGGGC

AAGCATTTCACTTGCTGTACTGACCACCAAGCACTTACGTACCTCTACAAAATGCAGGACACCTCTAACA

TGCTGACCCGTTGGGCCATCGCACTTCAGAATTTCGATTTCACTGTCAAACATGTCGCAGGCAAATTAAA

TGTTGTGCCTGATATGCTGTCTCGTGCCTTTAGCGAAGTCAACGGGGAACCCGTCCCATCCGAGCCCAGA

TTGGCCGCGATTTGTCGTAACGTACCGATAGACGGACCTTACTACCCACCTGGACCCCGCGATTACGAAT

TATCCGCGTCTAATTTGCACGATGTAGCACCTGTCGAAAGCGATCGTGAATTGTTCATGAGCGCAGTGTC

TGTCTTTCCTACTGTAGATCCAGCGAAACTTGTCGACCTACAGAAAGAAGAATTTTGTCAGTACTTTGAG

TATCTCCGTGCACCTGCATCGGCCCCACTTCCTTCTAGAGAATCTAAGAACACGATGAGCTATTTTTTCT

TGAATGAAGGACTG

>NODE_151_length_573_cov_7.494565

CGGTGCTGACCTTGGACGGTGGTGAGAAGAGCGGCGGTCCTGACTCTCGAACTTTCTTCAAACAGGTAGC

AGCGCGTTCGCCTTCCTCAGGACATGTAGAACTCAGCGGTGCTGACCTTGGACGGTGATGAGAACATATC

ATTCACACAAAACACCGACAAGCCTATTCGACAAACAAGACCTCGATAAGGAGCTCACCTTCGCCGACCC

GTTCGACACGCGCATGGCAGGTGCTTACCTTTTACTGTCAAGCCGCTACCGCCGTGATACACTACCATGT

TGCGTTTTATTACCCCCCACAAATCACAGAGTCCAAAGAGCAACCGCAAGCTAGTAATAATACCGATGGC

TGACATACCCCATAACACACTCGGAGCCCGTACTTACCTGCTTTTCCTCCTACCTCACCTGCGTCTGTGT

GAGACGTTGTTTCTCACTGTGCCTTCAACAACCATTTATGCGCAAGTTAGCATCCATCGACCCCTCTTGA

GTTAATCGAGATAGCTCTAGTCTCCTTGTACGGATATCATATTTGGATACTTCCTTCTCTCGATGATCGA

TGAAAATATGGAA

>NODE_152_length_572_cov_5.165154

TCGGGGTGACGTGTCTGTGCTATCTCTGTGGTCTTGTTTGTTTGAGTGTTTGTGCTTTTGGCTTGTCTCT

TTGTCTTTGTCTTTTTCCCTTCGCTGTTTTTGACCTTTTGCAGTTACACTGCTCGCAGGCCGGGGGTACA

TCAGTAGCACGGGACCCTAGTTCCCTTCCCCCGTATCATGTCTGCGTTTTATAATCGGTGTGGAGGACTC

GATTTGTTTGCCTGGCCGGCGTGAACCCGGCACGGAACACCCCGGCATTATAGGGGCACCAAGATAGAAT

CAGGGGGCGTATACCCCCGATTGGTGATGTTGACCTTTCCAGTTTGATTTTTACCTTCACATAGCGTGTA

GAGGGGAGCGGAGGGCTCCAGACTATCTGTTTTTGACTGACTTTGATCGCTTTGCGGACCCTTCGGGTTC

CACTTTTATTTCAGTATACCTTTTTGGGGTAAGTTAGCCTGTGACCACGGGAATCGACCGCGTCAAGTGT

GGATTAGTACGGACAGATGGAGGTACTGGATAGAGAGACCGGCACGGCTTTTAATGTGAAGACAACAACA

ACAACAACAACA

>NODE_153_length_571_cov_110.550909

TTGTAGTGTCATCACCCCGCTACCAGGACCAAAAAATCCAGTAACATGGAATGCACCAAAACCAAAACAT

AATAAACCAGATAAGAATAAATGTATACCAAAAATTTTTGGTAAATCTAAAGAAGGCTCACCTGTTCTTG

GATCACGGAATAATTCAAGATCCCAATAAACCCAATGCCAAACAGCAGCTAAGAAACATAAACCTGATAA

TATAATATGACTATAGGCTACTCCTTCGTAACACCATAAACTTACAAGTGGTTCAGATCTTTCTCCGGCA

ATATCCCATCCTGTCCATGAGTCAGAAACACCTAATCTAGTCATAAAAGGCATAACAAACATACCTTGAC

GCCACATTGGGTTTAAAATAGGGTCTGAGAAATCAAATATAGATAATTCATATAATGCCATTGAACCAGC

CCATCCTGCAACTAATCCTGTATGCATTAAATGAACTGCAATTAATCGGCCTGGGTCGTTAAGAACTACA

GTATGAACACGGTACCATGGTAATGCCATTTGTTATAAGCTCCTATTAAGTGAACATGTTTTTGACTTTC

TTACTATGCAA

>NODE_154_length_569_cov_29.662409

CACGCTTCCGTACATATCGCCAGCTTTGCGACAGCCAACGTTTCGTCATATATGCCACATACACTAAAAT

GTATATTTAAGTTGCTCTACGATGGGCCATTGTCCGGGTATACCGGCTTCAACTCGCAACCGTAAGGGGG

ATCGTCCCGACCCAGTCGATTTGGTTCTACGATTCCCCCAATAGACGAAAAATGCACTCTCAAACCACTG

TGTTCCTCCTGTGGACACCAAACATATCGGCCGACGACAATTTCAGTCAACTGTTCACCAACCCTAAGCA

CTTTGACCGTGCATCCAGTCACCAAATTTACCGTCACACAATACTCTGGACCGGTTGTATTCACCTCCAG

CACTGAAAATATCGGCAGTCATATTCCTGCGTCGGATCCAAAGCTTGAGTATACTACTCGCTTACGGCGA

CAGCACTCTAGGCGAGACAAAAAACGACCAAAAAAACGTAAAAAACAAAACCACGTGTCGGTGTCCCAAT

GCAACGATGTCCCAGGCCTTGAAGGAATCATTGGTGGTCTCACACCACCCAGAACTTCACCTCGGTGATG

CTCAGCCAG

>NODE_155_length_569_cov_10.589416

GTTGTTGTCTTCACATTAAACCCGTTCTTTCTACTCAACCACGTCAGTACACCCAGTGCCTGTCCTGATT

CCCAATCCTAGAGAGATACAAATTATGTGGTCACAGACTGTGAAACAGAGAAAAACCAAAACAAAATAAA

CGCAAATGAATGGGTAAAGCAAGTGAGATGCTTAACGTACTGGGTACGAGAACACAACGACACGAGCCGA

ACCAAAAAATAAGAACAAAAAGACAAATAAAAAGAATTCTGAAATGAAAACAGAATACAAATGACGCAAA

TGACGCAGGGAGACAGAAGTGCCCCTAGAAGAAGAACGCACATGCCCAAGAGGAACAGATAACTGCTGCA

CGGGCGAGGCAACACGAAAAAGAGGAAGACACCGGCATAGGGTGGAAACAAAGGGAGAAACAAGGCAAAA

AAGCAAGAAAGAAGACACGAGAGAGGAAAAGAGAAAAACCGTTCTCAAGGCAAGCTCAAGGCGATGGTAC

TAACAGGTTCCCGTCTCTGCCATAAAAACCGATACGGGGACCATGTCACACCTTCTGCCATCTCATTCCC

ACTCTAAAA

>NODE_156_length_566_cov_9.394495

ATTCCACTTTTTTTTTTTTTTGGTTGTTATGTTTTTCTCCAAGTTCACGAGTTTCGCTCCCGATTACAAC

CGAAAACCCCCTTGCGAGGGCCCCCGGCCGCGGCAGAGACCACTAAACGCCAAGCAAGCACCAACAAGCA

AGGCACAGAAGCAGGTACGGCCAGAAAGAGCAACCGCCTTTCCCCTATAGCGCGAGCATTCCACCCACCC

GTCCGACACATCCGTCCCGCCAACAAAGACAAGACTCAGCTAGACGAGAGCACGGCAGCACCGCCCAGGC

CAAAGACAATCGCCCTCCCGGGACAGGACAAGGGGCGCCTTTTTTCCATACAAAAGCAGGGGATAAAAAA

GCGCGCAACGACAATAAAAGCAGCCGCCGTGGGACCATGTATAACCGATGCAACCGTTCGCAACTTCGCT

GAAAAATCCAGCGAGAAACGCTACGGCTGCCGCGATTTCCACTACCCCGACACCAGCTAACGCTCCTACC

GCCACCGCCGCTACCCTTGTCCGCTCCCACAGCGCCGACGCCCCCCAGATCAGCGGTACCGCTTGAGGGA

GAGGGT

>NODE_157_length_564_cov_9.160221

ATTTTTTTTTTTTTTGAAATTTCTGTGAGCTTTGATGCCATTATCTCCTCCGAAGCGTGATGCTACGACG

GTGTGACTCTCGTGCACGGGTAAAACTCAGCACCATTTCAGCCGGGGGACAGTGTTCCCCAACACCCCCC

CAACCAGGAGAGACCATCGCCCCTTCCCTCTGACTGGGCGTCCATCTCGGTTACAACAACTCATAACGGC

GACATGGTCGGCCCACAGGGATGGAAGGGTGAGGGCCTAACAACGGCAACTGTGCGAACGGTGGGTTCAC

TCTTCTGCGGCTTCCTCGCCCTCCTCGTCCTCCCTCTCCACCTCCATGCCGACGTCCTCGACGGTGTTGT

CGCCGTGCATCTCCATGAGCTTCGTCAGGTCGAACTTGGGCTTCTTGAGCGCCTTGACCTTGCGGATGTA

GACGTTCTGCATCGGGTAGATGCTCTGCACAGCCTTGAGGATCGCCAAGGAGATGTTCTCGGGGATGAAC

TTGAGGACGAGCTCCTTGAGGTCGCACTTGGTGGCCTCCTCGGTGATGATGTCGCACATCTTCTTGCGCA

CCGC

>NODE_158_length_560_cov_10.259740

GATCTTGTACGCGTACAGCATACACAGATCTTGTACGCGTACAGCATCCCGTGACGGAAGGTTGGGATCT

AGATGGTCTACATGGTGATTTTGAACGAGGTGTGTGTGTAACTGTCAAATCGGAAACTCCGCTGCCACGG

CTACATGGCGCAACCTTGCAAACGTTTACGGCGACCGAACGGAAAGCGTTTCACCGGTAAACTACAAGTC

TACCTGTACACAACCCGGAAAAAGTACAGTGCCGTGTAAGATGAATATAGATCTTCAACGACAACCCAAA

ACAAAGCATATCCAACAACACTTCAAACAACCTTTTAGCAGAAAAGTGTAAATGCATACCTATATCAAGC

CCGCAAAGTACATGTGTATGTGCGTGTTCTGTGGGTTCGGACCTACCCCTTTATTGCGCCATCAAGGTGA

GTAACTCTACACGCATACGCCCTTCAGTAACTCGATCACAGCTCGGAACCTACACCTATCTCTCGCTCTC

AACCGCCGGAACTTACACCTATCTCGCGCGCCAGCCTAAGCGCGGCCACAACTCTACACGCATACGCCCT

>NODE_159_length_554_cov_3.202627

AGTTTTTGTACCTTGGTGGTCTTGTCGTCGCAAGCGCGGACATTATGCCAGAGATCAAACGACGCATCCG

ACTTGCATGGGCATGCTACAATCGGTTCAAGCGGGAGCTGTACGATATGGAGGATGCCCCGTTCACTCTA

AAGGTGCGCATGCTAAAGGCCGAGGTGATGGAGACACTGCTGTACGGGTGTGTGACGTGGACCCTTGGCC

AGGAGCACTTCGCTGAGCTCCGAACGGCACACCACAATTTCCTCCTACGGATCATTGGTTTCCAGCGCCG

ACAACGCACCGACCATCTCATGTCGTACGCCAAGGCTCTCAAGAAGGAACAATGCGAGAGCGTTGAGACG

ACCATCCGCAAGCGACGCCTCCTATTTGCAGGGGGCTTACAGCGCACGACCAATGAGCGGCTGACCCGGC

GGGTGATGTTTGGGACGATCGTCGGTGGGACGAACCCGGGACCTGGCCGTCCAGAAAAGAACTGGGCACA

ATGTCTAGCAGATGACCTCAGGGTTTTTCGAGCGACAGAGGGCTCCACAGAAAGCTCCGCGTCT

>NODE_160_length_553_cov_6.642857

CCAGCGAGTGGATGATGTCTCCGGCGTCGAGTGAGTAGAGGCGCTTGCCCTCCTTAAGGTCCCACAGCAT

GGCGGTTCCGTCCTTGCCTCCAGAGGCGCACAGAGACCCGTCAGGCGACACCGTGACCGTGTTCAGGTAC

CCGGTGTGGCCGATGAGGTTCGTGTGCAGCTTGCAGTTGGACAGAGCCCACACCTTGACCAGCTTGTCCC

ATCCCGCAGAGACGATCACGGGGGCAGACGGCGAAGGGGAGAATCGCACGCACGACACCCACTCCGTGTG

GCCGTCCTCCTGGATGGTGTACTTGCACTCTCCAAGGGTGTTCCACAACTTGATCGTCTTGTCACGGGAT

CCCGACACGATCTGCCTGTTGTCAGCCGAGAAGGCAACCGAGAGGACGTCCTTGGAGTGTCCAACGAACC

GGCGGGTGGTGCGGCCGGACTGGATCTCCCACAATCGGAGGGTTCCATCCCAAGACCCGGACAGAGCGAA

CTGTCCGTCGGAGGAGATCACGACCTCAGACACGAAGTGGGAGTGGCCCGTGAGCGAGCGGTG

>NODE_161_length_548_cov_9.034156

CTTCGCCCCCGACAAGTCCTTCAAGAACAAGATCAAGCTCACCTACGACCAGCGCAAGCAGAGGGTGGCC

GAGACCAAGGCGAGGCTCAGGGCCGAAAAGCTGGCTGCCGAAGAGTAGACGATGGCGTAGCTTTCATCCT

GAGTAGACAAACGCCGCCGCAGCAGCAGCAGCAGTACCTGGATACACTGGAGCAGTTTGTCGTGTGCACT

TTTAGCGGTTTTTGCGGTCTGGGGAGGGTTTGCTTTTTTTTTCACGTCGGCTGTGATACCTCTTTTGTGG

CCGGAGGTGATAGAAAAATGTCTGCCTTGCCTTTTTCGTATACTGCTTGTTTTAAGTGTAGGTTGGATGT

TCTGGTATTTTGGACGCTGGTGGCGGTGTGTGTGGGTCTTCTCGTAAGCTCTGTGTGGTCCTGTGGTGGG

ACTACTCAGATGCGACAACTTTGAATCCGAAGTGGTGGTTGTGAGGTGGGAGTTTGGGACGTTTTTCTCG

TTTTTTCTTCTTGCGACGCTAAACAAAGATCTGAGCTAAAAATCAAAAAAAAAAAAAA

>NODE_162_length_545_cov_7.061069

AGGGTGACAGGCGAATATCAGGTTGGTCAAGTAATTCAAAGTATGCTTTTTTAGGTGCTTTACGTTCTGC

TGCTCAGATGGTTTCTTATGAAATTTCAATGGGTATTAGTCTTATTAATGTTTGTTTGTGTGTAGGTACC

TTGAATTTAACGGAAATAGTAATTGCTCAATTAGATATCTGGCTTGTTTTTCCTTTATTACCAGTTAGCA

TAATGTTTTTTATAGGGTGTTTGGCTGAAACTAATCGTCATCCTTTTGATTTACCAGAAGCAGAAGCAGA

GTTGGTTTCGGGTTATAACGTAGAATATTCAGCTATGGGATTTGCTTTGTTTTTTTTAGGGGAATATGCT

AATATGTTGGTGATGGGGGGGTTAACTTCTATTTGCTTTTTTGGTGGGTGGTTATCTCCCTTTAATATAG

GTATTTTACCCGATGGTATCTGGTTTGGTTTAAAAATATGTTTTTTTGTTATTTTATTTATTGTGATGCG

GGCTATTTTACCTCGTTATCGATATGATCAACTAATGAAACTGGGTTGGAAATGT

>NODE_163_length_545_cov_3.927481

TGGAAAAATCATTTCTACTATTATACGATAGGCCGCGGTGAAAGTTAAGCGTCCGTTTTGTTTCAAAATA

CGTTTGTTTAGGGGTAGATGTAGAGGTCTTACGCTATAATCAAGTGGGGAGCCTAATAATAAACAGGTGG

TACCGCTGTGAGGCAAAAACGTCTGGCTCGGCTTGGTTTGAGTTGAGGTGGTTGGGTTGGGTTGGTTTGA

GTTGATTTGGGTTGGGTTGGGTTGGGCGGAATTGGGTTCGGTCGGGCTTTTGCCGCAATAGAGCGTTGAG

CTAGCACAGGGATGCGATCGTTGATCAGCGCTCAAAGGTTTTTCCAGAACGGCAAGGTTTGCCAATACCG

AACAAGGCGTTTTCGGAAATATCTGGTCAAGAGCCTGCCGAAAATACAATATTTGACTTCAGCACCAATA

CCGCTGTGGAGAAACGAGCGAGCGCTGAAAAATACAATATTTGCCTTGAGAAATTTTACCATTTGACAGC

TTGGGGAGGGGGGCGTGTACTCGATATCATGCATGGCCGTAGTCGTATGACCATA

>NODE_164_length_543_cov_7.808429

CCTCCCGCACCCCACTCCGCCTTCAAATTAATACCCTGGTAATAATCGGTGTTTTACTTCGACATCATCA

TCTTAACGAACTCCTCGTAGTTGATCTGGCCATCGCCGTCGATATCCGCCTCGCGGATCATCTCATCCAC

CTCCTCGTCCGTGAGCTTCTCTCCCAGGTTAGTCATGATGTGGCGTAGCTCGGCTGCCGAGATGAACCCG

TTCCCATCTTTGTCGAACACCTTGAACGCCTCAAGGATTTCCTCCTCGGAGTCGGTGTCCTTCATCTTCC

TCGCCATCATCGTCAAGAACTCGGGGAAGTCAATGGTTCCGTTTCCGTCCGCGTCGACCTCATTGATCAT

GTCGGCGAGCTCGGCCTCGGTCGGGTTCTGGCCGAGCGAGCGCATGACCGTGCCGAGCTCCTTGGTGGTG

ATCGTGCCATCCCCGTCCTTGTCGAACAAGCTGAAAGCTTCCTTGAACTCGGCGATTTGCTCCTCGGTAA

GTTGGTCCGCCATAGTACTAGGGGGTGGTGGGGCTGGTGTGAGAGGAAAGGGG

>NODE_165_length_541_cov_7.194231

CTTTGCGTTGCAAGCGGCATCGCTGCCCCCGCCATGACCGCATCTCTCTCCTACTTCGACTCCTACCGCC

GCGCCAACTTGCCAGCCAACCTTGTTCAGGCTCAGCGCGACTTCTTCGGAGCGCACACGTACGAGCGCGT

GGACAAGGAGGGAACCTTCCACTGCCTCTGGGACGACAGCCACAAGGACATCGGAGACGTCACCGGCCGC

ATGGCGGGCGAGCTGTAATATGGCAGGGTAGCAATCTTGGCCCGTTCTTCGTTTGGCGGATGGGGGTGTC

TGAATTGGAGCAGCACAGCCTTGTCTGTAGCGTACTGCGGTATGCTAAGTATTACATAGTGGGCCTGAGT

AATGCATGGGCGTTAGTGTGTATTTTATGTTATACAATGCGCGTGTTTTTTGTAGGGTGCTTGGGCCGTC

GATGGTGTAAGGTTGCCCGCGATAGTTGTGGAACAACGCTGATGTTGTGGGTTACAACAGATGGCTTCGA

GTACACTCGACAGTGCGAAGCCTGCTTGATCGAGTGTGCTGAGAGTGGTGC

>NODE_166_length_537_cov_12.127907

CCCGATGGACTCCGGCAACGCGCAGGAGGGCCCCGGAGAGGGCTTCATCCACGTTCAGCGCGGTATCCAC

GGAGAGGCGGACATCAGCACCCGCGTGTACGACTGGAGGAACCCCGTCGCCGAGGTCTTTGTCTTCGGCC

CGTAAAGCTATTAATAGCCTATTCGGAAACCTATTAATAGCCTATTCGGAGACCTATTAATAGCCTTGAC

GCCTAATCATAGACTTGATGAAGAGGCTTTTGTGAGACTTGGGCCGTCAGCGCGTGGGACTAGAGGAAGG

GACCCCTCAATAGACTCGGCCGTTTTTGGCGACACGTTACTAACTACTGTGGCAAGTCAGTACGAGCAGA

GCTGGAAGGCGTTTTCGAGCGTAGGCCTGGAGGAAGGAGCCCCGTCGCCAAACAAGCCTTCCGTCTTCGT

CTGGACAACCTAACAGTACTCTTGTCAATGGACTGGATCATTACTGACCATTGCTGTACGTTACTCTACG

TTGGCAAGTCAATAAGAGAGCAGGAAGGCGGCGGAGAGCGCGGGGTG

>NODE_167_length_537_cov_6.771318

TCCGAACGGCTAACTTCCCCTTCGTGGCAAGGCCTCCCTTAATTCCTACCACGAATGTCCGTTTCCTCCT

CCACGGAATCCCTACTTGGCTCGCCGATATCTCTACCTTGTCCACGATGTATGACTCTCGTTCCAAAGCA

GGGAGAATCTCGTCGTTCCATTCCGTCGTCCGCAGAAGGTTGCCCAAACATTCGAGTATAAAAACTCGCG

CGTCTGAGGCGATGGTAGCTTGGGTTAAGCTACTCGCTCCGGATACTCCTTCCTTCTCTCGTACCCCTTC

AAACCTTCTGCAGGCGTCACCGACCACCACGTCTGGTTTCACCTCTCTCAACATGCTTAGGCATTCCTGT

GACCCTGTTGTCACCCTGAGTAGCCTCGCACCGTTGTCCAGTCCGTTTGCTCTGATGCATTCCAGTGCTG

TCCAGTCCTGGTCTACCGCTCCTCTGACCGTCAGCAATCCCTCCGATGCAAGGATGCTGCCTCCCCCCCC

ACAGTCGAGCATGACCACCACTGGAAGCTCTGATGCATTCCAGTGCT

>NODE_168_length_536_cov_4.392233

GACTTCCTTGGCCCCCGTTAATTCCAAATCTTTACCCCCCCCCCCCGAACGTAAGATGATTCAGGAGCTC

CTCCGTGGCGCCTGGACGACGAATAACCCCCCAGACGTACTTCGACAGCATCACTTCCCGCAGCACTACA

TCCGGTCCAGGAAAGGACGGTCAACGCTCGGCAGACCTCCAATATATCATCCACACCATAGATACCGACT

AAGTGCAAACAATTTTTGTCTGGTAACAGCGCTGCCCAACTCAGAGTTCACCCCAATGGAGTATCGAGTT

AACTGTGCCAAAGGTCCCTACAGTAAATCCCAAACCTCGTGAACCACTGGGGTGTTTTTGCTTAGATCTA

CTCCCCGTGGAACAGCCACCACATCTGAGCAGATGTCTACTGCTATTTCGACTGCGGTCCAGATTGTAGA

AGCATAATCCAATATTTCTCAACACAAAGAGCATTTCTGAACTGAGCCGATTACCACGTTAGTGTTTTAT

CCGTGCTACACGACATCCAATATGTTGGTTACCGTCCCACGAAAAC

>NODE_169_length_534_cov_182.836257

GGCCCCGAGAATTCCAGGGTCTCAGAGTTTTGTGAGCTGCTCTTCTCTAGGAAAAGCTCTGGGCATCGGC

GCTCCATTCACAACGAGGGTGTTGCAGGAGGCATTATTTCCTGGGGTTATTTATCATCATTGACCCCCAA

GCCGCCCGCTTCCCTAGACTCAGTTAAGACCCTTTTGACAGTGATCTTTCCCTGGGTAGTTAGGCAGACG

TCACCGGGGGCAGCCGAAAATTCACTTCTACAGCTTCCGTCTCAACAGTTTCAGGTCTTTTGACTCTCTT

TTCAAAGTTCTTTTCATCTTTCCTTCACAGTACTTGTTCGCTATCGGTTTCCCATCAATATTTAGCCTTG

GGGGAAGTTTATCCCCCGATTTATGCAGCAATCCCAAGCTACATGACTCTGCGAGCACTTCTTCCGACTT

CTCAAACGCCAAATACGGGGGTTTCACCCTCTTTGCCGGACTATTCCAAAAATCCTTATGCGTTCGATCC

ATCTGCAGAATGTACTCCCCAGCTTACAACTCGCCACGCGAAGC

>NODE_170_length_533_cov_3.578125

TGATGGTCGCCTGAACCACTTGAGCTTTTATTGAACGTTAGCAGCGCAAGCCGCGACGCAACGGCTACAC

GTTGACAGCCAAAGCTCGGGAAAAGTGCACGGCGAGAAAAAAAACAGCCAGTGCCCTTTTAAAACTCAGC

ACCTTCCCCGCCCCCCGCTATCGCATGACCAAGAAAACACCAGCAAACCCCGCTACACTCCCACGGCAAA

AAAATGCCGCCTTGTGCAAGCAACGCCCCACCGATGCCCCCCTCATCCCGTGCAGTACATGCAGTGGCGT

TTCGGCACAGGGGACGGAACTGACAGGGAGGTTGCGGAGAAGAACATCCTGATGAAGATGACGGACGACG

AACAGGAACGCCAGAACACGGTGGGACAGGTCGAGGACATCTTGGGCCGTCGCCTCGAGGGCAGGGTGCT

CAAGTACGAGTGCTCCTTCGTCGGCATGTCTGCCAAGCACAACCGGTACATTGCCCGAGAGGAGCTCGAG

AAGCGCGGGCTGTCTAAGCTGGTGCAGCAGGCGGACGCTGGAA

>NODE_171_length_531_cov_23.572549

CTCCCCCTTGCCCCTCCTCCTCCCCGCCCAGTCATACCATCCGACATGATACATATGAAAACAAACATAA

CACGTGGGAGTTCGCTCTGTGAACGAAACAAATCCGGTAATTTTACGACGGACTTCCTGCCACGAACGAG

TTCATCGCGCATCAGCCAAGCGATCGGACAGGATGCTTGTCGCCTCGTTCTTAATGAGAGTTGAGTCCTT

GATTGGATTGGACCTCAGCCTCCTAGTCGGTTCGGCCTCGGACCCGTTCGCAGCAGGGAAGAGGTGGGCT

TGCGTCGGGTTTCCGGGGAGATAGGCGTTTACTTGTTGCCGCAGCCGCAGCATCCAGGCGCGGTGCAGCT

GCACGTGCCGTTGCCGCCGCACTTGCAGTTGTCCCCGCACGGGCAGTTGGAGGCGGAGTGAGAAGTCGAG

CCCGCCTTCATGGGTCCGCAGGATACGCCGCTTCCGGACATAGTAGAGAAGGTGGTGGAGTTGGTGTTAC

TTGGACAGGTTGAAATGGCACAGGTTGCTTTGCTTGAAGAT

>NODE_172_length_530_cov_9.734774

GACCCCGACCTTCGGGTCGTCCCAGGTGGACTTGCCGGTGCAGGTGTTGTGCCAGTAAGGTTTTCTCTTG

TTCAAATCGAAGTGCTCTTCCCATGTGTCCGCCACAAACTTCGAATCAGGGAAAAATTCGATGGACTGAT

AAACGTTGGGGTTAAGGCCGGGTGACAGTCCAGCCGAAGACAAGGGGTACTTGACCGTTCCGGAGATCTT

GTGGCGCTGGCAAATGCTCCCATACTGGTCGCGCCCTTGGATCCACATTGACATGTCTGAGCTCGCCATC

TTGACAACACCGCAGGTCCTCTACCTCTGCGACGAAACTCCTTGCTCCCTTGTGTCCTCCTGGTCCCGAA

GCAATATAGGGGCTTTATCGAGTAGTATATAAGAGCACCAGACCTTGGACTAGTTCTTCTTGTTGTGGTC

CACCGGCGTTGTCCTTAGGGGTCCAGAGTTTTGCCTTGTTGGCGATGGCTGCCGAAGAGTCGAAGAAACA

AACAAACGGAGAGGGGTCTTGTTTGTTTGGTTTTTCGCTG

>NODE_173_length_529_cov_3.915354

GGATAAAGGGAAACGCTACCCGACTTGATTTGATGGAGAGCACCGCCGTGTAAGCGTTTTTGATTAGAGT

CGTTTTGTTGGGATTGGACCGATTTGAGCGTGTGGCCGGTGTGGATTGTCGATGGGGGCATCGTGTAAAC

CTTTGATTGAATTCGTTTGTTGGGATCGAACCGAGTCGAGCGTGTGGCTGGTGAATTTAGGGATGGAGCT

TCGTGTGAACGGTTGATTGAGTCGTTTCGTTGGCATCGAACCTATTCGAGCAAGCGACCGGTGGGTTAGT

AATGAAGCATAGTGTAAACGTTTGCTTGAGTCGTTCGTTGGGATCTAAGCGATTTGAGTGTGTTACGACG

TAGTGGTGGATGGAGTATCGTGTAAACCTTTGAGTCGTTTTGTTGGGATGGAACCGATTCGAGGGCGTAG

CCGGTGGGTTATTTTTTTTTGAAAAAAACCCGATACGTTTCGATCCGATTTCTCGGTGGTGGTGGCGCAC

GAGGACGACACGAGGAGAGCTTGGAATTCTCGGTGCCAA

>NODE_174_length_528_cov_12.508876

CCTTGGCACGCGAGTATTCCATCGTCGAACGCGAGGCCCATGATGCCGTCAAACTTCCCGAGGATGAACG

CGAGGCCGAGACCCTTGGCGTCCTTGACCTCAGCGAAGGTCTGCTCCTCTAGGTCCATGCCACCCCACGT

GACAGTGTCCACCGAAAGCTCGCCGCTGACTGGGCCGCTCGCGTACGTGATCTCGAAGTCACGCCCGTCG

TCAGCGTATGTGGAGCTCTTGGCGGCCTTGTACCGGTGGTGCAGCCCGCACGACATGCCGCAAGCATTAC

CCGCCACCCAGAGGTTAGCGCTACCGGTGTCGAAGATCACGTCGAACGACTGAGGCGGCGTGCCAATCTC

CACCTGTCCGTAGTACTGAGCGTTCTGGTAGTCCTTCACGATCACCTTCCCCTCTCCGGGTGTCTTCCCG

ATCGAAAGGAGTTCCTCTATCTTGTCTATGGCGCTCTCCGCACCACGAAGGAGGCTACCGCGGGCGGTTT

CAAACTTCAGCTTAGGAGCGTCAGGTGGGGTTTCCGCA

>NODE_175_length_526_cov_73.413861

CCAGTACACCTCTGTCTGCTGCGCGGCCAGCCGTTCGGAGAGAAGACTGTCAGGCGTGGCGATAAAGATG

TTGTCGCTGTCGGCACCGTCGACCTGGTGTGCCGTGCCCCCAAGCGGGTTGGTCAGTTTCGCGCTGCCCC

CAGCCGGCACATTCTCGCACACGTTGATGTCGCTGGTGCGCACACAGTCCAGCAGCTTCTGGTAGTCGTT

CTTGTTCACCTGCCCATATTTGTTGTGCGGGAGGGTCTTGTGGTACTGTCCGGCGAGCATCAGCGTCTCG

TTGTCCTCGTCTCCATTGGTGAGGTGGCACGTCGGCCCGACCTTGAACTCGTCCTCGGCGATGGCCAGTC

GGTTCCTTCGGGTCACCTCGGCGCGCGCCACCGGGTCGAGGAAGTCCAGGGAGTCCCGGACGTTGCACAC

ATCCCCGCTGGCCGGAGACTTGTTCGATCGGTTTCCGTTAGAGTCCCCAGCGCCAAGGCAGGGAGGAAAC

GCAGCCAGAAGCGCTGTCAAGCCCACGAAAGTCTTC

>NODE_176_length_526_cov_8.970297

CTTGGCACCGAGAATTCCATATATATACGAAGTATATATATATAAGTGCCCCCGTTGCAGAGCTCTATCG

ATTGCGCCGTGCGTTTTGTTTGCATATATGCATGCATACGCATGCCAAGTTTGCTTGCCCTAATCGAGTT

GAGTACACACGAGGTACTTGTATTTGCAGGGATATGTAAGCGAGGTGAACGATTTGTTGGTGTGGAGTGC

GAGTCGCGATCCCGTTGGGTAATCGGGTACTGTCGCGTAAGTCGTAGGTTTATAGTGCGCATGTAGAGCT

CTACTTGCGTGGATGCTCTGGGGCTCTAAAAAGTTGGCTGAGCCCTACATATGCAGTAGCAACACATAGC

ACCCGGTACTATAGTGATGGGGCGGGGGTTTTGTTAGCACAGAGCCCTTTTATCTAAAGTACTCAAGTAA

AGAGAGGCAGAAGAAATCCCAGGTACTCGGGTGGGACTACCGGGCGGCGCAGGAGGCCTCACTCAATCCA

ATATACTTACATTCAGTGAAAAAAAAAAAAAATTTT

>NODE_177_length_526_cov_5.180198

CTTGGCACCCAGAATTCCAGTGTAGAGTACCTTCGTCCTCCCATACAGAGCTTCCTAACTCCATAATCCC

CTTCGGAATCTTCGGTTCCTCCAACTGTTCCGGTTCTTGCTCGTCTGTCGATGGTTTCACCGCGCAATAG

TGGTCCCACTCGGCATACATCGTTCCTGCTGCGGCTTCTCCGACCTCCTTCGGGGCGAATGGTACCCACG

ATGAGGATGATGTCGGCCGGAAACCCTCACCATCGAAACTGATGATACTTCGAGGATCCGTCATGAACGC

GGCTCCAAGCACCATCGCGTACGAGAGTCCTTTCACCACCCGGAAATTCACACGGAAGATCTGATCGACC

GGTGCAAAGACCAGCGGTAGCAAGCACGAGCCTACCACTTCCATCGGGGTTACGGCGCTGTTCTTTGGGT

TTGCTACGCTAAGTAGGCCCTTTCCCATTGGACTCACCAGCGATTTTCTCCAACCATCACGTTCCTCAAT

GGACTCGATGAACTGCCGTGACAGGTAGTTTCCTCC

>NODE_178_length_523_cov_8.884462

CCCGGTTGTTCGCAAATCCCATCGGTCAAGATGGTGAACGTCCCTAAAGAGAGGAGGACCTACTGCAAGG

GCAAGAAGTGCACCAAGCACACGGTGCACAAGGTCACCCAGTACAAGGCTGGCAAGGCCTCCAACTTCGC

TCAGGGCAAGCGTCGGTATGACCACAAGCAGAGGGGTTTCGGTGGCCAGACCAAGCCTATCTTCCACAAG

AAGGCCAAGACGACGAAGAAGGTCACCCTTCGCCTCGAGTGCAAGGAGTGCAAGGCGAAGAAGCAGCTCG

TCATCGCTCGCTGCAAGCACTTCGAGCTCGGCGAGAAGAAGCAGGCGACCGGTCACCAGTACTGAGCAAG

ACAACCCGGGAGCGGACGGGGGCTGGGGAGACTTTTTTTGTCGAGAGTGTTCGGACAGCGATGCTGGCGT

ATTGTAGTAGCGGCGTGGCCTGTGGCCGAGCTTAAACTACGGGCTTTCGTGTCGTGGCGACCTTGAATCT

AAAGCGGTTTCTCTTGAAAAAAAAAAAAAATTT

>NODE_179_length_522_cov_16.481038

CGAGGGGGGGGGTCTTTCTAGCGAGCAAGGCTAGACTCACCCTCAAGCGGACCGATACCTTCTTGCTGCT

GCGCATCCGCCGCGAAGCGTGTGCAGCGGTGTGGCTGGGTCGGCACTGGTGCGTCAGGCGCAAAAGTTCA

GATGCACGGGCCTTTTTGGCCGTCAGTGTAGACAGCAATGTCCATCTCACTTTTGTGCGCGGAGGCAGAC

GCCATCAGAAAGGTGGACGGACACTACGTACACGGCCCTAGGTGTGTTTGCTAGAGGGAGCTGTACTGTC

TCAGTAACCTGGCTTCGCCATCAAACGAGGAGAGGACGTGTGGCGACGAGCTGAAGTTGGGCTGGTTGAC

CCCTTCGGCGCGAGAATTTTTCTTTTCCAGACCAAGCACCATGAGTGCAACGAGGGAGAAAGCGAGGAAC

CGACTGGGAAGGACAGCAACGAGAGCGTGTACGGACGGAGCGGGTCGAGTGTGGTCGTGCTAGGACTGGC

CAGAGCAGCACCGTGGAAATCTCGGGAGCCAA

>NODE_180_length_520_cov_7.300601

TGCTTTATCAGGTAAAAATCTATCAGCTATGTATTGAGCACCTAAATTAGCCGCAGCAGTTAAAGCTTCA

TTTGTGATTTCTAATCTATGGTGCTGCTCATAACGTAGTCTTAAACCTTTTAGAATAGTTATAGTTTCTT

CTATACTAGGTTCATTTACCCAAACCGGTTGGAAACGACGCTCTAAAGCTGGGTCCTTCTCAATATGTTG

TCGATATTCATCAATTGTTGTAGCTCCTATACATTGTAATTCTCCACGTGCCAGGGCAGGTTTTAAAATA

TTCGCAGCATCTAATGCTCCTTCTGCTGCACCCGCACCAACTAATGTGTGGACTTCGTCGATCACAATAA

TTATATTTTTTTGTTCTTTTAGTTCTTGTACGATTCTTTTTAAACGTTCTTCAAATTCTCCACGATATTT

TGTTCCTGCTAATAATAATGTAATGTCCAAAACAAATACTTTATGGTCATGCATTTCACTAGGAACTTCA

CGTTGAATAATTCTTTGTGCTAAGCCTTCG

>NODE_181_length_519_cov_6.465863

TCCAAGAAGCGCCGCAAAGTTCATTTTGCCACCAGCACATCATACTTCAGAATAGTTAAAATACCTGGTA

TGATTTACAGTCGCAATGGAACTCTACGCATATATACGCACACCATCATTTATTCTTCCTTTTTTCACGA

CATCTACCACTCACTACCATGTAGCCCGGACGGAGTACGTCCACATGCAGATTGTTCATACATTTTTTTT

TTAAGGACCGGTCTGCTTGCTTTTTGCTCGCTCGCGGCCTACTGCTACTGCTACTGTACCAAAACCACCA

CCACCGTAACTTCCGACGAGCGGCAACATTACACCGTCGACAACCTAATTCCAACAAAAAAGACGATAAA

TGAAATGACCTACCCGGTCTACAGCCACACTACTTTTTAGCGTGATCAATCGCCATCCAACCCGCGCCCT

CAAACAAATCGCTGTCCTCGTCCGCAACAACGCGTACTGCCTGTAGAAGGCGGAAACAAAAAGCGGGAAC

ATGCAAGTTTTCCAGCCCCGTCGCCCCAG

>NODE_182_length_516_cov_3.076768

CGCGCAGCTGTAGCCGTCGGTTCCAGGCTGGCATTCACAGGCTCTTCCGTTCTCGTCGCATGTGGGAAAC

TCCGGGCATTCACACGTTGCGTCTGCGTCCAACTGCACATCCACAGTCACGGAGGCGGGTATCGCAGGCA

GGCACTGTAGATCCTCGTTACCGACCAAAAACAGTTCTCGTAGGGACGTCAGACCTTCGAAGCTACCAGC

TGCCAGGGTCTCCAAATCGTTATCGGACAACCTCAGTATTTCCACAGACGAAGCACCCTCGAAGCTTCCG

CGCTGAAGGTTCAGTTTGTTGCAGTCGAGATCGAGTTCGCGCAGAGACGTCAAACCCTCGAAGGAACCAA

TTGGCAGGGACGACATGGCGTTGTCGGCCAGGTTTAGTTCTTCCAAAGCTGCAGCACCCCTGAAGTTCTC

AGACTGCACGTTCAAGAAGTTGCCCTCGAAACTCAAGGTCAGAAGGGATCGAAACCCATCCAAGAAATCG

AGCGGCACGCTGGTGAGCTCGTTGAT

>NODE_183_length_515_cov_4.991903

AGCTTGGGTAAAAAGAGAACTTTCATAAGGTGTTAAATAAGGTAAAAAAAGAACCATATTACCTTTTCTG

TAGAGGTTACTAACTGAATCAAAAATTACAACTCCTTTAGACCCTGAGACACTCACTTTACCATTAATAC

TTGAACACCCAACACCTATAGGTAATCGAAATACTGGAATTGTCATGACGCGTATGCTAAAATTTCACCA

CCCTCCCCTTTACGTATACATTGTTCATGAGTCATAATCCCTTTTGTTGTAGATAAAATCAAAACACCGA

AATCATTTGATAAACGAGCAAGATCTACTAAAGAAAGATAAATACGATCACGTGGGCGAGAAATAATAAC

AAGACGAGTACAAATTGGAGACCCATCATGATACCTAAGAAAAACTGTGATGATACCCTCATTTGTTTTT

GTATGTCAACGTTTGTTTCGGAATTGGGGCGAGTGCAAGTATATTTGAAAACTATATGTAGGTTGTATGG

GGAGGACGCATTTAGAGTGGAAGTT

>NODE_184_length_514_cov_10.874239

GGTGGAGAATATTGGAATTAGCAGGCAAGCATCCCAGGCCCCGAACAGACCATATCTACTGCAGCTTCCA

AGACCCGAGGAATCAAACGCCAGCACCGCCCAAAAGTGATTCGAACGATGGAGACATCAATTCCTCTGCG

GCGAAAGGCCGTGTCTTAAGCTACTGCAGCCGAAAAAACACCTCCCCGCTCCTTCCATCCAGCACCAAAA

ACAACCCCCGCGACAGTCGGAGCTCGCAAGAGCCCTCAACAATCATGGAGACCGCCCCCCTTGGTTAAAG

CCACCTCCAACGAAAACCCTACAACCAAAAGCTGAAATAGCTGCAAGGTTCTCGTGTGCCTGTAGGCGAC

AAGAGCTCTAAGCTTCGTCTTTGACTCGCTTGGTCCCGTTGTACTTCTTCGCAAATTTCTGGTTGCGAAG

GAACTTAGGGTCCATCCCCTTCATACTCTTGTACTTGTGGTTAGGCGCCTTCTTGATGCCGTTGACGTGG

TCCTTGAAGGTGGCGTTACGCCCG

>NODE_185_length_513_cov_4.591463

AGCCCCACGGGGGAATAGCGCCAATCGCGTCCACTTGTATAGACGCCCAGGTTTGATCGAAACCGTTTGT

GTGCAGGTCTAAGCAAACAGAAATGTAGCCACGAGTTTTCACGTCCGACACACATATACCAAAACTGGAG

ATAAAACATATTACGCGCTGCTGGCCTACACGAACCCTGCGGAATAGATAATACCAAAGGTAGCCTGTCC

TCACACCGCTACAGCAGTACGAACGGCACAACAACAGTCTCACACTCCGTCCCCGGCCTTGGCTCACTAC

ACGCAACTATCTTAAGCACATATCGCAATGCTGACACTACGCAAGCAATCAAACACACGGACGAGCCCAA

CCCTTGTGTATACCGAACGAAGCAGGCATCTACGGACTCCCTCTGCCTTTGAGAGCGAGCGCAGGCTGTC

CTTGTTCAGCATGCCCTTGCGTGTTTCAGTATCCACCAACGGACGGCACCCACGATTCCGCCTCAAGCGT

CCGCACCTCCCCGTACCCAGGTT

>NODE_186_length_510_cov_8.603272

TTTCACCCACCCCTGCTGCACACAACTCAAACTCGAGCCTCTACACTCTAACTCTACACCACCAACAAGT

TTCCAAGTCCCCGAAGAAGCAACTCGCAACTCACACAGTCAAAATAGGGAACAAGAGGAGCAAGATGGGC

AGCGAAGTCGATTACGTCAATGAGATGGTGGCTTCCAAGCCTCTGGTGGTCTTCTCCAAGTCGTACTGCC

CGTACTGCACGAAGGCCAAGACGGCCCTCAAGGCCGTGGGCGCCAAGTACGAGCTGATCGAGTTGGACCA

GGTGAAGAACGGCAGCGCGATCCAGGACGCCCTCCAGCAGATCACGGGTCAACGATCGGTCCCGAACGTC

TTCATCGGTGGATCGTCCATCGGCGGCGGAGACGACACGGCCAGGCTGCACAAGTCGGGAGCATTGTTGA

CCAAGGTCACGGAGGCTGGCGCGCTTTGAGGACCTGCCCTGCCATATTATATTCCAGCTGGGGGCTTTGG

GGGGGGGGGGGCGTTTTTTT

>NODE_187_length_509_cov_5.065574

GTGGTCTGTGCGCGCAGACCCAGGCTGAGTCGCTTAAGTACAAGCTTCTGGGTGGACTCGCCGTGCGTCG

TGCGTGCTACGGTGTGCTCCGCTTCATCATGGAGTCTGGTGCTAAGGGCGCCGAGGTGATCGTGGCTGGT

AAGCTCCGCGCGCAGCGTGCGAAGTCCATGAAGTTCAGGGATGGCTACATGATCAAGAGCGGCCGCGTGA

GGGAGGACTACGTGGACCACGCTGTGCGCCACGTCATGATGCGCCAGGGCGTGCTCGGTGTGACCGTGTC

CATCATGCTTCCCCACGACCCGACGGGCAAGGAGGGGCCCAAGACTCCGCTTTCGGACGTTATCACCATC

CTCCCGCCCAAGGAGGAGATCGCTATGGAGGTCACCCCCAACTTCGGTTAATTTGCGTTATGCCGTGCCA

CTTTGGGGAGCGGCAACCATGACCGGATAGGGGGAGGGTGTGGGGGGGCACTGCTATGCCGCCAGTGACG

GTTAATGGAAGTGTTTTCT

>NODE_188_length_508_cov_12.410678

CTTGCTTGTTTCGTATGGCTTTGAAGATATGTGTTTGATCAGTGCGTGCAGGCATACGTTTTGCTCGTGC

CCTGTCTGTCCGTCGCACTGTCCGTTTGCTCGTTGTTCAATATCGCACACATGCATGTACTCTCGTGTGT

TAATAAATGACTGCAAGCATTTTGATCAAGCATACTGTCTTTAGAACCAAACGCAAACCACAGGTATCGA

TACAAGCCGAAGGTGATGTAACTACCACTCGCATAATGTGTGCTGATGGCCGGGCCGACGACATAACGAT

GTAGATATAGAGTGGACGCATGTCGTTATTGGTATGGTAAGTAAGCTCAGAAGGTCAGGTAAAAAGTAGA

TAACTCACGTCACGTAGTATATTCTATGCGTCTTGTCTTTGGACGGTATTAAGCCAGCCGTCATATGTCG

CGCGTCTCTAATTGGTATCGATAGCGACAGAACGAAGACGAGTGCACCCGGCAGAAAAACATAGCGATGC

ACGTTTGATCCTCTCGGT

>NODE_189_length_507_cov_3.985597

GTGTTACGCTTTTGTTTGACTTTGTCTCTTCTCTCTCTGTGTTTGTGTGTGTTTTTTTTGCTGTTGTTTT

CAGTTTTTTTTTTTCTTCTATTGCGGTCGTCTTATCCCTGCGTGCTATCGTTCTGCGTGCTTCCGTCCTC

TTCTTTTGTTGTGGTTTTTGTTTGAGTGCTAATCTTGTCTTCCCATGATACGATCGATCTCTCTGCTGTA

GTCTTCTTTTTTCTTTCTTTCTTCTATAAATCTATCTGTCTGTCTCTATCTACTAGTCTATATACAGTAT

CGCGGTTTGTTTGTTTGTAGTTGTGTGTGCGTGTGCGTGTCTTGTCTGTGTCGACCGGAATGAGCCGGCA

ACTCAGTAGTGTTATCACTCTTTGTTGAGAAAGCACTAGCTAGTCCTCGAAGGACGGACGGTGCTTTGCT

GCTGTTGTAGTAAACAAGGTGGTCGCTCGTCGGATTGCCGGTGACAAACCGACTTATCAGTCAGTCAGTG

TGCGGGGATGGAAGGCT

>NODE_190_length_505_cov_8.772727

CTCCCCGGGCGCCAGAAGATCCTCCAGTCCAACAAGTGGGGCTTCACCAAGTACCCCAGGGAGGAGTACG

TCCAGGGACGCAAGGACGGCTGGCTGCAGAACGACGGAAACTCCGTCAAGTACATCGCGCAGCACGGCCC

CCTCAGCAAGGCCATCCTCTGAGCTGTTTTTTAGACGGATGACTGCCCCTTCGCTGGGACGGATCTCAGT

GTGTTAGCATAAGCACCCCTCTGCCTGCCGAATGGAGTTACGAGTCGGGTTTCTCGTTTTTCAACGATTT

TCTGAGAAGAGACAGCGGTAGTTCTATTGGGTGGGGCTTTTTACCCTGGGTGTTGGGCTATTGCTGGAGA

GCTTGACTTGCCGCTCGGCGATACGAGGGGAAGGGATGTTAATGCACTGTTTCTGTTCTACGAAATAAGC

GAGTGGGCTATCGTAGAGGAGGATCATGGAGAGGGTCTTCAGCCGAATACAAACCTGCCATTAATGCCCA

AGCACACCTTTTTGT

>NODE_191_length_505_cov_5.663223

TTTTGTGTTGTGTTGTGTTGTTTTGTTTTGTTTTATTGTTGTTCATTGTCATAAGAGTTTTGTTTTCCTT

TTTTTTGTCGGACGTATCGCGGGCGTCGTGGTTTTGTTCGTTTTTTTTCACGCTCATGAGGGAGCTTTTG

ACGAAGGAACCCGGCGGCAGGTGGTACAACGGGTAATAAGGTGCAGAGTGGTATGATACGTGTGTACGCA

GCGATGTGACTGCTACATTGCCCGCCGCGTGGTATGGTGTACCAGGATCCGTCGCACTTGGGGTATGGTC

GGGCACCTGTTTTATTTTATTGTGCCGCCGATAGACGTCAGTAAGGCTGAGCCCGCCACGAGCCATATTA

TTTCGCAAGGGGAGTTTTTTTTCCCGCCTGCCTCCGCTTGGAGTTCGTCCGCCTGTTTTCGTGCCTTGCA

CTGCTTGCTGTTTGTCGCTTCTGGCTGTTATCGTAACGCTTCAAAACGAAGCTAACCACGCAAAAATAAA

CAAAGGTTATGTATA

>NODE_192_length_504_cov_12.801242

CCGGAGCAGAAACACCGCGGAGCCTACGGACGACATCAACCCGAGCCGCACGGACAAACATTATTTTTGC

TGACAACTATTTCGTTTTTTTCCGCTGAATGTTAGTGTTAGGTCTGTGTGGATGACGTCACTGGTTTTGT

AGAGAGACGCGGCAGAGATGGCGCGGACACCCCGGGTGCCTTAATTGTGTGCCTGATCGTGGAAGACCCC

TTCAATGTGGAACGATTCCTAATCAGTGTCGATGTTTATAATGCTGTGCCTAACACAGACGTTCAGGATA

GATTCGGGTAATTGTCAATTGGCGTGATGGCTTTGGTTGGCTGTTCTTTTTCACTGTGTGCCCGCGGACT

GTCATTCTGGCAGCACTTCAGACAGCAGCTGCGTGCTCCATCTTGTTCCCACGCCTACCGTGTGTTTTTG

TACAGACAGCACTTCAGACAGTAGTTACGCTGTAGTCGGCAAATTCATTGGTAAAGAGTGGGAGACGAGA

TCGAGCGTGTGGCC

>NODE_193_length_498_cov_9.628931

CCAACCATCCCAAGAGAGACGAAGCAAACCGATATTTGCTGTGGCCAGTTTTCGAGAAAATAGGCGTTGC

CGAAGAGGGAAGAAGAACCAGCAGCAGCGGCGGCAGCTAATCTACCGGTTAGCCTTGGCCACACGCTCGA

TCTCGTCCTTCTTCTTGATGGCGTAAGAGTTGGACGATCCGCGGGCGGCGTTGATGATCTCATCGGCGAG

GCACTCGGCGATGGTCTTGATGTTGCGGAACGAAGACTCGCGCGCTCCGGTGGTGATCAGGTAGAGAGCC

TGGTTGACACGGCGGAAGGGCGAAAGGTCGACGGCCTGGCGCCGCACCACACCAGCGGACCCCACGCGGG

TGGAGTCTTCCCGGGGTCCGGAGTTGATCACCGCGTCCACAAGCACCTGTACGGGGTTCTGGTCCGTGAG

GAGGTGGATGATCTCGAGGGTGTGCTTGACGATGCGAGCCGCCATAAGCTTCTTACCGCTGTTGCGTCCC

TTACGCAT

>NODE_194_length_497_cov_12.222689

TACATTTTTTATGTTAATATTGGTTACTGCCGATAATTTTGTTCAAATGTTTGTTGGTTGGGAGGGGGTT

GGTTTATGTTCTTATTTATTAATTAATTTTTGGTTTACTCGTATTCAAGCAAATAAAGCTGCGATTAAAG

CTATGCTGGTTAATAGAGTCGGGGATTTTGGATTAGCCCTAGGAATTTTTGGTATTTTTATTTGTTTTGG

TGCTGTCGATTATGCCACTGTTTTTGCTTTAGCCCCCCAACTATCCACATTTAGTTTAACGTTTTTTAAT

GTTGAATTCGGAGCCCTTAACTTTATAGGAATTTTGTTGTTCGTAGGTGCGGTAGGTAAATCTGCTCAGT

TAGGTTTACATACTTGGTTACCTGATGCTATGGAAGGGCCTACCCCAGTTTCTGCTCTTATTCATGCCGC

AACAATGGTTACTGCCGGTGTTTTTTTATTAGCCCGTTGTTCTCCTTTACTGGAATATTGTCCTGAAGCC

CTTACAG

>NODE_195_length_495_cov_19.002110

GGTGGTGGTGGTGGTGGTGGTTACATTGGAGGTGGTAGTGGTTTTTGTGTGAGTTGGTCCGCTTGGTGCT

AAGGCGGGTGGCAGCGGCGGTATAAGAAGAGGAGGAGATCGGTTGGACGATTAGAGAGGTCCATATCTGC

CGCCGACGGGCTGTCTTAACTTACGTGCGTGCGTTAAAAAATACTAAACTTGGTTTGATTTTGATTTTTG

ATGAACGTAGACTTTTATTTTTTTTAGACTTTTTTTTAGCGTTTTGTGAGGTAACAATAGCAGTAACTTG

TAGTTCCTGGTCGTATCGTACGTATTCTGCTGTAAAAGTAGCAGTTCAGACATTACAGCAGTAAGTAGAA

TTTGAGAGAAAGGGCTGCGAGCACAGCAGCAGCAGCGGTAATTTAGAGACTGGGGAGTACGACAAGGGGT

AACACTAGGTGGAGCGATAACCAGATTCATTTTTTTTACATCCAAAAAAAAAAAATGGAATACACGGGTG

CCAAG

>NODE_196_length_495_cov_7.867089

TGGCACCGAGAATTCCAGCCCCAGCTTCCGACGACCTCTGACAGTATAAATACGAGTTTAGTACCAACAA

AATCCATCCGACCTACACATATATTGTAACCTTCACACACCTTATCGCCATTCTATGTATCATACAGCTC

GCCGATCCTCATCATCGCTAGGCTTTGCGGTTCACAGAAACCTCACTATCAAGAGACAGGATGCCAGTAT

TTAAACCGCACTACTGCGCTGCCCCACAAAACTAAAGCTTCTGACCTCCCCCACCCGATAACCCTCAACG

ACAGTTTATCACAGCCTGGACTGCTTCGCTTTCTTTCCTGCGACTAGTCATTCCTTGATCTGCAGGTGCG

CCTCCTCCCAGTCGTAGTTCTCGTTGTACTTGTAACCCATCTTCTCCACTAGCTCGTCGCTCTCCTTGGG

GCCGCGAGACCCGAACTCGTAAGCCACGGGAGGGATCTCTCCGGCGTCGATTGCCTTAAGGACGGGCGTG

AAAAT

>NODE_197_length_494_cov_4.293869

TTGAGGACGGTGGCTTCAACACGGAGGCATGTCTTCCACGATTCGTAGTCGGGCAGAGCCGCCACGAACT

CTTCCTCCGAATCGAAGCGTACCATGTCACCGTCCTCATCCTTGACGGCGAAGACGACGAGCTCCGAAAA

CTGTGCGACCGGTAGCTCGGGGTAGAGGTCGTGGAGCGAGCGGAGCAGGAACTTGCAGAGGGTCAGGTGG

TAACCTTCCGCTTGGACATGGCCGAAACGACCCCTGGACACCATGCTGTTGTCGAAGCTGAATCCGAAGC

GCCTCAGCTCGGACGTTCCATCCTCGCGGGTGATCTTGACCTTTAGGGGTGCCGTAGTGAGAGACATCTC

GAAGCCTCTTGGTACAACCTCGTGCAGCTGTAGTAGTAGTTAAAAATGAAAACGCTTGAAAGTTGAGGTA

GACCTCGACAGTTAAACTTTTGTGTAGTCACCAGGACGTCGCACAACAGCAGTAGCAGCAGCAGCAGCAG

CAGC

>NODE_198_length_493_cov_4.286017

AGTCTTGTCGTGCCCCCTTCGCCTTATGCATCATGTTTTTCTTTTGGGTTCGTCTTCTGTATCCTGCCTG

TCCTTCGGATCGCTTCGATCATAACGAGTAGTCATCACCGCGATACCCCTCTGGACTGTGTTTTGTGTTC

CATGCTAAGTAGCCCCCATCGTTTCCACCAGGACTCGATGATCAGCCTCGCACTCGTATCTCCCTCAGGT

CCCTTCAAGACCCCTGGACATATGAGCCCCCGCACATACGGCTGTACCCTGAGTCTCGCTTTCTTCTACG

TTTGTCTTGGTACCTCTTGAGCTTCGTTTCGGGTTTATTGTGTCTGCTTTGTCTTGATTTCGACCCCTGC

TATTCCTGTTTTGGTCATGACCCCTCCCTCAGCGGTCGCTTCTGTCGACGACCCTATCCGTGATGCTTAG

GCCTCACTACGAGGAGTGGGGAAATCTATAGCCTCATTTTTGTTGTTTTGAATGGATTTCGTTTTTTCCA

TAG

>NODE_199_length_493_cov_2.985169

AAATTTTTTTTTTTTTCGAGTTAAAGATTATTGCATTCCTATTCTGACCCCATACCAAAGACACAAGACA

TATTATTTGGCTGCTTCGTCCTTCCTGCTGCTACAGTCGAAAACACGCCCTCCATCGTTCCCATGGCGGC

CAGGAGCCCCTCCCGCAACGTCCGCCAAAAAACATACACCAGCGTCACACCACATACATATGGCAATATC

GCACACACATATACCTGGTATACATATGCACACATGCTACGTACGTCATTTCCCTACAGTACATACACAA

TGACGTCACAATCCGGCACCACCCGCTTAGCGAACATGGCATGTGCCGCACAACGCCTAGCCCCACAAGC

AAAAACCATCAACACCAATCAAAGGCGTACGTACGTAGCCGAGCCTACTCCAATTAACTTCCGTTTTCGA

ACAGCCCTACGTAGCCTGTACCAAAAGTGTTCTTGGAAGCTTACACGTAGCGGACAGATCATGATATACC

TCT

>NODE_200_length_492_cov_7.585987

TTCCACATCCCTCCACTGGTGTGCGACGGTGCCGGTTACCTTACCGTAGTGCCAAAGACTAGTGCCCAAC

AATCATGATACAAAAACAATCCGAATATGACCCCCCATAGCGTGATGGGAGTCGGGTCTTACACGACTAA

CAATGAAGGACATCGCGCTTTTGGGTGCAAACCCCAAAATTGGCACGATATTTACAACGTGTGTGGAAAC

AATTGGAAGAATATTTACATTTTGGACGACTAAATACGCAAGACACTACATTCGGCTGAAGACATGTACA

CTACAAGGTGACTACAATAAAATTCTATACTGGGCACTGCGTATGTCGACGTGAAAACTAATGCGAGGGT

TCTCGCAACGACAGAATGTCGACAAAACGCTGACAAGTTGGCACGCACACTAAAACACACATACTCCTTT

TGGCTCTGACGCCAAGATAAAAGGAAGTCAACTTACATTACACAGGGTGTACGATATAGTGGTCACCAAA

CG

>NODE_201_length_491_cov_7.874468

GCCAGCATCGAGCTGTGCCACACAGTTATCTTTTTATTATTTTTCTCTGGGTGATCATCACCCGGGACAC

TCAGGACACGACTCGACTTGCGTGGCTGCTGTGCACTACGGTTACACTCGGACACACAAGCAGATACTGG

GGCGTCATCAGCAAACGTCACCCGCTTGCGCGGACGATTCGAGCTGCACCCCCCACTACTTCGACCTGAG

ACAAGGACACTCTTGACAGACAACAAACCACGATCAACTCGTTGGCTACTCCCATCATTGACAAAGACAC

TCGGTACAAGAGACTCGGTACCTTCTGCTACAGCGGAGAGTTTCCTCTTCGCGCTAGCCTTGCCTACCGT

ATTAGTAGCAAGGTGTGTCCCGACTGCTGCAAGCGCTACAGCAGCTGGGCCACGTCAGTTGCCGTCTCCG

TTGGTGCTGCTGGCCTGGCTCCTGCGGTCATCTTCGTCGTCGTCGTCGTCTACTGTAACACGACGCTGGA

A

>NODE_202_length_490_cov_8.179104

CCCCGTAGAAGGTCATAGTGCCGGCGTTGTACAGCGACGTGCCTTCGGAGCTCCTGCAGTTCGTGTTGCT

GAATGTGGCTGAACCTCGGAAGCCCATGTCGGAGCCGGAGTCGTTGGTGACGTCGCAGCCTTTAGCGCCC

TCCAATTGAGATCCGTTGTCCTCGAAATCCGCCCTCGCATCGAATCTCACAATCCCACCGTCGAAGTTGT

CGATACCGACACCATCGTTTCCTTTGATCGTCGAATCGCCGCCGAACTTTAGCGTGCCAGTGTTCTCCGG

GATCGCATTGATGTTGTTTTCGATGAGCGATGTCTTCTTGAACTTCGTTATCCCTTTGTTCCGAAGCAAT

CTGCCCACGTTTTCGATCTCCCGGATTGTGGTGTCCCCGTTGAACGTAAGGCTGGCACCGACGTTTACGT

CAAGCAGTGTTCCCCTAGCACGGTCGTGGTGTGACTCATCATTAGGCCCAAATAGGGTCTCGGGCATGTC

>NODE_203_length_489_cov_5.852564

TTTTTTTGTAGCACCGCTTACGTCTGGACCCCTTCTTTGTTTGTTTCTTACCGTGGCAGTGGTGGGAGTA

GTTCGGGAGTCGAGAACGTGTGTCGTCCGCGAAATCAATGTTTCGATTTCAGTAGCAAGGAGTGCCGGGA

CACACACACGAGGACAAAATGCGTTCTGACCAGCTGGCGCCGGCACTTCGTTCGGGCCCTCTTTGTTTCA

GCCGCACGCTTCTGCCTTTTGCATTGTGTCGCTCTTTTTTGTTTCGTTGCGTGGTATTGTGTGTGCTCGT

ATGAGCAGTTTAGTTTCGGATCGAGAAAACCCACTTCGTGTACGTACGTGGCTCTGCGAGATTTGTTTTT

ACGTGCATTTGTTTTTGCCCTCTAAGAACGTATTACGCGTCTGTCTGCTGTTAAGGAGCGTATCAGGGCT

TTTTATTTGATTTTTTTTTTACCCGTTTATTTATGTTTTCGTATACATAGGTTTTTGATATTTTTACTT

>NODE_204_length_489_cov_2.747863

TTACTTCAAGAAGTGCAAGGAGGCGAACGGCGGCGCTCCCATCTACAAGATGCACCCCGGTCTGGCGTGT

ATCGCCATCACCGACCATGCGAGCACCAAATGGTTCATGTCACAGCCTGAAACCGTCCTGGACCGACAGG

ATGGGGCGTACTTCGGACCGTTGAAGTGCTCGAAGGATTACATCGGCGACTCGCTCCCGCCCCTGATCAC

GAACATGAAGGAGTCCCACGCGGAGGTTCGCGCGTACATGATCGAAGTCATGATGGAGAGGGTTGCGGGC

ACGCAGAAGGCGCTCACGCACGCCTCGGACAACTTTTACGAGCACCTCAAGGCGAACGGCCTGGGAGAAC

ACCCCATGGTGTACGATACCTTCCTCAGGCAGTCTTACGCCATCATTCTTGAGTGGGCGCTAGGAACAGG

AGAGGAGGGTGGCCAGCCCCTCCCTCCCTTTCAGGACTTCGTGAACGTCAACCCTATAGACGTCAGCCT

>NODE_205_length_488_cov_4.301927

ATCCTTCGCGCTCTGGAACAGATCATCAACAGACGCGTGCTGCAATAGGCGTAGTACCAAAATCTGCCGA

GGTTCGTCCCGTGAGTTACTTTGTATCGCACCACCAAAGAAACCACACTACTGTATTGAGAAACGTTATG

GAAAGACCGATACACATACAAATGGGCTTCCAAATACAATATTCAATGCAACATGGAATGCGGTGGGAAT

GGGTAGCTCCGCCCAGACGCAATGGTTGCGCCTCCAAATCAGTAGGAACAGTTTGGGTTTTGGTTAGAGT

GCAACTCCTACCGCAGCCCCTGTACAACAACGACTCGCCCTTTCAACCATCGCTCCCGCCCAACAAGTTC

CTTCCGCTAGAGTAAGGTACCTGCTGCGTTCACACATTACACGTAGTGGACCTTTCCCAGACGATTTCTA

CTGTATTAGTCACTCGAGCCTCAGCAGGGGTAGAAGCAAACAGGGGCGCACTACAGGAACTAGAAAGC

>NODE_206_length_487_cov_3.624464

CCTCCTCTCCTTCTCCATCGGCGGCAAGGAGTACACACTGTCGGGAGACGAGTACGTGATCAACGCCGGG

AACAGCAAGGTGTGCATCCTGGCCATCATGGGCATGGACGTGCCCGAGCCTATGGGCCCCCTCTGGATCC

TCGGCGACGTGTTCATGCGCAAGTACTACACGGTTTTCGACTACGGGAACGCTCAGATCGGACTCGCGGC

CGCGGCCTAAAGAGGCCGAGTGGTCGGTCTCGCCCTTGGCCTCATTCAGAAGCTTCGCCCTCGGGTTGGT

TTTTATCAACTTTTTTTCTTGTAGCTGAAACATTGTTGGAGCTCTAAAATCCCACGTTTTTTTTCGTCTA

CAAAACGAGGATGCGGTCGTTGGTCAAAAGGGGTTATAATGCTTTACACCACTGAAAAAAAAATCGTCCC

ACACTATTTTCAAGTAGCATGTCCCAAAAAAAATGGTTGCAGTTCTTGAGGGGTTTAAGGAGCCGGT

>NODE_207_length_486_cov_8.823656

AAATATTGGATTGCTCCAATTGCGGTCAAGTGTTTTCGATTCACATCCTCAGTCGTTCCGCCCCACCCCT

ACCCCCCCGCCTTCCATGCAGCTTTTCCGCCAAATCTCCCCAGCAGCAGCACAAACCGCTACGAAAAAAC

TCACGAAAAAAAAAACTCCTGGAAAAACAAGAAAAAAGAGCGTAACTCACGACCAAAAACGAAATACAGA

ATCGTCCACAACAAGACACAGGCCACAAATCAGCCCCCAAAAGCCCCGCAGCTACCGGAAGTACGTTTTC

AAGAAAAAAAACCATCAGAATCAAAATCCTTCAGGTTTCCCTCTGTCCCTCGCTTTTGTCATCTTAAGGT

CTCCGTCGAAAAAAACGGAACGTTTTTAGGCCGACAAGAGGCACGCTCCGCCTGCCGCGCGGATCCTGTG

CTCCGCGAGCTTGCTGAAGATTTTCGCGCGGACGTTGTTTGGCACCTTGTGCAGTTGTCCCTTGCC

>NODE_208_length_486_cov_5.731183

ACATATGAAACCAATAGGGGTTTTTGGTGTGCTGTAACTTGTTGTTTGTTTTGACCCGATGTTTTGGTGT

GGTTTCCGGCTTGGGCTTGTGGGGGGGCGGGGTGTCAAACGGTTGTCCGTTGGAGAGAGCCGTTGTGCTT

ACCTCTTGCTAATCGTGTATGTTCCGTTTTTTTGCGGGAATGCGGAATGCGAACATAAGTGTGTCTGTGG

TGTCAGGCTCTGTGTAACGTGTAACGGTATGTTTACGGTAGGTCGGCAAGTTCACACAATGGAAATTGCT

TTTCGAGCACGAGGTTGTTGTGGCTCCTCTTTGGAGTACGAGTTATTGAGGTAGGCAAGTCTTGAGTGTA

ACTCTGGCGTAAGGAGACCGTATCATCCAACCGAAAACAGTGTAACAAAGGTTGTTGAGCGATATGGTCG

CGGGTTAGCCCGCTGCGTTGTGATGCGCCGATATCTTGTATCGAGCAACGAAGTTATTATTTCGAA

>NODE_209_length_485_cov_17.463362

GTTCCTTGGCTCCCGTGAATTGCAAATTTTTTTTTTTAGCACGTGAGTATGTTTATTCCAACAACAAATC

GATCAGTCCTGCCGGCGCCGACACACAAAGCTGACGACGAAAGGGGACCAACAATATTGTCCTCCCCATC

GCAAACCACTGGCACCAGCCGTACAAAAGCTGGAGGCGCACGCGCACAAGGTAACCAAGCAAGAAACACA

AAAGGAAAGCCAAAATAAAATCAGAACGCGCCCTGGATGATCAACAGTAACAAACGACACAGACCCGACC

AAGTCGAAAGAAAAACTAGAACCAGCAAAATTCGCGATTCGCTCCGTACGAAACAAGCCACGGGCCAGGA

GTTTTTCGCGACCCGAAGGGAAAATAGCCAGCTCGCTACAACTCAACCAGAAAATCCCCCGAAACCCCAA

GGGAAAAGCAGGAAACGCCAGGCATTCACTTACTCTCTTAACGGAGCAAGCGTCGCATCTACATG

>NODE_210_length_482_cov_4.156182

TTTCTCTCCACGAATCGCAAAAGATTGCCCTTTTGAAATTCAAAATGCCCTCTACACCGACGAAAAAAAC

TGTCCTCTCCACAATTCGCAAAAGACTGCCCTTTGAAATTCAAATGCCTATACACCGACGAAAACTCTTC

TCTGCACTAATCGCAAGATTTGCCCTTGAAATCCGAATACCTCTAACGCCGACGAAATCTCTTCTTCATA

ATCGAAAGATTTTGCCCCTCCCCTTGAAATCCGAATGTCAACTCACTACCGACGAAAATGCCCCTTTCTC

CATGATCTCGCCGTTGATATTTTCGAATTGCCGACCGACGGAAATTATGTTCCGACAAGATATCGCCCTC

GAAAATCAGTATGTGTGAGAGAAAAAAAAAAGACCATGATCAAGAGGATCGGGTATAAATTCTCCACAAA

AAAGGAGCTTCGGAGTTCCAATCCAGTTCCGAAGGCTTATAAACGTTGGGGTTTCACACACA

>NODE_211_length_481_cov_12.000000

GTCCAGGCACGACGTGTTGGGGATGGCCTTCCCGCTCTCGTACTCGCCCACCACTTGCGGTTTCTCGTTG

ATTTGCGTTGCAAGTTGCTTTTGGGTCATCTTCTTTCCTAGCCGTGCTTGTTGGATGGCCTGCGCAAGCC

CGCGGTCGACCCGTGTATGCTGGAACTCCTCCGTCTCGTCATCTAGCTTTTTCATGTTCATATGTCCGCG

GTGAGTGCTCTTATTCTCTCCCGCGGAAAACTTGCGTTCTGTCTCCAATGTTCCGCTACGTTGTGCGGCG

GCTGTCGCGGCATTTTTCTGACCACGTGTCGCCGTGGGGCGCGTTTTGGAGATCATAACCGTACTCCAGT

CCTGAGACATCTTGGGCAAAAGCTTTCGAAAGCGATGGGGTAGGGTGGTATCAAGTTGTCTGAAAGGTAC

CGAGGGTGGAGGTCACAGAGGGTGGCTTGGTCGTCGTGATGACAACCAGCTCAGCGGACCA

>NODE_212_length_481_cov_8.767391

GGCAGACTTTCCGTGCACCTGTAGAGTGATGCCGTCTTCAACATACAAGCTCTCGGTGAGCAGCCACGTC

CCTGTTATCGAGCTGCTCACGTCGCCGCTGCTCGATTTCACGGCGTACAGAGGGGCGCTGCCGTCCAGGT

CTTTCCAGATGTCCTCGAGCGTGACGCAGCCGCCTCGAGTCTTGCCGTCGGCCGACTCCAGGTAGAGGCG

CTTCGACGTCGAGGAGTACCGGATGGACACCTTGGTGCTAGACTTGCACGACGGTTTCGCGATGGCCTTG

TTCTTCGAAGCCGTGTTTGCCGCGTCGGCTCCCCTCGCTGCTAGCAGGGTCATGACCGCAAACGCCCGAA

GCATTGTCAGCATGGTGTCCTATCAGACGCGACACACTTGCTTGCTCTCGTTTGGGAGATAGCAGCAAGA

TGAGTGCTGGGCTGGGGTGTGATTTGGTGTGGTGTGTTGTATTGTGCGGGGGTGTGTGTGA

>NODE_213_length_479_cov_4.069869

TTGGCACCGAGAATTCCAGCGTTGAACGAGAGCATGCTACTCTCTCACCCACAACCCACAACGTACACCC

ACCAACCTATACTTGCCCATGAACCAAAAAAGGTTTCTGTGCTGACACAAATCGCTTAAAATACAAAAAG

ATCCAATCAATGACAATTGCAGGAACACAGCAAAAAGAGAAGAACCGCCAACGCAGTAGCTTAATGCAGT

TAGGAGTTTGACCACTTGAAGGCGTCTGTCACGCTCCTCATTCCCACCGTTCTTGTAATGAGGAGATGTT

GATGCGAGAAGGCCGCAGGCCTTCGAGCATCAATAGATGCTCTTCCTAATGGGGTTGCCTGTGTGTGCCT

GTGCGGGTGTACCCCCGTGGACGCGAATTGACACAAACGGACAAAAAACAGTTCTGAAAGCTGCCACAGC

GTGGCTTTATTTAAGGAATCCTTAATATTGAGGTTTGTCCTTCGTCTGTTCGCTTTGCT

>NODE_214_length_478_cov_15.006565

CAGGTCGGCCTGGGTGAGGGGGTCACCTCCAGAGGTGGTAGGGAAGAGGCGCGTGGCGAGAAGCGTCCAC

GAATCCTTTTTGCCGGACTCGTCGGTTGCGACTCCTCCTGTGTGGGCATCCTCTGTGAAGCGGTAGAACA

GTCCGTTGCTTTGGAAAGGGTGGTCCCTGGTCACATGGCTGAAGTAGTCCAGATCAATTAGGAGATTGCC

GATCGCCTCCGCATCTGCCGTCGTAGAAATTTTGGAGCTGTTATTGATGAGCCAATCCGCGGCGTCCTTC

CCGACGAAGCATTGCTTGTACGTTCTGAGTCGGTAGGTGCGGTCCTTGATTTCCAGTCCTCTGTGCATGT

GCTCCCCAAGGCGCTTGATCTCCTCCTCCATCAGCTCCTTCTCCATGTCAGCCATGCTTTTGTCGATTTG

GTCGCGGGGACGCGATGGGGCAACCGAGTTCTCAGCCTGTAATCGAGCGAGGGATGAG

>NODE_215_length_478_cov_5.603939

TCTCTGCCCCTTGCGTTTGCGTGGACGATCAACTTCCACTGGTAGCTCAGTAGTAGTGATTGTCTGCGTC

GGTGCATTCTCCATCTCCTCCTCCTCAGGTAACGGTGGTTCTGCCTCCCAGACAACCTCCGGTTCCTGGG

AAGTGGTCTTCGCCTGCGATCTCGTGACCGCTCTCGTCACCCCCAAGGTTTCTGCTGCTGCGAGGTAATC

CTCAAACACCACGCAGGCCAATCCTTCCAGTTCCACGACCTCTGGCACCGGTTTTCCCAGTAAGCCTTGA

ACAGCTAGTTCAACTAACACTGCTCGTAGCATTGGAGGAGACGTGAAATCTGCGATGACGTTCGTCGCTG

CGCTTTGGCTGACCGTCTGCGGGATCACATAGTGTTCCTGTCCTGTGGTGATCTTGGCGTAGTCGTTGAA

GTGCAACTCCTCAGCTTCTTCAAGGCGTGCCGTATCCGCTGGGTGTGAGACATACCCT

>NODE_216_length_478_cov_4.185996

GCTCGGCCTGTTGAGTGTCTGTCTCCGATGGGTCTGAGGGTGTGAATGCGACGGCTGCCAGCACTACAAG

ACTTTGAGTACTAGTTTCGTCCACACCTTCCACTCCTAAGGTTTCGAGGTGTACACCGTCGAGCAATGGC

CCCTGTGGCCCCTTGTAAACGTTTGGCAACGACTCGTCCCCTGGAAAAGAGTCGTCGAAGTCCTCGATCA

CGTTGCTCTTGTTCGGTAACCGCGACATGGCGTCCGGGAGTTGATGTTGTGTTCCCGGCCTCCACTTCAA

GATTATGTCGTGCTCCATCAACCTCAGAGCCCAGCGGTGGTATTTCGGTGTGAGGGATTGGCTTTTGAAG

AGCCAAATCAACGCCGAACAGTCCGTGATGAGCGTGAACGTCCGCCCCCAAAGATAGGGTTGGAACTTGT

TCACCGCCGACATGACCGCCATGCATTCACAGTCTGTTGCTGACCGTTTAGAGTCCCC

>NODE_217_length_478_cov_1.382932

GCACAGATTGGATGCATCAGATCCCCTACAGTGGGGCTTTTCAGGCAAGATAGATCTCCGATCTGTATAA

TCTGCGGGATCTAGCTCATGTTGTGGGGTGGGAGCCGAATGATCTGCATGGTCTAGCGCATGTTTCCTGC

GTTGGATCTGTACTGTACAGATAGATCCTGCACAACCTCTCACAACGGCAGGTGAGGAACTAGATGATCT

AGATCAGATCTGTGTGATCTATAGACGATGTAGACCATGGTACTTCCGGTGTGTGTTGAGAGGGGTCAGG

GTTTGGGTGTACATTTCTTTCGTTCCTTCCTTCGATCGTCTCGCCCTATTAATGGTGTCGCGTGGCAACG

CGCTCAGTTCAGTCAGTCATCGAGTCGATTTTTTTTAATCGCGTGAGGCTTGGTCCATGCTGAACTGTCG

GGTATTACGTAGGTCAAACAACTTAACAAGAAAAAACGAAGCTTCAAAAAAAAAATTT

>NODE_218_length_477_cov_12.890351

AACAACGGCCTCCTCGGGTAAGGCATCGGCCACAGTGGCTGCGTCGACAGAAGCAGCTTCGTCCGCAGGG

GTGGGGGTTGCGACCACCGCCACCTCCTCCTCACACGTCGCGTCTTCCACAACCGCCACGTCGATGTCCC

TGGTAACCGGAGTCTCAACAACGTCCACGACCGCAGATTCTACTGTCTCCACAGACACAGCCACACTCTC

GACGACAGCAGGTTCAACTTCAACGGCAGGAGTCTCGGGCGAATCCTCAGTGACGGCCGCCTCCTCAATC

GGAGCAGTATCCGGGGTCAGAGCGACGACCACCGTCTCGTCCACCTCTGGCTCCGCGTCTTCCACAATCT

CGGCCACGACCTCTCTGGTCACTGCAGAAGCGGCAACGTCCACGGCAGCTGGCTCTTCAGTCTCCACTGG

CTCAGCCACAGTCTCGACGGTAGGAGGTTCACTCTCAACAACGGCCTCCTCGGGTAA

>NODE_219_length_477_cov_6.938596

ACAACAAGTCCCACTCCGTAACGGCGATGAAGGCGAACCCCCTCGGTGGCTCTTCCATGGCCAAGGGCAT

CGTCTTGGAGAAGATCGGTATCGAGGCGAAACAGCCTAACTCCGCTATCCGGAAGTGCGTGCGGGTGCAG

CTCATCAAGAACGGGAAGAAGCTGGCGGCGTTCGTGCCCCGTGACGGTTGCCTCAACTTCACCGACGAGA

ACGACGAAGTTCTTATCTCCGGGTTCGGTCGCCGCGGTCACGCCGTGGGTGACATCCCCGGTGTCCGTTT

CAAGGTCGTCAAGGTGGCGGGTGTAGCACTGCTCGCCCTCTACAAGGACAAGAAGGAGAAGCCCCGCTCT

TAAAAAAAACGCCGTGCGTCTCTTCACAAAACGTTATGGAGTGGGAGGGGGGTAGTTACGGATCAACGAA

GGAACGTATTCGTATCTCTTTCGGGGTCTACTCCGGTGGTTGGTGCGGATAGCTGAC

>NODE_220_length_476_cov_18.134066

CTCCGCCCTTTTCTTGATTGCCTTTTTTACCTAACTTAAGAATTTTTACAAATACGGCATTGGTACGTAA

GGCTCAAGAATCCGTGATGGAATTGCTTCTTCTCCATCACCCCTTAGATTGTCCTGTGTGTGATCAGGGA

GGGGAATGCGAACTTCAAGAACAAAGTTTCAATTTTGGGTCGGACAGAGGTAGATTTTTATTTTTCAAAA

ATTCTTTAGGTGACACTTTTTGGGGGGGTCTTATAAAAACAGTATTACGTAGGTGTATTGTTTGTACACG

ATGTGTAAGATATACTCACCAAATTGGAGGAAATGATTCATTAGGAACGTCTAATAGAGGAGGGCATTCT

GAAATTGGCATGTTTAAAGAAAGTGTGTTAAAAAGTGAAACTTCGGGTGGTCGAATTCTATGGTGAAAAG

GAACTCCATTCAAAACTACATATCCAATGAACTCAACTCAAACAAAAAAAAAATAT

>NODE_221_length_475_cov_12.312775

ATCAACCAAGCAATGACGCTTATTGAGTAAAAGGAAATGGACTGCTTACTCAACCAACCAATGATTAGGT

TACCAGGACAACACGGATTTTAGGACAGGCTCAGGATGAGTATCACAACGCAATGTTGTAAAAAGGATAG

CGTCAGGAAGATGCGAAAAGGATAACAACACGGCTAAGTTGTTTAACAGGATGTTGAAGGAATAAATGGA

CAGGCCTTCAAGGCAACACAATGGATTGGTTAAAATAGGGATACACTAGGAAGTGTAATTAAGGATTTGG

TAATAGTATTTACCCTACAGGATGTTCAGGAATTTAAGGATACAGGCCCGGATGGTCTACATTACATAGA

ACTAAAAAGTAATATAGTAATAAACATGGATTAGTGAAAATTGGAGTTGATTAAGTTCAATTGCAGGATG

CAGAAAATTAGAAGATTCCGCAAGCGCGCCTAGAAATAGAGCGCGCTTTTCTGGA

>NODE_222_length_473_cov_4.685841

GTTTCCACCCCTCTTTGAGTACTCTTTAGGAGCTGTAGTTTCGGTAGATTCCCCCGTCGGCGTCGGAGTT

GATTTTCAGGCCCTTGGCCGTCGCAGGCCGCGCCTCCATCCTTTCCCGCCAGGTCGCGACGTTCTTGAAG

CTCGCCAGCTCAAGCGTCTCGTACGCCTTGTAGAACACGTCCAGGCACCGGACCCATGGGAGCCAAGCCA

TGTCAGCGATGGTGCACTCCTCCCCGCAGAGGTAGCTTTTACCCTCCAGCTGCTTGTCAAGCACGTCCAT

CGCGCGCTGTACTTCGGTGGAGTACCTGGTCACGGCGTAGTCGATCTTCATCGGCGCGTACCGCGTAAAG

TGGCCGAACTGCCCGAAGTATGGCGCCTCTCCGATGTTGAAGAAGAGCCAGTTGAGGCACTCAGCCTTCT

TGGCAGGGTCGGTCGGGAGGAACTTGCCAGTGAGCTCGGCAAGGTACAGCAGG

>NODE_223_length_471_cov_16.231111

GACAACAAGTACGGCATCAGGCTGTCCGTGGGGTCGGCCGACAACGTCTTCTCCAAGAACGACATCACCG

GCTCTTCCAAGTACAACGTTTACACGTACGAGGGGTCTGATGACCCGTGGGTGGCGGATTCCGGGCGGTC

GCAGGACAACGTCTTCTCGGACAACAAGATCAGCGGCGGCGAGGAGACCATCAAGCTCACCAAGGCGGAC

GGCACCGAGTTCATCGACAACACCTTCAAGAGCGCGAAGACGATACGCTTCGAGGACTGCCAGGGAACCC

TTATGTCGGGCAACAGTGGCCTCGGCAGTGCTTCGCTGAAAGTCACCGACGGATCTTGTTTCGACAAGAA

GTCGGACAGTGACTACAAGCCTGTTTGCTAAGCGGACAGCGGGAACGTCTATATAGGGCATTAGGCATGC

ATCGCCCCTCTTGCTCTGCCTTTCTCGGCTACCAGTGCCGATTGGTACGTT

>NODE_224_length_471_cov_2.671111

ATATAATACTAAGGTAAAAAAGTACATGCCAGTAATTAAAGTAAGAGAGAACGAACCGTTTGACGTAGCA

CTTCGTCGCTTCAAGCGTTCATGTGAAAAAGCAGGAATCCTTTCAGAAGTTCGTCGTCGCGAATCTTATG

AAAAGCCGACTTGGGAACGTAAACGTAAGAAAGCTGCTGCTGTAAAACGAACCACGTTATATTTCTGCGG

TGAATTATGAGTGAGTGTGATGCGTACTGTTTTGTCGTGAGATTTGACGAGCGTTGCGTGATATTATTTT

GCTGTGGTGTAACAAACAGCAGGTGCACACTGGCCCACATCATTACCTCTGCGTGGGGTACACAGAGGGG

ACACTAAACTTCGTCGATGGCTTTGTCTATTATCATTGCTGTTGTAGGATCTTTTAACGGTTGAAACCTC

CAATAATAACGACGGACGTTTTTTCTTTTAAAAAAAAAAAAAATTTTTTTT

>NODE_225_length_470_cov_6.971047

CTAACCATTGGACTACGAGGGCATTGTGGTCCACTTTCGGTACGTGAATTTGGAATGTGGGATATTGATT

TGGGACGATGATTTGTGGTTTTTGGGGACAGCTATGAGGTATGACCAAGCTATGATATTGTGATGCTTTA

TTGCCATACTGTTCACCATGCAGGGAGGATATTAGTTGATGGGACTGTACACCTGTTCGGTATGCATACT

GCGGATGTGTAATGTCGCACAGTATCGATTACAAAGCAGGGAGGATATTGATTGTGCTTGGTACTCGATA

CAGCAGGTATGATGACGGTTACATATATTGGGGAGTTCATATACTTGGTGAGGCTTTATTGAGGTATGAA

CGGATGCTCGGCTCGACTCTGCTGACTTGACTTGCTTGCGGCTTCGCCTTGATGGTGGCCTGCGAGGTCC

GACGATAATCTGTACCTCCGCGATATCGAGCGAAGTTTCAGCGACGACTA

>NODE_226_length_469_cov_5.125000

TCCAATCATCACGATCGGGCCGTCCAAAATGGCTTCCACAAACTCATTACCCTTCAAGTTGGGTCCTCCC

TCGGCATCATTCCGGTGAACGAACTTCTCCAGCCAGTTCAGCGTGTAGCGGTCTCCCGAGATTCGCAGCT

CGGAGCGGTAGGCTCTGACCGCGGCCATGGTGATCATCTCCTTTAGGAGCTTGTAGTTCTTGCCCCTGTA

CGGAGAGCTCACGTTGTTCAGAGGGTCGTGGTCGAAAACCATCTTCCTGCCCTTCTCCGCAGATTCATCG

TCGAGGAACATGGCGTCCTCGCGATGCCTCTCCAGTTCGTTGTTCTCTGCAGCGATAAGGCTCAGGTCCT

TCACCCACTCCCGGGAGATCTGCTCCCGCACCTGCATAATCTTCTTGGCGATCTGCTGGGGCTCGATGAG

CACGTTGTACTCCATCACCGACCGGTTGGCCTTCAGGAAGGGGTTGTTG

>NODE_227_length_468_cov_5.617450

CGGACCCGTTCGTCCGTCCATCTGCTGGCGCACACGCGAGTCACATCGCATCAAGGACACTGTATCCGCA

GCCTGGTTCCATCGGGCGGCTGCTTTAGTGCTCTTGTTGCGTTTGGGTTGGTGTGTGTTAGAGTTGGCCT

TTTTTTCTCTCTGCCGCTGCGGTGGTTTAGGGGGAAAAGGCGTCGGCTCTTTGCTGGTGGATTCTTCAGA

ACTCGTGTTTACCCGGCGGGTATTTTGCCTCTCGTGCGTGCGGCAAAACTGTCCCGTCTTCTGTTTTTCA

TGGAGCAATTCCGGTGCGCTCTCTGTTCCTTCTGATTCGTTCTTGGTCTGCCTTGTTTCCATCAGTTGGT

GTCTAGATTTTTAGGCACCGGCTTTATGGGGGTGGTCGAGGAAGTAGTTGAGGAAGGAGGGGGGGGTTCG

CTGGGCTCGTGCAAGTTCATCCGAATATTCGAAAAATCAAGCGGGGTT

>NODE_228_length_468_cov_1.932886

ATAACAGCGAGATCAGCATGCGTATTCGTGTTGAACGCGGTCGTGGTTATGTGCCGGCATCTAGTCGTTT

ATCCTCTGATGACGATGAGCGTCCAATTGGTCGTCTATTAGTAGACGCGACTTTCAGTCCTGTTGAGCGT

ATTGCTTACAGTGTTGAATCTGCTCGTGTTGAACAACGTACCGATTTAGACAAACTTGTTATCGATATGG

AAACAGATGGTACTTTAGAGCCTGAAGAAGCAATTCGTCGCGCAGCTACAATTTTAGCTGAGCAATTAGA

TGCATTCGTAGATTTACGTGATGTAACTGAGCCAGAAGAGAAAGAAGAGAAGCCAGAATTCGATCCTATT

CTACTTCGTCCAGTTGATGATCTTGAATTAACAGTACGCTCAGCAAATTGTTTGAAAGCCGAGCAAATTC

AATATATCGGTGATTTAGTTCAGCGCACTGAAGTTGAGCTTCTTAAAA

>NODE_229_length_464_cov_28.984199

CCCCAAGGAATTATTGCTCTGGCCCCGGGAGGGAAAATCGCAGATCAGAACCGGGGATCGGTTTCGGCTG

TCCCCTCTTCCGAAATGCCTTTTTTTCTCCCACTCCGGTCGGGCCGAGAGCCCAGACACCTTGGTACAAT

AATATTGGGTGAAGTGGTTTTGGTCGCTAGGGGGGAGTATTTCCCTCCTGACCATTCTCTCTCCTTTTCG

GAGGCGAGAGGCCCCATCCGCCGTCGTCCTTAAGGCTGCTCCTCATCCCAGGGGATAGGCGTTTTTAGAG

GGAGGGGGTAGGGGATGCCTGGAGAAAACGGGAAACTGTTGTTTCTGGGCGTCTTTTTTTCTTCTTCCTT

TTCTGCCGTAGGTGGGGCGCGCTCTTTTGTGTCGCCTCGCTGTCCCCTGGGGCCCGGGTTTCCACCCTGT

CTCGTTGAGAGCATACGAGTCATAACGATGGTACAAGTTGGAAT

>NODE_230_length_463_cov_9.893665

ATTTTGATAAAAACTCTATCATGCCCCCTAGATAATATAACTTCTTATTAGTGTACCCCCTATGCCGCTA

GTATAATTGAAGTCCCTAGTGGGTTGCGCGCTCGGCCTGCTCTCGTCTCATGGTACTTCGTTCGGTACAT

CGAAGGAAAGCGTAGGTGCACCCTTCAATCCGTTCTTATGTGTAACCTCATAATTGTTTCCCTATGGAAC

TATCTCACCGCTCGGAGTCGGAGTACCGTGGTGTTTGTGCCTTGCAGGGGCTTGCGCGTGTTCTCACTCT

CGATGTAGCTCGTGTACACGTGGGCCGTGTGTCAAAGCCAAAAGGGCTTGTGAACCAGTCCTCTTGCTCC

GCTTTTTCCTCTCTTTGCTTCCTTAGTTCGTTGTCGGTCCTGTAAACATTTACTCCCGAGCCGCGCTCGA

TGGGCATACAGCGCAGACAACAGCAAAACACCGAGAGGATCAA

>NODE_231_length_463_cov_4.110860

TATACCATCCAGGATGAACCTAGACGATATTCTACCTTCGTACCATTCACGACTTTCCCGGGTTAAAAAA

AAAAACATTCCATACCATTCAGAGATCCCTACCAAGTGAAAAACAATACATCATTCACCACTTACCCGGG

AAGAACGATAAATCCCATATAATTCAGAAATCCCTAACACGTCGCAAACGATATCTTTCATTCACAACTT

ACCCGGAAAGAACGATATCATACCATTCGTAAATCCTACCCAGTCGCAAATGATTAATCTCCAAATTCGT

ACGATCAAAAACATTGTTCAATTAATTTCGACCGACAGGTGTATCCAATCGTGAAAACAAAATTATCTTT

CTCACGAATACCCACCAGAAAGACCAGACGTTAACCCTCGCCCCGCGATGTTGCCGAATCACAAAAAGCT

CCGGCTGGCGATATCCACGTTTGTCCCCTCCACCCTTCCCGCT

>NODE_232_length_463_cov_3.373303

GGGGTCTGGACTGTGGTAGGCTCGGGTTACACTGCGTATGGTGCACGATCGGTAGGCGTGTCATGTATCG

GTCGTTCGCTCTCTCAAAATTTGATACGACGAGCGTTGAATTTTTGTACAGATTGGTCCGACCCCTGTGT

CGTCGACGAAACTGGCACAAAACCCGCATATAATTCTGGCGTGATTTCTTTCGGCGGTGCACATATTTTG

GGGGCGTTTTTTCGGTAGGGCGGGTGCGTGTGCGGGTGAAGTTCACGATATAAAAAGCAAGGAAGGGAGG

AGGGAGTCTGGTGGTCGTATTGAGCAAGAACCTACTTCCCTCCCTGCCCCGTGAATGATGGACGGCGGAT

TCTATATGCGTAAAACTCGGTCGGTGCGAGCTGGTATTTTTGGGCGGAACACGGGAGGGTTGGGCGCCTT

CATAACTCTCCAAGTATGTCTTAGGACGTTCGTGGGACGATGT

>NODE_233_length_463_cov_2.395928

ATAGATAAAGAGGAAGAAGCAAAAAAAATAAACTAGAGCTAGGAACAAAAAGAGAGGGTAAGCCTCTCTT

TTTATTTATAACAAAATAAATATTTACTTTAATTTTTACGCAAGTCTTGAGTAGTACTCAATAACTAGCA

TATCATTGATTTTAAACCTAAGATCTTTACGTTTTACTAAATTCAGTATTTTACCTGTTAATAATTTTTG

ATCAAATTCTAAGTGACTATTTAAAACATTATCATTTGAAGTTGCGTTTTCCCCTTGGTTTAAATTGGAT

TTGATTAAAGCTATTGCTTTAGGTTTTGCAGAGAAACTAATAGAATCATTTGGTCGACACTGGAAACTAG

GTATATTAACTCTTTTTTTATTTATTAGTACATGCCCGTGGTTAACAATTTGTCTTGCGGCAGGGATTGT

TGGAGCTAATCCTAAACGAAAAATGATATTATCTAGTCGCATT

>NODE_234_length_461_cov_10.440909

TACAAGTCTATATACAGTTGGCGTGATTGTCATTACCTTTAAACGACGGACTCACTCGGCATCACACACT

ACCTCTCGGTAGTACATATACATTATATACCAGTATATATCAGTAGGTTGAAGTACGACGCGAACTACAA

GTCTAGAAGTGGAAGTATTGCACGAACTACAAGTCTATATACAGTTGGAAGATGCCGAAACCAGTAAGTT

AGCCCCTTTACAACTATAAACCCGTTTTTGGGGGACAAATGACTCGGAATTATTACAGGGAGGGGTTTTG

GGGCTCTAAAGGGGTTAAAACAGGTGGTTAAAAACAGGTGTTAAAATCAGTTCGTCACAACGACGCGCTC

ACTCGGAATCATAAACTAACCTACGGTAGTGCCAGTAAGTGGAAGCACGCGATTTCAGGTCAATATGAAG

TACGTTTTCAAAGGAGTCCGCGGCTGGAATACACGGGTGCC

>NODE_235_length_458_cov_12.482838

TTTTTTTTTTGTTGAAGAATATTGAAATTGAAGCCGAGTTAACCATGACGGTAGCGAACGCCCAACCTAC

TAACCGACCTACCAGCAAACCACCCGGACGCTTCTTCGATCACAATCCAACTAATTCCAAAACAACGAAC

AAACCATCAACATATATTTCAAGAGAAACGCAACACATAAAGCTGTCGTCCAACGAAGCAGCCGGCAAGG

AGCCAACTCACACGCAGCAACTCTTGACAACACCTGCACATGTGCCAAAACCAACTGCTTGTGTTCAAGT

GTTGCTCAAAACCACACAAGCCGGGCACGGGTGAAATTCATCTACTGTATAGTAATCATACATAGCCGCA

CAACGCTCTTTAAACGCGGTGCGTCTAGCCACAGACTTGAAGCCTGTCGCTAAGCACCACGCTCCACCAG

GTATACTTGATGTCGGAGCCTGTCTTCACAATCCGGTT

>NODE_236_length_458_cov_3.670481

TCGCGAGACTGGATCGTTCGGCCAGGCCATAAGTTCTACCAGGTACCAGGTACCCGGTACCTACTGCTGT

ACGTACACGACGACTGTTTTCAACTGTTTTCGCCCTTTTTTTTGTTTGTTGGTTTTTTGTTCCTTGGTTT

GACTTCCGCCTCGTGTATCGTGTTACAGCAGTACGACGACAGCTACAGCAGCAGTGCGGCACAAGCCAAG

CCTGTCGTTTGTTGGTTGAACCTTTCTCCCCTTCCTTGCCTTTCTTTTGTTTGATTTTCGAACGGAGAAC

CGACGAAAATCCCGTTAAATGTTACGTCCCGAGCTTTCCGCTTTCCTCCGACGATGCCGCTGCTGTCAGG

CTGTAACAGCTCTCTCTGCAGCAGTGTCGGGTGGCTGGAGTATGCCCTGCTTCTACTTGTTGTTCGAGAG

GTCGACTTGTCTCTCAATTCTGAGTCAGAGTGAGATTT

>NODE_237_length_457_cov_6.857798

AAGATTATAAAAATAATAAATATTACGTTTCCCTAAAAGATAGGGGTGCATTGTCTTTGAAATTCGAACA

TCCTCATTAAAAGGGCGAGGAACTAAATATCCAGAAGATCCAATTAAAAAAGAAAAAAGACGAGAATGAT

CTGTTCTAACCATTAAGTTTTTTTACCTTCCTTACACACTATTATTTCATTTTCATATAAAACCCCAGCC

CCTTTATATGAGTCAGGGTAACGATATCTTCTTATACTACTTGCAAAATTTTGTAAAACACCTAAATTTG

CAGAAATTAATGAAAATGTTTTTTTGCCCAAATTTACTGAAACATCTACAGGAATATCAACTATAACAGC

CTTACTGTAACCTAACGAAAATATAAGTTTTTCTTTTTCTTTGCGTACCCTGTAACCAATCCCTTTAAGT

TTTAGTTCTATGGTATACCCTAAAACAACCCCAAAAA

>NODE_238_length_457_cov_5.974771

ACTCCGTGCTCGACTGCGCCACCGACTTCGGCATCAACATCCGGCCGTCCTTCATGGAGAAGGACCAGGC

CAAGAAGATGGAAATGCGCGAGGACCTGGCGGCGAACTTCATCCCCAAGTGGCTTGCCAGCATGGAGAAG

CAGCTTGCGTCAACTGGAGGCATCTACTTCGGCGAGAAGATGACCGTGGCTGACATCATGATCGCCCTCC

GGGTGAAGGGGCTCAAGGGCGGCGTTCTCGACGGTGTCCCCACCACCATCGCCGACTCCTACCCCAACCT

CATCTCCCTCCACGACAGCATCGTGGGAGAGCCCAAGATCGCGGCCTTCATCGCCAAGCACGCAAAGTAG

AAACACACACAGCGAACGATTTTTCGGAAGGGCCGCCCGCCCGCCTGCCTGCCTGCCTGCCCTGAACTCT

ATTGTTTTCTTGAGCTCCTGGAATTCTGGGTGCCAAG

>NODE_239_length_457_cov_3.188073

CTAGTTCGATCATCGTTTTTTTAATAGTTTTTTAAGAAGGATTAACACATGCGTCATCGTCAAAGTGGTC

GCCAGTTAAACCGTAATAGCAGTCATCGCCAAGCGATGTTCCGCAATATGGCAAGCTCTTTAGTTAAGCA

CGGCGTTATCAAAACGACTGTAGCTAAAGCTAAAGAACTACGTCGCGTTGTAGAGCCTTTGATTACATTG

GCTAAAACCGACAGTGTTGCAAATCGTCGTTTAGCATTTGCTCGTACACAAGACAAAGAAGTAGTAGGTA

TTTTATTTAACGAACTTGGTGCACGTTACCAAGAACGTCCAGGTGGTTACACTCGCATTTTAAAATGTGG

TTTCCGTACTGGTGATAAAGCGCCTATGGCTTATGTTGAATTAGTAGACCGCCCAGCAGTTGAAGACGCT

GTATGGAATACTCGCCAGCCAAGCAACACCAGTCACA

>NODE_240_length_456_cov_13.482759

ACTGCTGTCTGCCTTTGTCTTGTTTCTCGCTTCCGAAGCGCAGGCCGTATAATATACGATGATATGTAGG

TACTGGTGTATCGCACTCTTTTGTCTTGAGGATGTAGTAGGTGGATGAGACCATGGGTTGCTTGGATGAA

TTGAGTCGTGCAGCGACTCCTTCTCCTCAATCCCTGCTGGTACAGCAGTAGTGCTGGGATCTGGCATATG

CCCTGCTGAACTAGGTTGATCGTATATACGTGGTGCACGAGATACGATGGGTGTAACGGCCGGGGGAGTA

CGTAGAGACACTTGAGCATGTCCATCGAGGGGATACTTCGGTGTGTATTCCCGGTGGCTGTGGACAGCAA

GAGCTGAGAGCAGTGTTTTATCGATAAATCGCTGGCTGAGGCAACTGCCGCTAAAAACTGTAGGCGTGGG

TGTAGGTGTGCGGAATGCTTGGAATTCTTCGGGTGC

>NODE_241_length_455_cov_5.668203

ACAGCTGAACCAAAACCATGTTATCTTGTGCTCTCTCGCAGCAGGCATCCAAAAACCCGGTAAAAATATC

CCCCGCCCATTCCGTTCCGAATGGTGTTAATAGTATCAGATCATAGTATACGGACAAACACAAGCAGGAC

CAAGAATTGAAAGCTTACACTTATTATGCAACGATGGGAGGGAGTCTTACAAAGCATACATAAAATCCAA

ATGAAACCAACTGACACACTATACGCAATAGTTTCATCCTTGTTTTTATCCGAATTCTTACAGTTAAGTC

TTAATCGAGCGCGGTGGGGCATGGGGATCTTCCCACGCCAAATAAGCGCGCTCCCGCGCCAGTTAAACGC

GCGCGGTAATCGGCAATTCGAAAATCTGGACTCGCATAAACAGGGACAGACACCGATAAATGACAGGAGC

CAGAACGATACAGGCTATCCCTATCACGATGCACG

>NODE_242_length_454_cov_2.801386

CATACAGTCGACGAGTGGCTTCGAAGAACGGTCACACGCGAAGGGATTCTTCGCGAACACGAAGAGCAGG

AACAGCCTCACCAGGGTTTCGGCTTCCGAGGATGACTCCGGTTCGCAGGTGGACCTCAAAGTGGACTCGG

CGTCGGACGAGTTTTCTTGCCAAAACGCCTATTTCTTCGCCGAGATGTGCGAAGCCGCTTACTCGAACAG

AGAGGAAGTGAAGACCATATTCCAAGGCAACGAGACCGTGACTGGTATGGGCTTCGAGCACTTCTACTGG

TATGAGGCGGATAAGGAGGTATCGGAGAAGTCTCTGTTCGATGCCATCCAGGACACCGAGGCCTTCGTGG

TGGCGAACGATGACGTCATCCTCGTGGTTTTCCGCGGCACTTCTGAGCTTACGGACTGGATCACCAATCT

CAGCTTCCGAACCAGGAATATCCCAGCGGCGTGG

>NODE_243_length_454_cov_2.321016

CACAGTCACGAAAAAGGCACAGATCACACCCACAGCAGGCAGCACAGCGCCGTGCTGTGAGCCGTGTTGC

TGTGCTTTCTCTTCCTTTCGAATATAGAGCTTTGTTGTCCTGGAGCATGGAGCTAATGTGCATCCCGAGC

ATGTTTTGCATGCATGTCAAGTCCGCCGTTTCTTCGCTGTGCACCATGAATCTCTTGGCATTTTCAAGTC

GGTCCTTGCACCACATCGTGGACCTCTTTTGCCCCTTTCCTTTCGCGAGCGTAGCGGGCGGAGCGGAGCT

TTGCTTTCTTCGCGATCTTTTTTCGTGTTGGAAGATAGATATTCACTTTGTCCTGTATTTGCCGACATAC

AGATGGAGCTCTTGGCATTATCAAGTCGCCAGTTGCAGCTAATTTATGTTCCTCTTTTACCCTTTTCACA

TTATTTGGTTATATTCTTCCTTCCGCGAGCGTAG

>NODE_244_length_453_cov_3.037037

CGCGTGATACGCGGGATTGATTGTGGGAGTACTAGGCTGTTGGTTTTGGGCGGTTGTTTGTTAAGTACGC

TTGCGTGCTGCATGGGATTGTGTGTGTACGGTAGTTGTGCTCAGTTTTGTTTTTGTCGCGGTGCCCTATT

ACCCAAGAACATCGCGAGGTATATTCGTGGGATTGCGTGTGTGCATTGCATTGATTTGTTGCGTTGGTTA

CATGTTAACCAGAGGACGCTCGCGTGCTACCTGGGAATGTCTGTGGGGATGCCCTCTGTTTTGCTTGGCG

CGCTGGCCTTACGCACCACCCTTGCGTCTTACGTGAGATTGGGTGTGTGGGTGTACTCGTTTTTGTTGTG

TCGTGCTGGCCTGTTGCTCAGGATGCTCATCATGTCCCACGTGGGATTTTGTGTGGCTAGTACTCTGTTT

CGTTTTGTCGCGATGGTTTTATGTTGGGGTTTT

>NODE_245_length_452_cov_48.937355

ATCGAGGGTCTTGCCGCTGGACGTGATTGTGCTGTGGTGGATGCCGTTGTCGAATACATCCAGCGTCCTG

GTGTTCTCGTCCCCCTTGTGAAAGGCGATGCTCATGCTGACAAGGTCCTGGGCTACCTCAAAGGAGTACG

TGATGCAGCAACCGTCGTCCTGATCAGCGGTGTCGAGGATTTCACCCTTGCACGACCAGCGAGAGTTGGC

CCAGACATCGTTGTCCCGGGTGTTGTAGGCTCCGCAGCCCTTCGGGGTGCAGCCATCCTCAGACGTCCGC

GTGTCGTACCCATTGGCGCTCACGTCCACCCAGGTTCTGTCGCCCGCCTCAACGAGGGCTGAGGTCGTGC

CAACGGTGGCAGCGCAGATAGTGGCGAGGATGGCAAAGAAGCGCATGGTGTAAGAGCCTTGGGTAGAAGT

ATGTAGTGCGAGGTAGCTCAGGTGGTTAGAGC

>NODE_246_length_451_cov_12.816279

CTGCTGCTGCTGCTGCTGCTGGGGCGGCATCTCCTCTCCTCGTGCTCTCTCCTCGTTCTCCTCTTCTCTC

TCGTATCTAACTAAATCTACAGCAACTTTCATCACGACCTCAGCTTGAAGCTCATGCTCGATGCGGAGAT

CCTGAAGTCTCTCCGACATCGCCTGGACTCGCCATGACCAAAATCGATTCAAGGCGGGATCCTCTGCATA

CGACGCTTGCAGAGCTGCCTCTGCGAGTCTGATCTCCTTCTCGAGCTCCAATGCCGCGATCTCTATCTTC

AGCAGCTGTATCCTCTCCATACCAGCGTCCTCGTGACCTGGCATGGGCTTGCGCAACCTGTTCCCCTCTG

GGATTCCCCTCTGGGATGCAGCGCTCTCCGATAGGATCACCGGTTGCCCGGACGCACTAGTGTGTTGAAT

GATCTCTAAAGGTGTACACGCATTTGGCACA

>NODE_247_length_451_cov_5.027907

AGCTCGTCGTGTTTTTGGCCAGTGGTGCGGTGACGCGTGGTAGTACGACGCCCGCCGTATGTTAAGGCCC

CAAGATGGGACGCGCCGCTTGATTCACGACGGGAATGCAGCACACACACACCGGAACATGCCTTCAAAAC

GGTGCTGATGATACTCGTAGATGTTGGTTTGCTTACTGTTTTTGTTGTGCTGTATCGAAATATGTTGGGG

TGTTTTTGTGTGGTTCATAACAACATACATTCGACGTGTTCGATAAGATGTAAAGCGGTGAGATCCACCA

CAGGGCGTGTGTCTGCTGTGAACGTTTGTCGAGGGTTAGTGTTTTACGATTGATAGGAGGCACAACAAGT

GATCTCCCCCCAGGCTTAGGGTTAGGCAGGCCCTTCTGTGTCACCCCCGTGTGATTGTTGGGCGCTGTAC

TTTAATATACATGTGGAATTCTGGGTGCCAA

>NODE_248_length_450_cov_5.804196

CTTTTGTGAGGATGGCTGCAGCACCGGGTTGCCGGCCGGTATGTAGGTTGGTCGGTCGGTCCGTACGTAC

GTATGTTACTGGTCGGTAGTAGGTTGGTCGGTGGCTGGGTAGGCAACTGTAAGTCCTTGCATTATGTAAG

CTGTGTTTAAGGTATCGTGAGGTTTCCTGTCTTGGGCGGTGTGCCGTGTTTAATGGCCGTCTCAAGTCTC

AAGCGGTGCGTTTGGTTTTTCGTTGTGTAAAAGGGGATCGGGTGGATAGTAGATGACCTGGCATATCATA

GCGGTCGATTGTAAGACCCGGCTTGGCTTCATTAAGACTTTCGAGAGCTCGGGAGACTAAAATATTAGCG

TGGATGTTAGTTAACGTTCAATTTTCGGATAGTGACGTCTGATTTTCCGGCGCTCAAAAGTCCGAAACGC

GGACATTTTATGCCAAAGGGGGGGGGTTGG

>NODE_249_length_449_cov_6.535047

ATAATAGTGCACCAGAGCTACAAAATGCGAACATGACACGAACGTACAAGGACGATGTTATATTTGACGT

CCCTTTGATAGTATTATTAACCGCTGGTCGCTAACATACAAAACAGGAAGCAACAACAACAACAAAACTA

TGCCACCAAAAAGTAAAGCCACTTCTGGAAAACGACGACGATAACAGCACCCACCACCACCCTTAAGAAC

TAGCTTGACAGCACAACCAACCAGATAAAAGAGTCAGCGCTACTACTATAGGCGAAGGGTGCGTGAGAAA

CCACCAGGCGAGCCCCAAACCCACCGTCAGTCTGGTCCATTCCCTTCGTCTGCGAGTGTCTCAAAAAAGC

GAGCACTTCGATAATCTTTTCGAGTCAACACGTTTTTTTTAATATCGTCGTTGTTCCATCCGCGCCTCCA

TCACCCAACCCCCCCCCCCTCTCCCCCGA

>NODE_250_length_449_cov_5.764019

ACAAGATCCTTTGGAGGAACCACCGAAGGACACCAGCAGCCGTTCAAGCGGATCAGGGAAAGAGGCCAAG

GAGTCCCAGCGGTCGGAGCCGCCCATGCTGGAAAAGGTGACAGCTATTCTCCAAGATACGCCACAACTGA

TGCAGAGGCAGTGCGAAGATATGGTACTTGGCAGGATTCGGAAAGAACTCCAGGAAGTGGGAAACCGAGG

AGCAGGAGGTGGCGAATATGTGATGGACGACAATGGCCTCTTGTGGGTAGCTCCTCTGGGAAAGGTCCCA

CGACTTGCAGTTCCCCAATCTCTCGTTCCAGGGATTATGGCGCTGGCGCACAGCACCTATGGGCATCCAG

GGACGGCGAGAACTACAGCCTTGGTTAGTCGTCGGTATTGCTGGCCTACCCTCATCAGGGACGTGAGAGA

CTACGTCCTCTCGTGTGGGTGCAGGCGTC

>NODE_251_length_447_cov_2930.361502

ATTAAGAGGCTCGCCTCTTTATCTAAATCTATACCCGAACTCTTAACGAACTCGAGCGACCACTGGCCCG

AATACCGCCGAGAAGATGACACTGTGAAGGACCAAAACAGTCCCCCCTATCGTAGCCTCGATCCAAAACA

ATAATTTACCCGACATTGCCAGAGGGTCTGTTCGGTCAAATTATAACCCAACCGAGTTCCACATGGGCTG

GGAGTTGGAACGCGGCTGGTTTTACGTGGGCGCTGTTGGTTGATTTCCCCATCACAGCTCGATGTGGAGC

GACTGGGTGGCGTGTTCCTCGACGATGTCTTCAAACGAGTCGAACGCGTCAACTCGAATCGGGTCACAGA

AGTCGTCGCCGGTGAAAGCTCCCTTGCAGAGCTGGCCGTCCAGCCTGTACAAGTTGGCGCCGAGAAGCTG

CAGCCGCTTGCCGCTGTAAAGGCGGAA

>NODE_252_length_447_cov_37.436620

CATCGTCAAGCACAGTGCCACGTTCTCCTTGCACGCCGAGCTGTGAACAACTTCACCGACCCGAGGCCCT

CCCCACAAAGGCAGTTGCCTTCTATAGTGCTGTATGTTTCCGACGTATCCCGTTGGACCTTGCAGGATGC

TGGGCAATAAGACGCCCTCGGGCTCACCTAGCGTAGGGGGATGCGTCCACTGACGTGGGGGCGAGTTGGG

CACTATAGAGGCAACAAGAAATTGCGCCGATATGACTTGTATATCCCCGTCACGGGGGGAATTTCTGTTC

GTATCATCTCAGTCGTTATCTAAAATCTAAAACAGGGTTCGATATGTTTGTGCTTATATTTGAATGTAGT

TGCACGCTGCTGTATTATGTTTCTTCCGCCTCGTTGTAGCACGTCACGGTAGGCTGACGAGTGAGGGAGA

AGCTGTACATGTGTTCATTAGCGCACA

>NODE_253_length_447_cov_30.833333

TCACGGACACAAAAAGCCAACAAGAATAATAATATAAAGGTATACATATATGTAGTCAACTATAATAATA

GTTATTATTATTATATACTTTAGTAGTAACAAATAGATTGTTATTATGTTTTGATTATGATCTGAAAATA

AATGCAACGAGGAGTATTACTCTAATGGCTGTACCTAAAAAAAGGCGCTCAAAAGCAAAGGGAAGAACTA

GGTTATCAAATTGGAAGTCTAAAGGACGTACAGCAGCTGATAAAGCTTTAAATCTGGCAAAATCTATAGT

TAAAAAACGTTCTACGTTTCTATTCGACCCTAAAAAGAAAAAAGTTGAAGAGAATTCTAACCAAGATGAA

AATAAGAACGTTAAAATGGAACAAACTGCAGAAGATAATATTACTGAAGAATAATTTATCAGTAAAGATA

AAGAATAATTATTTTATTAGATTATAA

>NODE_254_length_446_cov_10.305882

GGCTGCCGCGAGGACCCTCTCCTTGTAATCCTCTTTCTGAGACTCTGAGTCCGAGAGCCTGTAGCCTTGG

CCGAACGAGTTAAGCACCTGGAAGCCGAGGTCGTTGTCGTCCAGCAATTCGACGCCCTCCAAGTTCGTGT

TGAACCTCTCGCCCATCGTCTTCCACATCTCATCTCCGGTTGCCTCGTACTGCAGCCAGAACATTCCAGC

GACCATACCACTCGTCCATACGGTGAGGTCGATGTCGTTCCCGTTGCTCTTGACGGTGTTCCACTCTCCG

TCGGTGTTGGTGTAGTCCGTGAAGGCACCGTCTTCTGCGAGCACATCGTTCCCCAGCACCTCCAGCTGAG

CGCCGGAGAACACGAATGCGCTGGCCACATCCACGTTCAGGACGTCGTCCGCGAAGGCTGTCGGGACGAC

GCCGACAAGAGCACACGCCAAGCCCG

>NODE_255_length_445_cov_11.634434

TTTTGTGACAGCTCGGTGTACGTTACGGGCGATATATATAAATATATATATATTTACAGTACGTAGGCGT

TGTTCTGCTCTTGTTTGATTGGTTGACTGTCTGGTCTCGCTGGCGATAAACTACCCGGTACATTTATACT

CTACGCTATGGACTAGACAGCCATACAGTCTGTCAGACGGCAGCCTGCTAATTTTATTTTTGTTGGTGGT

GGTGGTGTTTTTTTTTTTGTGAGAGAGGCGTTGGGTGTTTCAGCTCGCAGCCGGTTCCGTGCGCAGCTTT

TTCTGATTTTGTTGGGATCGGTTGTGTCTGCGGAATGGCGGGAGGGGAGATTGGTCTGCGGACGTTCCCC

CATTATAATGGAATGTTGAGCGCTGGTTTTAGTTCAGTCCGAAGATTTTCTGTTGGAAGACTTAATTTGA

CATGCTAACATCCTCAAAAAAAAAA

>NODE_256_length_445_cov_5.273585

CTTCTCCGTTTTCTACTACGAAATCCTCAACTCCCCAGAACGCGCCTGCAACTTGGCGAAGGCGGCGTTC

GACGACGCCATCGCGGAGCTCGACACACTCTCCGAGGAGTCCTACAAAGACTCCACCCTCATCATGCAGC

TGCTGCGCGACAACCTTACCCTGTGGACATCGGACCAGGGAGAGGCAGAAGAGGCCCCCGGAAACGCAGA

TGGGACTGTCGTGGAAGACCTCTAAAACCGAATTATCCTTTTTTGGGGGGGAGGGGGTATCCCGGGCGTA

TTATGTGCTTTGGTTGACGCGAGCAGCAAAAGAAGAGAAGAAACACACCCCATTTTACTATTTTTCGTGC

TGTGTTACCCAAATGCCGTTTTCTAGCGCTTGGCTGGTGTAGTAGTTGCATCTCTTTTGGGGAGTGGAGT

TTGGTTTTGCTGCTACACCCCCTTT

>NODE_257_length_445_cov_3.044811

GACGCGGTTTTGGGCGTTGTCTTGTGTGCGAGAGCTGCGTGGTATGTCAAGGATGTGTTTTTTTTTGTTG

GTTGATTTGGTTGGTTTGTTTGTGTGTGGGACATACGGTAAAATGGGTAGATGCGTGTGCCATAGCGGCT

GGGAGAGAGGCCGACAACAATTTTGATGAGCTTTTTGTTTTTGTTTTTGTTCCACGCTGCTGCTAATGTA

TAGCTGATGCATCGCCACACTGTTGTACGGGTCGAGACGGCGTGCGGAAAAAAGGGGGGTTCGGACACGA

AGGCGCCGACTTCAGCGACCCTAGCATAATGTAGAAATGGTTGACTTGATTCTCAGAGGGAGATGGGCTG

GGGGGAGGGGGGACAAGAGAGGGTGTGATTTGTCAGTTTAAATAGTATCGGACCCTGGCTGTGAATTTTT

ACGTTTTGACGGGAGGCGCCACATG

>NODE_258_length_443_cov_53.187204

TTCATTTTTGGCTGATGTTTTTAGGTGTAAATTTGACTTTTTTTCCAATGCATTTTTTAGGGCTTGCTGG

TATGCCACGACGTATACCAGATTATCCAGATTCTTATGCTGGGTGGAATGGTTTAGCCTCTTTTGGGTCA

ATTTTATCATCTATTGCGTCATTATTTTTTTTCTTTGTTGTGTACATTACTCTTACACGAGGATCTGTTG

AAGAAGCAAACCCTTGGGTAAAAGGTAGAGGCCTTGCTTTTCCTGTATTACCTTGTAGATCTTCAAAAGT

GGGGAATAAATAAGTATTAGTTATTTTGTTAAAATTGGAAATTACGATTTAGATAAAAGTTATCGAGTGT

GTTAAGGTGGTTGTGTCAGGGCTTGTAACTCAGCGGTAGAGTGTACACCTGATAAGCGTAAAGTCGATCG

TTCAATTCGATCCAGGCCCACCA

>NODE_259_length_443_cov_2.976303

TAGTAGCAAAGTACTGCTACTAATCCCGGGATATTTTACTGTCGTTGCTCCCGCACGAAAATTGTCCAGG

ATCGTGAATGTCTATTTTTGTCGTAATCCACAGCTGGCCAGCCACAAAGTCCGCGATGTGTAATGAACAT

GTCGTCTGTGAAAGGATTACTCACTTGTCATCGCTTACTATTTTTCCTGTCCGATCCTATGTCCGGATTT

AACATCCGCTTGTCAACGGTACTACAGCAGAAGGTACGTCCTGCTGTCATGTCTGACATTTTTATATGTC

TGGCAAAGATATTTATCTTGCCTTGTCAAAATGAGTGTATAAAAAAACTCCAGGCGTTAAGCGAATCGCC

ACTGGAAGTATACAATGTCAACTAGACGTGTTGTAACCCTTTGTCTCTGGTTCCATTACTTTCTAGTTTT

GGTGGAGTGCGAGCTCTATCACC

>NODE_260_length_438_cov_67.647482

TGGTAACGGGAAAGGGATTCGAACCCTTGACATTTGGGATATGATTCCAACATTCTAACCACTGAACTAC

ACCGCCTTAAAAAAACCTACAATATCTCTCTATAATAATTTTTTTTAATTGCAACTGGAGAATTTTCTTA

CAGAGCAAATAGAATCAAGAAAGCCATCATTAAAGCAAATAAGGCAATCGCTTCGGTTAAAGCGAAACCT

AAAATAGCAATTCTAAATAACTCATCTTTCAGAGAAGGATTACGTGCGGTACCCAAAACAAGAGCACCAA

ATACGGTTCCAATACCCACACCTGCTCCAGCTAAACCAATAGTAGCTAAACCAGCACCCAAAAGTTTAGC

AGCTTGAACTAACATTTTTAAAAGGGTAAAAAATCGAATAACGGGACACTTTAACGATAAATAAATTTTT

CTAAAGATTGTCGTTGAA

>NODE_261_length_438_cov_20.110312

TTGTGCATTGAAGTTGAACCGGTCCGTTCCCTTATCATAGCCAAAGCCCAGCTGATAGTCCAATCCACCA

GCAAAACGCTGGACGCGTCTCCCACATAACGCCAAACCAAATTCAACACAGTCACCAAACTAGGAATGAC

AATAAAATCCCCACACGATGACACTAACGCCAAGATAACATACAGCCGTCCAAAGAAGCCCGGCAGGTAG

CCGACGACAGACAACAAGCCCAATCATCTGCAGAAATTATCGACTAATCTGGAGTAGACATTTAGCAATA

CGATATAGTTGTTCCATAGCACTAGCCGGACACGGTTGCACTTAATGTACCATGAGTCCCCAGAGCCACA

CAATGTCATTGAGAGGTGGTGCCTGTAGCTAATGGCTTGAAGAACTTTCTTCAAGCAGTACGCTCCACCT

AGGCATACTTCATTTAAC

>NODE_262_length_438_cov_2.717026

AAAGGCGAAATGAACTTTGACGTTGTTGTTGCATCACCAGACGCTATGCGTGTTGTTGGTCAATTAGGTC

AAATCTTAGGTCCACGTGGTCTAATGCCTAACCCGAAAGTTGGTACTGTTACTACAGCACAACTTGAAGA

AATCGTTAAGACAAAAGAATCTGATTTAACTGCTGGTTCACTTGAAGCTGCATTGCGTACCATTGCGGGT

TCTGCTCGTTCAATGGGTTTAGTGGTAGAGGACTAATCAATGGCTAAATTATCAAAACGCGCTCGTCTTA

TCCGTGAAAAAGTAGACGTATTAAAAGAATATGAAATCAGCGAAGCAATTACTTTATTAAAAGAATTTGC

TACTGCTAACTTCCGTGAAAGTCTAAACGTACTTCAGTACTTCATCTCAACGCATGGTGCACGTAAAGGT

TTGGCCGATACAGCACTT

>NODE_263_length_437_cov_15.963942

TTTCTCCCACACATTCAAAGCACGCTTCTCCCTACGCCGGCATGCCCGGCACCGTCTTCTTACGTCGCAG

GCTGGCATACATCCCAGTTCCAGCAGCGACCAACGGCAGAGACAATGGCAGAGGAGTGCTCGGTGCTGGT

GAAGAGTCTGCAGGAGTGCAAGGAGGAGGTGCTGGCTCTCTGCGCGGAGAAGCACTGCAACCCTATCCTT

GTCCGCTTGGCGTGGCACGATGCCGGCACTTTCAACAAGGCCAGCTCGGAGCCATGGCCGAGGTGCGGAG

GGGCCAACGGATCTATCCGGTTCGAGCCCGAGATCAACCATGGTGCCAACGCCGGCCTGTCGATCGCGCT

GAAGCTTCTGCAGCCTATCAAGGACAAGCACCCCGAGGTTGGGTGGGCAGACCTCATCCAGCTCGCCTCC

GCCGCGGCGATTGGAAT

>NODE_264_length_437_cov_4.283654

ATAAAAACGCACAGGAGAACAGGCAAAGCCAGCAAGCACCCTCAGGCGTGCTAGGCCGCCTTGGCTCACG

ATCCCCAACTTTCAACGCCCCTGCTGCCCCACGAACTTGCACCCTCCATCCATCGTGGAGCGCATCAGGA

GCACGACCATCAGGAGCGCCATCAGCAGTGCATCAGGAGCCCGTGAGTTCTCCATGAGGAGAGCATCCGG

GTGTTTATCGCTTGATAGCCATGAGGTTGCGGAACAAGGAGCGCACCGCAGGCGTGAGCTTGGTCTTGTT

GTAGAGGTCGCCAGCCTGTCGGACCGACTCCGCCCTCGTAGGGTACGGGTGGATCACGCCCGCCAAGCTT

CCCAGCCCGATCTTGCTCTGCATAGCGACCGACAGCTCGCTGATCATGTCGCCAGCACCCTCGCCAACAA

TGGTCGCCCCAATGATG

>NODE_265_length_436_cov_6.033735

ACAAGGGCGTATCTCGATGCGAGGAGTAGTCTCGGACGTAGTACGTAGACGACTTGCTCCAACTATTCGT

GACGCCTAGTTTGAACTGGAGGCAGTGTTTTCGAGAGCAGCCACTCGAAATCAGTGGGAGGCTCGTTTCC

TGGTATTGACTCGTGTTTTTGGGCGAATTGCTGGGGCTGTTCGATCGAAAAGACGGTTTGATAGGACGTT

TGGACCGTACCCTCAAATTTAGCGGCAAACATAACCGGAATTAGAGTATCATCGCGTTTGGGCAGCTGTG

GCAGGGTTAACCTTAGTGGTGAGAGTTGGCTGACTAGAGGTCTCCAGCTAGACGTTAGTTCGATGGGTGG

TTGCGGTTAGTTGTGGGGTTAAAAATGCGGGATCCTGCATCTTTTTGTTCCGATCTGGCTCTATTTTGTG

ACCCCTTGCGAGTGTT

>NODE_266_length_434_cov_3.631961

GCTAGAGCACGTTCCGCTCTCTTTTGAACTCTTAAGCCTTCGTTCGTAACCCGAAAGGGCTGCTGATGTC

GGGGCACGTTGAGCCATTATTCGTTGGGCCAATATTCTAGTCAAGACGAGATAAGTCTTTGTACTTCACA

ATTGGAGGACCTCATAAATTCGCAACGTAAGTTGCGTCTTGAATCAGAAGGTAGTAGGTATACAAAACAA

GCGGCTGCACAATAAGCTGCACGTGATTGCGGAACGGACTCGCAGAGGGACGGCATACACGTCTGCTATA

CACGTGTCTTTTTGTCTTGATGTTAGGTTAAATGCCTCTAGTTTGTTTGGCTGGCATTTACCTATAGCGG

CATTGATGTCGCGTTGGATGGTATTTGCCCCTGTAGGTTTGGATCGTTCAGCCGTTGTGGCTGGTGGGCG

GGTAGATGGCTACA

>NODE_267_length_434_cov_3.569007

CAGCAGTTTACAGTCCTGCACATAACAGACAGATCACGATCTCAATCGCCTAGCTCCTACATCTGCCGTT

CGTTACGGGTGTATTGTGCACAAGAGCTTACACCATACATATCGCGCGTGGAAGACATACATGTTTGTGC

TACAGCGTCAGCAGTACGTAGACTATGATAGATCAGGGATGCATCTGCCCTAAAAATACCTAGATCATGA

AACGGGGATATGATCGGTCCGGTTGCATGCAATTCTTGCAGACGCCTCCGAGGGTTTCCTATTCCAAAAC

CGTGCATTACGCCCACGGTGAGGATATTGTCGACGTTATCTTCAGGAACCAGGGGTAATGCAAGCGCCAG

TCCATAGAGGGAGGCACTAGAGATACACCTTTCCACGGAGCTACGTACGAGTACTACAGCGGTATTTTTT

TTTAACAAGTGACG

>NODE_268_length_434_cov_3.387409

ATGTAGCCGGGAGGTAATTGTTGTGCTTTTTTTGCGCACCAGAGACAAGGGACGGGGGAGAGTAGACAGG

AGTTGCGGAGTTGCGTTTGAGTCGATTGAGAGACACACTCGCCTGTGTTGTTGATCAGTTGGCGTTGAAT

TGATTTTTAGGTGTATCAAGAGTTCTTTTTGTAGGCGATCGAATACCACCATGGATTGAGTCGGCTTTTC

GAACCGTCGTTACCCCTGCCTGGATGTGGCGGAGAGCGTTGGGAGATGTTGGTCACACAACCTGACTGAC

TGATCGCGATTCGTATTTTGGTAGTGTAAGATGGATTCTTGCGCCTTTCTAGGCCGCCGGTGTTCGGCCT

TTTGCGACGACTGACTGCAGTTTTGTTTTGTTTCGGTTTTATTTCTTGTTTTTGTTTATGCATTGAATTA

AAGGCCTTTTTGAG

>NODE_269_length_433_cov_4.364078

AGCCACAACAGAAAAATCGTAAAAACCTAGCTAAGTTAAGTTTTGTTGGTACCAGCACGGCCAAGAGCAA

ACCATAGACGATGGGCACAAACGAGACTCAGGAACAAGTGAGACTCGTTTGTCTTCAGTAGTTGTAACCG

CAATCTAACACTCGGCCACGAAGAAGGGGAGAACCCACTATCGTGGAGATGCGCAATGCGGTTTCAACTT

ACAGGGCTAGATTACACGTACCCTAGTGTACACGATACATCAAACTCGATGCGTCGGCCAATCTACAAGA

ACAAACGGCAACAGACACAAGTCTCTCGGCACAGCTAACAAACGGCACCCGCCGATCCTTACACTATGCG

CCTTACGGTGGGACAAAAGCGCCACGGGTGGAGCCGTGGCATTACCGCTAGTACACAATAGCGGGGAAGG

TTCAGATCCCACT

>NODE_270_length_431_cov_11.900000

CACTTGTAGTCCTTGCACACGATGTCGTCGGCGTCGTCTACTAGTTCGTAGTACTTTGGGCAGTCATAGC

TAGAGCAGACCTTGTCGCAGCACTGGGACTCCTTGCACTTGCCCCACTTACACTTGACGTCGTTGGCGTT

ATCGATTAGAGAGTAGCCGGAAGAACACTTATAGGAGTCGCAGTAGACTTTCTTGCTGTACGAAGTGGTA

CGCCTCCCGCGGAGGCCCTGCTCGGTGTCTCCTTCAACTGAGGCATGCGTCCACTGACCCTTGGAAGATG

GTTGAGCGGCGCAAGCTGTGAAACCGATAGTCGCCAAGGCCGCGACGAAGGGAACAAACCGCACCATCTT

GTACGGGCCGTATGTCTTGATAGAAGTACAACAGCAGTAGTCCGGTGGTGGGCGCTTTTGGCAGATGAGA

TGTTGCAGAAA

>NODE_271_length_431_cov_6.956098

TTTTTTGTTAATATAACCGTGGCAGGGTTCAACAAGAAAAAGCACAACGCAACCCTTCACCGCTACAGCA

AAGACGACGGCAAAAGCTGAGCGGGCCGACGACCTCTCTCAAAACCAGCGAGGAAAAGCCAGCCCAGCCA

ACGGCCCGCCAACGACACCATATACCGTAGGAGGCTTCGGTTGCTGCTCAGAAGGAGGCAAGCGCCGACA

GAAAAAATATAACAAATATTTTTCCTGCCGACTAGAAAGAGAGAAAGGAGACGCGCGCCCTCCCCGCTAT

CTCTCACGCACGGGCAATGGTTACGCCGACAACAAACGACGACAGACAGCAAGGCCAATAGGGCTGGGCT

GCTGCTTCCACATCGTCTTCGCCTACACCCCCCGCTGCCGCGGATCGGACTTCAGAGTTGCCGTCCGCCA

CTAAAGTTTTA

>NODE_272_length_431_cov_3.139024

GCCACCCTAACGCTCGCCAGGCGCCCATTCCACCTTAAGAACCCTCCGGGCACAACAAGGCCACGAGGGG

GCGGTCGTAGTTCGGGAGGTCTTTGGCGCACTGCCTGATAGCTTCTGCCAGGATCTTGTGGACCTCCGTG

GTCAGATGGAGGGAGTCCCAGAATATGTACGTCTCCGGTGTATCACAGACCGTGGCGGACGGGTTTCCCT

GGAAGCAGTTCTGACAGGGCGAAGTCAGATCGGCGCAGAGGGCGCACTCCTCCTGGCACCCGGCAGCGCG

TTCGGCCTGTGTCCCCATAATCTCAGCCGTTGAATCTATGGCAAAGTCAAAGGTCAGACAAGGTACCGTC

AAGGTCGTAAAGCCGAGGCTCTGGAAGATTTCAGGCTGTGTGAACAGGGCCAATGTCAAGTTGAAGAAGC

TGAGTACCCGG

>NODE_273_length_430_cov_7.518337

CTTGGGTCCCGAGATTTCCAGTGTAGGGGGTCGGGTAATAATTCCTCAGTAACCCCTCAAAACACCCAGC

AGCTCCCCTTCACCCCCCCCCTCCACTAAAAAAAGAGAAAAAGAGGACGGAGGGGAAACAGCAGCGCCCG

AATGCAGACGCCATCGCTAAATGGCTACACTATATGCCCCGGTCAAAGGTGATAGATAGCCTTAGTTGTT

CTGCGCCTTCTTAGCCTTGGCGGCCTTCTCCTCGAGGATGGTGAGCTTCACGCCCTTGCCCTTGGGGAGA

GACACCATGGCCCTGGTTCCGTGGCCCATGGAGAAGATGTTCTGCAGCCGGGTGGCGAACACGTGTCCAG

CTCCGTCCTTCACCGATACGATGTCGAACGATCCGGGGTGCCTCTCGATGTGCTGCATGATGCCCACGCG

GCCGGCGTTG

>NODE_274_length_429_cov_4.566176

TAAATGAATGAAAATATTCAAAAAAAATAACTATAGAAAAATAAAAACATTAAAACAAAAATGTTCGGGG

GGGGAGGATGGTGTGCGTGCGACCAGGAGGACTCGGTCTTGATGACGGGACCCCACGATTTTCTGAAGCA

GTAGGTAGCCCCTTTTAAGGGACCCTATTTATAGCGCGTTCGGGGTTTTGGAGGTCCTTATATACCGACG

TTGGTACCGGTCCCTTATGATCAGTAGTGGGCTAAAACGGTGCGCATAGTTTTGAGGTCCCTAATATACC

GTTTGCCCTTATCGCGGGCTCCCTTAAAAAGGGCTACCTCCTGTATTGGTGTTTTGATTGACGGCAACAC

TTAAGAAGCTGGGAGTTTCGAATTTTGGCTCAGCTTTTGCACAAAACGAAAAATCTATTGAAAACATAAA

AAAAAAATT

>NODE_275_length_428_cov_2445.813268

CGCTCAGCGAATGTTGTCAACACTTGATGTTCAGGCTCATAACCGTTTCGTTGCCATCATCACGAACGAC

CAGAACAACCAGCAGAACGATCTGCAAGGCGGTCACCATCGACTACAACCTGGGTGTCGTCCATACAAAC

AACTGACGAAACTCAATAGATAATCAGTCGCACACCTCGTGGGTGGGTGACAAATATCGAAAGGCCAACA

AAAACGTAGCGCAGATCCAGTAACCGCCTAACCGCAAATCATGCCGATGAGGAGTTCCAGCTCTCTAGCA

ACCCCGTTTGTCCAAAAGGACTAACATTATAAGTGTACATACAGTATCCGAATTAGCCCCAACAGAGGAC

CCGCGCGTGTTTCTCCGCGCCATACATTTGAGTATGTATTGATACCGAGCAAACGCTATTGGGGAGCTCA

CCGCCGTT

>NODE_276_length_428_cov_9.700246

GAAGGGGTGGGGACAAGAGTGAATTATCCTCGTCCTCCTCGGATAAGCGCAACAGGGGGGGGGGCGGCGT

ACAGACAGTATGGCAAAAGTAGAAGCCGAATGAACACTTCAAAATAAAACCGTTCCTGACACAATATGTA

TCCATCTCCGGGTCCTCATCCCCATTTTTTTGGGTCTCTATGCTACAAATACTCTCTCCTCCTCTCGCTG

GCCGTGCTTCGACATGTTGCCGTACTTCATTCAACGTGTGACCGTACCTTAATGTGTGGCCGTACACCAA

AAACACATGCTAGGGAGACCGCGCAGTTTTTGACCGCCCTTGACTCCGTGCCTCAACAATGTCTGACCGC

GTACTTTAACGAACGATGACCGACTTCAATCAATGCCCGACAGTATGTACTTTAACGTGCTGCCGTACTT

CAATGTCC

>NODE_277_length_428_cov_3.975430

GTAGCCCACCATACCTTTGGCGACACGAATCGCCTCTCTGGAGCATTGTGCGGCTCTCGTGTGACTACTG

GTGGGATCAACATCCTCCGTGTCCTCGACTGGTGTTTTTTTGCACACGATGATGTCGTATTGGTGCACGT

CTACCGTAGATTTGGCGAGCGTTTGCTGCATTCGAATCAGCCCGCTGTCTGTTGCCGGCTCCTTGCTGGG

CGCTTCGTTGGACGACAGCTGCCCATGTTGGTGTGTTTGCTTGGACGGGTATAGTGTTGGTTTGGCTGTT

GTGGTAAAATTGACTTGGGTTCCGGTGGGAGATGCGTCCGGTGGCTTGCTGGTAGGTTGGTTAGTTGGGT

AAAATGTCTCTATCATCATGAGTCGGTCGGTTTCAATCTAAATACACGAACCCTTTTAAGAACCTCAAAA

AAAAAATT

>NODE_278_length_428_cov_3.442260

CGTGTTTCAAGCAAGCAGCCTGTCTTGTTGTGGTAGTCGTGTGGTCGTGTAGCCTTCTTTTTTCGTGTGT

CGTTCGTTCGTTGCTCGGCCTTATGGCCTGCGCGGAGGGGGGAGGGGAGGCAGATCTCAATTTGTCGCCG

GCTCGTATATACAGTAGTAGTAGTATTTCTTCTTGTTTTGAATCCAGAGATCTTCTTGATTGTTTCTTTT

CATATTTGTTACATAGCACTGCGGTAACTAAGACCTCCGACAGGTTCTTGTCTCTCGGTCCAAAGCATGC

CATATAATTGCAGGCAGCGTTGGAAGAAGGGCGTAGTCGTTCAAGTCACCAAGGAAGGGGAAAATCGCGG

GGTTCCGAGACCCACCTGCCCACCTGTGGACACCATTAATAATGTATGCACCGTCGAACCAAAAAAAAAA

AAATTTTC

>NODE_279_length_426_cov_7.641975

TTCTGATTGGTCGGCGCTGATAGCGGTATGAACCATGTTTGGTGGTGCTGTTCTGATTGGTCGGCGCTGA

TCGCTAGCTAGCAGAGTGACCCTTGGATGGCGTTGTGACAGTCCTAAGCAAGCAACTCGACGGCTACATG

AAACAGTCGGCGGGAGGCGTTTCGCCGTTGGCGTTAACGTTGCACTCGCCTACTTCGTTTTTGGAGTCGG

GTAGTTTTTCAAGGAGTGCTGTTACCACGGTGTTGGTTTTTCGCGGAGCCCTGTAGGATCGTGTCGTGGG

TTGTGTTGGTGCTGTCGTTCCGGCTTCTTGCGGTGATTGCGTGGCTGCAGTAGTAGATGGACACCAACTG

CTTAAGAAATTTGATTTATTCGGGGGGAAAAATGTGTGAGTTTCTTTTTGCGCAAGCATGTTCGTGGTTC

ATGGTC

>NODE_280_length_426_cov_6.108642

ACACTCCGACAAGCATGCTCCTCCACACTAATTCCAAGTGAAAAAAGACATACAACTACAACAACTACAC

CCTACGACACCCTACACCCTACGCCAACAGCTTTCTAAATTACTTATATACAAGTCAGAGCGGCTTTCTC

ATTCTTCTCCCTCGCCTACAGACAAACAAAGAAAGCAAGCAGAGCTCTTCCTTCCCGCTTGCCCTCCTTC

TTTTCCACCGCTTGGCAATGCTCGTGCAAGATCCGGTCTACTGCTGTGGGCCTCCCTGCTGTACAATTGC

CTTACACAACACACGAAGAAAAAAACTTACAGCCGACACGGCAGAAAGAGCGAACGCATCACAAAGCCCA

CGTAAGACGGCGCACGCGGCAATCCAATCCCTCCTCCTGCTTTCCAACCCAACATGCTTCTCCCAAGCAA

ACCTTA

>NODE_281_length_426_cov_5.298765

CGGGTGGGAGACCGATAGCGATGGATGTCGAGGACAGCGAGATCTCCTTCCTCGGCAACGAAGGCGACTA

CCACAACGACATCAACAACGACTACGGCCTGGTGTGGAAGGTAGAGGGATACGACGACCGTTTCCCGGGA

GACCTGTCCCTCTTCTCCAGGATCGGCGTCTATGGCACACTGAAGCGCAACAAGATCCACGACATGTTCA

TCGGTGCATACTGCTACGGGATGAAAGGAAACACGCTCTGGACCGGTAACGAGATTTACGACAACTTCCT

GAACGGCATGAACCCTCAAGACAACAGCGACGGCATGACGATCTCCGACAACCACGTGCACGACAACGGA

TGGCACGGCATCGAATACAGCAAGCGTTGCAAAGGGGCGCTTGATTTGTTGTATTTGCGTTGTACTGGTT

TGCGCG

>NODE_282_length_426_cov_3.619753

AGTCAGACGTAACGGCCGGAATCGCAATAAACTCAACTCTCGAGCCTACAGGACCGCTATGTTACACACG

ATGCTGGGTGATAACTTGCCGACCCAAGGTTCCGCGGGCCAACAGCCAGCATGTTATATCTACTCGTAGT

ACTCTGTAGATGTAAACTACTGTACGTACTTTTCTGTGCACAAACAACGGGAGTCAAAACTACACTACTC

GTCATGAACTACATGACGATCTATCTAATATGTGATGATGCCGCACAAGGCCTGCCTCGCCTCGTGTCGA

CCAAAGTTTTCAAAGGAGAGCGTACGAGTATGACAGCAGAAACAGCAAAAGCAGTACAACAGCCTCACAC

ACCGAACACATCATTACGCTCGACGCCGCGAGCGCACGCACTCACTACTCACTACCCTTGCCCTCCCGTA

CCGAGA

>NODE_283_length_426_cov_3.128395

CAATTGGTTGGGTTACTTTCTATTGGCATTGGAAACATGTTACAATTTGGCAAGGTAATGCAGCTCAGTT

TAACGAGTCTTCTAACTATATTATGGGTTGGTTACGTGATTATTTATGGTTAAATTCATCTCCATTAATA

AACGGTTATAACCCTTATGGTATGAATAGTTTATCTGTATGGGCCTGGATGTTCTTATTTGGACATTTAA

TTTGGGCTACTGGGTTTATGTTCTTAATTTCATGGAGAGGGTACTGGCAAGAGCTTATAGAAACATTAGT

TTGGGCTCATGAGCGTACACCTCTTGCGAACTTAGTTCGTTGGCGTGACAAACCTGTTGCTCTATCAATT

GTTCAAGCAAGATTAGTTGGGTTAGTTCATTTTACAGTAGGTTATATTTTAACTTATGCTGCTTTTGTTA

TAGCTT

>NODE_284_length_425_cov_11.091584

GGAGGCAGTGTAGTGGAAAAGGTGGGAGTGGGGCGGCGGCGCTTGCTCCCCTCGAGAGGGAAATGGGCGC

TAGGAGGGGTGGTGTAGAACGAGGAACGAACACAAGACCCCACATATGGAGAGAGTGTCGTGGTTTCTTT

TTTGTTTGGTCCCCCTTTTTTCTGAATAGTGTTCGGAGTATTTTTCTTTTTTCGGCCCCATTTTTCTGAT

TTTGTCAATCTACACGTAGTCAGTGCTCTTTTTTCTTGAGAAGAAAAGGCATGTTTTTTTTTTCTTGTTT

TCTTCGTTTCGGGTGGAGGAAGAAGGGTGGAAGTGTGGAGGAGGGTGAGAAAAAAAGAGGATTCTCGATC

AGTCGGAGTAAATCCCCTGTTTTCTGCTAGGCCTCTTTGAGTGGAAAACTCCCCTCCCAACGAACACCAC

TCACA

>NODE_285_length_425_cov_10.353960

CAAGCCCGTCGAGTTCATGGCTATCACCCCGGAGATGGCCAGCGAGAACCTCCACAGCATCAGGCGCGTG

GCCCGCAACGAGTTCAAGTTGACTGGCATCCGCATCCGCCAGAAGGAGGAGAAGACGAAGAAGTAGAGAG

ACAAACCGATCACCCAAACGAAACGCCCCCTTACCTCTCTCTCGCCATCCACCAGGATGGCGGTGAGATG

GAAGTCGGGTTGTTGACGACGGAGGCTTGTTTTCTATCGGGGATGGAGGCTTGTTTTCTGTTTTGGGCGC

TTTGGTGTCCGTTGCGCGCTATTTTCGGCTTCTGCTCGTGCTGACTACGTGCGAGTAGCGACGAGCAATA

ACCTGCGGCGGGCCTACTACTCCTTTTCAGGGCACTTGAAAAAAAAATGATACTAAAAAAAAAAAAAATT

TTGGA

>NODE_286_length_425_cov_9.537129

CGCTCGAGGCGACATGGCGGAACTCGGAAACCCGGGAGGGAGGGGGTGTCTCATGGGTGGGGGGGACACC

TGAGACAGTCAGTTGAAAAAGTACTGATGTCGTGTGTAGGAAAGGGGGAAAGGGAAGCGGGCGAAGCTTA

GTCCTGAAGCTCCGCTGCGTCTGCTGTTGCTGCTGTAGGCCCGGTGGATGCGCAGCAGCACAACGCGGGG

TTTCGGGTTTCGGTTATCGCGCCTTGCGTTTCCAGTTACATATGTTGTTTGGAAGTGTTGTGACTGGTGT

GTGGGGGTGTGTCGAGCTCTCTGGCAGCTTACCCCCCGCCACGCGTTGGTGTGACTACCGATGGAGCCTG

CTGCTCTTTACGATTTCCTTTTTGGGCCTGTCTGGGTTGCTTTTCCGCAAGTACCGCTCGAGGCGACATG

GCGGA

>NODE_287_length_425_cov_4.569307

GTTGTTACTTGTGAATTGATTTTTGCAAATGTTAAGGTATGTTTTTTTTGAGTTGGTTGTTTCTGGATAA

TTTCTAGTTTGTAATGTATGTAACTTGTTCCTGCCGCTTCAGCCTCCACATTTTTTGTTTGTTTGATCGC

AGAGGGTATGGGGGAGGAGGGCGGATGTTCGAACGAGGGGCCGGTTGTGTGATTCGTTTACAATTATTTT

GTCCTCACGCTCGTGAGGATGAGATAATTTGCGAGAACGTTGCGGAGTGCTGTTGAAACTGACGAACAAC

GAACGGTGTTTCCTGAGCCTGCCTTTGGTGCCGAGTGCCTTAGGAGGGGGGGGGGGTAAGGCATTGTCCC

CCCTGTTTTTATGTTTTGTGTGGCGGCGTGGTTTTCTCGCCGAGAAGGATTTGTGTGGAGATCACGGGCG

TTTTT

>NODE_288_length_423_cov_19.910448

TTTTTTGTTTTGACTGGATTTGATTTGTAGGTGTGAATTGATTTTGTTGGTTCCCTTGATTTAATTTGTT

TTTTTTTGTTCAACATTTTTTTTTCGGAGCCAAGGAAGGAAAACAACACGGAACAATCACAGCATGCCGC

AGAAACCCACCGGGTTCACAGCACAACGAAGACCTTTGCCGAAGTCATAAGACGGGAATCTCCGAAAAAA

ATACACACGGGGACAGCCGCACGACGCACTCCTCCCTGAAAGGCAGGCAGGCAGGCAGCCTTTTTTTTTG

CTGTACGTAAAAAACCTGTTGTTGGACGTACACGGTGATTGATGACCATTGGCGCCTGTATTTGGTGGCT

TTGGGTTTGGTTCTGTATTGGTTTTGGGCTGCTTGGTTTGGGCTGGGTTGGCGTGGATGGAATTCTCGGT

GCC

>NODE_289_length_423_cov_9.427861

TATTGCGTTCGAAACACACCCCTTACCTCCACTGCACGAAGCGCGAGATTGATGTGTGCGAGCTTTCGGG

TGGCGTGTCGAATGCTGATTGCATGTTGGAAGGTGGCAAGGATTGGGTGCACGCAATACCGTGGTACTTG

GGGTGTTGGTGTGTCAGATAGATGGATGTTCGAACATGAGGGCACAAACGTTTTGGCACTGGTGACTGCC

CAGTTACCTATTGGTCAAATTTGTCCGAAGGAATTTTTTGATTGGTGGACAATCTATCGCTGGTAAAAAA

AAAAACGAAGGAAGATATTTTTCATCGGTGGAAAGAAAATACCCTGATCGACCGGTGTGCCGGGCTCAAC

AATTGCAGCGCCACACTCATTTTTGTTTCGCGTGTGCGTGTGTGTGCCGCTGAATAGCGATAGATGGGGG

GAG

>NODE_290_length_423_cov_8.134328

CTTGGCTCGCGAGAATTCCACGGGTTCCCTGGGTCTTGGTGGAGCGAGAGTAGCTCTTGCCGAGGATGGT

GCGGGTTGCGATGATCACCACGTGGCGCCCGCTGAACTTCTTCTCGAGCTCTCGCACCAAGCGCGTCTGG

ATCTTGTGGTAAGCCCTCAAGAGACGGTACGGTACGAAGATGATGATGGCCTTCTTGCCGCCCCCGACAT

CCACCTGCTTTGCGCCAGTGATGTAGAGGTCGCGGATTTCCGACTTGATGTCCGCCGCGGACATCTCAAG

GTTGAAGAGCTCCTGAGCTACCGTCTGCTCGAACTCATCGGCCTCCTGGCCCTCAGGCTTGACAATCTTG

GTTGCTCCCGTAAACATGGTGACCCAGCAATGCTCCCAATGCTCTCGCTCGGCTATATTCCTCACCCAGG

GTG

>NODE_291_length_422_cov_6.002494

GCTCCGTCTCCATTCCCTTGGACGCCACCCATACAACGGATACGTGGCACTTCCTACTCAGTTCCGTTTC

GTTTTCGTGCCACACACGAGAGTCCAGGAAAGAGAAGAGGAAATAACAAAAACAAAGTGAATGAACAAAC

AAACAGACAAAGGGAAAGAAAGAAAGACCACCAGAAATAACCAAGGAAAAACACGCAGTTCGCCATCCAC

CAAGCAAGCCAGGTAGTACTTCTTCCTCATCACAGGGATAAGAACGAAAGTAACAATGAAACTAAAAACG

AAACGAAAAACACAACAGGGCCGCGTTCGAAGCAGCTACATAGGTGCGCGGTACGGGTAGTCTAGACGGG

ACTTACTAGTCAGCCTGGTACCTTGCTATCCTGTCGGCTGTCTCCCTCCTGCTTGGAATTCTTCGGGTGC

CA

>NODE_292_length_421_cov_3352.190000

TATAGATTGAAGCGTTAGCGTAAGCGGGCGTGGACGAAGCAGAAGTGAGAATGTCGGCTTGAGTAGCGAA

AACATGGGTGAGAATCCGATGCCCTGAAAACCTAAGGTTTCCTCCGGAAGGCTCGTCCGCGGGGGGTAAG

TCAGGACCTAAGGCGAGGCTGAAAAGCGTAGTCGATGGACAATAGGTTAATATTCCTATACTATTCTTTA

TTGGCAACGAGGGACGAAGACAGCTAGACTAGCCAAATATTGGTTATTGGTTTAAGTGTACGAGGTGTTG

AGAAGTAGAGAAAATACTTTGAGCTGAGTCATGAATACGGAATAACGTGCGCGTTATATGAAGTAGTTGA

TGTTATACTTCCAAGAAAAGCTTGCACTGCCATAAATAAAGAATACCTGTACTCTAAACCGACACAGGTG

G

>NODE_293_length_419_cov_11.143216

GGAGTTCCTTGGCTCGCGAGATTTCCAGATCAGTCCCTGCGTGTTCTGGTAGTAGTGCCTCCACAGCGGA

CGGATCTTGTCCTGCCCTCCGACGTCCCAAACTGTGAAGCTAATGTTCTTGTATTCTACAGTCTCCACGT

TGAACCCGATGGTGGGAATCGTCGTCACCACCTCCCCAAGCTTGAGCTTGTAAAGGATAGTGGTTTTACC

GGCCGCGTCCAGACCCACCATGAGGATACGCATCTCCTTTTTCCCGAACATACGCTCCCACAGTCTCGAG

AATGCGAGCCCCATGGTTCCGCCTACCCCCACCAGACAAGGACTCCTGTCTCGTGTGCACTGCTCGGGCT

CCTGTTCGACTTGGCAGGGGCTCGGGAGCGCGGATGTTCTCAGTTTTCGGTCGGTCTCACGTGTGCTCT

>NODE_294_length_419_cov_6.487437

ACAGGTAGTGCGGCGATATCCGGTGGCTGAAGCATCATGCCGCGAGTGCTGGTTCGCCATATCGAAGAGC

CTAAGATCTACTCCTGCGCGACGTGCGAGACTCACCTTTGTACTGAGGGAAGGGTCCATTCCAGGGAGTT

CCACACCCAGAATGGGAGGGGATACCTCGTTGAGAACGTTATCAACGTTACCAACGGCCCTTCGAATGAG

AGAACATTGAGGACAGGTCTGCACACGTGCGAGGACGTCTACTGCGTGAAATGCAGCACCATGGTCGGAT

GGGAATACAAGTGGGCGTTCGAGGAAGAACAAAAGTACAAGGTTGGAAAGTTTGTGCTGGAGGAAGTCAA

GATCGCCAAGTACCACTGGGAATCCGACTGAAGCGGAGGATAGGGTCGCAAACAAGTTGCGTGCACCAT

>NODE_295_length_419_cov_2.670854

AAAAATTCGGCACTCGGTGGAACCCAGTAGAATAACTGGTACCCAGATTCACAACATCAGATATTCATAA

GAAGCGATCCAGCTCCATCGGTGTCACCAAGCCTCTTACATAATATAATATTTTGGAAGCACACCTACGG

TAACATGTCTTGAGAAGTATTTGTCCAAGACAACACAATGTAAGATGCTAGATGACAAATGTACTTTCTG

TCCGGACTCCTTGAAACTGCCACCGTTAGCAAGTGGGGGTGTAGGCGGCGTCCGAACCCGCCATGAAACA

AGACCCGTCGGTGACCTTCAGTTGGGCACCGTCCAGGCCGGTGTTGCCCGACATGAGGATGTCCTGGCAG

TCTTCGAAGCGGACAATGTCGGCGTTCACGAAAGTGTTGTCTAGGAACTCGGTGCCGTCGGCCGTGGTG

>NODE_296_length_418_cov_4.987406

TGAGCCAGATGTTTGCGTGTTTTGATATTGGAGTTTAGCGTTGGTAATTGTAACCAGTTCCAGTCATTTG

TACTCGTTTCTTACCACCCTCGCTGTCTCCTGCCCTCGTTCTGTTTTCAGTTAACCCACGCTACCAGTAC

TACTACTGCTGTGTACGGAAAATGTAAAAGCTACAGTTCGGTGTTTGTTTGGGTTTGGGAATTATTGAAA

TAAGTACTGCTGTTTGTGAGACGAAGCAGGAAATATATCGGAACTAAGGCTTCCCGTTTCATTTTCATAG

CACCTTAAGTGCCTGCGGATTAAGGTTTCTCGCCCGTCGAATAACGACGTAGTACCTGTGGACTTGAGTG

CCTGCGGATTGAGGTTTCCCGCCCGTCGAGTAACGACGTAGTACCTTTGGACTTGAGTGCCTGCGGAT

>NODE_297_length_418_cov_4.652393

CCTTGGCACCCGAAATTCCACGCCCACCAGGTACGTCTTCTCGAGCAGAGCGCTCTCCAGAGCACCCAGA

GCAGTCTTCAAGTCCTCCTTCGCCTTCGTCACCGCGGCATGGTTGTAAGGCATGTAGCCGATCACCGGGT

AGCACCACACGGTGGCGGGCAGCTCAATCTCGTGAGCGCAAAAATCGATCCAAGAGTCCACCTGGGCAGA

CTCGAAGAAGGAAACCCCGCACAGCTCCGAATCGCGTCGCATGCGTGCCACGTAGCGCGCGATAGCATTC

GACTCAAAAATAGAGCCCTGAGGTGTCTCCAGGACAGGGACGCGTCCGAGAGGCGACTTAGCCACAAATT

CCGGAGACTTGCTGTCCGTAGGGTGGTTAAACTCGGGCACCTCCAGCTCTACGCCATTGTACTCGGCG

>NODE_298_length_417_cov_4.681818

CGGGTGCGTTCGGACAGATTTTGAAAAATTTCTGTGCGTGCTTTGTTTCTTAAGGTCTCTTTTTTTTTTC

CAGCACTTGTTCTGGATTTGTCTCTCTTTTTTTCTTCTCTCGCTTAGCCCCTCCGTTCTCTAGGTGTGTC

GATCTTGTTTCGGAGCGGTTGCCAGCGTCGAGTGGCACGACAAAACGTCGGTTTTGTCGGATCTTGAGAA

GCCTTGGGCTACACGTCGTTTTTTCCCGTCCCAACTCGGGCGCCGTCCTGTTCTCTACGAAAACCTGTAT

GTTGCTTTTGCCGCTTGCTGGGTGCATCGCCGCCACACACGAGCAAACATCACGTACGTACGTTCGTTCG

TTGTCGGACAATATCATGTTTTTCGCCACGCTGTGTGTTGTGTTTCTTGTCACGAGATTTTTACTAT

>NODE_299_length_417_cov_4.171717

CCGTAGCAGTAGTAGTTTTTTTCTAAATCTGTGTGTTTTTTTTCGTGGTGGTCGTCGTGGAGTCGTCGTT

TTTTTTGTCGCGGGGTGAATTTTGAACGTGCGGATTATGTTCGGGTATCTTGGACTAAGATCATCGGTTC

CTGCTTCATGGTCTAGGTCTCGCGGAGCAAGATCGGTTTCCTGATCTGTATGATCTTGCTCATGGTGGCG

GGTGGGGGTTGACGCTTTTGTTTTGTCGTGGGGTGAATTTGAACGTGCGGCTTGTGTTCGGGGACGATCT

GAAACCTCGCACAGATCATCGGTTCGCCTGCGTCATGATCTAGGTCTTCCAGAGCAAGATCGGTTCCTGA

TATGTATGATCTATACGTGGCTCATGTTGCCTGATCGGGAGCAGTTATACTCTACATGGTCTAGGAC

>NODE_300_length_416_cov_4.873418

TTTTTTTCATCTTTTTTACTTTTTGCCCCCGCCCATCATGTACGCATATGCGTCGCAGCATCGCTCCGAC

AAATTATCGGAACGCGCGGACTTGAAGTTCAACTTGAAGTTAAAGTTGCGCGATTCGCTGCCCGCCTCTC

CGCAGCCCCCCTCACGCTACAGCAACAACGGGAAAAGGCAAGGCAAGGCAGACATAAGAGTCACCCCCAC

CCCACCCCGCCCTGGATGCCCCGACAAAACATGAGCAGAGTCCCGTATAGGCAAGCACCGTGCCGTTAGT

TAGCCGCAGGCGACAACGTATGGTCCTGTCCTGTCCTCTGCTGTCGCATCACGACGTCCTTAGCTTAGCG

CTTGGCGCCCGAGCGGCCCGTGGTGCCTCCGCCCTTCTGCTTGGACGCGGTCGGCTTGGCCTTCAA

>NODE_301_length_416_cov_4.096203

GCGGCCGGTGAGCGAGCTTTCGACGCCGAACTCGTACGCTGGAGCGTGTACGGACGAGTATTCGGCCATC

TGGAGGCAGGCGATGGAGAAGGAGTATCGTGGGCTGGCGGACGCGGGGACGTTTGGGGCGGAGTAGCAGC

CAAGCGGACTGAACGTCATCTCGGCGAAGTGGGTGTTCGCCTGGAAGTCCAACGAGCACGGCGAGGTGGT

TCGGGCGAAGGCGAGGCTGGTGGCACGAGGTTTTAAGCAGCGTGAGGGTATTGATTTCTTTGAGACTTTT

GCCCCCACCCCTGCGGCGTCTTGTTTTCGTTTGTTGGGTGCTATTGCGTGCGAGTTGGGTTTGGATTTGT

GTCATTTTGATGCTGAGCAAGCGTTCGTTCAGTCGAGTTTGGAAGAGGATGTCTTTATGAGACTGC

>NODE_302_length_415_cov_48.469543

GGTCTCCATCGGCGGCGGCGGCGGCAGCAGGAGCAAAAGCAGCCAAAACAACAGGTGCAGAGGGCGTTGA

GGCAGGTAGAAGACCTCTCGACTTAAAACCAATATGGACGGTTTCACGACGTCGGTTTGGGTGAAGGGAT

ACAACGCGAATGGCATCACCTGCTGGCGCCACAAAGTCACGGGAGTTTTGGTATACACAGACCCGGAGCG

GGCAGGGCGAGGACTGGCAGACGGCCCCAGCCCCAACGCCCAACAGACTAGCTGGAACTCCACCCAACCT

GGGGGGTCGGATACGGCATGGGAACAATACACGGATCCAAAGACGAAAAACAAATACTGGCACAACAAGG

TCACGAATGAGAGCACGTGGAAGGACCCGAAGCCCGCGACGAACACGCATGGAATTCTCGGTGCC

>NODE_303_length_415_cov_5.644670

CTGACTGCACCAGCCGGAAGGGTGGCCGCCGTGGTCGCCGTCTCTAAGACGTTGCATCACCACAGCAGCC

TTATACGCACCGGACCTCTCCAGGACGTTTTTTTCTGCCACGGACTACTTTGCCTTTTTTCGCGCGTGCG

CTGCTGGTACTGGGGGCTTCTTGCTTTTGCATGCAGCAACGGTTCGACCGCGTTTTTCGGATGAAAAGTG

TGCCGGCTCTGTTGCGTGTTTTATAGCAAGTGTCTGCGGCGGGCTCTTGGTGATTTGACATCATGAGCCA

CCCGGCAGCCTCGAGCAAGACTGATGAAAAATAAGATGGGTCGCCAACCGAGACAAAAAAAAAAGAGAAT

GTGCATATATAAAAAAAACAATTATCCGGTACGTACGTGTTCGAAGTCGGGCAAAGAAAAACACG

>NODE_304_length_413_cov_10.132653

GGCTCTCGTCGGAGGTCGGCTGGCCTCACGGGGAACTGATCAAGCGCCTCGAGGCCCAGCGCAAGATCAA

GTCTGAGGCGTACTACGTCGAGAAGAAGGGCGCGACGCAGATCAAGCAGAAGGCCGAGGCGGCGGCGGAC

TTGTCCGCTGTCACGCCTGTGCTCGAACAGTACGGCTTCTAAGTTGCTCAGCTCCAAGTCTTTCAGCCTT

CTTGCCTTTCGCGCCAGTAGTTGAGCAGGGCTACCATGTATGGTCGGTCCCTTTTCGAGTGAGCACATGA

TGTGCGCGTTTGGGAGACTACCGTAGTGGGTGTGGTAGGAGGACTGGCTGACTGCCTTGCGTTTTTTGGC

CGGGATCATGGTGGGACTGGCTGTCCGAACAAAGGTCGAGGAGATGAAAAGTCTTCTTTCCCC

>NODE_305_length_412_cov_19.815857

CACGCTTCCGTACATATCGCCTTCTTTGCGACAGCCAACCTTTTGTCATATATGGTATACACATAGGCGT

ATACTGGTATATACATACCAGGTATACTGTTTCTGAAGAGGTTTGGTTGAGCTCATTGAAACGAAAGCAA

GCTCAATCCACATCGCAGCCGTCAAAGGTGGTGACAGGTGGCAGAGCGCCGGCATCATATCGTGAATACC

GTCACGCTTTCGGTAGCAATATGTTCAAACTGTTCGAAATCATGCGCACCGATAAGGACCATGCGGCAAA

CAAGTCTGGGTACAAACATCTAGACTTGACTTCGTTCGAGGCTCCTGCATGATCATGGAAACGCCAGAAG

TAGTCGACATTATTCAACTGATTATCCTGCTGACCGCCCTTAACAGCAAGGCCGGGCGGGGG

>NODE_306_length_411_cov_25.797436

CGCTTTTTTTTTTATAGGAGATGTAGGTTTTAGTGTTGTTGGTGTGAATGGTTTTATTTGTTTGTGGTTA

ATTGTGTCATGCAAGGAGGCGGGGGGGGGTTGGGGCGGTGGGCATTGTATTCGTGACAGGTGGAAACTCG

AAAGCGGAAGGTTGCGCCGATTATTTTGGTGCTCCCTCAACCGACGCGTTTTTTTTTTAGTGTTGAGGAA

TAGTGCGCGAGAGTGTACTCTCGTCATGGCGAGGGATACATCTATCTCTGCCTTATTTCCCGTTGAACAT

AATGGTAAAACGTGGTTGGTGCTTTCGCTTGTTTGTTTTTAGCTTTTCCTTGGAGATGTTCGGTTACGGA

GCTCTTTTTTTTTTTTATTCATTTTTTTAATTTGGAATTAACGGGTGCCAAGGAACTCCAG

>NODE_307_length_411_cov_10.894872

CCACCAAACTCCGTTGTTGTCATGTCAGGTAGTAGTAGGTGGTGCTCGCATACTTCTACATCGACTAAAC

GAACTACGAGCCCTACAAAGTTAAGTCAGTCATGATACCCGCCCACAAAAACAACTACATCGCTAACCTA

CGAAAACGATGCGAACAGAAGTGCATGCATTGGGGTAGCCAGCCGCACTCTCCAATTCGCCGCAGCTCGA

AACAAAAGCGCAACTATCCCCAGAAACCACAAACTATCTAAAAAATCACACCTTGGGCGACTAATGTCGT

GCCCGTCATATATATATATTGTCGACCCCAAGTCTAACGAACAAGAGAAACACAAAAGAACACTGCCGTC

CGCGGCCCACTTTTTCAAACCTCAACCCTAGCCAAAAACAAAAAAAACGCCCCCCCCCCCC

>NODE_308_length_409_cov_29.677835

AGTTGTGACTGGAGTTTTTTTTATCACGATTATTGAATTAATTTGACTTTATTAGATAAACGAACTTTAA

CAACCTTGCGTCGGGCTGAATTAGGCTTTTTGGGGGAGCAAACAAATACTTTTAAACAAACACCTCTTTT

TTGGGGGCATCCACGAAGACCTACACTCTTCACTCTTGTTTGCTTTTTCCCGTGACTTTTGTTAAACCAT

TGGTTTATATTTGGCCTTGACATTACTAAGGAGGGGAGTTGAACCCCTATCCCGTTAAGGACTGGAACCT

AAACCCAGCGTGTATACCAATTCCACCACCTCAGCTTTTGGAGTCGAAGGACTCGAACCTCCGATCCCCG

TACCAAAAACGGGCGCCTTACCGGCTTGGCTAGACTCCATTTATCAAACACTAAAACTC

>NODE_309_length_408_cov_7.041344

CGGTTTTTTTTTTGAAGCAGATTATTGCATAATATATTTTGTTGTTTATGGTTTTTCTTTGCAGATGTTA

TTTCAAATGCTACTATGTTGTTTTTTTTTTCTTGTTGTTTTTATTTGTTTCTTTATTTATTTGTGTACTG

ATATTTAGTTTGGTTTCATAATTTTTTTTTTTAATGGTTTGTTAATGACCTTTATGCTATTTATTGCTTG

TCTTGTAGGTTGGTTACTATATAAATCTTTAACGTTATTTAACGAGGCAGATCATAAAGATGCTGTAGGC

TTTAACCATTCAACATTACTTGAAGTCGTTTGGACAATTATCCCAGCAGGAATATTAATGGTCATTAGTG

TACCATCATACAATTTATTGTATGCAATGGACGAGGTTGGTCCGCTGAGCTGGTTGTC

>NODE_310_length_407_cov_38.318653

TGGGAGTTTGGTGTTTGGATTACAGCAATCTGATTTTGATTGAATACGAGTATGGATGATGTTGGATTTT

TTCGAATTCTGAACTATGAATTGCTTGATTGAATGGTTGAAGGTTGTCTGAAGAATGTGTTGGCGTTTTC

CGAGGATACGTAAGCTATCATATGAAGTCGGTCGCCCCGATGGTCTAGAGGTAACCTCGAACGCTTCTGA

ACGTTGATGTCTGGAGTTCGAATCTCACGTGGGTCGGACTTTTGAGTTTATTTGCAAAAATAAATAAAAA

ACGGAATCAACTGCTGAGAGCGCCAATTAATAGTAGCGTGGGTAGGCACAATTCGATGCGAGTCGACGAG

GGAAGAAAGGGCTGAAGTCTTCTCGCGATAAAAATGCAAGGCAAGAACCGTAGTGGG

>NODE_311_length_404_cov_157.678851

ACTGGAGTTCCTTGGCACCCGGAATTCCAGCCCAGTGCGGCATCGTTGCCTTGGAGCCGTCCTTCAGCGT

AATCGCCAGCCAAGCGTCGGCGACTTTGGTGTTGAATTCCTTCAGTCGCGGGGTACCGACGGCCGAAAGC

ACTTCGATCCACAGGAAGACGTCTCCCTCTTCGCCGAAAGCCGGGCAGATGGGGGTGTCGCTGTGCTTAG

TAAAGCGCATCTCGAGGGCGACGTTCAGGGGCATGGTGTCCTTGTCCTCGGCGAAACTTTTGACGATGGC

GATAGTCTCTTGCACGGCTTTGTACACGTTTGTGAAGTCCTTATCACACTTGATGCACACCTCAGTGTCC

GACACCGGATAGCCTTCTGTGATGAACTTCTGCCAGTGGATAGCCTGCCCGAGA

>NODE_312_length_404_cov_4.078329

TTCTAAGAGTCGAGTTGCTTTTATTGACGTTGTAGAAGGCCTGGAATTCTAGGTTGAAAATTAACTCCAG

AAAAAACAAACAAAAATCGTGTTAGTCGTTCACATAACCTATTATTTCCTGTTTCTTATCAGAGACTTTG

ATCATGGGACATGATACTACAGTATATGTGGGCAATCTGGTGCTTCCAAATGGAACCCTTACAACTTTGA

CCCCTTGGGTCTATCTCTCACTGGCTCCTTCAACCTTTGGGTCTATCTGCTTAGACTAGAATAATAACAA

GGACAGCATTATATTCACGTTCGTATGATATCACATCAGAAAAGACATGTCACGACACCACAAAATGGGG

GTGGTTCGGGTACATAGCTCGTATATATTTCGTGTTAGTCCACACAACCTATTA

>NODE_313_length_404_cov_3.955614

CACAGAACCGACCAGAGACGGTTTGTTGCTGCCCAACTCTCAGGCCTTGTTTCCCGTGAGCTTGTCCGTC

TGCTGATGCCACAATGTGGAGTACATGCCCTTTCGCTTCAAGAGCTGGTCGTGCGTTCCCAGCTCCTGCA

CCCCACCCCCGTTGACGTTCTGCGTTTCCGTTCCGTGGAACGTCACAGCAGGCTCCCAGACCAGCGCTGC

TCCGTTCACGACCTCGAACCGAATCTCGAACAGTTCGATAAGCGGGAGAAATACAGTAGTAGATCCTTCG

GGAGGCTCCACCCTGAGAGTGTTCGAGCCCATGCTCATAGTGGTGAATTTGGCGCATTTGATTACGGCCG

AGCGGGATATCTGAACGACGACGTCCTACTCGGAAACACCCGGTAAGTGTTATC

>NODE_314_length_403_cov_9.062827

GTAACCATGGTGAAGGACAAGAAGGCGAGCGAGTTCGCAACCCGCGACTACACCATCCACCTCAGCAAGC

GGTTGTATGGCGTGACCTTCAAGAAGCGCGCGCCGCGGGCTGTGAGGGAGGTGAAGAAGTTTGCCCAGCA

GATGATGAAGACGAGCGACGTGCGCGTGGACGCGAAGTTGAACAAGTTCCTCTTCTCGAAGGGTGTGCGT

AACGTGCCCACAAGGGTCCGCGTGCGCCTTTCGCGCAAGCGTAACGAGGACGAGGAGGCAACGGAGAAGC

TGTACACCCTTGTCCAGCACGTGGAGGTCGCCAGCATCAAGGGCTTGCAGACGGAGAACTACGACCAGGC

TTAAGCGTGACAAACGTTGACGTGTTCGTACGCCGCCACGGCGCGCGATGAGC

>NODE_315_length_403_cov_3.714660

CTGCTGTGGGGTTCCTACGTTCTCACGCTGTCTTTTTTTGTTGTTGGCATCTTCGTGCTGTGGTGGGTGA

ATGTTTCTCAGATTTGTGTCTGAGGTTTTATAGCAAACCGGAATGGTCGCGGTCGTGGTGTCGATCGATG

TTCTGAATTTTCCCCTCCGCCTATCGAGGAGCGGAAACGTTTCCTTAACGTTTTGTTGGTTTCGGGCTGA

GACACACAGCAGTTTATGGTGTACCTGTATTATAACCAAGGGGGGTGTTGTGGTGGTCATGGCGATGCAA

CACATAGTCGGGCGGCAACCAGGCTAATGAACCAGTACGTGATACTGAAAAAAAAAAATTTCTACATATC

GGATCCCAAAAAACTAAATACACAACTACATAACGAAAGCCGTATTCTAATAA

>NODE_316_length_402_cov_17.136483

GGGTCCAGGACGCCCTTTAGATGATGCCGATCTTGTTGGCGATCTCGAGTGCGTCGAAGTCCTGGGTGAG

ACGTACGTACGCCTTCTTCTTGCCGTCAGGCCGGATGAGGGTGTTCACGTTCTGCGACTCGATGTCGTAC

AGGGCCTTGACGGCGTCCTTGATCTGGCGCTTGTTGGACTTCACGTCCACGATGAAGACGAGGGTGTTGT

TGTCCTCGATCTTCTTCATGGCGCTCTCGGTCGTCAGCGGGAACTTGATGATGCGGTGCTTGTTCATCAC

CGGCGCTCGCGGCACAGCCTTCGTCGGGTACTTGGGCGACCGCGCAAGGCTGAGAGTCTTAGGCCTGTGG

AAGTGCACGCTGGTGCGGACCTTCGCCTTCCGCTTGCCGCTGCCCTTGAGGA

>NODE_317_length_402_cov_8.068241

GGCGACTTTGATACGTTCCGTCGTTGGATGGTTATCGCTGATGATCAAAAGCAAGAATAGCGTCGGTAGA

CTGTAGTTTGTCAGTCAAATCAGCCATTTGGGTACCGTTTTGTTGTGCCGTGGTTCGTAGTTGAAGTGGT

GAGGGGTTTGCAGAGTGGTGGAGGGATTGCAGGTGTGTTGTTGGCGTTTAGGCTCGAGATCGGTGGGGAC

GCGCTTCCCGCGGACGCGTGAAGGGCAGTGTCAGGTATCGAAATGAGAGGTACACATCGCGGCGCGTCTG

GAGGACGATCGCGATTGTAATATATGCCGGTGTGGAAGTAGGAGAGATCATCAAGCATCAAGGCACGTCT

GAGTGACGATTGCTACGTAAATAAAATAATGGAACCATGAAAAAAAATTTTT

>NODE_318_length_401_cov_56.794737

GATTTCCGATTTTTGCGAGGCTGCAATCCGCGGGGCATTTTTGCCGCGTTGAGCGCCACGCTGGTAGGCC

TTGTCCCGTCTTGGCGGAGGACCAAGCGCCTTTTTTGTGATTGTTGTTGTTGGTGGTGTCGCTCTGTCGT

GGCGGGAGTACTTAACGGTGCGTTTTCGTGCCCTGTATTTTTGTTAGCGGCAGACATTCATCATATATAT

ATATTATGTGTTCGTGTCAATCGTAGTCCTGATATATGCCTCCGGTATGCAAGGATGAACTCCACGCGAG

CGTATGTTTTCAGGGGAGATGGAGTGCGTGTCGCCCTATGATTAAAGTCAGGTTACATCTACATTTTATT

CCCTGAGCTAAAATCATGCTGACAACACCCTGTCGACTTCCTTATGAGGGC

>NODE_319_length_401_cov_16.923684

GGGCGTGGGGATGGGGCATATATAGGGCTGAGAGGTTGCTTGGAACAGACACCCGCTTTTTTGCCCGTTC

CAGGTTTATTGGCAAAACCATAACCTAAATCATGTAACAAGGTAGACTACTGTACAGCCACTTCCGTTTT

TTGCCTTTGGGCGAGGGAATTTTTTTACACCTGGCGGTGGCGTTTTTTCATCGGTCGTCTAGAAGAAGAG

GTATCGGAACTAGAAGGAAGGAAGGAAGGAACGAATCACAAAACTCGCATTAGCTGTGTTTGCGCCCCCG

GGTGTACTGTGCCTCGCCGTGCGCGTGAGACAGGCCTCTACCAGACAGACCACACTGCTGCTGACTGCCA

CTTCGGGCAGGAAGTAGTAAGTGTGTCGACCTGACTGCCTGCCTGCCTTTC

>NODE_320_length_400_cov_38.113456

ACGCTGGCCCAGCTAATCGTGGGGAAGGCTACGGTCGATGCCCCTTGGTCCGTTTCATGCCAGCTATATC

AACCGAAAATTTGACACCCCGACAAAAGTTGAGGTGCCTTTTTTCACTCCGTTTTTGCCCGACCCTCGGT

AGTTAAAGAAGTCCCCTTATCGTCCTGTCTTAATCGCTCGATAACGTTGCACCCCGGAACCGTGGCCCTT

TAAAGCACCCGAAGCTCGAACCAAAATGGGGAGAGAGTTGGCGTGTGGGGCAGCGAGGATGGGGAATCAT

GCGATGCTTTTCTGTGAAGTTTTGGCGTCGAGACGAGTTTTTCATTTTGGGGAGGTCTTGCCGGGGCATG

TGGGGAGTCAGTACCCTGCTGCGCTGATTGGGTACGGAGACATTTTGGAA

>NODE_321_length_400_cov_7.559367

CACCCCGAGAATTCCAAAATGTGGGGAGACGACTGTCGACGGGCTACGAGAAAGGACTCGCGGCAAAACG

CCGCGCTTCTATCGCGAACGATAAAAGACGTTTTTAAGGGGTCCCGACGGTAATTTTCCTCCCAATGCAT

TAATCTGGCACAGGCCGCTGTGGCGCCCCAGCCAGGAGGTCATAATCGGACGCGATCGAGATGAAAGCGT

CGCGACAATCCACTCACTATTTTCTGAGAGGCAGCCAAGGCAGCCTTTTCTGCACCGTGCTCAGTTGTGT

ATGGGGTTTGGACGAGGGTAGCGGCGTTTTCGGGGCACGTCTTCCTTCATTGTTCGATGTGTGTTTCTGT

GTGTTGCAAGGCGAGGGTGAGGGAGGTTCCCGTCTGCAGCAGCAAGTAGT

>NODE_322_length_400_cov_5.564644

TCCGCCCCCCCCAGCCTTCGGAAACAGCAACCAGATGACTCTTCGGGAACGACGCGAGCGACGAGCTCTA

CGCGGACGAGGCATAGAGCCGAAACCCCCTAAACGCCTCCCCAAAAAGTTGAATCGTGGTCGGGTTTTAG

ACGATGGAACCGGCGTTGCTGGCGATACGGGGCCACAGGTCGGCGCACTCCTTGGCCACGGGGCCGTTGA

TCGCGGAACCCTTCATCTCGCCCTTGTTGTTGACGATCACTCCGCCGTTGTCCTCGAAGTAGATGAACAC

ACCGTCCTTCCGCCTCCAGGCCTTGCGCTGCCTCACCACCACGGCGGGCATCACTTTCTTACGGAGCTCA

GGCTTTCCCTTCCTCACCGTGGCCATCACCCGCCATTTCCGACGTTAGCT

>NODE_323_length_399_cov_26.423280

GTCTCGGCCGTTTCGGATGAAACGGGGGTTCTCGTTATCGAACAGGGTCTTCACGTCGGAATCGCCGTTC

TGCACGTCGAGCCAGTCGTCGAAAGACGTCATGTAGTCCATCTTCGCAACGAGGGGGACAGCCGTTGGCT

CAACCACGATGGAATCGATCATAAAGTTGGACAGCAGGAACTGAGACACGACGGGACCGGTGGTCACACC

CGGCCAGGTCGTGCGGAACAGATCAGTGACGGGGTCGATCTTCCCCCCTTTGCTTCTGGGGATGTTAAGC

CCCTTAAACGCCGGATTTTTCTGAAGGTTCTCTGCGGCAAGCGCGACGAGATCGTTGGTGGCGAACTGGG

AAAAGGGGATGTCCCTCGCCAGGGCCATCCAGTACACCTCTGTCTGCTG

>NODE_324_length_399_cov_7.277778

CCCCAAAAACGATGATTTTATCTCTCTTTTCAATATTATTTTGTGCAGGGGCAAAACGAGGTAGCGTAAA

CGTGGCAACGCACATTCTGCCACCAGCCCTCTCTCCCTGCATCCCTATCGACGAACCGCCTCCAACGGAA

CTCCTGCAAACCTCACTCAGCCACTCGCTACAGAAGCAGCCGCGCTACACCGGACAGCATGTCCACAGCA

GCAGAAGACCACCGAGGGCAAGCATCACGACGCAGGCACCTAACGAATCGACCACCGGCAAACTTTGATT

TGACGCCCGACCAAAGATAAGAAAACGCGACAAAGACCAATAGTCCACCGTTACCGACAAGAGCGCAACG

CAACAAGGCTGAAACAGGAGCCATCGAGGAAGTTGGCATGCCCATGAGG

>NODE_325_length_399_cov_5.759259

CAACAACAACAACAACAACAAAATACTAAGCGAAACAAAACGAAACGAATCGAAACGAAAACAAAAAAAA

GGCCCGCACGAAAAAAACAAATAAACAAACAAACACACCTCTTCCTACCTACGAACCACCAATCCATATA

TTCCAAGAGCCACAAGCAGTCGTACATATATCTTGCGTCGTGCGTAGAGCTTAGATCCCGACCCCCCCCT

CCCCTCTCTCCTCCTGCCTTCAACATCATCATCAACTCCGTCATGTGTTATCACTGCCCGTCTAGCCGCT

CCATAAAAAGAGTCGCCACACAACGACGAGTCATCCATACTTCTTGAGCTAAAATGAGCTAAAACAAACA

CAAAAGAGGGCAGATCGAGATCGAGATCGAGAGAGAGAGAGAGAGAGAG

>NODE_326_length_399_cov_5.523810

CCTTGGCACCCGTTTATTCCATTTTTTTTCACCAGTCAAGCCATCGTATTATTGAAGCCCAAGCTAAAGT

CGACAATATTCAGCCAACTTAGAAGAGCATCTCATGCGGCCGGTCGCCCTTCTTGAGGGAGAATTTGGCA

TTGAGGTACGCCTTCAACAGGGGCTCGGCAGCAACAGCGGCCAGGAGCCCAGCGTCGACGATCTTCTGGT

CCGCCTTGCGCTTGTCGGACACAGTGGAGGTCTTGGCTCCCTGCTCGAAGAACTGGTCCTCTTCGGTCTT

CTTAGCCGAAGCCACGCGCGTGAAGTACGAGTCGTCAGCGACCGGGATGGTCACCTTGGACAGGTCCACC

TTCATCGACGTAGCAATCACGTACGCCTGGTTCACGCGCCGCAAAGGGA

>NODE_327_length_398_cov_5.676393

GGCGCTATTTTCGTGCGGTCTTGCCGTTTATTTATTTCCTGCAACCCTCCCTTCACATCTCCCTTTTTGA

TACTTTTGTCTGTTCCATGCCTACCATTTTCAGTCCGTCTCACTTTGGCATTGATTTTCTCTCTTTATCG

CTCTCCTCTCCGCGTCATTGTCCCGCAATGCAGTAGCACTGTTGTGTGCGACTCTTTGGCGAAGCCTTAC

GACCAAAACCACACTTACTTATACGAGAGAGGCAGATACATGAGAGTCACAGAGAAAAGAGATTCGAAAG

TAAAAAAACAATCGATGTTTCGGTTTTCCCATCTACTTGCGGGGCGTAATCTTCGAGAAGGCCATGCCCG

TGGGGCCGGCTTTGGCGAGGAAAAGGTCTTGGTCGATCACCGTTGGCT

>NODE_328_length_398_cov_4.925729

CTTTACAAGGGCGGCCGACTTCGTGCCTGAAAGGTGGATCGAGGCGGAGCGCGAGGAGGCTCTGCTGGGC

GAAGGAGCCAAATACTCGGGCGAAAGCGTACGTCACGACGAGAAGGCCTTCCTCGCGTTTGGCGCCGGCC

CTAGGGTATGCCCCGGACAGGACATGGCGAAGGTCGAGGCGTCCATGATCATCGGCGCCTTGTGCACCCG

CTTCGACATTTCTCTGGAACCGGGTCACGACGACCCTCCGGAGGAGGTACATTCCTTCACTTCGGGCCCG

AAGTCTGTTAGGCTGGTGCTGACGAAGGCGAAGCTGTAACCATGCGACCACGACTAACCGATCGATCGCA

CCAACATATACCAACCAACATTACATTGAGGGGGGCATGGTGGCCGAT

>NODE_329_length_397_cov_21.976064

AACCGGATTGTGAAGACAGGCGCGACATCAATCCAGTATACCTTGGTGGAGCGTAGTGCTTTAAGACATT

CCTTCAAGTATGTGGCTACACACACCGCGTTATTTGTGCGCCTATGCATGATTACTACAGTAGATGAATT

CTCCCTTGCCCGGCTTGTGTTGTTTCGAGCTAGACTTGAACACAAGTAGTCGTTGTTGGTACATGTGCAG

ATATGTCGAGAGTTGCTGCGGCTGAGTTGGCTCCTTGCCGGTTGCTTTGTTGGACGACAGCTTCATATGT

TGGTGTTTCTCTTGAGAGATTAATTTGTGGTTTCTTCGTTGTTCTCGAATTGGTTAGATTGTGGTCGAAG

ACGCGTCCGGGTGGTTTGCTGGCAGCTCGGTTAGTAGAAAGGGCTTT

>NODE_330_length_396_cov_2.117333

TCCACGGGGTTGCCGAGATCGGTGAGCTCCTTAAGCTTGGCGGTGTACACCGACTTAGAGCTGTCGAAAC

CTTCGTCGGAGTACAGCCAGTCTTCTGCCGCTGTCAGACGCCCCGTGAAGGTGTCCGCCTCCTTCCCCTC

CACGTACGGTCGGAGGTTCCCAACGAGCTTGTCGCGCATGGTGTACACGTAGCTCTCTAGCTCGTTCCGC

TTGTCGGCGGTCTCCCTCAACACTCGGTCCTGGTTGGCCATCTGAGCCTCCTGCAACACAGCGGCGTCGA

TCTCGCTCTTGGTCCAAGCGGAAGTCTTCGGCTCAACCTTCAGCGGCACCTTGCTGTAGACCTTCTTCTT

AGGTGCGTCCGCATCCTTGGCAGACGGGGGCGACGACGAACCCTTG

>NODE_331_length_395_cov_8.729947

GCTGAGATACGAGACGAGTGTGGTGTTTCGCGGCCAGTGTACCATCAGTACCACTGGCGACAGCAAGAGT

TTAGCGTTTAGGCGTGGAGCCCCCGCGAGTCGTTGCTCGGTCTTTATTGGTTGACGAGGCGGAACATTGG

TTGGCCTCCGTTCGGTATTGGCCAATCGATGGCTAGTGAGGCGGTAGGAAGGAGGAGAGGGTTACTACAG

GGATTCTGATTTCCTTGGTTACCCCTGACCCTTTTTTATATCGCCGTGTCTGACACGCCACGAGGTTGAG

CTGCAACTATGGTCCCAATCCCACTAATCCATTGAGGAAATATACACCCCTCGGCAACGATTTTCATTTC

ATGAAATCATTCCCTTGTTTTTTTTTGTGCATTGAAGTTGAACCG

>NODE_332_length_395_cov_7.671123

GACAGACAGACAGACAGACAGCTTGGTACTCGTACACTTTGGGGGAGTTTTCCTGGTGTGTTGGGGGGGA

CTACTACGCAGGCGTACAGTCAGCAACAAGAGTTGAGAGGACAAAAAAACGTACAAAGTGCGCCGATCAG

GTAAGCAAGAGAGCCGGTGGACACGTCTCATTTCTTGCCCCACATCCTCCTCTTTTTGCTTTGATTGTTA

AACCGACCTACATATTATTATTTCATGAGGGTATACATACACCCCGTAGCACGCGTGACCGACACATCCC

ACGCAGCTTGCAATATGAGTGTCGACGAAATTCTCGATCTCACGGCTGCTGTGTCCTTGTTTGTTTTTAT

TTTTTGTAAGCTGTAAATGCCTCGCCGTGGCAACGTTTTTTTTTT

>NODE_333_length_394_cov_11.592493

GAGGAACCCTGTCGCCGAGGTCTTCGTCTTCGGCCCGTAAACTCAGTTATAATAGTCGACGAAGGCTGCG

CGCCAAACCTCAGGCGTCACGCGCTCAGGCATCACGCCGATATGTAGACCATGCCTCTCTCTCTCTCTGA

TGGTCTTATCAATAGCCTACGAATAGTCTACTATTTGTAATAGCCCTCTAGCAGTATCATCTTATGAATA

GTCTATCGGCAGGGGACGAAGGTGTGGCCCAACGGAATAGTCTTACTACTGCCAGTAGTCTTCTCTTGTG

GCAGTCAGTATGAGAGAAGGGAGGGTTTTTCGTGTACAACTGGAGGAACTAGGTCGCCGGCACCGGACCG

TTACTGATGACAACACTGGGTGTCGGTATGAGAGCAGGAAGGCT

>NODE_334_length_394_cov_6.863271

ACATCTCTACCGCCTGCTAATCTGAGCGCCATTATCTGGACACTTCTCCCTTCGACTTTTTTTAGTCTTT

ATTATTTTTATTATTATCAGTCCGTATTGTTATTATTTTTACGTTCCTTGTTACATTCCATTTTGTTTTT

ATTATTAAACTTCCATGAGTTTTTTATTATTATCGGCCATCCTGACACTTTTTTATTATTTCCTTTAGTA

GCACTAATCTTGTGCTATGTAGTCATGCCGCGTTACTTTCCTTGATAGTTATTTCCGTTTTGTTATTGTT

GTTTTTTTTCGTTTTTTTTTCCCCGATGTTTTCACGGACAGAAGCAGAGCCAACTATATAGCCATCCATC

CATGTGATGGACGGCCCGCTACCACCACTCCCATCCCCCATCCC

>NODE_335_length_394_cov_3.683646

GTCAGCAAGGAACGAACATCAGAACGCGCACACTTAAACAGGGGATCGTGTTGAACGAAACAACCCGCCC

CGTTCAGCACATACTCGACACGGGAGGTGTATCGTAGATTCTCTGGCGTGCCTGCCTCGAAACACCAAAA

ATACTGGCCCAGATGATTTTCCCCCCGGCGAGGAGCAACCATCCCAAACGTTCAACTAGGATGGTGAGTA

AGCCCTATTTAACGGCTAGCTACACCCTGTCCTACAACAATATGATATTTCGCATTACGTTTGAGTGCCA

AAAATGTCATTTTGCTGCGGGACTTCACTGACTATAGTAATTGATTGGGTGTCGGAAGTAAGACGACTAG

TTTGACTGGGCGCTTATCTCAACAGCAAAACACTCGCAAGGGGT

>NODE_336_length_390_cov_13.295393

ATTCCAAAATTTTTTTTTTTTCAAGTGTTTGGCCCCTTATTTATTATTTTGGAACGAGTTCTTCAACGCG

GGAAGACTGGTGTCTGCGAGAACCCTATACCCACATGGCATTCCTCGGCACGACTCTCCGTTACGGTCCA

AGACTACCCTTTATCTTGTTGTGGTTCGTCGGAACAGTTTATGAATTCCTGGAGCAACACCAGAGATGCT

ATCGGCGGAAATCTCGTCGTTGCTGCAGCGGTGGTGGACTACTAACAAACGATTGAAGAACGAAATTCCA

AAAACAAACCCTCCAGTGGAAAACAGTCACAATCCCGCTGTCAAATATCACCGCAGAGAGCACACCATCA

CACCCTATGGCACGACCACCAAGGAGATTGGCGGCCTTCA

>NODE_337_length_390_cov_3.666667

ATTCGGTTAGATGTCAAATCCAGTATACCCAAAGATGTGGCGCCCCTAAAGCTTCCGGGCTGTAGAATTA

GGTCGTTGTTCGTGAGAGTCAGTTCGCGTAGGGACGTCAGCCCTTCGAAGACACCAGGTGGCAGGGATTC

CATGCCGTTGTTGCGCAATTCTATATCTTCCAAGGCCGTAGCCCCGATGAAGTTTCCAGGCTGCAGAATA

AGGCTGTTCCTTTCGAAATTCACTCTCAGAAGCGCCCCGAAATCGTTGAAGAAGTCGAGTGGCAGGGTGG

TGAGTTGGTTGTTGAAGAGGATCAATGATTCGATCGTGTCTCGTCCAATGGTGTCCAAGCATGCCACGAG

ATCAGGAACATCTTCGTCGGTGAGTTCGCAGCCAGACAGA

>NODE_338_length_390_cov_3.449864

AGCTGCCTGTTGCAGCTATGTACAAATCTGTGTGTGTGTTCTTCACTTTCAACCCTTGACAGCCTTGACA

TGCCGTGACTGTCACCAAACAACTAGTTTTTAGTGTGGAACACTCCACCCTACACCCTTGACAGCGCATG

TTTTGCCTTGCTTTTTAACACTGGTTCTGTTTTATAGTGCTATTAAACACAGTCACTACCGCCTAGCTCT

CCGATCTCGGCACGGCTTTCGACACGAGCAGCACAGGACGGTCGGCACAAATAGCGGGCCTCTATCAGTC

CCGAGCGTGTCTGAATCGGTTCAGTGTACGCGTCGCGCGCCAAAAAAAAGCAAAGCGAACGAAGTGCGAA

GTGCGAAGTGCAGACAGACGAAGTGCGAAGTGCAGACAAA

>NODE_339_length_389_cov_6.456522

TTTCATTTGGGCATTTAGTTTAATGTTCTTATTTAGTGGTCGTGGCTATTGGCAAGAGCTTATTGAATCA

ATTGTTTGGGCTCATAACAAAATTAAAGTTGCACCAGCAATTCAACCTCGAGCATTAAGTATTACTCAAG

GTCGAGCTGTAGGGTTAGCGCATTATCTATTAGGTGGTATTGGGACAACATGGTCTTTCTTCTTAGCAAG

AATTATTTCTGTAGGATAAAATTAACGAGGTAAAATATTTATGGTAACTAAATTTCCAAAATTTAGCCAA

GCTTTAGCACAAGATCCGACAACTAGAAGAATTTGGTTTGGAATTGCAACAGCCCATGACTTTGAAACAC

ATGATGGGATGACAGAAGAAAATTTATATCAAAAAATCT

>NODE_340_length_388_cov_41.841962

AGGAAAGGGCAGCCGAAGCTGCCCTTAATTTAGTCTTTCTATCGAATCCGTGATAGTTGATGCTCCCTGC

AGTACATCCTTTTGACTAAGCCAAACTAGGCGTCCACTTCCCAAATCCTTTGGGTTCATCCTTGTTACAT

CCGTATAAGATAAACATCTTGTCTATCCGTCATCTCCCTATCCTTAGGAGGTGTCCTTCTTTGCCATCCT

GGCTTAATAACTACAATCCTAGTAGGTACTAAAAACTCCATCCTAGAGTGTCCTGATCATCCTTAATTAT

AAAACATCCTGTTTTATAAGCTCTCATCCTGAGAGGTCCAAATTGCCATCCTAGCAATACAAAGGTTCAA

ATTTTACATCCTGTAAATCATCTACGTTCCTTGTTTGC

>NODE_341_length_388_cov_10.798365

CCAAGTACGACGACCTCGCGGCCAAGCTTGAGGGGGTGGACTCGGTGGTTATCGCCAAGATGGACTCCAC

CGAGAACGAGATCGACATCGCGGGGGTGGAGGTGTCCGGGTTCCCTACCCTGTTCTTTTTCCCCGGCAAG

TCGAAGGACACCCCTCTCCAGTACCAGGGCGCCCGCGAGACAGAAGACCTCGCCAAGTACATCATGGAGA

AGGCGTCTACGCCGTTCACTTTGGAGGGTGTCGAGTCAGCGGAGGCACACGAGGAGCTATAAACAGCGCC

TGTAGCGGTGTGTAATCAAAAAGTGTTTGATACTGGGGTACGCGCCAAGCAAGAAAAGAGTAACGAGCAG

CGGCGGCAGAATCGGCGGGGAGTTGGTTCCCTGCTGAC

>NODE_342_length_387_cov_8.191257

GCCAGGAGCGCCGAGTTGTCGCCGTCCTCGGCCATGAGCTTGATCGCGTCGTTGGCGGCGAAGCCCTCCT

TGAACAGGGCCACGTCCCTCGTGTGAGCCACAACCAGCACGGGCGAGAAGGGCTGCAGGTAGGTCAGGTT

CCTCACGCGCACCGTGTACTCCGCCGCGTTGGCAAGGGAGAGCGCCGACAGCACAGCGAAAGCAGTAGTC

TTGACCATCTTGCTCTTGAAAAGTAGTGAAGGTAGAAGTAGTAGAGGTGGCGAGCTCGGTGATAATGGAG

AAAGAAAGAGGTGGTGGTGGAATGAGTAATGGAGTGCGGGGTACGACAACAAAAGAGACAGCAACAGCAA

CGTTCGTAGTTAGACCTCAGACCAATGCAGATGGTTC

>NODE_343_length_386_cov_388.339726

GCTACACACGTGCTACAATGGAAAGGACAACAAGTTGCGAGGGAGTAATCCAGAGCCAATCTTTAAACCT

TTTCTTAGTTCGGATTGGGGGCTGCAACTCGCCCCCATGAAGCTGGAATCACTAGTAATCGCGGATCAGG

ATGTCGCGGTGAATACGTCACTGGGCCTTGCACACACCGCCCGTCACGTCCTGGAAGTTGACTTGGCCAG

AAGATTTTGGAGTAAACCTTTCAAACATATCCTCATTTGGTAAAGATATTTATCTAGGTTAAGTGAGTTT

GGAAATTCCCTTTAAAGGACAGGTAAGCAACCGGGATGAAGTCGTAACAAGGTAGTCGTAGGGGAACCTG

CGGCTGGAATATATAATTAAGAGGCTCGCCTCTTTA

>NODE_344_length_386_cov_29.134247

ATAAAAAAAGTGCAACTTAAATAGCTGCACTCTTTTAACTAGCTTGTTATGCATCCTGCATCTTATTCAG

TGTTCTCATTCATCCTTAAAAACTTTCCTTAGTATATCCTTTGAGTAAACTCCTACTCATTGTCCTTAGC

TATAAATCCTTTTACTTCCTTGCAAATTATATCCTCAATTACTTTGAGGCTAATTCCATTTAGATGTGCT

TCCTAAGCAATATCTTCCTAATATCAATTATCCTTAATTGATCACATCCTTGTTCATTTATCCCTTAGCG

AAATCCTATTCGCCTGATTTATCCTATAAATCGTCCTAAAATATTCTATTTCTAGAATAGTATCTAGAAT

AATAAACATCCAATTAACTCAAGTCACACCTAAATC

>NODE_345_length_385_cov_6.098901

AGGTGACGCGACGTTCAGCAACACCGGATGCAGGCGCGCCGAAGGCTCACCGGTGTACAACGCCGGACTC

ATAAACTTCTACGGGAAGGTGCTCTTCAATGATAACGACTCCGACGGCAACGGTGGTGCCGTTCTCAACG

ATGCTGGATCAATCAACTTCAGGAGCGCTGTGGAGTTCAACAACAACGAAGGAAACGAAAACGGTGGTGG

TATCGCTGTCGACGGTGGAGACGTCACGTTCCGGAAGGCGGTAACGTTTAGTGGCAACCGTGCTGAATCT

GGTGGGGCCTTTGCTGTCCTAGTTCGCAGGGATTTAAGAACTGGCGAGATGGTCGAGTCAGGCGTCGGCT

CGCTTACCTTCGAGATGCCGTCCGCAGTGACATTC

>NODE_346_length_384_cov_17.980716

CAGAATTAGCTAATGTAAAATGGTTTGCAGCACGAGTATCAAGAAGTTTGGATACTCCTATGCAGGAGGC

TGTTAAACCGTTCAATCGCCAATTTTCTGATGTTTCAGTAATTGAGTCGAACAGTGCTTTAGTTAGTGAG

GATGGCTTAAATGGTGCCGACGCTTTTGGTGAGGATGGCAATTCAGGAAATAAAAAATCCAAAGACGAGA

TAAAAGAGATGTTGTCCAAATTAGAAAGTAGGGAAATTGATTATACCCGTATAGGTTTACGTAGTGAAGT

TTTACAAACCCTTAAAGAATTGATTGAAACTGTAGACCAAAATACTGCAGATGACATTAAAAATATGTTG

TATGAAGCTTTGCTTTTAATTACTAATGAGGAAG

>NODE_347_length_384_cov_11.798898

CAACGTGCCAAACCAATACAAAATCATTCTCGCACTGTCACGAGACGCCAGGAGAAGGAAAGACCGGCTC

GCCCATAGAAGTGTGACAATGTCGAGCGACAACTGAATATATCAGACGGCCCGACTCATACGTTGGATGG

AGGGATATCCGCTCGCGTAAGCATGGGTTCGAGGCCACTCTGGTCAGGAGACCATATCATACAACGTAAT

GGGCCTGCCAATCAGAAAGTCCACCCCCTATCATACATATATCTACCCGTTACTCGATCGTTTTTGACGT

GGACTATGGTACGCTACGATTGAAGGACAACGCAAAAGCCGAAATGAACACAACACACGAGTATGTGCGT

TATTGAAACCGCCGTCAAGTAGGACTGCGCACAG

>NODE_348_length_384_cov_6.837466

TCGATGTTGAGCTTCTCGCTCACGTAGACACCGTCGAGGAACTTACGGATATCCTTCCGCTTCACCGACG

TAGACTGCTGGATGCGTGCCGCGGTCATGGAAACGGCCTGGATGTCGTTGCCGCTGAGCTCGATCTGGTC

CTTCACGTCCGCCGTGCGCACGTACTCGACACCCGGGGCAAGCTCCACCCTGCGCACGCGCTTCTCGCCG

AGGAAGTTACGGATCTCCACAGTGCGTCCGGTGATGGACACGTTGATGGGGAAATGGGCGTAACAGAAGC

GCATCTTGTACAAGTAACCCTTGGTCACACCCGTGATCATGTTGTCCACGTGCGAGCACACGGTACGGAT

GCATGCGAGCTGCCCCGCCCTTGTTCTCTACTAT

>NODE_349_length_383_cov_30.033149

ATTCCATGGTGGAGCTGGGGGGATTTGAACCCCCGTCCGAAAAGCCTCGACCGTCGGTACTACATGTTTA

GTATTGTCATTTAATTAACCTTTAAGTCCCGGACAAACACGGTAAATAAAGGCGAGGCCGCTTAGTTTTA

GCGCTTCAACCCCGGCCAGGGTTTCCGTCGCGATCAGGTGTTAGGGTGACACTCCAAAATCCAGTCCACA

AGAATACATGGAAGGAGCGCTAGCCAGCCTAAGCTGCTAGAGAGTAGTTATCGTCGTTTGCGATTACTTT

AGGTTGCGGCTGTTTAACGAGGCAAACCGCACCTCGACATGCACCTTGGGCTTCTTGAATTCCGTCGAAT

CCTAATCAGCCCCAACGCTTCGCGGCCTGAATG

>NODE_350_length_383_cov_10.552486

ATCATTCCATTTTTTTTTTTGTGTTCCCTTTTTAAAGCCCTGGTGTTGACGTCGCCGCAGCTACAGCAAC

CGTCCTCCCAGCCCTCTCAAACCACCCCTTCCGCCGTCCCTACCGTACAACCAACCAAAGTGTTTCAAGT

CTCAAGCACCATTCCAAAAACATGTTCACTCAAGTACGATATGAAAAACGCCACAAGGAGGAACAAAACC

CGAACCGAGTCTTGGACACAACCCCCAAAAGCGTTTTTCGGTCGTGAAAAGCGAAGCATACCATCATGGA

GGGATCGAAACCCATCCATGGTTTACTGTATTGTAGGTGACCGAAACGGCTGTCGGATCGGGACTGATCA

GTACTGAGCCTTCTTGGCGAGCTCGGCCTCGAT

>NODE_351_length_383_cov_9.005525

AGTAATTGCTTATTCCACTTGTAGTCAATTAGGTTATATGGTTTTTGCTTGTGGTCTTTCTAATTATTCT

GTAGCGGTGTTTCATTTAGGTAATCATGCATTTTTTAAGGCTCTTCTATTTCTTGGGGCAGGTTCTGTAA

TTCATGCGGTTGGTGATGAACAGGATATGCGTAAAATGGGGGGGTTACGTCGTTTATTACCTTTTACATA

TGCTATGATGGTAATTGGTTCTTTAGCTCTTATGGGTATGCCCTTTTTAACAGGGTTTTATTCTAAAGAT

GTTATATTGGAGGTAGGTTATGCAACTTATAGCAATGCTTCACATTTTGCATATTGGTTAGGTAGTTTAG

CGGCTTTTTGTACTGCTTTTTATTCTATACGGC

>NODE_352_length_383_cov_6.143646

AGGAATCGTCACCCGGGGACGATAGGTTCCCATCCCCGAATCGGAAATCCCGCCATGCATCCACAACTCC

TCACCCGGGGACGATAGGTCCCCATCCCCGGTTCGGAAATCCCTCCATACATCTACAACTCCAACAGCCC

GGGGACGAAAACCGAGGTCCCCCGGGTCGACACCGCCTGGCATGGTGTGGAAACCCCAGCCACATCGGCC

GATCGCTGATTGGCTGGAGCTCGAAAATAAAGCGATAAACCAACCGGACCGGGTGTTACTTGCTTGCTTG

CTTGTTTGTTTGCTTGCTCGCTCGCTCGCTTGCTTGCGGTGCGGTGCGGTGCGGTTTTCAAGAGCGCCAA

AATCCTACGGTGCGGTTCGGTGCGGTTTTCAGA

>NODE_353_length_383_cov_3.991713

CTCGGTGATTGTTCACTCTCCCCGCTGCCGAGAAATTCGGGTTGGGAAAATCGTCGGAGGGGATGTCTGG

ACAAAGGACCGACTGCAACTACCTCGCCCATGCGCTGTCCATTGTGGTGGTAGGCGCCTCCGGCGACCTC

GCGAAGAAAAAGACCTTCCCGGCTTTGTTGGACCTCTACCGACACGACTTCTTGCCCAAGTTCGTGACGA

TATGCGGATACTCCCGGTCGGGCATGAGTGACGACGACCTCCGCGGGAAGCTGAGGCCGTACCTCACGAA

GAAGGGCGCGGAGCCAGACGACGTGGTCGAGGAGTTCCTGCAGCGTGTTTACTACCGGTCTGGCGGCTAC

GGGTCCGTGGAGGCGATGGGGAAGATGACCGCC

>NODE_354_length_382_cov_8.393352

TTGATTGATTGATTGATTGATTGATCGGTTGGTTGATTGAGTGTTTGGTTGACGGGTTGGATGTTCATTT

GTGTGTTTTTTCTGTTCGTTTGTTTTGTATGTCTGTTTGTCTGCGTGTTCGTACTTGCTGCTTGTTGCTG

CTGTTGTTATTAATAGTAGGCCCCTCCACCCGCGGGTATTGATGACTGCTACAGTACAATATCCATATTT

CACAGTATTAATAGTAGTGGAAGTTTCCGTACGTCCCTGTGAGAAGATATCGATCAAGCCATGTATGGAC

GACGATGGTGATGGATTTTGCTGATTGGATAATGGATGATGGATTATATAATGGCAATGAGCCGTTCGCT

CGTTGGTGCCACGCTCTCTGCTGTGTCGATTG

>NODE_355_length_382_cov_5.761773

AACTGCCCGTGCGGGCACAAGGGTAAATGCTCCGGCAACGGCACGTACAACGGCAACAAGTAGATTATTT

CTCCCTGGAAACCCGGCACAAGCTCTCCTCTTCCCTGCCGCGCACGGGGCCGAGGCCCAATCGACTCGAG

GCTGAAGTCCAATAAAATAAAACAATCAATTATCATCATAGCCGGAGGCGACAAGCATCCAATCAGATCG

CTTGGCTGAGGCTCGATGACCTCGTTCGTGGCAAGACACCCAATCAAGAGCCGAGCCCTTCATTTTTCAA

CCCGAGAGAAGACAAGACAGCCAGGCCCATTGTTGGGGTGTTGGCGGGGGTGGTCATGGTGGTAGTGGTG

GCTGCGATTGGGCCAACACCAGCGTGATTATT

>NODE_356_length_382_cov_3.961219

CGTACGGCGTTACGCCGTGTACAGCGTTTATTTCATCCACATCTTACGCTTAACTTGTATCCATGCTATT

TAAATGCGTACGGTACGTGTAACGTAGGGCGTAGAAGGGACAACGTGGAAGTGAGGTTCGCGATTCAAGC

TGTGTGTTTGGTTTTCGCTTCGAGCGACCGTGTCTGCTTTCCCTGCTACACGGACGGGCTTCTAGCTGGC

GGCGTAGTGGTTTAGATTCTTCATTTATGCCTTTGGCGCCTCTGGTAGAAGGGCGTTCGCAGAGTGTTGT

TTATTTTTGTAATTTTCCGCTGGATGGATTGTTTCTGGTATTTCCCGTGTCCTTGTTTGGTGGCTTCACA

ACCGCACCGCGGGGTATGGCATTTGCGATGTG

>NODE_357_length_381_cov_16.263889

CTTGGCACCCCGAGAATTCCAGCCGCCTGCAGCCGTTTGAAAAAAAATAAAGAAAAAATCAAACATTTTT

TTTTTTTCCCGATAAAAAAATGCATTTTTATGTGGACACCCTTCGATCCAGTAGTCAGACAACACACATG

CACGCATGCAAGTGAGTATGTGATGAAATGTTCGCTCTGCACTTCCTGCCTACACGCAATACAAAAAGTA

AAAATGAAGCCCTCGTCAACGTGGCTCCTCCCAACCATTGACCCCTCTCTGCCAGACCCAATCACCGCCC

GACACCCCTTGGGTTGGTGTCCGTCCCTGGTCGGACAAGCCGCCGCAAAGGTTGTTTAAACGCAGTTACG

TAGTTATTCTCTTGTCTTGAAACACAATTTG

>NODE_358_length_381_cov_11.411111

ATTCCATCCCACATGCAATTTCTTTACGGAATGTTATCCCAGGTTCTCTGTTGCCCCTTTTCAGAACAGA

CCGAACGAATCGGCCAGCCTTCACTTTCCTAACAAAACGATATAACGTTGTGCTCAGTGTCCGAATAACT

CTGTGTACGATTGCCGTGGGCACGAAATCCTTCCCCGCTCCAGCATCCAACGTTGAATGTTCATAGTGAA

TTCCGTTAGATCTTTGCTACGCCAACCAAGATCTGCACTAGAGGCCGCTGCACTCCCCCTCACGGGGTAA

GCTTCTTCACAACCTCCACGTACACCTACTCGTTAAGGCTTCGCATCTACCTTAACGGTGGAGTATAAGT

CACACGCTTGAGCGCCATCCATTTTCAGGGC

>NODE_359_length_381_cov_6.152778

CCCGTTCCCCCGGATAAAAGTCTCCTTTCATCCCCTGCTCCCGCCATAGATTCACAGATCGCGACGGTAA

TGCGATATAGCATACATACTTCTTCCCAAATAATTAATCTGGCGCTCAAACCACAGTACCCTAAATCATA

GTTGCTCGTGCTGGAAACATCGAACACTCTCGAAATTCATTGCCAGGAAATACGGCATTATCCTCAGAAT

TAGCCAAGGAAAAACGGTATTATTCTTCGGATTCATAACTCACATACCCATACGCTGCTGTACGCTTACG

CAACGACCGGCAATAAATACTCCATTCATAAACCAACATAAACCTCCTTGATTATCCCCGATTACCTCGT

ATACTAGTACCCAGTATGGGGCCTAGCTGAA

>NODE_360_length_381_cov_4.108333

AGAAGCCCCACAGGTACAGGCCCGGTACGGTTGCGCTGCGTGAGATCCGCCGCTACCAGAAGTCCACGGA

GCTCCTCATCCGTAAGCTCCCCTTCCAGCGCCTGGTGCGCGAGATCGCGCAGGACTTCAAGACCGACCTT

CGCTTCCAAAGCACGGCAATCCTCGCCCTGCAGGAGGCCGCTGAGGCCTACCTGGTCGGCCTGTTCGAAG

ACACGAACCTGTGCGCCAGTACGTTTAGCAGTATCAACGATCTCTGCTGTAGATTGATCAATCAAACGAT

GATCGAACGCTTTCAAACGAATGCGAATTCTTTGATTTGACATGAATCAAATCCCCATTTTAAAGAACAA

CAAAACACAGCCCATCACCTTTCAATATATT

>NODE_361_length_380_cov_28.055710

TAGGTTGGACGCCGACATGGTGTTACGTAGCATTTTAGTCCACGTTTCTACCACATTGGCGTCGACGGCA

GGACATGGTGCTTTCAGCAGGCGAGGTCGCTGGTTCGATCCTCGGCATCATTCCCCTAAAAAAGGAATTT

TCCATGTTCAAGAATCTGTAAAATCTCCCACTAAAACTAATACAATAACGATGAGAGCAAAGGGTTGCGT

ATACTGACTCCAATACCGAGAAATCGGGAGGAAAATTTGGAGGCAGAATCTCATTGGTGTTCCTGACATA

GTCTAGGAGACATATGTCAGGTTTAGTGTGGGAGAGTGTGGTGGAATTGGGGTTTAGAAGGGTCAATTCC

ACGTTGATCCGCGTTACGTAACATTTGGAA

>NODE_362_length_380_cov_10.665738

GCCCACGGGATAAGGAGGTCGCTGAACTTCGCCTTGCCGAAGAAAAGTGCGTTGCGGATGACGATCCTCG

CCATGAACTCGGCAGCGTGAGTGAACTGGAACTTGGTGCAGCAATCTCCCACAGCGTAGATGGCCTTGTT

GGTGGTCTGCAGCTGGTCGTTGACCTTAACGCCATCCTTGGCGTTGAACTCCACTCCAGCTTTTTCCAAG

CCGACGTTGTTGACATTGGGCTTTCGCCCGGTGGCCACGAGGAGCGCCTCGCACTCAATCACCTTCTCCT

GTCCCTTGTGTTCGACGACGACACGGATGGTGGGGAATTCCTCTTCGCAAGAGGCCGGTTCGTGCTCCAC

GCGGATGAACTTGAGGTTGAACAGCAGTTT

>NODE_363_length_379_cov_568.513966

CCGCTACCACTCTACCAAGAGCGGAGAGGAGATGACCTCGTTGGACGACTACGTGTCCAGGATGGACGAC

AGCCAGGCGGGCATCTACTACGTTACGGGGGAGTCGAAGATGGCGGTCGAGAACTCTCCGTTCCTTGAGA

AGCTCAAGAAGAAGGGCGTGGAGGTGCTCTTCATGGTGGACCCGATCGACGAGTACGCGGTGCAGCAGCT

GAAGGAGTTCGAGGGGAAGAAGCTCATCTGCGCGACGAAGGAGGGGATGAAGATCGACGAGTCGGACGAG

GAGTCGAAGAAGTACGAGGAGGCGAAGGCCGCGTCAGAGGGCTTGTGTCGAAAGACACAACCTGACCTTG

GAATTCTCGGTGCCAAGGAACTCCAGTCA

>NODE_364_length_379_cov_53.553073

ATGGGCTCATATGCGGTATTCGGTGGTCATGTGGGTGGCAACACCACCTGCCACCTAAAAAAATAATCAT

AGCCCGGGGTCCGGAAGTTTCGCGCGCGCAAAAGAGAAAAACAAAAAACAGAAAATCGAAAAAAGCAAAA

AAACGAGCGGAAGCGAAAACACGCCCGAAAAAGCGCAAAGAAGGCAAAATGCAACTCCTTGGGTAAAACG

AAAGCGAGATCACAGGACCAAAATGAGAGAAGCATGTCTTAGCTCAGTCGAGATAACCTTCTTCGAATGT

TTCGTCATCACCGCCCCGCGACGTGAGTAAATGTATGCGAGGGCGGTTAGAAGCGGCCTTAGTCTGTCTT

CACGGCGCTCGACGAATCAAAGAAGGAAA

>NODE_365_length_378_cov_4.647059

GTGTCAGAGAGAGGCATCCTATTCCCTGACCGCGGTTTTCGTATTGTTGGTGTATGACATTTCCTGGCCG

TTTTTCCCCCTATTGGTCCAACGTTTTTGAGAGGAAATTTCTGATTTTTGGACAGATAACATGTGCCTGG

AGGTATCAATCAGTGTCGACCTTCCCGTGCGCACTCGTAGATGTTTTTGGTATCCCCCCGTTTCTCTGCG

TAGTGCGCACAAGTAAATGTACTGTACGACTGGCCGGGTGGATGGGTAGTGTGTTGTCCTTGTTCTATGT

TCCTTGCCTGTTTTTGCATTTTTTGTAGGAGGAAAGGAATCGTGTTGTATCCAAAGATGATGCGATATGA

GACTTGTCGTTTTTTGGCACGTTAAGGC

>NODE_366_length_377_cov_7.339888

CAGTTCAAGGCCCCTTTCAAGGCCTACCGAGTCATCCTCGACAACAACGGAGAGGAGGAGGTGTACGAGT

GCGACTGCCTCATGAACGCTACCGGCCGTGTCCCTAACGTTTTCGGAGTGGGGCTGGACGAGGCTGGGGT

GGACTACAACAACCGACAGGGGGTGGTTATCGACGATTACTTCAGGACGACGTCCCCCAACATTTACGCG

TGCGGGGACTGCGCGTCTCCGTACAAGTTTACCCACGCGGCGGATTGGCAGGCGCGCGTGGCGATCAGGA

ACATGTTCCTAGGCGACAAGAACAAGCAGTCCGACCTCCTCATCCCATGGTGCACTTACACCGAGCCCGA

GGTGGCGCACGTGGGCAAGTACGTGGA

>NODE_367_length_377_cov_2.415730

CGGGCATGGTGTCCGTGGACGAAGAACTACGGGACAACTCGGTGCCCATGTGCCTCAGCGGGTTCGACGA

GAAAGGCCTGTTTGAACCAGACGGCGTGGGCGGAAAAGGAGGTTCGATGGACGTGTTCGATGGTGGCGAG

GTGGATGATTCGATGGCTCTCAAGCAGGAGTTCATGCACAAGTTTCACAATATCAGTCGCATCATGGACT

GCGTCGGCTGCGAAAAGTGCAAGCTTTGGGGAAAATTGGAAACCCTTGGGCTGGGCACGGCCTTGAAGAT

CATGCTCTTCTCTTCTGGGGCAGATGGGCAGACGGAGCTGGCTTCCCTGAGCAGAAACGAGATCATCGCG

CTTGTCAACACCGCGGCGCAGCTGGCC

>NODE_368_length_377_cov_2.101124

GCGGGAGTGATGACGGTAGCGAGAAGATTGATGTAGCTGTCCACCGGATCGGTGTTCAAGGAGCTTAAAC

TCAGTGTTCGTGTGTACTATGTTCATGACCTGTATGTTTCTGCCTCTTCGGGAAGGCACATATATGCCTA

AAAACACTCTCTTTTAATACCCGTCGAAGATGTGGGCTGGCACATTAGCTTTGGCAGGAGCTTAAGCTAT

AGTTGTAAAAGTCGATAAAACTATACTTATCAAGTACCAAAGGTTTGCTTTATATATAGCGCTTGCTACG

AAGATTGATAGGACGAACTGGTTGACTCTGGTATATTTCGCTGTATTCCCCTGTCATTGGTACTATGCAT

CGGACAACTATAGGTACCGAAAGTCAT

>NODE_369_length_376_cov_16.791549

ATTACAGGTGGCGTTGAGTAAACACCAGGTTGATGCCCATAGATAGGTTTCTACAGGCATACGATGGCGT

TCCGGTCTCGGGGACCCATCCTCTAACCTCCATTGCTTTCAGCAGAGAGCTCCGTACAGCTCGACCCCTG

CCCTCGCAATGAACCCTCTTCGCAGCGTTGGTCTCGGCAACCGCTACGCGTCTAAGTATTAACCCCTGGT

TCCCGCCGCGCTGTAACACCCCCCCTGTCCGACGCATAACTATCGTCGGCACAAAGCGGGCATCCCGCTG

ACAGCGTGGGCGCCTTCCTCCTCCATCCTTCATCGTCGGCCCGGCGTTGTAACACCCTCTCTGTCCGGCG

CATTATCGTCGTGGCAGGTTGGGCTT

>NODE_370_length_376_cov_4.104225

CATGAGGACGATTGATACCTGCCGGCGTTTCGAGGCGCTGACGGGAAGCTGTACGAGTACGCTCGCTGTA

GCTTCGGGTTGAAGACAGTGCCATCCGCGTTTGCCAACTACGTGGGAGGGCGACTGTTACCAGTCAAGGA

TAAAGGAATCAAGAACTGGTTGGACGACATCGCCATCCCGTCGAAAACCCTTGAAGATCATTGGGGGCTG

CTCCGAGAAACGCTAGAGTGTCTACGACAAGGACGTCTAACCGTCAATCTCCAGAAATCGCATTTCTGTC

AGTCGGTCATGGAGTTCGTCGGCACGATCGTGGATAGGCTTGGTACCAGGCCGGCCCCGAGCAAGATTGA

TGCTATCACACAGCTGTCAAGGCCGA

>NODE_371_length_376_cov_3.140845

CGGCCCACTATCCCCGGAACTGCACCCAAATGGCTTCGCGAAAATTGCCCTCTATTGGGGTGGTTTTTCT

TCTCTGATGATCCCTCGGTTACTGTGGTGAACAGTTGTTGTGCGCCGGAAAACTCTGATTTTTTCCTGTA

CGTGAACGGAATGTGTAGCACCAGGCTTCTGGCTCTCGACACGGGCAGGGTGCTGGCAGACATGTTTCAC

AGGGATATCACGGTGGTACACAACCCCACGGATTCTGTCATGGTGGACCTATTCGAGTGTGTCATCGGGA

AGCTGTGGACGGGGAAAAGCTGGTCGACGACCGTCCCCTGTGCTTGGGCGTTCGACGAGATCAAGCGTGC

TCTGAAAGACAGAAAAAAGACCAAGG

>NODE_372_length_375_cov_13.703390

CCTCGAGCCCACTGTCGACACCATTGATACTGAGGGAGGGGTCGTCTTGTCGGTAAACCGGCCAGAAGAT

TTCTCCCCATGAAGGATCGTAGTTTTCAAACGGACTGAGCTCGGTTGCCTCCGAGTCGATGGATCCCCAC

ACTCGTCTGTCTCCTCTCTAGCTCCGAACGAGGTGCTCAGCACCCGTTCCTCCCGAAGATGAGATTTAAG

TGCTTAATGTGTCAAAGGTTATGACTCGAGATTGGGTGCTCATATGTCTTGCACTGGGTCTGTGAGGCCG

TGTTCGACCTGTGTGGCTCCTGAAACTGATGTTATCTACCTTTTGAGCGAACAATAAAACGCACGCAAAA

TACTAATACTCTGAGGGAGTGGAAT

>NODE_373_length_375_cov_6.485876

CCTCGCCCTGCTAACCCCTCCACTGCTTCTCGCGACCCTGGTCTGTCTGTCGCTGCTGTCCACCATGATG

CAGAAGGAGTAGATGTAAGCTTGCGCGAACGTGTCGATCTGAAACCTAGACACGAGGCCCTTGTCGTAGC

TTTTACTGATTGTCTCCCTGTACAGGATTCTACCGTTGTAGTAGAACCTCGAATTATGTCTGAGCTTGAG

TTTTCAGAATCCGGTAGTTCGTCGGTGTTTGAGAAAATTATTGTCGCTCGCACCCTCGCTACGTGGTATG

CTGCTGACGGTTCAGTAGCAGTGCAACTTGCGAACCCGTCCTCTGATGGCGTTGCACTTCCGATTGGTTT

ATGTCTTGGTCAGTTATTTACTGTT

>NODE_374_length_374_cov_16.144476

ACCGAAAACGTCCATAGGTTAATAAATTGAAATAAAGAGTCCAAAAAAAAACAAATAAAAACAAAAAAAA

CAAATCCAAATCTCTTAAGACAACCAACTACATTTTAAGTTCAAATGTTCAAATCGTTACGTTGCAGTTA

GTCAGCCATATCAGAAACCAGGTATGTATCGTATCATATCCGTTTTATAAGAACCCTTTATTCCGATATT

CCAAATATATAGTGTCTGGAAGAGTTCCGGCGTTATTGGCCAAAATACCACCAGATACACTGTTTGCCAA

AGTCCACCTACATATCACGCGCATGCACACTCACTCTGCTTTGTATCAGGCATTTATGTACCCTACGTAC

TGTATACGAATAAGCAAACTATAT

>NODE_375_length_374_cov_10.524079

CGTTCATACATACACCCCACCCTAAGTAAACTTGCACGGGGTGCAGGAATATCACAATTTCTGCGCGTGC

CATTTGAAGGGTAGTACACTATGACAATGGCTAGAAGTGAGCAATTTCCACCTTTTGTCTAGGCTCCAAA

GCTCAACATTCGTCAGAGCAACGCCTTTATATCCTTTACTTGACGGTATACTTGCTACTACAACACACGT

TCGGTTATGACCCAATCCATTTCCATTTCCCTACAGAGATGGTCGATCAGTGCCAACGGCTCGTCTGGTT

GGGTGAAGTCAGCGATCTCGCCATTGTGTCAAGCGATCTCGCCATTGTGTATATAGCTAGGAACGCGCAA

CTTGGCCCGTGCACCACTCACAGC

>NODE_376_length_374_cov_8.651558

GTGGTGGTGGTGGTGGTGGTGGCGGTGGTGGTGGTGGGTACACGGAGAACATGAATTGAAGGAAGGACAT

TAAAGAAAAAAAAAAGGTCCCCCCCCCATACATCATTCTTGAGTCCATTCTCTCGTGTTCTTTGTACTCC

ACAAAATGAGTCCCTGCTTCATACTCCCACCCTCCCTCTCCTCCTAATATTCTCGGTTTGTGTTTTCCCG

CTGTTGCTGTAGGTAGCCAACGAAAAATGCGCGACAATCCACAACAAGAAGAACCGCTCGACGACAAAAC

AAAAACAAAACGAGGTATATTCGTATGGTATTTATTCATTTCTTGCCAACAAACAGTTGTAACACTCTCT

CGTACATCCCAGTCGTTGGAAGCG

>NODE_377_length_373_cov_17.946023

CCTTGTAACCCGAGAATTCCATTCGATGCCAACCGACCCATGATGATCGCCAATACCCAATCGACCAACA

ACCGGCCTACCAACAAGCCACGCGTCTTCTACCACAAGCTAATCAAGTTCATACCACCGAACAAAGCATC

AATGAAACCCTGAGGCGCAAGCACGAACACACGCAGCTGTCGTCGAGCGAAGCAACCGACAGGAGACAAA

GAACCAATCCATATGCAGCACCAACTCTCGACAAATCCATGCATGTACCAATCAATACCAAATAATTGCG

TTCAAATATTGTTCAAAATCACACACAGCTCGGGCATGAGTGACATTAATCTAGTCCTCACAGCCACACA

ATGCTCTTTGGAGGCGGGGCGAG

>NODE_378_length_373_cov_10.576705

GGAACTTCTGTGAGTTCTGTACGCCTGTGTCACAATACCCGGAACTTCTGTGAGTTCTGTAACACTTTCA

TACCCGTACCCGGAACTTCTGTAAGTTCTGTAACACTCCCATACCCGTACCCTAAATCCACAAACCCTAC

AGAACCTGACCTTGAACCGATCCACCCACCATCCAATCAGTATACAGTAGCACCAGTACGTACAGCAATA

GTAGTAGCCCTTATATAACCCATCTATCTACAGCAGCAATAGCAGGGACAGCAGCAGTCCAGCAGTACGA

CTCGTACGTACGTTCAGCGGACACAAACAAAACGCCGGCATGGGTGCGCAGCCCTTGTACGTACAGCAGT

GTAAACAAAACGCCGGCATGAAA

>NODE_379_length_372_cov_14.074074

AGCCAAGGGTGGATTAAAGGGCCGTCGTCGACAATTTGAAACCGATCTGTACAGTGTTCTGGATGTTGCA

ACTGCTCCGTCCTGTACCCATGTGCTGTTGATGGCTTATATGTATTATCCAGGGAGTAGTTCGTGGAGTA

CCTTCCGCCCTCGGGTATGCGATGGATTAGTTGTTTATCCCGTCCCTTTCGCAACACAAGCGTTTGGTGC

ATCCGCTGTCCGCGCAAACAGTGGTGTCAGCGTAGTCCACTAGGCTGTACTTATTCGGGCACTTGAAGCT

CGAGCAGAAGACGTCGTCCTTTTCGCAGCAGAGGTCTTTGGTGCACCCGGTGTACGCGCAGACAGTGGTG

GCAGCGTAGTCCACTAGGCTGT

>NODE_380_length_372_cov_7.344729

TTGGCTTGGTTTTTCAATGCATGATCAATGTGTTTGAAAATTACTGCTGATTTAAACTTGATCCTACAGC

AGTACATGATATTGATGTGTGAAAATCAGTGCTGATGCGTTGTAATGGTAAAAGTTATTTTTTTGTGGTC

ATGGGGAAGAAAGATCAACCAGAAATCGGTACTGACGTTGTTTTTGTCACTATTGTTGTGTTGGGAAGAA

TCACGAGGGGGTGTTTGATGGCATGATCCTACCCACAGACATAGTGCATAATGATGTGGTTCGTTATCTA

ACGATTTGACTTGGTTCCTGGCGCTGAAACTAACCACCCGATGCACCCTTTGCTCAACGAGTCAAACTGC

ACCCTCACTCTGCGGATTTATG

>NODE_381_length_372_cov_6.643875

GGAGCCAGGTCAGGCGGCACACCACGTAGATGTCGCCAATGGAGAGCTTGCCGCCGGAGAAGTAGCTGCC

GCCCGCCGCCTTGAGCGCCCTCTCCAGGTTGGACAGCCAACTAGGAAGGAAGGTGTCCGCCATCGCCTTC

CGCATCTCCAGCTTTTTGGTAGGGTCTTGCTCCTTCATCGTGGGCAGCAGGTGGAAGTGCATGTCGATCA

CAGCCTCCATTGCCGCGTCCGCGAAAGCCGCTTCGATCGGGTCTTCCGAGTACAGGCCTCCGATCTTTCC

GATGTATCTGATCTGGGCGAGGCTCTGGGGGATGAGAGTTCCGTCCACCTCCAGGGTGGGCAGCTGGTTG

TACGGGAGGCTGGGCTTCATGG

>NODE_382_length_371_cov_5.585714

CTGGAGTTCCTTGGCACCCGGTAATTCCAATGCGTCCAATGTCCCCCCAAAGTCATCCATCAATAAGCAC

ATTGTGGATTCAAGAATTTTGTTCAGCAGCAGGAAAACCTTGAATATAGTTTGTATCGACAGCAACCAAA

ACGCGTGTCAAGAGCGAAAACACTGGTCGCTACAGAAAAACAAGCCTTTCAAAACCGCGCCAAAAACCAG

CCCCGGCTCCGCTCCCCGCCGTCATGACAGCCGGCTACGGATGGCTGCTCTCTCGAAGACGAAAAAACAG

CTGTTTACGGAACCAAGGAGTTCAACGCAAAAGGCGTGCATGTTCGATGCGAAGAACAAGACGATGTCGA

CCACCGTTGTCCCTCCCCTCG

>NODE_383_length_370_cov_18.782235

TCCAAGGAAAGGGCAGCCTAAGCTGCCCTAATTTAGTCACACTATCAAATCCATGATAGAAGTTGCTCCA

TGCAATAAATCCATTTGACTACCGCCATACTAGGCATCCTCTTCCCAAATCCTTTGAGCTTCTTCCTTGT

TTCATATCCTTATCCGACAAACATCCAGTTCATCAATCCAACATCCTGTTGGTTTCTTCCATGACACATC

CATATTTAATAAACATCCTGCTTATTAACTGCTCCCTATCCTTGGAAGGTGTCCTTTCTACATCCTGTAA

TGTCTTCCGTGTTCTTCCTTAACGATAAACATCCTGCTTATCTAACTCCAATCCTTGAAGTGTCCTTATT

ACATCCTGTAATGTCTTCCG

>NODE_384_length_370_cov_4.111748

TGCTGCTGCTGCTGCTGCTGCGCACCAGTTATTGCTGCTGTGTACCAGGTATATTGGAAAGCACGACACA

GCAGATATAGCAGCTACTGCTGTGACGTACGTATGGATACAGCGGCAGCAGCAGTATAAATAAGCAGGTT

GTTGTTGCGCTGTGCGCTGCGCCTATACTCTGTACAATCAACATCCAACACCCGGCTCGAAAGCTCCAAT

CAGCACACATCATCACCACCATCATGCATGCATGTCTTCGTGAAGAACACACACAGCAGGAAAGACAGAG

ATGAAGTGAAAAATGGGAGATGGAATGAAAGCAGAACGATGATGAGAGATGGGAGAAATGGAACAAAAAA

TGGCTTGAATTTCCGGCCGA

>NODE_385_length_369_cov_10.563218

CACACCGACTTCCTGGCCAAGAACCCGGCCCCCAAGCCCACCGAGGCTTAAGCGTTAAACCTTAAACTTG

GAGTGGAGATTTGTCTGTCGGTAGCGGGGAGGAAAGCGTAAGGGCAAGTACGCTACGGTGCGTAGGTGTC

GACCTCCCATATAGCTTCCGACGCCCCTTTGTCGCATCTTTCGCGGGGTTTCCTAGCATGTTATGTGCTT

TATTCCGCGGGGTGTGTGGGTAGAGGGGTTGGTTGAGCGGCTGCGTGGCGATGTTATAGAGACAACGTGA

GGGTTTTCGGTGGCGTTTTTTCGGCATTTTCGAGAACGGGAAAGTTTTTTTCCGGCGAGGTTTGATTGGC

GGTTGGGAGCAGGGGGGGG

>NODE_386_length_369_cov_9.241379

GAAGGAGTCCACCCTCCACCTGGTGCTTCGCCTTCGCGGTGGCATGTAAAAAGTATCTCTTTGTTGAGGC

GCTTTTTGCTTTTAATTCCTTTGGGAGCTTTGTTGCTGGGGTGGCCACATACGGATCGGTTTTGCTGCCG

TTTAGTTGACGCTGGAGTCAAGGGTAGGCAGGGGTGGGACGTCTGGAGGCTTGTAGGCAGGGTTTGGCTG

GAGCGTGTTGTTAATAAGAGGGTTGGTAGTTTTGCCTTGAATGTTGTGTCGACGCGCGTTACGTCGCATT

GCTTGGGCTTTTGAGCTCGAGTGCTGCACGTTTTGCGAAACTGACGCTTGTGAACTAATATATATGATGT

AAAACAAAAAAAAATTTTG

>NODE_387_length_369_cov_4.204023

TCGAGCATCACTGCCAATCTATTGAGGGCTCTGCGTTGATAGTTGGCCACTGTGAGAGTTTGAAGAATAT

CAAGAAGTCACGGAGGGAGTCAGATTCGACATCTAAGCCGCATCATCAGCAGGGACGCAGACGTCGTCAC

CGGTGAAGCCAATCAGAGTCGCTCCGTCCGAGACCTCTCCGAAGGTGCAGCCATTGCCCAAATTTCCGCC

GATGACGGCGTTCCTGGAGATGACCCTTTCCGGTTTCTCGAATGTGAGGGAGCCACCAGCCCTGATGGCG

AAGGCCGAGCCCGATTCGGCGGCGTTTCCGTCAAAAGTCACGCCCTTGATGAAGGTGACATCGCCGCCTT

CCACGTTAACGCCTCCACC

>NODE_388_length_368_cov_7.994236

GAGCAAAGTCCTGCGAATCACGTGCTTGCATTGCACACACCCTGCATACTGCTACTGAAGTAGCCGCAAC

TGTCCGTCCGAGCATGTCGGAGGTGAGGTGGGCTGTCGAATGGTTTAGTCGTCTCCGCGGACAACGGCGC

CCGCGATGCGGTCAAGATCCTGCTGGCCAACGCGAGAGATCTGGCGTCCACCGTTCTCCATCTTCTCCAC

AACCTTGAGCGACTCAAGCTGCTGGAGCACGTGGCGGATGATTCCCGTGGAAGCCGGCTTGTGGGTGTTG

GTCATGGTCCCCCTGCGTGCCTTCGTGCCGAAAGCCTTGCTCAGTCCGCCAACGCCGCGCCCGGGGCGAA

GGTACACCTTGCGGGCGA

>NODE_389_length_366_cov_1826.344928

GCCTTCGCCATCGGTGGTCCTCCCAATCTCTACGCATTTCACCGCTCCACTGGAAATTCCCTTTACCCCT

ACTATCCTCAAGTCTAGTAGTTTCTATTACATTTTTGAAGTTGAGCTTCAAGATTTAACAATAGACTTAC

TAAACCACCTACAAACGCTTTACGCCCAGTGATTCCGGATAACACTTGCATCCTCCGTCTTACCGCGGCT

GCTGGCACGGAGTTAGCCGATGCTTATTCGTCAAGTAACGTCAGAACTTCTTCCTTGACAAAAGAGGTTT

ACAACCCTGTGGGCTTTCTTCCCTCACGCGGTATTGCTCTGTCAGGCTTTCGCCCATTGCAGAAAATTCC

CCACTGCTGCCTCCCG

>NODE_390_length_365_cov_120.517442

CAGTTTGTATTCGTACTGAAATTCAAAATCAAGTGAAACAACAAAAAAAAGAAAAAACTTGGGGCTGATT

CAGGATTCGACGAGATTCACGAAACCCAAGGTGCATGCCGAGGTGGCGGTTGGCCTCGTAAAAAGCCGCA

AAACTATAATTGCTAACGACGAAACGTTCGCACTAGCCGCTTAGGCTAGCCATCGCCTCAAAAATTCTTC

TGTTGTTTAGAGGATCGATGGTCACATTAAACGGAATAGCGAGGGAAGCATGCTTAAGGCTGAACCGCGA

AATAGTATTAAGCTCACCATGACGAAGCCTGTCGATTGGCGTCCAATTGGTTAAATAAATAATCGACTAA

GCATGTAGTACCGAG

>NODE_391_length_365_cov_4.720930

GTTGTCCTCCCAGTAACTGAACAGGCTGAGGGACCTTTCCAGGAACCTCGGCTCTCCCAGGTAGCGGTGG

ATCATAGCGTAACCGTAGATGGCCCACGTGTGTCCCCTCGTCCAAGTCGAGTTGTCGACCCAACCCTGGT

ACGTTCCTTGTTCCACGAGGTCGCCAGTGTCGAGGTCGTAGTCGGCGACGTGGAACGTGCTGTTGTCGTC

GCGCACGATGTCCTTCCAGGTCGTGTCGGCGTGCCCAACGACGTCTTCCGCGTACGACTCGTTCCCTCCG

TTAACGCTGGCCCACAGCATGATCTCCATGTTCATGATCATGTCCACGTTCACCGCCCGCAGCCACTCCG

GCCTGGAGCTAGGCT

>NODE_392_length_365_cov_4.232558

TCCTCAACCTTATTCAGCACCCTCGCTGTCATCGGTGTCTGGGTCGTCGTCCTTGTACGCTTGAGTGGCC

TCATCATCCTCAACCTTATTCAGCACCTTCTTGGCGAGCGGAGCGACGAACTTCCTCTTGAGCTCCAGGC

CACTCGCGAGGGTGGGGTCCCGGCAAGCATCTTGGAACCGCTTCTCGTTGACGAGTCGGTCCTCCACGTA

GTCGCAGTTGTTCAGCAAAATGTTGATTAACTTGTGCAGTCGGCCGCCCCTGGCAACTCGCTTGTTTGGG

ATACGGCACTCCTCGGCACGACACCAGGCAATGGTAACGTTGACATCCTTCTTACTCCACAAATCGAACT

CGAACTGGAGAGCGG

>NODE_393_length_364_cov_4.650146

GTGCTTCACCTTCTTACGGTCGCGCTCGGGCTCGTTGAGGGCCTTGAGGATAAGCGACGAAGCTGCGGGG

ATCACCGACACCGTCGCCTGACGGTTGACGATGGTGAGCTTGACGGTGATGGGGAGACCCTTCCACTCCT

TGGTGTTCTTCTGGATGTCCTCACCGATCTTCTTAGGCGAGAGACCCAGGGGACCAAGCTTGGGGGCCAG

AGCGGCCGTGTTTCCTACCTCGCCCCCGATAGCCCGAAGGATCACGTACTGCGCCTCGCCGGTCGGCGGA

GGAGGGCCCATTGTTTATAGAGGATGGATTGTCCCGCGAGGGATGGCAGAGGCAAGTTTGGGATGAGAAG

CTGAGGTAGCACTA

>NODE_394_length_363_cov_3.675439

GCCACTATGAAGCGGCCAAGTATTAGGGGAGCAACGCTAGTAAACGCAGCAAGGAGAGCCAATCAGAACC

GATGCACTAAATAATTCGCCAGAAAAAATATCCGAATGTGTAGACCTAAGTCGACTAGTTCGACGTGGAC

GAAATGTGTCCCTACCGTATAGCACGCACAAAAGTCAGCTATGGTAGCTACATGAGAGCGGAATCGGTCG

CGATAAGAAACGCGTTGGTAGGCGTTGGAAATGCGAAGACACAACAGAGGAAACAATGACGAAGATACCA

CAACAATTGAAGTCCATTGCTGTCCACTTACTGTGGAACGGACGCACCAACCGTGGGGACGGTAATGTGG

TGGTCTGGTAGCC

>NODE_395_length_361_cov_11.629412

AGCAAGCTACGTCCCCGCCCGACAAAGCAAGCACAAAGCAATCATGGCTGAACCTCAGGAGACCCCGGCT

GTCGAGGCTGAGACCTTCGCTTTCTCGGCCGACATCAACCAGCTCCTCTCCCTGATCATCAACACCTTTT

ACTCCAACAAGGAGATCTTCCTTCGCGAGCTCATCTCGAACTCGAGCGATGCGTTGGACAAGATCCGATA

CCAGTCCCTCACGGACAAGGCTGTATTGGATGCGGAGCCCAACATGGAGATCCGGATCATCCCCGACAAG

GCTAACAACACCTTGACGATCGAGGACTCCGGAATCGGCATGACCAAGGCCGACCTGGTAAACAACTTGG

GGACCTGGAAT

>NODE_396_length_360_cov_14.510324

CTATCGAAAGGGAATCAGGTTAATATTCCTGAACTACGTACATAAAGTGAAGGGGGGACATATTGGGATA

TAGGAGATTCTTAATGAATTGAATTACGAACGTAAAGTCTGATATATAGGAAAATCCGTGTATCGTGAAG

GATTGAGCTAAGTTAAGTTGATATCTTTAGCTTGCTAAAGATGACGACCGGCGAACGGGTGCGTAACGCG

TATAGAATCTACCTTATACTGGGGGATAGCCTTTGGAAACGAAGATTAATACCCCATAGTATTATTTTTT

GGCATCATTAAATAATTAAAGTTTCGGCGGTATAAGATGACTATGCGTTCTATTAGTTTGTTGGTAAGGT

AACGGCTTAC

>NODE_397_length_360_cov_13.799410

AGGTTGGGTTGGTGGATGCAATATAGCTCCGCGGTCTATGGTGCTCAGCTCCTCATATACGGCGCCCGGT

TCGATTCCACGCGGTGGCTGCGATTTCTTTTTTTTTTGCAATCGGCGATTTTACGAATTTGAGGAATTTT

GAACTGGAGCGGAAGACTCTACTGGAGGAGAGAGAGAGAGAGAACGGAATGAATGGTTGGTTGGTCGGTG

TCTGTTGAGGTTTGGGTTTGGTTGGAGGAGTGTGAGGGAAAAGGTGGTGATCGATTGAGAGGAAATAAAA

GGAAGGACAAGAGAGAGTGTGTGTGAGAGAGAGGAGGGAAGGGATGGTAAGTACTATAATACTAAACAAG

GCTCCTCTTC

>NODE_398_length_360_cov_5.734513

ACTGGAACTCGGCAAGATACCGGTGGTAGTCGCCCTTCATCTTGTAGTAGAACACCTTGCCCTCCTCCGA

GGTGGAGTTGGGGATGAGCTCGGCCTCAATGATCTTCAAGATGTCGTCGCAGATGTCCGTGAGTTCCGTC

TCGATCTTCGACTTGTAGTCGCGGATCAGGGGAAGCTGGTCGCTCACGGTGTCCTTGCCCTCGATGGAGC

TGATCACCCTCCACGACGCACGTCGGGAGCCGATAACGTTCTTGTATGCGACGGAGAGAAGGTTGCGCTC

CTCCACCGAAAGCTCTTTGGGCTGCTGCGCCACGGCCTTCATGTGGTCGACCATCTCATCGAAACGTTCG

GCCTGCTCAG

>NODE_399_length_359_cov_12.408284

GACCTTGCTGATCGCGCGAAGCCCTTGGTCTCCTTGGAACCCATCGCTCGAATCAGAGTCTTGACGCGCC

CAACCACATAACTTGTCGAAGAAACTCCTCGATTTGGCTGGTTTCACTTCCACGCCGTCACCGTCCACAG

CGGAAGCGCCAGTCAGGATAGTTTCCTCTTGATCGCAAAACTTCCGCAAAACAGATTTGCTGTGCTCGTC

CGACCATCTCTCGTTCCACTTGGGCGGCTGCTGTTCAGACATGGTCCTGGCTGTATTATTCGACGAGCGT

CGCTGCTTGCGAAACCGTTCCTTTTTTCTTTATCACGCACCTGGTGCAGCAGTACAGCAGCAACACAGCA

CAGCACATA

>NODE_400_length_359_cov_5.248521

CCGTACGTGATGTCCAAAACGCTGGCGGAGATGGCGGCTTGGGAACTGGTGAAGGATGAGGAGAACGAGG

GGTCTGTCGAGCTTGCCGTCGTCAACCCGAGCCTGGTGATTGGCCCCAGCCTAACCAAAAACACGGGCAC

CAGCATCGACATTTGCATCAGTATATATGGCCACGTTTAGGTTCCAGGTCAGCTCCCAGGAGCTTTTGCA

TTTAACGACCTTCTGATAAAGCCGTGATCACAGTCGATTTTCCCTCCCTCCCCCTGTGTATGCATTCGTT

TCAATCCGTACAGGGTCAAGCCTTCCCGCTGCTTCGTAATTTGCATCGAGGTTTGCGAATTCAAGCGTTC

GCGCTCTAA

>NODE_401_length_358_cov_10.270030

GTTTTTTTATTAATTATTTTACAAATCTATAATTCATCATTAATTTTAAATGATGTTGATGTAGATTGGC

AAAGTTTAGGTAATTAACTTAAATCGTATAGGTAATGTTTATTGTATTATAAATAAAGTAAAAACTATGG

AAACATCTACAATTTTAATAATTTTTGTATCTAGCTTATTAATAGGGATAACTGCGTATTCAACTTATAC

GGCGTTTGGGCCGGGGTCAAAATTTTTGAGAGATCCCTTTGAGGAACATGAAGATTAATACTTATAAATC

AACTTTATGGTTTATCTTTTCTATCTATATAAATAACATATAGGTAATGGTTGTAATTTAGTTTTCTCAT

CAATGCTG

>NODE_402_length_358_cov_7.326409

TTAAATAACCAGATACCATTGCAATTACAGCAAAACAGGTTAATTCTGTTAGTTTATTGGTTTTACTCGA

CGTATTGGGTAATCCTCGGGGTTTAAAGTATTTACGAAATTTTGTTTTTTTAGGTTGCATTTTTATTGTT

TTCCCAGTTACATATCCAAATTTTTAATCCTACACTCCCATAACCTGTTTGTGTTTGTGCATATTCGGCC

TGTATATTTTTATTTAACTGACTAATAGGCATTTGTCCCATTTGATATTTCCAAACCCTGCTCCTTCCTT

TTGCTTTATGCGTAAATCTCCCTTTTATTTGTATTTTTTTTGTTGATGAGTATTTGATAAAGGTTCTTAT

ATTATTGT

>NODE_403_length_358_cov_3.400593

ATTCATGGTGTTTACTACTATAGCTTTTTACGCCTGACATCGTACCCCTAAACGAGTAGTCTACACCAAT

AACCTATCAAGAGAGCCATACAGAGGGAAACATTGACGCATTCGGTACAGAGCTTTGCCTTGTTGCGCGT

TTCAGCATGGTTCTTCGTTTAGAGCAGGCCCTTCCTGCTGTGCCTTCAACCCCTGCACAAGCGCTTTGGC

TCGTTAATCTGGAAAACATGTGCAGTACGTAGGCCCTTCTTGTTCCAACCTTCGGAGTACCCCAAACCAA

CACAATCGAAATCGAATCCCCTCTTTAAGTCTTCGAGACTGGGACGTAGAGATATGCAACGTCCTAGAGC

GAGGAGTT

>NODE_404_length_357_cov_8.288690

ACAGGGCAGTCATGTTCTTATTATGTGGTCTCCCCCGTGCCTCCAGGGCAAGAAAAGTAACCGCCGCAGG

ACGGAAATAGACGCTAGCCACTCAGCAAGCGGCGCCATTGCGCAACTACAACCGGGCAGAGCCGCGACTA

CGACCTCGCAAGAGTTACATTCCGGGAAATTTCCGAAAACAACAAAGCGAGGCTTGTTGCACTCGGGCCC

CGGCTATAAGCTCCCTCCCCTCCAGTGCGTGCGGCTTCGACCCATAAAACCTCTCGAAGGCGATTGGCTT

AGAACATACGGAACTGCTTCCGCACCGACACCGCCGTGTTGATGCACGCCTCGTGCTCGAGGACGTTGAA

GCGCACC

>NODE_405_length_356_cov_15.570149

ACATTTACAATACTTAAGTAGCATTAATTTTTTAATCTTTTAAATTATCAAAATAGAATTTCTTTATAAG

AATTATTAGCTATGTCACATTCTGTAAATATTTATGATACTTGTATTGGTTGCACACAATGTGTTCGTGC

TTGTCCTACCGACGTATTAGAGATGGTTCCTTGGGATGGTTGTAAAGCAGGACAAATTGCTTCAGCTCCT

CGTACAGAAGATTGTGTTGGTTGTAAACGTTGTGAAACAGCTTGCCCAACAGATTTTTTAAGTGTTCGTG

TTTATTTAGCCGCTGAAACAACGCGTAGTATGGGATTAGCATACTAACAACATAATAACCAAGTATAGTT

TATATA

>NODE_406_length_355_cov_15.482036

CTTGGCACCCGAGTTTTCCACCTTGCGCCTTTAAAATCTACAACGCGGGGACCTCGATATATAGCGGCAA

AATGCGGGGGTTCTACATACCTGTTGTCGTGTTTCATCCCGCTCACTCACTCTCATCCACGCCGGCGGGG

GAAACGAGATACAGTAGGTGGGGCTATATAAGGCGGTATATATAGTTCCCCGAAATTGGGGCCGCCTTAT

ATCCAGTTGCAGCCTTGTTTAATACGTTATATGAGTGCAATTGTCCGGTAGTCCCGTGCTACGATAGATA

CAGTACCCTTTTGGCTGAACGCTTCTTGAATGACGGTGCTCCTTAACCCCTAAGGAACTGGTCCCACGAA

TTTGG

>NODE_407_length_355_cov_13.299401

AGCGGACAGTACAAATCCTGTTGTAATTTTCTTCCATTATAACCTTGACGAGACGATCCTGATCAACGAC

CGAGCAAAAATCCAGCAGACGACTACCACAGCTTTTACCGAAGGGTTATTTCCTTCAAGCGACAGCCAGT

TCGATCAACCCTACCTACGGAAACGACAAACCCATAGCAAAACAACGCCGCGGAGAACTTGGGAACACCA

AAAAAACGTGCTCGCGTGAGCCAATTTCGGTGCAAATTCTCCGCAAACGGTGCGCGGGTGCTACACGGCA

GTCACAGCCCCATTTCAAAAACTCGGTCCTCCAAAAATTTCGACCCTCTTGGGTACAATATAAGCTTAGA

AGCGG

>NODE_408_length_354_cov_3.360360

GTGGTACTATTGGTCACCCTGATGGTATTCAATCCGGTGCGACAGCAAACCGTGTTGCTTTAGAATCTAT

GGTTCTAGCTCGTAACGAAGGTCGTGACTATGTTGGAGAAGGTCCTGAAATCTTACGTAACGCAGCGACT

ACTTGTGGTCCTTTAAAAGCAGCGTTAGATTTATGGAAAGATATTACTTTCGATTACACTTCAACAGATA

CACCTGATTTCGTTGAAGTTGCAACTGAAAGCAAATAGTATACTGAAAACAAATTTATAAGAGATTAATA

TTAAATAATTTTAACTAATCAATAATCTGGTACAAAATAACTACATAAAAACAAAAAACTATTAACAACC

GACG

>NODE_409_length_353_cov_10.918675

TAGTTGGTTTTCTAGATTTATGTGTTAGTTGTGTTGCTTGTTTCAGTATTTTTTGTATCAATGATTTTTT

TTAGTTTAACCCCTCGTGGTGGAGGCAAGAAACCCGTGCCAGGCCAACAATCCGAAAGACGAAACGCCAA

CGCGGCGGACTACATAAAATCCGACGTCGTCAAACGTTCGTCATTCCAGCAAGCCAGCTACGACAGTCGG

CTACATCAGTAGTCGCCGCCGCCGGCCTCCTCTTCGCCGAACATGCCGCCGGCTCCGCCCATGTCAACCT

CCTCCTCTTCCTCCTCCTTAGGCTTGGCGGGCGACATTTTACCGGCCTTAGCCTCCTGGGTGTCTCTGAT

CTT

>NODE_410_length_353_cov_10.900602

CTTATGACTTATTATTATATGTTGGTTTGAATTGAGTTAGTTGGGAACAGATATTTGTTTTTTTGTGTTG

TGTGCCTTGCCACGTCAGTTTCAGAGTCCTCGTGTGCCTCTCGGTTTTCAGCCCTAGGGTAGAGAGAGGT

GCGAGGAGTGTTGTTTCTTGGCGCTCTGAAAAACACTTGATAGTTTGTGCATGGTTTTCCCCGCTAGCTA

CCCCGAAGGCTACAGATCGTACAACCAGCGTTGACGAATTGCTGCATTTGGTGTAGTTCAGTCAATTTTG

TTTTGGCGGTTATATAGACCGCAGCAGTGAGTTGGTTTTTTATTTCGTGGTCTTTGGCGCTTGGAATTCT

CGG

>NODE_411_length_353_cov_6.177711

CCCCGGCGGCGCGTGAATTCGCGTGATGGGCTTCCCCGTCAAGAAGTTTCCCGAGTTCTGGTCCACACCC

GAGGCGTACGCGTTCGACGAGAGCATGTTGGAGATAGGAGATGACGAGCCTCCCCTCGAGGCCGTCTGCG

GAGTGCTGGGCTCCGTGTCGTCGACTTCCATCCACTCCTGCTTCCGCCGATTCCAGACCATCTTTTTGCC

CTTGGGTGTGGCGGGTGTGTCCGTGGTGCCCAAACAAAGAGTCGTGTTGCTCACGTGCGAGCGGCTCATG

GTTGCAAGAGATCCGCGACGCGTAGGGGTAGTTCGTACAGGTGCAGAGCGATTACGGGTGCTCGGTGTTG

GTC

>NODE_412_length_352_cov_3.694864

ACCAGGCAAACGAGAAGCAAAAGCGTAAAAAAGGGGGCACTGTAGCTGACACAAAAGGAGAGCAGGAGAG

ACAAGAGCAACGCATATACACCCGTCAGAACCATGGCATGTTGTCTAATCTGTGTTAACGATGACCCGGT

GCCGGGGTCCAAGCTGGACGGTTACTATGCTAACCCGAACCAGTTCCAGATTGGAATGATGGAGGCGCCC

TGCAAGAACCCCTGCTGGTGCCTTTTCGGATGCCTGTGCACGTGCTGTGCCCAGTTCATCGTGCGGAAGA

AAGCCCTCGATGGGAACCTCGAAAACTACCTGTGCTGCCAGGGGTACTATCCCATGTGCTGCATGATGGA

AT

>NODE_413_length_351_cov_13.239394

ACCTATCGCCAGACGAGACCTACACCCATACCCTCCCATATACTCTGTAGAATATAAATGTACCAGAGTT

GAAATAAAAGTCACTATAATAAGCATAACGACAGTTAAACTGTCAAAGCAAAAAGCCCAATCAACATGAA

AAGAATCAGACTCCAACCAAGGCATTAAATAAAGGTAAGTTGAACTTCCTCCTAAACCTACCTCATAAAA

AGCAAGACCACTTAGAAAAAAACTAAGGCCAATACAAGATATAGTTACTAAACCAGCACCGCGGCCTCCA

AGGAAACGACCAAAAAATCCTGAAACAAGAGAACCAAGTAGGGGTAGAAAAACTATGTTTAAATACATCT

T

>NODE_414_length_351_cov_7.333333

ATGGCGACAGGCAGCAACTCGCACTCTGACAAGAACACCGCCACCGTCTCGCCGATGGAGCACGACAACG

ACGAAGAAGCCAGCCCCGAGGCAACCAGGCAGCAGAAGCTCATCAAGGTGGACGGCAAGGTCCGCACCGA

TGTCAACTTCCCCGCTGGTTTCATGGACGTGGTGCAGATCCAGAAGTCGGGCGACCTGTTCCGTCTGCTC

TACAACACCAAGGGACGCTTCGTGCTCCACCGGATTTCGGAGGAGGAGAGCAAGTTCAAGCTCTGCCGTG

TCAACAAGGTGGAGATGTCGAAGAACAAGATCCCCTACCTGGTCACCCAAGCCGACTTCGTTGACAAGGG

T

>NODE_415_length_351_cov_7.290909

CTCCCACAACGATGAAGAACAGAAAAAAAAAAAGTCCCGGCTATACGCCTAGCATGGAATCTAGATTATC

ACTCCACGAGCAAGAACGACGGCCTTGAAATTTGAACGCTAAACTCAACCCCCCCCCCACGACTCGCACA

TGAGAAGTGGCATTCATTCCCCTGCCGGAAGCGACGACCTTTCCTCCGTACAGTCGGGCAAAACAAAAAC

AACGGTCGGTAGACATTAACCAATCAATCACTCGGACGAAACACTCTCCCCATCAACAAGAGACTTCTAT

TTAGTCCCGCCGCTTGATGCAAGCAGCTCGTTGGTTTAGAAGCTTCGTATGGTACATAGAAGCCTCGAAA

A

>NODE_416_length_351_cov_5.433333

CGCACTTCTCGGCGGGAGCAGCACAGCAGTTCAACAACTTGGTCGGCGTGTTCTTAACGTTTGCCTCCTG

GGCTTCTCTGGTCGCCGGTTATACTGTTGGCGCCTACGTCTTGCTGGCTGTGCTTGCTGGCACGATGTTC

CTTCAGGGGTTCCTCAACTTCTGCGCGGGATCCTTCGTCCTCGACTACATGATGAAGTGGGGTTTGGTCA

GCAAGTCCGTCTACCGCGTGCACATCAAGACGAGGGCCAACCTTGAGCGCGCTTGGGCGAAGAGCGGCGC

CCCCTCCGAGAAGGGAGCGTTGACTGAAGCCGTGACGTACCCCACCGTGGGCCTTCCTCCTAACCCCACA

G

>NODE_417_length_351_cov_4.221212

AAATTTTTTTTTTTTTCAATTTCACAATTTTATTAACCTGGTTGCCGTCCGACAATGTGTAGCATTGCCA

TAACCCTCACATACCTTGACTAAAAGCAAAAGTACACCAACTATTCGTTGGGGGAACGTTTCCGCTCCTG

TTAAGACGGACGAGAAAACACAGAACACCGCTCGACAACGCGACTGCTGTAGTAGCTGTTGCAGTGGCTC

AAAACTATCGTGCGCAATCCGAGAAACTTCGCGCCCCAGCACAAAGCAGTCAACAAACAACACGGCGGGA

CAACGTGGAACAACCTACAGCACGTTTCAACAGAGACTTAAAACACACAGACACTGAGAGGCTGTCCCGC

G

>NODE_418_length_349_cov_7.478659

TACGCGGCCACGAAGGCGGGCGCCGACATGCTCATGAAGTACGCGGCGATCGAGGGTGCTGCGTCTGGGG

TGCGCGTTAACTCCGTCGCGCCTGGGCACGTAGAGACGCCCATCTACGGCGACATGCCTCTCGAGATGCT

AACCAGCGTGGCCAAATCTACCCAGCTCATCGGACGTCCCATCCAATCGGACGAGATTGCCAAGGTGGTG

ACGTTCCTGGCCTCCGACGACGCTGCGATGGTCACCGGCTCCGTTTATCTTATGGATGGCGGCTGGTCCA

TCAAGGCCTAGCGTGTTTAACCGTCCACACAGAAGCAGAACCACCCCAGAGCGAGATAGGGAGGACGGG

>NODE_419_length_349_cov_5.192073

TGCTAGTTCAGCTAATCCAGAAGTGAAAGATGAACAACAAACGAATTCTGATGAAACAGTTATCGATACT

GATTTTACTGACGATTAGTATTTCTATAGTAAGTTTATATATGTAAATTTAATTGCTTAATACTTACTAA

TTGATATATAATTATTAATAAATAATTTTATCGGAGGATTCTTATGCTTAGTTTATATTTTTTAAAACTA

TTAGTTTATGCCATTGTTACTGGGTTTAGTTCACTGTTTATATTTGGGTTTTTATCTAACGATCCGTCAA

GAAACCCAAACCGAAAAGATTTCGAATAAGACATTATATAAAGTAAAGTAACGTACCAACCTTATTTAT

>NODE_420_length_349_cov_2.222561

ATCACGGAGGGACCGATCTCCCCCAGAGCGTCCGCTATCCGTGCATCCTTAGGGGTGTTGGGGTCCTGGT

GTAGGAAGTAGTGCACGATGTGCACCATGTAGTCCACCACCAGTCCAACAGCCATGATCAACTCGATAAC

GGTGATGCTGTTCACGTCCAGCCCCCAATGGTAAACAAACCCGAGCAAGTCCACGTCTATGATCACCACG

GTGAAGCAGATCAGGGCGACGATTGCGATCTTCCCCAAGACGAAGACACTCAGCACGGCCACGGCAACGA

GAGCCAGCACGAAGTTCATGATCGAGGAGATCCTGAAGGACTTCGTCAACAGCAAGAAGGGCGGCGACT

>NODE_421_length_348_cov_6.678899

CACGGGCGACGCAAAAAAGCAGCAGCAGATTTCAAAGATGGTGTCATGTCCCCCTTTCGTTAACAAACAC

CGCTACCATTGCGTGCAGCGGCAAGCAGCAGTAGCGCACACAGAAAACCAACAGCAACCAAAAACTTCGG

GGTCCCTACTCCTCAAAGCCGGTTGCAGTGCTTATCGCAGCTGAAGAGCTACGCGCAACGGTTTTCAGTG

CCATCCGCAATCAAGCCGATTTGGTCGCACTCCCTAAGCGACGACCTTGGCGGAAGCCTTGCGAGAGCTA

CGCCTCTTGGAGATCATGGAGCGGCGGGCCTCCTTCTGCTCCTTCTGGCGCTGGGCGTACAGGCTAGC

>NODE_422_length_348_cov_6.553517

TTAAGAGAGGGAAAGGTAGCACTGCACGTTAAACGCTTTTGGTGGTCGGTGGTTGGGGACGCACGGGCGG

GGTCGGGGGACGGGTGGGGTTGTCGGCGAGAGCTGACGTCCAACTTACTACAAAAATAATAAAACAAAAA

AAACAAAAGCGTCTCCCAAGAGTAATATAAAGGCGTTCTACAAAACCGAGAAACAACATGTCAGGATACA

CAAAATCCAATAGACAAGGCAAACAAAAAGGGAGGATTAAACAAAAAACGAAGGGAAAAAAGCAAAAGAG

GGGAGATAATAAAAAAGATAACAACAAAATGACACAGAACACCACAAAAAAACATACAAAACTAAATC

>NODE_423_length_347_cov_15.058282

CCTGTGCATGTGCCAGGCGTCGTCCACCGAGCTCCCCCGCTACGGCGTGAAGGTCGGACTGAAGAACTAC

GCCGCCGCGTACTGCACGGGCCTCCTCATGGCCCGTCGCCTGCTCCGCAACCTGGGCCTCGACGAGAAGG

ACGAGGAGAAGAAGATGGAGAACCTCTACGTTGGACAGGGTGTCGGCGCGGACCCCGAAGAGGAGGGTGA

CGACGATGAGGACGAGGTCGACGGAAACGTTGTCAAGAGCGAGTCTAACGGACGCACGTACTACGTGGAC

GAAGTGGACGAGGAGAAGCGCCCCTTCAAGGCCCTCCTTGATGTTGGCATCCGCTCCACTACTGGAA

>NODE_424_length_347_cov_9.960123

TGGAGTTCCTTGGCATCCGAGTATTCCAAATCTGTCTGCCCAGCACATAATGCGCGAAAACTCGCAACCG

TTTGCCGTTCTTGGTCACTGCTCCCGCCATTCGACTCTCTCGATCAGCTGTCGTTTGGTTGTTCTCTTCG

ATCCTCCTTTCGCGTGCGTGCCTGCCTTCTCGGGTTTGCGTTGATAGGATGGGTCTAGCCGCCAAGGCCG

CCGACAAAAATAAGTATCGATGAGAACGGGGATCACTTGTGTTGCGATAACCTTGTGCACCGAATGCAAG

CTCGGAGTTGTTGTGGGGATCGGTGCCGGCGGTTTGATAGTTTGCTCCGCCCACGGAAGAAAGTGAT

>NODE_425_length_346_cov_15.150769

TTTTCCTCTCTATTGCTAAACATGGACGAGATAGACCGTGAGAGGCAAAGGGAGGGTCTGGCCGTTTAAA

GGACTAAAATTATGTGATGATTCTGTACTTGAGGGGGACCCGACGATTAAGGAACTGATTGAAGTTGTCG

TTGCTTTTCGTTGTTCACAATCATTCTGATATCTACAACTGGGGTTGAGGGGCTAGTAGAAGCGTCCACC

ACATTGGCTTATCAAAGCCCACTCTGAACCAACGTATCATATGGAATAGCATTGATGTTTTGGTCAAGGA

CTAACGTTACGCATTGCGTTTCTCAAACAGGGTATATAGCAGTGCCTAAATGGTGCCTAATACGAT

>NODE_426_length_346_cov_3.769231

CCCGTGATCTGCATGCATCCGCATGGGCTGGATGCTCTGCCCACGGATGAGCCACGCCCCCGACACCAGC

AGTTCATCTCCTTCCTTACCCGTGATCTGCATGCATCCGAATGCCCACTTCCTGATCTCGCCAGACCGCT

GGATGAATCCTCCCACCAGGTTCGACGTCATGAACAAACGAGAGTTCTCCGCGTTGTACTTGTACCGGCA

AATCCACAGGGACCATCCCTCGGCGTCGAAGATCTCCCAGAACTGCTTCATGCTGTCTTCGTACGAGTTG

TTGGAGTAAATCTTCTTCCACGTGTCCCCGGCGAATGGCGACTTGGCCTCCTTGTCCATGATGGCA

>NODE_427_length_345_cov_45.077160

TGGAGTTCCTTGTGACCCGAGTATTCCAAAGGTTAACCAATCGCCTACCCGAACTAAGCCTTATAACACT

CTCCCCCCCTCTGAACGGCGGGATATTTGGGACGTCGGTACCGTATACGGCCGTACGTTGACCTACCCGC

TTCGTTTCAATACCTGGGGAACCAGGGAGACGTGTTATTTGGGGACGCACGCGCAAGCGTTTCTGAAAAA

AGAAACGTTCGAAGTGAGATTCGAACCCGTGTCCTCGACGACGTCAAATGTGAGGGTATTACCTGCTGAG

CTATAGGGGTAAAAGATCGATCGATTACCCAGCCCAGGGTAATCCTTATAACACTCTCCCCCCCT

>NODE_428_length_344_cov_10.780186

TTAGTGGCTGCTTTAGGTGGCTTAAGTTTAACTTTTGGTGGAGTGCTATTTATGCATAACTATAAAGGGG

GGGGGGAATTACTTTGTTTAGGGGTTTTTACTATTTTATATGTTATGTTTACTTGGTGGAGAGATGTGGT

TCGTGAGGGGTTATTTGAAGGTCAACATACATCATCGGTTCAGCAAGGATTACGTATGGGTATGATACTT

TTTATTGTATCTGAAATTATGTTTTTTTTTGCTTTTTTTTGGGCTTTTTTTACTTCGTCAATTTCTCCTG

TTTTTAATATTGGAGGAGTTTGGCCTCCTGCGGGGATAGAGGCTTGGAATTCTCCGGGTGCCAA

>NODE_429_length_343_cov_1112.807453

CTCTCGTAGGTCCTGCAACGGAGACACCTCATGGTGTCTATTCAATACCTCGACCTCAATTTAGGCAAGG

TTACCCGGTGGACTTAAGCATATCATTAGCCGGAGGAAAAGAAACTAACTAGGATTCCCCCAGTAACGGC

GAGTGAAGCGGGAAGAGCTCAAATTTAAAATCTGGGGCCCTTATGGGTCTCCGAATTGTAGTTTTTGGAT

CGACAGTCACGGGGTGGCACAAGAGTATAAGTGTTTTGGAACATTGCATCAGAGAGGGTGAGAATCCCGT

ATGCGACTCTTGTGTCCATCTACTACGTGTCTGCCCTTTCACGAGTCGGGTTGTTTGGGAATG

>NODE_430_length_343_cov_67.984472

AAGGGCGAAGATAATAAATGGTGGAGGTGGTAGGATTCGAACCCACGCTACATTAAAAATGTAGATTGAT

TTAGCAAACCAAGGCTTTCAACCACTCAGCAACACCTCCAAAAGTTACCTTTCCAAGGGTTTAATATTAA

GTTCCTTATTACTATTATTAACCCCACACCCGAAGTTTGAGTTTTTATAATTATGTCTACACCAAGATAT

AAATTTTAAGTTATTATCTAGAATCAACTGTCACTTTTAAGTGTTTGATTCTAGATTTTAAACTAAAGGC

TAAAGAAGGTAATATAAAACGTACAACTACCAAATAAAAATTAAAAACACAAAAAAAAAAAAG

>NODE_431_length_343_cov_4.015528

ATTGGATATGTAGTTTTGAATTTATTTCCTTTTTACACGTGAATTGATGTCAAAAACCGCGTCGATGACG

GTACTTGGTCAACCGAGGTAAATCGAATTGACACACGTTTTGCTCAATACAACCCCGCAGACCCCGGCGC

AAATGCCGCAGCTGCCAACAAAAAGGAAAATCGATCTCCAGTCACTATCGCCTTATTACAGGGCGTGACA

TGAAACGCCACGTGATCGGTACGGCCAACAATCGAATGGTAAGCGCCACTTTTGGCCAGTGGGGCTTGTG

ATACCTCCAAAAATATTCTACGGAGGGGAAACGCGCGCCTCCCGTGAAAAAAAAAAAAAATTT

>NODE_432_length_343_cov_3.400621

TCGGAGTAGCGAAGAAACTGCTAGAAGTGATGCTTACTTTTGGGCTACCGCTGTATGTTCGCAGTGATCC

GGGATCCGAGTTCACGGCAGAGGTGATGCAACACTTATGCAAGTGGTTGAACGTGACCATCGCCTATGGG

CCTGCCGACCATCCTCGTGCGCAGGGGACGGTGGAGAGACTGGGGGGGTGGCTACACGAAGCACTAGGAG

AACTGTGCAAGACCTGGCCTCGCAGATGGGACGAGTACGTGCAGCCGGCACTGTGGATCCATCGTACCAC

GCCGGACCTGCGCTTGCCAGAGAAGCCAACTCCCTTCCGTCTTTTCTTCGGTCGTGATGCACG

>NODE_433_length_342_cov_22.987539

TGAGCCACGGAAACGCGGCAGGTAATCCCCGTTGAGTTGGCACCGGCGCACGAGGCACCGGCGCTGACTG

CATAACTATGCTGTGATGAAAACGACCCGAATGTCCGGCTGGGACTCCGAGAGCGGGGCGCGCCGAAGCG

CAAACCCAACTCCCATCTTCGCCGCAGTTGCCAGACGCTCGAGGATTTTTCCGGGTTCCGGATGGCACTC

TCTGCCCAAGTATCCGGGGAATTTCCCAAGGCTCGACACACCCCCTCCAAGCTCGGGACGCCCGTAGGCT

AATCCCTCGCTGGATGATTGGGGGTGCCGCCCGATAGCAGTCCGTGTCGTGTTTGCTTGAAT

>NODE_434_length_341_cov_9.146875

GCCAGACCCCTTTGAGGGTGATGCAGCCGCCCCGCTTGCTACCTCCCGACTCGACGTACAGGTTCTTGGA

CCCCGAGTTGTACCGGACCGACACCGTGGCACTGGAGTCGCATCTAGGCTTGGAGATGGCCTTGTTCGGT

GAAGCCGTGTCCGCCGCGTCGGATGGTCTGGCGGCCGCAGCCAGGAGGGTGACCACCGCAGCCGTTCGAA

GCAGCATCGTCGTGTTTTCTACAAATACCGGGTTGTTTTGCTGTAATACTCTCCCTGGGAGATATAGGGG

TTGGTGTGGTTATTGCTAGAGCTGGGTGGTTGACCCTGGGCGATATGACGAAGAGTGGGGT

>NODE_435_length_341_cov_8.746875

ATTATCATTTATTGTACAACGACCTGCAGACGAACCTGGTTTCTACTTAGAACGTAATGAAGTTGGTGGT

CGTAACATTCAATACACGATTTCAAGTTATGCTGTTCAAGCACGACCTGCTGGAGACCGTTATTAAGGAT

AAGACATCATTTTTAATTTTCTAGGAACTTACGAACTGTAGCCTTTATTAAAAGTCCTCAGTTCGTTTAG

TTTTCAGAAAAACTTTGGAAACTTAACATTATTTGTTATATTGATTAATTATTTTTTGTAGAAATACAAA

GTTAATTTAACTATTAGTTTCCTTTTCCGGTATAATATTTTATGTTTAGAAATAACTTCTA

>NODE_436_length_341_cov_7.262500

ACTGTAAAGCATTGAAATTGAGGGTAACATAAAAAAATTTCTATTGAACGTGTGTTTTTATGTTACCGCT

GAGCTATTGAATTAATCATTTCTTACCTATTTCATTTTTGCATTCTACCTTCATCTTTCTGGTCTATATC

CCACTGGCTAATAGACAGATGCTTCATTTCATCAATAAACATAATGATAGAGTGTATGGGCTTAATAATT

TTATAAATCATCATTTACTCCTTGTGCTAAATTGAGTAGTTATATTATTCAGAAAATATGCTGTCTGCAG

ATTCATCTACTTCAAGTATTTTGTTTTTCAATAATTGTAAGTTACCAGCAGTAATAGTGTT

>NODE_437_length_341_cov_3.471875

ATGTCAGGGAGGTCAACGTCAAGTGGCTCCGATCGCAGATCGGCTACGTGGGACAAGAACCGGTACTGTT

CCAGGGAACGATCCGCGAAAACATCGCCAAGGGCGACCCCGGGGCTACCGTCGAGCGCATCGAGGAGGCT

GCTAAGGCCGCCAAGGACGCCAAGAAGAAGATCGACAGGTCACAGATCGTGTTCGAAGTCAAGCCGTGGG

AGGCAGAGTCGGACTTGAAGGGTTTGTTCGATAAGATCCGCAAGGTTGAGATCGCCGGGCTCACATGGGG

GGAGGCACACAAGCTCGTACCCGTAGCGTTCGGCGTGAAGAAGTGGAATTCTCGGTGCCAA

>NODE_438_length_340_cov_8.479624

CTCCACTACGGATACGAGGTCGCCAGGAGCTTCCTCGACGGGGCCCACGACATGGACAACCACTTCCTCA

ACGAGCCCCCCCGCCAGAACATCCCGGTCCTGATGGGCCTCCTCGGGGTGTGGAACTCCACTTTCTTGGG

GCACGGGGCGCGGGCCCTCTTGCCGTACTCTCAGGCGCTGAACCGTTTCGCGGCGCACATCCAGCAGGTG

GACATGGAGAGCAACGGCAAGCGAGTCGCCCAAGACGGCTCCGTGCTCCCCTTCGAGGCGGGAGAGATCG

GATTCGGAGAGCCCGGCACCAACGGGCAGCACTCGTTCTACCAGCTCATCCACCAGGGCC

>NODE_439_length_339_cov_23.323899

TTCCTGCTGCAAGCGGCGGACCGCGCTAGTCTCTCCCATGAGCGCTCCGTACACGCCGTCCATCCTCCAG

TGGACCCCGAGGTGGGACCGTCCGATGAGGACGTTGGCGCTGACCTTGTTGAGCTCTCCCTCGTACGTGA

GGCCGTCAACCTTCTTCCCCTTGTCGTTGATGCACTCGCCCACAATCTCCCTTCCTGACGGCACGTAGGC

GATCCTCTCAGTTCCCTCGTTGTTCGACACCACGGGGTCGGGGAAGCACCGCTGCCCAACTTCGAAGCCG

AGGAACGCCTTAAGCGAGGTGATGTAAGCGCCGACGTTGATGGCGTGTCCGCTGGGGTA

>NODE_440_length_339_cov_14.974843

TGAAGACTGCTCATGTGTTGGATGTTTGTTCGGGCACAGGGAAGAAAGTAGCCGTAATCTTTTTCCTGGT

CCAGTGTTGTATATTTGTGGCGCAACACAGTAATAGTTGGGAACTCGTAAATTTTGTGCAAAAACATTCT

GTGTGGCAAGGTCGGCGGCGAACCGAGGAAGAGGGATGCGGCGCCGCAGAAAAGTAGAACATCGTTGACA

GCATTAAATGGCAGCAGTGGGTCATCGCGCGTCGAGATCTATTTAGCGGTAAACAGCAGTCACGTACGTA

CTGTTTTGATGTCGGCGCTTCCGTTGCTTTTGGCTGCGCTGGAAATCTCGGGACCCAAG

>NODE_441_length_339_cov_6.279874

AGGTGAAATTGAACCAGGCTTACATAATACTTGACCACGTTCAACATCTTCACGCTTAAGACCACGTAAA

AGAACACCACAGTTCTCGCCAGCACGACCTTCGTCAAGAAGCTTACGGAACATTTCAACACCTGTACAAG

TTGATTTTTGTGTATCACGGATACCAACAACTTCTACTTCTTCACCAACTTTGATGATACCACGTTCAAC

ACGACCCGTTACAACAGTACCACGACCAGCGATTGAGAATACATCTTCGATAGGCATAATGAATGCACCG

TCGATTGCACGCTCTGGCTCTGGAATGTAAGAATCTAATTGGTCAGCAAGTTCTACAAC

>NODE_442_length_338_cov_27.703470

GATGGCTTCTACCGACCGTGACGCACTGGTTGCCCTATATAACGCGACCGACGGGGCTAACTGGAATAAA

TCGACAAACTGGAACAAGGGTGCCGACCTCTCGCTATGGCACGGTATCAAAGTGAACGACCAGGGTCGCG

TGGTGGAGCTACGTCTACCCGAAAACAACGTTGTTTTCGGGTAGACGTAGCTCCACCACGCGACCCTGGT

CGTTCACTTTGATACCGTGCCATAGCGAGAGGTCGGCACCCTTGTTCCAGTTTGTCGATTTATTCCAGTT

AGCCCCGTCGGTCGCGTTATATAGGGCAACCAGTGCGTCACGGTCGGTAGAAGCCATC

>NODE_443_length_338_cov_4.766562

TTTTTATGGAATATAGAGCATTTCTTGCATCGTTGTTAGCACAGTGGAACATTACTCCTATATTTTATTA

ATAGGACGCAAATTTTGCCAAATGTCGAAGAAATTTTCCTGCGTATCCGAACAACTTGTCACTCCATCCC

ATTCGGTTACTGAACCCGTGCACGACGATAGTTTACCACAGACGCGTCGAAGACAAACCAAACCAAACCA

AACGAAATGAATGACGTACAGTAGAACTTTAAAGAAGCCGGACCCATCTTTCAAAGCAATACAGCCAGTT

TTTTCATCCGTTGACCCAACCGCGAAAACCAGTCAAATGCGGGATATTGTTGCCTAAA

>NODE_444_length_337_cov_77.781646

GTTCCTGCTGCAAGCGGCGGATGGCGCTAGTCTCCCCCATCATCGCTCCGTTAACACCGTCCATCCTGTA

GTGAACACCGAGGTGCGACCTGCCCAAGAGCACGTTGGCGCTGACCTTGTTGAGTTCTCCCTCGTACGTA

AGGCCGTCGACCTTTTTGCCGTCCTCGTCGATGCACTGGCCCACGATCTCCCTTCCGGTCGGCACGTAGG

GGATCCTCTTGGTTCCTTCGTCGTTCGAGATGACGGGTTCGGGGAAGCACCTCTGCCCCATCTCCAATCC

GAGGAACGCCTTAAGCGTGGTGATGTAGCTGCCGACGTTGATGGCGTGCCCGCTGGG

>NODE_445_length_336_cov_4.853968

TAATCACCCACCGACACGAACATCAGACACGGCGAGGCTCTACATACCTGCTACGCGGGTGCCGTTTCTA

CGACCTCCTGGGAGCGTTACGTACTCCTGCTGCTGTGGTGGCAGGCTGGGGCTGTGAAGAACAAGAACAA

CAGAAGCGCAAAGCGATCGCGACAACCACACACCGGCGGCACTTGTTTTCCTGCCACGATGGGTGCGGCC

TGCAGCGGCGCTAATTGTAAAACCGACCCTCGGCCTAGCCATCCCTACCACCCAGAGGGTATACCTAGCT

TACACGCCAAACCTATTAATGGCGCCAAAATACCGGAGAAGACTCGTACGAGCACC

>NODE_446_length_335_cov_13.904459

TTGCACCGCCGCCGCCGCAACATGTCCGGAAACAATTTTGTGACGAAGCATTTACTGAGGAGGAGGGTAT

CGGAGCACCGACCGTACGAGGAAGACAACGACAACAACGACAACTCTTCCTCCTCTTCGTCTGATGAAGA

CGAGGGAAGGAACAGGGGGAGCAAGGGTAGGGAGGACGGCAGGGGGAAGGACAAGAAGGACAAGAAGGGC

AAAAAGGACAAAAAGGACAAGGCCGAGTGGGTGCAGGTCCAGGACCCTGTTGGCCGCCTCTGTTGGCAAA

ACCGCAAAACCGGTCAGGTGACGTACGGCAACAATATCGCTGTGCCTCAGCAAGG

>NODE_447_length_335_cov_7.095541

TTGCTAAGGAGAAACTTAGGAATTATGCCTTGTTTTGGTGGTCCAAGTCGTATCGTAGGACTATTTTTTT

TGTTGGGCTCGATCGTCGCGATGCGTAGGCCGTCTGTGGGCGTACAGTAGTGGAGGGAGGCAGCCATACG

ATCTGTCTCTCTCAAAACCGGCGCCGAAGCGAAAAAACACGACGATTCTATTCACAAAATGCAAGAGTCA

AGTCCGTCCAATTTCAGTTTTGTTTTTACGGTACATATGTAGACATCGAAAATTGAACGTGGGGAATCTG

GGGTTGGTGGATCGATATCAAAATCTTAATATACAAAAAAAAAAAAATTTGGAAT

>NODE_448_length_334_cov_1564.124601

TGACACTTTTATATCCAAATTGCTTTTGCAACTCAGCTACAACTGTATCTTTATATAAATCATGCAGTTT

CGCCATCGTTTACTCCAATTATATTAATTCGTTATTAGATTTGAAAAAACGTACTTTCTTTCCGTTTTCA

TCTCTAAAGCCAACACGATCTGCTTTACTCGTTGCCGAGTTGTAGATCGCAACATTTGATACTTCGATCG

GTGCATTTTGTTCAATGATTCCGCCAGCTACGCCCGCTTGTGGGTTTGGCTTTTGGTGTTTCTTTTGAAT

GTTTACACCATAAACCAATTATCCGAACCAACGGTTCCTCTCGTACTAAGTTGG

>NODE_449_length_334_cov_18.744409

GTTACCACCCGTATTCCATCACGTTTTTTTAGTCAATCAAGTCGCGCGAACAACGAACACCTCGCTGTAC

CCAACCAATCAACGCATCCTGCTCTCTCGGCCTTCATGGCCACCACCACGGATTGATGAGTAAGTGTGCA

AGGAAGGACACTTCGCCAACTACATTTTCGGTTAGACAGTGGTGCACGATCGCCGACAACTTGCTACACT

CAATTCCATCTATCTCCAACTGCAACAACTGACCAACAACGACGCGTACGCCACAAGGCAATGCCTACGT

ACCTCTGCTACGTACTCCCTCATTGTAAAATTTACGGTGCGAAAGCACATCCCC

>NODE_450_length_333_cov_33.307692

CTTTTTTTTTTTGTTTATTCTTTTATATTTTCTTGTCCCCAGTCAGCGCCGAAACGCCTGTCGCACCTCA

AAACCCCTCTTTCGGTCAAAAGCAGCTTCAAAGAAACAACTCACACCCAAAAAAACAGATCCCAGCGCGA

TAACCCTCCCAGCACCCGAAAAAAACTTCCGAGTGGAAGCAAAGCGAGCAACAAATCTCGGCGACCTTGT

AGCAAAAAATCTCAGCCCCACCAAGGCAATAACGTCAGATACAAGCTGAAACCCCTGCAACATCGCCCGA

GGCAGGCTTAGAACTTGATGTCTCCCTCGAGGTTGGCCTTGAGGTAGAACTGC

>NODE_451_length_333_cov_6.926282

TGGCACCCCGAGAATTCCAAGCAGCCAGGAGATCTTTGGAGGTGAAGGCGGTGCGCCAGGATCATCTGGT

TCGTATTTGGGTTGATCTGGAGAATCGGCTGGAGCTGGCGAAAGCGGGTCTTCGCAAGGAGCTATGGGAG

GTAGGCCTTCGCATTGAGCACGATGCTTTCTGGACTGTGCTGGATGGTCTACCATTCGCGGAGCGTGAAG

CATTCTTCTCAAGGATGGATGTAGAGCATGCGCCGAAAGCAGCCAACGCGGCCGAGCGAGCAGGGGCGCA

AGAACGCAAAGTTGGGCCCGTCTTCTTTTTCGTCGGAAGCGCGACCTTCGGGT

>NODE_452_length_333_cov_4.615385

TGCGATGACCGGGATGCCCTTCTTCGCGAGGATGTCGAAGGTCGCGTGCCCGTCCATGCAAATCTTCTTG

AGGTCCGACTTGTCCTTGGTGGAGCTGATCATCCGGATGTCCGCGCCGGCAATGTAGTTGTCCGGTTTTC

CGGAGATGAAGACGACAGCCTTAACGGACGGGTCGTCGGCGATCTGTCCCAACCAGATGTCCTCGATCTC

CTGCCGGAACTCGCCCGAAATCGTGTTCATCTTCTCCGGCCCGTCGATTCTGATGATCGCGACTCCGTTG

TCCAGTTTCTCCACTGCGCGAAAAAACCTCGTGTTCGAGTTGACCTGACCTGC

>NODE_453_length_333_cov_4.551282

GGTCTTGGTTTCCCACGAAACGCCTAGGGTTGAACTGCCTGGCGTCTGGGCCGAACTTGAGCTCGTCCAT

CTGGCCAAGTGCCTGGTGGCACATCAGCATGTCTCCCTTTCTTATCTGGTACCGATCGCCGTTGCTGGTG

GGGACGACAACGTCCATCTTGGCCGTCTTGCAGTAGAAAGGAGGTGCGGGGAACATGCGGAGGATTTCCC

AGTGGAAGCTGTCCAGCAGGGGGAACTTTTCCACATCACCCACAGCGGCCATGGTCAGGTCCTGGCCATC

GAGTTCATCCTGCAGGTCCTTCAAGAAGTCGGGGTTGGCGTTTAAGGCAGCGA

>NODE_454_length_333_cov_3.173077

ACTGTCCATAATATGTTAGCTATGTATAGCAACGTGTCGTTCGATAACTGGGCAGAGCTGCTACCTTTCG

TCCAGTTAGCGCACAACACCGCTTATTCAACAACGCTCCAGGAAACACCCCATTTCCTCATGTTTGGACG

CGCAGCTGTACTGCCTGTCGATCTCATTTTAGGAGTGCCGTCCACCTCCGCACCGCAAACACAACTCGAT

TATTCGAAACAAACTGTCGAAAACTTGCAATTTGCATACGAGCTAGCCCGTCGTAATCTCAAAGAGCGAG

CGGATAAGCAAGCGGTTGTAAATGAGACCCTGTCATTTCCCAGTTTTAAAACT

>NODE_455_length_331_cov_7.777419

GATACTTGAAACGTTCCTTTTAAATGCATTAAACCGATGATGCCATGCAGCGCCTGCCACCCTCTCTCCA

GAAACCCGCTGTGAAACCTTACAGTACAAGCACCGGCGCACGAATTTTGATCATGATCGACACTTCCGAT

TATATCCGAGCACTTCAGATGGGTATACATGTATGTACGTACAGTAGGTGGACATGATACTCGACCACCG

TCGAGTTCTATCATATACCGTGCCAGGTATATAGCCCATTTCTATGCGTGATGTGCTTTGCTTTTTGTCA

TAGATCATGCATAGTATGCTTTGTTTTTCATCATGTATACATAACCTTTGT

>NODE_456_length_331_cov_3.848387

GCCGGCTGCCAGCTTGATGAAGGGGGCCGAGAGGAGGGGCTCGCTGAAGTCTTGGGAGATGCCGAGGCGA

GCGTTCACCTCCGGGTCGTTAGCTGTCTCCGCCGCGTTGAAGTACACGATGCGCACCTTGAATGTGCGGG

AGTAGTCCTGGAGCTGGGCCTTTTGAGTCTCGGTGACCGTAATCCCAACGTACACCACGTTGCACTTCCC

CGATGTGGCTGAGTTCGTGAGGTCGAGGGTGCCCGTCGTCGCGCTCGGTGTAAGCTGAGAGGCACCCCAG

TACCCCTTGTAGAAGACGGAGGTCAGGTACGCGTCCGGCGTGGCGGAATCG

>NODE_457_length_331_cov_3.619355

GATTATTCCATGTTTTTTTTTTTTAACCGGGGCGTTCTTTTCCGTTTCTTGGATCGTCGTGTACACAACA

AGGTACAGTAGTGTGCGGAGGGTGTTGTGATGAGAAATGTGGAAATGGCAGGTAGACTACTATTAGTGTG

CTATGGTGGGGTAGGAAAAGAGGTTTCTTTATTTTGTTTCTGGTAATACCACCTACCCTGGTAGGACTTT

TACTTGTAGGACAGCATCGGAAACCAAACTCGGCGCCCTTGGTAGTGATTAGATGGAACGAACGGTGCAT

GGGGTATAGGCAGAGGCTTCAAGACGCCAACCCAAGGCGACTGGATGAGCG

>NODE_458_length_331_cov_3.341935

CCAAAGAAAGGCCTAAGCAAGCAAGACTATGGCTGGAAGGACTACTACCGCCTATCTTCTACCGGCCGTC

ATGGCCCTTCTTTTCCTGCTCGTGGGGCTTGTGGCAGCTGCCGACGTGCGGCTCGCAAACGAGGAGGGTG

GAATGGGAACGATGAGGAGGGCCTTGATGACCTCGGCCTCTGAGTGTGCCGCCGAAAGATCAGCTTGTAC

TTTGGACACCTCGTGCACCACATGCTTCACCGAGTATTCGTCCAAAGTAGAAGGGTGCGGTTCAGAAGTC

GTTCCCTGCGGAGTCGCACAGGATGTCGTTTGTTGCGCGCTCGCCGACGAG

>NODE_459_length_330_cov_9.919094

GGTGACGACTACAGAATCTTAACCGCTATTAAGACATTGCCAGGTGCACCACGTAGACACTCGTCGTACT

GCTCGGAAGATTAGGGACACACAAGCACCAGACACAGGGAAGACGCGTGGCCGGCGCTAATCGGTACGGA

AGGGAACTGCAGTCGAGAACCGGAGTCAATAGGGTTTGGTGTGGACACCAAAGGTCGTGGGTAATTTCTC

GAGCGGAACAAACAGCAGTACAGCGTGCGCATTGCACTAAAGAACGTCAGGGTCCAGCTCCACGGCGCGT

GAATCGTGCCTTCGAATCGAGCGAGCTACCAGGTCCCTTCTACGATGGCT

>NODE_460_length_330_cov_3.673139

TATACGGCTTGAAACAGGCGTCACGAACCTGGCACAGACACCTGGTCGTCCGTCTGAAGAGTCTTGGATT

TGAGCAGAGCCCAGCTGATGCGTGTGTCATGCGATTGATCGAGTCGGGCTCTGTCTCCATTGTAACAGTG

GTTCATGTGGATGATATTTTTGCTGTAGGACTCAAGGCCAGATGCGATCAGTTCTGTGAAGACCTGAACC

GTCTCGTCCCGATCAACAACCTGGGCGAGTTACGGTGGTATGCTGGTTGTCGTTTTTTGAGGGATTGGGA

TGCTGGTACTTTGACGATTTCGCAGCAGGCTTGGAATTCTCGCGAGCCAA

>NODE_461_length_329_cov_6.467532

GCTTAGAAGCGGAGCTTGTCGAAGAAGTACTTGACGCCGGTGGCCTTCTTCTCGGACTTGACGTCCTTCT

GGTTCAGGTACCGGTCCTCGAAGATTTTCCTCACGGCCTTCTTCACGTCGGCAGGGGAAGCCTTGATGGC

GGACTCGTCCACCACCTTCTTGAGGTCGATGTCCACCTGGTAACGGGTGGGCATGAGGTGGTTCACGTTC

ACGAACTTGACGAACGGCTTCAGCTTGGTGCGCTTCTTGATGGTGTCCTTGCTCATCGCCTTGGTCACCC

TGCGAGGGTAGCGGTCGATCCCGGCGACGATGGCGTGGCCGAACTTGCG

>NODE_462_length_329_cov_4.889610

AGTGCTTTCTTTTTAAGAATGGTTGTATGTCTTTTTTCTAAAAACTTAATAGAGTACCATTTTTTATCTA

TAGATATACCGAGGCAATCAAGTTGCGAAGCTATTTCAGGCACGTTGTTCTGCAAGTCTTGTAACGTCTC

ATATATTATCATAATAATAGGACATCCTATACCAAGTTTGGGGGGGTTAAGGTACAAGGGTACAGAGTAT

AATTTTGCATTTTGCAATGATAATCGTATTTTTGATATTTGAGAAGATTTTAAATTACTGCCATTAAACA

AAGCAAAAGTGCGTTGTTCTGGTAAAAAAAATCTTTTTTTTCGGAGATG

>NODE_463_length_329_cov_3.633117

TCCACGCAAGCCTTGGCGAACAGGACGGGCGCCTCGGCCGAGATCATCCTCACATCCTCGTCCGACTTCA

TGATGCGCTTGATGCGCGCCAGGGGCAGGTCGTTGTGGTTCTTGAAGTCCTGCTCAGTGCCGATGTTCAG

GACTTGCATCTCCTGCATCTGGTCAGCCCAGAACCGACTCAATTGCTCCTGCATGTTCTGTGACTCGGGG

CTTTCCTCTTCCGGAGGGTGCAGTCGAACGTAGGCGTGGGCGTTAAAGCTCATCCGAGATGGAGCCTAGC

AGACCCAAAGCCCGAAATATTGAATCGCTATATACCTGGTACTACTAGT

>NODE_464_length_328_cov_4.228013

TCCTACGAAACATTTTGACCAATCAACGGATTTTCCTCCCCCGCAATACCCAATCAAAACCCCTTTCCGG

GGCACACTACCTTAAGGACGGAACTACTCTTGCCGCCAAGGGCGCGCAAAAACCTACATCCAACCATACA

CAGAAACCCCGCAGCGGGCCAGTAGGATCAGTATCGCAACATGCATCCTATATACCAACACAAAACAAAC

AAACATAAAAAACAAAACAGGCCTCAACAACATAGAAGTCTGATTTGTCATGCTGCATCCGCGCAGAAGC

CTCCGGTGCTACAGTACTGTACGCTCTTGTGGCCTTAATTAAGGCTCT

>NODE_465_length_327_cov_6.529412

AGGTTTAATTGAATCCGCAGTGGATTCTGCTTGGATAAAAACACTTCAAGCTGATGTGAATGCTTCTATT

TTAGATTATAAAGAGGCTTCACGTATGCTAAGTCATGCTCGTAAACTATTAGACAAAACTGCATATGATC

TTTCAGCTCCTTCGAAGTTTAGTGAAAACAATTCTATAGATAGCCTTTTAGCTCTTCAAACAGCAACAGA

TAATAGCTTAGAAAATATGTCAAATGGTATTAAAAACGCTTTAACCTTATCCTCGGCAATTGATACTATT

CTGGAACCAGGAAGTTTACAAGCTGGAATCTCGTGTGCCAAGGAACT

>NODE_466_length_327_cov_2.411765

TACATCTATACACGCAACCCCACGTGAACCGCAATCGACATGGCTCACAGGCCAACACGGCAACACGAAA

CAAATATACTCCTACAGGCCCAATCCCATGTAGCACGGGAGCGCCTTTAACAGGCTAATCAAAACCGAAC

ATAGTACATCTACACACGCAATTCTCGGTAGCACACGAGCGTCCTTGAAAGGTCAACACGGAGTTGAACC

TAAGACGCACCTACACACACAATACCACCCAGCACGAGAGTTTGCTGCCTTACTGGTCAGCGCGACGAAA

ACAACAGACGACACCTACACACGCAATTTTGTGTAGGGGCCAGCGTG

>NODE_467_length_326_cov_4.845902

CGACCCACCAGAACCCCAGCCCTGCTACCCCTCGGCCGACTTGCTTTCGACAGTTCCGTGAGCCCCGTAG

AAAAAAAGAGTTCCCAGAATTTTTTAGGAAGTACTAAAGCTCCGACCTACTGAACTGAGTACGAGAGCGC

AAGGTGGGCTGCAGTTCGAGCTTATTTTTAGGGCCAGGTGTCTCCGACTAGAGGGTTGCTGTGAGAGCGG

AGCAGCCTAAAGACGATGGCGCTGAAGCGTATCAACAAGGAGCTTCAGGACTTGGGCAAGGACCCACCGG

CGAACTGCAGCGCCGGGCCGGTCACGGACGACCTGTACCATTGGCA

>NODE_468_length_324_cov_567.752475

ATACGGCTTTTAGCTTTATGTTTTTTATCAGAACCAAATGGTAGTCGGTCATTATTGCTTTCGGCATCTG

AAGGGGGTTGGTCAATGGGGCTAGTTTTGGGTATATTGGCGTTGCCTAGTATGTTTATTGGTTATTTAGG

TCGCGACCTTTTTATAGGTTTGGGTACTGATTTTTGGGGAAATGCATTATTTTGTATGCCTAGTAATTTG

AGTATTATAGATGCAGAGTTTATGTCAACTGATATAAAACTGTTCCCCCTTTGTTGTAGTATTATTGGTG

GGTTGGCTTCATTGGATAATGAAGCGAACCGGGAGAACTGAAAC

>NODE_469_length_323_cov_90.658940

ATCACACCCGCGTACAACCTAGCCAAGAAGATCACCCCCAAGATCTCAGCCACAGAAGCCGCCGCTCTGG

AGGCAGGCACGATCGGGTTCGACCGGGACATCTTCGACGGAACCGCGAGCCTTGACGGGCTCAAGAAGAA

GTACCCCCTCGTGAGCCTCAACGAACGGGAGCAGAAGTTCATCGACAACGAGGTCGAGGAGCTATGCGAG

ATGCTCGACGACTACCAGATCTCCAAGGACAGGGATCTCCCCAAGAACGTGTGGGACTTCATGCGTGAGA

AGGGTACTAGCGTGGCCCTTCAACTCTGGAATTCTCGGTGCCA

>NODE_470_length_323_cov_16.897351

CAGTCGCTGCCTCCCGGCCTGCACAAGAAGCAAATCGATTAAAGCCGCCCATGTGGCGCCAAACTGGCTG

CTGGGCTCGGGGAAGGAGGTTGTCTGGATGGGTCGGGCTGGAAGGCTAGCTCTACTGTGCTAGTTTACCG

TAGTGCGATGCATCGCCCTGGGTGATTTCAATCGTCCGCGAGCATATATACCAGTTGACGTGCGGTGCCG

GACAAGGTGGTTTCCACGGATTTGGGGGGTGTGCTGTGCCATACTGGCTTCAGCTGAGCTGACTGCTGCT

CGAGAAAAGGAACTTTTTCAAAAAACATAAAAAAAAAAAAATT

>NODE_471_length_323_cov_12.337748

CCGTAGTGGAGAGCGAGGGGCAGCACTCCGACGGCGGAGCACACGCTGTACCTGCCACCAACCCAGTCCC

AGAAGCCGAAGATGTTCGCGGGGTTCACTCCGAAGTCTTGCACCAGAGCGTCGCTCGTGCTGACAGCCAC

CATGTGCTGGCCGGTGATCTCCGCCTCAGATGCCTTCCCCTTCATGGAGGTGAGCAGCCAGTCCTTCATG

GTGCGTGCGTTGAGCATGGTCTCGGCCGTGGTGAAGGTCTTGCTGACCACGATGACAAGGGTCTCCTCGG

GGTTAAGCCCGGTGATGGCACGGCTGACGTCAACAGGGTCAAC

>NODE_472_length_323_cov_10.903974

ATATATATATATTATATATATGTAAGTCAAAATCATAGTCTTCATACATCCAGTATGCAAGCACGGACTC

ACGCGAGAGTATGTTTCGTAGGTTGGGAGAGTGTGCTTGTGTATTTTCGGGCAGGCGAGGTTTTTTGTGT

AGGCACTGTAGCCACTGGGCGTCGTGCGCGTTGGCGCAGCCCTTTTATTTTTATACATTGGACGATGAAC

CCTTTGGCGATTGCCGCACGACATCCAATGAGACTAACGAGAGTACAAGTTTAGGACGGCAGCTCCCGAC

GCGAGAGGCGATCGTCAGGTGCTTAAAATGGGTACGAATAACC

>NODE_473_length_323_cov_4.890728

TATCTTGACCCTTTTGGAATTGAGCGCATAATAGGTTTGGTGGGCGGTAGGACTGACGATATTTTGCGGT

GTTTCTGGTACAATTATTGGTAGATATAATATGGTATCCCGAGACATGAGTTTGGATATGTTTTTTTTTA

GTGACAAAATATATTTCCTATAAAAATGAGGATTTGACATTGGAGACGTATAGTATCTGGGTAGTTTTCA

TTTGGTTCTTGGTTTTCGCGGTTGTTTATTTTGGAACTTGAAAAACAAATTTGGTGTCTCCACAGCAGAG

AATCGGCTGTTAGGTACTATACGTAGTGGGTTATGGAATGCTT

>NODE_474_length_323_cov_4.192053

CGCGGCCTAGCAGAAACCAGTTTTTGACGACTGTTGGCGTTGGTATCTACAGTATGTGTGTGTCCGATCT

AGCCGATTTGATCCTCAGGTGAGTCTCTGTGCGACACTTGTGCCGGTGAAGTCAGAAAGTATTTACAAGG

CAATTGTTCCGTCAGGGGCGAACACCACTTGTATGTGCGGCATATGCGTAAGTCGACGGCATGTTGAGGG

GGCGTTGGGTTTGTTGGTGCTGCCTGGGTAAGACTCGAGTGTGGCGCGGGGCTGTTGTTTAGTGCGTCGT

TTGTGTTTTGTAGGGCTGAATTCGGGGTGATGACACTACAAAT

>NODE_475_length_323_cov_2.652318

AGTGGTAGGCCTCTTCCTTATTTTTTGGAGCGGAGGCCCGCCACATAGTAGGATATATACAAGGGAACTT

CGCCGTCGGCCCAGGCCGCCGGCGAGAGGAATAATGTTAAGGTCCGCAATTACGATGCCAAGGGACAAAC

GCGTATCACAAGGATAGAAAATTTCAAAAATTCATCACGGCGGGGGGTAAGAGTACGGAGCGCGAGGCAC

GAGCGTCGAGAACTCAGGTTTTCCTGAGATATGAAAATGGCCCGAATGGCACGAGCGTCGAGAACTGACG

GTTTGCTGAGCTCATCTAATTATAAGAATATGGTATGTACGTA

>NODE_476_length_322_cov_3.521595

TGATGTAGTGATTGGTGTTGTAGGTGTTTCTTGAGTTTTTTGTGTGCGGTGATTTAAAGTATTGAAATTC

GACCGTGAGCGCACTCGTGTATCTCTAGGTATGAAGCAGTTAGGCGAAGATCCATGGGTAGCAATTGCTA

ACCGTTACCCAGAAGGTTCTAAACTTTCTGGTCGCGTAACTAACTTAACTGACTACGGTTGTTTCGTTGA

AATCCAAGAAGGCGTTGAAGGTTTAGTACACGTTTCTGAAATGGATTGGACTAACAAAAACATCCACCCA

TCTAAAGTTGTTAGCTTAGGTGATACTGTTGAAGTACTAGTT

>NODE_477_length_321_cov_6.456667

TTGGTCATGCCGCCGCCGTCGGCACGGGGGTTCTCATTGTCCTTGAAGCGCGCCTTTCCTTCAAACGTCA

TGGTTCCCTCGTTGAAAACGGCCGCGCCCTCATCACACTCGTTGTTCTGGAAAGTTGCATCTCCTCCGAA

GATGATCGTCGAAGTATCCAGGTTCGCCACACTGCACCCCTCCGCTCCGTCGAAGGCGAAAGCATTGTTA

TCAAAGGTTGCGTCTTTCGCAAAAGTGACGAATGCTGACTCCTCGTTGTCAAGGGCAAGGAAGGCGTTGT

CCTTGAAGGTAGCATCATCCTTGAACATCAGGACCCCAGTG

>NODE_478_length_321_cov_5.250000

CACTGTAGCGTCGTGCTGCGTGCGGTTGGTTTTGAAAACCTGTGCGATCTCTGGCACGAGCGGGTCGTCC

GGATTGGGGTCGGTCAGTAGCGAGCATATAGACAACAGCACCTTGGAGATTGTGAGAGCTGGAGACCACT

GGTCCTTGAGGATGTCTAAGCAGATGCCGCCGTTACTGTTGATGTTGCAGTGGTAGATCCTCGTCGTGAA

GTTCACTTTAGGAGGCTTGAAGGGGTAGTCCGCCGGGAAGTGGATGTTAAGGAAAAACACCCCGCCCGCG

TAGGGGCTGTCATCCGGCCCCATGATAGTTGCCTGCCAATG

>NODE_479_length_320_cov_36.133779

TCGCCGCAGAGAACTTGCAGGGCCTCCCGACGTTCAAGGGACTGAACATCCCCAGGAGCAAGGGGGGAAA

GATTGACCCCGTCACCGACCTGTTCAGGACGACCTGGCCGGGTGTGACCACTGGCCCCGTTGTGTCTCAG

TTCATGCTGTCGGACTTCCTGATCGACTCGATCGTCGTGACACCGAAGGCGGACCCCCTCACCCCATTCA

CGGACTACATGACGTCCTTCCAGCCTTGGCTCGACGTGCAGAACGGTGCTTCCGACGTGGACACCACGTT

CGACTCGGACAACCCCCGCTTCATCCGAAACGGCCGAGAC

>NODE_480_length_320_cov_5.003344

GGCCTCGCTCGTCGCGGTTGCCAGCTTCGCAGCTGCTAATTCTCTCCTCATCTCTTCCATCTCCTCACGG

AGGGCTGCCACTGCAGCCTCGTCCATCTCGTCTTCCTGCTGCCTCTAGCGGCCTGAAATCCTTAGCAACC

CGTGATCGCGTGGCAATGATCTTGCGCTATAGCGTGTTGATACGACCTCAGGCGCCAGACAGGTTCTTGC

GCAGACAGGCGAGCTCCAAGTGCTGTTATTCTCCAGCTGTAGACATAAACCACATATATATATATATATT

TGGCTTCGTTTGAGGCTCATGCATGAGCATCGAACGCCTG

>NODE_481_length_319_cov_5.674497

CCGAGATCGTGAAGGAGATCACGTCCATCTCCATCGAGCCCGGAGTAGAGGTGGAGGTGACCATCGCGGA

CGCCGCCTTCTAAACGTAGCTGGGTTGCTAGCTGTTGCTGGTACTTCCAGCTTAGGGCTAGATAGCGTGG

CCTGAATAGCTTGATAGCTATTACTGGGGCATTCTACTAGCTTTGGTTGGATTGCGTGGCAGTCGGTGGC

AAGCAATGGTGGTGGCGACGGTGGTGGTGGTTATCATACCGCCACCACTGGCGGGTGGCTTTCAATTGAG

ATGGCGCTGAGCTGAACATAAAAAAAAAAATTTTGGAAT

>NODE_482_length_319_cov_5.426174

ATGGCTTGCCGCGCAGGCAGTAACAACAGAGGGGGTCCTGCGAGGCAACTGTTTCCTGCCAACAGTTGTA

CATACTAAGTATTGAATGTGGTGCATATATAGAGATATCACTCGAAGTCCAACCGAACCCTGTGTGTGCG

CGCCTATGGTATGTACAAGTGCGGCCAAACGTCATCATGAATATTCTACTCAAGGATTTCCATCGATACT

TCAGCGCTACTGCACGGGTAGGGCGGCGCGCTTCCCTTCCTACACCGCAACGACTGACTGCGCAGGCCTA

CAGAGATGGGCCGAAGTATCATCTGTGACAAAGTGCCGT

>NODE_483_length_319_cov_4.768456

CTCCGCCCCGGAACACATCGCCTGGTTAGCTGAAATGCAGGCCTGGGTGACGGTGTGGGAGGGGACGGAA

ACAGGGATCCCCGCCCCCATGCCCGCCTCGCGGGCGATGTTGGAAGTCCTCGACTCCTGGATGACGGTGC

CGAACAAGATGTAGTCGATGTCCTTCGGGTCTAGCTCCGGGTTCCTGTTGAGCAACCCCCTCATCGCCAG

CTTGCCCAGGTCCACCGCGAGAAGATCCTTGTACGCGGTCCCGCTCATCACGAAGGGGATGCGTGCTCCG

TCGACGAAAACAGGTCTTCTGCTGCTGCTGCTGCTGCTG

>NODE_484_length_319_cov_1.536913

ATTCCACTCCCGCCCCTAATCCTCCTAACCCTCCTCCTCCTGCTGCTGCTCGTGCACACCCAGAAAGCGC

GCCAGCTTCGCTTTGCCTTTGATGCTGCCGCGGTGGTTCTCGTCTGCAGCAGCTGCCTTGCCTGGCAGGT

TTCCTATATATATCGACTGGAGGTAGCCTCGAGCCTGGTGCCTGTCTGACTCTCTTGACTGTCTGTCTGT

GTATGCCTCAGTACGAGGCGAGGCAACAAAGTGCGGAATATTCGCAAATGTCTCGATTTCGCGACGCGAA

CGCTCTGCACTCGTGCGTTGCCGTGCCAGGTGGTAGACT

>NODE_485_length_318_cov_5.444444

ATCACCATCCTCCTGATCTGATGCTGCTGCAGCAAGCACCACCTCCTGCTGCTGTTCCTCGTCCTCCTCC

TCCTCTTCTTCTTCCTCCTCCTCCCTCCTCCCGATCTTCGTCGTAGATTCATTTCAGCGCGACGAGGAAG

AAGGGAAGCCTTCCGTTCGCTATCAGAGGAAGAACAGTCGCCCCCGTCTCCACCACCACCACTGCCTCAA

TCTTGATCAAATCATCTCCTGTTCTTCCACGGGCCACCTTCGCGTACCCACTCATCATCCTCCTGGTGGC

CAAAGGCGCTCTTACGGCTGCGCTTGCGAGCAGCACCA

>NODE_486_length_318_cov_5.090909

GAGGCGGAGCAGCTAATGGAGTGCGTTCAGCGCCCTGGTGACGTCATGTACATCGCCGGCGGTTGGGGGC

ACACCTCGATGTTCATTGACAACAACACGTTCGGATACACTCTTGGACTGATCAACCAGAGGGACACGCT

CTCGTCGGTCGCTGGCGTGGCTTGCGAGCAGTAATATGCCGTATATTTATTGGCTGTATGTGCCCTATTT

GGAGCACAACTCTGGAGAACGGTGGAGGTACTGGTCGAACTGGATGCACTGTAGTACTGGTAGGTCGAAC

AATATCGGTGCAGGCTGCAACAACAACAACAACAACAA

>NODE_487_length_318_cov_3.131313

TTTCCGCGACTGTCGAACCCAATTTGTTAAAGAACGCATCCCCGACAAACAGACCGTTAATTGTGCCTTC

ATGTCCCAGATAGTTGTACGGCATGATGGCCTCGGCCCCGAACTCGTCAATGATCCCTGTCCAACGGTCA

CGGATTTCAGACAGCGCTTCGTCCCAGGCGATCCGTTCGAATTGGCCGCTGCCCTTTTCCCCCACGCGAC

GCATTGGGTAGAGGATGCGATCCGGGTTATAATGGTGTTCAGCGAAATCCTTTAGCTTCACACACAGACC

GCCACGCGTAATTGGGTGATCTTTGTTGCCAGTGACCT

>NODE_488_length_317_cov_94.912162

TTGGCCCCGAGAATTCCACTGTGCCCGTCGTGTCCTCTATCGTTATTCCCCCTTCGCTGAGCGTCTGCAT

CGTGCTCATCACCAATACCTGCACTCGCCACAATCCACCCATTTCGAGGGACAACATACTCGTCTTTGCA

CTACTAGGTGCTTCACGGACCCCCGCAACGAACAGGTCAGCAGCCTTGGACAGACGTAATGAGCATGTAC

TTCTTAGGGTCTAAGGATTGCTACCTGCCGTAACATGGCACCGAAGGAGGGGCTCGAACCCTCGACCACA

GGATTAAAAGTCCTGCGCTCTACCGACTGAGCTACTC

>NODE_489_length_317_cov_4.925676

AGCTACCGTGGTCTCCGTTTTCTGTATATTCATCATCTGTTTGTGATTATTCTCCATTTTGTGTTTCTTC

GTTTTTTGTGTGTGATTTGACACTACATCACGCTGCACGTTGCCAGGTGGCAAAAACAATACCAGGTGCG

TATAATACCAGCAAACAGCAAGTAAGTCAGTGCTAGAACCAAAATATTTTATTTCTGCCGGGCTCGCTCG

TGGGGCGGATGGTGAACGCTGCAAGAGAGAGGGATTTCGTGCCCGCAGTTTTCTGATTGGTCAATTTTTT

TGGTTGGAAATTTTGAATGGTGGAATTCTCGGTGCCA

>NODE_490_length_317_cov_2.530405

CGAGCTAAAGGTAGCTCGCTCAATCGCTCAAGTCGTCACTGTCAATATTTAGGTTGCCCGTGACGGTGTC

GTTGCGCCAGGCAGTCTATTCATACAGGGCCAAAAGAAGTGCACGTGAAACTTCACGCCTTCGGGACCCT

GCCAGGATCTAACGCTACGTACTATAAACAATGCCAAGAGATTCACGGAAAAACTGGCTGGTGGTCCATA

ATTAACACCAAAATGTCAAGTTTCCTTACGAAGTGGCAGTCTACTTGGCTGTGCGTTTCTGATCCGTGAT

GTCGTCGGTCTTTCGTCCGCACATCCCGCATATGCCT

>NODE_491_length_316_cov_7.247458

GCTAGCGATCTTGATCGCTTTTCTTCTTGGGATATGTGTGAGAAACTATTAAGATGGATTTCGTATTTCG

TTCGGAGTTGAGTGAGTGCGCGTGGACAAGCGTTGGTTTCGTTGAGGGTTAGAATGCCTAGACGGTACGT

GCGTTTCAAGGACTGCCACATAAAACGTTTTTGTTCACAGCGCTGACACCCGATAAATCCAATATGGACT

ATTGGTGTGCGTGCTTGATATATCGTCGTAGGCGAAGGCACACGAATCGGGTGGGAAATCCTCTCTCCAA

GGTCAAAGTATACTGTTGTCCTTCCCGTTAGGGGAG

>NODE_492_length_316_cov_4.064407

TTGCGGAGAGGATGGAGTGGCACTCACCATCCCACCTTCATCCGGGTTTTTGTGCGTGTTTATGCTTGGA

TGTTGTGTGGTGGAAAGACGGTGGCCAGTACCACCGCAAATTGGACGCAGATGTGCGTCGGAGATGCTGT

TTTCTATTGCCCATATACCTATAGTATCGGTCAAGGGTGATGAAATTGGAGTCATGTCGGCCTTATGGGA

ATGTGAGAGTAGTCAGTCGAGGTTCAAGTAGTATGATGTACCACTCCATGCGTTGGGTCTAACAACGGTT

TTGAAATCGGCGAAGGATTTCGTTGTTTGCCACTAA

>NODE_493_length_315_cov_8.074830

CAGAGCCGCAACGCTACCCCACCTTTTATTTGATAAGAATTGGACGCCTCTAAATTCCAAACAAGTCTTA

GTGACAACAAGAAACTAGTAAATCCATGTAGTCACGTTTAGAACTTAGTAGTTCGGAGCACGGGAGGAGA

GAAAGCGAGGACTGGATGCAAGGATTCAACCCTCGTTTTCTTCTTGTAGCAAATGCCACTGGAATGCAAA

ACAGATTCCTTGTAGATGACTTGGTACGCTGGGGCTACTGTACGCGGTTGAGATTGCGTGTTCAATCGAT

CCGCTGAGGTTTTGGCTCTTTTAGTTTGAATCTTT

>NODE_494_length_314_cov_11.081911

TTATCTTTAAGGATTATTTGAATAGTGATGAGAGTTACACAAGGATGTTTTTCGTTTTTACCAGATCTAA

ATGATGAGCAAATCAAACAACAAGTTGCTTATGCTATGTCAAAAGGATGGGCTGTTAGTGTAGAATGGAC

AGATGATCCACATCCACGTAATTCATATTGGGAATTATGGGGTCTTCCTTTATTTGACATTCAAGATCCT

GCTGCATTAATGTATGAACTTGGTGAATGTAGAAAAGTTAACCCAGAAGGTTACATCAAAATCAACGCGT

TTGATGCTAGTATCGGTACAGAAAGTTGTGTATT

>NODE_495_length_314_cov_5.904437

CGGGCGTAATGTAGTATCCATCATATATCTATCTCTCCCATCTTTTCCCCTCCGTGAGCTCTTTTCTTTC

TTTCTCCGCTGCTTTACTATTTTTGCGCCATTTGTTTGGTTGAATTACTACAATACCACTGCTGTACGTC

GGGTACTTGAGGTAGAAGTCTTGCGTGGTCTGCGTTGTTTCTATGGACGGGGGCGTGTAGCAGATACACT

ACGGTAATATTATGCAACCGTTGTCTTTGGGTGGATATAGATTATGGCGCAACGGCAGCGCGATGTACAA

AAAAAGCAAGGGCGTCGGTAAAATATGGTGCCTG

>NODE_496_length_314_cov_5.194539

GTTGATTGATCACGCGCTCCGAGTGTTACTAGATGCCATCCTCTCCTCCTCAGCCACCTCAGTCCAGAAG

GGGTGGTCCGTTGATAGTGCGGACACCTCGGGCTCGTTGAACTTGTCCTTGGCATACTTTATGTCGTCGC

CAATCGCCACAATGAGGTCATCGAGAGAGCTAAAGTTCAGCTCGTTGCGGATGAAGCCGCAGAGGAGCAG

GTGCAACTTCTCCCCGTAAAAATCGTCCTCGAACTTATACAACAGATGTGGCTCCACAGTCTTCTCCTTG

TTGTCGAAGTACGGGTTCCACCCCACCGATACCA

>NODE_497_length_314_cov_4.907850

GGCGAGCTGGCGACGTTCTGCGACTCGAAAGACCGGGGGGAGTGGAGGGGCATGTTCCTGCCGTCCGATG

TTATCCTGGACACAGTCACCTACGGGGGCGTCGTTAACTCAGCCTGCCATGGCACCGGTAAAACTCAGAC

CGTGTGTGACTACGTCGTCAAGACCAGCATAATCAAGTGGGACGGTACACGTGTGGACCTCACCAGAGAA

GAAGACCCCGAGGGCTTCGCGAAGAAGATTGTCCACTTCGGTCTCCTGGGGGTTACCGTATCATTGACCT

TCCAGTTGGACGCCACACGAACTGCCGTCTGGAA

>NODE_498_length_314_cov_4.464164

ATGAAATAAACGGTTCCAAAGCAAAACAATACACACAAAAAAAGAGCAAAACAACATCTACAGCTTGAAC

ACTAATACGAACAAGGTGAGAAACAAGAGAACAAAAACACGACAAACCAAAAAAAAAAACGGGCACCGTC

CTCACGATCACCACCCGCGCACGAAACACTCGAAGGAGTACTACGCGCATGCTACCCTACATAACCCCAC

TTCCAAATAACTCTGGGCACCTCCCGAGTGGCTGGCCCCTTCCGAGAGCCAAAAACAAACAAAAAAAAAT

AATAAAAAAATGATTCCAATTAACTCCAGACAAA

>NODE_499_length_314_cov_4.331058

CCCTTTTTTTTTTCAAGCTGATGAGGGCTTACGTTTTTTTGTTTTTTGTGGTTTTGCTTTTGATGGTATA

TTTCTTGTTTAATTTATGTTGTTGTGATTTTTGTTTATTTTTCTTTCGGTCAGATCGTCGCCGCGGCGAT

TTTAATAAGGCGTTGTTGAGGCTTCAGTGCTTCCGATGCCTTGTGCTGTGACTTTAGTGGAAAATTAAGA

TGTGGGCGAAGAGAACGCGCCGGGTGGAAGGGGAGTGGAGGGGGGGGTGGCCTGGTTCGCAGCCCTGTAA

AGACATGCTATCTCTCAAAAAAAAAAAAATTTTT

>NODE_500_length_314_cov_2.747440

CTTGGCTCGCGAGAATTCCATGGCGTCAGGGTTGTAGGTGGAAGCGTAGCGCTCCCCGTAGCTGAGGTCG

AGCTCGGACTGGATCGGCCTGTTTGCCAGGCCAGGCGCCTTGACGTTGGTCTTGAAGTAGATCGATTCTT

CGGGCTTGAGCTTCATGACCAGCTCGTTACGCGGGAGGGGGATGTCCTCGAACGCGAAAGACCCAGCAGG

GACGTCCTTGAATTGGATGCGCACCTCTGCCTTGGTTTCGTTGAGAGCCTTGCCCGCCTTGAAGATGAAC

GGCACTCCCGCCCAACGGCGGTTCTTGACGTAGA

>NODE_501_length_313_cov_5.126712

GGCTTCTACCGACCGTGACGTTGGACTGCAGAGCGTACGGTTGCTGCCCGAACAACCTACTCTCTTATCA

TGGCGCCCGTACCTAAGCAGATCGCGGGGAGCTCCACCACAAAGCGGGCGCAGAAGGCCAGGCCGTCCCA

CCACAAGGGGCGTCTGGGCAAGCGCGTGAAGATGGTGAGGGAGGTCGTGCGCGAGGTGGCCGGCCTGATG

CCGTACGAGAAGCGCATCCTCGACATGATCAAGACCGGCGGCGCCTCGGCCGAGAAGCGGATCTACAAGT

TCGCCAAGCGGAGGCTGGGCACCCACAAGCGTG

>NODE_502_length_313_cov_3.626712

GCAGCTGTCTTGATATATTTTCGGTGGACGTGTGCTATGTACGATAGAGTAAGCACTTTTTTGGCTGATA

TATTTTCTGAAAGACGGGGCGGGGCGCCTTAACCCAAATTCGTTGGAAAGTTCATTTTTTGTGTTGCAAT

ATTTTTCACAATACATACTCAGGCCATGGGATTGCTCATCGGGTGTATAGTGCTGTTATTTTCCTCCATA

ATCGATTTTTGGGTGGGGTCTAGAGCTACTGTAAGTGAGCAGCGTCAGAAAAAAACATGGTGCACCTCTA

TCGTAGCATGCGTCTGTAGATGATCTGTAGATG

>NODE_503_length_312_cov_5.635739

TCCTACTCCAGCTTCTGAGGCATCGGTATGGACAACGAACTCTCGAGAAAAGTCCGGGTATTTAAGAACT

GGCGCAGAAGTTAGGGCGCGTTTAGCGCGAGCAAAAGCTTCACGTTGCGCGGGCCCAAAAGCCTTCTTGA

AAGCAGTCTTTTTGACAAAATCCTTACGAGTGAGTTCAACCAGCGGTGCTGTGATTTCAGCATAACCGTC

TATAAAACGGCGGACGTAATTGAGTGTACCTAAAAATCCGCGATTATCTTTGATACACTCAGGCTCAGGC

AAGGCAAGGATGGCTTTGACACGGTCGGTACT

>NODE_504_length_312_cov_3.962199

TTTTTGAAGGATTAGTAGTTTTAGGTGTGATTTGGAATAAAAGACACCAATAGGTGGTGTCTGAAGTTTA

GAGTGGAATGCATAAACACGTCACCATGATTTTTGGGTGATGTATGTATCGTATCGTGGTAGACATATAA

AAGGTGCGTTGTGGGTGGTAGCAACCGCGACAGTTTGTTTGGGCCAGTACACACGTGCGTGAAGAACCGT

TCAATTAGCACCCCGGCCAACATGCCTACGACAAGCGGCCTTCGCATGCGGGGCACTTTAGCAGTATCCT

TGTAAGTGCACCTACAGTAGTTTAGACGTGCA

>NODE_505_length_312_cov_3.408935

GGACTTCTCGCACTTCCCATCTTTACATTCCTCCCAATCGGCGTCGTGGATCAGAGTGTAACCCCAGGGG

CAGTCGTAGGAATCGCAGTAGACCTTCTTGCTGTAAGAAGTGGTGCGCCTCCCGTGCAGGCCCTGCTCGA

CGTCTCCTTCGGCCTTGGAAGTCGGGACAGGGGCGCCAGCTGTGGCACCGAGAGTCACGAACGCCGCGAC

GAAGGCAACAAATCGCACCATCTTTTATTAGTCGTACCTCTTGATATGCCAGTACTACAGTAGGATTGGG

TGGTGGGTGCTTGTGCCGAAGGAGATGATGCA

>NODE_506_length_312_cov_1.745704

CAGCGGCTTCCTGTGATTTCCATCGAGCAGGCCATTCGGTCCATGTTCGTTGTGCTTTGGCCAGTAGACC

AGTAATGCTGGAAGTGTGCGTTGAGGTTGAGACGTGACAAAGATGGCCGCCAGGGTATCTACTGTGGAGA

CCCATTGCAGTATGGGATGGGTCTCTTGAGCGGGGATGATTATACAAAAAAACACGCCGGAGCGCGCACT

CAAAACCACTTTCTTGTGGTACTGTTTATACAGTATAGCTTTCGGCTAGATGTTGTTGTTGCTGTTGTAC

CGTGATGAAATTCCATCAATTTATCCATTGTT

>NODE_507_length_311_cov_12.696552

GAACGTTCTGGGAGGGGCGTTTTGGTCTAGGTTTTTTGAGGATTTCATCGGGGTAGCGTTTCTTACTCTC

TTTTTTTGTTGTGCTTGGCGCTGCGGTTGTTGTCGCTGCTGCTTCTGTCGTTCGGCCAGAGGTTGTTGGT

CGTCGTTCTCTGGACGGCGGCCATGAGGGGATCATTGACGTGGCCCCTTCCCCCTCCACGCCGGCTTGTT

TATGTCTCGTCGGCTGGGTGGGGTTGGGTCAAAAAAAGAGGGAATCTGAGGGCCCTGCACTAATGTCAAA

ACACCAAAACCATAAAAAAAAAAAAAGTGGA

>NODE_508_length_311_cov_5.765517

TACGTACTGCTGCTGCTGCTTTTGTATGAGCTTGTGGAAAAGGAGGAAGGAGGCGTTTTTGTTGGCTGGA

AGAGCGTCCGCTTGTCGCCTGCTGTCACGTCACGACACTTGGGAAGGGGGGCGGGGCTAACGCTGCGAGT

TTTGTCTAGGGGTGCTAATTTTTTCGGGCTCATCATACGACGCGATTGTTTCCGTCTGGAAGATGGTTAT

TTCGTGACCGTTTCCGTCGGTAGAGGAGGGCGTGCTTGATTTTTCCGTCGGTAGAATGGTGGCCCTCTCC

GGTTGGCGGTGTCTCCGATTTCCGAGTGGCA

>NODE_509_length_310_cov_9.570934

GTATTTTTTATTTGATTTAGTTGTCTGCATAGTATTTGTAGTTGTTTTTGTAGTTCTTTTTGTACATATA

ATTTTATTTTTTTTAGTGCTCTGGTGTTAATCCCCCCGTCGCCCCCGCCAAAAGGCGGTGGAGAAAGAAA

GGAAGCTTCTCGTCGCGTCTTGACCCGTATCCGAAAAACACGTGCGCAGCGCAGTCTACTCGTCCGGGAC

GACGATCTCGCCTTCGGAGATGCCGGCCTGGATGTCGCGGGGGGACTGTCCGTTGACGGTGCACCCGACG

GAGTTGCACGTGCCGAGCATCTCCTTCACG

>NODE_510_length_310_cov_2.854671

GTCCCTGGAGCGGCAACCCCTTCCACATGCACTCCGACGCCCTGAACATGGTGATCGCGGGCAGGAAGAG

GTGGTACTTGCCAAGTTTCAGCTTGGGATCATATATTTTTAGGTTTATTCTGGATGTACAACTCAATTTC

AGTAGTTATATTCCATTTCAGTTGGAAAATGCAATCTGATGTTTGGGGATCAGTAACACCAACAGGAACA

GTTTCTCATATCAGTGGTGGTAATTTTGCCCAAAGTGCAATTACTATTAACGGATGGTTAAGAGATTTTT

TATGGGCACAAGCATCACAAGTAATTCAAT

>NODE_511_length_310_cov_2.730104

GAAGGTTGCCGGCGTAACGGTTGTGCAGCAGCTGGCTGAGGAAGTAGAGGTTGGACACGAGCGCCGTCTG

GAGGATGACGGGCATGTTCGACGTGTAGAACAGCTTGATCGGGTACGTCCCCTGCTGTCCGCGGTACTTC

TGGTACTTCACCGGCAGGTCGACCCTCCAGCCCTGGAAGTAGATGACGACGATGAACACCAGCGCAGTCG

CGAGGAGGTTGGTGATGTTGGGCAGGTTCTGGCGATAGAACGCCTCTTGCAGGGCGCGCATCTTGTCTGG

GCGAGTGATCATGAGGTGGAACAACGCGAT

>NODE_512_length_310_cov_2.629758

GGTGGTAGATGCCGTTGAGCCGCTTGACGTAGGCGTCGCGCATGTCCTTGATCTTCCCCCAGTCGAAAGA

AGTCTCGCCGATCGTGTACCCGTAGTGCTTGGCGGCGTGCAACATCTCGTTGACGTGGGCTGCGTTGAAC

ATGACCTTCTTCGGCACGCATCCCACATTAACGCACGTCCCTCCCAGGTTGGGAGTGCCCTCGATCAGGA

CAACCTTGAAAAGGTAGTCGTATTCGTCCTCTTTTCCGGCCATGGCTGCGTGTGCTTCACCCCCAATCCG

CAGAGAGACAGGGTCAGTTCAGGGATGGCT

>NODE_513_length_310_cov_1.733564

ATTGGCGTTGGCATCGGCTTTACGCTCGAACCGTCGCATCTTCTCCTCTGCCGCGATCAAGTGGCCTCGT

ATACATCCTGCAATCTCGTCTGACTTCCTCCCGAGCTCGTCGAACTTCACGTTCATGTGATCCTCCAACG

CACTCTCTAGACGCGCCAGCATCACCTCGAGCTCCCCATCGGAACAGGCGCGCGTCTTCTCGGTCTTCCC

CTTACAGAACAGGCCAACGCCGTGGAATCCATAGACCTCGTCCTTGATGAGCCCGTCCAGCTCTTGGATG

ACTTCAGTCAGAGCCCGCCCTGCCTTCTCG

>NODE_514_length_309_cov_13.246528

GCCGTCGGTCGTGCGGACGTCAACCTGCGCCTCGATGAGCGTCTGCCACTTCTTGATGAGGGAGCAAAGC

TTGTCGCGGGTCAGGTCCATTCCGTGGAAGTTGGTCAGGACCGAAGTGCCCTGAACATCCTCGCAGCAAA

GCTTCACCTTCCGGTAGCCGAGGTCCTCGTCGTTGTTCAGGTCCCCAAGCGAGAACTCGAACACGCGCCC

CTTAAGTCCCTCGGAAGCGATCTTCGTTCCCTTGGTACGCGTGACGAGAGTCTTGCCCGGGTTGCGCACT

TGGAAGATGGAGGGCGCCTTGATGTCGTA

>NODE_515_length_309_cov_5.107639

TAGTTATTTTGGTGTGTATTTTTTTAATTGTTACTCTAGTATTAATTTTGTTTATTGTCTGTAGTGTATT

GCATGTAAATAAAGAGATGAGTGTCAGGAGACAACTATTATATATCCCTATCGATTACGCGTAGAAGCGG

AACATTTTAATAGGAAATAAGAACGCCGAATCATATGCGTGGTGCATACTAAGGGAAGCAACCGGTCGGC

CGTGCTGGTTGCCATACATGGCCGTAGTCGACGCCGTAGATAGACATTTCGCATCCATCAGTACAATACA

GGGATGACTGGAATTCTCGGTTGACAAGG

>NODE_516_length_309_cov_3.902778

GTTGATTGCCAGGAAGTGTCGGTTGACTGCCACCCGTATCCGTTGAGTATTGGCAGATGTATCCGTTCAC

TGTCAGTGGTGTTCGTTGACTGCTAGTCTACCCGTTGTCCGTAGACTGCTATTATCCGTTGTCTTTTGAC

TGCCAGAAGGTATCCGCTGGTTCTTAGTGGTGTCCGTTGAGTTCCAGGAGGTATTCGTTGATTGCCAGAG

GTATCCGTTGGCTGCCGCTATGTCACTTACAATGTTCCACCTTGAGACGTGCTGCAATTGTCGGTTGCTG

ATTGGCTGGCCGCGCCTTGGTGATAAGCA

>NODE_517_length_309_cov_2.809028

CGAGAGTTACGCATGCTCACATGGCATGTACGGCCACACAAGTCAGGTGTATACGCAGTTGAACAAAGCA

CCGATACACATCCGACAATGTTTGGCGTCTAGTCCCACGAGTCAATAGGGCTACAGGTATGTATATGTGT

TGGTTTGTCTATACCGCGTAAGATTAGGACGAGCTATGAAAGCATACCGGTATAAAGCAGAAATTTTCGA

TGGGAGAGCTGTGACCTAATGTGGGTTTACATTCTGTACGTTTCTAGTGGCGTACTGAGGAAACGAGAAT

ACGCATGATATACCAAGTACTTTTTCTAT

>NODE_518_length_308_cov_10.473868

ATTCCAGGTGGTGTTGAGTAAACACCAGGTTGGTTGTGATGCCTTCAGCCAGAAGAACTGATTGAAAGCA

CGCAGCTCGGAGAGATCGAGTTAAAGCTGCTGTAGTGGTCGTACGTAGGGGTAGACCGCTTTACTTCTTC

ACGACCTTCTTCACCACCTTCTTCACGATCTTCTTTTTCGGGGGGGGCTTCTTCGCCTCGGCAGAGAGCT

TGTACGAGCTCTTCACCTTGACGAACTTGCCGGCCTCGGTTCCCTTCTTGAGGGCGCTGCGCATCTGGTG

CGGCGCAACCGTCGAACAGGAGGATGTG

>NODE_519_length_308_cov_2.944251

GCACGACCGACGTATCGTAATCGTACGCAGCTTGACTAGAAAAAAAAATAACTTAAAAATTTTTATGATT

TTTTACCAAAAATTCTCGACCTCTCACGGTATATATATATACCTATAGACGTAGCTGACATTTTGTTTCC

GTATTTATACATATACCAAAAAATGTAAGGGAGGAAGGAAACCATAATCAGAGCCCCAACCCCCTCTTCT

TTCGCGAAAGGAAGGGGGTGAAAGCGGATCGGACTACCGTATAGATCAGAGGAAGTCCACGCTAACGCTG

ACGCTGATGAACTTGCTCACGAGAAGAG

>NODE_520_length_307_cov_2.923077

CCATGCGGCGATAGGACATCTTCGAAAAGGAGTCGGCGTCGATCTCCTCATCATCAGGGTTAACCTCGTG

CGAGTTCCAGCGGGATGCCATGGTCAAAGACGTCTGCAGGCTCGCTCGTGAAAAACGTACCGTATCGTTA

CCTCACGATGAGAGAGGTTGATTGGTTATGCACCCTACCCCGTACCGTGACGAGCCCCGCCAAAAAGAAC

GAGTTGTCAGACAAAAGCACAGAGAAGGTCCGGTGCGTTTCTATTCTCCTCCTCTATTTCCAAGGCTTGC

AAGCACTAGGCGCTCGGAGAAAGCTGG

>NODE_521_length_306_cov_32.368421

CATGATCAAACTCTTCAATTATAAATCGTTTGTGTAACCTCATCGCAAGCGATGTGTTACTGCTCAATGA

ATTCTGTCGTGTTACCTACATAAGTAGATAACTACATAAAACGTACTAATCAAATTAATGATTAATACTT

TTTTGTGTGACATTCAACATAAGCTGTTTTACGTCTTCCTTAGTCACGGTATAAATACCAGTAATATAAG

CCGACTATGTAAAATCAATCTTAATGTGAGTGTTCACACAAATTGCCTGATAACGAAATTAGAAGAAGTT

TGAACAAAATAGGTCTGTAGCTCAGC

>NODE_522_length_306_cov_23.940351

AGAAAATATAAAATTATGACACCATCTTTAACAAATTTTTTATCTAGCTTAATTGCTGGTGGGCTTATTG

TAGTTTTACCTATCAGTCTTGCATTATTATTTGTCAGTAAAAAAGATGCTTTAACTCGTGTACCACCAAA

AAAATAGTAATTAAAAACAAAAATTTTAAAAGTTAGTTTTTTAATTTTTGTTTTTTAGTTTTTGTTTTCG

TTTTTTTAATGACAATGGAAGGTGGACGCGCGCGCGCGCGGTGCCCGACTCGACGGTTGCCTTTGAAGAT

GAAATGGCCGTGCTAATCTGAGCAAA

>NODE_523_length_306_cov_5.778947

AAGCCGACGCATGCTACCTTAAGGCCAGAGGTGCATCACCATGCATTTTCTGAGAGTGGTCACTTGAGGA

GCTCTATACTCGGCCCAAAAACTGATTTTGGAGGAAAATAAAAGTGCTGCCGGATGCGAAATCTCATGAC

CGAAGTATACTCTGAAACATTATTCGTCACAAAAATGAAAGTCCGACGAATTTGGGTTAAGGAGCCCGAT

CGTTCAGAAAACGATTCAGCCAAAAAGTGCTACCTCTATCGTAGCACGCGTCGAAATACGTGCCGAACTT

AGAGCCTACCAGATACACTACTCTTT

>NODE_524_length_305_cov_19.429577

ACTGGAGTTCCTTGGGACCCGAGATTTCCAGCGCCGAAACGAAAGCCACAATATGATCAAATTCTGCACA

ATTGTTCAAGGGAGCGGTGTGTGCGATCCAAGATAGTTAACTTGTATTCGTTCCACCAACGCTAATCGAA

GAGACCTCTCAAAACCCTATGAAGTGTGCGTGGAGGGCAAATTTACTCCCTTGGGTCGAGGTTGTACAGC

GCACACAAACGCCTTTTGTCTTGGAAGCAATTGCTGAGATCGTTCCTTTCTTCGTACCTCCAGTAGGTAG

TAGCATCAGTGATAGTTGCAGCACT

>NODE_525_length_305_cov_7.929577

CCTGGTGGTGGAGCTTTCTTGCGGTACACGCAAAACCCCTCGTCAAGATGGTGCGCATGGGTGTGCTCGC

GGACACTCTGAAGACCATCTCCAACGCGGAGAAGCGCGGAAAGCGGCAGGTGCTCATCCGGCCCTGCTCC

AAGGTGGTCGTGAAGTTCCTGCAGGTTATGCAGAAGAACGGCTACATCGGGGAGTTCGAGATCGTCGACG

ACCACCGCGCGGGCAAGATCGTTATCGAGCTGACCGGGCGCATCAACAAGTGCGGCGTTATCTCGCCGCG

GTTCGACATGCCCCTCCACGACTGG

>NODE_526_length_305_cov_5.630282

AGTAGTAGTAGTAGTAGTAGCGTGCGTATATCAGTAGGCTGGAATCGGCTTCATTTGATAATAATAATAG

CGACTGGAGTTTGGTGTGGTTGCAGACACTATTGTAGCCCGTCCATGCAGTAGGTGCGTATAAGGACAGT

AGGCTGACTTGATTTCGATCATAGTAGCAACGATCGCAAGTTATGGTCGGTTGGGAGTTGCACGTAAATA

CTCGCACCTGGAGTTGTGAGTGGGCTGATTTTTGGCCCCAATAACAATAATAATAACAATGATATTGATC

GGAGTCCTCAAAAAAAAAAAAATTT

>NODE_527_length_305_cov_3.767606

TGTATTGGGCATAAGTTATTTATGTTTGATTTGATTTTTTTTTGTTTTGTGCAGCAATAATTGACGAATC

GATGGTGTTGTTATTTCGGCGGATAACCCCGGTCGGTGCGGGCAACCACCAGGTCAGGCAGAACACCAGA

GATGGGTAGCCCTTCGCCCACCACCACCACAACTATTCTACCACAACTTCAACGACCACCACCAGCAAGA

CGCTGAAGCAGCTAAGAGGCAGCAAGCCCTGTAGGCTATCGAAAACATAGAAACATATACACACTCCTTG

TAGGTTAACCCGAGTGGAAAACCAA

>NODE_528_length_305_cov_3.697183

CGCCCCTAGGATAAGAAGAGCACCCAGACAAAGACAGAACAGCACCCTGAGCAGAGAGAGAGAGGGTTTT

TCGGTACGGGGTTTTTGTATAGGTTTTGTAACGTGTGTTTTCAATCATTTTTTCGATACGAGTACTCCGT

CAATGAAAATCGTTACGCGGTGTGACGTAGAGAGGTGTGGGTTTTTCATTTTGACGCTGTACGTACATAT

TAGGGGGAAAGGCACCACTGGTTTGAATCGATAGGGACCTTTTGTCCTCTGTCATTCTTCGGATAGATGC

CTGGGTGCGAGAGCGGGTTGGGGAA

>NODE_529_length_305_cov_2.123239

CGACGTGCCCTACGGCAAGCGTGTCCACGTGCTACCCGTGGACGACACCATCGAGGGCGTAACGGGCAAC

CTGTTCGACGTTTACCTGAAGCCGTACTTTTTGGAGGCGTACCGCCCGGTCAAGAAGGGAGACCTCTTCC

TCGTCAGACAGGCGATGCACCCTGTAGAGTTCAAGGTAGTGGAGTGCGATCCTGCTCCCTATTGTATTGT

CGCACCGGACACGGTCATCCACTGCGAGGGAGAGCCCATCAAGCGCGAGGACGAGGAGCGGATGGACGAT

GTGGGGTACGATGACATCGGGGGCT

>NODE_530_length_304_cov_948.710247

ACTCTAAACCGACACAGGTGGGTTGGTAGAGTATACCAAAGGGCGCGAGATAACTCTCTCTAAGGAACTC

GGCAAAATAACCCTGTAACTTCGGGAGAAGGGGTACCTATTAGGTTGCAATAACAAGGTCCAAGCGACTG

TTTACCAAAAACACAGGTCTCCGCTAAGTTGTAAGACAACGTATGGGGGCTGACGCCTGCCCAGTGCCGG

AAGGTTAAAGAAGTTGGTTAGTTTTTTACGAAGCTGATAACTGAAGCCCCGGTGAACGGCGGCCGTAACA

ATAACGGTCCTAAGGTAGCGAAAT

>NODE_531_length_304_cov_6.424028

GGCGGCAAGCAGCAGCTCCTCCGATCCGGATGGCAGCGCGCAGTACGGGCTGCACGCGTGCGGCTTAAAC

AAGCTGTGGAGCTCTGTTTTCGAAGAGGAGCATGAAGATCTAGCTCGACTGCTGGATGGCGACGACGACG

ATGACGGGTGCTGACGATCTGTATGGAGAGGGCTTACCATCACGAGGAAGTTCGTAGCATCAAGGGAACG

GAACGGACCCTGCTCGGACTGCGGCAGGCAGAGCAGCTGAGCTCGAGAGGAACCAACTAACTGCTGCCGA

GGCACCGATGGAATTCTCGGTGCC

>NODE_532_length_304_cov_5.007067

CCTTGGCAGCCGAGATTTCCAGTGTGCCACACGTTTAGTGCCTTATCGACGGAGTACGAAAGGTATAAAC

AATTTATGATAATACTCCTCACTGAATATAGGTGGTAGTCGAGCCCACGTCACAGAAATTATGTTTAAAA

AAAATAATAATACGAGATTGGGAATTAGTTTTGAATAAACCCAAAATTCGCAAGTATTATATCAGGTCGT

GACAGATAATGGGCATAGGCAGTGCAAGGCCAGTCGTGCCAGTACATTCAAAAGAAGTACAGAAAACAAC

GGAGCTGCTGGCACCAGACTTGCC

>NODE_533_length_304_cov_4.540636

TCTGCATGGCGCCCCGCACCTTGTAGGAAGTTTCCTGGACAGCCGACTGAGCCTTGTTTGCGTTGGACTT

GGCCCACTTCAGGGCTTCGGGGTCGTCTATGTCCTTGGTTAGGTCGCGATCCGCGCGGCGCTTGCCGTAC

TCGCTCGCGTTAAGTTTGCGCAGCTCCTCGTCCCAGTCCACCGTCATCTCGAAGTCTGGGTCTTCCCGAG

GGTTGACCTGCATCACCCAGGAGCATCGCTGCCCGAGTGCAGAGCGTCCTTTCGGTGCCAGCGATCGCGA

GGGCGATCGATGAGCTGTTTGCAA

>NODE_534_length_304_cov_4.279152

CCGTCTGGTCTGGACCTGCCGGAAACTTTTATCCGCGGCGTCGACATCGACTCAACGGGTTCCGACCCCT

CTTCGTCATTTACACGGTGGGGTCTTGGTGACTACGAGTCTGACATTCTGCTGTGCACCTCGAATGAGCC

GGCGGAGAGCATTCACCCAGCGGAAGCCACTTGGTCTTGCAGCAGCGATGGCAGTGGAAACGAATTGGTT

TCTTCCGAGTGCGGTGGTAATTGTGAGGACAGCGACGCTGGCGAGACTACCTGTTCGAGCGAGACGTTTT

TCATAACGTCGTCGGCGTTCCCGG

>NODE_535_length_304_cov_4.021201

GGAGGAGAGCGAGACAGGTCGGAGATTGAAAATAGGGGTGGAAAGGGCGAATGAATGGTCGGGCGAACGG

CCTGAAATGGGTTACGTAGGGTTTTTGAGGGGAGGGCTTGGCGGCTGACGGCGTCATCAATCTGCACATG

TTCAACTTCACCTTCAGCTATTACTGCTTTTGTTTGAGCCACGGAGAGGGCTGGTGGCGTCATAAAATCC

CACAACTTCAACTTGAACTTCGAAATTCATCCGCTTTTGTCTGAGCTACGTACCTACTTGCGAGGAGTGA

TGGTGCCGTACATCCACTCAGCTT

>NODE_536_length_304_cov_3.770318

AACTTAATTACTTGTCAAACCAAAGATTATCTATTTTTAGGGGCTAGAACCATCACCATTTGACGACCTT

CCAGTTTAGGGAAAGATTCGACAGTAGAGATTTCAACTAAATCGTTTTTAACGCGAGTTAGTAGTTCTAA

ACCAAGTTCTTGGTGGGCCATTTCACGACCGCGGAAACGTAATGTTACCTTGATCTTGTCGCCATCCTCT

AAAAAGCGCTTCAGGCTGCGTAGTTTTACCTGATAATCGCCATCATCAGTCCCAGGACGGAATTTTATTT

CCTTCAACTGAACCTGTTTCTGAT

>NODE_537_length_304_cov_3.420495

TACGACGTACAGACTACCTGGACGTACATATACCGTATATGACACTGTATACCACACCCCCTCGGACGAA

TTTCAACGCTGATTTTCTCGACGACCGGTCGAGCGCTAACCCCGAGCGACCCGTCTATGGAAAGATCTCG

ACGAGATGCTCCGAAAGCCGCCTTTTTCGTTGTCTGTGCCCCCCTGGTTATGGAGAAAATTTGCTTTGAA

ATCCACCCCAGGGGGTGTGATATACACTGTCATATACGGTAGTGTAGTGTAGTCAGCGCTCTACTAAAAT

TAATGCCAGGTATTTTAATGCTAC

>NODE_538_length_304_cov_2.752650

CGCACTCGTTTCTCACAGCCTCCAGAAAATACCGAGAACATTCCGAGGGCACGCACTCGTTTCTCACAGC

CTTCAGAAAATACCGAGAAACGTTCCGAGGGACGCGGAAAGAGACAAGTCGTCGGCGTAGTATATATATT

TTTTATCCGGGATGGGATTCGTGTACTATTAATAGTTTTTGCCTGCATCACACGCTCATGTCTCATGTGT

CTTGTCACAAGAAGCAAGGCTAAACCAAAAACGCAGAGGCTCTAATCAACCCTCTCTGCTTTTGCACTTA

ACCCTGTCAGGACTGCAGACCCGT

>NODE_539_length_304_cov_2.646643

GTCAATGTTAAATCACCATTTATCTGGTTTATTAGGTTTAGGTTGTTTAGCTTGGTCTGGACACCAAATC

CATATAGCATTACCTATTAATAAATTATTAGATGCTGGTGTTGCTTCACAAGAAATCCCTTTACCTTATG

AATTCATTATTAATAGGGAATTAATCGGGCAGCTTTACCCAAGTTTCAAAAAAGGTTTAGTTCCTTTCTT

TTCATTAAATTGGGGTGAATATTCTGATTTTTTAACATTTAAAGGTGGATTAAACCCCGTAACAGGTGGG

CTTTGGTTAAGTGACTAAGCGTAT

>NODE_540_length_304_cov_1.893993

TGCTCGCTGGTGAAGGGCAACGACCAGTTCGTGAACAACGGGTGCGAGAACTGCCAGTTCCTGCAGATGG

AGAGCGACCAGGCGCGTGTACAGGACTGCACGACCACGTACTTCACCGGGCTCATCGCGATGGTGGACCC

AGAGACCAGCTGGGTTGGAAAATGGCAGCGAATCAACGGGTGTAGACCCGGAATGTACGCGATCGAGGTG

GTTGGAGAGCTCCCTCAGGAGGTGCTCGAGTTTCTGGAGGAGAAGCGCATCCCGTTCCACTGCAATCCGC

CAGAGACTGGCTCATAGAAACCAC

>NODE_541_length_303_cov_218.836879

TTGCTTGATCGGAGGTCCGAACGAAAGTGGTACGGTTTCCAACGCCGACCTTTTTTACCGGGTCGGCGTC

GGAGCGCCGGAATGAGGCAGGCATCCTGCGCACGTTTGGTGAGTCGAGAGTCGCCGCCGAAGCGAGCGCC

GTCAACAACTCTGTGCGGCAGGGCGCGAGAGGCTTCGTTCACGAGATTAGGTGCACTCGGAGCATGGGAA

CTCCGGAACACTCAAAGTCCGCCGCCGCGACCCCCTCGTCGTGAGACAGGAGAGAAGAGGGGCGAGTGGT

GTCAACAGACACTCCGACAAGCA

>NODE_542_length_303_cov_40.624113

AGCAGCAGCAGCAGCAGCAGTCTTTCTTCTCGCGTGCCTCCCATTCAGTGCCTTGACCCCCACCTTATCA

CACCAGTAAGCTCGAGAACAACAAGATCAGCTATCTACAGGCAGGTGTCGGCATCTCTGCTACTACAGCG

GCCAGCTGCAATCGAAAGAAGAAGATCGTACACACGCGCCTCGCCATGAAGAGTTTCGTCGGGTTGACTG

CCCTCCTGGCGGCCTTTCCTCCCTGCCTCGGCACCGGGGACCTCAACGAAGTCCAAGCAACACAGCCTCT

GCTCAGCGGGAGCGTGTGCAGGG

>NODE_543_length_303_cov_22.294326

AGCCATGACGAAGGGAACAAAGCGCACCATCTTGTTTTGAGCTTTGAATCTGATAGTGATGATGCTTAAA

GTGTTGTCGATGAACTGGGTGCCGTCCGACTCCTTGAGCTTGATGCTCTCCTCGCCTCCGACGATGGTGT

TGCCCTCGAAGATGTTGTTCTGGGAACGACCGGAGTCCACCACATCGGGCGCGTCGGATCCAAGGTACGA

GAAGGTGTTGTACTTGGAAGAGCCGCTGATGAGGTTGTTGGAGAAGACGTTGTCGCCGCATCCCACGGAG

AACCTGATGCCGTACTTGTTGTC

>NODE_544_length_303_cov_15.099291

AAGTTTCCCAACTTGTTCTCTTTATAATCTTAATATATATCAGGTTTATTAGCCTCCCCTTATATATGTC

CTACAACCCCGTGGTACCAAAAGCTACCTTGTACAGTATCCACAGTTTGGGCTCTTCCCGTTTCGCTCGT

CACTACTCAGGGAATCTCAATTGATTTCTTTTCCATTGCTACTTAGATATTTCAGTTCACAATGTGTCCT

TCTGTTTTACAGATAATATGAATTACTCATATTAGGTTCCCCCATTCAGAAATCTTGGAGTAATAACGCT

TTTTATCAACTCCCCTGGACTTA

>NODE_545_length_303_cov_7.042553

ATAAATTTACGGAGAAATAATCATATGGCTAATATAAGTAAAATATTTGGGGTTGACCTTGGGACGACCA

ACTCCGTTGCAGCAGTAATGGAAGGTGGACAACCCACAGTAGTTAACAATACGGAAGGATTAAGAACAAC

ACCATCAATTGTTGCTTATACTAAAAATAAAGATTTATTAGTTGGACAAATTGCTAAACGACAAGCTGTT

ATTAATCCTGAAAACACTTTCTCGTCGATCAAACGTTTTATGGGTTCCAAATTCAGTGAACTAAGCAAAG

AGTCAAAAGAGGTTTCTTATAAG

>NODE_546_length_303_cov_6.500000

CGAAATTGGTCACGCTTGACGGCAGGACGGAGCCGACGCAGCTAGTGGCGTCCCCAGTAAGAGTGTCCAA

GTCCTCCGTGATTTGGTTTAGCACGGTCAAGGTGTCGTTCGGCAGGTTCAGAATCCTCTGCAAGTCGCTG

GCGATTGTTTGGGTGCGCTGCTCGATGTTGGTCAAGCCGGTGTCCGTCTTCTCCAGGCCCCAGATCGCTC

CGCACGCGGTGGCTATGGCCGCCACGCTAGCCACCAACATTAGTAAGTTCCCTTTGTAGTGCATCTTGCG

CCCGAATTTCAACCTCTCGTTAA

>NODE_547_length_303_cov_3.893617

GTCAATGGTTGTGTTGCTCTCTACCAGGGTACAGTGTGTTTTGTGTCGAGTCCGTCGCAGCGATAGTCGC

AACGGCCAATGGCACCAGGAGTTCAGGGTCCCACACTACATGTGGTATGCGATCGTCAAGGTTTCCTACC

GATAGTTTTGCGAGGTTTGGGTGCTGTATGTATGTTTTGCAAGGTTTATGAGACGTGTAGTTTTGGTTGC

CCGGAGCTAGTTACGAAATCAGTCTGAGCGTGTTCTGGACTGGACGCTACAGTAAAATAAAACATTTTTT

TTATGAAAAAAAAAAAAAGTTTT

>NODE_548_length_303_cov_3.184397

ACTGCGTAACCGTGGTCTTTCACACACTGGTAGCGCTGTAAGCTTCGGTACTTTCGGATTGAAAGCAACT

GGCCGTGGCCGTATGACTGCTCGTCAAATCGAAGCAGCTCGTCGTGCTATGACGCGTCACGTGAAACGTC

AAGGTAAAATCTGGATACGCGTTTTCCCTGATAAGCCAATTACCAAAAAACCTCTTGAGGTTCGTATGGG

TAAAGGTAAGGGTTCAGTAGAATATTGGGTTTGCCAAATTCTTCCTGGCCGTGTTCTTGTCGCAGCTCCT

TCATGAACGTGTCGAACGTCTGG

>NODE_549_length_302_cov_16.644128

TCGCAGTCTCATTTGCGACAGGGGAGGGCCGGCGCGACCGTCGAAGCCCCTTCGGAGTGTTAGAGGCCCG

TCCCCTAGGGGGCGTGTGCCTCCGCTCCGGTGGTCCCATCGCGACAGCGCTAACCTTTGGCCCTTACCCT

TACTACAGACTGCAATACCAAACCCTTTTTGATTAGAGGGCTCGCCGTGTTGAGTTTGCCCGTCTCCGAT

TTTGTTTTTCCCTCCCCTTCACGGGGGAGAGTGGGAAGCAACTTCCACGGAATGCGAGCATAATTATTCG

GCGAGACATTTTGGAAATCTCG

>NODE_550_length_302_cov_7.992883

TCCTTTTCTCTCCGCCTTTTGTTGTTCCGCCGCATGTCTCTTTTTTTTTTCGCTTGGTAACGGGCGTCGT

TGCGCTAAAAGCTGGTGGATTCTCCGTCTTATTGTGCAGATTAAAAACAATTTGTACGATGGAATGGTTT

TTTCCTTTTTTTAAAAGCCTTTTTTGGTCCGCCCAACACTTTTTTCCAGTGACCCTCCCGCAGAAAAGAT

TGCGACATTAAATTTTTCGGTTTTATGGGGCTAATCCTACGCCACCGCCTTTTTCGACCATGGCGACTGG

CGCTTGAGCTCCATCCACATTT

>NODE_551_length_302_cov_3.854093

GCCCGTTCAGCCCCCAGTCGTACTCGAAATGGTTGATCTTCAACGCATCCTTGTGCTCGCTCACGTACGT

CTTGACGTCCTCGGGCATGTACGGCAAAAGGGTGTCGAGCACGGAGGCGTTTCCCACTTTGGTGGTGAAG

TCTTCCTCAAACCACAGAAACACCTTGGACAGAGTGATCACGCCAGCCGCCTTGTCAAGAGACAGTCCCT

TCGTGGGGTTCTGCAGGAAGAGCCTCATGTTCTCCTCCATGTCAGCCTGCAGCGTCTCGGGCTTATACGC

CTTGGGAGCGACGTCCGGGCAG

>NODE_552_length_302_cov_3.316726

GGCACCGAGAATTCCATCTTCTAACCCTTCCAACGAGCCGAAGAGGATCTGGGTGGCCTCATCCGATAAC

TCTAGCCCTAAGGTGCTCTCTGGCAACCCTTCCACGAGGCCCAAAGAAAAGTTGTTCAGGAGAGATGACG

ACATCTGTCGCACCTCACTCCGCTCGTGCTGCAGCCCTCTCAGCGACGCATCAAGCCACCCTGGAATCAC

GTCCGGGCGCACCATATACTTCCCCCCCCCGGGGGTCCCGAAAGCGTTCGCCGCCGTGCAAATCGCGATC

GACAAACTCTGTGGCGATAGAT

>NODE_553_length_302_cov_2.704626

TCCGGTATTCAAGAGTGGACGTTTTTTTTTTTTACCAGTTCAATTGTTCTCATCCATTAATTGTTCTTCT

CAGGACGAAGCTGCTTTGGATAATAACTTGTAAAGTCGAAGTTATTATTATGGTTCCAAATCTCTACAAC

AACAAAACTGACGATCGATCGACGCACCCGACACACCCGACCCCATGCCAAGCCCGCGGAGCCGAAAATA

ACGACTGGCTGCAATCCCAGCGTATGCCAACAGGTTTTACCCAACAATACCAGGTACAAATGAGAGTGTG

TAGCAATGTAGATAAGGGAAGT

>NODE_554_length_302_cov_2.540925

ACCGTGCGTCGTCTGTCTGGTCGCGCCGCTACTCCATCCGGTCACCTTAGCTCGTGCATCAGGTACCGTT

GCCTACACTCCAACTGAAATCAACTACAACATCGCTTGTGGCCACAAGGTATACATTTGTCCGGCAGAGC

TATCACACGCACACACCTACACGAAAACGGAAAAAAATATGCCGTCCGTAATCGCTCGTTATCCTTCCTG

CTTGCTGCTGCCGCTGCTTTTTCCACGAGGGGACAAGGTCAGAAATAAATCCGTACACTACCATACCATA

CAATACCATATGATACCATACC

>NODE_555_length_301_cov_4.696429

CCGCAAAGCCCCAGAACCGCCAGGCCTATGCCTGAATCTGTGGGTCTAGTTTGTGCAAGAGAAGAGGAAG

TTTATTGGCCGGCACGAGGAGCCGGAGGAGGGGAGGATACTTGGAAGCCGGGGGAGACGTTGCCCCCGAG

GAAGGAGGAGAAGGAGAGAGCGAAGGTGGGGGCTGGAGGAAGAGTCTGCATCGTCGGTTGGGTGGCAGAG

CACGGGCGCAAGAGGAGGCGCTGCCCGAAGCGAGAGAGAGTCAAGGAGAAATGATCACCACGCCGCCAAC

AAGGGCTTTCGCCCGCACGAG

>NODE_556_length_301_cov_4.678571

GCATAGCACTCAGCAGCATGGCTCCCAAGCTGGTCCTCACGTACTTCGACATCGCCGGCGCTGCCGAGGC

GGTTCGCTGGGCGCTGGAACAATCTGGTCTCGAGTGGGAGGACAAGAGGCTCACTAGGGACGAATTTGGC

GCTCTCAAGCCCAGCCTACCGAACGGGCAGCTGCCCGTGCTCGAGATCGACGGCTACGTCCTCCCCCAGA

GCATGGCCATCCTTCGCTACGTCGGCAAGCTCGGCGGACTGTACCCTTCCGACGACCTCGAGGCGGCGAA

GTGCGACTCCGTGCTCGACTG

>NODE_557_length_301_cov_3.960714

TGGCGATACGTTAACTGAAAATCACTTAGCATTAAACTGGGATGATATTGAAGATTCTGAAGGTAACGAC

TGGGCAACATTTGACCGTGACGTAAATAGTACTTTCACTGTAGATACTGATGGTGATACGACTGCAGACC

GTTTTGGTCGTTACTTAGTTGATGCGCCTCGTTTCATGCTTTACAACGATGTTGGTATCTTTACTATCAC

TAAGCAAACTATCGGTTATTCTGATACTGATGTAGCTCTAGATGGTGATGATGATGGTAATGATGATGAT

GATGACAATAATGAAAAAATT

>NODE_558_length_301_cov_2.210714

ACTCTCAATGGCGCTGTGCAACAGGCATTCGGGTTCGCGGGCGAGCAGCTAACGGTGCTAGCAGACGAGG

TGCTGGTCACGCAAGGGGCGTTCACCGACTACACGGATACCGATGGACAGTGGAATACGGTCAACGGGGA

CGACGAGCTGCTCGACAAGACCGTTTGGACCAGTGGCATGGTGGCCGGCATGTTCTGGTACCAGTACGAA

TACACTGGAGACGAGGCCTGGAAGGTCAAGGCGGAGAGGATGACCGAGGGCCTTGAGGGCGTAGAGGAAT

TGGACGACAACGACCTCGGCT

>NODE_559_length_300_cov_19.182796

TACGGCTTCGACCCCCACGACGACAGCGACTTCCTGACCATCCACGACAACGAGGTCTACAACAACGGGA

ACCACGGCATCATCGCCTCCAAGCGCTGCAACGGAGTGTCCATCCAGGGGAACGAGGTGTACGAGGGCGG

TAGTGGATCAGTCGGCATATTCCTGCACAGAAGCAGCGACGACGCCATTGTCAAGGACAACTACGTCCAC

GACAACGGCGACGCTGGGATAGCGCTGATGGAGACCTTCAACGCCGACGTGTCCAACAACAAGTTCGAGG

ACAACAAGTACGGCATCAGG

>NODE_560_length_300_cov_10.655914

TTTTTTGAAGAAGTATTAGGATTTGTAGTTGTTTGTGTTTTTGGTTTTCATTGGATAATTAGATTTTCAT

TTTTTTTTTCTTTTCTTCTTTCTTTTTTTTCTCTTTCTCATTGCTTTGGAATAGTCAGCTGGCTCTGCTT

TGGCTTTTGCCATTTCGCTCAGCTTTGCAACGGTGTCTTTTGTCCATGTAGAAGAAATCTCTGTGGATTT

CTCAACGGCGGACAATGCAACGCCGGAAAGTTTTTCGCTCAGCGCGGCGGATGTTTTGAAAGAAGCTTCC

ATGGATGTTGTGTCCACAGG

>NODE_561_length_300_cov_6.670251

TTCAAAACGCACTGCATTTTTTTGTATTCGCTCGGTTTCTTTCTTTCATTTTTTTTTTTTGGTTTTTGTC

TTTTTTCAGTGAGCTTTAATATTTTCCTTCCGACGAAGAGGAGGAGGAAACAAGTGAGCGCGGCAGATCA

CAAGGGGGGTCGCGGGACCGGGCTGGCACTCGTCAACAGCCTCACAACAAAAACAAACAGCCTATAACCC

CCACAGCCGAGACACGAAACGAAACGAAAGTCTCTCGATGATGGTGCCGCCTGCCATTTTTGTTGCTTGC

CCGCCAACTCCGTCCCCACC

>NODE_562_length_300_cov_3.817204

GTATGGTCGCAAGGCTGAAACCACCCGTACCCCACGGCGGCGGAGTGTATCCGACAGGCCGCCAACATGT

TCGTTAAGACCTACCGCACACCCGCAGTCAACAAGGCGCTCGAGATCGTCATGGCCGACCAGGCGTAACC

GTCGAAACGTAACTGTCAAACGGTCTCCCCATCGAAACGCAACCGTCAAACCGTAAACGTCGAAACGTAT

CCGTCGAAACGTAGCCGTCGAACGTGACCGTTAAAACCGTAGCCGTCGAAACGTAACCGTCAAATACGTA

TCCGTCGAAACGGTTGTGGA

>NODE_563_length_300_cov_2.401434

ATTCCACACCGGCGCACACGTAACGGCACAAGCACGTGCGCGACCGCATGCTGATGCCACGACTGGAAAA

TTCTATACACAGGTAACCGGGACCCCCTCCTTGCACATTAACGGGTTTCAACAAATGTTTGTGGAGACCT

TGAACAATGTAGCAAATCCAACAAGGCGTCGACGACCTCGTGGTACTGTAGCCTATCTACAACAGACGCG

TGTTCGTCGTCAAATGGCCTCCCGCCCTACGTACCCCGAATCAGCTCGCCTTCATGCGGCAGCCGCTACA

GCCCGTCAACATACGAAACA

>NODE_564_length_299_cov_10.248201

ACTTCGCTCAGTTATTATTATTATTATTATCTTTGCCATCCGACAATGTTGCAGCATGAGCTTCAACGTT

GATCACTCATAACATTCCCGAAAGCAGAAAACCACGGAAACAAAGGAAACCTACGCGCAAAGTCTAACTG

TTTGGCAGCTCTTTTATCTTTGGTTGTGTTTATGCTACGTATTATCTATCTATTATATATAGTTCGTCTG

ATGCCCAAAAAACAATACTTTAATCAAGTGATCTTTGATGCCCACTTTATCATAGCAACTTAGAGTGGTC

TGTCTCGAGCTCTTAACCT

>NODE_565_length_299_cov_3.859712

AAGATTCTTGTGCAGCGCCTGAAGGTGCCCCCCAGCATCAACCAGTTCAGCAAGACCCTGGACAAGAACC

AGGCAACCGAGCTGCTCAAGGTGCTCATGAAGTACCAGCCGGAAGACAAGAAGGCCAAGCAGCAGCGGCT

GAAGGAGATCGCCGCCGCTAAGGAGGCCGGCAAGGCCCCCGCCCCGTCCAAGGCCCCCAAGGTGATCAAG

TTCGGGCTCAAGCACGTGACGACACTCATCGAGGAGCAGAAGGCATCCCTCGTGGTGATCGCTCACGACG

TGAACCCCATCGAGCTCGT

>NODE_566_length_299_cov_1.593525

AGATCAGAATAAAAAGGCAATACGCGTTTTATCTTCAAGATATAACGCTGGAGAGAAGTTACCTGCACTC

GCGCCATCATCGTCGTCTGAAGATAATTCAAATGCTTGGTCATGGCTACTCATGCGATTTTCCATTTCCA

ACACATCTTTAGTGCTTACACCTAATGTTTCAGCAACTGTATTCACTTCATCGTTACTAAACCAACCTAA

ACGTTTTTTATTTTTTCTTGCTTGGGCTTTGTGCTTGTTCCTCTGGGTCTTTTTTTGTTTTTCTTGTTTT

GATTTGCTTGGGAGAATTG

>NODE_567_length_298_cov_673.750903

TGCAGAAAGGGGTCGAATCCGGTGACTTACCAGGTTTAACAAGAAAAGTAGTACTAAAAAAGCATAATGC

ACAAGGAAAAGAAACAGGTGACCAGAAAACCCCAAAAGCATCAGCTTATAAAATTAGATTGAATGAAAAA

CAAAAATTACGCTATAATTATGGGGTAACTGAATCACAATTATTAAGATATGTTAGGGAAGCAAGACGAA

GAAAGGGTTTAACAGGTTTTTTATTAATGCACCGCTTTCCCGCCCAGCCCACCCTATCGCCGGACAAGCC

GATTCCAGGGTCGGCCAA

>NODE_568_length_298_cov_81.180505

ATCAGCTTGGAAGGCTGAGGTAATAGCCAGTATACGACACCCGCTAAATTCTATCTGTTTATCTCTGTTA

AAAATGGTGGAGGGAGGTGGATTCGAACCACCGAAGGCGGAGCCGTCAGATTTACAGTCTGATCCCTTTG

GCCACTCGGGAACCCCTCCAGTATATTTACCTATTTCACTAACGAGTAGTGAATAGTATTAATGGTGCCG

ACTGCCGGAGTCGAACTGGCGACCTACTGATTACAAGTCAGTTGCTCTACCAACTGAGCTAAGTCGGCAC

GGAGGAGGTATGATCGAA

>NODE_569_length_298_cov_17.317690

ACCGAAGCGCAGCAATGAGCGACGGGGCAGCGGACGGGGCGGAGCGGAGACGCTAAGTTACTGAAGTGAA

GTAGAAATTCAGAGGTTCAGACGGCGGGGCGCGCTAGAAAGGTTTAGCGCATCTTGTTGGTTTTTGTCTC

TACGTTATTTGAAAATAGAATATCTGAAGAGATATGTGGGCGGTTTGGTCTATTTCGATGGATCAACGTC

TGTATATCGCGCTCTTAGGCTTTTGGCCGATGATAGAGTGTCAGCTTCACTGTTTGACGGACTTTCGGTT

AGAAGCATCAGAATGACC

>NODE_570_length_298_cov_8.844765

CATCCCACAACACTACCGCAACATTGCTCCACCCCCCAACCCTGCCGAAAGCTTCGACGGGGGGCAGGTA

GAACGTCGTCACACCAGAGACGAGAGAGAGCCAGAACGTCGCGTGTTGGCAAAAGGAGCTAGCTGGCCGA

CGTTTACTTGGACTTCTTCTTGGCGAGCATCTTCTGGTAGAACTCCAGCTCCTTGCCCTCGAGGATGTAC

CCGTCGCAGCGGCCAGACTGGCCGGGGCGGGAAGAGATGCATGCGATGAAGCGGCCCGACGCGAACTCGC

ACTGGTTGCCCTATATAA

>NODE_571_length_298_cov_5.436823

TGAAGTTGAAAAAATGATAAAAAATGCGGAAGAATCATCTAAAATAGATAAAGAGAAAAAAGAGAAAATC

GACTTGAAGAATCAGGCTGATTTAGTTTGTTACCAAAATGAGAAACAATTAAAAGAGTACGAAGATAAAA

TATCTTTAGAGAAAAAAGAACTTCTTTTAGAAGGTATTAAGGCAATTCGCGATATATTACAAAACGAAGA

GTTTGATAATGAAGATCTGAAAAATAAACTTGAAGAGCTACAAAAAATTTCCCAAACTATTGGAGAAGAT

ATTGCTAGTTCAGCTAAT

>NODE_572_length_298_cov_5.212996

AAAAAACGACGCTTAGCGCTACAGCCTTAGCCTCAAAATTACCTCGGGTCCAACTTCTATGACAGACTGC

TGTCAGTACAGTATGCCTGTCTGGCAACACACTTAACACGCGTGACAACTTCACGGAGCGGCGGGGGAGA

AAAACAGCGTCCCGCGACAATCATGGTTCTTATGGTCCTTCCTATGATATGTACAATGGTACCTATAATA

CTCGGAGTAGTAGAGGGAGTTACCACCCAATCGGCGTTCTCTGTATTGATAGTCGACACGATAATCAATC

GTAGGTTTGTTGACAGCA

>NODE_573_length_298_cov_3.306859

GCAAGATGATTTTCCTCATCTGGAGGTGCCCCGCCTCGGGTTTCGTCGACGCCACTGACACCTTCGTGTT

CAGCGACAACAAGAAGATATATCGCCAAAACGTGGCGTTCAGGACCGGCTAAATGATGCGGTATACCGAC

GGGGATTACGCTGCCTCGTCGTGATTGTCGGTACGGTGCGCGACTGTGCGCTCGGTACGTCGGTGAAACA

CAATTTTCTGTCTGTACGAGAGACCATCTCAATAGCTATAAATCGTATATATCAGACGTTGGATGGGTGG

ATGGATGGATGGATGGAT

>NODE_574_length_297_cov_23.184783

AGATTTTTTTTTTCTAGATGTAGTTGTGATTGGAGTTGTTTGTGTGACTTGATTTCCTTTTTATCTGAGA

TTTTCTGATTTTTTTCGGTCTTTCAGAGTGGTCGTATCTAGCGTCACAGGCTCGGAGCATGTACTTGCTC

CCAGTGTGTCGACGGTTTTGGAACTGGTTCGCCGTAGCGTAATGAACCAGTACCCCGGGCTCATGCCCAC

GCTTTTTCCGGGTGTTTTGGCCACTGTGCCCACCCCTTTTGTCTCGATTGATGCCATGTGGTTGTCTTTT

CAGACTCGAGTGTGGCT

>NODE_575_length_297_cov_9.865942

GGGTCCAGCGTGCGCTCCGCCTGGCGCTTGGCCATCTTGCGCTGCACGTGGTTGGACTTAGCCACGGCAT

CCTCCTCAGTCTTAGGGGGCGCCTTCTTGCCCTTCTTGATGCCCACGACGGTGCCGTAGTGGTTCTCGTA

CCACGACTTGAAAGGGTGGCCGTCGATCACGACGATGGCGTTCTTGACAAGGGTCTTGGTGCGGACCAGC

TCGTTGTTGCTGGCGTTGTAGATCACGTCGAGGATACGCGTCTTACGGGTGCACACCTCAGTGCCCCTGC

TGAAGTTGCCGGTGTCG

>NODE_576_length_297_cov_6.496377

TGGTGACCACCTGAAGGCGGTACGACCCAGCTTTTTAGCATTACTTTTGAAGGCTCGGGAGCGAATCCTT

GGTCCCGAAAAAACAAGCGGGCGGTTTCACGCACCCGCCAATCAAAAAGGAACCAAAAGGCATTCCACAA

AAGCAGAAAAAGGATATATATCCACCGACGAAAAACAGTCGAAAAGTCGAGGGGAAAAAAGGGGAAAAGT

CAAGAATCAAGAGCACAGCAAGGCCGGCACATCGCACCGCAGCGTCGCCCAACGCTTTCCCCACCGTGTC

TTTAGTCCTTTTCTCCC

>NODE_577_length_297_cov_3.568841

GCGGAAATCTGATTGAAGTAAGGCAGAAAAGTAGAGAGGGCGGCATAATAATGAGAAGAAGAGGCAGAAA

ATACACGAAAAACAAACAAAATCCAGAAAACCGGTGTCGCTATGGTCCAGAGATCTTATGGAGATGGACC

TGCGAGGCCGGAGATGGGATAGAACTGAGAATCCAAAACTAGTAACTCAGAAAATGACTCCATGACGAAG

ATAAATCAATAACGCCCGGATTGTAGCTACAAGCACAAGGTGGGCGTTAGCGATGGTGAAGGATTCCAAA

GTGAAGACCTAGCACAC

>NODE_578_length_296_cov_8.229091

ATTCCACTTTTTTTTTTTTTTCATTTTTGTAGTTGTTATCATTCATGATGACATCCTTTTGTCCCCGAGC

GGTGAAGTTTTAGGGGGGGTTACATCCCTGGTCCCCCCTTTCCCCCTCCGCCTCCCGAACGTAGCCCGAT

AAACCCCCTCTCCAACGAAAGAACGCGTTTTGTGAGGGGAAACCCCACAAAAAAACGGCGAAAGCCGAAC

CTGACAAGCCTCACAGTGTAAAGATCGAAACATTCGACTACACCCCGCACGCACCAACACCCTGAATCGA

AACTAGCACATAACAC

>NODE_579_length_296_cov_7.840000

CTTGGCAGGCGAGAATTCCAACTCTCGCGCTCTCTCTCTAATAATCGGCAAGTCGGGTTGTTACAAATAC

CCGCTAATATAAGAAGTCCGTCACTCGGTTAATCTACTTTTCTAAATCTACGGGTTGACCGCTTATGAGA

ATGTGCTCGTATAAGAACATTCCTATCTGCTTTATGAGAACTTATCTAGTTCCTATCGGGAGTTCCGACC

TTAAGGAACGTTCTAGTGGTTTGGAACGTTCTTCTAGGCTTGCGAAGCCTTCATGCCATTGTCTCTCTGA

TCTACCAAGGGGAACT

>NODE_580_length_296_cov_4.814545

CTTGGCTCGCGAGAATTGCACGCAGAGATCAACCGTCAGCTGAAGGGGGCCCACTGAAACTATGTCCGCA

ACCGATGGCCGCCCTTATGGCACACGTTAGCCGTAACAGCGTATGTTGGTCGTGTCTCCCTTCAGCGAGC

ACTGTCCGCCCTTCTCCAGATACTGCTGGTGGTACGTCTCCGCGCGATAGAATTGCGTTGCCTCTTTGAT

CTCCGTCACGATAGGCTTGACATGCTTGGCCTGCTCGACCGTGAATTCCGCCTGATAATTCTCGAGCTCG

TCCTCGGCGACAACTG

>NODE_581_length_296_cov_3.927273

CCCTCTCCCCTTGCTCAGCACCACCGGCAGCGGTAACTGCCTGCAGCAGGAAAGTAGATGGGACATCTTC

CCGCTTGGCGAAAAGGGCACATCTTCCCTCAAGCTAAAAAAGCGCACCGACCCCAAACGCCGCTATCTAT

CACGGAAGCAGCCATCATGCGCCGTCCTTTTTCCACACGGGGTGCGGAAAACCTAGGCGCACTGCTTGAG

GTTCTTGCGGCCGGTGCTGATCACCATGGGCTTCATGCCCCCGTTCGTTACACGAAGGGCGCAGTGCACC

TTGGACCAGCGGATGA

>NODE_582_length_296_cov_3.352727

TCGGCCTTGTGGGCGTAGTCCGCGTCCACGTACGTCTCCAAGTCGTACTCCAACTGCACTTCCTCTAGTT

TACTGTCCTTCCTGAACGTCAGACCTAAGTGAGCCGTAGCGCTCAGGTACTCGATTATCTTCCTAGCCGC

CTTAACATGTACCTCCTTAGGGTCATGAGAAAACCTCGCAACCGCCCGTACTGCGTTGGCGATGTCCGGC

CTCGTCTGATTCGCAATCCACATCAGACTCCCCACCATTTCGCGTATCTCGCGTCCACTGCAGGGTCCTC

ACCACTCACGTGCCTG

>NODE_583_length_295_cov_5.430657

GTGGTGGTGGTGGTGGTGGTGCTACTGCTGTCCGCTTGTACTGTTTCGACGATGCGATTATTTTTTCGGA

ACTGCCGACTTCTTTGGTTGCATTTGGTTGTGGGATTTGCCCTCTGTGCCTGGTTGCGGCTGGGCTGACC

TCCTTGCTTGCATTCAGTTTTGTGAGATTTTCCCCCTACAGTATGGCAGGTTGCTTGCTGCGCCCGATGC

CTGGTTGTGGGCTTTCTGCTTTGCCTGCATTCGGTTCGGTTGTTGGATTTGCCCCCAATGTCAGGTTGCT

TACAATCGATGGTGG

>NODE_584_length_295_cov_3.051095

CGGACATGGATCCCGACCGCCGCGTGAAGGAGGCCATGAACGAGATCAATGCCTCCAAGCGTCTCAGAGA

GGCCGCTACCAACAAAGCCGAGGCGGACAAGATCATGCAGGTCAAGGCGGCAGAGGCGGAGGCTGAGTCG

AAGTACCTGTCGGGCGTGGGTGTGTCGAGGCAGCGCAAGGCTATCGTGGACGGTCTGCGCGACTCCGTCC

AGACCTTCTCGGAAACAATCGACGGAACATCGCCGAAGGACGTGATGGACCTGCTCCTTCTTACCCAGTA

CTTCGACATTGGAAT

>NODE_585_length_295_cov_1.233577

GTTGAGGAGACGCTGCTTGTGAGCAAGGAGCTCATCCGAGTGGCCATCCTTTGGCACGAGCAGTGGCACG

AGGGACTGGAGGATGCTTCGAGGCTGTACCACGGGGATGGGAACGTTCCTGGCATGATGGATGTTGTCGT

CCCTTTGCACGAGGAGCTGGAGAGAGGCGCCACCACCGAGAACGAGGAGATCTTCCAGAAGACCTTCGGA

GAGGACTTGTCCCAGGCGTGGCTCTGCATCCAAACGTACCAGACACGCACGCAGGAGGCTCTGCGGCTGG

GATACGTCGCGCCCG

>NODE_586_length_294_cov_7.443223

AACCCGCTCGCCTTCTGGGGCTTCGAGGACTGCTTTACTTACCTGCGCAAGCACAACGTACCTTACCACC

CCCTCCACGACGTCGGATTCTCCTCGCTTGGCGACATGCAGAGCACCCGCAAGGTGCCTCTGGAGAACTG

GTTCAGCTACGGCGGGGAGCGGTCGGGGCGATTCGAGGGCATGAAGAACAAGGACGGCTCCGAGAAGACC

GAGTGCGGCATCCACAGCAAGTAGATAGCGTATAACAGCGCTGCCTGCGTTTCGCCTCGGCTGCAGTCGC

CAGGGGCGTAGACA

>NODE_587_length_294_cov_2.593407

CACCAGTGTGGTACGGGAGAGTGCTCTGTCTTATGTATATGAACTCGTATACTCGTACCTGCACATACAT

ACATAGGAAGTGCCCGGTAACATCTTGTCGGAGATCACCGTAAGGATCTCCCATGCAAGTATCACCAATT

CACCCTGTTCATAAGATCTCTGGTAAGTAGGCCCATGCAACATGACCAAGCTTCGCGAACTCGGGACGGC

TTTGAACACGAGCAGAGGCAGGAGGAGGGGATGTTATCTAGGGGCCTTCAGAGACCCCGTCCGTTTCTAA

AGCAGTGGTGCTTT

>NODE_588_length_293_cov_8.863971

GGTCGAACCTCGTCGAAGTCTTCTGGTCGATTCCGAAGAACGAGTCGTCGTCCTTCCCGATGGCGTTGTT

TGTCCGTTTAAATTTCTTCACGGCGCAGATATCGTTGTACTCGTCCCACGTTTGGTCCTCGAACTCTTTG

CTAAGGACGTCGGTGATGTCCAGGCTGGAGGCGGCAAAGTAGAAGATCTCCACTCCGTAGTTGTCTTCCC

AAAGCTTCAGCAACGGGTTGGTGTCCCTTTCCTGCCCCGGAGACTGCCCCCGTTCCAGGAAGATTTCCCT

TGTGCGGCACTTG

>NODE_589_length_293_cov_4.625000

GCTCCGCCGCATGACGCGGTTCGGTCTCCTCGGAGAGGACGAGAAGGAGCTCGACTTCGTGCTTCAGCTC

ACCACCGAGAAGATGCTCGAGCGGAGGCTTCAGACCAAGGTATTCAAGCAGGGACTGGCCAAGTCTATCC

ACCACGCCCGTGTGCTGATCAAGCAGCGCCACATCAGGGTGGGCAAGCAGCTCGTGAACGTGCCCCAGTT

CATGGTCCGCATGGACTCCGAGAAGCACATCGACTTCTCCATCACCTCCCCCTACGGGCAGGGCAGGCCC

GGACGCGTCGCAC

>NODE_590_length_292_cov_8.848708

TCCATTTTTTTTTTTTTTCATTGTATCAAAGCCCGGGTCGTATCATTATCATCGGCGACTGATCCTCGAA

GGAAGAAGCATCCACGCGAGGCATCCATAATCTCAGATCATAATGCCTCGTCACACCATATAAAACCCAA

TCTCCGTAGACCAGTATGAAACCCCCACATATAGGGAAACCAGTATCAAGCAACCGCATACCACAAACAA

AGTATACCGTGTTGGTTCGATCCCGCACCCGCGTCGCTTTTCGTGTTTTTTGTTTTTGTTTTTTTTCTCT

GTTTCACAGTCT

>NODE_591_length_292_cov_3.033210

ACAGGTCAAGAAGGCGACCCAGTTATTTACATTCAACCACGTCACTTAACACCAACTAATTCTGCTAACC

TTGCACAACGTTACACAGAATTTGATGCTTTTGAATTCTCTTCTAAATTAGTTGACGGTACTAACGATAA

CCCACTTTACACAGCTCCAGCATTTGCTGCTTGGTCTGACCGTGGTAATAGCTTAGGTGTTGCTTTCTCT

GAAACTATCGCTGTAGGTACTACTTCTCCAGTAGAATCTGCATCAGTTGACCTTACACTTACAGGTGTAT

TAAATGGTTTAA

>NODE_592_length_292_cov_2.147601

GCACCGAGAATTCCATACAGACTGTATCCGCTTGATCTCCTCCTTCGCGCGGATGAGTTCTCTCTTCAGG

TTCTTCATCTCGTCCTTGATGTACTCCTCCTGGATATCAAGGAACCCCATGTGCTTCTTGAGGGTTTTGA

GCTTGACGTAGACGTCCTGCGCTGCTGCCGTCTCTTCCACGGCTTGTCGTTCCTTCACAGCACCTCCGTC

TACCATCATCGTCATAACCTGTGCCGACGCCATGGCTCACTCTTGTCCCAGTTGCGCTTGTGCAAGCGGT

GCAGATCGCCTG

>NODE_593_length_291_cov_7.500000

AGACGCAACGACCAAGTCGGGCGGCTGCTCGAACAAGGTGGTCAAGTTCGACAACCTCATCAACCCCAAG

ACTCTGAAGGCCTTCGACAAGAAGAACCTGCGAAAAATCGGCAAGATCGCGGAAAAGCAGGGCATCCCCC

TCCCGGATTTCGTGAAGGCACTGAACCACCCCAATATTTTGCGACTGAAGGACGTCCACGAGAACCTGAC

TTACGTGAAGAAGAATGGACTGGAGAAAGAGGTGGTTATTTCGAAAGCTGTCCACATCTCTTGGAATTCT

CGGGCCAAGGA

>NODE_594_length_291_cov_6.792593

CACAAGTTCGACCTCATGTACGCCAAGCGCGCCTTCGTGCACTGGTACGTGGGCGAGGGCATGGAGGAGG

GAGAATTCTCCGAGGCTCGTGAGGACCTTGCCGCGCTGGAGAAGGACTACGAAGAGGTCGGCGCCGAGAC

GGCCGAGGGAGAGGGCGAGGAGGAGGACTTCGGCGAGGAGTACTAATCTAACTCCTCCGGCTCTCAAGCT

TGCGTGCGGCAACATCAATAGCGACGTCGACAACAGTTATAGCTGTAGTAGAAGCAGTATAATTAAATCT

AGCCAACCAAC

>NODE_595_length_291_cov_5.500000

GATCCAATTGCAGATGACACCGTATTAGTAGCTGACTCAGCGACAACGAACGAAGACACACCAATCACCA

TTGATGTAGTCGCGAACGATACTGACGTTGATAACGGTGAAGTATCCCCAGTTGATACGGTCACGCAAGG

CTCGAATGGTACAGTTGCGATCAACGATGATGGTACGGTAACTTATACACCAAACGCTGACTTCAATGGT

ACGGATACCTTTACTTATACCAATGTCGATGGCAACACTGAAACCGTGACGGTCACGGTTGATCCAATTG

CAGATGACACC

>NODE_596_length_291_cov_2.533333

TCAACGAAGACCAGATTTACCGCATCGACCACTACCTTGGAAAGGAGATGACGCAGAACCTGCTCGTCTT

GCGCTTCTCCAACACGATGTTCGAGCCCATCTGGAACCAGAACTACGTCCAGAGTGTGACGTTCACGTTC

AAGGAGGACTTCGGAACCGAAGGACGGGGGGGCTACTTCGACAGATACGGCATCATCCGTGACGTCATCC

AAAACCACCTCATGCAGGTCTTCACGCTGATCGCTATGGAACCGCCTATTCGCGTCTCGGGCAGCGGGAG

CGCCGACTACG

>NODE_597_length_290_cov_34.914498

CCCCCCCACAAAAGTAAGAAGGCCTCTTGTTCAGCGTGGAAGACGCACTCACACGTCGCGCTTATATAGT

CAGGCAGACGGTGGGCCGTCGGAACACGTCGGAACCCTTGTTTATTTTGCATCGTAGACGAGTACTAGAG

GTCGCGGTGTCTCGGGAAGGAATCGAGTCGATGAAAGCTACTCTCTTCTTTTGATGTAGAGAAGATTTCG

CGACGTGTGGTGAAAACAATAATCTACGGAACAAAAAAACAGAAAAAAAAAAAAAGTGTAATTATCGGGT

TCCAAGGAAC

>NODE_598_length_290_cov_11.944238

TGAAAAGAACTTTGAAAAGAGGGTTAAAAGATTTGAAACCGTTGAGAAGGAAGCTATGGGAGTTGGCCAA

TGTTAGACATGACCTACGAACGCAGAAAGGAAAGGACTCGCAAGGCCGGACCTGGAAGCATAGGAAAGTT

CTCTGCTAACAAAAGATAAGATGGCGTGCTGGCTGGGCGAAAAGCTGTGAAGATATTAGCTTCGCAGTGG

GGCTTGGCTGCGCGTGAGGGGCTTTTGCGATCTTGTCGAAACGACTTCTATAGACCAGTAATTCTCGGGT

CACAAGGAAC

>NODE_599_length_290_cov_5.591078

CTTGGCACCCAGAATTCCACTGCCGCCTCGTAGAGCGCGATGATGTTGTGTGCCCGCTCGAGAGTGGCCT

CCGTGTCGAAGCTCAGCCTGGCGTCCACCTCGGTGGAGACGTAGCCCTTGACGAGGGTCAGGATCTCCAC

ACCGAAGTTGACCGCGAGCTTGTCCATCGTCTTCGCCACTCGCTCGTCATCCGTCAAGCCTTCGAGGGCC

TTCCCGTACGCCAAGGCATCCTCTACGAGGTCCTGGTGCTCCGGAGCCGAGGACGCCTTGTAGATCAGGG

AGGGGTTGGT

>NODE_600_length_289_cov_243.731343

CACCGAGAATTCCAGGCAGTCAGACAGTACGAATAATAGTTCAGCTCCACCGCGTCGCTAAGAAAGCGGA

CGCGTCAGCCTATTTGGCGTATCTTCACACTGCCTGCTGCATCGACACGCCCTCATGCCCACGCAGCTGC

TGCGGCTTCACCAACGACGAAAGCGGCGGGGGCGTTACGCCAGCACCACCAAGCTCCTCCACAGGCTCAG

CAACATCAAGCTCCAACACGTCAGCCTCAGCAAAACCGAGTTCTTCCTCCTCAGCCTTGTGTGGAATTCT

CGGGGTGCC

>NODE_601_length_289_cov_4.570896

AGAAGCCGGACCCGTCCGTTCTTGGGTCGAAGATGGCGCCTCTGCAGGCGTGCGTCAAAGAGGCGGGGGA

CTGCCGAACTCGCGACACTTTTGAGAACCACACAAAGGCTGTCGGCGAGGCTATGGGGTGCCTCAGCTGG

GTGGTGATGTACCCGCCTGCGGGCATCCCGAAGGACATGGTTGAGCCGACCATCGGGGGGTCGGACTACT

GGGCCAACAAAGTCCGGATGCAGTACCGCAAGACCAACCCGGAGCAGGTGGAGTTCTGCGACGCCCTGAA

GGCGCTGCT

>NODE_602_length_289_cov_4.399254

TTGGTTTAAAACTAATGAGTCACGTTTAAACCACCATTTAACAGGTTTATTTGGAGTAAGCTCTTTAGCA

TGGACAGGACATTTAGTTCATGTTGCTATCCCAGCATCTCGTGGTATTCATGTTGGTTGGGATAATTTTT

TAACAACTTTACCGCATCCAGATGGTTTAACTCCATTTTTTGATGGTAATTGGAATGCTTATTCTCAAAA

CCCTGATACTGTAGAACATATTTTTGGTACAAATACGGGTGCGGGTACTGCTATTTTAACATTCTTAGGT

GGTTTCCAC

>NODE_603_length_288_cov_13.235955

GCAGGATGAGGAGAGGCTTGACACAGCTAGGCGAGAAAATGCGGAAGCAGGGGAGGGAGAGAGACAGAGG

CGCACGGTTCGTCTTGGACATGTACAGGCGCTTGAGCACTACCTTCGCGAAGGCGCTGTCCGTCCTCCTG

GCGAGGAACCTGAACAGCTTTACGAGCAGTCCAAGGTAGACATTGTTCGACACAGGTGCCGAGCGAACCT

TGTGGCCCACACGACGGCCACCGGCGACCAGATCAATACCCATGCTGAGAGAGAGAAGGCTCACACGCGT

GGGGTGTG

>NODE_604_length_288_cov_4.779026

AGGCGTTTTCTTGCTTGGTTTCCCAAAGTTTTTTTTTTTTTCCGTACGAGCTGCTTGTGAGAGAAAGATC

AATTAAACTACGTACAGCTTCTGATTATTACCCAGAGGGACAGGAAAACGTCTAATACTTGTAGGATTTT

TCCTTTCTTGGGTGGGTTCTCCGGGTTTTCTTATGTCCGTATTATAGTATACGTATCATAGCTGTTTGTG

AGATATCGGCTTGAGAGATTATTATGATGTGCCTTGTCGGCTGGTAAGACACGATAATGTTTTGACCACT

CCGGCAGG

>NODE_605_length_288_cov_4.483146

GAGGCGGAGTGCTGTACGAGCGTATACGTAGTCTACAGCAATAATAATGGCATTTTTTTTCGGGCGCTTC

CTCTGTACAACAACCGCCACCGCTACACCTGTCGCATAGTGAAACGGTACACCAATCACAGCCTATTGAG

TTGTAACCGCCTGCTCTGTGAAAGGGCACTTACTTCTACGTTTCCTTTTTAAGCCGACTGGTTTTACATG

TACAGCAAAATACAGCACCCCTTCCACCACAACCTTTTGCAAAATCCCGCCCCTCCCAATGCGACGACAG

CGCAAAAC

>NODE_606_length_288_cov_4.423221

CCATACATTGGACACATTGGACACAACAGATCTCCCTTTCTTTGATAATCTAGATCTTGGGGGCAGATAT

ATCCCTGATTTTATACGACCTACACGATCTACCGTAGTAGATCATGCTTATTGCGCGGTGGGATCTGTTT

GGCACGACTCTTGCCCACATTTTTGCCGGGAGGGATCTGTATGATACAGCTCTTGCACACCATGTCGCAA

CGGTAAATACAGGATTTCGATGATCTGTAGATGATCTATCTGTAGATGATCTAGATCATGATCTATCTGT

AGATGATC

>NODE_607_length_288_cov_3.423221

CAGCTTGATAGTAGACAACGCTCTCTAGCAGGCAGGCATGTAATGTTGGTGCCTAAAAAAAAAGGAAAAA

CTCAGCGGCAGACACGTATAGCAAGCGGCAAGCAGCAGCGGTTACAGCGAAAATAACCCCAACAAAACAT

CCACCGGCCGAGTCATATCTCGCTTCCAGCATGTACGTTGCATTTGGCCTTGTTCTTGTTGCAACGAGAG

CGGAGCGTACGTATCATTCTGCGTCGGGGATAGCTGTTGGTTCTCGGTATTATCTCATTGCTTGTTGTTT

GTATTAGA

>NODE_608_length_287_cov_9.853383

TTCCAATCTTTTTTTTTTTTATATTTTTTTTATAATTCTCTCGTTCTCAGGCAAGCGCCAAACAAACCAC

AAGACAAGCGGCAGCTCCGAACGCCGCACAAAAAAGTGCAGCACCCGAAACAAGTCGTGCGTGCTGCTAC

GCTGGTTTCAGCACCCAAAAAGCCACACACAGCCAAACGAAAAAAAAGACAACACAACCGCCCCCCCCGC

CCACATAAGGAAGGTCCAAAAGTGCTGATTTTTCCACCAAGCTATATACCGAAAATCTGCTCGTACTGCA

GGCAAAA

>NODE_609_length_287_cov_7.443609

TGCTGCTGCTGCTGCTGCTGCCGCTGCTGCTGGTGTTCACTGCTGGTGCTTGCTGCCGCGAGGGAAGACG

ATGAGGTTGGCCCGGTACTCTTTCAGGCGAGTCACGTTCGTCACGAACGACTCCTCCGACTTGTTGGTGC

GGCGGTGGTCGACGGCGATGCCGATGGTAAGCGCCACCTTCTTAGAGATGCCCGCCTCCTTGAGCTCTTC

AACGTTGAAGCCGCGTCCCAGGCGAACCTTGGAGTTGTACTTGACGGTCGGGCAGTGCACAGCCGGCCGC

AGGAGCT

>NODE_610_length_287_cov_3.007519

CAGCAGCAGCAGCAGCAGCAGTAGCAGCAGCCGCCGCCGTCAAGCGTCAAAGTCAAACCCCCAACCCATC

CCCAATAAACTACAGTAAGTGGTGGTATTATTAAAAGTAGTTGTTGTCTCGTTTTTTGTTGTTGTTCAGA

TTTGCGGGCTTGGCGCTGTAAGTAAGTGCGGCGTATCGATTCGCCGATGTCCTAGGTAACTGACTGTGGC

AGGACACACACTATCAAAAATGCTATATATACATATATAAGAACACCCAATCAACCGCTCAGTAAATAGT

AAATACA

>NODE_611_length_287_cov_2.721805

CGGAAACGTCCGTCTCCTCGTAGTCTAGCCTGGTTGCATGCACGAGGTTGTGCAGGTTGCAGTAGTTGAC

GCTGTTGTCAAGCACCGCGTACCAGGTCTGACTGGTTGAGAACGCCTGAGCCGAGCATGACCCTCCCTTT

TCGGTGAAGGTGAAGGTGAGGCTGTCCTTGTAGTGTTTCACAGACGTCTCGTGGACGAATTCAGCCTTAC

CGTTGGACACCTTGGACACCGTGTAACCGCACTTGTTTCCACCGTTGCAGCTGTCGAACATTGCGAGCTT

TTCATGT

>NODE_612_length_287_cov_2.466165

GAATCCACATCGTCCCCGAGCAGCGCATCCCCGGAAATAGGTGCCGATGACGAATCCACCTCATCGTCGA

GCAGCGCAGTCCCAGGACTTTTACCCTTGACGTATGCTGTGACCAGCGCCATCGCTCTCGCTTGCGCTCC

GCTCATCGTCAACTTATGTACGTAGAAGGCTGTGCAGTATCTGCAGTAGTTGAGGGTACAACTAATTGCC

GATATTGTAACGTGGTGATTCTTCGTGGTTCCGCGGTGCCGGTGTACAGCCCGTCGGTTGAGTGCATGTA

GGTGGGA

>NODE_613_length_286_cov_12.109434

AGATCCGGGTTCAATTCCCGGATCGACATCGACATTGACACCAAGGAGCTCCTCGACGCGCTTGACTTCA

AGGACCTGGCTGGCATCTCTGTGTCGGCCACACCGGCCCGTGTTTGATTTATTGACCAAATTGGGATGTT

AACAACCGCGCCTCTGGGACAAAAATGATTTTGACTGGTTTTTGGAAGGCTTCGTTGTCGGGCATCTCCT

GGGAGTGTAGTGTAGCCCAGCACGGTCTGCCAGCGCCCTCTTGTTTCGAGGCTCAAGAGAAGGAGCTTAC

GGGTGC

>NODE_614_length_286_cov_8.388679

AGCTTGGGAGGGGTCATCGAGGGCACTCCGTGGTACTCGCGCAGGTTGCGAGCGGCCCTCCTGTAGATCC

TCTTGGTCTCCTCGGGCTTATTAGCGTCCTCCTCCGTCTTGTCCTGCGGCTTGCCCGCGGCAGAGATCTC

CAGCGGGCTGGGGCACAGGCGCAGCGCCAGGCTCAGCTCGGAGTCCTTGTCGAACACGATGTCATCCCCG

TCCGAGTCCATGTAGGTGAAAGACAGGTCGCTCTCGGCGCCCCCAGCCTCAGAGCGAGCCGCGAGCTCCA

CCTTGA

>NODE_615_length_286_cov_7.294340

CGCCACGCTAAATGCGCGAAGGAGTTCATGGCACGTCCTGGAGCTGCGTTGCCCGTGGCAGGGCTTGGCT

GGTCTACTTGGCGGCCTTCTTCGCCTTGTGCTTCGCGATACGGCGACCCGTGCCGCGGATCTTCAGGCCC

CTGTTCTTGGCCTCCTTGATCTGCTTGCGGGAGGTCTCCTTCTTCTCCGTGAGACCGGCGCGAGCCAGGC

GGTGCTTGGGCTCGAACTTCTTCATGTCCTCCAGGCTGTCGTAGATGAGGCAGAACCCCGTGCTCTTGCC

GCCACC

>NODE_616_length_286_cov_6.750943

GGTGTTGCTCCCCACGGAGGCCAGGTGCTCCTGCACGCCCTCTAGCTCCGCGAGCGGCACCTTAAACTGC

TTGCGGATTCGCGCGTAGGCGCCGACGATGGAGGCGGTCATCTTCGACGCGGCTACAGACATGGCCGGGA

GAGAAATACCGCGGCCCTCGGCGAGGCAGTCCATGAGCATGTTCCACCCGAACCCCGCCCTTTCGCGGCC

TCCGATGATCTGGTCGAGGGGGATGAAGACCTCCTCTCCCTCCACAGTCCCGTTCATGAAGCTGCTGGCG

GATGGG

>NODE_617_length_286_cov_3.313208

GGTTCACGTGCGAGTCCAGCAACGAAGCGGACGCAAGTGTATCACCACCGTCCAGGGGCTCGCGGACGAC

CTCGACCTGAAGCGTATCCTCAAGGCCTTCAAGAAGAACTTCAGCTGTAACGGCGCGATCGTGAGCGACG

CCGAGGGTGGGGATGTAGTGCAGATGTCGGGGGACCAACGGACGAACATCCACGAGTTCCTTGTCGACCA

ACAGATCTGCGCGGCAAGCCAGGTGGTGCTCCACGGATTCTAAGGGTCAACCGCGGGCCGTAGCGCCACG

TGGAAT

>NODE_618_length_286_cov_1.969811

TGCTCTGCTAAACTGCTTACGGTTTCTTCTGTACGTCGAACGACATAAACAGGTTTACCACGCCATTCTG

CACGGATTAATTGACCTGGCTCTATTTTACCTACATTGACTTCTACTGGAGCACCCGCAGCTTTCGCGCG

AGCACTTGGATTCCACGAACCAATGAAAGGCACAGCCACACCGACCACACCGGCACCACCAACAACAGCA

GTAGCCGCCGTTAAGAAGCGTCTGCGGCCGTTATTTACAGGCACATTGCTCATTCACACTCTCCAATTCA

TATTGA

>NODE_619_length_285_cov_3.655303

AGGCGGTCGTCAACGCCACCGCGGTCATGGGTTTCCTCAACCGATTTATGGACGCCTCGGGCATGACCCT

TGAGATGGAGGCTATCGAGACGAGCCTGGCCAAGCTGGCGCCGAGCGGCTGGACTACTGGCGTGGCGTAC

GACAAAGATGCCGACCAGGAGTTGATGGAGGAGGACCGTATCGAGGCGCAGAAGAGGACGGAACGGACGA

AGAAGTCGGGTTTCTTTTCTTACTTGAAGATGATTTTCGGGGCGAAGTGGGCGGACTCCAAGTCGCTGGG

CAAGA

>NODE_620_length_284_cov_53.452471

TTAGGGACTTCTAAATTATTCCCTCATTAAAGGATAAAAACTATTAAGAGCTTGTTGCTGGCTTCGGAGA

ATCTCTATGTCTGAGAATGTACAGGAAAGAACTCGATTAAAAAGTGAGCGTTACGAATCTGGTGTAATTC

CGTATGCCAAAATGGGATATTGGGATGCTGATTATAACGTTAAAGACACTGATATTCTAGCTCTATTCCG

TATAACTCCACAACCAGGCGTAGATCCCGTAGAAGCTGCCGCTGCTGTTGCGGGTAATGCTGAGAACTCT

AAGG

>NODE_621_length_284_cov_10.045627

ATCGATCGAAGAGTGTAAAGGGATAAGGAAGCTGATGTGTCGGAGGCGCGGACGGAAGTGGACAGTTTTT

CCGGAGACCGCAGCGCTGTCCCACAAACATGGGTCGACAAGAAGTACACTGGTAGCCTCGGTGACCTCAA

GGCGATGGGCGCCCTCGGAGTACAACTCAACGGCCTCCCCCGTGGCAAGGCAGCGGTAGACGCTGAGAAA

GATCTCGATGAGTCGGCCCGAGAGTCCACGGATGCGGCCGTCGCCTTATTCCCTGGCCCGGAGGCGGCAT

TTCT

>NODE_622_length_284_cov_4.634981

CATCGCCAAGGCCGGCCTCCTCGACAAGATCCAGGTGGTCTTCGTCGACACCTACTTCCTCTTCCCCGAG

AGCGTCCAATTCCTGCACGAGATCGAGGAGCACTACGGCTTCAAGGCCAAGGTGTACCACTGCGTGGACT

GCGACACTCAGCAGGCGTTCTACGACAAGTACGGTGAGGACTACTGGATGGAGGACATCGACGAGTACGA

CAAGAAGTGCAAGGTGGAGCCCCTCATGCGCGCCCTTAAGGAGGCCGACACGGAGGCGTGGATCAACGGT

CGCC

>NODE_623_length_284_cov_3.235741

GTCCCCCGATGGAGAACGTCAAGTTCAGATCCGCGAACTCCTCCCTCATCGCCGCAACCATCGTCTCTCT

GATCTTGTGCATGTTGTCGTATTGCTCGAACGCATCTCGCTCCTGCTGGCTGCAGTTCCTGCCCACGGGA

GAGATGTTCAACATGCCTGTCCGGAACTCGATAAAGGTACCCCTCTTGACCGGAATGTCAATGCCCGACA

TGTACTTCAGCACCCAGTTGATGATCTTCTTCAAGTTATCCTCCCCCACGTGCTTGGAGATCGTCTGCTG

TCCG

>NODE_624_length_284_cov_3.220532

ACACCACAGCGAGGTAGCTGTTTTTCATTTTGTTTTCGTGGTATATGCGTGTGGTTGCGTTTGGTCACTA

CGGCATGCCTACTACTGCCTGAGATTCTGCCGGGGTTGCGTTCCGTTCATGCCGCTGATCGTGCGTTTTC

GTCGGGTGAGTTTCAGGGCTGTTGTCTAGAATCGGGGGTTGCTTAGAGCCTTCCATATTCCCATCAGAAC

CGGCACACGAGGCTCAAGAGGTACACGACAAAAATAGTGAAGTCATCGAAATCTTGTTCCTCTGGAAAGC

AGGC

>NODE_625_length_283_cov_36.213740

TGGCAACGACGCTGTAAAACACGCCACTCGCTGCGTGAGCTAGACAACCAGGGTGCCGTTTTTTTTTTTT

TGGCGGTCACCAAAGCTGGCGGTCGAGTAGATCATTTGTGCGGGGTGGAGAACCTTTGCTCCGCACAAGG

GGAATGGTCGTATCACGTAGTGTTAATATGGTAAGGCATCCGCGAGGCAGTTACATAGTGCTCCCATTGC

TGTCTCCGGAAGTGGCCACACCAGAAATTAAATTAACTCATTGAAGCTGTCGTCCATATGCAAGTCCGCC

TCA

>NODE_626_length_283_cov_4.595420

CTCCCGGTCACCACGACGACCACGGCGACCATCCCAGGTCGACGTTTTCCGTGCTGTGGGACTGGCTCTT

TGAGGACTCGTCGGACTCGTCGTCTTCTTCCGAAGACTCTTCCGTGGACTCTTCCGTGGACTCGTTCGAG

GACCGATTCGTGCCCAAGGTGGAGAGGTTGGCGGATAAGGAGGGGGTGTTCCGGCACCAGGTGATGCAGT

CGGGACCTATTGTCAAGCGCGCGCAGGAAAAGGTGGTGGAGGCGGAGGAGCGTATTGTCAAGCGCGCGCA

GGA

>NODE_627_length_282_cov_7.325670

TGCGCACTTTTCCGGGGGTTGAAATTAAATTCACTTGTTCATCAAGACCACGTATGTACAGTATGTACGT

ACGTACGCGTACGAGAAGGCAAATGAATCAGAATGACGAGATGAACTTTTGAACTTTTTTTGTGCTTTCT

TGCCTATCTGCTATGCGTTGAGAAGGTCGTACGAGTAAATAGCAGTCGGGATCGCAACGATGCACCTTCG

ATCGTTGTCTTGCAGATTGGTATGCGTGTGTTGTTGTGTCGAGTAGCAGCAGGCTTCGGCGTTGGCAAGG

GC

>NODE_628_length_282_cov_4.015326

TCGGCATGGCGCGCTTCGGTGCCTCTGCCCCTTTCAACGACATCTACCCCAAGTTCGGCTTCACCGTGGA

GAACCTGACCGTACAGGCTCAGAAGGTGGCTGACTTCTACAAGGCTAAGGGTGGCGCCCCGTCCCTCATC

AACACCGTCGACCTCAACCTCGCCGGCTACCACAACGGTCACTAAAGGGACAATAGAGGGACGCTTCTTT

TGCAGGCGGCAACTGTTCCTATTGTAGCTTTCTCGGGGGATATTTGGTTTTGATTGCTGCTGCGATACGG

GC

>NODE_629_length_281_cov_5.842308

AATAGGTTGCACGTTATCTAGCGGTATGCATCGGTACGGCAGATAATATTTGAACACGTGGTTTAAACTG

TAGTACCCCCAGTAGTCACGAAGATCGTGCATCCCTTCTGAAACCCGATGTCGGCGGTTTAAAACATCGC

GGTCGTTATGGCGACGGTGTTGGAGACAGTGCTGAAGCCACAGTGATGAGATCGACGTCGTTGCCATCGA

ATATTTCGCCACCCATCAAGATATAATACAAAAAACATCAACAAAAGAAGCAATTCAATTCTCCCAAGCA

A

>NODE_630_length_281_cov_4.284615

GCTTCTTGGAGTCAGTTTCGATCTTGTAGGTATGAAGAGTGTCGAATGGTGCATAGAGGAGGGTATAGAG

TGGGAGGTATAGCATGGGAGCTGGCTCGGCGAGGTATGAGTGAGTTAAGTGCGCGCGTACCGCAAGGCAA

CTTAAGGATGGACTTTTCCGCCTGGGTTGCGTCGAGCAGGGAGCAGCGAAGGGACTCTTGTATGCGTTTC

TTGTCGAGAGAGGCGTTACTTGTCGACTGAGGCGTTTCTTGTCGAAAGGGGCGTTGCTTGTCGAGAGAGG

C

>NODE_631_length_281_cov_2.157692

GCTAGTTCTAGATATGGAAACTAATGGTACATTAGATCCAGAAGAAGCTATCCGTCGTGCTTCAACTATT

CTTGCTGAACAGCTTGATGCGTTTGTTGAACTTCGTGATATTAAAGAAGTAGAGCAAGTGGAAGAAAAAC

CATTGTTCGACCCAATCTTACTTCGTCCGGTGGATGACTTAGAACTAACTGTTCGATCTGCTAATTGTTT

GAAAGCGGAAGCGATTCAATACATTGGTGATTTAGTACAACGTGCTGAAGTTGAGCTTCTTAAAACGCCT

A

>NODE_632_length_280_cov_8.996139

CAGCGGGTGGTGCGCGCTTTCTTGATCGAGGAGCAGAAGATCGTAAAGCAGGTGCTCCTGGAGAAGCAGA

AGAAGAAAAAGGCTTAGACGAAGCAGCTGTGCTTTTGCGGTGGTAGCCGTCTTGCTTCCTTTGGTCACAT

GTCGGGGCTTCAAATGATAGGAACATAGGTCAACTTCGGGCAATTCGACTTTGGCTTCCCCATCTACGAG

GCTCCTTCATGGCCGCTCCGCGCCCGAACTGGAAATAAATAAACTCTGGAAAAAAAAAAAATGGAATTAA

>NODE_633_length_280_cov_7.204633

CGATGGGCTGCCTGTCCGGGAGCCGTCCTTGGTTCCCTCAATCGCGGACGAGGAGACGATGCGATCTAGA

GCAGTAGGCGAGAGGCGGCGCCAAGGCAAGCTGGAGCTCAGGGGGGCGTGCTACAAGGACGCCAAATTTG

AGGAGTGCCACACGGGCCAAGGTAGAACGGCCAACGCGCTGCACCTGGAAAGATGCATGTCCAGTGGACC

CCGCCTGTCGAAGGCTTGCGCGCACGGGATCAAGCAGTGGAAGAAGCTGCTTGCCGAAATGTGGCATGAA

>NODE_634_length_279_cov_14.003876

AGGTTAATATTCCTATACTATATCAAGGAGCTGTTTTTTGAAGTGGTAACACAAATGATCTCAGAAACAT

CAGTAGAATAACCGGGAAGAGTTCTCTTTTCTTTTTAACAACCCCATTCAACTTACTGTAAAAAACCATG

GAATCAATTTATTTGGAGAAATGGTTTGAAGGTTGGTAGAGCAGTCTATTTTTTGGATTGTCCGGTTGTA

TTCTAATGATACATAAAAATCTGAGGGAAATATTTACTTTCTTGAATAGTCGTACTCATAACCGCATCA

>NODE_635_length_279_cov_3.492248

TAAGCTCAGGGAGGAAGGCTCGGCGAGTACGCACAAGAGCAAGCCCACCTCTTCTCCCCTACCTCAGAAA

ACAGTACTTGAGAAGCAGCACCTGTTAACATGATATGCTACCCCTCAACTTTGCCACGTACTGAGGTGCT

CATCCATTGCCTCTCGCACCCGGTCAGTCGGACGCAGATCCAGCGACCCAGCCTTCGTGGCCACGACCAG

AAAATGACGCCCGGTAACCGCAGCCTTGCGCACCACGCTGAACCCCGTGTCCGCTAGGAGGTGGTGTAT

>NODE_636_length_279_cov_3.058140

AATCATCAAAGATCCTGTTTCTACAGAACCGCTTGTTGATGCAAACTCTAATAAGTTCTGGCCATTACAC

TCAATGCCGTCGAAAATACGACGAATTTTTAATTTATTCATTTTTAAAAATGAACGCATGCGTTTCTTGT

CATCAGCAGCCACATACTCACATATGCTTTGTGCAATGTTCGTCGCTTGTGCTTTAGGTGTCGATACAAC

AGACGTTAAAGTTAAAATAGTGATGGTTGAAGCTATTAATAATTTTTTCATTTTAGTAACCTTTAATTT

>NODE_637_length_279_cov_3.007752

GACCACGGGTTGATCTACGAGCACAGCATCGACAACCGGATCAAGCGGGACATGTGGGAGTACTCTCACT

CGTGGATGGGGATGCTCAAGGCGGACCGCCCCCCCTCGGCGTAGTGTGTATGTACAGCCGGCAGTAGAAG

GGGCAATCGCGGGGTCTGACTGTTGCCTTAAAGGGGTGACCTGCGTAGGCTACTATTTCGGGTGGGCTTA

TGTAGTAAGAAGAACCCCGGACCCAAAAAGCATGTATTCCACGATGTTGACTGGAATTCTCGGGGTGCC

>NODE_638_length_278_cov_10.000000

TCGACGCATACTGCGCTTGCTTGGCTGTCTACTACTTCCCTTCCCAGTTTGCCGAGATGGGAGCTATTGT

ACTGCGCGCCGACGTTTTATTCAGGTCGGAGGTCGACGACGACGCAACCTACGTGCTCGACATAACCACC

CTCTACGCGGGGGAACCCGATGTCGAGTACGGCGATACCATATCTTTCGTGACGGCAGCCAGCGTTCACG

ACTGTGGAGTGGACCTCAAAATCGGGGGCGACTACCTCCTCGGCCTGACCCGCCGGGCGGCAAGCAGC

>NODE_639_length_278_cov_2.451362

GTTACTACCACTTCACAAGTGTTCCATTATCAGGTGTAGATGGTTCTAATGATCCTAACTCTGCATCAGC

TTACACTGTTACAATTACAGGCCCTAGCGGTTATGCTAGCGGTACAGTAAGTGTTACAACGAACAATATC

CAAATTGATTCTGGTAACAACGGTACTAACACTGAAGATACTACTAATAACGGTGAGCAAATCACTTGGT

TTGACGGTTTCTTAGCACAAGCAGAAACAGCTAACTTACCTATGTTCTCAGCTCAAGTTACAGGCGTT

>NODE_640_length_278_cov_1.692607

GTATACGAAGTATACGTTACGACGAAAAAACTTGACTTTATTTCCCGCACATGCCCCCAACTGTAGGCAT

GGCATGCGTGGTACCATACAAGTATGCTGCATGTATCGCGATTGGTGCATGCATGGGATGACGCCCTGTC

CATGGTGTGAGCGGAACTTCTACTGTAGCTCCGGCACCTTTGCTTTCGTGGTCGGCGCGGGGGTAGAGGG

CTTGGGTACAATACGAGGTAATTTTTTTTCTATTTGTTTTTTTATGATTGGGTCGTGCCGTGATAATT

>NODE_641_length_277_cov_14.625000

TCTTCTTTTTGGTATGGTATTAGGTCTTATCCCAGTAACTTTAATTGGTTTATTCGTTGCTGCTTTTTTA

CAATACCGTCGCGGAAATAAACTTGGTCTATAAGAAAGAGAGATAATATAATTATTATCTCTCCTGTTTT

AAATCATCAATTACCAACTAGCTTTCTTAAGACCTGGAACTTCACCACGCATCATAGTTTCGCGTAATTT

AATACGGCTTAAACCAAATTTGCGTAAGTAACCATGTGGACGTCCAGTAACATTACAACGATTACGT

>NODE_642_length_277_cov_8.390625

CCGCAAGGAAATTTATTCCACCGCGCCCCCCGATTTTTCTCACGGGGCGAAGGTCTATGAGCCCTTGCTA

CAGAAAGCGCAAGCGAAACCCAACTCACGTTGTTGTTTTCTTGCGCGGCGCGACTTCGCTTTTCGGGATC

GTCCTTTCCTGAACATGCACCAGTAAACGTTGAACCTGACACGCTATTCTTCAATGAGTGGTGGTGTTGG

GCGAGGCGTTTTTCGGTCGTTTCGGGTTTTTTCATGGACGTCTCCGACTGCGAGCCCGCTTACATTT

>NODE_643_length_277_cov_6.242188

CCGCCCTCCTGGGAAGCTTCGCGAGAGAGGAGGATGTAGCCGCCCATGCCCCAGTCCCCTCCCCAGGAGT

TCTTCACCTTCCAGTACTTGATCCCGTCCTCGGAAACGCCGTACCCAACTAGCAAAACCCCGTGGTCGAG

GTTGGTTCCGCAGCTGGCCGTCAAAACGCCTCCCGAGTACAGCTGGAAGCTCATCTCGTCCGCCTCGATA

GCGATGGAGACCGGCTGCTGGGCGACCGCCGCCATCAGCGCCTCTTGGGAGTCGTGGTCGACATCGA

>NODE_644_length_277_cov_3.464844

GGGATGAGGGGGCGGCAGGGAGGAGGCAATGCAGCGCAGCCCGGGTCCCGCTACCTCGTTGTGACCCCGC

CGACAAGGCACATCTCGACGCCGGAAGCGAATCCCGGACGTACGTGGGCGAGCTCATACGATAGGTTGTG

ACGCCTTTAGAGCTTATTTAGGGTAGTGGCATGGGGACGGATTATTATGACTTAGACTAGGACCTGGTGT

GCCGCAATCGAAAGGTTGACTCACCTTGACCGTTCCCCGGCGGTGGCTTTTCATAGCCGCGAACATT

>NODE_645_length_277_cov_2.738281

CCCTCTGCTCATTCCATTCCATCCTACTCTCTCTCGCACAGAGCGCTGCTAATAAGAAGAACCTTAACAA

CAAGAGGAGTGTCCACACTGAGTTCGTCATCTCTTGCAAAGCGTGACCTAAGCTTCACAAACCCCTTGCC

TACATTGTACAAGAACACGGGCGTGATTCCAACGATATTGAGGGCATTAGTTTTAGCTGAGTACAGTACG

AAGCCCCGCCCCATAAATCATAGCCATTTTGGCCAGAAGGTGGCACCTTCTTTTGCGACACGCCTCC

>NODE_646_length_277_cov_2.738281

AAGGCGGCTGTACGCGTTTTCTTATATCCAAATACTGACTGCAATTCCTTGACAAGTTGATTCTCAAATT

CAGTCCCCTGATCGGAATGCAGAGTTTCTGGAGGGCCAAACACCGAGAACACGCGTTCAATCAAATGACG

AACAATCGTGCGAGCCGCTTTATCTTTGATGGCTATCAAAATTACAAATCGAGTGAGATGGTCAATGACC

GAAAGGATGAATCGATTTCCTTGTGACAAACTCTTCTACTCGACAAGATAAACTGCAACGCATTGGA

>NODE_647_length_276_cov_1067.388235

TCGGACAGGATAACCGCTGAAGGCATCTAAGCGGGAAGCCCCCCTCAAAACAAGATATCCCTGAGAGCCG

AGGTAGACTACCTCGTCGATAGGCCAGAGATGTAAGCGTGGTAACACGTTCAGTTGACTGGTACTAATGG

CTCGATAGGCTTGATTTGATCCAGTAACAGAGTGACTTAACACCTCACTGTAACTAGACAATCAAAAGCA

TACACACACATCAGTTGTACTGACTTGGACATCTTTAGGAATTTATTTGGAATTCTGGGTGCCAAG

>NODE_648_length_276_cov_6.219608

CCAGAGAAGGACATGGCGGTTTCAGAGGCGGAAGCTATCAAGGACCTCGCGGACGAGGAAGTGGAGGACT

TGCGAGAGACGTTCCGTACATTCGACAAGGATGGGAGTGGGGAAATCGACCTGAACGAGCTGCGAACCGT

GATGACATCGCTCGGATATAGCCCCACGAACAAGCAGTTGGAGGACATGATGGCAAAGGTGGACCTTGAC

GGCAACGGTCTCATCAACTTCGCCGAGTTCGTAACGATGAAGCGCAAATGCAAGGTCGACACGGAT

>NODE_649_length_276_cov_6.121569

ATTCCAAATTTTTTTTTTTTTTGGAGTAGATAGAGTACTATTATTGTACATCGGGATTATATTTTGGTCG

AACGAATGTACGACTGTGCGCGTTCTTCCATATGTTGCCCCACCGATGCCCCCTTCCCTTCGCTAAACAT

CGCTACGAGTAGAGCGAATCGTTCATAATCCAGCCACGCTATTTATGTTCGAGCGGGCCAAAAACGCTAC

GTACGGGCACCCATCGAGTAAACAAATATCTAAAAAATAATGTTTGCCTATGCAAAATCCCAACGG

>NODE_650_length_276_cov_4.752941

GAAGAGGCTGGGACCGATCGAACCCTGCCTAAGTCGCTGGTCGCAGTAACTGGGACGTAAAAGCTCACTC

CGGGGGTCGAGGAACTAAGGGCTGTCGTTTCGCTCTCTCGGCCGCCGCGCCTCTATTGATTGTTGACTGT

TTCTGCGGAGAAATAAGCGGAAGCAGCAGTATGGCTCGGCAGGACTCTCCCACCGACTTCAGCTTTGTCC

ACGAGCGGACCGAACCCGTCCACGGGGAGCCTGTCCAAGGCGTGGACCTCGAGAACGACAGCGGTT

>NODE_651_length_276_cov_2.298039

ACCAGAACATAGATGGCTTAAGAAACCTCTTCGTAAAGAATTTAAAAAATGGTATAAAAAGACTATAGAA

TGGAAGCAGGAATTCCCATTACTACCAATCCCGGGAGATTATTCATTATTGCCGAAAACGGTTGATGATG

TTTTATTACCTCAAACTGTTATTAAAACATTAACAGATATTAGACCTTATGCAATAATTACTACAGATGT

TGGGCAACACCAAATGTGGGCTGCACAGTATACAAAATGTAAACGACGTAAATGGTTATCTAGTGC

>NODE_652_length_275_cov_22.468504

TCCTTGCTACTGACTGCATGAGTGACGACATCAAGGATAACAAAAACAAAAGAGCTAGTCAGTTGAAAAC

ACGCAATGCAATCCGCTGTTCTACAAACCAGGACCAACTAACGACGAGAAGGCCCGCGCATGCCATACGT

AAACATACATACATACACCCAAGTGTGTGTATATTTTAATGTATAGACCAACAAGCAGCCCATTCCGCCG

TCGCTACGGAAGTACTGTTAGTTGCGAAGATGGTGGCTGGTTTGGCTCTGCCCTAAAAACACAAA

>NODE_653_length_275_cov_9.696850

CCGACATCGTAACTGTTGTAACGGATCTATGGGCTCGTGCTGTCACGAACCGTGTCGAACGCGAAAGCAA

AACAGTCCCACGCGGGCCGTGGGGTTACCCGGGCAGTTTTTGCCCTACGGGTCTCTCTGCAGGCGTTCGC

GTACACTCCTCCCGAGAATAAACGAAGACAACATAGTACCCGCGGCGTCTCCTTTGCTGCTTCGCGCGTA

GCGCCTGTTAAGGGTGCCGTGCGTGAGGTGTGTATCGGATCCCACCGCCGTGTGTCTCCCATGAT

>NODE_654_length_275_cov_7.952756

GTGGTAATTACGGTGGTTTGGGTATTTGACGGGATGTGGGCTTGGAGGGAGGAGACAAATGATTGTTGAG

TTATTATAGGTAGAACGGCGAATCGATCGACCTAGGGAGTGAGTACCTTCATTATTTGGACTTCTTGACC

CCTTTAGTCACTTTGCGTTAGGTATATTTTTATGCCCGGAGAAAAACGGGAAGTGTGGCCGGAAGACAAA

AAATATAATTTTGGCGTGGTATTTTTGTCGCATGTATGATGCTTCACGCGAGAGTGGCATTTGGA

>NODE_655_length_275_cov_2.133858

TGCAGAACCCGGCAAATCATCGCGCATCACAGTGTAGATCCCTGTCCCCGTTTTCCGCCCCCTTCCCTCA

TGGTCATACGCTTACGGCCGTGCACGGCATGAGTACACCCCCCTTGAACCGAAACAAAAGCGCAACACGT

GCAAACTTCAACAGAAGAAACCACACGAGCTTCGAGAAAACACTAAGACACAAGGCAGAGAAACACCAAA

ACGGCGGACAGTTCGAGCTACTCTTTCAGCCATTTATCGACGGCGTTCCAGTTGCATCGCAGGAA

>NODE_656_length_274_cov_1494.478261

CATGCAAGTCGTACGAAAGTGTTAAAACTTTAGTGGCGGACGGGTGAGTAACACGTGAGAATTTACCTTT

AGGAGGGGAATAACAGTTGGAAACGATTGCTAATGCCGCATATCATTTATATGAAAGAAGAAATTCGCCT

AAAGAAAAGCTTGCGTCTGATTAGCTAGTTGGTAAGGGTAAAGGCCTACCAAGGCGACGATCAGTAGCTG

GTTTGAGAGGACGACCAGTCACACTGGAACTGAGACACGGTCCAGACTTGGAATTCTCGGTGCC

>NODE_657_length_274_cov_501.442688

CTGTCTCCAACATAGACTCAGCGAAATTGAATTACCGGTCAAGATGCCGGTTACCCGCGGTTAGACGGAA

AGACCCCGTGCACCTTTACTATAAGCAAATATTGTAGGTCAAAGACTCTAGTGTAGAATAGGTGGGAGCT

TATGATTCTTTCTTGTTAGAGATTGGTGAGGCAATAGTGAAATACCACCCTGAGGTCTGTTGGCTTACTA

ACTAAGGGACAGTGTTTGCTGGGTAGTTTGACTGGGGCGGTCGCCTCCTAAAGAGTAACGGAGG

>NODE_658_length_274_cov_27.284585

CATGCTTACAATACCCCGTTCATTAGTTAAGTCGTCTACAAAGGATTTTAAGACCAGCGATATTTGAAAT

TATGATTCGGTACGCTTGGCCAAGTAGGGATGAGTCGCCCTTGCTTGGCTGCAGCACCAGTTCTATGATT

TGGGACTCCGAAGAGCCTATCCTTATTATCGCCTAAATACGCTGGTTACTACTGTCGACATTCAGCATGG

ATTCTGGTTTAGAGGCGTTCAACCATTATCCTAAAGATGATAGCTTCTCGGCACTACCCGTTCG

>NODE_659_length_274_cov_10.134387

GCTCTTCAAGGGCGCAGGCGCCAACGCTCTCCGGACGGTCGGCTCCGCGCTCGTGTTGGTCATGTACGAC

CAGATGAAGACCATCATGAACCTCCACTAACCGGGACCCCGGGTCGAATTGGGTGGGATTGGTGATGGTG

CCGCTGCTGCTGCTGCCATTGATTGCGCTTGTCAAGAGACCGAAAGAACGAGACGCGTTGGATTCTGAGG

GTTCCTTACCATGATGGAAAGAATGAAACGCGTTGGATTTTTGGTGTTAATGGAAGGCTCTGGA

>NODE_660_length_274_cov_9.612648

GTTCCTTGGCACCCAGAATTCAATAAAATCAGTAGTCACCACCAATACCACATACCAGTACTACACCTCT

CAAGGTGCACATACGTACGTACCCTTATTTCTCCTAACCGTAAAAAATGTATGTTTCAGTTCCACCAGTC

TCCTTCTGGTCCTGTCTATACCAGAAAAACACTACAAGTTTTTCGGATTTACAAAGACTGTTTGATTTGA

ACATGAAACTACATTACAAAACTACAAGGCGAAAAACGGTTACGTACACCATCCAGAATGAATC

>NODE_661_length_274_cov_3.920949

TGATGATGATGATGATGACGACGACGATGATGATGATGATGCCGTCTGTGGCTTTCGACCTTTTAGCGGC

GTGGTCGCCGTGTTGTTTTTTTGCGGCCGTATTGGTGTTGTGTGCGTAGTACGTACGTACGGCACAGAGG

ACGGGACAAAATGATAGCGGTGACCCACGCCGGACTTGTGATACGAGTGAGTGGCGAACGGAAGTATTCG

TCTGTTTATAGCGGTCCCTACTAAAAAAAAAAGTTTCGCAGCGCGTGGTTTGGGTGTTTTCATG

>NODE_662_length_274_cov_2.245059

GCCCCGGACCCCAACACGGTGAAGTTATCCACGCGGGTCAGCTCCATGTAGTTTCCGGTCTCTTGTCTCG

ACTCCAAAAAGGAGATCGCACTGATGTTGCTGGAGTCCGTAAATCGCGCTGCGCCTGTTCTGCATTCCGC

CGGAGACATATCATCGAACATCAGGTCCTTGGAGTTCCTGACTTCTAAACAAAGGTTTCCAAGCGTGTTG

GGCACGATGTCCCCGGTCCAGTCCGTCTTTTGTAGCATATCGATGCTGTACACCGTTGCTCCCG

>NODE_663_length_274_cov_2.118577

GGATCGAGACCTACCATCGAGGAACATATTTTCTAGGTCAAATTTTGATCGTGAGCGAAATACAACTCTT

AAAACAATAACAAAGTACTAACACGATTTTCATTCTCATCGGAAAGGCAATTCTCAATTAGGAAAATCTG

CTGTTATATTATGTTCTTGTATACCCGGTACACTCGTACACATAGCGAGTAGTCTATAGTAGAACAACAT

ACATCGTACGAGTATTCGTAGCTACTTGTATAAGAAGAATCAATAAGTAATATCTTAGATATGA

>NODE_664_length_273_cov_6.805556

CCACAGGGGCGGGATGCGCGATGTTGATGCCCATCGAGTTTTCTTGTACTTGTTTGGAAGGATGGGGAGC

TGCGATCTGGTTGATTGGTTCGGGAGCTGGGATTTTGAGTGGTTGTGGTCTGGAGTAGGGATTCGATTGG

TTGGTCTGGATTTTTTTTGGTCGGTAAAGGTGCTCGTCGATGCGCCGCGTTTCTCGGTATTTGCGGGGGG

AGAAAAATGAGTGCCGATAAACCCGATCGTGTCTGTGTCTGGGGTCAAGGTCGCGGCGCCCTC

>NODE_665_length_273_cov_3.277778

GATGATAGAGGGTCGAAATAAATATAATAGGAATAAATAGGAAACAAATTGACGGCGAAGGTTCTTAATA

GGAGTACAAATGGTGAAGAAAATAGAAACGATGGGGATTGACTCCGAATCAATGCGGGGGGCTGGCTGCT

TCCTGAGAGCACGTTGTTTGTTGTTGTCGGTTGTCTCGCGGTCATCGCGAAAGGGGCTTGCAAAATCGCT

CGCGAGACTGGCTGTGCTGATGATCGAGATAATTTGACTCGTGCATGCATGCATGCATGCATG

>NODE_666_length_273_cov_2.690476

GAACAGAGTGCGTACTGTATCGTACCAGTACGTACTGACATATCATACCACCAGTAGATAATTGTCTCGC

GTCACAGCCCAAAGAACGACGAAAATACACCCCCTACGGCGCGCGACGAACCAGCGGGTCGGGCGTGGGG

AAACGTCCACGTTACCTCAAAAGACCTCGAACCCACTCCCAGCATCGCAACAGCCAAAAAAAGAAACACA

AAACAATTGGAATAAAATAAAACCAATCAACTCCATACACACCAAAATAAAGTATGAATACTA

>NODE_667_length_273_cov_2.527778

ATTCCATCTCAAAGTATTAGCTCTGGCACTATTGTTATTTAAAGAAAAGTTCAACATATTCTTCTCCTTA

TTGAACAGTAATAGTTAATTTGGCTACAGGGTTTAACCAACGGTGAATACTATTGTTTAAATCACTTTTA

CCGCCATCAACCATCTCATCACCTAAGTTTCCGCGATGAATATGTATAGTGGTATTGCTTTCTGTATCAG

TAACACCTGTTCCATTTGTACCTGCATCGCCACCTGGTGCCGCGGGAATACCTAACACACCTG

>NODE_668_length_272_cov_45.382470

AGGCCACCGTACGCCTCGGGCCTCAGCACGCGGTGCACTTGCCACTTCGCATACCACGCGTGGCGAGTAG

AAGAGCTGGCCGAGGCCATTTGGGTGAGGATGTGCGGCTCGCCGAAGGTAGCGAACCCCTGCTGTCGCGC

GGCACCCGTGTAAGGCCCGACGGAGCCCCCAAGCACCCCTTGCGTGAACAAGATCAGCGCGGTGCGGAAG

GCCTCCGTGTAGAGCTGATCGGTGAAGGAGAGGTTGGCCAAGTCTCGGCCGTTTCGGATGAA

>NODE_669_length_272_cov_7.876494

GAATAGCTTGTTTGCACCGAACCTTCACAATTTTCGACCTTACAATATCGGAGGTCATGCCACCCTCGGT

CTCACGTATGATTCCCAAAATGGCCTGCGTTGATTGAGAGTGGATACTCCGTGTGAAGGCGGAGGTTGAG

TTTGTACATCGATCTACCACGCGCTGTATGAAAGTCCTCTTTCTGCTTGGGCATCTTCCGGTTGAATAGA

GTTCTAGTTCGTATGACACATACAAGCGTAGTTGAGTCCCGTCGTACCAGGGTGGTGCCGCG

>NODE_670_length_272_cov_5.270916

TTTGCGGGGAAATCCCCGCGATTTTCGCACTTAAAACTGATACAAGCACGGCATGGTTCCTGTTTTCGAA

CGGGTGAAGTACGCCCTAGCCACACAGATCGGAGGGGGGAAGGTACATCGTTTTTGATCATCACATCGTC

CGTGAACGCCGGGGGTGATTGATTCGGCACCTACCACGTATACTTATTTTTCAGCTCTGCTGCAAAACTA

ACTGTAGATAGATGCCTATTTTTTCTCCCTCGTTTGTTTTGCACTTTTTTTCCCGCTTCCAA

>NODE_671_length_272_cov_4.219124

GAAACACCGGCCTCGACGACGCGCAGCTGAAGGTCTCCAACGGAGCCTGCTTCGACAAGAAGTCGGACAG

CGGCTTCAAACCTGTATGCTAATAAACGGACCTCATCTCTAATGAGCTCGCGATCTCGCGGGCGGAGGGA

GGCGTGGTACACACCAAAGTATTATATACTTCACCTCCCTTGTCCTCCTTGTCTCCGGGAATCTACGTAC

TGGAGATGATTGTGGTCGGGTATGATCTGTCGCGTAACTCCTTACGCATCTTTATGCGGAGG

>NODE_672_length_272_cov_4.099602

AACGTTAAAATTAAACGCTGTAGTACATCGCAAATTCTACTGGGTGAGTTGAAGTATTTAATAATTCACA

ATCAGCACGTTTAAGATCAATGTAAGCATCGATCATATCTTTATCCATAACACCGCCAGCAGTTAAGAAG

TCGTTATCTGCAGCTAAAGCATCTAATGCTTCAGTTAATGATGTTGCAACTTGTGGAATTTCTGCTGCTT

CTTCAGCCGGAAGGTCATAAAGATCTTTATCCATAGCATCGCCAGGGTGAATCTTGTTTTTG

>NODE_673_length_272_cov_3.884462

TCGGGGGCTGTATAGTTGCTATGCTGAAAAGCTGTCGTGCGGTATCGACGCGTCGTCTGTCCGTCGCTGC

GTTTGATTTGTTTTGGTGTTCCGCGATCGACTCGTTGCTCGTAGCCCGTTTGAAAACCTCGGCGCCGAGA

TCCGTTTTTCCTCGCTTGAAATGATACCCGACGAAAACTACCGGGCGGCCTTTGTGATAGCTGTGCCTGA

GGGGGCATGATCAACTTCTGTGATACGATGGCGCGTTCGTCGCGGGGCCGAGAGGGCTGCGC

>NODE_674_length_272_cov_3.665339

TTGGCACCCGAAATTCCAAACGGAGCATGCACGTACGGATGCTCCGGCCCTTCGCCACGGTAGCCTCGGT

GTCGTAGCTCAGGCGAGCGTCCACCTCGGTGGACACGTAGCCCGGCACAATCTTGAGGATCTCGAGCCCG

AAGTTGACCGAGAGCTTGTCCATGGCAACCTCCACGCGCTCCTTCTCCGAGCAGTCTGGGAGGCCGACGG

CGTGCTTCACCGCCTCGTCCACGACACCCTGGTACTCCTCCATCTGCGCGGCCGCGTAGATA

>NODE_675_length_272_cov_3.294821

AGTGGGTAAAAAAGCGAATTGCTGCGCCCTTGTGTAACTTGTCTTGAGGGCGGCGGGCAGACCGGGGACC

GTAATTTTTGTACTGGTAGAGTTTTTTTTTGTTTCTTGTGTCTTTTTGTTTGGTTTGTTGGCCTCGGCTC

TATCTATATCCGGGCAAAGCTTTTTACTTCTTGACTCTCCCTGTCTCTACGTGATGCTGATATCGTGGAG

CCTGCTTTTTTTTCCCCGCTATATAGGCGATGCAGGGTGTAATGCCGTTCGAGTTGCTGACT

>NODE_676_length_272_cov_3.243028

ATCACCGAGCTAATGGCGGACTTGAACTTGCGACGCGCGTTGAAGAGGCGGAGCTTGTCCATGTTCTTGC

AGAGGTCGTGCTTCTCGAGCTCGGCGGCATCTCCTACTATCCAGGGGTGCTTCAGTGCCTCCTCGGCTGT

CACTCGCTTGTCCTTGTCCATGGTTAGCAATCGCTTGATCAGGTCCTTCGCCTCGTCCGAAACAGAGGTC

CAGTACTGGGGGTGGAACGATACTTTCCCCTTCTTGATCTTCTGGTACAGCTTGGCCTGATT

>NODE_677_length_272_cov_2.956175

GAGGGCATGTCGTCGAAGATTCCCGTGGGCAAGGTGGCGATGTCATTGTTCTCCATTCTCAAAAATCTCA

GAGAAGGTGTACCATCGAAGACACCCGTCGGCAGGCTGGATAGAGCGTTTTCACTGAGCATCAGGTTGTC

TAGGTCCTCCAGTCCGTCGAAAAGGTTGGCAGGGAGGGTGCTAAGCTGGTTTATGTTTAGGCTCAGCACT

TGCAAAGCCGGCGTGCTCCTGAATATTTCCTCCGGAAGAGTTTCGATGTCGTTGCGGATAAA

>NODE_678_length_272_cov_2.808765

AATATCATAAAAGCTTCTTACTTTAATTATATAAACCAAAATAAATACAAATCGACGCACCGACCGCGAC

GGCTACCGACCCTCCTGGGTTCACTCTCCACAAGCGGCTTCCTCCCAACATCACCGGAAGACAAAACGCC

TCCACGGTCACCGCCTACACACTCTCGCCGCCGCGCACGCAAAAAACGCCAGCGCATATTTTTCGCGCTT

CACGCCAGACCGGCCTTAACCCCTTCAGAACTGGTCCCACATATTTGGGGACAAACTACTTG

>NODE_679_length_272_cov_2.800797

CACAGAAGACGGGGCGTAGGAGCTTGGGGCTTCGCAGGACACCAGCCCGAACGTGGTGCCGTACAGGAAT

AGATCGTTAATTTCGGCGACCAAGGTCGTCTTTGGATAACAACCGAGGCACGTAACAAACCGATTGGAGA

GAGGGGTGCTTGTCAATCCATCGTGTACCAAGACACCCTAAGCTCTCCCGCAGCTGCTTCGTTTTCTCCA

ACTCACTGGAGAGCATGCGACTATTTTTGGTCCAGCTACTGCTTCTGGCGGTGCCTCTCTTC

>NODE_680_length_271_cov_686.460000

GATAGCTGGTTCTCCTCGAAATCTATTTAGGTAGAGCCTCGCACGAACACCATCGGGGGTAGAGCACTGG

ATGGGTAATGGGGCCTCACCGGCTTACTGATCCTAACCAAACTCCGAATACCGAGGAGTACTAGTCGGCA

GACACACGGCGAATGCTAACGTCCGTCGTGAAGAGGGAAACAACCCTGACCTACAGCTAAGGCCCCTAAT

TCATGGCTAAGTGGGAAAGCAGGTGGGACGACCAAAACAACCAGGAGGTTGGCTTAGAAGC

>NODE_681_length_271_cov_13.740000

CAACAAACCCTGTTCCTTTACCTATAAGTCTTTCCACTTCTATTCTTACTTTTCTAATAGTAATTCCATA

TTCTCTGACAGCATTAGTAGTTACACCACAGCCTTCTCCTAATAAGCCTAAAAGTATTTGTTCCGTACCT

ACAAAATTATGTCCCAACCGCCTTGATTCTTCTTGTGACATCATAATTACTTGTAAAGCCCGCTCAGTAA

ACCTCTCAAACATAAATATACCTCCTAAATTGGTATTAATTAATAAAACTATAGGCTGCTT

>NODE_682_length_271_cov_4.592000

GCGACAGACCCATCTCGCTGTTCTGGGCGCAGAAGCGCATGTTCCCCGCGTCCGAGTTCGAGATCATCAT

CCACACGCCAGAGTTGCAGAAGTTGCACTCGCCGTCGAAGAGCACCACCGCGCTGTCTGCGCACTGGAAT

ACCGCGTGCTCCACCCCACCTCTCACGTGCGCATCGCCCCAAAACCAGCTGTCGAAGTCACTCCCTGACA

CGTTCGGGGTCCTCTCCGACTGCGGCGGCACCGTCGAGCTCATCCCCACCACTTGTCGGCA

>NODE_683_length_271_cov_3.936000

CCAAGGGGCAGGGTCTCGGGCGTCAGCTCGTTGTCGAAGAGCCGCAGTGTCTGCAGGCTTTCAAGGTCGG

AAAAGGCTCCCTCGGGCAAGCTGACGAGGTTGTTGAAGGCGATGTCCAAGAACTCGAGCTTGGTGAGACC

ATCGAACAATCCCTCGGACACGGTGACCAGAGCGTTGGAGAACAGGTCCAAGAAAGTCAGCTCGGACAGG

CCAGCAAATGTGTCATCCGGGTATGATGTAACTTCGAAGCCATTGTCGAGGAGTAGTCTCT

>NODE_684_length_271_cov_2.876000

AGCCGAATGCGCCAACGGAACCACCAACAGTTACGAGCCAGAATTCGTCGTCGTTGTCGTCACCGTAAAG

TGCGCCAACTGTGTAGTCGCCGAATGTGTAAGTCGCACCGATCTGCCATTCGTCACCGCCACCTGCGCGG

TTACCCAGCGCGTCTGTGCGGTTGTCGTCACGGCCGCTTTCGTCAACGTAGGAACCTGCAACTGTGAAGT

CGCCGATTGCGTAGTTAACCAGAACACCGTTGTCGCCTGAGCCTGCGCCGAATGCTGCGCC

>NODE_685_length_271_cov_2.288000

CACCCCGAGAATTCCAGAATACCTAAACCTCCTCGCATAGTGCCTACTCGTACGTACCCGTGGGACTGCT

CGTGGGTGATACCACGAGACGATCCCTGTAACTTACCTCCGTGACACATCGTATGTAACCGTAACTTATT

GCTAGGCTGTTCGTTGAGACAAGCTATATCGAACACCCTGCCTGGCGCTGGATCAACTAACAAGCCTCGA

TAACACAAACAAACACCATCCACAAACAACAAAAAAAAAATCGCCACCGCACCCCCTCAAC

>NODE_686_length_271_cov_1.384000

TGTTGAAGGTCCGAACTTCGAGTTCTCAACCGAGACCCGCGAAGAACTGTATTACGACAAGTCCAAATTG

TTAGAAAATGGCGACCGTTGGGAAGCCGAGATTGCCCGTAACCTAGAAATGGACGCGCCCTACCGATGAC

ACTCAAGGCCACCCAGTCAGGCAGCCAGAACCAAGACAGCGTTTTTACAGCGCTGTGGGGTGTGCGGTAT

GACTGATAAAAACCCATGGGAAGACATGATGCGTCAAGCGCAGGAAATCGCTAAGGCTTTT

>NODE_687_length_270_cov_32.642570

GGAGCAATGTTGTGGGAGCGGACAGAACACGGCGCGCGGTGTTGAGCTTTCTCGTTTCAACGGAGTGGAG

GCGTTTTTTGTGCCGTTTTCGTGCGCAATAATCCCGCCGCTTTTTTTGGTAGGGTTTGGTTTGAAACCGT

TTTTGCCGGTGTGTGCGATGGTTGGAAGGAGTGTTGCGGCAGGCGCGCTTCATGCTTTCGGGCAATGAAG

ACAAATGACAGGGCAACTAAATAAAAAAACCGGGAAACATAAAAAAAAAAAAAGTGGAAT

>NODE_688_length_270_cov_24.642570

TTCCTAAATATCGGAGACCACAGCCGGCGCCCCAGGGAAAAATTGCAGAGTTTCCGGGGCGTGGTGGAAA

AAAAAGCCGGGAAGATTTTTACTGTCCCGTGAATTTCAGCGGCTCCAGCCCTACGCGACGTTACCGAGCA

GGCTAATGAAAGCCAGATCAAAACGTGGAGGTATTATTCTTCGAGGACTTGGATGCGTCAAAGCCATGGG

GTAGGGCTAGACGCGGAAGACGAGTATGTTGTCCTCGTCTTCAAGTCCTCGATCTATTAG

>NODE_689_length_270_cov_12.044177

CGGCTGCCATCGTGCTGCGCGAGGCGTTTCGCCACCTTGTGGAGACGAGCGCCTGATTTATGACCTACCG

AGAAGGCCGCCTGTCAAAATGTGATACAACATATAACACGCGGGGAAAGGAAACTGTTAAGATGGTAACG

GTACCCGGGAAATAAAATTGTCAAGAGAATACGCAGCAAAGGGGACTGTCAAGATCGTGCGCGCGAAAGG

AAGCTGTCAAGATTACACGCGGTAAATTGAGCTGAGGGCCGGTGGAATTCTCGGGGCCAA

>NODE_690_length_270_cov_3.361446

GGGTCGCTATAGGGTCTAAACGAGGGGAAGCGCAGCTGCAGCTAGTAGCTTCGTAGTAGCGCGAGGAAGC

GGGAAGCACGTACTGTAGTTCGACTATGGTAAGTTGAACAGTTGACCAGCGAGCCAACGTGGTTGCAGTT

GAAGAGATTGGTTCGTTCAGTCGCGGTTTGAACCGCGGCGTATCAGGGTACAACAGGATACGCAGTAGCA

GTGGGTAGGCAGCCTCGGGGATCTTGCCCACTAGAGGCAGCTTCACGCAAAACAAGGACC

>NODE_691_length_270_cov_2.726908

ACGACAACCCCGGGTGCTCGACTCTTCCCGAGAAGGTTAACCACGCGGTGCTGGCGGTCGGTTACGGCAC

CACGGCCGAGGGAGTCGACTACTGGATCATCAAGAACTCTTGGGGGCCCGACTGGGGTTTGGACGGGTTC

TTCCACATGGTTCGGGGGGTGAACATGTGCGGGGTCGCGGACTGCGCGTCGTTCCCTTTCGTACCCTGAC

GAAACGAAAACTGGAACTGCTTTTGGCGCTTCATCCGCACCCGTACAGTAGCAAAGTAAG

>NODE_692_length_270_cov_2.060241

AATATAGAAAGGCCTATACGTACGCCTGCCGTACTACTGTAGAGTGGAAGCAATCGGCTGGAGTGCTCTG

GCTCAGGCTGTTCATTTTTCGGAAAGGGTGCGTCGCACCCTGATTTCGGTAACATATTTTCTGAATTTGT

GATATTTCTGTAAATCCCAGCCTGCCTGCCCTCGGTGTTGGTGCTCTTAAGATGATCCAATATCTGAACT

GCTGTACTACCCCACGCGACCGTTCCCAAACGGTATGCATGTGGTGACGTAGTGGTTGTA

>NODE_693_length_269_cov_3.995968

ACGTCTCCACCGTCAACAGCGATACCACCACCGTTACCGCGAGTGAAGTTGTTGTTGAACTGCACGTCGC

TCCGGAAGGTTATTGATCCAGCGTCGTTCAGAAGAGCGCCCCCGTTGTTCTGAGAGTCGTTATCTTCGAA

GAGCGCCTTCCCGTAGAAGTTCATGATTCCGACGTTGTACACCGGCGAGCCTTCGGAGCCCCTGCATCTG

GTGTTGATGAACGTCGCGTCTCCTCGAAAGCTCATTTTGGAGGTGGCATCATTCTGGAT

>NODE_694_length_269_cov_2.754032

CCTCGACTTCGACAGCTGCGCCGTCTGTGAGGAGGCCTACGACGCAGTGTGCGAAGGCGCGCTTTCCGTG

TGTGACTTGGAAGACTTCGGCAACCCGTTCAGCGCCGAGGCTTCTGCCTCCGTCGATATCGTGTGCGGGA

CCTTCGGAAGCGCATGCTCGAGGTTTGAAGCTAGCGTGGCTTGCCGGGATCAGTGCACGCCCGAAGAGGT

CGGCGGTGGCGGCGGTGGTGGTGGCGGCGGCGGTGGTGGGTGTGTTTGCAGCAATCGTT

>NODE_695_length_269_cov_1.919355

AACCATCTTCTACAGCGTCTAGAGCTACATCACAAACATCGCCAACTTCAACTTCAATTTCGCCAGCAGC

GTTCTTGAATTCTTCAACTGGGATAGCAGCTTCAGATTTAAGACCAGCGTCAACAATTACGTTGTCTTTG

GTTACTTTAACGATTGTGCCACGGATAATTGAACCCGGGCGAGTTTCGATTTCTTTTAAACTTTCTTCAA

AAAGTTGTGCAAAATTTTCAGTCATAGTTAAGTTATTTGTTTCATTTTGTTTGAATCGT

>NODE_696_length_268_cov_43.072874

CACGTCGATGGAGTCGTACTCGAAGTCGGACAGCAGGAACTGAGACACGACGGGACCGGTGGTCACACCC

GGCCAGTCCGTGCGGAAAAGGTTAGTGACGGGGTCGACCTTGGTCTCGTCACCGTCGCCATTTCCAACTT

TGGTCCTGGGGATGTTCAGCCCTTGGAATGCCTCGTTGTTTCCAAGGTTCTCTGCGGCGAGCCTGGTGAG

GTCGTCCGTGTCGAATTGCGAGAAGGGAATGTCCCTAGCCAGGGCCATCCAGTACACC

>NODE_697_length_268_cov_28.457490

AAACCACGTTGGGTTTTGCGGTTGAAGGTAGGGGCCTTCTGCCGCTGCTTGGACCAGAGCACGCTGTCTA

GCTTTGCAAGGTGTCTTCTGAACTTCCCAAACAGAGGACCAATACGCCAAGTTTCTCGTCCGCTCCTCTT

GAGTGCAGTGCTGTTGTTGTGCGCCGGCTTAGTACCTCTGGGACGTCAGGTATGGTATACGTCTCTTTCT

TCTCTGTCTCTGTGTGTTGTCTTTCGCCTTGTGGTCGTGTTATCTGTCGAGTGTCTGT

>NODE_698_length_268_cov_20.340081

AACCTGCTAGAAACAAGAGGCTGCGATAGATGGGAGTGCTCTTTTGCACGTCCGATCGTTGATCGCGACT

CTGAAACCGCAGGGAAATTATAAACCGTACACTCCCCTGCCTGCCTCCTGGCGCGTCTCGTCCTATGTCA

AAGCGCGCAACCATTTTATTTTGGTGGTGCAATTTGATTGGACGAGAGGCGTTGGGTAGGTAGAATGGTT

GGCGTGCGGCAACCATTGCTTACCGTGCGGAACTGCCTCTGGAATTTCGGGTGCCAAG

>NODE_699_length_268_cov_5.927126

TCGCTGTCGGGGTGTTTGCTTACCCACGCGATGCCCAGCCTCTCCTCTCCCTATGCTGCACGCATTGCGC

TCTCCGCTGTGGCGATGTTCGCGTTTAACAGCTTGGAGGACTCCGTCGCGATTGGACCCGTGCACCGTGC

GCTCTTCTACGCGTCCCTGTCTGTCGCTCTGGTTCTAGCGCCCTGCGTGTGGCCCTTGAACTTCTTCCAG

GAGGTGCAGTTCGCCATGACCGCCTGGGCCTTCGCCTGCCCCTGTGGCCCGCAAGTAT

>NODE_700_length_268_cov_5.566802

TCGAGCAGAGGAAGTGGTAGGGGCTGGCAGATTGTTCACACATTTCTTCGACTTTTCGGGACTGACTGCT

GTCTGTGGTACGGTGGGTATGTCCATGCGAGAAAAAGCCTGCGTCGCGGTATAGGGATGGATGGCGTACG

TACGGTTTTTGGTGTGATCGTTTCATCTGACTTGGCGCTTCAGAAGAAGGCTGTAGGAGGAAACGGGACA

GGGCACAGCGAGCGCGGCGCTTTGCTTGGCTGGACGGGCATGAAGGCTACGCTACGCT

>NODE_701_length_268_cov_3.534413

GTGATTGATTTCGTGTCGTCTGGGATCCTTGAAAAGGTGAAACAGCTTTGCTCTTCCTCGTCCTCGTCTC

CCGACGGTGTCAGCTGCGACCCCAACGAGACTGACCCCGCGTTCGGGACCACTCCGCTGCACCTGGCCGA

GCTTTGGGGCAGCCGCGATCTGGTGGAGTATCTCTTGTCTATCGGAGCCAACCCCGAGCTTTACGATTCG

GCGGGTCGTCAGCCTCGAGACATGGCCTACAGCGACTTCACCGCCAGCAGCAAGGATT

>NODE_702_length_268_cov_2.850202

TATAATTTGGAATAGGAATCTTGACGCTCTTTAAAAGATTTAGTATCCTTAGTATATGTAAATATACTCA

AATCTTTTAGCTTATTAAATACGTTTCAAACAAAATAGAATAAAATGTCTCATTACACATCTATTAAAAC

GAAATATACAAACCCCAATGTATTAAAGAAAGTTATAAATAAACTTGGATACCCTTATGTAGAACATAAT

AATAATAATATTATTGAGGTTCCTGTGATTTTTTCAAAGAGTTTAAAGTCTATGGAAT

>NODE_703_length_268_cov_2.287449

GCAACGACGACTATGCCCACGAGCTGTACACGGCGATGACGGACATGTGGGATGGGACCGGGATGTTCGT

GTTCACTTCTTCGGGAAGTGTCTTCGCGGAGAAGGACGGAGGGATGGTCAACGAGGAGTCGCCCTCATCC

GAAGCTTCGAAAACGTCACCTCTAGGAATCGCGGAGAGAGTTTGCCTGGACGGCGGGGGCACCGTAGTTC

GATTGGCGGGCCTCTACTCCGTCGGGAGAGGACCCCACGGCGTCTGGCTCCAAGACAA

>NODE_704_length_268_cov_1.805668

TCGGTGGATTGAAACGGTGCTTGGGATGCCGAAGCGCAACTTGGCGAAACGGATCTGAAAAAAAAAAAAT

GAAAATCCCAACGGAACCTTGAAAAAAACGCGCAACGGAACTTAGAGAAGTGAAACCGAATTTTTGCGAA

GCGAAACGAAACAAAACGTAACGTAAGAGGCTCCTAAGACCTACTGCTACTTTTTTTCTGAAGGGTACAC

GGAATATATACCGCCCCCGCGGTTGTCTGTTTTTCGCGTGGTCATTTGGGGTTGGGGA

>NODE_705_length_267_cov_1615.772358

CAAATGAGAACTTTGAAGACTGAAGTGGAGAAAGGTTTCATGGTAACTGCAGTAGATCATGAGTGAATCG

ATCCTAAGGTATAGATTAACCTCGTGAACGGCGACAATCGGGCCGGCATTAGCAGGCTCACCTCAGGTTG

TCCCCAATACCGAAAGGGAAGCTGGTTAATATTCCAGCATCAAGAGCCGGATTACAGACGGTAACGTAAG

TGAACTCGGAGACGCTAGCCGATGCCCTGGGAAGAGTTCTCTTTTCTATTTAACAGC

>NODE_706_length_267_cov_3.605691

TCCCCCCCCGCTACGATCTCAACATCCACCTCTGCCTCGACGTCCACTTCGTCGCACTCACTGTCCGCGT

CCTCCCCCTTATTTTCCGACTCGCATTCCGAATCACTGACGTCTACATCACCCTTCCACTTGAGTTTGCC

GTTTTTGTCCTTCTTCAGCTTCAGTTTGCAGCGGTCGCCCATTCCGACCAGTTTAAAGCTCGATTTCACG

ATGCCGTTGAACAGCTTGATGGTCGTGCCGTCCCTATCCCCCTTTCCGTCGCTGGCG

>NODE_707_length_267_cov_2.922764

TGCTCGAGTTGCTTCGGCTAGGCAGCTGCGAGTTTGTTTATGGATACTGTTCAACACATGCATGCCTGGT

TGTTGCAAAACCGGGGGCTACCGGAAATGCTGTAAGAAGCGGTGCACGTGTGCCGTGTCAACCCGATGAA

CAATGTTTTATAGCCAAAAGTGGCTTTTTGAGACAATAAGGATGTGCGTTTCGAAAACGAAGATATCCCT

TTGGGACAGAGCGAGTTGCCAGGTCGTATCACGCGTACACCGCGAATGGAAACCCCG

>NODE_708_length_267_cov_2.430894

ACGTCTATGTACTTATGTTAAACCACCAGAAACACCAGAAGGTCTTACTGGTGAACAACTAGTTAAAACT

CTTGATTCTACATTATTAGCTGTTAGCCCTCTTTGGGACAACGATGGTGAAGGCGCAGATCCAATTATCT

CTAACTTAAACGGTCAAGAAAGTACTGATGCTGGTCGTGATACTGGTGAACGTCTTGGTACTCGTTTTGG

TAAAACAGTAATTAATAATACTGCTGTTGTTGTTGTTGCTGCTGCTGCTGCTGCTGC

>NODE_709_length_267_cov_2.121951

CGGAGCTGGACCCCATCTTTCACGGATTTGAGGCTAAGGACATGGACCGCAAGCTGAACCTGATGAGTAA

CGCCTCGGGCGGGAACACGGGTTACCTGGAGGAGTTGGGCAGGCAACCTACGGTCACCCTCCGCCAGGTG

CACCAGAGGCTGGAGAGGACGTACTGCGGCACAATGGGGGTGGAGTACATGCACATGAACAGCAGGGAAA

AGTGCAACTGGATCCGAAGGAAGGTGGAAAACCCTGCGTGGCTCAAGTACACCAAGG

>NODE_710_length_266_cov_4.812245

GCTCGCCAAGTGCGCCGTGCACCACTACACGGGCTCCAACACTGACCTCGGCACCGCCTGCGGAAAGTAC

TTCCGCGTGTCGGTCCTGTCCATCACTGATGCGGGAGACTCCGACATCCTTCGCCAGGGCGCCGAGGCTT

AAGAACCGCCGATCGCAAGAGAGGCTATGGTCGCGCTTTGCAGGGGGGATGGAGTTTGAAGAAAAAAAAC

GGGAAATAGGGAGATGTTTTCATTGATGCATGCGCTCTCTTTCGTCGTCTTTTGAG

>NODE_711_length_266_cov_4.179592

TGACGTTACACCGATCCCACATAATGGTTGTCGTCCTCCTAAGAAACGTCGCGTTTAATCGTTTATTTAG

GATAGTTGGAGAAAGAAAATGGCTAGATATTTAGGTCCTAAGTTAAAGCTTAGTCGTCGCGAAGGAACAG

ATTTATTCCTTAAAAGCGGTGTAAGAGCAATTGACACTAAATGTAAAATCGAAACAATACCAGGTCAACA

CGGCGCTCGTCGCGGTCGTTTGTCAGACTACGGTGTTCAACTTCGTGAAAAACAAA

>NODE_712_length_266_cov_2.971429

CCTGATCCTTCGACCTGCTGGCCCTTGGGAAGAAGCGCGATGTCGGCATCAGCAATGTCCAGCACATACG

CCGAGTCCACCTTCTCGAGGATGCCGAGCCCGTTGAAGGGCCACTCGGTGACGATGCTGGAGCCGTACTT

GGAGTTGAGAGAGCTCTCGAAGGACGCGAACGCCGCGATGTCGCTGCCGCTCAGACGCTCCTCGGGGCCT

TCGTTCGTCACCCCGTCGAAAGAGAAGACTCCGGTCGAGAGGAAGAGGTCGTCCAC

>NODE_713_length_265_cov_48.426230

CCCACAACGATGAAGAACGCAAAAAAAAAGAGCTATTATCCATCGTCAATCAATCAGTCGTCCTGACAGG

GGGAGGAGAGAGTCCTTGCGGGTGGGGAGAGAGACCCCCCTCCGCCCACGACAGCACCCAGAAAATTCGC

GTAAGGCAAACCGGTTTTTGAAAACCAATATAATACGTGAGAGCCCTCCCTTCCACTGCCGAGCTCTGGG

AGCGGTCGCTCTTTAGGTCTACCCTTGATTTACAACACACATGTGGCGCCTCCCG

>NODE_714_length_265_cov_9.028689

TCCTCTGTGCTCGCACAGCGCCAGCGATTTCAGCACAAGGCTAGACCACCCGGCCCTCCACTCCATTCCT

ACCGACTACGACTCGTCCATTACTTGTCGGATTTTGGATGCCCCCAAGAAAAAAACACCCAAATCATCGC

CGTCAATGACAGCAGTAGTAGTCGAAAACAAACGGAATGACGTTACTGAGTAGCAGCACTCAGCAAACCA

ACAGACCAACTAGACCAACCAACAGACTGACCGACCGACCGACCGATATAAGTAC

>NODE_715_length_265_cov_5.299180

TGATCTACTGGGGAGAGGAGGCGTACAAGAGCCTAGACGCTTTGATGGCTGAGATGAAGAACCCCGAGAC

AGTGAAGGCCATGTTCAACAAGTTCGATGACGACGGCAACAACGTTTTGGACGCGAAAGAGCTGGGAAAC

CTCCTGGCGTCTTTGGGGGCGAAGCTGAGGCCTACGGAGGTTGAGGCAGCTGTGATGATCATGGACAGAA

ACCAGGACGGAAAGATCAGCCTCGACGAGTTCCAGGCTTGGTGGTCCCGTTACTG

>NODE_716_length_265_cov_3.655738

CCAAGGGTGTCGTGGTAGTGCTTGATGACGACGTGTCGTTTGGTGCTCATTTTCTGTGTGAGCATTTCGT

TCTCCTTCGTGAAGCCTTCTTCCAAGAGCTCGAAGCTTCGCGGGATCTCATATTTCTTGAAGCCTTCGAG

CGACAGGGCTATTTCTCCGGAGATCAAGTTCCTAACGCTTTCCAGGGCTGCGAGGTCCTCGTGTGACATG

GCAGGAGTGGCTCCACCCTTCGACTCCGCCCAGGAGCGCACCAGACCCCAGTCGA

>NODE_717_length_265_cov_1.581967

GTTTAGGTTATTTGATATTTATTTTTGAGTGTATTTTTTTGGATCTCGTGAATTTCTTTATTTTTGTGAT

GTTTTTTTGTGTTTTCTTCGTCCAGAGGCTTGCTGTGTCCTCGCTTGTCGACATCGTCGTGCCTCACGAT

GAAGCGGCTGCGGCCAGGCTGGCCTCGAAATTCTCTTTTCCCGAGCATACCTCGTCGTACGTCTTCCCGC

ACCACAGCTTGTGAACACCCCAGTCCTCTCTCTGGGGCACTGAGAGCGACAAAGA

>NODE_718_length_264_cov_46.181070

CCAGTGAGCTATTACGCACTCAAATGTCAGAGAAAGGTCTGGCCGAAAAATGCGTGCCTTTTCTCCCCTG

AACCTAATCAGAGGAGGCAGCAAGTACGACGGCACAAAATAGACCCAACCTCAGCGCAACCACACCATCT

CCCCCTCTCCCGGAGGAGGGAGGGAAAAGGCGCTGTGCGTTAATGTTTGCTCCATACCCGAGGATTCCTT

CCGTCCCCAGACGCAGTGCGCCTGGAGTCCAGCGCCCTTGAAACGAGGAGCATT

>NODE_719_length_264_cov_20.123457

TCAACATCCCACCAGCCAGGTCCTTCTCTCTTCTTCGCCTTTCCAGCACCTCCTAGCAGGAAGTAGCTCC

TAACACCTCGAGGGAGATCTGGGCATCTACTACTACTGCGTTCGGCTGCAATCAAGATCGCACGCACAGA

CGATATATCATCATGAAGACTTTCGTGGGCTTGACGGCGCTGCTAGCTGCATTTCCTCCCTGCCTCGGGT

ACGATGAGCCCCCTCAACCAACACACCCCCTAGTCAGCGGGAGCGTGTGCAGGG

>NODE_720_length_264_cov_12.637860

CGCGGACGCCGCAGGTCCAACGGATTCAAGATCTAAGGTCGTCCGGCAAGGTCTATGCGAGCAGAAGGCG

CATCAGATGCGTGATCTACCGTACATGCTGCTATTGGCGTGGGCGGGCGCTGATTGCTTGGGCTGGGGAG

AGAAAATTCGTGTGTTGGCTTGTCGGCGGTGGTTAGAGTGGGGTGCTTCGAGAGGGGCGTGTGGATGCTC

TTAGGCTGGTCGAAATGAAAACACAAGGATATGACAAAAAAAAAATTTTGGAAT

>NODE_721_length_264_cov_10.666667

CCCTTCGTGGGACGCATCTTGGACTTCTACAAGAAGAAGACGGGTAAGGCGTACACGGCCGCCGAGGACC

CGGGGTGCGTTAGCGTGTCGGAGATCTACGGGTACTACAAGAAGCACGGGCACGACACGATCGTCATGGG

GGCCAGCTTCCGCAGCAAGGAGGAGGTGCTGGAGCTGGCGGGGTGTGACCGGCTGACCATCGCGCCCAAG

TATCTCGAGGAGCTGCAAAGCAGCACCGATGACGTGACGCAGCGTCTGGAACAC

>NODE_722_length_264_cov_9.041152

ATTCCAATTTTTTTTTTCTAGAGAAACGTTGAACGTTTGTGTTTATATATATGATATGCGGGTCATTCAG

AGTAGGTTCCGGCAATGCCCAGCGTAATGCGCCACACGAACGAGCATCTGGCTACAAAAAAAAAAGGCAA

AAAACATCAGCACCAGTAGAAAAACAAAAAAAAACAGACCGGCTCCAAGGAAATCGACCGTTCCGACTAC

GTACGTAGGGTTTTTCCCGCTCCGTACCACACACGCCTTCACGCACATTGCAAG

>NODE_723_length_264_cov_6.164609

TCAAGATGTTCTTCTTCTAAAAAAGCTTGCTCTGATTGCTCTACATTACCTGCTGTTAATTGACCTGTAC

ATGATGAACAAGCTCCCGCACGGCAAGAATATGGAAGATTTAAGTCTTGCTCTTCAGCTGCTTCTAAAAT

AGTTTGGTCGTCAGGACAATCAATAACTTTATCAATACCTAAGTCTACGTTTATAATGTGTATATTAAAT

CCCATGTAATTAAATTTGGTTTAAAGTGCTTCTTCGTCGAATTCGCCAGTACGA

>NODE_724_length_264_cov_4.460905

GCCTGACGGATCATTCTTGGGCAAATTGTGTACCTGGTAGGTTGCATATTACCAAATGTTTTGCTTTGTG

CAACGATATGCAGAACCGAGGCCCCAACGTGCTACTATTATAGGTAAACCGGGGACAGAGTGCCCGCGCT

GTCTTAATTCAGTCTACGAACGAATGTTCTGCGGTACTCTCGGTAGAAGCTGGGGTCGATGTACGAGGCC

CTATTTATTACGTATTCCAGTCTGCATTCTCGACGGCCTGCACCAAACACCCCT

>NODE_725_length_264_cov_2.543210

GTCGGGCCACAGCAGCAGCAGAGAGGGACCTTCCTTCCCGGGGGTCATTTCAGAGTTTTTTATCGCACGC

GCTGTGTGTACTGTATTGTATGTACATCACGAACGAAGTATCCAAACGCATGTGCATGGGCTAGTAGGGC

GGTAATCGGGTGTTTGCCGGTTCGCCCCCACTAACATAATTCTGGTCGTTGAATATGGGGCCACACTAAC

TTTTTTCCGTAGAAATATAGAGCTGACACACCAAGTCGAAATAAGCGGGAGTTG

>NODE_726_length_264_cov_2.526749

AGCTCGTCGAATACAGTGACCGGCACCGAGGTGAAGTCGTTAACTCCAAGATCCAATTCTATGAGAGATC

CGATGCCATCGAATGTGCCCGCAGGAAGGGCAGTGAACACGTTGAGCGCTAAGTCCAAGCGTTCCAGTGC

CGGTGTTCCCTCGAAGATCCCTGCGCTCAGGGCGTCCAAGTCATTTCCCCTGAGCGACAACGATGTCAGG

GATGCCAAACCCCCCAGCAGACCCGATGGCAAGGAGGCTAGATCGTTGCTCGCA

>NODE_727_length_264_cov_1.876543

GTATTTGTAGCAACGGCTATGTCTATTGTTTCAGGTGCTGTAGCTGAACGTATGAAATTATGGGCATTTT

TAGCATTTACTGTTGTATTAACAGGCTTTATCTATCCAATGGAAGGTAGCTGGACTTGGAATGGCGATGC

TGTATTTGGTTTATACACTTTAGGTGACCTTGGTTTCTCTGATTTCGCAGGCTCTGGTATTGTACATATG

GCTGGTGCTTCAGCTGCATTAGCTGGTGTATTGCTATTAGGTGCTCGTAAAGGT

>NODE_728_length_264_cov_1.839506

CCATTACCCAGGGGTCGAGGCGAATACCCCGGGCTCGCTTCGAAAAAACTAGCGGACGGGGGGGTCAAAT

AGGTTTGTGTTCAATATAACCTGCTCGGGATAAGGGGTTTTTCATTTGTGTCAGGAATTCAGTTGGGATA

GAAGTATACCCAGCAACCCAGTTATAGGGATAATAGGTTGGGTGGAGAGTTTTGTGTTTTGTTGCGTACC

TGTATTTAGTGAAGGGGTATGGTAATAATGACAGTGCTACACACTGTCGTGGGC

>NODE_729_length_263_cov_4.900826

AATGTCTAGAAAAACACTCGCCGTATACTCAATCAATGCGCCTGTTCGCACGTATCATTCGTTCGCTCGG

TATTAATTATCAGCCCTTTTTCGGTCGCGCTAAGACTCTTCGCTTGCTCGTGTTGCCTTTTTTTGCTCCT

TTTTCTTCTCCGTTTTTGTTTCTGTTGTTTTTGCTTCTTTTTTTTTCTCGTTTTTGAACGTTTCTCGCCG

GAAAAAACCCGCGCACTCGCTGCATAGTGCCAGCCAGCCCAGCCCAGCTCAGC

>NODE_730_length_263_cov_4.615702

CGGGAAGGGCTTCTTGAGCAAAGACTTGCCCACCGACTTCTTGGGCTTGTTGCCGTTCTTCGACGACTTC

AAGGACATAGACGCGCAGGGGCGGCCCTTGACGTCCTGGTCGAACTCGATGCCGACCGCCTTGGTGTTGG

CGAGGCCGGAGTACTTGTACGAGTTGACGCTCATCAAGTTGTTGGGCTCAGAGGTGAGCACAACCTTTCC

GGACCGCTTGGTCTGGCCGTTCCTCTTGTAGAGGAAGCAGGTGTTACTCCGTA

>NODE_731_length_263_cov_4.103306

GAAAGTCCGAGCTTTGTACTCGTGATACTCAGGAAACGAGGGTATATGTATGAGTTCCGTGCCTTGGCGT

TGAAATGAGCGAGGTTGGATGATCGGGGCCTGGCTCGCACAGCAGATGAGAGCAGCAGTGTGCGCTCGCT

CTATGTAAAACATGTGCGATTACGGTTCTGAAGGTTTTGAGACATCCTCTCTGGCGATTGCCTTTGCAGA

TCGAGATATATCTCACACATGAATGATTCCGCACGGCCCCACGCCGTGTACAG

>NODE_732_length_263_cov_3.842975

GCTGTACCTTACGGCCAGGTCCCCTCGCACTCGCCGGTCCTTCTGGCTCGGCATCCCCTCTCCCTTGATC

ACAGTCTCGTGCGACGAGGATGATATCCCGGGTTCCCTTGCTGTCACTGTTCGCCCGTCCAGAGTTTCCA

CAGGGATACTGAACCCTGACAGCGCCTGCTCGAGCGTGATGAGCACCGTGCAGGAGAGGTTGTTCCCTTC

CCTCTTGAAACGCGGGTGGAGCTTGGTCTTGATCACGAACACGATGTCTGAGG

>NODE_733_length_263_cov_3.549587

GTCGCTCGTGCTCAACAAGCGCAACTCGTCCACATCGCCCCCGGCGGAGGTACCGTGCACCTGTAGAGTG

ATTCCGTCTTCAACGTACAGGGTCTCCGTGAGAAGCCACGTCCCTGTCGCGGTACTGCTCACGTCGCCAC

TGTCCGGGTCCACCGCGTACAAGGGAGGGCTGCCGTCCAGGTCTGCCCAGATTTCGGTGAGAGTCATGCA

GCCGCCTCGCGCTGAGCCTGCCTCCGACTCCAGGTACAGCCTCGCCGATGACG

>NODE_734_length_263_cov_2.727273

ACTGGAGTTCCTTGGCAGCGGTGAATTCCATTTTATTTTGCTGTGATTGCATTGGTCTGATCTTGCGATC

TGATTTTGTGTTGATTTTGTTGTGATGTTCCCGTGATCTTGCGCTCTGATTTTGCTGTGATTTTGCTGTA

GTCTTGCTGTGATTTTGCCATGATCGAGGAAGAGGTGATGTGCTGCATCCACACTGAATGTGACTGACCA

ACGTATACTATGTCTGGAATAATTTAAATATTTCAAAACCTTAAAAAAAAAAG

>NODE_735_length_263_cov_2.458678

GTGAATCTCAAGGATACAGCCATCGCGAACCACCAGCTCGTTGTTGAGGGTGTACTCAGTCCCGTCCTTC

GTTACAGCCTCCGGAAGCGCAAGGTGGATGGCCTCCATGGTCACACGGCATCGCGAGCCTCTGATTTCGA

TCACGTCATCCTCAACCGTGATCGAGGATGCATCGCAGTCGCTCGGGGGGCCGCAAACATCATCCCCGGT

GCAAATGTTCACCCCCCAGCAGTGGTGGTCACCGCACGCGTTGGCTGCACAGA

>NODE_736_length_262_cov_6.522822

AAAATTTTTTTTTTTTTAATAAGTATTTTAATTACTTTATTTTGCATTTTAAAAGGAATGCTAAGTATAG

AGTAAATTAATTTATTACACTTACTTTATTATTATATTTAAAAATAATTAATTTATGACAACCCAACAAT

TTTTTAACTTAATGCCCTACTATGGGGAAAACTCTGAACCAGAATTAAAATCAAAAGAAATAGAACCAAC

AGAAAAAGTAGAACAACAAATATATTATTTTTCAAAAAGCCCTTTTCGAAAT

>NODE_737_length_262_cov_5.120332

CGCCGCCTTCGCCTACTGCTCCGACATCGCGCTCACCATGGGCGGGAAGATCAAGTTCGCGGAGATGCTC

TCGGGGCGGTACGCTGACGTGTTGAGCGGCCTGTTCCTGGGGTACGCCACCCTCTGGTACACGTCCCACC

ACAAGAACGTGAAGGGGCTCGACAAGGTCACGGACTTCGCCATGCAGCGCACACTAGCGGACATCGAGGA

CGCTTTCTACGGCATCTTCGCCAACTTCCCCAGCAAGACCCTGTCGCTCGCC

>NODE_738_length_261_cov_28.429167

GTAGGTTTTAGTGTTGTTGGTTTGAATGGTTTTATTTGTATTTCTAGAATTCGATGTATTTGTTCCACTG

TAGTTGACTTTTACAAATCTAACTAAAACAGGGTAATTTGAAGCAACTTTTTCTACTATTACAACAGTTC

CAACATCGTTATACCAATATGATTCTTTTCTTAAAATACGTACTTTTGCTCCTTTTTCTACCATAGTATT

TTTGTCCCTTAGAAATAATTTAATTAACCCTTTACACAACTCCAATCTAAA

>NODE_739_length_261_cov_6.970833

CTTGTCGAGGACCCCGTGGACGTCGGGCATGACGTTCTTGCCGTCGACGGCCATGTTCGCGTCCTTGGGG

GCGCGGAGGGCGACGTGCATGACAGATCGGTCCTCGGTGTTGTTGATCTTCTCGCCGGTGTACATCGCCG

CGCGCTTTCCGGCGAGGTCAGCAGCCTCGGCCAGGTTGAACAGCATGTCCATGGTCTCGTTGGTGACCCG

TTGGCGGGAGTAGTCGAGCAGGATGCCCTCGCGCTCGGCCGTGAGGGCGGA

>NODE_740_length_261_cov_5.295833

GCGCACGTAATGACCATGGGCGACCACGAGGTGGAGACGATCCAGGCGTTGCCCTTCGTCGACCTTGTCT

GAGCGTGCTCGAGACTCTGGCGCTCCGTTTGCAGTACAGGCCTACGACGACTCCTGTGTTCGTGGATCGT

TTTGGGTTGGTGTTTCTCTCGGGGCGAGCTGGATCTGGGGATGCCAGTTATACGGGAGTCGTTGTTGCTG

ACGCCTTGTTGTATTCTTCTTCCCAGGCCCGGTTTGCTGGCTTCCTCACGT

>NODE_741_length_261_cov_3.804167

ATTCCATATGTTATTATCAGTAATATCAAATGGTTCGTTTGTCCATTATTTTTCTGTTTTTTTCGGAAAT

TTCGGTATTTTCTAAATTGGGTCCTTAACAAGTACGAGCTTCAGTTGCCTCGCTAATAATCGTCCACCAG

GCACAAAACCGTCACTACAGCAGCAGCGCTCGCGGTAACCATCCGTGTGGTGGTCCCACTGCGGTCGGCC

TGGCTCGTAGGCAGGTATCGGTGCCTCGGAGAGTGGAATCCATCACCAACT

>NODE_742_length_261_cov_3.612500

CCCATCATGGCTTGCATCTGCCTCTGCGCTTCCGCCATCATCTTCGGGTCCTTCATCAACTGTTCCACGT

TGGCTCCCGCCTCGGCCAATTGTGCCTTCATGGTCGGGTTGTCCATCATCGACCGCATCTGCCTCTGAAA

GTCGGGGTTCTTCATCGTTTCCTCCACCTGCTTCATGACGGCGGGGTCATTCATCATCTCCATCGCTTCC

TTCATAAACCCAGGGTCCATCATCCCCATCGCCGCCGCCTCGAGCCCCATG

>NODE_743_length_261_cov_2.958333

CAGTATTTGTCGTTCCCATCTGGACGTCCAGCGTGTCCGGCTGAAACTGCAAGTTACCGCCGAGGGCCTT

GTACTTGTTGACGATGTTGGTGGCTCCGACCACCTGGTTGCTCTCGATTGTCAGTGTCGATGTCGGCTGG

TACAGCGCCCCCAGCTGGTCCAGCTGTGCCCCGCCCGAGGCGAACTTGCTGTAAAAGTGCGCCACAAAAG

CTTTCGCCACATCCTCCCCCGACATGGCTCCCCGCTTATTTGCCGCCTCCG

>NODE_744_length_260_cov_13.389121

TTGATTACGTCCCTGCCCCTTGTACACACCGCCCGTCGCTGCGACCGATTGGATTGCGTGGTGAAGGCTT

CCGGAGGATGCGTTGGTAGCTTGCTGCCCTCTGCGTCCAAAGCTGTCTGAACCCTGCACTCTAGAGATTG

CAGAAGTCGTAACAAGGTAACCCTAGGGGAACCTGGGGTTGGATCACCTCCTTATCTTGAAGTAAAAACG

AATAATGGAAGCCTCGGCTTCAGAGTATTCACACAGGCTCCATGATTACC

>NODE_745_length_260_cov_11.309623

CAAGATGGCTATGGATATTGCTGGATGCAATAAGTGGCAGGAGCTACTTAAAAGGATTACCAAGGATGAT

GCTACGGTATGTAGCCAAACGATAGATTTCGTGGGTTTGCCAAGATGGCTATGGATATTGCAGGATGCAA

TGTGAGGGTAAACAAGGTTGTTTACTCGTTCAGGGGATGATAATACGGATTTAAGATAATGGATATCTAG

CAGGATGCAGTCCAATTGGGGCGTAACTTTAGTTACGCCTTAGTTTGAAC

>NODE_746_length_260_cov_10.393305

TCCAGGGCCACCTTGCGCACCTGGGCCTCCGGCTTGGGGTCGTTCAGGACGCTCTGGATCTCATCGGAGT

TGATGATGCGGTTGAGGTCGGTGTTGGTCATGCAGTGGCGGGGCAGCTGGAAGTTGCTCTTCACGTCGGA

CTTCTCGGTGTAGGTGCCGTAGATGGAGTCCAGCTTGTCGAACGCCGCCTGCGTCCAGACGCAGAAGCGG

CCCATGTGCCCGCCGGGCGCCAGGGTGAGCAGGTTAAGGCGGTCCACGCA

>NODE_747_length_260_cov_5.301255

AAATCCGTGGACGAGGCTATCGGCTTCTGCGACAGTGGGCCGATCCCCCTGGCGGCTTACGTGTTCGAGC

GAAACTCAGCCGTCCGGGAGAAGTGGCTGAGGGGAGTGGATTCCGGCGGCGCCTGCGTCAACGACTGCAT

GTATCACGTAGTCAACAACGGAGGAGGCTTTTCTGGCAAGGGGGAGAGCGGCATGGGCACGACGAGGGGT

CGAGCCGGGTTCGAGACGTTCTCGCACCGCAAGGTCGTCGGCATCCAGCC

>NODE_748_length_260_cov_4.280335

GAACAGGAACACCACGCAACTATAGAGCGCTTCGCCTGCCGGAGTAAGAGCGCACGGCACCTCTCAACAC

GAACTTCGCCCCGCCCGTACCACTCGGCCTCTTTCCTACTGAAACCCTGACTGAAAAATGATCAACCGAG

CAACAAACCAACGCAACGCAACGATCTCCCAAGATAAACCACCGGTAAGCCAGGGAAACAGGCATGAATA

CAGCAGTATGCAGTCTTCATCCAACCTACCATTGGTATTCGGAGTTTGCA

>NODE_749_length_260_cov_3.640167

AAGAAGCCCAAGGTCGCCGCGATCCGGCCAAGATGATCGCGAAGCTGGATACAGATAAAAGCGGGGGTGT

GAGCCTCGAAGAGTTTGCAAAGGCGCGTCACCATGGCAAAGGTAAAGGCCATGGTAAAGGTGAGCACCGC

GGCAAGCATAAGACAGACGCGAAAAAAGCTGACTGATCTTTGAAACGAACACATCGGGCGGCCCGTTTGG

GCCGTCTGGATCCGAAAGGGGCACTGTGAAAGATTTGCCAGAGCATATTG

>NODE_750_length_260_cov_2.029289

AACCGTACGTAACTATCTGTTGTATATGACTCTGCTTAGTTCTTCTCCACGCCCTACTTCTCGGCTTGTT

GAAATTTGAGAATATATAGGTAACCATAGCATACCATAGCATAGCATATTATATAATGACGGCGGGGACT

AAACGACACCTAGACAAATCCAGTCCCTTTTCTGCTCGAACGGCAGGGGAGGGGAGGGGGTAGTACAGGA

GAATTGATGGATAGATCATTGTGACCTGCTGTAGAGGTCGATCCGACCAT

>NODE_751_length_259_cov_20.609244

TCCCTCTCCGGAATCGAACCCTAATTCTCCGTTACCCGTGATAGCCAAGGTAGTCCAATACACTACCTTC

GAAAGCTGATAGGGCAGAAATTTGAATGAAACGTCGCCGACACAAGGTCATGCGATCTGTTCAGTTATTA

TGATTCACCATTAAAACCCGAAGGTATTGGTTTGTTATCTAATAAGTGCCAACCATTCCGAATACTAGTT

TGGTTGATTTGCATGTATTAGCTCTAGGTTACTACAGTTATCCATGTAG

>NODE_752_length_259_cov_16.781513

CACGTCGATGGAGTCGTACTCAAAGTCCGACAGCAGGAACTGAGACACAAGGGGCCCGGTGGTCACACCG

GGCCAGTCCGTGCGGAACAGGTCAGTGACGGGATCGACCCTTGTCTCATCACTTCCTTTGTTTCCAACCA

CGGTACTGGGGATGTTCAGCCCGCGGAATGCCTCGTTGCTCCCAAGGTTTTCCGCGGCGAGCCTGGTGAG

ATCGTTGTTGGCGAATTGCGAGAAGGGGATGTCCCTAGCCAGGGCCATC

>NODE_753_length_259_cov_5.966387

TTTATCTCCGAGCACTCAACACATCAGCCATCATGGCTTCCTCTTCTCTGCCCGCTTGCCTCCATGCCAA

GGAGGAGGACATCCAAAAGATGCTCTCCGCCCAGGTGCACATCGGGACTCGCAACAGCGACTCCCAGATG

AAGGACTACATCTGGCGCCGCCGCCATGATGGCATCCACATCCTCAACATCGGCAAGACCTGGGAGAAGA

TCGTCCTGGCCGCCCGTATCATCGTGGCCATCGAGAACCCCCAGGACGT

>NODE_754_length_259_cov_4.882353

TCCTCAGCACCAGCACCTAGCAGCGACGACCACCACCAATTCCACAGACAACAAAGCACCATGTCTGAAC

ATGTCGTGAAGAAGCAAAAGGTGGTCCTCACGCAGTACGAGCAGCTGGCGGCGTTGACGACGATCGTCGC

AGACACGGGAGAGATCGCGCAGATCAAGAAATTTGCGCCGACCGACGCTACTACCAACACTGTGTATGTC

ACGCTCCTTGGCACTTGTTTTTTCTCGACGTAAGTTTAACCCTTTCCCT

>NODE_755_length_259_cov_4.693277

GTTCCTTTGAATTCGATAATTCCAGTCGTCGTTGTTTTTTTTTTATATATTTTCTCTTTTTAAGCACAGC

ATATGATCCTCAAAACCACGATGTATTCTACCTTAACCGACCACACATCCAGTGGTCCCCCCTTGACTGC

GACGAGAGTGTGGAGGACACCACACAGATGCGTTTATCTGCGGCGACTTCCTCCCTCAACATATTACACA

CACTCCATTTCTACACCATTGACGATGGCGAACTGTTCGCTAGATATGC

>NODE_756_length_259_cov_4.294118

GGTCAACAACAGCCAGGATTCATCGGGCCAGCGGGAACTCAGGGGTGGCAGGCACCGTGGGGTGGTCAAC

CCATGATGCAGGGGTGGGGACCCCAGGGAGGACAACCACCCTTCCCACTAGGAACTGATGGATCGCGAAT

CGGACAGCTTAATCCGTTATACGACATAACCGGAGGGTTTTCTAAAGAAGCCCAACGCATGGTCGACCTC

CTAGTCAAGGAAATGGGGGTAGAAGGACGGAAGAGCGCTTACGAGTCGG

>NODE_757_length_259_cov_2.008403

TTCCTACTATACCTGTAGCATGGATTGATGGATTGTTCGTGTGATGAGTGCGTGGATGGGATGGGTGGTA

GAGTGGCCTGTGACAGCGAGAGAGATGGCGAACCGATCGTTGCAAGAGAAGTGGTTCTTCTCGGTGCCCT

ATATGCTACTACTATTTACGATGATGGCAGGCACCGGTTGTTCGTGATGTAGGGTGAATGAGTGATCTGT

GGCAAGCGATTGTTATCGCGAAGCCGATGGTGATTAGATGGCGGTTAGG

>NODE_758_length_258_cov_37.569620

TCGATAGCGCGGAGGTACAGATTCATTTATGTTATCCGCAAGAGGATCAAGCTCGACCCGGAGAAGGCCA

TCTTTATCTTCGTCAACGACACCATCCCACAGACCTCCGCGCTGATGTCTCAGGTTTACGAGCATCAGAA

GGACGAAGACGGGTTCCTGTACATCACATACAGCGGGGAGAACACTTTCGGGTAAAAAACAGAAGAAAAA

AGAGAAAAATCAGAGCATCGACCGCGTGTGTTTCCATCGACGTCCAAG

>NODE_759_length_258_cov_10.008439

AATTTTCTATAGTGAATTCTTTATTTTTTTGTATTTTCTTTTTTTTTTCTCCGTTTCCACGCTGGTGGTT

TCCCCTCTCATCCGCCAATGTCGCCCCTCAAAAGATCCCGCCTCCCTATCTCATTAAGAAAACCAAGAAG

ATCGTCTACTAAAAAAGGGGGGGCGGCACATTGGGACGCGAGCTTCGCAAAGCTGTATCGACACGGCGTG

TTATCGCATTGTGGTCAGCCTCTACGTGTCTGTAGGAGCCCCCGCCGT

>NODE_760_length_258_cov_3.843882

AAAGTCAGCGGTCTTCACGAACAACGTGGCCGGGGGAGACGGAGGGGCGATCGCCAGCCCTGCCTACGCC

ACGTTGGATCTGAACGGAGACGTCATCTTCTCGGTAAACTCGGTCACGGATGACGAAGGCAACTGTGACA

ACTGGTACCAGAAGACATCGGAAGACGGCACTGACTTCCTCTGTCTCTGGGACTAGGCTCATGAACATCA

GCTCCATTGCCGTGAAACATGTGGTCATGTCGCGGCAGAGAGATTGCT

>NODE_761_length_257_cov_2628.737288

AACAGACACTCCGACAAGCGAAGCTCATAGAGTATGGTCGCTTGAAGGTCTTCACTGTCCAGTCGAAGAG

CGCCGCAACTATAACTGGCGAATCTGCACACCCAGTCCCATCATACCTCTTTCGGAAAAAAAGCAATCAA

GAAAAGGGCGGAGACTCAAACCACGGCGGCGCCAAGTTTTTACGATCATCGCTTCGACTGGCTTTGTTAG

GCTAAGGAGGCGGTCAAATTTCTGAACCCCCCCTTTGGTCCGTAGAC

>NODE_762_length_257_cov_34.940678

CCTTGGCACCCGAATTCCAAAATGTCCGGACTCCTACGTCCCATAACCAGAATTGCGTCGTTTTGATCGC

TTCAAGCACGGGATAGCGAGGAGCCGATTCGGGGGAGGGGGAAAACTTAAAAGAGACAATCGATGGTTGA

AAACCAATCCCCTAAAAAAAATAGCCCTCCTCCCCCCTCTCTTGCTCTTGCTCGAAGTATTTGGGAGGTA

CAGGCACCGTTGAAAACCCCTGCATCCCAGCTAGACCCTCGAGCTGG

>NODE_763_length_257_cov_15.470339

GCTAGGCTCGCGTTTGCTTCGCCCCGCTCTGTTCGCTGAGGAAGACGAGACGGAGATAGGAGTCGTCTTC

GAGGCACTTAGTCTCTCGGCGCATCGAAGCGAGAAATCGTTTTTGGTGTTGCTATGGATTTCTGAGGCGT

GTTTGTTTCGCGTCTTGTGGGTCCGGGACACGGGGGACATTGCTGTTGGTCGTCACTGAACAGGCGACGG

CATTAATAAAAAAAAGCGGATTGGGAAAAAAAAAAAAAATTTTTGGA

>NODE_764_length_257_cov_11.665254

GCACCCGAAGAATTCCAAGCAACGACGTCAAACTATCTACGGCTCCTGCACGGGGAACTCGAAGACAACG

TCCTTGTTGGTGATCTTCTTGTAGACCGAGGAGAAAGTGTCCAACTTCGTCTCCACGTTCACCTGGTCCT

TCGGGTCGAGCTGGACCTTGAGGATCTTGGAGCCGTCCACCTTGCACCTTGTGCGCTTGCCGACAATCTC

CGTGGGGAACACGAGGTCCTCGAGCATGGCGTCCTGTACTGCGGTGA

>NODE_765_length_257_cov_10.169492

ACCAACAAATTAAATAACTAAAAAGCGTGCCTTATTTATTACAAAGATTTCGTAAATTTACCAACAAATA

AAGATATTTCAGTCTGTTCTTTAGGGCGCTTTCCTGTCCATACTTGATGGTTGTTAAGATCTAAACTTTT

ACAATTCAACTCTTTTTGAGAAAAATGGCTAGGCAAATCTAAAAAATTAAAACTGCCATCTGACAATAAA

CTACCAAATGTTTTTGCTTTTACTATTGATTTCATTTAATTTGTGCG

>NODE_766_length_257_cov_7.364407

TTGCCGTTGATGGGTTGAATTAGAGCTGATATTTTTGTTTTGTATTATTATTATTGTTAGTAGTATTGTC

ATTCCTTGTTCATTCAGTTCATTCATTCATTCAGGTCAATCAAGTCAAAGTCAGTTCATTCAATTAATTC

AGCCAGACAAGTTCATCCTTCAATATACATATTGCCTACTTAAAAGGAATGCATCAAGTTAACGCATACA

CCCCACCCACCACTCACTCGCTCGCTCGCTCGGCACGCCAAAAGAAA

>NODE_767_length_257_cov_5.228814

CGGAGTACATCGGGCACGCCGGCCCGGGCATGTGGCTGCACAGCTCGTTGTTGGTCTCGGTACCGGCGTC

AAAGGCGGGCGGGAAAGACTTCGAGCCGGAGTAGAGGTCCTCGGAGTCGATGCCCATGAAGGCGTCGTTC

GTGTTAATGAGCATGGAGACGGTCGAAAAGACAGCGTCTTCACCGCAGATGTCGGTGTCCACCTCCATGG

TGCCCCTCCACGTCTCTCCGGGGAACAAGGGCGCGTCGCCGACCGCG

>NODE_768_length_256_cov_183.153191

GGCACCCCGAGAATTCCAGGTACTTTTCGTGACGACCAAGGACCGGGTCTCAGCAGGCGTTGCCCCTCGA

TCGTTATGCATACTTCCCAAAATGGGAGGAGCACGGTTGGCGAGGGGTGTCATGAACCCGAAGAACAATA

CCTACGAGGTAGCCAGTTTTTAAGCCTGCCCCGCCTTCTCCCTATCCGTGCACAAAAAATTGCGCGCGGG

CAGGGAGGGGATCCGTGCGTGGTAAGACCTGTCGATTGGTACATAC

>NODE_769_length_256_cov_5.272340

CAGCTGGGCGACGGCGAGACTCTCGACTACGTCATTTGCAACGCCGGGATTTTGACCCCCGACACGCTCG

AAGACCCCAAGTACGACGCAGCCGTCAAGATGTACGAGGTGTGTGTGCTTGGGCCCCTTCGCGCGCTGCA

GTCTCTCACCGTCTCGGGGCTGTTGCGGGAGGGGTCCAAGATCGGGATGGTCACGTCTGAGGGAGGTTCC

GTAGGATTGAGGACGGAGATCGAGGGCGGTGGAATTTCGGGTGCCA

>NODE_770_length_256_cov_4.255319

CGAGGACCTCGATGGCTCGGGGGCTGTGCTGACGGCCACCGTCGACAACGTGTCCGGCGACATCTCTGGC

TTGAGCCTGACCGCTTCCGGCACTGCTGGCTACATCCCCTTCACCGTCCCTTCCAGCAGCTCGATCTCTA

CCTCTGGATTGACCGTGGGTAGCACTACCACCTACGGTTCGGACACGACCTACTACATCGCCACCTCGTC

GAACGCCATTCCTTCCCAGGCCTCTGCCCCTGCCGCAAGCGAGCTT

>NODE_771_length_255_cov_665.431624

GCCGGGAGGGGTCACGCCCACCTGTTACCGCGATGTGCGGGGTAGGATTCACTTGACTACAGTGTGATCG

CTAGATGTTCGTCCGCCGGAGCTGTGGAGTCGCCAGAAAGTTCTTACTGGAAAGAAAGATCTGCCTGCGC

GATAGAGTGGTGGTTACCGCCTCAGGAGGGCAACCTAACAGGGCGCCCGCCCGCTCGACAGGGTCCGTGG

ACCGCGAACTGAGATCCGTGGGATATGTCTCCGTCTCCCTCTTTT

>NODE_772_length_255_cov_271.679487

GGCAAAATAGATCCGTAACTTTGGGAAAAGGATTGGCTCTAAAGGTTGGGTATATGGGTCTATTAGAATA

AATTATTAGATAGTATAAATACGACGGCTTTCGCAAGATGGCTGTTGTGTTTGTATTAAAAGATAATGGA

ATATGATAGATGAAAGAAATTTCATACTGTATACAATGAACAACTAATTTAGAACTGTTACTGACCAGGG

GAATCCGACTGTTTAATTAAAACAAAGCATTGCGAGGGCCGGAGA

>NODE_773_length_255_cov_7.713675

GAGCGAGTCGGGCTCGGACAAATACGAAACGGAGCAAACAAGCTGTGCAATAAAGTCGGACACGCGCATT

ACGCACTGGCACACTCCTTGTTGTGTGGTAAATAACACAGCGAGCGGTACAATGTATGCGAACGATTGTT

TTTTGTTGTGTACATATTTCCATACCTTTTCTCCTGCTCTACTGTAGTAGCGATAAACACACCTGCCTAC

GGACAAGAAAAAACTCCTTCTGCCTAGCGGGGGTGCACAGTAGAG

>NODE_774_length_255_cov_3.350427

ACTCGCCACCGGGTAGTGCTGCTGTTGGCGCATGGCATGTTTTCACCATCCCTGTTCTCGTGGTGCCTGC

CGACGGCGGTAGGGGACCTTGGAGACTCCCTTGCGTTTCGTGGATGATAGCTGTACGTATGTTACCATTG

GGGGAGCGTTCCTGGGGGGCTGGCTGTTGTAGTTGTTAGAGTAGACGGCGCAGGGTGGCAGGGCGGACAA

GAGGGAGGGAGCTACGGCTTGGTTTTGTGTTGCCTGCCTACCTAT

>NODE_775_length_255_cov_2.854701

CTTGGCTCGCGAGAATTCCAACCTCTTCCTCGCCGCGGGTGATGGAGGCGAGGATGTTGTCCTTGGTGTC

ATTGATGCCGGCCTCCAGCGCCGAGAAACGCTCCGTGTTGGCCGTCCACAGCACCACCACCTTGTCCAGC

TTGTTCTTCTCCTTGAACTCGCGGATGTCCTTTCGGATCTGGTCCATCTGCTCCTGCTTGCTGCCCTTCA

GGACGTTGTCCGCGCGCTCGGACTGGTTGGCGGCGATGAAGTCGG

>NODE_776_length_255_cov_2.564103

CAGGCGCAGAAGCTGGACGAGATTCTGGCGACGAAGGGTGTCGCGATCGACAAGGTCATCAACCTCCAGA

TCGACGACAAGATGCTCATCCGGCGTGTGCTCGGCAGGTGGATCCACCCGGGGTCGGGACGTAGCTACAA

CGTTTTCTCTAGCCCGCCGAAGGTGCCTGGGCAAGATGATGTGACGGGCGAGCCTCTCACACGTCGCTCG

GACGATACCGTGGACAAGCTCAGGAGCCGTCTGAAAGCGTTCCAC

>NODE_777_length_255_cov_1.905983

ATACAACTGGTTTTGCATCACAAGTTGAATTAGATATCGCGCTTAATGAAGATGGTACTCCACAAGTAAC

TTTAGCTGCTGACGGTATGTCGTTAACAGTTGTACTTGCTGAAGCATTAGATGACGAGCAACGTTTCTCA

ATCTGGTTCCCACAATGGCAATACACTGATACAAGCGGTAACGTGATTGTAACTGGTGATGAAACTGGTT

CTACTGATACTGATGATCTTCTTAAAGATAATGGTACAGGTGATT

>NODE_778_length_254_cov_9.034335

AATAGGCCATACCGCTGCAAAATTAAGGCCCCAGGGTTTTCTCATTTGCAAGCTCTAAACTTTATGGCTA

ATTCCCATATGCTAGCTGATGTTGTAACTATAATTGGAACGCAAGATATTGTTTTTGGTGAGGTAGATCG

TTAATTTTTGTTTAAAACAAAAAACTTTAACAGTTAAAATAAGTTTAGGCATTTACTTTGGGTTTTATGC

GATTCGAGAGGTGACGCAGTGGTAGTGGAATTCTGGGTGCCAAG

>NODE_779_length_254_cov_6.171674

TAGGTCTTTGTAGAGTTTGTTGTTTTTGGTTTTCATTGGGTTGAGTTATTTAATGGTGTTTTGTATTTTG

TTTTATTTTTTTAGTATCTCCGTTATTGGCAGTTGACCATCACGACGTACGTGCTCTCAAAAGAAAACAA

AACCATCAGTTAAGAGTCGAAAAGCCTCCTCCGCCCGGCAGAGGCGAGACAAGAAAACACTTCCCCCGAC

CCTTCCACTCCTTTCGCTACCGGCTACTACGAAAGTAACAGGAC

>NODE_780_length_254_cov_5.819742

TGCAGCGCGGCGACCACCCCATCCAGCTCTACTCCTTGGCGACGCCGAACGGACAGAAGGTTACCGTGGC

GCTCGAGGAGATGGGGCTCAAATACGATGCCTGGTACACCGACATCATGGCGGGGGACCAATTCACCTCA

GGATTTGTCGAAATCAACCCCAACTCCAAGATCCCGGCTCTGGTGCACCGTGACGCTCCAGGTGGCAAGC

CTCTTCACCTGTTCGAGACAGGCAACATCCTGCTGTACCTTGCC

>NODE_781_length_254_cov_3.789700

AGGGGTTTGTCCGGCCTCGAAGAATGCGCTGTTGGTGGGTCGGGTATCATCTCAGGGCAGGACATCCTTG

AGGAGTTTGGGTTCTCGAACGTGAACGTCTGGGTGAACGTTGGCGCCCTCGCCGCAATAATCATCGGGTT

GAGATTCTTCACGTACCTCTTGATGCGGAGGTTTACGAAGACCAGAACCTGAGCTTGCGAGCGAGCTTGC

GAGCGAGCTAGTGTCGTGTCGTGTCGTCGCGGGTTTCTGTGAGC

>NODE_782_length_254_cov_3.098712

GCAAGGAATGGTACGTCAATGGCAAGAAACATTTTACGAAAGTCGATATTCTCACTCTAATATGGTAGAA

GGAGCACCAAAATTTGATGTACTTGCAAATGCCTATGGTATTAAAAGTATGTATACTGAACGCCAATCGG

AAATATGGCCCACATTACGCGATACCTTAGAGGGGCCTGAACCTGTCTTACTTGAATGTAATGTGCTAGA

AGATGAGAATTGTTACCCTATGGTTGAACCTTCAAAATTGGAAT

>NODE_783_length_254_cov_2.849785

ACACTTGATTGGAATGTTTGATGCAGCGTAAGGAGGGGACGACGTTTTACTCGTTGAACGCATTGAAGTG

ATTAGGCTGGTGTGTGGTCGGCAAGAGCGTCTAATCATATACTTCGTATACGTGCATGCATGCGCATTTG

GGTGTGTTTACGCACATACATAAAGCGGTGGTTGGGGTGTGGTTTGCATGCATGAGGACGTAGTTGTTTA

TTTTGCGGTCAAAATAATAAAACGATGGGAAAATCAAAAAATTT

>NODE_784_length_253_cov_121.849138

ATTCCATGGTGGAGCGGGGGGGAATCGAACCCCCGTCCAAAAAACCTACATCCTCGGTACTACATGCTTA

GTCAGTTATTTGGTTAACTCGTTAATCTCCAACTGACAGGATATTAAAAAGCGAGTCCGATACTATTTCG

CGGTTCACCCTCGAACAAGGTTCCCTCGCTATCCGATGTAAATGACCTTCCGAATCCCTACCCATTGGAA

AAGCGCGGGTGGAAGGCTAGCTAGCCTAAGCTGCTAGAGAGTA

>NODE_785_length_253_cov_6.534483

CTCGGCCGGCGAAGACGGTCTGCGTGCGGCTCTCTCCGGCTTCGTAGCCACCGACGAGGAGGGTCTCGTT

AGGGTAGTTCGAGTAAGCAGGGCTGCCGTAGAGGTAGTCAGCGCCCATTGCCGTGTAGGTCAGCTCAAGC

GCGGCGATGCTCTCGGTGTAGCTGTAGTCGTACGAGTAGTAGTCGTCGCTCATGTAGTCGTCCGTGTAGT

CGTCGGTCGTGTCGTAGCCGTACGAGTACGAGCCGCCGGAGCT

>NODE_786_length_253_cov_3.689655

TTCCAGGGCACGAACGGCGCCGATGGGGTGCGGGAGGTAGCACGTCCTGCCTCCTCTTGAGCGTTCGCCG

AGTCGGGCTCATTCAAGTCCGCATCCGGCACGACATCATAATCGTCGCCTGCAATCTCCTGATTTTCCAC

GACAACGAACCCCGCCTCGCCGAGAGGGGCGGCTGCGGATGTAGTACCGGGGTGCGCGCCACCACCGTCA

GGGTCGTTAATTTCTCCTCCCATGCCGCTTAGAGAGGTAAGTA

>NODE_787_length_253_cov_3.383621

GAAAAAACGGACACCGTGTATCGCACCAGTGATCAACCAAAAACGAACTACTGCAGTTTCCTTCGTCACG

AGCACTATTTATTGTTTCGTCGCCTCGGCCAAATTCGAAGAAACTTTTGAAACATAGAACGTCTTCCGCG

CCCTCCAGTTTTTTTTTCTTGTATCGCCCAATTTCCTATCGATGCCGAAGAGTGTTGAGTCAAAACGCAA

CTAGAGAGCTATGATAGCTGTCCATCGTCAGTGTGTTTTACCT

>NODE_788_length_253_cov_3.068966

ATGAATGTAGGTAAGATTATTAAATAGTTTTTAGGTTTGTTTTCCTTAAGTTCTTATTAAATAATAAAAA

TAAATTAATATAAATGTTATGACAGACACCTTAATGTTAATGGTAGTAGCATCATTCGAATGGCCGTCAC

TTAACCCAAATGATTATACAAGAGCAGAAATGTTAAATCTTTTGGTAACGGCTATGGTGGCAGGACTAAG

GCAGTATTATTGGATTTTAACTTTACGTTTATCAATACAATGG

>NODE_789_length_253_cov_2.853448

ATTCCCAAAACTTTACGGCAATAGATTTAACTTTTGAACCAAGACTCCAAGATTATTTAAATCATTTACA

ACAATTAAACAAAGCGCTAACTCTTAGCTTTCCTTGCCTAATTACCCTGAATATTGTCCAAATTTATACG

GATAGGCAAACGTGGGTTAAATATCGCGGAATAGCTTATATAACAGCTTTTTCTATAGCTGCTTTTATAA

CTCCACCAGATGTTTTAAGCCAAACTTTAATAGGATTACCTTT

>NODE_790_length_252_cov_59.069264

ATTCCAGTATGTCTGCTCAAAGATTCTAGGGCGGTAGCTCACGGATCTCTGGTCTGTGTTGCCACAAAAC

CAGAGGAAGCGCTGCGCCCACTCTCATCGAGCGAAGTCGGCCAAGGCTAGCGAGAGAGCCTCAAAACGAG

TGCTCCCCCTAAAGGTCGCCAAGGCCGGGGGTTTACTGTTCCACGGACGTAACGAGCGGACGGCACTTTT

TATTTCGGAGGCCAGGACAAGCACGCGCGCCGCGCAGGAGGC

>NODE_791_length_252_cov_9.277056

GAGGGGGGTACCGATGGGGCTACTAGGGGTCAAGCCATAAAATCAGTTCCTTTTCGGGTCTGCTTTGGGT

GATGTATTGTCAGGTCAAAAGTCTCGGAAGCAGACAGGACAGTAGTAGTATGGTGTGTTTCGTGTTTGTT

AGGTGTTTGTGGCAACCTCGGGAATATAAGGTGTTTTTATCAGTCTGATGGTAAATTGAACGTAAGGTGA

GATTGGGACGTTTTTTCGGCAGCGTCCCAAAGGTCCCTTCCG

>NODE_792_length_252_cov_8.047619

TATTCGATCAGACCGATACCTCCTATATACCTCGGGATGTATTAAGGCTCTGCGTTTGGACGTTTGCTTG

AGTTGGGCGCTCGTACGCCAACGAGCGTTCCTTTGGGAATGCTCCCGTCAATTGTTTTTTTCCCCCTCAA

GACTTTGGACACCGAAACATCGTGTGCTAGAAAGGCATTTCCATCTTGAGTGTTCATATGCTTGTTGTCC

TGCTGTTGTCGTAGCATGCCTGTACGTAGTTCTGTGTATGTA

>NODE_793_length_252_cov_7.926407

GTCACGTACTGAGAAAGTACCAAAACCAACTAAAGCAACTTGCTCACCACTTTGTAATTCTTCAGTTACA

GCGCCAATAAATGAATCTAATGCGCGACCTGCAGCAGCTTTTGAAATGTCAGCGCCTTCAGAAATTTTCT

CGATAAGTTGAGATTTATTCACAGTGTATACCCCTTCAATTGTTATTATTACAACGCTATTTACTTTTCT

TTAATAAAAGTTTTAATAAAAGTTCGTCCTAAAAAAATAAAG

>NODE_794_length_252_cov_5.987013

CTTGGCTCCCGAGATTTCCTGTCTCCATCTCCTCCGCATGCAGTACACGCTCGTTTGAGCGCTCTACAGG

CCGAGCTTCGTGCGGCGCCAGTGGCGGCGCTTGGCGTTGTAGCGGATCTTGTTGTCCGTGCGCAGGCGGA

TCCAGTGAGGGATGGGCCTGTTCTGCTTCATCTTCTTGCCAAGGGTCACCTTGGTGCGGAAGGTCTTGTG

GGAAGGCATGGTCCTCGTGCAGACAGTCAACACCCGGAGATT

>NODE_795_length_252_cov_4.000000

GGATGGGGGGCTACGGTTGGAGCGTTGACGGTGGGTGCAATGCTTTTCTTCTGTTCTGTGCTATCCTGTG

CGATGCAATGCAATCGCTATGGTTGTTGTTCCATGATTCTGCTGCGAGTACTGGAGCAAGGACAGAAGCC

CTTGGGGACTGAGGGGCGGCGATATGGTTCCATCATTTTACTGTGCTGAATACGAGGAGAGAGGTCAGAG

GACGCTGAGGAGAATATTGCTGGCGGGGGGGGCAGGTGAAAT

>NODE_796_length_252_cov_3.878788

ACAGGTGCCGCAGGTCGCTGAGAGCTCCCAGTTCCTTGGGTATCGAACCTGTCAGTTTGTTCATGCACAG

CCCAAGAGTCCTCAGCTCGGTGAGGGCTCCCAGCTCCTTGGGAATAGATCCTGTCAGTTTGTTCATGTAC

AGCGCAAGAGTCCTCAGCTCGGAGAGGGCTCCGAGCTCCTTGGGTATCGGCCCATCTAACTTGTTGTCGG

CGAGGTTGAGGAACTCGAGGGCGCCAAGCTGTCCGAGCTGCG

>NODE_797_length_252_cov_3.541126

CCCCATGCGGGGGGATCGGAGATGAGGGGTGTATTATGGTGCTTTAATGGTGGGGGAGCGATCCTGGATT

TTCGCTCGGCCGTTTTAGGCTGGGGAGCAAGCATCCCCTACACCTAATATTTTAGCGTTCCTTTCTTGTT

GTTGTCGTTGTTGTTATTGATGCGTTGTGCCACGCTTCCCTTTGTTCGTGTCGCGGACGGTATCTATCCC

ACTCATGGCGCGGTATCTAATCAAGACGGGGGCCGTTCGAGC

>NODE_798_length_252_cov_2.415584

TCTAAACATCTAACCTATCACGTGGTCGTCTACCACGAATCTCAAAAGAAAATTTATTTTAAGTTCCAAT

TCCTACTTAATATGCTTTCAGTAGTTATTCAGTTTATGCGTAGCTTCCCAACCATGCATTTGGTAATACA

ATTGGTAAACAAGAGGCATATCTTCAGCGATCCTTGCGTACTAACTTCAGATACTCTCAATTTTCTTACA

CTCACCGCAGATAGGGACCGAACTGTTTCACAACGTTCTAAA

>NODE_799_length_251_cov_9.208696

GTTCCTTGGCACCCGATTATTTCATTTTTTTTTTGAACCGATGGATTGATTAAGGCCTAGCCACGTAATA

ACCAAGAGTAGTGCCGCCCCAAGTACTCTCCCATCGCATTCTACTGCTGCCGCCAACCGCTGCCGTAGCT

AGGCAGCTCACTCCCATTACGTCGCAGCACCTACGAGAACTTGGTGAACCCAATGTCCTTAGCGTATACG

CGGAAGCACTGGCGGCACATGTTGAGTCCGTACTTGCGGAT

>NODE_800_length_251_cov_7.878261

GCGGGAGTGGCTCATGAGCGACTTGCGGAGGGTGATGCAACGCTTCTTGCAGCCAATGACGCAGCCCTTG

AGCATAAGCCAGTCCTCGTTGATCTCGCCGTAGTGGGGGAAGCCTCCGAGGGGGGTGATGTGCTTGACGG

TCAAGTCGGCTTCCGTGGAAGCGGACCCGACGTCTCCCTTCTTACCGATGCGGTAGATCTTCTTGTTCAT

CTCAGTACGGTGGTGGTATCCGTGCTGACCGGCGCGAGCGA

>NODE_801_length_251_cov_4.656522

CCTCGTGGGCGTTTTTTTCTACCGCATCAACCCCTTCACCAACCATGGCTTTTTGCTGCCAGTGCGTCCC

TAACTCTGAGATCGGCGTGGTCGAGCGGCTCGGAAAGTACACATCGCTCGCTCAGCCGGGCTTCAACTGG

ATTTGCTGGCCCTTTGACCAGATTGTCGCGAGATTGTCAACGCGCGTCCAGCAGCTGGATGTAAGGATGG

AGACTAAGACGAAGGACAACGTGTTCGTCACGGCGGTAGTG

>NODE_802_length_251_cov_3.839130

TGGAAAACGAAGTCTGGGGCGGGTGCGTATCGTTTGCCGGTTGTCGTCTCCGCAGGAACTGCAGCAGCTG

CCACCACGGTCGCGTCGTACTCGTTGAGCAGCATTCGCTCTTTTGAGGCTTCAAAGTTCTTCGCGTAAGA

CACGCTTTGGACGTCCACCTGGCGGTGTCCCCAGGATTCGCTGAAGGTGGCCGCATCCGGGTACTGAAGT

GTTGAGATACCTTCATCGCTTTCCAGCTCACGAGCATCAGC

>NODE_803_length_251_cov_3.113043

GGAATCTCACCAGTCCATCGTCTTGGCGTGTCTCTCTCAAACTGGAGCAATATGTTTTTGTTTTGTAGTC

CGCCGCTTTTCGTTTGGTACCTGAGTGAGTGGGATACGTACACATAATAAGCTGATACAAGTATAATCAG

GGCAATTTGTGAAGGACAGGTTTTGGGAAAGGTTCGTAGAATGATCGTTGCATATGTACCCATGGAAGAT

ACACCAAAAAAATAAAAAAACAATTAAATCTAGCCAACCAA

>NODE_804_length_251_cov_2.791304

TTTGTGGTTTGGCTTCCGCTTCCGGTATTAGTTGCAAGCGGATTATCGGTTTTACGTCCGGTTTGATTGC

CGGTTCGGTGGCCGGTTTGATTGCCGGTTTGGTTTCGGGTTTGGTTTCCGGTTTTATTTCCGCTGTTATC

TCCGGTTTGATCTCCGGTTTTATTTCCGGTTTGGTTTCCGGTTTGGCGACGACGGGTCTCGGTAGTAGAG

GGGGGTCTCTGTCCTCGAGCTGGCTTCCGTGAAAATTGGCC

>NODE_805_length_251_cov_2.669565

CATATTTTTTTTAATGATTCTTGTTTTCGTTGGTTGGTTGGCACCGGAGCCGTTGGATGGGGAGGAGGAC

CCGTGGCCGCGAGAAAGGGCGCACGGATTTGTAAGGAGTGCTGAAAACCCGAAAGAACGCTCGGGAGAGA

CTCGGCGGTGATCGCCGTTACCCCGACTTGTTGCGGACTGCGGACAAGCACGATTTAGAGGTGCACATAA

TTTACTATTATTCAAGAGTGAAAAAAAAAAAAATTTGGAAT

>NODE_806_length_251_cov_2.547826

GAGAGCAGCTGCATTCCCTGACCACTGGCATGGAGAGGAGGACAAAGGCCAACGAGGAGAGGAAGGTGAA

GGCGCAGCAACGAAAACGGGGAGACACGCGAGACATGATGGCAGAGAGCAAGGCGAAGGACGACCGGATC

AAGGGTCTAGAGTCCACCCTGATGCAAGAGAGGGCAAGACTTGAGGAAGAGCTCCGAGCGGCGCGTTCGG

ACTACGAAGAGGCCGCGAAGGAGAGCAGGCGCAAGGCTGGC

>NODE_807_length_251_cov_2.495652

ACTGCCTTTCCGCAGGAAACCGCATCCGCAGAGCTGTCACCACCGTCGCACCATCCAAACTGACCACGAT

CTCGTTGTCTTTCTTGGGAAAGTCCAATGCGAATCTCGTGGGAATTTCGACTTGAGACACGTCGTCATCA

CTGTCGTCGTTAAAAAATCTCAGATCGACATTAATCACAGGGGCAATTCTCGCGATGACGTCGTCCGTTC

TCTCCTCAAGGATCACGATCTCGCTGATGCTGAACCACTGG

>NODE_808_length_250_cov_154.655022

CCTTGGCACCCGAAATTCCATGGCTCCCCAAGTCGGACTTGAACCAACGACTTTATGGTTAACAGCCACA

TGCTCTACCAACTGAGCTATTGAAGAAGGCCGGAGAGGGGCTTGAACCCTCATTTATGGGTTTGCAACCC

ACCGCATTAGACCTTTATACTATCTAGCCAAGGCGGGCGAGGGGACTTGAACCCCCGTCTTTCAGAGCCA

CAACCTGACACTCTAACCAACTGAGTTACGCTCGCCATTT

>NODE_809_length_250_cov_19.650655

TTCCAAAAAGATATGCCTCGTGGGTCGCTTCAGACTCGGCACTGATCTCCGGGGCGAACCCCTCGCCGCG

CGCTAAATACTGGTCAGATAAGACAGGATAATCCACCTTCCCCCCCTTTTTTTTTTGTTTTCTTTTCCCA

CCCGCGAGAGGGAAAAGTGATGATCGCCGAGAATCGAACTCATCATTTATTAAATATATACGTTGGAAAA

GAAGGGGATAACCTGCTAGCCTACCCAGTCAGCGCTGACT

>NODE_810_length_250_cov_7.812227

TGCTATGGCTGTGTCATAGTGTCCCTTTTGCTGTGTGCGTTCGAACGCGGAGATTCTGACTATTTCCAAT

CGAGGATTGCATTTCAATATTTGTCGTTGAAGCTTTTGTCGGCGGTGAACGACCGAGGCAGGCAATGCGA

TCGCATCTCTACACCAATCGAGGACCTATTTTGACGAGTTCCAACCAGCTACATGATCGTCTGGGACCGT

AGCCGAGATTTGTGTCTCGAAGTTATGTGGGTCGAGGGCT

>NODE_811_length_250_cov_7.510917

GAACTTGACCATCGCCGGTGCCTGCGCCATGAACTCCTTGATCTCGGGCTGCGGGGTACCTTTCTCCTTC

AGCTTAGCACGTACACTCCGCACGTAGTCCTTAAGGTACTCCTTGAGCTCGGTTTTGCTGCCGAATGGTA

CTTCGGTGTACTTGAACGGGTCGATCACGTTGTTCACCTTCTCTACGGAGTCGTCGACGGCCTCGTCATC

ACCGCCGAAGGCGTTTCCGCACCCGATGTCGACGTTGTCG

>NODE_812_length_250_cov_6.209607

GCCGCCCCTTAATGTTTGTGTCCGTGAGAGGTTGCCTTTGGGGTGACGAGTCGCGAGCTCTTTGTGAGAG

TTGGCGGCGAGGAAGGAGCCTTGAGGGGTCTCGGTCGGTCGTTTGAGAGAGAGAGATGAAAGAGTTGGTC

GGGTAGAGGTGGGGTTGGCGTCGCAAGCAGCAGTCGTCTGCTTGAGGCCTGGGTTTGGTCACCCGGGTTG

GAACGCATCAGTCCGCCTCGTCTAATACAAAAACACTCGA

>NODE_813_length_250_cov_4.388646

GTATCCTTATATAACTTTAATGCAAATGAAATATAGGATAGAAAAGTTTATAAAAAGTTCAACAATCCCA

TTTACTATTTTTAAATACGCTGGCTTTTTTCAAGGACTTATCTATCAATATGCAATACCTATTTTAGAAC

AAAAATCTATTTTAGTTACATTAGAATCACCAACAATTGCTTATATAGATACTCAAGATGCTGCATTTTT

TTGTATAAAAAGTTTATCCATTAAAGAAACAGAAAATAAA

>NODE_814_length_250_cov_2.441048

CAGGGTAGGGTTCTTTCGGTTGGAGCTCTCGTTGAGGATTATGTCAATCCCGCCCCTGGCCGCCATCCCG

ATCTTGCAGCACTCATTGCCGAAGTCGATCCCGATCACCGACATGGTGCTCGCTTATTTGCTGCTCCTCC

CTATATAACCTCCCTTTCGCTCTAATCCTGTGTCGGCGTCTGTTTTTACGCCAACTCGCCGTTACGTTTC

AAAAGGAGTAGTTCCCAAAAAAGTTGCCTTGGGGAGAGGA

>NODE_815_length_250_cov_2.318777

GGTATGAACGACGCCGGGGGGGGTGGCGCCTGCCATCATTACCCGACGCTTGAAGCGTGGGTGGTCGGTG

AAGGCCTGTGATTGGTCGGGCCACATTTTCGGCGGTAGCGCCAGTTCTAGGTCCTTGAACAAAAAACTGG

CGATGTCACCCTCTTCCACTGCTCATGGCGGTCAGTATGATACCAAATATGTGACATTGGAGTTGCTATT

GCATGCGGCATTCGTGACTGACGAAATTTGCGGTGATATG

>NODE_816_length_249_cov_81.877193

ATCGTTTGTTTGAGCGCTCGACCACCCATGCGTCTCATTCCAGAAAAGGTTGGCCACAGTGATCTCTCAA

AGCTTCACTCGCAAGAGCTGAAGCTTTCGAAGGGCCGCTAAAATGACTCCCAGCGTCACGGTCGGAAAAT

ATTTTTATATATATATATATTTTTTTTGTCCATCCCATTTATTATCTTCGCCCTTCCTTTCTCTCTCCTT

CCTAGTCGTAACTCAGATCCGGGGTCACAAAGTAGGCTC

>NODE_817_length_249_cov_5.065789

TGGAGTTCCTTGGCTCCGAGATTTCCAGGGTTATTTTTGCGACGTCTTGATAAAATTTGTATAACACGTT

CAATTTCATTTTGCCTACCAACAACAGGATCTAATTTTGCTCGTTCAGCTGCTTCTGTTAAATCTGTTGT

ATACTCTAATAAGTTTGGTGCAGTTAAAGAAGCTCTATCTAAGTCATCAAAAAGAAATAAATAAACAACT

AAATAAATAAAGCCTTCAAATGAAATAAAAAAACAACTA

>NODE_818_length_249_cov_4.122807

GGAAAAAGTAGGTTTCGACGTCAATAGTATCTGAGGCGTTGTTTGGGTAGAGATGGGTCATCTCGTAGTA

GGCACGTGCATCAAGGCGCGAAGGCGATAGTACAGTTAGTTGTCAAACGTTATATGGTTGTGCATGGTCC

CCTCCGGTATGTGTGCGAACATTGAGAGGAAGAACATTTGATTGGTAGGGCCTTGCCGTAGAAAGCAGCG

AAACCTTTCGAGGAGGCGGCGGACCACGGTAGGATCAGC

>NODE_819_length_249_cov_3.793860

GAAGATCGAGGAGATCGACTAAGCTAGGGTGCTTGTGTTGGATATCGTATCGGAATTTTCGATATATCGT

TTCGGAGGTTTTGATTTTGATTTGTCGAACTAGATCTCCTTTCGGTATCCAACTCTGACTGAATAAAGGC

CTGGTCAACGTCCAAGTGCCTCAAGTCCCAGTCAAGTTTGCATGCCATTGCTGCGACAAGTCTGTTAGAC

GTGGTTGGTGCAGCGGGGGGCAAAAGTGTCAAAGTAATC

>NODE_820_length_249_cov_3.758772

TTGGCACCCAGAATTCCACATTACCTTGACGATCTGTAAGAGTCACGATTGTGTTGTTGAAAGATGCATG

GATATGAGCCATGCCATCAGCAACTTGTTTTTTTACGCGTTTACGCGTACGTACGGGTGTTTTAGCCATT

GTCTAGTTCCTCTTACTTCTTAATTGGCTTACGAGGACCTTTACGTGTGCGCGCATTAGTTTTAGTGCGT

TGACCACGTAGAGGAAGACTGCGACGATGGCGAATACCA

>NODE_821_length_249_cov_3.464912

GATGATTGAAACAGGAATATTCGCCGCAGGCTGCTTCTGGGGCGTTGAGCTCGCGTACCAGCGTATCCCT

GGCATCCTGGAGACCGAAGTCGGCTACATCGACGGACACAAGGTGGACCCCACCTACAAGGAGGTCTGCA

CGGGAACAACGGGGCACACTGAAGCTCTCAGGGTTCGATTCGATCCCCGCGTTGTGAAGTACAACGAGCT

GTTGACCGTGCTTTGGGATCGCATCGACCCCACCACGCT

>NODE_822_length_248_cov_131.991189

TTTGTCGTCTCTATATTATTGTACCTGCTTGAAACTTGAGGCTGTGGTAGCTTAATCGCTTTGACCACGA

CTCTGAAATTGGCAGGAACATTAAACAAGTAGTGGACGCCTCCCTCCCCGGTTTTCACTGGGCGAGCGGC

AGGTACGCGAAAAATCATTGCTTACCGTGCGGGCCTCTGCCTCTATTTTTAACCCCAAGGCTGGCTCCGC

CCCGAGGCGATGATATATTTTGGAATTCCGGGTGCCAA

>NODE_823_length_248_cov_6.295154

TTTTACACCCTTATCAAAAGGTGCATCCATGGTTCTAGTTTTAAGCGGTTTAACTATTATCTTTTTTATG

CTTAGTCCGAATATACTTTATCTGTTGACATACAAGGTAGGACTAGGTCTTATGTTTTAATAATTAGCTA

ATGTCTTTTACTACACATTCTAAACCTTTATCGCAATTATGGTCTCCATCGGTTATTAGCTCGTCATTGA

GTGGTTTCTCTTCTAGGTGGTTATTTTCCACCAACCAT

>NODE_824_length_248_cov_6.132159

TTGCGGGGGGAAGGGCGGGTAAATGGCGTTTTTTACATGGACTGAGGCGTGTCGTGTCCGTGCCGGCAGC

GGCGGAAGGGGATAAAAGGCGTGTGGTTTTCAGGATGACGTGCTTTTGAGGGGTACACGATTATGTGTAT

ACCCATTTTTTATCTCTTTCTGGTAGTCTGTCGGGCAGCGTGAGGAGGGACGCAACAAAATTGGAGAGGC

AACGCGAAGCTTGAAAAGAAAACACAAAAAAAGGTCGT

>NODE_825_length_248_cov_3.889868

TCCCAGGACCGGACTAGTGGGCTGCTGCCGTAGCAATCATGGTTTCTCTGAAGCTTCAAAAGCGGCTTGC

CGCCTCCGTACTCAAGTGCGGAAAGGGGAAGATCTGGCTCGACCCCAACGAGGTCAACGAGATCTCCCTC

GCCAACTCTCGCCAGAACATCCGCAAGCTGGTGAAGGATGGCCTGGTCATCCGGTTGCCGCAGGTCGTGC

ACTCCCGCGCCCGCGTGAACTTGCGCCACGAGGCCAAG

>NODE_826_length_248_cov_2.546256

AACAAAGACCATTCGATCCTATATAATTATGCATGTCCTGTCATAGCATAAAACGCTAAAACGAAAGAAA

TGCTTTTTAGGTATTATGCTCCATCACCAATACTTCTGGTTCCAACGTAAATATATATTATACGTGAATA

TATACTTCGAATATATCCGAAGACCATAGCCCACTACTGTAAGATCATTCAACATAGCCTCCCCAGAACG

GCCTCCCTGCGTCCAAAACTCTTATTTTTTCACAGTAA

>NODE_827_length_248_cov_2.361233

AGTTTGGGGGGAGGTGGGGAAATGAGCAGGGCGCCGATGCCGGCGCTGTACCGGAGTACAGTACGCGTAG

TCGGAAGTCGAAGAATGTCTGATGTAGTGTTCGTTTTGCACGGTTGTTGTTGACATTGTTGCCTATTTTT

GTCGGGCTTTGTCTTTTAAGTCTGTTTCATTTTTTTTTTCCGGTTGCGCGTGATTGCTCTGCAGGTACCT

ACTGTAAGTAAACGTTGCCTCTGCCAAGTTTGTGTATG

>NODE_828_length_248_cov_2.154185

TTCGCTTTGAATGACTTATACATACATCTGGGTGTGTATGGCCACATGCCACAAGCGGTAGATGGGTCGT

GTTCTCTAGATGGATACCCGGAGTAGAAGCGGGCTTAGACTGTGCTTCTTCCCAAGTACATCCGGAATTG

AGGCGCACGAGTTGTACCCACTTTACGCGTAGGCATTGGAGAATGGAGGGAATATGTTCGTGCTTCATGC

ATAATATTGATACTGGACACGCCGTGTTAGGAACGGCT

>NODE_829_length_248_cov_2.039648

TTTGGTATAGAATTTTGCGTGTCGTGGACGAGAGTGGGAAAGGAAGGAGGAGCCTGAAACATGAGCTCTA

AGCCAACTGTCGTCTAGGTATGTACCAGAGCAAGGAATCAGACCTGTATCGGGCTAGAACTCTAGAGGAC

GAGGCGGCACACACTATTATTCCAGTACTTGTACCTTGGTACGGCGGAGTAAGAGTTAGTAGGGGCGATT

TGGTACGGGACGGAGCAGAGCTGGTGTCGCTATCGAGC

>NODE_830_length_247_cov_45.407080

GGTACTGGTTCACTATCGGTCTCTCAGGAGTATTTAGCCTTATGGGATGGTCCCCACAGATTCATACAGG

GTTTCACGTGCCCCGCACTACTCAGGATACCACTATCTGTAACTGTCTTTACTTATACCGGGCTATCACC

GTCTACGGCTTGTCTTTCCAAACAATTCTAATTCATCCTGCTTCTAATATCGTGGTCCTACAACCCCAGC

ATTGCCGTAACAATACTGGTTTGGGCTAATCCGCGTT

>NODE_831_length_247_cov_8.433628

CGAAGTCTTTGAACCACTTCATCTGGGCGATGTCGGTCATCTCGGTCATGAACGCCTGCAGCTTTCAACC

ATTGCTGGGGCCTAGTTGTAGGTCTTCTTGGCGTCCGTCTGGCACGCCTGCAATGCGTCGAACACGAACT

GGCAAGATGTCACGTCGTTGTTGTTGTTCTGCAGGCACGACCCGAAGTTCTGCTTGTCCATGTCGCAGGG

GTCAACTTCAGGCTGCTGGTACGACTGCGTCGCCGCT

>NODE_832_length_247_cov_3.017699

AAGACCGTAGAAAGCGCAAGTGGGCTCGACATCTCTTACAAGGTGTACTCGTTCGACGATGATGTGGAGC

AACGCGGTTGGTTCTTGATCTTCTGCTTGTTCTGGTCTCTGAACTTCATCACGGCCATGGGGCAGATCGT

GATCGCTATGTCGGTCGCCCAGTGGTACTTCGCGAGGAACAAAAACTTGGTGGGATCGTTGACTGTGGTG

TCCTCGGTAGGCAGGACCTTCATCTTCCACGCCGGAA

>NODE_833_length_247_cov_2.646018

GACGAGCAATACTTCAGCTTCACGCCCCCGGCAGAGATTTTGGGTTATGAGAACTACTACAAGACCGTGC

GTCAGCTCGGCTTGCCGGAGACCTCCGACATCTCCGTGCCGAACCAGAACATGCTCCTGCTCCACAGTGC

CACCGAGCTCGCTCTCAAGCCGGAGCTTGTCCAATCTTTCGTGGACAAAGCGTACTGCAAGGGGTGGGGC

ACGATCTACGTTACCGACGACCTCTTTGACGCGGGAA

>NODE_834_length_247_cov_2.367257

TCTGTACGAACGTGCAAATGTGCTTCAAAATCAACGGGAGGAGCAGGTGATCGCTGGTACTACTATTACT

GTTGAGGTACGCACGTGGGTTTCAATCCACACGTCTCATCGTAAAATAGCACCTCGCAGGCAGTGTACGG

TATGATATCGTGATATCAAGAGGTACAAGAGAGGACAGGAAAGTAGCTAATGGGCTACGAAATGCTCTAC

TTCGTAGGTACTTGGGCTACATACACATACGCGAAGG

>NODE_835_length_247_cov_2.327434

GTCCATCTTCTCTCAAGAGAAATATCGGGCCAGCCAAGATAATCGAGCTTTGCCAGACACGGCCTTCCCA

CGTTGGGGAACCGATGCTTCTACCTAACAACATGAGTCCTACCAGTGCTCCACCATGGTGAGACCGTGGA

ACACCGTGAGCGGTGTGACTAACCGCGACGAAGCGCGTTCCTACTTACCAACTGCGTGAACTTCATTAAA

GAGTAACGGCACGCGGGCCACGTCCCGCGTGCCAAGG

>NODE_836_length_246_cov_25.484444

ATGCTACGATAGAGGTGCACCTGTATACAGGCCATTTTCTGAGAGTGAGTGGTCAATATTGAGCTCTATA

CCTCACCCAAAACCCAGATTGAGGGGGACAACCAAAGTTCTGTATACCCGATGAAAAATCCCATGGCCTG

TTTATTAAACGAAAACTAATGAAAAGAAATATTTAGATTCCGACGGTTTTGGGTTAACCCGTTCAGAACT

GAAAAAAAAAGTCCTACACTAATTCCAAGTAGTTTG

>NODE_837_length_246_cov_12.320000

TTACCGTCCCGTCCCTTCCCCCCTCAGTTGTCGGTCGGGAAACATGATATGTTCATGCGATTTTTTGTCT

TGCCATTTTGTTTTGTTGGGGATTAGGTTTCAGTGTTCCTGGCGTTGCCTGTTATGTTAACCAGCGCTGC

AGGAAGAGGACCTGTTGTTTTCCGCTATTTTCTGTGTTGGGGTCTCGTCAAACCCCGGCGCGAACGTTGT

TTATTCGGAGGCATTCTGAGTGTACGGGGGGGCGGG

>NODE_838_length_246_cov_7.897778

ATTGTTTTTTTTCAGTGTTCAACCGTTTTATTGTCTCCTGCAGCAGCCACACGCCAGCCCACAAGCCCGA

TTCATGAGCCAGCAACTCAGATTCTACAATCCTTCTTGTTCATCCGAAAATGTTAACCGGATCACCATCC

ATAACCACAACGCGTTACGTATTAACAGCGCCATCGGATTATCGCCAAACAAAGCCACAACACGCTACTG

ATATATTAGCGTCAACACGCTACTGATATCGCATTG

>NODE_839_length_246_cov_5.137778

GTCCGCGCCAATGTGGAAGTGGCTGCGGAATTCCTCCCACGACATGTGCGAGTAGGAGTTGTGGGCGAGG

GTGTAGCCCGCCTTTTCCGCGTCGTGAAGAGCGATCTTGTCGTCCGCCTTCGCGTAGTTCTTGAGCGCGT

GAAGCCCATCCTCCATGGACGACCATCCCGTCAGACCCTCCCCGAACGTGTACGTCCAGCGGAAGAAACC

CGTCAGGTAGTGCGACAGGGGGGTGCAGAAGCCCCG

>NODE_840_length_246_cov_4.364444

GGCGTCAAATTGAATCAGCCCGACAAAATATTAGACGTAAAGTTAAAAGGGAGGGTAAACTTGAAATTCT

TATTTTCCCTGACATTGCTATAAGCCGTAAGGCTTCCGCAGCCCGTATGGGAAAAGGAAAAGGAAAAGTT

GATCATTGGATTGCCTCGGTATCTGCTGGACAAACTTTGTTTTTACTTTATGGTATAGATGCTGAACAAG

GTATACCAGCTTTACACTCGGGAGCTAATAAGCTCC

>NODE_841_length_246_cov_3.128889

GACCACCGTTCCACGACCAAAAAACCGGGGTCCGATACGATTGTAACAACCCTGATCACGCGGGCTGGTT

GCGGAAGGAAAGCGCCTGGCTCAAGGACTGGCGACCGAGGTTCTTCGTGCTCAAGGGGAGCAACTTATTC

TTCGCGAAGTCCGAGTACGAAACGCCACACGGTAGGATTGACCTTAGTTCGTGCATGACGGTCAAGTCGG

CGGAGCAGAAGACCAGCAAGCGAAACGCCCTCGAGG

>NODE_842_length_246_cov_2.546667

CCTTTAGCTTAAGGAAGTCGGGTGTGTTCAGAATAAAAGCAGACGTTCATCGCCACACCTGACACTCCTC

CATTCCCTTCCCAACGCTCACCCGTTCACGACCACCTCGTCCGTCTCCAATTGCTTCATAATCCAGCACA

CCCCCATGCTCCACGCGCCATTTCGATCCGCTCGGGGTTGGACAGGAAGGTGTTGGTAATCCGCTCGATC

TCGGGGGCCATTGTGGCTGAGAAGAACAGCGTCTGG

>NODE_843_length_246_cov_2.475556

GCAGGTGGGTTACGTTGATGGGCAGTATGTGCGAGCACCGTACATGGTTGATTCGACGTTACCGCTTAGC

GTTGCAGCCTTACACAATGACGAGTTTTCAACATACGCTGTGCCTCCTCCCTTGAAGGAGTGGAGACTGT

AAGACTGTAGTCCGTATGGGCTCGCGCGTTCGGGTAATAACCATTCGATGATTACTAAATTAGCTGATAT

GGCCGATGAAGAGTTATTCGCAGAGTGGACTGAGCT

>NODE_844_length_246_cov_2.128889

ATGTATTTTGCTTGCTCTCTAAGGACCACCACCAGCGCAATCAATGCTCGTGTCAGTCAGACCGACGATT

CTCGAAAAACAGCCTGCCCACATTCTAGGAGCGGCAGCAGCACAAGCAGGTACCCCCGTGCAGAGGGTGC

AGCCGCCGTAAACAAAATAGACACCGGTTAAAGTATGTTACACTTTTACAGCCGTTGCTCGTGGCACTCA

AAATAAGGGAAGGGGACACAACCGACGATCATTGGG

>NODE_845_length_246_cov_1.568889

GGATCCGATCGAAATAAAAGTGTGAGACACAGCAACCAACCACACCGTAACTTACCTCGCATCCATATCT

TATCCTATCCCATCTCGTCTTGTCCGGGTGTGCGCGTTTTTCGTTCGGCGTGCCATAAAGCCTTAAAACA

GAACAGAGGCTGTGGTGTTTATTTAGGCGCTACCGGTACAGCAGTAGTAGGTACCGTTGCAGAGACTAAG

ACTAAGGGGTTGCGCTACCACTGCTGTGTATAATAT

>NODE_846_length_245_cov_372.151786

ACCCCCCAGGGTGATTACCGTCCCCGATACCGAGCGCCTATCGTTCGCCTTATCAGCATAGTCGGCATCA

CTATACGCATCCAAGCCCAGTCCCGAACCCCGCACAAAAGTCAATCCCAAAAACCTGGTCCCGTGAAGGT

ACTCCATAATCTTCAAAACTGCTCTCCAGTGCCTCTCAGTAGGGTTGTGAGAGTGACGCGCCACAGCACG

CACAGGCGCGTGCGGCCCAGGACATCTAAGGGCAT

>NODE_847_length_245_cov_26.196429

AGTTGATTATTTATACAAGTCTTACTGTTTATTTCTATGGCTTCACGCCAACCACTGAACCATTGATATT

GTACTTTTCTACTTGTCTACTCTAGTACATTGGTAAGGGCATTGTTCTAAGAACGTTAGGATATACCTAT

TTGGTTATTACCTATATGAGCCCTGTATTGGTGGTCTCGTTTTTGCCTTCTCATCATAAATGCACACTCC

TTTATTACCATTGATATTGGAACACACTTACCTTC

>NODE_848_length_245_cov_15.531250

TGTCATGTTGTGGTTGGTGCGTGGGCTGCGGTGTTGGCGTTGCCTTGGGCGGGGGAGGAGGCGGACGGAA

GTTTGTCCAGGAGGTGCGGTAGATCGTTTTAAGGTTTTTCGTTTTAGGGGTGAGACTTACAGTCAGTACT

GATACATACATACATATCATTGGTACTGCTGTGATGCGGACGGCAAGAACGTCGACGAACTTGAGAAATA

AAACAAACGAAATCATCGAGAAAAAAAATTTTAGT

>NODE_849_length_245_cov_7.491071

CCTTGGCTCACGAGAATTCCAGCTGCCCCGGTTCATTGTCGTGGATGTCTTGTCGCTGTTTTAGGGTTTC

TCTCGATTTTTTTGGGAACGAGAAGATATGTCGGTGATCAGCTGGAGCGGTCAACGACTGCGCGGATAAC

CCGTCCTGTAATGCACTGCCTCCGTCTCTCTCTCCGGGAATATATCTCTGTCTAACTCGTGTGAGGTCTT

CTCAAGCTTTTCGCTGGGATCAACGTCAACGTCAA

>NODE_850_length_245_cov_5.196429

GAAATCATATTACTAACAAAGTTACCAGAACTGTACTCAATGTATAGTCCAATTGTAGATATTTTACCAA

TTGTTCCAGTTCTTTTTTTATTACTTGCCTTCCTTTGGCAAGCTTCAGTTGGTTTTAGATAAACAACCTA

TAAATAGTGGTTTTGTAGAAATATATTAGTATCTAGAATATGGGGCTAATATCTTCTGATTATTTAAGCT

TATTAAACGCAATATCCTTAATCACTTTAATAATT

>NODE_851_length_245_cov_4.616071

CATTCTGCTGGAAAAAATATCTAGGAGATCTGGCCGCAGTCGGCGACGACCACGGGTGAGCTGGTCTTGC

CGCTGGAGGATCCGACTCCCTCGATGGCCTTGACGACGTCCATGCCCTTGGTCACCTTTCCGAACACCAC

GTGCTTGCCGTCCAACCAAGGGGTCGGCACGGTGCACATGAAGAACTGGGACCCGTTGGTGTTCTTGCCA

GCGTTGGCCATGGAGAGCACGCCAGGCTTGTCGTG

>NODE_852_length_245_cov_3.696429

GCGAGCGACTGACCGCTGCGAACCAGGAAAACACCGAGCCCAGGTTTTCAGGACCGGCCGCGTCATGGGT

CGCCGAAAGAAGAGCACGAAGAAGATCGTGGTGAAGAAGAATTCGAAGCTGGACAAGGTGTTCAAGTGCC

CCTTCTGCAACCACGACAAGGTCGTCGAGTGCACCATGGATCGGAAGGAAGGAACGGCAAGACTGCAATG

CCGCATGTGCGATATCAACTACGAGATGGGTATCA

>NODE_853_length_245_cov_1.433036

CAACCAAAAAAAAGTGTTTCTGAGACTTTTTTCGCTTTAGAGGAGACCTTAACCTCGTCCAGCACCCCTA

CACCGAGAAACAAGTACAAAAGCTAAAGTCAACAAGTACACAATCCATCCCACGGGCTGATGTAGGCACC

TAACGAAAACGATATTATATATACGAGTCGGGTCCTAATACAGCGAGGTGGGGGTATGATAGCTCCTCAG

GGCAGGCGCCCCTGCCCCGTCCGTGATGGCTGTTT

>NODE_854_length_244_cov_18.017937

TCCAAAATGTCCGCGCTGGCTTCAGCTGCTCATGCAATGATGGCTTTTCAGCCATTTTGTAAAAATCAAA

AGACTTTACTGTCACAGAGAGCCGCAGAACCAACGCCCCCCACCAACAAAACCACACCGCCTTCGCAGGC

GATAATGGCGTAGAGGGGGTTATTTAATGTATCCCCGAGTTTGACAACCACCAGGCCACTGGAAAACGTA

TTTCAGACCACGGCGGATAACACTACCGGGGACG

>NODE_855_length_244_cov_6.739910

TGTGTTCCTTGTATTCCTAGAATTCATATGAAATTAGTAAACTAATTCTGCACGACGGTTTTCAGCACGA

GCAGCTTCGTTATGTCCTAGGTTAGGCTATATCTCCCTGTCACGATGACAACAAACTACATGCCGGATAC

AAGCCGATGACGCCCAAAGTAGAAACAGATAAAACATGGTGTTTTCCCATCTTGACGGTGTCCAGGCCTT

GAATCTGTAATCATTGGAGGTTTTACTCCACCCA

>NODE_856_length_244_cov_6.251121

GGCACGATACGCGGCACGATGTCGTTGTTGTTCCTGCACCGGAAGTACTTGTCCTTAAGAGCGGTGCCGT

GGTTGTCCTTGTTGTCGAAAATCTCGGACAGCTGTCCGTCAAACACCCTTGGGCTGCCGATGGTGTAAAT

TCCGGCGATCTTCATGTCGTGTTCGAAGGCCAGTCGGGCTCCGGCGATGGTGGCGAGGGCGCCCCCGAGC

GAGTGTCCGGAGATATACAGCTTCCGGTTTTTGC

>NODE_857_length_244_cov_4.991031

CATTCCCACCTGTGTCGGGTTAGAAGGAGTTCGATTATACTCGAGGCGTGCTTGTTTTTCGGTCAACAAT

GCATATGTTTTCACAGCACAGCCGTAGTACCATAGGCGCTCTTTGCGGAAGTGACCTGTAGAGATCTCCA

GCAAGCGATACGAAAGGGAAGGGAGGGGCAACATTTGTTTCAGCTCCTCTATCGCGCGCCCGATACACTA

ATTTGCTTTTCTTCGTCGATTTGTTTTGCTTGTC

>NODE_858_length_244_cov_4.780269

CAAGGCCTACAAGCGCCGTCGATCGAAAGGAGGCCACCGTGAAGCACATCACATCACTTCTTCGCATCCG

CCGCTTCTTTCGCCGCGGTTTCTTTTTCCGCCGCCTTGCGAGCCTGCTTTTCCCTCATCGCCTGAGCGTC

TCGTTCTGCGCGCGCACGTGGGTCCCCTTCCTTGCTATTTTTTCCGTTCTTCTTCGCGTCTTTCGCCTGG

GCCTTCAGCCTGTCCCGGTCTCGCTGGTTTCCGC

>NODE_859_length_244_cov_4.556054

TTGGCACCGAGAATTCCAGATGCCGCCACCCCACTTCCTCTGGAAGCCCTCGTTGAACTGAGCGTTGAAG

TTGGACTTGAGGGTGGTGAGCTTGCTCTCGTCCTCCTTGTTGACGTTGGTGAGGCATATCACGGCGGCGT

TCTTCTGGTGCACGAGGGTGCCGAGGCGAGCCTTGCCCTTCATGATGCAGTAGGGGATGTCCATCTTGCG

GCAAAGTGCGGGCAGCCACACCACGAGCTCGATG

>NODE_860_length_244_cov_4.237668

GTGACTGGTGTTCCTTGGCACACGTGTATTCGACAGTTTGTGTTTTTTGTTTGCCGATGGGACGAAACAT

ACCGGTCACAGAAAGAATCGCACCCCAACGTCTAACACGCCACCGTCCGTACAAACACCTATGCATCTCC

ACTCTACGCATATGTGCCGTGGAGTTCGACTGATTGCATCTACAGTACGAAGCTTCCCTCTTGCTGCTGT

CACCGGCAGTTATGTACGGCTCCAGTGGGTTTAA

>NODE_861_length_244_cov_4.085202

CGACTGGAATATTCTCTCGGGTATATCTTATACCTGGCATTACCTACTATATATAATAGTTGGTGGATGT

TCTCATACGTACGTATTGGAAGACTAAAGTATGGTATGTAAGGTAAGGTCAGTATGTATGGCTTAGCCGC

GTGAGTGTCTTGTGTGGTACGATATAATTACCCATGTATGGGACGAGAGCGCTTGTCGCATTTTATGGTT

CGGGGACGAGGCAAGCTTGGGAGAGGGAGGGCTG

>NODE_862_length_244_cov_3.650224

TCCTTCCGCCTCCAAAAGGTATTTACCCAACACACCAAAAAAAAAGGGCACGAAAAAACGCCTCCTCGAG

ATACAGCAAACCAGCGAAACCCCGAAAAAAAAAAAAAAGGAAAAAAGACCGATCAGACCCAAGAAAGAAA

GAAGAGAACAGAGAAAACAAAAACCACCACGAGCGGCGGAGGTAGTTAGGAACACAAAGCACGCAGAAGT

AGACGAAGAGGCAACACCAGCGTAGTCATGCATA

>NODE_863_length_244_cov_2.811659

GTCAAGTGTAAAAAACAAAAGGGAGGAGGGACTAGTCGCTCCGTCGCAAGGGGTTTCCACTCCGCGCGGT

GCTCGGGGATCAGGGACCCCCCAAGTTTTTTTTCCAAGGTGAGGGGAAGACTCTACTGTAAGTAACTGAA

GGCGCTGAGAATCGGGGTTCGGGTATTTCTTCAGCGGTGTGTCGTCTTGATCAGGCGTCTTTTTGTGGTG

TTTAGAGGTTGTGTTGTCCGTCTGGTTAAATGGC

>NODE_864_length_244_cov_2.399103

CGTTTGCGGCCTGCCTTCACGCGTGCATGATGCCGTTGTTCATCGTGAACTTCGGCAGCACTCTTGACGA

GCTAGGCGAGCCCCTCGTCGAGGGCGAGGAAGCGTCCATCCTGGAAAGCGTGGAGAGCTTCGTGGTTATC

TTCGGGATCATCGGAGCCGTCGCGGGCTTGGCGGGATTTACCATGGTCTCCCTATGGAGCATCGCGGGCG

AGAGACAGGCGTTGCGAATGCGGCGAGCGTACGT

>NODE_865_length_244_cov_2.026906

GGCTCCCGAGTATTCCATTTTTTTTTCAGCTTTTTTTTCTTTTCTCATTCTCAAAATCTCAAAATCAGTA

TTTGGGTGAGGTACTATTTACTACTGTAGTACTACTGCTGTATTATTGCGTGCTGCCTGCTGCGGCACGA

CGAGAAGCGAAACTATACACAACTGTTGCGTGATGCCCTGCTGCTTGCTGCTCTGTAGCTCTGCTGTCGG

AGTACTCTACAAGGAACGCTGGAGGAAGAGAGGG

>NODE_866_length_244_cov_1.636771

ATTGGCATCCTTCTAAATCAATTGTCCTGGCAATTCCATTTAATAGACATCCTTCTCAACCAATTGTGCT

GGTAATTCCATTTAATTGACATCTTTCTAAATTAATTGCGCGGGCAATTCCATTTAGTTGGCATCTTTCT

ACATAATTGTCCTGGCACTTTCATTTATTTGGCAGGCTTCGAAAACAAATGTCCAGACAATCAGGCATTT

CCATTTAATCGACAATAATATGTTACCGGACAAT

>NODE_867_length_243_cov_880.711712

TGAAGTCGTAACAAGGTAGCCCTAGGGGAACCTGGGGCTGGATCACCTCCTTACGTAAAGAAACACATCA

TTTGTATTTCGGTACATTGGCGCAAGTAATAATGAGTGCCCACACAGATTGTTTGATTAGACTGATTAGA

AAGCAAAGTAGGTTCCGGGTCCCCATCGTCTAGAGGCCTAGGACATCTCCTTTTCACGGAGGGAACAGGG

ATTCGAATTCCCTTGGGGGTACCAGGAAATCTC

>NODE_868_length_243_cov_51.243243

CATGATCAAACTCTTCAATTAAAAATCGTTTGTGTTACGTCACCGATAAACGATGCATAACTGCTCAATG

AATTCTGTCGTGATACCAACCGAAGTTGATATCGCATTACATAGTCGCTACCTATCTCGCTAGAAATAAG

TAACTATATTTATTTGCATGAACATCATTCATTAAGCGTTTTTTTGTTCCGCTTCCCTAAGAAAGAAGAA

CTATGTAAAAACAAGCTAATGTGAGTATTCACA

>NODE_869_length_243_cov_14.869369

CACTGGGCAGAAATCACATTGTGTCAACACGTTTACACGCCATCACAATGCTATGTTTTTATTAAACAGT

TGGATTCCCCTTGTCCGCTTCAGTTCTGAGTTGATTGTTCCTTGTGTAGGGAGAGCCGATGAAGACGTCC

CAACAAATCCAGCCCCCACTAGCGCTAGCACACTCCGAAAAGCACGCAGGACACTTATGGTGACGTTCAT

TGAAGGGCCCCTACACCCAATCCTCAGAGCCAA

>NODE_870_length_243_cov_4.882883

ATGCCGCATTCGAGACCATCATTATTTCCATTACTGTAACAGTAGTAGCCCTAACCTCCGATCCCCAGAG

CGGATATGCTCTGTCTGCTGCCCCACCACCGCATGCGCGTGGAACCACACCCCCCCCCTCCTGTATAACT

GCAGCCAACCGACGAAAAGGCTTCGTCTCAAAAAAGACGGTAAAACCGCCGTCGGTAGAAACGGATCAAA

CTGCGAGATTGATCATACCGCGATATCGCAAAA

>NODE_871_length_243_cov_3.445946

TTAACCAAAGTCTTTTTTCAAGACGGTCCACATACACTATGATAAATCAACGGGCGAGTTACGTTCGATT

CGGATTATAGGTTTTACATCCCGTCATCGAGCTCCTCGTCCTCGTCGAAGTCGGCATCGTCCTCCGCGGT

GGCGTCCTGGTACTGCTGGTATTCGGACACGAGGTCGTTCATGTTTGACTCGGCCTCCGTGAACTCCATC

TCGTCCATACCCGGAAAAGAGGAGCCTCCAGAA

>NODE_872_length_243_cov_3.256757

GTCCTTAAGGGACTCGGTGAGCGGCTTGAAGACCTTCTTGTAGTACTTTTCTCTCTTCTTCACGTCCGCC

GCATCCTCATCTCCGAAGCTGAGGCCCTCCTTCGTCACGGACTGCAGTTTTTTGTCCTCGTACTCGGTGA

TCGACTGGATCGTCATCTCGTCCACAGCCTCGGTGAGGTAGACCACCTCCAGCCCCTTCTTCGACAACTT

CTCCAGGAAAGGCGACTTCTCAACCGCCTCCGT

>NODE_873_length_243_cov_2.027027

TTTGGTTGTATTGATAGGTTCAAGGAAGGGCGCGAGCGCGAGGGCCAGCGACTTCTCTGTCCGCGATTTT

GGCGTGTATACGTGAGGGCGGCAGAGATGTGTACAGGAGAAGTACGAGTAGAAGAAGAGTAGAGGGAGCA

TATATACGTGCGTGTGTGAGTGTAGGTTGGCGTACATGGTGTGCTGCTGTGAGGTCCGCCGCTGATCTGA

AATGTTTCTGGAAACAAACGTCTCGTGTTCTGT

>NODE_874_length_242_cov_108.000000

TTCGGCGAAGCTCATCAGAGACACGAGAGCTATTCATCTTCCGTCTCTGAGCCGGCCTCGCGCCATAAAC

CATGGCCTTTTCGACGTCCAAGCATTCTCGAAGAGCAACACCTCCCTCCTGACGTATTTTTCTTTAAATG

CGTTCAGCTGCGTAGAGGGCCTGACCCACCGTTTATTCCCACATGACTTCAGCCTTTGCCCTTGGCTGGG

GGATACGGGCTGTGGCTCCGGACATTTTGGAA

>NODE_875_length_242_cov_7.036199

CAGATGCGCTACGGTGTCCGGGAGCACGGCATGTCTTCCATCGTCAACGGCATGTTCGCGTACGGCGGCC

TCAGGCCTTTCGGAGCCACGTTCCTGAACTTCATCACCTACGCCTGGGGAGGGGTTCGCCTCAGCGCTCT

GTCGAGGTTCGGTATCCTTTACGTCATGACGCACGACTCCATCGGACTTGGAGAGGACGGACCGACGCAC

CAGCCTGTGGAGGTTGGAATTCCGGGTGCCAA

>NODE_876_length_242_cov_7.018100

CAAAAAAATCGCCCGCGTTCTCACCGTGTACCACCAGTCGCAGAAGCAGAGGTTCCGCGAGCACTACGCG

GACAAGAAGTACGTGCCGCTCGACCTCCGCCCGAGGAAGACCCGCGCCATCCGCCGCCAGCTTACGTCGG

ACCAGCTGTCGAAGACCACGACTCGCCAGAACAAGAAGAAGGCCAACCTGCCCACGCGCTGCTACGCCCT

CAAGGCCTAAGGCTGCTGTTGTCGGCATTGCT

>NODE_877_length_242_cov_5.411765

GCGCACGGGAGCGCCGATGGCCTTGGGCGGGAGGTTAGTCCTGAACTTGGCCCGTACCACTCCGTTGGTG

CCGTGCCCGCGGCACACCTTACCCCAAACTACGCGGTACTTCGTCCCGTTCTTGAGGGTCTTCGCCTTGT

AGATGTAGGCGATGCGCTTGCCGAGGTAGAACTCCGAGGCCTCCTTGTCTTCCACACCCTGGATCTTGAT

GAGGGAGGTGTGGTGGTACGTGTTCCGGAGAC

>NODE_878_length_242_cov_5.221719

TGGTGTGCGTGCACCCTTCCAACTTGCCAGCTTCAACAGCGCGAACCAACACGGTAATATTTAATCCGTA

CAACACCCTTTTCCCCCTCAACCCGTCTCCCTGGTTCGAAACGCTTGCTACAGCAGCAGTGTTACGTGTT

ATTCCAGACTACCGTCTACTTTTTTCTTGAAACAGCTCGATCTCCTGAGCGAGGGTTTGGCAATACAATC

AATAACAAAATCCCAACAACAGTAGGTAGCAG

>NODE_879_length_242_cov_4.443439

TCTTCGAAAAAAAACCGTACGATGTAGGAAAAGTAGGCAAGGGGTAAACAATCAAAAGCCAACTCCAGTA

CGAAGAAGAGAAGAGGAAAGGGTAAACGATTAAAAGCCAACTCCAGTTAACGCCCACCCTTCTGCCTTGG

CACTCACTTCTTCATAAAAAAGACTCAGCCACGCAATACCGAGCGCATCCGGCCCGTAGTGTGGTAACAA

AACGAGGTAGAAATACGCTTGTCTCCTCCAGA

>NODE_880_length_242_cov_3.764706

GCGGAACCATGAGGAACATATCATCGGTACACGTAGCATTCACCGCGGACAGTGAAACCGGACTACAGAT

CCTGGTGAATAACAATCAGGTCGTCGTTGAATCATGGACGGGAGAGAAGGGCTCGACGGATCTGCAAGAG

ATACCAGGCGTAGCGGGGCACTCCGCATATCAGCTGGAGCTGATGGCCGTTTCTAGCGAAGGCAGCTACA

TCGGAATCACAGAAACCGAGATCTACGTGGAA

>NODE_881_length_242_cov_2.312217

GAATGTTACGCCACCAGAACGCCAGATTGGACCACGAGTAGTACCGGCACGAGCTCGGCCTGTACCTTTT

TGACGCCATGGTTTGGCACCACCGCCGCTAACTTCTGAACGAGTTTTCTGAGCACGAGTACCTTGACGAG

CACCAGCTGCATATGCAACTACTACTTGGTGTACTAATGCTTCGTTAAACTCACGTCCAAAAGTAGCTTC

AGAAACTTCAAGACCGCCAGAAGCGTCTTTAA

>NODE_882_length_242_cov_1.705882

TATCTGTACAGGCGTGCAAAACACACGCATCCCGTAGCTGTCTGCGTTCGGACCTTGTTTTTTTTTGTTC

GGTTCAGTGACTGGTTGCTCTCTTCGCCTGTTGTTTTGTGGGTGCGAAGCAGGCAAAGCGAGAGCAAAAA

AAAGCAAGCTCCAGGGTTGGTTGGTTGATGGGCGAGTGAATGAATGAAAAATGACTGGTCGGTCGCTCTG

TGGAATGAAAAAAACGACTGGTCGGTCGCTCT

>NODE_883_length_241_cov_468.577273

CAGTGTTGCTGATCATCCTCTAAAACCAGCTATAGATCGTAGACTTGGTAGGCCATTACCCCACCAACTA

TCTAATCTAACGCGGGCTGATCCAAATCCGATAAATCTTTCCCCCGAAGGGCGTATACGGTATTACTCTC

AGTTTCCCGAGGCTATTCCGTAGATCTGGGTACATTCCCACGCGTTACTAACCCGTCCGCCGCTCGCCCC

GAAGGGTGCGCTCGACTTGCATGTGTTAGGC

>NODE_884_length_241_cov_7.622727

GCGGTCGAGCAACGCGCTCCCGGAGAGGAGTTCCACGTGAGCGCTACCATGACTACGCCTCCCCTGAGCG

GACGTTACATCGGGTACTGGCGCCTGCGCACGGCCGACGGCCAGAACTTTGGACACCGCATCTGGGCCGA

CGTGCTTGTCACGGGCGGTACCGCGGACCTCGTCATCGAAGGCAGCATGGACGGTGACTGGGACCTAGTG

AACATGCAGCGTCTTGGAATTCTCGGTGCCA

>NODE_885_length_241_cov_2.313636

TCTACCAAGAGCGCGGCCGAGCACGATGTTGGTGCTGGTTCTGGTGCGGTGGAAGAGGGCACCGCCCCCG

AAGACACAGCGCCGGTTGCTGACCCCGCCGCCACCGCCGATACTACTGCGTCGGCGACGCTGCTGGAGTC

AGAGGCGGGTGCGGAGGGAGGGCAGACCTCAGACTACCAAGACATGCTAGCGGCGTCGATGCGCTCTCTG

GCCTCGTCCATGACGGCGTCTCTGCCGGCCT

>NODE_886_length_241_cov_2.054545

CCCAAAGGATACTCGTGTACTCGTACTCAGTCACAGGCAGGCAGCGCTTCGGTGTTCGGACGAGAGTTGA

AGATGGTTGGTGTGGACGGAGAGCGCTTCAAGATGATTCGTCGAAACAGCCTGCGCAATGAGGTCGACGG

CTCGTTCGACGAGCTCGAAGAATTTGAAGACCGCAACGAAACCACCGTCAATGACAAGGTGACCCATCAG

GAGTGCGTGTTGCTGAACCGGGTGCTCTCGG

>NODE_887_length_241_cov_1.909091

TGGTATGGCAACCTAGTTTTAGGAGAGCTTTACAGCGCTAAAGAATTTCTGCAAGTAGAAGACTACACAC

ATGTTCAACGTTGGGCAAAAGCAATGTTAGCTCGCCCAGCTTTTCAACGTGGCCGTATCGTCAATAAAAC

TTGGGGAGAAGCGTGGGAACAACTTGAAAATCGTCACAATGCTGAAGATATTGATAAGGTGTTAGCGCTA

CAGCCTAAATAACCTGAGCTATTAATTTCGC

>NODE_888_length_240_cov_8.178082

TTTCAATATCAACCAATCGTGGTCTAAGGAATTCGGTTACAGAACCCTGCATATATGTCCTCTCTTAAAG

TGCAACTTTACTTAGAGTAAAGTTCAACAATCAACTGTTCATTAATTTCGGCAGATAAATCTGAGCGATC

AGGGATGCGTTTGAAAACACCTTCTAATTTCTTATTATCTACTTCAACCCAAACTGGCTTCTCACGTTGT

TCAGCTAATTCTAAAGCAGCGATGATACGT

>NODE_889_length_240_cov_6.237443

TTGGCACGCGAGTATTCCATCCCCGGCCTTCCGTGGCGAACAGGCTTGTACGTGATCGAGAACTCCCCGA

GGTAGTGGCCCACCATCTCCGGCTTGATCTCCACTCCGTTCATGGACTTGCCGTTGTATATTCCGACAAT

GGAGCCGATCATCTCCGGCACGATGAGCATGTTCCGGAGGTGGGTCTTGATACCCTTAGGCTTCTCGCCG

CTCGCGGCGTTCTTCTTGGCCTTCCGGAGG

>NODE_890_length_240_cov_4.287671

GCCCGTGTACTCTAGTCGCAAAGAAGAACGGCAGCCCTCGTTTCTGTGTTGATTATCGTCATACCCTCAA

CCGTCATATCGTCCGCAAGAGCTGGCCCCTTCCGAACCTGGATTCCTGTCTAGATGCAGTCGGAGGCGTC

AAGTTCATCTCTACAGCCGATGTCCTCAGCGCGTTTTGGCAACTGCCCGTTGCAGAAGAGCACATCGACC

GGACAGCATTCGTCACTCCTACCGGCAAGT

>NODE_891_length_240_cov_3.579909

CGGGGAGGAGAGCGGGTGTAAAGCGTGCGCAAAAAAGGCGGAGGAACCTCCCGGCAAGGAGCAACGAGGA

CGGGGGACGGGGGGTTGCGGGGTGTGTGCGACGAGGGGGTGGCTTCAGCGAGTGGGAGCGGCTCTGCTGT

GTGGAACGAAGCAGGAGGGAAGGAGGACTGGCTGGTCCAAGTGGGTGGACGGAGAAGAGAGAGGGAGAAC

GGGTGGGAGCTTGCTAGTAGGTGTTTTTGT

>NODE_892_length_240_cov_3.054795

AGCCTATGAAAAACCTATGAATAGAGAGTTCGATCGCGACAAATTGCGCAGGTCAAGTTCCCTCTTCCAC

CGGCCGGACGATTTTCAGCAAGGCATGGCAAGGCAAGGCTCCTCTCGCTGCCGGCAAGACGGCGCTCCAT

CGTCGTCGCTTGTGCGACCTATTTGTCGGCAAACACGACGTCCATGTTCTGGTATCTGCGGTAGTCCTTG

GTAGCGGCGTTCCAATCGGTTGTCATAGTG

>NODE_893_length_240_cov_2.858447

GTGGGGAACTGCCAGCGATGAATGTAATTGATGCTGTAGCACGATTAGTGCCGGGTGTATTGGGACATAA

AGAATCAGCAGAGCAGGACTCCTTCTCTAACGGTTTATTAGATTGTCCTCATTATACTAGGCCAGAAGTA

CTAACTACCGCTTCGGGGGATGAAATGTCAGTACCAAAAGTTTTGTTAAGTGGTAATCATGAACACATAC

GCTTATGGCGCCAAGAACAGTCTCTATTGA

>NODE_894_length_240_cov_2.063927

GGCGGCAGCTGGTCCAACCATTTGGTGGGGTTTCGGTTGAACATTGGCCAGTCCGGCACCACAAGAACAA

TAGACAGCACGACGCCCGCAGCCCACCCCATAAGCGTGTATGTGAAGTCCCCCATGACAAAGCCGATGAC

CCATCCCAAAGCCCCTAGCAACAGAACGATGTAGCAGTACAGGTTCTCCGCCAACAGTTGCCCCTTGAAG

TCCATCGTAGCTGCCGTGTTAGGGTAGGGA

>NODE_895_length_240_cov_1.876712

GCTGATGAAATCGGTCCCCGAGCTGAGCGGGCGGGACGAGGATTGGCTTCTGGGAACCTTCGGGTCGATC

TGGGCGGTCATCGACCCGAACCAGGACGGGAGCATTTCTGAACAGGAGTTCATGATGGAGGGAGGCCTTT

ACGAAACACTTCTGGCCAACTATTTCCGCCATCTCATGTGACACCACATCGGGGCGGGCTATTCGTAGAA

CGCACAGAAGGAAGACGCGCGGGGTGGAAT

>NODE_896_length_239_cov_7.559633

CCGGGTCTCATGAGTGAAAGTGCAGTCCTTGTACATCTCCTCAACATCGTCGAAGGTGATTTCGTTCTCG

ATGTAGGCCGCGAAGTGCTCCTTGTACTTGGTCTCATCTTCCTCCGTGAGGTACTCCATGTACTCCGCGA

TGTGGGCACCGTAGATGCGGTCGCGGTGGAGCTCGGGGTCGTACGACTTGCTGTCGCGGTCGTATCCGGG

GAAACGCTTCTCGTTGTGGGGCACGTCGA

>NODE_897_length_239_cov_7.013761

AGGAGAGGAGGGGCAGGTCGGAGGTGCACGGCATCAGGTACTCCCCGTTCATGAACTTAATCGCCCCCAC

TTTGCTGGCAATGGCCTTGACGTCCTCCGAGGGCCCGACGAGGAGCGAGGTGCCCGAGTCTAGGATGGCG

CTCTTCACCGTGGTCACCGACTCGCCCGAAACCGTGACGTCGTCCATGTCGATCTGCCAGTACCCCTTCT

TGGTCACGGGGACGTAGCTGATGTCGTGG

>NODE_898_length_239_cov_6.426606

ACCTGCACTTGGGAGATAAACACGCCCTCGGACGGGAAAGTTCCTCCGACCACGATGGCGCCCAAGAAGC

CCAACAGAATCCATGGCCAGTCGTCCTTGCTCAAGCCCCACAAGCGGGCCCTCGCCGGCTTGGGCAGCTT

CTCCTTCTCTTCCTCCTCATCTTTGTCAACCAAGGTTGAAAGGTCGATCCCTTCCGGGCCGAGGTTTGAG

CTCTGCCGACGGAGCGGGCCGCTAACGCT

>NODE_899_length_239_cov_4.032110

CGGACAAGGTGGTTTCCACTGATTTGGGGGGGGCGTTGGACGCTATAGAGGCTGTTACAAAGTGCGCCGA

AATGACCTGGGTATATCCCGATGTGACGGGGAATTTCCATGTGTGTGTTGTCCATCGGTGCCAACAGCAG

GACACGATAGGCTCGTGGCTGTAATAGGATATGTGCGCGCACATTTTTGTGGTGTTCGGTTCCTTCCGCC

TTCTCTTCGTAGCACGTCACTGGAGGCTG

>NODE_900_length_239_cov_1.944954

AAAACATAAAAGGAAACTGGGGCACTACTGAGAAGAGGGTTCTAACCGTTCGCCGGCATCGCGTACAGTA

GCGTTGGTACGTTTTATACCAGCAAAACTATGCCAGCATGCCGACCTTTCGCTTCCACACGAGTCACTAC

CGATCGTCCTACAATATACGATACCAACACAGCAGTCGTGCTAAATATACGTAAAGTAGCATTTCCCGTT

TCTTACCCCGTACGCTTACTTAGCGTGAC

>NODE_901_length_238_cov_952.391705

TAAAGATGCGCGTGATCAGTATGAGATTCGTACTCACAAGCGCTTGATTGATATCGTAGAACCAACTGAA

AAGACTGTAGACGTTGGGTCCATACCAGCGTCTACAGGGGATGAAGATCGTTTCACTTGGTCCGTGTTTT

GGATCAGCCGGTGTCAGACCCGTGCTATGACGCAAAAAGTGTCTCGAACAGAGAAATGCTGACCCTTTGG

AATAATCGGTTGCCAAGGAACTCCAGTC

>NODE_902_length_238_cov_6.691244

CAACGCACTTTGAGAATAATGTAAGCTATGCTCTGGGGGGGTAGGTATCTGACGAAACGTAACATAGTAC

GAGATGTATCGGGTAATATGTGTATTGGGGTGTAGAGGTTTGAGTGTTGAGAAGCTTCGTCCGTGTTGTT

GATACATGTATGGGCTCAGACGTGGTTCACCCAGTCGCCAGTACGTAGCCGATAGCTGGGTGGTTTACCT

GACAAAACAGTTGACTGAGGCAGCACCT

>NODE_903_length_238_cov_3.940092

CACCGTTCACCTGGTACGGGCCAGTGACCAGGAGCGTTCCCGACGGGAGCTGCTTCAGGAACACCACACG

CTTGCCGCGGAAGCGTCCAGAGAGGAGGATAAGGATAGTTCCGGGGGTGATGCTGCTGCGAAGCTTGGTA

GGGTTGTGCTTCTGCGGGGCGCCCTTGGCTTTCACATCGTCCCCAGCGTACCACTTGGGGGCTTTCACCT

GGGGCGCCGCTGCCTTAGTCATGGTGCT

>NODE_904_length_238_cov_2.645161

ACATGTCGTCGTCGGTGGTGTCGGGGTCCGGCGTGTCCTCTTCCACTCCACTGTCGACTGCTGCTGTTGC

TGAAGCTGCTGCTGCTGCCGAGGCTGCTGAAGCCGTTAACCGCTGGGGCTCTTCTGCTCCTCCTTCTGTG

TCTGGGACGGGTCCTGAAGATACCCCGGCGGCTTCACCGGCTGGGCCTTCTGGGTCTGCCGCCAACTCCG

CTACCGGCGCTGAAAGTAACCCCTCGGC

>NODE_905_length_238_cov_2.258065

GCTGCCGCTGAGCCCAATCACCAATCACCCAATGTGTTTCCTCGTCTGTTCTTGCTTAATTTGATTATCC

GATCTTCCAACACATGTTCCTCGTCTGAATAAGCGACGACAGACGCCGCTGCTATGATATCGCATCAGGC

TATCCAGTCACACGGACCATAATCCCTTCATCCTTTTTACTCCGACCACATGATCCAGCTAGGCCCTGGC

CAGTCCAGGCCAGGCCAGGCATCGACGA

>NODE_906_length_238_cov_1.944700

ACCGATCTTATGAGAAAGACCTGTGTAGAACAGAACACGTTCTGTTGTGGTGGTTTTGCCTGCATCTACG

TGAGCAACAATACCTATATTACGATAGCGCTCTATAGGGGTTATACGGGCCAATTTTAATTCCTCTTGCA

GTTATTAATTTACTTAAAAGTTAAATTAAATAGCTAAATAATTAATCGGGGACTGCAAGAGGAAGTTGTT

ATGATATTAAATCATATTTGGGGTGGAA

>NODE_907_length_238_cov_1.290323

GCAGGCGACCTGCTGTCGTAGTTGAGCTCGTTCTCGGACGACGTGCCCTCGTGCGCAAGAGGAAGAGGCC

CCGTGGGCTTCAGCGAATTATTTTTCTCCACGGGACGCGTGGGGGCCTCGTCTCCGACACGAAACTCAAC

CTCGTTGATGACGAGGCTTCCATTGCTGGGAGGGGTCACTTCGATCTTCACGTAGCGCGCCAACGCCCGA

ATTGCGCCGAATGTCTGCAATCCGTCGC

>NODE_908_length_237_cov_10.759259

AAGCATAACAGCATAATGGCATGATGGCCATCCAGACCAATCCAGGCAACCAAGCCAATACAATCAAGCC

CAGAGCAATCAAAAACCCAGATCAACCCAGCTCAAACCAGACACATCCATATTTCCAGTCCACACTACCA

GTCCAGACCGAACCACAATAACCCATTCCAATCGAAATCCATGTTTAAATCCATCGACAAAGAAGCCCGA

TAGATTGGAAACCCTGGAAGTCGGGGG

>NODE_909_length_237_cov_10.212963

GACCGATTGTCCATTTTTTTTACATGTTTAATGTCTCCAAGCGTCCACCCGCCTCCATCATCAGCAGTAC

ACAAATGAATACAGCAACCAAAATAACCCGTACGTCCTCAAAATAAGGGTGAGGATACGTTTATAGCATT

TTTGTGGTGCTGTTGTTCCTGTGCCTGCAAAAGATAAGAATCCACTCCGCCTGCCGAAAACGCGCTACGT

TCGTCCATCATTACTCCTGCCTACCCC

>NODE_910_length_237_cov_3.805556

CTCCGAACACCAGCGCCTGCGTGATGTACGTTGACGACGTCATGGCCGCCTCGATCGGTCCAGGCACAAC

ATACTTTCCGTTCTCCAGCTTGTACTGCTCCTTGATGCGCCCGGTAATGCGTAGGTAGGTCCCATCTTCC

ATGCAGCCCAGGTCTCCAGTACGGAAGTACCTTTGGCCGTCGAAGTCGAAGAAGACCTCCAAGAAGCAAA

CTGTGATGGAATTATCTGGTGCCAAGG

>NODE_911_length_237_cov_3.027778

TACTGTAATCGCACTAGTCGCACCACCCATGACAACACACGGGTGTTAGGTGGGCCTTGTGCCATTGTGA

TTAGTTGAATGTAGACGCTGCAAACACGAAAATAAACTTGTGAAGTATGCATAGCTCAAGGGATACCCAG

AGGAAGAATGGGTGAAGGGGGCAGCAAACCATTATAATATAGTACGAGTACAAGTAGTGGGTACCCCGAG

TAAACGCCCAAGTTAATAGCATTGAGC

>NODE_912_length_237_cov_2.861111

GCCTCCCGCCTATTCTACACCAGCCAAAATTTCCAATCGACTAACAATCGAAGAGACAACACCATGTGAG

AATAACCGGACGCGTCGCGTATGTACGATCGATCGCGCAGAATCAAATAACCGGGTGCCCCAACACCGGC

GGAACAGCCGCAGACCCAACCATCAACCATATTCGCTACCGCCAGCGGTACGCTTCTTTTCAGGCCAGCA

AGTGACTTTACATACAGCACCAGTACT

>NODE_913_length_237_cov_2.833333

ATCCATCCATCCATCCCTCCGCCCCCCCTATATCTCTCCTGTGAGCAACAACTACTGTCTCGAAAAAAAA

AAGTGGACGAAAAAGTCGGTATATATTTTCTTAGTTCCGCGAAAAACAGAACGAACGTCGATCGAGCCAG

CGCGCGGGAAAAAACAGACTGGTGAGATACCTAGCCCAACCCTGAACGAAACCAAAACCATAGCCCCTAT

AACGTGTAAGAGGTGTAGATACCTAAA

>NODE_914_length_236_cov_6.004651

GATTTTGTTGTGAGTTTTGTTGATTGGTTGTTTTTTTTTTTTGATCTTTCATATCATTATCACGATTGGC

TGCTTAATCGGAGCCATTTATAAACATACTATCTCCTCCCTCACCTCCTGCACAAAACACACACTCTGCC

TAAAACCAGCTCAAACCCGATCCCCAGCGCTCCCGACAAAACTTGGGACCTAGAAGCTTACGCTGATATA

CAGTCAGCACGGAACGAACATCAGAA

>NODE_915_length_236_cov_5.232558

TCCACTCGTATTTTTGTTTAGTGCCTATGTATGCCGTGTATTTATCGTTAAATTCTGGCGGCTGGAGTAG

CGCAAAAGGACGAGACTGGGATCCGCTTTGAGGGCGTTCGCTTTCCATCTCGTTTCATTATGAGGTCGGG

CCCGGCTAATGCTTTGCTTGCTTGCACAGACAGCAATGCTTGCTTGCATGCACAGCAATGTTTTGCCGTT

GTTTGTTTGCTTTACTTGCTTGCTTG

>NODE_916_length_236_cov_3.795349

CAAATTTTTCGTTAGCTTTAGCGAAATCTGTGAATAAACGTGCAGCAGCACCTGGATGTTCGTTTGAAAA

TGCAATTAGTGTAGGTCCTTTAAATGTGTCATCAACACATTCAAATTCAGTACCAGCTACCGCACGACGT

GCTAATGTATTACGGACAACTTTCATCCATACGCCATTCTCACGTGCTTGCTTACGTAGCACTGTAATTG

CTTCAACTGTTACACCGCGGGCATCC

>NODE_917_length_236_cov_2.953488

CTCCCCTCCCCTCCCGCTCTCAAGGGTATGCTGCCACTTCCTTCGTTGGTTCCACGATTTGGTGTATTTC

GTTTTGGTTCTCTCTGGTTAGAGCTTTCGTTCGGCCATTCGAGATCGTTCGTTCACTCGCTCGCTCGTCG

TGGTGGTTGCCGCTCCAGTCCTCTTTGCTCGGCTCTGCTCCGCGTTTCTCCGCGCTGCTCCGGTTCATCT

CTTGCCTTTTCTCCGCTTTTTCTTCG

>NODE_918_length_236_cov_2.637209

GCTACCACTGCCGATCTCAGCCAGAAATAGTCTATGCACACGTCATACAGTTAGTGCGCTGCTAGAGTGT

TGACCTCCGTGGAGTTCTGACAGATACTCGAGGCTTGCACTCGTGTTCTCAGTAGCCCTTGTAGCTGTCG

GATTTGTAGATGCTCATGCAGCCGACGCCTCCGCACACCCTGGAGCGGTCTCCAGCGCACTCGTAGTCGC

ACACGCCTCCCTTCTTGGTAGGCTTG

>NODE_919_length_236_cov_2.479070

AGTGCCGGTGTAATCTGCAGCAAGCAGTACCATACATACCCAGAGAACACAGAAAAAGGGAATGGATGTA

GATACTGGCGTGGATCATAGATAGATACACACGTTACGTACGTACGTAACGCGGTGGTGAGAGTGAAAGG

AGAGAAAAGGACGGGACAATGAATGATTCGAGAAAAGGACGAGAACGACCGGACCCTAAATGCCCTTTTG

AAGAAGGAAGGTCGGAGAAGAAAGAA

>NODE_920_length_236_cov_2.265116

AATGTCACTAGATGATATCACAACTAAGGTACAGTGGGCTCCCGGTGTTAAGCGACTTCAGTTCGGTGTC

CAGGCTCAACGACTTGTTTTGTTTCTGAAGTCGATCTAACTATTTTGTGTGGGATTTACAGTCGATCCCA

CACTGGTGAGCTGTGTTTTTTAAACCGATTTTTTCTACGTATTGTTCGTGCCAATCTGAGCTCATTTTTC

TCGACACGTTATTTCTTAACTGCAGG

>NODE_921_length_236_cov_2.125581

TTTGTTAATACTATGACACAAGCACCAACAGTTGATCAGTTATTGCCTTTACCTAAGTCTGACGATGTAG

ATATGAAGCATCGTTGGGATTACATATACGAACCTGATGCTAACGTGTTGTTAGATCAACTATTAGTTCG

TTATGTAGAATCGCAAGTGTACCAAGGCGTGGTTGAAAACCTAGCCTGTGAACAAGCTGCCCGTATGGTA

GCCATGAAAGCTGCTACAGATAATGC

>NODE_922_length_235_cov_25.299065

AGCGGAGGAAAAGAAACTAACTAGGATTGCCCCAGTAGCGGCGAGTGAACAGGCAAAAGCTCAAGTTGAA

AATCCTTGGCACAAAGTTAAGGAATTGTAATCTAGAGAGGATATTCGGAGGGGATGGTCGTAGCATAAGT

TCCTTGGAACAGGACGTCAAAGAGGGTGAAAACCCCGTAGTCGGCTACTACCGGATCTAACGAATGTTTT

CGAAGAGTCGGGTTGTTTGGGAATG

>NODE_923_length_235_cov_23.518692

GCTGCTGCTGCTGCTGCTGCTTGCTCTGCTGGTGGTGTCAAAAGTAGAAAAAAAAAGTGGTGTTTGCTCC

CTCTGTGCTGGGTGCTGTGAGAATCGAGGCTGCTGCAGTCCTGATGATGCACTGGCGATGAGATGCAATG

CTGTGCTTCCTGTCCTGTTCTTGTCTGGTGTGCAGTCCGCAGCGATGTTGTGTTGCGTGGAGTCGGGCAG

CAGTGTGTGCAGGTGTTGGGGGGAA

>NODE_924_length_235_cov_5.925234

CCATGATGGTAGCGAAAGCCCCACCTGCTAACCGTCCTACCACCAAACAATCCAACCCGTCTTTTACGAC

AAGGTGACCAATTCCAGCAAAAGGAACAACTCATCAGTGTATCCCTCAAGCGAACGCACCAATACATGCA

GCCGACTTCCAGCGAAGCAACCGGCAGGGAGCTGACAACTGACAATGAGCCGATTCATACAAGATAGGCA

ATAACTCTCGACAAAATGCTTGCAT

>NODE_925_length_235_cov_5.144860

CTCTTCGAAATGCCCTTTCTTCCAGAAACCCTCTTCGCTCGTGCCGACGCCGAAGTCGACCGTGAGCAGC

CGGTTGTCGTCGGTGTCCACGTAGGTGTACATCTGAGTTTCGTCCCAGTACAGGCCGAACGTGTGAAAGT

CGTCGTTGAAGGTCGCGCCGTCTAGCTTCATCGTCCCGTACGTCAAGTCCCACGAGTCCATGCTGTAGTG

GGGACCCCAGTGTACCGCACTCGAC

>NODE_926_length_235_cov_5.014019

ATTCCAATCGGGTTATAAAAACCCAATTCCTCTAATACTTTCCCATCTCTTTTTGTTCTAACATCCATTA

CCACAATTCTATAAAAAGGTTGTTTTTTACGGCCACCACGTTTAAATCGTATTTTTAACATTTCTAATAT

AAGTTTTATTATAATATATTTAATTTCTCTTTCTAAAAAATGGGAGGTTCTGTAAAAGTAATAGACTCTA

ATGTTCTTATTATCCACTCAAGGAA

>NODE_927_length_235_cov_4.883178

TTTATTTTCGCTCTCCGAAAACAGCCAGCATGCCGAACCCCAGCTTCTCGAACCACACCACTCTCTCGGA

CGAAGTCCTCGCGGAGTTCCGCAAGCTTCCCCAGAATGGCAAGATCATGGCCGAGTACATCTGGATCGGC

GGCTCCGGCCAGGACCTGCGCTGCAAGACCCGGACGTTCGACGAGAAGCCGAAGACCGCCGCTGACCTGC

CGAAGTGGAACTTCGACGGCTCTTC

>NODE_928_length_235_cov_4.200935

TGCTATCACGGATGATGGTCCGGGCCGTGAAGGCATTGCCGTACGCCTGATACCAACGTCCGAGGTATTT

GCTGACTATCAGTTCATCAACTGTCTCGATCGCAGAGGTTGGGTGCAGACAAGCGACGCCCAGGACCCCA

CCAGCTAAGAGCTGGAGCAGAGATCTCGCCATGTTATTCTCCAAGTTTTTAAATCCGATCACAGAAGTAG

TTTGTTGTGTGCACTGACGACGTGT

>NODE_929_length_235_cov_3.869159

GTAACATCGTTTTTGCCGGCGTCAGACTGAATGGCTGCATAGGGTATATTAATATTTCCGGCTGAACGGA

CTATGCTTGCGCAAGCGAGTGAGGAAATGTGACCGAGTTGCTCGCTTTCGTCTTGAGAGATGACCGGAAG

AAAATTGTAGCGTGGGTGGTTTCGGGGGCAGAACCCATGAGGTTTTGGTAATCACTGTTGTTTGTGCTTT

CACGTTCGTCAGGCAGCTCGCGGTG

>NODE_930_length_235_cov_3.742991

TTCAATCAATAAGTTGTATTGGGCTGGAGTGGAAAAGCACTGCCGGCTTTTTTCCGGTTCTTTCTTCTGT

AGACCCGCCGCCGACAGGCTGCAGTACATACAGTACATAGGTAGGCGACCCGTGACGCTGGCCATCCGAC

GTGTGCGGCCTTCTCGTCTCTCATCATCTCTCGTCATCTCCCATCATCTCTCATCTGATGGGACAAAAGG

GTGTCGCCATTTGGTATCACCATGT

>NODE_931_length_235_cov_2.042056

ATGGGTGAATATCATCGTTAGAAGTGATTAAACAAACTTCCATATCACCAAGGTTGTTTACGATACCTTC

AGAAATAGTGCTAACTGAAGTTGCTTGAAATTCTACAGAACTACCGTTTGCTGCAGCATAACCTTCAACT

GCTAGAGTTAAATCAGAGTCAGTAGGTAATGCATCAAAAGAGAAAGTACCATTTTCGCCAGTTACTGTAG

TAAAGGTTGGAACACCAAGGTCAAC

>NODE_932_length_234_cov_46.220657

GTACACGCGTGAAAGAGTGGGAGGATCCGCAGCCCAGCGGCCCCCCAGGGACGATACATGCGGAGAGCAA

TTTTGGATGCCTCGTCGTGGCCTTTGATACGGCCATGTCGGGTGTTCCGTGTTTTATTTTTTTAATTCAT

GAAGTCTGCCTTTCAAGATACCGTCCGTTTTTTTGCCTCGCGCTGTGAAGCGCACTGTGGCCGGCACGGT

ATAATCGGCTAGATACATACTGGA

>NODE_933_length_234_cov_35.769953

ACACGAGCTGGGATGCGAACGCGTGGTTTACACAGGCCGCGGCATCGCTCTCCTTGCCCTTAAGCGCGCT

CAGAGGTTTAAACCCGACGCCGACCTCTCTTCGCCCCTCTTTTGGGGCGGGGTGTCTGGAAGGTTGAGGG

CCTCGTTTCATGAGGCACAAGCGCGGCAGAAGGCTCAATATTAAACTCCAGTCGCACGCGTTGATGCTGC

TGTCTGGAATACTCGCGTGCCAAG

>NODE_934_length_234_cov_25.643192

TTGCACGCTTTTATCAAGCTCCCTGGCTCTTCAGGTGCATTATATGCGCCGATGAGATAGTCGGGCAGAA

GGTGGTGCTACATTAAGTCGAGGCGTGTTGCATTTGCTGCTCGTCTTGCGGCCCCGGCGTACGAGCAGGA

ACATTCAGAGAGAGGGGCGTAGCAGCGGAAGCGCAAGCTTGTGTCTGTCTCGGTGCTTTCGCTTCTGGTT

TCTGTATCGTCGGGGCGAACATTT

>NODE_935_length_234_cov_21.375587

GGCGACCTTGTTGAGCTCTCCCTCGTACGTTAGGCCGTCGACTTCCTTGCCGTACTCGTTGGTACACTTG

CCCACGATTTCGTGCTTTGTCGGCACGTACGAGATCCTGTCTGTCCCCTCGTCGTTCGAGATGACGGGGT

TGGGGAAGCACCTCTTGCCCATCTCCAGACCGAGGAATGCCTTAAGCGTGGTGATGTAAGAGCCGACGTT

GATGGCATGTCCACTGGGGTACGC

>NODE_936_length_234_cov_6.497653

TTCCGCCTAACGCAGATCTTGTTTTCGAGGTGGAATTGCACGCCATCAACGGTGTCGGGAAGTGATTTTG

ATGGAAGAGGAGAAGAAGGAGCAAATAGTTTGACGGCAAACGATGCTCAGATACGGACCGGACCTTTTTG

GACACGTCAAGACCTCTTTCGGCCTTAATATACTAGTTTTAGTGTAGGGGGGCGGTCAGTCGAGCGGTCA

AACCGCGGTCTCTTGTTTTGGCCT

>NODE_937_length_234_cov_6.159624

GTGGTCAAACACCGCCACACCTGAAACGGTGCGACGGCGCCAACAGACTGAACGGGACCCCGGCTCCAAT

AGAGAAACAAGCGCCCCTCCCGCTGTAGGACGAAACGCTAGACACAACTAACTACCCACGTGCGCCAATG

CCAATGCCAAGCCAAAAGCCTTGCCGACCATTATCCGTGGCCTAAACTAGCCAGCGAGAAACTGGACGAG

CCGCCCTGAAACTGAAAAATACAG

>NODE_938_length_234_cov_5.474178

ATCTTCGCGTGCGAGGCCGACAACGCGCAAAACGAATGCTTTGACGCAACCGAGCGGGACTGGTACATGG

CAGCTGTCGACGACGACACGGGGGTTTCCGACGAAACCGCCCTCGGGGATGTGATCGTTACGGATCCGTA

CATCGACGGCGTCGGGTCGGAGAAGGACTGGCTGGTGACCATCGCGCGCGCCGTCTACAGCGACACGGGC

TCGACGTCCGGACAACTCTTGGGA

>NODE_939_length_234_cov_5.201878

ACTGTAACTGTTTCACCTTCATATAAGGTTGTCACTTCAAGTTTTTGTTGATGTGGCTCTACAGTAATAG

TTGAATCTAGCGTCATAATTGACGGATAATGTTTAGAGATTTTCGCGAATTTACTCTCGATAGATTGCTT

AATACCTTCTGTAATTTCAACATGGTGTCCAGAAATTTTAATATTCATAATCTTACCTCTTGGTCATCTT

AATCGTAAACTAATCATTCTTTGC

>NODE_940_length_234_cov_5.084507

CATGATGATGAAGGTCGTAAAGAAGAGTGGGATCTCGCCCGATACGCTCTCATTCAACGCATCGTCGATC

GATCGGCCCGTCAAGTACAGGATGTTGAAAACGTCGTCGGCGTCTGACTCCTCCTCCAGAAGGTCCTGGA

ACTCCAGCTGCCACGCCGAGATGTCCTCATATGTCTCGGTGTTCTCTTCGCCATCATCGTCATCCTCTGA

CTGCAAGGCATAGGTCTGCGTTAA

>NODE_941_length_234_cov_3.413146

CCCGTCAAGTGTTACCCCGCTCTTGATGAAAATGGGTCCGTCCAGGAAGTACTCTCCTAGTTCGAATCGG

ATATGCACGGTGGAGTCGTCGAGTTCTCCGACGCAGTCTGTGCCGTCGAACGTGGCGCCCTTCGTCTCGC

AGATCAGCGTCATGATCTCCTGCTGCTCGGAAACCCCGTTTCCGATTACGTCCGGGACACTGCCCCATAG

AGGAACCGTTGTCGTCGCCATCTT

>NODE_942_length_234_cov_2.070423

TTACATGGCCGGGATCAAGGAAGGGGTCGCCAAGGGCGTCGGCAATGGCATGCTCTTCGTCTCTTTCTAC

ATGAGCTATGCTCTCGCTTTCTGGTTCGGCACCAAGCAGGTGGCTGACGGCGGTGGCCGAAGCGGGGGAG

ACGTCCTATCCTCCATCTTCGCGGTTCTGATGGGCTCCATGATGTTCGGGCAGACCGCTCCGGGTATCGC

AGCGCTGGGCACCGCGCGAGGGGC

>NODE_943_length_234_cov_1.563380

TTCTGGCGCTAACATCGTCGAGACGGACTTCGGAGACGGATCACGCTGCGGCCATCTGGACGAAGTGATT

TTCGACCAGGAGCTGATGACCCCCATCGTCGACAATCCCATGTTCCTGTCGAAGATGACCGCCGCCACTC

TCGAAGACATCAACTACATTGTCGACACAAGCATGGTGGACCCGTACGTGCTTCCTTCCGCTGGTGGTGG

TTCGATTGAAGCGCAGCCGGAGCC

>NODE_944_length_234_cov_1.422535

CCAAAACCTCTGATCTCAATACGATCTGAAGTAGATAATGAGCCGGCCATTTGTTCAAGAATTTCTTTTA

CAGCATTCTCAACATCTTTCACTGGAACGTGTGCGTGTTGTTCCGCAAGTGTTTCTATCAGTTCTGACTT

AGTCATAGCGTTTCCTTAAAAAGAGATATAGGAAGGGCAAAAAATGCCCTTCCAACTAAAAATTAATTAT

ATATTAGTCTTTTTGTGCATTTGC

>NODE_945_length_233_cov_1482.528302

TAGCAGAGCGTAGTGTGACATAGTTCCATGTGTCCTTATCCTCCTTCGGGTGGAATTGGACACAAGGAGC

TTTCTGTGAAGCCGGCCTGTGAGGGATCCGGTGGAGAGATCACTAGTGAGAATGATGACATGAGTAGCGA

CAAAGAGTGTGAGAGACACTCTCGCCGAAAATCCAAGGGTTCCTGCTTAAAGCTAATCTGAGCAGGGTAA

GCCGACCCCTAAGGCGAGGCCGA

>NODE_946_length_233_cov_4.476415

GGTCGTTGCGGATGGCGGTGTGGGACGGGTCCACCGTGATCACCTCGTAGTACTTGAACGTGGCGTCCTG

GTTGACCCAGTAGGAGTTGAGCACACGCATGTTCGAGAACTTGGCGCCCACGCGCTCCTCGGCCACAGAC

CTCAAGTTGCGCACGGCCTTGATCTTGTTGATACCGTGGTGTTTCGGCTTACCGTAAACGATACCCTTGG

CGTTAGGCCTCTTGCGCCCGCCG

>NODE_947_length_233_cov_1.363208

GTTAGGGCTGATAACTTCTACTCGTATAGGAGGGCAAGTGTCGAGGGCCCCGAAGTCGTTCTTGACGGAG

TGACGGATTTCATCCTCCAACAGTCGGAGATACAGGGAGCCATGATTAAGGACTCGGTCAACGTTTTGTT

CGAGGGTGTTCCAGAGAATCGCGATTTGAGGGGGTCCATCGAGGTCCCGGAGGGAGGAGATCAGTCCGCT

AACCTCGTCGTAACCGGAACTAC

>NODE_948_length_232_cov_1258.682464

GGCACCGAGAATTCCAGGCCCCCGTCAATTCATTTGAGTTTTAACCTTGCGGCCGTACTCCCCAGGCGGT

CTATTTAATGCGTTAGCTTTGAATCCCACGGCATATAGCCACAAAATTCTAATAGACATCGTTTACTGCG

TGGACTACCGGGGTATCTAATCCCGTTTGCTCCCCACGCCTTCGCGCCTCAGTGTCAGTCTTTGTCCAGG

TGGCCGCCTTCGCCACTGGTAT

>NODE_949_length_232_cov_137.876777

CCACCACCTGTCACACATACGGTCCAGCACACTCCGAGTCCGCTCAATGGGGTTGTGACTGAACAATTGG

CAGGGAGATTGATCGTAAAAATCGAGGCGGCGGGGGACGAAGACCGCGGGGGCGAGAGGAGAAAAAATCT

CTTCGCGCCTCGGTCCCTCGGTTCGGCTGATGCGTAAAATGCGCTTTCAAGCTTGTTTCCGTTGGGCTGT

CCCCGGCCGCCACATTTTGGAA

>NODE_950_length_232_cov_8.649289

AAGCCATCCTCAAGCAAAGCAACCTGATACATCTCAACCTTCAAGTAACCCCAACTCCACCACCTACTCT

ACTATGTCCGGAAACGGAGGATGCAAGATCTGGGCGGACTGCAGCTGCGGCTCGTCCTGCACCTGCGGAG

ACTCGTGCACCTGCGGCACCGTGAAGGCGGGCTCGAATTCTCACTCCGCCGCCAACTGCCCATGCGGCGA

CAAGTGCAGCTGCTCCGGCAAC

>NODE_951_length_232_cov_7.682464

AAAAAATCATGTTAGGTATATATACAATATGGGGTTCGTAGGGATCTTCTGAGTCTATCTGCGCGTTGAA

ACGTTCAGGTGGCACCGTCCATCTTTTGGCACCGTCCGTCGTTTTATGTAAGAAACTGCAAAAACCCGAC

CTGCATGAGGGTGTGGGTATAGCCGCACGAATGGGTGTGTGTGGTTGGTCGTGAGCTAGCTATCTATACA

TAGACGGCGAGGGTACTGTCTG

>NODE_952_length_232_cov_7.142180

TCAAGCACCCCGGATGACACCTCAAAGCGGACGTTCTTGAACTTGAAAGTGTGGTCGTGGTCGAATTCCC

CGTCGGCGATGAGCTGTAGACTGTGGTCCTTGACCTCTAGGGTTGTCCAATCCTCGCATTCTATCGTTCC

CGCGGTGATCACAGCGATGGTGTCGCCAGAGCTAGCGAGGTCTGCCGCTTGCTGCAGTTCAGCGCACGTA

CCTACGTCCACCGTGTTCTTTG

>NODE_953_length_232_cov_5.213270

AGCCAAGACCTCCTTGTGAGGCGGACTGAATGGACGAGCGATATATCGGAGTGGATTGTCGAGCTACGGA

CCGCAACATACGGCCTGTGTTCGGAAGAAGGAGGAGCATAACGGGAGCGCACACCAGCATTGTCTACTGC

GAGGTGTTGGAGGGGTCAGGATGGAGCCGGAAGGACGGCGGCTGACGTGATCAAAGAGAGAGAGGGAGAT

CGCGCAAATAGAAGACCACCCG

>NODE_954_length_232_cov_3.507109

TTGGCGATACGTGTAGCTCCTGCCGCAGGAGTCTCTCGTCGGAAAACCCTTCAAGCGCCGCGTGCAGGCA

ACCTCTCCCACGCTAGCGAACTCAGAGGACGTTTGCACGGCTGCTGCTGGGCTTGCTGGCTGTATTCGAA

GTCGGGGACTTTGCACTACAATACTGGGAGAGGTATCAATCAGCAAGCTTCGGTAACTACCGTCACCATC

TACTACTAGGGTCTGTATCATG

>NODE_955_length_232_cov_3.180095

CGAAAAACATCGACGGGGTGAAGCCCAGCTTTTCCGCGGCCCACTGGAACTGAGTCGGCGTCAGTCCCCT

CCCGAACTTTGAAGGGCGAAGGGTCATAATGCGTCGGCAGTCGGCGGGAGTGGTGAAGACGACCATCTTC

TTTCCGAACAGCGAGAGCTCGTAGTCTCCCCCGAACTCATCCGCCCACCCTTCCAACTGCTGGACCATCT

GGCGACAAGAACAGAGATTGGT

>NODE_956_length_232_cov_2.815166

TTGCTTTTGGTAAATCCATTGAGACTTGTTTTGCATTATAAATTGCTTGAAGAAATTCTTGATTATCTAC

TCCCGCAATCTCTACTGGGTACTGGAATGCTAAAAATAAACCTAAATGGGAGCGTAAATCTGGGTCTAAG

TCACAAATGTTTTGTCCTTGAAATAAAATTTCTCCTTCGGTTACCTCCTAAGATGGGTGACCGGCAATTA

CTTTTGAAAGAGTACTCTTCCC

>NODE_957_length_232_cov_2.075829

TTCTCTGTCTTGTTCTTGCCTGCTCTGAGGGGACACGCTACTCTGCTTCAAGACATTATTTCGTGAAAGC

TACAACATCGAAACGAGGAGGGGACAGGAGAAGATAAGGGAAAGGGGAAGATACAGAATCTGAACAAGAT

GGAGCTCGGCATGGGCATGGGACTGTACACCGACGCGGGGCTGGCGAACCTTCCGCAAATCATCTCCATG

ATCGGGGGCTCGTTCGTCGGGT

>NODE_958_length_232_cov_1.867299

TTCTTGGTTATTATTTTTTGTCGGTTCACCGGCACTTTTACGAGGACACAAGAGCAGTAGGGCGTAGGGG

GCACGAAATTAGGAACAGGGTATTGTGAGGATGACAGGTAATAGAGACAGGGTATCCCTAGCCAGGTCAG

CCTATGGAACGGGTACTCGTTGGTACTCACGTGTGTTGTTTGCTTAGAGATAGAAGAAGAGCAGCTTTGT

TCCTTTTCCCACGATGAAGACT

>NODE_959_length_231_cov_3.909524

CATCGCCCGTGCCATCCTGAACAACCCTCCCATTCTGCTCCTCGATGAGGCGACATCCGCCCTCGACAAT

GAGAGCGAGAAAGTGGTGCAAGAGGCGTTGGACATGACGTCCGCGCCTCCCTTCTCGTATTCGGTGTGGT

TCACGCCCATGACGTACATCGGGGTGTCATCCTTAGGGGGCGCCGAGATGAACACCTTCTTGGCTCCGCC

GTCGACGTGAGCCTTCGCCTT

>NODE_960_length_231_cov_3.400000

CCACCGAAAGCGTCAGTACCATACAGTTACGGGACACCTGTTATTTCCAATAAACAGGAATACTGACCGG

GCTGAATTTCTTATTAAGGGGAAAAAGGGTGTACTTGACACCCACAAGCAGCACACTGTAGCATATCGAC

TTCAAGATTAATTATTCCTCTGTAAACCTCAGCATGGATTTTCCATGGTGTTTGTTTTCCAGATAGTTGA

GTATTTTCAGATTTAGCCTTG

>NODE_961_length_231_cov_1.861905

TTTTTTGGAAAAAATATGTAAGACCATCTATTATGGGCGTTTGGGGGAAAAAATGACTCTCAAAACTGAA

TTTTAGGACGAATATACCCGTTTTACTCGGCTGAACTGACGAAAACACCGATACACTCAAGATCCACCCC

GTAGTTTTCCGAGGGCTTTCTGGAGCAGTTCTCCGTTTTCTCGCGAGTTGTAGATCCTAATGGCGCGTTC

CGAGAGTTCCACAACCTGATG

>NODE_962_length_231_cov_1.590476

TATGTTACAAGAATCTGTTGATGCACTTTTAGATAACGGTCGTCGTGGTCGTGCTATTACTGGTTCTAAC

AAACGTCCTCTTAAATCACTTGCCGATATGATTAAAGGTAAGCAAGGTCGTTTCCGTCAGAACTTACTTG

GTAAGCGTGTAGATTACTCAGGTCGTTCTGTAATCACAGTTGGTCCTACACTACGTTTACACCAGTGTGG

TTTACCTAAGAAAATGGCACT

>NODE_963_length_230_cov_14.923445

GGGAAGGGGGGCGGGTATTTATTGCGTTTGGCGTACTGGTTTGCAGAGAATAAAAAAACGTCGAGAAAAA

AAAAGTAGATTAGAAAATACTGGAAGTGGAATCCTGTGCTGACCACGTGCTTGATGGTTTGGCGAGGATC

GTTGAACAAGTACCGGCGTAGGAGAAGCCGTGAAGGAGGGGAGATGCAGCTTAACCCTTTCAGAACTGCA

AAACTTATCCCACACTAATT

>NODE_964_length_230_cov_14.315789

AAAGAGCCGGTCGCCCCGGTGGGCTAGCAGGTAACCACGAAGCCTGTCAGCGTCAAGGTCGCGAGTTCGA

GCCCCCCCGAGTGCATGTTCGTAGGGACTTTTTCTCGCAAAAATATCTTGACTAGCGGAAAGCGCGAGAG

CGTGAGTTAGCAAAATTCGATGAAAATCGACGAGCGTTGGGAAGGCTGAACCCTCTGCGCGATAAAAATC

AAGGCAAGAACCGTAGTGGG

>NODE_965_length_230_cov_9.377990

ATTCCATTTTTTTTTTTTCCTTCTCACTTGGTTCGCGATTTTCTTTTTTACCCAGCTTCCTTACTCGTGT

TCGAGGGGTCCCCAAGACTCGGGAAGTGTGCCTGCAGTAGTACTGTTCGCTAAAACGCACAAAATGCGGA

TAGGATACCAGAGCCCTGCCACAGCTCCCATACTTCAACACAAGCGTCGCTTGGTAAGTAAATAGCCCTA

CGAGAACGGGAGGGAGGGGA

>NODE_966_length_230_cov_4.688995

CAGCCACTTTTCTCTATTGCAAACCCTTCCCGTCGAAATCATCCCCGATCCCCGGTACACATGAAGGGCT

TCAGCAGTTCGCAAAGTTCTCGTTGGCGAACTCCCAGTTGACCAGGTTCCACCACGCCGCGATGTATCCA

GGGCGAGAGTTGCGGTAGTCGATGTAGTACGCGTGCTCCCAGACATCACACGTGAGGATGGGCGTGCCCA

GGCCCTCCTTGATGGGGTTG

>NODE_967_length_230_cov_3.511962

GTTCAAGAGGTCACTGCAGGTAGCCCACCGCCGCAGGAAGCTGCTCGAGGAGGGGAAGGTCGTGGGACCC

GCTCTGGAACTGCAGTACCGCCTCCTAGACAAGGCGGTTCTGCAGACGATCCGGGACAGGTTCGGTGGCA

ATCTCAAGGTCTCCTTCGTCGGGGGAGCGGCCACCTCCATGGAAGTGCTCAACTTCTTCGAGAACATCGA

CATCAAAATCCTCGAAGGCT

>NODE_968_length_230_cov_2.933014

AACGACTTCGACAGCATGCACATCGAGTTCAACGTCTTCAAGTCGGCGATCGCATACGACACTTCCATCA

AGACCCTCCCCAACTTGGCGAAGGATGGAGGAGAGGAGCGGATGGTGGAATACGAGTTTGAGCCCGACAC

CAAGGAAGAAGTCATGCAGGACATGTACGAGTACCTTGTGGCGTCGCAGGTATGGCACAGCCTTATGGAG

AGCGCCACTTCGGAGCAATG

>NODE_969_length_230_cov_2.464115

ATAATAAATACCTAAACTATACTTACGCTAAGTGAGTAACTATAAATAATAATAACAGGAATTAAGACAA

CGTAGATGCGCCCAACTCTGCAACTCCGGATAGGACAACAACTAACCATGACGCCACAACTGCAACAAGC

GATCAAGCTTTTGCAGTTATCAACGTTAGAACTTCAACAAGAAATTCAAGAAGTCGCGAATCGCGAAGCT

CGCCAGCCCGTCCACCCCAT

>NODE_970_length_230_cov_1.851675

TTGGCGCTGGCGTGATTTTCGTTGGCAGGGAGGTAAATTCCGCGGGCGTTTGCAGGACGAGTGGAAGAAG

AGGCAGCTTCGCGTTGGAACGGTGAATCGAAGCGGCCAATGTGCTCGTGGTATGATCTTCGCCACCGTGA

TCGTCGCTACTGCTGTGCTGCCGGCAGTTTTACTGCTGCGGCGGATTGGTTTGTCGACTAAATCTTTTAA

GTGCTCTGTTTTTGCAACAA

>NODE_971_length_229_cov_32.024038

GTCATTTGCGGATCATGGGGTCGAAACAGGATCGACGAACGTCTAAAGGGGTTATGCTTTATCTCGGCTG

TCTGCCACCGATATCGGCGAAACTTGTACAATTGCAAATGACAATCGTGCTCCAATGGCGATGGCCGCGT

AAGCGACCGGAACTATTGAAACCTTAAGTCCTAGCGTTTAGCGACCCTAGGCGGGGCCCGTCGGAGCCTG

GCAACAGAATCCGACACCT

>NODE_972_length_229_cov_4.168269

ATGTCGTCGGTGTTGATCAAGGCTTGATCGATGTTTGACGATAGGTTCTTGGTGTATGATAAACAAAAAA

AACGACGCGGCCGGGGCCTTGCGAGAAGGGGAGATAGATGGACTACTGTAGACACTCGAGTATATGTTCG

ATGGCGATACGTGTACGTATGTATTCGATGCCGTTTTTTTCGTGGAGGTGTACGTATACATCGGGAGCTT

TTTGTGTTTTTTATGGTCG

>NODE_973_length_229_cov_3.451923

TCCAAGATCCTCCGCGTGCTCAAGGCCAACGGTCTCGCCGCCGAAATCCCCGAGGACCTGTACATGCTTA

TCAAGAAGGCCGTGGCCGTCCGCAAGCACCTCGAGCGCAACCGCCGCGACAAGGACTCCAAGTTCCGCCT

CATTCTCATCGAGTCGCGCATCCACCGCCTCTCTCGCTACTACCGCACCAACCGCAAGCTCGCGGCCAAC

TGGAAGTACGAGAGCGCCT

>NODE_974_length_229_cov_2.673077

TGTAGATGATATCGTGATTGCAGGTTCGGACTAGTCATGCAGAGATTTCCATGCCGCGTTAGTAACGAAA

TTCCCCACGAATAACCTCGGAGAACTGACTTGGTACACTGGTTGTGCTTTCAAACGCGACTGGGAATTGG

GGACTTTGGAGATTACGCAGAAGGCGTTTATTGAGAGCATGTTGAATCGTTTTGGCGTTAACTCATCCTC

TGACATTCCCGCTACCCAA

>NODE_975_length_229_cov_2.581731

GACGACCCTTGATAGGGTGTTGAAGAGTAGTCCCCTTCAAATCAACTGTTACTAAGTTACGACGAGCCTT

TTCCATTGCTTTTTGGATTGCAGCAGGCACTTCACGTGCTTTACCGTAGCCAAAACCAACACGACCGTTA

CCGTCACCAACTACTGTTAGTGCTGTAAAACTGAATATGCGACCACCTTTAACCACTTTAGATACGCGGT

TAACGGCGATTAGCTTTTC

>NODE_976_length_229_cov_2.442308

GTAACCAAGTCGTTTTCCGCAATCTCCCCCTCGTCTTCTACGAAGATGTCCACCTTCACTTCCACATCCG

GGATTATGTCGCACACGCGATGCACTTCATCTTGCTGCGCGGTGGTCAGCGTGTCCAATCCCTTGCGGTC

TTCTCGCGGCAGCTTGATGTACTCCGCGATCCCCATCGCACCATCACTACCGTCTGAGATGGCCGCGGCC

TCCGCGTCCCCAATGTGCG

>NODE_977_length_229_cov_2.182692

CGACGATTACTCTCGAAAGGATTTCGACGAGCTGCCCGGGTTCTCCGCTGGCCTCGTGTTCACTTCCGGC

AGCCCGGACTGGGCTTACACGATCCGCGCCAACACCACCAAATCCGGCCTCGAGGCGGACACGTACTACG

CCTTCAACACCCCCGAGACGGATTCCTCCACGGAGAACAACTGCAAGTCGCCCACGGACTGCCCGGAAGA

CGACGAAGGTCGGGATATG

>NODE_978_length_229_cov_2.168269

ACCCGATGGACAGTCCCATGGCCTAGCGATACAACGAAACAGTTTTCGACACAGAAAATGCAATCACGAC

GGTTTTGAGTTAAGGAGCCGCCCCCTTTCATTCAGAAACGATTTTCGCCAAAAAAAGTGCTATGGTACCC

CTATCGTAGCACGCGTCCATATTTTTTTCTGTGGGTGGTCGCTTGAGGAGCTCTATCATGCCTCACCCCA

GAAACTGATTTCGGGTGAA

>NODE_979_length_228_cov_5628.106280

GAAAGAACTAAAGGATCGATAGGCCATGCTTTCGCAGCCGGGGGAAGTACTGAATCCCCCGGTCAAGCTC

ACTTTTATCCTTTTATTCTACGAGAGGTTTCCGTCCGCTCTGAGTCAACCTTTGGACACTTGTGTTATTG

TTTAACAAATGTGCCGCCCCAGCCAAACTCCCCACCTGACGTTGTCTCCCGCGTAGATCGCCCCAATAAA

GGGACTTCAGACCAGAAT

>NODE_980_length_228_cov_5.801932

ATCGCACATGCTCGATCTCTCGGAGGAGGTGGACGAGGAGAACATCAGCACCTGCAAGGCTTACCTCGAG

CGCATGGACAAGATGAAGATCTGGCTGGAGATGGAGATCGGTATCACGGGCGGCGAGGAAGACGGTGTCG

ACAACACTGGCATGGACCAGAACAAGCTTTACACCCAGCCCGAGCAGGTACAGCAGTATTACGGTACCAG

TACTGAGAACATCGCCGG

>NODE_981_length_228_cov_3.531401

CCCCAGAACTAATCGACCCGACGGCCTTGCAGATGCAGTCCGTGGGCAAGGAGATGTCGTCCAACGCCTT

CGGCATGGTCTTCGCGTAGTGCTCGGTACACTTCGCCCCGTAGTTCAGCCCAAACATATTCTTGATCCCG

TCCGGGCATGCCTCGAAATTTTCCCTGGTGGTCCGCACGTAGTTGGGGATCATGTTGGTGTCGGTAGTTC

CAGGGTTGATGATGCTCA

>NODE_982_length_228_cov_3.381643

CCCACTCCTGTAGGAGGAGGTCAGACAAATGATGGGAACAGTAGCACAGGGCGGACATCAGTGGTTGCTC

AAACTGATGCGACAGAAGAAAGCGCTAAAACAAAATAAGACAAAACAGGCCCATTGAAAGTGCTACGATA

CGTCCGCGGGGACAGGACGAGATTTCCGCGGAGGGGGGTTAAATGCTGAGGAGTCCGTCGACAATACTGC

AGCACGGTAGGAAGCAAT

>NODE_983_length_228_cov_2.753623

ATGCTGAACGCGGAGACCCACCGTGTGATCCGGTCGGAGCACGAGATCAAGCTCTGGTTCCCCGAGGGCG

TGTGCGAGTGGGACTCGTGCACCCAGGCCTGGGTGTACGAGAACTAAGGTGCCCAATTGGCAACCAACCA

TATGATGATGCCAGCCATCTGATGCTAGCTGGTGGGAGGGGGATGAGCGCGACTTTGCTGTTTTCAGCCA

TGTTTGCGGTGCTTTGTT

>NODE_984_length_228_cov_2.705314

ATGTGTCGCTTTGTAAGTTGTTGCCCAGAGGCTTGGCCGGGCCACTACTGATTATTTATTGTTTCTACGT

TGAGCCGAATTTCGCGCTCTGATCTGATGTGTGCCCCCTGCGAAATAGTGCTGCAAAGCTGATTGGCTCG

TTGTGGACTCTGATTGGCTCGTTGTGAACGCTGATTGGCTCATTTTGAAAGCTGACTGGTTCGTTTTTCA

AAAGCTGATTGGTTCGTT

>NODE_985_length_228_cov_2.449275

ACCATTTGTCTTGAGAGATATGCATCGTTCCATTACACTCTGTCTCAATACAGACATGAATGGACGATGT

CATACATTATGTGTCTGCTTCTTGCACACTTGTCTTGTCATCCACTGCACACAATTATATTTGTTGCATG

ACCGGGGCTTGCTCATCCAGGCCTGTCTGCATCAATGTCAATCAACTTGTCGCTTGTAAATCCAACTTAT

TTTTCTTCAGTATTTTTT

>NODE_986_length_228_cov_2.193237

GCCAGCGCGGATCCAGCCGACCACCCAATCATCATTGAAGAGTATTTTCTCGGTCACGTCCACATCTTCG

TCCATTCGCTCACCCCAATCGTTGAGGTCGCCTCCGTCTTGGATTGTTTTCGTTCCGCTGTTCCCGTCGG

TGGAATCCGAGTATCCCACTCCTTCTCCGCCAAGCGGAACAGCTTCGCCCTCGTCTTCCGCGTGCATTCC

TCCAGTTCTTCGCCATTC

>NODE_987_length_228_cov_1.888889

TCTCGTTTATATATCTCGAAAAATCGGCCATGGAACTGATACCGGTACTGGTATCGATACTGGTACTGAT

ATTGGCACTAGCACTGGTATTGGCGCTACTGTAGTATTGGCACTAAAATTGGTACTGGTATGATACTGGT

TCTTGGACTATTAATAGTACTAATACTGGTACTGGTACTGTAGGAACCAGCAAGACCGAAGTTGCTTGGT

GTCCGGCATCATATTGCC

>NODE_988_length_228_cov_1.772947

TATCTAGCCAAATGTCACAGATCCATATTTTTATACCAGGTACAACATGGCGTAGATGGATTGATGGGCT

ACAGAGCCAAATCGTGTTTGGAGTCGAACTAAGTAGGGGAGGGGATTACTAAAAAAAGATGGAATTAGAC

CTCGAAACAGAGTTGAAAAGCAGGGGATTTGGGAAATTAAATAATTTCCGCGACAGAGCCCAGTAACTAC

AGTCACTGGGAATACTCA

>NODE_989_length_228_cov_1.473430

CCGTCTTGTTTTCTTGCTCGACATTCTACAGAACGGTAAATCAGAGTGACACCTCCCCTACTCGCCACAG

CAAGACTGCCCCCTCCTGGCGGCATCAAGGGCAGCCTTCGCTATCACATACTTCAGTCCCCGGACTACCA

GACAATCTCCGAAGCACCCCATGCATCCCTTCAGATAGAAACCGCACTAGCATGCCAATAGACACCCAAC

CCCCTACTTACAATAATG

>NODE_990_length_227_cov_3.781553

CAGCTGCTGCCACAACTGCAATCAGCAGCAGTGGGATCAGGATGCCCGCGGAAGAGCTGTTTGCTTCTTC

TACGATGACCGGTGCTTCAATGATCGGATCTGCCAGCGAGCCAGCGGATGCGGAAGAAGCGGCAGCTGTC

AGAGCAGCGGCGAGAACGAGTTTTTTCATAGTATTACCTCCAGAAGATGGCGTGTCATGTTTCTCAGCAT

TACACACCTTACACTTA

>NODE_991_length_227_cov_1.684466

CTCCAGGTTCAATTGGCCAATGTCAAACACCAGGACGCGTATTTAAAGGCAAGAAAATGGCTGGTCATAT

GGGTGCTGAGCGTGTTACGACTCAAAACCTTGAAGTTGTTCGCGTAGACATTGAACGCAACTTACTGCTC

GTTAAAGGTGCAGTAGCTGGTGCGAAAAATGGCAACGTATTCATCAAACCTGCTGTTAAAGCATAGCCGA

GGAGTTAGTAATGGAAT

>NODE_992_length_227_cov_1.621359

TGTGGATGTGCGGTTTTCTTGTGGAGCAGAAGATGATTCGACCGCGGGGCAGCGGGGTTTGTGGGGGATG

TTTGTGTGTGTGTGTGCGTGCGTGCCTTTTTTTGTGGGAGAGATGAGCTCAAATTTTTTCCCACCCTCGA

CGATGTTGGCTCGGCTCGGCTCGGCAGCTCCAGAGCTATATACCAGGCTTGGGGCGGGGCTAGCAGCTTC

GTTTTCGTGGGGTGGAA

>NODE_993_length_227_cov_1.548544

TCGTGGAGGTGTACAGCTTCGCGGAGCTCTACCCCAAGGCTCTGCAGAAGATCGAGTTGTCGTCCAAAGA

GGCGGTGCTGAACGGTATCGCCTGGGACCCGGTGGAGGACGTGTTTCCGTACTGCAAGGATCTGGATCTG

GTGCTTGTCATGACCGTCGAGCCGGGGTTCGGTGGGCAGTCGTTCATGGCTGACCAGATGTTGAAGGTGA

AGGAGCTGCGGCGACGG

>NODE_994_length_226_cov_7.107317

AGCTGAGATGTACGGCATAATACCTAGAGCCAATACAGAGGCACGCTCCAGAGCACCACCAGAGAACATG

TTAAACATTTCTACAATGGTGCCCTTTTGTTGTTCAAAGAACTGAGCTAATACAGCGGCGTCAATACCAG

GGATTGGTACAAATGATCCTAAGCGTAGCACTACTAATGCAATGAATACGAACCATAATCTTTGTTTCAG

CTCAGACAAACCGCCT

>NODE_995_length_226_cov_5.502439

TCCAAGTTCCGCTTCGCCGTGTCCAATTTCCGCTTCCCTATGCCAAGTTCTGCTTCGCTTTATCCAAGTT

CTGCTTCGTTATGTTAAACCCCGCTTTGCCGTGCCCAAGTTCCGCTTTGCTATGCCCAAGTTCCGATTCG

GAATGTCAAGTTCCGCTTCGCTATGGCAAGTTCCGCTTCACCATGCCAAGTTCCGCTTCGCCATATCCAA

GTTCCGCTTCGCCGTG

>NODE_996_length_226_cov_4.039024

TGCAGGGTGCCAACTGCGTCGTGATCGAGAAGTGCTCGCCGGCCACCCGAAAGAAGCTCAAGATTCTAGA

CTCCGACCCTTCCTACGAGAGTTTGAAGACGAAGGTGGACTCTCTCTTGGCTGCGGGCAAACTGTTTGTT

GTCGACCAAGAACTCCTTAAGGACATGGAGCCGTCGCCTCTGGACGGAGAGATTACCCGGTACCTGACGC

CTAGCATTGCGCTCTT

>NODE_997_length_226_cov_3.034146

TTCCAAGGATCCGTTCCCCGCGGTGATATAGGTTTCGCTGGTGGATACACCACGAGGTAACGGGCCACGC

GCCCCGCCGCATTTCCTCCCCATCCTGCCTCCCCGTTTCTATGCTCCCAACACGAAGGATCAAACTATTG

TACCAGGTATCTAGCTAGCCCCTGCTTAGTTTCCGTCCTGGATAGAAAACTAACATCTATACAGTTATGA

GGGTATCCAGACACGC

>NODE_998_length_226_cov_2.736585

TTCACCTGCTTGTCCATGAGGTTTGCCCCGGACCACCTCTGGTGCGTCAGCAGGTCCACTTGTTCGTCTC

GCTCCGGCCTCTGCACCAGCTCGACATACTTGATCACGCCGTGCTTTCCGTACTCCTTAGCCAGGCGTGA

CACGATGAGCCCGTCCGAGTGAAAGCCTGCCAGGGTGATGAACTGCCACACGAACCCCAGGCGGCCGAGG

TCCCGGTTAAACGAGG

>NODE_999_length_226_cov_2.590244

GAGGAGGAGCGGAGGGCTGGGGGCGGGGTGCAGATGGTGGACGGGGATAAGATGACCGCCACGGATGACG

TTGAGGCTCAGAACGGCACACGGCCAGCCGTGGTGGAACACGCTGTTAGGATGAACGCGTAGGATAGCTA

CGAACTGCTGTAAGACTATATATACTGCTTGCTGTGCTGAACGCGTAGGCTAGCTAAGGCTTCCTGCTTG

CTGTGGTGTTGATGAT

>NODE_1000_length_226_cov_2.497561

TACATATATTCCCAAGTCGATGTAAGCAGGGAGCCGTCAGGTGGTTGGTGGCTGTGCTCACTCACGACAG

CCTAGGTTTGGATGTTTTGTGCAATTTATTTTGGCCGATATCGTATACCACTTGTATTGTGAGCATAATC

GGCCGGGTATGGGAATTATAGGTGCGGGACGGGGCCAGGGGTTGGCCGGACGGCAGTGTTGGAAATTTTG

CTTGGTTGGGTGATCT

>NODE_1001_length_226_cov_2.458537

TTTTCGCGTCGCCGCTGCGCTCGATTCTTTTATCATGTTGTTTCAGGGGATTGGGGCGGAACCTCTTTGT

AGTTTGTATTTGCGACGTTTTCCGAGTTTTCGGTTCGGACTGTTTTTCTTCTCTTTATCGCTTACCACTT

TCTGATAATTTTTCTGGGGCGGATTTGGGGGTAGTTCTATCGTAATTCGTCCTGCCGACCCCCCCCGCTC

AATGCCGGGGCTGGTC

>NODE_1002_length_226_cov_1.619512

TCAATTTTGTGATGTTATCAGTTTGCTAATGAGCTCCCGAAGGGCGAGAATTTAAAGGCTCATATACCGC

GTTATTGATTTTGACAAGGTTCCTACATGGATGTAGGACATTAAGGTAACGCAGGAGCAGTTACCGAGAA

TAACCATTCTCGGCTTTGTCTCATGAGTCGCTCTACCTCCACCATCCATGGTGTCGTCTTCAATCAATGC

CTCGCCTACATGGATG

>NODE_1003_length_225_cov_17.642157

CTGGAGTTCCTTGGCACGCGATATTTCCAGTTCTGCTCTTCCTGCCATGGAGCCGTCTCTCTGGATAATC

CAATTCCAAGGCGATAGGCAGTTAAAAAGGAAAGACAATCCCTCCCAGGGTTCATTCTGGCGTCTCCGAG

CTTGTTTGCGTTACCGCATAATATCCTCGATCGGCTACAGGAATATTAACCTGTTTCCCATCAGCTACGC

ATCTCTGCCTCGCCT

>NODE_1004_length_225_cov_9.269608

ACCACCGTTTGATTGACCAATCAACGGCAGAAATCGTGGAAACTGCGAAACGCACTGGCGCACAAGTACG

TGGTCCAATCCCACTACCTACACGCTTTGAACGTTTTACTGTACTAACTTCACCGCACGTTAATAAAGAT

GCGCGTGATCAGTACGAGATCCGCACCCATAAACGTCTGATAGACATCGTAGAACCAACTGACAAGACTG

TATGGAATTCTCGGT

>NODE_1005_length_225_cov_9.151961

TTAAGTTCCGGAGTTTGCCTCTCTCTCCCCCCATTCCTCGTCGAGTTTTCTCGAACCTGTTAGCCGTTTC

CGTCTCTCGAAGAGCGTGATGCGTTGCTAACGGGCTGGGGATTTTGCGAAAGCGAGGTTGTGGTGGAGAC

TGTAACGGTTGTTTTGTTGCGGATAAGGGGTTTCCCAAGACGACTTGTCTGCTGCGCTGCTTGGGGGCCG

GGGCCGGCCTTTGGC

>NODE_1006_length_225_cov_6.230392

ACGGAATCCAATTCTCAGCGAAGCACCCCGTTATGCCCAAAAGCCAGACCATGCCAACCTCACTCATCAC

TCACCGACTGGTCGCCTCGAAATGCGTGTCTGAGAAAAAGCCTATACAACGACTATAGACCTATTACTGC

TACGTACCAACCCTCACATCATGACATTTGTCCCGCGCCACAGACATAATAACCAAAAAATGAAAAAACG

ACAACTCGTAGACAA

>NODE_1007_length_225_cov_3.406863

CGGTTGAAGGCCCCCTTTACGGGCGAGGGGTCTTCGTTGGGCGTGAACCACCACGAAGCCACGGAGCCGT

TCACCGTGCACGCCACAACGTTCCCGATCACGACGCAGCCCCAGCTCATGCTCATGAGCCACACGATCGC

CAAGGGGATGCCGTACCCGCTGAAGTCGCAGCTTTCATCCGAAATCTTGTTGCCGTTGCACATGCACACG

GGCCCAAGCGCCGCT

>NODE_1008_length_225_cov_1.686275

TTACAGTTTGTATTTTGTATTGGTGTACATTGTATATATTCTGGAACCTCGTTTTTATGTTCTACTTGAA

CCCCTTCCACACCTATTGATAAGACGTGGTGCATGGTAAATATGCGCGTGCGTTGGGGTACGCTGGACGT

GTTTACAGTAGCTTTGAAGGTAGGCATGGGCGTGTGGGGTAACAATGCGTACCTGCTGACGGACGAGTGT

GTGGCCGCTGTAGCC

>NODE_1009_length_225_cov_1.127451

AGAAAGAGCCTTTTCGTCTTCGGACGACTAAATCCGATCAGCTGGAGCAAGACCCACGATTCCGGCTCTT

TTGCTGAAGAAGCTCGTCGAACTGGGTTCGCTTCTTGGTTGGCCCGCGAAGATTCGGTGGTCCGAACGCT

TAGCGTTGGGTAGGCCCCCGGCCTGTCAGCAAAGCCATAGGTCCAGAAAGAGGAGAAGCTCCAGGGAGGC

ATGTCGATATTGTTT

>NODE_1010_length_224_cov_16.029557

CTGGAGGCGTAGACTTGTACGAATAAATAAGCAGCCCTACACGCTTGCTGCTGATGCTGATGTTGGTTCA

TACCTGGAGGCGTAGACTTGTACGTCGGGGGGGGGTGATATTTTTAGGCCGAAATATTTTCCGACTTACT

GCGTATTACTGACGTATTAGTATACTCCTGAACTGAGGCTGAGGAACTACCACCCGGTGTGGTATATATA

TAATATATATATAT

>NODE_1011_length_224_cov_15.137931

CGCTTTCGTTCCTCAGCGTCAATATAGTTCCAGTAAGCTGCCTTCGCTTTTGGTGTTCCTCTAAGTATCT

ACGGATTGCACCCCTACACTTAGAATTCCGCTTACCTCTCCCTAATTCTAGATTAGCAGTTTCAAATACA

AACACGGAGTTGAGCTCCGGTCTTTCATGTCTGACTTACTAATCCGCCTACGAACCCTTTACACCCAGTG

ATTCCGGATAACAC

>NODE_1012_length_224_cov_9.857143

AGAGTTCGAATGAGAGTCGTTGCCGCAGAGATGAAAGGGAACCTGATGGGAGTTGGCCGGCAGCAGCAGG

AGGGACTTCATTGCTTTCGTATCTCTGGCCTCCTCGCTCTCCGTCATTCCCAAGAGCGCGCGCATGCACT

CCCTCCTCATTCCAATCTCCTCTTCCTGCTCGACATCCTCGTCCGCTCCGACCTTCTCCTCTCCGGGGAG

TTGGTGTCATGCGA

>NODE_1013_length_224_cov_8.926108

CGTCAAGGCACGCCCGTCAAGGTATGTTCCTTGACCCCGTCCGTCAAGGGGTAAAAGTGCCGTCTCCTGA

CATCTCTGCTGCTTTTTGTTTTTGTTTTTTTATCCTCCGGCGGATGGGTAAAAATAAAGTGTAAGCCTTC

CACACTTCCACGTCTCACTTCGAATACTGCTGTATGGTATATCAACAACGCCCCCCTGTCCCTGCCAAAT

CACCACCGCCGGAG

>NODE_1014_length_224_cov_8.197044

CGCCGCAAGATGCGCCAGCGCGCGCGTTAGAGCGGTCGTGTGGTCCTTCTGAATAGGAAGGGGTTCCACC

GCTTGCGACCGGGCTGAGCTACTCACTGCTACTGCTGCTGGTGCTGGTGCTGCTCTCATAGGCGTGTCAG

CTATTATATAGCTACAGTAGTTGTCTTGGACGTTTTTTTCTTGCGAGAGGAAATATCGCGTACCCCTTTT

TCTAGTTTTTTTCG

>NODE_1015_length_224_cov_5.729064

TTTGTGGCATAGGACCATCTGTAGCAGCAACTACTAAGATAGCGCCATCCATTTGTGCAGCACCAGTGAT

CATGTTTTTAACATAATCGGCATGTCCAGGACAGTCTACGTGTGCGTAGTGACGAGTTTCTGTATCGTAC

TCGATGTGAGAAGTATTGATTGTAATACCACGTTCACGCTCTTCTGGAGCATTATCGATTGCAGAGAAAT

CACGGATGTCGCCA

>NODE_1016_length_224_cov_4.788177

CCCTCCAGCGCATCGAGTTCGTCGTCCAGACACGCCAACTCCGCGTCGAGGTCATCCTCGTCAAGCTCCT

CTCCGACGTTGTACGAGCGGCCCATGATGTCGTTGATCTCGTTCATATCCTCCATCATGTCCGCCATGTC

GTCCGTGAGGTCCTCCACCTTGTTGATGTCGATCTTCTTCACTTCAGCCTTGAGCGTCTTGCTCGCGTCT

TTCATCGCCGCGAC

>NODE_1017_length_224_cov_4.778325

TGTCCAGCGTCAGCCAGTTCCACTCCATCACCACCACTTCCGGCACGAACGGTTTCTCCGGGTTCCAGAT

TGCCCCTCGCTACGCCACCTTCGTGTACTTCAACTGCGTCACCGGAGACTGCCTGGTGGCGGAGAACGAG

TACATCCGTCGTATCGTGTGTGGCTGCTCAAACCTCAACCCTGCGGAGGACAAGGGGTCTTGCAGCCTTT

GCACCGACATTCAG

>NODE_1018_length_224_cov_3.413793

GCGGCGCCGTTGGAGCTCTTGAGGTAGAGCACCGGTCCTTCCAGCTCGCGCAAGAGCTGGTACTTGACTC

CGTTCATCCTGTATCCGTGGTCCGGCCTGTTCGAGGTGGTCACGAAAGCAAGGGTGTCCATGGCCTCGTT

GATGTCCTTTGGCCTCTCCATGCCATCCTCGCACATGACGGTCTTGGTGTAAGCTCGGAGCTGGAAGTCT

CCGGCATTAGCGTA

>NODE_1019_length_224_cov_3.246305

ATGGCAATGCGTACCACAAGCAGGACGGTACAGTGTACTGCTGTGTCGGATTTTTTATTTTTCCGGTGTG

GACGCCCAGTAAACGAAGGGAATGTACAGGCAGTAGGGAGACACCGACGACTTTTGTGTTTTTCCCGCGA

GTAAACCGTATTTGTTTTTGCTTCTTCGTTGTTGTTTTGGTCCCCCGGCGGATAAACTAGGTTACAAATA

TACCCACAATAGGT

>NODE_1020_length_224_cov_2.694581

GTTGTTGGCAGATAACCCCGACGAACTTAGGGGTGTATCTCGTCAACAAGCCAGTTTCACTTAGGATCTC

CTTTCGGCGTGGAAACGGACACATACATCTACTGCAACAGCCGACGAGAGAAGGGTTTCGCTTTCCTAAA

GTTCAACGTGGGCTAAGTTTGCAAAGTGGTCAACCAACCGCTAACCATCGGGGGGTTGAGTGGATGAACA

TCGAGGGGTGCAGC

>NODE_1021_length_224_cov_2.546798

AACATAGGCACGTTTCGGGTATCACCATGAATACCCCAACGACCCCTATATTTCCGGACATTGAACCAAT

TCTTATATTTTTCCCGGCGCACGGTCCCTAATATTTTCTGGACAGAGTGTCACCATCACCCATGAACCAA

CTCATTCGCGAACGAAGTCGAAGCCCATGAGTTCCCGTTTGAACAGAACGTCCTTCTCCGCTACGTCCCT

CAGCTCCTGCTGGT

>NODE_1022_length_224_cov_2.334975

CTTGACCTTGAAGGACGTCAGGTTGTTGGGGTTGGGGAAAGTGATCTCTGCTACGCTTCCGCCATCCAAG

TCCGCAATATCCTTTTGAATACGGATCTCTCCTGGTGTTTTCTTCGAGCCAGTTGTCTTGGCACTACCTC

GCTTGGCCCTTCGTCCTCCAATTCCCAATACGTGAAGCTTCCCGCCGGCCTTCTCCTCGCCCCCCTTTGG

TTGAGCAGCTGCGT

>NODE_1023_length_224_cov_2.221675

ACGACAAGCTCGCCGAGTTCATCCGTTCGGCCATTTCGGGGGACATCACCGAGGTGCCCGGCATCGCGGC

TAGAGGGAAGGAGATCCTGGCGGAGGGAGAGGGAGACGACTGCGTCACCAACACACACCAGCTTATCGGA

AAGTTCTTGTCTCTCCGCGGCCCCGACAAGGACGATCACGAGGTGGACTCCGTGGAGCACTGCGACAAGT

TCTGGTACTGGTTG

>NODE_1024_length_224_cov_1.940887

CGGGGCGAACGTCGTCCCCGTTTCGCCGATGTCAGTCGGTGCCGTCATGAACGGCAGGAAGAAGAAGAAG

GAGAAGGGTCGGGCCGGCTGTCCTTTTCCACGGGAATGGCCGACCAGGAGACGCTGCGTCTGTTCGAGTC

CTGGAAGGAAGAGTACGGCGTGTCCTACCCGAGCGCGGAGGAGGAAACCAGGCGGCTGGCGACATTCGAG

GAGAGCCTCAACGC

>NODE_1025_length_223_cov_107.975248

CCAGAATGTGGGGCAATCGTCCAAGGCCTGAGCCACGAGTAGCGTCGCCGTCGACTCTTCAACCAATGGC

AAGCAATGCAAGCCAAAGGGCAAAGAGGAGGCAGGCGCAAAGCAAAGGCCTAACTATACGTTCGCTCCGG

CTTCCGTCTCGCATTCGTTAACAAAAATTGTTGACCGCTCAGGACCAAGGGATGAGGCCAAGACGGAGAT

ACCGATATGCTTG

>NODE_1026_length_223_cov_8.896040

GCCGTGACCACGGGGGCGGACATTGACACGGACGAGTTGGTCGACGAGACGGGAGTGGACCCCAAGGACA

TCGAGCTCGTCACGCTCCAGGCATCATGCTCGCGGTCTAAGGCAGTCAAGGCCTTGAAGAACAACGACGG

CGACATCGTCAACGCCATCATGGAGCTCACCATGGATTAGAGACAGAGACGCAGCAGAGTCGAACGGGGA

GGGGGAGGGCCGG

>NODE_1027_length_223_cov_5.292079

GGATGGTGGATTTATTTATTTATCTCCACCAAAAGTGCGTGTTTTGTTGATTTTGTTTTTGTGACGGCTA

AAGGAGAGAGGGTTGTGTTGAGGGCCGAACTCGCCGTTTTTCCCCGCATGATCCCGATCGGAGGGGGCTG

CCCGAAATTAATTTATTTTTATATTTTATTTCGGTGGTGCCCTCCGATTGGTCGGGAGATACGGGCGGCG

ACCTCGGACATTC

>NODE_1028_length_223_cov_3.262376

CCACAGTCGATGGCTACTTCTCTCAGGTCGCGCTCTGAGGAGCCGCCTCTTCAATGTATATACTCCATAG

GTTCAGTCCCGCCCGGCCGACCTACGTATCAATAAACACACAGCTCGCGGTCAGCGTACCCAAACTTCGC

CTTATTCTCATCAAACAGCCGCTTCAAAGCGTCCATGTAGGCCCCGTGTGCCTCCCGCAGCTCCTCGTCG

CTAACCCCTCGAA

>NODE_1029_length_223_cov_3.000000

GGGCGGAGACGGCGACTTTAGAGCAAGCAGCAACAGGAGCATTTACTTAAACATGACACGCCTTCAGCTA

CTACCCATGTTGCTCAATACGTAACCGATGCATATATATATCTATATATATCCGGTCATCGGTTCGCCAG

CTACCCATGTGCTCGATATATAGCCCCATGTCCATCAGGTGATCGATTCGCCAAATTCACTGATTCATCG

AGGGCTGCCGTGT

>NODE_1030_length_223_cov_2.896040

CATTTCTCTCGGCGTTTCACTCCCCGGCCACCGTGGACCACCGGGTAGTACGCGTCACCAGCATGATCAC

GTTCTTTATGGCGCTATTCGCGGGGGCCTTGGCCTGGCGCGACGAGGCCCAATGGCTGGTGCTCGGCATG

TTCGTCGACTGCTCTCTCAAGCTCGGCGGCTTCCCAAGCCCTGTTGACCTCCTCGCAGCCCTCCCCTTCG

CCGGCTCGAAGCC

>NODE_1031_length_223_cov_1.193069

CGGCGGGGCAGCAGTTTCGACCACTACTACTACTGAGGCGTGCATCCTGTACAACTCCCTACGGGGTGCC

CGAGTGCTTGCCTTCTCTGGGGAACTGCTGAGAACGTGTTTCACAGTTCAAGAAAACAACAACCTGACAA

CATCGAGCGCTCGTTTGTGGCAAGATGAACACAGGTTCGGCTGTCCAAGATCAGGAAGTGCTACTTGACT

TTTTCCGGTCGAC

>NODE_1032_length_222_cov_13.104478

TTGGCCCCGAGAATTCCACCACGGTGAAAGGGCCCTCCAAGATGTGGGCCAGAACGGAGAGCAGCCTCCA

CGGGTCGGTGCGAGAAGGGCGCATTGCTGCGCCACAGCCGCGGTGACCGGAGACCGCGCCTAGTGTCCGA

ACACGAAAAGGTTTAATATCCCAAAGCACCGGCTTCTATCACTCATGGTCAGGAGTGAAGAGAATTTGAC

AAGGTTTGGGTT

>NODE_1033_length_222_cov_3.960199

CTCAGCCAACCCAGCCAACCGTTAGCAAAACCACACATGTATCCTTGATTATCACCATTGGGAAGGCAAG

GCAAGGTTATCATACAGCGCACAACAGGAGGAGCACTTTTTCCTTATTTTTCATTTTTTTTTGTTCTCTC

TATCTCTCTTCTCTTCTCTCTACAGGAAACGCATAGCGATGGGGCAGTTGACGATGATGCCGCTCGACGC

GTACGGCGCGCC

>NODE_1034_length_222_cov_3.741294

CGAAGTTCGAGTACAAGGCGAACGTCAAGAAGGAGATCGGGATGATCGCTGGCGGCACCGGCATCACGCC

GATGTTGCAGGTCCTGCAGGAGCTGCTCTTCAACCCGGAGGACAAGACCAAGGTCACCCTGCTCTTCGCG

AACCAGACGCCGGCGGACATCATGCTCAAGTCCGAGATCGACAAGCTCGCGGCGGAGCACGACAACTTCG

AGGTCATCTACC

>NODE_1035_length_222_cov_3.487562

CGAGCATGGCGTAGTACTCGATCTCGGACTTGCGGAGGGGGGGGCAGTTGTTCGAGATGAGGACCATCTT

GGCGCGGGTGCAGCGCATGGACTTGATCGTCGACTTGTACCCGAGGGAAGCCTTTCCGGACTTCATCACC

AGCTGGAGCCGGGCGTTCATCGACTCGATGGCCTTCTTCGATTTCTTTCCAGGAGCCATGTTTGTCGTAG

GGTCCGGGTAGG

>NODE_1036_length_222_cov_2.960199

CCTATATTTTTAAATTTAGCGTCATTTAACTTAACTAAGAGATGGCGCAGTACGGAGTAGCTATGTACGC

GGTATTCCAAAGCGGTGGTAAACAGCATCGTGTAACCGAAGGACAAACAGTTCGCCTAGAAAAGCTTGAG

CTTGAAGTAGGCGCAGCAGTAGAATTTGATAACGTTCTTATGATCGCCGATGGCGAAAAAAAAGCAAAAA

CCCACCGTCCTC

>NODE_1037_length_222_cov_2.935323

AAAATCGGACGAGCAAAAAAAAAAAAGGAAAAACTGGAGGTAGGTAGCCTACCTTTTCATACCGCGCGCC

AGGTTTGGCGAGAAACTATGCGTCGAGGGTAGAGAGTTAGAAAAGGTCGGGCTGGCGGTCTTCTGACTCG

CTCTTGCGTTTTTTGTATATAAGCAGAGCGGTACGCAAAGCTAAGCATTGTCGTTTGTTGGAGAGCGAGA

GCTTTCGAGGGG

>NODE_1038_length_222_cov_1.830846

TGAAGGACGTGCAGGATCTTGCAGACTTGCTTCATCGCTTCCCTGCAGGTCATCTCGTCGAACTTCAACT

TCTCCATTTCCGTCTTCGCTCCCTGGCGACCCTTCCCGACCGCAGCGCCGAAGTACCGGTACATCACCCC

CGACGGTTCGATCACGTACAGCTCGTGCTTCTTCAGCTCCACATCGTACCCGGCGACAAGAACTGACGCC

CCAAATGGCCGG

>NODE_1039_length_221_cov_15.345000

ACTCAGTACCTTCAATCAAACCATTTAACTTTACCACGTAAGTGTTGTGATGTTTTCCGTAATGAAATGA

TAAAGTTTCTGCAGAAATATGTGGTTCCAGGGCATCTTGTTCGTAAGGTAATGCAGGTAATTCAATAGCC

ATTGTGCTCTCCAAGTTCTTTGTTTTAAAGTTACAGTGTATTCGTGAATAAAGGGTGTATTTTCCAATAA

AACTTGAGTAA

>NODE_1040_length_221_cov_9.095000

AACCTCTCTCTGGTGTCCTCGCAGCCTCCTCGTGATCAGCATCACAACGAGAACATAAAGAGCCGACACA

AACGCGCAAAGCAGCACAAGGCTCAGCACAACAAAGCTTCATCGGCAAGAACACCGACAGTCGGGAAATA

ATGAAGATCATGAGTTTCGCCGCCTCCATCCTCCTCGTCATCGCCTTCCTCCTGTCGGCCTCGGAGGCGG

GTCCCCTGGCA

>NODE_1041_length_221_cov_2.570000

CGGCCTCGATGCGCGACGTCGGCTACGTGACGGAGGAGAACGGGGTCATGGGCGTGCGCGCGACCTTCAA

GAAGCGCTACATCACGCTCGCGCCCGTGGCCGGCATCGTCGGCCTTGCGTTCAACCTCAAGGACCCTAAC

AACCTGCTCAAGGGCGACGGCAACGAGGGCATCACGGTGTGTCTGTTGGAGAGGGGGCACCCTGGCCTCC

GCCTCGGTGAC

>NODE_1042_length_221_cov_1.965000

TGCCAAAAAAGGGGAAACGGTTCGTCGATCATTTGATTCATTCGAGACAATGCTTCAAGATGGCTGGAAG

CTTAAAGATGGCTGGAAATCTGGTATGGCTATGGGGGAGTGGCACGGCGTTACCACCAACGCGGAGGGCA

GGGTGATCGGCCTGGATCTTCGCCATAACGACCTCAAAGGCATCATACCCCCAGAGCTGGACGACCTTGA

TGCGCTGGAGC

>NODE_1043_length_220_cov_16.447236

GCGGGAAAGGATGAAACTCATGAAATTCGGACGCAGTGACTGCCTTCCTGGCCTACGCGCGAAGGAACTT

AAAGGGGTCGGGTATGTGGGTTTGCAGGGCTGAACAAGCCTTGCCGTCCCTCCCCCCTTCCCCACCCCAG

CGTGTTTTGCCAGGGGGGTGTCTCGGCACCACAAAACCATGACGGTCTCTAAACGTTTACTAGGGCTGAT

CCCGTGGAAT

>NODE_1044_length_220_cov_5.944724

CTGTTGCCGCCGTCGAGACCATCATTATCTCCATAGCCGCCAGTCATCATCGCCATTTGGACGATGCCAG

CGAAGGCGATGAGAGGGATCGTCGCAATTGCCAACAGCCCGATCTGCCAAGAGAAGGCCAACCCGATCAC

GGTACCGACTATAAGCGTCATGCTGAGCTGGGTGGCGTGGGCGAGGTTGGTACCGGTGGCCCGCCGCACC

GCGGAAGCCT

>NODE_1045_length_220_cov_2.276382

CCAGACCAGGGCGTAATAAAGCGGAATCTAAAACATCAGCTCTGTTTGTAGCTGCAATAACAATTATACC

ACTTGTAGGCTTAAACCCATCCATCTCGGTTAAAATTTGGTTTAATGTTTGTTCTCTCTCATCATTAGTT

CCTCCAACACCTGTTCCCCTTTGTCTACCGATTGCATCAATTTCATCAATAAACAAAATACATGGAGAAT

TTCGCTCTGC

>NODE_1046_length_220_cov_1.944724

CTATCTTCGGCGACGGTCTGGTTCTGCGCGACAACACGCTTTGGGTTGTCGACAATGGTCAAAACGTCGG

CGGGCTGCAGCAAGTCACGGAGATCTCCCTCTCGGCAGACATGACGTGCGGCGCTGTTGTGCCCCGCGCT

TTGACGAGCGACCTGTTCGACACCCCGACTACTGCGGCACTAAAGGGCAACTCTCTGTACATCGTCAACG

CCAAGTTCGG

>NODE_1047_length_220_cov_1.592965

GCAGACGGAGTACGACTTCACGGGCGATTACAAAGTCGTCGGCAACATCATCAGCAGCCCGCGATCGTTT

TTTACCGCAGGACAACTGGTTCGGCTACGTGGCTCTCTCTACCGACCAGAAGGAGCTTTTGATCGTGTTC

CGCGGGACCCAGACGACCAAAGAGTGGCAAGAGAACGCCACCGGGGTCATGATGCAGCTTGAAGGCGAGC

CGGAGGTGTC

>NODE_1048_length_220_cov_1.396985

CAGTGCGGATCGGTATCGGGCAAGGGCGCTTTTGCAGCCTGAGCCATGCCTTTACAGTGCGGTAGGGGGG

GCACACGTTTGGGCCTAAATTGTTCGGCTGATGGAGCGTTCCACACTCCTTGATTTTCCCACGAGATATC

TGTGATTCCATGGCAGTCCCACGGGATGAAACTCACCCACTCCGATAAAAACCACAACACTCAACAGCAA

TTCTCCCGTC

>NODE_1049_length_219_cov_67.444444

GGCACCCCGAGAATTCCAAAATATCCTCGAAATTCCTATAACGATCGCCCGGAGGTGTACAGGCGACGGT

CTCAAACATTCGGGGACAGCCAAGTCCAGAGACAAGGCTGTTTGATGTAGGCAATGCGTTCGTTTTCATT

AGGAGGTGATCTGCATACGGTGCGGCTTTTGCAGAGCCAACACAGTCACCGCTGCTGCTGCCATGACGTC

GGACTTGTA

>NODE_1050_length_219_cov_43.323232

CCAGTACACCTCTGTCTGCTGGGCGGCCAGCCTTTCGGAGAGGAGGCTGTCGGGCGTGGTGATGAAGACG

TTGTCGCTGTCGGCACCAGTGACCTGGTGCGCGGTGCCCCCGAGCGGGTTGGTCAGCTTAGCACCACCAG

GTCCGGCGGCCCCAGAGGGGACCTCCTCGCAGACGTTGATGTCGCTCGTGAACACGCAGTCAAGCAACTT

CTGGTAGTC

>NODE_1051_length_219_cov_15.313131

CCCAAGCAGCCAGGTAGGACCAGAGTCACCCCTGCACGGCGTAACCATGGGTGGAGTTCAGCCCATCAAC

AAGGTCAATGTGGTCAAGAAGCGCAACAAGAAGTTTGCGCGGCATCAGTCGGACCGGTTTATGCGGATCA

AGAACTCGAGCTGGAGGAAGCCCAAGGGTATCGATGGACGTGTGCGCAGGCGATTCAAGGGTTGCCTTCC

CATGCCCAA

>NODE_1052_length_219_cov_4.702020

ATGCCGCCATAGTCAGAAACTTAAAGCAGGCCACCATGATGTTGATTTGAGCCGGGCCAGTGGGTGCAAG

AAGACGGGAGGGGTGGCACGGCGCTGCTCTTCTCGTTGACGATGACGGTACCCTTGCTTGTAGAAGGGAG

GGTGGGTGCTCTTTTTGGTTTTATGAAATATCTTGATGAGCTTGGTGGGTTTGTTCTTTTTGTCGATGGT

CCGGAGCTT

>NODE_1053_length_219_cov_4.353535

TTACGGATTTAGCTCCTTCTGTCTGTCGCCGAGAAATTGCAGCACTGATTGGCTTCGAGGGATACGTATA

CTAAGTGGTTGGTTGAATGGTTGATGGACCGATGGGATTGGTTGGATTGTTGGTCTGATGATTGGCTGGA

GGCTCGATCGACTGATGGGCTGATTTATCCACCGAATAGGCTGTAAAGGAGCGATGCACTGATTGGCCTA

GTATATACA

>NODE_1054_length_219_cov_4.131313

CGGGATCCGCTGTGCTGCTTTTTGACTCCATGTGGTGCATGTTGACTAAGAGCGGTAGGATATGAAGACA

GTTTGTTCGTTGTGGTTGGGTCTTTTGCGGCATGAGTGGCTTAAGGTTTTCATTAAAGTTTTGATAGGAG

GCGTGTTTTCCGTGGAGTTGAAAGGATTCGTTCCGAGAGTACCGTGCACATGGCGGCACCCTTCGGAAAA

GCACGTGTC

>NODE_1055_length_219_cov_2.272727

GACTGGACTTTGAACATGCGAATCGTCGATGACCTGAACCGCGAGAGGGACCCCGTGGTGCTGACCGAGG

TTGTTCGAGTCCTGCGCAAGAAGCTGGGCTCAGGAAATTTGCGCGTTCTGTCGGTTGCTCTCACCCTCGC

CGAAACGCTGGTGAAGAACTGTGGAGACCCCGTGCACCGAGAAATTGCGTCCGAGCGTTTCATGCTGGCC

ATGGCAAAG

>NODE_1056_length_219_cov_1.868687

ATTCCAGCAATGCTGCGTGAATCCGTCTATAGTCGAGGGTTTCAACCCGCCGATCCCTTCCCCAGCACAC

GAATCTTGAGCGCTGTTCTCGGTGGGAGGATGCCGCAACATCACACAGCAAAACGAAACAAAACAAAACA

AAATTATTATTTAAACCATCACCTCCAGCTTCTCTCAGCTCAACTAACTCTCGTCTACAGCGCCGGCCAT

GACAGACGA

>NODE_1057_length_219_cov_1.797980

AGTCGATCTGCAACCATCGTGACAACGTGGGCATGCTTGGCGCCAACGCCCCTGACCGGGTGAAAAAAGC

ATGGAACATCCAACTCGGAGGCAAGGTGATTTACCCCAACAAGATTAGGTGGGGGCATATGATCTCGTCG

CACTACCTCACCTCAAGGCCGGAGACTACAGCGAATCTAAGCTGCACAACTACATGAGCACGCCGGCGAA

CGCCTCGGG

>NODE_1058_length_218_cov_10.497462

GGGCGTAGTTGTCGAACACCGAGGCCTCGCGGATGTCACGGAGGGACGAGGCGTCTACGATGTTCCTGAC

GATGAAGCGCTTGATGGCCTTGTCCTTGGGGATCGCCTTAGCGGTGGACGCGCATCGCACACGCACGACG

TGTCCGCGTCCCTTCTTGTTCCTCCCGTGGTTCCTCCTTTTGACAGTCATCCTGAGGAAGGCTTGGGTGT

GCGGCTGC

>NODE_1059_length_218_cov_9.050761

GTCCAGCACGCTCGTCTGCGTGGTGAACGTATCCTTGAGGTATGCACCGATGTCCGGGCGTCCGAAGCTC

ACAGTCACGGGGTCGGTCGGCTTGTTGCTGATAGAGTTGATGCGACGAAGACGTCGAACCGAGGCCGTCG

AGAGGCCGATTGCATCCGAGATAACCGCCTCCCGCTCTGCAGGCAGGTGGAAGTTGACAGAAGTGTCGCC

GTAAACGG

>NODE_1060_length_218_cov_7.695431

GCGGGGTCTACCATTAATACTACCCAGTGTCGTAGGACGGGGTGGTACTGGGTTTTTGTGGAGAGTACTG

ATGTGATTGCAGAAGTGGAATCGGGTTATTAAAAAAAAAACGACACCATAACCGGCGTATCGCAAACTGA

CCTATATCGTGGGCTGACCGGTTCTTAAGCTGACCAATCTTGAGCTTGCCAGTCGCGAGCTGGCGAATCT

TGATGGAA

>NODE_1061_length_218_cov_5.918782

ACCTCGGTGAGACGAGCCCTCTCGTTCGCAAGCATGGCAACCCGTCCAGTACGCGCAGCCCAAGCTTCAT

AGTTCACGATGCGTTGAACCAGGGCGAAGCGCTCGGCATACAGCTTGTCGACAACTCCCGCCAGCTGCAC

CGTCTTCCATCTCTCAAATTTCTCGGCATCCATCTCGACCCGGAAGAAGCCACCCTCTCACTGCCTGATA

CCCAAAGC

>NODE_1062_length_218_cov_4.928934

CTGGCTCTTCCTGCCATCGGCTCCAGTGCTCTCGTTGACGCGAACACCAAGACGGCTATCTCCACGTGGG

CGTGGGCTCACGACGAAAAAACGCTGGACGCTGGGGGCTGCGACCCGGACGGGTGCACCTCCGAACTGAC

CCTTGACTCCGACACCCACCCTTCATCGCGGTGGTCGTGCAAGTACGACCTCGCGAACAAGCCGTGCAGG

CTCTTCTA

>NODE_1063_length_218_cov_3.451777

CCCTCCTCGAGCCCTTCCCCTGACCTGGCGAACAACGGCAGCTTTACGCCTACGTTCGACCCGCATCTCC

TCACATCGCAGTTTGCGCAGCAAGGCAAGGCTGGTAGTCATGATTTTGTAGAGGGGGATAAGAGGGAGGA

ATGGGTAGCGGATTCTGGTGCTACTTTTCATGTGACTGGTAACCCCTTTGGTATGGTAGACTGTTCCACC

CCTTCCCC

>NODE_1064_length_218_cov_2.796954

CAAATCAACCTGGCGGTAGGCCTGCGAATGCTCGCGGGAGGTTCCTCTCTGGGTGTCGCGTCGAGATTAA

GAATTCACACTCCTCGACATACACCATCATGTGGGAGGTCGTCGACGCCATCAACAACACCCCTGAAGTC

GGCCCGTTCTCCCCTCTGCAGACAGTGGCAGAGGCCCAAAGGCTTGCCACGCGATCCATGGTGAGGTTAC

GTTTGATG

>NODE_1065_length_218_cov_2.751269

TTTCTCGGTGCTGGGTTTTGTTGTTGGTTTTTGCTGTTGGTTTACTTTTTCTTTTGTTTTGTTTTCTTTC

CCTGCTTTTATTATTTTTGTTGGTTTACTTTCTTTTATGGTGCTGTATTTTTTTATTGACATGTGTTCTG

GTGGGGATATTTGCGTTGATTCATATCCATACGAGTAGTGCATCCGGCTCCGGGTGGATTTCTATTTGTG

TTGGTTAG

>NODE_1066_length_218_cov_2.238579

AGTCTCAAGGTCTGTAAAGGCTGGTGCAGGTCTATGGATAGGTAATGTTTCGTTACCAGTATCATCTACT

AAATTATCTACAGTTTGTCCTAAAACATTAAAAATACGGCCAAGAGTTGGTTTACCTACAGGAACTAGGA

TTGGAGATTTAGTATCAATAACTCTAACACCTCTTTGTAAACCATCAGTAGCACTCATTGCAATAGCTCT

AACTCGGT

>NODE_1067_length_218_cov_1.878173

GTCGGCCTTCCGCGACCTTTTCTCGTGGGCCGCGCCGAAATTCGTCTGCTCCTTCGGCAGCCGAGAGTTT

TGCGACTTGCAAACGCAACTGTTTATGGAAGAGGTGAGGCAGCAAATGTATCTACGATTTCGAGACGGGA

TCGTCTACGGGCTGGGGTACGAGACCATCCGAAAATTGTGCAAGATTTCTGGCAAGTACAAGCTCAACGA

GAGGTCCA

>NODE_1068_length_218_cov_1.629442

CACCCACGCCATGAGTAGAGCCCCGTACTAGAAGGTCTTCTTCCTTTCTATCTCAACCCTTCCCAGCCTC

TTCTTGTACTCGGCTACATAGCACCCGGTCGACGCAAATAAAAACCTTCCACCATTATGACATGGAAACC

CCCTAAGGGCGTGTTTAGTAGGGTGTATAGCCGGGTATGACCAACGCATCTACCCCTACCCGGGTATGGT

CAGCATAA

>NODE_1069_length_218_cov_1.385787

GAATGCAGGTATATCTGCTCCGGGTGCTGGTTTTGGACGTGATGGTGATGCAACACCGATTGTTCGTATT

GAACCTAAATATCCAATGCAAGCAGCGCGTGACGGTAAAGAAGGTTGGGTTAAGCTTTCATTTACAATTA

ACGAAATTGGTGGAGTTGAAGACGTAGAAGTTATTGAAGCTCAACCTAAGCGTGTATTTGACAGGGAAGC

GAAACGTG

>NODE_1070_length_218_cov_1.340102

TGTCATGTAGATCACCGTTTCGAAAGAAGCAGGGCTTCGAGAGCCTTCAAGCCCTGTCGCTGTCTGATGG

TGTGAACTAAAGCTTCTGACTCCGACACGAAATTCCAGTGAGACCGACGGATCTACGATGTAAAATGATC

GACGCTCTTTGCAACCCTTGATTAGGTAGTAAATTAGCATTACTGTAACAACCTCGTAGCGGCTGCAGCA

CCGACGGT

>NODE_1071_length_217_cov_18.918367

ATTCCAAAAATTTTTTTTTTTTTCATTCTATCAAAGCCCGGGTCGTATCATTATATCGGCGACTGCCTCT

CGGGGCAGCTCCCTCTCTCCATCCGCAAGGCATACTAAATAATGCCTCGCTACATAATACGAAACCCCAC

AATACATGAAACCCCCAAGACCGGAAAACCTGTTGTATCAAGCTACCGCATACCACGAACAGCCCCCCTA

TTCGGTA

>NODE_1072_length_217_cov_9.979592

TGGAGTTAGGGCCCAGGCGCTTGCCGCCCACACCGGTGACAACGTTGACGAAGCGTCGGTTGTACTGCAT

CCTCTTCTTCGCCCTTCCGCGAGGCTGTTTCTTCTTCTCGATCTTGTCGATCTTGGGCGTCTGCCCCTTC

ACCTTACCGGCACGGGCCAGCGATCCGTGAACCTTTCCCATGGTTTAGAGCGGTTGTATGTCCTTTCCCG

TAGAGAG

>NODE_1073_length_217_cov_7.765306

GTAGATGATCTGTGCCAACACGTACTTGTTCGTGAAGCGCACCACAAGCCTGTACTTAGGCGAGTTGTAC

TTGTTCTTGTCCTGGCATATCAACCTTTTGCGCGCCCGGTAGTCGGTCTTGCCCTCGCGCCTGCGGCGGA

ACTTGACTTGGTAGCGCTTGAAGTACGCCTTCGTCTTCTGCACCTTAACGAACGTCATGATGACGTGGGG

GTGGGTG

>NODE_1074_length_217_cov_4.494898

TGGTTCTCTCGTCTGGCGTGCAGTACCGTGCATATATAGTGGTTATATTATATGTTGTCGATATAGTTTT

TTCGCTGCAGGCTTTACCGAGGGTGCAGCAGTTAGAGTTGCGCCGGAGAAAATAGTAGTCAAGGACTTTC

TGACTTTTGCGCGAACGGGAGGTGGATTTCCCCGTGGGTCAGCACGCATGGAATTGAGTGTTTTCTTTTA

AGGGTTC

>NODE_1075_length_217_cov_4.423469

CACGGGTTCCCACCGAGGTCCGACCGATTTCAGCATTCCGACTCTCACGAGTCTTCTGTCTGTCATCATG

CCTTTTCAGTAGAGCCTCCCGTACATCCCGTACCTCCTTGTACGCTTGACGTTTGTCGGCCACGTAACTG

TGCATCCCGCCTCTGAAATCACCACCATCGATCTCGGGGTGCGTAGCATCAAGTTGCGTGCGTGTATCAC

GACCGAA

>NODE_1076_length_217_cov_4.326531

AAGTGGTTTTCCCCCAAATTCGTGGGACTAGTTCTCAAGGGGCTTAAGAAGTTCTACCTAGTGTAGTGTA

CCAGGGTATGCTATTTTCTCGTCTTGGCCTTCCAACATTCATTTTGTTTTGCTCGTCTGCCGTGATATGG

AGCTCTATTAATAGTACGAGCGTAGGGTCTTACCGGCATTTTTTGTGTCCTGCTTTTGTGTCGATTACAA

ACCCGAT

>NODE_1077_length_217_cov_4.198980

CCCCACCGACTTCGACGCTCCGGTGGACGCCGAACTTAACGGCAAGGACAGCTCCGCTTCTGCCATCGTT

GCCTCGGCCCTCTTCGAGCTCTTCATCATCACTGAGGACCCGACGTACCTGGTGAAGGCTCAGGAATACC

TGGCGGCTGTGCTCTCGCCCGACTACTACCTGCCGGAAGCCACCGACGGCTGGCAGTCCATCCTTCGCCG

CGCCAGC

>NODE_1078_length_217_cov_2.540816

GCCTCGCGGGAGATGGCGTGTTCTGGTCCAACGCAACCATTACCCCATGATGATGACCTGTAGGGAAAAG

GTCTCGACATGATCGTCCAGAGGGTAGAGAGGATATGTACGTCGCGAAATAGCCCTCTGGCTGTTGGGCG

ACGACTGAAGGCGTCGCCTGCTTTAACTCGTCACTTTTGAACGTAACACTATCCGATACCAAAAAAAAAA

AAATTTG

>NODE_1079_length_217_cov_2.285714

TCGTTATCAGATTTAAAGAAACGTACTTTTTTGCCGTCTTCAAATCTAAAACCAACGCGGTCAGCCTTGC

CCGACTTCGGGTTAAGGATTGCTACGTTAGATACTTGTAAAGGTGCTTCTTTCTCAATAATTCCACCAGG

CTGCTGTAACTGAGGTACAGGCTTAGTGTGTTTTTTGATTAAGTTTACGCCTTCTACGAATACTTTGCCG

TTTTCAA

>NODE_1080_length_216_cov_53.251282

GGCGTACATGTCGCCCCGGTAGACTAGTGGTCAGCTTCAAAGACTACTCGCATTGGTACCCAAGTTCGAA

TCCCACCGTGGTGAGAGATTTTGTCTTTATTTGAAAAAATAAAAAGGAACCAATTGCTGAGAGCGCCTTC

GTGGCGTGGGTAGGCGGAATCCGACGCAAGTCGACGAGGGAAGAAAGTGCTGAAATCTTCTCGCGATAAA

AATGCA

>NODE_1081_length_216_cov_47.569231

TTGGCACCCGGAATTCCATAATGTTCGACCCCAAACTTCCCAAAATAAAATCTGCGTCATTTTGATCGCG

CCTATAACGGGAGAGTAAGAAGTCTGGAGGTTGCATGCTAGCCGCTAAGGACCATCTGCTCCCTTCGTTG

CTTATTCTAGCTCAGAGCATTTGGGATGCACAGGCACTGTTTAAACCCCTGTGTCCCAGCTAGATTCTCG

AGCAAA

>NODE_1082_length_216_cov_17.312821

TAATTATATTGGAGAAATAGTTCATTTTTCTATAGGGAGAATCTATTAAATTTTTTATTAGGGATATTAG

TATGGATATACGTCTTATATTTGTTTTTTCTCCTGTGTTAGCAGCAGCATCTTGGGCATTATATAATATT

GGTCGAGTAGCATTACAACAATTCAACAAACTAGCTTCACGCTAAAATGAATAATTACTAAATAAAATGA

AAAGTG

>NODE_1083_length_216_cov_5.446154

ACCGGCTGCGACCCCGGACAGACCTCGTTCTTCCAAGTGCTCCAGGTCCCTACCAAGATCACCAAGGGTC

AAATCGAGATCACCGCTGAGGTGTTCCTCGTGCCCAAGGGCCAGAAGGTGGGCAGCTCCGAAGCGGCTTT

GCTGCAGAAGCTGAACATCCGCCCGTTCACGTACGGTCTGATGATCGACTGCGTGTACGACAACGGATCG

GTGTTC

>NODE_1084_length_216_cov_5.415385

TATTATTTATAGTCCGCGTCTATGATATAAAGGAACTACCGCAGTAGTAGGTATAAATCTCGTCTGTACG

TGCGCGTACGGCAAATCTGGGTATTTTTGGTCAATCAATGCTTGGTCGGTCTGTGTTTTGATCAAGTAAG

AAGTACGAGTGCGGCGCGGCGGACGAGTAGGATCGACAGCAGCAGTATCATAGGTATATTTTGCTTGTTA

CGTATA

>NODE_1085_length_216_cov_4.943590

CATGGGAGTCACCGTGGAAACGACGAGGGAGGGAGACAAGCAGACGTACCCTCGACCAGGAGACAGCGTT

TCGATGGACTACACTGGGACGCTGCTCGACGGGACGGTTTTCGACAGCTCCATCCGGAAGGGGAGGAGGT

TCAATTGCAGGATTGGTGTAGGTCAAGTTATCCAGGGATGGGATGAGGGCGTGCCTAAGATGTCGCTCGG

AGAGAA

>NODE_1086_length_216_cov_3.897436

ACTTCGAGTACCTCTACTCCGAGTGCTTCTTTCGTGGGGAGCGGGAAGGTGAACAAGAACTTGCTGGCTT

TGTCGACTGCCACCAGTAGAACTTTGTTGCCTGCGTCCGACTTCACCCCCATGTCCTGGATGTCGACTTC

GAGTACCTCCCCGGGTCGCAGAAACCTCGCCGGTAGCATGGCAACCCGTTGACTCGCTGATCGCTTGCGA

CGCCTG

>NODE_1087_length_216_cov_3.487179

TTGGGGAGCGCTCAAGAAACGGGTAGCATCAGCGAAGGGATATTTGGGAAGGGCTCTCGACGCCAGGACA

ACGGTAGGCTCGTGACAGCGTTCCTCGTGGTGAACTACATGATCGGGAGCGGGATCCTGAACTCGCCACA

GACGTTCAGGGACAGCGGGCTGGCGGCGACGACATTGCTATACTTCATCGCCTGCCCTGCTGTCTGGCTT

GGGGCG

>NODE_1088_length_216_cov_3.430769

ATTCCACCCAGAAAACCAGCCGCCCGAAGTTGTCGCTGCTCCAGTTGTAGCCCTGGTGTGCTGGTAGAGT

GTGCTCCCTAGCTGTGCGCCTGGACGATCCAGTGGGTGTTCGATGGGTCCAGTGCTGACGAGTCTGGCGG

TGATGGTCGACCTCCGATGGGTGTCCAACTCAGAACTTTCGTGGATGGCCGTTTGCTTTACCCGTTGTAC

ACACGT

>NODE_1089_length_216_cov_3.246154

TTGGTTTTGTCTCGCCCTGCTCCTCTTGCCAACGGGAGGCTATTATTGAAGGACTGGAGGGTTGACACCG

TACCCCGCAGTCGAGGGTAGGGGACTTTTTTTGCGATCGGTCGTCCCGTCGATCAGTTGCCGAAGACTAG

GTTCGAGGCGGTGCGTGTACGTGCGCGTTTGGGGACGGGAATGAACTTTTTCGAGAGTCTGGAGGAGGGG

AGGCGG

>NODE_1090_length_216_cov_2.917949

ATTCCATCAACCCCCGACCACTCCTCGAGGATGGGCCTCAGTCCCTCGAAGATCTGCTCCTTCAGCTTCC

CGTTCTCCGGCAGGTTCACCATGTAGCTCGGCGCCTCGACATTATTGGTGAAGACGTTGTTCTGGCCCCA

CTTCTCCTTCTTCGGCGACTCTTTGTTCTCCTCGTAGTAGGCCCTCGAGTGCAAAAGGGTAGTATCACCT

TGAAGA

>NODE_1091_length_216_cov_2.805128

TGTCGTATATCTTTGGCAAAAAGTTGCATGCGCTTATCCGGTGAAAGACGAGGTGAACTCGTCGCACACA

GCCGAGAGGTGCTTCTGCATGTCCGCGGTCAGCTCGTAGTTGTCCTTCTTGATGCCGTCCAGGATGTCGG

TCTTGTGCGAGGCCTTGATGTGGCTGAGCCACGCCTTCTCGTAGGCGGTGACCTTGTTCACGTCGACGCG

GTCGAT

>NODE_1092_length_216_cov_2.702564

GGAGCTGCTGCTCGCCGCCCCCGCCCCGCGAGCGCCAACAGCGTTAAGAATCTTCAGCCTAGGACCCTTT

GGGATGCCAATGTCGGCGAGGTCCCCCTCGGTCATAAGCCCGAGAGATTCCAAGTCCACCTCGTTCTCCG

CAAACTTCGAGCAGTACTTGCCGAGCCCCATCGACTGCAGTCTCCTCCCGAGGCGCGCCACCTTATCGGC

GATGTC

>NODE_1093_length_216_cov_2.646154

CGCCGCCTCCGTGCCCAGGGCAACGGACCGCAGCCAACCTTCGATGCCACCTCGGATGGCATCCCAGCTA

ACTTCGATGCCAGGGAGGCGTTCTCCGAATGTGCCTCCATTATTGGACGTGTTAGGGACCAGAGTGATTG

TGGGAGTTGCTGGGCGTTCGCCTCGACGGAGGCGTTTAACGATCGTCGATGCATCGCCGGTATTGGTGGT

GGCGGG

>NODE_1094_length_216_cov_2.625641

ACCGAGACACACGGCAAAACGAGATCTTACTATACAGGGGAAGAGGGGCGTAGCTGATGGTATCGTTCCA

CAGGCAACAATACAATTATTCTACCGACGGCATCATTTATCGACGACGGTATCAATCCACCAACGATATC

ACCCTGCTACCGACGACATAATCCCACCGAAGATTACCGTGCTTCTACCGACGGCATCATTTCACCCACG

GAATCA

>NODE_1095_length_216_cov_1.907692

GACAATACTTAATATTTCGGCCAAAGTGTGCTGCACGTTTCGTGGGGCTGCCCGAGAGATGCGACTAAAG

CACGATCAGGAGAGGTGTTTAGGTTTGCAGGCGAAACTTAGATAAACCGGATTTCCGTCCATACAAGCGT

TTGTGTAAACTCTAGACCGTTCCATGGGTACTACTCATTTTCCCTTAGACACAACAAAGGTGCTCCCAAA

AGGTAG

>NODE_1096_length_215_cov_2049.293814

CGTCAATTCCTTTGAGTTTCACTCTTGCGAGCATAGTTCCCAGGCGGGATACTTAACGCGTTAGCTACGA

TACTGCATGGATCGATACACACAACATCTAGTATCCATCGTTTACAGCTAGGACTACTGGGGTATCTAAT

CCCATTCGCTACCCTAGCTTTCGTCTCTTAGTGTCAGTGATAGCCCAGTAAAGTGCCTTCGCCATCGGTG

GTCCT

>NODE_1097_length_215_cov_4.737113

AGGCCCAAGGAGGTCTTGGTTACGTTGAACACGACGCCGGTGCGGCCGTGGTAGTACTTGAAGGGCATAC

CCTTGTGGATGGAAGGGTTGCACTTTATGTCCACGTAGTCTCCCCTCTTGTACTTCCTGAGGTAGATGGA

GAGCTCGTTGACTCCGTTCTTGCCGAAGTCACGCGCGAACATGTCGCGCGTACGCGCGCGGTAACCGAAA

GAGTG

>NODE_1098_length_215_cov_3.917526

CATGAAAACGGAGAACACCATGTTTAAAGAAACTATGACAGCATTCGCCATCGTTGCAGGTCTTGTGGCC

ACCAATGCCCAAGCCAAAGAAGGTGGCGATAAGATGGAAGCCCGCTTTACTCAAATGGACGTAGACGGCG

ACGGTCAAGTCACCGCAGCCGAGTTGGAAGCCCGCGCTGCTGCGCGCTTTGCTGAAACCGATACCAATGG

CGATG

>NODE_1099_length_215_cov_3.819588

TCCAAGACCCGCTACGAAAAAAGGTCGCGGCCTTGACGACATTTCATCATCATCATCGTCACTAGCGCTA

TGTACTACGCTTGCCTTTCCTGATAATACCCTCCATTCCTTTTCCTGTCCTGTCTGGCCGAAAACATCAC

AAAAAAACACCCCGTGTGCCTGGATACCCGAGGCCACTATCATATCACACCCACCCTTTATTTAATCCAA

CCCAA

>NODE_1100_length_215_cov_3.340206

CAAGCTTGGACAGGCCGCCCAAAGCACCGCCTTCTTCGAGGCCTGTATGCATTGAGCCCATCAGCACGCG

ATTCTTGAGCTGGGTATGGCCCAGGTCCAAGGGCTTCAGAAGATTGGCATACCTCGTCATTATGCCAAAG

ACGCTTCCCGGGATCGATCAGCGCTTCAGGGGTGTCGTTGGTAGGATGTGCTCTGATGGTGGATGTGGGG

TCGCG

>NODE_1101_length_215_cov_2.953608

CTAGTTTTGGAGCAGCTTTGTGATAATGGAACCAGCCACCAAATAACATTAATCCAGACATAATTAATCC

ACCAATGGCAGTCCAGTATAATTCAATTTCACTTGTTATACCTTCAGCACGCCATAATTGGAAAAATCCT

GATGTTATTTGAACACCTTGAAAATTACCACCAACGTCACCATTTAAGATTTCTTGACCAACAATGGGCC

ACACA

>NODE_1102_length_215_cov_2.716495

CAACCCAGAAACTGTTGACTTCGTTTACCGCATCCGGTCTGTCCTCCGACTGTCTGTACGCCTCCGAGGC

CGCTCCAGGCCCGTGGAACTTGCTGTGCAGCAGGCACTCGTCCATATCCAATACGACCGTCATCTTGCCA

GGCTTGTAAGGCGCCAGCCGAGAGTGCCTGGTGACAGTGATAGGGCCTCGTGGCCCTTCCTCTGCTGCCA

CGCAT

>NODE_1103_length_215_cov_1.840206

CGCAAGAACAGTTTCACTATTCGGAAATCCCCCAAGTCCGTTCGATGCGGGCAGCCAAGCATCGGACAAC

CACCGTTCATTAAAAAATATCGACCAGAGGTTTCGTCCTAACCGTATAGACTCCATGAAACGCTGTGCTC

GTAAATCTTCAAGATTGTCTACCTCAAAGTGTTTTTTGTCTATTTCTGCCTTTTTCGGACGCCCGTGAGT

TGATG

>NODE_1104_length_215_cov_1.768041

ATTTTTGCAACATTTCTTTGTGTTACAAGTACGTTATCAGCCCCTTCTTCGGTGCATCCTTTTCTACTTC

TTCTGCAAATATCATCCCACCACTACTGCTGTACGAGCACGTAGATCGTGCAGGTCGTACGAAATAGATC

AGGGATACAAGGCTCCCCTTCGGGATTGTGGTTGACTTGAATCCCGGGGGGTGTCGCAAGCTGAAAAGAA

AGTGG

>NODE_1105_length_215_cov_1.634021

CTTTTGATCGTGAGAGTGCCGCCTTCCACAACAATCGTCTCTATACTCGTGGCTGTATCCAGAATCTCGC

ACTCAATAGCCCCCACGCTCAAGTTTATGGTGGTGTCTCCCGTTAGGGGTGAAGGGACGTCCAAGAACAT

GTCGACGTCGTCGCAGACCGCCCTGGCAGTGGACAAGGCGCCCAAAGCCAGCCAACATCCCATAGCCCAA

GACTT

>NODE_1106_length_214_cov_17.238342

CCCCGAGAATTCCAACAACAACCCAAATCATCGCCAGCAACAAGAACACGTACGAGCTGCGCTACTTCTC

CATCATCAACAACGAGCCGGAGGCGGACGAAGAGTAGACTTTCACTAGCTGCTGTTCATGACATCGGTGG

CGTGTTGGTTCTTGTAGAAAACTTTGCGGCGGGGAGCAGGGGGCATTACTCTTTCATGAGTGTTGCCTTC

AGTT

>NODE_1107_length_214_cov_4.787565

TGATTCACGCAAATTCGTAGGTAACTGACTTTGCGCCATCACCGTCCCTTTCCATGGGCTCGATCGCTTC

TCTTCTTGTCCTTGTTCTTGGTATTCTTGTTTTTTACGTTGTCGTGGAACGTGCGACGTCAGGGCAGGCT

AGGGGCTCGAAGCCCGTGCACGACTTGCCGAGGACGTTGTAGTATCCCGTCCAGTTCACGGACGTGAGTC

CGAG

>NODE_1108_length_214_cov_2.989637

CGACAATGTCTTCTACCGACTAGGATACTGGGTGGCGGGGCACCCCAAGCGGACACTCCTGATCAGCCTC

GTGCTGGTCGTCCTGTGCTGTTTCGGCTTCGTGAACTTCGAAAACAGCGACGGCGATATCTGGGTCCCGA

GAACATCGTTGTCCAAGGACCAAGAGGAAATCGTGTCGGGAGCCTTCGACGACAGCGGAACTTTCGCCGT

GATA

>NODE_1109_length_214_cov_2.797927

TGGGACGACTGGCTCCGAATCAAACTGAAAATTGCACCTAAACATAGGTTGTTCGCGTTACGTAACAAAA

CCTCCAATGGCGGTGAATATAGAACAACAACAATCAAAAATATGGTTATGAGTTCAACGCAGTAAAGCAG

TTACTCGGCCGAGTAGCCCTGTGCAGGTGAAAAACAAGCACCTATACATGACCCCCCGACCGTAGCACCA

GTCT

>NODE_1110_length_214_cov_2.751295

CACGCTGGTGGCACTCACTCGCTTTCTTCCGTTCCATGATCGATCGCGATTCAGAGCTTTCGCCCGCAGA

ATCCACCAACAAGAGCGCCATCCAACAGCGAGACAAAAAACTCCACTCAAATCAGTCATCCGCACAGTCA

CACCATGGTCCATCATGTCAAACCAACGACAATTTCCCGACCAACTACACCACAATCCACCTCTGCCGCA

GCCT

>NODE_1111_length_214_cov_2.549223

CGATTCATTAGCGTAATCAATACCGCCTGAAATAGCTGCATCGCCCCCAGTTTGCTCTTGACCACGCCAT

AAGTAGTTTGATGTTGCCGCTGCGTTTGCTGTTACTTCAGCATGTACGGCTGTTGACGTTAAAGATGTTG

CGCCCATTAATGCTATAGCACTTATAGCGATGGTTTTTTTTAGTTTTAGCATGTCCATTTCCCGTTATAT

GATT

>NODE_1112_length_214_cov_1.901554

AAGATACGTAGGATGCTTCTAAGCTTACACATAGGTACGATGATTATATCCCCTTTAGATAGGATTGCAT

TTTGCTGGTGGCTTGATAATGACCGAACTTTGGTGGCTTACCGGTCTTGTTGTTGAGCGCGCCGGGGACT

CTTTTGCCCTTGTTAGCGTTAAGGAGGCCCTCGTAGGACCCGGGCTTTCCCTCCGCCTCCTCGAAGCAGA

ACAC

>NODE_1113_length_213_cov_67.796875

TGACGGAATAGTTCGTTGAAGTACCTACCTATAAGCTGCGCAGGCAAATCCACGCGTCTTGGGGAAATAC

GATAGTACTCGGAGTCTTCGGACAAAGAGATAGTGCGCCTAAGGGCTTCCAAGAAAAACCTCTAAACTTC

AGATTATAAGTACCCGTACCGCAAACCGACACAGGTAGTCGAGGAGAGAATCCTAAGGTGCTCGAGAGAT

TCA

>NODE_1114_length_213_cov_19.125000

AGGCATCAGGGATTGCGAGCTCGTCACAGCGACATCATCGGCACCGGGAGCAGACACTACAGCAGTTTCA

CATTGGTGACCACCGTGGGATTCGAACCCACGAGTACGAATACATGACGACCTGAACGCCACCCGATAGA

CCGCTCCGGCAGATGGTCCCCAAATCTCACGAATTATGTGTAAAGCTGGGAAACCGCTCCACTGGAAATT

CCC

>NODE_1115_length_213_cov_6.578125

CTCCCTCTTTTCTTTTGGGCCCGGACTCATTTCTTTTACTAACTGTTGTACTCGTATATACTATATACAT

CGTATATACAGAGTAGATAGGCACCAACAGGAGGCCAGAAACCACAGCCGGGCAAAGACAAGGGCCAAGC

AAGCCCCCTTTTACTGATACACCAGGTACCAGGTACGACGGATGTATTGATAATATACAGCAGTCTTCGT

AGT

>NODE_1116_length_213_cov_4.598958

CGATGGGGATGGCCATGTAGGGGTTTCCCATGAAGTTCGCCGGCCCGCTGCAGATGCCGGCCCCGCCCAG

GACGTCCATGCCGTCGTTGATGTTCTGGCGCATGCGCGTGGTCATCTGCTGCTTCATCACGGCCGACAGC

ACCGCCGGCTGCTCGTGCCCGTTGAGCATGGAGTTCATGAGGGCCTGCCCGGAGGTGACGAGTAGGGTGT

TGC

>NODE_1117_length_213_cov_3.958333

GCTTTGACGATTTTTCAGGATGCTACCAAGAAACTGGCCTCCTACGCTTGGGGCTCCCGTACTATACTCC

GTCTGGTCTGGACCTTCCCTCCACGTTTATCAGTGCATTCGAACTCGACGCAACGGGCTCCGACTCCTCT

TCGACAGTTACACCGTGGATAATTGGGGACCACGAGGCCGACGTGGCGTGCATCTCATTTGAGCTGGCCG

AGA

>NODE_1118_length_213_cov_3.427083

GAAAAAAGCTTCCGAGCTGCTTGCCTGCTGCTGCCGAGGTTTTTATTCCCCCAGGTCCCTACATACACGA

CACCGCACGCCCTCAGACTGGTGAGGGGGTAAACCAGTTAACAATGACACACAGCTACCCTTACACTCGG

CGAGGTACGCCGCTTTTTTTTCTTCTTCCTCCACTATCTCAGCCAGAGACACAAATTTGCGTTTCTCTGG

CCT

>NODE_1119_length_213_cov_3.031250

CCCAGTCACGCACGAGGTACACGGCCGGTCAGGTCACGGTTGTCCTCTACCATACACCATGCATCTTTCA

ATGCCATCCCATGTTCACTTGAGTTGTGGCACCCTCCGCGCCTCCGAGAGTTGCCATAAAATGTTCTCCC

AGCCCTCCTCCGATGATGTCTTTGCGCTCGCTCATGACTCCACGCATCCCAGACCCCATCTCGCTCGCCG

TGC

>NODE_1120_length_213_cov_2.703125

TCACAGATGGTCGATACTGCGCGTTGCGAGGCGTACACATATGTACCCCGAGAGTAATCGGTAGCGGATG

GCATGTTCACTCCGAATTGTCATTAGAACCTGTGGAACATGTTTTGTCCACAAAATGAGGGCGGCACGGC

GTTTATCCTGTCTAAGCGGCAGTTATCAACGGTTTTGAGCCGTGGTGCTCCGTACTGGTCTAATCAAAAA

CAT

>NODE_1121_length_212_cov_3575.523560

CGATAGCGCGGAGGTACAGAGAGCATTGATGAGAAAACTACATTCCAACTAAAACACCGAAGACCAAGCG

GAAGCCGGGCCCGACGGCGCAGCAATGCGCAGGAAAGGCCTTGCCCCGTGAATGAGCGACCCTTTGTTAA

CCAGACTCTTAGCTCTTGGAATTCTCTGGTTACAATGAACTCAAGTAACAACTACATCTCGTATTCCGTC

TT

>NODE_1122_length_212_cov_2.450262

CTTGGATGTGGATGTCGATGTGCCCTCGGTAGACGTTGACGCGTCTGCCCCGTCCGCCTCCATCGACATG

CCCTCTGGCTCTGTGGACGTGCCGTCTGGCTCCGTGGACATGCCGTCTGTGGACCTGTCCGGCAAGGCTC

CTGACATGCCGTCTGTTGGCGGGGAGATCTCCGGCGATTTACCATCAGTGGATGCTAAGCTGACCGGACC

AG

>NODE_1123_length_212_cov_2.356021

GTAAAGATCTTAGCTGCTGGTCACACAGAAATTGAAACTGAACGTAACCGTGCAAAAGAAGAACTGCGTA

AACAAGTCGCTATTCTTGCTATTGCCGGTGCCGAGAAAATTCTTGAACGTTCTATTGATGCCGCTGCACA

CAGAGACATCTTAGATAAATTCGTCGCTGAACTTTAAGAGGATAGAGCTATGTCTGAATTGACAACCATT

GC

>NODE_1124_length_212_cov_2.099476

CTCACCAGCGCTTACATACGTCCAACAACGGTACCGAAGTACGACAGCTACGCCAGTAGTCCTACAAGAG

TCGCTCAGGCGGCTATATTCTATCTCCCTACCTCCCGCTATCCGCTCCTCGCTTGGCTGTAGCCTATCAG

CAGACAGCAGCTATCCACTCTCAACAGTCTTGCTTCAACGCTCGCGCACGTTCAACAGTATTCCCTCCCA

AC

>NODE_1125_length_212_cov_1.952880

CTAGCCTAGTTTCGTCCTTGAAACGTGGAGCTAGGCCGGTCGTATCGGTATGCCTTGACCAGGCACTAGA

TTGAGTTGTGTGGATGTCAAGAGTGAACGCGTAAACTCGGGTTCTCGATCAAGTGACCAATTAAACGAGT

GTAGCGGGTTAAGCGGGGTAGTAGTGTAATGAAGAAAGTTAGTAGTTTTGTATCGCGGTAGGTTTGAGTT

CC

>NODE_1126_length_212_cov_1.591623

ACGTATTGGCAGAGTCAATTTAAAATTAACGCCAGAGTGTCGAGATTTTATTGAACATTACCCGTGGCCA

GGTAATGTGAGGCAGTTAGAAAATGTGTTAACCCGAGCAACCTCGTTGAAGTTCCCACATTGGGAACAAC

AGCACGGCGAAAAGCCATGTTTTTTTGTCTACTGTGAGATAGCTCGCTCACTCGCTGCTGAGTTGCTTAC

GG

>NODE_1127_length_212_cov_1.319372

GAAATCAAAGGGTTCGGCGCGCAGTCGGGTCATACAGAAATTTACCGCGGCGCTGAATACGTTGTGGATT

TCCTTCCAAAGGTCAAAGTCGAGATTGTTCTGGACGATGATCAAGTCGATGCCGCGATCGAAGCCATTGT

GAATGCTGCCAAAACCGAGAAAATCGGCGATGGCAAGATCTTCGTATCTCCCATTGAACAAACAATACGC

AT

>NODE_1128_length_211_cov_9.668421

AGTTTAAGTCCGTCAGCTAGGGGCTGAAGTAATCCCATATAACCAATAACGTTTGGACCACGCCTTCTTT

GAATTCCCGCCATAATTTTACGCTCCGCTAAGGTAAAGTAGGCTACAGCAATCAATAAAGGAATAACAAG

ATCAAGAATATTTAAAAAAATAGTTAGGAAAGTTAGCCACATAATAAATTAAATAATACAATAATATAAT

T

>NODE_1129_length_211_cov_7.084211

GGCACCCGAAATTCCAGCATCCAAACGCATCACATCCATCCCCTCCTCTTCTTCCAACGGTCATCAATCA

TCTCAACAAGCGCGTTTTTATTTCGTTTAATTATCCTTTTTCGTTTTTAATCACTTTTTTTTTTGCATAT

TTCGGGGCGCCCTTTTTACCTAAATCCGACGCCCTTAGGCGATGTACCCCCTCACTCTCAACCAACCCTA

C

>NODE_1130_length_211_cov_6.473684

GCGAGGTCGCGCCTTGAGTTATACTTGCATTATCTCCTTAAACACGGATTTTTGCTCCCGAGTCCCAGAG

TTTTCATGCCAAGGCTGACTTTTCAAGAGGTAAAATAAAAGTACCATCTCCTGACATCTTTAACCTGTAG

TTCTTTTTGAGGCCAAGCGTTACCAACCATTCTGCACAAATCAATCTGTCGTGGAAGCTTTCGATGCGAC

G

>NODE_1131_length_211_cov_5.873684

AGAGTGCAGGTCGAGGATGCGCTTGTGGATGCGCATCTCGAAGCGGTCCCAGGTGTTGGTTCCCTCGCCG

CAAGGGGACTTACGCGTGACGATCTTCAGCTTCTTGGTGGGCAGGCGCACGGGGCCGGACACCTTGAGCT

CCTTGTCCACGGCGCCCTTCTTCAAGTCCGCGCAAACCTTCTCGAGGTTCTTGACGTTCTTGGAGGTTAG

G

>NODE_1132_length_211_cov_5.163158

ACTACTGGGAATTTTTTAGTTCCTGATTCTGAAATTACTGTATCCCCAAGATAATAACATTCGTAAAATA

GAGATTTCATTATATATTATTTTATAAATTAAGAGGTATCAAACAACATGGACAATTTCTTAAAATATCT

TTCGACTGCACCTGTTTTATTTGTTGGATGGATGACATTTACTGCTGGTTTTATTATTGAAGCTAATCGT

T

>NODE_1133_length_211_cov_4.300000

GGTCGAATGCGCTCCCATGTTCGTGCCCGTGCGTGCGCACACTATTCATCCTTCATCCATCGATCTATTC

GCCACCGCAAGGGACGTGGACAAAAATTGCCTAACGACCTCCTACCTATACCTACCTACATTCTACCTTC

CTGCCTGTGGGTCGTGGTATGCAGCAGCAGCAGCAAGTAGCGAACGAAACGCACCCAACCCAACCTAACC

C

>NODE_1134_length_211_cov_3.994737

TAAGATTGGAGCGTTGGCGATTATCTAATAGGTTTAAGTTTGAACTTCATCTTCAACCAACACCAAAATG

GATACAGACAGATTACTCAAATTTAAGTTTTATTCTTTCCTCAGATGTTTGTGCCCCGGTTTTGTATCCT

TTTAGGGTAGACGTAGATGAAGCTCTTTGGTCTTCTAGGTATGGTTATTTATAGTTTTTCTCTAGGTGGG

T

>NODE_1135_length_211_cov_3.747368

AGCCGAGATACGGCTGACCGGCTGGCTGGCTTTTGACTTGGCTGGCTGCTTGCTCCACTCCGTCCACCGT

CGGTTTTACAAACGGTCCAGGTAGCGGTCCAGGGGGGGGATGTTCTCGCTCAGCCCCTTCCTCTTGCGCA

CCTCCCTGACGACGTCGTACGCCTTCGACCCCTCCTCCTGCGGGTTACCGCCCATCTCTTGCCAGTGGTC

A

>NODE_1136_length_211_cov_3.515789

AGGGCGCCGTCGAAGCGCAGCGAGGCCGTCAGCGACGAGATCACCTGCGCCACCAGGCGGTTCAGGTTGG

TGTACGTGGGCCGCTCGATGTCCAGGTTGCGGCGGCACACGTCGTACAGCGCCTCGTTGTCCAGGCAGAA

CGTGCAGTCCGTGTGCTCCAGCAGGGCGTGCGTCGACAGCACAGAGTTGTACGGCTCCACCACCGCCGTG

G

>NODE_1137_length_211_cov_2.689474

TTCTGGGTGCTGTAGGGTTATCGCTAGACCCAGCCGACGATGTTCAAGGCAGCCTGAATCAAGAAATAGC

ATAAAGCCTGTGAGGGTCTTGGCCACTACCACCACATACCGCAGGTGCATTCCACACTTCTAGCCGACGC

ATTCGCCCAGGGCCTCCCCGATAAAGGTATGGAGTTCCTCGCCTCGAACCAAGATGTATATACAATGCCA

C

>NODE_1138_length_211_cov_2.357895

ACATAATATGTCAACCGAAAGAAAATAAATTGAGGATAGGAAATACATGAATATTCATACTATAGCCGAT

CATTTAAATGGTTTAGGCGACAACTCAGAAACTGGAATGAACTTTGATTGTCAGCCTATTTCAGGTGAAG

TCGATGTACTACAGATTACTATTTTAGGTAGAGAAGAGTTACCTATTTTTGTTTCAGTAACGGATGATCA

A

>NODE_1139_length_211_cov_2.194737

TGTGCGGAGCAACGGCCTACCTCGTGGGTCTCTACGTCGACGCTCATGATCGAAGGAGCAACCCCCCAGT

CAAGATCGAGCAGAAGGTGGAGCCCTACAGCTTTCCGGATGTGGCCTTTTGTCCTTCCGCGGGAGACGGC

TGCGACTCCTCCAACCCTGAGGATTGTGCGCAAGATATGCAGCTGTTTGTTCCAGCTCTGGCGGGCTTCG

T

>NODE_1140_length_211_cov_2.057895

GAACGCGGTGCGAGCGGTAGCGAGGTATTCGAATAAGCCGAGGGAGGTACACTGGAAGACAGCGATAGGT

ATTTTAGAGTACGTGTTTTCTACGAGTGATTTTGGCATTACGTTCCAGAAGGGAAATGGGCTTGAACTGG

TAGCGTTTGCGGATGCAGACTATGCTAGCAAGGCCGCTGACAGGAGGTCAGTTTCCGGTGGGGCTATAAT

G

>NODE_1141_length_211_cov_1.689474

CGCATGGAGACATGTACACTGGAAAATCTCGACATCGAAAGCCAAAACAAAGCCCATAACAATCATATCA

AAGTCTGAATATATCCTCCGCGTGGCCAAGAACGCTGGAAAATCGCTGCAGCTTTTGTTATGGTCGAAAT

GAAAAACAAACCGCCCATAACTGCAGTCAAATCAAAATATGAATATATCCTCCGCATGGCCATGAACGCT

G

>NODE_1142_length_211_cov_1.505263

CCCCGTCCTAGGCATCAAGGACTGAAGGGCCGAAGGACCGAAGTACTTGTGATCCCTAAAATAGGTCCAT

GAAGCAGTCCTATTTAAGGCATGATCCAGGCATGATCCAGGCATGACCCAGGCATGATTAAGGCATGGTC

CAGGCATGATTGAGGCATGATCAAGGCATGGTCGAGATTGGTCCCGGAACAGACCAACACCCTCCCCCCG

T

>NODE_1143_length_210_cov_825.830688

AAGTGCTTGGTTGGTGTTGTCCCCGAGGGGACGGACCTCACAAGGCGGTAGTGCAATGCCGAGGAAATCC

AGCCGGAACACTGCAAGCAGCGCTAAACCCGGCAATTTCAAAAACAGGTGTGAAAATGACACTCGATGAG

TGTCAATCAGTTTTCACACTGATTAGAACAGATACTACACATGATCTTAGCCAAAAGGCCGAGAAGGTAT

>NODE_1144_length_210_cov_12.687831

CCAGCGAGTAGCCTGAGACTCGAAGGAGTGTTTCAAAACACTACGATCCACCAGGTGTACTCGATGTCGC

GTCTGTCGTCCCAGCCCGTGTCGTATAGGAGTTGGGGATAGTTCATACTCCAGATGCCGCATTGTATCTG

GAGAGGTCATCGTGCAGTCCATGCCGTCTCTGAATGCGAATTAGTGGTAGCGTCCGTCTTCGCAGCACAG

>NODE_1145_length_210_cov_6.100529

ACTTCCGAGTGGGCTGCCAATTGTTCGACGGTTCGCAAGAAAGACGGGACAGTACGGGTTGTGCAGGATT

TTCGAGGTGCCAATGCGCTTCTCAAGTCACAGAGTGGTGGACTCGGGGACCTGCAGCACATTATGGATGA

GATAGAAGGGCCGAAGTATTTCTCCATCATCGATTTGGCTTCAGGATTCTTACAACTCGAGATCCATGAA

>NODE_1146_length_210_cov_4.153439

TTTTTTTTTCTTTCCCCTTGACGCGTGTAGTTTTACAAGCTATCTTGAAAAACCAAAGCTCTACACATAC

GCGTACCTCTTGTCTTTCCACGCTCTCGAGTTTCCAGGCTACCTTGAAAAACAGAAGCCATGTACAACTC

TTGTCGTCCTTTCATACTCTGCGGGTTTTTATAAGCTGTCTTGAAAAACCGAAGTAGACATTGAGGACGA

>NODE_1147_length_210_cov_3.349206

GCTCAAACATGGTCATTTTCCCCGCGTTGTAGAAAGCTCCGCCCAGCTTCACCTCGTTAGAATTGTAGCC

GCAGTCACTAATGGCCAACTCCTCGCCCTTGAACACCATGTTCCCGGTCTGCCAATTGTATATTCCTCCT

CCGTTCCCGGCGTGAGCCACTGTGCCTCTCTGGCCGTTACAATTGGAGAGCGAGAGGCCGTTTTCGAACC

>NODE_1148_length_210_cov_3.142857

CAGCTGCTCATCACCGAGGGTGGTGCGGTTGACCGTCTTCAGGCCAACGACATCAACGACTCTTTGGACC

TCGTCCTCGCCCAGACGCTGGAGCGCATCAAGAAGGACTTCTCCCCCGAGGCCGACTTTGAAGCCCGCCA

AGTCACCAAGAAAGACTTCCCCGGTCGGTACATGTACCGCGACTTTGGGACCCAGTACTGGGAGGCCACG

>NODE_1149_length_210_cov_2.904762

CAAACACTGCCTCGCGAATTCTTCGTCCACCTGCTTCATCTCGCTGAAGATGTGGGCGAACTGCGGAAAC

AAGGTGACGAGGTAAATGTGGGCCTCTTCGTAGGGTTCGTCGCTGTCCTCTATCGTCGAAAGCATCTCCG

AGACAATGTCGCCGAGGCAACGGTCCGCTCCCACCAGCATCTCCCTCCGTTCTCGACGCTCGCGGTACTT

>NODE_1150_length_210_cov_2.867725

AGTCCACTAAGTGAACCTGTCGAGAAGGAAATAGCCCACACCATGCGGTTCACCAGTCTGCTAGGACCAC

TTCTAGCTGCCGTCAGCACTGCCCACGGTCTCAACTCATGCCTCACCGATGACCGCTGTGCCCGTGGAAC

TACCGACCTGAGCCTACGCAGCTGCGACCTTCTAGACGAAGATGTACCTGATATCTCGGCGTGCTTCGAC

>NODE_1151_length_209_cov_19.207447

GGAGGGAGGGAGGGCGGAAGAGCACACAAACAGCAGTCAATCTGCATCGCCGAAAAAATATTTTCACTCT

GTATCAATACTCCATCCATTCCATCGCATATCCATATCATATCACAACGTCCCAGAATTTTCCAACACCG

TACCAACCATGAGAACCTTATCTGAGCAGCAGTACGTTCAGTTGCTTTGTACACCTGTATCTGCACACC

>NODE_1152_length_209_cov_6.367021

TACCGTAGCTTCAATCAGGTCACCACACACTTCCTTTCCACAGGCAGCTAGCTGTTGTTGTGTCATGTGA

TTCGTTTTATCTCGGTCGTGGGAACAACAACTTCTTCCGTAGCACACAGCAGCACGAAAAACGTCGACTC

ACCCTGATCCTCTCTTCCCAGTGGAGTATCAAGCGAGGCGGAGATATCCACCCGAGGAGGGTAGCTCTC

>NODE_1153_length_209_cov_5.367021

CTGAGGTGTGGCAGACGCGACAAAAGGCACGTCTGACTAATAGAGTCGCACGGTGTTCGTATGCATATGC

CCCGCCCGTGCTGTAGATACACCACAAAGTTCACAAATAATGTCCTTGCTCATTCGAAGGTGAAAAAAAA

TCATCAAAGTCTTGAAGAATAAGTATATATAAAAAAGGTGTTGTATCGTAGGCTTTGGTGGATTTTTGC

>NODE_1154_length_209_cov_4.526596

AGCGCTGTCCTCAGAGCCGTCACCCTCCTCCCCGAAGCCGAGGTTTGAAATACTACGAAAACTACCACCG

ACCAAACGAGGTCGCTCAAACAGTGCAACCCGCCGGCCTAAGCCCGTGGGGCCCCTACCCCAAGTATCCG

GTTCAGCACATGGTGAGAGTGAGAACAACAAGACCCGTCTTAGGGACTTGGATTGGGGTGGGGGGTGAG

>NODE_1155_length_209_cov_3.585106

CGAGCCAGAGGAAGGGCTTCTCGAAGTTGTAGTTTGATTTCGCCGAGATGTCGTAGTACTGCAGGTTCTT

TTTGCGGTGGAAAGTGATCTGCTTCGCCTTCACCTTCCGGTCCTTGATCTCCACCTTGTTCCCGCACAAG

ACGATCGGGATGTGCTCGCAAACCCGGGTGAGGTCGCGGTGCCAGTTGGGGACGTTCTTGTATGTGATG

>NODE_1156_length_209_cov_2.047872

TTGTGGATACGGTGGTTAAGTCAAGTATGCTTGGTGGAGCGTAGTGCATGAAGAAAGTTCTCCAAGCCGT

TGGAAACAGGCACCACCTCTAGTGAGCATTGTGTGGCTCCGGGGTCTCATGGTAGATGATGTTCAGCCGT

GTCCGGCTAGTGCTATTGAATATGTGTATCGTACTGCTCAATGTCTACTGCAGCTTAGTCGACAAATGT

>NODE_1157_length_209_cov_1.441489

CGGCAGCGGCGGCAGTGGCAACGGCGGCCGAGTCAAGGGTAAAAATCGTGACGAGTGGACGTCGGGGAGG

TCGGGCACGGGACTGAGAAGGCTTGCGCCGGGAGGCGGTGCGATGGGGAGGGGATAGTTATGAGACACCC

CCCCCTTTTTTTCGTGATTAGAGGTAGCTTCTGGGACGAGGATGAAGAGAGGTGGTGCGCTGGGCAGCG

>NODE_1158_length_209_cov_1.170213

CCGACAAATTCGCGAGACAGGAAATTATGTGCACCCTTTATGAAGTTCGGCGTGTGGCCTCATAAGCTGG

TTTGTTGGCCGCGCTTGTTTTGGTTACGCCGTCTCACCAAAAAAATACCTTCGTTGGTTCAAAGTACTGC

TGTATGATATGTACAATTATTATGTACGAATTGTGATAGTATACCCCTCGGCTGCTCTAGGATACAGCA

>NODE_1159_length_208_cov_304.288770

GCGGGAATAGCTCAGTTGGTAGAGCGCAACCTTGCCAAGGTTGATGTCGCGCGTTCGAATCGCGTCACTC

GCTGCCGGTGAGGAAAAAAATCAAGATGATGAACTTCTCGATAGGGGCGTAACGGCTTAACGCCGCTTTC

TCCCCCATGTGAAGATCCAACGACTCCAATTGATCCATCCGAGGCTCTGGAATTCTCGGTAGCCAAGG

>NODE_1160_length_208_cov_5.930481

AGCTACGTACCTGCAGCAGGACGTACCCGTGTATAGTTGTGGCCCCAGCCGATATTTCGCCCTTCTGAGG

CGTCGCTCTACCGCCGATATATATACCGTGGTACGCATCTAGCATTCATAAAAAAAAATGCGCAACGACC

CATACAACACGGAAGGTAAACAGCCAAACCAAACCGGCTACCCGTACGTATCCCATACAGGTTATTCG

>NODE_1161_length_208_cov_5.518717

AGTAGTTCCTTCCGTGGATTCGCTGCTGCTGCTTCCTCCCTCTCGAGTCTCGAGGACCTTCTTGTTGCTT

TGAAACTCCTGGTTTGATTCTTTTTCTACAAATCACTCTTGAAGTATCGCCTAACGGTTTGAGAAAAACC

GTAGAACAAAGACAAGGACCGTATCACAACCGGCAATCTTGTAGCAACGAGCCCGCAACGGGTGAGAC

>NODE_1162_length_208_cov_4.657754

CCGAGCGTGGTTACTCGTTCACGACGACCGCCGAGCGCGAGATCGTGCGCGACATCAAGGAGAAGCTGAC

GTACGTGGCCCTCGACTTCGACCAGGAGATGAAGACCGCCGCCGAGTCGTCTCAGCTCGAGAAGTCGTAC

GAACTCCCCGACGGAAACGTCATCGTAATCGGAAACGAGCGCTTCCGTTGCCCCGAGGTGCTGTTCCA

>NODE_1163_length_208_cov_4.406417

CCAACCTGAAGACCGTGCGCGAGCTGGTCTACAAGCGCGGGTTCGGAAAGGTGCACAAGCAGCGCATCCC

AATCTCGGACAACTCGGTGGTCGAGGGCACCCTCGGCAAGACCGGCATCATCTGCGTGGAGGACCTTATC

CACGAGATCTACACCGTGGGCCCGAACTTCAAGCAGGCCAACAACTTCCTGTGGCCCTTCAAGCTGTC

>NODE_1164_length_208_cov_3.433155

CGTGAAGGTCCGCGCGAACGACCTCGTGGAGTTCGTCACCCCGTGCCCGATGATCTGAGTCAGGGCCTTG

TTGAAGCCCTTCATGTCATCCCCGTGCTGGATGGAGCGGATGATGTCCAGGAGGTGGGGGTGCGAGCGAG

TCAGCCCCTGCCCGAAGATAATGAGGCTGCGCGTGAGCGTGTTGGCTCCCTCGACGGTGATAGCGATG

>NODE_1165_length_208_cov_3.187166

CGCTCGTTGCCGACCCGACCAAGGAAATTTCTGCGGACTACGGGGTGCTGATACCTTCGCTGGGCATCGC

TCTCAGGGGTTTGTTCATCATCAACCCCGAGGGGGTATTGGAGCAGGTGACGATCAACAACCTCGGCATC

GGGCGGGACGTGGACGAAACTCTCCGGCTCATTCAGGCGCACCAGTTCCTCGCCAAGCACGGAGAGGT

>NODE_1166_length_208_cov_2.101604

AGGTGGTGGGAGTGGCAATAGTCCGCATCCTTCTCGTAGTTCTCCTCCACCTCGCTGTCCTCGAATCCGA

GCACGCCCTTCATCGGCCCCTCTGCGGCGGCCTTCATCGCGGCACAGATCTCCTCGTAGGTGCACTCCTT

CTCGAGCTTGGCCGTCAAGTCAACCACCGATACGTCCGGGACTGGTACGCGGAAGGCCATGCCCGTGA

>NODE_1167_length_208_cov_2.032086

GGTCATGCCCTTTAAGTCGAGAACGGCACCCAAAAATAAACTAAACTCCCAGCCACAAGGCTGTCCTACT

AATTTGTGGACCAAATACAAAACCTTCCGATAAAAACACCAGCCAGGAACGCTACAACAAGTAGTACCCC

TGACACGCCCGTGGGTGGTGCGGACGTCTAACGACCTCTAGAAAGGTCTCTCGAGAGAACCGTACCTT

>NODE_1168_length_208_cov_1.481283

TTCTCGAGGAAGTTTTGCATCCCTGAACTATTGGCAAAGCGCGGCAATTGCACTTTGCAAGCGTTAGCTA

GAAGGAAGCGAAGGATGGATCTCTCAACTCAACAAGAGTACCCAGGGCTGAAGCAAGGGGAGGTTTCGTC

CGGGACGACGATCATGGCCGTAACCTACGATGGCGGGGTCATCATGGGTGCGGACTCGCGGACTTCTA

>NODE_1169_length_208_cov_1.192513

CTGCTTGATTTCTTTCATTCCTGGTACACTCGTATACCAGTCGAAGCTAGACTTGAGACCACTCGGCGCA

AGCGAAAAGAGTCCCGGTTTGCGTTGTCATGAACTCAGCGAACCGCCCTTGGCCTCCAAACTGACCTGGT

CAATTATGACGGGCTTCCTTTTTTATTATGGGCGCGCCCCACGCTCGTCGTCTCCATTCTCCAGAGTT

>NODE_1170_length_207_cov_1242.887097

TGGCACCCCGAGAATTCCAGTATGTGCGGCGCTCCGAAAACCCAGGCGCCAAGGGGTAGGTGCGTCGAGT

TCCAAAACGTCACGGTTTTAATCTTGCTTGGTTGGCGTATTTAAGAAGGGTGAGCGGGGCCACGGGAAAG

GGCCCCGTTCTCTCCTGCCCCAGCGGTGCTACCGAATACGCCCTCGACGACGCGAGTCGACGAGGGT

>NODE_1171_length_207_cov_118.672043

GCACCCTCCGTATTACCGCGGCTGCTGGCACGGAGTTAGCCGGTGCTTATTCTTATGGTACCGTCATCAG

TATATACATATACCTTATTCTTCCCATATAAAAGTAGTTTACAACCCATAGGGCAGTCTTCCTACACGCG

GCATGGCTGGATCAGAGTTGCCTCCATTGTCCAATATTCCTCACTGCTGCCTCCCGTAGGAGTCTGG

>NODE_1172_length_207_cov_30.994624

AAATGTTCCAGAAAACGCCAACACGTAAACCGCAATTCCCGAAGGTGCGGTTTGCGCTAGTAGAGGGGCG

GGTATACCGGTTAATCTCGTGGACGAGTACTGTAGCAAAACTGCGAACCGAAAATAGAGATAGATTCCAC

CCAATACACAAACCCGAAGGCTGCGCGGGGGCTCTAAAGTGGTGACGTGTTTGCACCGCCACCTCTG

>NODE_1173_length_207_cov_11.801075

GCTCCATGTGTTGATGTTTCTCTTGCGAGATACAGCAGTGATGGTTTGTTCTCGTGGTTTTTGAATTGGT

TAGATTGCGGTAGAAGACGCATCCGGGCGGTTCCCTGGCCGGTCGGTTAGTAGAAAGGGCTTTCGCTACC

ATCATGGTTCACTCGGTTTCAATTTCAATGTGCTTCAACATTTCAACTTTCATGGGGTGATCGGTCG

>NODE_1174_length_207_cov_5.876344

GAAAGATCCTGGCGGCCGCTTGCTCCCCTTCGCCAACATCCTTCGCATCTCCGCTGATCTGTCCTACGCC

GAAGGCTCCGAATATGACAGCGAACATCGCGGTGAAGAAGTTGTCGTACTCCACCGTGCCGTCGTCCACA

AGGATAGCGCCAACGTAGAACAGGAGCGCGAACACCCAGAAGATGATGGCCTGTGCGTAGCCAAAGG

>NODE_1175_length_207_cov_4.333333

TGATTTATAGCGGAAAGGAGCAGGGCTTTTGGGAGACAGAGAGAGGTACTCATAGAAAGGGTCGTGTGCA

GTATATGCCGTGGGGTTGGTCGAACCAGTTTATTTCACACATGCCATGCATGGAAGGAGGAGCTTGTAGG

CTCTACGTACGGGCGGGAGGCGATGGTTAACGGGATCTGGAGGGAGCCGGGGGAAGACACAGAGGAG

>NODE_1176_length_207_cov_3.983871

CCGGGCCCGCCTGCCCTCTCGTCGTCCGACCCGCCGTTCATGTTCATCTCGTGCTGGTGGGTCTGCCCGA

ACGTGCTGAGGTCAACGCGCTCAAGCATGGGTTCCTCTTCGTCTCCGTCGTACTCCGTCTCGGGCCTCGG

TGGCAGGACCTTCTCCAACACTTCAACGGCAGACGCTGACAGCGTATCCGGGAACGTGACCTTGAAG

>NODE_1177_length_207_cov_3.806452

TCTAGACTAATTTTAATAAATTAAAGTCCTTCTAGATTAATATTTAAAAATCTTGCTAATTCAGATGCTT

CATTTTCCAAAATCTCTAAAGACCTTGGTTGTTCCACAGGTGTTAGTGGAATTTCTCGTTGATCCTTTGT

ACATAAATAAATAACACGTTTAGGATTAATACCATCTTGTATATTTAATTTAATACTTTTAATATTT

>NODE_1178_length_207_cov_2.763441

AGACCTTATATACCGATTATCATCCCGTGTGAACAGCACTCTCTGCCGTTCCCCTGGTACACATGTCCGC

CATTACGAAACGCCAAAGCTTATACTGAGTTCATACGCATAGACTACCACCTGGGTGAGCATCGACCGTT

GCCTGGCCGTCGTACATCTGGTCTCTCGGACCCCATAGCTAGTCCTTCCAACACGAGAAAGCTGGGT

>NODE_1179_length_207_cov_2.639785

TGAGGTCGCGGAATACTGCCTCGTTCGTCAAGTCCAGTTCGTCGGATTTGTAGTCCGCCAACACCCAGGG

AAAGACCGGGTACTGGCACAGGTCGTTGTAGCTCCGGCCTGCCAGGGTGTTGAGACCCATGAGGTACTCG

AAGTTGCTGATGTGGCGTCGCTGCCACGCCTTCAAGAGCCGGTTGGCCTCCGATCCGTCCGACTGTC

>NODE_1180_length_207_cov_2.575269

CTCTGCCGATCGAACCCCCAGCACACTCGGGGATTACAGACAGTTCCGTTCATTCGGATCTGCCTCCATC

CCCGACGTCGCTTCAGGGCTATAGCCCTTCTACACACCTCCCACTACATTCCGAACCCACGGAATCCCGG

GACAACTTCCATTTTTTCTGGCATTCCTCGTCCATCTTCTCGCAGTGGTACCCGTGCAACATCGACT

>NODE_1181_length_207_cov_2.440860

TTATTAGTAGTAGCAGTAGAGGGTGTCTGGAGAGGGTGCATGCAACCTTCTGCATCTTGTTCAAGGCGAG

CCAGCCGCGAGCCGAGTCTTGTCAACTGGAGATCCTCTCGTCCCCCCCGCTTTTCAGCTCTTCTTCTGAT

GTGTGGGAGGAGCCATTTATTTCTGTTGCTGTTGATGCGACACTCGCTACTCGTAGACGCGGTGCTA

>NODE_1182_length_207_cov_2.263441

TACTCCTCTCGAAAAGCCAGATCAGCTGTACATGAACTCGATAAACTCTTGGATGCTGATGGTCCCGTCC

CCGTCCTTGTCTGCCACCTTGACCAGGAGTTCGATCTCCTCATCCGATAGCTTGTCGCCGAGACCCCGCA

TAATTTGCGCAAGCTCCGACGGCGTCACCTCTCCGTCCCCGTCCTTGTCGAAAAACTTGAACGCCTC

>NODE_1183_length_207_cov_2.241935

TAGCAGTGGTTAGCCAGGTGAAATCTCGTCTTGGAGCAACGCCTGTTCCAATTCAGTTGCCTGTAGGCGC

TGAGGACGACTTTAAAGGTGTTATTGACCTTATTAAAATGAAAGTGATTAACTGGAACGAAGCGGACCAG

GGTATGACTTTCTCTTATGAGGCAATACCGGCAGAACTTCTGGAATTAGCTGAAGAATGGCGCTCTC

>NODE_1184_length_207_cov_1.838710

TGATGATCAAGGACCCCCGCACGGAGAAAACCCGCGTCATCGACCCGGGGCACATCGGGAGGCAACTCAT

GGGATTCCGCCAGCTGGAAGCCCAGCTTGTCGTTGAGGCCATGGAGTCCGTCCCGCACGACCACACGAGG

CTCAACAGGCAGTTCTTGGAGGAGCGGATGAGGGATCAGATGGAGGAGGTTGTGCTCACACAGCGCC

>NODE_1185_length_207_cov_1.559140

TAATAAAATAGATATGCCTATTATTCCTGGAATGTTGACAAGTGTTGACGTGATGGTCAGTAAAAGGACT

GTACTTGAATATATACTTAACCCGATATTAAGAGCCAAAGAAATGGCGTTGCGTGAATATTAAATTTAGA

TATGGACATGGAGATAGAATGAGACCATTAAAACGTGGAATAACTAAAAATTTTTCTTTAGTTGCTT

>NODE_1186_length_207_cov_1.193548

GCTTAGTCTACTGGCACTGCTCAGGCTACTGACACTGTTCAGACTGGTGACACTCCTCAGACTGTTGATA

CTGCTCAGACTACCGTACTCGCACCGATTTATCTTGCTCGCGCGCTCGCGGGGGCTGGGTTCGCTTGGAT

TGAATTGGCTTGGATTTGGTTAGGTTGGGTCTTGGGTAACTTGGCTTGTGTTGGTATGGTTTTGCTC

>NODE_1187_length_206_cov_39.318919

TTCCTCGTATATCTGAGGGCCTGCCCCCGGTCACGGGTCAAAACTTTTTATGCGAAAGTTCCGTGCTGAT

ATGGGCAGGTCTCTTCGTTCCAAGAGGTAAATAAAATTATCGGCCGAGATGCTGGAGAGTCGCGGCCCCG

GGTTTCCTATCGTGTTTGGAAACCCGGGGCGGTGGCGGCCTTCATTATCCCACTTTGCCGACATTC

>NODE_1188_length_206_cov_24.978378

TCCACAATACTACAATGATTATCATATATCAATTAAATATCAAGCACTTCTTTGAAGTACAAATTATATA

ATAAACATATGGCTGATAGCATTGCCTTGAGGCACAAGTTTGACGAGTCAATCCGATTCAGTTTATATAT

AATATGTTTCTTCTTAAATACAAGTATTTAAGCTTACCTCCATAAAAGGAGAACATCCAGGCTGTG

>NODE_1189_length_206_cov_19.178378

GCGCTCTAACCAACTGAGCTAATAGCCCTTTTTAGAAGTTTTTTAACTTCTTTTTTCGTTGTCAACAATG

AAAGGTTTGAACCTATAAAACATTAGAGACAACTTGAGAAAACAATTTTATCGACCAATTATGTTCCAAT

TAAGGAGGTGATCCAGCCGCACGTTCCCGTACGGCTACCTTGTTACGACTTCACCCCAGTCACTAG

>NODE_1190_length_206_cov_7.891892

TACACACCCTTCTTCATCTACAAACCTCTTGACGAGAGCAACACACTCTTTCGACTCGACCATCATCACC

ATGGCTTCCACCACCACCACCACATCGTCATCCTCTACCATGACCCTGACGTACTTCGATTTTGCTGGAC

CTGCTGAGCCTGTCCGATTGGCGTTGGCGATGACGGGCCAGCCTTGGGAGGACAAGAGGATTTCTT

>NODE_1191_length_206_cov_6.037838

AGAGGCCTGCTGACCCTGAGGCAGAAATCAAACCTCCGGTCCCGGCTCCGATGGCGATCAGGTTGTACGT

CCCATCTGGTTCGGCGTCCTTCCACTTTGCGGGGTGCACGTTGTCGAGGAGCGTCGTGTTGTACTCGTCG

ATGGGTGACACCCCGATTTCAACGAGGGTGTCGATCATGTCGCCATCGCGCGATACCACGTTTTCC

>NODE_1192_length_206_cov_4.037838

CTTTCTCATCCGAGAAGAACCGTGACAAGTTTCTCGCAAACCCAGCCAAATACGCACCTCAGTATGACGG

CCACTGCGCATATGGCGTTGCCCAAGGCGGCAAAGTTCCCGGCAACCCGACCCTGTGGCGCATCATTGAC

GACAAACTGTACCTCAACATCACCAAGAACGTTGTTGGCTTCTGGGAAGCGGATATTCCGGGCAAC

>NODE_1193_length_206_cov_3.821622

TGTCCCCCGGTCCCTTCTGCCGTATAGCAGACCTTGAGCATCACCATGCAAATCTTCGTGAAGGCGAACT

CTACCATCGCCTTGGGCGTGGAGCCATCCGACACTATCGATGGCGTCAAGCAGATGATCGAGAGCAGGGA

GGGTATCCCCGCGGGCGAGCAGCGTCTGATCTTCGCGGGCAGGCAGCTGGAGGGCGACCGCACGGT

>NODE_1194_length_206_cov_3.589189

CCCAACATTTAATCTGTTACGTGTATTACCGAACAAATTATTAGGTGTTTTAGCTATGGCTGCTGTCCCA

TTAGGCTTAATAACAGTACCTTTTATTGAAAATGTTAATAAATTCCAAAACCCATTTAGAAGACCTGTAG

CTTCAACTGTCTTTTTAATAGGAACATTTGTTGCTATATGGTTAGGAATCGGTGCTTGTTTACCTG

>NODE_1195_length_206_cov_3.194595

GAGCCCAAGCGCAAGACATACAACGCCCGTGTGCCTCTAAACCAGGACATCTGTATCTAAATTACCGTGT

ACAAGGGAAGGGAGGGGATCAATCGGTTAGCGAAGAAGTAGAGTTTGAAAATAGTGCAAGGCAAACGGGA

CGTTACGAGTGAGCTCTAAACCCTCTTGTTTGTTTCCGGAACGAACGACCTCGTCATGCTGAGGAT

>NODE_1196_length_206_cov_2.594595

GAGGTACGCCCTGAACAGTGTCTCGCTGTCAGGCTCTTCGAACAGTTTGGTGAACGGGAACTTCCCGATG

TCGTCGAAGTCGAAGCTCGCCGGGGCGTCAAACCTCCTGCCAGCGTCGGACGTTGAGTCGCCGAACACGA

CCATCTTGGGGTTTTGGCGCCATTTCTTGTAGTATGGTGGCCGCTCGGCACTCGAGTCGTTGCTCG

>NODE_1197_length_206_cov_2.502703

CGCCGAGGTCGCCAGCATGCTTCTTGTACAGCTTGGCCATTGCATCGGAATAATCAGCGTTCAGCTCCGG

TGGGTTGCCGAACATAAATTTTGCAGGGTCTATCGCGGCTTTGCTCTCGGCAGAGGACCGCGTTGCCATG

GCGTGGATCAGGTCCCTCTCCAGCTCCGAGTAGGGGGAGTCTTCGACGTCGACCTTCGCCACGGCC

>NODE_1198_length_206_cov_2.491892

CACTTAAAATGTAATCGCCAACCGACCATTCTTCGTCAATATCTGACGCTAATAATCGCTCGTCTATGCC

TTCATAACGACCTGCTATAAATATCAGGCAGTCATCTTGCGCTAATTCACTTACACCTGCTTGGTCCAGT

TTTCGTCCCTGCGGCGACAAATAAATAACCTTCGCTTTTCTACCATTTTCTTCTGCAGCTTTACGC

>NODE_1199_length_206_cov_2.351351

ATGACCGAGTTCATGACGGACAACGGTCGATCGGAGTCCAGCGGCGCTGAGGGCAGCGTTGCTGTAGGAG

AGAGGCCAGCAGAGGCAGGAACCACTGCTGTATTCACTGCTGTAAGAGAGACTACTGCTGCTGTTGGAGC

ATGCATTGGCCAATCGGAGTCCGGCAGCGCCGAGAGTAGCGCTGCTGTAGGGAATAGCCCAAGAAG

>NODE_1200_length_206_cov_2.135135

TCTCTTCTTTTGGTATTTTGCTTGATTCGCGTTGAATGCGGAACTACCAGCAGGAAAAATGTTTTTTGGC

TTCTCCGAACATTTTTGTCTTGTGTTTGTGTATATGTGTTGTGTATCATATGTTGTGGGTTTGCATATGT

ATGTGCGTGTTTAGTTTTAGGTGGAGTGGGTTTCGGGTAGGAAACAAAAAACGTTCGCTAGATGCG

>NODE_1201_length_206_cov_1.643243

AAGGTTGTCCTCCCAAAACACCCTGGGCGCCTTGCAGCCGGGGTAGCTTTGGTGGTAGCACCACTCGGGC

TTCCAGCTCATGGCGAATATGTACAGGCCGAAGTCGCCGGGGGTGCCTCCCCTGACAGCGAACGTCTCGC

TCTCGTCGACAACCCGGGGCTCGAGATGTGTCTTTCAACGACACCACCACAACAGCAACAACAACA

>NODE_1202_length_206_cov_1.562162

CGGGTGTTGGCTGGCCCAAAATGCGCAAGCAAGTGGGCGGACAAGGGGTTTCCTTTTTATCGAGGGGAAG

CGACGCAAGGGGGCACAACTTCCTACCAGATAATGATAATACCAAGGTATATATATACCAGGTATATACA

TATCATCTATTTGTGGGATTTTTGGGACAGTTGTTTTTGGTCGCCCATCGTCGTAGGCCTGCGATT

>NODE_1203_length_206_cov_1.535135

ACCGGTGGTTTTGTAGGTCTGGGTGCGGCATATCAGGCGACTGACAGCATCACAATCGGTGGTGAGATCC

TGCAGAACCGTTTCAACAACGTTGGCAGTTCGGACAACGATATTGATGCGACCACGTTCAACCTGCGTGC

AAGCTACCGCTTCTAAGAGTGTGACTGTGATCTGAAAGATGCGCCCTCAGCGGGGCGCATCCTTTT

>NODE_1204_length_206_cov_1.091892

ATCCAAGCCACGGACGGACAGTAAATGAGAGCATGCCCATCGCCATCCCGGGGAGCGTAAGCATGACTCG

GGAAGGAAGGAGGGAGCTCGGCAGTGGTTTCCAGATTATTACCCAGTGTCAGGATCAGCTCTTCGGCCAC

CGGGAGGTTAAAGAAAACTTTGCAAGCGAGCCCGGGAACCATTTCATGCACTTTACCAGCGAACCC

>NODE_1205_length_205_cov_5777.125000

TTACAAAAACTCTCGGCGAGGCGCCCCTCGCCCAACTTCGCATAACGAGTTCTTTCTCGGGGTACAGAGG

CCGCCGAAACCGGCGGCTTCACAGACAATTTATAGAGCGGGGGGGATTGAACGAACCCGCTTTCGGTAAT

GATCCTTCCGCAGGTTCACCTACGGAAACCTTGTTACGACTTCTCCTTCCTCTAAATGTTGAGGT

>NODE_1206_length_205_cov_268.505435

GGCACCCCGAGAATTCCAAGATGTGCGAGCTCCAACGTCCCGCGGCAAGACTGGTCAGTAACCGCAATCG

GAAACGACTTTCGAGCACAGGATAGCGAGGAACCACAGCAGACATTTTGTCCGCTTTAATCTTGCTCGAG

TATAGTGGGAAATACAGGCACTGTCCGGAGACGCCTGGGTCCCGGCTGGGGGCCGTTCGAGCATC

>NODE_1207_length_205_cov_69.195652

AACTCTGTGACAGGTGCTGCAAGGCTGTCGTCAGCACGTGTTATCAAATGTTCGGTTAAGTCCGGCAACG

AGCGCAACCCACATCTTTAGTTGCCAGCAGTTCGGCTGGGCACTCTAAAGAAACTGCCCGTGATAAGCGG

GAGGAAGGTGTGGATGACGTCAAGTCCTCATGGCCCTTATGGGTTGGGCTACACACGTGCTACAA

>NODE_1208_length_205_cov_56.586957

CGCTTTCCCAGCAGCTTTCCCATGGGGCTCAACCACCGCTGCAAAACGGTGGCCGTGCCATCGGGAGGCG

ACAGAATATACGGCGTACTGGTGTTTAATCTTGCTGTGCGAATGATAGGAGAGGGTCACGGAGGAGGACG

GAAGAAAGTTCCTTGGAAAATTCTCCCCGCCCCCTCGACCACCACCTGAGAATTTTTTCAGCTTT

>NODE_1209_length_205_cov_26.380435

ATCGAGGGTCTTGCCGCTGGATGTGATGGTGCTGTGGTGCTTGCCGTTGTCGAACACGTCCAGCGTCCTG

GTGTTCTCATCCCCCTTGTGGAAGGCGATGCTCATGCTGACGAGGTCCTGGGGTTCCTCGAAGGAGTAAG

TGATGCAGCACCCATCGTCCTCGTCGGAGCTGTCGAGGAGTTCTCCCTTGCACGACCAGCGAGAG

>NODE_1210_length_205_cov_17.282609

TCGGGCTCCCACTACTTGTACGGGTCCATGGGCAACTTCCTGCCAGAGGCCGTGTTCTGCCAGTACTTGG

TCCCTTCCGAGCTGTAGTGCCTCTGCCACGTCCCCGTCTTCTCGAGCCTCTTCCTCGACTCCTCTTCCTT

CCGCTTCTTCTCGGCGTGGATCTGCTTCTTCACGTAGTCCGGCACCGCAACGGCCACCGCGGGCC

>NODE_1211_length_205_cov_5.239130

TTGTTTTGGGATGTGTGTTTTTGATTTCTCCGTCCGTCCGTCGCCTTGGCGGTGCGTTATGCGTCGTCCT

TCCCTCCTTCCTACCTCACCGCCTTGTGGTGCATTTTTTTTCTCTGTTGGATCGTGGTTGGTCGTCGTTT

CGGTGGGCGACTGTGGAGGACAAAACGTCTCTCGTCGGCTGCGTTTGTTGTGATAGCTTCTCGGC

>NODE_1212_length_205_cov_5.217391

GAAGGACCAGGCCGGCGATGTCGCGCGTTCCACGGGAAAGGCCGGTCTAGCGGTCGTCGACAAGGGCAAG

GAGATCAACGAGCGTTACCGCGTCACCGACAAGGTGAAGGAAGCTGCGCAGGGAACGTACCAGAGGGCGA

AGGAGGTGGACGAAGAGCACAAAGTCGTGGACAAGGCCAAGGCGGGAGCGTCGAGGGCATGGATC

>NODE_1213_length_205_cov_3.885870

GCGCGCATCACGTTGATAATCGCCTGGCGCTGTGCGAACAGGGCGTTGACGACCGGAGTTCCACGGGGTA

CGAGGGGCGCCTTCAGCATGTACGACAGCAGGCTCAGCACCGAGTGGAAGCTTTCCCACTTGCCGCCCTC

CCTCTTGACCTGAATCCGTTCACACAGCTCGGCCATGATGACGAGATCGAAGATAAGAGGGCTGG

>NODE_1214_length_205_cov_2.652174

AGCTCTGCAGCAATAGCAGCAGTAGGAACTCCAGTAGTAGGCTATTGCCTGTGTTGAAAAAGGAAATATG

GCTTCGAGACCCCAAGACCCGGACCAGGTGGACAAGACGTCCACCCCCACCCTGTCGGCGTTTGTGATGA

GGAGCACCGGAGACCACGAGCTGGCCATCCTCATGAACGCGATTTCGATCTCTTGCAAGGCCATC

>NODE_1215_length_205_cov_2.152174

CGGTGGGCGATCGTAACCGTCGTTCTTGGCTTGTCCTTGTGTAGCTTGTCCAGGGCGGCTTGAACGACCT

TTTCGCTCTCGGAGTCGAGGGCGGATGTAGCTTCGTCGAGAAGAAGGATAGCCGGGTCTTTGATGATCGC

GCGGGCGATAGCGATACGCTGCTTCTGCCCTCCGGATAGCTGGAAGCCGCCTTCTCCCACACCCG

>NODE_1216_length_205_cov_1.701087

TGATTGATGGTATTATGGCCTGAGGCTGGTGCCTGCCTACATGGTTCCGTCTTTGAACAGATTCGATATG

TCGTTCGCTGGCATTGTTTGCAGCGCGCTCATGTTATTTAACGGGATTCACGTGTTTACTATTCGAAAGG

CTGCAGGTACTGTCCTAGCACATGCTTAAGAACAAAGATTAGGTTGTCCTTTCCCGCAGCCGTGG

>NODE_1217_length_205_cov_1.532609

ATTGCATTAGCACAAAAAGAAGTTGAACTTGATAATGCTACAGGCACATTAGGTGAAATGTTTGGTGTTT

CTCGTGCAGCAGCTGCTGGTGCTTACGGCCAAATCGCTACTTCAATTGTAAGTGCAGAATTTCCTGGTCG

TGGTGAGACATTAAACCGTATCGCAAATGCTAAAGAAATTCCTGCTTTAGAAGATCTTGAAGAGT

>NODE_1218_length_205_cov_1.184783

ATTCCAACCAACAGACCGAAAAAAGACGAGAGCTCTGCTAGCCAACGCCTGCTATATATACTATACATAT

ACCTAAGAGGTCTACCGTGAGGAGGCAGCACCAACAACAGGAGACAAAAAAAAAGCATGAACGGTCTGCT

ATCCAAACGCCTGCGGCCTACCGTGTAAGATTCGCAGCGGCCCGCAGCACCAAACAAGACACTAA

>NODE_1219_length_204_cov_2849.825137

TGCAGCACCTGTCACAGAGTTCCCGAAGGCACAAGTCTATCTCTAGTCTCTTCTCTGGATGTCAAGGGAT

GGTAAGGTTCTTCGCATGCTGTTGTTGTTCCGCGGAACAACAACAGCATGCGAAGAACCTTACCATCCCT

TGACATCCAGAGAAGAGACTAGAGATAGACTTGTGCCTTCGGGAACTCTGTGACAGGTGCTGCA

>NODE_1220_length_204_cov_5.032787

GGTTGCAGATCGACTCCATGTAGGGGTACCCATCCTCATTCTGTTCCATCGTTTGCGCTGCATTTGCGAA

GCTGTACATCTCCAGTTTCTCCATTGCCCTCTCGAGAATTTTGTCGTCCGCCGCATCATTCCACAGGCGA

CACAGGAGGTCGCTAGCGATGATGGTTCCCTGTGAGTGCGAGATCAGAACAACCTTGGTCTTTT

>NODE_1221_length_204_cov_3.956284

GGCCCCTCGCTTCTTCCAGCCAGCGTTCCAAGTCCTCCGGCGTTATCTCGGGCAGATTGGGATAGTACTT

TAGGCCCATGTCGTTTCCGGTCTGTTGTCTTCTTCAAATGTGTTGGCGTAGAATCAAGCGTAGAAGAGAA

GCAGCTTTTATTGTTCCCCGAGCCAGATGACATCTGGCTCGATGGGGTGCAATTCAAGTATGGC

>NODE_1222_length_204_cov_3.579235

GGCAGGCAGTTTCGGTTGTGGTCGCGATTGAGTTCGTTGCGGCGGTTGTGGTTGTGGTCGTGGTAGCTGC

TGCTCGTGTTTGTTTTCTTCTCGTCCTCCGTGTCCGTCGTCTCGGCGTTCTTCGTGCCGCTGTTTTTGTT

CGCCTTGTTCCTCTTTATCTCGCGGACGTTCTTGTTGCCCTCGCCTTCCTGCTGCCTCATTTTG

>NODE_1223_length_204_cov_3.169399

GCAGCAGACCCTCTCATAGACCCCGCTTCCCGCCGGTCTCCGGGATACGCGGTATTGTTTTCAAGGAACC

GAAAGCTGAACATATATTTTGGGCCTTCTTCTCTTTGCTTTGTACCAGGTACACGACAGGCACGAAGAGA

ACGTGAAGGAAAACCTCGTTCAGAACATGGCACTCGGGCATGGAGTAAGCTCCCAACCCTCCCC

>NODE_1224_length_204_cov_2.808743

GCAACGACGAAACCCGCAAACGTGCACGGTAGTCAGACGGGAACCTACAGCAGAGGGATGGAGACGGTAG

CAAATTGCCAAATACATTCTAAACATGTACGACGCCAATACATAGTACCAGTTCTTCAGCAACCCTGTGA

GTATGAACGAATACTCTCCGGCCACTATTCCAACATCTACTAATCGAGCGAGAGGCTGTGTCTC

>NODE_1225_length_204_cov_2.606557

TCTTATTTCCCCGGTGTACTATTTGTAGAATGGCTCAGCATTGTCCGGGTTTTTTTTTTTACCCGTTCAT

CGCGCTATTTGCGCGGTGGAATAGCATGGGACGGAATGGGATCGGATGGGACGGACGAGATGGGATCGAA

TGGGAAAATGGGGGACGATATGAAATCCGATGGGAATCGAGAAGAAATGGGATGGAAGGGAAGG

>NODE_1226_length_203_cov_2279.961538

AACCTGGCAGCTTCACGAGGCGCGGCTATATACCCTTTTGGTGGGAGTTGCGCCTTTTGGGTTTCGTGAA

AGCCTTCGGGTGGGATCGAAACCCGCTGCCTAGATTAAAATCAAAACCAGCTTTCCACACTCACCGAGCC

GGAGCTTCGGCGAGGATGTGCTTGGCCTTACACTGCTGGACATTTTGGAAATCTCGGGAGCCA

>NODE_1227_length_203_cov_29.824176

GGAGAACGTCATCCGCGACTCGGTCACCTACACTGAGCACGCTCGCCGGAAGACCGTGACCGCCATGGAC

GTGGTGTACGCCCTCAAGCGCCAGGGCCGTACCCTCTACGGTTTCGGCGGCTAAAGCTGGACGTCGCCCA

ACTCATCGTCTGCGAGCCACCTGACGCTCATCACAAAACCAACTGGTGTTTACTCAACACCAC

>NODE_1228_length_203_cov_10.142857

TGGATGCGCAGACAAACGCCGACGACCTCGACGCGGACCTGGGACCGGATGACGTCGAGACGCACATGTC

GGTGTTTACATTCCTGAAAATCGTGACTTTGTTCTCAGCGCTGTACATGGCTACGAACCCGTGGTTCTTG

TTCTACGACCCCGACAAACAAGAGATCTGCCCAACCACGGTCCTTTGCGCAACAACCTTCGTG

>NODE_1229_length_203_cov_4.642857

TACAAAACTCGTAAGAACAAGAAAACTGATCAGTATATTGTTCGTCGTCGCAATAAGAAATAAGTAGAGG

TATCGCCATGCCACGTTCTCTCAAGAAAGGTCCATTTATAGACCTGCACTTGTTGACGAAGGTAGAGAAA

GCGGTGGAAAGCGGGAATAAAAAGCCAATTAAGACTTGGTCCCGTCGTTCAATGATCATCCCT

>NODE_1230_length_203_cov_3.868132

TGCCGTTGTGAGGCGGTGACCAGTCCTGGGAAGCTTGGCAGGAGAAGAGAACACAAGAACACCTTGCCCA

TTGTCCGGATAGCTGAGGGTGCCATAGTGTCGCTCTAGTATAAATTTAGTCCGGTTGCCGCTGTCCGTGC

CTCTCTGTGCCGTGTGCCGCGAGCAGTTACTTGATCTTGTTATTTGGCGAAGTGGAGTCCTCG

>NODE_1231_length_203_cov_3.670330

CATTGCGGTGAAAACGTTCGCCACAAGATGATTGATGGCTTCTTTTCTCTTCCTTCCCTCCTTCTCCCGT

CCCCTACGCATATTTCATACGATGCCGTACAGCCTCCACACACCAAAACACACGCAGCAAGAACGAAAAG

ACATTACGTCGTACGAACTTCATGGCAGGAAAAAAAAAAGGGCGGAATTGTTCGTACGTACGG

>NODE_1232_length_203_cov_3.532967

GCAACGCAGCACGATGGTGCGGAACACAGACTTGTGATACATATACAACACCATTTCCCGCCACTACTAC

AAGAGAGCCCATCAAAGGAGCAATCTCAGACTTCACCACAATTCGCACTACCAACATCCAAGCGGCTTGC

ACATCAAACAAAGGATATGACCTTCCCGACACCAAAACATACAATACAACAGGCGCACTCGTC

>NODE_1233_length_203_cov_2.906593

AACCATGAGGATATTCTCAAGCGAGAACCCCAACATATACCTCTCTTTCACACCAAATAACGTAACTCTC

GTGCAATCAATAGACTACAGGCCGACTACAGTCGTACGCAATACCTACCTACAGCCATGGTTAGCGAAAC

ATTCATTATATATCAACACCGTCGTTCTTTCAGCGGTACCTACTGGGCATGGGCGATATCAAT

>NODE_1234_length_203_cov_2.730769

AAAAGCTTGGAAACCAAACCGTCAGAGCTCTTGAACTTGTCGTGTTTCACCTTCGACAACTTGGGGTGGA

TTACGCGCTTGCAGTAGCCGTTGTTGCTCTTGTTACGCCTGTAGTAGTCCTGGTGCCCCTTGTCCGCCGG

GTACCACGCGCCGGCCGTGTTCACTTCTGTTACGATGGGCCTCCCGAATTTGGGCTGTACCGC

>NODE_1235_length_203_cov_2.604396

TGGTGTTCGGCAACATCGGCGCAGCATACGGAACCGCTAAGTCCGGCGTGGGTATCTCGTCCATGGGTGT

CATGAACCCCGGACTGGTGATGCGTAACATCATCCCCGTTGTTATGGCGGGTGTGCTCGGCATCTACGGG

CTTATCGTCGCCGTTATCCTCTCGGGTTCAATTGGAATGCCAACGGAAGCCGGCAACACGTTC

>NODE_1236_length_203_cov_2.263736

CCCCCCCGCGCCTACTCCATTTCCCCTCTCCCCTCGATGTGATACCGCCAGGCTCTCTTTGTCTCTCCCT

TCCCCCGCCCGGTGTGATGGCCCAACTCACGACATGTACGATATCACGCCGCGGATATCTCTTCGCGTTG

ACCACTTGCACGCAGTTGCAGAAAGCGCTTGTGCTCATTCGTCGTGGCTTCGTTCGAAGAAAA

>NODE_1237_length_203_cov_2.230769

TCCAGCTCGACCGTCACGAACTCCCACTCGTGCGGGGACAGGCCGTCTGTGAGAGTGAAAGAGTACTTCT

CTGTGCCGTCGGCTTCCATCAACTTCAGGTATACCCTTCCCCTCTGGTTCCCGTATCCCTGATCCTTCCA

CCTGAACTGAACGGTTCCGGAAACCGGCTTCTTCTCCAAAGGCTTGGAGTTCCACTTCACTTG

>NODE_1238_length_203_cov_2.208791

CAGAACCAGAAGAACAATCAGCAGCAACAGCAGAAGCAGCAATTTGGGTACATTGACGGAGAGATCGTCT

GCACGTGTCCGAATACTTGTTTGTGTGCGTTGGAGAGCTGTTGTACAGGTCGCCCTCTGTTAGACGCCGG

CGGCGTTTTCAACGCGGTCGACCGATGGTCAGGTGTAAGCAATATTTGATTGAAGGTCCGAGC

>NODE_1239_length_203_cov_2.186813

TCTCCGGTGCTAGGGCCATGATGCAGACCTACAAGATGCACAAAGAGCTGCGCAAGACGACGGTCACTGG

AGCGCTCGTCAGACGCTGGCAGGGAGTCTTCGAGACACTGATGGAGCGAGAGACGGCCAACTTCGAGGAC

ATTTTCAACATCAAGTACTTGACCGGCAGATCGATCAGCGACGCGCTCAACGAGAGCTTGTCG

>NODE_1240_length_203_cov_2.005495

CGGCGTCCTCGAAGTGGCGGGGGAGGATCTCCGGCACCGTGTCCTCCTCCTCGACGTCTTCCATCGCGTC

CCCTGCCTCACCACGGAGGCGTTCGCGCTCCATGTCCTTCATGATGTTCTCCCGGATAGCGATCTTAGCG

GCACGCTGGCAGATCTCCGTCAGATCGGCACCCGTGAACTTCTCCATCTGGGCAGCAAGGTAC

>NODE_1241_length_203_cov_1.934066

TCGCGAGGAGCACCTTGCAGCCGCTCCCCTCCTTCTCCAGAAGGATCTTCTCCCTCTTCTCCAGCATCTC

GTCATCACACACCCTCAGCAGTTTAGCCTCCGTCGAGGGGTTGAGGTAGTTCGCCACTCGCAGCTTCTCC

GACTCCAATGCGTTCTCCGCCTTGATCATGTAGTCAGGCGTGGAGTCTGTCTCTATCCACTCT

>NODE_1242_length_203_cov_1.307692

GACTTCTACGTGCAAGTTTAAGTTTAGGCGCCAAATGGCTCTGTCGATGACGGCCTCCGCCTCTTCCTGG

CGACGTTTAAACAGCGTCGACATCAGAACCTGCGACACTTTGTTCCTGCTCTGGTAGATTTTGATTAGTT

GCACCATATCGGCGACGGCATCGTGCACGTCCTCAATCATTTCAGACTCCGACTCTTTTCTGG

>NODE_1243_length_202_cov_980.182320

GGCACCCGGAGAATTCCAACATGTTGCCAGCGTTCAGCCCGTCGGGTACAGGCAGGCGATGTGACCACCG

CCCTTTTACCCCACGGTTCCTATCATAATACGGGCAAGCTTCTATCTGCGCAGGTCATGGCAGACGAACG

CGCCAATGGTTTGGCGGAACTCTGTGGCAGATAGACCTGCGGGGTCCCAAGGCAGCCGCAAC

>NODE_1244_length_202_cov_11.292818

CCCCCTCTCCCCCGACTGCAAAACGTCACATTCGACCGCACCTCAGAAACAAAAACTTTTCAAAATCTCC

GCGACGCGGTGCCAATCAACGCCCGCTTCATCAAGACCGCCAGACCTTCTCCCAACATCACAAGACGTCG

CCCCACCGGGGCAAGGCAACGCGCAGGGCATCACATCCAGCCAGCCGCTTCAAATGTCCAAA

>NODE_1245_length_202_cov_5.541436

ATCTTGTGTTTTATAATATTTTACAAAATCACCTTTATAATCACCCCAAAGTAAAAGCGTCAATTCATTT

CTAGAATTTTTTTGTCGAGGAGCTGGAAATTTTACTTTTATTTCCATAGCTTGGCATTCCTTATAAATTA

AATGGACTGGTTTTTCAATAACTTTTACAACAAAAATAGCATAATTTAATTATGATATTATG

>NODE_1246_length_202_cov_4.093923

CGTGCCGGCCGCTCCGCAAGCGGACTTCCTCCCTCAGGACGACTACAACAAGAAGCTCGTCGGGAACGTC

GGCCCCAAGGACTACGTCAACCCGGAGCCTCTGGAGAAGTACGACCTGGTTGTTATCGGCTCGGGCGCGG

CCGGTTTGCTGAGCGTGATCATCGCCAACGGCCTTGGAAAGACGTGCGCTCTGGTGGAGCAG

>NODE_1247_length_202_cov_4.055249

AAGGATGGATGCATTTTTGACTCCGGCGCTGTCATTTCCCATGCCACCAAAACATCAACACGACCAAAGG

TGGCCACATCGATCCTCGAGGATTACAGCCGCGCCACGGCAAGGCTAGTACGTACACGCCCACCCAGTTT

ATATAACACAATAAAGTGACGTCCAGTTCCCAACCCCGCCCCCTTTCGCCTCGCCCTAACGT

>NODE_1248_length_202_cov_3.508287

TTTGTCATGTACTGCGATATCAGCCAAAATCTTACGATCGATTTCAACAGAAGCCTTTTTAAGACCATTG

ATGAAACGGCTGTAAGATAAACCATTTTGACGAGCTGCTGCGTTAATACGAGCGATCCAAAGCTGACGGA

ATTGACGTTTACGTTGACGACGATCGCGGTAAGCGTATTGACCTGCTTTAGTTACTGCTTGA

>NODE_1249_length_202_cov_3.033149

TTGAATGAAGGACTGCTATTTAAGTCGTACCTCCCTGGTCACCTAAGGAAGCGTAGCACATTCCGTGATC

AGCTGGTAGTACCTAAATCCCTTCGAAAACTTGTCATTAATTCGTGTCACGACCTACCAGCGTCCGGAGG

TCACTTAGCTTTTAAAGCTACCTTCGATACTATTCGAGATAGATTCTGGTGGCCGATGATGC

>NODE_1250_length_202_cov_2.895028

ACAACCAGGTGACGTTCGAGGTGGAGGGGAGCGCTCTCGCGAACGGCTCGTGGGAGAAGGTGACGAGCGT

GGCGTTCTTCTTCAGGGAGGGCGACAACCTCCCGGACACGGCCAGCACCATCGTGGCCGTCAACTGGCTC

GTCGTCTCCAACAAGCTCGTCCTCTAGAAAAGCAAAAACCACCACCACTGCACGCGGTCACC

>NODE_1251_length_202_cov_2.839779

TGCAAAATCGTAGTCTGGCCCAACACAGTGCGAGTAAGTCAACACGCACGCGGGCTTACTCACACTGACA

TGGTGTCTACACACACAATATACAATTCATGGGTACACACAACAAAAATCAAAACTCAAGGACACGGTCA

CGCAACACAAATAAGCGAGACACACAACGCAAATGAAGGGCACCCACACAAAACAACGAAAC

>NODE_1252_length_202_cov_2.359116

TTTTACAGTATCTTTTGTTTCCCAACGGACGTTTGAACGGAAGAAACGAACGAACAGGCCCCAAGCGAGG

TCGGCCTCGCCCGCGTTTGGCTTTTACTACATCCCGCGCGCTTTCAGGGACAGGGACGGTACACATTATT

TGTTTCTCGTGTCTTTCATGCGTACTCGTGTCACGAGTACGTCTGTTGCATGTATCAAAAGT

>NODE_1253_length_202_cov_2.287293

CGTCTCCTGCTGGCCCCGGGCCGAAGACCGATTCCATCTCTCCCGCGAAAGGTTGCGCGGGGGAGAGATA

CGAGTAGGTATCCATTGAAGGAAACAAATCCCCCCAGAGTTCAAGTTTTTTATATGAAAAAAAGAAGCGG

TGTTCTTGCTGTGCTCTCGTCCTCGTGCCTGTTTTTTGCTCTTCCTCGAGAGCTGCGCACGG

>NODE_1254_length_202_cov_2.276243

GCTTTACCGAGGGCTGAAGGAGATTGACACCTTGAAAATCGCTGACGAAATCACCGCAAGCATCATGTAT

TGAGAACGCCGTCGTCTTTACGCGCAAAGGGAGACGGCAGCAAAAGAAGCAAGGGGGGAGGGGGAATGCG

GCAGTTTGATGACGTCCTACTCTGCCCTACAGTTCCCTTTCACCTCACAAATTTTGGTTATC

>NODE_1255_length_202_cov_2.215470

ATCCATGCTTCCTCACATATCACCTGATATACGATAGCCAGGTATATGTGCTCGATATTCCACAGCCACA

AGTGTATATATGAAATCCTCTTCCATGGATCATTACCCGGACATACCTACCAGTTCGAACAAGTAAGCGT

AGGCGAAAACCTCCCGACTTCGCCGGTATGGTCCAACTATTTCCCCAACAAACGTAAAATCC

>NODE_1256_length_202_cov_2.127072

GCCTGTGAATGCAAATTAACAATACGCATTCATACCTGGGGTGTCCTGTGCCCCCTGGCCGAGGGAAGCA

TCTTGAGTGTGGTCCCAGAGACCTGCTAAGCAACAAGGGTGAACAGTCGGTGCGTAAGGGCCGGAAATCT

ACTAGAATAATGCAAGGTTGTGTCATGTAGGGTACGTACAGAACCATACTCGGGGTGTTTTC

>NODE_1257_length_202_cov_1.900552

CGACGGGAGCGCTGGAGTCGCTTCTGGATGAGGAAGACTTGAGGCAAACGGGGTGGGAGGATTGGCCCGG

GGAGTCGGTGCAACAGAACCTGGTGATGGAGCAAAGGTATGAGCAACTGCGGAGGAGCAACTCCGAGCGG

AACGAGGGCCTGCTCGAGCACACGAGAGGAGGCGACAAGAACAGATTGTAGTATACTGGAAT

>NODE_1258_length_202_cov_1.596685

CCCTTCGCCTGCTCAAAAAGCAATACAACATGCGATGAACAGGTACTCACGTGCGACACCAAGGTACCAA

CGTGTCAAATGCGTAACCCCAAGAAACTGTTACCCAACCGCTACACACCAGATGCCCTATATGCTGCTGT

AAGGGTACTAAACACTTTGGAACGTAGATGCGCGAAACAATCGAATATTTTATCGTCCACCG

>NODE_1259_length_202_cov_1.469613

ATCATTCATCCTGAATGATACACTCATCCTAAGTGGTTCCTTTTCCTTGTTCGTCACTTTCCAAAGTGAA

TGCCGTCTTCCCGACCGCCGTCCTATTTGCTTCCAATGCGCCCAATTCCGTTTAAGCAACTCGTCCTGAG

TATGTCCATTTCCTTGTGCTCAATATTGAGCGACTTCTTCCAGAAGTATCCTTGTCATCCTT

>NODE_1260_length_202_cov_1.458564

ACTTTGTCGGCGCGGAATCATAGGAGCAGTTCAAATGGCTATAGTAAAATGTAAACCTACTTCTCCGGGT

CGTCGTCACCTAGTTAAAGTGGTTAACTCTGATCTACATAAGGGTAAGCCATACGCACCACTTTTAGTGA

AAAACTCTAAATCAGGTGGTCGTAATAACAATGGTCGTATTACGGTTCGTCATATCGGTGGT

>NODE_1261_length_201_cov_11.400000

GGGGTTGCTGGGATATATAGTGTGGCAATTTTGTTTTTGTCGGCACTACGTTACGCACCTTAGCCTATAC

TGAGTACTCGTTTTCGTTTTTTTCTTCTTCTTGCAAGCGTTTCGGCTTTCGATGGGAATGGCAGCAACGC

TTTTGAGCGGTCATGAATGAACATAAAAAGAGAGATACAACAACCAAGCTACACGGGATAT

>NODE_1262_length_201_cov_6.394444

GCTTCCTTGGGCTGCGTCGGCGACCGCTTCAGCACCTTCTTCACGCCTCCCGCCCGTGCGTCCTTCTTAG

CCGCCTCGTTCTCCTTCACACGCCTCTTCAGCTCGTCCCTGCACTTAGACGGGGAAACGTGCTCCACCCG

TACGTGGATCTGCTTCGAGATGATGCGACCGTTCACCTGCTTGTTGACGCGAAGGCCCAAG

>NODE_1263_length_201_cov_3.483333

TTCCAGCCGAGTCTCGAGCCGCTGCCCGTTCTCGTGCATGTGAAGGCCGTGCGAAAAAACTGTCACTTCG

TTGTCCTCAAAGTTCTCCTCGAAGCAAGAGCTCGGGCACTGGAAGGAGTAAGCAGACTTCCCATCCGACA

GGGGTTCTTGGTCAAGAAGGACGAACGGATCGCCCAGCTGAATCACCCCCATGTTTATCGG

>NODE_1264_length_201_cov_3.222222

TCTTTTATAACTTTTGGTGTATTCTCAACTAATTCTTTAGCTTCTTTTAAACCTAGTTCTGTTATAGAGC

GAACTATTTTTAAGATAGGGATCTTTTTATCTTTAGGCACTTCATCTAAGCTTACATCAAACTCTGTTTT

TTCTTCAGCTGCTTCACCAGCGTCAGAAGCTGCTCCACCTTGGCTCATCATCATAACTCCG

>NODE_1265_length_201_cov_2.966667

GGCGCCGTGGGAGGCGCCCTAGCAAACTCTGTCGTGAACGGATTCTTGGGGTATACCGAGCTGTCCTCCA

GCCCTGTCGCTCGCGCCGCCGCTCCTCGTTTCCCGAGCCCCGTGATCATCAGCGAGAATTCGTGGGCGAG

GGTGCTGATCTCGTCTTCACCCCTGCAGAGCAAGTAGTCCACCGCGCGCGTTCTGCGGCTG

>NODE_1266_length_201_cov_2.594444

TGAACAAAATTGGCATTTTTTTAAATTATGTTCGCCTGTGACACAGAACAAAATAAGATGGTCCGATAAA

ATTGACTGTTTTGCCTGCAAGCACAGCTACGCAGAGCCCAACCACGACAACTGATAAGCGTTCGACCACG

GCAAGGAAGTACTCAATTGCGGCTACGTAGCACCCAAACACGGTTATTGCGCTCAATCACA

>NODE_1267_length_201_cov_1.561111

CTTCAGGACGCGGGTGAACGCTTTCGTGATCTCCTGATCCATCGAAGGGCATACCCGGTCCAGGAACTCG

ACAACCGTAACCTTCGAGCCCAGTCGAGCCCAAACAGATCCCATCTCCAAGCCAATAACGCCACCGCCGA

TCAGCACGAGGTTCTTCGGGACCTCCTTAAGGTCCAGCGCTCCCGTAGAGTCGACAATCTT

>NODE_1268_length_200_cov_4.966480

TACATAGTTCACCGTATTGGGGGGGGGCGCGTTTGACATTCGATAGCTGCACTTTTAGAGTTTTTTTTCG

TTTTTTGGTAGAGAGGTGTGTCTGGGATAATACCAGGCAATCGTTATGATAGCGGTCACCCTAGCCGGTA

ACCCGGCGTTTCTCTGTGGATTCCGACGAGCAGCCCACTCGAGCCAAGCTCGTCGTGTTT

>NODE_1269_length_200_cov_4.139665

AAAAAAACGTACGCGGGTTGTTGGTGACGGGAGTTCCAGTAGTTTGTGCCGAGGCTTTTTTTCTCTTGAC

TTAGTTCGGTTGCGGTTGTAGCATCCTGTGTTGTAACGCGTTGTTGAGGGCTTCGGCGGGTTTGTTTTGC

CGTCACTTGCTTCTGGTTTGGTGACCGGCGGGGGCAAAGAGGGAAAATACACTTTAACTC

>NODE_1270_length_200_cov_3.826816

GATAGCTGGACTCACGCAGCTATAAAAAATCATAGCAGGGCGAGCACTACACCAAAAGAGCGTTTGGTTT

GAGTCGAAGGGTCTAGCTAGTCCTAACATCAGAGACGCCAATCCACGTCTCAAAAAACAAAAACATTTCG

CCGCGTCCCGAGCCAGCAAAACAACGCGCACAAGCCCAAAAGCGCGGCTCCTTCGCAAGC

>NODE_1271_length_200_cov_2.770950

AGCGCAGAGACAGGTGCAACACTTCCACTACGGCGTCGGTATCCCCTCACGAGAAAGTGCCGGGGTTGAC

TATCCAGCTGGAGTGGTATTCAGCCAGAAACCAACCGAAGAGACAGGACAGAGATTCATTGGACTTGAAG

GGGCGAACTACGTTACATCGATCGAGGGGAAGGGGAGTTTCCCGTCAGCAGGGAGATTCT

>NODE_1272_length_200_cov_2.603352

GCCGCCTAACGCAGCTTGCCGACAAAAAAAAATGGAGCCGAACTCCACGGGATGTGATGACCGTGCACCA

GAAACAACACTCGTCGTGACGCTAGTGCTAGCAATCACATCACCCACCAGACGCTCAGGTAGCCAGCCGC

CCCTATGAATAACAATGCCTGCCAGGGTGGGAGCTTGGCTCACACGTCAGAAATGGCGGT

>NODE_1273_length_200_cov_2.402235

TGCAGCCAGCGAGCAAACACGCACAAACAATATCACCATATCAGACAAAGAGTCTAAAATAAAGGCAGCG

ATACGACGGTTACTCCTTCTGAATTATAATCATCAGATCACTTTTCACGCCCTCTCTGTTCGCCCTCTCT

CAAAACACCGGAAGCCAGTCAACAACCCCATCCAATCCCCCTCCCCCCCCTTGGATAAAA

>NODE_1274_length_200_cov_1.385475

GCGCCCGTCTCACCGGTCAGCACATTAAGACCCGGCTGGAAGTTCAACTCCAACCGCTCAATGATCAGCA

TATCCGAAATATCAAGACCGCGCAGCATTAGGTGCGCCCCGTCACAGGCCTGGCAGGTGCCACGTTAAAT

CCACTCGCCTTTCACCATTTCACGGTAAATTCCCGACAGCCAGTTATTGCCTAACGAGCG

>NODE_1275_length_199_cov_114.101124

TTCCTTGGCACCCGAATTCCATGGTACATTTTATAGGATTCGAACCTACGACCCACGATTTAGAAGACCG

ATGCTCTATCCACTGAGCTAAAAATGCTTAGATATAAAAATTTAAAAAATAATTCTTAACTACTGTTAAT

GAAGAACAATTGCGTCGTTTATATAAATACAACTTAAAATTGTAAAAACATAAGCTTGA

>NODE_1276_length_199_cov_20.432584

CTTGGCTCCCGAGTATTCCAATGTTCGGCCCGTATCTGTCCACCCGGCTGTTCTCTGCCCGCGTAGTCAT

CAAAGGTCGTTTGCGCGGTACGCTGCTCTTGGGGACGAGATGAAAAATAAGGCGTTTTTGAAGCGCCCTC

TCGTGGGGAGGCAGGGCGGGAGGCACGAAAGAAACCACCGCCTGCACTCCTTCCCCATT

>NODE_1277_length_199_cov_13.162921

ATGCATGCGCGCACGCGAATTGCTTTCTTGTCCGTCCCGCGATCCCACACCCTATTTCTTGCTAGGAGTA

CGAGATAGTCCTGACTGATTCTGAGCGATTGCGGCGGTTCGTTTCCTTCTGCGCTTTGTGTCTGTGGGAA

CTAACTGGGGCTGCTGAGACTAAACGAGGATTAACAATTAAAAAAAAAAAAAGTGGAAT

>NODE_1278_length_199_cov_4.870787

GGTTGCAGGAAGAAGGGAATGCAACCTCATTTCTCAGTCTCAACAGTCAGTCTCAGAAATGTAATCAGTC

GCGTTCGGTGTTCCCTGGATAGTCTCAGTCCTCAGGATAGTTTGTGAGATTTACGACAACAATCTGATGG

CGCTCGCTCGTGTTTTGACATCTGACACATACTACGTGCAGGAAGAAGGAAACGCTTGG

>NODE_1279_length_199_cov_2.516854

AGTGGATTTAATTGGAAGCTGTTATTTATTATTTATTTTTTCTGTTTTTTTCCACGGAAAATCGACTGTA

TACAGCAACTTTAGGCCGTATTTACTCTACAGTAGTATTAACAGTGTACGTTAAGCCCGTGGCAGTTCGT

CGAGCAGCATCAGGTTGCGATTGTTCTTTGTTCGCTTGCCATTGTAATAAATCATCCGC

>NODE_1280_length_199_cov_2.438202

GTGACGGTAGTACATGTTACTGCAACAAACGCATAATTGACGCGCATCCTGCGGTCTTCCTTGTCTGGAC

ACTTCATCTAACAGCTGACGACGACATCATGACACTGGTCACTGACACTATCCCCCCCCCTGGGTAGAGC

ATCTAGTCAGCCAAACAATACTCAAACAATACTCTTAACCGGTGGTGTGCACCGCCCCG

>NODE_1281_length_199_cov_2.359551

TGCTGACCCAGCGCACCCTGCGTGGCTTTGCACGGGGGATGCTGTAATTGCCTTCCATCGACTCGTACAA

CAGTTGTTTTCTCCCTGTGAAGACAAAAAACGGATACGGTGTATCATACCATCTATCATACGTACGTTAA

CCTCGTTCTACAATATGCAAATCCCTGTCATAGGGGACTATGAATTGACGAATACATCC

>NODE_1282_length_199_cov_1.983146

CGGGCGTTTTTGAATCAAACGTGGCTTCACCTTTTATTTCCCCTTTCCTTCGTTCCTTCCAAAAACCCAC

GACCTGAAAATATTCCGACAAACAAGAAGGCGGTGAAGATGGCGGCCACCGGGTACAATACGGCTCAGGC

TGCCGCGATAGTTTCTCGGGCGAACAAGAAGGAGAAGCTGTTTCCCGTGTAGAAGGCTG

>NODE_1283_length_199_cov_1.752809

TTTGGTCGTAGATCGACCAGGATCAGCGAGCGGACGAAGTGTATGTATGTATGTTGGTATTTACAGGTGT

GCGGAATAACTACAAGTGCTGCACAACACAGCACGCAGGCTAGATAATGGAAGGGCGGAAACAATAAAAA

AAAAAAATTAAACTAACCACAGTCCGCATTAACACAATAACCAACAATATCGCATATAT

>NODE_1284_length_199_cov_1.601124

CGCGACGACATTTATCGGGATCATGCCTCTAGCCTTTGCCAGCAACGCCATCTTCCGAGTGTTCTTCAGT

ATGTTCCTCATCATCATCAGCTTTGGGTTCTTCCATGGCTTGGTTTTTGTTCCTGTAGTGCTGTCCCTCA

TGCCCGAGAGCCGAGTCACGGCCAATAGTCCTACACTGCGGCGGTACGGCGTCTCCAAT

>NODE_1285_length_199_cov_1.589888

TGCAATAGAGATCCCCTTTACCCCTAGGGCAAAAAAGGTATTAGAGATGTCTATAAAACAATCTAAGGAG

CTAAACCATAGTTATATCAACACTGAACATATTTTTTTAGCATTATTAAATGATACAGATGGTATATGCT

CAAAAGTTTTACAAAATTTAGGTGCAAATATACCCCGAATTAAATCTTATGTACTTAAT

>NODE_1286_length_199_cov_1.421348

AGTGACGGTTCCCGGCAGGGATTTCAGAGACATGTACACCTCGATTCGGCACTTCCTCGACAACTACCGG

ATGGTTACGGACAGAATGGACGTCGATGAATCTGAGGTTCAGTTCTCGGCTTTGAGCTTCAGCGCTACGT

CTTTCTCCCCCAAGTTCCTTCAAGTCCCTTCTCGGCGAATGGCGGTGGTGCGCCTTGAA

>NODE_1287_length_199_cov_1.224719

AGAGCCAGCCGAGTTCAACAGGAAGGTCACGCGCAGCGGCTCTGATGATCAATTCCGGGTGCGCCTTGGC

ATCTCGCGGGTCTTCAACCTGAACTTCTAAGACATCCGGCCAGATCGCGTTTGTCGATCTGGTCGATGAT

TTGCCCTAGATTGCGTAGTAGTCACGATACCACTTCACAAATTGTCACTAATGCAAAGG

>NODE_1288_length_199_cov_1.196629

TCTGCCCTGAAATATCTAGACCATGACGTAAGAATCGATGGTCTTTCCGAGGTGTGAAACAGTTTACAAC

AGTGTATGTATACAGTACGCGACAGATGTTAAGAATCTGATAATTAAGAATTTGAGACTGAGATTTTCGT

GGGAAAAATCAGAACGCGACCTCTTTGCACTGAAAAAAGTCTGGGACACACCCAGTCTT

>NODE_1289_length_198_cov_2464.548023

TGCTTGTCGGAGTGTCTGTTATTGCGCATGTACTATGAGACTGGCAAAACCTTATTGCCAGCTTCAACAT

CAACTGCGCATAGAACAAACGGGAGTACTAAAAAAGAACTCCCAATCGGACTAAAGCACTCGTTTGCCCT

CGAATTTACGGGCAGCGGGCCGTTGCGTTCGCAGACATGCTGGAATTCTCGGGGTGCC

>NODE_1290_length_198_cov_64.344633

CGCATTTCGATTGGAGGAGCTGGTGGGACTCGAACCCACGTCCATTTAAAATAATCATCAAACTAATCAA

TGTAATCACAGATTTAGTTATCAATTAGTTTACTTTCGTTACATTTAAAACGTTAAATAAACGAATTTTT

AACAAGAAAGTTGATGAGTTTATGTTTATTTTATATTAATAAATAAAGCTTACAAATA

>NODE_1291_length_198_cov_38.954802

TGGCACCGAGAATTCCATCAAACTCCACAATTATCACAGCGGCTCAGCTTCTTGTTTTTTTTCCACGCGC

GCTCCGCCATCCCGCATAATTTCACAGCAGGACGACGAGCAGGGCACGCACTCAAAGATCATGTATAAGA

TGCGCCGTATCTGTGATTGTTCTGTGGCGCCATCATTGCCGCCAAGGAGATTGGCGGC

>NODE_1292_length_198_cov_30.429379

TGGCACCCCGAGAATTCCATAGCCCTCATGAAACCTTATATCTTCGGCATCGGCTCTTTCTCGCCTTTCA

CACACCAAGCAAAAGCAACGCCACCCTCAAAACCATCGCTGGCCATCAGGAGCGGAGCCTGCCGGTGATC

GAGAGCGAGCCCTGAGATAAGATCATGTAAATTTTGTAATCATCGGGCTCGTCCCTCC

>NODE_1293_length_198_cov_23.864407

TTTTTTTTTTTTTTGATTTTTTTTTCATTTTTTTATTGCTCATTGGTTCTGCTCTCGGATGTTTCGACTA

CGTCGAACCGATACCGTCACCCCCCCCCTCCCCTCCCTCTCCCTGCGTCCGCTGGCTGGCTCTTCGCCTA

TACTCGGTGCAACCAAAAAACAACGCAGCCCGCTTCCTAGCGAAGAACGTATGTATAA

>NODE_1294_length_198_cov_6.711864

ATAGTACTCATTGTGTAGTCATTCCGAAGCGGATTTGCCCTTCTAAATAACTCAATAGCAGAGGCAGGCC

CTCGCGACATGGTCTGTTGGTCCCTGGCTGTTACCACGTTGGTCGCTTTGTTGAGTTGCTCGAAGGCACG

AACGAGTGGAACACATCAAGGTGACGAATATGTTTATGTGGGGTGCGCGTGCCGCACT

>NODE_1295_length_198_cov_4.615819

CCGCCGACCTGGAGGTGTGGGAGAGTCAGACAGGCGAAGGGGACATCGCGAACCGTGTGTTTTACTTCGC

GATCCCTCCCAACGTCTTCCTGGACACCGCCGCGTCCATCAAGCAAGTCGGCATTAGCGCCACTGGATTC

ACGAGGCTCGTGGTGGAGAAGCCTTTCGGTCATGACTACGAGTCTGCTCTGCAGCTGG

>NODE_1296_length_198_cov_3.881356

GCACCAGTAGGCAATTCCTGGATGTCTGCCAGCCAAAGTGTAATCATAGTACACTGGGTTTAGTATTGAG

AGGGGTTTCAGGGTCTACGCACAAGAACGAGTGCATCTATACTCTTCTCCACTATCTCGCCAAACAACAC

CTGTGCGCAGTGCAGGGGAGGGTTCAGGGTGTACGCTCAAGAACAAGTCCTTATAGTT

>NODE_1297_length_198_cov_3.446328

TCCAACTTGTACACCATGTCGGAGGTGTACCCGAGGTTCTTGAGTACACCCGTCAACTCCCCCTGGCTCT

TGTCCTCGTTCTTAGGCCCGGAAATTGCGTTGACCATGCCGGGGGGGCCGCAGATGAAGATGACGTTGTC

GTCCGATGGCTTGGGCAGGTGGGTTTCGATCAAGGAGGCGCTAACGTACCCGGTCTTG

>NODE_1298_length_198_cov_3.033898

ATTTTTTTAAGGTTAAATTACTCAAATATATTAGGATATACACATGTCTACTACTACTGGTACAGTTAAA

TGGTTCAACGAATCTAAAGGTTTTGGTTTCATCGAGCAAGAAAACGGTCCTGACGTTTTCGCACATTTCT

CTGCTATCGCAAGCACTGGTTTTAAAACTCTTGCTGAAGGCCAAAAAGTACAGTTTAC

>NODE_1299_length_198_cov_2.197740

GTTTGCCTCGAACGATGTGACCGTGGGCTTGACGCAGCACGTGTCCCTCACATCTGCTGGCAAGACGGGG

ACAACTGAACGCAGTTACACTCAGACGACGACTACCGGGAGGCACACGCTTAGCTTTGAAGACCAGGATC

TCTTTGCGGGAGCGCTTCTTTACGCGACGTTTAGCATCGACGCCTTTGAGTACCAGTC

>NODE_1300_length_198_cov_2.101695

TCTGTATCGCTTCCAACAATTGCTGATTAGTTTTGGCCATGATCGGCACTGTAGTCTAGTTTCAATCGCG

CCCAGCAAAAAGCGGGTGGTCGTTAGCCCTGACCAGCCCTGTTGTCAAGTTTCTATCGATTCTATGATAT

GAAGTACACTGGTTGTCGTAAGGCGTGATCATAAAATTGATTTGCACTTCCGTTGAGG

>NODE_1301_length_198_cov_1.926554

TGGGGCAGGCGGGAGGGAGTGTACGGGGGTTGCTTCCCGCTTCGGAAACGCCGCTTTTAGATGTAACGCC

CCGGGGTGTAGGAAAAATGAAACCGGGCGGGTTCTGGGCGCGTAGTGAAATAAGAGATGGATGTGGATAG

GGCCACACGCACCGCGTCCGCGTGCAAGGTCTGTTGGGGAAAAGGACATGACGAGGTT

>NODE_1302_length_198_cov_1.694915

CCTGAATGAAATATCGTTTGTTAGTGAAAACCATGCGAAACCCATGCTACCACACGTCCATCCTGCAAGC

AAAAGTCCTACAGTCTAGTACTCATACGCGGGCTGAATTAAGTTCAGCTCGTATAATCAAGCGACCGAGA

TGCAGTGTGTGCGCTTTCAACCGATTTGCACCGATTGAGCATCTAGTATCATTACCAA

>NODE_1303_length_198_cov_1.621469

ACACGACATTAAAAATTCACCATTACAAAGACCATTTACAAACCAGAAAACAAAAACAAACCCTAACCAG

TGCCCAAAGCCCATAAAATATTTTCATAACCGCCTCCCGGTTGTGTACATAACAATTTTCAACCAACAAC

AGCAGTTTTTCTGTAGTCGGTACTACTGCTGTAGTGTAGAATACACAATATCCGGAAA

>NODE_1304_length_198_cov_1.259887

CAGTCAAAATCAGGTTGGTTGCCTGATTGCACCAGCGCAAGCGCCTCGTGCGCAGAGGAGGTCGTCGTCA

ATTTATGCGTGCCAACGATCTCGACAAACTGAGAGAGGAGCTCAAGGATGATTGGATCATCATCTACTGC

CAAGATCTTCATTTTATCTCCGTGCTTATAAGGCTGCCATTGGACAGCTTGTAAAAAT

>NODE_1305_length_197_cov_86.198864

TCAGCGGGAGCGTGTGCAGGGTCCGGGACTCCCTCGACTTCCTAGACCCAGTGGCGCGCGCGAAGGTGAC

CCTCGAGACGCGCCTGGCCATCGCCAAGAGCGAGTTCGCGGTCGGACCTACGTGCCACATCACCAATGGC

GACGAGGAAAATGTCCCGCTTTTCGCTGGGCAGTACCACAAGACGCTCCCGCACGAC

>NODE_1306_length_197_cov_24.931818

TGATGTCATGAGGCTCTCATAGCGACCACGTGTGGAAATGAAGGATCAGGCGAGCCAATGAATGGCCGCC

CGGTCGAACGAACGTTGGCCACGCCGAAGTGCCTCTCGTGGAATTTTCTCTAGCCCCCCCCTGAAATGCC

GCTACACCAACGCCGGAGCCGCACCCCCTCTTATCGATCGATCCTTTTCAAGACCGA

>NODE_1307_length_197_cov_23.034091

TTGGCAGGGGCAGCTTCTGACGCTGCCCTGGTCTGGGAAGGCGTTGGCTCGGTGCCCATCACTTGTTTTG

GTTGCATTGGCTCTGCGGGGCGGGTAGATGGTTAACGTGCTTCCTTGTTCTGGGATGGCGCTGCTACTGT

CGAAACTGTTGAAACGAGGTGGCCTTGGCTCGGGTGTGGCTGTGGTGGGTCACGGGG

>NODE_1308_length_197_cov_11.511364

CCCCCCAATACCTCCTCCATACGCTTCGGCGTTTCTTTCTTCGTGCTTGCTCTATCAATATCCGAGTCGA

ATATCTACCTCCACCGCCTCACCTATACACGCCAACGTCATCTCTTATTTAATGCCCTCTCGATGCCTGT

TTCTCTCCGTCCACGTCGAACACACACAACAATAGTACACGTCCAGGCTACACGCTA

>NODE_1309_length_197_cov_6.732955

GCGCGACTGACGTTACATTCCAGTGGCCTTAAGCAAATCCGAGGGAGCTGGCTCTGTTTTAGCCCTGGTT

AATTTAACCGGCACCCACCTGTACATACAGGTCCTCGGGAATACAACCGACCCGGCAACGGCGGTCCCGC

CACACTTTTGACCTAGGGCGCAGGCGCAAAATATCTCGCGCTTAGCGCACGCGCAAT

>NODE_1310_length_197_cov_6.619318

CACTCTCACTGCCCTCGGCGCCGCCGCTGAGGTCGAGAAGGTGAAGGACTCCAAGAAGATCCGCGCCGGC

AAGGGCAAGATGAGGAACAGGCGTTACGTCATGCGCCGTGGACCCCTCGTCATTTACGAGAAGTCCGAGG

GCATCAACCGCGCTTTCCGCAACCTCCCCGGTGTGGAGACGGCGTGCGTGGACCGCC

>NODE_1311_length_197_cov_5.170455

GACCCTCCACTGGAAGCTCTACCGCAACTTCGCCATTACCCACCCTCTCGACGAGGGGCCGGTCAACAAG

GCCGGAGACTTCTTGTTCCGGCAAATGATGGAGCTGACTCTGAGGGAGGACAAAGAAATAATCGAGAACA

TTTACCCCGAGTACCAGCGCGGCTTCATGAACGCGCGCTATGATAAGCAGGTGGAAT

>NODE_1312_length_197_cov_4.221591

GGGGGGATTTTGATGGCAGGGGTCATGTCCGGCACACCCGTCAGGGGTCCCAGGAGGTCCGCGATGGTTT

TGGTGCCCGCCTGCAGCTCAGCGATGATCGCCATGGCCTGAGCATCAACGCTCCCCTCGGAAATAGGGGT

CGACATTCTGGGTGAGAGCCTGTTATTCCTCGCTCTGCAATGAAAATTCGACAGGGG

>NODE_1313_length_197_cov_4.000000

ACGGAAACGTTTATAACGAGCCACTAATCTAGCTCGAATAGCAGGACGGTGGGACGGACAAAATTCAACA

AAATCCCCTTTTTGAAGGGTAAACCCACCATGCCCTATTTTTTTACCATTTACCAATACCTTACTATGAA

GAATAGCCTGGCGACTGTAATAAATTGAATGGAAAAAACCTAAACGGTATAAACTAA

>NODE_1314_length_197_cov_3.107955

CTGGGGTGTGGTGGCGATGGGGTCGAGGATTCTGGGGGCGAACGAGGAATCGACGACGCGATATCTCGGG

CTCATGAGGACGGCGCTCCGGGAGTGCTTCGACTGCAACGACAAGGAGGTGGTCCAGGCGTACCTCCTCC

TGTCGACGGTGGAGACGATGCGCGGCAACGAGGAAGCGGGCAAGCGCTACCTCGACC

>NODE_1315_length_197_cov_3.034091

TGCAGGTGGACGGAGGAGGTAGTGGCGAGGAAGAGGAGGGCAAAAAACCAAAAACGACGCCATACAATTT

ATAAAAAAGGCGGAACAATGCCGGCGCGTGAGAATGGAGTGCGAAGGCAACAAAGCGCGGCGTTATATCT

ACTCGGGCGGGAGGGAATGTTTGTTCACTTTTCCTCGCGAGAGGAATAAGGAGGGGG

>NODE_1316_length_197_cov_2.710227

TCCGTGGTACTATGCTTGCAGGTTGACTCTGACCATCTCATTTGACTCACAGACACAGCAGGAAAGGCAC

AGCTGCTCTTAAGTTGAAGCTGGTAAATAATGTACAGGAAATAGACTGGGCAGTCTAAACATGTAATGAT

TGCATGGGATGATTCAAGAAATTGCAGACTGCTGTTGCTGTATTATACTCATAGTAG

>NODE_1317_length_197_cov_2.522727

TGGCTGACGCTGGACACGAGGCTGACGGAGGTCTGCACGCCGTCGAAGGTGACGATGCCCATGTGGTCGG

GGGACTCTATCCGCGCCGGGCAGAACAAGGGGAGGGCGTACCTGGAAATGGAGAACGAGGAGGATACGGA

AACGGCAATCAGCCTCAGCGGGAGGTCTGTCGGGGAGCGCTGGATCGGCGTTGCCAG

>NODE_1318_length_197_cov_2.096591

TTGGTCCGGACACGCTTTTGACGGAAACCAGTACGTACGTAGCCCGGGCTGCTTTCTTTTTTGTGTGTGG

CCGCGTTGTTGCCGTATGTAATCGTTGTTTTGTTTTGTTTTTTCGTCGCTTTTCTCCTCCATACCACGGA

AAGAACACAAAGGTGTGTTCGGCTGTGGTAAGCGGTACGTAGCCTTTGTTCATGGTG

>NODE_1319_length_197_cov_1.863636

GACTCGTGCATGCATGCATACCGATTTTTTCGTGAAAAATATAATAGGGCACAAATACTACATACTCTGC

AGCAGTGATCGCTGGATGCGTCGTAGTTAGGCACATTGAACGCACGTACCATACGTACCATCTGATCCGT

ACGAGACTGTACTCTGTACCATGGTGTTACAGCAGCGCGCGTAAATCTGGAATTTCG

>NODE_1320_length_197_cov_1.767045

GAACTCTCGCATGATAACGTCACGCTCCCCCTGGCCCATGTCCCCGTGCATGCACGAGATGGTGAAGTCT

CGCTCCTGCATCTTCTCCGAGAGCCAATCTACACGCCGCCGGGTGTTGCAGTAGATGATGGCCTGTGTGA

TCGTCAGCGTCTCGTACAGGTCGCACAAGGTGTCCAACTTCCACTCCTCGCGGTCAA

>NODE_1321_length_197_cov_1.681818

GCTGGCCAAGACTATGTACCTTGCTTGGCATGTACATTTACCTACAAGCAAGTACAATTTACATGTATTG

GACATTACTTTTGGCATAGCGGGTTCCGAGAAACATGCTTTTTTTCCAGGAAGGGTTTCAACCTGCTTAT

ATTGGCGTTCGGGTTGTAGGAGGTTGTGGTGTGCTGTCTGTGTGGAGCCTGTAGGGC

>NODE_1322_length_197_cov_1.352273

GAGCTGGATCGTACGGTGTGGAACTTCAAAAGGAGAAAAAGTACGATAAGGAAGGGCTGTTGCAGTATCG

TCCACTGCTTGGTTGCGGGATGATACTGTTGTTGGCGTACAATGCGAGGACCTGTTGTAACTGTCGTTTT

TTATGGTCGCTACCTCTGGGGTCCTGTAATCGAGGCCCACGTATATACGATAAGAAA

>NODE_1323_length_196_cov_758.725714

AACCTTCCGGCACCGGGCAGGTGTCACACCGTATACGTCATCTTTCGATTTTGCACAGTGCTGTGTTTTT

AATAAACAGTCCCAGCCACCTGGTCACTGCGGCTCCCGTCCGCTTAGAGAGCAAGTCTCATCACAGATAG

GAGCGTACCTTCTCCCGAAGTTACGGTACGATTTTGCCTAGTTCCTTCACCCGAGT

>NODE_1324_length_196_cov_72.565714

TGGTGGGTAGTTTGACTGGGGTGGTCGCCTCCAAAAGAGTAACGGAGGCTTCTAAAGGTTCCCTCAATAC

GGTTGGCAATCGTGTGTAGAGTGCAATGGCATAAGGGAGCTTGACTGAGAGACCTACAAGTCGATCAGGT

ACGAAAGTAGAGCATAGTGATCCGGTGGTTCCGTATGGAAGGGCCATCGCTCAACG

>NODE_1325_length_196_cov_14.691429

CGGTCTCCCACTCATATTTAGTCTTAGATGCTCCTTACCACCTCCTTGGTGCTGTACTCCCAAACAACAC

GACTCGCCACAAACGAGAATAAAAAGAAACCCAAGTAGCCACGGGACTTTCACCCTCTATGGTCACGATT

TCCAACGTGTTCTAGTCGTCTCTCAGGCTCGGCTGATCCGCGTTCGCTCGCCACTA

>NODE_1326_length_196_cov_12.011429

CTCGCGACTTAGCAAATCCTGATAAACTTCGAATACTGTAAAAATTATACCGGGAGTCAGACGGTGAGAG

ATAAGTCCCATCGTCAAAAGGGAAACAGCCCAGATTACTGTCTAAGGTCCCTAATTAATAACTAAGTGGA

AAAGGATGTAAGATTGCTGAGACAACCAGGAGGTTGGCTTGGAAGCAGCCATCCTT

>NODE_1327_length_196_cov_10.245714

GGTTAGGTTATACAGAGTTTACAATTTTTTTGGGGTCTGGTATCAACAAAGGGCGTAGAGGAGTTGTAAG

AGGGCTTAGTAAAAAGGGGGTTAAAATTACTTTTTTGCATTTAGGTACACGAGCCCCCCATAATGGGTGT

CGTCCTGTTAAAGTTCGTCGTAAAAAATTTCGTACAAAGCCTAAACTATCTAGGTA

>NODE_1328_length_196_cov_6.200000

GTTCAAATTTCTGGCTCTGTAGACAATTATCCAGTGCCCGCAATCACTGTTGATACTGCTACTGATATTT

CTGTCGTCTCTCAAGCATGGTTGATGTCTCATCCAACCTTGCGTTCTGTCGTTATTCAACCTGTTCCCCC

GGCCGCCGTTGCCTTGCGAGCAGCGAATGGGTCGCCGTTGAATATTTTAGGGTTTG

>NODE_1329_length_196_cov_5.302857

TACATTATACCTGGTATACGTATATATATATATTTTTCTGGATGTAAATATTCCGATGCCTGTTTTCGTG

AGAGCGTTGTTGCTACAGTAGTCGAAGCAAGCAGGGCTGTAACTGTTTTTGTTGTTGGGTAGCTTCAGGG

ACGCTCATTTTGGGTTGTGGTCGCGGCAATTGTTGGCGTCTGGGCGCTTATGCATT

>NODE_1330_length_196_cov_3.548571

TTCCAAAAAAAAAAAAATGGTCCCACCACGCCACACTCCACAAGAGATCTCAAAAGATGCTCCAGACTCC

CCCCCCTTGAACCAGACATGAAAAGAAAACACTAAAATCGTGTGCAACCAGGAGAAAGACCAGGTGCACA

CGTACACACGTGCAAATGCACACGCATACGTAAGAAGAAATAAGCAGCCCCGTAGA

>NODE_1331_length_196_cov_2.971429

AGCCTCATCTTCGTGTTGTGAAGAAGGATCTTGGTACCGTATGCCAAGGTGAACTGGCAGTTGCAGAGCT

CCTCCGCATCTTCGTCGTCATCGTTCACTTCCTCCGCGACCATCATCTCGGCACACTTCGCTTTCAGAGC

CGCGTCCGCGCCGCTCGGGAGCTTCTTGGCCAGCAAGCTCAACAGCGTGCGGATCT

>NODE_1332_length_196_cov_2.508571

ACTGGGTATTTGAAGAAGATAAGAACAAGAACAAGAACATCACAATAGAGAATAGGGAACAACATGCGGA

GCGGCTCGTCGGGTCGATGAGGCGCTACCTCAACGCGAGCAACAACGCAGGAAGGGTCTGGTCGTTGAGG

CAAGAAGATGACGGGGAGGACGATGACGAGGAGGAGGAGGAGAAGGGTGTGTACTT

>NODE_1333_length_196_cov_2.360000

GTCGTTGAGCAGGCTCTCCCCCTCGATCAGTGTTCCCAGCTTCTCGCTTACGCCCAGTTCCTCGAGCAAA

GCCACCACTGCGACGGGGTCCGTGGCGGCCACCATCGACCCGAAGACCAGGCAGAGGTTCCAGTCCCAGT

CGTAGGGCAAGACGAACTTCACGAATGCCGCCGTCAGTCCGGCGCCGAGGATCACC

>NODE_1334_length_196_cov_2.302857

CTGTTATGAACAATATATAGTTGAACTTTCGAACGACTGTGACGGGCTTCCATATACCCTTGAACTGATT

CCCCACGCTAACCCCCCAAGCACCCTGTCCTCAAAGCACGTGAGTGCGTTATTTCTAGTTCACTCTTAAA

GACTACTGCAAGAAACGATCCGGGTCTTAACTTTTCATCAGGTATATATATAGATT

>NODE_1335_length_196_cov_2.211429

ATTCCAAATTTTTTTTTTTTAGAAAGAAAATACGTATTGAGAAGGAACAGGCCTCGCAGGAGAGAAATCA

CAAATAACAATAAAAAAAAAGAGTAACGAAACAAAAAAACAACCAGAAAAATGTACTATAAGAATGGATA

TTAAAATAAAAAAAGAGAGCTTGCGGACGGACAAAAAGACGGCGCAGGGAATCAAA

>NODE_1336_length_196_cov_2.000000

GGGCTTTTGCGCGTTCGATAGAGCCACGAGGAATAGCAACTTGCTGCCCATTACCACAACTTTGTTCATT

GTTATCATGAGGAAGACATACAGCTAACATTGCCCCGCCAGCGTCTGTTCCATAAAATTTAATTCTATCC

ATTTCCATAACTCTTTTACCGCCAACTAGCTCAAGTAAAGCATCGTAAAAAGAAAC

>NODE_1337_length_196_cov_1.422857

GGGCTGCTGGCGAACAGCTCGTCGGACTCCTTCGGCAGCTCGTCCTTGTTCTCCAGCTCGTAATGCCACC

CCTCCTGCTGCATCACCTGAGGGTCACTCACTGCCCTGGGGAAGAAAACGGCGTCAAGGTAGACGTTCAT

CAGGTTGTAGAAGTCCTTAAGGTTCTGGGAAGCGACGGGGTAGCAGGTTCGGTCGG

>NODE_1338_length_196_cov_1.354286

ACGTATTGTGGTTCTATTGGTGCTGAGTACATGCATATCACTTCAACAGATGAAAAACGTTGGTTACAAA

ATCGTTTGGAATCTGTTCAATCTAAGCCTGTATTAACGGTTGCCGAAAAAGAAGAAATTCTTAAAGGTTT

GATTGCTGCTGATGGTCTAGAGAAATACTTAGGTGCGAAATTCCCCGGTGCAAAAC

>NODE_1339_length_195_cov_517.074713

TATAACGAGCGCAACCCCTGTTGTTAGTTGCCAGCGAGTCATGTCGGGGACTCTAGAGAAACTGCCTGCG

TAAGCAGAGAGGAAGGTGGGGACGACGTCAAGTCATCATGCCCCTTACGGGATGGGCTACACACGTGCTA

CAATGGCAGTGACAATGGGATAATCCCCAAAAACTGTCTCAGTTCGGATTGGGGT

>NODE_1340_length_195_cov_20.362069

TGCATGAGCATCGAACGCCTGAAGTAGTCAACATTATCCAACCGCGTATCCAGCTGACCGCCGTTAACTC

CAAGGGCGGGGGATGGGGGGAACACTACGCGCACCCGCGCTCTGCGATGACGTCTCCCCCTCTGTGCGAC

CGACACCCCATCACGAAGTTCAAAACAAAATATCCACGCTTCCGTACATATCGCC

>NODE_1341_length_195_cov_17.919540

AAATTTTTTTTTTTTTTCACGCGTGGATCATTTTTGTATTAGTATACGTTATTGTACATGTTTCTAGGTG

CGCTGAGAGCCCCCTGCGGTTCCATTCCAATCCTACAAAGCAAGCAGTCATTCGTCCTGTGCCTCGAGAA

GCGCACAGCATGACGTTTGGAAACTGGGCTACGCAGGCAGAGCTATAGCTGGCTT

>NODE_1342_length_195_cov_8.793103

TCGTTCCAAGCTGGCGTGTTTCGTGATTTTGGGGAGAAGGTGGTGATTGTGAGGATGAATGTGCATGGTC

GGTAACATGGTTGATAATATTGTGCGTGTGAGTGCGTCGTGGGTTTTTTTTTGTGTCGTGTTGAGAGAGT

TGGAATGAGGCGTGTTTCACTAGTACTACCAGGTATACACTAAGTTCGGGCTCCC

>NODE_1343_length_195_cov_6.022989

GATCGCCTCCCCCCAAAAGTGTTTTTTCGTTCTGTCCGTCCATGAGACTCCTCTTTTACGAGGGTTAGAA

ATTGGTATGGCACATGGTTATTTATTACTAGGACCTTTTGAAAAATTAGGACCATTAAGAGCATCTGAAA

TTTCACTTCTAATTGGGTTTCTATCGTCTGTAGCTTTAGTAACAATACTATGGAA

>NODE_1344_length_195_cov_4.367816

GGCGGTGGCGGCATCGTGCGCGGGGCGGAGGGGCTACCACGACAACATCGTAGAGCACTACGAGAACCCT

CGCAACGTTGGCTCCTTGGACAAGACGGACAAGTTCGTCGGAACGGGCCTCGTGGGTGCGCCGGCCTGTG

GCGATGTCATGAAGCTGCAGATCAGGGTGGATGAAAACGGGGTGATCACGGAGTC

>NODE_1345_length_195_cov_3.988506

CAGCTTGGTGAAGCGCTCCGCGCTGTCGTTGTGAGAGATCTCAGCTCTGAGGCGACGAAGCCGCTCCATC

ACCTTGCCGAAGTTGACGGTGACATATCCCGTGATTTCCACTCCGTAAGATTCGCTGTTCTTGACGGTGT

GGGCCACTCGTGCGGCGGAGATGAGGGCCTTGGACGGTACGCAGCCCACGTTGAG

>NODE_1346_length_195_cov_3.747126

GTCTCCGTTGGACCGCACGTTTGTTTGTCCACGTGTCCCCGCTGTGGTTGTTCGTTGGTTGCAGGTACAA

TATGCTGTGCGTTGAAGTTTCTCAGATTTGGCTGGGCGGTGTTGAGGGACCGACACAGCCAGCCGTGTTG

TCGATCGATGCTCTGTTTTTTCTCCTCCCTGCCTTTATAGGAGCGGGAAGGTCCG

>NODE_1347_length_195_cov_2.988506

CACCACGGACGACAGCGAAAAGGAAGGCTCCGAGCACAACTTGGTCCGGGCCATCCAGGCGGCAAGCACG

GCAAGATCAAGTACACGGAGGTCGGTCGGACGGAGACCATCGGAAACACTTCGGACCCCGAGTTCACACG

GCAATTCCTTTTTGACTATTCCTTCAACGAGACTCAGCACATCAAGGTGAAGATC

>NODE_1348_length_195_cov_2.890805

AATTCGACGAGCTTCCAGCTAAAACGATGAGCGTAGTGTAGGTAAATCCCTGCATCAGGCTGCTTCTGTT

CTTGATCGCGCGTGTTCTCCTCGTCCATCATGGGAGATAACCACTGTGTGGTCGGAATATGCTTCAGGTA

CTGTTTTACTTCTTTGTTTCTGGTGTTCCACAGGCAGCCCGTGAGCTATCCGTCT

>NODE_1349_length_195_cov_2.517241

GTGTCGATTTACGCCCGCTTTTTCCCGTGTGCCTCGAAGAGCTTCAGAGTGTTCTGAAGGAGGAGTGCTA

CTGTCATCGGGCCGACGCCCCCCGGCACAGGGGTGATGTAAGAGGCCTTAGAGCTGGCCTTGGCGAAGTC

CACGTCCCCGACGACCTTCCTCTTGCCGGGGGGTGTGACGCCTTGAGTGTTCATT

>NODE_1350_length_195_cov_2.413793

GCAGCGCGCTCGGATGCCCGATCACGACGAAGGACGGTGACTAGACTGGTGCCTTCGACGAACGTCACCA

CATCTTCGGCCGTAGGCGTACGCCAGAAGAGGGCACTCCAGAGCCCCAGCAAGACCGCAAAGGGGAACTT

GATGAAGTACATTGGATTTTGTAGGCAGTAGTGGGCCAGATATCGCATCACGGGC

>NODE_1351_length_195_cov_2.408046

CAGGCCTGATGGGTTGGGCGGTCGCGTGTGGGCCACGATTCCTACCTCCACGGTATATATACAATGATTT

TTCAACAGTGGAGCAGCTGGTATTAACAGAAACGGTGTAGGATACCCACCAGGACGGAGTATTTCTGTCG

CAGAAAGGCCATCGACTGATCAATGTTGACAACAATTCCAATCTAAAAAAAAAAG

>NODE_1352_length_195_cov_2.091954

GTAGACGACACTCGAAACAAATTGGCTATACAAATCCGACTGATAGATCTCACTCGGAATAATTTGACTG

GGCAATTCCAATTAATAGAAATCACTCGAAATCATTTAGTTGGACAATTGCATTTAATAGATAGCATTTG

AAATCAAGTGTCTCGGCAATTCCAATTAACAGCCAGCACCCGAAATCAATTGTAG

>NODE_1353_length_195_cov_2.051724

CCGTGAGATAAAGCCCGACGGTTTGCTCTTTGACGAATAAGGTCTCACCGTTATGGCATTGTTTATATGT

AGAGAGGTGTTGGCACACTAAATCGAGTTTGCCGAAGTTCTTCAATCACATTGTAGCGTTCTTGGTGATA

AGAGTTGGTGTTCCTCAAACACAGGCGAGTGGACGAGGTGGGTGGCGTCGATACC

>NODE_1354_length_195_cov_1.626437

CGTTTGCCTGCAGACTAGGAAGTATAATTTATCAGCTAGACTGTCGATACCCACCGGGCCCATACAACCG

CAACCGTGCGCTGCTTTGCGTTTGCGCCCTGTAAACGTTTGACAGTTCCTTGACGTGTGTGTTTGTTGTT

TGTGAGCGTGTGATACAGCGTATACACGAAACCCGAGAGGGGGTTCTTTGGTGCC

>NODE_1355_length_195_cov_1.517241

TAGGATGGGATGGCCTCGTACCCTTGGTCTGCGATTGGTCTACAGATGGTATCGGTGGTCTTGTGATTGG

TGGGCTGGTGAGGGTAAAACAAGGGATTTGTCCCGGTCTGTGATTGGTGGACTGGCGAAAAATGTCCGTG

GTCCGGTGATTGGTGGACTGGTCAATCGACATGTTCAGCTGGGATGTCGTCTTGT

>NODE_1356_length_195_cov_1.390805

CAAGGAAGCTGTGCATCTTGGTTGGCAACTGGCACGAGTGGTAATGGTGGCTGAAGATGGCGAGAGCAAG

GAACTGCCTCACACCATCAAGTTGCTGGACCTGGGGAAGCGCTACAACGTGCATCTTTCCGAGGAAAAGC

TGACAACTGTTAGCGAGGATGTTGGAACTTGGTGTTGGCATGTACATGTGACCAC

>NODE_1357_length_195_cov_1.201149

CAAGCTGTGTGCCACGGGTATCGAGAAGGCTGTGGCCGGTACGCCCATCATGGTCGTGGGACCTGAGGAT

GACGAGGAGGACATTAAGGAGGAGGTGATGCGCGACTTCACGGACCTGATGAAGATGGACCTGGAGCCGA

AGGGCGTCATGGTGCAGGCTAGCACCCTGGGCGCGTTGGAGGCGCTGATGCAGTT

>NODE_1358_length_195_cov_1.183908

CCCATCAACACGCTTCTGCCAGTCCGCGGCGCATGGCGTGGGTGGGACGGACGTTGGTAAAGCCTGACAC

CCGCGTACAACTGTGTTTAACGCAAATATATGCAGCTTTGTTCTGGCGGAAGGGATCTCGCTTGGAGAGA

CGCTGCAGCATATAGCACGGGCGGTAGAACACACCCACCTACCTGCGGGTGTACC

>NODE_1359_length_194_cov_51.676301

GGAACTCCCAAATACCGTTATCCTTCTCGTGCCAGGTTTGGCGCGGGATGGAGCAGTCTGGTAGCTCGTC

GGGCTCATAACCCGAAGGTCGTTGGTTCAAATCCAACCCCCGCAACCAGCTGTGGTAGCTCAGTGGTAGA

GCGCGTCATTGGTAATGACGAGGTCGGGAGTTCAATTCTCCCTCGCGGCACCAA

>NODE_1360_length_194_cov_36.763006

ATTCCAAAATGTAGCTGACCGTTTATGAACCCCGGCAATGGCGCTTGTACAGCGCATGCGAACGGCGAAC

ATCTTGAAGGTGAGCCAACTATCGAATCGATCCCGGAGGACCGAATGTCAAATGTTATTCTTGGCTGCGT

AACGTCCCTGGGGGACCGATGGGTTTCGGACTCTCCCACTCTTTCGCGCCTATA

>NODE_1361_length_194_cov_17.433526

TAGCAAGTCCAGTCGGCGACAAGCAAAGCATGTTTTAATTTCACGCACATTGCGATGCCGCCGTCTGCCT

TGTTTCAATACGAGGGTAGCACTTGCGAGAAGAAAAACCGTCGATAGTTCTGGTACCCCGCCTTTCTTTT

CCGTGCGGTGTTGTGCCTTTGATACGACGTAACATTCTGGAATTCCGGGTGCCA

>NODE_1362_length_194_cov_7.797688

TAGACCGCCTCGATAAGCAGCCCACATTCGCCCCCCTCCGCCTCCTCGGAGTCCGCCCTTTCCAGCAGGA

TGTAACCGTCGAGGCCCCAGGTCTCGCCCCACGAGTTCTTCACCTTCCAATAGTCCGTCCCATCCTCCGA

CGTGCCGTAGCCGACCACGAGAACTCCGTGGTCCAGCTTCTCCCCGCACGGTGC

>NODE_1363_length_194_cov_7.647399

CCCCCTCCCCTGCCTGCGTTCGTTCATGTTTGATCAAAATTGTGGGTCTCTATTTTCGCCCCCGTCTTCG

GTTTGACTTGGAATGAAGAGCATCCTCCTTCGGTGTATGAAGGTCCTCCAATCAGCACCACGCCTCCCTA

ACAACTTCATCCTCTCGTCACTATCAATCGGACCGGGTGCTTTCGCCCCGCCCC

>NODE_1364_length_194_cov_7.612717

GATAGAGTGCTGCGGGAAGTGATATGTCGAAGTACATCTGGAGGTGTAGTCGTCGTCCAGGCTCCACGGA

GGAGCACCCCCTGGACCATTCTACGTATGGGAAAAGGGCGGAGGGATTTCGAATTGAAGTGTTTCAGACT

GATGGAGCTATGATAATACAAATTACACCGGGGAACTGCCCAATATACTTGGAA

>NODE_1365_length_194_cov_7.098266

TCGAGGTGATGAGAAAGCCTTCTTTCCCCAGCCGGCCATGTACCCCGGTGTAATACCCCCCAGCAACACA

GATTCAGCATATCTACGTCCTTGGGCAGCTATTATGTGGTTGACACCAATTCCACGCTCTCCCTGTTCTT

TCCCTGAGCATGCTGCTACTGCGTTGTTGCCCCCGTTGGGCGCTGTATGAATGT

>NODE_1366_length_194_cov_3.722543

GTTTACAGGACGACACCCTGCGTTGCTTTGCATTAATACGCTCAAGACGGCGGAAGATCGGCACAGGTAG

ACACTATTTATCGCAACTTATCGAGGAACGCGATAACGTAAGTCACCACGGGATCAAGTCATTCCCAGAA

CCTTGGCAGAGCCAGTACTTCAGGGTACACCACCCCAGTATCTCAGAGTACGCG

>NODE_1367_length_194_cov_3.277457

CCCTCGGTGGCAGTCACGGTGGGCGCCATGGTGCTCTCGAAAGCACCCTCTACTGTTGGCGAAGGGGTCG

GGGCAGGGGTGCCCTCCTCCTGCATCTTCCGCTTCATGCCGTCCTTGGCGTGGAGGAGGTCGACGAGGTC

GGCGTCCAGGTTGTCCAAGTTGTTGAAATCAGCGTGGACGGGGGCGACCAGGTT

>NODE_1368_length_194_cov_3.132948

GTAATACACGGTCCATGCGATCCCCTCGATACGAAGCGTGTACACATCAAAATCCCAATCCCACTACAAT

ATGTACCAAACGTATCTACACCGTCCGAATCTGACAAATCACTACAAATGGACGAGAATATTCTACGGTG

TATAGTTGATGCATTTGCAAGCAACCTCCCCGCCCACCGTTACCGTTTTCTCGG

>NODE_1369_length_194_cov_2.838150

GTAAAAGCCCCGCAATCAGCGGGAAAAAAAGAAGAAGAAAGTATTGAAAATGTATTGGTGGGTAGATAGA

TGCCTGGTATTAATAGTTGGCTCAGTTCACGAGGAAGCTCCGTCAGCCAGGATTGGTTCGGCCCACAGCT

ACCCCCCTCTGCCATATCACGACTCGGGAGGAAAGGGGGGTCGGCGGCGGGGGT

>NODE_1370_length_194_cov_2.745665

ATGATTTCCGCTCTCACGGAATCGCACATGTGCGCGGTGCTTGTGTCCGAGGAGAACGAGGAGCCAATCA

TCATCGAGAACAAGAAGTCAGGCAAGTACTGCGTGGCTTTCGACCCCCTGGACGGGTCCTCGAACATCGA

CTGCAACGTCTCCACCGGCACCATCTTCGCCGTGTACGACAAAATTTCTGCGGT

>NODE_1371_length_194_cov_2.745665

ACAAAATATCACGCGTGAGACATGTCCGACATTCATGATATTTTTGCTGTTCTCGTGCCACAAAATAAAT

TCCAAGTTTCTAGACGCTGTCTACCAACACGCCTCAACGACCTCGGTAATGCAGTTGTACACTTCTACAT

CGTACCTTACCTCTGACTTTCTCCTGCCACAGCGTGAGACTTCCTCCGGTCAAC

>NODE_1372_length_194_cov_2.450867

TATACTCCTTAAGAGCGAGACCTATCATCCCCTTGGCTGCTGTATGATGGTGATGAACGATTTGCCACCT

GGAACCATCTCCTATGACGGCGAGCATATCCGAGCTCTCACACCCTGGTACCCATTTCCTGCGTGGTTCA

TCGTAGTGTTTGGGTCGACAGAGAACGACTGCTGCCTCATAGCCTTGCGACGCA

>NODE_1373_length_194_cov_2.092486

CAGGACCTTTTTGGTCTCAGTCTCTTCCACCAGGAAGCCCAGGGCCGCGTGCGTCAAGGCGTACGACCCC

CAGAAGCCTCCAGCGGCATTCAAAATATCGTCCCCGGCGTTCGACGGAAGTGCTGCCTCCAGTAGAAATA

CTTTCACAATCGCCCCGTTATCCACAACATCGATCGAGTTGGAGTCCATCCTCT

>NODE_1374_length_194_cov_1.635838

AAAATTTCTTCGCAAGCCAAGCCCACATCCTCTTCCCCCTCCGAGAAAATTTCTCGAACAGCCTCGGTGC

AGTCCTCGTTGGACATGACATCCCAGATTCCATCGCACGCCACAATTAGGAACTCGTCCGACTTTGGCGA

ACGCGGGATGCTCTGGGTCTCGGGGGCACAGGTAACTTTCCACTCTTCTGCCGG

>NODE_1375_length_194_cov_1.635838

TTGAAACTGATGAGAATGAAGAGTTTATTGCCAAATGGAAAAAATATATTGGTGATGACAAGCGTGTCAC

TAATGATCCTATGGAAGCCACTTATATTGGTTTCAAAATGTGGGCAAAAGCAGTAGAGAAAGTAGGTACT

ACTGACGTAGATGCTGTTGAGCAAGGTTTAAAAGGCTTAGAGATCAAAGAAATC

>NODE_1376_length_194_cov_1.526012

GTCCTCTTTGTAGCAGCCCTCGTGGAAGTCTCCGCATCGGAACTGTTTCTTGCCCCACCCATCGGCCATA

CCTCCTTTCCACGCTCCTTGTAAACGTCCCCGTTGCCCATCTTCTTCACTCCGTGACCGTGCATCTTTCC

CGCCCTCCAGTCGCCCTCGTACATGTCTCCCTTGCTCCACATGAAGGTCCCCTT

>NODE_1377_length_194_cov_1.352601

GAGATTTGGTCCAGTCGGAGGAAGCATTCTTCGAGGAGCTGTATGGCAACGACGCCAATGACAGCGCTGT

GGCGGGAGTCATTGACAGGGGTATGCTCTACGTGGCGATTAACACAGGAAACGTAGATATTGGGGGCGAC

CCGTTCGGTGGGACGGACCTGGTTGTTAAGATCTTCGACGAAGAAGATCTGGAG

>NODE_1378_length_194_cov_1.190751

TGACATCCATTGGTCCGCCATGTCGGGCCATGTCAGCACGTTCATCGTGAACGGCGGGAAATACGACAAA

ATCATCTGGACCGAGAGCTTTAACGACGGGATGCGCGATGTGCTTGATGCGAATCATCGGCGTGTCCCCC

ACCTTCTCCAACGCATTCTCCAGAATCTTGCCCCTTTTAGCCCTCCCCGCCATG

>NODE_1379_length_193_cov_261.098837

AGACCCCGTGCACCTTTACTACAACTTCACACTGGCATTAGGCCGAACATGTGCAGGATAGGTGGTGGGC

TTTGAAGCAGGGACGCCAGTCCCTGTGGAGCCTCCCTTGAGATACCACCCTTGTTCTGCTTGATGTCTAA

CCGCGGTCCGTTATCCGGATCCGGGACCCTGTGTGGTGGGTAGTTTGACTGGG

>NODE_1380_length_193_cov_16.273256

TGCCACCCTACGCGCATCTGCGGTGCAGATACGCTTCCTGAAGAGATATGAGGACGGCTCGGTCCAAAGA

CCATCTTATGATGATCCAAATATGAACAGCTTTGTTTGCTGTTCTGCTAAGTGTTCCACGAGTAAAAGAA

CAAGTTCTTATGCTGCTAGGAACATCCTTAGAAAGGAGGTGATCCAGCCGCAC

>NODE_1381_length_193_cov_7.389535

ACACTTTGCGCGAACACGATAATACCGGCCTCACTCTCCGTCGCCGGCCTTGCCTCACTACCTGAAAATG

CCCGAATCGCGAATTGGAGTACTAACATCATGACACAAATGAAATGGGGGAGCCAAATCCCATACATACC

AAACAACACACAGGCATCCGCTGGTGGAGCCGCCTGCCTTTGAGGCGAGATAT

>NODE_1382_length_193_cov_5.505814

ATGTCGCGGTTGCTGTCGAGGAGTAGCCTCTGGATGTTGGTTTTCCCGAACTCCTCGAAGCATGCCGGCA

AAAATGGCACATCGTCGTCCGTTAGGTCGCAGTCCGTGAACTCTATAGTCGTGTCTCCCGAAAAGGTGCA

CTTGAAGTCCGCACTCGCGGGGTCGAGACACGAGTTGCTGCTGCCGCCACCAC

>NODE_1383_length_193_cov_4.633721

GGAACTTGACCTCTCCGATGCCCCTCTCGACCCACTCCTTCTGTCCCGTACCCTTGTTCAGCAAAGTTTC

TCCGAAAATGAACAGCTTGGCCCGCATCTTCCACAGGACCTCCTCGTCTTCCTCGTGCGTCTTGACTTCT

ACCTCGTCCAGCTGGACGACAGGCTCAAACTGGGCCGTGCTCTCTTCCTCGTG

>NODE_1384_length_193_cov_3.941860

TATATAAAAACATCATTCCTGGAATTACAAATTCCACAGCATGATCCCATCTCCCATCTGTTATAAGATG

AGGTAAACCTTCTTTACCACATAATAACTCAGAGTTTGAGTAACGAGTAAATCTTTGTTTAGTCCTAGTA

ATTTGTTGTTCTAAAGCTAATGCTGGTGGAGACCCAGGCTCATATTTTTTAAT

>NODE_1385_length_193_cov_3.441860

AGGAAGCGAAGGACCCTTCCTCGGGCGGCCCCCTGAGCAACGGGCTCGCACTTCAAAGCCTCGAACACGA

ACTCGGGGCCGAGGTAAGAACCGCCGATGCCGATGGAGACGATGTTGACGAGCGGCTTCCCGGTGCAGCC

CTTGAAGGCGCCGGAGCGGACCCGGGTGGAGAAGTCGTCGATCTTGTCGAGGA

>NODE_1386_length_193_cov_3.180233

AGTTCATTGGTTTGGCGAGTTTGGTGCTGGGAGTGCATCGAAGGGAGCGCGAGTGCACGACGCGCGCGCG

GGAGAAAGAGAAAGATAACGAGAACGAGAGCGAATGAAAAATAAGAATATGAGATGGAAGAGATAGCGGA

GGAGAAATACCGAGTCTGTGGGAGCGTGTGCGTGGTTGGTTATAGCCTAGAAC

>NODE_1387_length_193_cov_2.953488

CAGGTCGCTTCCACAAAGAGCGACTATGGTACTGCGGCTGTACTGTGAAAACATATGCATTATTGACCGA

AAAACAAACACGCCTCGGGTATATTCCAACTCGTTCTAAACCGAGGCTGGTGTTGCAAGAAGCAGGGGAC

AGGTTCAGCCCCACCCCGGGTGGGTGCCGCCTCCTGCGAAGCCATATAGCGTG

>NODE_1388_length_193_cov_2.709302

TCGCGATGTCCGCTAAGCTGCTTGGCGCCAACTTCTTCACCTCTCCAACCTTTTTCTTGCTGTCCGTGAG

CACGTTGAACTGCACCCCGCAGCAGTCCCCGATGATGCACGGCCCCTTCACTACCAACTCCCTGTCCCCC

TTCCTGCTCAACACGTTGAGCTTCGGGGTGCAGCACCCCCCGCAGCACGACTG

>NODE_1389_length_193_cov_2.424419

CCCGCTCCAAACCAAGCATCGACTTGTTATTTCCCGTTGCGATTCGCCCTCGGGCCGTGCACTCCAAGTT

GTTGCTGACGTCTGACCACTACCCGTGAAAAAAGACGGCATCACAGCCATTTACTTTCACGTACATCCGG

CGTGTTTCCCCCAGCATATGCTTGAACCCGATTGCAACTCAACCAACCCGTTG

>NODE_1390_length_193_cov_2.412791

GAAGTGCCGGGCGATGCATGGACCGACTTCCTGCCGGCATCTGCTGTGCTGACGTCGGATTGGAAAGAAC

TGGGCTGCGGAATGTACAACACCGAAACCAAGACCTGCGTTCAGCTGACGTCGAACTACTAAGATATTTG

TAAAAATATCGGCGCGAGGCAGGCATTCGCTTTGCCCCGCGCAACCTTAATTA

>NODE_1391_length_193_cov_2.313953

AGTCTCTCCTCGGGATGGAGTTCATCGGCCACGAGGACTCGTTCGTGAGCATGTGCCACTCTGTCATCCA

GCACGGGATCGTGGAGAAGACGCCGAAGTACGCCGAATGGGAAAAGGAGAGCGCGGACGGTGGAAAGTAG

AGACAACGCGGTATACGCTCCCCGGCTTGAGTTGAGAGCCGAAGCACACGCCC

>NODE_1392_length_193_cov_2.197674

CTACCACAGTCCGTCCACCACAACACACCGAGACAGCGTTTCCACTGGGAAATACACTGTGTGGGGTGGC

GGTCGGAATGTGTTGGGTGGTTCATAGCCAGCCATAGCATGGCTTGACTTGTGGAGACAACAAACAAAAA

GTTGCATGCAGAGCGCCGCGGGTATCATCACTCTCTAACGAAACCTACTGAGC

>NODE_1393_length_193_cov_2.029070

GAAGAAGAATTACTAGAATTACCAGTCCTTGGTAAGGTTAAGGTATGCGGGTGAGTAAATTGACCTCCGC

TCGTTGCTACTGTATGTTACTTTTTATCTATAGCAAAACCGTGCGCAGTCTGCGCTGCCGTACAGAATAC

ACGGTGGCCTTGCCTCACTCTCGGCTGTCGCCTTCCTACACCATCTACTTTTG

>NODE_1394_length_193_cov_1.633721

GAAGGCATACGCCCCCGTAAGGAGCTCCCACGCGCAGCCCGTATTTGCGGCTCAGGTAGCACTCCGCGGT

CGAAGACACCAGGTACAAAACCTTCTGTGCGTCGATGGTGGGGTCCAGCTCGTGCGCCAGGCCGTAGATG

CAGTTTGCGAAAGAGATCTTGAGCCCGTTAGCATAGTTATGCACGAGCTTCAT

>NODE_1395_length_193_cov_1.616279

GTCCGTTACATCTGGGCAACTTTCCACGCATATCCCTGAACCCTCAGTGTTGACGTAGTACAAGTTTGGC

AGGTCCTCAAAGTCCGTATCCCAACCGCAAAGGTTTCCTTCGTAGTCTGTGGGCGACAACAGCTTCTGCA

CATCGCCCTCTCGAACGCTGGCAACGCCCACGGCAGTCATGGCTATCCACATG

>NODE_1396_length_193_cov_1.534884

ATGAGGTACTTGGATTTAGAGTGGTCTTTTTTGCAGTATGTAGCGGAAGGGTAAATGAAACGAACACGAA

GCACAAATTGGCCAATCGGCGGCGAGAGGCTAGCTGTTTTCATGCCTGTTGATGTTTTTTTTTTGGTCTT

CAACAACGTGCATTTTTCTCATTTAAGCAAAGCTCACTGTATCCATAGTCCGG

>NODE_1397_length_193_cov_1.430233

GCAAACTGCGCTAAACTGCGGGGCATGTGGGCGATCGAGATGGCGAAGGACGAGCCGAACCTGGACGTGA

AGTACCAGTACGCCCTGACGCTCATCAAGACCCGATCCAATGACAAGAACCGGGAGGGTCTGTCTCTCTT

GCAAGGTCTTGCCGACTGCAGCTACCACGTGGAGGAGTGTTACTACGGCATGG

>NODE_1398_length_192_cov_86.210526

ACAGCGGCCGCCTGACTGTCGAGAGGATGACCGGGGGTCGAGTAATATACCTGTGTCTGCTTCTGTCGTA

TTTCGTGGGCGCTCGTGCACACTGCGTGATTACAACAATACAGAGCGCGACGTGTTTCGGCCGGTACCCC

TCGGGGCGCCGACCCAGACCCCGTGGGCTCATGAGATTTCTGCCTGACAGGA

>NODE_1399_length_192_cov_26.883041

GATTGTCGGGCGTGGTTTACACCCCTGAGACACGAGGGGAGCGGGACGCAATGAAGCACCCCAAACCCCT

CCTCAGGCATGCTTGTTAGAGCGCCCACCGCTTCGCCGAGGGAGCAAATAAATGCACCCCCGGGTCTACA

AATTTCTATTGGCCGCCTTGTGCGCCGCAAACGCCATTTCCCCGGGGCTGAG

>NODE_1400_length_192_cov_20.181287

AGTGGGTAATCATTAGTGCAAGGTAAACTCCACCCGAAGCAAAACCAAATAGGCTTCCTGAGGCATGGCC

CATGCTGGAAGCGGGTAGGTTGCTTGAGCCTTAGAGCGATTTAAGGCCTAGACGAATGATTGTCACTTCC

TTTGGAAGATACAAAACCCGGCTTATGTGTTAGCTCATAAATTTTAGTAACC

>NODE_1401_length_192_cov_14.824561

TGGCTCCCGAGATTTCCAAGCCGAGGCCGACGTTCTTCACGAAACGCATGCCCTTGTTCTTGGCGACCTT

CTTGCCGAGCACACGCTTCGCTCCCACGAAGACGGCATCCTGCTTCTGGTACGCCTTCTCGTGCTGCTCA

GCCTCCCGGTAGTCCTGGAGTCCCATGGTGTCGGAGGTGCAAGGAAGCTGAT

>NODE_1402_length_192_cov_10.538012

TTGACAGACAACCCGATAATGGTAGGGGACATGAACGTGTCCCAGGAGGAATTGGAGCAGAGGCACAAGC

CGCAAGAGATTGCTGATGCGCGGTTTGTGTATGGTTGGTGCTTCAAGAGTGTAAGAGTTCATTATGGTTG

AGATGTGTTCATCCTGCGTTGTGGTGCCTTATCATGCACAGAGCAGGGGGGA

>NODE_1403_length_192_cov_9.081871

GTGTATGTATAGTCCCATAGTCCCGAGATATTTTTCGTCTCGTTGAACGGTCAAACGTCACGAATTGAAG

TACATAAAAAACTACAAAGCTGCAAAGCTGTGTCAGGAACCAGCCCTTTTGTCGGGCCCCGGCTTCGGCC

TCAAGTTTGCACACGGAACCATGATTCACTTGAACAACAAAAAAAAAAAATT

>NODE_1404_length_192_cov_7.099415

GGGCAGGGGCGGCTGTACTGATACAGCTGTCGACGAATGCAGAAACGAAATGGTAGATCGTGCGGCAAGG

TATCCGCTCGACTAGTCCTTGAATTCTATTTTTGTTTCCTGTTGATTTGTTCACTGGGGGGGGCGCGGCA

GAGGTAGTACCGGCTTTTGACACTGTTTTTGTTTTTAGATTGGGAATGAGAT

>NODE_1405_length_192_cov_5.339181

TATGTTTTAATGATGGCAACAGCTTTTATTGGTTATGTTTTGCCTTGGGGTCAAATGAGTTTTTGGGGTG

CTACTGTTATTACAAGTTTATTTTCTATTTTCCCTTTTATAGGTAAAGAAGTTGTTATGTGGTTATGGGG

AGGGTTTACAGTTGGTAACCCAACTTTAAATCGTTTTTTTATTTTTTCAAAT

>NODE_1406_length_192_cov_4.502924

CGCCTAAGCCCAGCTAGCTGATGTGGTCGATTGATTGGCTGATTGGTTGATTGACCTGTTGGTTGGTGGA

TCGATCGGTCGTTGATTCAATCGATTGATCGGTCGTTGCGGAAGTCGTCGTCGTGCACAGCGGGTTTTTG

GTTTTTTGTCGAGACCTTATGTTTCCCCCTTCTTTTTCCTGACTTTTGCGGG

>NODE_1407_length_192_cov_3.918129

CCACACGCTGTCGAACATGGCATTGAGCTCATGGATGGCCTTCTTCCCGTGTTCACCTCTAGCTTGATGA

ATTAGAGTCACCGTCATCACGAACAAGATGACGAAACGGACCAAGTATCCCCAACCCACAGAGCTTTTCT

CTGCTGGAGCCCCAACAATATGATCTTTGAACGGCTGCTCCATGAAGGACTT

>NODE_1408_length_192_cov_3.380117

GTATTGATCTGCTTCTTGCCCCGACAGCCTTTGTTTTAGGCGGGCTTGGCCCGGAGAGGAGACCAGGCGG

AAAGGGTGCGGTGGCAAAAGCGAAGGAGTGCTTGTCCTTGAACGATAATCGGACCACAAAACAATAAAAC

CCGGAGGAGGAAGGAGCGTGCACGTGCACGGATCACGGCGGCGTGGCTGGAA

>NODE_1409_length_192_cov_3.374269

TCGGAGCTAAGTTCTCTGGACGTGTCGGTTTCCCTTCTCGTCAAGTACGAACAGGCTCGACACTGGGAAG

ACTGGGAGGTTGGAGTGCACGGAATTGTGATCAAATTTACGGATGATCGCGGGGCGGGCTTCAGCGCGAC

CTTCCTCCCAGAAGTGGCAGCAGAGCAAGGCTGGACGAAGAAGGTGACCCTT

>NODE_1410_length_192_cov_3.058480

GCCCTCACGCGCCTGCCATCGTGAAGGCGGATCTGCAGCGTTGTGGAGGGTTGGGAGGCGTCCACTTCCG

GGACTTCCTCGGCCTCGGGGGCGCTTGTCGCCGACATCACAGCGCCTTCTGCTACGGTTGACCCCATGGT

CTGGCCTTCTCCGCCATATGCGACGTACGCGGGCGCAACATAGTCCTCGTTC

>NODE_1411_length_192_cov_3.011696

TTTGCGTGGAGGGGTGGACGAACCTTGATAGTAGATCTTTACGGGGGTTTTGTGCTTGGCGCCTGAAGCC

GACCCTCCGACTCGAACCCTAACCCTAACCCGAACCCTATGCCAGTGGTCAGTGCTTGCTTGGCCAATCA

GAGTGCCTAATTAGCTACCTTAGGAAGATCCACTCACAGTGTTTGTCCACTA

>NODE_1412_length_192_cov_2.918129

TTTCGAAAACGTCTTCGAAGTCGTCGACGGAGTCCCGACACTTACTTTTGCGCCCTTCATGAGGGAAGTC

GTGAATGCGTGGCAGGAGGGAGTCGACAGGCTGCTCGAGCGCGGGGTGACCGGAAGGATCCTACTCGCCA

ACCTCGTGTCGACAGACTTCTTCTCGGGTTCCGACTCCAGTGGGCTAGAAGT

>NODE_1413_length_192_cov_2.730994

TCGTGCAAAAAACAATCAAAATAAAAATGTAATTATGTTACACTATGTTAATTACATGATAAATAATTGT

ATTTATTTATGTGATTCATCTTTAAATCAACCATCTTTAAACCAACTTAGTAAGGCAGGACGTAGAGTAA

TGGCTCTCATGAGAAATTATCATATATCAACAAACCTTCCCCCTCCTAGATC

>NODE_1414_length_192_cov_2.695906

ACCTTCTTGTGACGTGCACGAGCTTGTACACCACGTTTTACTCTAGCCATTTTCTATATCTCCTATTAAC

CGTGACGTAAAAGTTTTTTAACTGATTTAATATCAGAAGCAGCGATCATGCATTTAGCACGCAAATGACG

CTTAACTTTAGTACGACGCTTAGTCAAGATATGACGAAGGCCAGCTTGTTTG

>NODE_1415_length_192_cov_1.947368

CCGTCTGTGGATCCTGCCAAAGACTCGCCCGAAACGTCCGATGATACGGAAGAACTAGCTGTTGTCGAGG

CTGTGGCTGGGCCAGTGGAGGCTGAAGAGCCGACCGCTGTAGACGTTGACGATGCTCTGGCGACCAAGGA

TACCGAGGACCGGGTTGTGGAAGACGCGGGTCATGAACAGGAGGTGTTGGCT

>NODE_1416_length_192_cov_1.824561

ACCGTGTACGATGTCAAGCGAATCATCGGGCGTTCGTGGAATGACCAGGTTGTCCGCGACGACGCCAAGT

CTTTCCCCTTCCGAGTGGTGGACGGTGGCAACGGCCGCCCCAAGGTGGAGGTCTCCTGGCTGGGAACGAA

GAAGCAGATGGCGCCCGAGGAGGTCTCCGCCATGGTGCTCGTCGAGCTCATG

>NODE_1417_length_192_cov_1.742690

CAAGGTGCAGACCCATGCCTGACAGCACTTCCTTGATCTCGTTCAAGGATTTGCGACCAAAGTTCGGCGT

GCGCAACATTTCGGCTTCGGTTTTCTGGATCAAATCGCCGATGTAGACAATGTTGTCGTTCTTCAGGCAG

TTCGCGGAACGTACAGACAGTTCCAGCTCGTCCACTTTCTTGAGCAGAAGCG

>NODE_1418_length_192_cov_1.625731

TTTCTTTTTTTTTGCATAGCGGGCTTGAGATTCAGCCCGCTATGGCCGCGCAATAACTAGCGACTTTTTA

CGACTTGATTTCTAGCAGTTCGACTTCGAAGACCAGAGTCTCGCCGGGTCCGATCGCCTGGCCTGCACCG

CGCTCACCATAAGCCAAGTCAGAAGGAATAACGAAGCGGTACTTGCTGCCAA

>NODE_1419_length_192_cov_1.619883

GCTTGCACCCCAACGCCCGCCCCTTCCTTCGCTCCTCCCGACATTATTACAGACAGTATCTACAGTACAC

TGCTTGTTGTGCACTGTGCTGTAATATCTGCTCTTTCTTGACTTCCAAGACTTGTTCTTGTATCTCCGTG

TGACTCGATGGGGTAGTTAAAGTCGGCTTTGAGATCCAGCAGTATGCCCTTG

>NODE_1420_length_192_cov_1.532164

TTTTGTCATTTAAGTAACATTTTTTAATTTGGGTTTTTTAATGAAAACTTTTGTAGCTAAACCAGCAAGC

GTACAACGCGAATGGTTCTTAGTGGACGCCGAAGATAAAACTTTAGGTCGTATCGCTACTGAAATTGCAA

GCCGTTTACGCGGTAAGCATAAACCAGAATATACACCTCATGTTGATACTGG

>NODE_1421_length_192_cov_1.374269

TTTCGTCATGCGCATGAAGACGGAGGTCGCGACACTCTCACAGGCACAACGCCTTCCAGGGCTGCCCTCT

TCCATGCTGGGGCTGGAGACCGTCCTGGGCCGAGACGAGACCGTGGAGTTCGAGGACTACCTCGATGCAC

CCGCCGACAGACCGGAGGCTCCACGCTTCCGAGTCCACGAGGCCATGGAAAT

>NODE_1422_length_191_cov_23.423529

AGATGTTGTCACTGTCTGCGCCGGTGACCTGGTGCGCAGTGCCCCCGAGAGGGTTGGTCAGCTTCGCTCC

ATTCTTTTTCGCCCCAGAAGGCACGTCATCGCACACGTTGATGTCGCTTGTGAAAATACACTCCTGCAGC

TTCTTGTAGGCGTCTTCGTCCACCTGTCCAAACTTGTCGTGCGGGAGCGTC

>NODE_1423_length_191_cov_18.064706

CCGTAATGGGTTTTAAGATGATATAATTTTAAGCATAAGGATGTAATAATTTATGGGTGGAGAAATTTTA

TCAATCGGTGTTTTATGTTTTAGTATGGTGCTAATCGGATTATCTCTTGGGTTTCTATTATTAAAAGTCC

AAGGAGAATAAAAAAGTAAATTAAAATAATCAATAATACTCTGCAAACAAA

>NODE_1424_length_191_cov_3.617647

ATAAATCGGAATTACCAATACCATCCAAATCGTTACCAAGCAAATCACGTTCTCCCATTCCCATCCAAAT

CGTTACCGAGAACATCATGTTATCCCATTCCCATCCAAACAATTACCGTGCAAATATCGGCATTTCTGTG

GTTTCTCGTTACCATCCAACCAAATACCGCGCAAACGTACAGTTACCAATT

>NODE_1425_length_191_cov_3.347059

TGCGCGCCACAAGTGCGGCAGATTTTAAACACTCAAGTGTCGAAAAATGGGCTGGCTACAGAATATACAA

TGCGTTGAAATTTACGCTTCCTCGCAATGTTATCTTTTGGGCGCAACAGTGAGAAGATTGAGTCCACGAC

GACCAATCGTTCGCGCCCCCCGGAGCGATGATAGAGACTCTGCCAAAGCGA

>NODE_1426_length_191_cov_2.976471

TCGCGAGTAAAGAAAGTTACAGTAACGCGCCGTGGAGTTATCAAAATCAAGTTCGAACCCACCACCGACA

CTACATCGTTCCACCCTCTCAGCTAGCCCCAACGAAGGTTTCGATCAGAGGGAGACACAAGAGCTCGGAA

ACAACCAAACAAATCGCGTGCGAGATAATACAAATGAACATATTTTCAAGG

>NODE_1427_length_191_cov_2.741176

TTGACAAGGATGTGGTGGAAGCATTGAATGAAATCAGGGTGGTCGATTTTCAGTAGCTCGGCGGGAAGGT

CGTCTGGTCCTACCGCTTTCCAGTTAGGCATTGTCTTGACGGCTCCCATCATCTCGTCCACAGATGGCTC

GTCGCCAAGTGATTGTTCAAGCGGTCGCGGTGGCAATGGGTCGATGATAGT

>NODE_1428_length_191_cov_2.552941

AAGTTTCATGTTCACAAAAAAAAAACTCGCGAGAACGCGGCGTGTGCCCAGCATTACACCGTTCTCCAGA

GGCAACAAAAAACTGATCGCCCTCTGCCCAGCGATTGATGTATACTGTAATGCAGAGCCAATTCACGGCG

TGCTGTTCGAACATCAACAACACCCGTTGTTCAGCGCTTGACGTATACATA

>NODE_1429_length_191_cov_2.494118

CAAGTAGACGCGAAAGGTTTCGTGATACCTAGGCGAATACCGGTAGGATTAACTTTTTGACCCATACTAA

TATACTCCTAAGAATCAGACACAATCACAGTAATGTGACTAGTGCGCTTGATGATGCGATCGGCTCGGCC

TTTCGCTCGTGGCATAATACGTTTCATTGTTGGACCATCGTCAATCATAAT

>NODE_1430_length_191_cov_2.458824

AAAGGTCCTTTAGGTCGTGAAATGTTCCGCAAATTAAAAGTGTATGCAGGCTCTGAGCATTCGCATACTG

CACAACAACCACAACTTTTGGAGCTTTAAGCAATGGCTGATAATCAATATTACGGTACTGGTCGTCGCAA

GAGCTCAACTGCTCGTGTGTTTATGAAAGCTGGTAACGGTGCAATTACAAT

>NODE_1431_length_191_cov_2.123529

ACGTTTTTTTTTTGTTGGTTTTTATTTTTGTTGTTTTTCCCGCTGTTCTTGTCCTTGCTCTTTCTCTTGT

TCTTGTCTTCTTTCTCCTCTCGATCGCCGCCGATTTTGCTGGCGGTCGAAGAACTTGCCGAAGAAATTTC

CGAAGAACTTTTCACAGAACCCTCCACAGACCCGCCCGCGGATTGTTTCCC

>NODE_1432_length_191_cov_2.123529

CGCGTGAGTGACAAGATCACGTCGTTGCACGACTACATCTACGCCTGTCGCTCTGGATCGGCCGCCGATA

CCCAGGCCGTCGCGGACTACGTGCGCTACTTCTTGGCGTCGCACAGTTACGGCCACGTGGATGCAGAGTA

CAGGGAGAACATGACGAAGGAGGAAGCGCAACAGTTCGTCCGCAAGGCTAT

>NODE_1433_length_191_cov_2.047059

CTCCCTCTTGATCTTCTTCTTCTTCACGGGCTTGAGGATTTTGATCACGCACTTGGTGTTGTTCAGCACG

TTCATGCCCTCGAACACCTCGCTGTACTTGCCCCTCCCGATCTTCCTCACCACCTCATAGTCCTCCTGGT

CCCCCCAGTTCACCACGAGGCTCTCGTAGTCCCAGTAGTCTCGTGGCCGGT

>NODE_1434_length_191_cov_1.976471

CCGATCACGGCAAGCCAGGCGCACGCGATTGGCATGGTGGACGACTTGTTGCAGCACTCTCCGAGTTCTT

CCTTCAACGAGGAGTTGTGCGCCATCCGAGACCAAGCGGAGGTCCTCGTGGCGAGTGGTCTCGCTGACAA

GATCATCGCGGAGAAGAAGCAAGACAGGGACGCTGCCTGGTTTGAGAAGGT

>NODE_1435_length_191_cov_1.770588

CGATGGCATGGTTGGTTTGGAAACATTGGATCTTGAAGAAAACCGGCTTAGCACGCTGCCTGACAACATC

TTCGACGACTTGGTGTCGCTTAAAGAGCTGAATGTTCCAAGCAATGAGCTTATCGCACTGCCTGATAACG

TATTTGACCACCTGGTGTCGCTTCAGAAACTGAACCTTCGGATCAATAACC

>NODE_1436_length_191_cov_1.582353

CGGATAACTTTCATCCAAACACCATTTGCACGAGCTTCTTTACGAAGGGCAGTGATTGCGCCAACTGTTA

CACCACGAGAATCTGCAACAACTGCAGATAGAGCACCATTGGCTGCTTCGTTGACTTCAGCAACAATAGC

TTGTTTGTCTTCGAGTCTTAATGCCATTTGGTTTTACTCCTGGATTCCACT

>NODE_1437_length_191_cov_1.194118

TCTGAATGGGTTAACTACACATTGCTCTTATCTGTAGTGGGCACAATATGTAGGCTTTAGTGTAGGCGGG

ACATGCCGATTTGACCGCTCTTAAATCCCGAATGTTCTTATATTAGGGCGCTCTTAAATCGGGGTACCCA

CTGCAATAGGAGTATCCCTAGTCTGTACCTTTTATATACTGCTGCATTTTT

>NODE_1438_length_190_cov_4.532544

CCACCGCGAATATGGGAGTGCGTAGGGTGGCCAGCTTATTGTCTTGGACCCCGATGTAGATGACGTTGTA

AATGAACCTGGAGATGAGCCAAACCGCTGCGACAGCGCTCATTTTGTCTTCGTCTACTTCCATCACGGCT

GCCAGGAGGACTGCCGCCACCCATACGATCATAGCCTCCCAGCCATTCTG

>NODE_1439_length_190_cov_3.893491

ACAAAGAGGGCGGATTAAGTTCAAGCACGCCTAGCCAGATAGGTAGTCGCGGATTACCGACCGTTTTATG

TTTCATTCGTTCCCGAGAAGAACTGGATCAGTCACCCATTGATCGGCAAGGCATCAAAAACGAGCGCATT

TTTTTTTGTGGCTTTTTTTTGTGGCAGGAAAGGGTCCCCTCTTTTTCCTA

>NODE_1440_length_190_cov_3.189349

GACGAGGCGTCTAAGACTGAATACGAGGTTTGGGGTAGGTATGCATGTGTAGCGGCCATTCTTATGTGCT

GCTGTTTCCAGTAGCTGGATCATCGATAACTATTTCGATTCCCGAGTATTATGGTCGTCGTGTGATCAGC

ATGTTTTCAACTGTCCAGGTTAAGCTGACTGCGGTGTGGAATTACGCGTC

>NODE_1441_length_190_cov_2.514793

GATTATTCTAAAACCCGGAGTATTTTTGTCGGGTTGCGGGCGAAACAAATCCCCGGGATTATTTCCTTCG

GAGATATCAGGAGGAATTTCCTGGGCGTTTTTTTCGCTTGCAGAGCGAAATCGCACGTTAAGTGTGTTTT

TCGTCTGTAAAATGTCGGGTGGTATGGCTAGGCGGGGGGAAGGATGGAAT

>NODE_1442_length_190_cov_2.366864

ATTCTCCGGCTGCCACCTGCGCCCGCAGACCGTCCTCCACAGGCTCCACGGACTCCCGGTTCTCGCTCTC

GAAGTGATCCAGCAGCTTCTTGTTGTCCGCCATTGCCGAGCTGTTGGCCAAGCCGGGGCACCAGTAACCC

TTGGTGAAGATAAACCACGGCTTCTGCGCGGTACCCCACTCCGAGGGCAG

>NODE_1443_length_190_cov_2.112426

CTGGTTTAGGTACGATGGGGTTTGGCTTACCTGCTGCTATTGGTGCAGCTGTGGCTTACCCAAAAGAAGA

TATTATTTGTATTACTGGGGACTCTAGTTTCCAAATGAATTTACAAGAATTAGGAACAATTTCTAAATAT

AAGTTACGAATTAAAATTATTATTATTAATAATCATTGGCAAGGAATGGT

>NODE_1444_length_190_cov_1.822485

TATAATTATCTCTCCTCTATTTTACGTGTAGATGATTGGTGATTTTGATTGATGACGACGATGAAAATAA

TGGCGATAATTGAAGAGCTTGGTAGGGGACGGTAGATCATCTTATGGCGGAACGATTGGTTGGTGGTTTT

AGTCTGTGATTGTTCATCCTTCTCACTCTTGATGGTTTGGATTTTTTTGC

>NODE_1445_length_190_cov_1.621302

TAATCTTCAGTAGGTTCGATATCTGTAGGGGTAATACTTGCTGCATGATGACCTTTAGGTCCTGCATCTA

TTTCATAATTAACGTCTTGTCCCGCTTTTAACGTGCGGTAACCTTCCATTTGAATAGTTGAGTAATGAGC

AAATATATCTTCTCCACCTTCTTCTCGAACTCGGGCATGTTCAAGCACAT

>NODE_1446_length_190_cov_1.224852

ATGCTAAATATGAAGAAAAAAGAACTTCATAAAGACTTAAGTAAAGCTGTTGAATTTGATCAAAGTCAAA

CATTTAAGAAAATCTATGAATCTGAATTTGGTACTCCAGGTGGCGAACCTTACGGTACTATCGTAGGTGA

CTTTGAGTTTACTAATCATCCTGAAGACGTTGAAACATTAAGCCCTTCTC

>NODE_1447_length_190_cov_1.136095

CCCGGAAAATGTTGTGTAGGTTCGTGCAGATCATGTGGATCGTGGATATACATTCCCTGAGTGATCTAGA

TCATGAAAATGGGGATCTGGTCCGTCTCATGTGGGAAAAGATTGGCAGGCAGGCTTTTACAGGTATATAC

GTAGGCGTGCTTTTGATTTTGCGGATTATGTAGACAGGCTGTTTCCTCCT

>NODE_1448_length_189_cov_62.529762

GAATATACCCGGCTTTCAATCCCACCGAGTTGTTGGGGTTACAATCTAGTCTCGGAAGGTCAGAGGCCGG

GCGCTACATGTGGTCTCCTTTTCCCGGAGGATAGGGAGTTCCACATGCGCTTAATTTTCGCCGCATTCTA

TACCCAGGGGGTCATACCCGGAGGTCCCCTCGGGCTGTTGATGCGCGAA

>NODE_1449_length_189_cov_62.517857

TTTAGTTCGTTTCTTGTTGCGATGGGGGGACGAAGAAGGCTAGGTGAGCCAGGCGTTGGTTGTCCTGGTG

AAAGCATGTAGGCCGAGGAATCAGGCAAATCCGGTTCCTCTGAGGTCGAGATGCGAGACGAACTGACTAC

GGTCAGGAAGTTGCCGATGCCACGCTTCCAGGAAAAGCCTCTATGGAAT

>NODE_1450_length_189_cov_10.886905

CGCGCTTGCTGCCGCCGCGGTCACCATGTACGGTCTCTCTCCATACAGCTCCATGCAGGTCCTCTATGTG

GTGACATTGTCGGTTGCTTCCGTGGTGAACGTGGTCAACGCGCTGTACACTCTGGCAGCCTTGAAAGCTA

AGAGGCTGTTCACCCCGGGGGTGAAGTGGTCGCCCCTTTCTTTCGTGGC

>NODE_1451_length_189_cov_8.750000

GAACGGACACTAAAGGAATCGCCAGGATGGCGCTCAGGAAGAGACGTCGGCAAGGAGCCGATCTGGAACC

ACACAGGAGTGTGACCTCAGGAAGAGGCAGGAACGGCAGTCAGGATGACACAAAGGATGCCGGGCACCAC

GCAAGGACAGTTGTGCAGAAGGAACAGGGAACGGACACTAAAGGAATCG

>NODE_1452_length_189_cov_7.577381

AGGATAAACAGGGTCTCTGCCTCGGGTCCACTGACGATAGCCCACTCGTACAACCCATCGGGGCCAAAGC

TAGCAGGCCCGAGTCCCACAATCCAATAGTCACCATCGAACGGAACCTGCCCAAGGTGCACTTTCAGTTG

CCCCGGTTCAGACTTGTCCACGCCGTATGCGTACCCCGAGATGGTGTTG

>NODE_1453_length_189_cov_5.571429

ATGCCGAAGCAAATCCACGACATCAGGGACTTCCTGCAGAAGGCTCGCCGCCGCGATGCCAAGTCTGTGA

AGATCAAGAACAACAGGGCGACGCACTCGACCAAGTTCAAGATCCGATGCTCGAGGTACCTGTACACTCT

TTGTGTCACTGACAAGGAGAAGGCCGACAAGCTCACGCAGTCGCTGCCT

>NODE_1454_length_189_cov_3.601190

GGGTGATGGATGATTCGTTGGGTTGGGACCGTTTTTTTATAGTTGGATTTGCACCGAATACATTGTGTAG

AAGTAGTATATATATATATACTTAAGAGTAATTATCCCGCCTTTTTTTCGAACTAAATATGCATCTGTAC

AGTATAGGGCCCACAAATAAGAAATATCCTGAGAAGAATGTAACGCTCA

>NODE_1455_length_189_cov_3.220238

TTTTGCTACGTACTACGCGTTCACACTCTTACTTAACAGACGATTGACCAAAATCGTGGCTAGGAACCAT

TTCAGAACTCGCAATATATACATGGGGTCGCGACAGGGTCATGCTCACAACCCGCCCTTGATAAATAAAG

TACGACAATTCACATGGGCCAAAAAATGATCAAGTTGGCGTCTTTCCCT

>NODE_1456_length_189_cov_2.744048

CGCGTGTGCACTTTATGTTGTGCAGCGTGGCACTGAGAAGCTCCTACGTAGTCGATGGATTGGCATAACA

TGGCCGGCGGGTGTAGGGTCTCGGAGGGCATGGACAGAGCCGTAATTTGTGTGGGTCGCTGCGGGTGTGC

AAACGAGTGTTCCCGGTACCGGCTGGAATAGCGAGTTCTCCTGCACCGT

>NODE_1457_length_189_cov_2.690476

ACCCTATCTGTTCCCGGCCTCACGGATGCCGTCGCATCTCTCCGCCTCCGAATCGGAAACCAAGCCACAG

CAAGCGACGGGCGACTTGTGAGCCTTCCACCATAATACAACATGTCGAGTCTCAACATCCCTTACTCCAT

ACCCTCTCTCCTCCTTACCCTCCCCGGCGTGCTCCCCAAGAGCATTATA

>NODE_1458_length_189_cov_2.458333

TTAACCAGGTACAGTATTTCAAAGCATCATTAACGCACCACGCTACGTATGGTGCCCTGAATTTCACGCC

AATAATAAATAATACGCGGTAGACCAAAGCAGCAGCAAGCAAAAAGCATACCGGCCGTACCTATATTATA

AATACACGTCACATAACTCTCTCTTCTTCTCTCTATCGAATAATCGACC

>NODE_1459_length_189_cov_2.363095

TCACGTAAATCTACCCCTTTTGCTGCGCAAGTAGCTGCAGATCGCGCAGGCGCTACTGCAAAAGAGTTTG

GTTTGAAGAATATTGAAGTGTTCGTTAAAGGTCCAGGTCCAGGTCGTGAATCTGCTGTACGTGCTTTGAA

TGCTGCTGGTTACCGTATTACCAACATAACTGACGTGACGCCAATACCA

>NODE_1460_length_189_cov_2.321429

TTCCAGGCATTATATTCTCACATCCCCCCTAAGGGCAGTACTGTCGCTATGATGTCCTTTCCATCCTAGA

ATCAAAAGCATCAAGTCGTAACCCCCGAACGTAGGCGATATGTTTATAACTGTTCAGTACTGTTCTCCAA

ATTATACATACGTCCAATGATATGTCACTAATTATTCCCCAATAGTTTT

>NODE_1461_length_189_cov_2.226190

ATGCTGATCAGATGCGATTATACTACAGTAGTAGCAGTTGTTGTTGTGCCACGTTATTCGCTCACAAGTC

TGACGAGCTCCTATAAGTCTCTAAAAGTTTACAACAGGCAATTAGTGCACGCGCACTGCAACTAAATTCA

TACATATTATGGTGCCCGCCCCATGCATGTCTGTCACTCCCATCTCTCG

>NODE_1462_length_189_cov_2.101190

AAGTACACGGGCCTCCTCCTACTCCTCCTTGTTTGCGATGATGTTGCTTCTGCCACTCGACACATACAAG

GTCCATGCCAATCCGACTCGACCCTCCATGACCAGGGAGCCAACACACATTGCTACCACAAGATTTGTCG

AATCTCACAGCCGGTGCGTAGCTGAGGGCGTTTTTTTTTTGTTTTCTTA

>NODE_1463_length_189_cov_1.922619

TCGGGATGGTCGCAGCCGTGAGCTACTTCGCGTTTAACGACGATGGATCTGGCGGTAATAGTGGTAGTGG

CGGCGGCAGCGGAGGTGGACGAAGTCGACGGGACAGACTAGGTTCAAACATAAGAGGTATGGACGACCTT

CCCAAGCCCCCTCCGTCCAGCTGACGAGGGCCGGGCGCGTGCTAAGCCT

>NODE_1464_length_189_cov_1.863095

AGGTTTCATTAATTAAGTGAACTATTTTATACGTAGAAAAAATTGCCAAATACAAGGCATTTATTTCAAT

AACTAGTTGTTCTAATTATTAAATAAATAACGATGTAGTTGGTGATTTTAGCAAGTAGAAATGATCACAT

AGTTAGTGAGATTGGTATAAAGCACATATACTTCACAAGAGTGCAACAA

>NODE_1465_length_189_cov_1.684524

CTGGGTCATCTGCACCATGCCCCACAGGGACGCGATGAATAGGACATAGAGCAACCAGGTATAGGAGTTC

TGCATCCATTCCGTGTTCTTGTTAGAGAGCCCCCCTGCCGAAATGTCCCCCTGCGGCGAATTCTCGCCGG

TTCTATCTACCTTTGGGCGTGCGGTGAAAAGGGGGACGAACAAGACTGG

>NODE_1466_length_189_cov_1.357143

ATTCCAGCTTGTACTTCTCGAAGATGAGCTTGGTCTCTTCCATGTTCCGAATGTCTCCGTCCTTGCTGCT

AAGGAAGATGTACTCCTCCTTCTCCTTCGCCTCGGGGTCGCTCTCCACGAACTCCTTGATTCCGCAGCCC

ACCAGGCCAGTACCACCGGTGACCATCACCACCTTCTTGGGCGCGGCAT

>NODE_1467_length_189_cov_1.047619

GCAGTCCCCCCACAGATCCCAGGCGAGCCAATAGAACAGAATCTCGTGGTGCTCATGCTCGTGAAGATTT

CCAAGATAGAGATGATGCAAATTGGTTATGTCATTCAATTTACACTCCTGAGACGCAAGATATGTCTAAG

CGTGATGTAAACATGGCACCAGTTCACCGTGAAGCTTTTCCACCTAAAG

>NODE_1468_length_189_cov_1.029762

TCTCTCTGAATCACTCGCTCTGCGATCTCTCCGCCTGCTGCCAATACGGTGGCGGTCCCTATTCTTAGCG

AGTGGAGGGAGACTCGTTCTGGATCTAGCCCCGCTTGACTTGCAATCTGGCGTAATGCCCGAGTTGCTTG

CCTGTATGCCCACACCTTCACAGTCCCGCCTACCCGAAACGCTGCCAGA

>NODE_1469_length_188_cov_75.856287

GTTTCTGGGGTGTTTGTATGTTGACTATGTCCCTGAGCCGATAAGTTTTAATTCAGGAAATTGATTAAAC

TCTATTGTGCAAGCCCGGCTCGTATCGGGAAGCCTACGGTTGAAATGATACTTCCGCCCTGAACACACGG

TGTTCAGGACTACGGTTTACTGCGGCACTCTGGAAGAACCTCTCTATC

>NODE_1470_length_188_cov_49.586826

GCCTCATGCTCCCCGGAGAGCTGGCCAAGCACGCCGTGTCCGAGGGCACCAAGGCCGTGACCAAGTTCAC

CTCGTCTTACTCCGCCTAAACCTGCGACATCTCGGGACGACTCGACACGAAGATGAGTGGGCCACCCACC

CACCGCTGCGGTGTGTCTACTCAATCAAACCAACCTGGTGTTTACTCA

>NODE_1471_length_188_cov_22.227545

AAAGTATACTGAGGTAATTTTTTCGTATTAATTTATTCGTCTTGATCAAATTAACGATTACTGATGTAAA

GCGTTACTTCAAATCCTAAACGCATTTCTTCAAAGTTAGGTTTAGTCCACATAATTTATTCCTCGCTAGT

TGATGATATTAATTTATGTTCTTGCTTACATTGCACTCATAAGCACAC

>NODE_1472_length_188_cov_6.113772

CTGCGGTGTGGCGGTGGTTTGAATTATTGGTTTGGTACAAGCAAGCGTGGATTGGGTCAAGAAGGCTACG

CACGGAACGAGGCTTTGAATTTATTAGTAATAAGTACTGATAACTAAAAAATGTGTAGAATTCGCAGTAC

TTTTGGGGCTGGCCTGACCCCCGGCTTGGTTACTTTTTTGTTTGGTGC

>NODE_1473_length_188_cov_4.916168

GTTCGACCGATGCGATGTGTTCAAACTCGTCCTCGCGTTAACGCTTCCCTTTCCGTTGCATGCACCCTCA

TCTGTGGGTGCAGCCCTGCACGATCAAAAACCATTCTACCGTACCCTTAAAATACCAAGCTCTATACAAA

AAACCGTCGGAAAAGTCGCAAACCTTTATTGCTCACCAAGTCACAGGC

>NODE_1474_length_188_cov_4.616766

GTGGCGACTGCGCTCTTCCTGCACGCTTCGTGCCGACCATGGTTCGGTTGGAGCGGTGAATTAATGAAGG

GTCTCTGCCTCATACTTGGATTGAATGAGCCAAGCCAAAGAGTTCTACTGCTGGTGCTGAATGGTGCTAT

TCGATACGGTGGCTAGACTAAGCTGTTTGGTAACCTCCTTGCCTGGAA

>NODE_1475_length_188_cov_4.251497

ATCGGGCTGTTCGGGCGAGGCCTGAGGACGAAGATCATGGCGAACGGCATGCAGGGCGTCATGTTCTCGG

TGCTCTGGAAGGCGATCGAGGAGAAGCTGAACACGTAAATGATGGAATGGTACAGTAGTCATGGGGTGAT

TGATGTAAGGGGGAGCAAGGAGACTGCCTTGTCTGGAAGCGCCTAGGT

>NODE_1476_length_188_cov_3.736527

GACGACAAGCGAGTAGCTCTGGAAGTCGGTGGTGAGCGACTTGGCCCACACGGACACTGTCCATGTACCG

CTTTCGACGTCAGAGGCGTCCACAATAACCCGCTCATTCACATTTGAATCGTCAGCCACGCCCGACTGCC

ACATTGTGTGCCTGCTTCCGCTGGGTGAGCTCACGGCCAGGTCTAGGT

>NODE_1477_length_188_cov_3.401198

TCCAAATTAAATAATGCCACAGCAGTAAAAGTTTTATAATTAATAGAATAAATGTCCAAAGTCCAAGGCA

AACCTTTGCGTAACGCTCATGATAGCTCCTCGCCGCCAGTGCATGCACGCAAAAAAAAACACAATAAAAA

GCGATCGTAGCGATAAGCGCTACGGCCGCAAGGTCCACCATCGCGTAG

>NODE_1478_length_188_cov_3.245509

CAGACAGACAGACTACTACGGAAGTTTCGCTCTTCTTTTTTTTCGTTTTGTGCGAGCATGTTGAGGAGGA

CTGTTGTGAAAGGCATGGATGGCCTTGTTTGTGCCAGCGTAAGTTGGCGCTCTCGTTTTGTTCTGTGAAA

AACATTTTTTTCCCCTTTGTGATCGGTCTTAAGGGATATATATCTCTT

>NODE_1479_length_188_cov_3.155689

TGGTAAATTTCTCAGGGCAACAGCGAAGAACCTTGAACGTAATCCGACTTTACCCCTGACGACCAACGCA

AAAGAAATTCCGTCCCTAAGGCTATACCGGTGCCACTCCACCATCACGAATGTGCCCATCACAAACCGAC

AAACAAATGTCGCCAACATGAGCACAGCCGTGGCCCTCGAAAGCTTCG

>NODE_1480_length_188_cov_2.814371

CGCTTCATCCCCGGGTTTGCATTTTTGGTTGCTTTCCAGGTGTTGATGGTGACGAAGATGTCGGGAGTTG

TTTCCACTTTTGTTGGATGATTTTGATAATAAGGAAAACATGAGCCTCGTGTGATACCGTGGCGTCGGAG

AGAGAAAGAAAGAAAGGGTTTTGTACTATATGTTCTGCCTTGTGAGGG

>NODE_1481_length_188_cov_2.634731

CCAATGCGTGCGTAGCGACGGTGCGAACCACCAAGAACCTTTATACATTGTACACGTCGAGCGCCGCTGT

TATCAGCAACTTCCAGCTGAGGATGGCGGACTTCGGATTATCGTCTATCATGGAGGACGAGAACGGGGTG

TGCTATACCGAGTGTGGTACTCGAAGCTACATGGCGCCGTAAGTGCTG

>NODE_1482_length_188_cov_2.574850

CCCTCCGACAATTGTGCAGCGTGGTCTTTAACGTTGACCACTCACAAACATTCCCATGTAGTGGTCGCTT

GGTGGTTAGCCGGGTGGGTACAGGCCACCACCGACGGAAATACAGGTACCTGAACCAACTGCCTAGACAA

TCCAGGCTGATCTTGGGAGTTTCAGGGCTGCCCGTAACGTGTGGAGTG

>NODE_1483_length_188_cov_1.970060

GCGTCCAAGGACGGAGGTGCGCGAACGCTACATTCATTCAGCTACTATCGTAACACAACCCAGCACAGAC

AAAAAAAGACAAGAAATAATCCAAGGAGCTGGGCTGTGCTGTGCTGGCGGGCCACCCGAGAGGAAGACCT

GCTTATACACGACACTCACTCCTAATTAAAGCAAGCCTACACAGCAGG

>NODE_1484_length_188_cov_1.928144

ATTCAAAGTATGTAACGTTCGTGCCCGTTATATGTGTGCTGCCAAAGGTCGCAAACGCACGAGAAATGAA

CAAAACAAAAGGGAGCAGAAGATACCGGGGTGAATTGTTGTGGAGGTGTTGATGCAATTGTACTGGACGT

GCTTGCTGTAGAACAGGACACATTAGCGTACAAGTATGGGTGCTGGTT

>NODE_1485_length_188_cov_1.682635

CCAAGGGTGCCGAAGAATCCCGCCCACCGCTGGAGGATGTCTTCCTTGCTCCGGAGCAGCACGCCATTCT

CGTCCGTGACGAACTGCTGCCCACCCGCTTGTCTCCCGCTGAGGCCCACTGATCTCTTAAGGTGTTTGTA

CAGCCCTCTCATGTCGCGGTCCTTGTAGACCGCTTCGAGCTCCGCCAC

>NODE_1486_length_188_cov_1.419162

TCCTGCTCCCTCCTGCTTATCCCGAACCTGCCCGACAAACGGTCGCTGCTGTTTCCCATCACCTCTCCGG

TGGTGTAGTTAGCAATAGCAGGGGTTTCCGGGGCTAGATCCTTAAGCTTGAGCCCTTTGAGAAGCCCCAA

AATACCCGGCAAGCCTTTCTTCTGGGCTTTCCCAAGATTGATCAGGCG

>NODE_1487_length_188_cov_1.401198

ATTCCACGACAATGCCAAAGGTAACGCCAACGGGAATCCAACGATAACGCTTACCGAAACGCCAACGGTA

AGTTTGCTACTAAGCATAGAAGGCGGACCTTTACTTGACCCTATAGAAAAAGCAGGTCTTGCTTCACTTA

CTGCAGGCTTGATGAATGAAGGAACAACTAATTATAGTAAAGAAGAAC

>NODE_1488_length_188_cov_1.365269

CTTTTTCGCTCGTATCCCCGCCAGGTAGTATTGCGACTCAAACAATTGCTGTCACGTTCCTCTAGTCCGG

GGTAGATCAACCTGGTACATGCGTTGTACTTCTCGTTTTTTGTGAATTTCTTGTGTAAACTTAAGCTTTT

TTGTGTTTTTCGTTTGTAACTTATGTGAATCTATGTCATATTGTTTGC

>NODE_1489_length_187_cov_109.638554

TCTCCCACCTATCCTACACATTACTTATCCAAAGCCAATACGAAGCTATAGTAAAGGTGCACGGGGTCTT

TTCGTCCCACTGCGGGTAACCGGCATCTTCACCGATACTACAATTTCACCGAGCTCATGGCTGAGACAGT

GTCCAGATCGTTACACCATTCGTGCAGGTCGGAACTTACCCGACAAG

>NODE_1490_length_187_cov_6.530120

TGCGCCACCGGCTGCAGGCTCGATGCCGTACTCGTCTTTGAGGATTGTTTTCAGTTCTTGTGCTTCAAGC

AGTGTCAGACCAACGATGTCTTCTGCCAGTTTTTTCAGATCAGCCATGTTTTAGCTCTTTCCGTTAACGA

TATGTGTGTTCCAACGTGTAGTTTCAACTAACACGCCGATTTGACGA

>NODE_1491_length_187_cov_4.903614

AAGAACGGTGTTCCACTAGTCGAAAACACAGGCTACGGGAGTATATATACTCCAGATTGTAGGGTGTTTC

GCGTGCTTGGTCTACCTCTCCTCCTATTACCTCGACCATTCCAATCAATCTACTTTCGAAAATACGCTAT

TCTTACAAAGACAAAAAAAAAACGCAACAAGAGCGTCGTCAACCAAA

>NODE_1492_length_187_cov_4.656627

CCCCACCCGGAGCTTTGAAGCAACCCAGGCACAGCCATGGCGACCAACATGTCCAAGAAGCGCAAGTTCG

TGGCCGACGGTGTTTTCTACGCCGAGCTGAACGAGCTCCTAACGCAAGAGCTTGCGGAGGAGGGGTACGC

TGGAGTGGAGGTTCGCGTAACGCCTCTGCGCACCGAGATCATCATCA

>NODE_1493_length_187_cov_4.469880

AACTAAACTCCCTTCAAATGTAGGCGCCGTAATTTTGTATTTCCTGTACTCTAAGTCCCCGACCTTCACA

GGCCTGATACGTGAAATATTGTCGTCATACGTCTGGGGCGTTCCGCGCAACTGAAGGGTCCCTGCAAGTC

CCTGCACAACTCTACCTATCTATGGCGTTGTGCACGCGTAGCAACCT

>NODE_1494_length_187_cov_4.469880

AGGAAGAAGCCTTCCGCCATCTGGTACGGTACGAAGCTGTTTGGTAGTTTGGTGGTTTGGATCGCGTTAC

GATCGACTGGGAAGAGGCTCGCCTGAGGCTTTAGGGCAGAGGCAGCGATCTCAGATGTTAGTATGATAAA

AAGAACGGTAAATAAATACAGCATTTAAACAAAAAAAATATCAACTC

>NODE_1495_length_187_cov_4.313253

ACTGTAGACTGACAACTCCGCTGTCCCGCCGGTCAACACGACCACTACAGCAGCATACATACTGACAACA

CACACCAAGCGACAGAACATTTCCACGGTAGACGTACAGAGCCAAGACGTTCAAAGTTGGCTTTACTGTC

GTATATGACCCGAACACAACCAGGTGTGTACGCGGAAACCCCCGATA

>NODE_1496_length_187_cov_4.246988

TTCCAAGTACCGCCCATACAACAAGCCAACTTGCTTCCCGGTCAAAAATCTTCTCTAAATCCCCCATCTT

CCACGCACCGGAAGGATATCGACGTGCGTGTTTCATTTCCTCTCCAGCCTCCATACTACTGAACCCCCTC

CTCCTACCCCCAGGGAACCGCTGCAACGACAAACATGCCTCCATGTC

>NODE_1497_length_187_cov_3.680723

TGCCAGTTAATGCTAGTGCGCCTACCATTGCCGTAGCGATGAGGCTTTTCTTACTTACTTTGGACATAGT

TCCCAACCAATTGCTATTTAAATTTTGATAACAAAGATTTTTGTTACCGCAAATATAAAAATCTGTCGTT

TTATTATTTGTAACTTTTTTTTATACAACCTATCAATAATCCCAAAT

>NODE_1498_length_187_cov_3.445783

AATACAGCCCTCAGATCTACTAAATACCCCGAATACGAATGCCATCCAAGTTATACGAAAACGGACCAAT

TACCTATACAGAGGTAATTACCGGCCTTCCTAGCCCATACCATTACACTAGCCCACCCTGATGAGCCGTG

TATGCAGGAATCTGGGAAATTCGGGAAAATCCCAATTCCACGAACTA

>NODE_1499_length_187_cov_2.873494

CGTTCTTACTAGACGCTCCACCGGTGTACCAATTTCTTTGGCAAGTTCTGCTACAGTTATATCTGCCATT

TACACTATTCTCCCGGTGTTATTTATTCTTCGTTAAACCAACAAATGTTACGTGCAGCCATTATTAGCTC

ACCCGCTTTGGTTTCAGTTAATTCTTCTGATGCTAATTTTTCTAATT

>NODE_1500_length_187_cov_2.849398

CGAACAGCGCGATGCTCGGGGTCAGGTACCTCGGAAAACCGTCCACAGGCGTCGCCTCCATCCCCGCGAG

CAGCTCGTGGTCAACGACGAACAGTTTCCCGGCCTTCATGAGCGAGTCCACTTTCTTCTTGAGGACCTTG

TAGGCAGGGTCAGAGTCCAACACTTTCAGCTTGGCGCGGGTGGCCTC

>NODE_1501_length_187_cov_2.807229

ACCGGCCAATCAACCATCTCCCCTTCCGGAAACAATTACACAGGAGTTCTACATGATTGCGTTTGGGGAT

CGCCTCGTCGCGAGAGAGGTCGATCTGTGTCCCCGCTACCTCCAACTTCGAATCTCCTTTTCCTATATGT

TTATGTAGACTGCCGCTTGGCACCATTCAAAAGTAAAATGTCGAATT

>NODE_1502_length_187_cov_2.765060

GCTGGGAAACAAAAGAGGGAGGGGAAGTACAATCGTAGTCTCGTGGAATTGAATTAGACCCGAAGCCAAG

TGTCTGTGGGCACCTACCTATCTATCTATATGTTGATGTACCTCTGGATTATCCTTTTGGTATTTGGTAT

TTTTGTAATTTGGGTGTTGGGCGTGCTGTTGGTTTTCATGCGATCGG

>NODE_1503_length_187_cov_2.536145

GCTTCTGCTGGCCAGAGCTATCAGTACCTACTAGCCCTGGCCCTAGCAATTCCAGACAGAGCCGTACGGT

CGCCGAAGAGAGAGGAAAATGACAAGGAGGCGCCACAAACCAAACCATATCGGCCACGCAAACCTCTCAA

GCCCATGGGCAGGCGGGCGAAGCATTACTGAGACCTCGAAGACGGCT

>NODE_1504_length_187_cov_2.500000

TCAACAAAGGTCGCCGCCAAGGACAAAGGATGGGCTATCGCCTTCAAGATTAACGTCGTGATGACTGTGC

TTGCCGCAGTATGGTTCGGCAAGATGGCCGTAGACGCTGTGGCGAACGACTCCTTGTCTGGAGCGGGGCG

CCGTCGCCTTCAGGCTCACGGGGGCTCCGAAGACGAGGATGGGACCG

>NODE_1505_length_187_cov_2.409639

TGTCTTGTTCTCACCTTCTCCTCGAGCGCCATGACCTGCCGGTCTCTCGTGCGAATGTGCAGTGCCATGT

CGCCCATTCTTGCGTCTTTCTCCTTGATTTTCTCATCAAATTTGTCTTCGTTCTGCTTCATCTCCCTTTC

AAGCCTGTTACGATCGGCTGCTTGCTTCTCAAGCATAAGCTCCGACT

>NODE_1506_length_187_cov_2.132530

TTGGTCGACTCTTCCCGGTGCGGAGGGACTCGAGGTGAAGGACGGCTTCAACATGTTCGCGGGCTACATC

AACGTCGACCAGGAGAACGGTCGGAGAATTTTCTACTGGTTCATGGAGGCCCAAGAGAATGCCGAGGACG

CCCCGGTGATCCTTTGGACGAACGGAGGTCCGGGGTGCAGTGGCATG

>NODE_1507_length_187_cov_2.036145

CTCTGCCGCTGTTGCTCCGAGGTACAAACGGTCTCCAGCATCACCGATGCCGGGTACCAGGTCCAGTCCA

TTCTCTGAGAGCTCGTCAATGGAGGCAAGGTGCACCTCCACGTCTGGGTGCATTTTGCAAAGGGCCTCCA

GTCCGCTCCGCGATGCCACCAATGTGACCACCTTAATCTTCTTCGCT

>NODE_1508_length_187_cov_2.018072

TTGCTGCTGATCCTACTTTTCTGCCGGTGCGGGTAGTGTATTGTTTGTTTGTTGGATGGTTATTTCGTGG

TCGTGCTGTGGTCGTGTAGGTCGTCCTCGTCTCTTGGGCTATGGGATGGGGTGGACAGGAAGGAAGGAAT

GACACTGGGTGGATGAAGCGGGTGCCGCTATACATAATAGCCGGAGC

>NODE_1509_length_187_cov_1.066265

CGTCCCAAGGCTCGTACCTAAACCCGGGTATTGCGTCGACGTGTATACCCGTACCCGGGTAAAGGTACCC

GGGTACATACCCGAGTACAGCAACGGTCCCTCTATAATGCCGGTAGGGCAGCAGCTTTTTGGACCTATAT

CGGTACTCCCACCCATCATTCCCACTACTCTTTGCAAGGGTACGTTT

>NODE_1510_length_187_cov_0.975904

TCCATCGTACCCGCAGAATTGCCCCCATCCACCGCCAATGAGGTGCCAGTGATGTTTTTGGCCGCCGAAG

ACGCAAGATAAAGCGCGGCCTGCCCGATGTCCTCTGACTTGGTTTCGCTGTTAAGCGGTATGGCCCCCAT

GAACAGCTTCAAGGGCGGCCAGGATTCCATTGTACAGCGTTTCGTCC

>NODE_1511_length_186_cov_8904.406061

TGGCACCGAGAATTCCAAAGCTCAACAGGGTCTTCTTTCCCCGCTGATTATTCCAAGCCCGTTCCCTTGG

CTGTGGGTTCGCTAGACAGTAGATAGGGACAGTGGGAATCTCATTAATCCATTCATGCGCGTCACTAGTT

AGATGACGAGGCATTTGGCTACCTTAAGAGAGTCATAGTTACTCCC

>NODE_1512_length_186_cov_6.309091

GGTGCACTTTGGATAAAAAGGCTCAACATCAGTTATAATCGTTCGAAAATAATTGTACAACGGTAGTGTA

GTGCATCTAATGTAAAACATGAAATGTATGATAGATAGGATGATTATGATTGGCAAGTCAAACTTATGCC

TCTAGGCTGTGATGTTGGTGATATGGATGTTATGTTAGATGGACTA

>NODE_1513_length_186_cov_5.781818

TACACCCTAAATGGACAAATAGCGAGGGTTCCTGTCAGGTTGTGGTTCACGTCTAACCTGTGTATAGTAT

ATAGTGTCACTACTTGCGCATTGATTCGTCCATTCGTAAATCTGAATATCTACACACTACTTTTTCTGTG

CGCCTGAAGTCGGGTATCTGATATTAGCGTTGGCCTACTATTAGCG

>NODE_1514_length_186_cov_5.745455

ATTTGTGACGCCATCAGCCCTCTCTCCTTAATAGCTCAGAAAAATCAGTCGTTGGAAGTTGAAGTTTGTG

TGTGGAATAGTGACGCCAGCAGCCCTCTCCGCGGCTCAATAGAAACGGTAGTTGAAGTTGAAGTTGAAGT

TTATGTCCAAGTTGAAGTTGTGTGGATTTGTGACGCCATCAGCCCT

>NODE_1515_length_186_cov_4.072727

ACGTTGGACCCATTATGAACCAGAGTCACGCCAAGACGGTCGCGGCCGAGTACATCCGGCAGCATCCCGA

GTTAAAGTGGACTGGACACTGGAACACTGTGGGACCGGGAAAATCGGTCATCAACGTGCGTGCCGCGACG

CCTTCTTACCCCGCGTATGATGTAGACGTTGGACCCATTATGAACC

>NODE_1516_length_186_cov_3.357576

TTGCTTGGACGGTGATGGAGGATGCTCCGATTTTCGCAGTTGGGGTGGTGTAGTCATAGATTAGGCGATG

TGTGATGTGAGAGACAATTTGACGAGGTTGGAATGGTAAGATTCGGCGAATTAATGGTGACCTTGGTGTG

CGAGGTTGGGCCATGCATTTTGGAGGACTAGATTAAGTGACTGGAA

>NODE_1517_length_186_cov_3.169697

TGATTGTTTGCTCCGTCGTAAATATCGACGGACTGTTCTCGCCGCTTGTGGTAGACCTGCACATGGTCGA

ACACGTCTATTCCGATTTGCCATCCGTCCGGCGAACGCACGTCCTTGGTCAGTAGCCCTCTCGTCACTCC

TCCTTTCATGTAGTGCACACCCAACAGCACTACACCCTGGTCCACC

>NODE_1518_length_186_cov_3.163636

GGTTCTTGACGTAGAGCACGCACGTCGCGAACGTGGGGCAGTTGGAGTCGTTCGGCACACCGTCATCATC

CTTGTATCCGGGCTGCGAGCCGTCTTCGTTCCCGACGTACTGTCCAAGAACGACGTTGTCCGGGGTGACA

GGCTCGATACACTTCAACACGGCCACCTTGGCGTCCCTAACGTAGT

>NODE_1519_length_186_cov_2.715152

TGCTCCGTCCTCGGAGCGGTGTTCGCGGGAGTACTTCTTATCCTCCTGATCCAAGTAACCTTGACGCTGT

CCGCCGTGCCGGACTGGGGCGCGGACGCGAAGCAAGCCATGATCGACCTCGAGATGGAGAACATCTTGAA

GCTGGCCTCGGACAAGGCCGAATTCGTCACGGAAATTTTCGGGCGA

>NODE_1520_length_186_cov_2.339394

AAAAGCAGAGGACTACGTATTACCAAATCGCCCGGTAACATCTTGGATCAATGTACGGGCGTAGAACCCC

TAACCCTAACCCCAAACCCGACCACACTATTAGTGACCTGTAAAATTCCCAGAAATATCCTGCCGGACGA

AATGCTAATACTACACGATAACATACATACGAGTGTCGACGCAATT

>NODE_1521_length_186_cov_2.012121

ACTATTGAGTTCGAAACGATTATGGTCTTGCTCCGGAGGAGAAATGCAAAAAAAATACACGGCTTTCGCC

GGCCAAAGCCACACAAGGACAGAGCGAGACGCCAAAACCGAGCAGCTAACATGAAACTATGGATACTTTG

CCCCCTAAAGCTCTGCAGAAGCTATTAATAGCTGCTGTGCCAGCAG

>NODE_1522_length_186_cov_1.775758

TTCCAACCTTCTTGTTCTTCGCGGCATCTGCAGTGCGCTGTGCCTCCGTGCACTTCACGCTCTTCTCAAC

GGTGGTGATGTATTTCTGCACGTTCTTGCTCTTGTTCTTGTTCTTCAGGCCGAAGGTCTTGTCCTCGATG

AGCTGATTCTGCTTCTTGGCAGCGGCTTTTTTGCCTTGCTGCTGCT

>NODE_1523_length_186_cov_1.266667

CCAAGTCGCTGAAACGCACGACTCGCGCAGCTTCGTGTGCAGACATGGTTGTCCAGTTGACCGCGCGGTC

TGCTAGCTTCATCGTCGGCCGCAAGGTACCAATCACATCGGAATGACTGTAGTTCAGAGGCCTGGGAGGA

ACACCGTGCTGGAGCCGTTCCAGGGCAGCCTTCACTCCGTCCAGAG

>NODE_1524_length_185_cov_81.439024

TCTCGGCCGTTTCGGATGAAGAGCGGCTCATCCGGGTCGACGAATATAGTCTCCACGGTGGAAGCACCGT

TCTGCACATCGAGCCACGGCTGGAAGGCCGTCATGTAGTCCATCCGCGGGAAGAGGGTGACGGCCTTAGG

CGGCACCACGATAGAGTCGATCGCGAAGTCCGACAGCATGAACTG

>NODE_1525_length_185_cov_16.432927

TAGTCGATCCTAAGAGATAGGGTAACTCCGTATCAAAGTGCGCGATCGCCTTCGGGTTGTTTCGCGCCGC

CTATCGAAAGGGAATCAGGTTCATATTCCTGAACCAGGACGTGGATACTGTATGGTAACATAAGTGAACT

CGAAGACGCCAGCGGAAGGTCCGAGAAGAGTTATCTTTTCTTTTT

>NODE_1526_length_185_cov_8.871951

TCTTGATCTCCTCCAGGGAGGCACCAACGATCGCGCGGTTGATCTTCGTGGTCTTACGCGTGCGGCGGCG

AGTCACTTCCTCGACCTTGATCTTTTTGTTCAGGCGCCTCCATGCCTGGGTCCAAGTGAGCTTGGCCGGC

TTCTTCCTCTGCTTGAGGAGGCTCTTGCACTTGGCAGAGAAGAGC

>NODE_1527_length_185_cov_7.000000

TTTTTTGTTTTTCCCCCTGTAAATGTCAGCCTGAGAGTCTTCATGCGCCCGTTTTGTTACGAGGTTAGTA

GGTGTCCCGTGATGTATGTCATCACGCCGCGGGGTCGGATGGACACACGCGTATCTTTGCCGTTGGTCGC

GTATCTTGGCCGTTGGTGCGACCTTTAGGCGCTCGATCTTTTCGC

>NODE_1528_length_185_cov_5.359756

GAGACAAGGCCGGCAATCTTATCCGCTCGGATCAGCTCTTCGATTTCCTTCAGGTTCTTCTTTAGGTCGG

CGTTGCGCGGCTCCACACACAACCCACGCTTCAGAACGTCAGCTGCAGCCTCCAACTCGTTCAGGTTCTT

CTTCGCAGTCGCCAGTCGGAAGTAGCCTGACAGAAAAATGCGTGC

>NODE_1529_length_185_cov_4.957317

CCAGGCATCTTTGTTTTCAACGCCACCGGATCAGGTTACTGTAGAAGAATCATCTTCTGCACCTCGTCCG

GCCGCTCTCTTTGTTAGCCAGTCCCCCTTCAACCATGTCCGGTTACGCCAAGGTGAAGGGCCACGGCCAG

GAGGAGGCGCGTGTGCACCGCATCCGCATCACCCTTACCTCCAAG

>NODE_1530_length_185_cov_4.067073

ATCTTCTCTTTCTGAGTTGTGCTCTCGTGTCGGATACGGTCTGTATCGCTCCGCATGATGCGGATCTCGT

TGTCCAGAAGTCGCATGCGCTGTCGCAGCTCCTCAGAACCCATCTTCATAATCTCTTCATCCAGTTGATC

AGCCTCGTCCCCCCAGATCTCCTTCGATTCCAAGTCGGCCGCTGC

>NODE_1531_length_185_cov_3.932927

GTGTTAAGACAGTATTAAATCAGAAGGCAGGTGCGTAATGAGCGAAAAAAATACTCGTACTCTTCAAGGT

AAAGTAATCAGTGCGAAGATGGATAAGTCTATCGTTGTAATGATCGAACGTCGCGTTAAGCACCCTATGT

ATGGTAAGTACATCACGCGTTCAACTAAGTTGAAAGCACATGATG

>NODE_1532_length_185_cov_3.890244

AGATTGCAGTTCGACTTAAAACGTTGTAAGGGTATTGAGTCAGCTAGTGAGTGTGTGAGGTTCTGTTATC

AGCTTGTCAGGCTCTGGGGTTGGGTGGTGACTTGTTGTATTGGAGCATATATATAGATATATGCATGTAT

GGTAAAGGAAACGTTGGCACCGCAGATGGTCAATACTGCGCGTTG

>NODE_1533_length_185_cov_3.835366

TCCCCCGTAGCCAACGCCGCGGTCGAGGTCCTCACGAGGCTGAAGGGAGGAGGGCCTACGAAGGAACAGA

AGGAGTCCGTCAAGGTCCTTCTCGACTCCATCACCTCCTCCAAGGTCTGGCCGAGGTTCAAGGAGGTGCT

GGAGAAGAACAAGGTGCCCATCACCGATCCCGAGCGGTCTTTCAT

>NODE_1534_length_185_cov_2.865854

CGCTGAACGAGACGACGGTGAAATCAACGCCGCCCTCCGACACCGACAGCGTTTCCCTAACCGCATCGGA

AACACTAATACTGGCCAGCGCCACACCCAATAGAGAGACGCGACCCGCACTGATCATCTTCATGGCTGGC

GATCCTGTTAGCGAAGTTCGCACGAGTGTCTGGGGTGGCTATGAT

>NODE_1535_length_185_cov_2.817073

TGGCCTGATTCAGGGAACAAGGAAAAGGGAAGTTTCATTTGTCTCGCGTCTTTGTCTTGTCTTTGAACGT

GTTCTACGATGAACCTCTTTGGCAAAGCGAAGAGCAAGCCCCAGGCCGCGGACACTACGGGCACCATTCT

CAAGCTTCGGGAGACGCTGGACTCGTTGGACAAGAGAGAAACTCA

>NODE_1536_length_185_cov_2.640244

CAGACCACTCGGCCGCTTTCCTCGACAGCAAGTCGCTGGGAACCCTGCACAACGTGAAGCATGGCAAGTC

CGGAAACACGCCTATTTGGCTGGAAACGGGGCTGGCGGGACGCCTGAAGAGGATTCGAGACGTGTTCACT

GGCCCCAAGGACTCACCTGTCGTTCACGGATACTCGGTCGGCATG

>NODE_1537_length_185_cov_2.012195

TGGCGCATGCCGCACATCGTTCGTTGCGGAGGTATGATAGCTCTCAGGAGCGCAGGGCAGGTGCTATAGA

GGACAGATGGATGGTTGACTGCCACGATCATAAAATGTGCTTGGCTTCTGTTCGGGTATCTTGGTTGGTA

TACCTTTGGCAGGTGGTACAGCACAGGCTTATTGCGCTGGTGGTG

>NODE_1538_length_185_cov_1.963415

GTATCGGATTCGCCTCTCAATTGTGCACGGGTAAATTTTTCTCGGCGTGTGTTGCCCCAGCACAGGAAGG

CGAGTCTGCCAAACGATCAGACGCGGTGTGTGGGTGTGTTTTGTTTTTGTTTTCGGTTGTCATGGCACAG

GAGCGCCGGCCGACGGGTATGGTCTATAGCAGGCAGCCAGGCCAG

>NODE_1539_length_185_cov_1.926829

GGCATAGCGGCCAACCGCCACATTGCGGCTGGCGAAGGCGCGAGCTCTGATCATGGCTCTACTTCTCGCG

AAGACTTTCAACGGCAGCAGAGGTACACTGCTTGTAGACTACAATACAACCTCCAAAACATATCAGCGAA

TGCCGGCAAGATTCTTGCCGAAAGGGAGATCCATCGCGCGTGCGG

>NODE_1540_length_185_cov_1.829268

TACCGCCAACATGGCCAAGGTGCAGGACGGTCTCGGCAGGAAAGTGGGCGACATCGTGCTCAATGGGCTC

AGCGGGATCGCGCTACTCGCTACCGCAATCCTCATCAACTGGGAGCTTGGTCTGGTCATGCTGGCGTGCG

TACCCTTCATAGGCGCTAGCGTGGCTATCCTGTCCAAGCTTATGT

>NODE_1541_length_185_cov_1.567073

GCATCGGTGAAACCGGCAACCGCGATAGAAATGCGGTAAGCCTCATCGTCGAGCTTTTCGATGTTGTAGG

GCGGATACGTCGGCTGGGACGTTTCAGTGGTCAGCACACGGTCCATCAGATTGGCGATTTGATCAAAACC

AACAGTGGAGCGGTAAAGCGGTGCAAAATCATAAGTACGCATAGT

>NODE_1542_length_185_cov_1.463415

GTGTCTGTGTTTTGTGTACATGAATATGGTGGTGGGAGAGAAGGGTATGTCGGTGTTTTGGTTGTTTAGA

TCGAGGCGTGTTACGAAGCGCGCGATGCATACGCGTATACAGACAATGTTTTAGGTCGACATACGCTTAA

CCCGGGACCGGGGGGGGGGGAGCCTTGCACATGCAGAAGAAGGAC

>NODE_1543_length_184_cov_1475.613497

GTCCGCTGAGCTGGTTGTCTCAGGGGGCAATTAGTTCAGGGGGCGATGAAGATTGCGATCAACGACTCGA

GCGGTCAGGTCCACTCGATACCGTTTCGTAAAAGACGGGGACGGGCCGCGGTGATGAGCGTCGCGAACTA

AAATGTCATCCTTACTCTGAGCTCTTGGAATACTCGGCTGCCAA

>NODE_1544_length_184_cov_1156.368098

GCCTGGGACTCGGCGGCTTTCACCTTCCTGTCACCAGCACTCCTACACCTTTGATGGTTGAACTGTCGCA

GTACAATCACTACTTCCATACGCTTCCCTGTTAACAATTTCAAGCACTTTTTAACTCTCTTTTCAAAGTT

CTTTTCACCTTTCCTTCACAGTACTTGTTCGCTATCGGTCTCCC

>NODE_1545_length_184_cov_47.368098

AATCTCTAAGCATTTCACCGCTACACTACATATTCTAACCACTTCACAATAATTCAAGACATACAGTTTC

AATGGCAATTCTATGGTTGAGCCACAGACTTTCACCACTGACTTATATGCCCGCCTACGGACCCTTTAAA

CCCAATGATTCCGGATAACGCTTGCACCCTCCGTATTACCGCGG

>NODE_1546_length_184_cov_19.754601

ATAAAAAAAGCCCGTAAGGGCTTTTTTAGTTTCATATCTACCGTCACCGATATGAGCAATGTAGCAAGTA

AAATAGTAGGGCGGTTACTCATCATCAACATCAAGGGGCGTTGCGGTTCATAACAAGTAACCATAGCTGT

TTACGTGCGCCACACGTTGTTTCAGGGTAGAGAAACAAATCAAT

>NODE_1547_length_184_cov_15.963190

ATCTTGAATGGATCCGTGCATTTCACCGCTAACCATCCAATATAAATACCCCCAACTGTCCCTCTTAATC

ATTATTTAATTCATTCAAACCAACAAATATATTGACCTAAATCCTATTCTATTATCCCATGCTAATGTAT

TCAGAGCCCCAAGAGGCTGCTTTAAACACTCTAATTTTTTCACA

>NODE_1548_length_184_cov_12.828221

GTTCCCTGGGATATCGGAGCTTGGTTAAGTCCCTGAGCCGATAAATTTTATACCAGGATTTCTGTTCGTC

TACTATTGTGCAAGCTTGGCATGTACCGAGAAGCCTACGGTAAACCGCAGATTTCCGCCCTGAAACCTTC

TGGTTTCAGGACTGAAACCGAGTACCGTATCTTGGGGAACACCT

>NODE_1549_length_184_cov_8.196319

GCGTAAACTAACGCTCCTGCTACTTCCTGTAGCGTCATCCGTGATTATCCTAAAGTAACTCCTAGCTACT

AAGTGCTTCATGCACTTCCTTATTCTTCCTTGTATCCGTTTTAGCTTGTCCTTTATGCTTCCTGCTAGCG

TCCGCTATGTTGCCTATTCCGTATCCTTGGCAGTAACTCAATCA

>NODE_1550_length_184_cov_6.000000

CTGTACGGAACCCCGTGGGGACTCGGGGTTAGCTTTGCTACCAGAGCCTGGGAGAGTTCGACATAAAGTC

CAAGGCTGTTGTGGAAAGGACGTAGAAACGGCCCTTGGCAGATACAACTGCTGGTGGAACCGCAACGAGT

TATGATATATTGCACGGGTGAAATTCGAGCGATGGCGGGGTGTG

>NODE_1551_length_184_cov_4.079755

CACATCACAAAGACGAACGGACCTGGTTGGCCAGCTTCTTGCCGAGCTCCACACCCCATTGGTCGAACGA

GTTCAGCCCCCACACGAAGCCCTGTACCGCAGTGCGGTGCTCGTAGATGGCCAGGAGCTGGCCAGCCGAA

AAGGCGTCGAGCTTAGGGAAGAGAAGGGACAGCGAAGGCCTGTT

>NODE_1552_length_184_cov_3.993865

AGAGATCCAAGGTTCAGAACCAAGAGGGCGAGGAAAGCGAAAGTTCCGTGGGGAGAAGAGCAGAACAAAG

CTTTTAATGCGCTTATCGAAGCTTTGACGTCACCTCCAATTCTAGCCCTGCCGGTGTGGACAGAACCGTT

CTCGCTGTACACCGATGCCAGTGAGACAGGAGCAGGGGCGGTGC

>NODE_1553_length_184_cov_3.846626

GCCGTTTTCGAACCGAACTGTGCCCTCGTTGTGGATGCAGCCTCCAGTCCAACTGCCGAACGCGAAGAAG

GAGCCCTCGCCAGAAAGGACGGCGATCGTTACGTCGGACATTAAAGTTCTCGAGTTGAAATGGGCGTAGC

CCTCTTCGCCGACACGAAGGAAGCCACCATGCACTCCCTGGGTC

>NODE_1554_length_184_cov_3.220859

TGATATATTGACGGTGGGGCAGCTGGAAGGCTGTACTAGGTGCTGCTCGATCGTAGCTTGTGCGTAGATG

GAACTGGCGCAAAATACACACATCATGGTATGTGCATGTACGCGCCGGGAGGGAGAAGAACATGGGCGGG

AGTGGTAGTATTCGGTGTGATGGTTGCCTTGAATGGGACTACAC

>NODE_1555_length_184_cov_2.785276

AGCACCTCCCTGGCGAGGAGCCAGTAGAGGAGACCGATGGACAGCTTGCCCTTGTTGTTGGACGGGATGG

CCACATCTACGTTCTGCAGCGGGGAGTCGCTGTCGCAGAAGGCGATCACCGGCACGTTCACGTACGAGGC

CTCCTTCACGGCCTGGGCGTCAGGGCGGGGGTCGGTGATGATGA

>NODE_1556_length_184_cov_2.730061

TTCTCCTTGGCCTTCGTAAAGTCGTCCATCGTCACCTTCATCCGACGTTCTCGCAGTGCCAACAGTCCCG

ACTCCGTGCAGATGGCCTTGATGTCCGCCCCGGACAGCTCATCCTTAGCCATGATGAACACCTCCAGGTC

TACATCGTCCGCCAATGTCATGCGCCCCGTGTGGATGCTGAAGA

>NODE_1557_length_184_cov_2.638037

CAATTTTAAAAAAAAAAAAACGACAACAGAAAATATTTTTTTTCTGATTTCTGATTACAGCCGCCGCGGG

TTCAAGCCACCCTCCTTACAAGAAAAAGAGGTATGTTTTTCTGTCTCTGATTACAGGCGCGGCGCGTTCA

ATCCACCCTCCTTGCCATGGGGTTTGCACAACCATAATGGCATG

>NODE_1558_length_184_cov_2.552147

ATTCCAAACCTCCATAGTTACTTACTTAAGTAAAGTAAGTAACTATAGATTAGAGGGAGGGTCTACTTAG

CGCTGAAGCTGTCTAAGTAGGCACGGTCTGAGAAGTTTGTCCAAGCACGTACTTGTTCTTGGTAGCTTTT

GATAGAGTCTAGAATCTCTTGAGTTTTTGGATCTTTACTTGCAA

>NODE_1559_length_184_cov_2.030675

CAGTCCCCGCAGGCTCTGCGTATGCTCTCCACGATAGACAGAGGTAAGCGCATCCTGTTCGTGTAGCTCC

CGCCCCACTCGTCTTCCCTCTTATTCGTGTGCTTGACGAGGAATTGGTTGATCAGGTAGCCCTCGGACCC

CATCACCTCCACCCCGTCGTATCCACCCTCCTTCGCCAGCATCG

>NODE_1560_length_184_cov_1.779141

GAAGCAATCACTCCTCCTCGTCGCTGACGTCGCTGCCATGGGTGTTAAGCGCGGCCTGTACCATAGTAGC

CGTCTGCTTGAAGGTGACCCCGGCCAATATTCCCAGTTTTTGCTTGCGCGGGACAGCCGCCTCCCACATC

CACAAGAGCATGATCACGCCAAGGAAAGCCCCCATGGCAAACCC

>NODE_1561_length_184_cov_1.711656

CTGGTGCGACGATTTCGGGTTTGATGCGATTATCCCGGGTAGGGCCGTAGGAGCTAAAGACAGCCACCGT

GTCAATGTCAGCGGATCCGTTTGTGCCGTCCGCCTCTGTACCATCTTCAGCGGTGGCCGTCTTTCGCGTT

TGACCCGACGATGTAGCCCCGACTGTAAGAACATTCTTACCAAT

>NODE_1562_length_184_cov_1.570552

TTTGGATTTGAATTTGGGATTAGTCTAGGAACTGCTGGCCGGTGGGTGCTCTGCTTGGCCGCCGCGATGT

GTCGTAAAGGCTTGGGACGTTGGCGGATAGGGGCCGGGATGTTTCTGTGGGGAGGTTTGCGAGTTCGTTC

GTTCGCTCGCTCGCTTGCATCGCTAGCGTGTTGGTGGTTTCGGC

>NODE_1563_length_184_cov_1.570552

GCTCATTCTCTGCGTATATACAGCCACCGACGTACGCGACGTAATTGAGTTTATCCGCGTACTGAATCTG

AAGCGGCTGTGGTGCCGCCGAGGAGCACTGCCTGTACGAGACCGCAAAGTGGTCTGATGTGTCTTCCGAT

GTGAAGTCGATGCTCGCTCTCGCGTCCGGCTCAATAACAATCGT

>NODE_1564_length_184_cov_1.478528

TTCGGAACTCTCGGTGTTCTGGGACAACGAGGACAACGTGATTGGTATCGACACCACCAACCCCGCGGTC

GTCGTTGCCTCAGGAGGCGAGGCTGACGTTCCCAACTCCGTTGTGTTCTACGGAAACGACGGGTCTGAGT

GCAGGTCGGTGTTCTTCGAGGAAGACGAGACGTGCCTTTGAAGC

>NODE_1565_length_184_cov_1.128834

TATTCCAGAAAAAAGTGCAAAAGCACATCGTGGTGAATGGTTAAAATCAGCAGTAGGCTCTTTTGAGGCT

CGTGGTAAAACACTCGGTATTATTGGTTACGGACATATTGGAACTCAACTTGGTATTTTAGCCGAAACAT

TAGGTATGCGTGTTAGATTTTACGACATAGAAACTAAATTACCT

>NODE_1566_length_183_cov_911.432099

AGTCAAACTACCCACCAGACAGTGTCCCCAAGCCCGATTAGGGCCCTAGGTTAGAACATCACGCATACAA

GGGTGGTATTTCAAGGTTGGCTCCACGTCATCTGGCGACAACGCTTCAAAGCCTCCCACCTATCCTACAC

ATGTAGGAGCAATGTTCACTGTCAAGCTATAGTAAAGGTTCAC

>NODE_1567_length_183_cov_216.962963

GGAGCCGTAGCGAAAGCGAGTCTGAATAGGGCGTTTTAGTTAGTTTGAGTAGACGCGAAACCGTGTGATC

TACCCATGGGCAGGTTGAAGGTTGAGTAACATCAACTGGAGGACCGAACCCACTAATGTTGAAAAATTAG

GGGATGACCTGTGGGTAGGGGTGAAAGGCCAATCAAACTCGGA

>NODE_1568_length_183_cov_204.734568

TTGGACGGCAGGCTGGTGTCTGTTCCTTAGACTCTACCTACTTGTTGTATTTATTTTGTTCACCCTTTAA

AGGCAGCATTGGATAAGTGGTCGAGTGGTTTATGGCACCAGTTTTGAAAACTGACGTAGTTAAAGCTACC

GTGGGTTCAAATCCCACCTTATCCTCTGGAATTCTCGGGGTGC

>NODE_1569_length_183_cov_104.388889

GGCACCGAGAATTCCAAGCCCTCAGTTCTCATAAGCATTTGGCACTCATCGGCTTATCAACTGCCAAGCT

CCTCAGGCCAGGCAAGCTTCCAGGCACCCTCACATTGAGGCATGCGGGTCGCCGGCGCCCGATCTACAAA

GCTGGCCAAGCCTAGCGAACGTGTTTGGTTTACGTCATTGTGC

>NODE_1570_length_183_cov_32.833333

GCTCAAGGCATTCGTCGGATTCGAGGCCGGGCAGAAGTGCTTCCCCAAGCCCGTTGTCTCCAACGACGCG

GGGACTAAGAGGGTTGCCTTTGTGCCGTCCGGAACGGAGTTCCTGGACGAGTGCATCGACGAGAGCGGCC

AGACTACCACCGGCCTCACTATCGAGGGAGAGCTCAACAAGGT

>NODE_1571_length_183_cov_7.475309

ATTCCACAGCGTAGACGCTTCCTTCTTGATATCGCAAGTACTCTCTCCTCGCCTTCCGCACAAGGGGTGC

CGTTTAGTGTGTCCGAAGAGGAACTACTGCAGGAGTTCCAGTTCTTGTTCTGAAATTCTTCGAGGGTTTC

TTCAGTCCTTGGAGCTGCTGCAGACGGCCGAGGGAGGGCACGG

>NODE_1572_length_183_cov_6.308642

AAACCAGCAGTCAAGGAGAAAGCATTAGACACGACACCTGACGAAGAATCCCATGGGCTCGGAAAATAAA

AACCGGCAATCCGAAGACGTCGGGCAGTTCCGGCCTTGACAAGTCCACGTATATACTTATATATGCGTAT

ACCCGTCGTTTTAGGAGCCTTGTTGTAGATAGGATTATGTGTA

>NODE_1573_length_183_cov_5.296296

GTCGTTAAAAGCGGCACGGAGGATACCACCAGCAAATTCACGCGCTCAGTCCAGCAAGGCGTTCATGGCC

CGTTCCTGCATGCACTGAGAGGTGAATCACAGCCCAACTAACATTTTCCCACACACTTATACTTCAATTC

AACACAGACGCCTCACGTTGCAGTACGTACTACTATTATTGTT

>NODE_1574_length_183_cov_4.740741

AGCCTACAGTAGGTACACCACGAAAAATGTTATCGTAAAGCGTCAACTTGTGTTTCCAACGATACTGCGA

TAGTTTCTTGCCTCTGTCGAGGGGCATGCTACAGTAATAAGTCAGTCAGAAACATTTTGTCCGAAATGTG

CTCTACCCCTGTCGTAGCACGCCTCGGGGATGTAACGCCCGTA

>NODE_1575_length_183_cov_4.703704

ACTAACCCGCCACGAAGGCGGATTCCACTGCGTCGTTTGGTTCTCCGTGTCCACATAATATTTTCTGCCG

TCCTCGTTTGTCCTCTGTTGCCAGCCCGGCGGCAAGCTCCCGCCATCAGTCGTTCCGTTTGTAGAGCTTT

GCACACCGCCGGCCCCCGAGTCTAGGGGGTCTTGCGCCACCCG

>NODE_1576_length_183_cov_4.697531

TGGGTGCTTGGATTTGCATTGGTTGCTTCTGGCAATTAGATGTAAACGAAGTACCTAGCTTAAGCTTTAA

TTATCTGTTTTCATCTTCGCTCTTCGGAGACTGGATGTTAATAGTTTAGCTTGGGTTCGGACCTTCGTTC

AAAAAGTTAGTGCGGTGAAGACCGCTCTGTTTATCTCCTCGGA

>NODE_1577_length_183_cov_3.623457

TCCTCGCTCCAGATCGGGGCAATTTTTTGTAATTTCTCTTCGCCAGCTTGTTCAACTTCAATTACGCTCA

ACTTCGCTCCAGCCCAGGCAACTCTCGGAACGCCCACCACTGCTGTGTACGATGCAAACGACAGCTACGC

TGAGACTGGGAATATCGTCGCTCGCCCGCGCTAGAGCCTGGAA

>NODE_1578_length_183_cov_3.203704

TCCCCCCTGCACCAGAACAGACATCAGGCGGAGGTGCGCAAGGCCAGAACAACGGCGGTGAGACTGTGTG

TTGTCTTGCGAAATGCATGCTTAGTACGAATGATGACTCGTGTGATGAGGAACTTAGTGAAGGCATGGGT

GAGAAATGGATAGCGGATAGTGGCGCATCCTTTCATATGACCC

>NODE_1579_length_183_cov_2.734568

GTTAGTATTGGCATTAGCGTTGGCGTTAGCGTTAGCACTCGCGTTGGCATTGGTGTTGGCGTTAGCGTTT

TAGTTGGAGTTGGAGTTGGAGTTAGCGTTGACGTTGGCGTTGACGTTTGCGAGCATTAGGGTTTGATTTA

GTGTTATAGCCTCGGCGTTGACGTTAGTATTGGCATTAGCGTT

>NODE_1580_length_183_cov_2.561728

AGTGGTTATGGTGGAAGGCTCGGTCGCGCTGTGTGACGGAGCCCTTCTGGTGGACACGGACTCTGCCTTG

CTGCGACACGAGCTGGCCTACATTCTAGGACAAATGCAGAACGAGGCGGCATGCGAAACTCTGGAGAAAG

TTCTTGGAAACACGTCTGACGACTGCATGGTCAGACACGAGGT

>NODE_1581_length_183_cov_2.432099

TATACCCTCATAAAACAAATTGAAAAAACAAATTGACGACTCCCTGACGTCCCTAGCTCGATGCCCGGTT

CTCTACAGCTCGTGCAGGGCCTCGCCCGTTAGCTATCTCCCCTGTTACTACGCTGCCTGACGACTCCCTT

CTTCCACCGACTCTACGGTTACTTGACTTGCATCTGATTTCCG

>NODE_1582_length_183_cov_2.172840

ACCAGAGCGGCAGCAGACGCAGCCAGGGCTGAGGGCATAATCGTGTTTGCTGTGGGAGTAGGCAGTGGAA

CTGGCGCTGGTGAACTTCTCGCGATCGGAGGGGACGATGCCAACGTCTTTCAGGTCGCAGAATTCGACCT

GCTCAACGCGGCCTTGGAAAGCATCCTCTCGTCAAGCGGAGGC

>NODE_1583_length_183_cov_1.962963

TCTACTGGGGACGTGGCCGTGTCTTTGGTGCCCCAACCCATGTTTGAACCGGACCATCCGTTCCCTCCGC

CAGTGCCGACCGATGCCGTCGCCTGTGGCTGTTCGTCCGCCCCTCTCACATCCGGTACGCCGAACGATAG

CTGCCCGAAGCTCAAGAACGACTCGTCGTTGGCGATGGTGTCG

>NODE_1584_length_183_cov_1.833333

TGGCGTTGAACGTCTGCATGTTCATGTTCGACCCCTCCAGCGCGATGATCTGGCTTTCCAGAGTCATCCG

CGCTCCGTTAAGCTTGTTGACCTCCCCCTCGTACATCTTCTTCCTCTTCAGGGCGAAGAGGGCGCCGCGC

TTGTCCTTTGCGCTGCTCTTCGCCTTGGCGTCCAGCAACTGCG

>NODE_1585_length_183_cov_1.833333

GGCACCGCTAACTCCTCCGAGGGCGAAGTCCACCAAAAAGTTGGACTTTGCCTTGGGTTCTCCCATGGTG

CTGAGGGTATGCGGATTCGGAGGAAAACGCTAACACTCAACGGTGTGGGGCTGCAGCGTTTTGACGGCTG

TCTGAGGCGCCTGGAGGAGGAGAGCGGTCGCGAGCGTGCAGAG

>NODE_1586_length_183_cov_1.759259

AACCGCATTGAGTGTTTGCTGGTTTTCAACGGCTCTAGAGTTATTCGTAGCTTAACGATACTCCTCGAGA

TGCGGATCATGACTGGTGACGTAACGATAATGCGATTCCTCTGTTTGCTGGCAGAAGCAATCCACCTGTC

GTCGTTATTTCGAAACTGACCAAAGCGTCATCTGAAAGAGCAG

>NODE_1587_length_183_cov_1.679012

GGTGCTAGCAGTCCAAATAAGGCCAAATAAAACTGCTCTATTAGAAAGGGTTAGTTTCATTTTTTTTCCG

ATTCTCTTTATTTACAGTTCGCTATATATAATTAATACATAGCGATTAGTAAATAGTTTTTAGTATACAA

TTACCGCACGGCGATTTTTAGAGAAAGCTTCTTCTGATCTATC

>NODE_1588_length_183_cov_1.561728

AGGGCGGGAAGCCCATGCCGAACACCGCACCGATGTCGCCGTCGACGGGCGACGAGATGATCCCCTCTTG

CAGACAAAGCACGGCTTCGTTCACGAACCTGCACATCATGCGGTCTTGGATGACGTCGGTCGGCAAGGAG

GAACCGTTTCCGTCCGCCTTGAGGTGCGCCTTGATCAGAGACA

>NODE_1589_length_183_cov_1.296296

ATCTATAGACAGGTTCACGCACGCACGCGGTTTTTCACCTGAGGATATGCGGAGTAGATCCAGAAGAAGT

GGTCTTAGATTTTTGCAGTCACAGGCGTTCGAGCAGTGCACTTGAAGCTACAGCCCCTTGTCTTTCCACC

TAAAGCTCTATCGGGCAGTACGTAAATCACAAAGGTCTGGACC

>NODE_1590_length_182_cov_9.273292

CTTGTACTTGGTATCATAAGAACAATAAGCACCAAAGGACCCCCAACTTTTTCCATGTCGAATTCATCAC

ATCTCCGACAGATGGTCTACAGATAGATCAGATAGATAATCGACCTCCACTTGACCGAGCTAGATCTTTC

AAAGCAGCAGCTACGAGATAGGTCCCTGATCTGAATTATCTA

>NODE_1591_length_182_cov_6.546584

ACTACTGTTGTAGAGGCTGCCCCAAATAACTAGATCGAGGCTTGAGCGTTTATAGGACCCCCTTTCACAC

ACTCTCTGCTGCTGGGCTGTGTCTTCCGACCGACATGGGGAGAAGGACGTTGCTGCTGGGCGCTACTGCT

GGTGCCGGCCTCTTCTTTGGCATCGTAGCTGGCTCGAGCAAC

>NODE_1592_length_182_cov_4.913043

GCTTGGACAGAGACGCAAGGAAATTCGAGCCACGGAAACGAGAGCCAAAGGAGGCAGCGAAGAACAGCAG

CGAGAACCCACCAGATTGCAACTATGGCTGAGGAAGGGAAAAAGGAAGGGGGGAGCAAGAAGCGGTCGCC

CAACCGCATGGTTGTGGATGAGGCCACCTCTGACGACAACTC

>NODE_1593_length_182_cov_3.546584

AAGTCAATTTTTATTTTATTTTATGTTTCAGTTCCTTTTTAGCCTGTAGCGCAAGGTAACAAATAGTCAC

TCCGTAGATCGGAGAACGAGGCACCTACTGCTCCATTTTTAGCTGCAGGCAACTTTCCTTCGCGTAGGTG

GTCAGAGTCTCACTACGTAGCCCCGATATATGTATGTAGCCC

>NODE_1594_length_182_cov_3.173913

AGGTCATGCAGGCCTACGATGACGAGACGGACAGCGAGGATGAGGACGGTTCCGCACCAAAAAAGATTGC

GAATGGAGAGGAGGAGGACGACGACAGCGAGAGCGAGGATGAGAGTGACGAGGACGAGGACTGATGCTCA

CGCCGGCAAAACCTAGTTTCACGTCAGCCAAAAGTTGTATTT

>NODE_1595_length_182_cov_3.167702

GCTAATTGGCACTCCTTCTGTGTCTCGCAAGGGATGCATCATTAGAGACGGTCGGCATGCGAGGATTTTG

ACACATTTCCGAGTCATGTTTAAGGGAGAAAGGGGGTGGCGATAGAGTTTGTGACTCACGGCAATATTTT

TTTATCGACTCCCATTAACTGGCTAGAGCACGTTTCGCTCTC

>NODE_1596_length_182_cov_2.956522

CCCCGAGGCAGCATGAAACACGGTGGTAGTGACTGCCGAATATAAGTATACTAACTGTTCAAATTTACTT

CTAAATGTGGTGTGCATTGGCTGGCTAATACATGATGCTTGTTGTCCATATGTACTACTGTACTACTTAC

CACTATACTGTGGTTGTACTCGCCTTGTTTCCTTGCAATTGT

>NODE_1597_length_182_cov_2.931677

CGTGCTGCAACTTACCGCGTGTATCGGTGAAACGTATGTCGACGTATTCGATGTCCTCGTCTTTGATCCG

TTTGATCAGGTCTGCATTAGCGCTCATGGCGTTGTCCTTTTACTTTTTCGAATGGTCACGGAAAGGGTTA

CAGCGCGTCCGAGCCGGTTTCACCGGTGCTGATGCGGATTGT

>NODE_1598_length_182_cov_2.807453

GTGCATTAGCCTGGCGCCGTCGTCGGCGAACAGTGGGGTCCACGAGTGGCTCGAGCCCATGCACCGAATT

CTAGGCGGGGGGTCCACCCCCCTGGCCTTTTTAAGCACTTGCAGTACATCTTCCACAGACTTCGGCCGCA

GCACCTTGGCAGGCGCTGTCTGAACCGTAGCGGCCCAGTTTC

>NODE_1599_length_182_cov_2.770186

GGTACGCGCGAGTATCGAGACTATATCCAAGAGTAGTGAGCCAGCTGTTTCCACTCGGTACACAAGTAGA

GCATCAGACAGTACCCACGGAGCTATATAGCCTACAGTAGTTACCCAATCACTCCCATTGGGTACAGGTA

CATCAAGTCGAGTATATCAGAATATCAAGACCATCAAGGAAG

>NODE_1600_length_182_cov_2.726708

AAACAACTCGGCAAGATGCACGCTCGACGGCTACATGTACCGTACGGGGCATGGCACTGACAGACCCTCA

GCAGTACTCCTTGACGTTTGCTTGTTGACATTGATGTTTGTCCGTACGAGGCACTGATACTTGCTCGTGG

TAACGTAGTTACGTTGTACCGTTGTCATCACCCCGACGCTAT

>NODE_1601_length_182_cov_2.422360

AGAGGTTTTCCCAATGAGAAAAGAAATTCACCCAGAATACAAATTAGTTACTGCTACTTGTTCATGTGGT

AACGTTATTAAAGTTGGTTCTACGCTTGGTAAAGAAGAGTTAACTTTAGATATCTGTTCAGCTTGTCACC

CATTCTACACTGGTACTCAACGTGTTGTTGATACTGGCGGTC

>NODE_1602_length_182_cov_2.409938

ATGTCGCGCACATAAAATGTGAACGACGGTTGTCTGCCCCAGCACTCAGCACACCCTCTCCCTTGGCCGT

TGTGCCAACACTGGTAACAGAAAGAGCAGTCAGTGTTGCAGAGCCACGTCCGTCCTAAATTTCAAGGCAA

GGGCCTATTAATATGATACCAGGTACCCGGCCCACACCGAAA

>NODE_1603_length_182_cov_2.285714

GCCCCTCTCCGCCACGTTGTTGACGAACACCGTCGGGACGATCACGATCGACTTGCTACCCTTCTCCTGC

AGTTCCTTCTCTAGGATGGTGTTTTCCTGGTCTTCCTCAGTTCCCCCTGCCTCCGACATGCACTTGTTCA

CCGCATTCCTATCCACTCCTGCCGCCCTCATGGCCCCGTCCA

>NODE_1604_length_182_cov_2.279503

AGGGATTGATGTACCGGCAGTTGTCCACTTCGGGGATTACTGTATAATACGTATTGTAACGTGACAGTTT

AGTAATAGAAGTGTGGCGTACAACAGGCTATAGTTCAAAGCGTTCGGAGAAAAAGAGAAACCCCCGAGAG

AGAGAAAAAATGGTTCGTTATCTGGGCGTATACCCGTAAAGT

>NODE_1605_length_182_cov_2.204969

ACCACCGAGGAAGACACGCCTGTTATCGTTGACCTTTTGGGCAACGACAGCGATCCGGACAACACGCTGG

ACGAGCTGACGATTTCCGACGTAACCGTGGACCCTGCCGAGGGCACGGTTGTGGACAATGGCGACGGCAC

AGTAACCTTCACGCCAGCCGAAGACTTCACCGGCGAAGCGTG

>NODE_1606_length_182_cov_1.850932

TAGCGGGTCTTTATCTTGTTACCTCGACCGATCTCCTCTATTCTATCGCGGCGGATTGGATCTCTCTGGT

GACGGGCGGGGTTGGAACGATCTGTATCGTGAACGATCTGTATCGTGACCGATCTGTATCGTAACCGCTC

TCTGGTCGTAATATAGCCGATCTGCGTTGGGGTTGGAACGAT

>NODE_1607_length_182_cov_1.770186

TCAGTCAGTCCATGTCGCCGTGGGGTTGAGATTCTTAAACAAGCGCACCCTTACGCCTCGCGTCTTTTGG

AGAAACTTGGGGAAGGTCGGGGATGGGGACGAGGACGGGACTGTCGAGTATTCGAAAGCCACATTTTCGC

GTACGCGTTGCAGAGAGGGGGCCGCATTCGTCACGGGCTGTG

>NODE_1608_length_182_cov_1.757764

GAGCAGAGCAGCGTCGGCGTGCGTCAGGAGGTATGGCCCGGAGCTCTTCGATAGGCCGGGAGCACCGTTT

CGACTCAAGACCCGCCTGCTTCAAGCGGAGGCGATGGAGGCACTGCTGTACGGTTGCATGACATGGGCCC

CACGTCGCGACCACTACCGGCTGCTGCGGACGATACACCACC

>NODE_1609_length_182_cov_1.708075

GGCTTGTAAGAGGCGTCTACCTTCTAGCTGCATGGTTCCGATGCGGGCGGCCTATCGCAAGTAAAATCCA

ACAGAGGACCCGAAAAGGAAGAAGAGGTTCAGTACCGGCCTCATGGTGATGATGCGAGCGGAACAGTCGG

TGATGAGAAGGTGTAGACTGACACAACGCTTAGAAAAGGAGT

>NODE_1610_length_182_cov_1.422360

TCGAAGGCTCGAGAGAGGTGGCCACCTGCACTTGCTCTGCATTATTTCGCAATAAAAATACGGACGAGAG

CCGGCGGCGAGACCGGTTTTTTGGAGGCACGAAGTCATCAAAAATGAGGGTCGCGGCGAGATCGCTGGCT

GCAGCTTTGGGATGGAGACTAGCAGCTTCTGAGAGCATGGCC

>NODE_1611_length_182_cov_1.422360

CGGCAGGGTGCGGCTTGAGCGAGGCTAGCTACTTACTCAAGAGAGCAAGGATGTTGATGATCGAAGCGCA

TGCATCGAAGCCATCGCGGCAGGCTGATGTTAGGGTTTTTTTTTGACGCGTAGAACAATGAGCCGTCCCT

CACGCGGTTACTGCTGTAGTTTGCTGGTATCGGATCGATCTT

>NODE_1612_length_181_cov_2362.281250

GCACCAGGACAGAAAGACCCTATGAAGCTTTACTGTAACTTGAAATTGGTTTCGGGTTTTATTTGCGCAG

CATAGGTGGGAGGCTTTGACGTTACTCTTACGGGAGTAATTGAGCCATCAGTGAGATACCACTCTAATAA

AACTAGAATTCTAACCCTGTATCTGGAATTCTCGGGGTGCC

>NODE_1613_length_181_cov_1365.100000

AAAACTTTCTGCCATCTTATATCGTTTGTTTGAGCGCTCGATTCCCCACGCACCTCATTCCAGAAACGAT

GGCAACATAAAATTTCACAATACGGTCGCTTAAAATTGCGCTGCCCGCTTTTGCGGTTTTGCCCTAAAGT

TGACCCCCATGGCGCAGCGCGTTAGCCGCGAACCGACATTC

>NODE_1614_length_181_cov_613.075000

GTTATATGGTCAAGCCTCACGGGCAATTAGTACAGGTTAGCTTCACACATTACTGCGCTTCCACACCCTG

CCTATCAACCTTGTGGTCTTCAAGGGCCCTTCAGGAATCTCAAGGATTCAGTGAGAACTCATCTTGGGAG

GGGCTTCCCGCTTAGATGCTTTCAGCGGTTATCCTGTCCGA

>NODE_1615_length_181_cov_282.200000

CGGCTACCTTGTTTCGACTTCACCCCAGTCGCTGATCCTACCGTGGTCCGCTGCCTCCTCGAAAGGTTGG

CGCACGGCCGTCGGGTAGAACCAACTCCCATGGTGTGACGGGCGGTGTGTACAAGGCCCGGGAACGTATT

CACCGCGTCATGCTGTTACGCGATTACTAGCGATTCCGACT

>NODE_1616_length_181_cov_18.762500

TAAAAGGGTGAGCATTATTAATGCTCACCTTTTTTTGTTGTCTTCATTCCATTCCCTTGAAATTAACCGT

ATCCTTGTGATTCCATGACAATGCTTTCCTAAGCAAATCCCCTTGCCAATCCTTTGGCGGTTCCTTATCC

TTGTAAACAGTTCCACTGCATCAATGCGTCTTCCTCTACTT

>NODE_1617_length_181_cov_10.256250

CTTTTTTTTGTTTTTATGTTAATATTTTCTTTTGTAGCAGCACGCTGTGGCGCCGACGGTTGTCCAAGCG

AAAAGCTAGGAAGAAGAAAAGAACATCGTCAACTCGGAACCGACGAAAGCAGAGTGGCCCAGATGCCCCG

ACTTCCGCGGGCGAAAAAGGCTAAACCGCGAGCAGCGCGTG

>NODE_1618_length_181_cov_7.356250

GCAATCCGCTACTATTACTGGTACTCCATATTACTGGTTGTATTTCAACACTTGTAACCCAATCCCGACA

AATCTGCCTTGCATCATAACGAAATCCCAAACAACGTGAAATTTTAACACCCATACGAACTAAACAACTG

CATGCAATATTGTTACCCAAAATCCCCTGCGTTAATTCAAT

>NODE_1619_length_181_cov_4.425000

ACCCTAGGCCGAATGAGACGTTGTCCGAGCCCACCGAAGAGAGGTCGGTGTCGTAATAGCAATCCGTCTC

AAACCGGTCCCCTTTCTGAATAGTCACGCTGTCGTTGTTGGTTATGATGTGGAAGCCCGCCTGCAAGAAG

GAGTAGTACTCGACCTCGGCAGAGTGGACCAGGACCTCCTC

>NODE_1620_length_181_cov_4.006250

GCCTCCATAAAAGTCTTGCCAGGTACTGGTACTACTTCTTAACGTTACTACCCCGGTATTTTTTGTTTTA

GTTGGTCGTCCACTCGACTTTTCTTTACTCTTCTTCTTTTATTTGCTGTGGCTAGTCCGCCTCCTCATTT

TACATGGGTAACGCGGGTCAGCAGCACCGGTTATAGTTATA

>NODE_1621_length_181_cov_3.418750

TCTCCAGGTCGAGCTGCCCGTCGTCGGGCAGCCCCTCGACGCCCGTCAAGTCCCAGCTGTTCGTGTGCTT

GGCGTCTTTGCTGACGCTGTGGTAGTCCGCCAGTATATCCGAGAAGAAGGGTCTAAACCTGTCGTAATCC

GTCGGCTGCATGGCGTACACGCCCACCTCGGAGTCTGGGTT

>NODE_1622_length_181_cov_3.162500

ATCCCTTTCCCCTCCCTGCCCGCCTCGCAGGCCAAGTAAACCCCGAAACGGCTTCGTGTTCACTTGTCGG

AGCCCGACCGACCGACTGACTCTCGACCTTCGTCCGAACATCCGACAAAAATAAAATAAACAGCACCGGA

CAGAAACTATTCCACCGCCACTGCCGCCACACACCGCCGCT

>NODE_1623_length_181_cov_2.925000

GACCGTAGCAAAGCTTTCAGAAAAGCGGGCGAAGAGAATTCCAACCGGGAAGCACACGCTCCAGGCAATG

ACCATTAACCAGGCGTGCGCGTGCTGCTTGGCGTTGCCGGAGTTCACCGCATTTCGGACGTCGCAGGAGT

CCAGACTTAACGCGAAAGCGCCTTCGACGGTGTGTTGCACG

>NODE_1624_length_181_cov_2.618750

TTTTTTTTGAAACCACGACAAGCGCACAGAGGTGGCTCCGAGCAAGGACCGCAAAAGTAGTAACGCGCGG

TCAAAAACCGACCGAACAAGGAACAAACACACACAAGCCGACAAAGAAAGGCGATCGCGCATGAGATCTG

CCATCATCATCGTGTACCAGGTAATGTGTTAATGCGCCGAA

>NODE_1625_length_181_cov_2.543750

AGTGTACTGAACTTCCTTCATACAAACACTTGCACGTTCAAATATATGTTTTGTGGAGGCTTTTTTAGCG

GCACGGTACAGATTTTGCTTTTGCGTACGAGGGTGCCGCCGTTTCTGTTCTCTTGGAATTCTTCTTTTGT

GTCTTGGAGACACTCGGACTTTCCGATGTTTTTGCTCCCGA

>NODE_1626_length_181_cov_2.293750

GCTGCGCTGCTAACTCAATGCTTAATGTTGATACTTTATTTAGCGTTACTTTACTTGAATCTGAAAAAGT

AAATGTACGTGTATTTACTACACAAATTAATATACCTCGTTCACTCTACCGCCCTCCTATGTCTGCTTAA

TAGCAGGATAGGTGTATGTAAAAATTGCGGTAACCCCGTTT

>NODE_1627_length_181_cov_2.168750

TACACTGGGTTTGCACGTTAAGTTTGCCACTACTATTACATAGTCCGAAATAAACATACCTGGTAGTACT

CTTTTTAAATATACAGTTATTCGTATTGTGTTCTTGGTCATAGATACTGACAATTTCCCCGCCGTGGAGG

TTATACAGGGGTTTCGTGGCTCCTCATACCATACCAGGTAT

>NODE_1628_length_181_cov_2.118750

TCCACTTCTCGTGGAACTCCCCCTTGGCGGACTTGTAGATGCCGTACGCCAGCGCGTCGGGCTGAGCGAA

GAAGTTGCTCATCAGCTCGTCGTGGTTCGCCACCGGCTCTCCCTCCAGGAGCACCGGGCACTGCGACTCG

CAGTAGCCCACGAAGTCGGCCGGGATGACGCGGCCCTGGTG

>NODE_1629_length_181_cov_2.031250

GAGAAGGGGAACATGTCTGATAAGGCAGTTACGGAGGCTATTCAAAAGGATTGGAAGGAGCGAGAGATGA

TCGAGATTGTCCATCTCAATATCCTCAAGATCACCAAGTTTTTGAACGAGTTTGAGGTGTCGGTGCGGTA

CAAGCTGTCAACCCTAGCGGAGCGCGTTGGGTCCTTGGTGG

>NODE_1630_length_181_cov_1.631250

CTCGTCCACATCCGTGAGCTTGAGGTGGTTGGGGCAGTCCTCGCTCGCGACGCCGCCCTCCTTGTACACG

GCAGCGTTCCCCATCCTCAACGCGAGGTCCTTGACAGCCACGGCTGTCTCAAGGTCTACGCCACGACCGA

CCACAAACGCCGTGGGCCTGTCTAGCAACCCCGCGACGGCC

>NODE_1631_length_181_cov_1.587500

GAGGTTTCTGCCTGCCCAAGCGACTTCGCGGTGACCTTCCCTTCCAGCTTCATGATCCTTCCCCGCACCA

GGATGATGTCTTCGTCAGGCGCCATGCTGATGTCCTCTTCCCCCGCCTCTCCTGTCTTGCCCACCTCGAA

GGTAGCTCGGTCCATGAGGCTGTGGCAGCTCTGCAGCTTGT

>NODE_1632_length_181_cov_1.525000

GTACAAAGACTACAGATGCAATGGGGGAGATGCCTTGCCCGGGTGCCGGGACAGCGGCGATTACGCGGAC

TGCCGCACGTGCTACATCAACAAGGACGCTTACGTGGAGGAGACTGGAATTGACGATGGTTGGCCGGAGT

GTCCGTGCTGCGTCGCGGAAGGGCTTGATTTGCCCCCTCAG

>NODE_1633_length_181_cov_1.487500

CAGGCATGCCCGCAATTGGGCTGTTAGGATCATCATTAGCCGCAGGATTCACAGTATCATTAGCACCAAT

AACCAACACCACATCAGTGTTAGAAAAATCATCATTTATTTCATCCATTCCTAGAACTATATCGTATGGA

ACTTTAGCTTCGGCGAGTAATACATTCATGTGCCCAGGTAA

>NODE_1634_length_181_cov_1.325000

CGGGTCAGAAAGCGAGAAAGAAAAGAGGCCTCCGGACCGCTCTTACAGCCAAGCCTGTCTATGGCGTTGA

GGGTAGCGACTCTTACGCCAGCCCCAGTGTGGCGTTGTGATTGGTACACAGAGCAAGAAGCGAGATGAGT

GGGTCTACGGCCCGCACTCGTAGTCAAGCCTTCGTATCTTG

>NODE_1635_length_181_cov_1.125000

GAGATATTATAGGTTCTGGAGAAGAGAGATGGGGAGGGGTGCTCGCACTGCTGTCGCTTTACGGGACACG

TACATAGACTGTCGAGTTTTCGTTGTTGTTTTTGGTTTTGGTTTTGGTTTTCGCCGTCGTATTCGTTTTT

GTTTTCGTAGGCTATGGGATCACGCATCGGGGGGCGCTCTT

>NODE_1636_length_180_cov_224035.779874

CTCTCGAATCAAAGCGCACCCCACATTTCAACCCATTAAACTCTGAATCTGAACTCAAAGGGGCGCTGCG

CTAGCCGCGGCTCCCCCAACCTTTAACGTTGTAAAACTTTCAGCGACGGATGTCTTGGCTCCCACAACGA

TGAAGAACGCAGCGAAATGCGATTGGAATTCTCGGTGCCA

>NODE_1637_length_180_cov_16.106918

TTTGCTCAGATTAGCACGGCCGTTTCATCGTCAAAGGCAAATCGTCAAGTCGAGCAACCAACACCTCTCC

GCCTTTAGTTAAAAACAGCGAGACGAAAGCGCTCGAAACTAGACGCAGCCTGAACGGCTCGCCTCCGATA

TATTTTTTGAGAAGTTTTTAAGGCTGTCGTCATAGCAATT

>NODE_1638_length_180_cov_6.289308

CCTCTAAGCCTGCCCCCAAGGCATTCTCCGCCGCTTCCAAGGCAGGGCCGAAGGGTCCCCCGAAGTTGGA

GTTCGTGCAGCAGGGCATGAAGTGGATCGTCGAGAACCAGGGGTCGGACAGCGGAGTGGTGACGGTCGAG

GTCATCGACAAGAAGCACCAGGTGTACATCTACGGGTGCA

>NODE_1639_length_180_cov_4.100629

TGTTTTATTCGCCTCGCCGGAAGAAATGAAGAGCCGTGGGCGAGGTTGTCCTAGGTCACGAGAGGTTATA

ACACGTTCGTCCGTGGCGATGGTGGTGGTAGTTGTGATGGTTGTGGTGATGTTGGTGACGTGGAAGGGTT

GCACGAGGCCACGATATGTGTGTACACATCGTTGTGAGTT

>NODE_1640_length_180_cov_3.081761

CTGACATGTACGCGGCCATCGATCTCCTGACAGACAAACTGGACAGACAACTGATCCGTCACAAGGAAAA

ACTCAAGTCTCACCGACGTACAGGTGACACTGCGATCTTCGAATAATCATCTGACGCTTTTAATCCGACT

TGCGAGAGCGCACAATAGCGCTTTCGCAACGACCTGGAAT

>NODE_1641_length_180_cov_2.861635

ACCACCGGAAATCTCAAAATCATTACATGGTTTCCGAATCCAGACACGAATGCCTACACCCTACTCTTAC

CTTACCCCATCAATCCTCCACCTGTAACTCAACCGAAGTTCAATCACAGCCCTCACTAATATCGCAGCCT

CGATTGCGGCTTAGAGGCCTTCAGCCGTTATCGAACACGG

>NODE_1642_length_180_cov_2.849057

TGAATCGTGTGCCAAATAAAAATGTATCCCATCTTCCATCCGTCCATGCTGGCCAAACCTTCCGGCCGAG

TACAACAACTTACAACTCTCCCCCGTACCCATACAAATCCTACAATAAAAAATACAGCAGGACTACAATC

CCTATAATACCATCGTACTGTATATCGCCACTCACTCGTA

>NODE_1643_length_180_cov_2.817610

CTCTTTTGTGATTTTTTTTTTCTTGTCAACGTAGGACCACACGTCGACCCTCCTACCCTTATTTCTACCT

TACTTGTACATGGTTATATACGTACGTATTGTATGTATGTATGCATTTTGTTTCCAGTTCAGAGTGAACA

AAAAAAAAGCAAGCATGGCGTCGTTCGTCGTTGTCGTGAA

>NODE_1644_length_180_cov_2.811321

ATCCAAACAATATTGATGTTCCCAGCCCGAATCCAATTAGCGCCTAAAAGGTCCAGTACATGGTATAACC

TGAACGTTCCTTCTCTTCTTCCGCGTGCTCCGACCTCGCTGTACCATGCACCGTCACCAGAAGTTGATTC

CGCTTCAGAAATTTATTCTATCACCAGAAATGCATTCCGT

>NODE_1645_length_180_cov_2.415094

TTTTTTTCCCTGTCCCCTAAGCCAATAACGCCTACATATCTATATGGAAACCCCGTTATGTATGGACATG

GACACACAGCACACAGCACTCTCTCGTACACTGGCGACGGTGACTGTTCACCCTAGCCAAGGCCACAGTC

GTTGAGCAAGATCGTTTATATTTACGCAGGGATGTGTTTA

>NODE_1646_length_180_cov_1.823899

TCGTCGTGGGAGAAGGCCAGATCCTGTCTCAGGTCCGGCAGAGCTATCTCCGAGGAACGGAGGAGGAAGG

CTCGGCGGGGAAAGTGGTGGCGCGATTGTTGAACACGGCGGTGTCTGCTGGAAAGCGAGCTAGGTCGGAG

ACGAGCATTAGCAAGGGCGCGGTTTCAATTTCATCTGCAG

>NODE_1647_length_180_cov_1.698113

GTTCGAAACCATGTACGACGAAAATGGTGTTGGTTTAGCTGCAATTGGCCCTGGAATTGGTCAAGGTACT

GCCGCTGGTCAAGCTGTTGAAGGTATTGCTCGTCAACCAGAAGCAGAAAATAAAATCCGTGGTACTTTAC

TTTTAAGTTTCGCCTTCATGGAAGCTTTAACAATTTATGG

>NODE_1648_length_180_cov_1.471698

GAGGCAATCTCCGTGTCGATGTAGAGATCGAAGGTGCTGTCCGTCTTGACGTGCAGTGTGTACACGTGGA

AACCGTTGTCGTCCTTAATCTTTGGCGCGTCCTTGTAGTGCTTCTCTTCCCACTCGTTGGTGAGCGGGTT

CTCGTGCTGGAAGATGAAGTGCACCTTGTTGGTATCGCGG

>NODE_1649_length_180_cov_1.415094

CTTTGTTTTTTACTTTTACGACTTACCACCATCACCCATAACTTTTTGTTGCTGGCTCTTCTTTCTCACA

TGTATTTGTATTTGCTCTTGTCTGCTTCATAATCTTCTCGTTCATATTGTTCTTGCTGCTTCTTCGCGTG

CTTCCTCTGGTTGTTGTGGTGCTTCTTTTTTTGTTGATCT

>NODE_1650_length_180_cov_1.163522

AAGTAGGTTTTTGTGTGTAGCGTAGGTCCTATGTACTATGGTCCCCCATAATAACTGACCTCATCAGAGT

GCTTGTTTCGAGTCAAGCGTAATAAGACTACAGTAAAGTAGGTGAGACACAGAGTATCCCCTTGCAGGTT

TGTCTATTAAGCATCCTTGGTGTTACTCACTTTCCGCGTA

>NODE_1651_length_179_cov_60779.056962

TGGTAGAGCGTCTGCCTTACAAGCAGAGGGTCACTGGTTCAAGTCCAGTACTGCCCAATTCTTACTTCCT

TACCTTAATGGGCTCATCGTCTAATGGATAAAGACCGGAACCTTCTAAGTTTCTAATGTAGGTTCAATTC

CTTCTGGGCTGGAATTCTGGGTGCCAAGGAACTCCAGTC

>NODE_1652_length_179_cov_7273.139241

TTGTGCTGTGATGGAGAACATGTCCAACACAACGTCTGAGCTTTCCTCTTGGGCGAAAAAAACCAAGGGG

CGGGAGGCAACGGTCGAAGAGGCAAAATGCCGACTCGCCAGCCTCCCGAGCCTGATGACTTGGACCGGGT

GAACAATCTCTGAGCATATGGAATTCTCTGGTGACAAGG

>NODE_1653_length_179_cov_1533.968354

CTACGCATTTCACCGCTACACCTGAAATTCTACCACCCTCTACAAAACTCTAGTTAACCAGTTTCAAATG

CAGTTCCAAGGTTGAGCCCTGGGATTTCACATCTGACTTAATTAACCGCCTACGCGCGCTTTACGCCCAG

TAATTCCGATTAACGCTCGCACCCTCCGTATTACCGCGG

>NODE_1654_length_179_cov_726.556962

ATACTTACCTGGCGGGGGCTTGCCAGTGGTCAACATGACTGGTGACCCCAGGGGGAGCAGGCTCTTGCAC

TGGGCTAGACCCTGGAGTGCTGTAATCCCTGGAGAATATTCCTTCGGGGGTATCTCTACCGCATAATTTT

TTTCCGGGGAGGTGTCGCTTGCGGCCCTCCCAATGGAAT

>NODE_1655_length_179_cov_30.550633

ATTTTGGTGTGTATTTTTTTATTTTTTTCAAGTTTCATTTTTTTTTTGACGTGCCTCTGGTGCGTGCAAC

AGCCCCACCCTCATTCCGTGGCTAGCGGGAGAGAGAAGCCTCCACGACCCCCAAGAAGGAAGAAAAAAAC

GCCGCAAAAACCACAAAAACCAACATAAAGAATTACGTC

>NODE_1656_length_179_cov_15.772152

AAAATTTTGCAACTTTTTCCTCTCTATTGCTAAACTGGCTCTTCTCTCTGTCCTAGTCCGGTTCGCGCCT

TGGTTTCCATTGACAGGGTGCAGCCATCCCAGCCAACGCCGACACTCCCGTCCTTCCTGTCGCGTATGCG

TACCCACGCGATGGCCAGCCTTGGAATTCTCGGGTTGCC

>NODE_1657_length_179_cov_6.734177

CGTACTGCAGGCAAAAAGCCCAGCCCTTTCGACTCGAAGGGCCCACTGCTGGTCACATCTCAGCCGCTAA

CAAATCGTCGCCCAAGCATGAAAACCCAAAACCACACCAACAACCACAGCTGTCAAAAGACCAACACAGG

GTGTTGCGTTGCAGAAGGAACGCATGCCCGGGGGGGGGG

>NODE_1658_length_179_cov_4.531646

AAGTATTTCGCGCTCTTTTTTGGTTTCCCCTTGGGTTTGTGGAAGAGAGGGAGGCGTTTGTTTTGGGTCT

AACCATGGCCCACTCTCTGGCCGGTGCCTACGGTCTTGAGGTGGACATGGAAGGCGTGTTGGTGGGTTGC

TATTGCTACTCGCGGGTGTCGCTTTGCCCTCTTTAAGAG

>NODE_1659_length_179_cov_3.689873

CTTGTTGATTTGCTACACGGAATAAACTTGAATGAGCACCACAAAATCCTTTTGCAAGTTTTAAGATTTT

GTTTCTACGGTTCCGAGCGACATTACCTCGCTTAACTCTAACCATTAATTTTCTTATCGTTATAGACTAA

CAAACTAACATTTTTCTTATTGCTTTGGTATCTGATTTA

>NODE_1660_length_179_cov_3.544304

GCTCTACATCGTTTTCGACAAACACGAGCTCGTTGAAGGGGACGCGCTGACATTGTCACAGAGTGATAAT

AAATTATTTTCCTTCACCGTCGACGGGAAGCAGGACTTCAGAGATTCCCCCACGGGCTCTCCGACGGACT

TAGGGACCAAGGAAGCTCCCCTGGAATTCTCTTGTAAGC

>NODE_1661_length_179_cov_3.240506

CGTCTTCACGCGGAGGATCGCAGGCTCCAGCAGCTCGATGGGGCAACCGTTTACCTTGATCAGCCCGCGT

CCGCGCTTGCAGTAAGCCACGGCCGTGGCGGTCTTCTTGCGACCGAACACTTGTACGCTCTCCAGAACGG

TCTCGTTACCCATGGCTGTAAGGGAGGTTGTCCTGAGAC

>NODE_1662_length_179_cov_3.183544

CCATGTGACTAAATTTTATTTAAGATCCATCTCTTGGATGATCTCGTGTGCGATTTCAGGTGTAGTGCTA

ACAGGCTTTTCGTGTAAGTTTGCTTTTATCTGCCCTTTTCGATGTTTTAGTGCCGCATCGAGTTTTTTTA

GCTGCGCTTGCAATGGCTGGGTACATAACTTCGTTTTCC

>NODE_1663_length_179_cov_2.924051

ATATGACCTTAGTTGGTGAATTAATCGCATTTATCGTATTTGTAATTTTTTGTATGAAGTTTGTATGGCC

ACCGCTTATTGGTGCCATAGAAGCACGTCAAGCGACTATCGCTGACGGACTTGCTGCTTCAGACCGTGCT

GCTAAAGAACTTGAGCTTGCTCAAGAGAAAGCAACAGCA

>NODE_1664_length_179_cov_2.854430

TTGGCTGCACCACGGGCTTCTTCACGGTACCCTGGGCCCCCTGACCTCGATCAGGACCGAGGGAAACAAG

TGGATGGGGCTGGGATCGCCGACATCTCGGGACTCTCTTGTGGCGGAATCGCGCAACATGTCGGCTCTTG

GCGGCGCGGACCACGGTCGGGCCGACCGATGGAGTGAGC

>NODE_1665_length_179_cov_2.601266

CAGGCGAAGGTTGGGGAGCTTGGGGGGGGTCAGATGATCCCCTTCAGCGTGGAGTTCGAGCAGGAGTGGA

TGGACGAGGACATCGGAGGAACGCTGGAAGCCTACAAGGCAGCCAACCCCACCCACAAGAGCGTCATGGT

GCGCATTCTAAAGGCTGGCTACCACGCGCTCCAGCTGAT

>NODE_1666_length_179_cov_2.594937

GCCCTCGGTAAAGCTGCTTGTAGAACGGCCTGGAAGTGCACCATATTTTGTTGTGGTAGAACGGGTGCAG

AGTCGGAGCCATCCACATGAAACTCCAGCTGGCGCTTTCGCAGTGGTCAGTCCAAGTGGTCATCTCGCTG

TTGAAGTCGATTTTGTACTTGATCCCCCAGAGAATCCTT

>NODE_1667_length_179_cov_2.506329

TACACGGACGCGCACGCTCAGTATGTCAGCAATCAAACCACCTCTCGAGTGTCACATCCAAAAGTTGAAA

GTTGTTACGTTACTACAACCCTGCCGAATCTCCACCAAAACTCTATGCGATAAATACCGATAGCAAGAAA

AATATCGAGATGGGCTGACCAGCCAGCCCCCACCATCTC

>NODE_1668_length_179_cov_2.417722

GTTTTGTTTTTTCCATTTCCATTCCCCGAATCCTTACTTGATGCTGCCGTCGTCGTTTGCCACAATGCCA

TTCGCGTTGACCATCCCCTCTAAGGTGCGGAAGTTCTCAGTAAGCGTGATGGTGTTGGCGGCGGTGATCA

CCACATCCACCAGCTCCCCACTATCCGCGTCGAGCGGGA

>NODE_1669_length_179_cov_2.310127

AGGGGAAAGGGATCAGGGGAGAGAGACGGGTGGAAAGGAAGAAAGCACAGGAAAGGGAAGGCAGGGCAAG

AAAGGGGAGGTGGCTTGGAGGATTTGCGGTAGACAGCCGCGGGAAATAATACATCGTCTCCCCCGCTCTC

GGCGGAGACGACTTTTTGATTTCACGCTCGTGACGAGGA

>NODE_1670_length_179_cov_2.291139

GCAGAGCGGTGACGAGGATGGCAGGGTCCACGCGCAGCACCAAGTCTATCAAGGATCCGAGAGAAAATCC

CTGCAAGAGCCCGAAGAGAAGCAGAATGCCGGTCCGCTTGGAGAAGTTCTCCTTGTCTTGGTCAAACGCG

AGCCACATCATTGCTCCGAAGCCTGCCAACATCGTGGCG

>NODE_1671_length_179_cov_2.246835

CTCTAGGCCAAAAAAAAAAGGTAGTTATGGGTAATGTACGATTATTTTGGAGTTTTTTATCTACAGCTAA

TCCTAAATAGTTTTTGCTCCAAAGAATCTTAAAGACATAGACACTCTTATTTAACATCAATAAATTACCT

ATATTTTATATTATTTTTCTTTAAAACCCAGAAGACTTT

>NODE_1672_length_179_cov_2.177215

GTTGAATACGGCTTACATGCTTAAATATGACTCAACGCACGGTCGTTTCAACGGTACTGTTGAAGTTAAA

GACGGTAACCTAATCGTAAACGGTAAAACTGTACGTGTAACTGCTGAACGTGATCCAGCTGACCTTAAAT

GGGATGCTGTAAATGTAGACGTAGTAGCTGAAGCAACTG

>NODE_1673_length_179_cov_2.069620

GTGACACCCGCTCAAGTTGTTTAATGTTTCAAGGACATACAATCACTTGATCTGCCGTGATGGCCACAGG

TCTAGCTTAATTTCACAACCTGCTCATACTCTGGCTTACGATATTCAAAGGAGTAGACCATGACACTGAC

ACGCCGCCAATTTACAGGGCTCTCGATCGCCGCCACTGC

>NODE_1674_length_179_cov_1.993671

AAGGCGCTTTGTCTGCGTTGCTGCTGCAGCATGTCCGAAAATATAATGGTCGTTTATTACTGTAAAAGAA

CATTCAAGCGGTGCTTATTTATCGGAACAAATATTATTTACAGAGAGCCCGGCGGTTTGAACACCATCAT

GATGGAGCGGGCGAACGAGGGCCACGGTGAGCACCGGGC

>NODE_1675_length_179_cov_1.949367

ACACAGTATAGGTATAAAGGATAAAACGGTCACAGCAATTGGAGCGTTATTCCATCTACATCCACTGCCA

TGCCAAGAACAGGCCAACACATACATCAGAGAGATTACCCAATAGCCAAGACCCCATCCAACCAGCACAG

AAGGTACACGAGAATCAGCACAGCTGTTCCACCACTCTT

>NODE_1676_length_179_cov_1.949367

TATTTTCTCCAGGCTTTCATCCTGTTAGCATTGAATTGTTGGATTTTGTCTTCGTCAGTCTTCTTTTTGG

CCCGACCGGTATTCGTTAGCGAGCGATTTACCCTTTCTTCGCTCCTACATCGTCGCCACTCCTATTCTTC

TTTGCCGGTGTCCCTCGTACCTCTTCCTGTTCCCTTTTT

>NODE_1677_length_179_cov_1.772152

GGGCAGTAATGTAAAGCCATCGTGTAGTAGTCCACCGCGTCTTCGAAGTCTCCGTTCTTGTAGTGCTCGT

TCCCCGAGGCCTTCATCTCGCGAGCCTTGCCAGCGTCCTTGACCAGGTCTGTAGCGCTGGCGTAGGCATC

GTGGAAGTCGTCTTCGCCATCGCTATCGTCCCCGCCTCT

>NODE_1678_length_179_cov_1.563291

CCTTGGATATATGCAAGAGGAAGGCCCCCAGACGCCACAGCACCCCCCCCAAAGTAACCCACGGGCCCGA

AGAGACTTGTGGCAACATGGCAAAAACAAAAGGAAGGAACAGACGAAATATCCACGGCAAAAATAAGTCC

TAAAACAATCACCTAAATGTACTAAACGCAACCTGAAAG

>NODE_1679_length_179_cov_1.550633

TAATTGCTGCACCTCGTATGTGTGCAGTGTGCAGGTGAAACAGTTTTGGTGGAACAATTTTGATATGAGA

TTGTATATGCGTGGATCGTGTGTGCTGCAGTCGTTGTCAGAGATATGCGCTGTAGTCAGGACGTCGCACG

CAATAACGAACAGCAAGCTGGTCTGCCCTCCTGGCTGTT

>NODE_1680_length_179_cov_1.531646

GGTGGGGACGGCTCCACATCGCCGCCGCGAGAACGAGCTTAACCTTCTGCCCTCCCGAGAGCCCGCCGAT

CTTCCCGTAGACGCTGAACTCCTTGGCGAGGGCGAAGTCGTCCAGGTGGCCCTGAATCTCCTTCGTCGTC

ACCGGTCGGTTGTCGGTACCCGCGGCCAAGGCAGCCATC

>NODE_1681_length_178_cov_3448.305732

ACGGGTTAATATTCCCGTACCCTTTATTATTGCGATGGGAGGACGGAGAAGGCTAGGTGGGCCTGGCGAT

GGTTGTCCAGGTTCAAGTATGTAGGTGGGAGACTTAGGTAAATCCGGGTTTCTACTAACACTGAGATACG

ATGTCGAGGCACTAAGGTACTGAAGTCATTGATGCCAT

>NODE_1682_length_178_cov_644.783439

GTGTAGGATTGTTCACCCACTAACAGGGAACGTGACTCACGTGATGTGATGCGAAACGTTCGTTTTGTGG

AACACCATCGGTCCTGCACTCCGAGATCGAGACACTCCCACTGATTTAAGCTCGTACGGACTGAGGCTCG

TTTCGGAGAACGAAACCATTTCTCACTGAATGGGGAAA

>NODE_1683_length_178_cov_127.445860

GTACTGGTTCACTATCGGTCAATTAGGAGTATTTAGCCTTGGAGGATGGTCCCCCCATGTTCAGTCAACG

TTTCACGTGTGCCGACCTACTCGATTTCATGGTAAGTTTATTTTCGTGTACGGGGCTATCACCCTGTATC

GCTCCACTTTCCAGTAAATTCCACTAGCGCCCAATAAG

>NODE_1684_length_178_cov_30.146497

AGAAGAAGCCCGCCGCTAAGAAGGCTGCCCCCAAGAAGAAGCCTGCCGCCAAGAAGACCGCCCCCGCCAA

GAAAGATTAAGCTCGGACCTCATCATCTCAGTGAGCTACAATCGCAGTCTCTCGTCTCGACGGCGTGGCC

AAAAAACCAACCTGGTGTTTTCTAAACACCACCTGGAA

>NODE_1685_length_178_cov_29.433121

CTTATAACACTCTCCCCCCCTGGACCGGCGGGATATTCAGGACAGACCGGATATACGGGTCGTCGTGTCA

AAGCCAATTGGCCTCGCGGTTCCAATGTTATGGGGACCCAGGGAGAGTATTTTTCAGGTTCTTGACATGA

GGAAATTCACTTTTCTTTAAAAAGCGTTTTCATGGTGG

>NODE_1686_length_178_cov_20.828025

TTCCACTTTTTTTTTTTTTTCCACCTTTGAATATCCTTTAATATTGTGTCCATGGTCACCGCCGTGCGGT

GTGCAGCATGATCAAAACCACCATACACCTTGACTCAATACACATAAGTACACCACAAAACACAACACAA

CACGACACCTATAGCCACCAAATAACTTGGGGGAGGGG

>NODE_1687_length_178_cov_12.076433

ACCTGTGTCGGTTTACGGTACGGGCAATTATAACCTGAAGCTTAGAAGTTTTTCTTGGGAGCCTGTTTAG

CGTATTTCCGTCACCATAATGGTTCTGTTATCACTCAGCCTTTAGCAAGATAGCGGATTTGCCTACTAAA

CCTAACTACATCCTTCCACCTGGACAACCATCGCCAGG

>NODE_1688_length_178_cov_7.057325

TTGACCACGCATTCCTTTTGAAACGGTAGGAGGGGCAGCGGGGGGCAATTGCCTTCTGCTACCTGACCTG

GGTCTGACTTACGAGAAGGCGTGGGGTGGATAGGTAGGGCGACGGCAACATAAGGACGACGACAATGTCA

GGGGTTTGACATACATTGTACCATTTTGTATATGTGTA

>NODE_1689_length_178_cov_5.732484

AAGTTCGCACCGGCACGTACCGCCAGCTGTATCACCCGGAGCAGATCATATCGGGCAAGGAGGACGCGGC

CAACAACTACGCCCGCGGACACTACACCATCGGCAAGGAGATCGTGGACCTCGTGCTAGACCGCATCCGC

AAGCTGGCGGACAACTGCACGGGGCTGCAGGGCTTCCT

>NODE_1690_length_178_cov_5.528662

GGTTCTGCTTGCCTCCCTTCGCCTCCGCCGAGTCTTCGCCACCACCCTGCTCCTCCAGCTCCGGGATGTC

ATCATCGGAGTCTCCCTCTACGTCCTCGACCTCCGGGCCGGCCACAACCTTTGCCTCTTCGAACTCCGAC

ATACTAGGTGGGAGAGGTGAGGATAGAACGAACACGGG

>NODE_1691_length_178_cov_5.363057

ACAGTTTGCACTTCCAGCAAGGGACAAAAGAGTAAAAAACCGAATTGCTTATTTTATACCTGGAACCCTG

TAACAGAGACAGAAGAGACAGAAAAAGTGTTTTTAAGATAATCAGTGCCTAGAACCAGATTATCGAAGTT

TTCATTTTGCGAAGAACCAGAGAAATTTATCATTTCCC

>NODE_1692_length_178_cov_5.286624

CCCCGCTTGGGCACAACGCAATCCTCCATGGCGTTCCGTTCCATTCCGTGTCACAGTCTCTGAAGAAAGC

CCCGCCAGAATATACCGGCCGCTGCCCGACTTAGTTCTTCCTTTCTTGTTCTTGTTCTTCTTCGCCGCCC

CCTCCGCCAGGTGCGAAGCTCCACTCCCCTCAGCGTAT

>NODE_1693_length_178_cov_4.031847

GATCGCGAACACGACGTATAGCAGAACTTCACAGTGCAACGCCACAACACAAGCAGGACATTGTCCTCCA

AACATTGGGACACTACATGATCGGCAGCCAAACAGATGTGCGTGTTATCGTTCCTCTATCCAGTCCAGGG

TAGCAACGCAAGGCTATAACAGTTCCGGGGAACCTACC

>NODE_1694_length_178_cov_3.808917

TCCGGAGGGGATCACGATCGCCAAGAAATCTACCGCCTATGTGGGCTCCCTCGTTGATGGGTCGATTTGG

AAGGGAGACATTGGCACTGGCGAAGGAGGCGTCGTCATTTCGGGCGTCGGTGGCCCATCGATTGGCCTGG

ATCATGACGGCCGGTCGGGCTATCTCTTCGTTGCTGTG

>NODE_1695_length_178_cov_3.789809

AAAAATGCAGAAATAAAATACTCAACAGTCCAAAATTGGTATGCAGGTGACAAAAATGGAATCGGTGGAA

TTTATAACTTTGTTACTAAACGTGGTTTATGTATAGGGGAAAACTCGAAAATATCTTGGACACAAGTTGA

AACAGGATCTGCTAATACTTGGAAAAACCCTAGTTGAG

>NODE_1696_length_178_cov_3.707006

GCGAATTTTTACTACTACGTACTACTACTGTATACTGTATTGGGATTTGGATATATGATTGCTATGAGTA

TACCGTATTGGGATTTGGATATGATTGATATGATTTGGGTTGAACTGGATTTAGGGTGTTACTATTAGCT

TCTACGACGTATGCGTGTACCCGGTAGCAGAGGTGGAA

>NODE_1697_length_178_cov_3.127389

AAGGGGGTGCCGTTGGCTACACCAAAAGTCTGCAGCCGCTGGAACGCAGCAGTTCCGTCTCCATGGTCAA

AGTGGCTGTGGATGAGGCGAAGGAGGTCATGGCTGACATCGTTAAAGAAGGAAAACACCGACCGGTGCAT

GAACATGACGCCAGGGCGCGCAAGCCGCGGCTCGTGGA

>NODE_1698_length_178_cov_2.694268

TCTGCAAGAACAACAGAGTCATCAGGAGCGGAATCCGTGGCACAGCAGCGACCATGATGGTGCTGCCTGC

TGCTTCTCTCTTGCAATTTCCTTCTGGTGGATAATAAATAACGATCTGACGCTGTATCCACGAAGGCAAT

GCAGCTGTAGGCGGAGGTTTGCTGGGGAGGGAGAGTAG

>NODE_1699_length_178_cov_2.503185

ATTCCACGTACACACAAAAAATACTCACACGGCTATGCCATAAATACTGGCCTACCTTATGGTACCACAA

ATTACTCTATCTGCACGCACGTATGAGATCATCCACTACCTACCGACCTATGGAAAACGGTATCCCATTG

TTATACGCATATACCACACAAGATGACTCACATCCATG

>NODE_1700_length_178_cov_2.210191

TATATTTCGCCTGCGAATACGGTGTCGACTGCGCCTATGATCGTAACAAACCACTCTATCGGAGACGAAC

CAACTTCCACCGTCATCCCAAGCTCTTCGCGTGCGTTCCGCAGTTTCTGGAGCTCTTTCTTTATCCTGCG

CTCAGCTACGCTCGCCGTCATGGTTGGTGGTCGCCGCG

>NODE_1701_length_178_cov_2.197452

AGGCACCTTGGTCTGGGTCTTGAAGTGGAGAGTCTGGCTCACCGGTGTCACGATGTTGCCCTCCACCAGA

GTCGTCTGGTAGGTGTACTTGGAGACCATCTCGTCGTCCGTGTAGGTCACGTCCGGGCTCTGCACCACGA

AAGGCTTGTCGTTGAACGTGGTGGAAGTCATCTTTCTC

>NODE_1702_length_178_cov_2.127389

TCCGGTCGACCACTTGTCCGAACTACTTCTGTAGCCTAATTTCGGTCCACCATGAGTCCGAATCAAAATC

TACAATTATTTCGGTACCATCATCAATTCGCTCTGTATTGTTAGTAGGGCCCAACGTCAGGCCGAACTAC

TGCAGTAGCCTCATTTCGGTCCACCAATCACAAAATTT

>NODE_1703_length_178_cov_2.044586

ATCGACAAGTTGGTGATTCGTCGTGGGCGCGCCTGAAGCAAGACATCACATCGTAGAGGACAACCCACTC

AACACTCAACTGCAACACTACGTCAACAACAGCAGTACAGTACGTAGCGTGCGGGGTGTAGAGCCAGGTG

CGAGGTGCCAGCCAGGGTACGTAGTCTAGTCAGCTAGC

>NODE_1704_length_178_cov_1.980892

CCCCCAGAGTTCCCAGTGGAGATAGCCGCGTCCGTCTGGATGAAGGTGTCTCCAGCAGGGTTCGCGTTCG

ATCTCTCGTTCGCCAATAAGTCACTCTTCCGCCTCGCGACGGCGCTGACGATTCCCAGAGTGCACGTTTC

CACCAGGCCCATGGGGCTTCCCATCGCCACTACGAACT

>NODE_1705_length_178_cov_1.834395

CTAGCTTTTAGCACGCCTCTCCACACCAACCACCACAACAGTACTTCCCACACAACTTGAGTTGTCATCA

CATAATAATTTCGTTTGTAACCTCAAATTTCAAGCAATTTCCCGATCCGAGATCCCAGCCAAGCAAGCCA

CAAAACCTGTGCGGCTCGTTTTTCGCTTCAGCAACTTA

>NODE_1706_length_178_cov_1.770701

CTTTTCGAGCCTTTTGCCTGCTTTTGGCGTGGTTTCCCTTTCTTTTATTTTTCCGCTCGTTCGTCACCAA

GTGCTACTCGTAGCAACCAGATGTCCGACTACGCGACGGAGCTGCTTCGTCGGCAATTGCAAGAGCTGAA

CAAGAACCCTCCTGAGGGTGTCTCCGTGGGTCTGACGG

>NODE_1707_length_178_cov_1.681529

CGGACCCGATATTCCGACGGGGGTGACAGTCGGCGTTTGCCTCGTGACATTTTTTACGTTCGGCAACGCC

CAGCGGCGAATAGCGCAGGCCCTCCGGAGAAAAGAAGACACGAAGAAACCTTGAACACGACGGCAAAGGG

CTTGCACTATTTATAGCGCCGAAACTTTGATTGAAAGG

>NODE_1708_length_178_cov_1.554140

AGGGTTTAGGGTGTGCGTGTTTGTCGGGTTTCTATTGTATGGGTGCATTCCTCCCCTGCGCGGCTATGCA

TGTGCGTGTTTTGGTCTGCTAGGGAAAATTTTGACGAGGGAAGAGGGAGGGGGGAGAGAGAGAGGTAAAC

TAAACATTCAATATCCCCCCAACAGTTGAAATGGAGAG

>NODE_1709_length_178_cov_1.503185

TCTTAATAATAACACTGCGGAATACAAAACATGACTCAAAGTAATACGTCAGCAGTAGACTACAACTTCA

ATGTTGTACGTCAATTTACTGTAATGACCGTAATCTGGGGGATCGTTGGTACATCAGTAGGTGTGCTTAT

CGCGGCTCAATTAATTTGGCCTGCGATAAATTTTGAAA

>NODE_1710_length_178_cov_1.050955

TGTCCAGATCAATCATCAGCTCCCCTGCACCGCTGATATCAGCCGTCAGATCCAGGCTCTTGCTGGCCAC

CGAAAAGTCAACATTCAGATCAAACGGCTTTTCCAGTTCGATGGTGCCCGTCCTCAGGTTGAAACCTGAC

AACAGAGCGTCGGTACCCGATACAGCATCTTTCCAGCT

>NODE_1711_length_177_cov_96.314103

TCAGCACTTTCTTCCCTCGTCAACTTTCGTCGAATTGTGCTTACCCACGCTATAATATGCGCTCTCGACA

GTCGGTGTCATTTGAGAATAAAAACCTCCATGGGGAGAATCGAACCCTCGACATCCGGCCCAAAAGGCTT

CGTGGCTACCAGTTAGACCACCGGGGCGACCGGCTAC

>NODE_1712_length_177_cov_6.416667

TTATTTCATCCTTCCCTTCCTTCCTTCCTTCCGTTCCCGCCAGCCGAAAAAAAAAACCCGCAGCAGGCGC

GTAAGAGGCTTGCTTCGCTCATCCGAGTCCCCGGTTCGACTCTCTCGTCCATCCACACACCAGCGATCGT

CCAACGAAAAACCAAATAGCAACCTAGCGCTCCCTCC

>NODE_1713_length_177_cov_4.961538

ACCGCTTTTTTACCTTTAATTGCTTCACTTAACGTAAAAACGTCAACAATTTCACCACTACCTAATACTG

CTGCTGCAGTAAAATCTGGTGCTGGACGACCAACTAATACACTCATTTTATGCTCCTTTCTATTAATAAT

AATTAAGAATTATTATTGAAGACAATGAATAACAACT

>NODE_1714_length_177_cov_3.961538

TTAACAACGTAATCAATAGCAGATTGAACAGTAGTGATTTTCTCTGCTTCGTCATCTGGGATCTCTGTAT

CGAATTCTTCTTCAAGAGCCATTACAAGCTCAACTGTGTCTAGTGAATCAGCACCAAGATCGTCAACGAA

AGAAGCTTCGTTTTTAACTTCGTCAAGTTGAACACCT

>NODE_1715_length_177_cov_3.782051

GATCGTCGCCTTCGTGGGGACAGCCATTTCCAGCCCCAGCGACTTATCCTTGCGCTCCTGGTACGCCTCG

GACAGAGTCTTGCCGCTCGGTAGAGTCTGCGACGACGTCGTCATGTCCAGGATCCAATCCGCCGGGTTCG

TCTCTGGCGGGAGCTGCAGACCCAAGCTGTCGAAGTA

>NODE_1716_length_177_cov_3.737179

TATGGCGGAAGAGAGGAGGGACGAGGGACCTGGGGGCGACGGAGGCCACAAGCCTCTCACCGAGGATGAC

ATCGAGACGAACTGGAATGAGTGCATCGAGACGTTCGATGCGATGGAGCTGCGGGAGGAGCTCCTCCGTG

GCATCTACGCGTACGGTTTCGAGAAGCCGTCGGCGAT

>NODE_1717_length_177_cov_3.474359

GAGTGCACAAAAGTAGGCTGTAGCGTCACTGTTGTAAGGCGTCCCGTTGTGTTGCTGCCATGTCAACCTA

CGTACCAGGTTCGGTTTGGTTTTGTTTGGTTGTTTTTGGACGTCTCGCGGTGTACATTTGTCCAATTTGT

GTACGACGGGGCTCGGACCTTGGACCCTTGGTGTCTC

>NODE_1718_length_177_cov_3.314103

CATCTTCGACGACTTAGCATCGCTTGAGGATTTGGACATTCGAGGCAATTCGTTTTCTACTCTGCCAGAC

GGCATCTTCGGGGAACTGCAGATGCTCCTCGAATTGCGTCTTGATCAGAACGACTTCGTCTCACTTCCGC

CGGATATGTTGGACGGATTGGTCAGCCTGCAAAGCGT

>NODE_1719_length_177_cov_2.852564

GTACCACGAAAACACGTAGGTCTGTACTGGAGCGAAATTCATGCAACAAACTATTCGGTTCGGACCGACT

CGATCGCCATTGGGATCTGACGATCAAACGGGTTTGTCTACCGATATTTTCATGGGGCCTCCTACCGGCT

ACAAGTCTCGTGCTGCTGGTTGGTGCAACACAACACC

>NODE_1720_length_177_cov_2.775641

CTCGCCCACTCGCTCGCTGCAAGGCTTACTTTCTCATGATGTTGAGCACGACGGCGGAGGGCACCATGAA

CATAACGACCACCCCGAACACCGCCAGGACTTTACACCACGTCCGGTACGCGGCGACGTTCTTCCCGTCA

ACGTCGTTTCGGTCTGCCATCCCCTTGGTCGCCTGCA

>NODE_1721_length_177_cov_2.769231

CTACGGGTGCATGGACGCTACGATCGACATTAAGGGCAAGTGCAAGATGGTCGCCATCGACGGCTGCAAG

AAGACACAGGTGCTCTTGGACGACGCCATCTCGTCCGTGGAGCTCGTGAACTGCCAGCGCATCAAGGTGC

AGGTCCGCGGTTCCGTTCCGTCGTTCTGTTCTGTTCT

>NODE_1722_length_177_cov_2.397436

GCATGCCGTTTTGTTTTCTGACCAGCAGGTACACTTTTTCTGCGCATCATAATGGTGTTACGATGTTGTA

GTTGGGGGTGGAGTGGAGTGGTGAACGGCAGTTTGGTGTTAAGTTGGTCTTTGTTGGCTCCCTTTTCAAG

GAGGGTTTCATAGAGACTTTCTGAAGCTGCTGCCTCT

>NODE_1723_length_177_cov_2.346154

TTTTATGATGAAGACGATGACGATGCGCCGGCTAGACTGATCGACTGGTGCTGGTGAGGTAACACCGAGA

GAGACTACTTGATTGGTTGACTTGACTTCAACTTGAACTTGAACGAATGACAAAAAAAATGGAATTGCAA

GGTTGTTTTGTTGGCTTTGGCAGCATTGTTTTCTTGG

>NODE_1724_length_177_cov_2.262821

CAAGCCCGCGCCGACCATGGCGGCGAAGATGCTCGAGAGCTGGGCGTGGTTCATCATCGACAGCGGCTTC

CCTTTGACCCAGGGAACGTCCTGAGGCCAGAACAGGGGCCGCTTCATGCTCACCCCGTACTGAAGCCTTA

GGCCGAGCTTGGAGCGGATGAGGGCGGACAGCTCCTT

>NODE_1725_length_177_cov_2.198718

CCGTGAACTTCGGGTTCTGGTCGTTCGAAATTATCTCCGTTCGCCCTATTTCCCTCCAAGTGTCGTCAGT

GTCCGTTTTCCACTTCACTACAACGAACGGATCCGACTTGCTTCGACGGTCCATTTTTCTGAGGTCTTTG

CACGATATGCTCGATGTGACTCTGCTCCCCAGACCCT

>NODE_1726_length_177_cov_2.128205

CCATCCCCATCCCATTCCCAAGCTCGATCCCAGCCGCCCTCATGCCACCTCCCATGTGCTTCATGTCCAT

CTCCACGAAGTTATCGGTCGCCACAGCAGCCGTAACGGGGATGAAGTCCGGAACCGTCTTGACGAATTTT

CCCTTCATCCTCGGGCGCACCCTGGCCGCCTTCGGCC

>NODE_1727_length_177_cov_2.076923

TGATGATGAGCTCTTTGATGCGCCCAGTGATGTGCATGAAGCCACTGGGGGATGGGATCCTCTTGTCCTC

GTCCTGGTCGAAAGTGGCGATGTCCCCGCTGTGCATCCAGCCATCCTCGTCGATCGTCTCGGCCGTCTTC

TCGGGGGCGTGCATGTACCCCATGAAGATGTGCCGGC

>NODE_1728_length_177_cov_2.051282

AGCGCCAATATATTCGTGGAACTCTCCGGATCTAAAGTTCAGTCACCAACCTACCAATAAACACCAACCG

ACAGTAGTAGCAGCAGCATCATCAGCAGCAGTGCTTTTGCCTTCTGCGGGCGGTTTCACCTAGTAACTCA

GCAGCCGCACTACACGCCTTACGGCCACTCTTCGAGA

>NODE_1729_length_177_cov_1.948718

GCGTGGGGTCTCTCTCCTCGGCTCGCACGGCGCCCTCGCCTAGAAGGTGTTCTTCGTCAAGCCTTGCCAA

GGCGTGGTCACTAGCGGGGGTCGCTGGTGCTTGGTACTCTTGGAGCGCCAGCGCGTTGTGGTTCTGGTGC

TGGATACTGAGCGGTATGTCGTGCAGGAGCGAGTCGA

>NODE_1730_length_177_cov_1.673077

TGGTGATAGACAAACTGGAAAAACTTCTATCGCTTTAGATACGATTATTAACCAAAGAGGTCAAGATGTT

GTTTGTGTTTATGTTGCAATTGGTCAAAAAGCCTCTTCTGTTGCACAAGCTGTAACTAATTTACAAGAAA

GAGAAGCTTTAAGTTATACTGTAATCGTTGCTGCAAA

>NODE_1731_length_177_cov_1.673077

GCTCAGCGTCGTCATCAAATTTAGCAGCTTCTTCGTAATACTTAGCAGCTTTTTCTAGATCCTTAGAGCG

ATGATAGTTACTAGCAATCGCGCTAACATTCTGCTTAGTACGTTTTACCTTTTCACTCTTAATGGCCTTT

TCTAGCGTTACAGCTGCTTTATAAGGGATCTCATTTA

>NODE_1732_length_177_cov_1.352564

GCGGTAGAGAGTCGGCTCGGTCGGAGGAAAGGTTGGGCATGTCGGTGGGATCGAGACTTACGCGGCTGCG

CGTTGCTGTCTGAGCGAGTGATGCCCAACGTAGAAGTATGATTGCTTCGGTTCCGTTTTGTGTAATACTT

CGAAGTATATACTTTTGTTAAATCCTGTTGACTCCTG

>NODE_1733_length_176_cov_7.445161

TTTCTCGCGCATGCGTTTTCGCCCCATGCCGCCCACCTAGATGGAACTGGGGGTGACCGCTGTTTTGTTT

TGGTTTTCCTTTTGCAGCGTGCGTTTTCGCCCAACGTTAGTCCCTGGGGATGTGGCGGTACAGTAGCTTG

TGTTTTTGTTTTTCATGGATATATATTCTCCCCGCT

>NODE_1734_length_176_cov_5.038710

GTTTCTTGGATTGGTTTGGTTTTTGCGTTCCTCGTTTCGTTGGCGTCGTCGGCGTTTTTCGTTCGTCGTT

CATTGGTTTGGTCGCTGGTTGGCGCGCATAATGTAACGTTCGGCACTCGATCGCTACCCTGGTCAAGACA

CGATGAATGGCATTGAATGAAAATATGCACGCCTCT

>NODE_1735_length_176_cov_4.632258

TCTCTTGCTTGTTGTTGAAGGCCCAGGAGATCTTGTTGTTGTTACCGGTGCCGAGATCGATCATGATGTG

CTTGTTTCGGAAGAAGAACATCACGGTGCAGGGGTCGTACAGCTCGTACATGTCGTTGAAGTCGGGCACC

TCCGATACGTCGACGACGTAGATAGCGCAGAAATTC

>NODE_1736_length_176_cov_3.419355

TGTGAGAGTTAGTTGTAAAGAGACTTTTGGGTACGAGTAGCCTGTCTGTATTATGTACGGTTTAGTTGTG

GATCATTGTGTATGCCATTTTTTTCTGGCTGTTTGTATCGTGGTTGTGGTTTGGGAAGTTTATAAAGCGG

CGAGATTGTCGGCATCTGCCGAAAACAAAACGTGTT

>NODE_1737_length_176_cov_3.419355

GTGGGTATAGTTGGGGAAGGCTGACAGATCGGAGGTGTTCCAGAGATATATGGGCGTTTATGTTAGGGTT

TGGTTTATCTCTCGTGCGTGGATCACATTTCGTACGGGACGTGACGGTAGTTCCGTAACGTAGGGTAGAG

AGGCAGGTAGAAGGAGCAGCGATGCAAAGCGAGGGC

>NODE_1738_length_176_cov_3.161290

CCCGGCTTCGTGCCAAACAGCGGCCGGTTGCCGAAGCTCTGGCGAGTGACCAGGTGCATATCGACGAGGG

TATCGTAGTCCATGTAGTGCGGGGTGGCGTGGATGGGGTGGTCAACAGCTGCTGTCTTTGAACCAAATTG

CCTCAACGGCACCGCATTCGCGAATCCATGTCGTGA

>NODE_1739_length_176_cov_3.122581

GACAAGCCCGTGGCGCCAGCATAACGACGCGGCGCCACTAGCGACTTCCTGCCCCACAGTTCAACTGTGA

TCGCAGAAAACGACCGAGGTGTGGACAGGACGTTGACGTCCTGCCCACTACCTTCATCTTCCCACCGACT

ACTATCGGATGGGTTGACAAGCCCGTGGCGCCAGCA

>NODE_1740_length_176_cov_2.600000

GGCACGATAGTCACGGAAGCGACTCTCACCAACGCAGTGGGCAGCGACGTCGTCTGGAACGTTCTGGAGA

ACAATCGCGAGTATATGGAGAAGAAGTTCGGCAAGGTGACGTGGCTGTCTCCCGAGGAGAAGAAGAAGAA

GCGGGGTTTCCTTGGTTTCGGCGGCGTTGGTATCGG

>NODE_1741_length_176_cov_2.503226

ATCACCAAATAAATTACCTAAAAGTAACCCACGTGTATAAAGATTTTCCCGCTCATTATACTCGTTTTCA

TACGAAGGTACCCTTAAAGAACAACTAGTTTTTACTTTTTTTTCCGCGGTATCCATCAAAGTACAAAATA

CATTTCTTTTTGTTGATTTGATAATAATAATACCTT

>NODE_1742_length_176_cov_2.374194

ATCGACTTCTCGCTCGCCTACGGCGGCAAGATCTTGCTGCACAACACCAGGCTCAGGCTGCTCGCCGGAC

ACAGGTACGGACTGCTTGGCCCTAACGGAGCGGGAAAGACGACCTTGATGAGGAACATCGCGAACGGAGC

CATCGACGGCTTGCCCACCGAGCTCAGGACCGTCTT

>NODE_1743_length_176_cov_2.116129

CCCCCGACCCTCCCATCAAGCAACCCCGAAAGTGCAGCATATCCTACCGCCCTGGTAGTCTCGTTCACGC

GCAGACAGGGGGCAGCGCGTCACCACGAAACCAGCCAAGGGAGACAAAAACAAACAAGCGCACAAGCAAA

GACATAGCACCCGTATTTTCAACACGATTCACCTCT

>NODE_1744_length_176_cov_1.903226

TGCGGCTAGTTGGCTCGCGTACGAGCGTCACTCTGCCGTTCTCTACTCGTCTGGTCGTGCTTCGTTTTTC

GCTTGCTATCGGTTGTCCGGCATCGCCCCAAGTGGTATATACACAGTCCTCCTACAATAAGCAATATCCG

CGACGACCTAATCATCCACGACGAAAACCCGCCCAC

>NODE_1745_length_176_cov_1.883871

CACGCAAGGGGCTGGTCTTACTATTAATAGCATTATTGTCCTAACCCGAGGCTGGCTGTACGTAGGCATG

AATAAGAGCGTTTGTCTAGTCTTCCCCATCGGCGACTGTCGGTAGCTGTTGCGATTTGTGTCTTGGTGTT

TTCTCCCCCAGTTGATGGATGGTGGTGGGCTGGAAT

>NODE_1746_length_176_cov_1.741935

ATCAAGCATCCTTCCTTCCATTTGCAACACCTTCGGCGACTAATGTCCGCTGCTATATATATGTATTCAA

GTATAGTATATTATCTCCCCTACGGATTAGCCCGAACCTGATCAGCGCCCCTCCCCTATTCGTTCAACGG

CTAGTAAGCAGATCTCCCTCCGCCTGCAACACCCCT

>NODE_1747_length_176_cov_1.729032

TGCAAAAAAAAATCCTAGTGATTAATCGGCCGCCAAACATTTTGTCCGAGATATTTAATTTAATGGGCAT

CCTCCGAAATGTGTTGTCTGGTCGATTCCAATTAATAGACTGCTTCATGAATCAATTGGTCCGGCAATTC

CAATCGCTAGACAGCCTCACGAATCGTTTGCCTTTG

>NODE_1748_length_176_cov_1.180645

GAAACATGCATCGATAATAAATATCGGAACGCGCGTCGATAACAGCTGCTACAGACTTCGATATGCCGTA

ACAAGCATCGACATAAATATACATATCGGAACACGCATCGATATATATCGGAACCTGGGTCGACATGTGT

GTATCATATCGGGAACCTGCATCGACGTATTTATCG

>NODE_1749_length_175_cov_1866.272727

AGACGAATTAGTTGGAATGCTAAATCCTAGAAAGTGAAAATCTTGTAGTCGAAAACATTAAACCCTTTAG

ATTGTATCCCGAGTAGCATGGGGCACGTGAAATCCCGTGTGAATCTACGAGGACCACCTCGTAAGGCTAA

ATACTACTGAATGACCTGGAATTCTCGGGGTGCCA

>NODE_1750_length_175_cov_508.707792

GGAGCCGTAGCGAAAGCGAGTGTTAACTGCGCGTTTAGTTGCAAGGTATAGACCCGAAACCCGGCGATCT

ACCCATGGGCAGGTTGAAGGTTGAGTAACATCAACTGGAGGACCGAACACACGTATGTTGAAAAATGCGG

TGATGACTTGTGGGTCGGAGTGAAAGGCTAATCAA

>NODE_1751_length_175_cov_94.201299

GACAGGTTCCTTCTGGCAGCACTGCTTTTTACTGCACTTGTAGTCCTTGCACACGGTGGAGTCAGCGTCC

TTCACCAGCTCGTAGTAGGTAGGGCAATCGTAGCTCGAGCACACCTTGTTGCAGCACTGTGACTTCTCGC

ATTTGCCGTTCTTGCATTCCTCCTTGTCGGCGTTC

>NODE_1752_length_175_cov_4.668831

TCTCAACAGAAGACTCCTCTAGGGTGCCGTCAGAATTTTTCTCGGAGACGTCAAACTTCTTGAGCCGCGC

CAGCTCAGCTCAGAATTTTCCTCTTTTAATAGAACACTCCTCCAGGGCGCCGTCAGAAATTTCCTAGAGT

ACGCAAAAGTTCTTGAGCCGCGTCCCCAGCTCAGC

>NODE_1753_length_175_cov_4.168831

CTCGGTAGAGGATCCCAGGGCCCTGCAAACATGCAAATCTTCGGGAAGACCCTCACGGGCAGGACCATCA

CGTTGGATGTTGACTCCTCCGACACCATCGTGGGCGTGAAGCAGCTCATCCAGGACAGGGAGGGCATCCC

TGCGTGCCAGCAGCGGTTGGTCCTCGGAGCCAAGC

>NODE_1754_length_175_cov_3.623377

TATAGATGTAGTCACTTTTCGTTTTCGCGAGGGTCTCTTAACCAGGGTGCTTGCGTGTTAGGTGTGATTT

GTGTATGCAGTACTCGGTGTTTCTTTTGTCGCGATGACCTGTTAAGGACGCGCGCTTGTTACGTGGGATT

GTGTGTGTGTGCTTGTGGTCTTCAGATATGTCGAG

>NODE_1755_length_175_cov_2.525974

CGAACGTCACCTCCACCTTGGGTTCCCCCTTACGCGCCCGCTCCACACCAGTGATGGTGAATTCCCCCAG

CTTGTTGCACGTGTCCACGGAACGGCGCTCGCCCTCGTACACGACTACGTCGATCGACGTCTGCCAGTCC

TCGACCGTGGTGTATGACCGCGACTTGCGGCACGG

>NODE_1756_length_175_cov_2.512987

CCAAAAAGGGTCAATATTTTTTATCCTTGTTCCAAGTTTTTCTGTTTTGCTATTTTTCCGGCCCGCTGCG

GGAATCGAGTCAGAGGATGTCCGTTGATACTACTACGTGGCGAAAATAACCAATCAAAACCGTCCCATGC

ACACACATCAGAGGCAAAGATATTCCCGCCCCGGC

>NODE_1757_length_175_cov_2.461039

AATCCTGTTTACTAGTTTCCGAACCCATTCCGAGCGATAATATGATGGTGTAAAACGTTACTCAAGCTCG

CTGTTTCGTCCGGGCTAGCAAAATCCTACGAGTAGTGAGCCGATGCTCAAACTCCAACTACATCTGTGTA

GTGCTACTAGTCACGATAAATGAGATGCACAAAAC

>NODE_1758_length_175_cov_2.357143

TCACGCAGCGAACCAACTGATCAACCGACCCGATGAAATCAAAGCAAGCATATCAGTCAATCAAGAAAAT

CAAGCAATCAAGCAAATCAGTCAATCAAGAACTCAGTCCATCAATCAACCAGTCAATCAATCAATTCAGT

CAAACCAATCAGTCAACCAACCAATCTATCTATCA

>NODE_1759_length_175_cov_2.292208

AATATAGCGGCACGCTGCACAACTTTCCACGAATGTACCGACATGACGTACTGTTCTGGAAAATGTATCG

ACACTTGCCCGTTATTGTTATTTATGGATTCCCTTCCTCGTTTGTGCGGTTAGATTGCCACTGTACCGGC

ATAATGCTACCTTATGCCATTGTACCGGCTGGAAT

>NODE_1760_length_175_cov_2.136364

AGTCCTCTCCTGACGAAAGTGTTATCACGCCGCCCTTGTTTCCGCAGTAGTTTCCCGTGAGCACGCAGTC

CGACAGCACAAGCGTGATATCCCTGCCGACGGCAAAGGCACCCCCTTCTCCCTGCGGATCGTCCTCGATA

AGAAAACCCGTGGTGGAGTTGGCCACCAAGCATCG

>NODE_1761_length_175_cov_2.012987

GTATTTTTGATCAAGCATTGCGAGAGTATCATATAGTAGTAGAATAGTAGTAAGCGAAACGATCCGCAAC

TGAAAGTCCTTCCTTCCTTGCTCCCGTCTGCAGTGCGGGACAGGAGGAGGAGGAATGCACGAGAACCGAA

AACGAAGTAATCTAAATCGAAAAAATGAAGAAAAT

>NODE_1762_length_175_cov_1.980519

GCAGACACCAAAGTGAAGATCAAGTGCAAGAAGAATGGCTGTGGGGGCATCTTGAGGCACACCGATCCGG

CCGGCGGTCGCAAAGCCGCAGGCGTTAAGTGACTGGAATACACGCGAGCCGGACTCGTGTACAGTACTCA

CAGTCGTGTGGAGTGGAGTTCCCCGTTTCCTGGCG

>NODE_1763_length_175_cov_1.974026

AGTAGTGAGTAGAAAAAGACATTTCGTGCCCGCGGTAGTATTGTCTGGACCGTGATTGGCGCCGTCCGTC

GCGACTTGTGGGAAACGCGCGCACGCTGTCCTTTACGGAACGTTCCCAGAACCGGTGTTTTCTCTTGTTT

TGCTGCATACCATCAACATGTATCTGCTCCTGAGA

>NODE_1764_length_175_cov_1.688312

TGATCTGCTCACGTTCTTGCGGAATTCCGTCAAGCCGCTTCCGGACAACCTCGCCAGACGTATCATCGGG

GACGTTTGCGCAGGAATGGCGTTCTTGCATGCCAGGGAAGCCGTCCATGGGGACCTCAAGTCCGCGAACG

TAATGCTGGATGGAGACGGCCGGGCGAAGATCGGA

>NODE_1765_length_175_cov_1.571429

ACTTTTTCATACAAATGTTCTGCTACTTCTGCTTTAGTAAGCGCCATATCTAGTCCCTCAACGATGCATT

AAACTGTTCGGCCAATTCGGCCACCACGTTATCAACTACTAGATTGATATCTTTCTCTTCGAGTGTTCTA

TCAACCGCTTGCAGTGTTAGAGCAATCGCCAATCT

>NODE_1766_length_175_cov_1.551948

CTGTCGGCGCGCTCAAAGGAGGACAATGTCCTGTCTACACATACTCACGCAGGGTCGCCTGCGCAAAACG

GTGCGGTCATAATGGCCGGCCAAACGCAAGCAACCAGCGATCTGCTGCTGGAGACTGTTCTTTCTTCCCT

GAACGACGACAAAGCCGAAGACATCGTGCAGATTG

>NODE_1767_length_175_cov_1.272727

CAGCAGGGAAGCACGGAGAAACAATGGAGTGCACGCACGCAATCCCTCTCCGAGAGAGAGCAGAAAAAAA

AACAAATATCACAAAGATTGTGGGGGCAGCTGGCCTCTCGTCGATCAAAGCAGTAGATGCACCGCTTTGT

TTTAGTTACGATGATCTGATACAAAAACGAAAAAC

>NODE_1768_length_174_cov_2694.732026

GACTGTAGTTCCTTGGCACCCTAGATTTCCAGCGGATCAGTCGAATCTAGTAAAGGCATCGACATGAGCC

GAGTCGCCCCCGGAGGAACGCCCGGCTTGCGACGAGACTAAGCGCCATTTTTCAGAGTAGCTGGTTATTG

ACATCATATCCGCTAATATCTCACAGCTTCGGCA

>NODE_1769_length_174_cov_189.045752

CGGGAGGCAGCAGTGGGGAATATTGGACAATGGGGGCAACCCTGATCCAGCAATACCGCGTGTGTGAAGA

AGGCCTTCGGGTTGTAAAGCACTTTCAGTTGTGAGGAAAGGTTAGTAGTTAATACCTGCTAGCTGTGACG

TTACTGACAGAAGAAGCACCGGCTAACTCCGTGC

>NODE_1770_length_174_cov_5.457516

CTGCAAGTCAACCTCCCCAAAGATGTGTTCAAAATTAGCGTAGAACACAGGAGATCTTTGAGAAATCCTT

GGGTCGGGATCATCCGAACGTTGCGACAATTCTCAACAACCGGGCGCTGTTGTTGGAGAGTCAGGGGAAG

TTTGAGGAAGCTGAACAAATCTTCGGAACAAGAG

>NODE_1771_length_174_cov_5.078431

CCAACCGCAAAGCCACCCACCATCATCAGTGACGATGCCTCGCTCACGTTCAAGAAGCCTGATCAAGTTA

TTTCCAGGAACGCCGGCCTCGACGAATTTTCGGACGAATTCAGCTGCACCTTTGGATTGGTCTTGGACGG

AGCGACCCTCGTTGGCTTCACCGGCGATGATTGG

>NODE_1772_length_174_cov_4.627451

TCCTCGGCACGCTCGACGGGATGGTTGTCCCCGACGAGGATGCCTCCTGTTTCAGTTTTCTCTCGCCGTG

TGAAGTGCCCTTTTTCCGATGAGGCAGTGCCGGTGTGGGGGTGATCTCGAGATACCCCATGTTGGGAAAG

GAAATTTGGTCGGCCGTGTTGGTGTTAGGGTCCG

>NODE_1773_length_174_cov_4.326797

TATCCGGACAGATTTTCCGCAACGGCGAGGTAGTCGAATCTGGCGTCGGCTCTCTTACCTTCGAGAAGCC

GTCCGAAGTGACATTTGAGGACAACACCGTCGATACCGAATTCAGGGAGGACGACGACTGCACCGTGGGC

GAAGTTGAGGAGGGAACCACGCTGATTGGCTTCC

>NODE_1774_length_174_cov_3.954248

AATTTCTAGTGTTATAGTTTCCATTTTTTGTTCTTGAGCAAATTTTTCGGTGTTTCGTTTAATCTTACCT

CTAATAAATCCTGGGATTTTTGTTAATTCGGCCTGTGATTCTAAATTCCATTTTATTTCTGAGTCTTGTG

AAGTTACAGATAACCCTGCACTCAAAACTTCTTT

>NODE_1775_length_174_cov_3.830065

GCGCGCGGCCAACTGATGCATTCGCATTTATTTCAGCTCAGCTCTTGCTTTTTGAAGGAGAAGAAGCTCT

GCCATTGTGATGCAGAGCAAATAGAACTGCCACGGCTATGATACCAACACAGGTCTACTAAACTGGATCC

TATCAATAGCGAAGAACAAGTAGGCCTGGATCGT

>NODE_1776_length_174_cov_3.496732

CTTCCCGATCATACGGCGGAAGGTGCAACGCTGCGATCGCGTCCGCGTTGCGGGCTGTAATACCTCCCCG

ACCATACGCATCGGCTTCTTCGGCATGCATCGAAGAGCCACTCGTCAGCTTAGAAGAGCTACTCATCACC

TCGACGTCCAGGTCATGGGCTTCATGCCACATTT

>NODE_1777_length_174_cov_3.477124

TTCCAGACATCGTCATCCGTAGCAAGATCAACCACGCAGTCCTTCAGGTTGGTTGCCTTGAAAACGCGGT

TCAGGTCAAGAGTGTCCAACGTCCAAAACTTAACGAACCAGCCCTCGATCGCATTGAACAGGTCCAGCAC

CAGCCCGCTTCCCTGGTAATCACTCCGGACGGCA

>NODE_1778_length_174_cov_2.836601

AGCCTCGTGGTACAGGTGCGACGAATGGAGCAAAGCCTTGGAAGGGATGCATCCCACGTTCAGGCAAGTC

CCGCCCAGTTTGCCTCTCGACTCCACGACGGCAGTTTTCAGTCCCAGTTGACTCGACTTGATGGCAGCTA

CATAGCCGCCGGGGCCTCCTCCCACGACAACGAC

>NODE_1779_length_174_cov_2.359477

CAAGTTGCACCGCAGAATCCAGGCCCTGACGTTCTGCTGGATCACCTTCACCTCGTCTTCCAAATCGCTC

AGGTCCGAAGGTCCCATCATCGACATGATCGCTCGGAGGTCATTCTCTCGCCACCCCCTCTTCTCGCCAC

CCGCGAGAACCTTCGACATGCCGGCATGCTCGGA

>NODE_1780_length_174_cov_2.359477

GGGCAGTTGATATGTAGTGTGGCACTCGCGATATAGAGCGTAGTCTTGGAAGAATGCATGGCTGGCTGGC

TGTATGCCGACGTCCGTCTTCTTTCTTTTTTTTTGTGATTGTACTGCTGCTAATAACGATGGCCTGCTTT

CATTTAATTCACTACGCGAATCGACTTTTAGTTT

>NODE_1781_length_174_cov_2.313725

CGGACCGCAAGATTCCGTTCCGCGAGAAAGTGCTGTGGACGGTTATCACTTTGTTCATCTTCCTGGTGTG

CTGCCAGATCCCCATCTACGGCGTCCAGTCCGCCAAGTCCTCGGACCCGTTCTACTGGATGCGAGTCATC

CTCGCCTCCAACCGTGGCACCCTTATGGGGCTCT

>NODE_1782_length_174_cov_2.222222

CCAGCGAGAAGTAGGCAACAACGTCATAACCTGAGGCGGCATAGCCGGTCTCGTCTACATATTGGTCGCC

GGCAAAGGCTGCACTGGCAAAGCTCAGCGCGGTGGCTGCGGCTGCGATGGTGATTGCAAAACGGTTCATT

GTCTTCTCCTTAGTGAGGGCACTGGTGGCCGATT

>NODE_1783_length_174_cov_1.712418

CTGGAGGCATGGACGATCAGCACCCTGGACACGATTTTCGCACCTCGCTTGGTGTTGAGACATCTTATGG

TCGCCGTAGCGCTGCAAACGGGACTGTTGACCTGGAACGGGCTGGTGACCGCGGCAAATCGACTGCTGTC

TCGATTCAGCACGCGGCACGGAGAGGCCAAGGCC

>NODE_1784_length_174_cov_1.653595

GATCGACGTCGCCGTCATGGGCTCGACCACCACGGGCAACCCCCTCGAGCTCTACCTGGGCTTCCTCCAC

TGATCTAATTGACGGAATAACCGAACGGGACGCGTTACCGGGAACACCCCCGTCACGGGTAGCCTAGCGG

GGAACCTGACGGGAAGTCGACGGACACCTAACGG

>NODE_1785_length_174_cov_1.522876

AGCGCTCCCGATCGTTCCCGCGTCATAAATGGCGTTGTTGGTATAGTATCTCCTGCAGTTTTCCACCATG

AGACGGAGGTCCGCCACCATACCCCCCTCGTCCTTGTACTCCCCGTCCAACAGCTTTACCACTACCGTCG

CCATATCCATCGGGTGTTTCACCTTTTCCGCGTA

>NODE_1786_length_174_cov_1.516340

GTTGTCGGGACAAATTCGACAGCAGTCGCGTCGCCCAGAACAGCCGCGGCAACTGCGCGGCATACGGAAA

CGTCAAAACCCTGCCATTCGCCGTTTGCGTCCGGTGCAGCAAAGCCGACCAGACCTGTTGTCACGCCGCA

GTTCAGCTTGCCGCGCGCTTTGACGTCATCCAGC

>NODE_1787_length_174_cov_1.405229

GAACATGATCAAAGTCATCAGAAGGAACAACCCCACCTGGAAGATGGCGATCATCAATGCCAAGCCGGGC

CCCTCGAAAGTCCATGGCCTGACGGGGCAGTTGTTTTGGTCAATACTCGAGTATGTACCCGGGTAATCCC

AAGAGTATATACCAGGTACCCTAGTATTTACCCG

>NODE_1788_length_174_cov_1.313725

CAGAAGCGCCAATAACTAAGGTATAGTACTGCTGTACGCACCCAAACCCAACGTACCTACTACTAGCGAG

CGAGTCCGGAGGACGACCTTTCCGTGCATGTACACGTGTAGGGGGAATCCCGGCACGCGATGCAGAAGCG

GCAACAATGTACTGTATGACTGCTCGTACCACCT

>NODE_1789_length_174_cov_1.261438

CATGGACCAGCATAGACCACAGTATATAGGTTGTAAGACCGTTCTTCTTCGTGCAGTTTCTCGACGGCCG

GCATTTGTACAGCTCCAGGGAGAGACACCAGCATACGGACTATCAGATAGAAGAGCTCTTGATGTACTGC

TGCTGTACAGCCGGCATTTTATGAGCGTATAGAG

>NODE_1790_length_174_cov_1.156863

GCCTCGGGCCCGAGGGTGGCTCGTTGTAACGTATTGTCTTCCTGTTGCTGCTGCTGCTGCTTCCCGTGCT

ACCGTCCTGCCCCTCCTCTCGTCTTGTTAACTCTCACACCGATGGAATAATAATAATACTCGTGCCCCGC

CCCTCCCGATTTTTTTGTTTTTCGTTTTTTTGTT

>NODE_1791_length_174_cov_1.078431

AGATCAACGAATGCGCCGTAATCAGTAAGGTTCTTAACAATACCTTTAACTTCTTGACCTTCTTGAAGAT

TTTCAAGAAGCTCATCACGTTCAACACTGCTTTCAGATTCGATAACAGCACGACGAGAGACGACAACGTT

ATTACGTTTAGCATCTAACTTGATAACTTTAAAT

>NODE_1792_length_173_cov_1284.434211

AATTAGTATCAGTTAGCTCAACGCCTCACAACGCTTACACACCTGACCTATCAACCTGGTGGTCTTCCAG

GGCCCTACAGGGAACTCAAGGTTCCAGTGAGATCTCATCTTGAAGGAGGCTTCCCGCTTAGATGCTTTCA

GCGGTTATCCCGTCCGAACGTAGCTACCGGGCA

>NODE_1793_length_173_cov_526.894737

GATTTCGCATCCTCATTATTTGATCAGACTTGCAGAATCGTTCATCATCACTGAAGACTTCTCAGGTCAA

AGTGAAGGATTTCGCATCCTCATTATTTTATCGGACTTGCAGAATCGTTGATCATCACTGAAGTCCGAAA

AAGTTCAAAGCCGAACAAATCTTCGGAACAAGA

>NODE_1794_length_173_cov_472.440789

GTTCGACACTGAGAAACGGCATGTGCAGGGATGATCCACGAAGGCCTACTCCTGCACGCCATTTGCGTTC

AAAAATTCGGTAATTGATTGATTCTGCAATTGATACCACTTATCGCACTCTGCTGCGTTCTTCAACGAAA

TAGGAAGCCAAGTCATCCATCGCGACACGTTGT

>NODE_1795_length_173_cov_48.763158

CAGCGCACTATTAATCACTTCTCAAGGCTTGACTATACCCCCACGCCCGAGGAGTTTCATGCCAGTGCCA

CATAAACGCCAGGCCCTTTTGGTTGAGGGCCTGGGCGACTCACTTTGTCCCCGAGCAACGCCATCACAAG

GCAGCATGGGGACCATTGTGTCGCCCTACATTT

>NODE_1796_length_173_cov_7.986842

AAAACCATCGATTTTCGCTTATTTCGTGTAACACACAGGCACCCTGCGGTTTGGGCTTCTCTGGACAAGC

GAAAGGAACTACATAGAGGGACGGATCAACGCAATATTAGCAGAAGGCGACACCTGGGCTCTCGGCGCTT

GAGACGTTTCCGACCCAGAAAAGAAAATGAGGC

>NODE_1797_length_173_cov_4.375000

GGGAATTGAATTTTGCCAATTGCGTCAGACGAGTCGTGCCAAGCTTTATGAGAGAAATGTGTGTCAGTTG

ATACAGCGTATACTTCAACACCCAATTTTTGTAACTCTTCATATTGGTTTGCCATGTCTTCTAATTCAGT

AGGACATACAAACGTAAAATCAGCTGGGTAAAA

>NODE_1798_length_173_cov_4.118421

AGTGCAGAGCAACCTGTACAGGGAAGACACGGACAAGATGCGCCAGATTGGTAACCGGTGGGCCCTCGGG

TTCGTTGCGCTCGGATTCTGCGCTCTGATTGGGCACGTGACTCTGGCCACTGGGTTCGCTGTATCGGGGG

AGAGGGTAACCAGGACACTGAGGAACATGGCCT

>NODE_1799_length_173_cov_3.532895

ATATAGCGCTTCGTGGTGCACGTCTCGAGACTGGCTATTATAAGTCTGTGGTTGTTACCGACTCGCTGTC

AGCGTGCAGCGCCCCGTACTTGCAATCGTTCGCATATAGAAACGACCGAAACATGAGACACGTGGTGTAC

TGTAACACTGCTGCACTACAGTAGTGGTACTCT

>NODE_1800_length_173_cov_3.217105

TTTAATCGTAACAGATAATGGTTTTGATGCATTATTTGCTGCAAATCGAATCGCTGCTTCAGTAAGAGAA

AAAGCTAGAACACATGCATTAAGACTTGCGGGACTTATCGGTAATAGAACAGCTACTCGAGATTTAATTG

ATAAATATATTGATGCAGTACCAATGCCAGTTT

>NODE_1801_length_173_cov_3.125000

GTGCTGATCTCCACTTCATACTCCCTTGCCAGTTCCGTATAGTCACGTGCGTAGTGGTCATCGGGTGCAA

GGTCCAATACGTCGAATCCCTCCGTCGCCTGAGTGACGCCATATATCCCTGCAGCCAGGATGGCTGCGGT

TCCGAAAAGCACCAACGCCTTGCCCTTTGTCGA

>NODE_1802_length_173_cov_2.960526

GGGGAGCTACGTTGGTATTCGGGGTGTTTTTATGAGAGGGATTGGGAGAAGGGTGCATTGAAGATCTCCC

AACGAACATTCGCCGAACAATTGGCGGATGAATACGGGATAGAGTATGGTAGGAGTGTACCGCTTTCTAT

TGGCACCAAGCTTGCAGAATTCGACAAGAATGA

>NODE_1803_length_173_cov_2.828947

CTCGCGAACTCCCGCCCACGAGGTTCTGGACCTGCCTGACTCAGTCACCGTTATGGTCGACCTCCCCGGC

TGCAAGAAGCAAGACGTCGACGCACAGATCTCCCAAGACTCCGGAACCAAGACCCTGAAGATCACAGCGA

CGAGGAAAAGGCTTGTCCAGCCGCCGTCTCCTT

>NODE_1804_length_173_cov_2.736842

GATGTTGCGACACTACCCGATGAGCTGGAAGAATGGATAGATATTAATGAGGAACGTTCTCAAATTTTAC

CCGTCATTTCTGAAGCACAAGCACCGTTATCAACGGCAGAACACAGAAAAAGTGAATTAGGATTTTAATT

AAACCTTATAATCGTCCTCCAGATATTGGAGGA

>NODE_1805_length_173_cov_2.644737

GGTGGTGGCTTACTTTTTTCTGAGTTGGTCGCATGAGGAGCTCAATATCTCACCCGAAACATGACTTCGT

AGACATTCGCTCGCGCAACCCGACCAAGCCTTCCAATGCGGTAGTCCAAACAAAATACCTGTAAAATAGT

AATAGTATCCGGTGGTTATTCACGTGAAAGGCA

>NODE_1806_length_173_cov_2.598684

GTGTGGTCTGTCTGGGCGGTGAACGACGTGACCTCGCAGAAGTCATCCATCGTCGGTAAGGATGACTCTC

GGAACGTTCTCGGAATATTCCTTGATTGCTCTCGGAATGTATTGATTGTCATGCGCCCGAGATTGAATCA

ACAGAATGTTATTGTGATATTCTCGAAATATCC

>NODE_1807_length_173_cov_2.210526

TCTAAACGAATGAAGTTAGCTACAGTGATACCTTTTTCTTTCAATATTTCACCTACAGTTTTCTTAGGCT

CCATGATGAAAGCTTGGCCAGTTAGAGAAACTTCGCCAGTGAATTTTTTCATACGACCAGTAACCATTTT

CTCAGCGATTTCTTCAGGCTTACCTTCGTTCAT

>NODE_1808_length_173_cov_2.190789

TGTGTAGGCATTCGGGTTCGTAATTCGTAACTGCCAAAATCTCGCTCGATCTTGTGATCAAAGATGTCGG

TTGACGGTCAAGTGTCAAGTTTGTCAAGCGTCAAGGCTGTCTGTCAAGGTGTTTGGTGAATTGTTGTCTG

TCAAGGGGTTCAAAGTCTGACAACCGACATCTT

>NODE_1809_length_173_cov_2.105263

AGTGTCGGAATATGCTCCAGCCAACGAAAGAAAGCAACCTCACTGGCCCAATCAATCGCCTTTTCCCTCT

GTCCTAAGCAGCGACCGCCTCCACCCAGCTGAAACCGGTAGGTCTCGCGATACCATCCGGCTCGCCAGAC

CAAAACGCCTGCCAACCCAACGGAAACCGGTAG

>NODE_1810_length_173_cov_2.006579

GGTTGTAAATCTGGTACCCGGCGGGGCGGTCTAGCGAGCGGATGACGCCGTCAACGATGTCGGAGATGTA

CGTGTAGTCGCGAGAGGTAGAGCCGTCGCCGAACTGCATGATCTCCTGACCCCTGCACACCAGGTCGATA

AACTTGTAGGGAGCCATGTCGGGGCGTCCGCGG

>NODE_1811_length_173_cov_1.940789

CGCTTACAGCGTTCTCTTCTACAGTCATCGTGACATCATCGGTTGCTTTGAAGGTCCGGACCCCTTTGGG

GCCCTTCACTTCGATGGTTTGGCCGCTGACAGAGGCAGTAACGCCACTGGGCAGATCGACCGCTTTTTTA

CCAATACGAGACATCAATTTCCCCTTAGAATAC

>NODE_1812_length_173_cov_1.888158

CCTAACGTTATCACTAGTAGCTACAGGTGACTTGTGCAGGGCACAGGCTCTTCGCCTGATCATAGCAGCC

GCTTCCCTTCCGGGAGTCAACACAAGGTCAGTTTTCCAATTGACCCTCGCTATAATTTCTGCAGTTTTTC

CACAGCCCGGCACCCCATCGACTAAAGTTATAT

>NODE_1813_length_173_cov_1.875000

TATGACAAAAAGAACATTAGGTGGAACGAATAGAAAAGTTATTGCTGTTTCTGGTTTTCGTGCTCGAATG

AAAACTAAGCAAGGGTGCAAAGTAATAAACAACCGTAGAAGAAAAAAAAGAAAAAATTTAAGTATTTAGA

CAATAGATTTATAGAAAATTAATTTATATAAAT

>NODE_1814_length_173_cov_1.309211

AGCTCTCTCCTCAAGCATCTGCGGTTGTCTATTCGATATGCCTGTAAGGCTGTAAAACCGGGTTTAATCC

TATAGGGTACGTCATAAAATATGTAAAATGGAAAATGTACAAAACACAAAGCGATCGCCGTGTATCAGCC

GACAACAATACTACAAGCTTCCATCCACATCGC

>NODE_1815_length_173_cov_1.256579

GGAATCGGACAGCTGGGGTTTGTTTGGCTAAGTGGCAGCCCAAGGGTAAGGCTGACCGCACCCTTGGCCC

GGTCGGATGTGCTGGGTTGTGCGCTTGACGTGGATTCTGGAATTGCGTGGTTTTCGGTGAACGGGTTGTG

GGCGAGCGGAGTCACGATGGAATCGTTTGGGTG

>NODE_1816_length_173_cov_1.046053

ATTCCATGCAGAGGGTCGAAGCTGAAGCTTTCGCCAAGGCTGAGCCACCGGCGTGTCAGAAGTGGTGGCT

CGTGTTGGGTCTAAGTTTGGCCATGTGAATAAACTGCCCGCGCTCGCTCAGGAGGGCCACCTCCGAAATA

TCCCGAAGTAAACCAGTCGTCCGGGGGTGTTGC

>NODE_1817_length_172_cov_12.523179

CATCCAGGCTCCGGTTCACCTTGGTGTCTTCAGGGCCTCCACGTCGTCCGAGTCGTCCGGCTCCTTCTTT

GTTGCTCCCTTCGCCGGGCGAGGGGAGCCGCTGCTGCTATCGCTATTCTCCTCCCGACGGTTCTTCTCCG

GTCCCGCTCGATGTTGGCGTGAGGAGCTGGAT

>NODE_1818_length_172_cov_6.629139

AATAAACCTCCTTGTACAAGAGGCGGACGTCGACGGAGATGGCCAGGTGGACATCAACGAGTTCCTCAAG

ATTATGTATAGCTGAAGGAAAACACTCTGGGCCGACCGAGCTGCTATATTAGGTGTAGATTTTAAGGGCT

GCAACAACGCCTGCGGGTTTTCAACATTCCCC

>NODE_1819_length_172_cov_5.635762

CGAAGGCCCTAATGAACAGGCTCATCGAGGACAACCCGGCCGGGCGATCTGTCATGTGGAAGAAGGTCGG

AGAAAAAGTTGCCAAGAGCACCGGCGGCAACTACCCGAATGCCTTGGCTATCGTGGACTGCATTAAGTTC

GGCCTGACCTCGTCCAAACAGGCCGCGCTTGA

>NODE_1820_length_172_cov_5.092715

ATAGGTGTCCAAACTAATTAATCTGTGGGCGCTCAGGCGCAGCACTGGGCAAAGGCTGTCGACTGAGCGT

TCTCCTACGAGCGGATTTCCGACCACCATTCCAATTAGCTCGAGGCCTTCCAACCCGCCTGTTCCTCTAT

GAAATGTACCACCAGTGTAAAATCAAAATGTC

>NODE_1821_length_172_cov_4.635762

TCCAGGCTCTGGCTTACGGTATTCACGAATGTTCGCTGCAGTTTGACCAACTAAATGCTTATCAGCACCT

TTCAGTGTGATTTCAGTTTGGCTAGGAGTTTCAACAGTAATTCCTTCAGGAATTGCGTGACTGATCGGAT

GTGAAAAACCTAATGATAAATCTACAGATTTA

>NODE_1822_length_172_cov_3.847682

GCTTCTTCGCTGGTCTCATCTACTACATGGCGTCCGGACCGGTGGTGTGCATGGTGTGGACCGGCGTGAA

CGTGGTGCTCGAGGGGCGCAAGATGCTGGGCGCCACCAAGCCGTCCGACTCTGACATGGGCACCATCCGC

GGTGACTTCTGCGTCGAGGTGGGACGCAACGT

>NODE_1823_length_172_cov_3.708609

TTAAAAATTATTGGCAGCAGTTGGAACTTGCGGGGAATACCGAAAGGAGTGTGGCATGAATTATAAGTTC

GGGCTGGGAGGACGAAGTAAGAAATGAAAAAAGGAATGGAAATGCAAGGAAGAAATGAAAAAAGGAATAG

AAATGCGTTATGGAAATCAAAAAAAATTTGGA

>NODE_1824_length_172_cov_3.000000

CCGAGTTTTTGTCGAGGCGGGCTAAGCCCAGCGCTGGTGTCATTTTTTCCCAGTTAGATGTGTGGGACTA

TACTTCTTATACTGTTATTGTCTGTGGGTTGGTGGTGAGAGTTGATGAGTGTTTATTAGTCGAGACAGGT

GAGAGAGCCTGTTTCGGGGATTTGTAGGGGGG

>NODE_1825_length_172_cov_2.900662

CGGAGAAAAGATGGGAAAGGCCGGGCTCACGGTGCACAAGAAGTCTGAGGTGGAGAAGGTCACCAAGGAC

CCCGTGACCGGTAAGCTGTCCCTGCACAAGAAGGATGGGGAGGTGCACGACGACTTCGAGGTCATCTTGT

TGGCTGTGGGGCGAGAACCGAACGTGGCTTCG

>NODE_1826_length_172_cov_2.841060

CCCATGTGATGCTCCCATACGAAAGCCCAACGCAGCAGAAAGAGCAGTGTCTACCCATGTGAAGCTTCGC

ATGAAGCTGGCACAGCAGAAATAGCCACGTCTACTCATCTGATATGAAGCTGACACAGCAGAAATCGCAA

CGTCTACTCATCTGACATAAAGCTGACACCGC

>NODE_1827_length_172_cov_2.516556

CAGGACCCCAGTGTTGCCCCTCCAGACGTTCTGGTTGCCTTGGAACACCACAGGCTTCTCAAACTCCATG

GTGCCCTCGTTCCGGAAAAGGCCGGATACGTTGGTAACCCCCTCCGCTAGAACTTGCCCCAGGAACGTAG

CGCTCGCGCCCTGCTCAACTCTCGTAGCAAAG

>NODE_1828_length_172_cov_2.291391

AAATCTAGTAATATTAAATTTTAAAAGTTTTTTTTACATCATCAATTGCACTTTTTAAAATAGTCTCAGC

TTCACCAGTTAATTGTTTTGAAGAATTTACAATTTCCAGGTATTCTGGTTTAGAGTTTGTTATATAGTCA

CGAAGACTTTTTATATACGGTGAAATTTCACT

>NODE_1829_length_172_cov_2.251656

ATGTGACCTTTCTGCATGTGACTGGCAGTGAAACCGAAGCCTTGGCGATGGGTGACAGCGTTGCAGTGGT

ATGATATCGACGTTGCCCTATCGAAGCGTACGAATATCTTGCCCTCGGGAACAAGACCCGGCGATCGCGG

TGGTCGGAGCTATATAACGCCCATCCAACATA

>NODE_1830_length_172_cov_2.231788

CCGTTGACGCTAAGCTGACCGGGCCGGACGTGGATGTGAAGGGTAGTGGCCCTTCTCTCGGGGCGGGACT

GGCTGCGGGCGTGGCTGCGGGTGTCGGTGCTATGGCTGTCGGTCTTGGCTTGTCGGGCAAGGCTGACAAG

CCGGAAGGGGAGGTGCCGGACGTCAACCTGTC

>NODE_1831_length_172_cov_2.205298

TGCGGGCAGCCTCGTTTTTTCCTTGCTCTTTTGTTGGTGTACGTGTTTCGTCGGCAGACTGTGGGGGGTG

AAGAAAAGCCTTGTAGTTCGATGCGGAATCCGTGGCAATTTTGGCACTCAGCGTAACTTTCGGTTTTGGT

GGTTTTTGCAAGACGTGGCGCAATACGATTGC

>NODE_1832_length_172_cov_2.039735

GCCGATGATCTGGACCTCTACCTCTCTGTTGCTCCGGTCTTGCTGTACCACAGCAGTGAGTGGCAGGGCC

TTCAGAGTAAACGGCGGCGCAACGGTATACAGCAAACAACAGCAGGGGGTAGGCTTCTCTTCTGTGGCAG

AGTGTACCCGGCCCGTGTTCCTCCTGGCAAGA

>NODE_1833_length_172_cov_2.033113

GACCAATAAAAAAAGCGGCAGCAGCAACAACGCGTCCTCCGACCAATCAAAAAGCGGCAGCAGCAACTAC

ACGTCCTCCGACCAATCAAAGAGCGGCGACACTCCTTCCGACCCATACAATAAAAAAAGCGACAGCAGCA

TTTCCTCCGGCCAATAAAAAAGTGGAGAGATT

>NODE_1834_length_172_cov_1.874172

CACCATAATCACCAACGTGGGGGGTCTTGAGGGTCATTCTTTCGTGGCCATTTATGACGGGCATGGGGGG

GCGCTGTGCGCTGCTTACGCGGGCGAACACATGATGCGGCACATCATGGAAACCGAGGAGTTTGCGGCCT

ACGCGGAGTCGGCCGAGAAGGACACCGGGTTG

>NODE_1835_length_172_cov_1.754967

TGTTGTTCCGTCCGCCATGCTTGCTAAGACCCTCTGTGAGGGCTTTAACTGGGCGGCCTTTCCAAAGCTC

CGAACGGTCAATCAGTACCAGCCCGCGCTGGCCTGGCGTCGTCGGTTTGTACGACTTGAGTGCCATGTTC

TCTGTCTTCCGTTTGCTTGTATGCGGCCCCGT

>NODE_1836_length_172_cov_1.549669

GCTCCATCCGAAATTAAGGGCGGACACGCGCTACGATAGCGGGTGCACCTGCTTGTGTCGTGGGCCACCC

ACTCGAGGAGTTCCATCCCATTACTGTAGCAATACTATACCAAAACTGACGCTCGTGAAAACATGGAAGT

GCATCCCGATCGCCTAGCCTAGGCGTACCACG

>NODE_1837_length_172_cov_1.529801

CCCTGATACGGATGCGGATATGGAGAAATGGGTTGAGTTCAATCGCAAGTACTCCGAGCTGTGGCCCGTG

ATCGTTACCAAGAAGGACCAGCTGCCCAACGCCGAAGAGATGGACGGCAAACCGGGCAAGATGGATCTGT

TCTCCGATGTACCTGGCGAAGGCGGTTAAGCC

>NODE_1838_length_172_cov_1.423841

GCCTGTCATGACCACCACCACCGCCACTAGTCTATATATACGCGTCGCATCGGCGTTTGGCGTACGGCGC

GATGTCGGTATCCAAAGCAGTCACGGTGAAGACCTTCTTGAACAGCCCAATGGAATAAACCTTGTCCTGG

AACTATTGCCACTGCATGCCTTCGCGCTTGAT

>NODE_1839_length_172_cov_1.350993

GCATCGGGGAGGGACGATGAGGTGGATGGGGTGGTTGGCATAGACCTAGAGATGCAGGGACGTTGTAGCG

CGGCTACTGTAGCTGCACCCCGTTTGGAAAATCAAGGAAATTTGGGAACGCTTAACATTTTCGATACTGT

AATGTTACAAAGGGGGTGGGCCATAGTCTCCT

>NODE_1840_length_171_cov_567.193333

CGCTCAACGGATAAAAGGTACTCCGGGGATAACAGGTTTGGTGATCATAGCGTAAGCAAAACACCCGGTC

CCATCCCGAACCCGGCAGTTAAGTGCTACTGCGCCAATGGTACTGCGTCTTAAGGCGTGGGAGAGTAGGT

CATCGCCAAACCTGGAATTCTGGGTGCCAAG

>NODE_1841_length_171_cov_12.460000

GCTGCTGCTGCTGTTGCTGCAGCTCCTGTTCTTCCTGCCGCCCAAAATCCTCGAAAACCATCTCCAGCTG

CTGCTGCTGCTGTCCATAATCCTCAAAATGAAAGACCATCTCCTATAGTCAATCCTCCCCGTACCAACTC

CCCCTCTCACACAGTATAGCGCTTCAGCGAG

>NODE_1842_length_170_cov_82.147651

AAATCAAATTAAAACTTTCAACAACGGATCTCTTGGCTCTCGCATCGATGAAGAACGTAGCGAAATGCGA

TACTTGGTGTGAATTGCAGGATCCCGTGAACCACCGAGTCTTTGAACGCAAGTTGCGCCCGAAGCCTTCT

GGCCGAGGGCACGTCTGCCTGGGTGTCACG

>NODE_1843_length_170_cov_24.812081

TTGGCACCCAGAATTCCAAAATGTACGGCTCTTCTAGTGTCTCCTCGCTGTAGATGATCAGCAAGGCTTC

CGGAAATCGGATATGACGAAGACGCCGGTTTATGTCGCAAACGAAAAAAACTTCCTCCATCGTCGCCCCG

AAGGACTAGGGGAAGGGAGTGTGCTTGCTT

>NODE_1844_length_169_cov_429.344595

TGGAGTTCCTTGGCACCCCGAGAATTCCACGCCTCGATCTCGTACTTCTCGGACGCTTCGGAGTGCTCGT

CCACATCCACCTTGATAAAGACCACATCCTCCATGGTTTCGGACAGCTTGGCCAGGAACGGGGCGACTCG

CTTGCAAGGGCCGCACCAAGTGGCGTGGA

>NODE_1845_length_169_cov_18.195946

TAGTGGCGAGCGAACGCGGATTAGCCCAAACCAATGTTGTTACGGCAATGTTGGGGTTGTAGGACCACGA

CATTTTATGCACAAGGAATTAGAATCTACTGGAAAGTAGGACCATAGAGGGTGATAGTCCCGTATAGGAA

GTTTTGATGGACGTATTAAGTAGGGCGGA

>NODE_1846_length_168_cov_10.034014

GGTCGAGCGTTTTCAGCGCCTGCAGCTTCCCGAGCTCCGAAGGGATGCTCCCTTCAAGCCCGCATCCTGA

AAGCTTGAGCTCCTCCAACGCTTTCAGGTCCCCCAGCTCGGGGGGTATGACGCCTCCGAGCGATGGGTTC

CTCCTCAGGTCGAGCGTTTTCAGCGCCT

>NODE_1847_length_167_cov_1189.239726

CTCGTGCATGTTTGCGCTCTTGGCGCAGGTGAGGCAGAATCGAACTGCTGCGACGCTAATCGCCTACGGA

TTTGAAGTCCGCACCCCGCACCAGCGAGGATCACCCCTGCGGGATGTGGTGCGATCGACGTGGTATGGCC

GTTCCTTCAAGCCGCTACTCAACCACA

>NODE_1848_length_167_cov_190.684932

TCCGTCGCTGAAAGTTTTAACGCTCAGCGTCGCAGAATAAACTTCAGAGAAATGCAGTAGGCTGCAGCAA

ACCATTTTGGTCATTAGAGCTCAAAAGCATTCAAATCAGGGACGAATAATCGCCATTTTGTTGTGTCATC

AACGCGCACAACAAACTGACCGGTCCA

>NODE_1849_length_167_cov_176.075342

TTCCACTCGTTCAGAGTTGGTCGAGCTAGGTGCGGCAACGCGTGTGTTTTACAGGCACACATGCCTTTTT

TTGAATTTCGGGACGTAAAACATCATAACGGTTTTTTTATAAAAAAAAAAAAATATTTTATATTTTTTTA

AACAATAATGAGTTTGATCCTGGCTCA

>NODE_1850_length_166_cov_113.434483

ACAGACACTCCGACAAGCATGGTGGAGATAAGCAGACTCGAACTGCTGACATTTGCCTTGCAAAGGCAAC

GCTCTACCAACTGAGCTATACCCCCAAATGGGCCATCCTGGATTTGAACCAGGGCCCTCACCCTTATCAG

GGGTGAGGTTAAGAGCTCGAGACAGA

>NODE_1851_length_165_cov_2518.263889

CTGAAATTCTACCACCCTCTATCACACTCTAGTTTGCCAGTTCGAAATGCAGTTCCCAGGTTGAGCCCGG

GGCTTTCACATCTAGCTTAAATAACCGCCTACGCGCGCTTTACGCCCAGTAATTCCGATTAACGCTTGCA

CCCCTCGTATTACCGCGGCTGCTGG

>NODE_1852_length_165_cov_512.395833

CGTGATAAGCTGCGAAAAGCTACGGGGATCTGCACACAAGAATTGATCCGTAGATATCCGAATGGGGCAA

CCCGGTATATTGAAGATATATCACCCGCAAGGGGGCAAACCCGGAGAACTGAAACATCTAAGTACCCGGA

GGAGAAGAAAACAAAAGTGATTCCG

>NODE_1853_length_165_cov_24.972222

GGCACCGAGAATTCCAGTTCAGCCTGTAAGATCTCGGTCTAATGTAATCCGGATTTACCTAAATTACAAC

CTACATCCTTCGACCAGCTATTCCATCAGCCAGCTCGATTAACTCTATGCGTCCCCACATCACGCTCTAT

TATAAGTATAGGAATATTAACCTAT

>NODE_1854_length_165_cov_21.673611

TTCCAAAATTTTTTTTTTTTTAGTCAACAATCCTCGATTCCTTTTTTATGTGCGACACGAAGGAAGTTCA

GTTTTTTTTTTGTTTCTGTAGTTGGTTTTCTAGATTTATTTGTTAATTGAGTTTCTTTGAAGCATTGTAT

TAGTAGTTTAAGTTATGATCTTTTT

>NODE_1855_length_165_cov_10.861111

CGACAGTTCCATCACCTTGACCGCCTGTAGGTTGCCAAGCTCCGCTGGGATTGTTCCGCTTAGTTGATTA

GTATGAACACGCAGGTCCTCAAGTAGAGACAGCTTACCCAGCTCCTTAGGGATCGTACCGGACAGCTGGT

TGTCCGACAGTTCCATCACCTTGAC

>NODE_1856_length_165_cov_1.791667

TCCCAGCTCCGGCGGGATGGGACCGGTGAGCTTGTTGCCCCAGAGTGAGAGTGTCTCCAGCTTGCTCAGA

CCTCCCAGCTCCTTCGGGATGCTTCCTGTGAGCTGGTTGTCGTCGAGCCACAGCCCCTTCAGCTCTCTGA

GAGCTCCCAGCTCCGGCGGGATGGG

>NODE_1857_length_164_cov_6614.307692

TGGCCCCGAGAATTCCACCTATACAAGATCGAAAGCAAGAGCTCAGGATCTAGTTTCGAGACCTGTTCGG

TAGGTCCTCTCGTCATCTGTCAGCCAACTACGGCTTTCTCAGAGAGATTCCCATTTCGTGACTCATCAAC

GCGCTCTGTACCTCCGCGCTATCG

>NODE_1858_length_164_cov_93.727273

GAACTGCCGACCTCACCCTTACCAAGGGTGCGCTCTACCAACTGAGCCATATCAGCATACGATTTAAAGA

TTGGAGCGGGTGGCGGGAATCGAACCCGCTTCATCAGCTTGGAAGGCTGAGGTCTTACCATTACACAACG

CCCGCAGCAGAGTCGTTCGTGGTA

>NODE_1859_length_163_cov_847.838028

TGGCACCGAGAATTCCAATTTTTGGGACGACAGTGATAACTTAGATGATGAGTGGTGGAAGCAGGAAAAT

ATTGGGGTGAGTGCAGAGGCGGAGGAGTCGATGGCGAAGCTGCCGGGAATTGAACCCGGATCTGATCCGC

GACAGGGATCAATTCTAACCATT

>NODE_1860_length_163_cov_179.535211

GTGTTTTTGATAAACAGTCGCCTGGACCTCTTCACTGCGGCCCATCCGAAGATGGGCGACCCTTCTCCCG

AAGTTACGGGTCTATTTTGCCTAGTTCCTTAGCCATGAATCTCTCGAGCACCTTAGAATTCTCATCCCAA

CTACCTGTGTCGGTTTACGGTAC

>NODE_1861_length_163_cov_129.640845

AATAGATATATTAATGATACTATAGAGTTCACTTATCGGTTAAGTACGATATAAAAAGCATCTTAATTGC

GGACGCGTAGCTCAGCTGGATAGAGTACCTGACTGCGAATCAGGGGGTCGGGGGTTCGAATCCTCCTCGG

TCCGCCAGGGGAAACTCTGGTGG

>NODE_1862_length_162_cov_84.354610

TGATAAGCGCTGAAAGCATCTAAGCACGAAGCCAACTCCAAGATGAACTTTCCCTGAAGATCCCAGCAAG

ACTAGCTGGTTGATAGGATAGATGTGTAAGTGCAGTAATGCATTCAGCTGACTATTACTAATAGATCGTT

TGGCTTTGGAATTCTCGGTGCC

>NODE_1863_length_162_cov_12.099291

GTAAAGCGAATGATTAGAGGCATCGGGGGCGCAACGCCCTCGACCTATTCTCAAACTTTAAATAGGTAGG

ACGGCGTGGCTGCTTTGTTGAGCCGCGCCACGGAATCGAGAGCTCCAAGTGGGCCATTTTTGGTAAGCAG

AACTGGCGATGTGGGATGAACC

>NODE_1864_length_160_cov_2459.316547

CTTGGGGGAGATCAGCCTGTTATCCTTAGAGTAACTTTTATCCGTTGAGTGACGACTTTTCCACTCAATA

TCGCCAGATCACTAAGACCGACTTTCGTCTCTGTTCGACTTGTAGGTCTCACAGTCAAGCTTCCTTATGC

CTTTACACTCTTCGATCGAT

>NODE_1865_length_160_cov_534.719424

GCAGAGTAGCTACGTTGGGAAGGGATAAGCGCTGAAAGCATATAAGCGCGAAACCCACCACAAGATGAGA

TTTCTTTAAAGGGTCGTGGGAGACTACCACGTTGATAGGCTATAGGTGTAAAGGCAGTAATGTCATAGCC

GAGTAGTACTAATAACCCGT

>NODE_1866_length_160_cov_294.899281

CTCTTCGGAGGTACTCAGAAAAGCTAGATTTTTAAACTGAAGAGTTTGATCCTGGCTCAGGATGAACGCT

AGCGGCAGGCTTAACACATGCAAGTCGAGGGGTAACAGGAGTAGCTTGCTATTTGCTGACGACCGGCGCA

CGGGTGCGTAACGCGTATAG

>NODE_1867_length_160_cov_192.517986

CTCCTACGGGAGGCAGCAGTGGGGAATCTTAGACAATGGGCGCAAGCCTGATCTAGCCATGCCGCGTGTG

TGATGAAGGCCTTAGGGTCGTAAAGCACTTTCGCCAGAGATGATAATGACAGTATCTGGTAAAGAAACCC

CGGCTAACTCCGTGCCAGCA

>NODE_1868_length_160_cov_16.589928

CTGGAGTTCCTTGGAACCCGAGTATTCCAAGTCCTTCCTCCTCCTTTCTTCTCTGTCTTCTTCTTGTTCT

TAGTTATTAATAGTTGTCTTCTTTCTTCGCCTCTCCTCGTGTATATAGCCTCTCCTCCTGTTTTGTTGCT

GTTGTTGTTGTTGTTGTTGT

>NODE_1869_length_159_cov_2771.463768

TTGGCAACCCGAGAATTCCAGTCCCAACATGCACGCTCACACTCAAACCTCGCCCCCGGAGGGACATGGT

CGGTCGATGGTGCCACCTTGCGGCTTCCCACCTGCATTCACTTTCACTGCGCACGGGGGTTTGAACACCC

CTGTACTCGCGGGCATGTT

>NODE_1870_length_159_cov_381.282609

GACTGTTTATTAAAAACACAGGTCTCTGCTAACTCGTAAGAGGATGTATAGGGACTGACACCTGCCCGGT

GCCGGTAGGTGAAGCTTTGATGTCTACGCATTGAGGTCAAACCCCGGTAAACGGCGGCTGTAACTATGAC

GGTCCTAAGGTAGCGAAAT

>NODE_1871_length_159_cov_154.289855

GGCCCCGAGAATTCCATGGTGAGCTAGGAGGGATTCGAACCCCCGACACCGTGGTTCGCAGCCACGCGCT

CTAGTCCACTGAGCTACAAGCCCTATATGAATATAATACATTACTTTACTATTTCATAATATAAGAAACA

CTCATAAAATTCACTTATG

>NODE_1872_length_159_cov_117.173913

CGGCGGACGGGTGAGTAATGCATGGGAATCTGCCCGATAGTGGGGGACAACCTGGGGAAACTCAGGCTAA

TACCGCATACGTCCTACGGGAGAAAGCAGGGGATCTTCGGACCTTGCGCTATCGGATGAGCCCATGTCGG

ATTAGCTAGTTGGTGAGGT

>NODE_1873_length_159_cov_48.891304

TGGAGTTCCTTGGCACCAGAGATTTCCAAAATGTTGGCCAGGTACATTCCGCATTCAGGCACGTGAAAGC

CCGGTGCGGCCAAGCCCGGCCGGGGGAGCCAGACGAACGAATGTTAGGCGCCCCCCCGGATTGTGTCTCG

CCTCGCGTAAAACAAGTTA

>NODE_1874_length_159_cov_37.094203

TGGCTCCCGAGTATTCCAGGGGATCAGATTACTAATGATATTAAAAGTCGTCCTGCTTGTGCGCCCCTCG

CAAAAAAAAACCGCGAACCAAAACAATCCCCCCCGCACACACACCGGCCGCTCGCCGGAGACCGTCACAT

TTACGCCCCGCATTCAAAA

>NODE_1875_length_159_cov_29.731884

CCCAGCGGGCACGCCATCAACCTCGGCGGTTACATCACCGCGCTCAAGGCATTCGTCGGATTTGAGGCCG

GGCAAAAGTGCTTCCCCAAACCCGTAGTCTCCAACGACGCAGGGACTAAGAGGATTGCCTTTGTGCCTTC

CGGAACGGAGTTCCTGGAC

>NODE_1876_length_159_cov_28.355072

GGTGTTGAGTAAACACCAGGTGTAGTTCAGGGAAGCCTCAAGATAGAGGGACCTCCAGATGATTCGAGCC

CATAAGTTCGAGAACGCGAAGCAACGCTTACGAGCGCTCACCGCGGATGCGGCGGGCCAGCTGGATGTCC

TTGGGCATGATGGTCACGC

>NODE_1877_length_158_cov_69.635036

GTAGTAGACTATACTGCGTTAAATGATGAAGCCAAATGATAAAAATATGAAAGTGTATAGAGTTAGATCC

TTTGCATACGACTTAAATACAAAACGGGGTATTGTAAGGAGTAGAGTAATCTTGTTATTACGATCTTCTG

AGATTAAGCCTCTTGTTC

>NODE_1878_length_158_cov_39.861314

AGCGCCTTGGTATTCTCTACCTGTCCACCTGTGTTGGTTTAGGGTACGGTCTTAGGGGACGCTATTTCCA

GGGACTTCCAGGCTGCTGCTCCAATCCAATAAGGAACAACAACTTTCGAAATCCGTCACACTTCCCCCGG

CCCACGAATATTAACGTG

>NODE_1879_length_158_cov_18.598540

CCACAGAGATGGTGCGGGCATATGTTGACGTCTTGAGAAGGAACCGGACCCGCGCTGGACAGGAGTTAAC

GCGGACATCTGTTGATGTCTAGGACGTAACCGAGAACAAGATGTACCCGGCACCCGACTTGAGTTCGTGC

GGGCATCTGTTGACGTCC

>NODE_1880_length_157_cov_48.845588

GGGGAGCAGGAAGGTCTTCTCGCCGTCTGGGTTCTGTGCCGCATTGTGCGTCACCACGCGGGAGAGAAGA

TCAGTGTTCCCAAGGATCGATGCCGGCAACGGGGTGATGACATCCCCCGTGAGCGTGTTGTGCAGCAGGC

CACCGTACGCCTCGGGC

>NODE_1881_length_156_cov_821.022222

CTCGGTTGATTTCTTTTCCTAAGGGTACTTAGATGTTTCAGTTCCCCTCGTTTGCTTCGCTTAGCTATGT

ATTCACTAAACGATGACACTAAGTGCCGGGTTGCCCCATTCGGAAATTCCAGGATATAACGCTTGTTATC

AACTCCCCTGGACTTA

>NODE_1882_length_156_cov_146.437037

CATAACGTCTACGGACCAAAGAGGGGGATCTTCGGACCTCTCACTATTGGGTGAGCCTAAGCCGGATTAG

CTAGTTGGTGGGGTAATGGCCTACCAAGGCAACGATCCGTAGCTGGTCTGAGAGGACGATCAGCCACACT

GGGACTGAGACACGGC

>NODE_1883_length_156_cov_63.222222

GGGAGCAGGAAGGTCTTCTCCCCGTCTCCGTTCATCTTTTGGTTGTGCACCTCCACTCGGTTCAGCAGCT

CCGTGTTCTTCAGGATGGAGTCGGGCAGCGGGGTGATGACGTTCTTCATGAGCGTGTTGTGTACGAGGCC

ACCGTACGCCTCGGGC

>NODE_1884_length_156_cov_57.022222

GGGGAGCAGGAAGGTCTTCTCCTTGCCTCCGTTCATCTTCTCGTTGTGGGGGCCCAACTTTCGGAGCAGA

TCCAGGTTGTTCAGGATCGAGTCCGGGAGAGGAGTAATGACGTCCTTCATGAGCGTGTTGTGGACCAGGC

CACCGTACGCCTCGGG

>NODE_1885_length_156_cov_39.814815

AATGTATTAAGATCCATTCGTCCGATTAGTATTACTCGGCTGAGAGTGTCACCACTCTTACACCTGTAAC

CTATCAACCATGTAGTCTCCATGGGGACTTATTGCTCGAGGCAGAGAAATCTAATCTTGAAGTTAGCTTC

CCACTTAGATGCTTTC

>NODE_1886_length_156_cov_26.844444

GCATACTTCATTTAACAACCGCATCCACAACTCGTGTCGTACAGTTGGGCGACATCACGCTCGGGGTGCC

TCCGTATATCTTGATATGTCATATTCCAGTCATTGCCGTCTTCGGTGGCAAACTAGTGGTGGCGACCGTC

TTCGCAGCACAGGTAG

>NODE_1887_length_155_cov_179.358209

ATCTTATTAAGAAAAGGTCATTCGGACGCGGGATGGAGCAGCCTGGTAGCTCGTCGGGCTCATAACCCGA

AGGTCGTCAGTTCAAATCTGGCTCCCGCAACCAATTTTTATTATTAAAAATATAATGATACAACTAAAAA

AATACACACCAAAAT

>NODE_1888_length_155_cov_32.977612

TAGCCTGGTGACCATAGCGAGTCGGTACCACCCCTTCCCATCCCGAACAGGACCGTGAAACGACTCTACG

CCGATGATAGTGCGGATTCCCGTGTGAAAGTAGGTCATCGCCAGGCTTACAGCTGGACGACGGTACCTTG

GAATTCTCGGTGCCA

>NODE_1889_length_155_cov_18.044776

GTGGCGGACGGGTGAGTAACATTTAGGAATCTACCTAGTAGTGGGGGATAGCTCGGGGAAACTCGAATTA

ATACCGCATACGACCTACGGGTGAAAGGGGGCGCAAGCTCTTGCTATTAGATGAGCCTAAATCAGATTAG

CTAGTTGGTGGGGTA

>NODE_1890_length_154_cov_1150.330827

ATGTAGTTGTGATTGGAGTTCCTTGTGACCAGAGAATTCCAGGTCAGTGTGGATTCACGCGAGGTGTAAC

AAGGTATTTGGGGCCCTCGCGTCTTCTCATTTTTGTGAGATTGATCCGCTCATCATAAAAATGAACTCAG

AAGAGCCATTACCT

>NODE_1891_length_154_cov_471.669173

GGGAGTACAATTTGCGTTCAAAATTTCGATGAATCACAAGCCTGCAGATCACCATAGTTATCGCAGTTTG

CTGCGTTCTTCATCGAGTAGCAAGCCTAGACATCCACCACCTACGGTCGTAAATTTTCTATTTTTTCGAG

TTGAATTTTCACGG

>NODE_1892_length_153_cov_1493.462121

ATAGGTCAGGTGTGTAAGGGTATTCCGTGGCCAGATGTGTGTTGGTCTATGATCGAAATTGTGAAAGAAC

CTGAATGGTTACCAACTACTTATGTTGACCATCGTTTTACCACACACCCATTGACTGAGACCATGGAATT

CTTCGGGTGCCAA

>NODE_1893_length_153_cov_59.734848

TCAGAGCCAATCCTTTTCCCGTTGCCCCAAGGGGCATCAGAAATTGGTGGCAACCGTTAGGTTATCCACA

CGGGATTGGAGTAGTTTTCAACTAGCAATCACCACGCATCAGCAAAGCTTCACGCACAATGGTATTGCCC

GAGGAGCAAAGAT

>NODE_1894_length_152_cov_1235.885496

ACGGGTAACGGGGAATTGGGGGTGAGATGGCTGAGTGGCTTAAAGCGTTAGATTGCTAATCTATTGTACA

AAGCTCGTGTAGCACAGTGGTTAATGCACCTCTCTTGTAAAGAGGAGATCGTGAGTTCGATTCTCACCTC

GAGCTCCATGGA

>NODE_1895_length_152_cov_553.198473

TCGAACGGGGTGAAAAAAGTGATCTCCGATCCTCTCGCATCACGAATTTTACCCTCGCCTTTTACTAGGA

TAAACCGTGCTGAGAGGATCCTGAAGGAGTTCTTTCGATTTTTGACCCCACTTGTGGGGTATGGAATTCT

CGGTGCCAAGGA

>NODE_1896_length_152_cov_217.389313

ACTAGGGGCAACTCCTCTCAAGTATCCTACACCCACGGAAGATAGGGACCGAACTGTCTCACGACGTTCT

AAACCCAGCTCACGTACCTCTTTAAACGGCGAACAGCCGTACCCTTGGGACCTGCTCCAGCCGTACAAAG

GGCAGGGACGTA

>NODE_1897_length_152_cov_98.496183

TGCTTGTCGGAGTGTCTGTTTCGAACGCGACTCTGAGCATGTCTCGACATAGCTCAGCCCGGTAGAGCGA

CAGACTGTAGTGGTCACCGTGGTTATCTGTTGGTCCTTGGTTCGAATCCGAGTGTCGAGACCATGGAATT

CTCGGTGCCAAG

>NODE_1898_length_152_cov_62.022901

TCCCCTTCCAGGGAATTGCCACTCGAGTTTCTCCTCCTCTCCATCTGCTCACCACAACCCCATTCTTTTG

AATGAATGGGTTTCAGGCTTGGACTCCTTCCTCTTCATTCGCCATTACTGGGGAAATCCTCGTTAGTTTC

TTTTCCTCCGCT

>NODE_1899_length_152_cov_50.488550

AGCAGGCAGCCCTTCTCTCTGCCTCGCCTTTCCCCTGCTATCGTAAAGAAACGAACTCCGAAAACCTCAA

GGCGGAATTCGGCACCTACTACTGCGTTCGGCTGCAATCAACGTCGTGCACATCCGCAGCAATGAAGACT

TTCGTGGGCTTG

>NODE_1900_length_151_cov_280.638462

ATTTCAAGGTTGGCTCCACAACATCTAGCGACATCGCTTCAAAGCCTCCCAGCTATCCTACACAAGCCAA

TTCAAAGTTCACTGTGAAGCTGTAGTAAAGGTTCACGGGGTCTTTCCGTCTAGCCGCGGGTATACGGCAT

CTTAACCGCAA

>NODE_1901_length_151_cov_104.930769

CTTTTGTTTTCTTCTCCTCCGCCTACTTAGATGTTTCAGTTCAGCGGGTTTGCCCACCTATCGGTGTACT

ATGTCTTCAACATAGTGGGTTGCCCCATTCAGGTATCTACGGATCAATCGATGTGTGCTCGTCCCCGTAG

CTTTTCGCAGC

>NODE_1902_length_151_cov_61.784615

GAAAGCATCTAAGCGGGAAACTTGCCTGAAGATGAGGATTCCCTGGAGACTTGATCTCCCTGAAGGGTCG

TTCGAGACCAGGACGTTGATAGGCTGGGTGTGGAAGCGCAGCGATGCGTGGAGCTTACCAGTACTAATTG

CTCGTGGAATT

>NODE_1903_length_150_cov_246.224806

GGACAGATACTGACGCTGAGGTACGAAAGCGTGGGGAGCAAACGGGATTAGATACCCCGGTAGTCCACGC

CGTAAACGATGTCTACTAGAAGCTCGGAGCCTCGGTTCTGTTTTTCAAAGCTAACGCATTAAGTAGACCG

CCTGGGGAGT

>NODE_1904_length_150_cov_18.534884

TTGACCTACGCCATACTGGTCCTACTGTTGCGAGTTCAATTCAAACGAAGGTTTGATATGGCGCCATGGT

TCGGTGGTTTGACCTACGCCATACTGGTCGTACTGTCGCGAGTTCAATTCTCGCTGCGTACATTTTGGAA

TTCTCGGTGC

>NODE_1905_length_149_cov_24.187500

GGCTTGGCTTGGCTTGGCTTGGCTTGCTTTTGGCCTGGCTTTTTTTTTTGGCTTGGGTTTGAGTTGGGTT

GGAGTTTCGGGTTGGAGTTTCGGGTTGGGTTCAGGCTGGGTTCGGGTTGTTATAGGTTCGGGTTGGCGAC

GCGTGTGCT

>NODE_1906_length_149_cov_13.968750

AAGAGAGACTTGAACTCTCACGGCCTAAGCCACAGCCCCCTCAAGGCTGCGTGTCTACCAATTCCACCAT

ATCCGCTTTTTAAAAATACTACTCTGGGATTTGAGTAGATGTGTCAACTTCTACTGGTACTTCAGTAGTT

TGTTCAACA

>NODE_1907_length_148_cov_130.007874

AGTTGCAGTACAGCACCGGCTGCTTGTACGAAGAAGCGCGCCTCCCGTGGATGCCTTTCTCGGTGTTGTC

AGCGGTGGCCTGGGACGATCGGACATGGGCACCAGCCGTGACGCCGAGAGTCACCAGAGCCATGACGAAG

GGAACAAA

>NODE_1908_length_148_cov_29.023622

GGCACCGAGAATTCCAAGAGATCAGACAGCGGCCTAGAGGCGCGCGGCGTTTCGGCAAGGCTAGGCGCGT

GCTCGCGAGTGGGTTCAATTACCCGCGACAAACGCGCCGCCTAGGCTATCTAGGTGCCGATATTGCATCA

AGTCTCTC

>NODE_1909_length_148_cov_22.070866

TGGCACCGAGAATTCCACGCCAGCTCCACCCTAGGTATGCAACTTCAGCTTACAACGCGCCCGCAAATGC

GAGAGCTTTACAGCTTGAGCTCTTCCGTGTTCGTTCGCCACTACTAAGGGAATCTCTGTTGATTTCTTTT

CCTAAGGG

>NODE_1910_length_147_cov_17036.587302

CTTCCGTTTGTCTAGTTCCTTCTTTGCGAAGGAAGGCTGGATGTCGAGATGAGGAAGTAAGATTGTTCAG

AAAAAGCACACTTTGGTTCGACCAGGTCGTGCACGTTGAAGACTCCTCAAGCATTCACTTTTCCGATCGG

CTGGAAT

>NODE_1911_length_147_cov_175.015873

ATATAGGGAGGAAAGAAATGGAAAGCACCAAGAGGAAGTGGAAGCCATATGCAAATAAGGTTGCTACTCC

CTCACTACTGGATGCCATCGTCTTGAAACCTATGAAGATCCGGCGTTTGATGTAGTTTCTATGTCCGACG

GTCGCGA

>NODE_1912_length_147_cov_128.896825

TGGCAGCCGTGAATTCCAAAGCCTCTTGACAGGTTTTGGCATGTTTTTTGTTCCAGTTGGCGGGCCCAAG

AATGGCGCGAGAACGCCAGTGCAAGGAGTCCCGCTGGTCAAAGCCCCTGGAGAGGTCATGTTTGTCATCA

TCGGCCA

>NODE_1913_length_146_cov_3428.784000

TGGTGAATCATAATCACTTGCGGATCGCACGCTTCGGCGGCGACGTTTCATTCAAGTTTCTGCCCTATCA

GCTTTGGATGGTAGGGTATTGGCCTACCATGGCTTTAACGGGTAACGGGGAATTGGGGTTCGATTCCGGA

GAGGGA

>NODE_1914_length_146_cov_219.848000

TGGCACCCGTGTATTCCAACGAGTCAGGCTGTTTTTTATCCTTGGTGTACATCATTTATCCCACGGGGAT

AACCGGGGACTCTTGAAGGGGAATCCATTTAAGATCGTCATTCCCGACTCGTCCCCTTCGCCAGTTAACC

CCCACC

>NODE_1915_length_146_cov_176.104000

TAGTTGGTCGTCGCTGAAGCTGCGTGATGATGACTTTATTCCCATAATCATTCGCAGTGAATCAGCCGAC

GCTCACACACCCTGCCGGGCAGTGTTCGAGCCTGGGGCCCCAACTTCCTTGGATGTCTGAGCGCTTGGAA

TTCTCG

>NODE_1916_length_146_cov_103.424000

GTACAGGAATATTAACCTGTTTTCCATCGACTACGCCTTTCGGCCTCGCCTTAGGGGTCGACTTACCCTA

CCCTGATTAACATGGGATAGGAACCCTTGGTCTTCCGGCGGGGAGGTTTTTCACCCCCCTTATCGTTACT

CATGTC

>NODE_1917_length_146_cov_25.296000

CCGGGGTCACAAAGTAGGCTCATATGTGTGTGTGTGTGTATTCTCTCACCCATCCTTGCGAGACGACGAA

ATGTTATTGTCATTATCGTCATCTTCATCATCATCGTCATCGTGGATCGGAAAAAACTACAAGCAGTGTA

TCAAAA

>NODE_1918_length_145_cov_1189.379032

GGCACCGAGAATTCCAAGTTTGTTTAGCCTTTCACCCCTATCCACAGCTCATCCCCTAACTTTTCAACGT

TAGTGGGTTCGGTCCTCCAGTACGTGTTACCGCACCTTCAACCTGGCCATGCCTAGATCACTCGGTTTCG

GGTCT

>NODE_1919_length_145_cov_647.911290

CACGAGAAATCCGGATTGAATCTGGGAGGACCACCTTCCAAGGCTAAATAGCATTCGAGATCGATAGTGA

ACTAGTACCATGAGGGAAAGGTGAAAAGAACCCCGGGAGGGGAGTGAAATAGAACCTGAAACCGCATACG

TACAA

>NODE_1920_length_145_cov_71.588710

TAGATTAAAGATCAAATTCTAAGAAAAACTCCTGTAAAAATCAACTCCAAACAAACAAAAATCACAAATA

CATAAATAATCAAGAAAAATATCTTACAAATTAAAAACCTAAAAACCAACATATAAAAAAACATCAAAAT

AAATA

>NODE_1921_length_145_cov_7.088710

TCTTGTCCCGCCCTAGTTTAAGTCAGTCGAGACCCCCCAAGTTCGATAATATAGAAGCCCACTTTGGTAT

GTGGCGCAGTAAGTTTCTGGCTTACATGTCATCGCTTGGTTGCCTGCACGTACTCAAGTTGACAGACAAC

CCGAT

>NODE_1922_length_144_cov_10745.178862

TGGCACCCGGAATTCCAAAAGCTCAGAGTCTGGTTAACATGGATCGCTCATTCCAGGGGCGTAACCTCCC

CGCGCATTGCTGCGCGGTCGGGCCCTTAGCCCCCTTGTTTTCTTCGGTGTTTCAGTTGGAATGTAGTTTT

CTCA

>NODE_1923_length_144_cov_2070.170732

CTAGAATTCTAACCCTGTATCGTTAGCCGGTCAGGGGACAGTTTCAGGCGGGCAGTTTGACTGGGGCGGT

CGCCTCCTAAAAGGTAACGGAGGCGTACAAAGGTTTCCTCAGATCGGTCAGAAATCGATCGAAGAGTGTA

AAGG

>NODE_1924_length_144_cov_400.365854

GTGAGGAAAGGATGTTCGGGGATGATTACACCGAACTTCCTGGAGTCCCATCTTGGCCCAGTTCCTGGGC

CACGGGACGATGCCGGGCTATAAATGTCATAGTTACACTGACTCAATCCAATTCTCGGCAGCCAACGAAC

TCCA

>NODE_1925_length_143_cov_18379.409836

CGGATAATTGGTTTATGGGGTTGCCTGAACAGGCAGTGCCCCGAAGCTACCATCCGTAGGATTATGACTG

AACGCCTCTAAGTCAGAATCCATGCTGAAAAGTGACGATGCACTTGCTCCCTATGTCACGCGAATAGCGA

TAG

>NODE_1926_length_143_cov_54.131148

TCTTGGAAGCATGGCATCAACCACTTCACTCTATAAAGAGCTCGTCATCACGTCTCAGGTAAAGCGACCC

CGGATTTGCCTAAGGTCACACCCTACTCGCTTAAACCGGACACCAACTGCCGGCTGGCCTAGCCTTCTCC

GTC

>NODE_1927_length_143_cov_29.672131

GGCACCGAGAATTCCAGTTTTTTTTAAGCTTCCACGGCGTCCTTCAGTCTCCTTTCACCACCGCCTTAGT

AGCAGCAGCAGCTAGCTTACTCCTTCTCCGTGCGGATGCCCCTCAGGCGGCCAGTGCGGCGGGCGGCGAT

GAG

>NODE_1928_length_142_cov_2697.966942

CTTGTCGGGTAAGTTCCGACCCGCACGAAAGGCGTAACGATTTGGACACTGTCTCAGAGAGAGACTCGGT

GAAATAGACTTGTCTGTGAAGATGCGGACTACCTGCACCAGGACAGAAAGACCCTTGGAATTTCGGGTGC

CA

>NODE_1929_length_142_cov_432.239669

CCTCCTAAAGAGTAACGGAGGCGCGCGAAGGTTGGCTCAGAGCGGTCGGAAATCGCTCGTTGAGTGCAAT

GGCAGAAGCCAGCCTGACTGCGAGACTGACAAGTCGAGCAGAGACGAAAGTCGGTCATAGTGATCCGGTG

GT

>NODE_1930_length_142_cov_111.909091

ACGTTGCAATCCGCACGGGTTGCGACAGTTATGTATGCTTTTGATCTGGAAACAGGGATAAACAGTTTCT

TACCAGCTTCTGTTGTCCACGAGAGACGTTAAGTCTTTCTCTTTCTGGATCAAATCAAGCGCGAAAAGGG

CG

>NODE_1931_length_142_cov_47.024793

TGACTGGAGTTCCTTGGCACAAGAGAATTCCATACAGACCAAACTTTACAGATATATATAATGCTTTTTT

TATACTGTGCCATATATTTAAATAGATGCTTAACAAATAACCCTAAAATAATTGATAGTGTACAATCTCC

CT

>NODE_1932_length_142_cov_16.528926

TGCACCAGATGCGTAAGATGAAGAAGACCACGGGAGAGATCCTCGACATCATCGAGATCACGGAGAAGAA

CGCCCGCATCATCAAGAACTACGGCATCTGGATGCGCTTGGAATTCTCGGCTTCCAAGGAACTCCAGTCA

CA

>NODE_1933_length_141_cov_2927.475000

CTGGGCTACACACGTGCTACATTGGGCAAGACAAAAAGTTGCGAATTCGTGAGGATAAGCTAATCTTTAA

ACTTGCTCTAAGTTCGGATTGAAGGCTGCAACTCGCCTTCATGAAGATGGAATCGCTAGTAATCGCTGGT

C

>NODE_1934_length_141_cov_47.966667

AATAAATTTAGTTTGTTAAAAAAAGTTAAATTTTTTATTGCGATTTTGGTGAAATTGGTAGACACGTCAG

ACTTAAAATCTGTTTCTATTTAAAGAGTATCGGTTCAAATCCGATAAGTCGCACCATGGAATTCTCGGTG

C

>NODE_1935_length_141_cov_36.608333

TAAAAAGCGTTTTCATGGTGGAGGTGAGATTCGAACTCACGGCCAGCAACTCATGTCCTGCCGTCGTCGC

CATATGTTAAGGCATTAGTCCGTTGGGCTATACTGGTAAAAGGTTAACCGATCGCCTACCCGAACTAAGC

C

>NODE_1936_length_141_cov_13.575000

TATCATGATGTAGGCTCTCTTTTTTTTCTTCTTTTGCATCTTGATGCAGTACTTCCTCAAGATGCTCAAG

AAGAAGAACGGAACCGACATCAGCGGCGACAACCGCGCCTTGCAGAAGCTGCGCCGCGAGGTGGAGCGAG

T

>NODE_1937_length_140_cov_176.529412

GACTCCTTCTGGATGTTGTAGTCCGACAGGGTGCGGCCGTCCTCCAGCTGCTTCCCCGCGAAGATCAAGC

GCTGCTGGTCGGGCGGGATGCCCTCCTTGTCCTGGATCTTCTGCTTCACGCCGTCGATCGTGTCCGAGGG

>NODE_1938_length_140_cov_67.789916

GGGGAGCAGGAAGGTCTTCTCAGTGCCTCCGTTCATCTCCTGGTTATGCGACTCCACTCGGTTGAGAAGC

TCCGTGTTCTCCAGGATTGAGTCCGGCAGCGGAATGTTGACGTTCTTCTTGAGCGTGTTGTGGACCAGGC

>NODE_1939_length_140_cov_21.050420

ATGAGTGTAGAAACTTTTAGGCGTAATGAAAGTAAGATGGTAAGAAGGCTGTAAAAAGCTAGCATTATCG

AACAATCCGGATCCTTGAGTGAAGGAATTGATTAGGAGCATATCTGCTAGGACCCGAAAGATGGTGAACT

>NODE_1940_length_140_cov_16.420168

ATCGTGAAAGTCGGGAAAACGGGGCCCGTAGGCTAATGGATAAGCCGCTGCCCTTCGAAGGCAGATATTG

TCGGTTCGAGTCCGGCCGGGCTCGCCATTCCTTGGGCCGAAATACCTCGGCAAATAGGCATCAAGTAGAA

>NODE_1941_length_140_cov_8.571429

TTCCACTCTTCTTCACTTCTCTGTTGATCGTAAACCTCTTACCATGTTACCGTACGTACAAGGTTCCACT

CTTCTTCACTTCTCATCAAATATGTTTCCCGGGTACAGCGTGGCAGATCAACACAGTCTCTCTATGATTT

>NODE_1942_length_139_cov_9290.169492

TTCCAAACGCTCAGTTGTTACTTTTCGTCCGCACCGGCCAGCGGAAAATCGAGTAGCCGAGGCCTGAGGG

GTCATCATCCTTTTTGAGTGTCTTTCGACATTCACGCATCATAGCATTTCCACGTAAACGTCCGCCTCA

>NODE_1943_length_139_cov_1532.144068

GATAGCGTAGCAGCTCACTGGTCTAAATAAGTTGTCCTGCGGCGAAGATGTAACGGGGCTCAAGCCATGA

GCCGAAGCTTAGGATGCATTTATTGCATGGTAGCAGAGCGTAGTGTGACATGGAATTCTGGGTGCCAAG

>NODE_1944_length_139_cov_446.720339

TTAGATACCCTGGTAGTCCACACCGTAAACGATGAATGCCAGTCGTCGGGCAGTATACTGTTCGGTGACA

CACCTAACGGATTAAGCATTCCGCCTGGGGAGTACGGTCGCAAGATTAAAACTCAAAGGAATTGACGGG

>NODE_1945_length_139_cov_339.161017

GGCACCGAGAATTCCAAGTCACCGTCGTCACTTCTCGGTGTTAAGATCCCGGATTTGCCTAAGATCTCCA

CCTACAAGCTTAAACACGGACAACCAACGCCGTGCTGGCCTAGCCTTCTCCGTCCCCCCGTCGCAATAT

>NODE_1946_length_139_cov_210.161017

TTGGCACCCGAAGAATTCCAAACCCGGCGATGACCTACTTTCACATGGGGAAGCCCCACACTATCATTGG

CGCGGAATGGTTTCACTTCTGAGTTCGAGATGGGATCAGGTGGTACCCAAACGCTGTGGTCGCCGGGAG

>NODE_1947_length_139_cov_94.508475

AGTCTAAAGATTCACAACCATACATTACAAAAAAGTAAATAATGGCGGAGAGGGAGGGATTCGAACCCTC

GATCCGGGATAAACCAGATACTCCCTTAGCAGGGGAGCGCCTTCGGCCACTCGGCCACCTCTCCAATGG

>NODE_1948_length_139_cov_57.508475

CCTTCGGGGCGGGAGACATATTGCCCCCACCAAAGGGCAGCACTGCGCGGTTGTAGCTCAGCTGGTTAGA

GTACCGGCCTGTCACGCCGGGGGTCGCGGGTTCGAGTCCCGTCCAGACCGCCATGGAATTCTCGGTGCC

>NODE_1949_length_139_cov_15.271186

TTGGCGTTTCGACGGATCGCGAGCTGGCAGTTACGTGGCCACCGAAAGGTGGCTAGACCCACACTGTCAG

CGTTGATCCGTCTCCAGCGCCTAGATCAACCGTCGGCCGCTTGCCCTCGATACTTGGAATTCTCGGTGC

>NODE_1950_length_138_cov_949.820513

GAAGGCCGCCAATCTCCTTGGCGGCCCGTGATGTGAACCTTTGCCGACTTCTGAACCCAGAGTGAAGACG

CAAACAAACCTAATATATCTACTTTGCTGAGGCCGTGGAATTCTCGTCTGCCAAGGAACTCCAGTCAC

>NODE_1951_length_138_cov_920.401709

GAGATTTGGGTATAGGGGCGAGCGCAAAGATGACTTTTATTCCCATAATTCATTCGCAGTGAAACCGCCG

ACGCTCACGCACCCAGCCGGGTCTGCGTTCGAGCCTGGGGCCCCAATCTTCCTTGGATGTCTGAGCGC

>NODE_1952_length_138_cov_30.957265

CAGTAGTAGGAGTCGCAGTAGACAGTGTCCTTCTCGCAACATTGGTCTTTGGTGCACTTGTCGTCCTTGC

ACACGGTAGTGTCGGCATCATCCACCAGCTCATAGTATTTAGGGCAGTCGTAGCTCGAGCAGACCTTG

>NODE_1953_length_137_cov_4585.948276

GGTAATTAGTACAAGTTAGCTCAATACATTACTGCACTTACACACCTTGCCTATCAACGTTGTAGTCTCC

AACGGCCCTTTAGGGAACTTAAAGTTCCAGTGAGAACTCATCTTGAGGCTCGCTTCCCGCTTAGATG

>NODE_1954_length_137_cov_1395.267241

GCACCGAGAATTCCAAAGCCTGGTAATGTCCTACTTTCACATAGCAAATGCTACACTATCATCGGCGCTG

TTTCGTTTCACTACTGAGTTCGGCATGGAGTCAGGTGGGTCCAAAACGCTATTGTCACCAAGCAAAT

>NODE_1955_length_137_cov_341.103448

TAGCGACAATAGCGCTGTGGTACCACCTGATCCCATGCCGAACTCAGAAGTGAAACGCAGCTGCGCCGAT

GATAGTGTGAGGTCTCCTCATGTGAAAGTAGGTCATTGCCAAGCGCCGCGAAAAGGGCGTTTGGTGA

>NODE_1956_length_136_cov_752.339130

CCCGCAACGAGCGCAACCCTTGTTTCTAGTTGCTTTACAAAAGGTATCTTGAAAGACTGCCGGTTATAAA

CCGGAGGAAGGTGAGGATGACGTCAAGTCATCATGCCCCTTATACCCTGGGCTACACACGTGCTAC

>NODE_1957_length_136_cov_221.713043

TTCCCAACGTAGCTACTCTGCGGTGCCCCTGGCGGGACAACAGATACACCAGCGGTTAGTCCAATTCGGT

CCTCTCGTACTAGAATCAGATCCACTCAAATTTCTTGCGCCCACAGTAGATAGAGACCGAACTGTC

>NODE_1958_length_136_cov_94.921739

TGGACCGGTCAGTTTGTTGTCAGGCTATGATGACAACAAATGCATAGTTCATCCGATTGCTCAGTGAAGA

ACAATCTTGTTACCATCTTTTGTTACCATCTTAAACGTATCTGAGCCTCTGGAATTCTCGGTGCCA

>NODE_1959_length_136_cov_54.556522

TGCGCTGCAACAGCCCTCTGGTCACATCTTCCAGCTGGGCCTGCGGCGATATAGCCAAGTGGTAAGGCCG

TGGATTGCAAATCCTCTATCCCCAGTTCGAATCTGGGTGTCGCCTCCATGGAATTCTCGGGGTGCC

>NODE_1960_length_135_cov_195.833333

CACATCGTTTCCCACTTAACTGTAACTTTGGGACCTTAGCTGACGGTCTGGGTTGTTTCCCTTTCCACGA

CGGACGTTAGCACCCGCCGTGTGTCTCCCATGATTGCACTTGTTGGTATTCGGAGTTTGCAAAGG

>NODE_1961_length_135_cov_183.798246

TTGGCACCGAGAATTCCAGGGGCTCAGTCTCGCCTAACTTTTCTCATGCCCGACCCCAGCGGCGCCCGCG

AAAGCCCAAAGCCCAAACCACGTCCCGACGTCTGGCCGCCGTCTTCGCGCCAACGCGCGTCAAAA

>NODE_1962_length_135_cov_44.526316

TGAGGAAAAAAAACTGGTTAGTTTAGTAATTTCATGTCTATGTCCGGTTTGTGTTTTGATATGAGGAAAA

AAAACTGGTTAGCTCAGTAGTTTCATGTCTAGAGTCTGGTGGCTTGTTTTACAGCGTCGTTGCCA

>NODE_1963_length_135_cov_13.868421

GTCCATCCGCGCCTCTCCCATCTCTTTCTTCGCCCTGCCCTCGCACGTCTGGCCGTCTAGCTTCTCCGAG

ACGGCGCTGATGTACGACCTGCCATCTTCTTCGTTTTTGTCGGGGGCGTTCTTGGATGTGTCCCA

>NODE_1964_length_135_cov_2.719298

CGTCGTCCTTCTCGCAGCAAAGGCGTTTGGTGCATCCACTGTCGGCGCAGATAGTGGTGTCAGCGTATTC

CACTGGCCTGAACTTGTTGGGGCACGTGAAGCTTGAGCAGACCACGTCGTCCTTCTCGCAGCAAA

>NODE_1965_length_134_cov_1105.752212

GGCAACCCGAGAATTCCAAGAGCTCAGACATTGAAACTAGCTATCGGCATCAGCCTTGTCTGCTTTTTGC

AGGGGCCTAATAGTGAGCTGGGTAGTGTATTTCCTCACAGCTCACTATACCCGACCCTGGCCAC

>NODE_1966_length_134_cov_391.628319

GGTGTGTGTGGAGTTCGTTGGTACGCTAGATTTACTTGGAGCGGCTAACGAGACTCGAACTCGTGACATT

CACGTTGGCAACGTGATGCTCTACCAGCTGAGCTACAGCCGCTTCATCTGCTGACTCCCTGTCA

>NODE_1967_length_134_cov_75.893805

TGTGGTAACCAAATCAGCCACACCTCGGCACATGAGTCCTTTGCTGCAGTTGCTTCCTTCCGGACCTGGC

CGAGTTCACAGAGTATCATTGCGAGGGGACCAAAAGGGTCACCCCGCGCCACAAATCCAACTAC

>NODE_1968_length_134_cov_63.778761

GCACCCGGAGAATTCCAAGGGTAGAGAAAATCGCGGTAGAGGTGTTTCACTTTCGGAGCAAGCTCCTCCC

TCCTACGCTACACCCACAATCTCTCCACCCGGATCCAAGCCCCAGTCAAGCTCAACAGGGTCTT

>NODE_1969_length_134_cov_19.230088

GACACGCCCACGGGCCGGGACGTGGTGCACCGGGCCCTGGGCCTGGACACAGCGGACCTGGACGCGGTGG

ACCAGGACACAAATGAGGTCACGGCAAGTAAACCTACAAATACCTCTACTCTACTCTACTCTAC

>NODE_1970_length_134_cov_6.336283

TAACCATTATACCACAGCTCCCGACAGAACTCGTCGGAAGCAAGACCTCAGTAAGCGCAACCAACGCTCG

TCTGGTACTTCCAAGCGCGACCGTTCTTCGTCTCCGCGTGGCAACCTCTGGAATTCTCGGTGCC

>NODE_1971_length_133_cov_6244.214286

GATTAGAGGCTGCGGGGAAGCAACTTCCTCGCCCTATTCTCAAACTTTAAATGGGTAAGAGACCGGGGTC

GCTTAATTGGACCCTGTGTTGGAATGAAGGTTGCTAGTGGGCTGGAATTCTCGGTGCCAAGGA

>NODE_1972_length_133_cov_4177.125000

GATAGGTCAGGTGTGTAAGGGACGCAGTGACGATAAAAAGTTATCCCTGTCTTACCCGGGAGACTATTAA

TAGCTTTCCCTCCATGGTGACAGATACCTGGTAAGCTTCCGAGCGCCTGGAATTCCGGGTGCC

>NODE_1973_length_133_cov_69.598214

TAATCTGCGAAAAGCTCAGATAAGGTGATAAAAACCGTTATAGACTGCGATATCCGAATGGGGAAACCCA

CCCGTCGTAAGGCGGGTATCAACAAGTGAATACATAGCTTGTTGAGGCGAACCGGGAGAACTG

>NODE_1974_length_133_cov_17.839286

GCGATAGAACTCTACCTCACGAAATAAGTAAATTATCTTAGAATGTAGTGGTCTTTCAATGTTGGTTGCC

CTCCCACCTAGTCTACACCATTCCAGATAATTCACAAAGTCAGACTAGAGTCAAGCTCAACAG

>NODE_1975_length_132_cov_1829.495495

TCATTCGGACTGTGCCTCGCGAGGGCGTTGATGTGAGCAAACGATTGTTTATCCTATGGCCCTTGTAGTG

AGGTGCCGGATGTGAGGATTCACCGTGTGGAATCTCATTAGCTGAGCCCTTGGAATTCTCGG

>NODE_1976_length_132_cov_315.801802

GCTGTGTTTTTAATAAACAGTTGCAGCCATCTGGTATCTTCGGCCTCCAACAGCTCAATCCGCAAGGGAC

GTCACCGTCGGAGGCGTACCTTCTCCCGAAGTTACGGTACTATTTTGCCTAGTTCCTTCACC

>NODE_1977_length_132_cov_312.297297

GACGTCGTCCCCACCTTCCTCCGGTTTATCACCGGCAGTCTCCTTAGAGTTCTCGCCATTACGCGCTAGC

AAATAAGGATAGGGGTTGCGCTCGTTGCGGGACTTAACCCAACATCTCACGACACGAGCTGA

>NODE_1978_length_132_cov_160.459459

ATAATATCATAGACTGTTAGGGTCGATGCCCGAGTGGTTAAGGGGGGCGGATTGTAAATCCGCTGGTTTT

CCTACGTTGGTTCAAATCCAACTCGGCCTACCATCGAAAACACCGGTGCCAAGGAACTCCAG

>NODE_1979_length_132_cov_153.288288

AATTACATGATAACAAAATTGAAAGAAACCCTGATGGGGCTATAGCTCAGCTGGGAGAGCGCCTGCCTTG

CACGCAGGAGGTCAGCAGTTCGATCCTGCTTAGCTCCACCATGGAATTCTCGTGCCAAGGAA

>NODE_1980_length_132_cov_88.891892

GCGGTGAAATGCTTAGAGATTACATGGAATACCAATTGCGAAGGCAGGTTACTACTAATGGATTGACGCT

GATGGACGAAAGCGTAGGTAGCGAACAGGATTAGATACCCTGGTAGTCCACGCCGTAAACGA

>NODE_1981_length_132_cov_19.324324

CAGGCGTTCGATGCTCATGCAGGAGCCTCGAACGAAGCCAAATCTATATTTTTGTACCCGGACTTGTTTG

CCGCACCGTCTCTATCCGCGCGCGTTATTTGGTTTGGTTTGAAACCATTGCTGCCAAAGGCG

>NODE_1982_length_132_cov_3.567568

TGAATGGACCAAACTGGTGAACTATTCTGGTCGACAAAGTCTGGTAGACTAAACTGGTGCCCAAAACTGG

AACTCCCGCGCCTCGTCATCTCCAAAATCGGCTCGGTCGACTGAATGGACCAAACTGGTGAA

>NODE_1983_length_131_cov_125.545455

TTGCCTATGATTAAAACAGGTTGACCGTTTTTAGGCGAGTTCAGGCAGTCTTCTTCGGGAGTTGTTCTGG

CTTTCCGATGGGGAATTATCCAACCTTCCTGTGTCACTGAGGCTCTGGAATTCTCGGTGCC

>NODE_1984_length_131_cov_117.836364

CTACTTGATGCCTATTTGCAGACTCGAACTCCCAACCATCGGTTTTGGAGACCGCTGTTCTACCAATTGG

AACTACACCCCTACAATGCATTTAGCATTAAAAACGTTTGCTAAAGCAATAGCTACACTTA

>NODE_1985_length_131_cov_32.581818

CCTGAAACCATACGCTTACAAGCGGTCGGAGCTCTTCGGACTGACGGCGTGCCTTTTGCATAATGAGCCT

ACGAGTTACTTTAACTAGCAAGGTTAAGACTTTTAGAGTCGGAGCCGTAGCGAAAGCGAGT

>NODE_1986_length_130_cov_3071.944954

TTGGCACCCGGAATTCCAAGTGCTCGGTGTGAGGGTTTCATCTTCACCAAGAAGGCACAGAGTGCAGTCT

TTCTAGAAGAAAAACAAGTATTAAGGATTTTCCTCATTGCACCCTCTGTACCTCCGCGCT

>NODE_1987_length_130_cov_53.651376

GAGCAGGAAGGTCTTCTCGCCATCTGGGTTCTGTCCCGCATTGTGCGTCTCCACCCTGGAGAGAAGGTCC

ACGTTCTTCAGGATGGATTCTGGCAACGGGGTGATGACGTCCCCCGTGAGCGTGTTGTGC

>NODE_1988_length_130_cov_26.440367

CCGACGTTGTGAAATGACTTAAGAGGTGTAGAATAAGTGGGAGCGTAAGCAACGATGAAATACCACTACT

CTTGACTACCCGCTACTTATTCTGTGAAACACTGTCCCAGGAATTCACGGGAGCCAAGGA

>NODE_1989_length_130_cov_8.596330

GGAGTTCCTTGGCTCCCGAGTTTTCGAATCGTTAGGGTTTTTTCGGAGGGTTGGTGGTACTTGATGGTTG

GTTTTGTCTTTCTTAAGGTGCTTTTGTCATTGTTTATTCTACCGAATAGGGGGGCTGTTC

>NODE_1990_length_130_cov_1.504587

TCGTGCATCTATCAATGTCCGAAAATATTTCCGTACCCAAATGTTTAGTAGCATCTCAACTTACTCGTAC

GTACTTGTAACATACAAAAATGTAGTGTGATTTCTGCACTCGTGCATCTATCAATGTCCG

>NODE_1991_length_129_cov_17818.120370

TAGGATAGCGCGGGGTTTCACGTCCTGTGATTACAAATCTATGCACCACTCCAGAAAGCATTCTGTGCAC

GTGTGGAGAAAACACCTGATCACCATCTTTCGTATGAGGACGTGGAATTCTCGGTGCCA

>NODE_1992_length_129_cov_3410.833333

TGTGGTTGAGTAGTGGCTTGAAACCTGTGAAGACAAAAAAGCTCCCTCTACTGATGAGCAATCACGTGAG

GAAAGGCTATCATCCGGAATCAACATGAAAAGTCAGAGGTCTTGGAATTCTCGGGGCCA

>NODE_1993_length_129_cov_1536.185185

TCCCATCAGAAATTAAAACATATTAAAAGTCCCCAAAGTCGAGCGGGCGCGGGAAACGGAAATACTCCGC

TGATTTCCCCCGCCCATAGTGCCCATAAATTCGGCTTGAGGGTTTTCGTCACGCAGAGA

>NODE_1994_length_129_cov_302.833333

AGATCACTCGGTTTCGGGTCTGATCCATCTAACTCATTCGCCCTATTAAGACTCGCTTTCGCTGCGCCTA

CACCTAACGGCTTAAGCTTGCTTGAGAGACCAAGTCGATGACCCATTATACAAAAGGTA

>NODE_1995_length_129_cov_176.157407

CGTGTCATCCTTACTCCAGGGGCCATGCTAATCTTCTCTGTATCGTTCCAATTTTTACGGATGATGACTT

TCGTCATTCAAAAGTTTCGATACACCTCTCGATTTATGCGTGTCATCCTTACTCCAGGG

>NODE_1996_length_129_cov_140.703704

ACTGGAGTTCCTTGGCTCCCGTGAATTCCAGAGGCTCAGCAAGTGAAGGAGTATTCATCACGTGGCCCTC

GAGAGGACCTTAGATTATGAATAGCGTATATTAATGTATGTTTTCTTCATCAGTGCCCC

>NODE_1997_length_129_cov_101.777778

CCTGACTTCACCGGGCTCAGCATCCCCAGGGGCGCGGACGGGAAGATCGACCCTGTTACCCAACTGTTCC

GGACGGAATGGCCGGGTGTAACCACCGGTCCCGTGGTCTCTCAGTTCATGCTGTCGGAC

>NODE_1998_length_129_cov_13.324074

AGGTCGCTGACTTGTTAAAAAGAAAAGATAACTCTTCCCGGGCTCCCCGCCGGCGTCTCCGAGTTCACTT

ATGTTGCCATACAATATCCACCTCCTGGTTCGGGAATATTAACCCGATTCCCTTTCGAT

>NODE_1999_length_128_cov_21242.859813

AAGCGCAATATGCGTTCAAAGATTTGATGATTCACTGAGTTCTGCAATTCACATTACTTATCACATTTCG

CTGCGTTCTTCATCGTTATAGGAACCTAGGTATCCGTTGTTAACAGTTATTTCTTTGT

>NODE_2000_length_128_cov_1946.691589

GCTCCCGAGATTTCCAATCCGTCATAGATATTTTGGGAATGGCTGTTGTCATCCAATCGGTTAGAGGATC

AGTGGATTAGCTCTGATAATTCAGGTAATCATCCGGGACCCCTAACCACAGCTCATCT

>NODE_2001_length_128_cov_355.588785

TGGCTGGGGTAGGAGGATTCGAACCTACGAAGGCGGAGTCAAAGTCCACTGCCTTACCACTTGGCTATAC

CCCAAAAAGGATCCATAAATCAATCTATAATATTCTCTGAGCTAGGAGGGATTCGAAC

>NODE_2002_length_128_cov_32.373832

GGTCTCTATCTACTGTGGGCGTTAGAAATTTGAGTGGATCTGACTCTAGTACGAGAGGACCGAGTTGGAC

GAACCTCTGGTGTACCTGTTGTTCCGCCAGGAGCATTGCAGAGTAGCTACGTTGGGAA

>NODE_2003_length_128_cov_16.822430

TAAAACTTCCAAGGCTGTATGGCCACATGACGAAAGCCTAGCCTGGCAGCGGGCGCTTCCGACCCTTCTC

CTTTTCAAGGGATGGTCAATGGGGTTCTAAAGCAATCTCTTTTCAAAGCCCGATGGTC

>NODE_2004_length_127_cov_7172.358491

CGTTGCGGCTGCCTTGGGACTGCTGCCTCACGGCGGCGGACCCTCGGCGGCTGCTCCGGCGACGGCCTTG

GAACGGCCTCTCTTACGATTGGCCTTTCCCCGTGCGTTGAACAACCGGCTCAGAACT

>NODE_2005_length_127_cov_4761.830189

TTCGACACGATGGCTTTCTCAGAGCCATGATGACGCTTATTTTTTAACGCTAACTACGTTGTGACTGAAC

CAATTATGTCGACTATCTCTGAGGTTCTGAGAGCTCTAGGAATACTCGGGAGCCAAG

>NODE_2006_length_127_cov_312.047170

GTCTCAGTCCCAGTGTGGCTGGTCGTCCTCTCAGACCAGCTATAGATCGTCGCCTTGGTGAGCCATTACC

TCACCAACTAGCTAATCTAACGCGGGTTGATCCTTCACCGATAAATCTTTCCCCCGA

>NODE_2007_length_127_cov_307.160377

GTTTGATTAGCCTTTCACTCCGATCCACAGCTCATCCCCTCATTTTTCAACATAAGTGGGTTCGGTCCTC

CAGTTGATGTTACTCAACCTTCAACCTGGCCATGGATAGATCACTCGGTTTCGGGTC

>NODE_2008_length_127_cov_71.292453

CATAGAACCACTGTCCATCTTCTCCTTCTGCACCAGAGCTCTTCGTCGACAATACTGTTGTAGTACAACT

ACTATCAAGACAAAAAACTCAAACAGGATGGTGCGATTTGTTCCCTTCGTCATGGCT

>NODE_2009_length_127_cov_57.405660

CACACAGATTGTTTGATTAGAAAGTGAAGAGACGTATAGATAGACCTGTTTGGGTCTGTAGCTCAGTTGG

TTAGAGCGCACCCCTGATAAGGGTGAGGTCGGCAGTTCGAATCTGCCCAGACCCACC

>NODE_2010_length_127_cov_32.915094

TGCGTCCTGTGGGGCGCAGGGAGAAAGGTGCGCCCTTGAACAGGGGAGGTGTTGGGTGACGGGCATGCGC

GCGGTTATCTTTGATGTGGCCGAAACATTCATTTTTTCTGAGCGCTTGGAATTCTCG

>NODE_2011_length_126_cov_2740.314286

GTGCTCAAGGATGAAAGATACGTCGGGCGGACTGTACAAAACCCCGGCCGCTGAGAAGCGGCAGGGGGCC

ATGAGGACGCGACGCTGTACAAAAGACCACTGAGAGTTTGGAATTCTCGGGGCCAA

>NODE_2012_length_126_cov_1479.314286

AATCTGTCCGAACGCACCCGAAACGGTAGTGATGATGTTTTTCACAGGCACGTATGTGTCCTTGATCCCC

CATGAAGGCAATTTTTTAACTGCAGTCTGAGCCGTGGTAGTGATGATGTTTTTCAC

>NODE_2013_length_126_cov_41.038095

GGGAGCAGGAAGGTCTTCTCACCACGGTCGTTCATGTACTTGTTGTGCTCCGCAACGCGGCTGAGCAGAT

CCGTGTTCTTCAATAGTGAGTCCGGCAGCGCGGTGATGACGTCCTTCATGAGCGTG

>NODE_2014_length_126_cov_23.190476

GATAGGTGGGAGACTTTGAAGTGGCGTCGCTAGGCGTTGTGGAGTCATTGTTGAAATACCACCCTTTACT

TATTCGGAGTCTAACTTCTGTGAGAAGAACAGTGCGTGGTGGGTAGTTTGACTGGG

>NODE_2015_length_125_cov_1917.538462

CCGAAGCTGTGAGATATTATCTCCTGTGAGGAATTGCTTCAAGACTGGAATTACCGTCTAAGAACAACCC

TGCGGAAACAACTTCGAAGCGGCTAAGCCTGAGGAGATGGAATTCTCGGGGTGCC

>NODE_2016_length_125_cov_810.509615

CGAATGAGGAAACTTTTTAAACTATCTGCTGAATTCATAGGCAGAAAAGAGCGAACCTAGCGAATTGAAA

CATCTTAGTAGCTAGAGGAAGAGAAAGTAAAAACGATTCCCTTAGTAGCGGCGAG

>NODE_2017_length_125_cov_489.298077

CAGGGAGGTAGTGACAATAATGGCGTAATGATGGCCATGCTGTCTCTACCCGAGACTCAGTGAAGTTGAA

ATCGCAGTGAAGATGCTGTGTACCCGCGGCTAGACGTGGAATTTCGGGTGCCAAG

>NODE_2018_length_125_cov_121.125000

TCAATTAAAAATCGTTTGTGTAACCTCACCATTCTTACCTAAGTAAAAGTGATGTGTTACTGCTCAATGA

ATTCTGTCGTGATACCAGCACTCCCTAAAGAGAATTGATATCGCATTACATAGTC

>NODE_2019_length_125_cov_49.567308

GTCAACAAGCCGGCATAGCTCAGTGGTAGTAGCAGCACCTTCGTAAGGTGTAGGTCGAGAGTTCAATCCT

CTCTGTCGGCTCCATCTCGTGCAATGTTGTCTACTCTGGAAATCACGGGTGCCAA

>NODE_2020_length_124_cov_7708.495146

GGTGGGCTTCCCACTTAGATGCTTTCAGCGGTTATCCTTTCCATACGTAGCTACCCAGCGTTTACCATTG

GTATGATAACTGGTACACCAGCGGTATGTTCTTCTCGGTCCTCTCGTACTAAAG

>NODE_2021_length_124_cov_3814.135922

TTTGCAGGCGTTGGAGAGTTAGGGCTTCATCGAGATATGTTATCTTCCAACCTCCAATGAATGGAGATAT

TCAGTTCTCAAACTTTGTATGGATCATCAGCCCCACGAAGAAGCCCAGTTTCAT

>NODE_2022_length_124_cov_2724.135922

CGAGAATTCCAGTTCCTCAGGATCGCCGGCAGAGTCGCTCTACTCCGAAGAGAAGAGCGCAGGTCATCGA

AACAGATCAAGTGGGCACAAGGCCACACCAAAATCTGAAAACCATCGCTGCTCT

>NODE_2023_length_124_cov_1906.766990

AAAAGCCGTGACGAAAAGCTAATTCCTCGTTTCCGACGCGCTTCGCTGCAAAAGCGGGACGCGCATGAGA

CCAAACGTTGCTTATCCAGCACTTCTGAGCTTTTGGAAATCTCGGGAGCCAAGG

>NODE_2024_length_124_cov_1640.834951

TGGCACCGAGAATTCCAACTTTCGTACGACTTGCATGTGTAAAGCATACCGCCAGCGTTCATCCTGAGCC

AGGATCAAACTCTCGATAGTTTCCTGAATATTTAAAAGAGAGATAAAGAGTGCC

>NODE_2025_length_124_cov_9.135922

AAAAAAACGCCCCCCCCCCCCACTTTTTTTTCTTGAGTCAACGCAACTAGTCGATCTCCTCGATCTTCGG

GCCCGAGTCACCCTCCGCCTCGGGAGTCGGGGCACCTCCGGCACCGCTCATGTG

>NODE_2026_length_123_cov_1383.980392

GTCCACGCCGTAAACGATGTCAACTAGCCGTTTGTGGCCTTGAGACGTGAGTGGCGCAGCTAACGCACTA

AGTTGACCGCCTGGGGAGTACGCCGGCAACGGTAAAACTCAAAGGAATTGACG

>NODE_2027_length_123_cov_1372.872549

CTTACCCACCCACAACCGAGTCTTCCATATGGCAGCGCTAGATTACCGTCTGACAGCGGCGGGCTTCTGG

GCATTCCTGGTCTGGCTTAATATCCCACGACGATCCGGACAATCAGAGCCCGC

>NODE_2028_length_123_cov_390.117647

TGACTGGAGTTCCTTGGCAGCTGAGTATTCCATCGGCTCCGACAGACGATGAAATATGTTTGCTGCGCTC

ACTTTTGGACAGAATCCTCGCGATTGACTGGCAGTTTTTGTCGTCATGGCCGA

>NODE_2029_length_123_cov_44.372549

CGTGCTCTACCAACTGAGCTAGTACCGCTTCATAATTTAGTTTCATATATAGTTTGGAGCGGTACACGAG

ACTCGAACTCGTGACCCCGACCTTGGCAAGGTCGTGCTCTACCAACTGAGCTA

>NODE_2030_length_123_cov_23.186275

CACAGTTTGCACTGGCAGTCTGGCTAGAGTTCCCGACATTACTCGCTGGCAACTAACCACAGGGGTTGCG

CTCGTTATAGGACTTAACCTGACACCTCACGGCACGAGCTGACGACAGCCATG

>NODE_2031_length_122_cov_16679.782178

TTGGCACCCCGAGAATTCCAGTCCTGGCATAAGCTATCTTCCCAAAGGACTGCTCCTTAAGTATTGTAAC

TGTTATAGCGTTTCACCATTGAGTTCGAAATGGATCAATGTGGTTCCACTAT

>NODE_2032_length_122_cov_10331.475248

TGGGCGGGAAAGCGGGGCACTTTTTGCCCCGTCCGCGTGCGCTCTCGACGGCCCTTGAAAATCCCATGAA

CGTGTTTTATTCTCACCCCCGGTCGTACTCATATGGAATTCTCGGGGCCAAG

>NODE_2033_length_122_cov_2804.138614

GTAAAGCGTTCGTAGGCGGTTTATTAAGCAAGATGTGAAAGCCCAGGGCTCAACCTTGGAACTGCATTTT

GAACTGGGTAACTAGAGTACTGTAGAGGGTGGTGGAATTTCCAGTGTAGCGG

>NODE_2034_length_122_cov_426.336634

CCGAGAATTCCAGGCGCTCAGGTGTAATTTGTCCGCACCGACACCTCGGCAGAATCGAGAAGCCGAGGTG

CCTGAGGGAAGTCATCATCCTTTTGACCGTGTGTACACGCATCATGGCGCCG

>NODE_2035_length_122_cov_127.178218

CTTGGCACCCGAAATTCCAAAGCCTCAGAAACAAATTTGATGGAGTTTTCGACAGTCAGCCGTTATCAGC

GGCGTTCCGCGTGTCACAGAAGTCATATTTGTGAGCAAATCTTCATCGACTG

>NODE_2036_length_122_cov_112.554455

TCCAGCGGCAAGACCCTCGATTACCAGGGCTTCAACCTCTACACCGACGAAACCAAGACTCTCAAGCTTT

GCCTCGACGACCCTAAGTGGTACACTACCGTCTGGCTGAGCATCACCGAGGT

>NODE_2037_length_122_cov_49.782178

ATGGGAGAGGCGCGGATGGACATCGAGGACAGCGAGATGGCCTACCTCGGCTATCATGGCGCCGAGGCCT

ACGGTCTGACATGGAAGGTTAGAGGGTACTGCAAGGACGAGAGCAACCCCGG

>NODE_2038_length_122_cov_40.495050

GGGGAGCGGTTGATGGAGGAAGCGGAAATGGTTGCGTTGTGCCGGACAGGTTTCCCTCAGCTAGTATTGC

TGGGAAGGAGAAGTATTGCGGCACGAACATTCACGGGTGGGAGACCGATAGC

>NODE_2039_length_122_cov_37.762376

ATTCCGACGATTTTGGGTTAACCCCTTAAGAACTGGTCCCACGTATTTGGGGACAAACTACTTGGAATTA

GTGTGGGATAAGTTTTTCAGTTCTGAATGGGTTAAGAACTGGTCCCACGAAT

>NODE_2040_length_122_cov_16.772277

GGTAGGCCTCTTGGTTTCTCGCATTGATAACTTATTAACCTGGTCAGAGCCGGAAGGCAGCAGCCATAGT

AGGTGATTCAAGTGCCGAGAGTTTGCTGGGAGGTCTTGGAATTCTCGGTGCC

>NODE_2041_length_121_cov_2153.510000

ACAACAAACTGAAGGGGTCCAACAACTGCTCGAACTCCAACAACAACGCGACTAACACCAAGAACACCCT

CCTTGATGCTCTTTTCCAAGAGACTTCGCCCCCGGCTGCGGCTGGCCACGC

>NODE_2042_length_121_cov_2079.970000

TGGCACCCGGAGAATTCCAATCGCTCAGTCGAGTTCTTGCTATTCACATGACGCGTTTTGCGTTCAATCT

CTTCCTTTCATTCGTTGACATCCTTGCGATGACAACAAACTGAAGGGGTCC

>NODE_2043_length_121_cov_821.280000

GCACCCCGAGAATTCCATACTCAGAGCGTTGTTTAAAATCGTCCACATAGTTCTTCCGAATTCAGGACGA

TAACACAGTATTGATGGCTCATCATCGAAGCAGTGAGAACTCATCTCAAAG

>NODE_2044_length_121_cov_228.520000

CTTGGCCCCGAGAATTCCATCTGATCAGCACGAATTAGCGTGTGTTTGTCACCAGAGCCCCTCCATTTGG

AGAGGCATATGACTGGGGCGGCATCCACAGCGATTAGCTCATCACCTCAGA

>NODE_2045_length_121_cov_134.170000

GAATACCAACAAGTGCAATCAAGGGAGACACACAGCGGGTGCTAACGTCCGTTGTGAAGAGGGAAACAAC

CCAGACCGCCAGCTAAGGTCCCAAAGTACTAGTTAAGTGGGAAACGATGTG

>NODE_2046_length_121_cov_35.670000

ACTAAGGATAGTTGTTAAGGATGATATTTAAAAATTAATAACTTAGAAGAGTTATTGCCACGGAAGGTAT

GAAGGACAACGAAGGGTTCGTTTAAGGAATACTAAGGATAGTTGTTAAGGA

>NODE_2047_length_121_cov_31.420000

GCTGAGCCCGGTGAAGTCAGGATTGGCCTGCAAGTTCTCTGCGGCGACTTGGACGAGACCGTCGTTGCCG

TATTGCGAGAAGGGGACGTCCCTGGTCAGAGCCATCCAGTACACCTCTGTC

>NODE_2048_length_121_cov_23.220000

AGGCCGGAGATCGCTCCCGCGTCCTTAGTCGCCTGTCTCTGCGAGTCGTTGAAGTACGCGGGCACGGTGA

TCACCGCGTTTTTCACCTCCTTGCCCAGGTAAGCCTCCGCGATCTCCTTCA

>NODE_2049_length_120_cov_2878.707071

TGGCACCGAGAATTCCAGCTCGTCAGGGAATTGTAGTCTTTTCGTAAGCATCAACGCGCCAAAAACGGCG

CTCACGAAAGGGTTTCGTCTTCATCCAAGCGACTCCCTTACAATACCCAA

>NODE_2050_length_120_cov_1395.101010

TAGTTAGCCCCGTCGGTCGCGGGGCGGTGACGAAATATAACGGACTAGAGTTTCTGAAATCAGTGAAAGC

TGCCTGAAGATACACCCTTGGCTGTCCGAGCCCTTGGAATTCTCGGGGTG

>NODE_2051_length_120_cov_781.212121

TGTGTAGGATAGGTGGGAGGCGTTGAAACAGAGTCGCTAGATTTTGTGGAGCCAATCTTGAAATACCACC

CTTGTATGCGTGATGTTCTAACCTAGGGCTGGAATTCTCGGGTTGCCAAG

>NODE_2052_length_120_cov_711.636364

TGGCCCCGAGAATTCCAGCTGGTCAGGGAATTGTAGTCTTTTCGTAAGCATCAGCGCGCGTTTGACGGCG

CTCACGAAAGGTTTTTGTCTTCATCCCAGGCGAGGCACAGTCCGAATGAC

>NODE_2053_length_120_cov_532.636364

CTTGGCACCCGGAATTCCAAGAGCTCAGGTTTGATCCTTGTCTAAGCACAGATGCCTTTTTGGCTGTCAT

ACGGCACTTCCTGTTTACTGGATTGTGTTCATCATAGCTCTCACCTTCCT

>NODE_2054_length_120_cov_319.303030

TTTTGAAGCTGATGACGGCTTACGAGATGTTGGTGTGTCTGGAGTTCCTTGGCTCCCGAGTATTCCACCC

CCCTTTTTTTTTTCAAGCAGATGACGGCTTACGAGATGTAGGTGTGACTG

>NODE_2055_length_120_cov_71.515152

ACCAGAACTATCGACGGTTTTCTTTCTCGCAAGTAGTTACCCTATAAAAAGGTAGACGGCGGCAGCGCAA

TGAGCTTGAAAAAAAAACGAGCGTTGCGTGCCGCTGACTGGACTTGCTTT

>NODE_2056_length_120_cov_68.272727

TCATAACCTGAAGCTTAGAGGATTTTCTTGGAAGCATGGCATCAACTACTTCGTCTCTTACGAGACTCGT

CATCAACTCTCAGCATTCTCATTAAAGAGAGCAACCCGGATTTACCTAAG

>NODE_2057_length_120_cov_22.101010

TTGTACACACCGCCCGTCGCTCCTACCGATTGAATGGTCCGGTGAAGCCTCGGGATTGCGAATGCTTCGT

GCGGACGCGAGAACTTGGCTAAACCTTACCATTTAGAGGAAGGTGAAGTC

>NODE_2058_length_120_cov_12.040404

AATGGGCCGATTTTGATTTTGACCGAAGTTCTGGAGATCCATATTTTCACATCTGGGGGACTCTTTAGGA

AAAAATTACTTGATTTTAGTATGGCCAGGAAAATTACTTGATTTTAGTAT

>NODE_2059_length_119_cov_6861.938776

ACATCTCTTTCATTCCTTGGTAACGGAATGATGAAAACCTTTCACAAGGAATGGCTTGATTTACTCTCCA

TGAGAGCCAATGGCTGTAGGTTCTCTGACTGTTTGGAATTCTGGGTGCC

>NODE_2060_length_119_cov_5577.693878

CCATCTAGTAGCTGGTTCCCTCCGAAATTTCCCTTAGGATAGCTGCAACCTGCGTTCAGTTTTATCAGGT

AAAGCGAATGATTAGAGGCTGCGGGGAAGCTGGAATTCTCGGGGCCAAG

>NODE_2061_length_119_cov_827.285714

TTGGCTCCCGAGTATTCCAAGTCCTCATTGGAAAGAAGTATTTATCTTCCACAGGTTTGCCTTCTGGAAT

TTTAATCAGCATACTTTTTAATCACAGGACAGGGTTAGAATTCTAGTTT

>NODE_2062_length_119_cov_169.091837

TACCTGAAACCGAACGCCTACGGGGTTGTATCTCAATTTGGTTAGAGTATCACCCTGTCACGGTGAAAGT

TGCGGGTTCGAGTCCCGTCAATCCCGCCAGGAGCGGTAAATCCTTTGTA

>NODE_2063_length_119_cov_111.408163

CCGTCCTGCTGTCTAAATGAAAAACAACTCAAGCTAAGTCACCACCGAACTTGAGTTGCCTTTCACACTG

ACAACATCTTCACACACAATATAAAGATGATCAACGATGAGTTCTTCCA

>NODE_2064_length_119_cov_64.510204

TTGGCACCGAGAATTCCATGGCGTCGACGATGAGATTCGAACTCACGCGGGCAAAGCCCAACGGCTTAGC

AGGCCGTCGCCTTAACCACTCGGCCACGTCGACGTTTTCCTTTGATGAT

>NODE_2065_length_119_cov_25.642857

AGATCACTCGGTTTCGGGTCTATTCCCAGCGACTATGTCGCCCATTTAAGACTCGGTTTCCCTACGCCTC

CGCTAAGCACTTAAGCTTGCCACTGAGAATAAGTCGCTGACCCATTATA

>NODE_2066_length_119_cov_20.295918

CCACGTCCTTCCCCTGTACGCCGGAGTCAATACACCATGGCGACCGAACTGAACCCTGACGGATGGGATG

GGGGAGAGCCCGTTGTGGAACCCGTGGTCGTGGAATTCTCGGGGCCAAG

>NODE_2067_length_119_cov_5.520408

GAGCATCCCCGCATGGTGTATATGCTAATTTGATGCCGAACCAAAACTGAATATCGAAATCGTTTCTGAA

AAAAACGATCCTTTGCCGTCCATCCCCCGAGCATCCCCGCATGGTGTAT

>NODE_2068_length_118_cov_7896.783505

TTGGCACCCGGAATTCCAGCCGTTTACCCGCGCTTGGTTGAATTCCTTCACTTTGACATTCAGAGCACTG

GGCAGAAATCACATTGCGTCAACACCGTCTCCGGCCCTCGCAATGCTT

>NODE_2069_length_118_cov_4145.670103

ATACAGAAAGTACTATGCTCGTATCTACAGCGTGAGTTCGGAACAGATCTTGGTTTTGTAGAGGTAATCA

CAGGGCCTGGAGTTCTTCCTCGCCGCGAGCAGCGTGATTCTCGGCGAA

>NODE_2070_length_118_cov_2784.020619

CGTTGCGGCTGCCTTGGGACTAAGCGCTGATGCACTCTATACAAACATGTTAGACTTCTGACAGCATCGT

GCTGACAGCATTCCGCAACAGAGCTGAGCTTTTGGAATTCTCGGTGCC

>NODE_2071_length_118_cov_274.639175

CGAACTGTCTCACGACGTTCTGAACCCAGCTCACGTACCGCTTTCATGGGCGAACAGCCCAACCCTTGGG

ACGTACTACCGCCCCAGGATGCGATGAGCCGACATCGAGGTGCCAAAC

>NODE_2072_length_118_cov_122.938144

TTGCGGCCATCTAGCAGAAAGCACCGTTTCCCGTCCGATCAACTGTAGTTAAGCTGCTAAGAGCGAGACC

GAGTAGTGTAGTGGGAGACCATACGCGAAACTCTCGTGCTGCAATCTT

>NODE_2073_length_118_cov_53.484536

AATCTGGGCAACCAGGCTTGAGGGGCACAGACCAGGGGGTTGCGACTGTTTATCAAAAACACAGGGCTCT

GCAAAATCGAAAGATGAAGTATAGGGCCTGACACCTGCCCGGTGCCGG

>NODE_2074_length_118_cov_43.041237

GGAAAGCACCCTAACTCGTCAGATCTAGGAAGTTAAAACTGGGAGGGCTCAGTTAGTAGTGAGGTTGGTG

ACAACTTGCGAACCCTGAGTGCTGTAATCTTGGAATTCTGGGTGCCAA

>NODE_2075_length_118_cov_15.268041

CGGTTATAAGATCCCTCAAAGATGATGAGGTTAATAGGTTCGAGGTGGAAGCATAGCGATATGTGGAGCT

GACGAATACTAATCGATCGAAGACTTAATCTGGAATTCTCGGGGCCAA

>NODE_2076_length_118_cov_2.020619

TGTGTGTGAGAAGTAGGATTGTTTTTGTTCCAGACATATACTGGGTGTCGATGGTATTTCACTGCATGAC

CGTCTGTGTGGATTATGATCGGGTTTGTGTGTGTGAGAAGTAGGATTG

>NODE_2077_length_117_cov_15202.927083

TGGACCGGTCAGTTTGTTGTCGCGGGATGAAGACATCATGCAATACCATCTTTCGAGTCTGAAAACACCA

TGACTGACAGAACTTCTGACAAACGGACCGCCTGGAATTCTCGGTGC

>NODE_2078_length_117_cov_46.593750

TCGGCATGCACCTTGGGTTTCAAGAATCTCGTCGAGTCCTAAATCATCCCCACAGTGGTCAGTGTAAAAA

CACCAACCTGTTAAGTTTAGCAAAGGTCACTGTATCAAGATAGTATT

>NODE_2079_length_117_cov_46.322917

AGAACTCTAAGGAGACTGCCGTCGTTGTAGCTCAGTGGTTAGAGCACCTGTCTTACTCACAGGGGGTCGA

GGGTTCGATCCCCTCCAATGACTCCACTTTGGGTTCCGGGGGGAGTA

>NODE_2080_length_117_cov_21.177083

CGATGGTGGCGACGTCGCAAAGACCACCGACCACTCCGTTCACAGCATTCACTGTGAACGGAACGGAGAA

TCGTGAGACTGATAGATGGAATTCTAGCGTGCCAAGGAACTCCAGTC

>NODE_2081_length_117_cov_18.956522

CTAATAAGAAAGCCAGGCTGAGATTTAGAGGAAAATAAGAAATTCGGGAGTCGTTATCTTGAAATTAGTG

TAGAAAGCACCCTCTCTCAAATAAGAAATCCAGCCTGGACCAGATTT

>NODE_2082_length_117_cov_15.239583

TGGAGTTCCTTGGCTCCAGAGTATTGCAGGGGTTTGGCTGGCGGTGGTTCTTGGCCATGTTGGTGTGCAC

AAAGTTCACGATGTCCGGGCGGATGGGCACCGTGAACACGGCGGGCA

>NODE_2083_length_116_cov_27720.115789

CACCCCGAGAATTCCAACAGCTCAGAGGGATAACTGGCATGTTCGTTAATTTCGCTCACAAATGAACAGC

GACTTAACGTATATCATCATTGCCGTTCCTTAAGTCTGTTCCACGA

>NODE_2084_length_116_cov_23506.010526

GTGAGGCTGCAATTCCTCTAAGAAGTTCCGAGCCTCACGGCTATGACGAGAAACTCACTCTTGCGCCGAG

ACGCTAACCAGACAGTATTCTGTCGGTGAGGCTGCAATTCCTCTAA

>NODE_2085_length_116_cov_7249.957895

CTTATGGTGGGGGACAACAGTCCCTCGTAGTCCAATGGTTAGGATATATGACTCTCACTCATGAGACCCG

GGTTCAATTCCCGGCGAGGGAGCCAAAAACTTTCAGCGACGGATGT

>NODE_2086_length_116_cov_33.547368

GTTATTATGTCACTAATAGTGGGTTGCTAACTCAATGGTAGAGTAATCGGCTTTTAACCGAAAAGTTCTG

GGTTCAAGTCCCAGGTGACCCACCATACCACGAACGACTCTGCTGC

>NODE_2087_length_116_cov_30.105263

GATGATGATGATGATGATGATAATGTTATCACATTATGACAGCTCATGTCCCTGAAATCCGGCCCGTGGT

CACAATTCCGGCTCAACAAAAAAAGCGTCTCGATGGAAGAAAATAT

>NODE_2088_length_115_cov_3354.925532

CTTGGCACCCGAAATTCCAACCGCTCGAGAAATGGATGACGCTTAATTCGTCCTCAGTGTTTGCGACTAA

GATTAAAGCTTGTCGTCATCGCGGAACTCGAAGGCGTCATTCCCA

>NODE_2089_length_115_cov_1623.563830

CTACTAACGAAATCTCGGTTGATTTCTTTTCCTCGGGGTACTTAGATGTTTCAGTTCTCCCGGTTCGCCT

CGTTAACCTATGTATTCAGTTAACGATACCTAGCTTATGCTAAGT

>NODE_2090_length_115_cov_1552.627660

CGCAGCGAACTGCGATACGTCATGTGAATTGCAGATTCCGTGAATCATCGAAATCTTGAACGCAAATTGT

ACTCCCTTGAAAGAGGGAGTGTGCCTGATTGAGTGTCGGCTGGAA

>NODE_2091_length_115_cov_97.574468

GGCCGTAGTCGTATGACCATAGACCCTCGCATCCCTACAATGCCGGGACGGAGCACGTTGGGTTTTCACC

AACCAGGCAGACATTGCTTGCACCAAGCGCGAAGCGCCGTGAGGT

>NODE_2092_length_115_cov_73.797872

GCACACGGTGGAGTCAGCGTCGTTCACGAGCTCGTAGTATTTAGGGCAATCGTAGCTGGAGCAGACCTTG

TTGCAGCACTGCGACTTCTCGCAGTTTCCGTTCTTACATTCCTCC

>NODE_2093_length_115_cov_43.851064

TAGAGCACTGGATGGGTAATGTCAATCATTCCGCCCATGTAGGCGGGAGTGGCGGAATTGGTAGACGCGC

TGGATTTAGGTTCCAGTATCTTAGGATGTGAGAGTTCAAGTCTCT

>NODE_2094_length_115_cov_3.329787

ACGGGGCTCCTTAACCCAAAATCGTCGGAATTTCAGTATTTCTTGTGTTGCAACGTTTTTCATTTTACAG

TTGGGAGGCGGGATTGGTCATCGGGTAGCACTTTTATTTTCCTCC

>NODE_2095_length_114_cov_25961.107527

GACGGGTGGGTTTCCCCATTCAGAAATCTCCGGGTCAAAGGTTGTTCGCACCTCACCGACGCATATCGCA

GCGTACCACGTCCTTCATCGCCTCTTACTGCCAAGGCATCCACC

>NODE_2096_length_114_cov_612.494624

TTCAGCGGTTATCCTGTCCGATCATAGCTACTCTGCACTGCGGCTGGCGCCACAACAGATCCACCAGTGG

ATCGTTCACCCCGGTCCTCTCGTACTAGGGGCAACTCCTCTCAA

>NODE_2097_length_114_cov_373.440860

TGTGGATACGAGCATTGTACTATTCAGTAAGTAGTGCAGGACGATGTGCAGAGGTTCGACGTCAAGGATA

TGATCGTAAGATCATTTCAACTTGAGAGTTTGATCCTGGCTCAG

>NODE_2098_length_114_cov_340.677419

ACATCCAAAAAGCATACAGGATGGTTCCGTAGCTCAGTGGCAGAGCGTGCGACTGTTAATCGCGAGGTCG

AGGGTTCGATGCCCTCCGGAACCGCCATGGAATTCTCGGGGTGC

>NODE_2099_length_114_cov_16.064516

ATAGATAAATCAAATACTCATTAACCTTAAGTTTAGAAAATTATGACTACTAACCTGAACAACCGACATC

ACCATCATATGGAACTGCCAGTAGGAATTCTCTGGTGACAAGGA

>NODE_2100_length_113_cov_3988.641304

AGTCGGCCCCTAAGGCGAGGCGGAAACGCGTAGTCGATGGGAAACAGATTAATATTTCTGTACTTCTATA

TATTGCGAAGGAGGGACGGAGTAGGCTGGAATTCTCGGTGCCA

>NODE_2101_length_113_cov_249.217391

GGAATAATGAGATAGGGCCACACCGCATGACGAAAGCTTAGCCTGGCAGCAGGCGCTTCCGACTCTCTCC

TCTCGAGGGGACAGTGGGGTCCAAAGCAATCTCTTTTCAAAGC

>NODE_2102_length_113_cov_156.923913

GAGAGGTTGGTGCGGGCAGGCGCCTCGCCAACCTGGTCAGATCCGGAAGGAAGCAGCCACAACGAGTCCC

GCTTGGGTCGTTATCAATCTCTCACCTGGAATTCTCGGTGCCA

>NODE_2103_length_113_cov_57.228261

CTCCACCAGAGTTTCCCCTGGTGATCCTACTGGGAAACGAACCCAGGGCGGGAGATTCAGAGTCTCCAGT

GTTACCGACTACACCATAGGATCGAGCATAGCGGCCAAAAGAT

>NODE_2104_length_113_cov_42.304348

ATCAGTGATAGTTGCAGCACTTGTCGGTAGTGCATCCGGAGTGGTGGCATTTGATCTTGTCAGAGGAATC

CTTCGCCGCGTAGTGCTTAGGGCAGTAGTAGGAGTCGCAGTAG

>NODE_2105_length_113_cov_19.630435

ATGTGGGTTCGGTCCTCCAGTGCATGTTACTGCACCTTCACCCTGCCCATATCTAGATCACTTGCCTTCG

GGTCTAATGCTACGTACTCAGTCGCCCTATTCAGACTCGCTTT

>NODE_2106_length_112_cov_26846.527473

TGACTGGAGTTCCTTGTCACCGGAGAATTCCAATATCTCAGCGATCTTGGTGGTAAGTGTCGTCATCCCG

ACGGTCTAGAGTAGCAAATATATTTTGCATCCTTGATATCCA

>NODE_2107_length_112_cov_3414.043956

ATAGCGCGGAGGTACAGAGGTTGACCTCAAATCAGGTAGGACTACCCGCTGAACTTAAGCATATCAATAA

GCGGAGGAAAAGAAACTAACAAGGATTCCCCTAGTAACGGCG

>NODE_2108_length_112_cov_310.945055

GAGCGCGGTGATGATTACACAAGTATTTCATCGTCACTGACAGCTTGCGCAAGCAATTGCAACTGCTTCT

GATGACAAATACAGAGTTTTCCTGAGCGCTTGGAATTCTCGG

>NODE_2109_length_112_cov_268.000000

TGGTGGAGAGAGTCGGATTCGAACCCACGGAACGCGTAAACGTTCATCTGATTTCAAATCAGACGCAATC

GACCATCTCTGCCATCTCTCCGACAACCAGCTCAGCGGACCA

>NODE_2110_length_112_cov_266.538462

TAGAGAGCAAGTCTCATCACAAACAGCGGCGTACCTTCTCCCGAAGTTACGGTACGATTTTGCCGAGTTC

CTTCACCCGAGTTCTCTCAAGCGCCTTAGTATTCTCTACCTG

>NODE_2111_length_112_cov_158.593407

CTATAATTAGAATGTAAGGCTCTGTAGCTCAGTGGTTAGAGTAGGGGACTCATAAGCCCTTGGTCGCAGG

TTCAAATCCAGCCAGAGTCACCAGAGCCATGATGACCCTTAA

>NODE_2112_length_112_cov_112.692308

TTTGTGTGCGGCCTCGATGGGGGCTGTGACGAAGCAAAATGCATAGTTCATCTGATTTTCCAGTGAAGAA

CTAGTCTTAAGCAAATGTCTTTCGCTCCTATCCGAGCCCCTG

>NODE_2113_length_112_cov_54.219780

TAGGTGTTCGGGTTACAAGCGCGGCCGTTAGCTCAGTTGGTAGAGCAGTTGGCTTTTAACCAATTTGTCG

AAGGTTCAAATCCTTCACGGCCTGGAATTCTGGGTGCCAAGG

>NODE_2114_length_112_cov_31.626374

TTGGCACCGAGAATTCCAGACTTCACCCCAGTCATTAACCACAAAGTGGTGAGCGTCCTCCCGAAGGTTA

GACTACCCACTTCTTTTGCAGCCAACTCCCATGGTGTGACGG

>NODE_2115_length_112_cov_2.252747

TTCTGTTTTGAGCATGGGAAGGCGTGTGGCTGTTACGTGCTTCGCTGAAACTCCACTCCCATCTGTGGAG

GTTTGTGGCGGTTCTGGCTGGTTCTGTTTTGAGCATGGGAAG

>NODE_2116_length_111_cov_125155.266667

TTGGCACCAGATAATTCCAAGTTCAAGTCATCGATCTATTAGTACATATCAGCTTAATATATTACTATAC

TTCTACTTGATGCCTATTTGCAGTTCAAGTCCTCGATCTAT

>NODE_2117_length_111_cov_395.033333

GGGGCTGTATTGGTTTCGACATTTATAATTCTATTAATCAATAAGCAGATTTAAATACTTAAAACATACA

GTAATTTAACTTGAAATAAACGGTTCCAAAGAAAAACAATA

>NODE_2118_length_111_cov_308.422222

CAAACGCCCTTTTCGCGTGGGGTGAATGACGGGACTTGAACCCGCGGATGGCGGAATCACAACCCACTGC

CTTAACCACTTGGCTACACCCACCTTGATATTTCTCCTACA

>NODE_2119_length_111_cov_140.855556

AGGCCGCCAATCTCCTTGGCGTCCTTAGTTCAGTTGGTAGAACATAGGTCTCCAAAACCTAGTGTCGAGG

GTTCAACTCCTTCAGGGCGCGCCAGGGGAAACTCTGGTGGA

>NODE_2120_length_111_cov_34.888889

GCCCTCATAAGGAAGTCGACAAGGTGTTTGTCAGCACATGATGCTTCAGGGGTAATATTGTAGATATGTC

CTGGGTCAAATCATCGGGCGCTGCAAGTCAACCTCCCCAAA

>NODE_2121_length_111_cov_14.377778

GCCCGTCGCTCCTACCGATTGGGTTTAGATGTGAGCTGTTTGGATTGAGGGGGATTGTGGGTGACTACGG

TTCTTTGAGAGAAATTCCGCTAATTTTTAAACCTAGAGGAA

>NODE_2122_length_111_cov_13.766667

AGGCTTTGCCCTAGATCGTACCCGAACGTAACTGGCACTTTTCCCCGTTAGAATCTATTGTAGACGACTT

TGTAGAGACCGGGTATTGTAAGCTTGGAATTCCGGGTGCCA

>NODE_2123_length_110_cov_5530.213483

CACGGTGGATGCCTTGGCAGTAGGAGGCGATGAAGGACGTATTAACTTGCGATAAGCCTAGTCAAGCTAG

TAAAAAGCGCTTGAGACTAGGATTTCCGAATGGGGAAACC

>NODE_2124_length_110_cov_503.202247

CATCTAAGTACCCGGAGGAAAAGAAATCAACCGAGATTCCCTTAGTAGCGGCGAGCGAACGGGGACCAGC

CCTAAACCCTTAGATGTGATAGTGGAATGCTCTGGAAAGT

>NODE_2125_length_110_cov_43.370787

CTTGGCACCCAGAATTCCATGGCGGACAGACAGGGATTCGAACCCTGGAGACGGTCTCCCGCCTACACAC

TTTCCAGGCGTGCGCCTTCAGCCACTCAGCCATCTCTCCA

>NODE_2126_length_110_cov_31.898876

GTACCCGTTAATCTTGACTAAGTCCCTGAGCCGATAAGCATTACCCTGGTGAATTTGTTCGGATACTATT

GTGTAAGCTCGACTCGTATCTTTAAAGAAAGCGTAATAGC

>NODE_2127_length_110_cov_25.584270

AAGGGAACAAATCGCACCATCTTGTTTTAGTCTTGTTTCTTGATCGTAATCCTAACAAGGTATTTGTCGA

TGAAGAGGTCTGGCGCAGATGGAGAAGAAGAGGAACAGTG

>NODE_2128_length_110_cov_14.966292

CAAATACGGCATCGGCTTCTGGAAACCCGGATCAGGCATTATCCACCAGATTTGCGGGCACAGCCGATGT

GCTCGCATGACACCAACTCCCCTTGGAATTCTCGGTGCCA

>NODE_2129_length_110_cov_2.337079

GACAGTCCCAGGAACGGTTCCCGGTATCCGCCTTCGCGGGTCGTAGTTATCCTATGAATACCCTTCAAAG

TCTGTCAGTAGACCATCGGGACAGTCCCAGGAACGGTTCC

>NODE_2130_length_109_cov_1036.670455

AACTTCCAAGGCTGTATGGTCTAATTAGTATAGTGGTCAGTACACCACATTGTGGCTGTGGAAACCCGAG

TTCGATTCTCGGATTGGACACCATGGAATTCTCGGGGTG

>NODE_2131_length_109_cov_1024.409091

AAGCGGGCGCGTCTTTTGTCGCTCTTTCGAGAGACCGACTACGCCAATCTGTTTCTTCTTCACGGCCAAG

AGCTAACCGAACACATGTATGTATTAGCTCTAGAATTAC

>NODE_2132_length_109_cov_383.102273

ATTCCAGTGGCTCATTCGAGCGTGTGTCACCATGGGTTAGTATGACGGGTAAACGGCTAAGTTTTTTGCT

GAGATATAATCATCACCAGCAAGCCGCTCAGATCCAACA

>NODE_2133_length_109_cov_211.784091

GTCGCAATGAGGAAAGCCTTAACTTCCGCCGTTTCTGAAGGCCGTTCTCTATTTGTTTCGTTTTGAAGCA

AATGCGGCCCAATGCTGGACAGTGAAGTTCCTGAGCGCA

>NODE_2134_length_109_cov_98.704545

TTGGCTGCCGAGAATTCCAAGGTCTCGGTATAAAAGCAAAATAATCAAATGTTGGCCTTTTGCCTTTCAT

ATGACCGTCGTGAAACTTGCGATAAATCGTCATCGACCT

>NODE_2135_length_109_cov_76.545455

ATCCCTCACACTGTTTGGTACGTAGTTTAAATCTCGCGTTCCAAATAAAAACAGTTCAAGCTAAGTCACC

ACCTAACTTGAACCGCTTTTTCACACTGACAACATCTTC

>NODE_2136_length_109_cov_52.784091

AACGGCAAGTGCGAGGTGTCCCAGTGCTGCGACCTGTTCTGCAACTCCATCGGCTGCCCTGGCGGCTACA

TTCCGATCAAGAACGCCAACAAGGTTGAGTGCTACGACG

>NODE_2137_length_109_cov_24.636364

GAGGGGGGCTGGCAGCGCTGACGGATGGACTCTTTTGTTGCTGCTGTTGTTCTGCTTCGGGCGACTCAAT

AAAAACTGGCTGAGACTCCTTGAAAAAAAAAAAAAATTT

>NODE_2138_length_109_cov_22.602273

AACCCGACGAAACTCTTCATGACGAGGCGCGTGTGTAGATCTTCTTCTTTCGATTGCAGCTGGCCGCTGT

AGTAGCAGAGATGCCGAAACCTGCCTGTAGATAGCTGAT

>NODE_2139_length_108_cov_67759.195402

AGCATGCTCCCAGGAGTATCCGTCGCTGAAAGTTTTACAAACAGACACTCCGACAAGCATGCTCCCAGGA

GTTTCCCGGAAGCGCATGATGCGTTCAAAGTTTTGATG

>NODE_2140_length_108_cov_12495.747126

CTTGGCACGAGAGAATTCCAGTCGCTCAGCGAATGTCGTCTTCACTTGATGTTCAGGCTCATAACCGTTT

CGTTTGCCATCATTACGATGGCACGGTATCAAAGTGAA

>NODE_2141_length_108_cov_8907.000000

TTTCTAAGAGTCGAGTTGCTTGGGATTGCAGCTCAAAGCGGGTGGTAAATTCCATCTAAAGCTAAACACG

GGTGTGGAATTCTCGGGGCCAAGGAACTCCAGTCACAC

>NODE_2142_length_108_cov_6935.816092

CCAGGCAGTCAGACAGTACGATTACGATGTAGTTACAGCTCCACAGCGCAGTTAAGGCGTCGGCCTACTT

GGTCATCTTCACTCTGCCTGACCTAAGTTCTAACCATT

>NODE_2143_length_108_cov_6859.172414

CTGTCCGGTGAGCTTCAAATTTAATTAAACTCTGAGTCTCGGCACTGCAGAAAATATGTCGAAATCAGTG

AGGACGTGGAATTATCTGGTGCCAAGGAACTCCAGTCA

>NODE_2144_length_108_cov_5774.896552

CGGGTCAGATGATGTCATAAAATTATCTACATTGTCGGACTGCTGAGTGAGGACAAAACTTACGCGCCGT

CTAGGCGGCTGATGATCGTCAGATGATGTCATAAAATT

>NODE_2145_length_108_cov_2061.586207

ACACCGCCCGTCACACCATGGAAGCTGGTTATACCCGAAGTCGTTATTCTAACCTAATATACTTGGAGGA

AGGCGCCTAAGGTAAGGCTAGTGACTGGGGTGAAGTCG

>NODE_2146_length_108_cov_1082.517241

CTTGTCACCCGATAATTCCAGGTCCTCACCCATGTTTTTATCATCACAGAGTTCAGGCGTAAATTACCTT

ATGTTAAGCTCACTGGACCTGTACCTCCGCGCTATCGA

>NODE_2147_length_108_cov_254.471264

GAGGGAAAGGCGAAAAGAACCCCTGTGAGGGGAGTGAAATAGAACCTGAAACCGTGTACGTACAAGCAGT

AGGAGCGGATTTATTCCGTGACTGCGTACCTTTTGTAT

>NODE_2148_length_108_cov_52.965517

CTTACAGAACGCTCCTCTACCGAGCACAACAAGTTGTGCTCCCGTAGCTTCGGTATATTGCTTGAGCCCC

GATAAATCTTCCGCGCAAGCCGACTAGACCAGTGAGCT

>NODE_2149_length_108_cov_36.126437

TTAAGCTAGATGTGAAAGCCCTGGGCTCAACCTGGGAATTGCATTTAGAACTGGCAGGCTAGAGTCTTGG

AGAGGGGAGTGGAATTCCAGGTGTAGCGGTGAAATGCG

>NODE_2150_length_108_cov_30.425287

TAGTCCACGCCGTAAACGATGGATACTAGTTGTTTGGTCTTTGGACTGAGTGACTAAGCGAAAGTGATAA

GTATCCCACCTGGGGAGTACGTTCGCAAGAATGAAACT

>NODE_2151_length_108_cov_26.114943

TCCACATACCAAATACGCTTCGGTAGTGGTGATGGTGGTGTCGTTTTGTATCTTCAGGACTATGCCGGTT

TCGGTATGCCCCGCGGGAGACTGGAATTCTCGGTGCCA

>NODE_2152_length_108_cov_17.551724

GAAAAGGATTGGCTCTGAGGGCTGGGCACGGGGGTCCCAGTCCCGAACCCGTCGGCTGTTGGTGGACTGC

TCGAGCTGCTTCCGCGGCGAGAGCGGGTCGCCGCGTGC

>NODE_2153_length_107_cov_7495.151163

TTGGCACCCAGAATTCCAGTCCATCAGATAAAATATAGTGATAAATTGTTTGTCTTCATCGATTTCACTC

GGTGGATTGACAGAAAGTTTTCATCGTCACTTGGGAC

>NODE_2154_length_107_cov_857.872093

TATATATCTGAAGCTTAGAAGCTTTTCCTGGAAGTATGGCATCAACTGCTTCATCACCGTAGTGACTCGT

CTCGTATCTCAGAATTGACAACCCGGATTTACCTAAG

>NODE_2155_length_107_cov_714.546512

AGTCGAAGAGCGCCGCAACTAAAACTGGCGAATCTACACACCCATCTCCATCGTGCTTCTTTAAGAAAAG

CAGCAATCAAGAGAAGAGCGGAGACTCAAACCACGGC

>NODE_2156_length_107_cov_438.360465

AAGACGTTATGCGGTATTAGCCATCGTTTCCAATGGTTATCCCCCACACAAAGGCATATTCCTAAGTATT

ACTCACCCGTCCGCCACTCGTCATCTTCTAGCAAGCT

>NODE_2157_length_107_cov_112.465116

AGTTAAAAAGCTCGTAGTTGGCCACTATGATGGCTAAAATGATATAATGAGCCTGAAACACATCATGTGG

CCAACTCTAGCTGAGTGACTGGAATTCTGGGTGCCAA

>NODE_2158_length_107_cov_17.360465

GGCCAGGAGATGGTGACGGATCCTCAGGGTTGTTCGATCTTAACGGATTGATCGGGCAGCTGCGGGGTCC

CTGGAAATAGCCCTCAGGAATACTCGGCTGCCAAGGA

>NODE_2159_length_107_cov_15.383721

TTCCAAGCAATCAGCCAGTGGTGGTGGGGTTTCCACACCTTGCGGGGCGGTATCGACCCGGGGGATGTCC

GTTTTCGTCCCCGGGTTGTTGGAGTTGTAGATGTATG

>NODE_2160_length_107_cov_1.720930

CCGTCTTGCAACGACGTGCCATGTGTTTTGGCCACGGTACGTGCAAGGAGAAGGGTTGAATGGAGCCGAT

TGAGTGGGGGGGGGGGCCGTCTTGCAACGACGTGCCA

>NODE_2161_length_106_cov_5865.788235

GGGTGGGTTTCCCCATTCGGACATCTCAGACTATAACGGTTTTTATCACCTCATCTGAGCTTTTCGCAGA

TTAACACGTCCTTCATCGCCTCTAACTGCCAAGGCA

>NODE_2162_length_106_cov_5605.564706

TCCGAACGTCAGCTGCTGGAGCTCTGTGATTACCTCTACAAAACCAAGATCTGTTCCGAACTCATTGTGT

AGATACGAGCATTGTACTTTCTGTATGAGGGCCTGG

>NODE_2163_length_106_cov_1730.247059

ACCTTGGTATTCAGAGGCGAAGAAGGACGTGTTTACCGGCGAAACGCTTCGGGGAGCTGGAAACAAGCTA

TGATCCGGAGATATCCGAATGAGGAAACTTTTTAAA

>NODE_2164_length_106_cov_1343.023529

CACCGAAGCTGCGGATTCAGAATTTATTCTGAGTGGTAGGGGAGCGTTCTGTAAGCGGTTGAAGGTGTAC

CGGGAGGTATGCTGGACGTATCAGAAGTGCGAATAC

>NODE_2165_length_106_cov_500.094118

AACAACAATCTCAGCGGAACAAAACAACAATCTCAGCGGAGCAGACAACCAGCTCAGCGGACCAGACAAC

AAACTGACCGGTCCAACAACAAACTGACCGGTCCAC

>NODE_2166_length_106_cov_408.352941

TGGTACGAACGGAGAGACTTGAACTCTCACAAGAAACCTTACTAGAACCTAAATCTAGCGCGTCTGCCAA

TTCCGCCACGTTCGTGCCGTCGGAAGTGGTGGAAAA

>NODE_2167_length_106_cov_126.388235

ACAGCAGGACGGTGGCCATGGGTCTAGTGATGATGATAATGATGTTACCACATTAGCCCATACTGGTATA

CCTGAAATCTGAAGCCTTGGAATACTCGGGTTCCAA

>NODE_2168_length_106_cov_126.129412

TTGGCTCCCGAGTATTCCAAGCCCGTCATATTGAAGACTGAAGTTGCGATTTGGTTCCACATCGTGTCTT

CGGAACATATTTTTTGCTGAAATAATCACGGGGCTC

>NODE_2169_length_106_cov_105.270588

AGGTCGGATGAAGATAGCCCAACGAAACATTATGCTGACTTATATGCCATGATCATATTAATCAAAAGCA

ACGTTGCTATTCTGACGATTTGGAATTCTCGGTGCC

>NODE_2170_length_106_cov_97.117647

TGTCGGTTTGGGGTACGGTTCGATATATCATAAGTTTAGAGGCTTTTCCTGGAAGCAGGGTATTTGCAAC

TTCAACTCCGTAGAGTCTCGTCTCGTGTCTCAGGCA

>NODE_2171_length_106_cov_92.164706

CACCCGTCCGCCGCTCGTCATGGTGCCCAGAAGAGGACTCGAACCTCCACACCGTTGCCAGTACTAGCAC

CTGAAGCTAGCGCGTCTACCATTCCGCCACCTGGGC

>NODE_2172_length_106_cov_92.000000

GGAACTTGATTCCCCTAAAGGGCCGTTGAAGACTACGACGTTGATAGGTTGGGTGTGGAAGCGCTGTGAG

GCGTTGAGCTAACCAATACTAATTGCCCGTGAGGCT

>NODE_2173_length_106_cov_79.847059

TGGCAGCCGAGATTTCCATGGTGGACGATACTGGGCTTGAACCAGTGACCCCCGCCTTGTAAGGGCGGTG

CTCTCCCAACTGAGCTAATCGTCCGTCTCTGTGGGT

>NODE_2174_length_106_cov_23.611765

CTTGCCGCTGTAAAGGCGGAAGTCGTACGTCGCCGGGGGGATGTCGCCGCTGCTAGGGTCCTCGACAACG

CGAGCCTCCGGAAGGCCGGGCAGTTCCTGCTGCAAG

>NODE_2175_length_106_cov_21.505882

TGGAATTAACTTCAAAAAATTAAATCAACTCACACCAAAAAAACGTATCCAATCAACTCCATTAAAAAAA

AAAACACAAAAATCTAATAATACTCTGCAAACAAAC

>NODE_2176_length_105_cov_14510.333333

CTTTTTGATTCTTCGATGTCGTGTCGGTGATGCCGAAACTTAAAATTATGCTGACCTGACTCTGAGGAAT

CAAAACAGACTACGATGGACTGAAGGCACTGGAAT

>NODE_2177_length_105_cov_3593.452381

AAAACTTCCAAGGCTGTATGGATATGTGATGAAACTCACTGACACCTCTTGTTTGAGGGCCAAAAAGGCC

TTGTGAACGCAACCCTACAATGAAACTGATATCCT

>NODE_2178_length_105_cov_705.952381

TTGATTGGCCTTTCACCCCTAGGCACAGCTCATCCCGATCTTTTTCAACAGATGTGGGTTCGGTCCTCCA

GTTAGTGTTACCTAACCTTCAACCTGGCCATGCCT

>NODE_2179_length_105_cov_497.380952

TTTGTTCGTTTCTTGTTGCGGCCCGATGATGCTCTACATGAGATTACCGAACCGCTTTGCGTTTCGTTTG

GTGGCATTGGCGAGGAAACTGAAACGCTGAGGACC

>NODE_2180_length_105_cov_110.583333

CCTGCTCCGTCCCCCCATCGCTGGTGGGGGTGGAGGGACTTGAACCCTCACGATTTGCATCAACGGATTT

TAAGTCCGTTGCGTCTACCATTCCGCCACACCCCT

>NODE_2181_length_105_cov_107.416667

GCTGTTTAAAAGTTAGTGGAATGCTCTGGAAAGTGCAACCATAGAGAGTGATAGTCTCGTACACGAAATC

TATTTTAAGGTTATTCGAGTAGGTCGGGACACGTG

>NODE_2182_length_105_cov_80.511905

GCTCGCGTACCACTTTAAATGTGGTACGCATACGGAGAATCGAACTCCGGGCAAAACCTTGGAAGGGTTT

TATGTTACCACTACACCACATGCGTCTCAAGCGAT

>NODE_2183_length_105_cov_17.357143

ATTCCAATCCCGAGACCACAACCCTCACGTCCTCTCTCCCTCGAACTCATCCGTCCGATTGGAATCACCA

CCAGCACCACCTCACGTAGGTGGCCTGTGGCTAGA

>NODE_2184_length_104_cov_3149.626506

ATGACCCGCTGACGATGACTCTTAATACAAAAACCACTGAATTTCTGTTTATATGTGTGGATATTCAAAG

AGACACCTGTGTTTCTGAGGGTTTGGAATTCTCG

>NODE_2185_length_104_cov_882.096386

GACATGAGTAACGATAAGGGGAGTGAAAAACTCCCCCGCCGAAAGACCAAGGTTTCCTGTCCCATGTTAA

TCAGGGCAGGGTAAGTCGGCCCCTAAGGCGAGGC

>NODE_2186_length_104_cov_403.385542

CATTCCGTCAGGAAAAAATTAGAGTGTTTGTAAGCTTAAATGGTCCTCACCAAGATTGGACTGCTTTTCT

TCATCACTCGGAATTACACACCTGACCTATCAAC

>NODE_2187_length_104_cov_227.722892

TTTCAAATGATGTGAACACGCAATAATTCTGACGTTAGTGAGGAATAATTTCGCGAGCTTTTTAACACTG

ATGAAATGGAAATCTCGTGTGCCAAGGAACTCCA

>NODE_2188_length_104_cov_80.301205

CTTTCCGTCTAGCCGCGGGTACGCGGCATCTTAACCGCGAATTCAATTTCACTGAGTCTATGTTGGAGAC

AGCGGGGAAGTCGTTACGCCATTCGTGCAGGTCG

>NODE_2189_length_104_cov_72.626506

TATTCTTGCACTTTCTCTTCTTCAGTGAAGGGGGGACGGAGAAGGCTAAGCAAGCTTGGTGTTGGTTATC

CAAGTGAAAGTGCGTAGGGTTGAATCTTAGGTAA

>NODE_2190_length_104_cov_64.759036

CTGCCTCCCGTAGGAGTCTGGTCCGTGTCTCAGTACCAGTGTGGGGGATCTCCCTCTCAGGACCCCTACC

CATCGTTGCCTTGGTAAGCCGTTACCTTACCAAC

>NODE_2191_length_104_cov_51.855422

AGTAACACGTAAGTACGTGATGTTTTCTATTCATGTCAGTCTTCATACAATTCACTTCATAGTTCTAAAA

ACTACTTGGGTGTTGTATGGTTAAGCCTCACGGG

>NODE_2192_length_104_cov_46.867470

ATTCCAAATGACACTCAAACAGGCATGCTCCTCGGAATACCAAAGAGCGCAAGGTGCGTTCAAAGATTCG

ATGATTCACTGAATTCTGCAATTCACATTACTTA

>NODE_2193_length_104_cov_27.481928

TTTCTCGGGAATAGTAATTTCGAATTATTATCGACGCAAATCGTTTGTAGACGACTTAATTGGGACCGAG

TACTGTAAGTGTGAGATGGAAATCTCGGGAGCCA

>NODE_2194_length_104_cov_16.180723

TGAGGCTATTACCAAGGTTAATCATGACCGTGGGTACCGAGGACGTCGAGATGAAGGAGGAGGTGGCTGC

CGAGGAGGTAGCCAAAACTTTCAGCGACGGATGT

>NODE_2195_length_103_cov_14556.402439

GCAGTCAGACAGTACGAATACTATGTAGTTTAGCCCCACCGCGCAGTTAAGGCGTCGGCCTACCTGTTTG

TCTTCACTCCGCCGACCTGGTGGATGGATATCA

>NODE_2196_length_103_cov_11704.475610

TTGATAGGTCAGGTGTGTAAGTCCGATGATATTTACATACTTTGTCCGAATCTCGCATGTGGTACTTTAT

CGCTACGTATGAGGACTTGGAATTCTCGGGGCC

>NODE_2197_length_103_cov_1089.487805

CCTTGGAAGCCGAGAATTCCATAGAAGCTTTTCCTGGAAGTATGGCATCAACTGCTTCATCACCGTAGTG

ACTCGTCTCGTGTCTCAGAATTGACAACCCGGA

>NODE_2198_length_103_cov_42.817073

CTACGATTGCCCTAAATACTATGAGCTAGTGGACGACGCTTCCTCTACCGTGTGCAAGGACGACAAGTGC

AGTAAAACACAATGCTGCCAGAAGGAACCTGTC

>NODE_2199_length_102_cov_9933.888889

TAGGTGTTGTTGGTGTGTGTGTTGTTGGTTTGCAGCGGGGTGATGACACTACAAATACCAACTTTCGAGT

CTGAAAACATCATGACTGACAGAACCTCTGAC

>NODE_2200_length_102_cov_6355.382716

AGAGCAGCGATGGTTTTCAGACTCGAGTGTGGCTTTTGCCCATTTGATCTGTTTCGATGACCTGCGCTTT

CTCTTTCGAGAGAGAGCGACTCTGTTGGCGAG

>NODE_2201_length_102_cov_4127.246914

ATTTTTTCACAGTAAACGACGCTAACGCTTCAATCTATAACAGAACGCTCCCCTACCGATATAATTATTT

ATTATTATTTTTAATATCTCACAGCTTCGGCA

>NODE_2202_length_102_cov_549.111111

GTTACGACTTCACCCCAGTCATGAAACACAAAGTGGTGACCGTCCTCCCGAAGGTTAAACTAGCCACTTC

TTTTGCATCCCACTCCCATGGTGTGACGGGCG

>NODE_2203_length_102_cov_404.308642

GACCTGCACGAATGGTGTAACCATGGCCACGCTGTCTCCACCCGAGACTCAGTGAAATTGAAATCGCAGT

GAAGATGCTGTGTACCCGCGGCTAGACGGAAA

>NODE_2204_length_102_cov_314.259259

AGTCACCGAAGCTGCGGATTTGAACTTAGGTTCAAGTGGTAGGGGAGCGTTCTGTAAGCCGTTGAAGGTG

TGTTGTAAAGCATGCTGGAGGTATCAGAAGTG

>NODE_2205_length_102_cov_187.975309

GCCCAAGTGGGATTAGCTAGTTGGTAAGGTAATGGCTTACCAAGGCGACGATCCCTAGTTGGTCTTAGAG

GATGACCAGCCACACTGTGGAATTCTCGGTGC

>NODE_2206_length_102_cov_95.246914

TGAAGTCGGCAAGGACTGCACGAAGACCTCTCAAGATACAAGGCGGTATCCCGAGTACGAACTCGCCCCA

ACTCCTGTACGAACCGGATTGTGAAGACAGGC

>NODE_2207_length_102_cov_23.469136

CTTGGCACCCGGAATTCCAACGCGTAGCTAGTTGGTTGCAAGGTCAAGGATTTTAAGTGATGCGTGAGCT

CACGTCGCACTTGTGTGGAATTCTCGGTGCCA

>NODE_2208_length_102_cov_12.469136

CTCGCACCTCGGCTTAACCATGGCGGTCGGCAAGACGAACAAGAAGGCCGGAAGGAAGGGCAACAAGAAG

AAGACGGTGGACCCCTTGGAATTCTCGGTGCC

>NODE_2209_length_101_cov_6110.012500

GGCACCGAGAATTCCACACTTCCACATCTTGCCTATCAACGTTGTAGTCTTCAACGGCCCTTCAGAGACT

TTAAAAGTCTAGTGAGAACTCATCTCGAGGC

>NODE_2210_length_101_cov_1085.312500

GGAGTTCCTTGGCTCCCGTGTATTCCAGGGTTTGAACAGCCCTGTACTCGCGGGCATGTTAGACTCCTTG

GTCCGTGTTTCAAGACGGGGTAAAGAAAACC

>NODE_2211_length_101_cov_355.162500

ATTGGAGTCGTTGGATCTTCATAGAGGGGGCACAGAGCCGTTGAGCTGCCATACCCCTTCCTTCGAGAAG

TTCATCATCTGATTTTTTTCCTCACCGGTTT

>NODE_2212_length_101_cov_78.825000

GGAGCATGCTTGTCGGAGTGTTCGGTGGTGGAGCGGATGCGGTTGCGGGCACAGCCGATGTGTCCGCTTC

CGCGTACTTACCCTTGGAATTCTCGGTGCCA

>NODE_2213_length_101_cov_67.712500

GGACACCTCGCACTTGCCGTTCTTGCACTCCACGTTCTGGGCGTCGTAGATCGGAGCGTATCCGTTGGGG

CAGCCGATGAAGTTGCAGTACAGCACCGGCT

>NODE_2214_length_101_cov_50.375000

TTCACCATCTTTCGGGTCCCATCATTTAAGCTCTACCGCAGTCCGCTCCGATGAGCAGCCGGGCGATGCT

GCCAGCCCCCTAAAGGGACCTTGGCACCTCT

>NODE_2215_length_101_cov_43.725000

GGCACCCCGAGAATTCCAACATTTAGTCCAAGGCGCATGCTGCGTTCGAAAATCTGATGTCTCACGGATT

CTGCAATTCGCATTACGTATCGCATTTCGCT

>NODE_2216_length_101_cov_17.450000

TTGGCACCCAGAATTCCAACACCAGACGCAACCTCACTAACCGCAGCAGTTCTGCGATCCATCTCCTCGA

AAATCTGCGTCTTGGAATTCTCGGGTTGCCA

>NODE_2217_length_101_cov_16.400000

GGCACCCCGAGAATTCCAGCGTCACACCCTATACGTCCACTTGCGTGTTTGCAGAGTGCTATGTTTTTAG

TAAACAGTCGCAGCCACCTGGTCACTGCGGC

>NODE_2218_length_100_cov_910.189873

ACTTCGGGAGAAGGTACGCTCTCTAGTGTGAAGGACTTGCTCCGTAAGCACATGAGAGTCGAAGTAACCA

GGTGGCTGGAACTGTTTATTAAAAACACAG

>NODE_2219_length_100_cov_171.405063

AAGTGAAATCGAGTAGGTCGGAGCACGTGAAACTTTGACTGAATATAGGTGGACCATCATCTAAGGCTAA

ATACTCCCAACTGACCGATAGTGAACCAGT

>NODE_2220_length_100_cov_107.974684

AACTGCGATGGAGGGACGGAGAAGTGAAATGTATGCGTGCTGACAGAATAGCACGTCTAAGGCTGTAGGT

ATAAGGTAGTAGGCAAATCCGCTATTGGAA

>NODE_2221_length_100_cov_95.683544

CCACAGCGCTATTGTCGCTAAACAAAAAGGTTACAATCTTAGAAAGCTATAAATATCGTTTTTTCTTTAT

TCTGATTCAAGTAACACGTAAGTACGTGAT

>NODE_2222_length_100_cov_29.177215

TTATTATTATTATTATTATTATTAATATTAGTATATCACGATTGGTCTGACATGCGACAATGCTCTGCGT

TGTTATCATAATCACCATAAACCTATCCCC

>NODE_2223_length_100_cov_18.392405

TCGGACCTCCGATCAAGCAGGAGGACCCGCTGAACTTAAGCATATCACTAAGCGGAGGAAAAGAAATCAA

CTGAGATTTCCCTAGTAGCGGCGAGCGAAC
