## Supplemental Document SD3 for "Extreme GC3 Codon Bias in a Novel Brown Seaweed Virus Results in Pseudoambigrammatic Characteristics"

### Supplemental Document SD3: SalaUV-NL1 variant sequences

**>SalaUV-NL1-1-9_SL**

ACACAATGTCTTCTACTTATTAATACTATCATGCTGACCAACCTGAAGATGAACATCAGCCTGCTCCTTGCACAGGAACTACAAGATGGACAGCAAGCTCCTCCTCTTGTCCTGGACCTTGGTCACCATCCTGTCCAAGGCCACCGCCCTCTTCGTGTTGAAGGCGATGTCCAGCTTGATGAGGGAGTTGAGAGTGGGCGCCTTCTCCCCGATGGACGAGAAGATCTCCGTGACCAAGCCCGTCGTGACCATCTGCACCATGACGACGAGGGTGTCCGAGGCGGAGTTGTACCCCTTGTTCTCCTTGAAGTCCCTCAGGGCGTCCATCTGGGCCAGGCCGGACATGCCGGTGGTGGGGTTGATCGTCCCGGTCATCTTCATGTAGACGGCCGTGTACACCATGTTGCACACCACCGCCAGCTTCCTCTTCTTCTCGTCCCCGATGAAGTTCCTCTGGATGTCGTTGACCGTGTTCATCACCAGCCTCTGGTCGAACGAGTAGAACTTCTCGAAGACCCTCTTCACTGAGGTGTAGACCGTGCCCATGTACTCCCTCAAGGACATCCCCTCCTCCAGGGGCTCCTCCTCCATGTGGGCGGTGGTGTCGATCTCCTTCAACCCCGAGAAGGCCCCGGCCAGCCCGGCGAGGGAGGTGAAGTCCATGGACATCCCGCCCATCTCCTCCTTCCCGATGGAGGAGAACATCTCGGCGTTCAGCTTGGTGGCCATGGAGGACGAGGAGATCCCGAAGTCCGTGGCGCTGATGGCGCCCATGACGTTCTCGACGCTGTAGTCCTCCACGGCCTGGGACACGTCCATCATCTCGTATATCGTGGAGTCCACCTCCGCCTCGAACTCGGCCTCCTTGATCCTGTTGTCCGGGGCGAACATGGAGTTGTCCGCTATCCCCAGGTCGTTCAGGATCTCCTCCACGTCCGACAAGTCCAGCTTGCTCGAGACGACGAGCTCGGAGAACTTCCTTATGTTGGGGTTCTCAGAGTCCTCGTTGAACGGCATCTTGTAGCTGTTGGTGCCGATGTGGAAGAACACCTTCCCCCCCAGCTCCCTCTCGTTGAACATGGCCAACACGGTCCACTTGGTCATCCTCCTGGTCAGGATCAGCTCCGTCTTGGGCCAGATCCTGTAGACCTTGATCCACAGGTTCCCGATCTTGGCGTTCTTCTCCACCCCCAAGTTGTCCGTCCTGGGGGCCCAGGGCCTGATCTGGATGGTGGAGAGGTCCTTGTGCATCAGCCTGGTGATGTTCCCCAACTCCACGTCGGAGCTGCTGACCTTGTACATCCTCACCGAGACGGTGTGCGGGGACGCCTGCCTGTCCTCGTGGATGATCCCGAGGTGCACCAAGTCGGTGTAGTACCTGGTCTTCATCTTCCCGTTCACGTCGACCTTGTAGAACACCTTGAGGGACTTGGCCCCCTGGAAGGAGTACTTGTCCTCGTCGGTGGCCGAGACGGCCATCTTCGCGAACCTCACCTCCAAGGAGTCCGACGGGATCACGGTCAGCAAGGGGTTGTCGATGCCCCAGGGATCCCTCACCAAGTTCTCCATCGGGCCCTCCGTGAGGTCCCCCCTGAACTCCTCCACCTCGAAGCGATCCACCCTCCCCCTCATCGACAAGGCGCTGTTGAACCTCTCCTCCGACACCACCCTGAGGGACTTCTTCGGGACGAACATCTGCAAGATCCCGCGGTTCGAGTACTTCTCCAAGTAGAGCATCCTCATGTTGGTGAGCGCCTCGGAGTTGTCCGGGAAGGAGACCATCATCTCCATGTTGAGGTCGGAGGTCATCTTCAGCCAGTACATGATGTAGCTCTTCAGCATCGTCAGGGGCCTCCTCACCCCCGGGAAGGCCTTCTTGATCTGGTATATGGGGTACTCGATGAACTTCTCCAACTTGACCCCCAGCGACTCGGAGATCTTGTTGGCCTCCGTCATGGCCGAGTTGCTAGCGTTGGAGGTGGCCCTCATCATGTACCTGAAGACCTCCATGTCGGACACCTTCACCCTGTCCTTCATCGGGGTGAAGTTGATCTTCCTGGACTTGTTGTGGCTGTAGGAGTGGGAGGGGATGGCGATCTCGGCCTCCTTCTCCGCGATCTCCCCCATCTCGCAGATCTCCTTGAAGGGGGCGGTGAGGTGGAACTTGGACTTGGTGTCGTCGTTCTTCAGGATCTTGTCCACCACCCCCACGAAGTCCGTGGCCACCACCCTGGAGGAGGAGATCAACGCGATGTTCTCCCCCGCCATGGCCAAGGCCCTGATCAAGATGTGGACCCTGATGGAGTCGTTGAACCCGAACTTCCTGTCCATCTGGAACATGTAGGTGTCCATGTTGGAGAAGAAGCTGGCCGAGTCGCTGCGGTCGGAGAAGGACAAGGCCGTGTTCCTCTCGGCCAGCTTGTCGATGTCCTCCCTCGTGATGGACCTCTTGTTCATCAGCCTCTTCTTCATCTCCTTGATCTTCGTGTCGAACCTGAAGGGGAGGGTCATCTTCGTCTTGCCGTTCATGGAGGCCATGTAGTCCCTGTCCTCCTCCATCTCCTCCTCGGTGTAGTCCGGGGAGGCGGCCATCTCGACGTCGGTCATGTAGGCGGCGTAGTACTTGACGTAGAACTCCATCACCTTCTCGGAGTTGCCCTCCTTGTACATCAGCACCTCCGGGCCGGCGATCATCTGCTCCACCACCCTGTCGGTCGACATGTACCCGAGCTGGAAGGGCAGGGAGTTCCTGTCGCACTCCAGCTTCCTCATGATCTCCTCCTTCCTCTCCCCGTCGATGGCGTAGATGGACGAGAGCGCCACGGACTGCAGCTCGAACATCACCTTGATCGTGGGGATCTTGACCCCGTTCTCGAACGCGTTCCGCATCTGGGATATGGAGGACTTCACCGCCTTCTCCGGCTCGGTCATGTCCACGATGTTCTGGGAGTTGTAGACGGCCTTCAGCACGCTGTCCGTGGCCCTCTTCCCGGTGGAGAAGTAGGAGTTGAACTCGGTCAAGATGGACAAGGCGGACTTCTTCCAGTTGGTCGTGATGTTGGCCATCCTGTTGGTCATGTCGTGCACCTGGAGCATGGACACCATGAAGGCCGTCAAGTCGGACATCTTCGTGTACCTGACCAGGAAGATCTTCGTGAGGTCGTCCGAGGAGATGAGGGTGGTGCACGCGGCGTCCCCGTACGACTTGGAGAACAGGAAGCACATCACCGAGTCCATGACGTCGTCCTTCGCGCAGTGGAACAAGGAGGAGAAGTAGTGGAGCATGCCCTGCCCCATCCCGGACTTGAATATGGCGGTCCCCCTCGAGGAGACGAAGTTCTCCTTGAGCCAGTTTATGTTCTTGTCGTCGGTCTTCTCGTTCCACAGCTTGTCCGTCTGCCACCTCCTGGTCAGGGTGGGCGGGAGGAACATGGTCTTCGTCGCGAAGGCCCGGATCACGGTGACGCAGAACTCCTTCAGGTTCACCGGCAAGGGCATCATCAAGCAGAAGTAGATGAAGCTATCCATCACGAACCCGGGGGACCACCTGGAGGCGTCCGTGTTGAAGGACAGGATGATGCCCGAGGGGGAGGTCTCCGACTCGTCCTTGTGCTTGTTCATCTCCAAGATCATCTTCTTGTACACGCTGGTGGCGTTGGACTGGAGGTCCTTCTTCACGTCCGCGTGCGTGAGCATCTCCTTCTCGTGCAAGGAGCAGAGCTCGTAGAACATCTTCTCCAGGAACTTCACGTGCAACCTCAACTTGGCCGCCTGTATCAGGATCTCCCTCGGCCCCCCGATCTGGGACTTCGGGAAGAGCTTGAACACGGCGTTCACCACGTCCGTGCAGGTGACCATGTCCATCAGGAAGATGGACTCCAACTCCTCCACCTGCTCGCACATCGTCTCGAAGGCGGTCGCCGTGGTCAGCTTGTCCGAGTAGTCCACCGACTCCTCGTACAGGGGGCACGACTTCAGGGACGACCTCATGGACATCGTCTCCCTTATGGTGGTGGTCATCGCCCTCATCAGGCCCTTGTTCAAGTCGGGGGAGTGGATTATCTTCATCCTGAGCTGGTAGCCCATGTACATGACGAACTTCTTGTTGAAGGTGTGGGTCTGCATCTTGTTCTTCGCGAGGGCGTCGAAGTCCGCGTCGCCCGTGGAGGCCCCGGGGAACAGCTCCGACATGGACAAGTAGTGCCTCTCCTGCTCGGACATCTTGGTCATGATGGGCACCAGCCTGTGGCCCGACGTCCCCTTCTCCTTGTTGAAGAGGTTGCTCATGTAGATGTCGTTCATCACCTCCGCGAACTCGACCAGCATCCCGGCCACCTTGAACGAGGGCATGTATATCCTGTCGTAGTCCGCGAACTCCGAGGCCATCTTCTCCGCGTACTGCTTTATGATGTCCGGGGCGGCCATGGTCAGCCTCTCGGCGTAGTCCATCTGCATCGCCCTGATGGAGGCCTCGAACATCGTCCTGACGGGGTCCTTCATGATGTTCTTCACCATGCTCACCTTATCCGAGGCGAACGCCGTGAGGCCGTGCATGATGTACCTGTTGTACTGGAGCGAGGTGGACACCCCCCTCTTGTGGCACATCATGACCAGGATGGGGACCATCACCATGGCGGACCTCCTGTACTCCTCCCTGGTCATCTTCTGGGCCCTGGTGCTCTCGAACTCCTCCCTGTCCGCCATCACCTGCGAGTGGATGGCCGCGCTCCTCTCGTAGAGCATGGAGTAGTGCTCCACGTCCGCGACCGAGACGGTGAGCCACTCCGTCTGGTACACCTCGGAGTACTCCTTCGTGTACTGGTTCCTGGCGATGGTCACGTCCATGTAGGACTTGCAGTCCTCCTTGCTCTTCAGCTTCTGGCACTCCATCCCCTCCATGTGCTTGTAGGGGGAGAATATCTTGACCCTGATCTGCTTCTCCTGGGTCAGGGCGGACCCGGCCTTCACGGCCAAGCAGTAGTCCCCGAACTCCTTGATGACCGTGAAGCGGGAGACCTTCTTGCCCGAGTCCCCGTTCCTCTTGGCCCTCCTCCTCCCCTCCAGCCTGGCGATGTTCCTGTAGAGGTCCGTCCAGAAGTGGGTCACCTCCGCCACGGCCAACTCGTTGAACATCTTCATCTGGGGGTCCTTGTACACCTTGGCCATGTTGTTGGTGAACTCGAAGGTGGCGTCTCTGGACATCTTCTCCACCATGTTCTCCATGAACTTCCTGTCCAAGAGGTTGGCCCCGATCCCCGCCATGTTCTCCTTCTTCTTGGTGTTCCCCGCCTTGGTGCAGTCGTTCAAGAAGATGGTGCCGTCGAAGAGGGAGCCCCCCTTCGAGTCCGTCACGTGCATCTTGACGTCCTTCATGTCGTTCAACTTCTGGGGGATCGGCATCGGGAAGGACTTCCCGTAGTCCTTCTTGGCCCTATTCATCTCCTCCGTGGGGGACAGCTCGTCCATCTTGTTCAACTCGGAGACGATCTCCTCCTGCGAGGTGCAGACCGCCTTCCTGATCTCCTCCGTGTACTTCGCGTTGGTGAACATGTCCTTGAAGTCCGAGACCAGCTCCTCCCTGAAGGCCTCCCCGTCGAAGTTCTTGCAGTCCTCCTTCGCGTCGGCCACCATCTTGTCCTCCTTGTTCTTCACGAACTCCTCGCTCCTGTAGAAGGTCTTGGCCTCCGCCCTCATCTTCTTGGCCCTCATCTCGTCCAAGGACTCCACCTCCGAGAAGATGTCGGTGCACACCTCGATCATCCTCTGCACCTTCTCGTCGGCCTTCTCGTCCACCTCCTCCCCCTCCCACTGGGTGTGCAAGGCCCTGTACCCCTTGTAGGAGCTGAAGCCCTTCATCATGATGTCCGCCACCTGGATCTTGCTGGCGGCGGACTCCAAGGAGGCCATCACGTGGGCCGTCTTGTACTTGGCGGGGCACATGATCTCGTTGGTCCTGGTGGAGTACACCAGCACGTCCACGGTCCCCCTGGTGCAGATGTTCTGGGCCACCATATCCTTCATGACGGGGTCATACTTCTCCCTCTTGGACAGGGCCCTGTCGGAGCAGATGCCGTCCGTGACGGCCACGTCGATCATGTCCAGGCAGTTCCCGGGGGCCGAGAAGAGCATGTCGGGGGTCTTGTTCGCGAACCTGCCGTGGTAGTTGACCTTCTTCAAGTCGACGTCCATCTCCCCCCTCTCGCACCCCACCGAGGTCATCCCCAGGGAGTCCAGGAAGGCCTTCCCCAACAGGTCGTGCCTGGACCTGAGGTAGACGTCCACCACCTCCACGGACATCTCCGGGCCGTCCTTCTTGTTGATGGAGTCCACGTACTCCACCAGCTCGTCCAGGCTCTCCATGGGCTTCACCTCCTTCCTGACCTCCTTCACCTTGTCCCTGGACATGTGCTCCTCCACCTCCTCCAGCAACTCGCACACCCTCTCCACCTTCAACATCTTCTGCATCGCCCTGGTCCTCTCCTCCGTGTTCGTGCCCACCCTGCCGGAGAAGGGGTACAGGATCTTGTCCCCCTTCATGCCGGACATCTTCTGCGCGTAGAACTCCTCCCCCGTGATCACGAACCTGGCCATGTAGGTCCCCCTGATGGTCTTCGGGGTAGAGAACAGCTCCAAGTACTCCTTGTAGGAGATCTCCTCGAACTCGTCCGTCTCCATGTCGGTCAACTCGAAGGCGTTGGGGGAGCGGCTGAAGGACTGCTCGGAGTTGGGCATCTCTATGGCGGTCTTCATGGACTCCGTCAGCATCCCGACCACCTTCTCCTTCGAGTGGTTCTCCAGCATGTTGGTGTAGATGCGGGAGGGGGTCACCTGTATGTCCCCCCCCATGGAGTTGATCACGCCCGAGAGGCCCCTGTGGAAGGTGATCCTATAGGGGACCGACTCCATGACGATGGAGTAGGAGACCTTCATCAGGAAGGGGTGCATCGGCCCCTGGAGGAACTCCTTCATGTTGATGCACGGTGATCTCCACGATCGTGAAGCTGGTGTTCGACCTCTTGTTCCTGGTTGAGATGGTGATATTATGATTTAGTAGAAACGTTGTGTA

**>SalaUV-NL1-1-11_SL**

ACACAATGTCTTCTACTTATTAATACTATCATGCTGACCAACCTGAAGATGAACATCAGCCTGCTCCTTGCACAGGAACTACAAGATGGACAGCAAGCTCCTCCTCTTGTCCTGGACCTTGGTCACCATCCTGTCCAAGGCCACCGCCCTCTTCGTGTTGAAAGCGATGTCCAACTTGATGAGGGAGTTGAGGGTGGGGGTCTTCTCCCCGATGGACGAGAAGATCTCCGTGACCAAGCCCGTCGTGACCATCTGCACCATGACGACGAGGGTGTCCGAGGCGGAGTTGTACCCCTTGTTCTCCTTGAAGTCCCTCAGGGCGTCCATCTGGGCCAGGCCGGACATGCCGGTGGTGGGGTTGATCGTCCCGGTCATCTTCATGTAGACGGCCGTGTACACCATGTTGCACACCACCGCCAGCTTCCTCTTCTTCTCGTCCCCGATGAAGTTCCTCTGGATGTCGTTGACCGTGTTCATCACCAGCCTCTGGTCGAACGAGTAGAACTTCTCGAAGACCCTCTTCACTGAGGTGTAGACCGTGCCCATGTACTCCCTCAAGGACATCCCCTCCTCCAGGGGCTCCTCCTCCATGTGGGCGGTGGTGTCGATCTCCTTCAACCCCGAGAAGGCCCCGGCCAGCCCGGCGAGGGAGGTGAAGTCCATGGACATCCCGCCCATCTCCTCCTTCCCGATGGAGGAGAACATCTCGGCGTTCAGCTTGGTGGCCATGGAGGACGAGGAGATCCCGAAGTCCGTGGCGCTGATGGCGCCCATGACGTTCTCGACGCTGTAGTCCTCCACGGCCTGGGACACGTCCATCATCTCGTATATCGTGGAGTCCACCTCCGCCTCGAACTCGGCCTCCTTGATCCTGTTGTCCGGGGCGAACATGGAGTTGTCCGCTATCCCCAGGTCGTTCAGGATCTCCTCCACGTCCGACAAGTCCAGCTTGCTCGAGACGACGAGCTCGGAGAACTTCCTTATGTTGGGGTTCTCAGAGTCCTCGTTGAACGGCATCTTGTAGCTGTTGGTGCCGATGTGGAAGAACACCTTCCCCCCCAGCTCCCTCTCGTTGAACATGGCCAACACGGTCCACTTGGTCATCCTCCTGGTCAGGATCAGCTCCGTCTTGGGCCAGATCCTGTAGACCTTGATCCACAGGTTCCCGATCTTGGCGTTCTTCTCCACCCCCAAGTTGTCCGTCCTGGGGGCCCAGGGCCTGATCTGGATGGTGGAGAGGTCCTTGTGCATCAGCCTGGTGATGTTCCCCAACTCCACGTCGGAGCTGCTGACCTTGTACATCCTCACCGAGACGGTGTGCGGGGACGCCCCCCTGTCCTCGTGGATGATCCCGAGGTGCACCAAGTCGGTGTAGTACCTGGTCTTCATCTTCCCGTTCACGTCGACCTTGTAGAACACCTTGAGGGACTTGGCCCCCTGGAAGGAGTACTTGTCCTCGTCGGTGGCCGAGACGGCCATCTTCGCGAACCTCACCTCCAAGGAGTCCGACGGGATCACGGTCAGCAAGGGGTTGTCGATGCCCCAGGGATCCCTCACCAAGTTCTCCATCGGGCCCTCCGTGAGGTCCCCCCTGAACTCCTCCACCTCGAAGCGATCCACCCTCCCCCTCATCGACAAGGCGCTGTTGAACCTCTCCTCCGACACCACCCTGAGGGACTTCTTCGGGACGAACATCTGCAAGATCCCGCGGTTCGAGTACTTCTCCAAGTAGAGCATCCTCATGTTGGTGAGCGCCTCGGAGTTGTCCGGGAAGGAGACCATCATCTCCATGTTGAGGTCGGAGGTCATCTTCAGCCAGTACATGATGTAGCTCTTCAGCATCGTCAGGGGCCTCCTCACCCCCGGGAAGGCCTTCTTGATCTGGTATATGGGGTACTCGATGAACTTCTCCAACTTGACCCCCAGCGACTCGGAGATCTTGTTGGCCTCCGTCATGGCCGAGTTGCTAGCGTTGGAGGTGGCCCTCATCATGTACCTGAAGACCTCCATGTCGGACACCTTCACCCTGTCCTTCATCGGGGTGAAGTTGATCTTCCTGGACTTGTTGTGGCTGTAGGAGTGGGAGGGGATGGCGATCTCGGCCTCCTTCTCCGCGATCTCCCCCATCTCGCAGATCTCCTTGAAGGGGGCGGTGAGGTGGAACTTGGACTTGGTGTCGTCGTTCTTCAGGATCTTGTCCACCACCCCCACGAAGTCCGTGGCCACCACCCTGGAGGAGGAGATCAACGCGATGTTCTCCCCCGCCATGGCCAAGGCCCTGATCAAGATGTGGACCCTGATGGAGTCGTTGAACCCGAACTTCCTGTCCATCTGGAACATGTAGGTGTCCATGTTGGAGAAGAAGCTGGCCGAGTCGCTGCGGTCGGAGAAGGACAAGGCCGTGTTCCTCTCGGCCAGCTTGTCGATGTCCTCCCTCGTGATGGACCTCTTGTTCATCAGCCTCTTCTTCATCTCCTTGATCTTCGTGTCGAACCTGAAGGGGAGGGTCATCTTCGTCTTGCCGTTCATGGAGGCCATGTAGTCCCTGTCCTCCTCCATCTCCTCCTCGGTGTAGTCCGGGGAGGCGGCCATCTCGACGTCGGTCATGTAGGCGGCGTAGTACTTGACGTAGAACTCCATCACCTTCTCGGAGTTGCCCTCCTTGTACATCAGCACCTCCGGGCCGGCGATCATCTGCTCCACCACCCTGTCGGTCGACATGTACCCGAGCTGGAAGGGCAGGGAGTTCCTGTCGCACTCCAGCTTCCTCATGATCTCCTCCTTCCTCTCCCCGTCGATGGCGTAGATGGACGAGAGCGCCACGGACTGCAGCTCGAACATCACCTTGATCGTGGGGATCTTGACCCCGTTCTCGAACGCGTTCCGCATCTGGGATATGGAGGACTTCACCGCCTTCTCCGGCTCGGTCATGTCCACGATGTTCTGGGAGTTGTAGACGGCCTTCAGCACGCTGTCCGTGGCCCTCTTCCCGGTGGAGAAGTAGGAGTTGAACTCGGTCAAGATGGACAAGGCGGACTTCTTCCAGTTGGTCGTGATGTTGGCCATCCTGTTGGTCATGTCGTGCACCTGGAGCATGGACACCATGAAGGCCGTCAAGTCGGACATCTTCGTGTACCTGACCAGGAAGATCTTCGTGAGGTCGTCCGAGGAGATGAGGGTGGTGCACGCGGCGTCCCCGTACGACTTGGAGAACAGGAAGCACATCACCGAGTCCATGACGTCGTCCTTCGCGCAGTGGAACAAGGAGGAGAAGTAGTGGAGCATGCCCTGCCCCATCCCGGACTTGAATATGGCGGTCCCCCTCGAGGAGACGAAGTTCTCCTTGAGCCAGTTTATGTTCTTGTCGTCGGTCTTCTCGTTCCACAGCTTGTCCGTCTGCCACCTCCTGGTCAGGGTGGGCGGGAGGAACATGGTCTTCGTCGCGAAGGCCCGGATCACGGTGACGCAGAACTCCTTCAGGTTCACCGGCAAGGGCATCATCAAGCAGAAGTAGATGAAGCTATCCATCACGAACCCGGGGGACCACCTGGAGGCGTCCGTGTTGAAGGACAGGATGATGCCCGAGGGGGAGGTCTCCGACTCGTCCTTGTGCTTGTTCATCTCCAAGATCATCTTCTTGTACACGCTGGTGGCGTTGGACTGGAGGTCCTTCTTCACGTCCGCGTGCGTGAGCATCTCCTTCTCGTGCAAGGAGCAGAGCTCGTAGAACATCTTCTCCAGGAACTTCACGTGCAACCTCAACTTGGCCGCCTGTATCAGGATCTCCCTCGGCCCCCCGATCTGGGACTTCGGGAAGAGCTTGAACACGGCGTTCACCACGTCCGTGCAGGTGACCATGTCCATCAGGAAGATGGACTCCAACTCCTCCACCTGCTCGCACATCGTCTCGAAGGCGGTCGCCGTGGTCAGCTTGTCCGAGTAGTCCACCGACTCCTCGTACAGGGGGCACGACTTCAGGGACGACCTCATGGACATCGTCTCCCTTATGGTGGTGGTCATCGCCCTCATCAGGCCCTTGTTCAAGTCGGGGGAGTGGATTATCTTCATCCTGAGCTGGTAGCCCATGTACATGACGAACTTCTTGTTGAAGGTGTGGGTCTGCATCTTGTTCTTCGCGAGGGCGTCGAAGTCCGCGTCGCCCGTGGAGGCCCCGGGGAACAGCTCCGACATGGACAAGTAGTGCCTCTCCTGCTCGGACATCTTGGTCATGATGGGCACCAGCCTGTGGCCCGACGTCCCCTTCTCCTTGTTGAAGAGGTTGCTCATGTAGATGTCGTTCATCACCTCCGCGAACTCGACCAGCATCCCGGCCACCTTGAACGAGGGCATGTATATCCTGTCGTAGTCCGCGAACTCCGAGGCCATCTTCTCCGCGTACTGCTTTATGATGTCCGGGGCGGCCATGGTCAGCCTCTCGGCGTAGTCCATCTGCATCGCCCTGATGGAGGCCTCGAACATCGTCCTGACGGGGTCCTTCATGATGTTCTTCACCATGCTCACCTTATCCGAGGCGAACGCCGTGAGGCCGTGCATGATGTACCTGTTGTACTGGAGCGAGGTGGACACCCCCCTCTTGTGGCACATCATGACCAGGATGGGGACCATCACCATGGCGGACCTCCTGTACTCCTCCCTGGTCATCTTCTGGGCCCTGGTGCTCTCGAACTCCTCCCTGTCCGCCATCACCTGCGAGTGGATGGCCGCGCTCCTCTCGTAGAGCATGGAGTAGTGCTCCACGTCCGCGACCGAGACGGTGAGCCACTCCGTCTGGTACACCTCGGAGTACTCCTTCGTGTACTGGTTCCTGGCGATGGTCACGTCCATGTAGGACTTGCAGTCCTCCTTGCTCTTCAGCTTCTGGCACTCCATCCCCTCCATGTGCTTGTAGGGGGAGAATATCTTGACCCTGATCTGCTTCTCCTGGGTCAGGGCGGACCCGGCCTTCACGGCCAAGCAGTAGTCCCCGAACTCCTTGATGACCGTGAAGCGGGAGACCTTCTTGCCCGAGTCCCCGTTCCTCTTGGCCCTCCTCCTCCCCTCCAGCCTGGCGATGTTCCTGTAGAGGTCCGTCCAGAAGTGGGTCACCTCCGCCACGGCCAACTCGTTGAACATCTTCATCTGGGGGTCCTTGTACACCTTGGCCATGTTGTTGGTGAACTCGAAGGTGGCGTCTCTGGACATCTTCTCCACCATGTTCTCCATGAACTTCCTGTCCAAGAGGTTGGCCCCGATCCCCGCCATGTTCTCCTTCTTCTTGGTGTTCCCCGCCTTGGTGCAGTCGTTCAAGAAGATGGTGCCGTCGAAGAGGGAGCCCCCCTTCGAGTCCGTCACGTGCATCTTGACGTCCTTCATGTCGTTCAACTTCTGGGGGATCGGCATCGGGAAGGACTTCCCGTAGTCCTTCTTGGCCCTATTCATCTCCTCCGTGGGGGACAGCTCGTCCATCTTGTTCAACTCGGAGACGATCTCCTCCTGCGAGGTGCAGACCGCCTTCCTGATCTCCTCCGTGTACTTCGCGTTGGTGAACATGTCCTTGAAGTCCGAGACCAGCTCCTCCCTGAAGGCCTCCCCGTCGAAGTTCTTGCAGTCCTCCTTCGCGTCGGCCACCATCTTGTCCTCCTTGTTCTTCACGAACTCCTCGCTCCTGTAGAAGGTCTTGGCCTCCGCCCTCATCTTCTTGGCCCTCATCTCGTCCAAGGACTCCACCTCCGAGAAGATGTCGGTGCACACCTCGATCATCCTCTGCACCTTCTCGTCGGCCTTCTCGTCCACCTCCTCCCCCTCCCACTGGGTGTGCAAGGCCCTGTACCCCTTGTAGGAGCTGAAGCCCTTCATCATGATGTCCGCCACCTGGATCTTGCTGGCGGCGGACTCCAAGGAGGCCATCACGTGGGCCGTCTTGTACTTGGCGGGGCACATGATCTCGTTGGTCCTGGTGGAGTACACCAGCACGTCCACGGTCCCCCTGGTGCAGATGTTCTGGGCCACCATATCCTTCATGACGGGGTCATACTTCTCCCTCTTGGACAGGGCCCTGTCGGAGCAGATGCCGTCCGTGACGGCCACGTCGATCATGTCCAGGCAGTTCCCGGGGGCCGAGAAGAGCATGTCGGGGGTCTTGTTCGCGAACCTGCCGTGGTAGCCGACCTTCTTCAAGTCGACGTCCATCTCCCCCCTCTCGCACCCCACCGAGGTCATCCCCAAGGAGTCCAGGAAGGCCTTCCCCAGCAAGTCGTGCCTGGACCTGAGGTAGACGTCCACCACCTCCACGGACATCTCCGGGCCGTCCTTCTTGTTGATGGAGTCCACGTACTCCACCAGCTCGTCCAGGCTCTCCATGGGCTTCACCTCCTTCCTGACCTCCTTCACCTTGTCCCTGGACATGTGCTCCTCCACCTCCTCCAGCAACTCGCACACCCTCTCCACCTTCAACATCTTCTGCATCGCCCTGGTCCTCTCCTCCGTGTTCGTGCCCACCCTGCCGGAGAAGGGGTACAGGATCTTGTCCCCCTTCATGCCGGACATCTTCTGCGCGTAGAACTCCTCCCCCGTGATCACGAACCTGGCCATGTAGGTCCCCCTGATGGTCTTCGGGGTAGAGAACAGCTCCAAGTACTCCTTGTAGGAGATCTCCTCGAACTCGTCCGTCTCCATGTCGGTCAACTCGAAGGCGTTGGGGGAGCGGCTGAAGGACTGCTCGGAGTTGGGCATCTCTATGGCGGTCTTCATGGACTCCGTCAGCATCCCGACCACCTTCTCCTTCGAGTGGTTCTCCAGCATGTTGGTGTAGATGCGGGAGGGGGTCACCTGTATGTCCCCCCCCATGGAGTTGATCACGCCCGAGAGGCCCCTGTGGAAGGTGATCCTATAGGGGACCGACTCCATGACGATGGAGTAGGAGACCTTCATCAGGAAGGGGTGCATCGGCCCCTGGAGGAACTCCTTCATGTTGATGCACGGTGATCTCCACGATCGTGAAGCTGGTGTTCGACCTCTTGTTCCTGGTTGAGATGGTGATATTATGATTTAGTAGAAACTTTGTGTA

**>SalaUV-NL1-1-12_SL**

ACACAATGTCTTCTACTTATTAATACTATCATGCTGACCAACCTGAAGATGAACATCAGCCTGCTCCTTGCACAGGAACTACAAGATGGACAGCAAGCTCCTCCTCTTGTCCTGGACCTTGGTCACCATCCTGTCCAAGGCCACCGCCCTCTTCGTGTTGAAAGCGATGTCCAACTTGATGAGGGAGTTGAGGGTGGGGGTCTTCTCCCCGATGGACGAGAAGATCTCCGTGACCAAGCCCGTCGTGACCATCTGCACCATGACGACGAGGGTGTCCGAGGCGGAGTTGTACCCCTTGTTCTCCTTGAAGTCCCTCAGGGCGTCCATCTGGGCCAGGCCGGACATGCCGGTGGTGGGGTTGATCGTCCCGGTCATCTTCATGTAGACGGCCGTGTACACCATGTTGCACACCACCGCCAGCTTCCTCTTCTTCTCGTCCCCGATGAAGTTCCTCTGGATGTCGTTGACCGTGTTCATCACCAGCCTCTGGTCGAACGAGTAGAACTTCTCGAAGACCCTCTTCACTGAGGTGTAGACCGTGCCCATGTACTCCCTCAAGGACATCCCCTCCTCCAGGGGCTCCTCCTCCATGTGGGCGGTGGTGTCGATCTCCTTCAACCCCGAGAAGGCCCCGGCCAGCCCGGCGAGGGAGGTGAAGTCCATGGACATCCCGCCCATCTCCTCCTTCCCGATGGAGGAGAACATCTCGGCGTTCAGCTTGGTGGCCATGGAGGACGAGGAGATCCCGAAGTCCGTGGCGCTGATGGCGCCCATGACGTTCTCGACGCTGTAGTCCTCCACGGCCTGGGACACGTCCATCATCTCGTATATCGTGGAGTCCACCTCCGCCTCGAACTCGGCCTCCTTGATCCTGTTGTCCGGGGCGAACATGGAGTTGTCCGCTATCCCCAGGTCGTTCAGGATCTCCTCCACGTCCGACAAGTCCAGCTTGCTCGAGACGACGAGCTCGGAGAACTTCCTTATGTTGGGGTTCTCAGAGTCCTCGTTGAACGGCATCTTGTAGCTGTTGGTGCCGATGTGGAAGAACACCTTCCCCCCCAGCTCCCTCTCGTTGAACATGGCCAACACGGTCCACTTGGTCATCCTCCTGGTCAGGATCAGCTCCGTCTTGGGCCAGATCCTGTAGACCTTGATCCACAGGTTCCCGATCTTGGCGTTCTTCTCCACCCCCAAGTTGTCCGTCCTGGGGGCCCAGGGCCTGATCTGGATGGTGGAGAGGTCCTTGTGCATCAGCCTGGTGATGTTCCCCAACTCCACGTCGGAGCTGCTGACCTTGTACATCCTCACCGAGACGGTGTGCGGGGACGCCTGCCTGTCCTCGTGGATGATCCCGAGGTGCACCAAGTCGGTGTAGTACCTGGTCTTCATCTTCCCGTTCACGTCGACCTTGTAGAACACCTTGAGGGACTTGGCCCCCTGGAAGGAGTACTTGTCCTCGTCGGTGGCCGAGACGGCCATCTTCGCGAACCTCACCTCCAAGGAGTCCGACGGGATCACGGTCAGCAAGGGGTTGTCGATGCCCCAGGGATCCCTCACCAAGTTCTCCATCGGGCCCTCCGTGAGGTCCCCCCTGAACTCCTCCACCTCGAAGCGATCCACCCTCCCCCTCATCGACAAGGCGCTGTTGAACCTCTCCTCCGACACCACCCTGAGGGACTTCTTCGGGACGAACATCTGCAAGATCCCGCGGTTCGAGTACTTCTCCAAGTAGAGCATCCTCATGTTGGTGAGCGCCTCGGAGTTGTCCGGGAAGGAGACCATCATCTCCATGTTGAGGTCGGAGGTCATCTTCAGCCAGTACATGATGTAGCTCTTCAGCATCGTCAGGGGCCTCCTCACCCCCGGGAAGGCCTTCTTGATCTGGTATATGGGGTACTCGATGAACTTCTCCAACTTGACCCCCAGCGACTCGGAGATCTTGTTGGCCTCCGTCATGGCCGAGTTGCTAGCGTTGGAGGTGGCCCTCATCATGTACCTGAAGACCTCCATGTCGGACACCTTCACCCTGTCCTTCATCGGGGTGAAGTTGATCTTCCTGGACTTGTTGTGGCTGTAGGAGTGGGAGGGGATGGCGATCTCGGCCTCCTTCTCCGCGATCTCCCCCATCTCGCAGATCTCCTTGAAGGGGGCGGTGAGGTGGAACTTGGACTTGGTGTCGTCGTTCTTCAGGATCTTGTCCACCACCCCCACGAAGTCCGTGGCCACCACCCTGGAGGAGGAGATCAACGCGATGTTCTCCCCCGCCATGGCCAAGGCCCTGATCAAGATGTGGACCCTGATGGAGTCGTTGAACCCGAACTTCCTGTCCATCTGGAACATGTAGGTGTCCATGTTGGAGAAGAAGCTGGCCGAGTCGCTGCGGTCGGAGAAGGACAAGGCCGTGTTCCTCTCGGCCAGCTTGTCGATGTCCTCCCTCGTGATGGACCTCTTGTTCATCAGCCTCTTCTTCATCTCCTTGATCTTCGTGTCGAACCTGAAGGGGAGGGTCATCTTCGTCTTGCCGTTCATGGAGGCCATGTAGTCCCTGTCCTCCTCCATCTCCTCCTCGGTGTAGTCCGGGGAGGCGGCCATCTCGACGTCGGTCATGTAGGCGGCGTAGTACTTGACGTAGAACTCCATCACCTTCTCGGAGTTGCCCTCCTTGTACATCAGCACCTCCGGGCCGGCGATCATCTGCTCCACCACCCTGTCGGTCGACATGTACCCGAGCTGGAAGGGCAGGGAGTTCCTGTCGCACTCCAGCTTCCTCATGATCTCCTCCTTCCTCTCCCCGTCGATGGCGTAGATGGACGAGAGCGCCACGGACTGCAGCTCGAACATCACCTTGATCGTGGGGATCTTGACCCCGTTCTCGAACGCGTTCCGCATCTGGGATATGGAGGACTTCACCGCCTTCTCCGGCTCGGTCATGTCCACGATGTTCTGGGAGTTGTAGACGGCCTTCAGCACGCTGTCCGTGGCCCTCTTCCCGGTGGAGAAGTAGGAGTTGAACTCGGTCAAGATGGACAAGGCGGACTTCTTCCAGTTGGTCGTGATGTTGGCCATCCTGTTGGTCATGTCGTGCACCTGGAGCATGGACACCATGAAGGCCGTCAAGTCGGACATCTTCGTGTACCTGACCAGGAAGATCTTCGTGAGGTCGTCCGAGGAGATGAGGGTGGTGCACGCGGCGTCCCCGTACGACTTGGAGAACAGGAAGCACATCACCGAGTCCATGACGTCGTCCTTCGCGCAGTGGAACAAGGAGGAGAAGTAGTGGAGCATGCCCTGCCCCATCCCGGACTTGAATATGGCGGTCCCCCTCGAGGAGACGAAGTTCTCCTTGAGCCAGTTTATGTTCTTGTCGTCGGTCTTCTCGTTCCACAGCTTGTCCGTCTGCCACCTCCTGGTCAGGGTGGGCGGGAGGAACATGGTCTTCGTCGCGAAGGCCCGGATCACGGTGACGCAGAACTCCTTCAACTTCACCGGCAAGGGCATCATCAAGCAGAAGTAGATGAAGCTATCCATCACGAACCCGGGGGACCACCTGGAGGCGTCCGTGTTGAAGGACAGGATGATGCCCGAGGGGGAGGTCTCCGACTCGTCCTTGTGCTTGTTCATCTCCAAGATCATCTTCTTGTACACGCTGGTGGCGTTGGACTGGAGGTCCTTCTTCACGTCCGCGTGCGTGAGCATCTCCTTCTCGTGCAAGGAGCAGAGCTCGTAGAACATCTTCTCCAGGAACTTCACGTGCAACCTCAACTTGGCCGCCTGTATCAGGATCTCCCTCGGCCCCCCGATCTGGGACTTCGGGAAGAGCTTGAACACGGCGTTCACCACGTCCGTGCAGGTGACCATGTCCATCAGGAAGATGGACTCCAACTCCTCCACCTGCTCGCACATCGTCTCGAAGGCGGTCGCCGTGGTCAGCTTGTCCGAGTAGTCCACCGACTCCTCGTACAGGGGGCACGACTTCAGGGACGACCTCATGGACATCGTCTCCCTTATGGTGGTGGTCATCGCCCTCATCAGGCCCTTGTTCAAGTCGGGGGAGTGGATTATCTTCATCCTGAGCTGGTAGCCCATGTACATGACGAACTTCTTGTTGAAGGTGTGGGTCTGCATCTTGTTCTTCGCGAGGGCGTCGAAGTCCGCGTCGCCCGTGGAGGCCCCGGGGAACAGCTCCGACATGGACAAGTAGTGCCTCTCCTGCTCGGACATCTTGGTCATGATGGGCACCAGCCTGTGGCCCGACGTCCCCTTCTCCTTGTTGAAGAGGTTGCTCATGTAGATGTCGTTCATCACCTCCGCGAACTCGACCAGCATCCCGGCCACCTTGAACGAGGGCATGTATATCCTGTCGTAGTCCGCGAACTCCGAGGCCATCTTCTCCGCGTACTGCTTTATGATGTCCGGGGCGGCCATGGTCAGCCTCTCGGCGTAGTCCATCTGCATCGCCCTGATGGAGGCCTCGAACATCGTCCTGACGGGGTCCTTCATGATGTTCTTCACCATGCTCACCTTATCCGAGGCGAACGCCGTGAGGCCGTGCATGATGTACCTGTTGTACTGGAGCGAGGTGGACACCCCCCTCTTGTGGCACATCATGACCAGGATGGGGACCATCACCATGGCGGACCTCCTGTACTCCTCCCTGGTCATCTTCTGGGCCCTGGTGCTCTCGAACTCCTCCCTGTCCGCCATCACCTGCGAGTGGATGGCCGCGCTCCTCTCGTAGAGCATGGAGTAGTGCTCCACGTCCGCGACCGAGACGGTGAGCCACTCCGTCTGGTACACCTCGGAGTACTCCTTCGTGTACTGGTTCCTGGCGATGGTCACGTCCATGTAGGACTTGCAGTCCTCCTTGCTCTTCAGCTTCTGGCACTCCATCCCCTCCATGTGCTTGTAGGGGGAGAATATCTTGACCCTGATCTGCTTCTCCTGGGTCAGGGCGGACCCGGCCTTCACGGCCAAGCAGTAGTCCCCGAACTCCTTGATGACCGTGAAGCGGGAGACCTTCTTGCCCGAGTCCCCGTTCCTCTTGGCCCTCCTCCTCCCCTCCAGCCTGGCGATGTTCCTGTAGAGGTCCGTCCAGAAGTGGGTCACCTCCGCCACGGCCAACTCGTTGAACATCTTCATCTGGGGGTCCTTGTACACCTTGGCCATGTTGTTGGTGAACTCGAAGGTGGCGTCTCTGGACATCTTCTCCACCATGTTCTCCATGAACTTCCTGTCCAAGAGGTTGGCCCCGATCCCCGCCATGTTCTCCTTCTTCTTGGTGTTCCCCGCCTTGGTGCAGTCGTTCAAGAAGATGGTGCCGTCGAAGAGGGAGCCCCCCTTCGAGTCCGTCACGTGCATCTTGACGTCCTTCATGTCGTTCAACTTCTGGGGGATCGGCATCGGGAAGGACTTCCCGTAGTCCTTCTTGGCCCTATTCATCTCCTCCGTGGGGGACAGCTCGTCCATCTTGTTCAACTCGGAGACGATCTCCTCCTGCGAGGTGCAGACCGCCTTCCTGATCTCCTCCGTGTACTTCGCGTTGGTGAACATGTCCTTGAAGTCCGAGACCAGCTCCTCCCTGAAGGCCTCCCCGTCGAAGTTCTTGCAGTCCTCCTTCGCGTCGGCCACCATCTTGTCCTCCTTGTTCTTCACGAACTCCTCGCTCCTGTAGAAGGTCTTGGCCTCCGCCCTCATCTTCTTGGCCCTCATCTCGTCCAAGGACTCCACCTCCGAGAAGATGTCGGTGCACACCTCGATCATCCTCTGCACCTTCTCGTCGGCCTTCTCGTCCACCTCCTCCCCCTCCCACTGGGTGTGCAAGGCCCTGTACCCCTTGTAGGAGCTGAAGCCCTTCATCATGATGTCCGCCACCTGGATCTTGCTGGCGGCGGACTCCAGGGAGGCCATCACGTGGGCCGTCTTGTACTTGGCGGGGCACATGATCTCGTTGGTCCTGGTGGAGTACACCAGCACGTCCACGGTCCCCCTGGTGCAGATGTTCTGGGCCACCATATCCTTCATGACGGGGTCATACTTCTCCCTCTTGGACAGGGCCCTGTCGGAGCAGATGCCGTCCGTGACGGCCACGTCGATCATGTCCAGGCAGTTCCCGGGGGCCGAGAAGAGCATGTCGGGGGTCTTGTTCGCGAACCTGCCGTGGTAGTTGACCTTCTTCAAGTCGACGTCCATCTCCCCCCTCTCGCACCCCACCGAGGTCATCCCCAGGGAGTCCAGGAAGGCCTTCCCCAGCAAGTCGTGCCTGGACCTGAGGTAGACGTCCACCACCTCCACGGACATCTCCGGGCCGTCCTTCTTGTTGATGGAGTCCACGTACTCCACCAGCTCGTCCAGGCTCTCCATGGGCTTCACCTCCTTCCTGACCTCCTTCACCTTGTCCCTGGACATGTGCTCCTCCACCTCCTCCAGCAACTCGCACACCCTCTCCACCTTCAACATCTTCTGCATCGCCCTGGTCCTCTCCTCCGTGTTCGTGCCCACCCTGCCGGAGAAGGGGTACAGGATCTTGTCCCCCTTCATGCCGGACATCTTCTGCGCGTAGAACTCCTCCCCCGTGATCACGAACCTGGCCATGTAGGTCCCCCTGATGGTCTTCGGGGTAGAGAACAGCTCCAAGTACTCCTTGTAGGAGATCTCCTCGAACTCGTCCGTCTCCATGTCGGTCAACTCGAAGGCGTTGGGGGAGCGGCTGAAGGACTGCTCGGAGTTGGGCATCTCTATGGCGGTCTTCATGGACTCCGTCAGCATCCCGACCACCTTCTCCTTCGAGTGGTTCTCCAGCATGTTGGTGTAGATGCGGGAGGGGGTCACCTGTATGTCCCCCCCCATGGAGTTGATCACGCCCGAGAGGCCCCTGTGGAAGGTGATCCTATAGGGGACCGACTCCATGACGATGGAGTAGGAGACCTTCATCAGGAAGGGGTGCATCGGCCCCTGGAGGAACTCCTTCATGTTGATGCACGGTGATCTCCACGATCGTGAAGCTGGTGTTCGACCTCTTGTTCCTGGTTGAGATGATGATAGTATGATTTAGTAGAAACGTTGTGT

**>SalaUV-NL1-1-14_SL**

ACACAATGTCTTCTACTTATTAATACTATCATGCTGACCAACCTGAAGATGAACATCAGCCTGCTCCTTGCACAGGAACTACAAGATGGACAGCAAGCTCCTCCTCTTGTCCTGGACCTTGGTCACCATCCTGTCCAAGGCCACCGCCCTCTTCGTGTTGAAAGCGATGTCCAACTTGATGAGGGAGTTGAGGGTGGGGGTCTTCTCCCCGATGGACGAGAAGATCTCCGTGACCAAGCCCGTCGTGACCATCTGCACCATGACGACGAGGGTGTCCGAGGCGGAGTTGTACCCCTTGTTCTCCTTGAAGTCCCTCAGGGCGTCCATCTGGGCCAGGCCGGACATGCCGGTGGTGGGGTTGATCGTCCCGGTCATCTTCATGTAGACGGCCGTGTACACCATGTTGCACACCACCGCCAGCTTCCTCTTCTTCTCGTCCCCGATGAAGTTCCTCTGGATGTCGTTGACCGTGTTCATCACCAGCCTCTGGTCGAACGAGTAGAACTTCTCGAAGACCCTCTTCACTGAGGTGTAGACCGTGCCCATGTACTCCCTCAAGGACATCCCCTCCTCCAGGGGCTCCTCCTCCATGTGGGCGGTGGTGTCGATCTCCTTCAACCCCGAGAAGGCCCCGGCCAGCCCGGCGAGGGAGGTGAAGTCCATGGACATCCCGCCCATCTCCTCCTTCCCGATGGAGGAGAACATCTCGGCGTTCAGCTTGGTGGCCATGGAGGACGAGGAGATCCCGAAGTCCGTGGCGCTGATGGCGCCCATGACGTTCTCGACGCTGTAGTCCTCCACGGCCTGGGACACGTCCATCATCTCGTATATCGTGGAGTCCACCTCCGCCTCGAACTCGGCCTCCTTGATCCTGTTGTCCGGGGCGAACATGGAGTTGTCCGCTATCCCCAGGTCGTTCAGGATCTCCTCCACGTCCGACAAGTCCAGCTTGCTCGAGACGACGAGCTCGGAGAACTTCCTTATGTTGGGGTTCTCAGAGTCCTCGTTGAACGGCATCTTGTAGCTGTTGGTGCCGATGTGGAAGAACACCTTCCCCCCCAGCTCCCTCTCGTTGAACATGGCCAACACGGTCCACTTGGTCATCCTCCTGGTCAGGATCAGCTCCGTCTTGGGCCAGATCCTGTAGACCTTGATCCACAGGTTCCCGATCTTGGCGTTCTTCTCCACCCCCAAGTTGTCCGTCCTGGGGGCCCAGGGCCTGATCTGGATGGTGGAGAGGTCCTTGTGCATCAGCCTGGTGATGTTCCCCAACTCCACGTCGGAGCTGCTGACCTTGTACATCCTCACCGAGACGGTGTGCGGGGACGCCTGCCTGTCCTCGTGGATGATCCCGAGGTGCACCAAGTCGGTGTAGTACCTGGTCTTCATCTTCCCGTTCACGTCGACCTTGTAGAACACCTTGAGGGACTTGGCCCCCTGGAAGGAGTACTTGTCCTCGTCGGTGGCCGAGACGGCCATCTTCGCGAACCTCACCTCCAAGGAGTCCGACGGGATCACGGTCAGCAAGGGGTTGTCGATGCCCCAGGGATCCCTCACCAAGTTCTCCATCGGGCCCTCCGTGAGGTCCCCCCTGAACTCCTCCACCTCGAAGCGATCCACCCTCCCCCTCATCGACAAGGCGCTGTTGAACCTCTCCTCCGACACCACCCTGAGGGACTTCTTCGGGACGAACATCTGCAAGATCCCGCGGTTCGAGTACTTCTCCAAGTAGAGCATCCTCATGTTGGTGAGCGCCTCGGAGTTGTCCGGGAAGGAGACCATCATCTCCATGTTGAGGTCGGAGGTCATCTTCAGCCAGTACATGATGTAGCTCTTCAGCATCGTCAGGGGCCTCCTCACCCCCGGGAAGGCCTTCTTGATCTGGTATATGGGGTACTCGATGAACTTCTCCAACTTGACCCCCAGCGACTCGGAGATCTTGTTGGCCTCCGTCATGGCCGAGTTGCTAGCGTTGGAGGTGGCCCTCATCATGTACCTGAAGACCTCCATGTCGGACACCTTCACCCTGTCCTTCATCGGGGTGAAGTTGATCTTCCTGGACTTGTTGTGGCTGTAGGAGTGGGAGGGGATGGCGATCTCGGCCTCCTTCTCCGCGATCTCCCCCATCTCGCAGATCTCCTTGAAGGGGGCGGTGAGGTGGAACTTGGACTTGGTGTCGTCGTTCTTCAGGATCTTGTCCACCACCCCCACGAAGTCCGTGGCCACCACCCTGGAGGAGGAGATCAACGCGATGTTCTCCCCCGCCATGGCCAAGGCCCTGATCAAGATGTGGACCCTGATGGAGTCGTTGAACCCGAACTTCCTGTCCATCTGGAACATGTAGGTGTCCATGTTGGAGAAGAAGCTGGCCGAGTCGCTGCGGTCGGAGAAGGACAAGGCCGTGTTCCTCTCGGCCAGCTTGTCGATGTCCTCCCTCGTGATGGACCTCTTGTTCATCAGCCTCTTCTTCATCTCCTTGATCTTCGTGTCGAACCTGAAGGGGAGGGTCATCTTCGTCTTGCCGTTCATGGAGGCCATGTAGTCCCTGTCCTCCTCCATCTCCTCCTCGGTGTAGTCCGGGGAGGCGGCCATCTCGACGTCGGTCATGTAGGCGGCGTAGTACTTGACGTAGAACTCCATCACCTTCTCGGAGTTGCCCTCCTTGTACATCAGCACCTCCGGGCCGGCGATCATCTGCTCCACCACCCTGTCGGTCGACATGTACCCGAGCTGGAAGGGCAGGGAGTTCCTGTCGCACTCCAGCTTCCTCATGATCTCCTCCTTCCTCTCCCCGTCGATGGCGTAGATGGACGAGAGCGCCACGGACTGCAGCTCGAACATCACCTTGATCGTGGGGATCTTGACCCCGTTCTCGAACGCGTTCCGCATCTGGGATATGGAGGACTTCACCGCCTTCTCCGGCTCGGTCATGTCCACGATGTTCTGGGAGTTGTAGACGGCCTTCAGCACGCTGTCCGTGGCCCTCTTCCCGGTGGAGAAGTAGGAGTTGAACTCGGTCAAGATGGACAAGGCGGACTTCTTCCAGTTGGTCGTGATGTTGGCCATCCTGTTGGTCATGTCGTGCACCTGGAGCATGGACACCATGAAGGCCGTCAAGTCGGACATCTTCGTGTACCTGACCAGGAAGATCTTCGTGAGGTCGTCCGAGGAGATGAGGGTGGTGCACGCGGCGTCCCCGTACGACTTGGAGAACAGGAAGCACATCACCGAGTCCATGACGTCGTCCTTCGCGCAGTGGAACAAGGAGGAGAAGTAGTGGAGCATGCCCTGCCCCATCCCGGACTTGAATATGGCGGTCCCCCTCGAGGAGACGAAGTTCTCCTTGAGCCAGTTTATGTTCTTGTCGTCGGTCTTCTCGTTCCACAGCTTGTCCGTCTGCCACCTCCTGGTCAGGGTGGGCGGGAGGAACATGGTCTTCGTCGCGAAGGCCCGGATCACGGTGACGCAGAACTCCTTCAGGTTCACCGGCAAGGGCATCATCAAGCAGAAGTAGATGAAGCTATCCATCACGAACCCGGGGGACCACCTGGAGGCGTCCGTGTTGAAGGACAGGATGATGCCCGAGGGGGAGGTCTCCGACTCGTCCTTGTGCTTGTTCATCTCCAAGATCATCTTCTTGTACACGCTGGTGGCGTTGGACTGGAGGTCCTTCTTCACGTCCGCGTGCGTGAGCATCTCCTTCTCGTGCAAGGAGCAGAGCTCGTAGAACATCTTCTCCAGGAACTTCACGTGCAACCTCAACTTGGCCGCCTGTATCAGGATCTCCCTCGGCCCCCCGATCTGGGACTTCGGGAAGAGCTTGAACACGGCGTTCACCACGTCCGTGCAGGTGACCATGTCCATCAGGAAGATGGACTCCAACTCCTCCACCTGCTCGCACATCGTCTCGAAGGCGGTCGCCGTGGTCAGCTTGTCCGAGTAGTCCACCGACTCCTCGTACAGGGGGCACGACTTCAGGGACGACCTCATGGACATCGTCTCCCTTATGGTGGTGGTCATCGCCCTCATCAGGCCCTTGTTCAAGTCGGGGGAGTGGATTATCTTCATCCTGAGCTGGTAGCCCATGTACATGACGAACTTCTTGTTGAAGGTGTGGGTCTGCATCTTGTTCTTCGCGAGGGCGTCGAAGTCCGCGTCGCCCGTGGAGGCCCCGGGGAACAGCTCCGACATGGACAAGTAGTGCCTCTCCTGCTCGGACATCTTGGTCATGATGGGCACCAGCCTGTGGCCCGACGTCCCCTTCTCCTTGTTGAAGAGGTTGCTCATGTAGATGTCGTTCATCACCTCCGCGAACTCGACCAGCATCCCGGCCACCTTGAACGAGGGCATGTATATCCTGTCGTAGTCCGCGAACTCCGAGGCCATCTTCTCCGCGTACTGCTTTATGATGTCCGGGGCGGCCATGGTCAGCCTCTCGGCGTAGTCCATCTGCATCGCCCTGATGGAGGCCTCGAACATCGTCCTGACGGGGTCCTTCATGATGTTCTTCACCATGCTCACCTTATCCGAGGCGAACGCCGTGAGGCCGTGCATGATGTACCTGTTGTACTGGAGCGAGGTGGACACCCCCCTCTTGTGGCACATCATGACCAGGATGGGGACCATCACCATGGCGGACCTCCTGTACTCCTCCCTGGTCATCTTCTGGGCCCTGGTGCTCTCGAACTCCTCCCTGTCCGCCATCACCTGCGAGTGGATGGCCGCGCTCCTCTCGTAGAGCATGGAGTAGTGCTCCACGTCCGCGACCGAGACGGTGAGCCACTCCGTCTGGTACACCTCGGAGTACTCCTTCGTGTACTGGTTCCTGGCGATGGTCACGTCCATGTAGGACTTGCAGTCCTCCTTGCTCTTCAGCTTCTGGCACTCCATCCCCTCCATGTGCTTGTAGGGGGAGAATATCTTGACCCTGATCTGCTTCTCCTGGGTCAGGGCGGACCCGGCCTTCACGGCCAAGCAGTAGTCCCCGAACTCCTTGATGACCGTGAAGCGGGAGACCTTCTTGCCCGAGTCCCCGTTCCTCTTGGCCCTCCTCCTCCCCTCCAGCCTGGCGATGTTCCTGTAGAGGTCCGTCCAGAAGTGGGTCACCTCCGCCACGGCCAACTCGTTGAACATCTTCATCTGGGGGTCCTTGTACACCTTGGCCATGTTGTTGGTGAACTCGAAGGTGGCGTCTCTGGACATCTTCTCCACCATGTTCTCCATGAACTTCCTGTCCAAGAGGTTGGCCCCGATCCCCGCCATGTTCTCCTTCTTCTTGGTGTTCCCCGCCTTGGTGCAGTCGTTCAAGAAGATGGTGCCGTCGAAGAGGGAGCCCCCCTTCGAGTCCGTCACGTGCATCTTGACGTCCTTCATGTCGTTCAACTTCTGGGGGATCGGCATCGGGAAGGACTTCCCGTAGTCCTTCTTGGCCCTATTCATCTCCTCCGTGGGGGACAGCTCGTCCATCTTGTTCAACTCGGAGACGATCTCCTCCTGCGAGGTGCAGACCGCCTTCCTGATCTCCTCCGTGTACTTCGCGTTGGTGAACATGTCCTTGAAGTCCGAGACCAGCTCCTCCCTGAAGGCCTCCCCGTCGAAGTTCTTGCAGTCCTCCTTCGCGTCGGCCACCATCTTGTCCTCCTTGTTCTTCACGAACTCCTCGCTCCTGTAGAAGGTCTTGGCCTCCGCCCTCATCTTCTTGGCCCTCATCTCGTCCAAGGACTCCACCTCCGAGAAGATGTCGGTGCACACCTCGATCATCCTCTGCACCTTCTCGTCGGCCTTCTCGTCCACCTCCTCCCCCTCCCACTGGGTGTGCAAGGCCCTGTACCCCTTGTAGGAGCTGAAGCCCTTCATCATGATGTCCGCCACCTGGATCTTGCTGGCGGCGGACTCCAAGGAGGCCATCACGTGGGCCGTCTTGTACTTGGCGGGGCACATGATCTCGTTGGTCCTGGTGGAGTACACCAGCACGTCCACGGTCCCCCTGGTGCAGATGTTCTGGGCCACCATATCCTTCATGACGGGGTCATACTTCTCCCTCTTGGACAGGGCCCTGTCGGAGCAGATGCCGTCCGTGACGGCCACGTCGATCATGTCCAGGCAGTTCCCGGGGGCCGAGAAGAGCATGTCGGGGGTCTTGTTCGCGAACCTGCCGTGGTAGTTGACCTTCTTCAAGTCGACGTCCATCTCCCCCCTCTCGCACCCCACCGAGGTCATCCCCAAGGAGTCCAGGAAGGCCTTCCCCAGCAAGTCGTGCCTGGACCTGAGGTAGACGTCCACCACCTCCACGGACATCTCCGGGCCGTCCTTCTTGTTGATGGAGTCCACGTACTCCACCAGCTCGTCCAGGCTCTCCATGGGCTTCACCTCCTTCCTGACCTCCTTCACCTTGTCCCTGGACATGTGCTCCTCCACCTCCTCCAGCAACTCGCACACCCTCTCCACCTTCAACATCTTCTGCATCGCCCTGGTCCTCTCCTCCGTGTTCGTGCCCACCCTGCCGGAGAAGGGGTACAGGATCTTGTCCCCCTTCATGCCGGACATCTTCTGCGCGTAGAACTCCTCCCCCGTGATCACGAACCTGGCCATGTAGGTCCCCCTGATGGTCTTCGGGGTAGAGAACAGCTCCAAGTACTCCTTGTAGGAGATCTCCTCGAACTCGTCCGTCTCCATGTCGGTCAACTCGAAGGCGTTGGGGGAGCGGCTGAAGGACTGCTCGGAGTTGGGCATCTCTATGGCGGTCTTCATGGACTCCGTCAGCATCCCGACCACCTTCTCCTTCGAGTGGTTCTCCAGCATGTTGGTGTAGATGCGGGAGGGGGTCACCTGTATGTCCCCCCCCATGGAGTTGATCACGCCCGAGAGGCCCCTGTGGAAGGTGATCCTATAGGGGACCGACTCCATGACGACAGAGTAGGAGACCTTCATCAGGAAGGGGTGCATCGGCCCCTGGAGGAACTCCTTCATGTTGATGCACGGTGATCTCCACGATCGTGAAGCTGGTGTTCGACCTCTTGTTCCTGGTTGAGATGGTGATATTATGATTTAGTAGAAACGTTGTGT

**>SalaUV-NL1-1-15_SL**

ACACAATGTCTTCTACTTATTAATACTATCATGCTGACCAACCTGAAGATGAACATCAGCCTGCTCCTTGCACAGGAACTACAAGATGGACAGCAAGCTCCTCCTCTTGTCCTGGACCTTGGTCACCATCCTGTCCAAGGCCACCGCCCTCTTCGTGTTGAAAGCGATGTCCAACTTGATGAGGGAGTTGAGGGTGGGGGTCTTCTCCCCGATGGACGAGAAGATCTCCGTGACCAAGCCCGTCGTGACCATCTGCACCATGACGACGAGGGTGTCCGAGGCGGAGTTGTACCCCTTGTTCTCCTTGAAGTCCCTCAAGGCGTCCATCTGGGCCAGGCCGGACATGCCGGTGGTGGGGTTGATCGTCCCGGTCATCTTCATGTAGACGGCCGTGTACACCATGTTGCACACCACCGCCAGCTTCCTCTTCTTCTCGTCCCCGATGAAGTTCCTCTGGATGTCGTTGACCGTGTTCATCACCAGCCTCTGGTCGAACGAGTAGAACTTCTCGAAGACCCTCTTCACTGAGGTGTAGACCGTGCCCATGTACTCCCTCAAGGACATCCCCTCCTCCAGGGGCTCCTCCTCCATGTGGGCGGTGGTGTCGATCTCCTTCAACCCCGAGAAGGCCCCGGCCAGCCCGGCGAGGGAGGTGAAGTCCATGGACATCCCGCCCATCTCCTCCTTCCCGATGGAGGAGAACATCTCGGCGTTCAGCTTGGTGGCCATGGAGGACGAGGAGATCCCGAAGTCCGTGGCGCTGATGGCGCCCATGACGTTCTCGACGCTGTAGTCCTCCACGGCCTGGGACACGTCCATCATCTCGTATATCGTGGAGTCCACCTCCGCCTCGAACTCGGCCTCCTTGATCCTGTTGTCCGGGGCGAACATGGAGTTGTCCGCTATCCCCAGGTCGTTCAGGATCTCCTCCACGTCCGACAAGTCCAGCTTGCTCGAGACGACGAGCTCGGAGAACTTCCTTATGTTGGGGTTCTCAGAGTCCTCGTTGAACGGCATCTTGTAGCTGTTGGTGCCGATGTGGAAGAACACCTTCCCCCCCAGCTCCCTCTCGTTGAACATGGCCAACACGGTCCATTTGGTCATCCTCCTGGTCAGGATCAGCTCCGTCTTGGGCCAGATCCTGTAGACCTTGATCCACAGGTTCCCGATCTTGGCGTTCTTCTCCACCCCCAAGTTGTCCGTCCTGGGGGCCCAGGGCCTGATCTGGATGGTGGAGAGGTCCTTGTGCATCAGCCTGGTGATGTTCCCCAACTCCACGTCGGAGCTGCTGACCTTGTACATCCTCACCGAGACGGTGTGCGGGGACGCCTGCCTGTCCTCGTGGATGATCCCGAGGTGCACCAAGTCGGTGTAGTACCTGGTCTTCATCTTCCCGTTCACGTCGACCTTGTAGAACACCTTGAGGGACTTGGCCCCCTGGAAGGAGTACTTGTCCTCGTCGGTGGCCGAGACGGCCATCTTCGCGAACCTCACCTCCAAGGAGTCCGACGGGATCACGGTCAGCAAGGGGTTGTCGATGCCCCAGGGATCCCTCACCAAGTTCTCCATCGGGCCCTCCGTGAGGTCCCCCCTGAACTCCTCCACCTCGAAGCGATCCACCCTCCCCCTCATCGACAAGGCGCTGTTGAACCTCTCCTCCGACACCACCCTGAGGGACTTCTTCGGGACGAACATCTGCAAGATCCCGCGGTTCGAGTACTTCTCCAAGTAGAGCATCCTCATGTTGGTGAGCGCCTCGGAGTTGTCCGGGAAGGAGACCATCATCTCCATGTTGAGGTCGGAGGTCATCTTCAGCCAGTACATGATGTAGCTCTTCAGCATCGTCAAGGGTCTCCTCACCCCCGGGAAGGCCTTCTTGATCTGGTATATGGGGTACTCGATGAACTTCTCCAACTTGACCCCCAGCGACTCGGAGATCTTGTTGGCCTCCGTCATGGCCGAGTTGCTAGCGTTGGAGGTGGCCCTCATCATGTACCTGAAGACCTCCATGTCGGACACCTTCACCCTGTCCTTCATCGGGGTGAAGTTGATCTTCCTGGACTTGTTGTGGCTGTAGGAGTGGGAGGGGATGGCGATCTCGGCCTCCTTCTCCGCGATCTCCCCCATCTCGCAGATCTCCTTGAAGGGGGCGGTGAGGTGGAACTTGGACTTGGTGTCGTCGTTCTTCAGGATCTTGTCCACCACCCCCACGAAGTCCGTGGCCACCACCCTGGAGGAGGAGATCAACGCGATGTTCTCCCCCGCCATGGCCAAGGCCCTGATCAAGATGTGGACCCTGATGGAGTCGTTGAACCCGAACTTCCTGTCCATCTGGAACATGTAGGTGTCCATGTTGGAGAAGAAGCTGGCCGAGTCGCTGCGGTCGGAGAAGGACAAGGCCGTGTTCCTCTCGGCCAGCTTGTCGATGTCCTCCCTCGTGATGGACCTCTTGTTCATCAGCCTCTTCTTCATCTCCTTGATCTTCGTGTCGAACCTGAAGGGGAGGGTCATCTTCGTCTTGCCGTTCATGGAGGCCATGTAGTCCCTGTCCTCCTCCATCTCCTCCTCGGTGTAGTCCGGGGAGGCGGCCATCTCGACGTCGGTCATGTAGGCGGCGTAGTACTTGACGTAGAACTCCATCACCTTCTCGGAGTTGCCCTCCTTGTACATCAGCACCTCCGGGCCGGCGATCATCTGCTCCACCACCCTGTCGGTCGACATGTACCCGAGCTGGAAGGGCAGGGAGTTCCTGTCGCACTCCAGCTTCCTCATGATCTCCTCCTTCCTCTCCCCGTCGATGGCGTAGATGGACGAGAGCGCCACGGACTGCAGCTCGAACATCACCTTGATCGTGGGGATCTTGACCCCGTTCTCGAACGCGTTCCGCATCTGGGATATGGAGGACTTCACCGCCTTCTCCGGCTCGGTCATGTCCACGATGTTCTGGGAGTTGTAGACGGCCTTCAGCACGCTGTCCGTGGCCCTCTTCCCGGTGGAGAAGTAGGAGTTGAACTCGGTCAAGATGGACAAGGCGGACTTCTTCCAGTTGGTCGTGATGTTGGCCATCCTGTTGGTCATGTCGTGCACCTGGAGCATGGACACCATGAAGGCCGTCAAGTCGGACATCTTCGTGTACCTGACCAGGAAGATCTTCGTGAGGTCGTCCGAGGAGATGAGGGTGGTGCACGCGGCGTCCCCGTACGACTTGGAGAACAGGAAGCACATCACCGAGTCCATGACGTCGTCCTTCGCGCAGTGGAACAAGGAGGAGAAGTAGTGGAGCATGCCCTGCCCCATCCCGGACTTGAATATGGCGGTCCCCCTCGAGGAGACGAAGTTCTCCTTGAGCCAGTTTATGTTCTTGTCGTCGGTCTTCTCGTTCCACAGCTTGTCCGTCTGCCACCTCCTGGTCAGGGTGGGCGGGAGGAACATGGTCTTCGTCGCGAAGGCCCGGATCACGGTGACGCAGAACTCCTTCAGGTTCACCGGCAAGGGCATCATCAAGCAGAAGTAGATGAAGCTATCCATCACGAACCCGGGGGACCACCTGGAGGCGTCCGTGTTGAAGGACAGGATGATGCCCGAGGGGGAGGTCTCCGACTCGTCCTTGTGCTTGTTCATCTCCAAGATCATCTTCTTGTACACGCTGGTGGTGTTGGACTGGAGGTCCTTCTTCACGTCCGCGTGCGTGAGCATCTCCTTCTCGTGCAAGGAGCAGAGCTCGTAGAACATCTTCTCCAGGAACTTCACGTGCAACCTCAACTTGGCCGCCTGTATCAGGATCTCCCTCGGCCCCCCGATCTGGGACTTCGGGAAGAGCTTGAACACGGCGTTCACCACGTCCGTGCAGGTGACCATGTCCATCAGGAAGATGGACTCCAACTCCTCCACCTGCTCGCACATCGTCTCGAAGGCGGTCGCCGTGGTCAGCTTGTCCGAGTAGTCCACCGACTCCTCGTACAGGGGGCACGACTTCAGGGACGACCTCATGGACATCGTCTCCCTTATGGTGGTGGTCATCGCCCTCATCAGGCCCTTGTTCAAGTCGGGGGAGTGGATTATCTTCATCCTGAGCTGGTAGCCCATGTACATGACGAACTTCTTGTTGAAGGTGTGGGTCTGCATCTTGTTCTTCGCGAGGGCGTCGAAGTCCGCGTCGCCCGTGGAGGCCCCGGGGAACAGCTCCGACATGGACAAGTAGTGCCTCTCCTGCTCGGACATCTTGGTCATGATGGGCACCAGCCTGTGGCCCGACGTCCCCTTCTCCTTGTTGAAGAGGTTGCTCATGTAGATGTCGTTCATCACCTCCGCGAACTCGACCAGCATCCCGGCCACCTTGAACGAGGGCATGTATATCCTGTCGTAGTCCGCGAACTCCGAGGCCATCTTCTCCGCGTACTGCTTTATGATGTCCGGGGCGGCCATGGTCAGCCTCTCGGCGTAGTCCATCTGCATCGCCCTGATGGAGGCCTCGAACATCGTCCTGACGGGGTCCTTCATGATGTTCTTCACCATGCTCACCTTATCCGAGGCGAACGCCGTGAGGCCGTGCATGATGTACCTGTTGTACTGGAGCGAGGTGGACACCCCCCTCTTGTGGCACATCATGACCAGGATGGGGACCATCACCATGGCGGACCTCCTGTACTCCTCCCTGGTCATCTTCTGGGCCCTGGTGCTCTCGAACTCCTCCCTGTCCGCCATCACCTGCGAGTGGATGGCCGCGCTCCTCTCGTAGAGCATGGAGTAGTGCTCCACGTCCGCGACCGAGACGGTGAGCCACTCCGTCTGGTACACCTCGGAGTACTCCTTCGTGTACTGGTTCCTGGCGATGGTCACGTCCATGTAGGACTTGCAGTCCTCCTTGCTCTTCAGCTTCTGGCACTCCATCCCCTCCATGTGCTTGTAGGGGGAGAATATCTTGACCCTGATCTGCTTCTCCTGGGTCAGGGCGGACCCGGCCTTCACGGCCAAGCAGTAGTCCCCGAACTCCTTGATGACCGTGAAGCGGGAGACCTTCTTGCCCGAGTCCCCGTTCCTCTTGGCCCTCCTCCTCCCCTCCAGCCTGGCGATGTTCCTGTAGAGGTCCGTCCAGAAGTGGGTCACCTCCGCCACGGCCAACTCGTTGAACATCTTCATCTGGGGGTCCTTGTACACCTTGGCCATGTTGTTGGTGAACTCGAAGGTGGCGTCTCTGGACATCTTCTCCACCATGTTCTCCATGAACTTCCTGTCCAAGAGGTTGGCCCCGATCCCCGCCATGTTCTCCTTCTTCTTGGTGTTCCCCGCCTTGGTGCAGTCGTTCAAGAAGATGGTGCCGTCGAAGAGGGAGCCCCCCTTCGAGTCCGTCACGTGCATCTTGACGTCCTTCATGTCGTTCAACTTCTGGGGGATCGGCATCGGGAAGGACTTCCCGTAGTCCTTCTTGGCCCTATTCATCTCCTCCGTGGGGGACAGCTCGTCCATCTTGTTCAACTCGGAGACGATCTCCTCCTGCGAGGTGCAGACCGCCTTCCTGATCTCCTCCGTGTACTTCGCGTTGGTGAACATGTCCTTGAAGTCCGAGACCAGCTCCTCCCTGAAGGCCTCCCCGTCGAAGTTCTTGCAGTCCTCCTTCGCGTCGGCCACCATCTTGTCCTCCTTGTTCTTCACGAACTCCTCGCTCCTGTAGAAGGTCTTGGCCTCCGCCCTCATCTTCTTGGCCCTCATCTCGTCCAAGGACTCCACCTCCGAGAAGATGTCGGTGCACACCTCGATCATCCTCTGCACCTTCTCGTCGGCCTTCTCGTCCACCTCCTCCCCCTCCCACTGGGTGTGCAAGGCCCTGTACCCCTTGTAGGAGCTGAAGCCCTTCATCATGATGTCCGCCACCTGGATCTTGCTGGCGGCGGACTCCAAGGAGGCCATCACGTGGGCCGTCTTGTACTTGGCGGGGCACATGATCTCGTTGGTCCTGGTGGAGTACACCAGCACGTCCACGGTCCCCCTGGTGCAGATGTTCTGGGCCACCATATCCTTCATGACGGGGTCATACTTCTCCCTCTTGGACAGGGCCCTGTCGGAGCAGATGCCGTCCGTGACGGCCACGTCGATCATGTCCAGGCAGTTCCCGGGGGCCGAGAAGAGCATGTCGGGGGTCTTGTTCGCGAACCTGCCGTGGTAGTTGACCTTCTTCAAGTCGACGTCCATCTCCCCCCTCTCGCACCCCACCGAGGTCATCCCCAGGGAGTCCAGGAAGGCCTTCCCCAACAGGTCGTGCCTGGACCTGAGGTAGACGTCCACCACCTCCACGGACATCTCCGGGCCGTCCTTCTTGTTGATGGAGTCCACGTACTCCACCAGCTCGTCCAGGCTCTCCATGGGCTTCACCTCCTTCCTGACCTCCTTCACCTTGTCCCTGGACATGTGCTCCTCCACCTCCTCCAGCAACTCGCACACCCTCTCCACCTTCAACATCTTCTGCATCGCCCTGGTCCTCTCCTCCGTGTTCGTGCCCACCCTGCCGGAGAAGGGGTACAGGATCTTGTCCCCCTTCATGCCGGACATCTTCTGCGCGTAGAACTCCTCCCCCGTGATCACGAACCTGGCCATGTAGGTCCCCCTGATGGTCTTCGGGGTAGAGAACAGCTCCAAGTACTCCTTGTAGGAGATCTCCTCGAACTCGTCCGTCTCCATGTCGGTCAACTCGAAGGCGTTGGGGGAGCGGCTGAAGGACTGCTCGGAGTTGGGCATCTCTATGGCGGTCTTCATGGACTCCGTCAGCATCCCGACCACCTTCTCCTTCGAGTGGTTCTCCAGCATGTTGGTGTAGATGCGGGAGGGGGTCACCTGTATGTCCCCCCCCATGGAGTTGATCACGCCCGAGAGGCCCCTGTGGAAGGTGATCCTATAGGGGACCGACTCCATGACGATGGAGTAGGAGACCTTCATCAGGAAGGGGTGCATCGGCCCCTGGAGGAACTCCTTCATGTTGATGCACGGTGATCTCCACGATCGTGAAGCTGGTGTTCGACCTCTTGTTCCTGGTTGAGATGGTGATATTATGATTTAGTAGAAACGTTGTGT

**>SalaUV-NL1-1-16_SL**

ACACAATGTCTTCTACTTATTAATACTATCATGCTGACCAACCTGAAGATGAACATCAGCCTGCTCCTTGCACAGGAACTACAAGATGGACAGCAAGCTCCTCCTCTTGTCCTGGACCTTGGTCACCATCCTGTCCAAGGCCACCGCCCTCTTCGTGTTGAAAGCGATGTCCAACTTGATGAGGGAGTTGAGGGTGGGGGTCTTCTCCCCGATGGACGAGAAGATCTCCGTGACCAAGCCCGTCGTGACCATCTGCACCATGACGACGAGGGTGTCCGAGGCGGAGTTGTACCCCTTGTTCTCCTTGAAGTCCCTCAGGGCGTCCATCTGGGCCAGGCCGGACATGCCGGTGGTGGGGTTGATCGTCCCGGTCATCTTCATGTAGACGGCCGTGTACACCATGTTGCACACCACCGCCAGCTTCCTCTTCTTCTCGTCCCCGATGAAGTTCCTCTGGATGTCGTTGACCGTGTTCATCACCAGCCTCTGGTCGAACGAGTAGAACTTCTCGAAGACCCTCTTCACTGAGGTGTAGACCGTGCCCATGTACTCCCTCAAGGACATCCCCTCCTCCAGGGGCTCCTCCTCCATGTGGGCGGTGGTGTCGATCTCCTTCAACCCCGAGAAGGCCCCGGCCAGCCCGGCGAGGGAGGTGAAGTCCATGGACATCCCGCCCATCTCCTCCTTCCCGATGGAGGAGAACATCTCGGCGTTCAGCTTGGTGGCCATGGAGGACGAGGAGATCCCGAAGTCCGTGGCGCTGATGGCGCCCATGACGTTCTCGACGCTGTAGTCCTCCACGGCCTGGGACACGTCCATCATCTCGTATATCGTGGAGTCCACCTCCGCCTCGAACTCGGCCTCCTTGATCCTGTTGTCCGGGGCGAACATGGAGTTGTCCGCTATCCCCAGGTCGTTCAGGATCTCCTCCACGTCCGACAAGTCCAGCTTGCTCGAGACGACGAGCTCGGAGAACTTCCTTATGTTGGGGTTCTCAGAGTCCTCGTTGAACGGCATCTTGTAGCTGTTGGTGCCGATGTGGAAGAACACCTTCCCCCCCAGCTCCCTCTCGTTGAACATGGCCAACACGGTCCACTTGGTCATCCTCCTGGTCAGGATCAGCTCCGTCTTGGGCCAGATCCTGTAGACCTTGATCCACAGGTTCCCGATCTTGGCGTTCTTCTCCACCCCCAAGTTGTCCGTCCTGGGGGCCCAGGGCCTGATCTGGATGGTGGAGAGGTCCTTGTGCATCAGCCTGGTGATGTTCCCCAACTCCACGTCGGAGCTGCTGACCTTGTACATCCTCACCGAGACGGTGTGCGGGGACGCCTGCCTGTCCTCGTGGATGATCCCGAGGTGCACCAAGTCGGTGTAGTACCTGGTCTTCATCTTCCCGTTCACGTCGACCTTGTAGAACACCTTGAGGGACTTGGCCCCCTGGAAGGAGTACTTGTCCTCGTCGGTGGCCGAGACGGCCATCTTCGCGAACCTCACCTCCAAGGAGTCCGACGGGATCACGGTCAGCAAGGGGTTGTCGATGCCCCAGGGATCCCTCACCAAGTTCTCCATCGGGCCCTCCGTGAGGTCCCCCCTGAACTCCTCCACCTCGAAGCGATCCACCCTCCCCCTCATCGACAAGGCGCTGTTGAACCTCTCCTCCGACACCACCCTGAGGGACTTCTTCGGGACGAACATCTGCAAGATCCCGCGGTTCGAGTACTTCTCCAAGTAGAGCATCCTCATGTTGGTGAGCGCCTCGGAGTTGTCCGGGAAGGAGACCATCATCTCCATGTTGAGGTCGGAGGTCATCTTCAGCCAGTACATGATGTAGCTCTTCAGCATCGTCAGGGGCCTCCTCACCCCCGGGAAGGCCTTCTTGATCTGGTATATGGGGTACTCGATGAACTTCTCCAACTTGACCCCCAGCGACTCGGAGATCTTGTTGGCCTCCGTCATGGCCGAGTTGCTAGCGTTGGAGGTGGCCCTCATCATGTACCTGAAGACCTCCATGTCGGACACCTTCACCCTGTCCTTCATCGGGGTGAAGTTGATCTTCCTGGACTTGTTGTGGCTGTAGGAGTGGGAGGGGATGGCGATCTCGGCCTCCTTCTCCGCGATCTCCCCCATCTCGCAGATCTCCTTGAAGGGGGCGGTGAGGTGGAACTTGGACTTGGTGTCGTCGTTCTTCAGGATCTTGTCCACCACCCCCACGAAGTCCGTGGCCACCACCCTGGAGGAGGAGATCAACGCGATGTTCTCCCCCGCCATGGCCAAGGCCCTGATCAAGATGTGGACCCTGATGGAGTCGTTGAACCCGAACTTCCTGTCCATCTGGAACATGTAGGTGTCCATGTTGGAGAAGAAGCTGGCCGAGTCGCTGCGGTCGGAGAAGGACAAGGCCGTGTTCCTCTCGGCCAGCTTGTCGATGTCCTCCCTCGTGATGGACCTCTTGTTCATCAGCCTCTTCTTCATCTCCTTGATCTTCGTGTCGAACCTGAAGGGGAGGGTCATCTTCGTCTTGCCGTTCATGGAGGCCATGTAGTCCCTGTCCTCCTCCATCTCCTCCTCGGTGTAGTCCGGGGAGGCGGCCATCTCGACGTCGGTCATGTAGGCGGCGTAGTACTTGACGTAGAACTCCATCACCTTCTCGGAGTTGCCCTCCTTGTACATCAGCACCTCCGGGCCGGCGATCATCTGCTCCACCACCCTGTCGGTCGACATGTACCCGAGCTGGAAGGGCAGGGAGTTCCTGTCGCACTCCAGCTTCCTCATGATCTCCTCCTTCCTCTCCCCGTCGATGGCGTAGATGGACGAGAGCGCCACGGACTGCAGCTCGAACATCACCTTGATCGTGGGGATCTTGACCCCGTTCTCGAACGCGTTCCGCATCTGGGATATGGAGGACTTCACCGCCTTCTCCGGCTCGGTCATGTCCACGATGTTCTGGGAGTTGTAGACGGCCTTCAGCACGCTGTCCGTGGCCCTCTTCCCGGTGGAGAAGTAGGAGTTGAACTCGGTCAAGATGGACAAGGCGGACTTCTTCCAGTTGGTCGTGATGTTGGCCATCCTGTTGGTCATGTCGTGCACCTGGAGCATGGACACCATGAAGGCCGTCAAGTCGGACATCTTCGTGTACCTGACCAGGAAGATCTTCGTGAGGTCGTCCGAGGAGATGAGGGTGGTGCACGCGGCGTCCCCGTACGACTTGGAGAACAGGAAGCACATCACCGAGTCCATGACGTCGTCCTTCGCGCAGTGGAACAAGGAGGAGAAGTAGTGGAGCATGCCCTGCCCCATCCCGGACTTGAATATGGCGGTCCCCCTCGAGGAGACGAAGTTCTCCTTGAGCCAGTTTATGTTCTTGTCGTCGGTCTTCTCGTTCCACAGCTTGTCCGTCTGCCACCTCCTGGTCAGGGTGGGCGGGAGGAACATGGTCTTCGTCGCGAAGGCCCGGATCACGGTGACGCAGAACTCCTTCAGGTTCACCGGCAAGGGCATCATCAAGCAGAAGTAGATGAAGCTATCCATCACGAACCCGGGGGACCACCTGGAGGCGTCCGTGTTGAAGGACAGGATGATGCCCGAGGGGGAGGTCTCCGACTCGTCCTTGTGCTTGTTCATCTCCAAGATCATCTTCTTGTACACGCTGGTGGCGTTGGACTGGAGGTCCTTCTTCACGTCCGCGTGCGTGAGCATCTCCTTCTCGTGCAAGGAGCAGAGCTCGTAGAACATCTTCTCCAGGAACTTCACGTGCAACCTCAACTTGGCCGCCTGTATCAGGATCTCCCTCGGCCCCCCGATCTGGGACTTCGGGAAGAGCTTGAACACGGCGTTCACCACGTCCGTGCAGGTGACCATGTCCATCAGGAAGATGGACTCCAACTCCTCCACCTGCTCGCACATCGTCTCGAAGGCGGTCGCCGTGGTCAGCTTGTCCGAGTAGTCCACCGACTCCTCGTACAGGGGGCACGACTTCAGGGACGACCTCATGGACATCGTCTCCCTTATGGTGGTGGTCATCGCCCTCATCAGGCCCTTGTTCAAGTCGGGGGAGTGGATTATCTTCATCCTGAGCTGGTAGCCCATGTACATGACGAACTTCTTGTTGAAGGTGTGGGTCTGCATCTTGTTCTTCGCGAGGGCGTCGAAGTCCGCGTCGCCCGTGGAGGCCCCGGGGAACAGCTCCGACATGGACAAGTAGTGCCTCTCCTGCTCGGACATCTTGGTCATGATGGGCACCAGCCTGTGGCCCGACGTCCCCTTCTCCTTGTTGAAGAGGTTGCTCATGTAGATGTCGTTCATCACCTCCGCGAACTCGACCAGCATCCCGGCCACCTTGAACGAGGGCATGTATATCCTGTCGTAGTCCGCGAACTCCGAGGCCATCTTCTCCGCGTACTGCTTTATGATGTCCGGGGCGGCCATGGTCAGCCTCTCGGCGTAGTCCATCTGCATCGCCCTGATGGAGGCCTCGAACATCGTCCTGACGGGGTCCTTCATGATGTTCTTCACCATGCTCACCTTATCCGAGGCGAACGCCGTGAGGCCGTGCATGATGTACCTGTTGTACTGGAGCGAGGTGGACACCCCCCTCTTGTGGCACATCATGACCAGGATGGGGACCATCACCATGGCGGACCTCCTGTACTCCTCCCTGGTCATCTTCTGGGCCCTGGTGCTCTCGAACTCCTCCCTGTCCGCCATCACCTGCGAGTGGATGGCCGCGCTCCTCTCGTAGAGCATGGAGTAGTGCTCCACGTCCGCGACCGAGACGGTGAGCCACTCCGTCTGGTACACCTCGGAGTACTCCTTCGTGTACTGGTTCCTGGCGATGGTCACGTCCATGTAGGACTTGCAGTCCTCCTTGCTCTTCAGCTTCTGGCACTCCATCCCCTCCATGTGCTTGTAGGGGGAGAATATCTTGACCCTGATCTGCTTCTCCTGGGTCAGGGCGGACCCGGCCTTCACGGCCAAGCAGTAGTCCCCGAACTCCTTGATGACCGTGAAGCGGGAGACCTTCTTGCCCGAGTCCCCGTTCCTCTTGGCCCTCCTCCTCCCCTCCAGCCTGGCGATGTTCCTGTAGAGGTCCGTCCAGAAGTGGGTCACCTCCGCCACGGCCAACTCGTTGAACATCTTCATCTGGGGGTCCTTGTACACCTTGGCCATGTTGTTGGTGAACTCGAAGGTGGCGTCTCTGGACATCTTCTCCACCATGTTCTCCATGAACTTCCTGTCCAAGAGGTTGGCCCCGATCCCCGCCATGTTCTCCTTCTTCTTGGTGTTCCCCGCCTTGGTGCAGTCGTTCAAGAAGATGGTGCCGTCGAAGAGGGAGCCCCCCTTCGAGTCCGTCACGTGCATCTTGACGTCCTTCATGTCGTTCAACTTCTGGGGGATCGGCATCGGGAAGGACTTCCCGTAGTCCTTCTTGGCCCTATTCATCTCCTCCGTGGGGGACAGCTCGTCCATCTTGTTCAACTCGGAGACGATCTCCTCCTGCGAGGTGCAGACCGCCTTCCTGATCTCCTCCGTGTACTTCGCGTTGGTGAACATGTCCTTGAAGTCCGAGACCAGCTCCTCCCTGAAGGCCTCCCCGTCGAAGTTCTTGCAGTCCTCCTTCGCGTCGGCCACCATCTTGTCCTCCTTGTTCTTCACGAACTCCTCGCTCCTGTAGAAGGTCTTGGCCTCCGCCCTCATCTTCTTGGCCCTCATCTCGTCCAAGGACTCCACCTCCGAGAAGATGTCGGTGCACACCTCGATCATCCTCTGCACCTTCTCGTCGGCCTTCTCGTCCACCTCCTCCCCCTCCCACTGGGTGTGCAAGGCCCTGTACCCCTTGTAGGAGCTGAAGCCCTTCATCATGATGTCCGCCACCTGGATCTTGCTGGCGGCGGACTCCAAGGAGGCCATCACGTGGGCCGTCTTGTACTTGGCGGGGCACATGATCTCGTTGGTCCTGGTGGAGTACACCAGCACGTCCACGGTCCCCCTGGTGCAGATGTTCTGGGCCACCATATCCTTCATGACGGGGTCATACTTCTCCCTCTTGGACAGGGCCCTGTCGGAGCAGATGCCGTCCGTGACGGCCACGTCGATCATGTCCAGGCAGTTCCCGGGGGCCGAGAAGAGCATGTCGGGGGTCTTGTTCGCGAACCTGCCGTGGTAGTTGGCCTTCTTCAAGTCGACGTCCATCTCCCCCCTCTCGCACCCCACCGAGGTCATCCCCAAGGAGTCCAGGAAGGCCTTCCCCAGCAAGTCGTGCCTGGACCTGAGGTAGACGTCCACCACCTCCACGGACATCTCCGGGCCGTCCTTCTTGTTGATGGAGTCCACGTACTCCACCAGCTCGTCCAGGCTCTCCATGGGCTTCACCTCCTTCCTGACCTCCTTCACCTTGTCCCTGGACATGTGCTCCTCCACCTCCTCCAGCAACTCGCACACCCTCTCCACCTTCAACATCTTCTGCATCGCCCTGGTCCTCTCCTCCGTGTTCGTGCCCACCCTGCCGGAGAAGGGGTACAGGATCTTGTCCCCCTTCATGCCGGACATCTTCTGCGCGTAGAACTCCTCCCCCGTGATCACGAACCTGGCCATGTAGGTCCCCCTGATGGTCTTCGGGGTAGAGAACAGCTCCAAGTACTCCTTGTAGGAGATCTCCTCGAACTCGTCCGTCTCCATGTCGGTCAACTCGAAGGCGTTGGGGGAGCGGCTGAAGGACTGCTCGGAGTTGGGCATCTCTATGGCGGTCTTCATGGACTCCGTCAGCATCCCGACCACCTTCTCCTTCGAGTGGTTCTCCAGCATGTTGGTGTAGATGCGGGAGGGGGTCACCTGTATGTCCCCCCCCATGGAGTTGATCACGCCCGAGAGGCCCCTGTGGAAGGTGATCCTATAGGGGACCGACTCCATGACGATGGAGTAGGAGACCTTCATCAGGAAGGGGTGCATCGGCCCCTGGAGGAACTCCTTCATGTTGATGCACGGTGATCTCCACGATCGTGAAGCTGGTGTTCGACCTCTTGTTCCTGGTTGAGATGGTGATATTATGATTTAGTAGAAACGTTGTGTA

**>SalaUV-NL1-1-26_SL**

ACACAATGTCTTCTACTTTATAATACTATCATGTTGGCCAACCAGAAGATGAACATCAGCCTGCTCCTCGTACAGGAACTACAAGATGGACAGCAAGCTCCTCCCCTTGTCCTGAACCTTGGTCACCATCCTGTCCAAGGCCACCGCCCTCTTCGTGTTGAACGCGATGTCCAGCTTGATGAGGGAGTTGAGGGTGGGGGCCTTCTCCCCGATGGACGAGAAGATCTCCGTGATCAAGCCCGTCGTCACCATCTGCACCATGACGACGAGGGTGTCCGAGGCGGAGTTGTACCCCTTGTTCTCCTTGAAGTCCCTCAGGGCGTCCATCTGGGCCAGGCCCGACATGCCCGTGGTGGGGTTGACCGTCCCGGTCATCTTCATGTAGACGGCCGTGTACACCATGTTGCACACCACCGCCAGCTTCCTCTTCTTCTCGTCCCCGATGAAGTTCCTCCGGATGTCGTTGACCGTGTTCATCACCAGCCTCTGGTCGAACGAGTAGAACTTCTCGAAGACCCTCTTCACCGAGGTGTAGACCGTGCCCATGTACTCCCTCAAGGACATGCCCTCCTCCAAGGGCTCCTCCTCCATGTGGGCGGTGGTGTCGATCTCCTTCAACCCCGAGAAGGCCCCGGCCAGCCCGGCGAGGGAGGTGAAGTCCACGGCCATCCCGCCCATCTCCTCCTTCCCGATGGAGGAGAACATCTCGGCGTTCAGCTTGGTGGCCATGGAGGACGAGGAGATCCCGAAGTCCGTGGCGCTGATGGCGCCCATGACGTTCTCGACGCTGTAGTCCTCCACGGCCTGGGACACATCCATCATCTCGTATATCGTGGAGTCCACCTCCGCCTCGAACTCGGCCTCCTTGACCCTGTTGTCCGGGGCGAACATGGAGTTGTCCGCTATCCCCAGGTCGTTCAGGATCTCCTCCACGTCCGACAAGTCCAGCTTGCTCGAGACGACGAGCTCGGAGAACTTCCTGATGTTGGAGTTCTCGGAGTCCTCGTTGAACGGCATCTTGTAGCTGTTGGTGCCGATGTGGAAGAACACCTTCCCCCCCAACTCCCTCTCGTTGAACATGGCCAACACGGTCCACTTGGTCATCCTCCTGGTCAGGATCAGCTCCGTCTTGGGCCAGATCCTGTAGACCTTGATCCACAGGTTCCCGATCTTGGCGTTCTTGGCCACCTCCAAGTTGTCCGTCCTGGGGGCCCAGGGCCTGATCTGGATGGTGGAGAGGTCCTTGTGCATCAGCCTTGTGATGTTCCCCAACTCCACGTCGGAGCTGCTGACCTTGTACATCCTCACCGAGACGGTGTGCGGGGACGCCCCCCTGTCCTCGTGGATGATCCCGAGGTGCACCAAGTCGGTGTAGTACCTGGTCTTCATCTTCCCGTTCACGTCGACCTTGTAGAACACCTTGAAGGACTTCGCCCCCTGGAAGGAGTACTTGTCCTCGTCGGTGGCCGAGACGGCCATCTTCGCGAACCTCACCTCCAAGGAGTCCGACGGGATCACGGTCAGCAGGGGGTTGTCGGTGCCCCAGGGGTCCCTCACCAGGTTCTCCATCGGCCCCTCCGTGAGGTCCCCCCTGAACTCCTCCACCTCGAAGCGGTCCACCCTCCCCCTCATCGACAAGGCGCTGTTGAACCTCTCCTCCGAGACCACCCTGAGGGACTTCTTCGGGACGAACATCTGCAAGATCCCGCTGTTCGAGTACTTCTCCAAGTAGAGCATCCTCATGTTGGTGAGCGCCTCGGAGTTGTCCGGGAAGGAGACCATCATCTCCATGTTGAGGTCGGAGGTCATCTTCAGCCAGTACATGATGTAGCTCTTCAGCATCGTCAAGGGCCTCCTCACCCCCGGGAAGGCCTTCTTGATCTGGTATATGGGGTACTCGATGAACTTCTCCAACTTGACCCCCAGCGACTCGGAGATCTTGTTGGCCTCCGTCATGGCCGAATTGCTGGCGTTGGAGGTGGCCCTCATCATGTACCTAAAGACCTCCATGTCGGACACCTTCACCCTGTCCTTCATCGGGGTGAAGTTGATCCTCCTAGACTTGTTGTGGCTGTAGGAGTGGGAGGGGATGGCGATCTCCGCCTCCTTCTCCGCGATCTCCCCCATCTCGCAGATCTCCTTGAAGGGGGCCGTGAGGTGGAACTTGGACTTGGTGTCGTCGTTCTTCAAGATCTTGTCCACCACTCCCACGAAGTCCGTGGCCACCACCCTGGAGGAGGAGATCAGCGCGATGTCCTCCCCCGCCATGGCCAAGGCCCTGATCAAGATGTGGACCCTGATGGCGTCGTTGAACCCGAACTTCCTGTCCATCTGGAACATGTAGGTGTCCATGTTGGAGAAGAAGCTGGCCGAGTCGCTGCGGTCGGAGAAGGACAAGGCCGTGTTCCTCTCGGCCAGCTTGTCGATGTCCTCCCTCGTGATGGACCTCTTGTGCATCAGCCTCTTCTTCATCTCCTTGATCTTGGTGTCGAACCTGAAGGGAAGGGTCATCTTCGTCTTCCCGTTCATGGAGGCCATGTAGTCCCTGTCCTCCTCCATCTCCTCCTCGGTGTAGTCCAGGGAGGCGGCCATCTCGACGTCGGTCATGTAGGCGGCGTAGTACTTGACGTAGAACTCCATCACCTTCTCGGAGTTGCCCTCCTTGTACATCAGCACCTCCGGGCCGGCGATCATCTGCTCCACCACCCTGTCGGTCGACATGTACCCGAGCTGGAAGGGCAGGGAGTTCCTGTCGCAGTCCAGCTTCCTCATGATCTCCTCCTTCCTCTCCCCGTCGATGGCGTAGATGGACGAGAGCGCCACGGACTGCAGCTCGAACATCACCTTGATCGTGGGGATCTTGACCCCGTTCTCGAACGCGTTCCTCATCTGGGATATGGAGGACTTCACCGCCTTCTCCGGCTCGGTCATGTCCACGATGTTCTGGGAGTTGTAGACAGCCTTCAGCACGCTGTCGGTGGCCCTCTTCCCGGTGGAGAAGTAGGAGTTGAACTCGGTCAAGATGGACAAGGCGGACTTCTTCCAGTTGGTCGTGATGTTGGCCATCCTGTTGGTCATGTCGTGCACCTGGAGCATGGACACCATGAAGGCCGTCAAGTCGGACATCTTCGTGTACCTGACCAGGAAGATCTTCGTGAGGTCGTCGGAGGAGATGAGTGTGGTGCACGCGGCGTCCCCGTATGACTTGGAGAACAGGAAGCACATCACCGAGTCCATGACGTCGTCCTTCGCGCAGTGGAACAAGGAGGAGAAGTAGTGGAGCATGCCCTGCCCCATCCCGGACTTGAATATGGCGGTCCCCCTCGAGGAGACGAAGTTCTCCTTGAGCCAGTTTATGTTCTTGTCGTCGGTCTTCTCGTTCCACAGCTTGTCCGTCTGCCACCTCCTGGTCAGGGTGGGCGGGAGGAACATGGTCTTCGTCGCGAAGGCCCTGATCACGGTGACGCAGAACTCCTTCAGCTTCACCGGCAAGGGCATCATCAAGCAGAAGTAGATGAAGCTGTCCATCACGAACCCGGGGGACCACCTGGAGGCGTCCGTGTTGAAGGACAGGATGATGCCCGAGGGGGAGGTCTCCGACTCGTCCTTGTGCTTGTTCATCTCCAAGATCATCTTCTTGTACACGCTGGTGGCGTTGGACTGGAGGTCCTTCTTCACGTCCGCGTGCGTGAGCATCTCCTTCTCGTGCAAGGAGCAGAGCTCGTAGAACATCTTCTCCAAGAACTTCACGTGCAACCTCAACTTGGCCGCCTGTATCAGGATCTCCCTCGGCCCCCCAATCTGGGACTTCGGGAAGAGCTTGAAGACGGCGTTCACCACGTCCGTGCAGGTGACCATGTCCATCAGGAAGATGGACTCCAACTCCTCCACCTGCTCGCACATCGTCTCGAAGGCGGTCGCCGTGGTGAGCTTGTCCGAGTAGTCCACCGACTCCTCGTACAGGGGGCACGACTTCAGGGAGGACCTCATGGACATCGTCTCCCTTATGGTGGTGGTCATCGCCCTCATCAGGCCCTTGTTCAAGTCGGGGGAGTGGATTATCTTCATCCTGAGCTGGTAGCCCATATACATGACGAACTTCTTGTTGAAGGTGTGGGTCTGCATCTTGTTCTTCGCGAGAGCGTCGAAGTCCGCGTCGCCCGTGGAGGCCCCGGGGAACAGCTCCGACATGGACAAGTAGTGCCTCTCCTGCTCCGACATCTTGGTCATGATGGGGACCAGCCTGTGGCCCGCCGTCCCCTTCTCCTTGTTGAAGAGGTTGCTCATGTAGATGTCGTTCATCACCTCCGCGAACTCGACCAACATCCCCGCCACCTTGAACGAGGGCATGTATATCCTGTCGTAGTCCGCGAACTCCGAAGCCATCTTCTCCGCGTACTGTTTTATGATGTCCGGGGCGGCCATGGTCAGCCTCTCGGCGTAGTCCATCTGCATCGCCCTGATGGAGGCCTCGAACATCGTCCTGACGGGGTCCTTCATGATGTTCTTCACCATGCTCACCTTGTCCGAGGCGAACGCCGTGAGGCCGTGCATGATGTACCTGTTGTACTGGAGCGAGGTGGACACCCCCCTCTTGTGGCACATCATGACCAGGATGGGGACCATCACCATGGCGGACCTCCTGTACTCCTCCCTGGTCATCTTCTGGGCCCTGGTGCTCTCGAACTCCTCCCTGTCCGCCATCACCTGCGAGTGGATGGCCGCGCTCCTCTCGTAGAGCATGGAGTAGTGCTCCACGTCCGCGACCGAGACGGTGAGCCACTCCGTCTGGTACACCTCGGAGTACTCCTTCGTGTACTTGTCCCTGGCGATGGTCACGTCCATGTAGGACTTGCAGTCCTCCTTGCTCTTCAGCTTCTGGCACTCCATCCCCTCCATGTGCTTGTAGGGGGAGAATATCTTGACCCTGATCTGCTTGTCCTGGGTCAGGGCGGACCCGGCCTTCACGGCCAAGCAGTAGTCCCCGAACTCCTTTATGACCGTAAAGCGGGACACCTTCTTGCCCGAGTCCCCGTTCCTCTTGGCCCTCCTCCTCCCCTCCAGCCTGGCGATGTTCCTGTAGAGGTCCGTCCAGAAGTGGGTCACCTCCGCCACGGCCAACTCGTTGAACATCTTCATCTGGGGGTCCTTGTACACCTTGGCCATGTTGTTGGTGAACTCGAAGGTGGCGTCCCTGGACATCTTCTCCACCATGTTCTCCATGAACTTCCTGTCCAAGAGGTTGGCCCCGATGCCCGCCATGTTCTCCTTCTTCTTGGTGTTCCCCGCCTTGGTGCAGTCGTTCAAGAAGATGGTGCCGTCGAAGAGGGAGCCCCCCTTCGAGTCGGTGACGTGCATCTTGACGTCCTTCATGTCGTTCAGCTTCTGGGGGATCGGCATCGGGAAGGACTTCCCGTAGTCCTTCTTGGCCCTGTTCATCTCCTCCGTGGGGGACAGCTCGTCCATCTTGTTCAACTCGGAGACGATCTCCTCCTGCGAGGTGCAGACCGCCTTCTTGATCTCCTCCGTGTACTTCGCGTTGGTGAACATGTCCTTGAAGTCCGAGACCAGCTCCTCCCTGAAGGCCTCCCCGTCGAAGTTCTTGCAGTCCTCCTTCGCGTCGGCTACCATCTTGTCCTCCTTGTTCTTCACGAACTCCTCGTTCCTGTAGAAGGTCTTGGCCTCCGCCCTCATCTTCTTGGCCCTCATCTCGTCCAGGGACTCCACCTCCGAGAAGATGTCGGTGCACACCTCGATCATCCTCTGCACCTTCTCGTCGGCCTTCTCGTCCACCTCCTCCCCCTCCCACTGGGTGTGCAAGGCCCTGTACCCCTTGTAGGAGTTGAAGCCCCTCATCATGATGTCCGCCACCTGGATCTTGCTGGCGGCGGACTCCAGGGAGGCCATCACGTGGGCCGTCTTGTACTTCTCGGGGCACATGATCTCGTTGGTCCTGGTGGAGTACACCAGCACGTCCACGGTCCCCCTGGTGCAGATGTTCTGGGCCACCATATCCTTCATCACGGGGTCGTACTTCTCCTTCTTGGACAGGGCCCTGTCGGAGCAGATGCCGTCCGTGACGGCCACGTCGATCATGTCCAAACAGTTCCCGGGGGCCGAGAAGAGGATGTCGGGGGTCTTGTTCGCGAACTTGCCGTGGTAGTTGGCCTTCTTCAAGTCGACGTCCATCTCCCCCCTCTCGCACCCCACCGAGGTCATCCCCAGGGAGTCCAGGAAGGCCTTCCCCAACAGGTCGTGCCTGGACCTGAGGTAGACATCCACCACCTCCACGGACATCTCCGGGCCGTCCTTCTTGTTGATGGAGTCCACGTACTCCACCAGCTCGTCCAGGCTCTCCATGGGCTTCACCTCCTTCCTGACCTCCTTCACCTTGTCCCTGGACATGTGCTCCTCCACCTCCTCCAGCAACTCGCACACCCTCTCCACCTTCAACATCTTCTGCATCGCCCTGGCCCTCTCCTCCGTGTTCGTACCCACCCTGCCGGAGAAGGGGTACAGGATCTTGTCCCCCTTCATGCCGGACATCTTCTGCGCGTAGAACTCCTCCCCCGTGATCACGAACCTGGCCATGTAGGTCCCCCTGATGGTTTTCGGGGTGGAGAACAGCTCCAAGTACTCCTTGTAGGAAATCTCCTCGAACTCGTCCGTCTCCATGTCGGTCAACTCGAACGCGTTGGGCGAGCGGCTGAAGGACTGCTCGGAGTTGGGCATCTCTATGGCGGTCTTCATGGACTCCGTCAACATCTTGACCACCTTCTCCTTCGAGTGGTTCTCCAGCATGTTAGTGTAGATGCGGGAGGGGGTCACCTGTATGTCCCCGCCCATGGAGTTGATCACGCCCGAGAGGCCCCTGTGGAAGGTGACCCTGTAGGGGACCGACTCCATGACGACAGAGTAGGAGATCTTCATCAGGAAGGGGTGCATCGGCCCCTGAAGGAACTCCTTCATGCTGATGCACGGTGAGCTCCACGATCGCGAAGCTGGTGTTCGACCTCTTGTTCCTGTTCTAGATGATGATAGTATGATTTAGTAGAAACGTTGTGTA

**>SalaUV-NL1-1-27_SL**

ACACAATGTCTTCTACTTATTAATACTATCATGTTGTCCAACCCGAAGATGAACATTAGCCTGCTCCCTGTACAGGAACTACAAGATGGACAGCAAGCTCCTCCCCTTGTCCTGAACCTTGGTCACCATCCTGTCCAAGGCCACCGCCCTCTTCGTGTTGAAGGCGATGTCCAGCTTGATGAGGGAGTTGAGAGTGGGCGCCTTCTCCCCGATGGACGAGAAGATCTCCGTGATCAAGCCCGTCGTGACCATCTGCACCATGACGACGAGGGTGTCCGAGGCGGAGTTGTACCCCTTGTTCTCCTTGAAGTCCCTCAAGGCGTCCATCTGGGCCAGGCCGGACATGCCCGTGGTGGGGTTGACCGTCCCGGTCATCTTCATGTAGACGGCCGTGTACACCATGTTGCACACCACCGCCAGCTTCCTCTTCTTCTCGTCCCCGATGAAGTTCCTCTGGATGTCGTTGACCGTGTTCATCACCAGCCTCTGGTCGAAGGAGTAGAACTTCTCGAAGACCCTCTTCACCGAGGTGTAGACCGTGCCCATGTACTCCCTCAAGGACATCCCCTCCTCCAGGGGCTCCTCCTCCATGTAGGCGGTGGTGTCGATCTCCTTCAACCCCGAGAAGGCCCCGGCCAGCCCGGCGAGGGAGGTGAAGTCGATGGCCATCCCGCCCATCTCCTCCTTCCCGATGGAGGAGAACATCTCAGCGTTCAGCTTGGTGGCCATGGAGGACGAGGATATCCCGAAGTCCGTGGCGCTGATGGCGCCCATGACGTTCTCGACGCTGTAGTCCTCCACGGCCTGGGACACGTCCATCATCTCGTATATCGTGGAGTCCACCTCCGCCTCGAACTCGGCCTCCTTGACCCTGTTGTCCGGGGCGAACATGGAGTTGTCCGCTATCCCCAAGTCGTTCAGGATCTCCTCCACGTCCGACAAGTCCAACTTGCTCGAGACGACGAGCTCCGAGAACTTCCTGATGTTGGGGTTCTCGGAGTCCTCGTTGAACGGCATCTTGTAGCTGTTGGTGCCGATGTGGAAGAACACCTTCCCCCCCAACTCCCTCTCGTTGAACATGGCCAACACGGTCCACTTCGTCATCCTCCTGGTCAGGATGAGCTCCGTCTTGGGCCAGATCCTGTAGACCTTGATCCACAGGTTCCCGATCTTGGCATTCTTGTCCACCCCCAAGTTGTCCGTCCTGGGGGCCCAGGGCCTGATCTGGATGGTGGAGAGGTCCTTGTGCATCAGCCTTGTGATGTTCCCCAACTCCACGTCGGAGCTGCTGACCTTGTACATCCTCACCGAGACGGTGTGCGGGGACGCCCCCCTGTCCTCGTGGATGATCCCGAGGTGCACCAAGTCCGTGTAGTACCTGGTCTTCATCTTCCCGTTCACGTCGACCTTGTAGAACACCTTGAGGGACTTGGCCCCCTGGAAGGAGTACTTGTCCTCGTCGGTGGCCGAGACGGCCATCTTCGCGAACCTCACCTCCAAGGAGTCCGACGGGATCACGGTCAGCAGGGGGTTGTCGGTGCCCCAGGGGTCCCTCTCCAAGTTCTCCATCGGGCCCTCCGTGAGGTCCCCCCTGAACTCCCCCACCTCGAAGCGGTCCACCCTCCCCCTCATCGACAAGGCGCTGTTGAACCTCTCCTCCGACACCACCCTGAGGGACTTCTTCGGGACGAACATCTGCAAGATCCCGCAGTTCGAGTACTTCTCCAAGTAGAGCATCCTCATGTTGGTGAGCGCCTCGGAGTTGTCCGGGAAGGAGACCATCATCTCCATGTTGAGGTCAGAGGTCATCTTCAGCCAGTACATGATGTAGCTCTTCAGCATCGTCAAGGGTCTCCTCACCCCCGGGAAGGCCTTCTTGATCTGGTATATGGGGTACTCGATGAACTTCTCCAACTTGACCCCCAACGACTCGGAGATCTTGTTGGCCTCCGTCATGGCCGAGTTGCTGGCGTTGGAGGTGGCCCTCATCATGTACCTGAACACCTCCATGTCGGACACCTTCACCCTGTCCTTCATCGGGGTGAAGTTGATCCTCCTGGACTTGTTGTGGCTGTAGGAGTGGGAGGGGATGGCGATCTCCGCCTCCTTCTCCGAGATCTCCCCCATCTCGCAGATCTCCTTGAAGGGGGCGGTGAGGTGGAACTTGGACTTGGTGTCGTCGTTCTTCAGGATCTTGTCCACCACTCCCACGAAGTCCGTGGCCACCACCCTGGAGGAGGAGATCAACGCGATGTCCTCCCCCGCCATGGCCAAGGCCCTGATCAAGATGTGGACCCTGATGGCGTCGTTGAACCCGAACTTCCTGTCCATCTGGAACATGTAGGTGTCCATGTTGGAGAAGAAGCTGGCCGAGTCGCTGCGGTCGGAGAAGGACAAGGCGGTGTTCCTCTCGGCCAGCTTGTCGATGTCCTCCCTCGTGATGGACCTCTTGTTCATCAGCCTCTTCTTCATCTCCTTGATCTTCGTGTCGAACCTGAAGGGAAGGGTCATCTTGGTCTTGCCGTTCATGGAGGCCATGTAGTCCCTGTCCTCCTCCATCTCCTCCTCGGTGTAGTCCAGGGAGGCGGCCATCTCGACGTCGGTCATGTACGCGGCGTAGTACTTGACGTAGAACTCCATCACCTTCTCGGAGTTGCCCTCCTTGTACATCAGCACCTCCGGGCCGGCGATCATCTGCTCCACCACCCTGTCGGTCGACATGTACCCGAGTTGGAAGGGCAGGGAGTTCCTGTCGCAGTCCAGCTTCCTCATGATCTCCTCCTTCCTCTCCCCGTCGATGGCGTAGATGGACGAGAGCGCCACGGACTGCAGCTCGAACATCACCTTGATCGTGGGGATCTTGACCCCGTTCTCGAACGCGTTCCTCATCTGGGATATGGAGGACTTCACCGCCTTCTCCGGCTCGGTCATGTCCACGATGTTCTGGGAGTTGTAGACGGCCTTCAGCACGCTGTCCGTGGCCCTCTTCCCGGTGGAGAAGTAGGAGTTGAACTCGGTCAAGATGGACAAGGCGGACTTCTTCCAGTTGGTCGTGATGTTGGCCATCCTGTTGGTCATGTCGTGCACCTGGAGCATGGACACCATGAAGGCCGTCAAGTCGGACATCCTCGTGTACCTGACCAGGAAGATCTTCGTGAGGTCGTCGGAGGAGATGAGCGTGGTGCACGCGGCGTCCCCGTACGACTTGGAGAACAGGAAGCACATCACCGAGTCCATGACGTCGTCCTTCGCGCAGTGGAACAGGGAGGAGAAGTAGTGGAGCATGCCCTGCCCCATCCCGGACTTGAATATGGCGGTCCCCCTCGAGGAGACGAAGTTCTCCTTGAGCCAGTTTATGTTCTTGTCGTCGGTCTTGTCGTTCCACAGCTTGTCCGTCTGCCACCTCCTGGTCAGGGTGGGCGGGAGGAACATGGTCTTCGTGGCGAAGGCCCTGATCACGGTGACGCAGAACTCCTTCAACTTCACCGGCAAGGGCATCATCAAGCAGAAGTAGATGAAGCTATCCATCACGAACCCGGGGGACCACCTGGAGGCGTCCGTGTTGAAGGACAGGATGATGCCCGAGGGGGAGGTCTCCGACTCGTCCTTGTGCTTGTTCATCTCCAGGATCATCTTCTTGTACACGCTGGTGGTGTTGGACTGGAGGTCCTTCTTCACGTCCGCGTGCGTAAGCATCTCCTTCTCGTGCAAGGAGCAGAGCTCGAAGAACATCTTCTCCAAGAACTTCACGTGCAACCTCAACTTGGCCGCCTGTATCAGGATCTCCCTCGGCCCCCCAATCTGGGACTTCGGGAAGAGCTTGAAGACGGCGTTCACCACGTCCGTGCAGGTGACCATGTCCATCAGGAAGATGGACTCCAGCTCCTCCACCTGCTCGCACATCGTCTCGAAGGCGGTCGCCGTGGTCAGCTTGTCCGAGTAGTCCACCGACTCCTCGTACAGGGGGCACGACTTCAAGGACGACCTCATGGACATCGTCTCCCTTATGGTGGTGGTCATCGCCCTCATCAGGCCCTTGTTCAAGTCGGGGGAGTGGATTATCTTCATCCTGAGCTGGTAGCCCATGTACATGACGAACTTCTTGTTGAAGGTGTGGGTCTGCATCTTGTTCTTCGCGAGGGCGTCGAAGTCCGCGTCGCCCGTGGAGGCCCCGGGGAACAGCTCCGACATGGACAGGTAGTGCCTCTCCTGCTCGGACATCTTGGTCATGATGGGGACCAACCTGTGGCCCGCCGTCCCCTTCTCCTTGTTGAAGAGGTTGCTCATGTAGATGTCGTTCATCACCTCCGCGAACTCGACCAACATCCCGGCCACCTTGAACGAGGGCATGTATATCCTGTCGTAGTCCGCGAACTCCGAGGCCATCTTCTCCGCGTACTGCTTTATGATATCCGGGGCGGCCATGGTCAGCCTCTCGGCGTAGTCCATCTGCATCGCCCTGATGGAGGCCTCGAACATCGTCCTGACGGGGTCCTTCATGATGTTCTTCACCATGCTCACCTTGTCCGAGGCGAACGCCGTGAGGCCGTGCATGATGTACCTGTTGTACTGGAGCGAGGTGGACACCCCCCTCTTGTGGCACATCATGACCAGGATGGGGACCATCACCATGGCGGACCTCCTGTACTCCTCCCTGGTCATCTTCTGGGCCCTGGTGCTCTCGAACTCCTCCCTGTCCGCCATCACCTGCGAGTGGATGGCCGCGCTCCTCTCGTAGAGCATGGAGTAGTGCTCCACGTCCGCGACCGAGACGGTGAGCCACTCCGTCTGGTACACCTCGGAGTACTCCTTCGTGTACTGGTTCCTGGCGATGGTCACGTCCATGTAGGACTTGCAGTCCTCCTTGCTCTTCAGCTTCTGGCACTCCATCCCCTCCATGTGCTTGTAGGGGGAGAATATCTTGACCCTGATCTGCTTGTCCTGGGTCAGGGCGGACCCGGCCTTCACGGCCAAGCAGTAGTCCCCGAACTCCTTGATGACCGTAAAGCGGGACACCTTCTTGCCCGAGTCCCCGTTCCTCTTGGCCCTCCTCCTCCCCTCCAGCCTGGCGATGTTCCTGTAGAGGTCCGTCCAGAAGTGGGTCACCTCCGCCACGGCCAGCTCGTTGAACATCTTCATCTGGGGGTCCTTGTACACCTTGGCCATGTTGTTGGTGAACTCGAACGTGGCGTCCCTGGACATCTTCTCCACCATGTTCTCCATGAACTTCCTGTCCAAGAGGTTGGCCCCGATGCCCGCCATGTTCTCCTTCTTCTTGGTGTTCCCCGCCTTGGTGCAGTCATCCAAGAAGATGGTGCCGTCGAAGAGGGAGCCCCCCTTCGAGTCGGTCACGTGCATCTTGACGTCCTTCATGTCGTTCAGCTTCTGGGGGATCGGCATCGGGAAGGATTTCCCGTAGTCTTTCTTGGCCCTGTTCATCTCCTCCGTGGGGGACAGCTCGTCCATCTTGTTCAACTCGGAGACGATCTCCTCCTGCGAGGTGCAGACCGCCTTCTTGATCTCCTCCGTGTACTCCGCGTTGGTGAACATGTCCTTGAAGTCCGAGACCAGCTCCTCCCTGAAGGCCTCCCCGTCGAAGTTCTTGCAGTCCTCCTTCGCGTCGGCCACCATCTTGTCCTCCTTGTTCTTGACGAACTCCTCGCTCCTGTAGAAGGTCTTGGCCTCCGCCCTCATCTTCTTGGCCCTCATCTCGTCCAGGGACTCCACCTCCGAGAAGATGTCGGTGCACACCTCGATCATCCTCTGCACCTTCTCGTCGGCCTTCTCGTCCACCTCCTCCCCCTCCCACTGGGTGTGCAAGGCCCTGTACCCCTTGTAGGAGTTGAAGCCCCTCATCATGATGTCCGCCACCTGGATCTTGCTGGCGGCGGACTCCAGGGAGGCCATCACGTGGGCCGTCTTGTACTTCCCGGGGCACATGATCTCGTTGGTCTTGGTGGAGTACACCAGCACGTCCACGGTCCCCCTGGTGCAGATGTTCTGGGCCACCATATCCTTCATCACGGGGTCGTACTTCTCCCTCTTGGACAGGGCCCTGTCAGAGCAGATGCCGTCCGTGACGGCCACGTCGATCATGTCCAGGCAGTTCCCGGGGGCCGAGAAGAGGATGTCGGGGGTCTTGTTCGCGAACTTGCCGTGGTAGTTGGCCTTCTTCAAGTCAACGTCCATCTCCCCCCTCTCGCACCCCACCGAGGTCATCCCCAGGGAGTCCAGGAAGGCCTTCCCCAACAGGTCGTGCCTGGACCTGAGGTAGACGTCCACCACCTCCACGGACATCTCCGGGCCGTCCTTCTTGTTGATGGAGTCCACGTACTCCACCAGCTCGTCCAGGCTCTCCATGGGTTTCACCTCCTTCCTGACTTCCTTCACCTTGTCCCTGGACATGTGCTCCTCCACCTCCTCCAGCAGCTCGCACACCTTCTCCACCTTCAACATCTTCTGCATCGCCCTGGCCCTCTCCTCCGTGTTCGTACCCACCCTGCCGGAGAAGGGGTACAGGATCTTGTCCCCCTTCATGCCAGACATCTTCTGCGCGTAGAACTCCTCCCCCGTTATCACGAACCTGGCCATGTAGGTCCCCCTGATAGTCGTCGGGGTGGAGAACAGCTCCAAGTACTCCTTGTAGGAGATCTCCTCGAACTCGTCCGTCTCCATCTCGGTCAACTCGAACGCGTTGGGCGAGCGGCTGAAGGACTGCTCGGAGTTGGGCATCTCTATAGCGGTCTTCATGGACTCCGTCAACATCCTGACCACCTTCTCCTTCGAGTGGTTCTCCAGCATGTTGGTGTAGATGCGGGAGGGGGTCACCTGTATGTCCCCGCCCATGGAGTTGATCACGCCCGAGAGGCCCCTGTGGAAGGTGACCCTGTAGGGGACCGACTCCATGACGACAGAGTAGGAGACCTTCATCAGGAACGGGTGCATCGGCCCCTGAAGGAACTCCTTCATGTTGATGCACGGTGATCTCCACGATCGCGAAGCTGGTGTTCGACTTCTTGTTCCTGTTCGAGATGATGATATTATGATTTAGTAGAAACTTTGTGT

**>SalaUV-NL1-1-9_SS**

AAACACAAAGTCTTCTACTTATTATATTATCAGCATCTCGAACATGGAGACCTACTGCATCTTACTGTAGAAGACCTTGTGCGCCAGGGCGGACTACCTGGGCTTAAACTAGTGGGACCTGAATTACATGTTCCACCTCCAGTCCGTGTAGGTTACGGTGAACAAGACCCATTCCAACTCGACCTACTTCCCGAGGTCCTCCTCCTCCTGCTTGGCCATCCTGGCGCTGACCATCTCGAAGAGGGCGCCGAGGTCGTCGGTGGGGCAGATCGCCACCTCCACGTCGGAGGACCTCAGCCACTTGACCAAGCTGGGGCCCGTGTAGCTGGTCCTGACCATGTCGTCGGGCTTCATGTAGACGATCTCCATGTCCTGGACGACGGGCAAGAGGTACCTGTCCCCCTGCTTGGTCTTGTAGTACCTGGCCTGGAAGCCCCTCTCGAAGACGTTGGGGTCCGGGTTGTTCGACGTGGCGATGGTGATGTTCCAGTAGGACTGCTCCCACCAGAGGTGCGCGGTCTCGATCTCCTCGTCGAACCTGAATTGCGCCGCCCACTTGTTGGCCAGGAAGTTCCCCACCGTCCTCTCCTCGTACGGCGTGATCATCTTCCACACCAAGGCCGCCATGGGGGGGGAGGAGGAGCAGATGTTCTTCACGCAGAACTTGGTGGGGGAGTTGTTGGCCTCCTTCTTCCACTGCACCAAGTGCATGTTGACGAAGTCCTGGAGGCTCTTCTTCATGCCCACCTCGAAGGAGCACTCGGCCCCGTTCAGCCAGATCTTCATCCTGTCCACGTCCTCGATCCTCACGGCCATGGCGAACAGGGCCAGCTGGGTGGAGGGGTCCATGCCCGCGGCCTTGATCAGGCTGAAGAGCTTGGCCCTGTGGGTGTGGCTGGTGTAGATCCCCATGTCGCTCGGGAAGTACTCGGGCTTCAGCTTGGTGAGGTTCGCCATGGCGTTCTTCAAGCTGGAGACCATCTCCTTGCAGAAGAAGGCGGCCTGGGTGCCCAACTCCGACGCGTCCGAGATGATGTCCATCTCCCCCTTCTCCGGCTTGTAGGAGGGGAGCACCGCCTGCGGGGCCGCTATGACGGGGGCCGACAGCCTCCCCGAGGAGTCGACCTGGAACTGGAAGTCGGCCAGCCTCTTATCCATGTCCGTCGCCATCTAGTACAGCGACCTGAACTTCTTCAGGAGCCTGGCCCTCTTCTCCGCCTCCGTCTCCTTCTCCTCCTTGCTCTCCTGCACGCCCCCCTTCCCCTTCAACCCCTCCTCCTCGTCCCTGGCCTCCTTCTCCCTCTTCTCCGCCTCCCTCTTCTCCTCCTCCCTCTCCTCCTTCTCCCTCCTCTTCTCCTCCTCCTTCTTCTTCTCCCCCTCCCTCTTCCTCTCCCTGTTCGCCTCCTCCTCCTTCTTCTTCTCCTCCTCCCTCTTCCTGTTGGCCAGCTCCTGCGCCGACTTCTTGTGCTTCTCCTGCTCGGCCGCGAAGATCTCCTGCAACTTGGCGGAGTTCTCCTTGGCGAACGAGGCCCCCTGCGCCTGGCTCAGGAAGGAAAGCTTGGACAAGATGTTGTCCAACTTCTCGGAGACCTCTTCCAGCTTCCTCGCGTTGTCCTCGACCCTGCTGGCCAACACCTCGCTGCACACGACGGACGCGGTCCTCCCGCTGAAGTTCTCCCCCAGGTCGACCCCCTCCTCCACCGTCTGGGACCTCAACGACCTGGCCGTGGTCCTCTGGCCCTCCTCCTCCTCCCTGTCGTAGGTCTCCTCCTCCCTGTTGGCGGCCTCCACCTCCCTGAGCCTGGTCGCCTCCAACGCCTTGGCCAACGCCTTCTCCTCCTCCTCCTTCTTCTCCTTCTGCTTCCTCTTCTTCGCGCTCTTGCTGAGGGTGGTGGAGGACGCCTTGCTCGCCGTCTCCGACCTGTCGCTCAAGCTCGTCTTCCCCGCCACCTTGACCCCCTCCGACGACACCTCCTCCCTCTCCTCCGTCTCCTTCACCACTTTCTTCCTCGACTTCGCGCTCAACAAGATGACCACCGTGATGGCCAAGCTGATGTAGGTGTTCCACTTCGTCTCCCGGTTGGTGATGATGATAGTATGATTTAGTAGAAACGTTGTGTA

**>SalaUV-NL1-1-11_SS**

AAACACAAAGTCTTCTACTTATTATATTATCAGCATCTCGAACATGGAGACCTACTGCATCTTACTGTAGAAGACCTTGTGCGCCAGGGCGGACTACCTGGGCTTAAACTAGTGGGACCTGAATTACATGTTCCACCTCCAGTCCGTGTAGGTTACGGTGAACAAGACCCATTCCAACTCGACCTACTTCCCGAGGTCCTCCTCCTCCTGCTTGGCCATCCTGGCGCTGACCATCTCGAAGAGGGCGCCGAGGTCGTCGGTGGGGCAGATCGCCACCTCCACGTCGGAGGACCTCAGCCACTTGACCAAGCTGGGGCCCGTGTAGCTGGTCCTGACCATGTCGTCGGGCTTCATGTAGACGATCTCCATGTCCTGGACGACGGGCAAGAGGTACCTGTCCCCCTGCTTGGTCTTGTAGTACCTGGCCTGGAAGCCCCTCTCGAAGACGTTGGGGTCCGGGTTGTTCGACGTGGCGATGGTGATGTTCCAGTAGGACTGCTCCCACCAGAGGTGCGCGGTCTCGATCTCCTCGTCGAACCTGAATTGCGCCGCCCACTTGTTGGCCAGGAAGTTCCCCACCGTCCTCTCCTCGTACGGCGTGATCATCTTCCACACCAAAGCTGCCATGGGGGGGGAGGAGGAGCAGATGTTCTTCACGCAGAACTTGGTGGGGGAGTTGTTGGCCTCCTTCTTCCACTGCACCAAGTGCATGTTGACGAAGTCCTGGAGGCTCTTCTTCATGCCCACCTCGAAGGAGCACTCGGCCCCGTTCAGCCAGATCTTCATCCTGTCCACGTCCTCGATCCTCACGGCCATGGCGAACAGGGCCAGCTGGGTGGAGGGGTCCATGCCCGCGGCCTTGATCAGGCTGAAGAGCTTGGCCCTGTGGGTGTGGCTGGTGTAGATCCCCATGTCGCTCGGGAAGTACTCGGGCTTCAGCTTGGTGAGGTTCGCCATGGCGTTCTTCAAGCTGGAGACCATCTCCTTGCAGAAGAAGGCGGCCTGGGTGCCCAACTCCGACGCGTCCGAGATGATGTCCATCTCCCCCTTCTCCGGCTTGTAGGAGGGGAGCACCGCCTGCGGGGCCGCTATGACGGGGGCCGACAGCCTCCCCGAGGAGTCGACCTGGAACTGGAAGTCGGCCAGCCTCTTATCCATGTCCGTCGCCATCTAGTACAGCGACCTGAACTTCTTCAGGAGCCTGGCCCTCTTCTCCGCCTCCGTCTCCTTCTCCTCCTTGCTCTCCTGCACGCCCCCCTTCCCCTTCAACCCCTCCTCCTCGTCCCTGGCCTCCTTCTCCCTCTTCTCCGCCTCCCTCTTCTCCTCCTCCCTCTCCTCCTTCTCCCTCCTCTTCTCCTCCTCCTTCTTCTTCTCCCCCTCCGCTTTCCTCTCCCTGTTCGCCTCCTCCTCCCTCTTCTTCTCCTCCTCCCTCTTCCTGTTGGCCAGCTCCTGCGCCGACTTCTTGTGCTTCTCCTGCTCGGCCGCGAAGATCTCCTGCAACTGGGAGGTGTTCTCCTTGGCGAACGAGGCCTCCTGCGCCTGGCTCAGGAAGGAAAGCTTGGACAAGATGTTGTCCAACTTCTCGGAGACCTCTTCCAGCTTCCTCGCGTTGTCCTCGACCCTGCTGGCCAACACCTCGCTGCACACGACGGACGCGGTCCTCCCGCTGAAGTTCTCCCCCAGGTCGACCCCCTCCTCCACCGTCTGGGACCTCAACGACCTGGCCGTGGTCCTCTGGGCCCCCTCCTCCTCCCTGTCGTAGGTCTCCTCCTCCCTGTTGGTGATCTCCACCTCCCTGAGCCTGGTCGCCTCCAACGCCTTGGCCAACGCCTTCTCCTCCTCCTCCTTCTTCTCCTTCTGCTTCCTCTTCTTCGCGCTCTTGCTGAGGGTGGTGGAGGACGCCTTGCTCGCCGTCTCCGACCTGTCGCTCAAGCTCGTCTTCCCCGCCACCTTGACCCCCTCCGACGACACCTCCTCCCTCTCCTCCGTCTCCTTCACCACTTTCTTCCTCGACTTCGCGCTCAACAAGATGACCACCGTGATGGCCAAGCTGATGTAGGTGTTCCACTTCGTCTCCCGGTTGGTGATGATGATAGTATGATTTAGTAGAAACGTTGTGTA

**>SalaUV-NL1-1-12_SS**

AAACACAAAGTCTTCTACTTATTATATTATCAGCATCTCGAACATGGAGACCTACTGCATCTTACTGTAGAAGACCTTGTGCGCCAGGGCGGACTACCTGGGCTTAAACTAGTGGGACCTGAATTACATGTTCCACCTCCAGTCCGTGTAGGTTACGGTGAACAAGACCCATTCCAACTCGACCTACTTCCCGAGGTCCTCCTCCTCCTGCTTGGCCATCCTGGCGCTGACCATCTCGAAGAGGGCGCCGAGGTCGTCGGTGGGGCAGATCGCCACCTCCACGTCGGAGGACCTCAGCCACTTGACCAAGCTGGGGCCCGTGTAGCTGGTCCTGACCATGTCGTCGGGCTTCATGTAGACGATCTCCATGTCCTGGACGACGGGCAAGAGGTACCTGTCCCCCTGCTTGGTCTTGTAGTACCTGGCCTGGAAGCCCCTCTCGAAGACGTTGGGGTCCGGGTTGTTCGACGTGGCGATGGTGATGTTCCAGTAGGACTGCTCCCACCAGAGGTGCGCGGTCTCGATCTCCTCGTCGAACCTGAATTGCGCCGCCCACTTGTTGGCCAGGAAGTTCCCCACCGTCCTCTCCTCGTACGGCGTGATCATCTTCCACACCAAGGCCGCCATGGGGGGGGAGGAGGAGCAGATGTTCTTCACGCAGAACTTGGTGGGGGAGTTGTTGGCCTCCTTCTTCCACTGCACCAAGTGCATGTTGACGAAGTCCTGGAGGCTCTTCTTCATGCCCACCTCGAAGGAGCACTCGGCCCCGTTCAGCCAGATCTTCATCCTGTCCACGTCCTCGATCCTCACGGCCATGGCGAACAGGGCCAGCTGGGTGGAGGGGTCCATGCCCGCGGCCTTGATCAGGCTGAAGAGCTTGGCCCTGTGGGTGTGGCTGGTGTAGATCCCCATGTCGCTCGGGAAGTACTCGGGCTTCAGCTTGGTGAGGTTCGCCATGGCGTTCTTCAAGCTGGAGACCATCTCCTTGCAGAAGAAGGCGGCCTGGGTGCCCAACTCCGACGCGTCCGAGATGATGTCCATCTCCCCCTTCTCCGGCTTGTAGGAGGGGAGCACCGCCTGCGGGGCCGCTATGACGGGGGCCGACAGCCTCCCCGAGGAGTCGACCTGGAACTGGAAGTCGGCCAGCCTCTTATCCATGTCCGTCGCCATCTAGTACAGCGACCTGAACTTCTTCAGGAGCCTGGCCCTCTTCTCCGCCTCCGTCTCCTTCTCCTCCTTGCTCTCCTGCACGCCCCCCTTCCCCTTCAACCCCTCCTCCTCGTCCCTGGCCTCCTTCTCCCTCTTCTCCGCCTCCCTCTTCTCCTCCTCCCTCTCCTCCTTCTCCCTCCTCTTCTCCTCCTCCTTCTTCTTCTCCCCCTCCCTCTTCCTCTCCCTGTTCGCCTCCTCCTCCCTCTTCTTCTCCTCCTCCCTCTTCCTGTTGGCCAGCTCCTGCGCCGACTTCTTGTGCTTCTCCTGCTCGGCCGCGAAGATCTCCTGCAACTTGGCGGAGTTCTCCTTGGCGAACGAGGCCCCCTGCGCCTGGCTCAGGAAGGAAAGCTTGGACAAGATGTTGTCCAACTTCTCGGAGACCTCTTCCAGCTTCCTCGCGTTGTCCTCGACCCTGCTGGCCAACACCTCGCTGCACACGACGGACGCGGTCCTCCCGCTGAAGTTCTCCCCCAGGTCGACCCCCTCCTCCACCGTCTGGGACCTCAACGACCTGGCCGTGGTCCTCTGGGCCCCCTCCTCCTCCCTGTCGTAGGTCTCCTCCTCCCTGTTGGTGATCTCCACCTCCCTGAGCCTGGTCGCCTCCAACGCCTTGGCCAACGCCTTCTCCTCCTCCTCCTTCTTCTCCTTCTGCTTCCTCTTCTTCGCGCTCTTGCTGAGGGTGGTGGAGGACGCCTTGCTCGCCGTCTCCGACCTGTCGCTCAAGCTCGTCTTCCCCGCCACCTTGACCCCCTCCGACGACACCTCCTCCCTCTCCTCCGTCTCCTTCACCACTTTCTTCCTCGACTTCGCGCTCAACAAGATGACCACCGTGATGGCCAAGCTGATGTAGGTGTTCCACTTCGTCTCCCGGTTGGTGATGATGATAGTATGATTTAGTAGAAACGTTGTGTA

**>SalaUV-NL1-1-14_SS**

ATACACAAAGTCTTCTACTTATTATATTATCAGCATCTCGAACATGGAGACCTACTGCATCTTACTGTAGAAGACCTTGTGCGCCAGGGCGGACTACCTGGGCTTAAACTAGTGGGACCTGAATTACATGTTCCACCTCCAGTCCGTGTAGGTTACGGTGAACAAGACCCATTCCAACTCGACCTACTTCCCGAGGTCCTCCTCCTCCTGCTTGGCCATCCTGGCGCTGACCATCTCGAAGAGGGCGCCGAGGTCGTCGGTGGGGCAGATCGCCACCTCCACGTCGGAGGACCTCAGCCACTTGACCAAGCTGGGGCCCGTGTAGCTGGTCCTGACCATGTCGTCGGGCTTCATGTAGACGATCTCCATGTCCTGGACGACGGGCAAGAGGTACCTGTCCCCCTGCTTGGTCTTGTAGTACCTGGCCTGGAAGCCCCTCTCGAAGACGTTGGGGTCCGGGTTGTTCGACGTGGCGATGGTGATGTTCCAGTAGGACTGCTCCCACCAGAGGTGCGCGGTCTCGATCTCCTCGTCGAACCTGAATTGCGCCGCCCACTTGTTGGCCAGGAAGTTCCCCACCGTCCTCTCCTCGTACGGCGTGATCATCTTCCACACCAAGGCCGCCATGGGGGGGGAGGAGGAGCAGATGTTCTTCACGCAGAACTTGGTGGGGGAGTTGTTGGCCTCCTTCTTCCACTGCACCAAGTGCATGTTGACGAAGTCCTGGAGGCTCTTCTTCATGCCCACCTCGAAGGAGCACTCGGCCCCGTTCAGCCAGATCTTCATCCTGTCCACGTCCTCGATCCTCACGGCCATGGCGAACAGGGCCAGCTGGGTGGAGGGGTCCATGCCCGCGGCCTTGATCAGGCTGAAGAGCTTGGCCCTGTGGGTGTGGCTGGTGTAGATCCCCATGTCGCTCGGGAAGTACTCGGGCTTCAGCTTGGTGAGGTTCGCCATGGCGTTCTTCAAGCTGGAGACCATCTCCTTGCAGAAGAAGGCGGCCTGGGTGCCCAACTCCGACGCGTCCGAGATGATGTCCATCTCCCCCTTCTCCGGCTTGTAGGAGGGGAGCACCGCCTGCGGGGCCGCTATGACGGGGGCCGACAGCCTCCCCGAGGAGTCGACCTGGAACTGGAAGTCGGCCAGCCTCTTATCCATGTCCGTCGCCATCTAGTACAGCGACCTGAACTTCTTCAGGAGCCTGGCCCTCTTCTCCGCCTCCGTCTCCTTCTCCTCCTTGCTCTCCTGCACGCCCCCCTCCTCCTTCAACCCCTCCTCCTCGTCCCTGGCCTCCTTCTCCCTCTTCTCCGCCTCCCTCTTCTCCTCCTCCCTCTCCTCCTTCTCCCTCCTCTTCTCCTCCTCCTTCTTCTTCTCCCCCTCCCTCTTCCTCTCCCTGTTCGCCTCCTCCTCCCTCTTCTTCTCCTCCTCCCTCTTCCTGTTGGCCAGCTCCTGCGCCGACTTCTTGTGCTTCTCCTGCTCGGCCGCGAAGATCTCCTGCAACTGGGAGGTGTTCTCCTTGGCGAACGAGGCCCCCTGCGCCTGGCTCAGGAAGGAAAGCTTGGACAAGATGTTGTCCAACTTCTCGGAGACCTCTTCCAGCTTCCTCGCGTTGTCCTCGACCCTGCTGGCCAACACCTCGCTGCACACGACGGACGCGGTCCTCCCGCTGAAGTTCTCCCCCAGGTCGACCCCCTCCTCCACCGTCTGGGACCTCAACGACCTGGCCGTGGTCCTCTGGGCCCCCTCCTCCTCCCTGTCGTAGGTCTCCTCCTCCCTGTTGGTGATCTCCACCTCCCTGAGCCTGGTCGCCTCCAACGCCTTGGCCAACGCCTTCTCCTCCTCCTCCTTCTTCTCCTTCTGCTTCCTCTTCTTCGCGCTCTTGCTGAGGGTGGTGGAGGACGCCTTGCTCGCCGTCTCCGACCTGTCGCTCAAGCTCGTCTTCCCCGCCACCTTGACCCCCTCCGACGACACCTCCTCCCTCTCCTCCGTCTCCTTCACCACTTTCTTCCTCGACTTCGCGCTCAACAAGATGACCACCGTGATGGCCAAGCTGATGTAGGTGTTCCACTTCGTCTCCCGGTTGGTGATGATGATAGTATGATTTAGTAGAAACGTTGTGTA

**>SalaUV-NL1-1-15_SS**

AAACACAAAGTCTTCTACTTATTATATTATCAGCATCTCGAACATGGAGACCTACTGCATCTTACTGTAGAAGACCTTGTGCGCCAGGGCGGACTACCTGGGCTTAAACTAGTGGGACCTGAATTACATGTTCCACCTCCAGTCCGTGTAGGTTACGGTGAACAAGACCCATTCCAACTCGACCTACTTCCCGAGGTCCTCCTCCTCCTGCTTGGCCATCCTGGCGCTGACCATCTCGAAGAGGGCGCCGAGGTCGTCGGTGGGGCAGATCGCCACCTCCACGTCGGAGGACCTCAGCCACTTGACCAAGCTGGGGCCCGTGTAGCTGGTCCTGACCATGTCGTCGGGCTTCATGTAGACGATCTCCATGTCCTGGACGACGGGCAAGAGGTACCTGTCCCCCTGCTTGGTCTTGTAGTACCTGGCCTGGAAGCCCCTCTCGAAGACGTTGGGGTCCGGGTTGTTCGACGTGGCGATGGTGATGTTCCAGTAGGACTGCTCCCACCAGAGGTGCGCGGTCTCGATCTCCTCGTCGAACCTGAATTGCGCCGCCCACTTGTTGGCCAGGAAGTTCCCCACCGTCCGCTCCTCGTACGGCGTGATCATCTTCCACACCAAGGCCGCCATGGGGGGGGAGGAGGAGCAGATGTTCTTCACGCAGAACTTGGTGGGGGAGTTGTTGGCCTCCTTCTTCCACTGCACCAAGTGCATGTTGACGAAGTCCTGGAGGCTCTTCTTCATGCCCACCTCGAAGGAGCACTCGGCCCCGTTCAGCCAGATCTTCATCCTGTCCACGTCCTCGATCCTCACGGCCATGGCGAACAGGGCCAGCTGGGTGGAGGGGTCCATGCCCGCGGCCTTGATCAGGCTGAAGAGCTTGGCCCTGTGGGTGTGGCTGGTGTAGATCCCCATGTCGCTCGGGAAGTACTCGGGCTTCAGCTTGGTGAGGTTCGCCATGGCGTTCTTCAAGCTGGAGACCATCTCCTTGCAGAAGAAGGCGGCCTGGGTGCCCAACTCCGACGCGTCCGAGATGATGTCCATCTCCCCCTTCTCCGGCTTGTAGGAGGGGAGCACCGCCTGCGGGGACGCTATGACGGGGGCCGACAGCCTCCCCGAGGAGTCGACCTGGAACTGGAAGTCGGCCAGCCTCTTATCCATGTCCGTCGCCATCTAGTACAGCGACCTGAACTTCTTCAGGAGCCTGGCCCTCTTCTCCGCCTCCGTCTCCTTCTCCTCCTTGCTCTCCTGCACGCCCCCCTTCCCCTTCAACCCCTCCTCCTCGTCCCTGGCCTCCTTCTCCCTCTTCTCCGCCTCCCTCTTCTCCTCCTCCCTCTCCTCCTTCTCCCTCCTCTTCTCCTCCTCCTTCTTCTTCTCCCCCTCCGCTTTCCTCTCCCTGTTCGCCTCCTCCTCCCTCTTCTTCTCCTCCTCCCTCTTCCTGTTGGCCAGCTCCTGCGCCGACTTCTTGTGCTTCTCCTGCTCGGCCGCGAAGATCTCCTGCAACTTGGCGGAGTTCTCCTTGGCGAACGAGGCCCCCTGCGCCTGGCTCAGGAAGGAAAGCTTGGACAAGATGTTGTCCAACTTCTCGGAGACCTCTTCCAGCTTCCTCGCGTTGTCCTCGACCCTGCTGGCCAACACCTCGCTGCACACGACGGACGCGGTCCTCCCGCTGAAGTTCTCCCCCAGGTCGACCCCCTCCTCCACCGTCTGGGACCTCAACGACCTGGCCGTGGTCCTCTGGGCCCCCTCCTCCTCCCTGTCGTAGGTCTCCTCCTCCCTGTTGGTGATCTCCACCTCCCTGAGCCTGGTCGCCTCCAACGCCTTGGCCAACGCCTTCTCCTCCTCCTCCTTCTTCTCCTTCTGCTTCCTCTTCTTCGCGCTCTTGCTGAGGGTGGTGGAGGACGCCTTGCTCGCCGTCTCCGACCTGTCGCTCAAGCTCGTCTTCCCCGCCACCTTGACCCCCTCCGACGACACCTCCTCCCTCTCCTCCGTCTCCTTCACCACTTTCTTCCTCGACTTCGCGCTCAACAAGATGACCACCGTGATGGCCAAGCTGATGTAGGTGTTCCACTTCGTCTCCCGGTTGGTGATGATGATAGTATGATTTAGTAGAAACGTTGTGTA

**>SalaUV-NL1-1-16_SS**

AAACACAAAGTCTTCTACTTATTATATTATCAGCATCTCGAACATGGAGACCTACTGCATCTTACTGTAGAAGACCTTGTGCGCCAGGGCGGACTACCTGGGCTTAAACTAGTGGGACCTGAATTACATGTTCCACCTCCAGTCCGTGTAGGTTACGGTGAACAAGACCCATTCCAACTCGACCTACTTCCCGAGGTCCTCCTCCTCCTGCTTGGCCATCCTGGCGCTGACCATCTCGAAGAGGGCGCCGAGGTCGTCGGTGGGGCAGATCGCCACCTCCACGTCGGAGGACCTCAGCCACTTGACCAAGCTGGGGCCCGTGTAGCTGGTCCTGACCATGTCGTCGGGCTTCATGTAGACGATCTCCATGTCCTGGACGACGGGCAAGAGGTACCTGTCCCCCTGCTTGGTCTTGTAGTACCTGGCCTGGAAGCCCCTCTCGAAGACGTTGGGGTCCGGGTTGTTCGACGTGGCGATGGTGATGTTCCAGTAGGACTGCTCCCACCAGAGGTGCGCGGTCTCGATCTCCTCGTCGAACCTGAATTGCGCCGCCCACTTGTTGGCCAGGAAGTTCCCCACCGTCCTCTCCTCGTACGGCGTGATCATCTTCCACACCAAGGCCGCCATGGGGGGGGAGGAGGAGCAGATGTTCTTCACGCAGAACTTGGTGGGGGAGTTGTTGGCCTCCTTCTTCCACTGCACCAAGTGCATGTTGACGAAGTCCTGGAGGCTCTTCTTCATGCCCACCTCGAAGGAGCACTCGGCCCCGTTCAGCCAGATCTTCATCCTGTCCACGTCCTCGATCCTCACGGCCATGGCGAACAGGGCCAGCTGGGTGGAGGGGTCCATGCCCGCGGCCTTGATCAGGCTGAAGAGCTTGGCCCTGTGGGTGTGGCTGGTGTAGATCCCCATGTCGCTCGGGAAGTACTCGGGCTTCAGCTTGGTGAGGTTCGCCATGGCGTTCTTCAAGCTGGAGACCATCTCCTTGCAGAAGAAGGCGGCCTGGGTGCCCAACTCCGACGCGTCCGAGATGATGTCCATCTCCCCCTTCTCCGGCTTGTAGGAGGGGAGCACCGCCTGCGGGGCCGCTATGACGGGGGCCGACAGCCTCCCCGAGGAGTCGACCTGGAACTGGAAGTCGGCCAGCCTCTTATCCATGTCCGTCGCCATCTAGTACAGCGACCTGAACTTCTTCAGGAGCCTGGCCCTCTTCTCCGCCTCCGTCTCCTTCTCCTCCTTGCTCTCCTGCACGCCCCCCTTCCCCTTCAACCCCTCCTCCTCGTCCCTGGCCTCCTTCTCCCTCTTCTCCGCCTCCCTCTTCTCCTCCTCCCTCTCCTCCTTCTCCCTCCTCTTCTCCTCCTCCTTCTTCTTCTCCCCCTCCGCTTTCCTCTCCCTGTTCGCCTCCTCCTCCCTCTTCTTCTCCCCCTCCCTCTTCCTGTTGGCCAGCTCCTGCGCCGACTTCTTGTGCTTCTCCTGCTCGGCCGCGAAGATCTCCTGCAACTGGGAGGTGTTCTCCTTGGCGAACGAGGCCCCCTGCGCCTGGCTCAGGAAGGAAAGCTTGGACAAGATGTTGTCCAACTTCTCGGAGACCTCTTCCAGCTTCCTCGCGTTGTCCTCGACCCTGCTGGCCAACACCTCGCTGCACACGACGGACGCGGTCCTCCCGCTGAAGTTCTCCCCCAGGTCGACCCCCTCCTCCACCGTCTGGGACCTCAACGACCTGGCCGTGGTCCTCTGGGCCCCCTCCTCCTCCCTGTCGTAGGTCTCCTCCTCCCTGTTGGTGATCTCCACCTCCCTGAGCCTGGTCGCCTCCAACGCCTTGGCCAACGCCTTCTCCTCCTCCTCCTTCTTCTCCTTCTGCTTCCTCTTCTTCGCGCTCTTGCTGAGGGTGGTGGAGGACGCCTTGCTCGCCGTCTCCGACCTGTCGCTCAAGCTCGTCTTCCCCGCCACCTTGACCCCCTCCGACGACACCTCCTCCCTCTCCTCCGTCTCCTTCACCACTTTCTTCCTCGACTTCGCGCTCAACAAGATGACCACCGTGATGGCCAAGCTGATGTAGGTGTTCCACTTCGTCTCCCGGTTGGTGATGATGATAGTATGATTTAGTAGAAACGTTGTGTA

**>SalaUV-NL1-1-26_SS**

AAACACAAAGTCTTCTACTTATTATATTATCAGCATCTCGAACATGGAGAACTACTGCATCTTACCGTAGAAGACCTTGTGCGCCAGGGCGGACTACCTGGGCTTAAACTAGTGGGACCTAAATTACATGTTCCACCTCCAGTCCGTGTAGGTTACGGTGAACAAGACCCATTCCAACTCGGCCTACTTCCCGAGGTCCTCCTCCTCCTGCTTGGCCATCCTAGCGCTGACCATCTCGAAGAGGGCGCTGAGGTCGTCGGTGGGGCAGATCGCCACCTCCACGTCGGAGGACCTCAGCCACTTGACCAGGCTGGAGCCCGTGTAGCTGGTCCTGACCATGTCGTCGGGCTTCATGTAGACGATCTCCATGTCCTGGACGACGGGCAAGAGGTACCTGTCCCCCTGCTTGGTCTTGTAGTACCTGGCCTGGAAGCCCCTCTCGAAGACGTTGGGGTCCGGGTTGTTCGACGTGGCGATGGTGACGTTCCAGTAGGACTGCTCCCACCAGAGGTGCGCGGCCTCGATCTCCTCGTCGAACCTGAACTGCGCCGCCCACTTGTTGGCCAGGAAGTTCCCCACCGTCCGCTCCTCGTACGGCGTGATCATCTTCCACACCAAGGCCGCCATGGGGGGGGAGGAGGAGCAGATGTTCTTCACGCAGAACTTGGTGGGGGAGTTGTTGGCCTCCTTCTTCCACTGTACCAAGTGCATGTTGACGAAGTCCTGGAGGCTCTTCTTCATCCCCACTTCGAAGGAGCACTCGGCCCCGTTCAGCCAGATCTTCATCCTGTCCACGTCCTCGATCCTCACGGCCATGGCGAACAGGGCCAGCTGGGTGGAGGGGTCCATGCCCGCGCCCTTGATCAGGCTGAAGAGCTTCGCCCTGTGGGTGTGGCTGGTGTAGATCCCCATGTCGCTCGGGAAGTACTCGGGCTTCAGCTTGGTGAGGTTTGCCATGGCGCCCTTCAAGCCGGTGACCATCTCCTTGCAGAAGAAGGCGGCCTGGGTGCCCAACTCCGACGCGTCGGAGATGATGTCCATCTCCCCCTTCTTCGGCTTGTAGGAGGGGAGCACCGCCTGCGGGGACGCTATGACGGGGGCCGACAGCCTCCCCGAGGAGTCGACCTGGAACTCGAAGTCGGACAACCTCTTGTCCATCTCCGTCGCCATCTAGTACAGCGACCTGAACTTCTTCAGGAGCCTGGCCCTCTTCTCCGCCTCCGTCTCCTTCTCCCCCTTGCTCTCCTGCACGCCCTTCTCCTCCTTCAACCCCTCCTCCTCGTCCCTGGCCTCCTCCTCCCTCTTCTCCGCCTCCCTCTCCTCCTCCTCCTTCTCCTCCTTCTCCCTCTTCCTCTCCTCCTCCTTCTTCTTCTCCTCCTCCGCCTTCCTCTCCCTCTTCGCCTCCTCCTCCTTCTTCTTCTCCACCTCCCTCTTCCTGTTGGCCAGCTCCTGCGCTGCCTTCCTGTGCTTCTCCTGCTCGGCCGCGAAGATCTCCTGCAACTGGGAGGTGTTCTCCTTGGCGAACGAGGCCCCCTGCGCCTGGCTCAGCAAGGAAAGCTTGGACAAGATGTTGTCCAGCTTCTCGGAGACCTCCTCCAGCTTCCTTGCGTTGTCCTCGACCCTGCTGGCCAACACCTCGCTGCACACGACGGACGCGGTCCTCCCGCTGAAGTTCTCCCCCAGGTCCACCCCCTCCTCCACCGTCTGGGACCTCAACGACCTGGCCGTGGTCCTCTGGCCCCCCTCCTCCTCCCTGTCGTAGGTCTCCTCCTCCTTGTTGGCGACCTCCACCTCCTTGAGCCTGGTCGCCTCCAACGCCTTGGCCAACGCCTTCTCCTCCTCCTCCTTCTTCTCCCTCTGCTTCCTCTTCTTCGCGCTCTTGCTGAGGGTGGTGGAGGACGTCTTGCTCGCCGTCTCTGACTTGTCGCTGAAGCCCGTCTTGCCCGCCACCTTGACCTCCTCCGACAACGCCTCCTCTTTCTCCTCCGTCTCCTTCGCCACCTTCTTCCTCGACTTCGCGCTCAGCAAGATGGCCACGGTGATGGCCAAGCTGATGTAGGTGTTCCACTTCGTCTCCCTGTTGGTGATGATGATAGTATGATTTAGTAGAAACGTTGTGTA
**>SalaUV-NL1-1-27_SS**

AAACACAAAGTCTTCTACTTATTATATTATCAGCATCTCGATCATGGAGACCTACTGCATCTTACCGTAGAAGACCTTGTGCGCCAGGGCGGACTACCTGGGCTTAAACTAGTGGGACCTAAATTACATGTTCCACCTCCAGTCCGTGTAGATTACGGTGAACAAGACCCATTCCAACTCGACCTACTTCCCGAGGTCCTCCTCCTCCTGCTTGGCCATCCTGTCGCTGACCATCTCGAAGAGGGCGTCGAGGTCGTCGGTGGGGCAGAGCGCCACCTCCACGTCGGAGGACCTCAGCCACTTGACCAGGCTGGAGCCCGTGTAGCTGGTCCTGACCATGTCGTCGGGCTTCATGTAGACGATCTCCGTGTCCTTGACGACGGGCAAGAGGTACCTGTCCCCCTGCTTGGTCTTGTAGTACTTGGCCTGGAAGCCCCTCTCGAAGACGTCGGGGTCCGGGTTGTTCGACGTGGCGATGGTGACGTTCCAGTAGGACTGCTCCCACCAGAGGTGCGCGGTCTCGATCTCCTCGTCGAACCTAAACTGCGCCGCCCACTTGTTGGCCAGGAAGTTCCCCACCGTCCTCTCCTCGTACGGCGTGATCATCTTCCACACCAAAGCTGCCATGGGGGGGGAGGAGGAGCAGATATTCTTCACGCAGAACTTGGTGGGGGAGTTGTTGGCCTCCTTCTTCCACTGCACCAAGCGCATGTTGACGAAGTCCTGGAGGCTCTTCTTCATCCCCACGTCGAAGGAGCACTCGGCCCCGTTCAGCCAGATCTTCATCCTGTCCACGTCCTCGATCCTCACGGCCATGGCGAACAGGGCCAGCTGGGTGGAGGGGTCCATGCCCTCGGCCTTGATCAGGTTGAAGAGCTTCGCCCTGTGGGTGTGGCTGGTGTAGATCCCCATGTCGCTCGGGAAGTACTCGGGCTTCAGCCTGGTGAGGTTCGCCATGGCGTTCTTCAAGCTGGAGACCATCTCCTTGCAGAAGAAGGCGGCCTGGGTGCCCAACTCCGACGCGTCCGAGATGATGTCCATGTCCCCCTTCTTCGGCTTGTAGGAGGGGAGCACCGTCTGCGGGGACGCTATGACGGGGGCCGACAGCCTCCCCGAGGAGTCGACCTGGAACTGGAAGTCGGACAACCTCTTGTCCATCTCCGTCGCCATCTAGCACAGCGACCTGAACTTCTTCAGGAGCCTGGCCCTCTTCTCCGCCTCCGTCTCCTTCTCCTCCTTGCTCTCCTGCACGCCCCTCTCCTCCTTCAAGCCCTCCTCCTCGTCCCTGGCCTCCTCCTCCCTCTTCTCCGCCTCCCTCTTCTCCTCCTCCCTCTTCTCCTTCTCCCTCCTCTTCTCCTCCTCCTTCTTCTTCTCCTCCTCCGCTTTCCTCTCCCTCTTCGCCTCCTCCTCCCTCTTCTTCTCCACCTCCCTCTTCCTGTTGACCAGCTCCTGCGCCGCCTTCCTGTGCTTCTCCTGCTCGGCCGCGAAGATCTCCTGCAACTGGGAGGTGTTCTCCTTGGCGAACGAGGCCCCCTGCGCCTGGCTCAGCAAGGAAAGCTTGGACAAGATGTTGTCCAACTTCTCGGAGACCTCTTCCAGCTTCCTTGCGTTGTCCTCGACCCTGCTCGCCAACACCTCGCTGCACACGACGGACGCGGTCCTCCCGCTGAAGCTCTCCCCCAGGTCCACCCCCTCCTCCACCGTCTGGGACCTCAACGACCTGGCCGTGGTCCTCTGGCCCTCCTCCTCCTCCCTGTCGTGGGTCTCCTCCTCCTTGTTGGCGGCCTCCACCTCCTTGAGCCTGGTCGCCTCCAACGCCTTGGCCAACGCCTTCTCCTCCTCCTCCTTCTTCTCCTTCTGCTTCCTCTTCTTCGCGCTCTTGCTGAGGGTGGTGGAGGACGTCTTGCTCGCGGTATCCGACTTGTCGCTCAAGCCCGTCTTGCCCGCCACCTTGACCCCCTCCGACGACACCTCCTCTCTCTCCTCCGTCTCCTTCACCACCTTCTTCCTCGACTTCGAGCTCAACAAGATGGCCACGGTGATGGCCAAGCTGATGTAGGTGTTCCACTTCGTCTCCCGGTTGGTGATGATGATAGTATGATTTAGTAGAAACGTTGTGTA

**>SalaUV-NL1-1-9_SL_RdRp**

MKEFLQGPMHPFLMKVSYSIVMESVPYRITFHRGLSGVINSMGGDIQVTPSRIYTNMLENHSKEKVVGMLTESMKTAIEMPNSEQSFSRSPNAFELTDMETDEFEEISYKEYLELFSTPKTIRGTYMARFVITGEEFYAQKMSGMKGDKILYPFSGRVGTNTEERTRAMQKMLKVERVCELLEEVEEHMSRDKVKEVRKEVKPMESLDELVEYVDSINKKDGPEMSVEVVDVYLRSRHDLLGKAFLDSLGMTSVGCERGEMDVDLKKVNYHGRFANKTPDMLFSAPGNCLDMIDVAVTDGICSDRALSKREKYDPVMKDMVAQNICTRGTVDVLVYSTRTNEIMCPAKYKTAHVMASLESAASKIQVADIMMKGFSSYKGYRALHTQWEGEEVDEKADEKVQRMIEVCTDIFSEVESLDEMRAKKMRAEAKTFYRSEEFVKNKEDKMVADAKEDCKNFDGEAFREELVSDFKDMFTNAKYTEEIRKAVCTSQEEIVSELNKMDELSPTEEMNRAKKDYGKSFPMPIPQKLNDMKDVKMHVTDSKGGSLFDGTIFLNDCTKAGNTKKKENMAGIGANLLDRKFMENMVEKMSRDATFEFTNNMAKVYKDPQMKMFNELAVAEVTHFWTDLYRNIARLEGRRRAKRNGDSGKKVSRFTVIKEFGDYCLAVKAGSALTQEKQIRVKIFSPYKHMEGMECQKLKSKEDCKSYMDVTIARNQYTKEYSEVYQTEWLTVSVADVEHYSMLYERSAAIHSQVMADREEFESTRAQKMTREEYRRSAMVMVPILVMMCHKRGVSTSLQYNRYIMHGLTAFASDKVSMVKNIMKDPVRTMFEASIRAMQMDYAERLTMAAPDIIKQYAEKMASEFADYDRIYMPSFKVAGMLVEFAEVMNDIYMSNLFNKEKGTSGHRLVPIMTKMSEQERHYLSMSELFPGASTGDADFDALAKNKMQTHTFNKKFVMYMGYQLRMKIIHSPDLNKGLMRAMTTTIRETMSMRSSLKSCPLYEESVDYSDKLTTATAFETMCEQVEELESIFLMDMVTCTDVVNAVFKLFPKSQIGGPREILIQAAKLRLHVKFLEKMFYELCSLHEKEMLTHADVKKDLQSNATSVYKKMILEMNKHKDESETSPSGIILSFNTDASRWSPGFVMDSFIYFCLMMPLPVNLKEFCVTVIRAFATKTMFLPPTLTRRWQTDKLWNEKTDDKNINWLKENFVSSRGTAIFKSGMGQGMLHYFSSLFHCAKDDVMDSVMCFLFSKSYGDAACTTLISSDDLTKIFLVRYTKMSDLTAFMVSMLQVHDMTNRMANITTNWKKSALSILTEFNSYFSTGKRATDSVLKAVYNSQNIVDMTEPEKAVKSSISQMRNAFENGVKIPTIKVMFELQSVALSSIYAIDGERKEEIMRKLECDRNSLPFQLGYMSTDRVVEQMIAGPEVLMYKEGNSEKVMEFYVKYYAAYMTDVEMAASPDYTEEEMEEDRDYMASMNGKTKMTLPFRFDTKIKEMKKRLMNKRSITREDIDKLAERNTALSFSDRSDSASFFSNMDTYMFQMDRKFGFNDSIRVHILIRALAMAGENIALISSSRVVATDFVGVVDKILKNDDTKSKFHLTAPFKEICEMGEIAEKEAEIAIPSHSYSHNKSRKINFTPMKDRVKVSDMEVFRYMMRATSNASNSAMTEANKISESLGVKLEKFIEYPIYQIKKAFPGVRRPLTMLKSYIMYWLKMTSDLNMEMMVSFPDNSEALTNMRMLYLEKYSNRGILQMFVPKKSLRVVSEERFNSALSMRGRVDRFEVEEFRGDLTEGPMENLVRDPWGIDNPLLTVIPSDSLEVRFAKMAVSATDEDKYSFQGAKSLKVFYKVDVNGKMKTRYYTDLVHLGIIHEDRQASPHTVSVRMYKVSSSDVELGNITRLMHKDLSTIQIRPWAPRTDNLGVEKNAKIGNLWIKVYRIWPKTELILTRRMTKWTVLAMFNERELGGKVFFHIGTNSYKMPFNEDSENPNIRKFSELVVSSKLDLSDVEEILNDLGIADNSMFAPDNRIKEAEFEAEVDSTIYEMMDVSQAVEDYSVENVMGAISATDFGISSSSMATKLNAEMFSSIGKEEMGGMSMDFTSLAGLAGAFSGLKEIDTTAHMEEEPLEEGMSLREYMGTVYTSVKRVFEKFYSFDQRLVMNTVNDIQRNFIGDEKKRKLAVVCNMVYTAVYMKMTGTINPTTGMSGLAQMDALRDFKENKGYNSASDTLVVMVQMVTTGLVTEIFSSIGEKAPTLNSLIKLDIAFNTKRAVALDRMVTKVQDKRRSLLSIL

**>SalaUV-NL1-1-11_SL_RdRp**

MKEFLQGPMHPFLMKVSYSIVMESVPYRITFHRGLSGVINSMGGDIQVTPSRIYTNMLENHSKEKVVGMLTESMKTAIEMPNSEQSFSRSPNAFELTDMETDEFEEISYKEYLELFSTPKTIRGTYMARFVITGEEFYAQKMSGMKGDKILYPFSGRVGTNTEERTRAMQKMLKVERVCELLEEVEEHMSRDKVKEVRKEVKPMESLDELVEYVDSINKKDGPEMSVEVVDVYLRSRHDLLGKAFLDSLGMTSVGCERGEMDVDLKKVGYHGRFANKTPDMLFSAPGNCLDMIDVAVTDGICSDRALSKREKYDPVMKDMVAQNICTRGTVDVLVYSTRTNEIMCPAKYKTAHVMASLESAASKIQVADIMMKGFSSYKGYRALHTQWEGEEVDEKADEKVQRMIEVCTDIFSEVESLDEMRAKKMRAEAKTFYRSEEFVKNKEDKMVADAKEDCKNFDGEAFREELVSDFKDMFTNAKYTEEIRKAVCTSQEEIVSELNKMDELSPTEEMNRAKKDYGKSFPMPIPQKLNDMKDVKMHVTDSKGGSLFDGTIFLNDCTKAGNTKKKENMAGIGANLLDRKFMENMVEKMSRDATFEFTNNMAKVYKDPQMKMFNELAVAEVTHFWTDLYRNIARLEGRRRAKRNGDSGKKVSRFTVIKEFGDYCLAVKAGSALTQEKQIRVKIFSPYKHMEGMECQKLKSKEDCKSYMDVTIARNQYTKEYSEVYQTEWLTVSVADVEHYSMLYERSAAIHSQVMADREEFESTRAQKMTREEYRRSAMVMVPILVMMCHKRGVSTSLQYNRYIMHGLTAFASDKVSMVKNIMKDPVRTMFEASIRAMQMDYAERLTMAAPDIIKQYAEKMASEFADYDRIYMPSFKVAGMLVEFAEVMNDIYMSNLFNKEKGTSGHRLVPIMTKMSEQERHYLSMSELFPGASTGDADFDALAKNKMQTHTFNKKFVMYMGYQLRMKIIHSPDLNKGLMRAMTTTIRETMSMRSSLKSCPLYEESVDYSDKLTTATAFETMCEQVEELESIFLMDMVTCTDVVNAVFKLFPKSQIGGPREILIQAAKLRLHVKFLEKMFYELCSLHEKEMLTHADVKKDLQSNATSVYKKMILEMNKHKDESETSPSGIILSFNTDASRWSPGFVMDSFIYFCLMMPLPVNLKEFCVTVIRAFATKTMFLPPTLTRRWQTDKLWNEKTDDKNINWLKENFVSSRGTAIFKSGMGQGMLHYFSSLFHCAKDDVMDSVMCFLFSKSYGDAACTTLISSDDLTKIFLVRYTKMSDLTAFMVSMLQVHDMTNRMANITTNWKKSALSILTEFNSYFSTGKRATDSVLKAVYNSQNIVDMTEPEKAVKSSISQMRNAFENGVKIPTIKVMFELQSVALSSIYAIDGERKEEIMRKLECDRNSLPFQLGYMSTDRVVEQMIAGPEVLMYKEGNSEKVMEFYVKYYAAYMTDVEMAASPDYTEEEMEEDRDYMASMNGKTKMTLPFRFDTKIKEMKKRLMNKRSITREDIDKLAERNTALSFSDRSDSASFFSNMDTYMFQMDRKFGFNDSIRVHILIRALAMAGENIALISSSRVVATDFVGVVDKILKNDDTKSKFHLTAPFKEICEMGEIAEKEAEIAIPSHSYSHNKSRKINFTPMKDRVKVSDMEVFRYMMRATSNASNSAMTEANKISESLGVKLEKFIEYPIYQIKKAFPGVRRPLTMLKSYIMYWLKMTSDLNMEMMVSFPDNSEALTNMRMLYLEKYSNRGILQMFVPKKSLRVVSEERFNSALSMRGRVDRFEVEEFRGDLTEGPMENLVRDPWGIDNPLLTVIPSDSLEVRFAKMAVSATDEDKYSFQGAKSLKVFYKVDVNGKMKTRYYTDLVHLGIIHEDRGASPHTVSVRMYKVSSSDVELGNITRLMHKDLSTIQIRPWAPRTDNLGVEKNAKIGNLWIKVYRIWPKTELILTRRMTKWTVLAMFNERELGGKVFFHIGTNSYKMPFNEDSENPNIRKFSELVVSSKLDLSDVEEILNDLGIADNSMFAPDNRIKEAEFEAEVDSTIYEMMDVSQAVEDYSVENVMGAISATDFGISSSSMATKLNAEMFSSIGKEEMGGMSMDFTSLAGLAGAFSGLKEIDTTAHMEEEPLEEGMSLREYMGTVYTSVKRVFEKFYSFDQRLVMNTVNDIQRNFIGDEKKRKLAVVCNMVYTAVYMKMTGTINPTTGMSGLAQMDALRDFKENKGYNSASDTLVVMVQMVTTGLVTEIFSSIGEKTPTLNSLIKLDIAFNTKRAVALDRMVTKVQDKRRSLLSIL

**>SalaUV-NL1-1-12_SL_RdRp**

MKEFLQGPMHPFLMKVSYSIVMESVPYRITFHRGLSGVINSMGGDIQVTPSRIYTNMLENHSKEKVVGMLTESMKTAIEMPNSEQSFSRSPNAFELTDMETDEFEEISYKEYLELFSTPKTIRGTYMARFVITGEEFYAQKMSGMKGDKILYPFSGRVGTNTEERTRAMQKMLKVERVCELLEEVEEHMSRDKVKEVRKEVKPMESLDELVEYVDSINKKDGPEMSVEVVDVYLRSRHDLLGKAFLDSLGMTSVGCERGEMDVDLKKVNYHGRFANKTPDMLFSAPGNCLDMIDVAVTDGICSDRALSKREKYDPVMKDMVAQNICTRGTVDVLVYSTRTNEIMCPAKYKTAHVMASLESAASKIQVADIMMKGFSSYKGYRALHTQWEGEEVDEKADEKVQRMIEVCTDIFSEVESLDEMRAKKMRAEAKTFYRSEEFVKNKEDKMVADAKEDCKNFDGEAFREELVSDFKDMFTNAKYTEEIRKAVCTSQEEIVSELNKMDELSPTEEMNRAKKDYGKSFPMPIPQKLNDMKDVKMHVTDSKGGSLFDGTIFLNDCTKAGNTKKKENMAGIGANLLDRKFMENMVEKMSRDATFEFTNNMAKVYKDPQMKMFNELAVAEVTHFWTDLYRNIARLEGRRRAKRNGDSGKKVSRFTVIKEFGDYCLAVKAGSALTQEKQIRVKIFSPYKHMEGMECQKLKSKEDCKSYMDVTIARNQYTKEYSEVYQTEWLTVSVADVEHYSMLYERSAAIHSQVMADREEFESTRAQKMTREEYRRSAMVMVPILVMMCHKRGVSTSLQYNRYIMHGLTAFASDKVSMVKNIMKDPVRTMFEASIRAMQMDYAERLTMAAPDIIKQYAEKMASEFADYDRIYMPSFKVAGMLVEFAEVMNDIYMSNLFNKEKGTSGHRLVPIMTKMSEQERHYLSMSELFPGASTGDADFDALAKNKMQTHTFNKKFVMYMGYQLRMKIIHSPDLNKGLMRAMTTTIRETMSMRSSLKSCPLYEESVDYSDKLTTATAFETMCEQVEELESIFLMDMVTCTDVVNAVFKLFPKSQIGGPREILIQAAKLRLHVKFLEKMFYELCSLHEKEMLTHADVKKDLQSNATSVYKKMILEMNKHKDESETSPSGIILSFNTDASRWSPGFVMDSFIYFCLMMPLPVKLKEFCVTVIRAFATKTMFLPPTLTRRWQTDKLWNEKTDDKNINWLKENFVSSRGTAIFKSGMGQGMLHYFSSLFHCAKDDVMDSVMCFLFSKSYGDAACTTLISSDDLTKIFLVRYTKMSDLTAFMVSMLQVHDMTNRMANITTNWKKSALSILTEFNSYFSTGKRATDSVLKAVYNSQNIVDMTEPEKAVKSSISQMRNAFENGVKIPTIKVMFELQSVALSSIYAIDGERKEEIMRKLECDRNSLPFQLGYMSTDRVVEQMIAGPEVLMYKEGNSEKVMEFYVKYYAAYMTDVEMAASPDYTEEEMEEDRDYMASMNGKTKMTLPFRFDTKIKEMKKRLMNKRSITREDIDKLAERNTALSFSDRSDSASFFSNMDTYMFQMDRKFGFNDSIRVHILIRALAMAGENIALISSSRVVATDFVGVVDKILKNDDTKSKFHLTAPFKEICEMGEIAEKEAEIAIPSHSYSHNKSRKINFTPMKDRVKVSDMEVFRYMMRATSNASNSAMTEANKISESLGVKLEKFIEYPIYQIKKAFPGVRRPLTMLKSYIMYWLKMTSDLNMEMMVSFPDNSEALTNMRMLYLEKYSNRGILQMFVPKKSLRVVSEERFNSALSMRGRVDRFEVEEFRGDLTEGPMENLVRDPWGIDNPLLTVIPSDSLEVRFAKMAVSATDEDKYSFQGAKSLKVFYKVDVNGKMKTRYYTDLVHLGIIHEDRQASPHTVSVRMYKVSSSDVELGNITRLMHKDLSTIQIRPWAPRTDNLGVEKNAKIGNLWIKVYRIWPKTELILTRRMTKWTVLAMFNERELGGKVFFHIGTNSYKMPFNEDSENPNIRKFSELVVSSKLDLSDVEEILNDLGIADNSMFAPDNRIKEAEFEAEVDSTIYEMMDVSQAVEDYSVENVMGAISATDFGISSSSMATKLNAEMFSSIGKEEMGGMSMDFTSLAGLAGAFSGLKEIDTTAHMEEEPLEEGMSLREYMGTVYTSVKRVFEKFYSFDQRLVMNTVNDIQRNFIGDEKKRKLAVVCNMVYTAVYMKMTGTINPTTGMSGLAQMDALRDFKENKGYNSASDTLVVMVQMVTTGLVTEIFSSIGEKTPTLNSLIKLDIAFNTKRAVALDRMVTKVQDKRRSLLSIL
**>SalaUV-NL1-1-14_SL_RdRp**

MKEFLQGPMHPFLMKVSYSVVMESVPYRITFHRGLSGVINSMGGDIQVTPSRIYTNMLENHSKEKVVGMLTESMKTAIEMPNSEQSFSRSPNAFELTDMETDEFEEISYKEYLELFSTPKTIRGTYMARFVITGEEFYAQKMSGMKGDKILYPFSGRVGTNTEERTRAMQKMLKVERVCELLEEVEEHMSRDKVKEVRKEVKPMESLDELVEYVDSINKKDGPEMSVEVVDVYLRSRHDLLGKAFLDSLGMTSVGCERGEMDVDLKKVNYHGRFANKTPDMLFSAPGNCLDMIDVAVTDGICSDRALSKREKYDPVMKDMVAQNICTRGTVDVLVYSTRTNEIMCPAKYKTAHVMASLESAASKIQVADIMMKGFSSYKGYRALHTQWEGEEVDEKADEKVQRMIEVCTDIFSEVESLDEMRAKKMRAEAKTFYRSEEFVKNKEDKMVADAKEDCKNFDGEAFREELVSDFKDMFTNAKYTEEIRKAVCTSQEEIVSELNKMDELSPTEEMNRAKKDYGKSFPMPIPQKLNDMKDVKMHVTDSKGGSLFDGTIFLNDCTKAGNTKKKENMAGIGANLLDRKFMENMVEKMSRDATFEFTNNMAKVYKDPQMKMFNELAVAEVTHFWTDLYRNIARLEGRRRAKRNGDSGKKVSRFTVIKEFGDYCLAVKAGSALTQEKQIRVKIFSPYKHMEGMECQKLKSKEDCKSYMDVTIARNQYTKEYSEVYQTEWLTVSVADVEHYSMLYERSAAIHSQVMADREEFESTRAQKMTREEYRRSAMVMVPILVMMCHKRGVSTSLQYNRYIMHGLTAFASDKVSMVKNIMKDPVRTMFEASIRAMQMDYAERLTMAAPDIIKQYAEKMASEFADYDRIYMPSFKVAGMLVEFAEVMNDIYMSNLFNKEKGTSGHRLVPIMTKMSEQERHYLSMSELFPGASTGDADFDALAKNKMQTHTFNKKFVMYMGYQLRMKIIHSPDLNKGLMRAMTTTIRETMSMRSSLKSCPLYEESVDYSDKLTTATAFETMCEQVEELESIFLMDMVTCTDVVNAVFKLFPKSQIGGPREILIQAAKLRLHVKFLEKMFYELCSLHEKEMLTHADVKKDLQSNATSVYKKMILEMNKHKDESETSPSGIILSFNTDASRWSPGFVMDSFIYFCLMMPLPVNLKEFCVTVIRAFATKTMFLPPTLTRRWQTDKLWNEKTDDKNINWLKENFVSSRGTAIFKSGMGQGMLHYFSSLFHCAKDDVMDSVMCFLFSKSYGDAACTTLISSDDLTKIFLVRYTKMSDLTAFMVSMLQVHDMTNRMANITTNWKKSALSILTEFNSYFSTGKRATDSVLKAVYNSQNIVDMTEPEKAVKSSISQMRNAFENGVKIPTIKVMFELQSVALSSIYAIDGERKEEIMRKLECDRNSLPFQLGYMSTDRVVEQMIAGPEVLMYKEGNSEKVMEFYVKYYAAYMTDVEMAASPDYTEEEMEEDRDYMASMNGKTKMTLPFRFDTKIKEMKKRLMNKRSITREDIDKLAERNTALSFSDRSDSASFFSNMDTYMFQMDRKFGFNDSIRVHILIRALAMAGENIALISSSRVVATDFVGVVDKILKNDDTKSKFHLTAPFKEICEMGEIAEKEAEIAIPSHSYSHNKSRKINFTPMKDRVKVSDMEVFRYMMRATSNASNSAMTEANKISESLGVKLEKFIEYPIYQIKKAFPGVRRPLTMLKSYIMYWLKMTSDLNMEMMVSFPDNSEALTNMRMLYLEKYSNRGILQMFVPKKSLRVVSEERFNSALSMRGRVDRFEVEEFRGDLTEGPMENLVRDPWGIDNPLLTVIPSDSLEVRFAKMAVSATDEDKYSFQGAKSLKVFYKVDVNGKMKTRYYTDLVHLGIIHEDRQASPHTVSVRMYKVSSSDVELGNITRLMHKDLSTIQIRPWAPRTDNLGVEKNAKIGNLWIKVYRIWPKTELILTRRMTKWTVLAMFNERELGGKVFFHIGTNSYKMPFNEDSENPNIRKFSELVVSSKLDLSDVEEILNDLGIADNSMFAPDNRIKEAEFEAEVDSTIYEMMDVSQAVEDYSVENVMGAISATDFGISSSSMATKLNAEMFSSIGKEEMGGMSMDFTSLAGLAGAFSGLKEIDTTAHMEEEPLEEGMSLREYMGTVYTSVKRVFEKFYSFDQRLVMNTVNDIQRNFIGDEKKRKLAVVCNMVYTAVYMKMTGTINPTTGMSGLAQMDALRDFKENKGYNSASDTLVVMVQMVTTGLVTEIFSSIGEKTPTLNSLIKLDIAFNTKRAVALDRMVTKVQDKRRSLLSIL

**>SalaUV-NL1-1-15_SL_RdRp**

MKEFLQGPMHPFLMKVSYSIVMESVPYRITFHRGLSGVINSMGGDIQVTPSRIYTNMLENHSKEKVVGMLTESMKTAIEMPNSEQSFSRSPNAFELTDMETDEFEEISYKEYLELFSTPKTIRGTYMARFVITGEEFYAQKMSGMKGDKILYPFSGRVGTNTEERTRAMQKMLKVERVCELLEEVEEHMSRDKVKEVRKEVKPMESLDELVEYVDSINKKDGPEMSVEVVDVYLRSRHDLLGKAFLDSLGMTSVGCERGEMDVDLKKVNYHGRFANKTPDMLFSAPGNCLDMIDVAVTDGICSDRALSKREKYDPVMKDMVAQNICTRGTVDVLVYSTRTNEIMCPAKYKTAHVMASLESAASKIQVADIMMKGFSSYKGYRALHTQWEGEEVDEKADEKVQRMIEVCTDIFSEVESLDEMRAKKMRAEAKTFYRSEEFVKNKEDKMVADAKEDCKNFDGEAFREELVSDFKDMFTNAKYTEEIRKAVCTSQEEIVSELNKMDELSPTEEMNRAKKDYGKSFPMPIPQKLNDMKDVKMHVTDSKGGSLFDGTIFLNDCTKAGNTKKKENMAGIGANLLDRKFMENMVEKMSRDATFEFTNNMAKVYKDPQMKMFNELAVAEVTHFWTDLYRNIARLEGRRRAKRNGDSGKKVSRFTVIKEFGDYCLAVKAGSALTQEKQIRVKIFSPYKHMEGMECQKLKSKEDCKSYMDVTIARNQYTKEYSEVYQTEWLTVSVADVEHYSMLYERSAAIHSQVMADREEFESTRAQKMTREEYRRSAMVMVPILVMMCHKRGVSTSLQYNRYIMHGLTAFASDKVSMVKNIMKDPVRTMFEASIRAMQMDYAERLTMAAPDIIKQYAEKMASEFADYDRIYMPSFKVAGMLVEFAEVMNDIYMSNLFNKEKGTSGHRLVPIMTKMSEQERHYLSMSELFPGASTGDADFDALAKNKMQTHTFNKKFVMYMGYQLRMKIIHSPDLNKGLMRAMTTTIRETMSMRSSLKSCPLYEESVDYSDKLTTATAFETMCEQVEELESIFLMDMVTCTDVVNAVFKLFPKSQIGGPREILIQAAKLRLHVKFLEKMFYELCSLHEKEMLTHADVKKDLQSNTTSVYKKMILEMNKHKDESETSPSGIILSFNTDASRWSPGFVMDSFIYFCLMMPLPVNLKEFCVTVIRAFATKTMFLPPTLTRRWQTDKLWNEKTDDKNINWLKENFVSSRGTAIFKSGMGQGMLHYFSSLFHCAKDDVMDSVMCFLFSKSYGDAACTTLISSDDLTKIFLVRYTKMSDLTAFMVSMLQVHDMTNRMANITTNWKKSALSILTEFNSYFSTGKRATDSVLKAVYNSQNIVDMTEPEKAVKSSISQMRNAFENGVKIPTIKVMFELQSVALSSIYAIDGERKEEIMRKLECDRNSLPFQLGYMSTDRVVEQMIAGPEVLMYKEGNSEKVMEFYVKYYAAYMTDVEMAASPDYTEEEMEEDRDYMASMNGKTKMTLPFRFDTKIKEMKKRLMNKRSITREDIDKLAERNTALSFSDRSDSASFFSNMDTYMFQMDRKFGFNDSIRVHILIRALAMAGENIALISSSRVVATDFVGVVDKILKNDDTKSKFHLTAPFKEICEMGEIAEKEAEIAIPSHSYSHNKSRKINFTPMKDRVKVSDMEVFRYMMRATSNASNSAMTEANKISESLGVKLEKFIEYPIYQIKKAFPGVRRPLTMLKSYIMYWLKMTSDLNMEMMVSFPDNSEALTNMRMLYLEKYSNRGILQMFVPKKSLRVVSEERFNSALSMRGRVDRFEVEEFRGDLTEGPMENLVRDPWGIDNPLLTVIPSDSLEVRFAKMAVSATDEDKYSFQGAKSLKVFYKVDVNGKMKTRYYTDLVHLGIIHEDRQASPHTVSVRMYKVSSSDVELGNITRLMHKDLSTIQIRPWAPRTDNLGVEKNAKIGNLWIKVYRIWPKTELILTRRMTKWTVLAMFNERELGGKVFFHIGTNSYKMPFNEDSENPNIRKFSELVVSSKLDLSDVEEILNDLGIADNSMFAPDNRIKEAEFEAEVDSTIYEMMDVSQAVEDYSVENVMGAISATDFGISSSSMATKLNAEMFSSIGKEEMGGMSMDFTSLAGLAGAFSGLKEIDTTAHMEEEPLEEGMSLREYMGTVYTSVKRVFEKFYSFDQRLVMNTVNDIQRNFIGDEKKRKLAVVCNMVYTAVYMKMTGTINPTTGMSGLAQMDALRDFKENKGYNSASDTLVVMVQMVTTGLVTEIFSSIGEKTPTLNSLIKLDIAFNTKRAVALDRMVTKVQDKRRSLLSIL

**>SalaUV-NL1-1-16_SL_RdRp**

MKEFLQGPMHPFLMKVSYSIVMESVPYRITFHRGLSGVINSMGGDIQVTPSRIYTNMLENHSKEKVVGMLTESMKTAIEMPNSEQSFSRSPNAFELTDMETDEFEEISYKEYLELFSTPKTIRGTYMARFVITGEEFYAQKMSGMKGDKILYPFSGRVGTNTEERTRAMQKMLKVERVCELLEEVEEHMSRDKVKEVRKEVKPMESLDELVEYVDSINKKDGPEMSVEVVDVYLRSRHDLLGKAFLDSLGMTSVGCERGEMDVDLKKANYHGRFANKTPDMLFSAPGNCLDMIDVAVTDGICSDRALSKREKYDPVMKDMVAQNICTRGTVDVLVYSTRTNEIMCPAKYKTAHVMASLESAASKIQVADIMMKGFSSYKGYRALHTQWEGEEVDEKADEKVQRMIEVCTDIFSEVESLDEMRAKKMRAEAKTFYRSEEFVKNKEDKMVADAKEDCKNFDGEAFREELVSDFKDMFTNAKYTEEIRKAVCTSQEEIVSELNKMDELSPTEEMNRAKKDYGKSFPMPIPQKLNDMKDVKMHVTDSKGGSLFDGTIFLNDCTKAGNTKKKENMAGIGANLLDRKFMENMVEKMSRDATFEFTNNMAKVYKDPQMKMFNELAVAEVTHFWTDLYRNIARLEGRRRAKRNGDSGKKVSRFTVIKEFGDYCLAVKAGSALTQEKQIRVKIFSPYKHMEGMECQKLKSKEDCKSYMDVTIARNQYTKEYSEVYQTEWLTVSVADVEHYSMLYERSAAIHSQVMADREEFESTRAQKMTREEYRRSAMVMVPILVMMCHKRGVSTSLQYNRYIMHGLTAFASDKVSMVKNIMKDPVRTMFEASIRAMQMDYAERLTMAAPDIIKQYAEKMASEFADYDRIYMPSFKVAGMLVEFAEVMNDIYMSNLFNKEKGTSGHRLVPIMTKMSEQERHYLSMSELFPGASTGDADFDALAKNKMQTHTFNKKFVMYMGYQLRMKIIHSPDLNKGLMRAMTTTIRETMSMRSSLKSCPLYEESVDYSDKLTTATAFETMCEQVEELESIFLMDMVTCTDVVNAVFKLFPKSQIGGPREILIQAAKLRLHVKFLEKMFYELCSLHEKEMLTHADVKKDLQSNATSVYKKMILEMNKHKDESETSPSGIILSFNTDASRWSPGFVMDSFIYFCLMMPLPVNLKEFCVTVIRAFATKTMFLPPTLTRRWQTDKLWNEKTDDKNINWLKENFVSSRGTAIFKSGMGQGMLHYFSSLFHCAKDDVMDSVMCFLFSKSYGDAACTTLISSDDLTKIFLVRYTKMSDLTAFMVSMLQVHDMTNRMANITTNWKKSALSILTEFNSYFSTGKRATDSVLKAVYNSQNIVDMTEPEKAVKSSISQMRNAFENGVKIPTIKVMFELQSVALSSIYAIDGERKEEIMRKLECDRNSLPFQLGYMSTDRVVEQMIAGPEVLMYKEGNSEKVMEFYVKYYAAYMTDVEMAASPDYTEEEMEEDRDYMASMNGKTKMTLPFRFDTKIKEMKKRLMNKRSITREDIDKLAERNTALSFSDRSDSASFFSNMDTYMFQMDRKFGFNDSIRVHILIRALAMAGENIALISSSRVVATDFVGVVDKILKNDDTKSKFHLTAPFKEICEMGEIAEKEAEIAIPSHSYSHNKSRKINFTPMKDRVKVSDMEVFRYMMRATSNASNSAMTEANKISESLGVKLEKFIEYPIYQIKKAFPGVRRPLTMLKSYIMYWLKMTSDLNMEMMVSFPDNSEALTNMRMLYLEKYSNRGILQMFVPKKSLRVVSEERFNSALSMRGRVDRFEVEEFRGDLTEGPMENLVRDPWGIDNPLLTVIPSDSLEVRFAKMAVSATDEDKYSFQGAKSLKVFYKVDVNGKMKTRYYTDLVHLGIIHEDRQASPHTVSVRMYKVSSSDVELGNITRLMHKDLSTIQIRPWAPRTDNLGVEKNAKIGNLWIKVYRIWPKTELILTRRMTKWTVLAMFNERELGGKVFFHIGTNSYKMPFNEDSENPNIRKFSELVVSSKLDLSDVEEILNDLGIADNSMFAPDNRIKEAEFEAEVDSTIYEMMDVSQAVEDYSVENVMGAISATDFGISSSSMATKLNAEMFSSIGKEEMGGMSMDFTSLAGLAGAFSGLKEIDTTAHMEEEPLEEGMSLREYMGTVYTSVKRVFEKFYSFDQRLVMNTVNDIQRNFIGDEKKRKLAVVCNMVYTAVYMKMTGTINPTTGMSGLAQMDALRDFKENKGYNSASDTLVVMVQMVTTGLVTEIFSSIGEKTPTLNSLIKLDIAFNTKRAVALDRMVTKVQDKRRSLLSIL

**>SalaUV-NL1-1-26_SL_RdRp**

MKEFLQGPMHPFLMKISYSVVMESVPYRVTFHRGLSGVINSMGGDIQVTPSRIYTNMLENHSKEKVVKMLTESMKTAIEMPNSEQSFSRSPNAFELTDMETDEFEEISYKEYLELFSTPKTIRGTYMARFVITGEEFYAQKMSGMKGDKILYPFSGRVGTNTEERARAMQKMLKVERVCELLEEVEEHMSRDKVKEVRKEVKPMESLDELVEYVDSINKKDGPEMSVEVVDVYLRSRHDLLGKAFLDSLGMTSVGCERGEMDVDLKKANYHGKFANKTPDILFSAPGNCLDMIDVAVTDGICSDRALSKKEKYDPVMKDMVAQNICTRGTVDVLVYSTRTNEIMCPEKYKTAHVMASLESAASKIQVADIMMRGFNSYKGYRALHTQWEGEEVDEKADEKVQRMIEVCTDIFSEVESLDEMRAKKMRAEAKTFYRNEEFVKNKEDKMVADAKEDCKNFDGEAFREELVSDFKDMFTNAKYTEEIKKAVCTSQEEIVSELNKMDELSPTEEMNRAKKDYGKSFPMPIPQKLNDMKDVKMHVTDSKGGSLFDGTIFLNDCTKAGNTKKKENMAGIGANLLDRKFMENMVEKMSRDATFEFTNNMAKVYKDPQMKMFNELAVAEVTHFWTDLYRNIARLEGRRRAKRNGDSGKKVSRFTVIKEFGDYCLAVKAGSALTQDKQIRVKIFSPYKHMEGMECQKLKSKEDCKSYMDVTIARDKYTKEYSEVYQTEWLTVSVADVEHYSMLYERSAAIHSQVMADREEFESTRAQKMTREEYRRSAMVMVPILVMMCHKRGVSTSLQYNRYIMHGLTAFASDKVSMVKNIMKDPVRTMFEASIRAMQMDYAERLTMAAPDIIKQYAEKMASEFADYDRIYMPSFKVAGMLVEFAEVMNDIYMSNLFNKEKGTAGHRLVPIMTKMSEQERHYLSMSELFPGASTGDADFDALAKNKMQTHTFNKKFVMYMGYQLRMKIIHSPDLNKGLMRAMTTTIRETMSMRSSLKSCPLYEESVDYSDKLTTATAFETMCEQVEELESIFLMDMVTCTDVVNAVFKLFPKSQIGGPREILIQAAKLRLHVKFLEKMFYELCSLHEKEMLTHADVKKDLQSNATSVYKKMILEMNKHKDESETSPSGIILSFNTDASRWSPGFVMDSFIYFCLMMPLPVKLKEFCVTVIRAFATKTMFLPPTLTRRWQTDKLWNEKTDDKNINWLKENFVSSRGTAIFKSGMGQGMLHYFSSLFHCAKDDVMDSVMCFLFSKSYGDAACTTLISSDDLTKIFLVRYTKMSDLTAFMVSMLQVHDMTNRMANITTNWKKSALSILTEFNSYFSTGKRATDSVLKAVYNSQNIVDMTEPEKAVKSSISQMRNAFENGVKIPTIKVMFELQSVALSSIYAIDGERKEEIMRKLDCDRNSLPFQLGYMSTDRVVEQMIAGPEVLMYKEGNSEKVMEFYVKYYAAYMTDVEMAASLDYTEEEMEEDRDYMASMNGKTKMTLPFRFDTKIKEMKKRLMHKRSITREDIDKLAERNTALSFSDRSDSASFFSNMDTYMFQMDRKFGFNDAIRVHILIRALAMAGEDIALISSSRVVATDFVGVVDKILKNDDTKSKFHLTAPFKEICEMGEIAEKEAEIAIPSHSYSHNKSRRINFTPMKDRVKVSDMEVFRYMMRATSNASNSAMTEANKISESLGVKLEKFIEYPIYQIKKAFPGVRRPLTMLKSYIMYWLKMTSDLNMEMMVSFPDNSEALTNMRMLYLEKYSNSGILQMFVPKKSLRVVSEERFNSALSMRGRVDRFEVEEFRGDLTEGPMENLVRDPWGTDNPLLTVIPSDSLEVRFAKMAVSATDEDKYSFQGAKSFKVFYKVDVNGKMKTRYYTDLVHLGIIHEDRGASPHTVSVRMYKVSSSDVELGNITRLMHKDLSTIQIRPWAPRTDNLEVAKNAKIGNLWIKVYRIWPKTELILTRRMTKWTVLAMFNERELGGKVFFHIGTNSYKMPFNEDSENSNIRKFSELVVSSKLDLSDVEEILNDLGIADNSMFAPDNRVKEAEFEAEVDSTIYEMMDVSQAVEDYSVENVMGAISATDFGISSSSMATKLNAEMFSSIGKEEMGGMAVDFTSLAGLAGAFSGLKEIDTTAHMEEEPLEEGMSLREYMGTVYTSVKRVFEKFYSFDQRLVMNTVNDIRRNFIGDEKKRKLAVVCNMVYTAVYMKMTGTVNPTTGMSGLAQMDALRDFKENKGYNSASDTLVVMVQMVTTGLITEIFSSIGEKAPTLNSLIKLDIAFNTKRAVALDRMVTKVQDKGRSLLSIL

**>SalaUV-NL1-1-27_SL_RdRp**

MKEFLQGPMHPFLMKVSYSVVMESVPYRVTFHRGLSGVINSMGGDIQVTPSRIYTNMLENHSKEKVVRMLTESMKTAIEMPNSEQSFSRSPNAFELTEMETDEFEEISYKEYLELFSTPTTIRGTYMARFVITGEEFYAQKMSGMKGDKILYPFSGRVGTNTEERARAMQKMLKVEKVCELLEEVEEHMSRDKVKEVRKEVKPMESLDELVEYVDSINKKDGPEMSVEVVDVYLRSRHDLLGKAFLDSLGMTSVGCERGEMDVDLKKANYHGKFANKTPDILFSAPGNCLDMIDVAVTDGICSDRALSKREKYDPVMKDMVAQNICTRGTVDVLVYSTKTNEIMCPGKYKTAHVMASLESAASKIQVADIMMRGFNSYKGYRALHTQWEGEEVDEKADEKVQRMIEVCTDIFSEVESLDEMRAKKMRAEAKTFYRSEEFVKNKEDKMVADAKEDCKNFDGEAFREELVSDFKDMFTNAEYTEEIKKAVCTSQEEIVSELNKMDELSPTEEMNRAKKDYGKSFPMPIPQKLNDMKDVKMHVTDSKGGSLFDGTIFLDDCTKAGNTKKKENMAGIGANLLDRKFMENMVEKMSRDATFEFTNNMAKVYKDPQMKMFNELAVAEVTHFWTDLYRNIARLEGRRRAKRNGDSGKKVSRFTVIKEFGDYCLAVKAGSALTQDKQIRVKIFSPYKHMEGMECQKLKSKEDCKSYMDVTIARNQYTKEYSEVYQTEWLTVSVADVEHYSMLYERSAAIHSQVMADREEFESTRAQKMTREEYRRSAMVMVPILVMMCHKRGVSTSLQYNRYIMHGLTAFASDKVSMVKNIMKDPVRTMFEASIRAMQMDYAERLTMAAPDIIKQYAEKMASEFADYDRIYMPSFKVAGMLVEFAEVMNDIYMSNLFNKEKGTAGHRLVPIMTKMSEQERHYLSMSELFPGASTGDADFDALAKNKMQTHTFNKKFVMYMGYQLRMKIIHSPDLNKGLMRAMTTTIRETMSMRSSLKSCPLYEESVDYSDKLTTATAFETMCEQVEELESIFLMDMVTCTDVVNAVFKLFPKSQIGGPREILIQAAKLRLHVKFLEKMFFELCSLHEKEMLTHADVKKDLQSNTTSVYKKMILEMNKHKDESETSPSGIILSFNTDASRWSPGFVMDSFIYFCLMMPLPVKLKEFCVTVIRAFATKTMFLPPTLTRRWQTDKLWNDKTDDKNINWLKENFVSSRGTAIFKSGMGQGMLHYFSSLFHCAKDDVMDSVMCFLFSKSYGDAACTTLISSDDLTKIFLVRYTRMSDLTAFMVSMLQVHDMTNRMANITTNWKKSALSILTEFNSYFSTGKRATDSVLKAVYNSQNIVDMTEPEKAVKSSISQMRNAFENGVKIPTIKVMFELQSVALSSIYAIDGERKEEIMRKLDCDRNSLPFQLGYMSTDRVVEQMIAGPEVLMYKEGNSEKVMEFYVKYYAAYMTDVEMAASLDYTEEEMEEDRDYMASMNGKTKMTLPFRFDTKIKEMKKRLMNKRSITREDIDKLAERNTALSFSDRSDSASFFSNMDTYMFQMDRKFGFNDAIRVHILIRALAMAGEDIALISSSRVVATDFVGVVDKILKNDDTKSKFHLTAPFKEICEMGEISEKEAEIAIPSHSYSHNKSRRINFTPMKDRVKVSDMEVFRYMMRATSNASNSAMTEANKISESLGVKLEKFIEYPIYQIKKAFPGVRRPLTMLKSYIMYWLKMTSDLNMEMMVSFPDNSEALTNMRMLYLEKYSNCGILQMFVPKKSLRVVSEERFNSALSMRGRVDRFEVGEFRGDLTEGPMENLERDPWGTDNPLLTVIPSDSLEVRFAKMAVSATDEDKYSFQGAKSLKVFYKVDVNGKMKTRYYTDLVHLGIIHEDRGASPHTVSVRMYKVSSSDVELGNITRLMHKDLSTIQIRPWAPRTDNLGVDKNAKIGNLWIKVYRIWPKTELILTRRMTKWTVLAMFNERELGGKVFFHIGTNSYKMPFNEDSENPNIRKFSELVVSSKLDLSDVEEILNDLGIADNSMFAPDNRVKEAEFEAEVDSTIYEMMDVSQAVEDYSVENVMGAISATDFGISSSSMATKLNAEMFSSIGKEEMGGMAIDFTSLAGLAGAFSGLKEIDTTAYMEEEPLEEGMSLREYMGTVYTSVKRVFEKFYSFDQRLVMNTVNDIQRNFIGDEKKRKLAVVCNMVYTAVYMKMTGTVNPTTGMSGLAQMDALRDFKENKGYNSASDTLVVMVQMVTTGLITEIFSSIGEKAPTLNSLIKLDIAFNTKRAVALDRMVTKVQDKGRSLLSIL

**>SalaUV-NL1-1-9_SS_Upp-SS1**

MATDMDKRLADFQFQVDSSGRLSAPVIAAPQAVLPSYKPEKGEMDIISDASELGTQAAFFCKEMVSSLKNAMANLTKLKPEYFPSDMGIYTSHTHRAKLFSLIKAAGMDPSTQLALFAMAVRIEDVDRMKIWLNGAECSFEVGMKKSLQDFVNMHLVQWKKEANNSPTKFCVKNICSSSPPMAALVWKMITPYEERTVGNFLANKWAAQFRFDEEIETAHLWWEQSYWNITIATSNNPDPNVFERGFQARYYKTKQGDRYLLPVVQDMEIVYMKPDDMVRTSYTGPSLVKWLRSSDVEVAICPTDDLGALFEMVSARMAKQEEEDLGK

**>SalaUV-NL1-1-11_SS_Upp-SS1**

MATDMDKRLADFQFQVDSSGRLSAPVIAAPQAVLPSYKPEKGEMDIISDASELGTQAAFFCKEMVSSLKNAMANLTKLKPEYFPSDMGIYTSHTHRAKLFSLIKAAGMDPSTQLALFAMAVRIEDVDRMKIWLNGAECSFEVGMKKSLQDFVNMHLVQWKKEANNSPTKFCVKNICSSSPPMAALVWKMITPYEERTVGNFLANKWAAQFRFDEEIETAHLWWEQSYWNITIATSNNPDPNVFERGFQARYYKTKQGDRYLLPVVQDMEIVYMKPDDMVRTSYTGPSLVKWLRSSDVEVAICPTDDLGALFEMVSARMAKQEEEDLGK

**>SalaUV-NL1-1-12_SS_Upp-SS1**

MATDMDKRLADFQFQVDSSGRLSAPVIAAPQAVLPSYKPEKGEMDIISDASELGTQAAFFCKEMVSSLKNAMANLTKLKPEYFPSDMGIYTSHTHRAKLFSLIKAAGMDPSTQLALFAMAVRIEDVDRMKIWLNGAECSFEVGMKKSLQDFVNMHLVQWKKEANNSPTKFCVKNICSSSPPMAALVWKMITPYEERTVGNFLANKWAAQFRFDEEIETAHLWWEQSYWNITIATSNNPDPNVFERGFQARYYKTKQGDRYLLPVVQDMEIVYMKPDDMVRTSYTGPSLVKWLRSSDVEVAICPTDDLGALFEMVSARMAKQEEEDLGK

**>SalaUV-NL1-1-14_SS_Upp-SS1**

MATDMDKRLADFQFQVDSSGRLSAPVIAAPQAVLPSYKPEKGEMDIISDASELGTQAAFFCKEMVSSLKNAMANLTKLKPEYFPSDMGIYTSHTHRAKLFSLIKAAGMDPSTQLALFAMAVRIEDVDRMKIWLNGAECSFEVGMKKSLQDFVNMHLVQWKKEANNSPTKFCVKNICSSSPPMAALVWKMITPYEERTVGNFLANKWAAQFRFDEEIETAHLWWEQSYWNITIATSNNPDPNVFERGFQARYYKTKQGDRYLLPVVQDMEIVYMKPDDMVRTSYTGPSLVKWLRSSDVEVAICPTDDLGALFEMVSARMAKQEEEDLGK

**>SalaUV-NL1-1-15_SS_Upp-SS1**

MATDMDKRLADFQFQVDSSGRLSAPVIASPQAVLPSYKPEKGEMDIISDASELGTQAAFFCKEMVSSLKNAMANLTKLKPEYFPSDMGIYTSHTHRAKLFSLIKAAGMDPSTQLALFAMAVRIEDVDRMKIWLNGAECSFEVGMKKSLQDFVNMHLVQWKKEANNSPTKFCVKNICSSSPPMAALVWKMITPYEERTVGNFLANKWAAQFRFDEEIETAHLWWEQSYWNITIATSNNPDPNVFERGFQARYYKTKQGDRYLLPVVQDMEIVYMKPDDMVRTSYTGPSLVKWLRSSDVEVAICPTDDLGALFEMVSARMAKQEEEDLGK

**>SalaUV-NL1-1-16_SS_Upp-SS1**

MATDMDKRLADFQFQVDSSGRLSAPVIAAPQAVLPSYKPEKGEMDIISDASELGTQAAFFCKEMVSSLKNAMANLTKLKPEYFPSDMGIYTSHTHRAKLFSLIKAAGMDPSTQLALFAMAVRIEDVDRMKIWLNGAECSFEVGMKKSLQDFVNMHLVQWKKEANNSPTKFCVKNICSSSPPMAALVWKMITPYEERTVGNFLANKWAAQFRFDEEIETAHLWWEQSYWNITIATSNNPDPNVFERGFQARYYKTKQGDRYLLPVVQDMEIVYMKPDDMVRTSYTGPSLVKWLRSSDVEVAICPTDDLGALFEMVSARMAKQEEEDLGK

**>SalaUV-NL1-1-26_SS_Upp-SS1**

MATEMDKRLSDFEFQVDSSGRLSAPVIASPQAVLPSYKPKKGEMDIISDASELGTQAAFFCKEMVTGLKGAMANLTKLKPEYFPSDMGIYTSHTHRAKLFSLIKGAGMDPSTQLALFAMAVRIEDVDRMKIWLNGAECSFEVGMKKSLQDFVNMHLVQWKKEANNSPTKFCVKNICSSSPPMAALVWKMITPYEERTVGNFLANKWAAQFRFDEEIEAAHLWWEQSYWNVTIATSNNPDPNVFERGFQARYYKTKQGDRYLLPVVQDMEIVYMKPDDMVRTSYTGSSLVKWLRSSDVEVAICPTDDLSALFEMVSARMAKQEEEDLGK

**>SalaUV-NL1-1-27_SS_Upp-SS1**

MATEMDKRLSDFQFQVDSSGRLSAPVIASPQTVLPSYKPKKGDMDIISDASELGTQAAFFCKEMVSSLKNAMANLTRLKPEYFPSDMGIYTSHTHRAKLFNLIKAEGMDPSTQLALFAMAVRIEDVDRMKIWLNGAECSFDVGMKKSLQDFVNMRLVQWKKEANNSPTKFCVKNICSSSPPMAALVWKMITPYEERTVGNFLANKWAAQFRFDEEIETAHLWWEQSYWNVTIATSNNPDPDVFERGFQAKYYKTKQGDRYLLPVVKDTEIVYMKPDDMVRTSYTGSSLVKWLRSSDVEVALCPTDDLDALFEMVSDRMAKQEEEDLGK

**>SalaUV-NL1-1-9_SS_Upp-SS0**

YTTFLLNHTIIITNRETKWNTYISLAITVVILLSAKSRKKVVKETEEREEVSSEGVKVAGKTSLSDRSETASKASSTTLSKSAKKRKQKEKKEEEEKALAKALEATRLREVEAANREEETYDREEEEGQRTTARSLRSQTVEEGVDLGENFSGRTASVVCSEVLASRVEDNARKLEEVSEKLDNILSKLSFLSQAQGASFAKENSAKLQEIFAAEQEKHKKSAQELANRKREEEKKKEEEANRERKREGEKKKEEEKRREKEEREEEKREAEKREKEARDEEEGLKGKGGVQESKEEKETEAEKRARLLKKFRSLY

**>SalaUV-NL1-1-11_SS_Upp-SS0**

YTTFLLNHTIIITNRETKWNTYISLAITVVILLSAKSRKKVVKETEEREEVSSEGVKVAGKTSLSDRSETASKASSTTLSKSAKKRKQKEKKEEEEKALAKALEATRLREVEITNREEETYDREEEGAQRTTARSLRSQTVEEGVDLGENFSGRTASVVCSEVLASRVEDNARKLEEVSEKLDNILSKLSFLSQAQEASFAKENTSQLQEIFAAEQEKHKKSAQELANRKREEEKKREEEANRERKAEGEKKKEEEKRREKEEREEEKREAEKREKEARDEEEGLKGKGGVQESKEEKETEAEKRARLLKKFRSLY

**>SalaUV-NL1-1-12_SS_Upp-SS0**

YTTFLLNHTIIITNRETKWNTYISLAITVVILLSAKSRKKVVKETEEREEVSSEGVKVAGKTSLSDRSETASKASSTTLSKSAKKRKQKEKKEEEEKALAKALEATRLREVEITNREEETYDREEEGAQRTTARSLRSQTVEEGVDLGENFSGRTASVVCSEVLASRVEDNARKLEEVSEKLDNILSKLSFLSQAQGASFAKENSAKLQEIFAAEQEKHKKSAQELANRKREEEKKREEEANRERKREGEKKKEEEKRREKEEREEEKREAEKREKEARDEEEGLKGKGGVQESKEEKETEAEKRARLLKKFRSLY

**>SalaUV-NL1-1-14_SS_Upp-SS0**

YTTFLLNHTIIITNRETKWNTYISLAITVVILLSAKSRKKVVKETEEREEVSSEGVKVAGKTSLSDRSETASKASSTTLSKSAKKRKQKEKKEEEEKALAKALEATRLREVEITNREEETYDREEEGAQRTTARSLRSQTVEEGVDLGENFSGRTASVVCSEVLASRVEDNARKLEEVSEKLDNILSKLSFLSQAQGASFAKENTSQLQEIFAAEQEKHKKSAQELANRKREEEKKREEEANRERKREGEKKKEEEKRREKEEREEEKREAEKREKEARDEEEGLKEEGGVQESKEEKETEAEKRARLLKKFRSLYK

**>SalaUV-NL1-1-15_SS_Upp-SS0**

YTTFLLNHTIIITNRETKWNTYISLAITVVILLSAKSRKKVVKETEEREEVSSEGVKVAGKTSLSDRSETASKASSTTLSKSAKKRKQKEKKEEEEKALAKALEATRLREVEITNREEETYDREEEGAQRTTARSLRSQTVEEGVDLGENFSGRTASVVCSEVLASRVEDNARKLEEVSEKLDNILSKLSFLSQAQGASFAKENSAKLQEIFAAEQEKHKKSAQELANRKREEEKKREEEANRERKAEGEKKKEEEKRREKEEREEEKREAEKREKEARDEEEGLKGKGGVQESKEEKETEAEKRARLLKKFRSLY

**>SalaUV-NL1-1-16_SS_Upp-SS0**

YTTFLLNHTIIITNRETKWNTYISLAITVVILLSAKSRKKVVKETEEREEVSSEGVKVAGKTSLSDRSETASKASSTTLSKSAKKRKQKEKKEEEEKALAKALEATRLREVEITNREEETYDREEEGAQRTTARSLRSQTVEEGVDLGENFSGRTASVVCSEVLASRVEDNARKLEEVSEKLDNILSKLSFLSQAQGASFAKENTSQLQEIFAAEQEKHKKSAQELANRKREGEKKREEEANRERKAEGEKKKEEEKRREKEEREEEKREAEKREKEARDEEEGLKGKGGVQESKEEKETEAEKRARLLKKFRSLY

**>SalaUV-NL1-1-26_SS_Upp-SS0**

YTTFLLNHTIIITNRETKWNTYISLAITVAILLSAKSRKKVAKETEEKEEALSEEVKVAGKTGFSDKSETASKTSSTTLSKSAKKRKQREKKEEEEKALAKALEATRLKEVEVANKEEETYDREEEGGQRTTARSLRSQTVEEGVDLGENFSGRTASVVCSEVLASRVEDNARKLEEVSEKLDNILSKLSLLSQAQGASFAKENTSQLQEIFAAEQEKHRKAAQELANRKREVEKKKEEEAKRERKAEEEKKKEEERKREKEEKEEEEREAEKREEEARDEEEGLKEEKGVQESKGEKETEAEKRARLLKKFRSLY

**>SalaUV-NL1-1-27_SS_Upp-SS0**

YTTFLLNHTIIITNRETKWNTYISLAITVAILLSSKSRKKVVKETEEREEVSSEGVKVAGKTGLSDKSDTASKTSSTTLSKSAKKRKQKEKKEEEEKALAKALEATRLKEVEAANKEEETHDREEEEGQRTTARSLRSQTVEEGVDLGESFSGRTASVVCSEVLASRVEDNARKLEEVSEKLDNILSKLSLLSQAQGASFAKENTSQLQEIFAAEQEKHRKAAQELVNRKREVEKKREEEAKRERKAEEEKKKEEEKRREKEKREEEKREAEKREEEARDEEEGLKEERGVQESKEEKETEAEKRARLLKKFRSLC
