## Supplemental Figure SF1 for "Extreme GC3 Codon Bias in a Novel Brown Seaweed Virus Results in Pseudoambigrammatic Characteristics"

### Slide 1
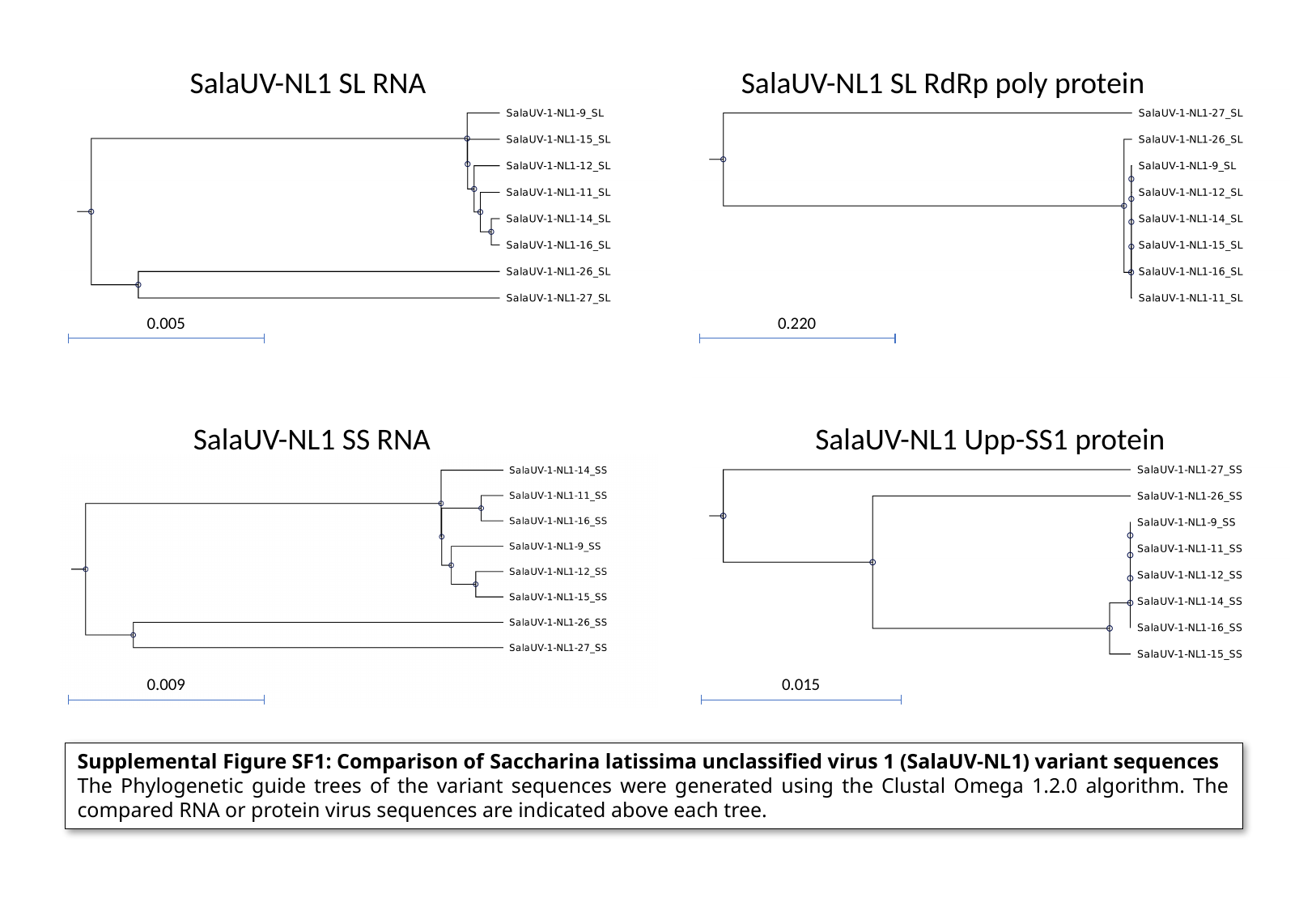

SalaUV-NL1 SL RNA
SalaUV-NL1 SL RdRp poly protein
0.005
0.220
SalaUV-NL1 SS RNA
SalaUV-NL1 Upp-SS1 protein
0.009
0.015
Supplemental Figure SF1: Comparison of Saccharina latissima unclassified virus 1 (SalaUV-NL1) variant sequences
The Phylogenetic guide trees of the variant sequences were generated using the Clustal Omega 1.2.0 algorithm. The compared RNA or protein virus sequences are indicated above each tree.
