## Supplemental Figure SF2 for "Extreme GC3 Codon Bias in a Novel Brown Seaweed Virus Results in Pseudoambigrammatic Characteristics"

### Slide 1
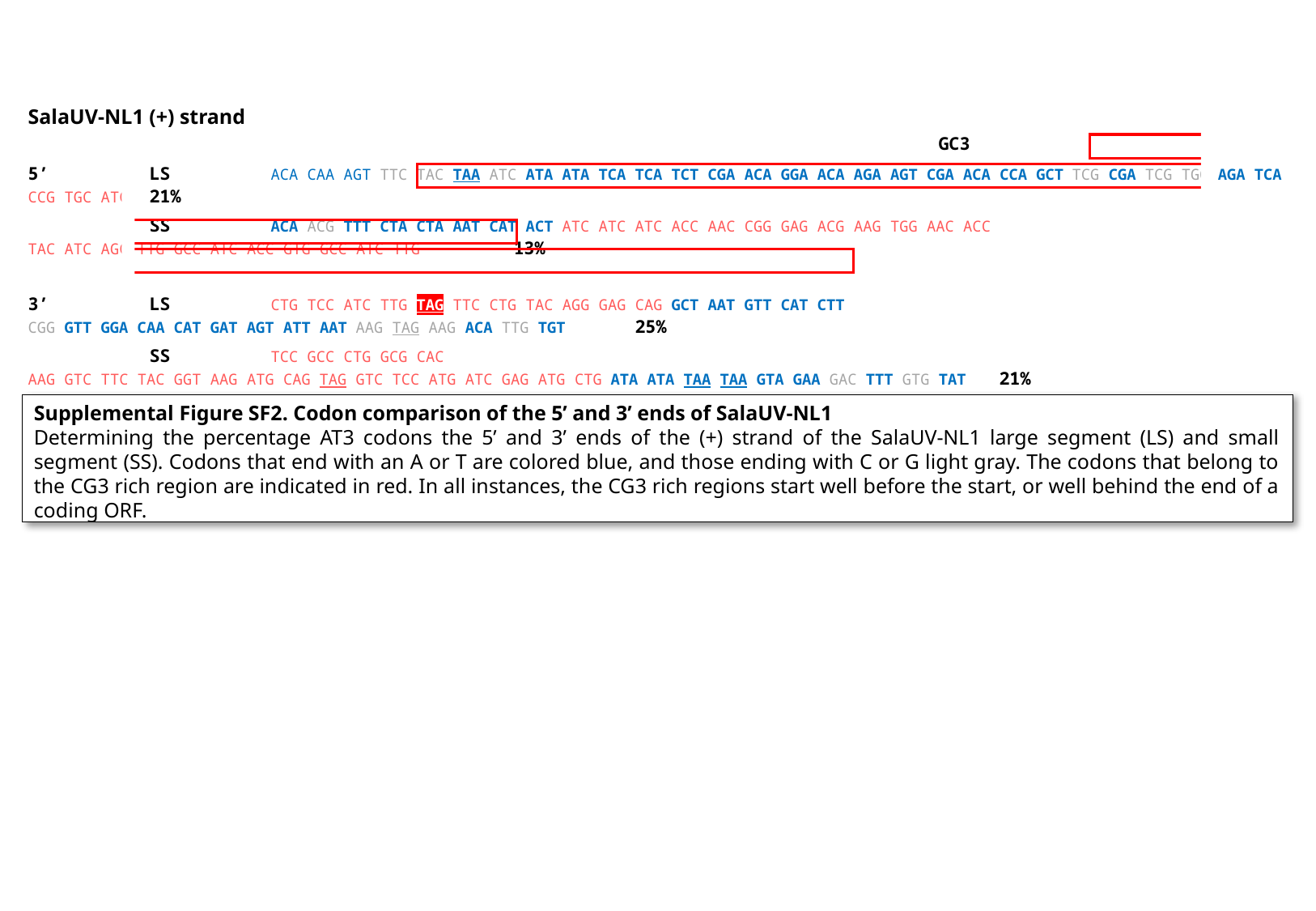

SalaUV-NL1 (+) strand																 GC3
5’	LS	ACA CAA AGT ttc tac taa atc ata atA tca tCa tct cga aca gga aca aga agt cga aca cca gct tcg cga tcg tgg aga tca ccg tgc atc	21%
	SS	aca acg ttt cta cta aat cat act atc atC atc acc aac cgg gag acg aag tgg aac acc tac atc agc ttg gcc atc acc gtg gcc atc ttg	13%
3’	LS	ctg tcc atc ttg tag ttc ctg tac agg gag cag gct aat gtt cat ctt cgg gtt gga caa cat gat agt att aat aag tag aag aca ttg tgt	25%
	SS	tcc gcc ctg gcg cac aag gtc ttc tac ggt aag atg cag tag gtc tcc atg atc gag atg ctg ata ATA taa taA gta gaa gac ttt gtg tat	21%
Supplemental Figure SF2. Codon comparison of the 5’ and 3’ ends of SalaUV-NL1
Determining the percentage AT3 codons the 5’ and 3’ ends of the (+) strand of the SalaUV-NL1 large segment (LS) and small segment (SS). Codons that end with an A or T are colored blue, and those ending with C or G light gray. The codons that belong to the CG3 rich region are indicated in red. In all instances, the CG3 rich regions start well before the start, or well behind the end of a coding ORF.
